# Synthetic gene circuits enable systems-level biosensor discovery at the host-microbe interface

**DOI:** 10.1101/557934

**Authors:** Alexander D Naydich, Shannon N Nangle, Johannes J Bues, Disha Trivedi, Nabeel Nissar, Mara C Inniss, Matthew J Neiderhuber, Jeffrey C Way, Pamela A Silver, David T Riglar

**Affiliations:** Department of Systems Biology, Harvard Medical School, Boston, MA 02215; Wyss Institute for Biologically Inspired Engineering, Boston, MA 02215

**Author notes:** Present addresses: J.J.B.: Ecole polytechnique fédérale de Lausanne, Lausanne, Switzerland; N.N.: Littleton MA, 01460; M.C.I.: Obsidian Therapeutics, Cambridge, MA, 02138; M.J.N.: UNC Chapel Hill, Chapel Hill, NC 27599.

## Abstract

The composition and function of the gut microbiota are strongly associated with human health, and dysbiosis is linked to an array of diseases, ranging from obesity and diabetes to infection and inflammation. Engineering synthetic circuits into gut bacteria to sense, record and respond to *in vivo* signals is a promising new approach for the diagnosis, treatment and prevention of disease. Here, we repurpose a synthetic bacterial memory circuit to rapidly screen for and discover new *in vivo*-responsive biosensors in commensal gut *Escherichia coli*. We develop a pipeline for rapid systems-level library construction and screening, using next-generation sequencing and computational analysis, which demonstrates remarkably robust identification of responsive biosensors from pooled libraries. By testing both genome-wide and curated libraries of potential biosensor triggers—each consisting of a promoter and ribosome binding site (RBS)—and using RBS variation to augment the range of trigger sensitivity, we identify and validate triggers that selectively activate our synthetic memory circuit during transit through the gut. We further identify biosensors with increased response in the inflamed gut through comparative screening of our libraries in healthy mice and those with intestinal inflammation. Our results demonstrate the power of systems-level screening for the identification of novel biosensors in the gut and provide a platform for disease-specific screening using synthetic circuits, capable of contributing to both the understanding and clinical management of intestinal illness.

**IMPORTANCE:** The gut is a largely obscure and inaccessible environment. The use of live, engineered probiotics to detect and respond to disease signals *in vivo* represents a new frontier in the management of gut diseases. Engineered probiotics have also shown promise as a novel mechanism for drug delivery. However, the design and construction of effective strains that respond to the *in vivo* environment is hindered by our limited understanding of bacterial behavior in the gut. Our work expands the pool of potential biosensors for the healthy and diseased gut, providing insight into host-microbe interactions and enabling future development of increasingly complex synthetic circuits. This method also provides a framework for rapid prototyping of engineered systems and for application across bacterial strains and disease models.

## INTRODUCTION

Recent advances in our understanding of both the human microbiota and biological engineering techniques have created myriad possibilities for the development of synthetic microbes for *in vivo* clinical applications (1,2). Bacteria residing in the gut are uniquely positioned to monitor a variety of host, microbial, and environmental factors and to respond to changes in intestinal homeostasis. Engineered gut bacteria also offer the potential for *in vivo* production and delivery of therapeutics (2).

Environment- and disease-responsive functions, which could minimize both the metabolic burden of engineered systems on the bacteria and off-target effects on the patient, offer exciting prospects for clinical applications. To this end, recent *in vivo* approaches have developed sensors responding to inflammation (3,4), intestinal bleeding (5), and pathogen quorum-sensing systems (6,7). However, the construction of disease-responsive circuits in bacteria has been hindered by the limited number of characterized bacterial systems that can be reliably employed as sensors.

Mining the genomes of native gut bacteria is a promising approach for discovering new sensors that respond under conditions of interest, such as in the healthy or diseased gut. To date, these efforts have largely relied on transcriptome sequencing and proteomics of fecal samples. However, to obtain an instantaneous snapshot of bacterial behavior inside the gut using these techniques, invasive sampling is required (i.e., colonoscopy and biopsy). Furthermore, transient or low-abundance signals may not be detected, and any responsive genetic elements identified with these techniques may not function predictably when employed in synthetic circuits. Approaches such as *in vivo* expression technology (IVET) and recombinase-based IVET (RIVET) have also been used to track *in vivo*-expressed genes non-invasively, but detect only constitutive expression (for IVET) and may have high false-positive rates (8). Nevertheless, these technologies show the potential for systems-level approaches to interrogate the behavior of the microbiota.

We have previously developed an approach for non-invasive measurement of bacterial responses in the gut, based on a robust synthetic memory circuit, which records environmental stimuli via a transcriptional trigger (3,9). When activated, the trigger turns on a memory switch, which can retain the memory-on state for over a week in the gut (9). After the bacteria pass through the host, their memory state can be determined via reporter gene expression, enabling non-invasive readout of transient signals within the gut. The circuit can maintain functional and genetic stability during six months’ colonization of the mouse gut, demonstrating its suitability for longitudinal studies and its potential to support the development of stable, engineered biosensors for *in vivo* deployment (3).

Here, we adapt this memory circuit for parallel, high-throughput screening of hundreds of potential triggers. We apply this method to identify new biosensors responding specifically to the *in vivo* gut environment. Through comparison between healthy mice and those suffering from inflammation, we also identify triggers that respond differentially during disease. Together, these results provide a platform for *in vivo* non-invasive biosensor discovery and longitudinal testing.

## RESULTS

### Bacterial memory as a biosensor screening tool

To enable screening of new potential biosensors in parallel, we modified our previously-developed *E. coli* memory circuit, which is based on the λ phage lysis–lysogeny switch (Fig. S1A) (9). This modified circuit is referred to as the high-throughput memory system (HTMS) (Fig. 1A). Both the original memory circuit and the HTMS consist of a trigger—based on a transcriptional promoter activated in the presence of a certain stimulus—and a memory switch. The memory-on and memory-off states of the switch correspond to the mutually-repressive proteins Cro and CI, respectively. Additionally, a β-galactosidase (LacZ) reporter is produced in the memory-on state.

**FIG 1.**
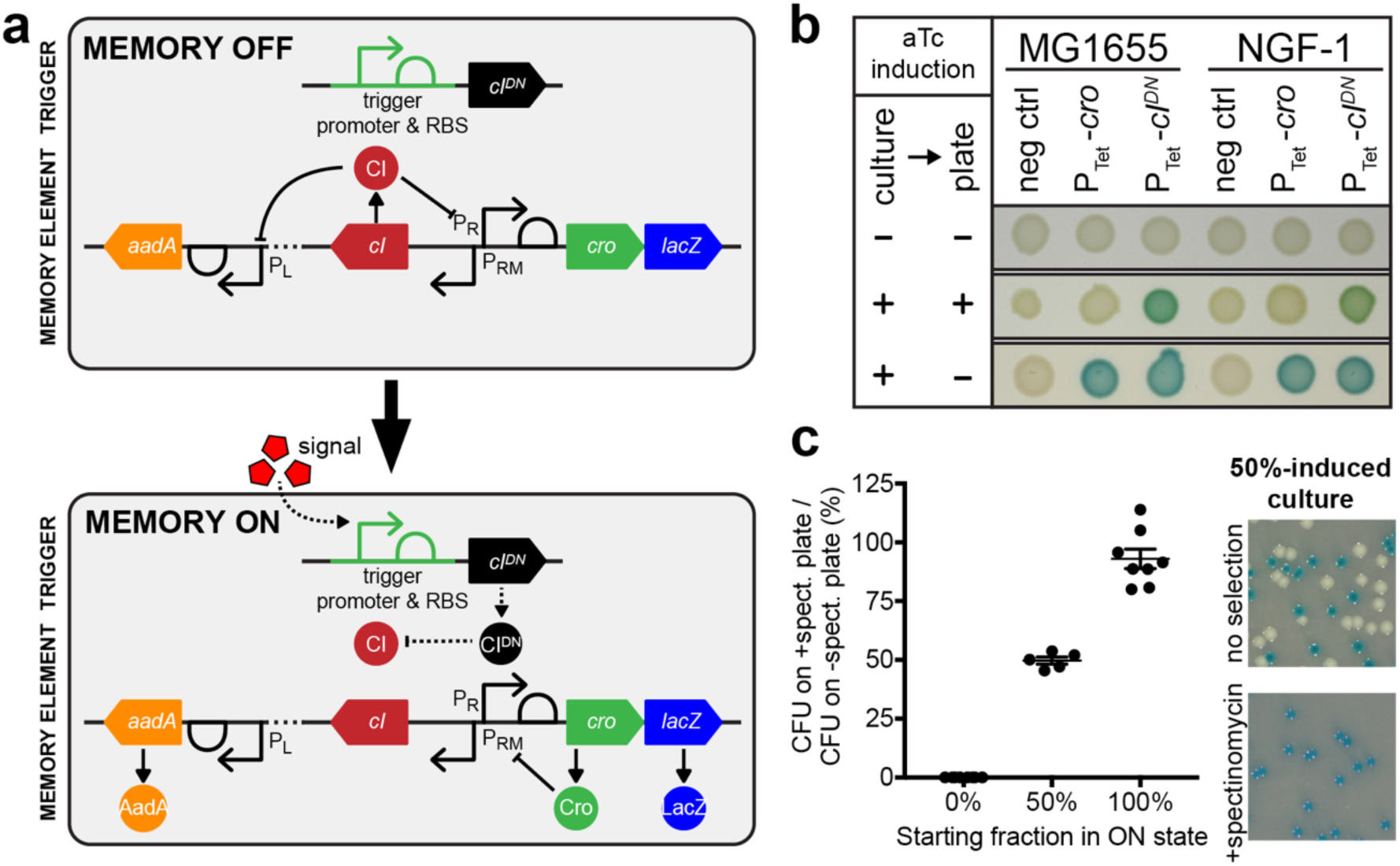
Design of a High-Throughput Memory System (HTMS). (A) HTMS circuit design in memory-off and memory-on states. (B) Comparison of memory element induction with *cro* and *cIDN* triggers illustrates differences in induction dynamics. Control and memory strains with P_tet_ triggers (PAS132, PAS133, PAS807, PAS808) were grown in liquid media, then spotted on indicator plates, each with or without aTc induction (100 ng/ml). (C) Selection of memory-on HTMS with spectinomycin. Memory-off, memory-on, and 50-50 mixed cultures of PAS810 were plated on indicator plates with or without spectinomycin (inset photo). Graph shows comparison of CFU counts between +spectinomycin and -spectinomycin plates. Error bars represent SE of eight biological replicates (for 0% and 100%) and five biological replicates (for 50%).

One key modification for the HTMS is the triggering of memory using a dominant-negative mutant of the *cI* gene (*cIDN*), instead of *cro* used in the original trigger. When the trigger promoter is activated by a stimulus, CIDN monomers, which have an N55K mutation in the DNA binding region (10) dimerize with wild-type (WT) CI monomers expressed in the memory-off state, creating heterodimers that are deficient in DNA-binding. This leads to derepression of P_R_ and transition to the memory-on state. As with the CI used in the memory element, CIDN carries a mutation to prevent RecA-mediated cleavage (ind-) (11).

Use of CIDN in the trigger ensures that there is no delay of switching to the memory-on state in the case of high, or constant, expression of the trigger promoter. To test this, a P_tet_ trigger driving *cIDN* or *cro* was integrated into *E. coli* K-12 MG1655 and NGF-1 strains containing a memory element. When grown in the presence of a high concentration (100 ng/ml) of anhydrotetracycline (aTc) inducer, *cIDN-*triggered strains showed switching to the memory-on state, while *cro*-triggered strains switched only after a subsequent period of growth in the absence of aTc (Fig. 1B).

The original memory circuit expresses a *lacZ* reporter gene for screening on indicator plates (9). To analyze pooled libraries containing many strains with varied trigger promoters, the HTMS also expresses a spectinomycin-selectable resistance gene (*aadA)* in the memory-on state.

This antibiotic-selectable memory maintains response characteristics similar to the original memory element. To test this, a P_tet_ trigger driving *cIDN* was integrated into strains containing *lacZ* (original) or *aadA+lacZ* (HTMS) memory elements, creating PAS809 and PAS810, respectively (see Table 1 for strain list). Strains were induced by aTc (0-100 ng/ml) and the response quantified by plating cultures on indicator plates containing 5-bromo-4-chloro-3-indolyl-β-d-galactopyranoside (X-gal), which turns blue in the presence of LacZ, indicating a memory-on state (Fig. S1B). Both strains responded similarly to aTc (*original memory* EC50: 4.1-4.6 ng/ml, 95% CI; *HTMS* EC50: 4.0-4.3 ng/ml, 95% CI), confirming the circuit’s modularity to additional reporters in the memory-on state.

**TABLE 1.** Key memory strains used in this study.

| Strain | E. coli strain | Memory | Trigger Promoter/RBS | Trigger Gene | Source |
| --- | --- | --- | --- | --- | --- |
| PAS132 | K-12 MG1655 | original | tet | *cro* | (9) |
| PAS133 | NGF-1 | original | tet | *cro* | (9) |
| PAS807 | K-12 MG1655 | original | tet | *cIDN* | this study |
| PAS808 | NGF-1 | original | tet | *cIDN* | this study |
| PAS809 | NGF-1 | original | tet | *cIDN* | this study |
| PAS810 | NGF-1 | HTMS | tet | *cIDN* | this study |
| PAS811 | NGF-1 | HTMS | - | - | this study |
| PAS812 | NGF-1 | HTMS | *fabR* | *cIDN* | this study |
| PAS813 | NGF-1 | HTMS | *ydiL* | *cIDN* | this study |
| PAS814 | NGF-1 | HTMS | *ydjL* | *cIDN* | this study |
| PAS815 | NGF-1 | HTMS | - | *cIDN* | this study |
| PAS816 | NGF-1 | HTMS | *ynfE15* | *cIDN* | this study |
| PAS817 | NGF-1 | HTMS | *yeaRWT* | *cIDN* | this study |
| PAS818 | NGF-1 | HTMS | *ynfEWT* | *cIDN* | this study |
| PAS819 | NGF-1 | HTMS | *ynfE17* | *cIDN* | this study |

The HTMS allows faithful selection of memory-on colonies with spectinomycin treatment. Plating of fully memory-off, fully memory-on, and 50-50 mixed cultures of PAS810, on indicator plates with and without spectinomycin further demonstrated that all spectinomycin-selected colonies were also *lacZ* positive (Fig. 1C). Spectinomycin did not yield false-positive results by inducing memory switching (fully memory-off: 0% +/−0% SE, n = 8), nor excessive false-negative results through inhibition of memory-on bacterial growth (fully memory-on: 93.0% +/-4.2% SE, n = 8; 50-50 mix: 49.7% +/-1.5% SE, n = 5) (Fig. 1C). Together these results demonstrate the ability of the HTMS to measure biosensor response and allow selection for downstream pooled analyses.

### Biosensor library construction

To build biosensor libraries for genomic integration, we adapted a Tn7 transposon genome insertion plasmid (12) for rapid Golden Gate assembly (13) of bacterial promoters upstream of the *cIDN* trigger and insertion into the genome of memory bacteria (Fig. 2A and S2). The modularity of this cloning strategy allows for adjustment of trigger sensitivity through incorporation of ribosomal binding site (RBS) variants, which vary the translation rate of mRNA transcripts (Fig. 2A). To test this concept, triggers consisting of a P_tet_ promoter combined with nine synthetic RBS sequences—previously demonstrated to vary widely in their translation strength (14) (Fig. 2A)—were constructed and inserted into the genome of HTMS bacteria, and the HTMS response to varying concentrations of aTc (0-100 ng/ml) was characterized (Fig. 2B). The RBS variants differed in their extent of memory induction at 0.1-10 ng/ml aTc (EC50 ranging from 0.5 to 4.1 ng/ml for responsive strains), illustrating our ability to tune trigger sensitivity.

**FIG 2.**
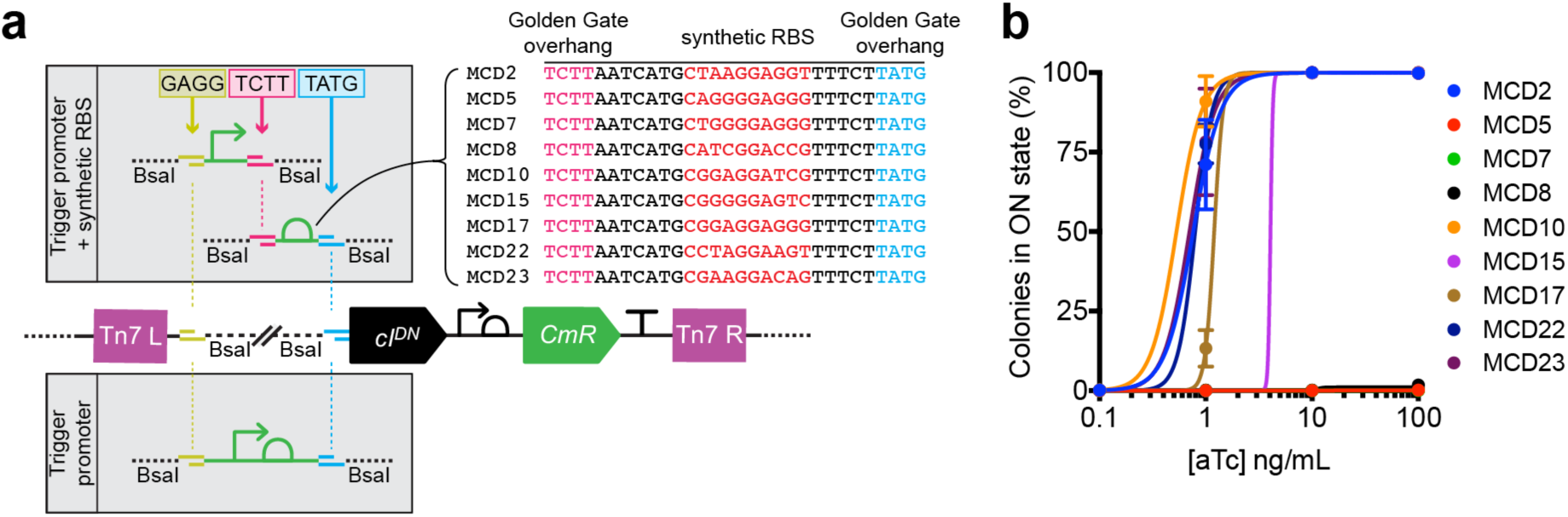
Tuning of trigger sensitivity is achieved through modular incorporation of RBS variants. (A) Strategy for insertion of trigger promoter and RBS variants (14) by Golden Gate assembly (13) into a Tn7 transposon plasmid containing the *cIDN* gene (Fig. S2). (B) Response curves for RBS variants of a Ptet-*cIDN* trigger in the HTMS chassis strain PAS811 induced with varying concentrations of aTc. Memory response was assessed by plating of bacteria on indicator plates after induction with aTc in liquid culture. Error bars represent SE of three biological replicates.

We explored two approaches for generating biosensor libraries: 1) a genome-wide collection of trigger promoters that would enable screening of a bacterium’s entire range of transcriptional responses (MG1655 library), and 2) a curated collection of promoters with sensitivity variants aimed at detecting inflammation (Nissle 1917 library). Both libraries were assembled into an HTMS-containing *E. coli* NGF-1 strain (PAS811), as NGF-1 has proven to be an efficient and persistent colonizer in the mouse gut (3,9,15). The genome-wide library was sourced from a previously published collection of 1600 unique promoters from *E. coli* K-12 MG1655 (16). Promoters and their wild-type RBSs were amplified by PCR from this collection, assembled into our transposon plasmid, and integrated as triggers into the genome of PAS811. Because our method focuses on detecting off-to-on sensor transitions, the resultant library was further subsampled by pooling 500 colonies that were LacZ*-*negative under routine *in vitro* culture. Sequencing confirmed the presence of 155 unique strains in this final genome-wide library.

Our second library was constructed with a subset of promoters sourced from the human probiotic *E. coli* Nissle 1917, which are involved in anaerobic respiration of sulfur- or nitrogen-oxides or nitrate, produced by the gut epithelium during inflammation (17,18). For each promoter, a trigger with its wild-type RBS, as well as with five different synthetic RBSs (MCD5, MCD10, MCD15, MCD17 and MCD23) (14) were included to tune sensitivity. Sequencing confirmed that the assembled library contained 61 unique strains out of 66 total designed constructs.

### Parallel analysis faithfully reports biosensor response

To screen for biosensor response, HTMS libraries are exposed to a condition of interest (Fig. 3A), and put through a processing, sequencing and analysis pipeline (Fig. 3B). After exposure, HTMS bacteria are recovered and cultured. The initial culture is split into two and back-diluted, and one of the two new cultures is subjected to spectinomycin selection. Following selection, the trigger regions of both cultures are sequenced and analyzed to produce an odds ratio for each trigger promoter in the library, corresponding to that trigger’s memory state. To calculate odds ratios, results are normalized to a positive normalization strain (PAS812) which remained in a memory-on state (Fig. 3B).

**FIG 3.**
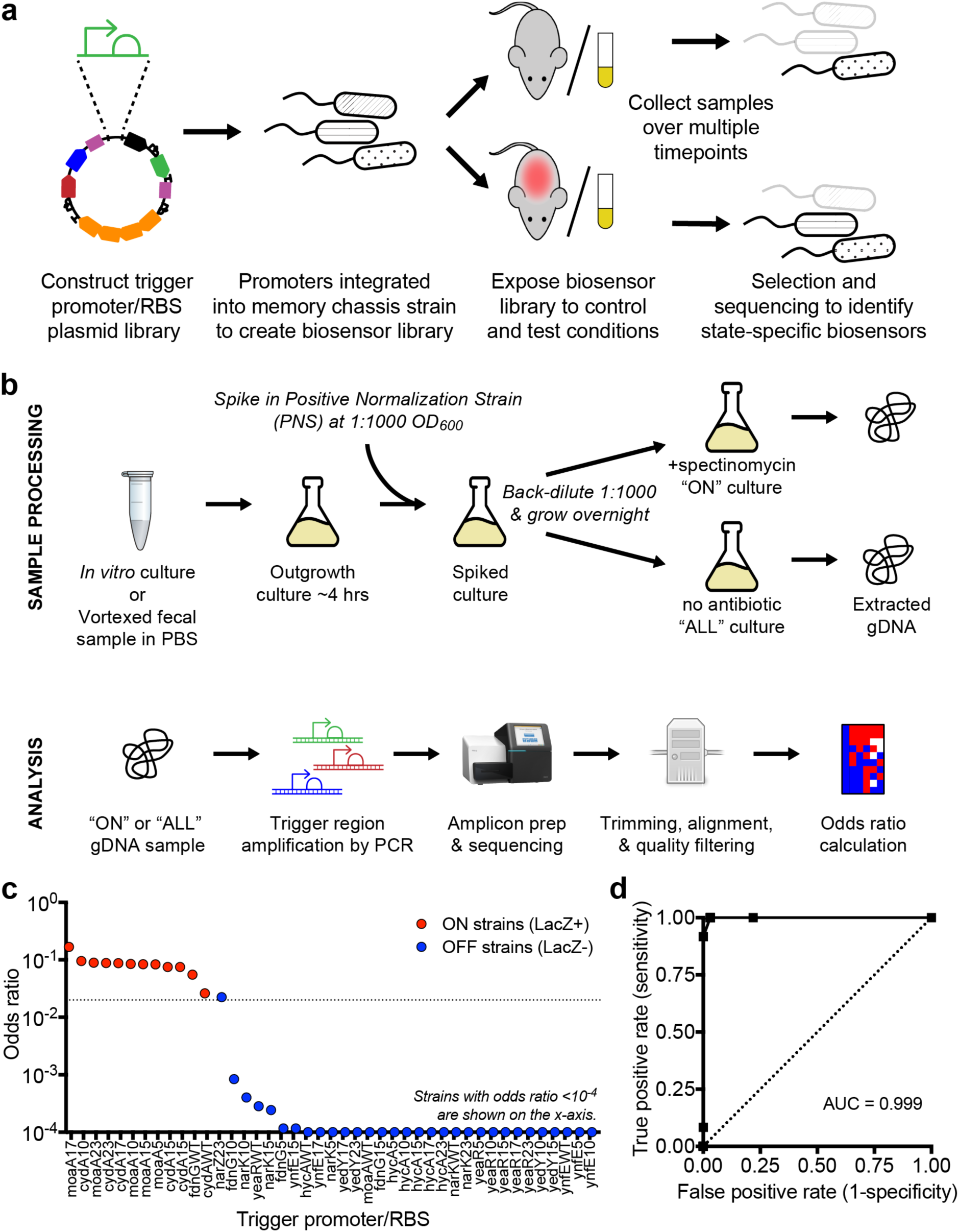
Biosensor library screening and analysis. (A) Libraries were constructed as plasmids, integrated into the genome at single copy, and screened as a pool for differential response to growth in control and test environments. (B) Post-exposure library sample processing, selection for memory-on strains, sequencing and analysis. The consistently memory-on positive normalization strain (PAS812) is spiked in prior to spectinomycin selection and used for calculation of odds ratios. (C) Calculated odds ratio from *in vitro* pooled growth in LB medium vs. memory state assessed by plating individual library strains on LB agar indicator plates. 44 strains subsampled from the Nissle 1917 library were tested. The on-off odds ratio cutoff used in the subsequent *in vivo* screens (0.02) is indicated by the dotted line. (D) ROC curve for varying odds ratio cutoffs as an indicator of memory state.

Pooled library analysis is predictive of the on/off state of HTMS bacteria. The Nissle 1917 library was cultured aerobically in liquid media and analyzed to obtain odds ratios as described above. Concurrently, individual strains from this library were grown on indicator plates to assess each strain’s *in vitro* memory state directly. Both tests showed strong agreement, with strains that were LacZ-positive also displaying higher odds ratios (Fig. 3C). Receiver operating characteristic analysis confirmed efficient distinction between memory-on and memory-off states, with an odds ratio of approximately 0.02 delimiting the boundary (Fig. 3D). This confirmed our sequencing method as a reliable indicator of biosensor memory state.

### Differential biosensor response in the healthy mouse gut

To screen for biosensor response to growth within the murine gut, the MG1655 library was administered to specific-pathogen free (SPF) mice by oral gavage (∼10^7^ bacteria/mouse), and fecal samples were collected over one (n=2) or seven (n=3) days. High library diversity was maintained in both experiments (92% and 82% of strains identified in gavage samples present at experiment endpoint, respectively). Analysis of HTMS strains recovered from gavage suspension and fecal samples identified 23 unique strains that responded specifically to growth within the gut (gavage: odds ratio < 0.02; fecal samples: ≥ 1 timepoint odds ratio ≥ 0.02 and p < 0.05) (Fig. 4A and 4B; Data Set S1 and S2). Five strains (containing ydiL, ydjL, gatY, gcvA and ubiG triggers) were detected in the memory-on state in at least 4 of 5 mice. The two most consistent responders (ydjL and ydiL triggers) were selected for follow-up testing.

**FIG 4.**
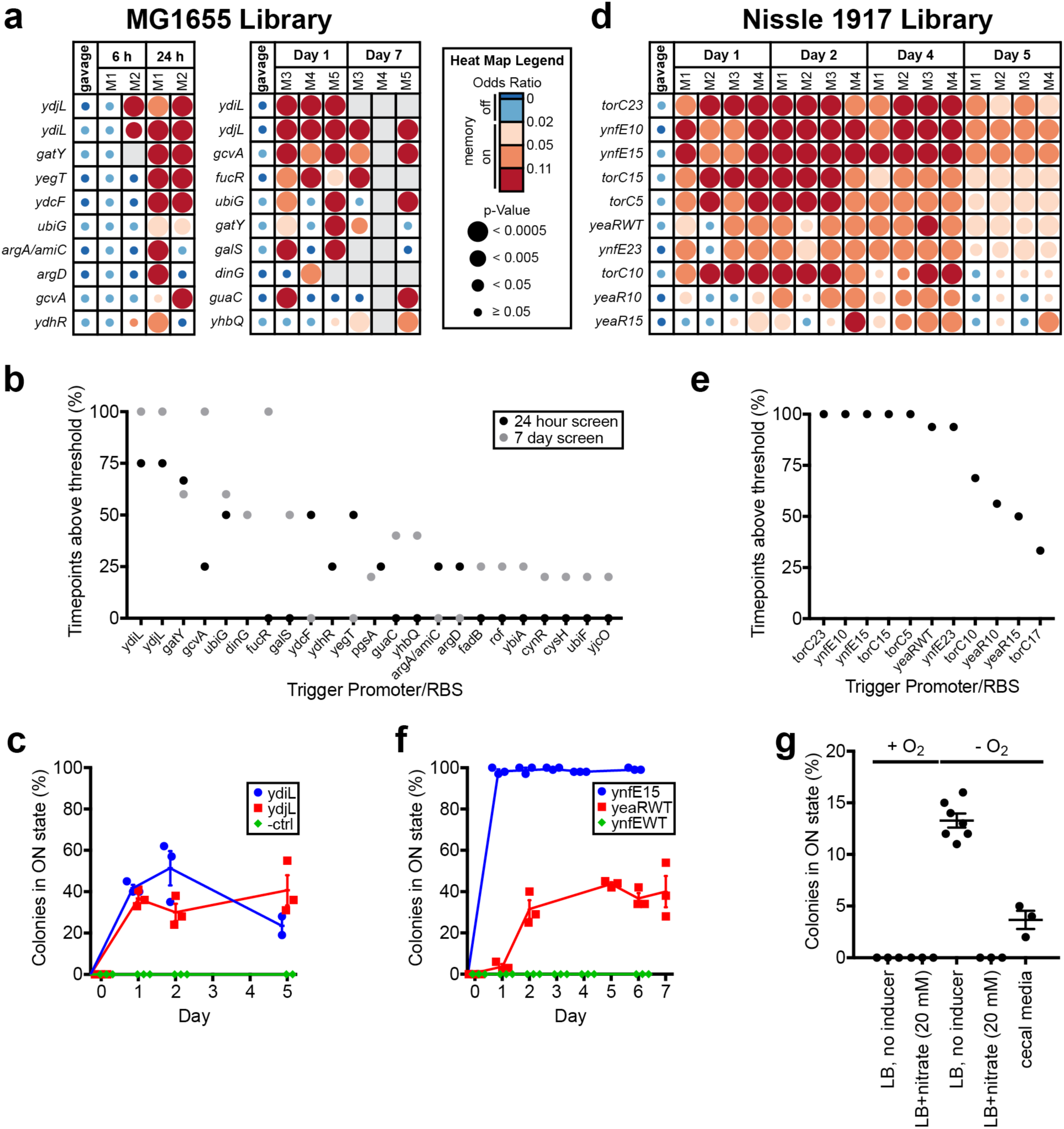
Library screening and individual sensor testing identifies biosensors of the *in vivo* gut environment. (A) Screen of MG1655 library in BALB/c mice over one day (left; n = 2) and over seven days (right; n = 3). Odds ratio heat maps of the top 10 hits sorted by percentage of positive timepoints (odds ratio ≥ 0.02 and p < 0.05) over the course of the experiment. Blank spaces on heat maps represent insufficient sequencing coverage. See Data Set S1 and S2 for full results. (B) Percentage of positive timepoints (odds ratio ≥ 0.02 and p < 0.05) for all positive hits from two MG1655 library screens (top 10 shown in Fig. 4A). (C) Response of individual strains from MG1655 library in healthy mice. HTMS strains containing triggers ydiL (PAS813), ydjL (PAS814) and an empty trigger (PAS815) were administered as monocultures to BALB/c mice (n = 3). Memory response was assessed by plating of HTMS bacteria recovered from fecal samples on indicator plates. Error bars represent SE. (D) Screen of Nissle 1917 library in C57BL/6J mice (n = 4) over five days. Odds ratio heat map of the top 10 hits sorted by percentage of positive timepoints (odds ratio ≥ 0.02 and p < 0.05) over the course of the experiment. Blank spaces on heat map represent insufficient sequencing coverage. See Data Set S3 for full results. (E) Percentage of positive timepoints (odds ratio ≥ 0.02 and p < 0.05) for all hits from Nissle 1917 library screen in healthy mice (top 10 shown in Fig. 4D). (F) Response of individual strains from Nissle 1917 library in healthy mice. HTMS strains containing the ynfE trigger with MCD15 (PAS816), the yeaR trigger with its WT RBS (PAS817) and the ynfE trigger with its WT RBS (PAS818) were administered as monocultures to C57BL/6J mice (n = 3). Memory response was assessed by plating of HTMS bacteria recovered from fecal samples on indicator plates. Error bars represent SE. (G) Response of the ynfE15 trigger strain (PAS816) to *in vitro* growth in rich media with and without 20 mM nitrate, with and without oxygen, and in the presence of mouse cecum fluid medium. Memory response was assessed by plating on indicator plates after growth in liquid culture. Error bars represent SE. (n = 7 for −O_2_, LB, no inducer condition; n = 3 for all other conditions.)

To validate the response of the ydiL and ydjL triggers during gut transit, memory bacteria containing these triggers (ydiL: PAS813; ydjL: PAS814) or a promoterless *cIDN* gene (negative control: PAS815) were administered to SPF mice as monocultures. Fecal samples were collected and analyzed over the subsequent five days. Culture on indicator plates demonstrated an absence of memory activation in all three strains prior to gavage. However, when recovered from fecal samples PAS813 and PAS814 colonies were consistently memory-on, confirming activation during gut transit (at Day 2, PAS813: 51% +/-8% SE; PAS814: 30% +/-4% SE; negative control: 0% +/-0% SE; n = 3 per strain) (Fig. 4C).

The Nissle 1917 library was also screened to discover promoters responding in the healthy mouse gut. Testing of the Nissle 1917 library over five days following gavage (∼10^7^ bacteria/mouse) identified 11 strains that specifically responded to *in vivo* growth (Data Set S3). Ten of these, derived from three unique promoters (*ynfEFGH, torCAD*, and *yeaR-yoaG* operons) registered a memory-on state in the majority of timepoints and all mice tested (n=4) (Fig. 4D and 4E). Promoter response was similar during parallel analysis in the inflamed mouse gut (n = 4; see below and Fig. 5 for experimental details), further validating these results (Fig. S3 and Data Set S3).

**FIG 5.**
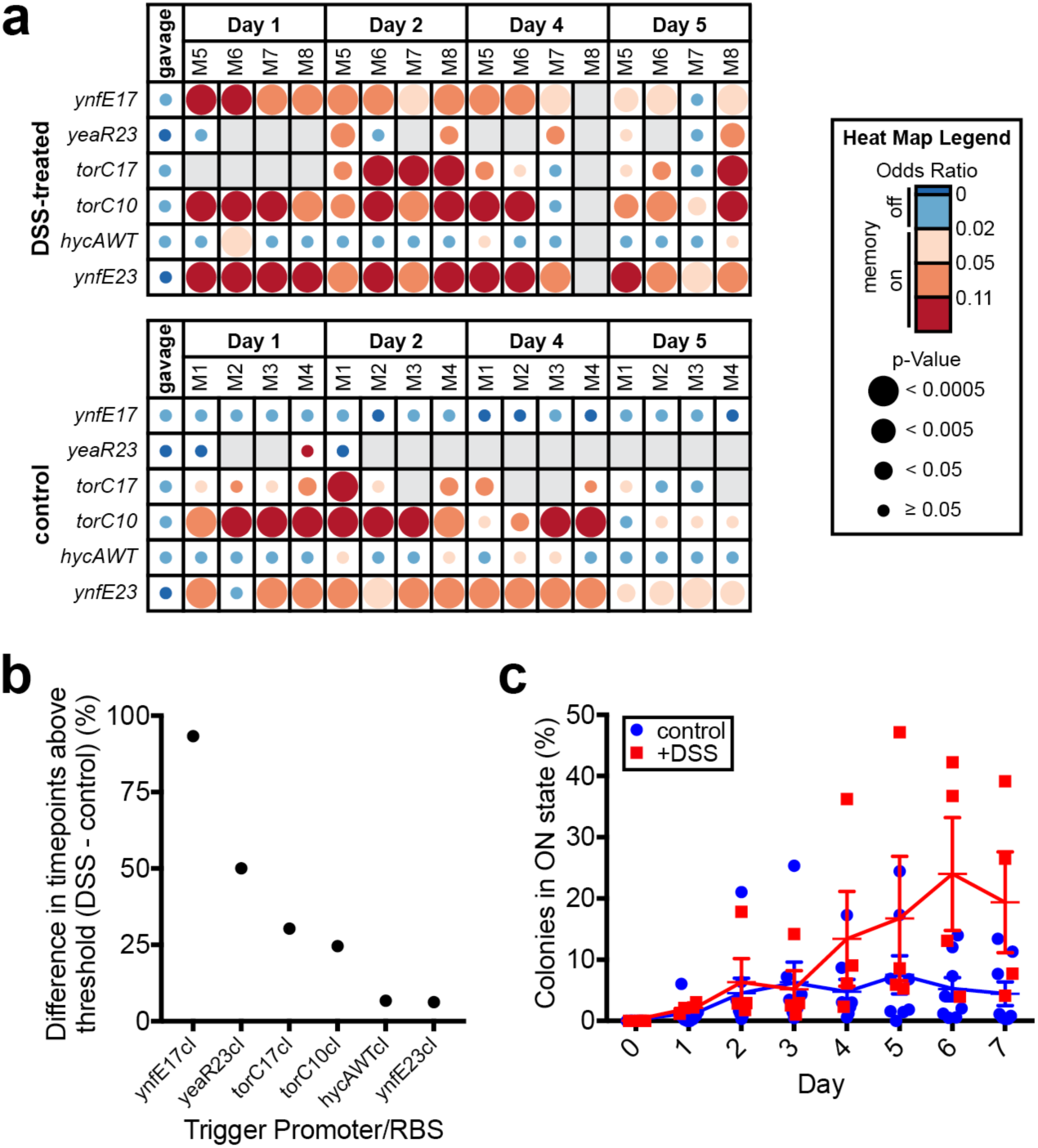
Library screening and individual sensor testing identifies biosensors with increased response during inflammation. (A) Screen of Nissle 1917 library in DSS-treated C57BL/6J mice (n = 4) over five days and comparison with results of previous screen in healthy C57BL/6J mice (n = 4; control group; also see Fig. 4D and 4E). To identify sensors responding more strongly to the inflamed state, the fraction of timepoints registering memory-on (odds ratio ≥ 0.02 and p < 0.05) in the control group was subtracted from the fraction in the DSS-treated group for each strain in the library, and strains were sorted according to the greatest difference between the two groups. Odds ratio heat map of the six strains registering a positive difference between the two groups is shown. Blank spaces on heat map represent insufficient sequencing coverage. See Data Set S3 for full results. (B) Differences in percentage of positive timepoints (odds ratio ≥ 0.02 and p < 0.05) between DSS-treated and control group mice for strains showing a positive difference between the two groups (heat map shown in Fig. 5A). (C) Response of HTMS strain containing the ynfE trigger with MCD17 (PAS819) in DSS-treated mice (n = 4) and healthy mice (n = 8). Memory response was assessed by plating of HTMS bacteria recovered from fecal samples on indicator plates. Error bars represent SE.

RBS variation to adjust trigger sensitivity affects *in vivo* sensing capacity. Variation in sensor response based on trigger RBS was most notably observed with the *ynfEFGH* promoter: WT, MCD5 and MCD17 RBSs showed no response throughout the screening experiment, whereas MCD10, MCD15, and MCD23 registered as memory-on in 100%, 100% and 94% of timepoints, respectively. (Note: Nissle 1917 library strains are referred to below by an abbreviation consisting of the first gene of the operon from which they are derived and the number of their synthetic RBS. For instance, “ynfE15” denotes the trigger consisting of the *ynfEFGH* promoter with MCD15.) To validate these findings, ynfE15 (PAS816), yeaRWT (PAS817), and ynfEWT (PAS818, used here as a negative control) strains were administered to SPF mice as monocultures, with memory status determined by culture on indicator plates (at Day 2, ynfE15: 99% +/-1% SE; yeaRWT: 31% +/-4% SE; ynfEWT: 0% +/-0% SE; n = 3) (Fig. 4F). Of note, changing the ynfE trigger RBS from WT to MCD15 increased memory-on rate in the gut from 0% to nearly 100%.

Together these results demonstrate the ability for HTMS analysis to rapidly identify biosensors *in vivo* and the power of varying trigger sensitivity to tune the strength of biosensor response.

### *In vitro* induction of *in* vivo-responding biosensor strains

The Nissle 1917 library includes some promoters with previously characterized induction conditions. We tested induction of *ynfEFGH* promoter trigger memory strains, which derive from an operon known to respond to low nitrate (through repression by phosphorylated NarL), and anaerobic (through FNR activation) conditions (19). When grown anaerobically in rich media, PAS816 (ynfE15), produced a memory response (13.3% +/-0.7% SE; n = 7). With added nitrate (20 mM), or in aerobic conditions, the response was zero (−O_2_/+nitrate: 0% +/-0% SE; n = 3; +O_2_/+nitrate: 0% +/-0% SE; n = 3; +O_2_: 0% +/-0% SE; n = 3) (Fig. 4G), consistent with previous literature reports of *ynfEFGH* promoter behavior (19). Additionally, no response was observed from PAS818 (ynfEWT) in any condition (n = 3), suggesting that it is less sensitive than PAS816 (ynfE15), consistent with our *in vivo* results. In addition to growth in rich media, strains were also cultured anaerobically in cecal contents. As in rich media with no inducer, only ynfE15 responded to growth in cecal media (3.7% +/-0.9% SE; n = 3) (Fig. 4G).

### Identification of disease-specific biosensors

To look for sensors responding differentially to disease, we compared the response of the Nissle 1917 library in healthy mice (n = 4; as previously displayed in Fig. 4D, 4E and S3, and Data Set S3) to a murine intestinal inflammation model (Fig. 5A, 5B and S4, and Data Set S3). After library gavage, SPF mice were provided water containing 4% w/v dextran sulfate sodium (DSS) *ad libitum* for five days, and HTMS analysis was performed on fecal samples throughout. Weight loss (Fig. S4A), colon length reduction at endpoint (Fig. S4B), and increased *E. coli* CFU counts (Fig. S4C) were all consistent with increasing inflammation throughout the experiment. Six strains registered memory-on at more timepoints in the DSS-treated group than in the control group (Fig. 5A and 5B). In particular, the ynfE17 trigger strain (PAS819) responded specifically in DSS-treated mice (control: no response; DSS-treated: 93% of timepoints with odds ratio ≥ 0.02 and p < 0.05) (Fig. 5A and 5B, and Data Set S3).

To validate ynfE17 (PAS819) response to inflammation, the strain was administered to SPF mice as a monoculture, after which a subset of the mice was provided water containing 4% DSS. Fecal samples were cultured for memory bacteria on indicator plates for 7 days following gavage. As above, body weights (Fig. S4D), post-dissection colon lengths (Fig. S4E) and CFU counts of HTMS bacteria (Fig. S4F) reflected increased inflammation in DSS-treated mice. Confirming the screen results, ynfE17 showed increased response in DSS-treated mice compared to untreated controls, with the greatest difference between groups at days 6 and 7 (at Day 6, +DSS: 24% +/-9% SE, n = 4; control: 5% +/-2% SE, n = 8) (Fig. 5C). The strong response of PAS819 in the absence of DSS in one of the control group mice indicates that *in vivo* conditions other than DSS treatment can activate the ynfE17 trigger. *In vitro* anaerobic growth both in rich media and in cecal media did not induce ynfE17 (0 +/-0% SE; n = 7; 0 +/-0% SE, n = 3, respectively), in contrast to ynfE15 (Fig. 4G), suggesting a lower nitrate threshold for ynfE17 activation and that individual bacteria may experience low nitrate conditions within the inflamed mouse gut.

## DISCUSSION

Here, we have expanded the use of a robust genetic memory circuit to assess the *in vivo* responses of multiple bacterial sensors in parallel. The original memory circuit (9) was altered to allow off-to-on transitions in the presence of constant induction, and to enable selection of memory-on strains from pooled libraries using spectinomycin. We developed a screening, sequencing and analysis pipeline to efficiently identify *in vivo*-responding trigger–RBS combinations. Tests conducted with both genome-wide and curated libraries containing hundreds of sensors demonstrated that our method is an effective, non-invasive way to identify new biosensors responding in the gut. We identified and validated biosensors responding to growth in the healthy mouse gut and preferentially in inflamed conditions. Together, these results demonstrate the power of tuning trigger sensitivity to physiological responses and for the HTMS to assess unique features of the mammalian gut environment *in vivo*.

One advantage of our method is its ability to discover sensors that could not be rationally designed based on existing knowledge, presenting an opportunity to apply the rapidly increasing but largely uncharacterized genetic diversity identified through microbiome sequencing. For instance, the two validated healthy-gut sensors from our *E. coli* MG1655 library (PAS813 and PAS814) are derived from operons with largely uncharacterized function and regulation. PAS814 is triggered by the promoter of the *ydjLKJIHG* operon, which putatively includes a kinase, a transporter protein, two dehydrogenases, an aldolase and an aldo-keto reductase. Only the activity of the aldo-keto reductase, YdjG, has been confirmed through reduction of methylglyoxal (20,21). Interestingly, a previous analysis of *E.coli* protein expression in germ-free mice showed that YdjG was expressed 3.5-fold higher in the cecum than *in vitro* (22). Another gene which has been studied in this operon, *ydjK*, may play a role in osmotolerance, showing a 50% increased growth rate in high-salt media when overexpressed in *E. coli* (23). It is not known whether methylglyoxal or osmotic stress can directly trigger transcription of the *ydjLKJIHG* operon. However, methylglyoxal occurs in many foods and is also produced by intestinal bacteria (24); it can also inhibit bacterial growth, suggesting a possible motivation for expression of *ydjLKJIHG* in the gut.

Promoters derived from three unique Nissle 1917 operons (*ynfEFGH, torCAD*, and *yeaR-yoaG*) showed robust memory response in the healthy mouse gut (Fig. 4D and 4E). The *ynfEFGH* operon encodes a DMSO reductase which has also been shown to reduce selenate (25,26). It is activated by FNR in anaerobic conditions and repressed by phosphorylated NarL in the presence of nitrate (19), which was further confirmed by our *in vitro* tests (Fig. 4G).

Tuning of trigger sensitivity (e.g., by RBS modulation) is important for generating responses to physiological conditions of interest and for successful application in synthetic engineered circuits. As we observed, RBS tuning can be used to increase the response of the ynfE promoter to as high as 100% in healthy mice (PAS816; Fig. 4F), and to adjust the response to distinguish the inflamed gut state (PAS819; Fig. 5C). Importantly, the sensors we identify can be used directly in downstream applications with the memory circuit. This provides an engineering advantage over any responsive genetic elements identified through analysis in their native context, for which incorporation into synthetic circuits would routinely require additional optimization.

Inflammation leads to an increase in nitric oxide produced by the host, which generates nitrate in the intestine (27). However, because the ynfE promoter is activated by a decrease in nitrate, our results suggest that DSS-induced inflammation may lead to lower levels of free nitrate available to *E. coli* NGF-1, possibly due to increased local competition for nitrate via respiration by NGF-1 and other *Enterobacteriaceae*. This idea is supported by our observation of increasingly higher NGF-1 bacterial loads in fecal samples of DSS-treated mice (Fig. S4C and S4F), suggesting a bloom of *E. coli*—and potentially other *Enterobacteriaceae* capable of nitrate respiration—correlated with increasing duration of DSS treatment. This is consistent with previous descriptions of *E. coli* experiencing a growth advantage due to anaerobic respiration of host-derived nitrate (27). Thus, we hypothesize that PAS819 responds in DSS-treated mice specifically through sensing inflammation-induced changes in its own microenvironment.

The HTMS enables both the recording of transient signals and the amplification of low-abundance signals through antibiotic selection. These features serve as a useful complement to other techniques, such as meta-transcriptomic or -proteomic studies which capture an instantaneous snapshot of total RNA or protein content. Screening of genome-wide libraries increases the chances of discovery of new, uncharacterized systems, while curating libraries to focus on a subset of sensors can allow greater fine-tuning and increase the chances of identifying a response for the condition of interest. Our use of the *E. coli* NGF-1 strain as a chassis allows robust colonization of the mouse gut without requiring antibiotic maintenance, leading to retention of high bacterial loads and high library complexity in fecal samples even after long periods in the gut.

An expanded arsenal of characterized sensors presents opportunities to construct more complex disease-responsive circuits. For instance, the combination of multiple redundant sensors would increase response accuracy under variable *in vivo* conditions, while complementary sensors may allow “fingerprinting” of different disease states. An exciting possibility is the use of more complex logic and signal processing within a single engineered strain, which may sense multiple inputs and produce anti-inflammatory, antimicrobial or other therapeutic proteins only when a precise set of conditions is satisfied (2). Sensors responding differentially based on localization within the intestine may create opportunities for more targeted drug delivery or for the construction of new safety and containment mechanisms—another important consideration in the deployment of engineered organisms.

The potential to engineer synthetic circuits into commensal gut bacteria is a promising new approach to the management of intestinal disease. Synthetic biology is just beginning to tap into the evolutionary breadth of capabilities found in natural systems, and our method represents a practical means for expanding the toolkit of useful sensors for *in vivo* application.

## MATERIALS AND METHODS

### Media, culture conditions

Unless otherwise mentioned, bacterial cultures were grown at 37°C in LB broth or agar (10 g/L NaCl, 5 g/L yeast extract, 10 g/L tryptone). Mixed liquid cultures (i.e., libraries) were grown in LBPS, which contains Peptone Special (Sigma) instead of tryptone. To quantify memory response on indicator plates, agar was supplemented with 60 µg/ml X-gal.

### High-throughput memory system (HTMS) construction

The spectinomycin resistance gene, *aadA*, was added downstream of the P_L_ promoter in the original memory element (9) by overlap extension PCR and genomically integrated by λ Red recombineering (28) into *E. coli* TB10 (29) between *mhpR* and *lacZ*, driving endogenous *lacZ* as a memory-on reporter. From TB10, transfer into streptomycin-resistant *E. coli* NGF-1 was by P1*vir* transduction.

### Biosensor strain and library construction

All triggers were cloned into pDR07 (Fig. S2), a Tn7 transposon insertion plasmid derived from pGRG36 (12). BsaI sites directly upstream of cIDN allow modular insertion of promoter–RBS sequences via Golden Gate assembly (13) (Fig. 2A). Assembled trigger plasmids were electroporated into PAS811. After recovery (90 min, 30°C in SOC medium) transformants were selected overnight (30°C in LB-ampicillin (100 µg/ml)). Cultures were then back-diluted 1:100 into LB-chloramphenicol (25 µg/ml) + 0.1% arabinose to induce transposase genes. After > 6 h at 30°C, temperature-sensitive pDR07 plasmids were cured from integrants by 2x 1:100 back-dilution into LB-chloramphenicol and > 6 h growth at 42°C. Plasmid loss was confirmed by restreaking on LB ampicillin agar.

For individual strains, post-cure cultures were plated on LB-chloramphenicol agar and attTn7 integrations confirmed by PCR and Sanger sequencing. For pooled libraries, library composition was confirmed by Illumina MiSeq sequencing of pooled PCRs of trigger regions.

### Assessment of memory state by LacZ assay

Cultures or fecal supernatants containing memory bacteria were plated on agar plates containing streptomycin (200 mg/ml), chloramphenicol (34 mg/ml), and X-gal (60 mg/ml). The percentage of memory-on colonies was assessed by counting blue (on) and white (off) colonies.

### *In vitro* induction

Overnight liquid cultures were back-diluted 1:100 into fresh media containing inducer, before 4 hours growth and plating on X-gal agar. For induction in cecal contents, contents of ceca from three female SPF C57BL/6J mice and suspended at 10% w/v in phospho-buffered saline (PBS). Suspensions were vortexed 90 sec and centrifuged for 3 min at 4300 rcf. The supernatant was recovered, supplemented with 200 µg/ml streptomycin and used for growth of HTMS bacteria.

For anaerobic inductions, pre-reduced anaerobic medium was used, and growth occurred in an anaerobic chamber (Coy Laboratory Products) under 7% H_2_, 20% CO_2_, 73% N_2_.

### *In vivo* induction of strains and libraries

The Harvard Medical School Animal Care and Use Committee approved all animal protocols. Experiments were conducted in female 7- to 14-week-old BALB/c (Charles River; MG1655 library) or C57BL/6J (Jackson; Nissle 1917 library) mice. Before experiments, all mice were confirmed to be absent of native streptomycin- and chloramphenicol-resistant flora. Food and water were removed ∼4 h before each gavage; water was replaced immediately, and food was replaced < 2 h following gavage.

One day prior to bacterial gavage, mice were provided streptomycin (20 mg in PBS) by oral gavage. The next day, overnight cultures of memory strains or libraries were washed once, then diluted 10-fold in PBS and administered by gavage (100µl; ∼10^7^ bacteria/mouse).

Gavage suspension and fecal samples were plated to track bacterial load and, for individual strains, to assess memory state. Libraries were processed according to the post-exposure processing protocol below. To plate fecal bacteria, samples were suspended at 100 mg/ml in PBS, vortexed 5 min, and centrifuged 20 min at 4 rcf to obtain fecal supernatant.

For inflammation experiments, water containing 4% DSS (36,000-50,000 M.W.; MP Biomedicals 160110) was substituted 2 h after bacteria administration. Mice were dissected at the end of the experiment to measure colon length.

### Post-exposure library processing

Fecal supernatant or *in vitro* culture was diluted 1:100 into LBPS-chloramphenicol (25 µg/ml) to achieve ∼10^6^ CFU/ml. Concurrently, an overnight culture of the positive normalization strain, PAS812 was back-diluted 1:100 into LBPS-chloramphenicol. Cultures were grown 4 h, or until OD_600_∼1. PAS812 OD_600_ was adjusted to match the library culture, then diluted 1:1000 into the library culture. The resulting mix was back-diluted 1:1000 into 50 ml LBPS-chloramphenicol, and immediately split into two 25 ml volumes. Spectinomycin (50 µg/ml) was added to one culture, and both were grown overnight before centrifugation to collect bacterial pellets, which were stored at −80°C.

### Library sequencing and odds ratio calculation

Genomic DNA was extracted from frozen cell pellets using a Qiagen DNEasy Blood & Tissue Kit. Using genomic DNA as a template, trigger regions from HTMS libraries were amplified by PCR and sheared with a Covaris M220 ultrasonicator to 200-600 bp fragments. Sheared products were prepared using the New England Biolabs NEBNext Ultra II Prep Kit and sequenced by Illumina MiSeq.

Raw reads were trimmed using Trimmomatic 0.36 (30) and aligned to a reference file (Data Sets S4 and S5 for MG1655 and Nissle 1917 libraries, respectively) using BWA mem 0.7.8 (31). The number of uniquely-mapped reads for each trigger was counted.

Odds ratio is expressed as (T_x-spect_/PNS_spect_)/(T_x_/PNS), where T_x_ and T_x-spect_ are the number of mapped reads for a particular trigger in the untreated and spectinomycin-treated culture, respectively, and PNS and PNS_spect_ are the number of mapped reads for the positive normalization strain (PAS812) in the untreated and spectinomycin-treated culture, respectively. Triggers with < 5 reads in the gavage suspension were discarded, unless they registered > 20 reads at any subsequent timepoint. For each pair of untreated and spectinomycin-treated cultures (from a single fecal sample), odds ratios were calculated for each trigger with ≥ 5 reads in the untreated culture. Statistical significance was assessed with a one-tailed Fisher’s exact test (*H*0: odds ratio = 0.02; *H*a: odds ratio > 0.02). The odds ratio calculation compares each trigger only with itself (between spectinomycin-treated and untreated cultures), normalizing any sequencing length bias between triggers. It also normalizes to the positive normalization strain (PAS812) in each sample, negating read depth disparities between samples.

## ACKNOWLEDGEMENTS

We thank Bryan Hsu, Andrew Verdegaal and Joseph Paulson for helpful discussions. A.D.N. was supported by the National Science Foundation Graduate Research Fellowship (DGE1144152 & DGE1745303). S.N.N. was supported by the Ruth L. Kirschstein Individual Postdoctoral NRSA F32 (F32 DK112640-03). D.T.R. was supported by a Human Frontier Science Program Long-Term Fellowship and the NHMRC/RG Menzies Early Career Fellowship from the Menzies Foundation through the Australian National Health and Medical Research Council. This work was supported by the Center for Microbiome Informatics and Therapeutics at MIT, the Defense Advanced Research Projects Agency (HR001115C0094), a Harvard Medical School Innovation Grant in the Basic and Social Sciences, and the Wyss Institute for Biologically Inspired Engineering.

## Author contributions

A.D.N., D.T.R., J.J.B, S.N.N., and P.A.S. designed experiments. A.D.N., S.N.N., J.J.B and D.T.R. constructed libraries. A.D.N., S.N.N., D.T.R, D.T., N.N., M.J.N., and M.C.I. performed in vitro characterization. A.D.N. and D.T.R. performed mouse experiments. A.D.N., J.J.B. and D.T.R. performed and analyzed library sequencing. A.D.N., D.T.R. and P.A.S. wrote the manuscript.

## SUPPLEMENTAL MATERIAL LEGENDS

**FIG S1** (A) Original memory circuit (9) in memory-off and memory-on states. (B) Response curves of P_tet_-*cIDN* trigger original memory (PAS809) & an aadA memory strain (PAS810), induced with varying concentrations of aTc. Memory response was assessed by plating on indicator plates after induction with aTc in liquid culture. Error bars represent SE of three biological replicates.

**FIG S2** Plasmid map of Tn7 transposon trigger integration plasmid, pDR07, containing BsaI sites upstream of *cIDN* gene for modular insertion of trigger promoter and RBS variants via Golden Gate assembly (see Fig. 2A). Tn7 left and right attachment sites are shown in pink.

**FIG S3** Heatmap for hits from Nissle 1917 library screen in healthy C57BL/6J mice (Fig. 4D and 4E), when screened in C57BL/6J mice treated with 4% w/v DSS in water. Blank spaces on heat map represent insufficient sequencing coverage. See Data Set S3 for full heat map.

**FIG S4** Body weights, CFU counts, and colon lengths of mice in Nissle 1917 library screen and ynfE17 (PAS819) biosensor strain validation experiments. All error bars represent SE. (A) Percentage of starting body weight for DSS-treated (n = 4) and healthy control (n = 4) mice used in Nissle 1917 library screens. (B) Colon lengths at dissection for DSS-treated (n = 4) and healthy control (n = 4) mice used in Nissle 1917 library screens. (C) CFU of HTMS bacteria in fecal samples of DSS-treated (n = 4) and healthy control (n = 4) mice used in Nissle 1917 library screens. (D) Percentage of starting body weight for DSS-treated (n = 4) and healthy control (n = 8) mice used in PAS819 validation experiment. (E) Colon lengths at dissection for DSS-treated (n = 4) and healthy control (n = 8) mice used in PAS819 validation experiment. (F) CFU of HTMS bacteria in fecal samples of DSS-treated (n = 4) and healthy control (n = 8) mice used in PAS819 validation experiment.

**DATA SET S1.** Screen of MG1655 library in 2 BALB/c mice over 24 hours (full heat map corresponding to Fig. 4A). See Fig. 4 for heat map legend.

**DATA SET S2.** Screen of MG1655 library in 3 BALB/c mice over seven days (full heat map corresponding to Fig. 4B). See Fig. 4 for heat map legend.

**DATA SET S3.** Screen of Nissle 1917 library in 8 C57BL/6J mice over five days, with and without DSS treatment (full heat map corresponding to Fig. 4D and 5A). See Fig. 4 or Fig. 5 for heat map legend.

**DATA SET S4.** Sequence reference file (fasta) for aligning MG1655 library reads.

**DATA SET S5.** Sequence reference file (fasta) for aligning Nissle 1917 library reads.

