## Supplementary Materials for "Synthetic gene circuits enable systems-level biosensor discovery at the host-microbe interface"

### Figure S1

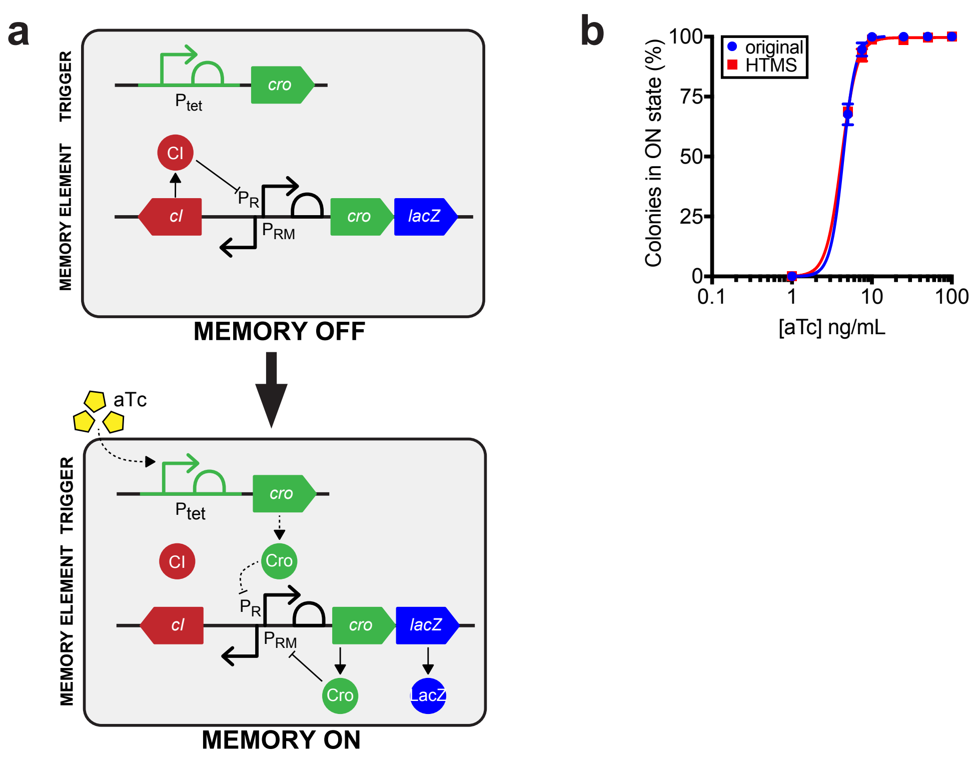
a
TRIGGER
MEMORY ELEMENT
Ptet
cro
CI
PR
PRM
cI
lacZ
MEMORY OFF
aTc
Cro
LacZ
MEMORY ON
b
Colonies in ON state (%)
original
HTMS
100
75
50
25
0
0.1
1
10
100
[aTc] ng/mL

### Figure S2

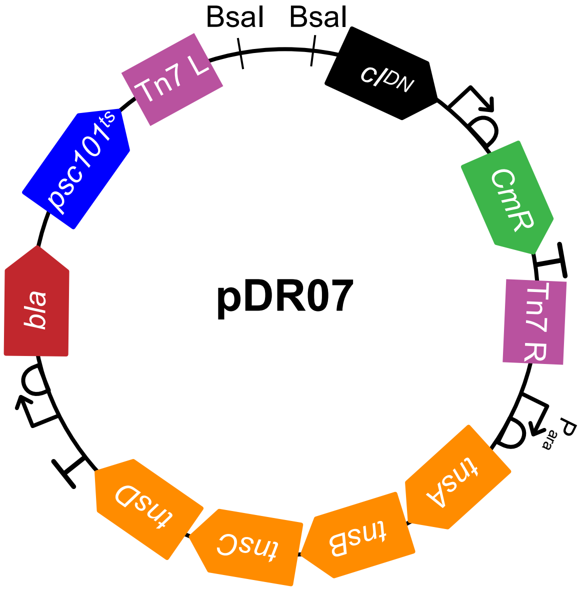
BsaI
BsaI
*cI*DN
CmR
Tn7 R
P ara
tnsA
tnsB
tnsC
tnsD
*bla*
psc101ts
Tn7 L
pDR07

### FIGURE S3

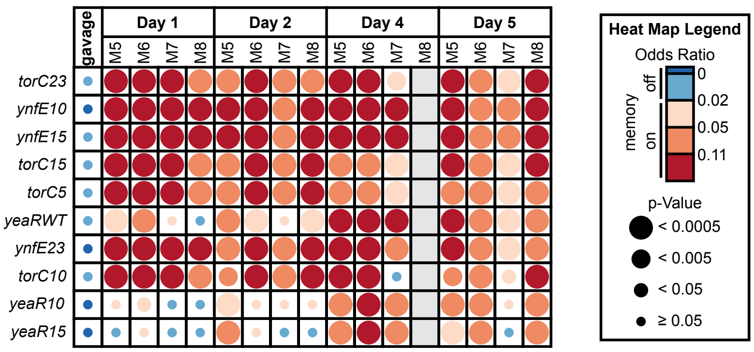
gavage
Day 1
Day 2
Day 4
Day 5
M5
M6
M7
M8
torC23
ynfE10
ynfE15
torC15
torC5
yeaRWT
ynfE23
torC10
yeaR10
yeaR15
Heat Map Legend
Odds Ratio
memory
off
on
0
0.02
0.05
0.11
p-Value
< 0.0005
< 0.005
< 0.05
≥ 0.05

### FIGURE S4

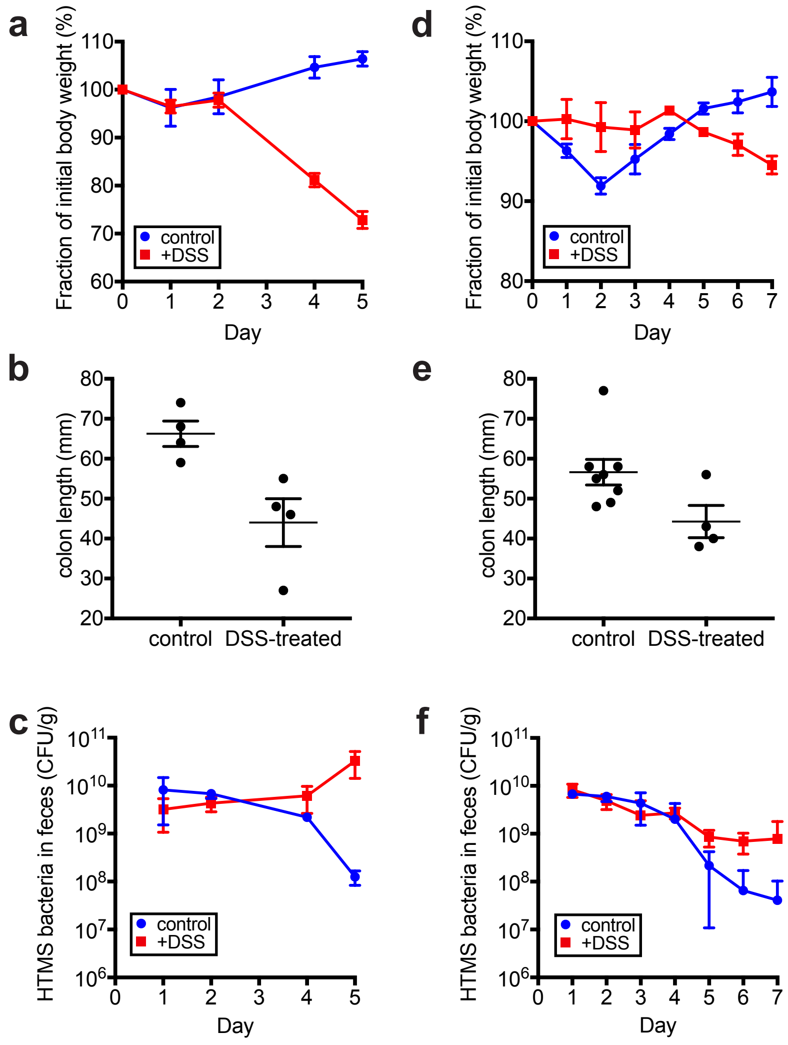
a
Fraction of initial body weight (%)
Day
control
+DSS
b
colon length (mm)
control
DSS-treated
c
HTMS bacteria in feces (CFU/g)
Day
control
+DSS
d
Fraction of initial body weight (%)
Day
control
+DSS
e
colon length (mm)
control
DSS-treated
f
HTMS bacteria in feces (CFU/g)
Day
control
+DSS

Data Set S1 - MG1655 library 24 h experiment
Odds Ratios

| | Promoter | gavage | M1-6h | M2-6h | M1-24h | M2-24h | #(OR>.02 & p<.05) / #non-N/A | AVG OR |
|---|---|---|---|---|---|---|---|---|
| 81 | ydjL | 0.0000 | 0.0020 | 0.5198 | 0.1035 | 2.1894 | 0.75 | 0.7037 |
| 60 | ydiL | 0.0001 | 0.0002 | 0.2171 | 0.1367 | 0.1918 | 0.75 | 0.1365 |
| 4 | gatY | 0.0003 | 0.0028 | N/A | 0.3054 | 0.1105 | 0.666667 | 0.1395 |
| 53 | yegT | 0.0000 | 0.0006 | 0.0000 | 2.3923 | 0.4726 | 0.5 | 0.7164 |
| 33 | ydcF | 0.0000 | 0.0052 | 0.0000 | 0.6003 | 0.2749 | 0.5 | 0.2201 |
| 65 | ubiG | 0.0002 | 0.0005 | 0.0002 | 0.0331 | 0.0360 | 0.5 | 0.0175 |
| 42 | argA_amiC | 0.0000 | 0.0002 | 0.0000 | 2.5280 | 0.0024 | 0.25 | 0.6326 |
| 43 | argD | 0.0000 | 0.0001 | 0.0000 | 2.3652 | 0.0000 | 0.25 | 0.5913 |
| 7 | gcvA | 0.0015 | 0.0039 | 0.0010 | 0.0309 | 0.6070 | 0.25 | 0.1607 |
| 73 | ydhR | 0.0089 | 0.0158 | 0.0576 | 0.1076 | 0.0000 | 0.25 | 0.0453 |
| 9 | pgsA | 0.0001 | 0.0002 | 0.0000 | 0.0054 | 0.0865 | 0.25 | 0.0230 |
| 27 | rof | 0.0000 | 0.0000 | N/A | N/A | 0.0423 | 0 | 0.0212 |
| 55 | cueR | 0.0000 | 0.0000 | 0.0000 | 0.0740 | 0.0000 | 0 | 0.0185 |
| 88 | rumA | 0.0000 | 0.0000 | N/A | 0.0444 | 0.0000 | 0 | 0.0148 |
| 18 | yebF | 0.0000 | 0.0000 | 0.0000 | 0.0371 | 0.0000 | 0 | 0.0093 |
| 86 | yhbQ | 0.0000 | 0.0000 | 0.0000 | 0.0371 | 0.0000 | 0 | 0.0093 |
| 80 | mutS | 0.0000 | 0.0000 | 0.0000 | 0.0222 | 0.0095 | 0 | 0.0079 |
| 24 | phnQ | 0.0000 | 0.0000 | 0.0000 | 0.0253 | 0.0048 | 0 | 0.0075 |
| 19 | ygjG | 0.0000 | 0.0000 | N/A | 0.0222 | 0.0000 | 0 | 0.0074 |
| 38 | ybgE | 0.0000 | 0.0000 | N/A | 0.0202 | 0.0009 | 0 | 0.0070 |
| 22 | cynR | 0.0000 | 0.0001 | 0.0000 | 0.0227 | 0.0003 | 0 | 0.0058 |
| 64 | ubiF | 0.0001 | 0.0001 | 0.0001 | 0.0125 | 0.0084 | 0 | 0.0053 |
| 69 | yjcO | 0.0000 | 0.0001 | 0.0000 | 0.0181 | 0.0004 | 0 | 0.0046 |
| 68 | hrpB | 0.0000 | 0.0002 | 0.0000 | 0.0171 | 0.0006 | 0 | 0.0045 |
| 47 | ynbA | 0.0000 | 0.0001 | 0.0000 | 0.0171 | 0.0000 | 0 | 0.0043 |
| 85 | yhbW | 0.0000 | 0.0000 | 0.0000 | 0.0171 | 0.0000 | 0 | 0.0043 |
| 45 | yjeB | 0.0014 | 0.0033 | N/A | N/A | N/A | 0 | 0.0033 |
| 91 | yaaU | 0.0000 | 0.0000 | 0.0000 | 0.0106 | 0.0000 | 0 | 0.0026 |
| 17 | flgH | 0.0000 | 0.0000 | N/A | 0.0076 | 0.0000 | 0 | 0.0025 |
| 51 | fumB | 0.0000 | 0.0000 | 0.0000 | 0.0097 | 0.0000 | 0 | 0.0024 |
| 67 | galS | 0.0004 | 0.0008 | 0.0000 | 0.0071 | 0.0000 | 0 | 0.0020 |
| 32 | mdtO | 0.0000 | 0.0000 | N/A | 0.0054 | 0.0000 | 0 | 0.0018 |
| 44 | yiiP | 0.0000 | 0.0000 | 0.0000 | 0.0071 | 0.0000 | 0 | 0.0018 |
| 5 | fadB | 0.0002 | 0.0006 | N/A | N/A | N/A | 0 | 0.0006 |
| 15 | yjdF | 0.0000 | 0.0000 | 0.0000 | 0.0000 | 0.0014 | 0 | 0.0004 |
| 62 | guaC | 0.0000 | 0.0002 | 0.0000 | 0.0000 | 0.0000 | 0 | 0.0000 |
| 54 | rihC | 0.0000 | 0.0001 | 0.0000 | N/A | 0.0000 | 0 | 0.0000 |
| 25 | msbB | 0.0000 | 0.0001 | 0.0000 | 0.0000 | 0.0000 | 0 | 0.0000 |
| 3 | b0671 | 0.0000 | 0.0001 | N/A | 0.0000 | 0.0000 | 0 | 0.0000 |
| 2 | eco | 0.0000 | 0.0000 | N/A | 0.0000 | N/A | 0 | 0.0000 |
| 6 | gcvT | 0.0000 | 0.0000 | N/A | 0.0000 | 0.0000 | 0 | 0.0000 |
| 8 | aniC | 0.0000 | 0.0000 | N/A | 0.0000 | 0.0000 | 0 | 0.0000 |
| 10 | dcuS | 0.0000 | 0.0000 | 0.0000 | N/A | 0.0000 | 0 | 0.0000 |
| 11 | ybbB | 0.0000 | 0.0000 | 0.0000 | 0.0000 | 0.0000 | 0 | 0.0000 |

Data Set S1 - MG1655 library 24 h experiment
Odds Ratios

| | | | | | | | | | |
|---|---|---|---|---|---|---|---|---|---|
| 12 | gst | 0.0000 | 0.0000 | N/A | 0.0000 | 0.0000 | | 0 | 0.0000 |
| 13 | pdhR | 0.0000 | 0.0000 | 0.0000 | 0.0000 | 0.0000 | | 0 | 0.0000 |
| 14 | yhhT | 0.0000 | 0.0000 | 0.0000 | 0.0000 | 0.0000 | | 0 | 0.0000 |
| 16 | yjdI | 0.0000 | 0.0000 | 0.0000 | N/A | 0.0000 | | 0 | 0.0000 |
| 20 | yijO | 0.0000 | 0.0000 | N/A | 0.0000 | 0.0000 | | 0 | 0.0000 |
| 21 | yfcC | 0.0000 | 0.0000 | 0.0000 | 0.0000 | 0.0000 | | 0 | 0.0000 |
| 23 | nudC | 0.0000 | 0.0000 | N/A | 0.0000 | 0.0000 | | 0 | 0.0000 |
| 26 | idnT | 0.0000 | 0.0000 | 0.0000 | N/A | N/A | | 0 | 0.0000 |
| 28 | yfbJ | 0.0000 | 0.0000 | 0.0000 | 0.0000 | 0.0000 | | 0 | 0.0000 |
| 29 | cysH | 0.0000 | 0.0000 | 0.0000 | 0.0000 | 0.0000 | | 0 | 0.0000 |
| 30 | cysQ | 0.0000 | 0.0000 | 0.0000 | N/A | 0.0000 | | 0 | 0.0000 |
| 31 | yihD | 0.0000 | 0.0000 | 0.0000 | N/A | 0.0000 | | 0 | 0.0000 |
| 35 | rrlG | 0.0000 | 0.0000 | 0.0000 | 0.0000 | 0.0000 | | 0 | 0.0000 |
| 36 | tktB | 0.0000 | 0.0000 | N/A | N/A | 0.0000 | | 0 | 0.0000 |
| 37 | yqgD | 0.0000 | 0.0000 | N/A | N/A | N/A | | 0 | 0.0000 |
| 39 | yeaU | 0.0000 | 0.0000 | 0.0000 | 0.0000 | 0.0000 | | 0 | 0.0000 |
| 40 | nupX | 0.0000 | 0.0000 | 0.0000 | 0.0000 | 0.0000 | | 0 | 0.0000 |
| 41 | argZ_argQ | 0.0000 | 0.0000 | N/A | N/A | N/A | | 0 | 0.0000 |
| 46 | grxB | 0.0000 | 0.0000 | N/A | 0.0000 | 0.0000 | | 0 | 0.0000 |
| 49 | yidK | 0.0000 | 0.0000 | N/A | 0.0000 | 0.0000 | | 0 | 0.0000 |
| 50 | yidL | 0.0000 | 0.0000 | N/A | 0.0000 | 0.0000 | | 0 | 0.0000 |
| 52 | ynaI | 0.0000 | 0.0000 | N/A | N/A | N/A | | 0 | 0.0000 |
| 56 | ybjK | 0.0000 | 0.0000 | 0.0000 | 0.0000 | 0.0000 | | 0 | 0.0000 |
| 58 | yqgA | 0.0000 | 0.0000 | N/A | 0.0000 | 0.0000 | | 0 | 0.0000 |
| 59 | ybiA | 0.0000 | 0.0000 | 0.0000 | 0.0000 | 0.0000 | | 0 | 0.0000 |
| 63 | yceF | 0.0000 | 0.0000 | 0.0000 | 0.0000 | 0.0000 | | 0 | 0.0000 |
| 66 | bglA | 0.0000 | 0.0000 | N/A | 0.0000 | 0.0000 | | 0 | 0.0000 |
| 70 | barA | 0.0000 | 0.0000 | N/A | 0.0000 | 0.0000 | | 0 | 0.0000 |
| 71 | yicC | 0.0000 | 0.0000 | 0.0000 | N/A | 0.0000 | | 0 | 0.0000 |
| 72 | mobA | 0.0000 | 0.0000 | 0.0000 | N/A | 0.0000 | | 0 | 0.0000 |
| 74 | mreC | 0.0000 | 0.0000 | 0.0000 | 0.0000 | 0.0000 | | 0 | 0.0000 |
| 76 | pabC | 0.0000 | 0.0000 | 0.0000 | 0.0000 | 0.0000 | | 0 | 0.0000 |
| 78 | yjaB | 0.0000 | 0.0000 | N/A | N/A | 0.0000 | | 0 | 0.0000 |
| 79 | fabF | 0.0000 | 0.0000 | N/A | 0.0000 | 0.0000 | | 0 | 0.0000 |
| 82 | yfjN | 0.0000 | 0.0000 | 0.0000 | N/A | 0.0000 | | 0 | 0.0000 |
| 83 | yfjR | 0.0000 | 0.0000 | N/A | 0.0000 | 0.0000 | | 0 | 0.0000 |
| 84 | metK | 0.0000 | 0.0000 | N/A | N/A | N/A | | 0 | 0.0000 |
| 87 | uidA | 0.0000 | 0.0000 | 0.0000 | 0.0000 | 0.0000 | | 0 | 0.0000 |
| 89 | rph | 0.0000 | 0.0000 | 0.0000 | N/A | 0.0000 | | 0 | 0.0000 |
| 92 | yjfY | 0.0000 | 0.0000 | 0.0000 | 0.0000 | 0.0000 | | 0 | 0.0000 |
| 34 | fucR | 0.0001 | 0.0000 | N/A | 0.0000 | 0.0000 | | 0 | 0.0000 |
| 48 | rluF | 0.0001 | 0.0000 | N/A | N/A | 0.0000 | | 0 | 0.0000 |
| 75 | ychH | 0.0912 | 0.2634 | N/A | 1.4328 | 1.6043 | | 1 | 1.1001 |
| 90 | fabR | N/A | 1.0000 | N/A | 1.0000 | 1.0000 | | 1 | 1.0000 |
| 1 | speE | N/A | N/A | N/A | N/A | N/A | NaN | | #DIV/0! |
| 57 | yahN | N/A | N/A | N/A | N/A | N/A | NaN | | #DIV/0! |
| 61 | dinG | N/A | N/A | N/A | N/A | N/A | NaN | | #DIV/0! |

| | | | | | | | | |
|---|---|---|---|---|---|---|---|---|
| 77 | edd | N/A | N/A | N/A | N/A | N/A | NaN | #DIV/0! |

### Data Set S1 - MG1655 library 24 h experiment
#### Odds Ratios & p-Values

| | Promoter | gavage | gavage_pValu | M1-6h | M1-6h_pValu | M2-6h | M2-6h_pValu | M1-24h | M1-24h_pVal | M2-24h | M2-24h_pValue |
|---|---|---|---|---|---|---|---|---|---|---|---|
| 1 | speE | N/A | 0.22767435 | N/A | 0.00021926 | N/A | 1 | N/A | 1 | N/A | 1 |
| 2 | eco | 0 | 1 | 0 | 1 | N/A | 1 | 0 | 1 | N/A | 1 |
| 3 | b0671 | 0 | 1 | 6.10E-05 | 1 | N/A | 1 | 0 | 1 | 0 | 1 |
| 4 | gatY | 0.00031195 | 1 | 0.00275519 | 1 | N/A | 1 | 0.30535751 | 2.80E-40 | 0.11047816 | 2.77E-10 |
| 5 | fadB | 0.0002215 | 0.99999999 | 0.00058426 | 1 | N/A | 1 | N/A | 1 | N/A | 1 |
| 6 | gcvT | 0 | 1 | 0 | 1 | N/A | 1 | 0 | 1 | 0 | 1 |
| 7 | gcvA | 0.00152604 | 1 | 0.00387389 | 1 | 0.00102993 | 0.99999825 | 0.03085246 | 0.24905002 | 0.60697363 | 1.22E-29 |
| 8 | aniC | 0 | 1 | 0 | 1 | N/A | 1 | 0 | 1 | 0 | 1 |
| 9 | pgsA | 0.00014612 | 1 | 0.00018776 | 1 | 0 | 1 | 0.00543663 | 0.9730744 | 0.08654078 | 2.32E-08 |
| 10 | dcuS | 0 | 1 | 0 | 1 | 0 | 1 | N/A | 0.35034773 | 0 | 1 |
| 11 | ybbB | 0 | 1 | 0 | 1 | 0 | 1 | 0 | 1 | 0 | 1 |
| 12 | gst | 0 | 1 | 0 | 1 | N/A | 1 | 0 | 1 | 0 | 1 |
| 13 | pdhR | 0 | 1 | 0 | 1 | 0 | 1 | 0 | 1 | 0 | 1 |
| 14 | yhhT | 0 | 1 | 0 | 1 | 0 | 1 | 0 | 1 | 0 | 1 |
| 15 | yjdF | 0 | 1 | 0 | 1 | 0 | 1 | 0 | 1 | 0.00143635 | 0.99998553 |
| 16 | yjdI | 0 | 1 | 0 | 1 | 0 | 1 | N/A | 0.22804067 | 0 | 1 |
| 17 | flgH | 0 | 1 | 0 | 1 | N/A | 1 | 0.00764368 | 0.92451716 | 0 | 1 |
| 18 | yebF | 0 | 1 | 0 | 1 | 0 | 1 | 0.03706919 | 0.45325307 | 0 | 1 |
| 19 | ygjG | 0 | 1 | 0 | 1 | N/A | 1 | 0.02222309 | 0.61269452 | 0 | 1 |
| 20 | yijO | 0 | 1 | 0 | 1 | N/A | 1 | 0 | 1 | 0 | 1 |
| 21 | yfcC | 0 | 1 | 0 | 1 | 0 | 1 | 0 | 1 | 0 | 1 |
| 22 | cynR | 0 | 1 | 0.00012023 | 1 | 0 | 1 | 0.02272149 | 0.29664926 | 0.00028551 | 1 |
| 23 | nudC | 0 | 1 | 0 | 1 | N/A | 1 | 0 | 1 | 0 | 1 |
| 24 | phnQ | 0 | 1 | 0 | 1 | 0 | 1 | 0.02525015 | 0.38140826 | 0.00482286 | 0.99942256 |
| 25 | msbB | 0 | 1 | 9.32E-05 | 1 | 0 | 1 | 0 | 1 | 0 | 1 |
| 26 | idnT | 0 | 1 | 0 | 1 | 0 | 1 | N/A | 0.01939485 | N/A | 1 |
| 27 | rof | 0 | 1 | 0 | 1 | N/A | 1 | N/A | 1 | 0.04234523 | 0.06978779 |
| 28 | yfbJ | 0 | 1 | 0 | 1 | 0 | 1 | 0 | 1 | 0 | 1 |
| 29 | cysH | 0 | 1 | 0 | 1 | 0 | 1 | 0 | 1 | 0 | 1 |
| 30 | cysQ | 0 | 1 | 0 | 1 | 0 | 1 | N/A | 1 | 0 | 1 |
| 31 | yihD | 0 | 1 | 0 | 1 | 0 | 1 | N/A | 0.22804067 | 0 | 1 |
| 32 | mdtO | 0 | 1 | 0 | 1 | N/A | 1 | 0.00543663 | 0.9730744 | 0 | 1 |
| 33 | ydcF | 0 | 1 | 0.00518026 | 0.99999988 | 0 | 1 | 0.60028605 | 1.11E-275 | 0.27489742 | 6.27E-21 |
| 34 | fucR | 6.19E-05 | 1 | 0 | 1 | N/A | 1 | 0 | 1 | 0 | 1 |
| 35 | rrlG | 0 | 1 | 0 | 1 | 0 | 1 | 0 | 1 | 0 | 1 |
| 36 | tktB | 0 | 1 | 0 | 1 | N/A | 1 | N/A | 1 | 0 | 1 |
| 37 | yqgD | 0 | 1 | 0 | 1 | N/A | 1 | N/A | 1 | N/A | 1 |
| 38 | ybgE | 0 | 1 | 0 | 1 | N/A | 1 | 0.02020376 | 0.64466971 | 0.00090725 | 0.99999996 |
| 39 | yeaU | 0 | 1 | 0 | 1 | 0 | 1 | 0 | 1 | 0 | 1 |
| 40 | nupX | 0 | 1 | 0 | 1 | 0 | 1 | 0 | 1 | 0 | 1 |
| 41 | argZ_argQ | 0 | 1 | 0 | 1 | N/A | 1 | N/A | 1 | N/A | 1 |
| 42 | argA_amiC | 0 | 1 | 0.0002276 | 1 | 0 | 1 | 2.52795944 | 9.34E-185 | 0.0023877 | 0.99927415 |
| 43 | argD | 0 | 1 | 7.71E-05 | 1 | 0 | 1 | 2.36515765 | 1.00E-265 | 0 | 1 |
| 44 | yiiP | 0 | 1 | 0 | 1 | 0 | 1 | 0.00714239 | 0.93643941 | 0 | 1 |
| 45 | yjeB | 0.00144872 | 0.9999697 | 0.00325755 | 0.99999043 | N/A | 0.34118907 | N/A | 0.00936346 | N/A | 0.01111834 |
| 46 | grxB | 0 | 1 | 0 | 1 | N/A | 1 | 0 | 1 | 0 | 1 |
| 47 | ynbA | 0 | 1 | 6.64E-05 | 1 | 0 | 1 | 0.01709627 | 0.68515063 | 0 | 1 |
| 48 | rluF | 9.65E-05 | 1 | 0 | 1 | N/A | 1 | N/A | 1 | 0 | 1 |
| 49 | yidK | 0 | 1 | 0 | 1 | N/A | 1 | 0 | 1 | 0 | 1 |
| 50 | yidL | 0 | 1 | 0 | 1 | N/A | 1 | 0 | 1 | 0 | 1 |
| 51 | fumB | 0 | 1 | 0 | 1 | 0 | 1 | 0.00966395 | 0.87354351 | 0 | 1 |
| 52 | ynaI | 0 | 1 | 0 | 1 | N/A | 1 | N/A | 1 | N/A | 1 |
| 53 | yegT | 0 | 1 | 0.00058013 | 1 | 0 | 1 | 2.39230495 | 7.52E-91 | 0.47257591 | 7.37E-24 |
| 54 | rihC | 0 | 1 | 8.92E-05 | 1 | 0 | 1 | N/A | 1 | 0 | 1 |
| 55 | cueR | 0 | 1 | 0 | 1 | 0 | 1 | 0.07404049 | 0.13713265 | 0 | 1 |
| 56 | ybjK | 0 | 1 | 0 | 1 | 0 | 1 | 0 | 1 | 0 | 1 |
| 57 | yahN | N/A | 1 | N/A | 1 | N/A | 0.01497678 | N/A | 1 | N/A | 5.15E-12 |
| 58 | yqgA | 0 | 1 | 0 | 1 | N/A | 1 | 0 | 1 | 0 | 1 |
| 59 | ybiA | 0 | 1 | 0 | 1 | 0 | 1 | 0 | 1 | 0 | 1 |
| 60 | ydiL | 0.00010062 | 1 | 0.00018091 | 1 | 0.2170857 | 0.00310964 | 0.13672717 | 1.09E-07 | 0.19183722 | 7.92E-15 |
| 61 | dinG | N/A | 1 | N/A | 1 | N/A | 1 | N/A | 1 | N/A | 1 |
| 62 | guaC | 0 | 1 | 0.00017545 | 1 | 0 | 1 | 0 | 1 | 0 | 1 |
| 63 | yceF | 0 | 1 | 0 | 1 | 0 | 1 | 0 | 1 | 0 | 1 |
| 64 | ubiF | 6.10E-05 | 1 | 7.90E-05 | 1 | 7.35E-05 | 1 | 0.01254851 | 0.99999765 | 0.00842189 | 0.99999946 |
| 65 | ubiG | 0.00023091 | 1 | 0.00050231 | 1 | 0.00019768 | 1 | 0.03307928 | 0.00020978 | 0.03603367 | 0.00449131 |
| 66 | bglA | 0 | 1 | 0 | 1 | N/A | 1 | 0 | 1 | 0 | 1 |
| 67 | galS | 0.00040577 | 1 | 0.0008455 | 1 | 0 | 1 | 0.00714239 | 0.93643941 | 0 | 1 |
| 68 | hrpB | 0 | 1 | 0.00018372 | 1 | 0 | 1 | 0.01709635 | 0.68495368 | 0.00059802 | 1 |
| 69 | yjcO | 0 | 1 | 8.76E-05 | 1 | 0 | 1 | 0.01807317 | 0.73502021 | 0.00042366 | 1 |
| 70 | barA | 0 | 1 | 0 | 1 | N/A | 1 | 0 | 1 | 0 | 1 |

Data Set S1 - MG1655 library 24 h experiment
Odds Ratios & p-Values

| | | | | | | | | | | | |
|---|---|---|---|---|---|---|---|---|---|---|---|
| 71 | yicC | 0 | 1 | 0 | 1 | 0 | 1 | N/A | 1 | 0 | 1 |
| 72 | mobA | 0 | 1 | 0 | 1 | 0 | 1 | N/A | 1 | 0 | 1 |
| 73 | ydhR | 0.00892603 | 0.9773347 | 0.01582939 | 0.79834133 | 0.05764369 | 0.08969248 | 0.10761049 | 6.20E-21 | 0 | 1 |
| 74 | mreC | 0 | 1 | 0 | 1 | 0 | 1 | 0 | 1 | 0 | 1 |
| 75 | ychH | 0.09120944 | 0.01363596 | 0.26338952 | 6.86E-08 | N/A | 1 | 1.43277382 | 2.47E-178 | 1.60428551 | 1.23E-40 |
| 76 | pabC | 0 | 1 | 0 | 1 | 0 | 1 | 0 | 1 | 0 | 1 |
| 77 | edd | N/A | 1 | N/A | 1 | N/A | 1 | N/A | 1 | N/A | 0.17447634 |
| 78 | yjaB | 0 | 1 | 0 | 1 | N/A | 1 | N/A | 1 | 0 | 1 |
| 79 | fabF | 0 | 1 | 0 | 1 | N/A | 1 | 0 | 1 | 0 | 1 |
| 80 | mutS | 0 | 1 | 0 | 1 | 0 | 1 | 0.02222309 | 0.61269452 | 0.00950564 | 0.87767014 |
| 81 | ydjL | 0 | 1 | 0.0020441 | 1 | 0.51980294 | 0.00024974 | 0.10351892 | 4.14E-14 | 2.18941939 | 8.31E-44 |
| 82 | yfjN | 0 | 1 | 0 | 1 | 0 | 1 | N/A | 1 | 0 | 1 |
| 83 | yfjR | 0 | 1 | 0 | 1 | N/A | 1 | 0 | 1 | 0 | 1 |
| 84 | metK | 0 | 1 | 0 | 1 | N/A | 1 | N/A | 1 | N/A | 1 |
| 85 | yhbW | 0 | 1 | 0 | 1 | 0 | 1 | 0.01709642 | 0.7009089 | 0 | 1 |
| 86 | yhbQ | 0 | 1 | 0 | 1 | 0 | 1 | 0.03706919 | 0.45325307 | 0 | 1 |
| 87 | uidA | 0 | 1 | 0 | 1 | 0 | 1 | 0 | 1 | 0 | 1 |
| 88 | rumA | 0 | 1 | 0 | 1 | N/A | 1 | 0.04444672 | 0.40402053 | 0 | 1 |
| 89 | rph | 0 | 1 | 0 | 1 | 0 | 1 | N/A | 1 | 0 | 1 |
| 90 | fabR | N/A | 9.76E-06 | 1 | 1.85E-11 | N/A | 0.00015059 | 1 | 0 | 1 | 7.17E-31 |
| 91 | yaaU | 0 | 1 | 0 | 1 | 0 | 1 | 0.01058425 | 0.84979785 | 0 | 1 |
| 92 | yjfY | 0 | 1 | 0 | 1 | 0 | 1 | 0 | 1 | 0 | 1 |

Data Set S1 - MG1655 library 24 h experiment
Aligned Reads

| | Promoter | gavage-ALL_S | gavage-ON_S1 | M1-24h-ALL_S | M1-24h-ON_S | M1-6h-ALL_S | M1-6h-ON_S3 | M2-24h-ALL_S | M2-24h-ON_S | M2-6h-ALL_S | M2-6h-ON_S6 |
|---|---|---|---|---|---|---|---|---|---|---|---|
| 1 | xapA | 0 | 0 | 0 | 0 | 0 | 0 | 0 | 0 | 0 | 0 |
| 2 | abrB | 0 | 0 | 0 | 0 | 0 | 0 | 0 | 0 | 0 | 0 |
| 3 | speF | 0 | 0 | 0 | 0 | 0 | 0 | 0 | 0 | 0 | 0 |
| 4 | xseA | 0 | 0 | 0 | 0 | 0 | 0 | 0 | 0 | 0 | 0 |
| 5 | xseB | 0 | 0 | 0 | 0 | 0 | 0 | 0 | 0 | 0 | 0 |
| 6 | speE | 1 | 54 | 0 | 0 | 1 | 80 | 0 | 0 | 0 | 0 |
| 7 | speB | 0 | 0 | 0 | 0 | 0 | 1 | 0 | 0 | 0 | 0 |
| 8 | speC | 3 | 0 | 0 | 0 | 10 | 0 | 0 | 0 | 0 | 0 |
| 9 | ybcL | 0 | 0 | 0 | 0 | 0 | 0 | 0 | 0 | 0 | 0 |
| 10 | ytfE | 1 | 1 | 1 | 3 | 6 | 1 | 0 | 3 | 0 | 1 |
| 11 | ybcK | 0 | 0 | 0 | 0 | 0 | 0 | 0 | 0 | 0 | 0 |
| 12 | frdA | 0 | 0 | 0 | 0 | 0 | 0 | 0 | 0 | 1 | 0 |
| 13 | ytfF | 0 | 0 | 0 | 0 | 0 | 0 | 0 | 0 | 0 | 0 |
| 14 | ybcY | 0 | 0 | 0 | 0 | 0 | 0 | 0 | 0 | 0 | 0 |
| 15 | pgpB | 0 | 0 | 0 | 0 | 0 | 0 | 0 | 0 | 0 | 0 |
| 16 | ybcV | 0 | 0 | 0 | 0 | 0 | 0 | 0 | 0 | 0 | 0 |
| 17 | ybcW | 0 | 0 | 0 | 0 | 0 | 0 | 0 | 0 | 0 | 0 |
| 18 | kbaZ | 0 | 0 | 0 | 0 | 0 | 0 | 0 | 0 | 0 | 0 |
| 19 | ucpA | 0 | 0 | 0 | 0 | 0 | 0 | 0 | 0 | 0 | 0 |
| 20 | cmk | 0 | 0 | 0 | 0 | 0 | 0 | 0 | 0 | 0 | 0 |
| 21 | cmr | 0 | 0 | 0 | 0 | 0 | 0 | 0 | 0 | 0 | 0 |
| 22 | upp | 0 | 0 | 0 | 0 | 0 | 0 | 0 | 0 | 0 | 0 |
| 23 | ydgR | 0 | 0 | 0 | 0 | 0 | 0 | 0 | 0 | 0 | 0 |
| 24 | ydgK | 0 | 0 | 0 | 0 | 0 | 0 | 0 | 0 | 0 | 0 |
| 25 | ydgH | 0 | 0 | 0 | 0 | 0 | 0 | 0 | 0 | 0 | 0 |
| 26 | ydgI | 0 | 0 | 0 | 0 | 0 | 0 | 0 | 0 | 0 | 0 |
| 27 | ydgL | 0 | 0 | 0 | 0 | 0 | 0 | 0 | 0 | 0 | 0 |
| 28 | tolC | 0 | 0 | 0 | 0 | 0 | 0 | 0 | 0 | 0 | 0 |
| 29 | tolB | 0 | 0 | 0 | 0 | 0 | 0 | 0 | 0 | 0 | 0 |
| 30 | tolA | 0 | 0 | 0 | 0 | 0 | 0 | 0 | 0 | 0 | 0 |
| 31 | ydgA | 0 | 0 | 0 | 0 | 0 | 0 | 0 | 0 | 0 | 0 |
| 32 | ydgD | 0 | 0 | 0 | 0 | 0 | 0 | 0 | 0 | 0 | 0 |
| 33 | entC | 0 | 0 | 0 | 0 | 0 | 0 | 0 | 0 | 0 | 0 |
| 34 | entD | 0 | 0 | 0 | 0 | 0 | 1 | 0 | 0 | 0 | 0 |
| 35 | entF | 0 | 0 | 0 | 0 | 0 | 0 | 0 | 0 | 0 | 0 |
| 36 | eco | 6 | 0 | 6 | 0 | 20 | 0 | 4 | 0 | 0 | 0 |
| 37 | entS | 0 | 0 | 0 | 0 | 0 | 0 | 0 | 0 | 0 | 0 |
| 38 | yacG | 0 | 0 | 0 | 0 | 0 | 0 | 0 | 0 | 0 | 0 |
| 39 | fecA | 0 | 0 | 0 | 0 | 0 | 0 | 0 | 0 | 0 | 0 |
| 40 | yacH | 0 | 0 | 0 | 0 | 0 | 0 | 0 | 0 | 0 | 0 |
| 41 | yhbO | 0 | 0 | 0 | 0 | 0 | 0 | 0 | 0 | 0 | 0 |
| 42 | fecI | 0 | 0 | 0 | 0 | 0 | 0 | 0 | 0 | 0 | 0 |
| 43 | rcsA | 0 | 0 | 0 | 0 | 0 | 0 | 0 | 0 | 0 | 0 |
| 44 | b2394 | 0 | 0 | 0 | 0 | 0 | 0 | 0 | 0 | 0 | 0 |
| 45 | rcsC | 0 | 0 | 0 | 0 | 0 | 0 | 0 | 0 | 0 | 0 |
| 46 | rcsF | 0 | 0 | 0 | 0 | 0 | 0 | 0 | 0 | 0 | 0 |
| 47 | uspA | 0 | 0 | 0 | 1 | 0 | 0 | 0 | 0 | 0 | 0 |
| 48 | uspB | 0 | 0 | 0 | 0 | 0 | 0 | 0 | 0 | 0 | 0 |
| 49 | uspE | 0 | 0 | 0 | 0 | 0 | 0 | 0 | 0 | 0 | 0 |
| 50 | uspF | 0 | 0 | 0 | 0 | 0 | 0 | 0 | 0 | 0 | 0 |
| 51 | uspG | 0 | 0 | 0 | 0 | 0 | 0 | 0 | 0 | 0 | 0 |
| 52 | gltX | 0 | 0 | 1 | 0 | 0 | 0 | 0 | 0 | 0 | 0 |
| 53 | yifM2 | 0 | 0 | 0 | 0 | 0 | 0 | 1 | 0 | 0 | 0 |
| 54 | gltJ | 0 | 0 | 0 | 0 | 0 | 0 | 1 | 0 | 0 | 0 |
| 55 | cspI | 0 | 0 | 0 | 0 | 0 | 0 | 0 | 0 | 0 | 0 |
| 56 | gltA | 0 | 0 | 0 | 0 | 0 | 0 | 0 | 0 | 0 | 0 |
| 57 | cspG | 0 | 0 | 0 | 0 | 0 | 0 | 0 | 0 | 0 | 0 |
| 58 | cspD | 0 | 0 | 0 | 0 | 0 | 0 | 0 | 0 | 0 | 0 |
| 59 | gltB | 0 | 0 | 0 | 0 | 0 | 0 | 0 | 0 | 0 | 0 |
| 60 | cspB | 0 | 0 | 0 | 0 | 0 | 0 | 0 | 0 | 0 | 0 |
| 61 | cspA | 0 | 0 | 0 | 0 | 0 | 0 | 1 | 0 | 0 | 0 |
| 62 | yhgN | 0 | 0 | 0 | 0 | 0 | 0 | 0 | 0 | 0 | 0 |
| 63 | yfeC | 0 | 0 | 0 | 0 | 0 | 0 | 0 | 0 | 0 | 0 |
| 64 | maeB | 0 | 0 | 0 | 0 | 0 | 0 | 0 | 0 | 0 | 0 |
| 65 | yhgF | 0 | 0 | 0 | 0 | 0 | 0 | 0 | 0 | 0 | 0 |

| | | | | | | | | | | | |
|---|---|---|---|---|---|---|---|---|---|---|---|
| 66 | yfeK | 0 | 0 | 0 | 0 | 0 | 0 | 0 | 0 | 0 | 0 |
| 67 | yhgE | 0 | 0 | 0 | 0 | 0 | 0 | 0 | 0 | 0 | 0 |
| 68 | yfeN | 0 | 0 | 0 | 0 | 0 | 0 | 0 | 0 | 0 | 0 |
| 69 | yhgA | 0 | 0 | 0 | 0 | 0 | 0 | 0 | 1 | 0 | 0 |
| 70 | yfeY | 0 | 0 | 0 | 0 | 0 | 0 | 0 | 0 | 0 | 0 |
| 71 | b0671 | 78 | 0 | 43 | 0 | 128 | 1 | 40 | 0 | 1 | 0 |
| 72 | b0298 | 0 | 0 | 0 | 0 | 0 | 0 | 0 | 0 | 0 | 0 |
| 73 | yjiX | 0 | 0 | 0 | 0 | 0 | 0 | 0 | 0 | 0 | 0 |
| 74 | rssA | 0 | 0 | 0 | 0 | 0 | 0 | 0 | 0 | 0 | 0 |
| 75 | rssB | 0 | 0 | 0 | 0 | 0 | 0 | 0 | 0 | 0 | 0 |
| 76 | ddlA | 0 | 0 | 0 | 0 | 0 | 0 | 0 | 0 | 0 | 0 |
| 77 | gatY | 155 | 34 | 48 | 66 | 270 | 146 | 108 | 143 | 0 | 0 |
| 78 | yjiW | 0 | 0 | 0 | 0 | 0 | 0 | 0 | 0 | 0 | 0 |
| 79 | ytjB | 0 | 0 | 0 | 0 | 0 | 0 | 0 | 0 | 0 | 0 |
| 80 | ygiF | 0 | 0 | 0 | 0 | 0 | 0 | 0 | 0 | 0 | 0 |
| 81 | ygiD | 0 | 0 | 0 | 0 | 0 | 0 | 0 | 0 | 0 | 0 |
| 82 | ygiH | 0 | 0 | 0 | 0 | 0 | 0 | 0 | 0 | 0 | 0 |
| 83 | eno | 0 | 0 | 0 | 0 | 0 | 0 | 0 | 0 | 0 | 0 |
| 84 | ygiS | 0 | 0 | 0 | 0 | 0 | 0 | 0 | 0 | 0 | 0 |
| 85 | ygiR | 0 | 0 | 0 | 0 | 0 | 0 | 0 | 0 | 0 | 0 |
| 86 | ygiP | 0 | 0 | 0 | 0 | 0 | 0 | 0 | 0 | 0 | 0 |
| 87 | ygiW | 0 | 0 | 0 | 0 | 0 | 0 | 0 | 0 | 0 | 0 |
| 88 | ygiU | 0 | 0 | 0 | 0 | 0 | 0 | 0 | 0 | 0 | 0 |
| 89 | ygiZ | 0 | 0 | 0 | 0 | 0 | 0 | 0 | 0 | 0 | 0 |
| 90 | smtA | 0 | 0 | 0 | 0 | 0 | 0 | 0 | 0 | 0 | 0 |
| 91 | b0582 | 0 | 0 | 0 | 0 | 0 | 0 | 0 | 0 | 0 | 0 |
| 92 | amiA | 0 | 0 | 0 | 0 | 0 | 0 | 0 | 0 | 0 | 0 |
| 93 | yhiD | 0 | 0 | 0 | 0 | 0 | 0 | 0 | 0 | 0 | 0 |
| 94 | yjiA | 0 | 0 | 0 | 0 | 0 | 0 | 0 | 0 | 0 | 0 |
| 95 | hslJ | 0 | 0 | 0 | 0 | 0 | 0 | 0 | 0 | 0 | 0 |
| 96 | yhiI | 0 | 0 | 0 | 0 | 0 | 0 | 0 | 0 | 0 | 0 |
| 97 | yhiJ | 0 | 0 | 0 | 0 | 0 | 0 | 0 | 0 | 0 | 0 |
| 98 | adiA | 0 | 0 | 0 | 0 | 0 | 0 | 0 | 0 | 0 | 0 |
| 99 | yhiP | 0 | 0 | 1 | 0 | 0 | 0 | 0 | 0 | 0 | 0 |
| 100 | hslV | 0 | 0 | 0 | 0 | 0 | 0 | 0 | 0 | 0 | 0 |
| 101 | arcB | 0 | 0 | 0 | 0 | 0 | 0 | 0 | 0 | 0 | 0 |
| 102 | yjeN | 0 | 0 | 0 | 0 | 0 | 0 | 0 | 0 | 0 | 0 |
| 103 | yjeK | 0 | 0 | 0 | 0 | 0 | 0 | 0 | 0 | 0 | 0 |
| 104 | yjeI | 0 | 0 | 0 | 0 | 0 | 0 | 0 | 0 | 0 | 0 |
| 105 | fadB | 7 | 1 | 1 | 0 | 28 | 3 | 3 | 0 | 0 | 0 |
| 106 | fdrA | 0 | 0 | 1 | 0 | 0 | 0 | 0 | 0 | 0 | 0 |
| 107 | fadE | 0 | 0 | 0 | 0 | 0 | 0 | 0 | 0 | 0 | 0 |
| 108 | ypeA | 0 | 0 | 0 | 0 | 0 | 0 | 0 | 0 | 0 | 0 |
| 109 | creA | 0 | 0 | 0 | 0 | 0 | 0 | 0 | 0 | 0 | 0 |
| 110 | creB | 0 | 0 | 0 | 0 | 0 | 0 | 0 | 0 | 0 | 0 |
| 111 | inaA | 0 | 0 | 0 | 0 | 0 | 0 | 0 | 0 | 0 | 0 |
| 112 | ompT | 0 | 1 | 0 | 0 | 0 | 0 | 0 | 0 | 0 | 0 |
| 113 | ompW | 0 | 0 | 0 | 0 | 0 | 0 | 0 | 0 | 0 | 0 |
| 114 | ompR | 0 | 0 | 0 | 0 | 0 | 0 | 0 | 0 | 0 | 0 |
| 115 | fiu | 0 | 0 | 0 | 0 | 0 | 0 | 0 | 0 | 0 | 0 |
| 116 | cld | 0 | 0 | 0 | 0 | 0 | 0 | 0 | 0 | 0 | 0 |
| 117 | ompX | 0 | 0 | 0 | 0 | 0 | 0 | 0 | 0 | 0 | 0 |
| 118 | yggE | 0 | 0 | 0 | 0 | 0 | 0 | 0 | 0 | 0 | 0 |
| 119 | ompG | 0 | 0 | 0 | 0 | 0 | 0 | 0 | 0 | 0 | 0 |
| 120 | ompA | 0 | 0 | 0 | 0 | 0 | 0 | 0 | 0 | 0 | 0 |
| 121 | ompC | 0 | 0 | 0 | 0 | 0 | 0 | 0 | 0 | 0 | 0 |
| 122 | ompN | 0 | 0 | 0 | 0 | 0 | 0 | 0 | 0 | 0 | 0 |
| 123 | htrL | 0 | 0 | 0 | 0 | 0 | 0 | 0 | 0 | 0 | 0 |
| 124 | cls | 0 | 0 | 0 | 0 | 0 | 0 | 0 | 0 | 0 | 0 |
| 125 | gcvP | 0 | 0 | 0 | 0 | 0 | 0 | 0 | 0 | 0 | 0 |
| 126 | gcvT | 26 | 0 | 34 | 0 | 54 | 0 | 66 | 0 | 0 | 0 |
| 127 | tnaC | 0 | 0 | 0 | 0 | 0 | 0 | 0 | 0 | 0 | 0 |
| 128 | srlR | 0 | 0 | 0 | 0 | 0 | 0 | 0 | 0 | 0 | 0 |
| 129 | gcvA | 264 | 301 | 36 | 5 | 446 | 341 | 416 | 3039 | 36 | 2 |
| 130 | ftsN | 0 | 0 | 0 | 0 | 0 | 0 | 0 | 0 | 0 | 0 |
| 131 | ftsK | 0 | 0 | 0 | 0 | 0 | 0 | 0 | 0 | 0 | 0 |

| | | | | | | | | | | | |
|---|---|---|---|---|---|---|---|---|---|---|---|
| 132 | aniC | 29 | 0 | 6 | 0 | 47 | 0 | 32 | 0 | 0 | 0 |
| 133 | ftsH | 0 | 0 | 0 | 0 | 0 | 0 | 0 | 0 | 0 | 0 |
| 134 | yieE | 0 | 1 | 0 | 0 | 0 | 0 | 0 | 0 | 0 | 0 |
| 135 | ftsQ | 0 | 1 | 0 | 0 | 0 | 0 | 0 | 0 | 0 | 0 |
| 136 | ftsZ | 0 | 0 | 0 | 0 | 0 | 0 | 0 | 0 | 0 | 0 |
| 137 | ftsY | 0 | 0 | 0 | 0 | 0 | 0 | 0 | 0 | 0 | 0 |
| 138 | hemA | 0 | 0 | 0 | 0 | 0 | 0 | 0 | 0 | 0 | 0 |
| 139 | yrfF | 0 | 0 | 0 | 0 | 0 | 0 | 0 | 0 | 0 | 0 |
| 140 | pgsA | 166 | 14 | 41 | 1 | 336 | 12 | 136 | 141 | 84 | 0 |
| 141 | fepB | 0 | 0 | 0 | 0 | 0 | 0 | 0 | 0 | 0 | 0 |
| 142 | fepA | 0 | 0 | 0 | 0 | 0 | 0 | 0 | 0 | 0 | 0 |
| 143 | fepD | 0 | 0 | 0 | 0 | 0 | 0 | 0 | 0 | 0 | 0 |
| 144 | yiiR | 0 | 0 | 0 | 0 | 0 | 0 | 0 | 0 | 0 | 0 |
| 145 | ligA | 0 | 0 | 0 | 0 | 0 | 0 | 0 | 0 | 0 | 0 |
| 146 | ligB | 0 | 0 | 0 | 0 | 0 | 0 | 0 | 0 | 0 | 0 |
| 147 | aidB | 0 | 0 | 0 | 0 | 0 | 0 | 0 | 0 | 0 | 0 |
| 148 | ligT | 0 | 0 | 0 | 0 | 0 | 0 | 0 | 0 | 0 | 0 |
| 149 | smf | 0 | 0 | 0 | 0 | 0 | 0 | 0 | 0 | 0 | 0 |
| 150 | prfB | 0 | 0 | 0 | 0 | 0 | 0 | 0 | 0 | 0 | 0 |
| 151 | prfC | 0 | 0 | 0 | 0 | 0 | 0 | 0 | 0 | 0 | 0 |
| 152 | dcuA | 0 | 0 | 0 | 0 | 0 | 0 | 0 | 0 | 0 | 0 |
| 153 | sodB | 0 | 0 | 0 | 0 | 0 | 0 | 0 | 0 | 0 | 0 |
| 154 | sodC | 0 | 0 | 0 | 0 | 0 | 0 | 0 | 0 | 0 | 0 |
| 155 | sodA | 0 | 0 | 0 | 0 | 0 | 0 | 0 | 0 | 0 | 0 |
| 156 | erfK | 0 | 0 | 0 | 0 | 0 | 0 | 0 | 0 | 0 | 1 |
| 157 | dcuS | 5 | 0 | 4 | 1 | 12 | 0 | 7 | 0 | 101 | 0 |
| 158 | ybbL | 0 | 0 | 0 | 0 | 0 | 0 | 0 | 0 | 0 | 0 |
| 159 | oxyR | 0 | 0 | 0 | 0 | 0 | 0 | 0 | 0 | 0 | 0 |
| 160 | umuD | 0 | 0 | 0 | 0 | 0 | 0 | 0 | 0 | 0 | 0 |
| 161 | ybbC | 0 | 0 | 0 | 0 | 0 | 0 | 0 | 0 | 0 | 0 |
| 162 | ybbB | 41 | 0 | 31 | 0 | 101 | 0 | 25 | 0 | 49 | 0 |
| 163 | yphG | 0 | 0 | 1 | 0 | 0 | 0 | 0 | 0 | 0 | 0 |
| 164 | ybdG | 0 | 0 | 0 | 0 | 0 | 0 | 0 | 0 | 0 | 0 |
| 165 | fixC | 0 | 0 | 0 | 0 | 0 | 0 | 0 | 0 | 0 | 0 |
| 166 | glyQ | 1 | 1 | 0 | 0 | 2 | 0 | 5 | 1 | 0 | 0 |
| 167 | glyU | 0 | 0 | 0 | 0 | 0 | 0 | 0 | 0 | 0 | 0 |
| 168 | aqpZ | 0 | 0 | 0 | 0 | 0 | 0 | 0 | 0 | 0 | 0 |
| 169 | glyA | 0 | 0 | 0 | 0 | 0 | 1 | 0 | 0 | 0 | 0 |
| 170 | trkA | 0 | 0 | 0 | 0 | 0 | 0 | 0 | 0 | 0 | 0 |
| 171 | ynfM | 0 | 0 | 0 | 0 | 0 | 0 | 0 | 0 | 0 | 0 |
| 172 | ynfL | 0 | 0 | 0 | 0 | 0 | 0 | 0 | 0 | 0 | 0 |
| 173 | ynfK | 0 | 0 | 0 | 0 | 0 | 0 | 0 | 0 | 0 | 0 |
| 174 | ynfE | 0 | 0 | 0 | 0 | 0 | 0 | 0 | 0 | 0 | 0 |
| 175 | ynfC | 0 | 0 | 0 | 0 | 0 | 0 | 1 | 0 | 0 | 0 |
| 176 | ynfB | 0 | 0 | 0 | 0 | 0 | 0 | 0 | 0 | 0 | 0 |
| 177 | ynfA | 0 | 0 | 0 | 0 | 0 | 0 | 0 | 0 | 0 | 0 |
| 178 | treA | 0 | 0 | 0 | 0 | 0 | 0 | 0 | 0 | 0 | 0 |
| 179 | qseB | 0 | 0 | 0 | 0 | 0 | 0 | 0 | 0 | 0 | 0 |
| 180 | treB | 0 | 1 | 0 | 0 | 0 | 0 | 0 | 0 | 0 | 0 |
| 181 | treF | 0 | 1 | 0 | 0 | 0 | 0 | 0 | 0 | 0 | 0 |
| 182 | pgmI | 0 | 1 | 0 | 0 | 0 | 0 | 0 | 2 | 0 | 0 |
| 183 | gst | 55 | 0 | 15 | 0 | 81 | 0 | 28 | 0 | 0 | 0 |
| 184 | ydfE | 0 | 0 | 0 | 0 | 0 | 0 | 0 | 0 | 0 | 0 |
| 185 | ydfA | 0 | 0 | 0 | 0 | 0 | 0 | 0 | 0 | 0 | 0 |
| 186 | chbA | 0 | 0 | 0 | 0 | 0 | 0 | 0 | 0 | 0 | 0 |
| 187 | ydfO | 0 | 0 | 0 | 0 | 0 | 0 | 0 | 0 | 0 | 0 |
| 188 | ydfI | 0 | 0 | 0 | 0 | 0 | 0 | 0 | 0 | 0 | 0 |
| 189 | ydfH | 0 | 1 | 0 | 0 | 0 | 0 | 0 | 0 | 0 | 0 |
| 190 | ydfV | 0 | 0 | 0 | 0 | 0 | 0 | 0 | 0 | 0 | 0 |
| 191 | ydfZ | 0 | 0 | 0 | 0 | 0 | 0 | 0 | 0 | 0 | 0 |
| 192 | serW | 0 | 0 | 0 | 0 | 0 | 0 | 0 | 0 | 0 | 0 |
| 193 | pdhR | 6 | 0 | 6 | 0 | 12 | 0 | 6 | 0 | 30 | 0 |
| 194 | ydgC | 0 | 0 | 0 | 0 | 0 | 0 | 0 | 0 | 0 | 0 |
| 195 | pqiA | 0 | 0 | 0 | 0 | 0 | 0 | 0 | 0 | 0 | 0 |
| 196 | yhhF | 0 | 0 | 0 | 0 | 0 | 0 | 0 | 0 | 0 | 0 |
| 197 | flgB | 0 | 0 | 0 | 0 | 0 | 0 | 0 | 0 | 0 | 0 |

| | | | | | | | | | | | |
|---|---|---|---|---|---|---|---|---|---|---|---|
| 198 | yhhN | 0 | 0 | 0 | 0 | 0 | 0 | 0 | 0 | 0 | 0 |
| 199 | yhhM | 0 | 0 | 0 | 0 | 0 | 0 | 0 | 0 | 0 | 0 |
| 200 | minC | 0 | 0 | 0 | 0 | 0 | 0 | 0 | 0 | 0 | 0 |
| 201 | flgA | 0 | 0 | 0 | 0 | 0 | 0 | 0 | 0 | 0 | 0 |
| 202 | yhhW | 0 | 0 | 0 | 0 | 0 | 0 | 0 | 0 | 0 | 0 |
| 203 | yhhT | 47 | 0 | 13 | 0 | 86 | 0 | 48 | 0 | 133 | 0 |
| 204 | adhE | 0 | 0 | 0 | 0 | 0 | 0 | 1 | 0 | 0 | 0 |
| 205 | yjdF | 52 | 0 | 11 | 0 | 73 | 0 | 59 | 1 | 175 | 0 |
| 206 | yjdI | 17 | 0 | 2 | 1 | 25 | 0 | 17 | 0 | 177 | 0 |
| 207 | slyB | 0 | 0 | 0 | 0 | 0 | 0 | 0 | 0 | 0 | 0 |
| 208 | yjdK | 0 | 0 | 0 | 0 | 1 | 0 | 0 | 0 | 0 | 0 |
| 209 | yhhZ | 0 | 0 | 0 | 0 | 0 | 0 | 0 | 0 | 0 | 0 |
| 210 | yhhY | 0 | 0 | 0 | 0 | 0 | 0 | 0 | 0 | 0 | 0 |
| 211 | yfdL | 0 | 0 | 0 | 0 | 0 | 0 | 0 | 0 | 0 | 0 |
| 212 | yhfK | 0 | 0 | 0 | 0 | 0 | 0 | 0 | 0 | 0 | 0 |
| 213 | yhfL | 0 | 0 | 0 | 0 | 0 | 0 | 0 | 0 | 0 | 0 |
| 214 | yhfA | 0 | 0 | 0 | 0 | 0 | 0 | 0 | 0 | 0 | 0 |
| 215 | yfdG | 0 | 0 | 0 | 0 | 0 | 0 | 0 | 0 | 0 | 0 |
| 216 | sgcC | 0 | 0 | 0 | 0 | 0 | 0 | 0 | 0 | 0 | 0 |
| 217 | yfdC | 0 | 0 | 0 | 0 | 0 | 0 | 0 | 0 | 0 | 0 |
| 218 | yhfX | 0 | 0 | 0 | 0 | 0 | 0 | 0 | 0 | 0 | 0 |
| 219 | yhfZ | 0 | 0 | 1 | 0 | 0 | 0 | 0 | 0 | 0 | 0 |
| 220 | yfdZ | 0 | 0 | 0 | 0 | 0 | 0 | 0 | 0 | 0 | 0 |
| 221 | yfdU | 0 | 0 | 0 | 0 | 0 | 0 | 0 | 0 | 0 | 0 |
| 222 | yhfP | 0 | 0 | 0 | 0 | 0 | 0 | 0 | 0 | 0 | 0 |
| 223 | yfdV | 0 | 0 | 0 | 0 | 0 | 0 | 0 | 0 | 0 | 0 |
| 224 | mcrB | 0 | 0 | 0 | 0 | 0 | 0 | 0 | 0 | 0 | 0 |
| 225 | gsk | 0 | 0 | 0 | 0 | 0 | 0 | 0 | 0 | 0 | 0 |
| 226 | mcrA | 0 | 0 | 0 | 0 | 0 | 0 | 0 | 0 | 0 | 0 |
| 227 | flgH | 58 | 0 | 29 | 1 | 81 | 0 | 30 | 0 | 0 | 0 |
| 228 | gloA | 0 | 0 | 0 | 0 | 0 | 0 | 0 | 0 | 0 | 0 |
| 229 | lpp | 0 | 0 | 0 | 0 | 0 | 0 | 0 | 0 | 0 | 0 |
| 230 | lpd | 0 | 0 | 0 | 0 | 0 | 0 | 0 | 0 | 0 | 0 |
| 231 | arpA | 0 | 0 | 0 | 0 | 0 | 0 | 0 | 0 | 0 | 0 |
| 232 | aldH | 0 | 0 | 0 | 0 | 0 | 0 | 0 | 0 | 0 | 0 |
| 233 | menD | 0 | 0 | 0 | 0 | 0 | 0 | 0 | 1 | 0 | 0 |
| 234 | sieB | 0 | 0 | 0 | 0 | 0 | 0 | 0 | 0 | 0 | 0 |
| 235 | menF | 0 | 0 | 0 | 0 | 0 | 0 | 0 | 0 | 0 | 0 |
| 236 | menG | 0 | 0 | 0 | 0 | 0 | 0 | 0 | 0 | 0 | 0 |
| 237 | aldB | 0 | 0 | 0 | 0 | 0 | 0 | 0 | 0 | 0 | 0 |
| 238 | aldA | 0 | 0 | 1 | 0 | 0 | 0 | 0 | 0 | 0 | 0 |
| 239 | sbcB | 0 | 0 | 0 | 0 | 0 | 0 | 0 | 0 | 0 | 0 |
| 240 | priA | 0 | 0 | 0 | 0 | 0 | 0 | 0 | 0 | 0 | 0 |
| 241 | priC | 0 | 0 | 0 | 0 | 0 | 0 | 0 | 0 | 0 | 0 |
| 242 | yojI | 0 | 1 | 0 | 0 | 0 | 0 | 0 | 0 | 0 | 0 |
| 243 | yebU | 0 | 0 | 0 | 0 | 0 | 0 | 0 | 0 | 0 | 0 |
| 244 | yebQ | 0 | 0 | 0 | 0 | 0 | 0 | 0 | 0 | 0 | 0 |
| 245 | yebR | 0 | 0 | 0 | 0 | 0 | 0 | 0 | 0 | 0 | 0 |
| 246 | yebS | 0 | 0 | 0 | 0 | 0 | 0 | 0 | 0 | 0 | 0 |
| 247 | yebO | 0 | 0 | 0 | 0 | 0 | 0 | 0 | 0 | 0 | 0 |
| 248 | yebE | 0 | 0 | 1 | 0 | 0 | 0 | 1 | 0 | 0 | 0 |
| 249 | yebF | 12 | 0 | 6 | 1 | 33 | 0 | 8 | 0 | 16 | 0 |
| 250 | yebG | 0 | 0 | 0 | 0 | 0 | 0 | 0 | 0 | 0 | 0 |
| 251 | yebA | 0 | 0 | 0 | 0 | 0 | 0 | 1 | 0 | 0 | 0 |
| 252 | yebB | 0 | 0 | 0 | 0 | 0 | 0 | 0 | 0 | 0 | 0 |
| 253 | ygjN | 0 | 0 | 0 | 0 | 0 | 0 | 1 | 0 | 0 | 0 |
| 254 | ygjO | 0 | 1 | 0 | 0 | 0 | 0 | 0 | 0 | 0 | 0 |
| 255 | ygjJ | 0 | 0 | 0 | 0 | 0 | 0 | 0 | 0 | 0 | 0 |
| 256 | ygjH | 0 | 0 | 0 | 0 | 0 | 0 | 0 | 0 | 0 | 0 |
| 257 | ygjI | 0 | 0 | 0 | 0 | 0 | 0 | 0 | 0 | 0 | 0 |
| 258 | ygjG | 29 | 0 | 10 | 1 | 45 | 0 | 54 | 0 | 0 | 0 |
| 259 | ygjD | 0 | 0 | 0 | 0 | 0 | 0 | 0 | 0 | 0 | 0 |
| 260 | bax | 0 | 0 | 0 | 0 | 1 | 0 | 0 | 0 | 0 | 0 |
| 261 | ygjV | 0 | 0 | 0 | 0 | 0 | 0 | 0 | 0 | 0 | 0 |
| 262 | yacL | 0 | 0 | 0 | 0 | 0 | 0 | 0 | 0 | 0 | 0 |
| 263 | ygjR | 0 | 0 | 0 | 0 | 0 | 0 | 0 | 0 | 0 | 0 |

| | | | | | | | | | | | |
|---|---|---|---|---|---|---|---|---|---|---|---|
| 264 | ygjQ | 0 | 0 | 0 | 0 | 0 | 0 | 0 | 0 | 0 | 0 |
| 265 | ecfG | 0 | 0 | 0 | 0 | 0 | 0 | 0 | 0 | 0 | 0 |
| 266 | ecfD | 0 | 0 | 0 | 0 | 0 | 0 | 0 | 1 | 0 | 0 |
| 267 | ecfE | 0 | 0 | 0 | 0 | 0 | 0 | 0 | 0 | 0 | 0 |
| 268 | ecfK | 0 | 0 | 0 | 0 | 0 | 0 | 0 | 0 | 0 | 0 |
| 269 | ecfI | 0 | 0 | 0 | 0 | 0 | 0 | 0 | 0 | 0 | 0 |
| 270 | dctA | 0 | 0 | 0 | 0 | 0 | 0 | 0 | 0 | 0 | 0 |
| 271 | pmrD | 0 | 0 | 0 | 0 | 0 | 0 | 0 | 0 | 0 | 0 |
| 272 | ybaK | 0 | 0 | 0 | 0 | 0 | 0 | 0 | 0 | 0 | 0 |
| 273 | ybaN | 0 | 0 | 0 | 0 | 0 | 0 | 0 | 0 | 0 | 0 |
| 274 | ybaO | 0 | 0 | 0 | 0 | 0 | 0 | 0 | 0 | 0 | 0 |
| 275 | ybaL | 0 | 0 | 0 | 0 | 0 | 0 | 0 | 0 | 0 | 0 |
| 276 | ybaB | 0 | 0 | 0 | 0 | 0 | 0 | 0 | 0 | 0 | 0 |
| 277 | ybaA | 0 | 0 | 0 | 0 | 0 | 0 | 0 | 0 | 0 | 0 |
| 278 | ybaD | 0 | 0 | 0 | 0 | 0 | 0 | 0 | 0 | 0 | 0 |
| 279 | ybaE | 0 | 0 | 1 | 0 | 0 | 0 | 0 | 1 | 0 | 0 |
| 280 | mug | 0 | 0 | 0 | 0 | 0 | 0 | 0 | 0 | 0 | 0 |
| 281 | ybaY | 0 | 0 | 0 | 0 | 0 | 0 | 0 | 0 | 0 | 0 |
| 282 | yzgL | 0 | 0 | 0 | 0 | 0 | 0 | 0 | 0 | 0 | 0 |
| 283 | ybaP | 0 | 0 | 0 | 0 | 0 | 0 | 0 | 0 | 0 | 0 |
| 284 | ybaQ | 0 | 0 | 0 | 0 | 0 | 0 | 0 | 0 | 0 | 0 |
| 285 | ybaV | 0 | 0 | 0 | 0 | 0 | 0 | 0 | 0 | 0 | 0 |
| 286 | ybaT | 0 | 0 | 0 | 0 | 0 | 0 | 0 | 0 | 0 | 0 |
| 287 | yijF | 0 | 0 | 0 | 0 | 0 | 0 | 0 | 0 | 0 | 0 |
| 288 | ygdL | 0 | 0 | 0 | 0 | 0 | 0 | 0 | 0 | 0 | 0 |
| 289 | ygdI_csdA | 0 | 0 | 0 | 0 | 0 | 0 | 0 | 0 | 0 | 0 |
| 290 | yijO | 16 | 0 | 10 | 0 | 16 | 0 | 15 | 0 | 0 | 0 |
| 291 | ygdQ | 0 | 0 | 0 | 0 | 0 | 0 | 0 | 0 | 0 | 0 |
| 292 | ygdR | 0 | 0 | 0 | 0 | 0 | 0 | 0 | 0 | 0 | 0 |
| 293 | pitA | 0 | 0 | 1 | 0 | 0 | 0 | 2 | 0 | 0 | 0 |
| 294 | b2974 | 0 | 0 | 1 | 0 | 0 | 0 | 0 | 0 | 0 | 0 |
| 295 | pitB | 0 | 0 | 0 | 0 | 0 | 0 | 0 | 0 | 0 | 0 |
| 296 | ung | 0 | 0 | 1 | 0 | 0 | 0 | 1 | 0 | 0 | 0 |
| 297 | rimL | 0 | 0 | 1 | 0 | 0 | 0 | 0 | 0 | 0 | 0 |
| 298 | rimJ | 0 | 0 | 0 | 0 | 0 | 0 | 0 | 0 | 0 | 0 |
| 299 | insA2 | 0 | 0 | 0 | 0 | 0 | 0 | 0 | 0 | 0 | 0 |
| 300 | tonB | 0 | 0 | 0 | 0 | 0 | 0 | 0 | 0 | 0 | 0 |
| 301 | ydaG | 0 | 0 | 0 | 0 | 0 | 0 | 0 | 0 | 0 | 0 |
| 302 | crcA | 0 | 0 | 0 | 1 | 0 | 0 | 0 | 0 | 0 | 0 |
| 303 | yaaW | 0 | 0 | 0 | 0 | 0 | 0 | 0 | 0 | 0 | 0 |
| 304 | ydaO | 0 | 0 | 0 | 0 | 0 | 0 | 0 | 0 | 0 | 0 |
| 305 | yaaI | 0 | 0 | 0 | 0 | 0 | 0 | 0 | 0 | 0 | 0 |
| 306 | frr | 0 | 0 | 0 | 0 | 0 | 0 | 0 | 0 | 0 | 0 |
| 307 | yaaJ | 0 | 0 | 0 | 0 | 0 | 0 | 0 | 0 | 0 | 0 |
| 308 | yaaA | 0 | 0 | 0 | 0 | 0 | 0 | 0 | 0 | 0 | 0 |
| 309 | lctP | 0 | 0 | 0 | 0 | 1 | 0 | 0 | 0 | 0 | 0 |
| 310 | znuC | 0 | 0 | 0 | 0 | 0 | 0 | 0 | 0 | 0 | 0 |
| 311 | znuA | 0 | 0 | 0 | 0 | 0 | 0 | 0 | 0 | 0 | 0 |
| 312 | slt | 0 | 0 | 0 | 0 | 0 | 0 | 0 | 0 | 0 | 0 |
| 313 | ais | 0 | 0 | 0 | 0 | 0 | 0 | 0 | 0 | 0 | 0 |
| 314 | hepA | 0 | 0 | 0 | 0 | 0 | 0 | 0 | 0 | 0 | 0 |
| 315 | gnd | 0 | 0 | 0 | 0 | 0 | 0 | 0 | 0 | 0 | 0 |
| 316 | lepA | 0 | 0 | 0 | 0 | 0 | 0 | 0 | 0 | 0 | 0 |
| 317 | hupA | 0 | 0 | 3 | 0 | 0 | 0 | 0 | 0 | 0 | 0 |
| 318 | yfcI | 0 | 0 | 0 | 0 | 0 | 0 | 0 | 0 | 0 | 0 |
| 319 | yfcJ | 0 | 0 | 0 | 0 | 0 | 0 | 0 | 0 | 1 | 0 |
| 320 | hupB | 0 | 0 | 0 | 0 | 0 | 0 | 0 | 0 | 0 | 0 |
| 321 | yfcC | 67 | 1 | 7 | 0 | 145 | 0 | 68 | 0 | 82 | 0 |
| 322 | yfcD | 0 | 0 | 0 | 0 | 0 | 0 | 0 | 0 | 0 | 0 |
| 323 | yfcE | 0 | 0 | 0 | 0 | 0 | 0 | 0 | 0 | 0 | 0 |
| 324 | yfcG | 0 | 0 | 0 | 0 | 0 | 0 | 0 | 1 | 0 | 0 |
| 325 | yfcU | 0 | 0 | 0 | 0 | 0 | 0 | 0 | 0 | 0 | 0 |
| 326 | yfcV | 0 | 0 | 0 | 0 | 0 | 0 | 0 | 0 | 0 | 0 |
| 327 | glnK | 0 | 0 | 0 | 0 | 0 | 0 | 0 | 0 | 0 | 0 |
| 328 | glnH | 0 | 0 | 0 | 0 | 0 | 0 | 0 | 0 | 0 | 0 |
| 329 | glnL | 0 | 0 | 0 | 0 | 0 | 0 | 1 | 0 | 0 | 0 |

| | | | | | | | | | | | |
|---|---|---|---|---|---|---|---|---|---|---|---|
| 330 | glnB | 0 | 0 | 0 | 0 | 0 | 0 | 0 | 0 | 0 | 0 |
| 331 | glnA | 0 | 0 | 0 | 0 | 0 | 0 | 0 | 0 | 0 | 0 |
| 332 | glnW | 0 | 1 | 0 | 0 | 0 | 0 | 0 | 0 | 0 | 0 |
| 333 | nudE | 0 | 0 | 0 | 0 | 0 | 0 | 0 | 0 | 0 | 0 |
| 334 | glnU | 0 | 0 | 0 | 0 | 0 | 0 | 0 | 0 | 0 | 0 |
| 335 | cynT | 1 | 0 | 0 | 0 | 0 | 0 | 1 | 0 | 1 | 0 |
| 336 | cynR | 1340 | 25 | 264 | 27 | 2426 | 68 | 1604 | 5 | 1309 | 2 |
| 337 | rne | 0 | 0 | 0 | 0 | 0 | 0 | 0 | 0 | 0 | 0 |
| 338 | glxK | 0 | 0 | 0 | 0 | 0 | 0 | 0 | 0 | 0 | 0 |
| 339 | kil | 0 | 0 | 0 | 0 | 0 | 0 | 0 | 0 | 0 | 0 |
| 340 | gssA | 0 | 0 | 0 | 0 | 0 | 0 | 0 | 0 | 0 | 0 |
| 341 | yneJ | 0 | 0 | 0 | 0 | 0 | 0 | 0 | 0 | 0 | 0 |
| 342 | yneK | 0 | 0 | 0 | 0 | 0 | 0 | 0 | 0 | 0 | 0 |
| 343 | yneH | 0 | 0 | 0 | 0 | 0 | 0 | 0 | 0 | 0 | 0 |
| 344 | yneI | 0 | 0 | 0 | 0 | 0 | 0 | 0 | 0 | 0 | 0 |
| 345 | amyA | 0 | 0 | 0 | 0 | 0 | 0 | 0 | 0 | 0 | 0 |
| 346 | yneE | 0 | 0 | 0 | 0 | 0 | 0 | 0 | 1 | 0 | 0 |
| 347 | yecM | 0 | 0 | 0 | 0 | 0 | 0 | 0 | 0 | 0 | 0 |
| 348 | yfeA | 0 | 0 | 0 | 0 | 0 | 0 | 0 | 0 | 0 | 0 |
| 349 | yecH | 0 | 0 | 0 | 0 | 0 | 0 | 0 | 0 | 0 | 0 |
| 350 | yecF | 0 | 0 | 0 | 0 | 0 | 0 | 0 | 0 | 0 | 0 |
| 351 | fadI | 0 | 0 | 0 | 5 | 0 | 0 | 1 | 0 | 0 | 0 |
| 352 | yjjU | 0 | 0 | 0 | 0 | 0 | 0 | 0 | 0 | 0 | 0 |
| 353 | ychJ | 0 | 0 | 0 | 0 | 0 | 0 | 0 | 0 | 0 | 0 |
| 354 | rnk | 0 | 0 | 0 | 0 | 0 | 0 | 1 | 0 | 0 | 0 |
| 355 | astC | 0 | 0 | 0 | 0 | 0 | 0 | 0 | 0 | 0 | 0 |
| 356 | nudH | 0 | 0 | 0 | 0 | 0 | 0 | 0 | 0 | 0 | 0 |
| 357 | nudG | 0 | 0 | 0 | 0 | 0 | 0 | 0 | 0 | 0 | 0 |
| 358 | nudF | 0 | 0 | 0 | 0 | 0 | 0 | 0 | 0 | 0 | 0 |
| 359 | rna | 0 | 0 | 0 | 0 | 0 | 0 | 0 | 0 | 0 | 0 |
| 360 | yliG | 0 | 0 | 0 | 0 | 0 | 0 | 0 | 0 | 0 | 0 |
| 361 | nudC | 24 | 0 | 5 | 0 | 63 | 0 | 10 | 0 | 1 | 0 |
| 362 | nudB | 0 | 0 | 0 | 0 | 0 | 0 | 2 | 0 | 0 | 0 |
| 363 | gadB | 0 | 0 | 0 | 0 | 0 | 0 | 0 | 0 | 0 | 0 |
| 364 | rnd | 0 | 0 | 0 | 0 | 0 | 0 | 0 | 0 | 0 | 0 |
| 365 | gadX | 0 | 0 | 0 | 0 | 0 | 0 | 0 | 0 | 0 | 0 |
| 366 | gadW | 0 | 0 | 0 | 0 | 0 | 0 | 0 | 0 | 0 | 0 |
| 367 | rnt | 0 | 0 | 0 | 0 | 0 | 0 | 0 | 0 | 0 | 0 |
| 368 | stpA | 0 | 0 | 0 | 0 | 0 | 0 | 0 | 0 | 0 | 0 |
| 369 | araB | 0 | 0 | 0 | 0 | 0 | 0 | 0 | 0 | 0 | 0 |
| 370 | araC | 0 | 0 | 0 | 0 | 0 | 0 | 0 | 0 | 0 | 0 |
| 371 | araF | 0 | 0 | 0 | 0 | 0 | 0 | 0 | 0 | 0 | 0 |
| 372 | araD | 0 | 0 | 1 | 0 | 0 | 0 | 0 | 0 | 0 | 0 |
| 373 | araE | 0 | 0 | 0 | 0 | 0 | 0 | 0 | 1 | 0 | 0 |
| 374 | araJ | 0 | 0 | 0 | 0 | 0 | 1 | 0 | 0 | 0 | 0 |
| 375 | ygeG | 0 | 0 | 0 | 0 | 0 | 0 | 0 | 0 | 0 | 0 |
| 376 | ygeF | 1 | 0 | 0 | 0 | 0 | 0 | 1 | 0 | 0 | 0 |
| 377 | ygeA | 0 | 0 | 0 | 0 | 0 | 0 | 0 | 0 | 0 | 0 |
| 378 | ygeI | 0 | 0 | 0 | 0 | 0 | 0 | 0 | 0 | 0 | 0 |
| 379 | cirA | 0 | 0 | 0 | 0 | 0 | 0 | 0 | 0 | 0 | 0 |
| 380 | ygeR | 0 | 0 | 0 | 1 | 0 | 0 | 0 | 0 | 0 | 0 |
| 381 | ygeQ | 0 | 0 | 0 | 0 | 0 | 0 | 0 | 0 | 0 | 0 |
| 382 | fruL | 0 | 0 | 0 | 0 | 0 | 0 | 0 | 0 | 0 | 0 |
| 383 | fruB | 0 | 0 | 0 | 0 | 0 | 0 | 0 | 0 | 0 | 0 |
| 384 | ygeY | 0 | 0 | 0 | 0 | 0 | 0 | 0 | 0 | 0 | 0 |
| 385 | ygeX | 0 | 0 | 0 | 0 | 0 | 0 | 0 | 0 | 0 | 0 |
| 386 | rffC | 0 | 0 | 0 | 0 | 0 | 0 | 0 | 0 | 0 | 0 |
| 387 | ypfG | 0 | 0 | 0 | 0 | 0 | 0 | 0 | 1 | 0 | 0 |
| 388 | ypfH | 0 | 0 | 0 | 0 | 0 | 0 | 0 | 0 | 0 | 0 |
| 389 | ypfJ | 0 | 0 | 0 | 0 | 0 | 0 | 0 | 0 | 0 | 0 |
| 390 | deaD | 0 | 0 | 0 | 0 | 0 | 0 | 0 | 0 | 0 | 0 |
| 391 | icd | 0 | 0 | 0 | 0 | 0 | 0 | 0 | 0 | 0 | 0 |
| 392 | borD | 0 | 0 | 0 | 0 | 0 | 0 | 0 | 0 | 0 | 0 |
| 393 | hdeD | 0 | 0 | 0 | 0 | 0 | 0 | 0 | 0 | 0 | 0 |
| 394 | hdeB | 0 | 0 | 0 | 0 | 0 | 0 | 0 | 0 | 0 | 0 |
| 395 | hdeA | 0 | 0 | 0 | 0 | 0 | 0 | 0 | 0 | 0 | 0 |

| | | | | | | | | | | | |
|---|---|---|---|---|---|---|---|---|---|---|---|
| 396 | ribA | 0 | 0 | 0 | 0 | 0 | 0 | 0 | 0 | 0 | 0 |
| 397 | ribC | 0 | 0 | 0 | 0 | 0 | 0 | 0 | 0 | 0 | 0 |
| 398 | ribB | 0 | 0 | 0 | 0 | 0 | 0 | 0 | 0 | 0 | 0 |
| 399 | ribE | 0 | 0 | 0 | 0 | 0 | 0 | 0 | 0 | 0 | 0 |
| 400 | htpX | 0 | 0 | 0 | 0 | 0 | 0 | 0 | 0 | 0 | 0 |
| 401 | yqjI | 0 | 0 | 0 | 0 | 0 | 0 | 0 | 0 | 0 | 0 |
| 402 | corA | 0 | 1 | 0 | 0 | 0 | 0 | 0 | 0 | 0 | 0 |
| 403 | yqjC | 0 | 0 | 0 | 0 | 0 | 0 | 0 | 0 | 0 | 0 |
| 404 | yqjF | 0 | 0 | 0 | 0 | 0 | 0 | 0 | 0 | 0 | 0 |
| 405 | yqjG | 0 | 0 | 0 | 0 | 0 | 0 | 0 | 0 | 0 | 0 |
| 406 | pheS | 0 | 0 | 0 | 0 | 0 | 0 | 0 | 0 | 0 | 0 |
| 407 | pheP | 0 | 0 | 0 | 0 | 0 | 0 | 0 | 0 | 0 | 0 |
| 408 | ileX | 0 | 0 | 0 | 0 | 0 | 0 | 0 | 0 | 0 | 0 |
| 409 | htpG | 0 | 0 | 0 | 0 | 0 | 0 | 0 | 0 | 0 | 0 |
| 410 | ppa | 0 | 0 | 0 | 0 | 0 | 0 | 0 | 0 | 0 | 0 |
| 411 | dkgB | 0 | 0 | 0 | 0 | 0 | 0 | 0 | 0 | 0 | 0 |
| 412 | dkgA | 0 | 0 | 0 | 0 | 0 | 0 | 0 | 0 | 0 | 0 |
| 413 | relB | 0 | 0 | 0 | 0 | 0 | 0 | 0 | 0 | 0 | 0 |
| 414 | relA | 0 | 0 | 0 | 0 | 0 | 0 | 0 | 0 | 0 | 0 |
| 415 | phnQ | 126 | 1 | 44 | 5 | 205 | 0 | 71 | 4 | 11 | 0 |
| 416 | pykF | 0 | 0 | 0 | 0 | 0 | 0 | 0 | 0 | 0 | 0 |
| 417 | sfmA | 0 | 0 | 0 | 0 | 0 | 0 | 0 | 0 | 0 | 0 |
| 418 | sfmC | 0 | 0 | 0 | 0 | 0 | 0 | 0 | 0 | 0 | 0 |
| 419 | b1997_b1998 | 0 | 0 | 0 | 0 | 0 | 0 | 0 | 0 | 0 | 0 |
| 420 | pphA | 0 | 0 | 0 | 0 | 0 | 0 | 0 | 0 | 0 | 0 |
| 421 | mtr | 0 | 0 | 0 | 0 | 0 | 0 | 0 | 0 | 0 | 0 |
| 422 | thrU | 0 | 0 | 0 | 0 | 0 | 0 | 0 | 0 | 0 | 0 |
| 423 | msbB | 44 | 0 | 24 | 0 | 83 | 1 | 58 | 0 | 30 | 0 |
| 424 | dacA | 0 | 0 | 0 | 0 | 0 | 0 | 0 | 0 | 0 | 0 |
| 425 | dacB | 0 | 0 | 0 | 0 | 0 | 0 | 0 | 0 | 0 | 0 |
| 426 | tesB | 0 | 0 | 0 | 0 | 0 | 0 | 0 | 0 | 0 | 0 |
| 427 | yigI | 0 | 0 | 0 | 0 | 0 | 0 | 0 | 0 | 0 | 0 |
| 428 | cyoA | 0 | 0 | 0 | 0 | 0 | 0 | 0 | 0 | 0 | 0 |
| 429 | dgkA | 0 | 0 | 0 | 0 | 1 | 0 | 0 | 0 | 0 | 0 |
| 430 | shiA | 0 | 0 | 0 | 0 | 0 | 0 | 1 | 0 | 0 | 0 |
| 431 | ybfG | 0 | 0 | 0 | 0 | 0 | 0 | 0 | 0 | 0 | 0 |
| 432 | tig | 0 | 0 | 0 | 0 | 0 | 1 | 0 | 0 | 0 | 0 |
| 433 | ykgD | 0 | 0 | 0 | 0 | 0 | 0 | 0 | 0 | 0 | 0 |
| 434 | sdaA | 0 | 0 | 0 | 0 | 0 | 0 | 0 | 0 | 0 | 0 |
| 435 | nmpC | 0 | 0 | 0 | 0 | 0 | 0 | 0 | 0 | 0 | 0 |
| 436 | idnT | 11 | 0 | 1 | 2 | 25 | 0 | 0 | 0 | 64 | 0 |
| 437 | idnR | 0 | 0 | 0 | 0 | 0 | 0 | 0 | 0 | 0 | 0 |
| 438 | sbmA | 0 | 0 | 0 | 0 | 0 | 0 | 0 | 0 | 0 | 0 |
| 439 | sbmC | 0 | 0 | 0 | 0 | 0 | 0 | 0 | 0 | 0 | 0 |
| 440 | idnK | 0 | 0 | 0 | 0 | 0 | 0 | 0 | 0 | 0 | 0 |
| 441 | mltC | 0 | 0 | 0 | 0 | 0 | 0 | 0 | 0 | 0 | 0 |
| 442 | mltD | 0 | 0 | 0 | 0 | 0 | 0 | 0 | 0 | 0 | 0 |
| 443 | pppA | 0 | 0 | 0 | 0 | 0 | 0 | 0 | 0 | 0 | 0 |
| 444 | xthA | 0 | 0 | 0 | 0 | 0 | 0 | 0 | 0 | 0 | 0 |
| 445 | fsr | 0 | 0 | 0 | 0 | 0 | 0 | 0 | 0 | 0 | 0 |
| 446 | rhaT | 0 | 0 | 0 | 0 | 0 | 0 | 0 | 0 | 0 | 0 |
| 447 | aspA | 0 | 0 | 0 | 0 | 0 | 0 | 0 | 0 | 0 | 0 |
| 448 | rhaS | 0 | 0 | 0 | 0 | 0 | 0 | 0 | 0 | 0 | 0 |
| 449 | aspC | 0 | 0 | 0 | 0 | 0 | 0 | 0 | 0 | 0 | 0 |
| 450 | rob | 0 | 0 | 0 | 0 | 0 | 0 | 0 | 0 | 0 | 0 |
| 451 | grxA | 0 | 0 | 0 | 0 | 0 | 0 | 0 | 0 | 0 | 0 |
| 452 | rof | 23 | 0 | 1 | 0 | 37 | 0 | 20 | 10 | 3 | 0 |
| 453 | aspU | 0 | 0 | 0 | 0 | 0 | 0 | 0 | 0 | 0 | 0 |
| 454 | rhaB | 0 | 0 | 0 | 0 | 0 | 0 | 0 | 0 | 0 | 0 |
| 455 | yhjK | 0 | 0 | 0 | 0 | 0 | 0 | 0 | 0 | 0 | 0 |
| 456 | yhjJ | 0 | 0 | 0 | 0 | 0 | 0 | 0 | 0 | 0 | 0 |
| 457 | yjjV | 0 | 0 | 0 | 0 | 0 | 0 | 0 | 0 | 0 | 0 |
| 458 | yhjG | 0 | 0 | 0 | 0 | 0 | 0 | 0 | 0 | 0 | 0 |
| 459 | dsdC_dsdX | 0 | 0 | 0 | 0 | 0 | 0 | 0 | 0 | 0 | 0 |
| 460 | yhjC | 0 | 0 | 0 | 0 | 0 | 0 | 0 | 0 | 0 | 0 |
| 461 | yhjB | 0 | 0 | 0 | 0 | 0 | 0 | 0 | 0 | 0 | 0 |

| | | | | | | | | | | | |
|---|---|---|---|---|---|---|---|---|---|---|---|
| 462 | yjjL | 0 | 0 | 0 | 0 | 0 | 0 | 0 | 0 | 0 | 0 |
| 463 | yhjY | 0 | 0 | 0 | 0 | 0 | 0 | 0 | 1 | 0 | 1 |
| 464 | pldA | 0 | 0 | 0 | 0 | 0 | 0 | 0 | 0 | 0 | 0 |
| 465 | yjjI | 0 | 0 | 0 | 0 | 0 | 0 | 0 | 0 | 0 | 0 |
| 466 | yhjR | 0 | 0 | 0 | 0 | 0 | 0 | 0 | 0 | 0 | 0 |
| 467 | yfbJ | 41 | 0 | 8 | 0 | 86 | 0 | 65 | 0 | 76 | 0 |
| 468 | yfbM | 0 | 0 | 0 | 0 | 0 | 0 | 0 | 0 | 0 | 0 |
| 469 | yfbE | 0 | 0 | 0 | 0 | 0 | 0 | 0 | 0 | 0 | 0 |
| 470 | apbE | 0 | 0 | 0 | 0 | 0 | 0 | 0 | 0 | 0 | 0 |
| 471 | yfbS | 0 | 0 | 0 | 0 | 0 | 0 | 0 | 0 | 0 | 0 |
| 472 | yfbR | 0 | 0 | 0 | 0 | 0 | 0 | 0 | 0 | 0 | 0 |
| 473 | yfbV | 0 | 0 | 0 | 0 | 0 | 0 | 0 | 0 | 0 | 0 |
| 474 | yfbU | 0 | 0 | 0 | 0 | 0 | 0 | 0 | 0 | 0 | 0 |
| 475 | glmU | 0 | 0 | 0 | 0 | 0 | 0 | 0 | 0 | 0 | 0 |
| 476 | eutB | 0 | 0 | 0 | 0 | 0 | 0 | 0 | 0 | 0 | 0 |
| 477 | cyaA_hemC | 0 | 0 | 0 | 0 | 0 | 0 | 0 | 0 | 0 | 0 |
| 478 | der | 0 | 0 | 0 | 0 | 0 | 0 | 0 | 0 | 0 | 0 |
| 479 | hdfR | 0 | 0 | 0 | 0 | 0 | 0 | 0 | 0 | 0 | 0 |
| 480 | hscC | 0 | 0 | 0 | 0 | 0 | 0 | 0 | 0 | 0 | 0 |
| 481 | acpP | 0 | 0 | 0 | 0 | 0 | 0 | 0 | 0 | 0 | 0 |
| 482 | seqA | 0 | 0 | 0 | 0 | 0 | 0 | 0 | 0 | 0 | 0 |
| 483 | nupG | 0 | 0 | 0 | 0 | 0 | 0 | 0 | 0 | 0 | 0 |
| 484 | pmbA | 0 | 0 | 0 | 0 | 0 | 0 | 0 | 0 | 0 | 0 |
| 485 | yodB | 0 | 0 | 0 | 0 | 0 | 0 | 0 | 0 | 0 | 0 |
| 486 | nlpC | 0 | 0 | 0 | 0 | 0 | 0 | 0 | 0 | 0 | 0 |
| 487 | nlpA | 0 | 0 | 0 | 0 | 0 | 0 | 0 | 0 | 1 | 0 |
| 488 | cysH | 28 | 0 | 12 | 0 | 50 | 0 | 20 | 0 | 28 | 0 |
| 489 | nlpD | 0 | 0 | 0 | 0 | 0 | 0 | 0 | 0 | 0 | 0 |
| 490 | cysJ | 0 | 0 | 0 | 0 | 0 | 0 | 0 | 0 | 0 | 0 |
| 491 | cysD | 0 | 0 | 0 | 0 | 0 | 0 | 0 | 0 | 0 | 0 |
| 492 | nlpI | 0 | 0 | 0 | 0 | 0 | 0 | 0 | 0 | 0 | 0 |
| 493 | cysB | 0 | 0 | 0 | 0 | 0 | 0 | 0 | 0 | 0 | 0 |
| 494 | uxuR | 0 | 0 | 0 | 1 | 0 | 0 | 0 | 0 | 0 | 0 |
| 495 | cysZ | 0 | 0 | 0 | 0 | 0 | 0 | 0 | 1 | 0 | 0 |
| 496 | cysT | 0 | 0 | 0 | 0 | 0 | 0 | 0 | 0 | 0 | 0 |
| 497 | cysW | 0 | 0 | 1 | 0 | 0 | 0 | 0 | 0 | 0 | 0 |
| 498 | cysQ | 12 | 0 | 2 | 0 | 44 | 0 | 21 | 0 | 43 | 0 |
| 499 | cysP | 0 | 0 | 0 | 0 | 0 | 0 | 0 | 0 | 0 | 0 |
| 500 | cysS | 0 | 0 | 0 | 0 | 0 | 0 | 0 | 0 | 0 | 0 |
| 501 | def | 0 | 0 | 0 | 0 | 0 | 0 | 0 | 0 | 0 | 0 |
| 502 | perR | 0 | 0 | 0 | 0 | 0 | 0 | 0 | 0 | 0 | 0 |
| 503 | ppiD | 0 | 0 | 0 | 0 | 0 | 0 | 0 | 0 | 0 | 0 |
| 504 | ppiC | 0 | 0 | 0 | 0 | 0 | 0 | 0 | 0 | 0 | 0 |
| 505 | ppiB | 0 | 0 | 0 | 0 | 0 | 0 | 0 | 0 | 0 | 0 |
| 506 | ppiA | 0 | 0 | 0 | 0 | 0 | 0 | 0 | 0 | 0 | 0 |
| 507 | ybgQ | 0 | 0 | 0 | 0 | 0 | 0 | 0 | 0 | 0 | 0 |
| 508 | ybgS | 0 | 0 | 0 | 0 | 0 | 0 | 0 | 0 | 0 | 0 |
| 509 | yifB | 0 | 0 | 0 | 0 | 0 | 0 | 0 | 0 | 0 | 0 |
| 510 | mscS | 0 | 0 | 0 | 0 | 0 | 0 | 0 | 0 | 0 | 0 |
| 511 | ykfG | 0 | 0 | 0 | 0 | 0 | 0 | 0 | 0 | 0 | 0 |
| 512 | ybgC | 0 | 0 | 1 | 0 | 0 | 0 | 0 | 0 | 0 | 0 |
| 513 | ybgD | 0 | 0 | 0 | 0 | 0 | 0 | 0 | 0 | 0 | 0 |
| 514 | mscL | 0 | 0 | 0 | 0 | 0 | 0 | 0 | 0 | 0 | 0 |
| 515 | ykfA | 0 | 0 | 0 | 0 | 0 | 0 | 0 | 0 | 0 | 0 |
| 516 | ybgH | 0 | 0 | 0 | 0 | 0 | 0 | 0 | 0 | 0 | 0 |
| 517 | ybgI | 0 | 0 | 0 | 0 | 0 | 0 | 0 | 0 | 0 | 0 |
| 518 | ykfJ | 0 | 0 | 0 | 0 | 0 | 0 | 0 | 0 | 0 | 0 |
| 519 | ykfI | 0 | 0 | 0 | 0 | 0 | 0 | 0 | 0 | 0 | 0 |
| 520 | yihN | 0 | 0 | 0 | 0 | 0 | 0 | 0 | 0 | 0 | 0 |
| 521 | bolA | 0 | 0 | 0 | 0 | 0 | 0 | 0 | 0 | 0 | 0 |
| 522 | yihL | 0 | 0 | 0 | 0 | 0 | 0 | 0 | 0 | 0 | 0 |
| 523 | ygfF | 0 | 0 | 0 | 0 | 0 | 0 | 0 | 0 | 0 | 0 |
| 524 | yihD | 18 | 0 | 2 | 1 | 21 | 0 | 6 | 0 | 19 | 0 |
| 525 | ygfI | 0 | 0 | 0 | 0 | 0 | 0 | 0 | 0 | 0 | 0 |
| 526 | mdtG | 0 | 0 | 0 | 0 | 0 | 0 | 0 | 0 | 0 | 0 |
| 527 | yihX | 0 | 0 | 1 | 0 | 0 | 0 | 0 | 0 | 0 | 0 |

| | | | | | | | | | | | |
|---|---|---|---|---|---|---|---|---|---|---|---|
| 528 | ygfU | 0 | 0 | 0 | 0 | 0 | 0 | 0 | 0 | 0 | 0 |
| 529 | mdtK | 0 | 0 | 0 | 0 | 0 | 0 | 0 | 0 | 0 | 0 |
| 530 | focA | 0 | 0 | 0 | 0 | 0 | 0 | 0 | 0 | 0 | 0 |
| 531 | yihU | 0 | 0 | 0 | 0 | 0 | 0 | 0 | 0 | 0 | 0 |
| 532 | mdtO | 47 | 0 | 41 | 1 | 101 | 0 | 63 | 0 | 0 | 0 |
| 533 | mdtN | 0 | 0 | 0 | 0 | 0 | 0 | 0 | 0 | 0 | 0 |
| 534 | hycB | 0 | 0 | 2 | 0 | 0 | 0 | 0 | 0 | 0 | 0 |
| 535 | flk | 0 | 0 | 0 | 0 | 0 | 0 | 0 | 0 | 0 | 0 |
| 536 | spy | 0 | 0 | 0 | 0 | 0 | 0 | 0 | 0 | 0 | 0 |
| 537 | pinH | 0 | 0 | 0 | 0 | 0 | 0 | 0 | 0 | 0 | 0 |
| 538 | purT | 0 | 0 | 0 | 0 | 0 | 0 | 0 | 0 | 0 | 0 |
| 539 | mpaA | 0 | 0 | 0 | 0 | 0 | 0 | 0 | 0 | 0 | 0 |
| 540 | spr | 0 | 0 | 0 | 0 | 0 | 0 | 0 | 0 | 0 | 0 |
| 541 | dxr | 0 | 0 | 0 | 0 | 1 | 0 | 0 | 0 | 0 | 0 |
| 542 | tilS | 0 | 0 | 0 | 0 | 0 | 0 | 0 | 0 | 0 | 0 |
| 543 | trxC | 0 | 0 | 0 | 0 | 0 | 0 | 0 | 0 | 0 | 0 |
| 544 | trxA | 0 | 0 | 0 | 0 | 0 | 0 | 0 | 0 | 0 | 0 |
| 545 | pheM | 0 | 0 | 0 | 0 | 0 | 0 | 0 | 0 | 0 | 0 |
| 546 | pinR | 0 | 0 | 0 | 0 | 0 | 0 | 0 | 0 | 0 | 0 |
| 547 | ylaB | 0 | 1 | 0 | 0 | 0 | 0 | 0 | 0 | 0 | 0 |
| 548 | ylaC | 0 | 0 | 0 | 0 | 0 | 0 | 0 | 0 | 0 | 0 |
| 549 | purM | 0 | 0 | 0 | 0 | 0 | 0 | 0 | 0 | 0 | 0 |
| 550 | ydcL | 0 | 0 | 0 | 0 | 0 | 0 | 0 | 0 | 0 | 0 |
| 551 | ydcM | 0 | 0 | 0 | 0 | 0 | 0 | 0 | 0 | 0 | 0 |
| 552 | ydcJ | 0 | 0 | 0 | 0 | 0 | 0 | 0 | 0 | 0 | 0 |
| 553 | ydcK | 0 | 0 | 0 | 0 | 0 | 0 | 0 | 0 | 0 | 0 |
| 554 | ydcI | 0 | 0 | 0 | 0 | 0 | 0 | 0 | 0 | 0 | 0 |
| 555 | ydcF | 508 | 2 | 162 | 438 | 938 | 957 | 569 | 1882 | 479 | 0 |
| 556 | ydcG | 0 | 0 | 0 | 0 | 0 | 0 | 0 | 0 | 0 | 0 |
| 557 | ydcA | 0 | 0 | 0 | 0 | 0 | 0 | 0 | 0 | 0 | 0 |
| 558 | fucR | 19 | 1 | 9 | 0 | 59 | 0 | 14 | 0 | 4 | 0 |
| 559 | ydcX | 0 | 0 | 0 | 0 | 0 | 0 | 0 | 0 | 0 | 0 |
| 560 | fpr | 0 | 0 | 0 | 0 | 0 | 0 | 0 | 0 | 0 | 0 |
| 561 | chiA | 0 | 0 | 0 | 0 | 0 | 0 | 0 | 0 | 0 | 0 |
| 562 | ydcR | 0 | 0 | 0 | 0 | 0 | 0 | 0 | 1 | 0 | 0 |
| 563 | ydcS | 0 | 0 | 0 | 0 | 0 | 0 | 0 | 0 | 0 | 0 |
| 564 | map | 0 | 0 | 0 | 0 | 0 | 0 | 0 | 0 | 0 | 0 |
| 565 | rrlG | 38 | 0 | 5 | 0 | 87 | 0 | 27 | 0 | 27 | 0 |
| 566 | cdaR | 0 | 0 | 0 | 0 | 0 | 0 | 0 | 0 | 0 | 0 |
| 567 | cpdA | 0 | 0 | 0 | 0 | 0 | 0 | 0 | 0 | 0 | 0 |
| 568 | cpdB | 0 | 0 | 0 | 0 | 0 | 0 | 0 | 0 | 0 | 0 |
| 569 | talA | 0 | 0 | 0 | 0 | 0 | 0 | 0 | 0 | 0 | 0 |
| 570 | talB | 0 | 0 | 0 | 0 | 0 | 0 | 0 | 0 | 0 | 0 |
| 571 | yncA | 0 | 0 | 0 | 0 | 0 | 0 | 0 | 0 | 0 | 0 |
| 572 | yncB | 0 | 0 | 0 | 0 | 0 | 0 | 0 | 0 | 0 | 0 |
| 573 | yncC | 0 | 0 | 0 | 0 | 0 | 0 | 0 | 0 | 0 | 0 |
| 574 | yncD | 0 | 0 | 0 | 0 | 0 | 0 | 0 | 0 | 0 | 0 |
| 575 | yncE | 0 | 0 | 0 | 0 | 0 | 0 | 0 | 0 | 0 | 0 |
| 576 | uvrY | 0 | 0 | 0 | 0 | 0 | 0 | 0 | 0 | 0 | 0 |
| 577 | yncG | 0 | 0 | 0 | 0 | 0 | 0 | 0 | 0 | 0 | 0 |
| 578 | yncJ | 0 | 0 | 0 | 0 | 0 | 0 | 0 | 0 | 0 | 0 |
| 579 | pncB | 0 | 0 | 0 | 0 | 0 | 0 | 0 | 0 | 0 | 0 |
| 580 | yfaY | 0 | 0 | 0 | 0 | 0 | 0 | 0 | 0 | 0 | 0 |
| 581 | torT | 0 | 0 | 0 | 0 | 0 | 0 | 0 | 0 | 0 | 0 |
| 582 | yfaD | 0 | 0 | 0 | 0 | 0 | 0 | 0 | 0 | 0 | 0 |
| 583 | torS | 0 | 0 | 0 | 0 | 0 | 0 | 0 | 0 | 0 | 0 |
| 584 | torR | 0 | 0 | 0 | 0 | 0 | 0 | 0 | 0 | 0 | 0 |
| 585 | yfaL | 0 | 0 | 0 | 0 | 0 | 0 | 0 | 0 | 1 | 0 |
| 586 | ygiL | 0 | 0 | 0 | 0 | 0 | 0 | 0 | 0 | 0 | 0 |
| 587 | torY | 0 | 0 | 0 | 0 | 0 | 0 | 0 | 0 | 0 | 0 |
| 588 | uvrA | 0 | 0 | 0 | 0 | 0 | 0 | 0 | 0 | 0 | 0 |
| 589 | ycaO | 3 | 0 | 2 | 0 | 2 | 0 | 2 | 0 | 0 | 0 |
| 590 | ycaN | 0 | 0 | 0 | 0 | 0 | 0 | 0 | 0 | 0 | 0 |
| 591 | ycaL | 0 | 0 | 0 | 0 | 0 | 0 | 0 | 0 | 0 | 0 |
| 592 | ycaK | 0 | 0 | 0 | 0 | 0 | 0 | 0 | 0 | 0 | 0 |
| 593 | cutA | 0 | 0 | 0 | 0 | 0 | 0 | 0 | 0 | 0 | 0 |

| | | | | | | | | | | | |
|---|---|---|---|---|---|---|---|---|---|---|---|
| 594 | ycaI | 0 | 0 | 1 | 0 | 0 | 0 | 0 | 0 | 0 | 0 |
| 595 | ycaD | 0 | 0 | 0 | 0 | 0 | 0 | 0 | 0 | 0 | 0 |
| 596 | ycaC | 0 | 0 | 0 | 0 | 0 | 0 | 0 | 0 | 0 | 0 |
| 597 | cybC | 0 | 0 | 0 | 0 | 0 | 0 | 0 | 0 | 0 | 0 |
| 598 | tktB | 35 | 0 | 4 | 0 | 49 | 0 | 27 | 0 | 0 | 0 |
| 599 | ssb | 0 | 0 | 0 | 0 | 0 | 0 | 0 | 0 | 0 | 0 |
| 600 | ycaP | 0 | 0 | 0 | 0 | 0 | 0 | 0 | 0 | 0 | 0 |
| 601 | b0540 | 0 | 0 | 0 | 0 | 0 | 0 | 0 | 0 | 0 | 0 |
| 602 | yqgD | 10 | 0 | 0 | 0 | 12 | 0 | 1 | 0 | 0 | 0 |
| 603 | lexA | 0 | 0 | 0 | 0 | 0 | 0 | 1 | 0 | 0 | 0 |
| 604 | cobU | 0 | 0 | 0 | 0 | 0 | 0 | 0 | 0 | 0 | 0 |
| 605 | ybgE | 39 | 0 | 11 | 1 | 79 | 0 | 97 | 1 | 0 | 0 |
| 606 | ymcE | 0 | 0 | 0 | 0 | 0 | 0 | 0 | 0 | 0 | 0 |
| 607 | ymcC | 0 | 0 | 0 | 0 | 0 | 0 | 0 | 0 | 0 | 0 |
| 608 | b0309 | 0 | 0 | 0 | 0 | 0 | 0 | 0 | 0 | 0 | 0 |
| 609 | ilvY | 0 | 0 | 0 | 0 | 0 | 0 | 0 | 0 | 0 | 0 |
| 610 | cytR | 0 | 0 | 0 | 0 | 0 | 0 | 0 | 0 | 0 | 0 |
| 611 | nupC | 0 | 0 | 0 | 0 | 0 | 0 | 0 | 0 | 0 | 0 |
| 612 | hcp | 0 | 0 | 0 | 0 | 0 | 0 | 0 | 0 | 0 | 0 |
| 613 | ytfQ | 0 | 0 | 0 | 0 | 0 | 0 | 0 | 0 | 0 | 0 |
| 614 | ytfR | 0 | 0 | 0 | 0 | 0 | 0 | 0 | 0 | 0 | 0 |
| 615 | yeaM | 0 | 0 | 0 | 0 | 0 | 0 | 0 | 0 | 0 | 0 |
| 616 | yeaL | 0 | 0 | 0 | 0 | 0 | 0 | 0 | 0 | 0 | 0 |
| 617 | rhtB | 0 | 0 | 0 | 0 | 0 | 0 | 0 | 0 | 0 | 0 |
| 618 | yeaH | 0 | 0 | 0 | 0 | 0 | 0 | 0 | 0 | 0 | 0 |
| 619 | yeaU | 21 | 0 | 7 | 0 | 46 | 0 | 11 | 0 | 28 | 0 |
| 620 | yeaT | 0 | 0 | 1 | 0 | 0 | 0 | 0 | 0 | 0 | 0 |
| 621 | yeaV | 0 | 0 | 0 | 0 | 0 | 0 | 0 | 0 | 0 | 0 |
| 622 | yeaP | 0 | 0 | 0 | 0 | 0 | 0 | 0 | 0 | 0 | 0 |
| 623 | yeaS | 0 | 0 | 0 | 0 | 0 | 0 | 0 | 0 | 0 | 0 |
| 624 | ytfB | 0 | 0 | 0 | 0 | 0 | 0 | 0 | 0 | 0 | 0 |
| 625 | ytfM | 0 | 0 | 0 | 0 | 0 | 0 | 0 | 0 | 0 | 0 |
| 626 | ytfL | 0 | 0 | 0 | 0 | 0 | 0 | 0 | 0 | 0 | 0 |
| 627 | nupX | 40 | 0 | 8 | 0 | 81 | 0 | 35 | 0 | 70 | 0 |
| 628 | ytfI | 0 | 0 | 0 | 0 | 0 | 0 | 0 | 0 | 0 | 0 |
| 629 | yeaX | 0 | 0 | 0 | 0 | 0 | 0 | 0 | 0 | 0 | 0 |
| 630 | blc | 0 | 0 | 0 | 0 | 0 | 0 | 0 | 0 | 0 | 0 |
| 631 | yeaZ | 0 | 0 | 0 | 0 | 0 | 0 | 0 | 0 | 0 | 0 |
| 632 | argZ_argQ | 11 | 0 | 2 | 0 | 14 | 0 | 3 | 0 | 0 | 0 |
| 633 | yjiY | 0 | 0 | 0 | 0 | 0 | 0 | 0 | 0 | 0 | 0 |
| 634 | ilvL | 0 | 0 | 0 | 0 | 0 | 0 | 0 | 0 | 0 | 0 |
| 635 | argP | 0 | 0 | 1 | 0 | 0 | 0 | 0 | 0 | 0 | 0 |
| 636 | intB | 0 | 0 | 0 | 0 | 0 | 0 | 0 | 0 | 0 | 0 |
| 637 | argU | 0 | 0 | 0 | 0 | 0 | 0 | 0 | 0 | 0 | 0 |
| 638 | yjiT | 0 | 0 | 0 | 0 | 0 | 0 | 0 | 0 | 0 | 0 |
| 639 | argW | 0 | 0 | 0 | 0 | 0 | 0 | 0 | 0 | 0 | 0 |
| 640 | yjiJ | 0 | 0 | 0 | 0 | 0 | 0 | 0 | 0 | 0 | 0 |
| 641 | argI | 0 | 0 | 0 | 0 | 0 | 0 | 1 | 0 | 0 | 0 |
| 642 | yjiH | 0 | 0 | 0 | 0 | 0 | 0 | 0 | 0 | 0 | 0 |
| 643 | yjiM | 0 | 0 | 0 | 0 | 0 | 0 | 0 | 0 | 0 | 0 |
| 644 | argA_amiC | 56 | 0 | 18 | 205 | 72 | 3 | 36 | 1 | 38 | 0 |
| 645 | argC | 0 | 0 | 0 | 0 | 0 | 0 | 0 | 0 | 0 | 0 |
| 646 | argD | 60 | 0 | 29 | 309 | 99 | 1 | 62 | 0 | 57 | 0 |
| 647 | argE | 0 | 0 | 0 | 0 | 0 | 0 | 0 | 0 | 0 | 0 |
| 648 | argF | 0 | 0 | 0 | 0 | 0 | 0 | 0 | 0 | 0 | 0 |
| 649 | argG | 0 | 0 | 0 | 0 | 0 | 0 | 0 | 0 | 0 | 0 |
| 650 | yggM | 0 | 0 | 0 | 0 | 0 | 0 | 0 | 0 | 0 | 0 |
| 651 | tyrS | 0 | 0 | 1 | 0 | 0 | 0 | 0 | 0 | 0 | 0 |
| 652 | yggH | 0 | 0 | 0 | 0 | 0 | 0 | 0 | 0 | 0 | 0 |
| 653 | htrA | 0 | 0 | 0 | 0 | 0 | 0 | 0 | 0 | 0 | 0 |
| 654 | iaaA | 0 | 0 | 0 | 0 | 0 | 0 | 0 | 0 | 0 | 0 |
| 655 | yggD | 0 | 0 | 0 | 0 | 0 | 0 | 0 | 0 | 0 | 0 |
| 656 | citT | 0 | 0 | 0 | 0 | 0 | 0 | 0 | 0 | 0 | 0 |
| 657 | ysgA | 0 | 0 | 0 | 0 | 0 | 0 | 0 | 0 | 0 | 0 |
| 658 | sstT | 0 | 0 | 0 | 0 | 0 | 0 | 0 | 0 | 0 | 0 |
| 659 | yiiF | 0 | 0 | 0 | 0 | 0 | 0 | 1 | 0 | 0 | 0 |

| | | | | | | | | | | | |
|---|---|---|---|---|---|---|---|---|---|---|---|
| 660 | yiiQ | 0 | 0 | 0 | 0 | 0 | 0 | 0 | 0 | 0 | 0 |
| 661 | yiiP | 168 | 0 | 31 | 1 | 286 | 0 | 104 | 0 | 51 | 0 |
| 662 | citC | 0 | 0 | 0 | 0 | 0 | 0 | 0 | 0 | 0 | 0 |
| 663 | yiiU | 0 | 0 | 0 | 0 | 0 | 0 | 0 | 0 | 0 | 0 |
| 664 | tyrV | 0 | 0 | 0 | 0 | 0 | 0 | 0 | 0 | 0 | 0 |
| 665 | degQ | 0 | 0 | 1 | 0 | 0 | 0 | 0 | 0 | 0 | 0 |
| 666 | yphH | 0 | 0 | 0 | 0 | 0 | 0 | 0 | 0 | 0 | 0 |
| 667 | hybO | 0 | 0 | 0 | 0 | 0 | 0 | 0 | 0 | 0 | 0 |
| 668 | yphF | 0 | 0 | 0 | 0 | 0 | 0 | 0 | 0 | 0 | 0 |
| 669 | accB | 0 | 0 | 0 | 0 | 0 | 0 | 0 | 0 | 0 | 0 |
| 670 | iap | 0 | 0 | 0 | 1 | 0 | 0 | 0 | 0 | 0 | 0 |
| 671 | smpA | 0 | 0 | 0 | 0 | 0 | 0 | 0 | 0 | 0 | 0 |
| 672 | smpB | 0 | 0 | 0 | 0 | 0 | 0 | 0 | 0 | 0 | 0 |
| 673 | serU | 0 | 0 | 0 | 0 | 0 | 0 | 0 | 0 | 0 | 0 |
| 674 | pnp | 0 | 0 | 0 | 0 | 0 | 0 | 0 | 0 | 0 | 0 |
| 675 | serB | 0 | 0 | 0 | 0 | 0 | 0 | 0 | 0 | 0 | 0 |
| 676 | serC | 0 | 0 | 0 | 0 | 0 | 0 | 0 | 0 | 0 | 0 |
| 677 | serA | 0 | 0 | 0 | 0 | 0 | 0 | 0 | 0 | 0 | 0 |
| 678 | alaS | 0 | 0 | 0 | 0 | 0 | 0 | 0 | 0 | 0 | 0 |
| 679 | yqhH | 0 | 0 | 0 | 0 | 0 | 0 | 0 | 0 | 0 | 0 |
| 680 | rrmA | 0 | 0 | 0 | 0 | 0 | 0 | 0 | 0 | 0 | 0 |
| 681 | aroL | 0 | 0 | 0 | 0 | 0 | 0 | 0 | 0 | 0 | 0 |
| 682 | yjeB | 6 | 6 | 3 | 3 | 21 | 13 | 3 | 5 | 0 | 2 |
| 683 | sulA | 0 | 0 | 0 | 0 | 0 | 0 | 0 | 0 | 0 | 0 |
| 684 | gyrB | 0 | 0 | 0 | 0 | 0 | 0 | 0 | 0 | 0 | 0 |
| 685 | yhiR | 0 | 0 | 0 | 0 | 0 | 0 | 0 | 0 | 0 | 0 |
| 686 | sohB | 0 | 0 | 0 | 0 | 0 | 0 | 0 | 0 | 0 | 0 |
| 687 | sohA | 0 | 0 | 0 | 0 | 0 | 0 | 0 | 1 | 0 | 0 |
| 688 | brnQ | 0 | 0 | 0 | 0 | 0 | 0 | 0 | 0 | 0 | 0 |
| 689 | betT | 0 | 0 | 0 | 0 | 0 | 0 | 0 | 0 | 0 | 0 |
| 690 | mrp | 0 | 0 | 0 | 0 | 0 | 0 | 0 | 0 | 0 | 0 |
| 691 | mrr | 0 | 0 | 0 | 0 | 0 | 0 | 0 | 0 | 0 | 0 |
| 692 | ykgC | 0 | 0 | 0 | 0 | 0 | 0 | 0 | 0 | 0 | 0 |
| 693 | ykgE | 0 | 0 | 0 | 0 | 0 | 0 | 0 | 0 | 0 | 0 |
| 694 | ybfF | 0 | 0 | 0 | 0 | 0 | 0 | 0 | 0 | 0 | 0 |
| 695 | ybfE | 0 | 0 | 0 | 0 | 0 | 0 | 0 | 0 | 0 | 0 |
| 696 | ykgF | 0 | 0 | 0 | 0 | 0 | 0 | 0 | 0 | 0 | 0 |
| 697 | ykgI | 0 | 0 | 0 | 0 | 0 | 0 | 0 | 0 | 0 | 0 |
| 698 | ykgH | 0 | 0 | 0 | 0 | 0 | 0 | 0 | 0 | 0 | 0 |
| 699 | ykgJ | 0 | 0 | 0 | 0 | 0 | 0 | 0 | 0 | 0 | 0 |
| 700 | ykgM | 0 | 0 | 0 | 0 | 0 | 0 | 0 | 0 | 0 | 0 |
| 701 | grxC | 0 | 0 | 0 | 0 | 0 | 0 | 0 | 0 | 0 | 0 |
| 702 | grxB | 51 | 0 | 28 | 0 | 88 | 0 | 33 | 0 | 0 | 0 |
| 703 | dicB | 0 | 0 | 0 | 0 | 0 | 0 | 0 | 0 | 0 | 0 |
| 704 | elaD | 0 | 0 | 0 | 0 | 0 | 0 | 0 | 0 | 0 | 0 |
| 705 | elaA | 0 | 0 | 0 | 0 | 0 | 0 | 0 | 0 | 0 | 0 |
| 706 | elaB | 0 | 0 | 0 | 0 | 0 | 0 | 0 | 0 | 0 | 0 |
| 707 | hcaT | 0 | 0 | 0 | 0 | 0 | 0 | 0 | 0 | 0 | 0 |
| 708 | osmC | 0 | 0 | 0 | 0 | 0 | 0 | 0 | 0 | 0 | 0 |
| 709 | osmB | 0 | 0 | 0 | 0 | 0 | 0 | 0 | 0 | 0 | 0 |
| 710 | osmE | 0 | 0 | 1 | 0 | 0 | 0 | 0 | 0 | 0 | 0 |
| 711 | hcaR | 0 | 0 | 0 | 0 | 0 | 0 | 0 | 0 | 0 | 0 |
| 712 | leuA | 0 | 0 | 0 | 0 | 0 | 0 | 0 | 0 | 0 | 0 |
| 713 | asnV | 0 | 0 | 0 | 0 | 0 | 0 | 0 | 0 | 0 | 0 |
| 714 | asnS | 0 | 0 | 1 | 0 | 0 | 0 | 0 | 0 | 0 | 0 |
| 715 | leuS | 0 | 0 | 0 | 0 | 0 | 0 | 0 | 0 | 0 | 0 |
| 716 | asnC | 0 | 0 | 0 | 0 | 0 | 0 | 0 | 0 | 0 | 0 |
| 717 | asnA | 0 | 0 | 0 | 0 | 0 | 0 | 0 | 0 | 0 | 0 |
| 718 | ydbH | 0 | 0 | 0 | 0 | 0 | 0 | 0 | 0 | 0 | 0 |
| 719 | ydbC | 0 | 0 | 0 | 0 | 0 | 0 | 0 | 0 | 0 | 0 |
| 720 | ydbD | 0 | 0 | 0 | 0 | 0 | 0 | 0 | 0 | 0 | 0 |
| 721 | dksA | 0 | 0 | 0 | 0 | 0 | 0 | 0 | 0 | 0 | 0 |
| 722 | b1228 | 0 | 0 | 0 | 0 | 0 | 0 | 0 | 0 | 0 | 0 |
| 723 | hisL | 0 | 0 | 0 | 0 | 0 | 0 | 0 | 0 | 0 | 0 |
| 724 | hisJ | 0 | 0 | 0 | 0 | 0 | 0 | 0 | 0 | 0 | 0 |
| 725 | rmf | 0 | 0 | 0 | 0 | 0 | 0 | 0 | 0 | 0 | 0 |

| | | | | | | | | | | | |
|---|---|---|---|---|---|---|---|---|---|---|---|
| 726 | hisQ | 0 | 0 | 0 | 0 | 0 | 0 | 0 | 0 | 0 | 0 |
| 727 | hisS | 0 | 0 | 0 | 0 | 0 | 0 | 0 | 0 | 0 | 0 |
| 728 | ypdA | 0 | 0 | 0 | 0 | 0 | 0 | 0 | 0 | 0 | 0 |
| 729 | fadL | 0 | 0 | 0 | 0 | 0 | 0 | 0 | 0 | 0 | 0 |
| 730 | ynbA | 68 | 1 | 39 | 3 | 114 | 1 | 62 | 0 | 174 | 0 |
| 731 | manX | 0 | 0 | 0 | 0 | 0 | 0 | 0 | 0 | 0 | 0 |
| 732 | manY | 0 | 0 | 0 | 0 | 0 | 0 | 0 | 0 | 0 | 0 |
| 733 | yajR | 0 | 0 | 0 | 0 | 0 | 0 | 0 | 0 | 0 | 0 |
| 734 | yibK | 0 | 1 | 0 | 0 | 0 | 0 | 0 | 0 | 0 | 0 |
| 735 | yajO | 0 | 0 | 0 | 0 | 0 | 0 | 0 | 0 | 0 | 0 |
| 736 | yajB | 0 | 0 | 0 | 0 | 0 | 0 | 0 | 0 | 0 | 0 |
| 737 | yajC | 0 | 0 | 0 | 0 | 0 | 0 | 0 | 0 | 0 | 0 |
| 738 | b0374 | 0 | 0 | 0 | 0 | 0 | 0 | 0 | 0 | 0 | 0 |
| 739 | yajG | 0 | 0 | 0 | 0 | 0 | 0 | 0 | 0 | 0 | 0 |
| 740 | ycbK | 0 | 0 | 0 | 0 | 0 | 0 | 0 | 0 | 0 | 0 |
| 741 | ycbB | 0 | 0 | 0 | 0 | 0 | 0 | 0 | 0 | 0 | 0 |
| 742 | ycbC | 0 | 0 | 0 | 0 | 0 | 0 | 0 | 0 | 0 | 0 |
| 743 | lpxC | 0 | 0 | 0 | 0 | 0 | 0 | 0 | 0 | 0 | 0 |
| 744 | ycbZ | 1 | 0 | 0 | 0 | 1 | 0 | 0 | 0 | 0 | 0 |
| 745 | csiE | 0 | 0 | 0 | 0 | 0 | 0 | 0 | 0 | 0 | 0 |
| 746 | csiD | 0 | 0 | 0 | 0 | 0 | 0 | 0 | 0 | 0 | 0 |
| 747 | ycbR | 0 | 0 | 0 | 0 | 0 | 0 | 0 | 0 | 0 | 0 |
| 748 | ycbW | 0 | 0 | 0 | 0 | 0 | 0 | 0 | 1 | 0 | 0 |
| 749 | glcB | 0 | 0 | 0 | 1 | 0 | 0 | 0 | 0 | 0 | 0 |
| 750 | glcC | 0 | 0 | 0 | 0 | 0 | 0 | 0 | 0 | 0 | 0 |
| 751 | artP | 0 | 0 | 0 | 0 | 0 | 0 | 0 | 0 | 0 | 0 |
| 752 | marC_marR | 0 | 0 | 0 | 0 | 0 | 0 | 0 | 0 | 0 | 0 |
| 753 | rluE | 0 | 0 | 0 | 0 | 0 | 0 | 0 | 0 | 0 | 0 |
| 754 | rluD | 0 | 0 | 0 | 0 | 0 | 0 | 0 | 0 | 0 | 0 |
| 755 | rluF | 23 | 1 | 1 | 0 | 32 | 0 | 60 | 0 | 0 | 0 |
| 756 | rluC | 0 | 0 | 0 | 0 | 0 | 0 | 0 | 0 | 0 | 0 |
| 757 | rluB | 0 | 0 | 0 | 0 | 0 | 0 | 0 | 0 | 0 | 0 |
| 758 | b0373 | 0 | 0 | 0 | 0 | 0 | 0 | 0 | 0 | 0 | 0 |
| 759 | helD | 0 | 0 | 0 | 0 | 0 | 0 | 0 | 0 | 0 | 0 |
| 760 | yqiC | 0 | 0 | 0 | 0 | 0 | 0 | 0 | 0 | 0 | 0 |
| 761 | polB | 0 | 0 | 0 | 0 | 0 | 0 | 1 | 0 | 0 | 0 |
| 762 | polA | 0 | 0 | 1 | 0 | 0 | 0 | 0 | 0 | 0 | 0 |
| 763 | yqiJ | 0 | 0 | 0 | 0 | 0 | 0 | 0 | 0 | 0 | 0 |
| 764 | ecpD | 0 | 0 | 0 | 0 | 0 | 0 | 0 | 0 | 0 | 0 |
| 765 | yjhF | 0 | 0 | 0 | 0 | 0 | 0 | 0 | 0 | 0 | 0 |
| 766 | yjhI | 0 | 0 | 0 | 0 | 0 | 0 | 0 | 0 | 0 | 0 |
| 767 | yjhU | 0 | 0 | 0 | 0 | 0 | 0 | 0 | 0 | 0 | 0 |
| 768 | yjhS | 0 | 0 | 0 | 0 | 0 | 0 | 0 | 0 | 0 | 0 |
| 769 | hda | 0 | 0 | 0 | 0 | 0 | 0 | 0 | 0 | 0 | 0 |
| 770 | pepN | 0 | 0 | 0 | 0 | 0 | 0 | 0 | 0 | 0 | 0 |
| 771 | pepB | 0 | 0 | 0 | 0 | 0 | 0 | 0 | 0 | 0 | 0 |
| 772 | pepE | 0 | 0 | 0 | 0 | 0 | 0 | 0 | 0 | 0 | 0 |
| 773 | pepD | 0 | 0 | 0 | 0 | 0 | 0 | 0 | 0 | 0 | 0 |
| 774 | mqo | 0 | 0 | 0 | 0 | 0 | 0 | 1 | 0 | 0 | 0 |
| 775 | pepQ | 2 | 0 | 0 | 0 | 1 | 0 | 0 | 0 | 0 | 0 |
| 776 | pepT | 0 | 0 | 0 | 0 | 0 | 0 | 0 | 0 | 0 | 0 |
| 777 | yidC | 0 | 0 | 0 | 0 | 1 | 0 | 0 | 0 | 0 | 0 |
| 778 | ybeZ | 0 | 0 | 0 | 0 | 0 | 0 | 0 | 0 | 0 | 0 |
| 779 | yidE | 0 | 0 | 0 | 0 | 0 | 0 | 0 | 0 | 0 | 0 |
| 780 | yidK | 19 | 0 | 23 | 0 | 32 | 0 | 6 | 0 | 0 | 0 |
| 781 | yidH | 0 | 0 | 0 | 0 | 0 | 0 | 0 | 0 | 0 | 0 |
| 782 | ybeU | 0 | 0 | 0 | 0 | 0 | 0 | 0 | 0 | 0 | 0 |
| 783 | ybeR | 0 | 0 | 0 | 0 | 0 | 0 | 0 | 0 | 0 | 0 |
| 784 | yidL | 20 | 0 | 24 | 0 | 37 | 0 | 7 | 0 | 0 | 0 |
| 785 | ybeQ | 0 | 0 | 0 | 0 | 0 | 0 | 0 | 0 | 0 | 0 |
| 786 | yidR | 0 | 0 | 0 | 1 | 0 | 0 | 0 | 0 | 0 | 0 |
| 787 | yidS | 0 | 0 | 0 | 0 | 0 | 0 | 0 | 0 | 0 | 0 |
| 788 | ybeL | 0 | 0 | 0 | 0 | 0 | 0 | 0 | 0 | 0 | 0 |
| 789 | ybeF | 0 | 0 | 0 | 0 | 0 | 0 | 0 | 0 | 0 | 0 |
| 790 | yidX | 0 | 0 | 0 | 0 | 0 | 0 | 0 | 0 | 0 | 0 |
| 791 | ybeB | 0 | 1 | 0 | 0 | 0 | 0 | 0 | 0 | 0 | 0 |

| | | | | | | | | | | | |
|---|---|---|---|---|---|---|---|---|---|---|---|
| 792 | chaB | 0 | 0 | 0 | 0 | 0 | 0 | 0 | 0 | 0 | 0 |
| 793 | cvpA | 0 | 0 | 0 | 0 | 0 | 0 | 1 | 0 | 0 | 0 |
| 794 | ycgX | 0 | 0 | 0 | 0 | 0 | 0 | 0 | 0 | 0 | 0 |
| 795 | kdgR | 0 | 0 | 0 | 0 | 0 | 0 | 0 | 0 | 0 | 0 |
| 796 | lpxP | 0 | 0 | 0 | 0 | 0 | 0 | 0 | 0 | 0 | 0 |
| 797 | ssuE | 0 | 0 | 0 | 0 | 0 | 0 | 0 | 0 | 0 | 0 |
| 798 | hyaA | 0 | 0 | 0 | 0 | 0 | 0 | 0 | 0 | 0 | 0 |
| 799 | yaaH | 0 | 0 | 1 | 0 | 0 | 0 | 0 | 0 | 0 | 0 |
| 800 | fimD | 0 | 0 | 0 | 0 | 0 | 0 | 0 | 0 | 0 | 0 |
| 801 | cca | 0 | 0 | 0 | 0 | 0 | 0 | 0 | 0 | 0 | 0 |
| 802 | ylcE | 0 | 0 | 0 | 0 | 0 | 0 | 0 | 0 | 0 | 0 |
| 803 | fumB | 84 | 0 | 23 | 1 | 171 | 0 | 78 | 0 | 53 | 0 |
| 804 | ycgL | 0 | 0 | 0 | 0 | 0 | 0 | 0 | 0 | 0 | 0 |
| 805 | rrnB | 0 | 0 | 0 | 0 | 0 | 0 | 0 | 0 | 0 | 0 |
| 806 | rrnC | 0 | 0 | 0 | 0 | 0 | 0 | 0 | 0 | 0 | 0 |
| 807 | rrnA | 0 | 0 | 0 | 0 | 0 | 0 | 0 | 0 | 0 | 0 |
| 808 | rrnG | 0 | 0 | 0 | 0 | 0 | 0 | 0 | 0 | 0 | 0 |
| 809 | rrnD | 0 | 0 | 0 | 0 | 0 | 0 | 0 | 0 | 0 | 0 |
| 810 | amn | 0 | 0 | 0 | 0 | 0 | 0 | 0 | 0 | 0 | 0 |
| 811 | rrnH | 0 | 0 | 0 | 0 | 0 | 0 | 0 | 0 | 0 | 0 |
| 812 | psd | 0 | 0 | 0 | 0 | 0 | 0 | 0 | 0 | 0 | 0 |
| 813 | b1142 | 0 | 0 | 0 | 0 | 0 | 0 | 0 | 0 | 0 | 0 |
| 814 | mog | 0 | 0 | 0 | 0 | 0 | 0 | 0 | 0 | 0 | 0 |
| 815 | ycgE | 0 | 0 | 0 | 0 | 0 | 0 | 0 | 0 | 0 | 0 |
| 816 | wrbA | 0 | 0 | 0 | 0 | 0 | 0 | 1 | 0 | 0 | 0 |
| 817 | fldA | 0 | 0 | 0 | 0 | 0 | 0 | 0 | 0 | 0 | 0 |
| 818 | ynaI | 7 | 0 | 0 | 0 | 6 | 0 | 1 | 0 | 0 | 0 |
| 819 | b1367 | 0 | 0 | 0 | 0 | 0 | 0 | 0 | 0 | 0 | 0 |
| 820 | ftsJ | 0 | 0 | 0 | 0 | 0 | 0 | 0 | 0 | 0 | 0 |
| 821 | yccR | 0 | 0 | 0 | 0 | 0 | 0 | 0 | 0 | 0 | 0 |
| 822 | b1331 | 0 | 0 | 0 | 0 | 0 | 0 | 0 | 0 | 0 | 0 |
| 823 | yccE | 0 | 0 | 0 | 0 | 0 | 0 | 0 | 0 | 0 | 0 |
| 824 | yccF | 0 | 0 | 0 | 0 | 0 | 0 | 0 | 0 | 0 | 0 |
| 825 | yccA | 0 | 0 | 0 | 0 | 0 | 0 | 0 | 0 | 0 | 0 |
| 826 | b1966 | 0 | 0 | 0 | 0 | 0 | 0 | 0 | 0 | 0 | 0 |
| 827 | b1788 | 0 | 0 | 0 | 0 | 0 | 0 | 0 | 0 | 0 | 0 |
| 828 | yccM | 0 | 0 | 0 | 0 | 0 | 0 | 0 | 0 | 0 | 0 |
| 829 | cydD | 0 | 1 | 0 | 0 | 0 | 0 | 0 | 0 | 0 | 0 |
| 830 | ushA | 0 | 0 | 0 | 0 | 0 | 0 | 0 | 0 | 0 | 0 |
| 831 | rmuC | 0 | 0 | 0 | 0 | 0 | 0 | 0 | 0 | 0 | 0 |
| 832 | dpiB | 0 | 0 | 0 | 0 | 0 | 0 | 0 | 0 | 0 | 0 |
| 833 | dapA | 0 | 0 | 0 | 0 | 0 | 0 | 0 | 0 | 0 | 0 |
| 834 | mngR | 0 | 0 | 0 | 0 | 0 | 0 | 0 | 0 | 0 | 0 |
| 835 | lolB | 0 | 0 | 0 | 0 | 0 | 0 | 0 | 0 | 0 | 0 |
| 836 | b0360 | 0 | 0 | 0 | 0 | 0 | 0 | 0 | 0 | 0 | 0 |
| 837 | rnhA | 0 | 0 | 0 | 0 | 0 | 0 | 0 | 0 | 0 | 0 |
| 838 | yegE | 0 | 0 | 1 | 0 | 0 | 0 | 0 | 0 | 0 | 0 |
| 839 | garD | 0 | 0 | 0 | 0 | 0 | 0 | 0 | 0 | 0 | 0 |
| 840 | ybiU | 0 | 0 | 0 | 0 | 0 | 0 | 0 | 0 | 0 | 0 |
| 841 | lon | 0 | 0 | 0 | 0 | 0 | 0 | 0 | 0 | 0 | 0 |
| 842 | nrfA | 0 | 0 | 0 | 0 | 0 | 0 | 0 | 0 | 0 | 0 |
| 843 | yegP | 0 | 0 | 0 | 0 | 0 | 0 | 0 | 0 | 0 | 0 |
| 844 | yegW | 0 | 0 | 0 | 0 | 0 | 0 | 0 | 0 | 0 | 0 |
| 845 | yegT | 41 | 0 | 9 | 97 | 67 | 7 | 44 | 250 | 27 | 0 |
| 846 | ygaD | 0 | 0 | 0 | 0 | 0 | 0 | 0 | 0 | 0 | 0 |
| 847 | ygaZ | 0 | 0 | 0 | 0 | 0 | 0 | 0 | 0 | 0 | 0 |
| 848 | zntA | 0 | 0 | 0 | 0 | 0 | 0 | 0 | 0 | 0 | 0 |
| 849 | soxR | 0 | 0 | 0 | 0 | 0 | 0 | 0 | 0 | 0 | 0 |
| 850 | soxS | 0 | 0 | 0 | 0 | 0 | 0 | 0 | 0 | 0 | 0 |
| 851 | ygaQ | 0 | 0 | 0 | 0 | 0 | 0 | 0 | 0 | 0 | 0 |
| 852 | aceK | 0 | 0 | 0 | 0 | 0 | 0 | 0 | 0 | 0 | 0 |
| 853 | ssrA | 0 | 0 | 0 | 0 | 0 | 0 | 0 | 0 | 0 | 0 |
| 854 | aceB | 0 | 0 | 0 | 0 | 0 | 0 | 0 | 0 | 0 | 0 |
| 855 | fryB | 0 | 0 | 0 | 0 | 0 | 0 | 0 | 0 | 1 | 0 |
| 856 | ypjD | 0 | 0 | 0 | 0 | 0 | 0 | 0 | 1 | 0 | 0 |
| 857 | ypjA | 0 | 0 | 0 | 0 | 0 | 0 | 0 | 0 | 0 | 0 |

| | | | | | | | | | | | |
|---|---|---|---|---|---|---|---|---|---|---|---|
| 858 | rihC | 29 | 0 | 4 | 0 | 54 | 1 | 7 | 0 | 38 | 0 |
| 859 | rihA | 0 | 0 | 0 | 0 | 0 | 0 | 0 | 0 | 0 | 0 |
| 860 | hsdM | 0 | 0 | 0 | 0 | 0 | 0 | 0 | 0 | 0 | 0 |
| 861 | lysA | 0 | 0 | 0 | 0 | 0 | 0 | 0 | 0 | 0 | 0 |
| 862 | hsdR | 0 | 0 | 0 | 0 | 0 | 0 | 0 | 0 | 0 | 0 |
| 863 | lysR | 0 | 0 | 0 | 0 | 0 | 0 | 0 | 0 | 0 | 0 |
| 864 | lysU | 0 | 0 | 0 | 0 | 0 | 0 | 0 | 0 | 0 | 0 |
| 865 | eaeH | 0 | 0 | 0 | 0 | 0 | 0 | 0 | 0 | 0 | 0 |
| 866 | ansA | 0 | 0 | 0 | 0 | 0 | 0 | 0 | 0 | 0 | 0 |
| 867 | ansB | 0 | 0 | 0 | 0 | 0 | 0 | 1 | 0 | 0 | 0 |
| 868 | mgsA | 0 | 0 | 0 | 0 | 0 | 0 | 0 | 0 | 0 | 0 |
| 869 | yqfA | 0 | 0 | 0 | 0 | 0 | 0 | 0 | 0 | 0 | 0 |
| 870 | norR | 0 | 0 | 0 | 0 | 0 | 0 | 0 | 0 | 0 | 0 |
| 871 | norV | 0 | 0 | 0 | 0 | 0 | 0 | 0 | 0 | 0 | 0 |
| 872 | pth | 0 | 0 | 0 | 0 | 0 | 0 | 0 | 1 | 0 | 0 |
| 873 | lacA | 0 | 0 | 0 | 0 | 0 | 0 | 0 | 0 | 0 | 0 |
| 874 | lacI | 0 | 0 | 0 | 0 | 0 | 0 | 0 | 0 | 0 | 0 |
| 875 | lacY | 0 | 0 | 0 | 1 | 0 | 0 | 0 | 0 | 0 | 0 |
| 876 | lacZ | 0 | 0 | 0 | 0 | 1 | 0 | 0 | 1 | 0 | 0 |
| 877 | atpI | 0 | 0 | 0 | 0 | 0 | 0 | 0 | 0 | 0 | 0 |
| 878 | sanA | 0 | 0 | 0 | 0 | 0 | 0 | 0 | 0 | 0 | 0 |
| 879 | pfkB | 0 | 0 | 0 | 0 | 0 | 0 | 0 | 0 | 0 | 0 |
| 880 | pfkA | 0 | 0 | 0 | 0 | 0 | 0 | 0 | 0 | 0 | 0 |
| 881 | hemH | 0 | 0 | 0 | 0 | 0 | 0 | 0 | 0 | 0 | 0 |
| 882 | hemN | 0 | 0 | 0 | 0 | 0 | 0 | 0 | 0 | 0 | 0 |
| 883 | yieH | 0 | 0 | 0 | 0 | 0 | 0 | 0 | 0 | 0 | 0 |
| 884 | hemL | 0 | 0 | 0 | 0 | 0 | 0 | 0 | 0 | 0 | 0 |
| 885 | rhaD | 0 | 0 | 0 | 0 | 0 | 0 | 0 | 0 | 0 | 0 |
| 886 | hemB | 0 | 0 | 0 | 0 | 0 | 0 | 0 | 0 | 0 | 0 |
| 887 | yieG | 0 | 0 | 0 | 0 | 0 | 0 | 0 | 0 | 0 | 0 |
| 888 | ybdR | 0 | 0 | 0 | 0 | 0 | 0 | 0 | 0 | 0 | 0 |
| 889 | hemG | 0 | 0 | 0 | 0 | 0 | 0 | 0 | 0 | 0 | 0 |
| 890 | ybdH | 0 | 0 | 0 | 0 | 0 | 0 | 0 | 0 | 0 | 0 |
| 891 | ybdK | 0 | 0 | 0 | 0 | 0 | 0 | 0 | 0 | 0 | 0 |
| 892 | ybdL | 0 | 0 | 0 | 0 | 0 | 0 | 0 | 0 | 0 | 0 |
| 893 | ybdN | 0 | 0 | 0 | 0 | 0 | 0 | 0 | 0 | 0 | 0 |
| 894 | ykiA | 0 | 0 | 0 | 0 | 0 | 0 | 0 | 0 | 0 | 0 |
| 895 | ybdF | 0 | 0 | 0 | 0 | 0 | 0 | 0 | 0 | 0 | 0 |
| 896 | cueR | 28 | 0 | 6 | 2 | 37 | 0 | 21 | 0 | 44 | 0 |
| 897 | frlA | 0 | 0 | 0 | 0 | 0 | 0 | 0 | 0 | 0 | 0 |
| 898 | nanA | 0 | 0 | 0 | 0 | 0 | 0 | 0 | 0 | 0 | 0 |
| 899 | cueO | 0 | 0 | 0 | 0 | 0 | 0 | 0 | 0 | 0 | 0 |
| 900 | nanR | 0 | 0 | 0 | 0 | 0 | 0 | 0 | 0 | 0 | 0 |
| 901 | nanT | 0 | 0 | 0 | 0 | 0 | 0 | 0 | 0 | 0 | 0 |
| 902 | trmU | 0 | 0 | 0 | 0 | 0 | 0 | 0 | 0 | 0 | 0 |
| 903 | trmA | 0 | 0 | 0 | 0 | 0 | 0 | 0 | 0 | 0 | 0 |
| 904 | trmE | 0 | 0 | 0 | 0 | 0 | 0 | 0 | 0 | 0 | 0 |
| 905 | dgoR | 0 | 0 | 0 | 0 | 0 | 0 | 0 | 0 | 0 | 0 |
| 906 | ggt | 0 | 0 | 0 | 0 | 0 | 0 | 0 | 0 | 0 | 0 |
| 907 | ybjS | 0 | 0 | 0 | 0 | 0 | 0 | 0 | 0 | 0 | 0 |
| 908 | ybjQ | 0 | 0 | 0 | 0 | 0 | 0 | 0 | 0 | 0 | 0 |
| 909 | ybjP | 0 | 0 | 0 | 0 | 0 | 0 | 0 | 0 | 0 | 0 |
| 910 | ybjG | 0 | 0 | 0 | 0 | 0 | 0 | 0 | 0 | 0 | 0 |
| 911 | ybjE | 0 | 0 | 0 | 0 | 0 | 0 | 1 | 0 | 0 | 0 |
| 912 | ybjC | 0 | 0 | 0 | 0 | 0 | 0 | 0 | 0 | 0 | 0 |
| 913 | ybjO | 0 | 0 | 0 | 0 | 0 | 0 | 0 | 0 | 0 | 0 |
| 914 | ybjN | 0 | 0 | 0 | 0 | 0 | 0 | 0 | 0 | 0 | 0 |
| 915 | ybjM | 0 | 0 | 0 | 0 | 0 | 0 | 0 | 0 | 0 | 0 |
| 916 | ybjL | 0 | 0 | 0 | 0 | 0 | 0 | 0 | 0 | 0 | 0 |
| 917 | ybjK | 27 | 0 | 10 | 0 | 48 | 0 | 27 | 0 | 22 | 0 |
| 918 | ybjJ | 1 | 0 | 0 | 0 | 0 | 0 | 0 | 0 | 0 | 0 |
| 919 | ybjH | 0 | 0 | 0 | 0 | 0 | 0 | 0 | 0 | 0 | 0 |
| 920 | ylbE | 0 | 0 | 0 | 0 | 0 | 0 | 0 | 0 | 0 | 0 |
| 921 | yjjY | 0 | 0 | 0 | 0 | 0 | 0 | 0 | 0 | 0 | 0 |
| 922 | ylbG | 0 | 0 | 0 | 0 | 0 | 0 | 0 | 0 | 0 | 0 |
| 923 | ylbF | 0 | 0 | 0 | 0 | 0 | 0 | 0 | 0 | 0 | 0 |

| | | | | | | | | | | | |
|---|---|---|---|---|---|---|---|---|---|---|---|
| 924 | fklB | 0 | 0 | 0 | 0 | 0 | 0 | 0 | 0 | 0 | 0 |
| 925 | valV | 0 | 0 | 0 | 0 | 0 | 0 | 0 | 0 | 0 | 0 |
| 926 | valT | 0 | 0 | 1 | 0 | 0 | 0 | 0 | 1 | 0 | 0 |
| 927 | zraS | 0 | 0 | 0 | 0 | 0 | 0 | 0 | 0 | 0 | 0 |
| 928 | zraP | 0 | 0 | 0 | 0 | 0 | 0 | 0 | 0 | 0 | 0 |
| 929 | pntA | 0 | 0 | 0 | 0 | 0 | 0 | 0 | 0 | 0 | 0 |
| 930 | aphA | 0 | 0 | 0 | 0 | 0 | 0 | 0 | 0 | 0 | 0 |
| 931 | yjjK | 0 | 0 | 0 | 0 | 0 | 0 | 0 | 0 | 0 | 0 |
| 932 | yjjJ | 0 | 0 | 0 | 0 | 0 | 0 | 0 | 0 | 0 | 0 |
| 933 | tpiA | 0 | 0 | 0 | 0 | 0 | 0 | 0 | 0 | 0 | 0 |
| 934 | pldB | 0 | 0 | 0 | 0 | 0 | 0 | 0 | 0 | 0 | 0 |
| 935 | malX | 0 | 0 | 0 | 0 | 0 | 0 | 0 | 0 | 0 | 0 |
| 936 | malS | 0 | 0 | 0 | 0 | 0 | 0 | 0 | 0 | 0 | 0 |
| 937 | malI | 0 | 0 | 0 | 0 | 0 | 0 | 0 | 0 | 0 | 0 |
| 938 | malF | 0 | 0 | 0 | 0 | 0 | 0 | 0 | 0 | 0 | 0 |
| 939 | malE | 0 | 0 | 0 | 0 | 0 | 0 | 0 | 0 | 0 | 0 |
| 940 | crcB | 0 | 0 | 0 | 0 | 0 | 0 | 0 | 0 | 0 | 0 |
| 941 | yahD | 0 | 0 | 0 | 0 | 0 | 0 | 0 | 0 | 0 | 0 |
| 942 | yahB | 0 | 0 | 0 | 0 | 0 | 0 | 0 | 0 | 0 | 0 |
| 943 | yahC | 0 | 0 | 0 | 0 | 0 | 0 | 0 | 0 | 0 | 0 |
| 944 | yahN | 2 | 0 | 0 | 0 | 1 | 0 | 1 | 21 | 1 | 21 |
| 945 | yahO | 0 | 0 | 0 | 0 | 0 | 0 | 0 | 0 | 0 | 0 |
| 946 | ycdZ | 0 | 0 | 0 | 0 | 0 | 0 | 0 | 0 | 0 | 0 |
| 947 | yahM | 0 | 0 | 0 | 0 | 0 | 0 | 0 | 0 | 0 | 0 |
| 948 | yahK | 0 | 0 | 0 | 0 | 0 | 0 | 0 | 0 | 0 | 0 |
| 949 | yahI | 0 | 0 | 0 | 0 | 0 | 0 | 0 | 0 | 0 | 0 |
| 950 | ycdC | 0 | 0 | 0 | 0 | 0 | 0 | 0 | 0 | 0 | 0 |
| 951 | ibpB | 0 | 0 | 0 | 0 | 0 | 0 | 0 | 0 | 0 | 0 |
| 952 | eutR | 0 | 0 | 0 | 0 | 0 | 0 | 0 | 0 | 0 | 0 |
| 953 | eutS | 0 | 0 | 0 | 0 | 0 | 0 | 0 | 0 | 0 | 0 |
| 954 | csgC | 0 | 0 | 0 | 0 | 0 | 0 | 0 | 0 | 0 | 0 |
| 955 | ycdM | 0 | 0 | 0 | 0 | 0 | 0 | 0 | 0 | 0 | 0 |
| 956 | ycdN | 0 | 0 | 0 | 0 | 0 | 0 | 0 | 0 | 0 | 0 |
| 957 | ycdO | 0 | 0 | 0 | 0 | 0 | 0 | 0 | 0 | 0 | 0 |
| 958 | dsrA | 0 | 0 | 0 | 0 | 0 | 0 | 0 | 0 | 0 | 0 |
| 959 | sseA | 0 | 0 | 0 | 0 | 0 | 0 | 0 | 0 | 0 | 0 |
| 960 | sseB | 0 | 0 | 0 | 0 | 0 | 0 | 0 | 0 | 0 | 0 |
| 961 | mntR | 0 | 0 | 0 | 0 | 0 | 0 | 0 | 0 | 0 | 0 |
| 962 | yqgA | 9 | 0 | 8 | 0 | 22 | 0 | 6 | 0 | 0 | 0 |
| 963 | secA | 0 | 0 | 0 | 0 | 0 | 0 | 0 | 0 | 1 | 0 |
| 964 | secB | 0 | 0 | 0 | 0 | 0 | 0 | 0 | 0 | 0 | 0 |
| 965 | secE | 0 | 1 | 0 | 0 | 0 | 0 | 1 | 0 | 0 | 0 |
| 966 | secM | 0 | 0 | 0 | 0 | 0 | 0 | 0 | 0 | 0 | 0 |
| 967 | ymfI | 0 | 0 | 0 | 0 | 0 | 0 | 0 | 0 | 0 | 0 |
| 968 | ilvI | 0 | 0 | 1 | 0 | 0 | 0 | 0 | 0 | 0 | 0 |
| 969 | coaE | 0 | 0 | 0 | 0 | 0 | 0 | 0 | 0 | 0 | 0 |
| 970 | ilvD | 0 | 0 | 0 | 0 | 0 | 0 | 0 | 0 | 0 | 0 |
| 971 | ymfE | 0 | 0 | 0 | 0 | 0 | 0 | 0 | 0 | 0 | 0 |
| 972 | ilvC | 0 | 0 | 0 | 0 | 0 | 0 | 0 | 0 | 0 | 0 |
| 973 | hfq | 0 | 0 | 0 | 0 | 0 | 0 | 0 | 0 | 0 | 0 |
| 974 | ego | 0 | 0 | 0 | 0 | 0 | 0 | 0 | 0 | 0 | 0 |
| 975 | yedX | 0 | 0 | 0 | 0 | 0 | 0 | 0 | 0 | 0 | 0 |
| 976 | yedY | 0 | 0 | 0 | 0 | 0 | 0 | 0 | 0 | 0 | 0 |
| 977 | yedW | 0 | 0 | 0 | 0 | 0 | 0 | 0 | 0 | 0 | 0 |
| 978 | ybeT | 0 | 0 | 0 | 0 | 0 | 0 | 0 | 0 | 0 | 0 |
| 979 | yedP | 0 | 0 | 0 | 0 | 0 | 0 | 0 | 0 | 0 | 0 |
| 980 | yedQ | 0 | 0 | 0 | 0 | 0 | 0 | 0 | 0 | 0 | 0 |
| 981 | allD | 0 | 0 | 0 | 0 | 0 | 0 | 0 | 0 | 0 | 0 |
| 982 | allA | 0 | 0 | 0 | 0 | 0 | 0 | 0 | 0 | 0 | 0 |
| 983 | poxB | 0 | 0 | 0 | 0 | 0 | 0 | 0 | 0 | 0 | 0 |
| 984 | emrR | 0 | 0 | 0 | 0 | 0 | 0 | 0 | 0 | 0 | 0 |
| 985 | ugpA | 0 | 0 | 0 | 0 | 0 | 0 | 0 | 0 | 0 | 0 |
| 986 | ugpB | 0 | 0 | 0 | 0 | 0 | 0 | 0 | 0 | 0 | 0 |
| 987 | allS | 0 | 0 | 0 | 0 | 0 | 0 | 0 | 0 | 0 | 0 |
| 988 | allR | 0 | 0 | 0 | 0 | 0 | 0 | 0 | 0 | 0 | 0 |
| 989 | emrA | 0 | 0 | 0 | 0 | 0 | 0 | 0 | 0 | 0 | 0 |

| | | | | | | | | | | | |
|---|---|---|---|---|---|---|---|---|---|---|---|
| 990 | emrE | 0 | 0 | 0 | 0 | 0 | 0 | 0 | 0 | 0 | 0 |
| 991 | pcnB | 0 | 0 | 0 | 0 | 0 | 0 | 0 | 0 | 0 | 0 |
| 992 | yibL | 0 | 0 | 0 | 0 | 0 | 0 | 0 | 0 | 0 | 0 |
| 993 | yibF | 0 | 0 | 0 | 0 | 0 | 0 | 0 | 0 | 0 | 0 |
| 994 | panB | 0 | 0 | 0 | 0 | 0 | 0 | 0 | 0 | 0 | 0 |
| 995 | panE | 0 | 0 | 0 | 0 | 0 | 0 | 0 | 0 | 0 | 0 |
| 996 | panD | 0 | 0 | 0 | 0 | 0 | 0 | 0 | 0 | 0 | 0 |
| 997 | yejG | 0 | 0 | 0 | 0 | 0 | 0 | 0 | 0 | 0 | 0 |
| 998 | yejA | 0 | 0 | 0 | 0 | 0 | 0 | 0 | 0 | 0 | 0 |
| 999 | yejL | 0 | 0 | 0 | 0 | 0 | 0 | 0 | 0 | 0 | 0 |
| 1000 | yejO | 0 | 0 | 0 | 0 | 0 | 0 | 0 | 0 | 0 | 0 |
| 1001 | yejH | 0 | 0 | 0 | 0 | 0 | 0 | 0 | 0 | 0 | 0 |
| 1002 | rbsD | 0 | 0 | 0 | 0 | 0 | 0 | 0 | 0 | 0 | 0 |
| 1003 | yejK | 0 | 0 | 0 | 0 | 0 | 0 | 0 | 0 | 0 | 0 |
| 1004 | lsrR | 0 | 0 | 0 | 0 | 0 | 0 | 0 | 0 | 0 | 0 |
| 1005 | ydiF | 0 | 0 | 0 | 0 | 0 | 0 | 0 | 0 | 0 | 0 |
| 1006 | ygbA | 0 | 0 | 0 | 0 | 0 | 0 | 0 | 0 | 0 | 0 |
| 1007 | ygbN | 0 | 0 | 0 | 0 | 0 | 0 | 0 | 0 | 0 | 0 |
| 1008 | ygbJ | 0 | 0 | 0 | 0 | 0 | 0 | 0 | 0 | 0 | 0 |
| 1009 | ygbI | 0 | 0 | 0 | 0 | 0 | 0 | 0 | 0 | 0 | 0 |
| 1010 | adrA | 0 | 0 | 0 | 0 | 0 | 0 | 0 | 0 | 0 | 0 |
| 1011 | dedA | 0 | 0 | 1 | 0 | 0 | 0 | 0 | 0 | 0 | 0 |
| 1012 | lolC | 0 | 0 | 0 | 0 | 0 | 0 | 0 | 0 | 0 | 0 |
| 1013 | ybiS | 0 | 0 | 0 | 0 | 0 | 0 | 0 | 0 | 0 | 0 |
| 1014 | lolA | 0 | 0 | 0 | 0 | 0 | 0 | 0 | 0 | 0 | 0 |
| 1015 | ybiV | 0 | 0 | 0 | 0 | 0 | 0 | 0 | 0 | 0 | 0 |
| 1016 | dapF | 0 | 0 | 0 | 0 | 0 | 0 | 0 | 0 | 0 | 0 |
| 1017 | ybiA | 18 | 0 | 12 | 0 | 46 | 0 | 25 | 0 | 11 | 0 |
| 1018 | ybiF | 0 | 0 | 0 | 0 | 0 | 0 | 0 | 0 | 0 | 0 |
| 1019 | ybiJ | 0 | 0 | 0 | 0 | 0 | 0 | 0 | 0 | 0 | 0 |
| 1020 | ybiH | 0 | 0 | 0 | 0 | 0 | 0 | 0 | 0 | 0 | 0 |
| 1021 | ybiN | 0 | 0 | 0 | 0 | 0 | 0 | 0 | 0 | 0 | 0 |
| 1022 | ybiO | 0 | 0 | 0 | 0 | 0 | 0 | 0 | 0 | 0 | 0 |
| 1023 | ybiM | 0 | 0 | 0 | 0 | 0 | 0 | 0 | 0 | 0 | 0 |
| 1024 | nac | 0 | 0 | 0 | 0 | 0 | 0 | 0 | 0 | 0 | 0 |
| 1025 | b2680 | 0 | 0 | 1 | 0 | 0 | 0 | 0 | 0 | 0 | 0 |
| 1026 | asr | 0 | 0 | 0 | 0 | 0 | 0 | 0 | 0 | 1 | 0 |
| 1027 | oppA | 0 | 0 | 0 | 0 | 0 | 0 | 0 | 0 | 0 | 0 |
| 1028 | oppB | 0 | 0 | 0 | 0 | 0 | 0 | 0 | 0 | 0 | 0 |
| 1029 | asd | 0 | 0 | 0 | 0 | 0 | 0 | 0 | 0 | 0 | 0 |
| 1030 | yaiE | 0 | 0 | 0 | 0 | 0 | 0 | 0 | 0 | 0 | 0 |
| 1031 | dinB | 0 | 0 | 0 | 0 | 0 | 0 | 0 | 0 | 0 | 0 |
| 1032 | recA | 0 | 0 | 0 | 0 | 0 | 0 | 1 | 0 | 0 | 0 |
| 1033 | recE | 0 | 0 | 0 | 0 | 0 | 0 | 0 | 0 | 0 | 0 |
| 1034 | recX | 0 | 0 | 0 | 0 | 0 | 0 | 0 | 0 | 0 | 0 |
| 1035 | ispB | 0 | 0 | 0 | 0 | 0 | 0 | 0 | 0 | 0 | 0 |
| 1036 | hscB | 0 | 0 | 0 | 0 | 0 | 0 | 0 | 0 | 0 | 0 |
| 1037 | ydiL | 54 | 2 | 26 | 16 | 120 | 4 | 63 | 145 | 98 | 1375 |
| 1038 | ydiM | 0 | 0 | 0 | 0 | 0 | 0 | 0 | 0 | 0 | 0 |
| 1039 | tatE | 0 | 0 | 0 | 0 | 0 | 0 | 0 | 0 | 0 | 0 |
| 1040 | pspF | 0 | 0 | 0 | 0 | 0 | 0 | 0 | 0 | 0 | 0 |
| 1041 | pspE | 0 | 0 | 0 | 0 | 0 | 0 | 0 | 0 | 0 | 0 |
| 1042 | pspB | 0 | 0 | 0 | 0 | 0 | 0 | 0 | 0 | 0 | 0 |
| 1043 | moaA | 0 | 0 | 0 | 0 | 0 | 0 | 0 | 0 | 0 | 0 |
| 1044 | dinJ | 0 | 0 | 0 | 0 | 0 | 0 | 0 | 0 | 0 | 0 |
| 1045 | ydiY | 0 | 0 | 0 | 0 | 0 | 0 | 0 | 0 | 0 | 0 |
| 1046 | ydiP | 0 | 0 | 0 | 0 | 0 | 0 | 0 | 0 | 0 | 0 |
| 1047 | ydiQ | 0 | 0 | 0 | 0 | 0 | 0 | 0 | 0 | 0 | 0 |
| 1048 | ydiS | 0 | 0 | 0 | 0 | 0 | 0 | 0 | 0 | 0 | 0 |
| 1049 | ydiU | 0 | 0 | 0 | 0 | 0 | 0 | 0 | 0 | 0 | 0 |
| 1050 | ydiV | 0 | 0 | 0 | 0 | 0 | 0 | 0 | 0 | 0 | 0 |
| 1051 | dinG | 2 | 0 | 1 | 0 | 3 | 0 | 0 | 0 | 1 | 0 |
| 1052 | yceP | 0 | 0 | 0 | 0 | 0 | 0 | 0 | 0 | 0 | 0 |
| 1053 | guaC | 25 | 0 | 6 | 0 | 73 | 2 | 31 | 0 | 30 | 0 |
| 1054 | guaB | 0 | 0 | 0 | 0 | 0 | 0 | 0 | 0 | 0 | 0 |
| 1055 | guaA | 0 | 0 | 0 | 0 | 0 | 0 | 0 | 0 | 0 | 0 |

| | | | | | | | | | | | |
|---|---|---|---|---|---|---|---|---|---|---|---|
| 1056 | cadA | 0 | 0 | 0 | 0 | 0 | 0 | 0 | 0 | 0 | 0 |
| 1057 | yaiI | 0 | 0 | 0 | 0 | 0 | 0 | 0 | 0 | 0 | 0 |
| 1058 | yaiM | 0 | 0 | 0 | 0 | 0 | 0 | 0 | 0 | 0 | 0 |
| 1059 | dfp | 0 | 0 | 0 | 0 | 0 | 0 | 0 | 0 | 0 | 0 |
| 1060 | yaiN | 0 | 0 | 0 | 0 | 0 | 0 | 0 | 0 | 0 | 0 |
| 1061 | yaiP | 0 | 0 | 0 | 0 | 0 | 0 | 0 | 0 | 0 | 0 |
| 1062 | cadB | 0 | 0 | 1 | 0 | 0 | 0 | 0 | 0 | 0 | 0 |
| 1063 | yceF | 25 | 0 | 9 | 0 | 55 | 0 | 20 | 0 | 29 | 0 |
| 1064 | yaiW | 0 | 0 | 0 | 0 | 0 | 0 | 0 | 0 | 0 | 0 |
| 1065 | gpt | 0 | 0 | 0 | 0 | 0 | 0 | 0 | 0 | 0 | 0 |
| 1066 | yaiX | 0 | 0 | 0 | 0 | 0 | 0 | 0 | 0 | 0 | 0 |
| 1067 | yaiZ | 0 | 0 | 0 | 0 | 0 | 0 | 0 | 0 | 0 | 0 |
| 1068 | gpp | 0 | 0 | 0 | 0 | 0 | 0 | 0 | 0 | 0 | 0 |
| 1069 | gntT | 0 | 0 | 0 | 0 | 0 | 0 | 0 | 0 | 0 | 0 |
| 1070 | ubiX | 0 | 0 | 0 | 0 | 0 | 0 | 0 | 0 | 0 | 0 |
| 1071 | b0501 | 0 | 0 | 0 | 0 | 0 | 0 | 0 | 0 | 0 | 0 |
| 1072 | yceK | 0 | 0 | 0 | 0 | 0 | 0 | 0 | 0 | 0 | 0 |
| 1073 | greA | 0 | 0 | 0 | 0 | 0 | 0 | 0 | 0 | 0 | 0 |
| 1074 | greB | 0 | 0 | 0 | 0 | 0 | 0 | 0 | 0 | 0 | 0 |
| 1075 | gntR | 0 | 0 | 0 | 0 | 0 | 0 | 0 | 0 | 0 | 0 |
| 1076 | ubiF | 9427 | 246 | 1857 | 105 | 16038 | 235 | 12965 | 1314 | 8732 | 22 |
| 1077 | ubiG | 2768 | 419 | 477 | 71 | 4844 | 476 | 3587 | 1555 | 2614 | 35 |
| 1078 | ubiE | 0 | 0 | 0 | 0 | 0 | 0 | 0 | 0 | 0 | 0 |
| 1079 | ubiC | 0 | 0 | 0 | 0 | 0 | 0 | 0 | 0 | 0 | 0 |
| 1080 | gntP | 0 | 0 | 0 | 0 | 0 | 0 | 0 | 3 | 0 | 2 |
| 1081 | parC | 0 | 0 | 0 | 0 | 0 | 0 | 0 | 0 | 0 | 0 |
| 1082 | bglA | 51 | 0 | 24 | 0 | 80 | 0 | 33 | 0 | 0 | 0 |
| 1083 | ascG | 0 | 0 | 0 | 0 | 0 | 0 | 0 | 0 | 1 | 0 |
| 1084 | b0165 | 0 | 0 | 0 | 0 | 0 | 0 | 1 | 0 | 0 | 0 |
| 1085 | selD | 0 | 0 | 0 | 0 | 0 | 0 | 0 | 0 | 0 | 0 |
| 1086 | alsR | 0 | 0 | 0 | 0 | 0 | 0 | 0 | 0 | 0 | 0 |
| 1087 | selA | 0 | 0 | 0 | 0 | 0 | 0 | 0 | 0 | 0 | 0 |
| 1088 | selC | 0 | 0 | 0 | 0 | 0 | 0 | 0 | 0 | 0 | 0 |
| 1089 | alsI | 0 | 0 | 0 | 0 | 0 | 0 | 0 | 0 | 0 | 0 |
| 1090 | alsB | 0 | 0 | 0 | 0 | 0 | 0 | 0 | 0 | 0 | 0 |
| 1091 | ymgE | 0 | 0 | 0 | 0 | 0 | 0 | 0 | 0 | 0 | 0 |
| 1092 | alsA | 0 | 0 | 0 | 0 | 0 | 0 | 0 | 0 | 0 | 0 |
| 1093 | yeeI | 0 | 0 | 0 | 0 | 0 | 0 | 0 | 0 | 0 | 0 |
| 1094 | galU | 0 | 0 | 0 | 0 | 0 | 0 | 0 | 0 | 0 | 0 |
| 1095 | yeeJ | 0 | 0 | 1 | 0 | 0 | 0 | 0 | 0 | 0 | 0 |
| 1096 | galP | 0 | 0 | 0 | 0 | 0 | 0 | 0 | 0 | 0 | 0 |
| 1097 | galR | 0 | 0 | 0 | 0 | 0 | 0 | 0 | 0 | 0 | 0 |
| 1098 | galS | 38 | 11 | 31 | 1 | 102 | 16 | 44 | 0 | 37 | 0 |
| 1099 | yeeA | 0 | 0 | 0 | 0 | 0 | 0 | 0 | 0 | 0 | 0 |
| 1100 | yeeE | 0 | 0 | 0 | 0 | 0 | 0 | 1 | 0 | 0 | 0 |
| 1101 | lit | 0 | 0 | 0 | 0 | 0 | 0 | 0 | 0 | 0 | 0 |
| 1102 | galE | 0 | 0 | 0 | 0 | 0 | 0 | 0 | 0 | 0 | 0 |
| 1103 | yeeR | 0 | 0 | 0 | 0 | 0 | 0 | 0 | 0 | 0 | 0 |
| 1104 | yeeU | 0 | 0 | 0 | 0 | 0 | 0 | 0 | 0 | 0 | 0 |
| 1105 | yeeV | 1 | 0 | 0 | 0 | 0 | 0 | 0 | 0 | 0 | 0 |
| 1106 | hipB | 0 | 0 | 0 | 0 | 0 | 0 | 0 | 0 | 0 | 0 |
| 1107 | rho | 0 | 0 | 0 | 0 | 0 | 0 | 0 | 0 | 0 | 0 |
| 1108 | msrA | 0 | 0 | 0 | 0 | 0 | 0 | 0 | 0 | 0 | 0 |
| 1109 | hofM | 0 | 0 | 0 | 0 | 0 | 0 | 1 | 0 | 0 | 0 |
| 1110 | yhaH | 0 | 0 | 0 | 0 | 0 | 0 | 0 | 0 | 0 | 0 |
| 1111 | yhaJ | 0 | 0 | 0 | 0 | 0 | 0 | 0 | 0 | 0 | 0 |
| 1112 | yhaK | 0 | 0 | 0 | 0 | 0 | 0 | 0 | 0 | 0 | 0 |
| 1113 | cdd | 0 | 0 | 0 | 0 | 0 | 0 | 0 | 0 | 0 | 0 |
| 1114 | yoaB | 0 | 0 | 0 | 0 | 0 | 0 | 0 | 0 | 0 | 0 |
| 1115 | yoaA | 0 | 0 | 0 | 0 | 0 | 0 | 0 | 0 | 0 | 0 |
| 1116 | prpB | 0 | 0 | 0 | 0 | 0 | 0 | 0 | 0 | 0 | 0 |
| 1117 | prpC | 0 | 0 | 0 | 0 | 0 | 0 | 0 | 0 | 0 | 0 |
| 1118 | yoaE | 0 | 0 | 0 | 0 | 0 | 0 | 0 | 0 | 0 | 0 |
| 1119 | yoaD | 0 | 0 | 0 | 0 | 0 | 0 | 0 | 0 | 0 | 0 |
| 1120 | yoaH | 0 | 0 | 0 | 0 | 0 | 0 | 0 | 0 | 0 | 0 |
| 1121 | ugd | 0 | 0 | 0 | 0 | 0 | 0 | 0 | 0 | 0 | 0 |

| | | | | | | | | | | | |
|---|---|---|---|---|---|---|---|---|---|---|---|
| 1122 | ygcB | 0 | 0 | 0 | 0 | 0 | 0 | 0 | 0 | 0 | 0 |
| 1123 | prpR | 0 | 0 | 0 | 0 | 0 | 0 | 0 | 0 | 0 | 0 |
| 1124 | ygcF | 0 | 0 | 0 | 0 | 0 | 0 | 0 | 0 | 0 | 0 |
| 1125 | clpX | 0 | 0 | 0 | 0 | 0 | 0 | 0 | 0 | 0 | 0 |
| 1126 | stfE | 0 | 0 | 0 | 0 | 0 | 0 | 0 | 0 | 0 | 0 |
| 1127 | bfd | 0 | 0 | 0 | 0 | 0 | 0 | 0 | 0 | 0 | 0 |
| 1128 | clpP | 0 | 0 | 0 | 0 | 0 | 0 | 0 | 0 | 0 | 0 |
| 1129 | clpS | 0 | 0 | 0 | 0 | 0 | 0 | 0 | 0 | 0 | 0 |
| 1130 | gshA | 0 | 0 | 0 | 0 | 0 | 1 | 0 | 0 | 0 | 0 |
| 1131 | sspA | 1 | 0 | 1 | 0 | 0 | 0 | 0 | 0 | 0 | 0 |
| 1132 | stfR | 0 | 0 | 0 | 0 | 0 | 0 | 0 | 0 | 0 | 0 |
| 1133 | hyfR | 0 | 0 | 0 | 0 | 0 | 0 | 0 | 0 | 0 | 0 |
| 1134 | clpB | 0 | 1 | 0 | 0 | 0 | 0 | 0 | 0 | 0 | 0 |
| 1135 | sthA | 0 | 1 | 0 | 0 | 0 | 0 | 0 | 0 | 0 | 0 |
| 1136 | plsX | 0 | 0 | 0 | 0 | 0 | 0 | 0 | 0 | 0 | 0 |
| 1137 | hrpB | 162 | 2 | 26 | 2 | 316 | 11 | 147 | 1 | 277 | 0 |
| 1138 | hrpA | 0 | 0 | 1 | 0 | 0 | 0 | 0 | 0 | 0 | 0 |
| 1139 | plsC | 0 | 0 | 0 | 0 | 0 | 0 | 0 | 0 | 0 | 0 |
| 1140 | plsB | 0 | 0 | 0 | 0 | 0 | 0 | 0 | 0 | 0 | 0 |
| 1141 | rpiA | 0 | 0 | 0 | 0 | 0 | 0 | 0 | 0 | 0 | 0 |
| 1142 | pck | 0 | 0 | 0 | 0 | 0 | 0 | 0 | 0 | 0 | 0 |
| 1143 | ldhA | 0 | 0 | 0 | 0 | 0 | 0 | 0 | 0 | 0 | 0 |
| 1144 | yjcD | 0 | 0 | 0 | 0 | 0 | 0 | 0 | 0 | 0 | 0 |
| 1145 | yjcE | 0 | 0 | 0 | 0 | 0 | 0 | 0 | 0 | 0 | 0 |
| 1146 | crp | 0 | 0 | 0 | 0 | 0 | 0 | 0 | 0 | 0 | 0 |
| 1147 | fsaA | 0 | 0 | 0 | 0 | 0 | 0 | 0 | 0 | 0 | 0 |
| 1148 | fkpB | 0 | 0 | 0 | 0 | 0 | 0 | 0 | 0 | 0 | 0 |
| 1149 | yjcO | 1596 | 9 | 418 | 34 | 2883 | 16 | 1811 | 9 | 1779 | 2 |
| 1150 | yjcH | 1 | 0 | 0 | 0 | 1 | 0 | 1 | 0 | 0 | 0 |
| 1151 | rfe | 0 | 0 | 0 | 0 | 0 | 0 | 0 | 0 | 0 | 0 |
| 1152 | aaeR | 0 | 0 | 0 | 0 | 0 | 0 | 0 | 0 | 0 | 0 |
| 1153 | crl | 0 | 0 | 0 | 0 | 0 | 0 | 0 | 0 | 0 | 0 |
| 1154 | agn43 | 0 | 0 | 0 | 0 | 0 | 0 | 0 | 0 | 0 | 0 |
| 1155 | uxuA | 0 | 0 | 0 | 0 | 0 | 0 | 0 | 0 | 0 | 0 |
| 1156 | cysK | 0 | 0 | 0 | 0 | 0 | 0 | 0 | 0 | 0 | 0 |
| 1157 | barA | 33 | 0 | 12 | 0 | 74 | 0 | 42 | 0 | 0 | 0 |
| 1158 | cbpA | 0 | 0 | 0 | 0 | 0 | 0 | 0 | 0 | 0 | 0 |
| 1159 | imp | 0 | 0 | 0 | 1 | 0 | 0 | 0 | 0 | 0 | 0 |
| 1160 | mhpR | 0 | 0 | 0 | 0 | 0 | 0 | 0 | 0 | 0 | 0 |
| 1161 | yicH | 0 | 0 | 0 | 0 | 0 | 0 | 0 | 0 | 0 | 0 |
| 1162 | yicC | 19 | 0 | 4 | 0 | 17 | 0 | 10 | 0 | 23 | 0 |
| 1163 | solA | 0 | 0 | 1 | 0 | 0 | 0 | 0 | 0 | 0 | 0 |
| 1164 | fdhF | 0 | 0 | 0 | 0 | 0 | 0 | 0 | 0 | 0 | 0 |
| 1165 | fdhD | 0 | 0 | 0 | 0 | 0 | 0 | 0 | 0 | 0 | 0 |
| 1166 | tgt | 0 | 0 | 0 | 0 | 0 | 0 | 0 | 0 | 0 | 0 |
| 1167 | napF | 0 | 0 | 0 | 0 | 0 | 0 | 0 | 0 | 0 | 0 |
| 1168 | sppA | 0 | 0 | 0 | 0 | 0 | 0 | 0 | 0 | 0 | 0 |
| 1169 | ybhQ | 0 | 0 | 0 | 0 | 0 | 1 | 0 | 0 | 0 | 0 |
| 1170 | ybhM | 0 | 0 | 0 | 1 | 0 | 0 | 0 | 0 | 0 | 0 |
| 1171 | ybhL | 0 | 0 | 1 | 0 | 0 | 0 | 0 | 0 | 0 | 0 |
| 1172 | ybhI | 0 | 0 | 0 | 0 | 0 | 0 | 0 | 0 | 0 | 0 |
| 1173 | ybhK | 0 | 0 | 0 | 0 | 0 | 0 | 0 | 0 | 0 | 0 |
| 1174 | ybhJ | 0 | 0 | 0 | 0 | 0 | 0 | 0 | 0 | 0 | 0 |
| 1175 | ybhE | 0 | 0 | 0 | 0 | 0 | 0 | 0 | 0 | 0 | 0 |
| 1176 | ybhD | 0 | 0 | 0 | 0 | 0 | 0 | 0 | 0 | 0 | 0 |
| 1177 | ybhA | 0 | 0 | 0 | 0 | 0 | 0 | 0 | 0 | 0 | 0 |
| 1178 | ybhC | 0 | 0 | 13 | 0 | 0 | 0 | 0 | 0 | 0 | 0 |
| 1179 | mlc | 0 | 0 | 0 | 0 | 0 | 0 | 0 | 0 | 0 | 0 |
| 1180 | fur | 0 | 0 | 0 | 0 | 0 | 0 | 0 | 0 | 0 | 0 |
| 1181 | tdcB | 0 | 0 | 0 | 0 | 0 | 0 | 0 | 0 | 0 | 0 |
| 1182 | b1172 | 0 | 0 | 0 | 0 | 0 | 0 | 0 | 0 | 0 | 0 |
| 1183 | glgB | 0 | 0 | 0 | 0 | 0 | 0 | 0 | 0 | 0 | 0 |
| 1184 | glgS | 0 | 0 | 0 | 0 | 0 | 0 | 0 | 0 | 0 | 0 |
| 1185 | motA | 0 | 0 | 0 | 0 | 0 | 0 | 0 | 0 | 0 | 0 |
| 1186 | ptrB | 0 | 0 | 0 | 0 | 0 | 0 | 0 | 0 | 0 | 0 |
| 1187 | dsbG | 0 | 0 | 0 | 0 | 0 | 0 | 0 | 0 | 0 | 0 |

| | | | | | | | | | | | |
|---|---|---|---|---|---|---|---|---|---|---|---|
| 1188 | ampH | 0 | 0 | 0 | 0 | 0 | 0 | 0 | 0 | 0 | 0 |
| 1189 | ydhH | 0 | 0 | 0 | 0 | 0 | 0 | 0 | 0 | 0 | 0 |
| 1190 | ydhO | 0 | 0 | 0 | 0 | 0 | 0 | 0 | 0 | 0 | 0 |
| 1191 | syd | 0 | 0 | 0 | 0 | 0 | 0 | 0 | 0 | 0 | 0 |
| 1192 | ydhB | 0 | 0 | 1 | 0 | 0 | 0 | 0 | 1 | 0 | 0 |
| 1193 | mobA | 11 | 0 | 2 | 0 | 11 | 0 | 8 | 0 | 26 | 0 |
| 1194 | ampD | 0 | 0 | 0 | 0 | 0 | 0 | 0 | 0 | 0 | 0 |
| 1195 | ydhD | 0 | 0 | 0 | 0 | 0 | 0 | 0 | 0 | 0 | 0 |
| 1196 | ydhZ | 0 | 0 | 0 | 0 | 0 | 0 | 0 | 0 | 0 | 0 |
| 1197 | dgt | 0 | 0 | 0 | 1 | 0 | 0 | 0 | 0 | 0 | 0 |
| 1198 | ydhS | 0 | 0 | 0 | 0 | 0 | 0 | 0 | 0 | 0 | 0 |
| 1199 | ydhR | 384 | 2588 | 128 | 62 | 515 | 1607 | 265 | 0 | 1847 | 6884 |
| 1200 | ydhQ | 0 | 1 | 0 | 0 | 0 | 0 | 0 | 0 | 0 | 0 |
| 1201 | ydhP | 0 | 0 | 0 | 0 | 0 | 0 | 0 | 0 | 0 | 0 |
| 1202 | yafL | 0 | 0 | 0 | 0 | 0 | 0 | 0 | 0 | 0 | 0 |
| 1203 | yafM | 0 | 0 | 0 | 0 | 0 | 0 | 0 | 0 | 0 | 0 |
| 1204 | yafN | 0 | 0 | 1 | 0 | 0 | 0 | 0 | 0 | 0 | 0 |
| 1205 | zitB | 0 | 0 | 0 | 0 | 0 | 0 | 0 | 0 | 0 | 0 |
| 1206 | yafJ | 0 | 0 | 1 | 0 | 0 | 0 | 0 | 0 | 0 | 0 |
| 1207 | yafK | 0 | 0 | 0 | 0 | 0 | 0 | 0 | 0 | 1 | 0 |
| 1208 | yafD | 0 | 0 | 0 | 0 | 0 | 0 | 0 | 0 | 0 | 0 |
| 1209 | ycfT | 0 | 0 | 0 | 0 | 0 | 0 | 0 | 0 | 0 | 0 |
| 1210 | mipA | 0 | 0 | 0 | 0 | 0 | 0 | 0 | 0 | 0 | 0 |
| 1211 | ycfR | 0 | 0 | 0 | 0 | 0 | 0 | 0 | 0 | 0 | 0 |
| 1212 | ycfS | 0 | 0 | 0 | 0 | 0 | 0 | 0 | 0 | 0 | 0 |
| 1213 | slyX | 0 | 0 | 0 | 0 | 0 | 0 | 0 | 0 | 0 | 0 |
| 1214 | yafC | 0 | 0 | 0 | 0 | 0 | 0 | 0 | 0 | 0 | 0 |
| 1215 | efp | 0 | 0 | 0 | 0 | 0 | 0 | 0 | 0 | 0 | 0 |
| 1216 | slyD | 0 | 0 | 0 | 0 | 0 | 0 | 0 | 0 | 0 | 0 |
| 1217 | ycfJ | 0 | 0 | 0 | 0 | 0 | 0 | 0 | 0 | 0 | 0 |
| 1218 | yafZ | 0 | 0 | 0 | 0 | 0 | 0 | 0 | 0 | 0 | 0 |
| 1219 | slyA | 0 | 0 | 0 | 0 | 0 | 0 | 0 | 0 | 0 | 0 |
| 1220 | yafT | 0 | 0 | 0 | 0 | 0 | 0 | 0 | 0 | 0 | 0 |
| 1221 | yafU | 0 | 0 | 0 | 0 | 0 | 0 | 0 | 0 | 0 | 0 |
| 1222 | yafV | 0 | 0 | 0 | 0 | 0 | 0 | 0 | 0 | 0 | 0 |
| 1223 | uxaC | 0 | 0 | 0 | 0 | 0 | 0 | 0 | 0 | 0 | 0 |
| 1224 | flgM | 0 | 0 | 0 | 0 | 0 | 0 | 0 | 0 | 0 | 0 |
| 1225 | yafS | 0 | 0 | 0 | 0 | 0 | 0 | 0 | 0 | 0 | 0 |
| 1226 | sscR | 0 | 0 | 0 | 0 | 0 | 0 | 0 | 0 | 0 | 0 |
| 1227 | dhaM | 0 | 0 | 0 | 0 | 0 | 0 | 0 | 0 | 0 | 0 |
| 1228 | dhaR | 0 | 0 | 0 | 0 | 0 | 0 | 0 | 0 | 0 | 0 |
| 1229 | yjbF | 0 | 0 | 0 | 0 | 0 | 0 | 0 | 0 | 0 | 0 |
| 1230 | yjbE | 0 | 0 | 0 | 0 | 0 | 0 | 0 | 0 | 0 | 0 |
| 1231 | yjbB | 0 | 0 | 0 | 0 | 0 | 0 | 0 | 0 | 0 | 0 |
| 1232 | yeeX | 0 | 0 | 0 | 0 | 0 | 0 | 0 | 0 | 0 | 0 |
| 1233 | mreC | 34 | 0 | 19 | 0 | 48 | 0 | 22 | 0 | 60 | 0 |
| 1234 | yjbJ | 0 | 0 | 0 | 0 | 0 | 0 | 0 | 0 | 0 | 0 |
| 1235 | yjbI | 0 | 0 | 0 | 0 | 0 | 0 | 0 | 0 | 0 | 0 |
| 1236 | yjdB | 0 | 1 | 0 | 0 | 0 | 0 | 0 | 0 | 0 | 0 |
| 1237 | yjbQ | 0 | 0 | 0 | 0 | 0 | 0 | 0 | 0 | 0 | 0 |
| 1238 | deoR | 0 | 0 | 0 | 0 | 0 | 0 | 0 | 0 | 0 | 0 |
| 1239 | yhhQ | 0 | 0 | 0 | 0 | 0 | 0 | 0 | 0 | 0 | 0 |
| 1240 | ychF | 0 | 0 | 0 | 0 | 0 | 0 | 0 | 0 | 0 | 0 |
| 1241 | ptsA | 0 | 0 | 0 | 0 | 0 | 0 | 0 | 0 | 0 | 0 |
| 1242 | ychM | 0 | 0 | 0 | 0 | 0 | 0 | 0 | 0 | 0 | 0 |
| 1243 | ychH | 70 | 4823 | 33 | 213 | 119 | 6188 | 291 | 5620 | 4 | 0 |
| 1244 | ptsG | 0 | 0 | 0 | 0 | 0 | 1 | 0 | 2 | 0 | 0 |
| 1245 | ymdB | 0 | 0 | 0 | 0 | 0 | 0 | 0 | 0 | 0 | 0 |
| 1246 | dppA | 0 | 0 | 0 | 0 | 0 | 0 | 0 | 0 | 0 | 0 |
| 1247 | dppB | 0 | 0 | 0 | 0 | 0 | 0 | 0 | 0 | 0 | 0 |
| 1248 | uhpT | 0 | 0 | 0 | 0 | 0 | 0 | 0 | 0 | 0 | 0 |
| 1249 | fhuF | 0 | 0 | 1 | 0 | 0 | 0 | 0 | 0 | 0 | 0 |
| 1250 | fhuA | 0 | 0 | 0 | 0 | 0 | 0 | 0 | 0 | 0 | 0 |
| 1251 | fhuC | 0 | 0 | 0 | 0 | 0 | 0 | 0 | 0 | 0 | 0 |
| 1252 | uhpA | 0 | 1 | 0 | 0 | 0 | 0 | 0 | 0 | 0 | 0 |
| 1253 | nuoM | 0 | 0 | 0 | 0 | 0 | 0 | 0 | 0 | 1 | 0 |

| | | | | | | | | | | | |
|---|---|---|---|---|---|---|---|---|---|---|---|
| 1254 | proX | 0 | 0 | 0 | 0 | 0 | 0 | 0 | 0 | 0 | 0 |
| 1255 | proS | 0 | 0 | 0 | 0 | 0 | 0 | 0 | 0 | 0 | 0 |
| 1256 | proQ | 0 | 0 | 0 | 0 | 0 | 0 | 0 | 0 | 0 | 0 |
| 1257 | proL | 0 | 0 | 0 | 0 | 0 | 0 | 0 | 0 | 0 | 0 |
| 1258 | phnC | 0 | 0 | 0 | 0 | 0 | 0 | 0 | 0 | 0 | 0 |
| 1259 | luxS | 0 | 0 | 0 | 0 | 0 | 0 | 0 | 0 | 0 | 0 |
| 1260 | phnL | 0 | 0 | 0 | 0 | 0 | 0 | 0 | 0 | 0 | 0 |
| 1261 | proB | 0 | 0 | 0 | 0 | 0 | 0 | 0 | 0 | 0 | 0 |
| 1262 | lipA | 0 | 0 | 0 | 0 | 0 | 0 | 0 | 0 | 0 | 0 |
| 1263 | lipB | 0 | 0 | 0 | 0 | 0 | 0 | 0 | 0 | 0 | 0 |
| 1264 | bacA | 0 | 0 | 0 | 0 | 0 | 0 | 0 | 0 | 0 | 0 |
| 1265 | somA_macA | 0 | 0 | 0 | 0 | 0 | 0 | 0 | 0 | 0 | 0 |
| 1266 | bcsE | 0 | 0 | 0 | 0 | 0 | 0 | 0 | 0 | 0 | 0 |
| 1267 | b3044 | 0 | 0 | 6 | 0 | 0 | 0 | 0 | 0 | 0 | 0 |
| 1268 | yqeG | 0 | 0 | 0 | 0 | 0 | 0 | 0 | 0 | 0 | 0 |
| 1269 | yehD | 0 | 0 | 0 | 0 | 0 | 0 | 0 | 0 | 0 | 0 |
| 1270 | yifE | 0 | 0 | 0 | 0 | 0 | 0 | 0 | 0 | 0 | 0 |
| 1271 | yobF | 0 | 0 | 0 | 0 | 0 | 0 | 0 | 0 | 0 | 0 |
| 1272 | b2641 | 0 | 0 | 0 | 0 | 0 | 0 | 0 | 0 | 0 | 0 |
| 1273 | yqeI | 0 | 0 | 0 | 0 | 0 | 0 | 0 | 0 | 0 | 0 |
| 1274 | yqeH | 0 | 0 | 0 | 0 | 0 | 0 | 0 | 0 | 0 | 0 |
| 1275 | yqeK | 0 | 0 | 0 | 0 | 0 | 0 | 0 | 0 | 0 | 0 |
| 1276 | yehS | 0 | 0 | 0 | 0 | 0 | 0 | 0 | 0 | 0 | 0 |
| 1277 | yhfG | 0 | 0 | 0 | 0 | 0 | 0 | 0 | 0 | 0 | 0 |
| 1278 | yehZ | 0 | 0 | 0 | 1 | 0 | 0 | 0 | 0 | 0 | 0 |
| 1279 | bisC | 0 | 0 | 0 | 0 | 0 | 0 | 0 | 0 | 0 | 0 |
| 1280 | pbpG | 0 | 0 | 0 | 0 | 0 | 0 | 0 | 0 | 0 | 0 |
| 1281 | idi | 0 | 0 | 0 | 0 | 0 | 0 | 0 | 0 | 0 | 0 |
| 1282 | bioB | 0 | 0 | 0 | 0 | 0 | 0 | 0 | 0 | 0 | 0 |
| 1283 | b0332 | 0 | 0 | 0 | 0 | 0 | 0 | 0 | 0 | 0 | 0 |
| 1284 | pabC | 50 | 0 | 13 | 0 | 112 | 0 | 8 | 0 | 59 | 0 |
| 1285 | yeeO | 0 | 0 | 0 | 0 | 0 | 0 | 0 | 0 | 0 | 0 |
| 1286 | intF | 0 | 0 | 0 | 0 | 0 | 0 | 0 | 0 | 0 | 0 |
| 1287 | intA | 0 | 0 | 0 | 0 | 0 | 0 | 0 | 0 | 0 | 0 |
| 1288 | ybgA | 0 | 0 | 0 | 0 | 0 | 0 | 0 | 0 | 0 | 0 |
| 1289 | yfdQ | 0 | 0 | 0 | 0 | 0 | 0 | 0 | 0 | 0 | 0 |
| 1290 | intS | 0 | 0 | 0 | 0 | 0 | 0 | 0 | 0 | 0 | 0 |
| 1291 | intZ | 0 | 0 | 0 | 0 | 0 | 0 | 0 | 0 | 0 | 0 |
| 1292 | infA | 0 | 0 | 0 | 0 | 0 | 0 | 0 | 0 | 0 | 0 |
| 1293 | rlmB | 0 | 0 | 0 | 0 | 0 | 0 | 0 | 0 | 0 | 0 |
| 1294 | ihfB | 0 | 0 | 0 | 0 | 0 | 0 | 0 | 0 | 0 | 0 |
| 1295 | atoS | 0 | 0 | 0 | 0 | 0 | 0 | 0 | 0 | 0 | 0 |
| 1296 | yhcG | 0 | 0 | 0 | 0 | 0 | 0 | 0 | 0 | 0 | 0 |
| 1297 | exuR | 0 | 0 | 0 | 0 | 0 | 0 | 0 | 0 | 0 | 0 |
| 1298 | tufB | 0 | 0 | 0 | 0 | 0 | 0 | 0 | 0 | 0 | 0 |
| 1299 | tufA | 0 | 0 | 0 | 0 | 0 | 0 | 0 | 0 | 0 | 0 |
| 1300 | ycgY | 0 | 0 | 0 | 0 | 0 | 0 | 0 | 0 | 0 | 0 |
| 1301 | chaC | 0 | 0 | 0 | 0 | 0 | 0 | 0 | 0 | 0 | 0 |
| 1302 | yagM | 0 | 0 | 0 | 0 | 0 | 0 | 0 | 0 | 0 | 0 |
| 1303 | yagI | 0 | 0 | 0 | 0 | 0 | 0 | 0 | 0 | 0 | 0 |
| 1304 | yagG | 0 | 0 | 0 | 0 | 0 | 0 | 0 | 0 | 0 | 0 |
| 1305 | yagE | 0 | 0 | 0 | 0 | 0 | 0 | 0 | 0 | 0 | 0 |
| 1306 | ycgR | 0 | 0 | 0 | 0 | 0 | 0 | 0 | 0 | 0 | 0 |
| 1307 | yagB | 0 | 0 | 0 | 0 | 0 | 0 | 0 | 0 | 0 | 0 |
| 1308 | ycgI | 0 | 0 | 0 | 0 | 0 | 0 | 0 | 0 | 0 | 0 |
| 1309 | ycgH | 0 | 0 | 0 | 0 | 0 | 0 | 0 | 0 | 0 | 0 |
| 1310 | ycgK | 0 | 0 | 0 | 0 | 0 | 0 | 0 | 0 | 0 | 0 |
| 1311 | ycgJ | 0 | 0 | 0 | 0 | 0 | 0 | 0 | 0 | 0 | 0 |
| 1312 | ycgM | 0 | 0 | 0 | 0 | 0 | 0 | 0 | 0 | 0 | 0 |
| 1313 | yagZ | 0 | 0 | 0 | 0 | 0 | 0 | 0 | 0 | 0 | 0 |
| 1314 | yagY | 0 | 0 | 0 | 0 | 0 | 0 | 0 | 0 | 0 | 0 |
| 1315 | ycgN | 0 | 0 | 0 | 0 | 0 | 0 | 0 | 0 | 0 | 0 |
| 1316 | yagU | 0 | 0 | 0 | 0 | 0 | 0 | 0 | 0 | 0 | 0 |
| 1317 | yagT | 0 | 0 | 0 | 0 | 0 | 0 | 0 | 0 | 0 | 0 |
| 1318 | fldB | 0 | 0 | 0 | 0 | 0 | 0 | 0 | 0 | 0 | 0 |
| 1319 | ycgF | 0 | 0 | 0 | 0 | 0 | 0 | 0 | 0 | 0 | 0 |

| | | | | | | | | | | | |
|---|---|---|---|---|---|---|---|---|---|---|---|
| 1320 | yhbP | 0 | 0 | 0 | 0 | 0 | 0 | 0 | 0 | 0 | 0 |
| 1321 | pyrH | 0 | 0 | 0 | 0 | 0 | 0 | 0 | 0 | 0 | 0 |
| 1322 | pyrG | 0 | 0 | 0 | 0 | 0 | 0 | 1 | 0 | 0 | 0 |
| 1323 | pyrC | 0 | 0 | 0 | 0 | 0 | 0 | 0 | 0 | 0 | 0 |
| 1324 | pyrB | 0 | 0 | 0 | 0 | 0 | 0 | 0 | 0 | 0 | 0 |
| 1325 | glpX | 0 | 0 | 0 | 0 | 0 | 0 | 0 | 0 | 0 | 0 |
| 1326 | glpE | 0 | 0 | 0 | 0 | 0 | 0 | 0 | 0 | 0 | 0 |
| 1327 | glpD | 0 | 0 | 0 | 0 | 0 | 0 | 0 | 0 | 0 | 0 |
| 1328 | glpA | 0 | 0 | 0 | 0 | 0 | 0 | 0 | 0 | 0 | 0 |
| 1329 | yciW | 0 | 0 | 0 | 0 | 0 | 0 | 0 | 0 | 0 | 0 |
| 1330 | fxsA | 0 | 0 | 0 | 0 | 0 | 0 | 0 | 0 | 0 | 0 |
| 1331 | yciT | 0 | 0 | 0 | 0 | 0 | 0 | 0 | 0 | 0 | 0 |
| 1332 | yciS | 0 | 0 | 0 | 0 | 0 | 0 | 0 | 0 | 0 | 0 |
| 1333 | edd | 1 | 0 | 0 | 0 | 1 | 0 | 2 | 2 | 3 | 0 |
| 1334 | yciG | 0 | 0 | 0 | 0 | 0 | 0 | 0 | 0 | 0 | 0 |
| 1335 | b1437 | 0 | 0 | 0 | 0 | 0 | 0 | 0 | 0 | 0 | 0 |
| 1336 | yciC | 0 | 0 | 1 | 0 | 0 | 0 | 0 | 0 | 0 | 0 |
| 1337 | yciA | 0 | 0 | 0 | 0 | 0 | 0 | 0 | 0 | 0 | 0 |
| 1338 | phoH | 0 | 0 | 0 | 0 | 0 | 0 | 0 | 0 | 0 | 0 |
| 1339 | ygcU | 0 | 0 | 0 | 1 | 0 | 0 | 0 | 0 | 0 | 0 |
| 1340 | yciK | 0 | 0 | 0 | 0 | 0 | 0 | 0 | 0 | 0 | 0 |
| 1341 | yciI | 0 | 0 | 0 | 0 | 0 | 0 | 0 | 0 | 0 | 0 |
| 1342 | yoaF | 0 | 0 | 0 | 0 | 0 | 0 | 0 | 0 | 0 | 0 |
| 1343 | mdtH | 0 | 0 | 0 | 0 | 0 | 0 | 0 | 0 | 0 | 0 |
| 1344 | codB | 0 | 0 | 0 | 0 | 0 | 0 | 1 | 0 | 0 | 0 |
| 1345 | ygcW | 0 | 0 | 0 | 0 | 0 | 0 | 0 | 0 | 0 | 0 |
| 1346 | yihR | 0 | 0 | 0 | 0 | 0 | 0 | 0 | 0 | 0 | 0 |
| 1347 | zupT | 0 | 0 | 0 | 0 | 0 | 0 | 0 | 0 | 0 | 0 |
| 1348 | katE | 0 | 0 | 0 | 0 | 0 | 0 | 0 | 0 | 0 | 0 |
| 1349 | katG | 0 | 0 | 1 | 0 | 0 | 0 | 0 | 0 | 0 | 0 |
| 1350 | racR | 0 | 0 | 0 | 0 | 0 | 0 | 0 | 0 | 0 | 0 |
| 1351 | acnB | 0 | 0 | 0 | 0 | 0 | 0 | 0 | 0 | 0 | 0 |
| 1352 | kch | 0 | 0 | 0 | 0 | 0 | 0 | 0 | 0 | 0 | 0 |
| 1353 | b0259 | 0 | 0 | 0 | 0 | 0 | 0 | 0 | 0 | 0 | 0 |
| 1354 | racC | 0 | 0 | 0 | 0 | 0 | 0 | 0 | 0 | 0 | 0 |
| 1355 | b0257 | 0 | 0 | 0 | 0 | 0 | 0 | 0 | 0 | 0 | 0 |
| 1356 | b0255 | 0 | 0 | 0 | 0 | 0 | 0 | 0 | 0 | 0 | 0 |
| 1357 | yhcO | 0 | 0 | 0 | 0 | 0 | 0 | 0 | 0 | 0 | 0 |
| 1358 | yhcM | 0 | 0 | 0 | 0 | 0 | 0 | 0 | 0 | 0 | 0 |
| 1359 | yhcB | 0 | 0 | 0 | 0 | 0 | 0 | 0 | 0 | 0 | 0 |
| 1360 | yhcA | 0 | 0 | 0 | 0 | 0 | 0 | 0 | 0 | 0 | 0 |
| 1361 | yhjX | 0 | 0 | 0 | 0 | 0 | 0 | 0 | 0 | 0 | 0 |
| 1362 | yeiM | 0 | 0 | 0 | 0 | 0 | 0 | 0 | 0 | 0 | 0 |
| 1363 | yeiE | 0 | 0 | 0 | 0 | 0 | 0 | 0 | 0 | 0 | 0 |
| 1364 | yeiG | 0 | 0 | 0 | 0 | 0 | 0 | 0 | 0 | 0 | 0 |
| 1365 | sapA | 0 | 0 | 0 | 0 | 0 | 0 | 0 | 0 | 0 | 0 |
| 1366 | yeiC | 0 | 0 | 0 | 0 | 0 | 0 | 0 | 0 | 0 | 0 |
| 1367 | nhaA | 0 | 0 | 0 | 0 | 0 | 0 | 0 | 0 | 0 | 0 |
| 1368 | yeiT | 0 | 0 | 0 | 0 | 0 | 0 | 0 | 0 | 0 | 0 |
| 1369 | yeiP | 0 | 0 | 0 | 0 | 0 | 0 | 0 | 0 | 0 | 0 |
| 1370 | nfrB | 0 | 0 | 0 | 0 | 0 | 0 | 0 | 0 | 0 | 0 |
| 1371 | folX | 0 | 0 | 0 | 0 | 0 | 0 | 0 | 0 | 0 | 0 |
| 1372 | folE | 0 | 0 | 0 | 0 | 0 | 0 | 0 | 0 | 0 | 0 |
| 1373 | folD | 0 | 0 | 0 | 0 | 0 | 0 | 0 | 0 | 0 | 0 |
| 1374 | folA | 0 | 1 | 1 | 0 | 0 | 0 | 0 | 0 | 0 | 0 |
| 1375 | folC | 0 | 0 | 0 | 0 | 0 | 0 | 0 | 0 | 0 | 0 |
| 1376 | folB | 0 | 0 | 0 | 0 | 0 | 0 | 0 | 0 | 0 | 0 |
| 1377 | hydN | 0 | 0 | 0 | 0 | 0 | 0 | 0 | 0 | 0 | 0 |
| 1378 | rpoS | 0 | 0 | 0 | 0 | 0 | 0 | 0 | 0 | 0 | 0 |
| 1379 | kdtA | 0 | 0 | 0 | 0 | 0 | 0 | 0 | 0 | 0 | 0 |
| 1380 | rpoE | 0 | 0 | 0 | 0 | 0 | 0 | 0 | 0 | 0 | 0 |
| 1381 | rpoD | 0 | 0 | 0 | 0 | 0 | 0 | 0 | 0 | 0 | 0 |
| 1382 | rpoH | 0 | 0 | 0 | 0 | 0 | 0 | 0 | 0 | 1 | 0 |
| 1383 | yjaB | 18 | 0 | 0 | 0 | 42 | 0 | 30 | 0 | 0 | 0 |
| 1384 | yjaH | 0 | 0 | 0 | 0 | 0 | 0 | 0 | 0 | 0 | 0 |
| 1385 | nohB | 0 | 0 | 0 | 0 | 0 | 0 | 0 | 0 | 0 | 0 |

| | | | | | | | | | | | |
|---|---|---|---|---|---|---|---|---|---|---|---|
| 1386 | rpsG | 0 | 0 | 0 | 0 | 0 | 0 | 0 | 0 | 0 | 0 |
| 1387 | rpsB | 0 | 0 | 0 | 0 | 0 | 0 | 0 | 0 | 0 | 0 |
| 1388 | rpsA | 0 | 0 | 0 | 0 | 0 | 0 | 0 | 0 | 0 | 0 |
| 1389 | fdnG | 0 | 0 | 0 | 0 | 0 | 0 | 0 | 0 | 0 | 0 |
| 1390 | rpsO | 0 | 0 | 0 | 0 | 1 | 0 | 0 | 0 | 0 | 0 |
| 1391 | rpsM | 0 | 0 | 0 | 0 | 0 | 0 | 0 | 0 | 0 | 0 |
| 1392 | rpsL | 0 | 0 | 0 | 0 | 0 | 0 | 0 | 0 | 0 | 0 |
| 1393 | rpsJ | 0 | 0 | 0 | 0 | 0 | 0 | 0 | 0 | 0 | 0 |
| 1394 | apt | 0 | 0 | 0 | 0 | 0 | 0 | 0 | 0 | 0 | 0 |
| 1395 | rpsU | 0 | 0 | 0 | 0 | 0 | 0 | 0 | 0 | 0 | 0 |
| 1396 | rpsT | 1 | 0 | 0 | 0 | 0 | 0 | 0 | 0 | 0 | 0 |
| 1397 | rpsP | 0 | 0 | 0 | 0 | 0 | 0 | 0 | 0 | 0 | 0 |
| 1398 | deoA | 0 | 0 | 0 | 0 | 0 | 0 | 0 | 0 | 0 | 0 |
| 1399 | deoB | 0 | 0 | 0 | 0 | 0 | 0 | 0 | 0 | 0 | 0 |
| 1400 | deoC | 0 | 0 | 0 | 0 | 0 | 0 | 0 | 0 | 0 | 0 |
| 1401 | deoD | 0 | 0 | 0 | 0 | 0 | 0 | 0 | 0 | 0 | 0 |
| 1402 | usg | 0 | 0 | 0 | 0 | 0 | 0 | 0 | 0 | 0 | 0 |
| 1403 | sirA | 0 | 0 | 2 | 0 | 2 | 0 | 1 | 0 | 0 | 0 |
| 1404 | fes | 0 | 0 | 0 | 0 | 0 | 0 | 0 | 0 | 0 | 0 |
| 1405 | xasA | 0 | 0 | 0 | 0 | 0 | 0 | 0 | 0 | 0 | 0 |
| 1406 | pheL | 0 | 0 | 0 | 0 | 0 | 0 | 0 | 0 | 0 | 0 |
| 1407 | phoP | 0 | 0 | 0 | 0 | 0 | 0 | 0 | 0 | 0 | 0 |
| 1408 | phoR | 0 | 0 | 0 | 0 | 0 | 0 | 0 | 0 | 0 | 0 |
| 1409 | ndh | 0 | 0 | 0 | 0 | 0 | 0 | 0 | 1 | 0 | 1 |
| 1410 | yfhK | 0 | 0 | 0 | 0 | 0 | 0 | 0 | 0 | 0 | 0 |
| 1411 | phoA | 0 | 1 | 0 | 0 | 0 | 0 | 0 | 0 | 0 | 0 |
| 1412 | thiI | 0 | 0 | 0 | 0 | 0 | 0 | 0 | 0 | 0 | 0 |
| 1413 | phoB | 0 | 0 | 0 | 0 | 0 | 0 | 0 | 0 | 0 | 0 |
| 1414 | pinE | 0 | 0 | 0 | 0 | 0 | 0 | 0 | 0 | 0 | 0 |
| 1415 | thiM | 0 | 0 | 0 | 0 | 0 | 0 | 0 | 0 | 0 | 0 |
| 1416 | thiL | 0 | 0 | 0 | 0 | 0 | 0 | 0 | 0 | 0 | 0 |
| 1417 | thiC | 0 | 0 | 0 | 0 | 0 | 0 | 0 | 0 | 0 | 0 |
| 1418 | prlC | 0 | 0 | 0 | 0 | 0 | 0 | 0 | 0 | 0 | 0 |
| 1419 | thiG | 0 | 0 | 0 | 0 | 0 | 0 | 0 | 0 | 0 | 0 |
| 1420 | yiaU | 0 | 0 | 0 | 0 | 0 | 0 | 0 | 0 | 0 | 0 |
| 1421 | yiaT | 0 | 0 | 0 | 0 | 0 | 0 | 0 | 0 | 0 | 0 |
| 1422 | yiaW | 0 | 0 | 0 | 0 | 0 | 0 | 0 | 0 | 0 | 0 |
| 1423 | fabF | 55 | 0 | 12 | 0 | 80 | 0 | 53 | 0 | 0 | 0 |
| 1424 | yiaD | 0 | 0 | 0 | 0 | 0 | 0 | 0 | 0 | 0 | 0 |
| 1425 | yiaG | 0 | 0 | 0 | 0 | 0 | 0 | 0 | 0 | 0 | 0 |
| 1426 | yiaF | 0 | 0 | 0 | 0 | 0 | 0 | 0 | 0 | 0 | 0 |
| 1427 | yiaJ | 0 | 0 | 0 | 0 | 0 | 0 | 0 | 0 | 0 | 0 |
| 1428 | yqjH | 0 | 0 | 0 | 0 | 0 | 0 | 0 | 0 | 0 | 0 |
| 1429 | narX | 0 | 0 | 0 | 0 | 0 | 0 | 1 | 0 | 0 | 0 |
| 1430 | tar | 0 | 0 | 0 | 0 | 0 | 0 | 0 | 0 | 0 | 0 |
| 1431 | narZ | 0 | 0 | 0 | 0 | 0 | 0 | 0 | 0 | 0 | 0 |
| 1432 | narU | 0 | 0 | 0 | 0 | 0 | 0 | 0 | 0 | 0 | 0 |
| 1433 | cbdA | 0 | 0 | 0 | 0 | 0 | 0 | 0 | 0 | 0 | 0 |
| 1434 | narQ | 0 | 0 | 0 | 0 | 0 | 0 | 0 | 0 | 0 | 0 |
| 1435 | tam | 0 | 0 | 0 | 0 | 0 | 0 | 0 | 0 | 0 | 0 |
| 1436 | gcl | 0 | 0 | 0 | 0 | 0 | 0 | 0 | 0 | 0 | 0 |
| 1437 | yneF | 0 | 0 | 0 | 0 | 0 | 0 | 0 | 0 | 0 | 0 |
| 1438 | paaX | 0 | 0 | 0 | 0 | 0 | 0 | 0 | 0 | 0 | 0 |
| 1439 | yrbL | 0 | 0 | 0 | 0 | 0 | 0 | 0 | 0 | 0 | 0 |
| 1440 | yrbG | 0 | 0 | 0 | 0 | 0 | 0 | 0 | 0 | 0 | 0 |
| 1441 | yrbF | 0 | 0 | 0 | 0 | 0 | 0 | 0 | 0 | 0 | 0 |
| 1442 | yrbA | 0 | 0 | 0 | 0 | 0 | 0 | 0 | 0 | 0 | 0 |
| 1443 | kbl | 0 | 0 | 0 | 0 | 0 | 0 | 0 | 0 | 0 | 0 |
| 1444 | dmsA | 0 | 0 | 0 | 0 | 0 | 0 | 0 | 0 | 0 | 0 |
| 1445 | ruvA | 0 | 0 | 0 | 0 | 0 | 0 | 0 | 0 | 0 | 0 |
| 1446 | tsf | 0 | 0 | 0 | 0 | 0 | 0 | 0 | 0 | 0 | 0 |
| 1447 | mutH | 0 | 0 | 0 | 0 | 0 | 0 | 0 | 0 | 0 | 0 |
| 1448 | ynjF | 0 | 0 | 0 | 0 | 0 | 0 | 0 | 0 | 0 | 0 |
| 1449 | mutM | 0 | 0 | 0 | 0 | 0 | 0 | 0 | 0 | 0 | 0 |
| 1450 | mutS | 21 | 0 | 10 | 1 | 41 | 0 | 9 | 1 | 38 | 0 |
| 1451 | mutT | 0 | 0 | 0 | 0 | 0 | 0 | 0 | 0 | 0 | 0 |

| | | | | | | | | | | | |
|---|---|---|---|---|---|---|---|---|---|---|---|
| 1452 | mutY | 0 | 0 | 0 | 0 | 0 | 0 | 0 | 0 | 0 | 0 |
| 1453 | ydjX | 0 | 0 | 0 | 0 | 0 | 0 | 0 | 0 | 0 | 0 |
| 1454 | ydjA | 0 | 0 | 0 | 0 | 0 | 0 | 0 | 0 | 0 | 0 |
| 1455 | ydjE | 0 | 0 | 0 | 0 | 0 | 0 | 0 | 0 | 0 | 0 |
| 1456 | b1420 | 0 | 0 | 0 | 0 | 1 | 0 | 2 | 0 | 0 | 0 |
| 1457 | ydjF | 0 | 0 | 0 | 0 | 0 | 0 | 0 | 0 | 0 | 0 |
| 1458 | ydjL | 85 | 0 | 88 | 41 | 178 | 71 | 159 | 4191 | 183 | 6151 |
| 1459 | ydjN | 0 | 0 | 0 | 0 | 0 | 0 | 0 | 0 | 0 | 0 |
| 1460 | nadK | 0 | 0 | 0 | 0 | 0 | 0 | 0 | 0 | 0 | 0 |
| 1461 | yadK | 0 | 0 | 0 | 0 | 0 | 0 | 0 | 0 | 0 | 0 |
| 1462 | cusR | 0 | 0 | 0 | 0 | 0 | 0 | 0 | 0 | 0 | 0 |
| 1463 | yadN | 0 | 0 | 0 | 0 | 0 | 0 | 0 | 0 | 0 | 0 |
| 1464 | nfo | 0 | 0 | 1 | 0 | 0 | 0 | 1 | 0 | 0 | 0 |
| 1465 | yadC | 0 | 0 | 0 | 0 | 0 | 0 | 0 | 0 | 0 | 0 |
| 1466 | yfjI | 0 | 0 | 0 | 0 | 0 | 0 | 0 | 0 | 0 | 0 |
| 1467 | yadF | 0 | 0 | 0 | 0 | 0 | 0 | 0 | 0 | 0 | 0 |
| 1468 | yfjN | 13 | 0 | 4 | 0 | 46 | 0 | 20 | 0 | 21 | 0 |
| 1469 | yfjM | 0 | 0 | 0 | 0 | 0 | 0 | 0 | 0 | 0 | 0 |
| 1470 | yfjL | 0 | 0 | 0 | 0 | 0 | 0 | 0 | 0 | 0 | 0 |
| 1471 | yfjR | 65 | 0 | 15 | 0 | 108 | 0 | 21 | 0 | 0 | 0 |
| 1472 | cusC | 0 | 0 | 0 | 0 | 0 | 0 | 1 | 0 | 0 | 0 |
| 1473 | yfjP | 0 | 0 | 0 | 0 | 0 | 0 | 0 | 0 | 0 | 0 |
| 1474 | yfjW | 0 | 0 | 0 | 0 | 0 | 0 | 0 | 0 | 0 | 0 |
| 1475 | yadR | 0 | 0 | 1 | 0 | 0 | 0 | 0 | 0 | 0 | 0 |
| 1476 | yadS | 0 | 0 | 0 | 0 | 0 | 0 | 0 | 0 | 0 | 0 |
| 1477 | psiE | 0 | 0 | 1 | 0 | 0 | 0 | 0 | 0 | 0 | 0 |
| 1478 | psiF | 0 | 0 | 0 | 0 | 0 | 0 | 0 | 0 | 0 | 0 |
| 1479 | rplY | 0 | 0 | 0 | 0 | 0 | 0 | 0 | 0 | 0 | 0 |
| 1480 | rplT | 0 | 0 | 0 | 0 | 0 | 0 | 0 | 0 | 0 | 0 |
| 1481 | hlyE | 0 | 0 | 0 | 0 | 0 | 0 | 0 | 0 | 0 | 0 |
| 1482 | rplN | 0 | 0 | 0 | 0 | 0 | 0 | 0 | 0 | 0 | 0 |
| 1483 | kdsB | 0 | 0 | 0 | 0 | 0 | 0 | 0 | 0 | 0 | 0 |
| 1484 | metZ | 0 | 0 | 0 | 0 | 0 | 0 | 0 | 0 | 0 | 0 |
| 1485 | metB | 0 | 0 | 0 | 0 | 0 | 0 | 0 | 0 | 0 | 0 |
| 1486 | metC | 0 | 0 | 0 | 0 | 0 | 0 | 0 | 0 | 0 | 0 |
| 1487 | mrcB | 0 | 0 | 0 | 0 | 0 | 0 | 0 | 0 | 0 | 0 |
| 1488 | metA | 0 | 0 | 0 | 0 | 0 | 0 | 0 | 0 | 0 | 0 |
| 1489 | metF | 0 | 0 | 0 | 0 | 0 | 0 | 0 | 0 | 0 | 0 |
| 1490 | metG | 0 | 0 | 0 | 0 | 0 | 0 | 0 | 0 | 0 | 0 |
| 1491 | metJ | 0 | 0 | 0 | 0 | 0 | 0 | 0 | 0 | 0 | 0 |
| 1492 | metK | 15 | 0 | 2 | 0 | 23 | 0 | 3 | 0 | 0 | 0 |
| 1493 | yhbE | 0 | 0 | 0 | 0 | 0 | 0 | 0 | 0 | 0 | 0 |
| 1494 | ycjW | 0 | 0 | 0 | 0 | 0 | 0 | 0 | 0 | 0 | 0 |
| 1495 | yhbC | 0 | 0 | 0 | 0 | 0 | 0 | 1 | 0 | 0 | 0 |
| 1496 | ycjZ | 0 | 0 | 0 | 0 | 0 | 0 | 0 | 0 | 0 | 0 |
| 1497 | ycjX | 0 | 0 | 0 | 0 | 0 | 0 | 0 | 0 | 0 | 0 |
| 1498 | ycjY | 0 | 0 | 0 | 0 | 0 | 0 | 0 | 0 | 0 | 0 |
| 1499 | yhbU | 0 | 0 | 0 | 0 | 0 | 0 | 0 | 0 | 0 | 0 |
| 1500 | yhbT | 0 | 0 | 0 | 0 | 0 | 0 | 0 | 0 | 0 | 0 |
| 1501 | yhbW | 54 | 0 | 13 | 1 | 103 | 0 | 69 | 0 | 53 | 0 |
| 1502 | yhbQ | 46 | 0 | 6 | 1 | 92 | 0 | 41 | 0 | 16 | 0 |
| 1503 | ycjG | 0 | 0 | 0 | 0 | 0 | 0 | 0 | 0 | 0 | 0 |
| 1504 | ycjK | 0 | 0 | 0 | 0 | 0 | 0 | 0 | 0 | 0 | 0 |
| 1505 | yhbX | 0 | 0 | 0 | 0 | 0 | 0 | 0 | 0 | 1 | 0 |
| 1506 | ycjL | 0 | 0 | 0 | 0 | 0 | 0 | 0 | 0 | 0 | 0 |
| 1507 | ycjM | 0 | 0 | 0 | 0 | 0 | 0 | 0 | 0 | 0 | 0 |
| 1508 | ymjA | 0 | 0 | 0 | 0 | 0 | 0 | 0 | 0 | 0 | 0 |
| 1509 | rhoL | 0 | 0 | 0 | 0 | 0 | 0 | 0 | 0 | 0 | 0 |
| 1510 | nrfE | 0 | 0 | 0 | 0 | 0 | 0 | 0 | 0 | 0 | 0 |
| 1511 | pgk | 0 | 0 | 0 | 0 | 0 | 0 | 0 | 0 | 0 | 0 |
| 1512 | pgi | 2 | 0 | 1 | 0 | 5 | 0 | 5 | 0 | 0 | 0 |
| 1513 | potF | 0 | 1 | 0 | 0 | 0 | 0 | 0 | 0 | 0 | 0 |
| 1514 | potA | 0 | 0 | 0 | 0 | 0 | 0 | 0 | 0 | 0 | 0 |
| 1515 | mviN | 0 | 0 | 0 | 0 | 0 | 0 | 0 | 0 | 0 | 0 |
| 1516 | gatR1 | 0 | 0 | 0 | 0 | 0 | 0 | 0 | 0 | 0 | 0 |
| 1517 | pflA | 0 | 0 | 0 | 0 | 0 | 0 | 0 | 0 | 0 | 0 |

| | | | | | | | | | | | |
|---|---|---|---|---|---|---|---|---|---|---|---|
| 1518 | ahpF | 1 | 1 | 1 | 3 | 1 | 0 | 0 | 0 | 1 | 1 |
| 1519 | ahpC | 0 | 0 | 0 | 0 | 0 | 0 | 0 | 0 | 0 | 0 |
| 1520 | udp | 0 | 0 | 0 | 0 | 0 | 0 | 0 | 0 | 0 | 0 |
| 1521 | yqcE | 0 | 0 | 0 | 0 | 0 | 0 | 0 | 0 | 0 | 0 |
| 1522 | yqcD | 0 | 0 | 0 | 0 | 0 | 0 | 0 | 0 | 0 | 0 |
| 1523 | rsd | 1 | 0 | 0 | 0 | 0 | 0 | 0 | 0 | 0 | 0 |
| 1524 | udk | 0 | 0 | 0 | 0 | 0 | 0 | 0 | 0 | 0 | 0 |
| 1525 | xylB | 0 | 0 | 0 | 0 | 0 | 0 | 0 | 0 | 0 | 0 |
| 1526 | xylE | 0 | 0 | 0 | 0 | 0 | 0 | 0 | 0 | 0 | 0 |
| 1527 | xylG | 0 | 0 | 0 | 0 | 0 | 0 | 0 | 0 | 0 | 0 |
| 1528 | xylF | 0 | 0 | 0 | 0 | 0 | 0 | 0 | 0 | 0 | 0 |
| 1529 | dnaQ | 0 | 0 | 0 | 0 | 0 | 0 | 0 | 0 | 0 | 0 |
| 1530 | dnaX | 0 | 0 | 0 | 0 | 0 | 0 | 0 | 0 | 0 | 0 |
| 1531 | rdoA | 1 | 0 | 0 | 0 | 0 | 0 | 0 | 0 | 0 | 0 |
| 1532 | dnaK | 0 | 0 | 0 | 0 | 0 | 0 | 0 | 0 | 0 | 0 |
| 1533 | dnaJ | 0 | 0 | 0 | 0 | 0 | 0 | 0 | 0 | 0 | 0 |
| 1534 | sdhC | 0 | 0 | 0 | 0 | 0 | 0 | 0 | 0 | 0 | 0 |
| 1535 | trpL | 0 | 0 | 0 | 0 | 0 | 0 | 0 | 0 | 0 | 0 |
| 1536 | yraR | 0 | 0 | 0 | 0 | 0 | 0 | 0 | 0 | 0 | 0 |
| 1537 | yraQ | 0 | 0 | 0 | 0 | 0 | 0 | 0 | 0 | 0 | 0 |
| 1538 | vacJ | 0 | 0 | 0 | 0 | 0 | 0 | 0 | 0 | 0 | 0 |
| 1539 | yliE | 0 | 0 | 0 | 0 | 0 | 0 | 0 | 0 | 0 | 0 |
| 1540 | yraL | 0 | 0 | 0 | 0 | 0 | 0 | 0 | 0 | 0 | 0 |
| 1541 | yraM | 0 | 0 | 0 | 0 | 0 | 0 | 0 | 0 | 0 | 0 |
| 1542 | trpR | 0 | 0 | 0 | 0 | 0 | 0 | 0 | 0 | 0 | 0 |
| 1543 | yliI | 0 | 0 | 0 | 0 | 0 | 0 | 0 | 0 | 0 | 0 |
| 1544 | fdoG | 0 | 0 | 0 | 0 | 0 | 0 | 0 | 0 | 0 | 0 |
| 1545 | rfaQ | 0 | 0 | 0 | 0 | 0 | 0 | 0 | 0 | 0 | 0 |
| 1546 | rfaZ | 0 | 0 | 0 | 0 | 0 | 0 | 0 | 0 | 0 | 0 |
| 1547 | rfaD | 0 | 0 | 0 | 0 | 0 | 0 | 0 | 0 | 0 | 0 |
| 1548 | rfaH | 0 | 0 | 0 | 0 | 0 | 0 | 0 | 0 | 0 | 0 |
| 1549 | pstS | 0 | 0 | 0 | 0 | 0 | 0 | 0 | 0 | 0 | 0 |
| 1550 | evgA | 0 | 0 | 0 | 0 | 0 | 0 | 0 | 0 | 0 | 0 |
| 1551 | envY | 0 | 0 | 0 | 0 | 0 | 0 | 0 | 0 | 0 | 0 |
| 1552 | ydeV | 0 | 0 | 0 | 0 | 0 | 0 | 0 | 0 | 0 | 0 |
| 1553 | ydeP | 0 | 0 | 0 | 0 | 0 | 0 | 0 | 0 | 0 | 0 |
| 1554 | ydeQ | 0 | 0 | 0 | 0 | 0 | 0 | 0 | 0 | 0 | 0 |
| 1555 | btuC | 0 | 0 | 0 | 0 | 0 | 0 | 0 | 0 | 0 | 0 |
| 1556 | btuB | 0 | 0 | 0 | 0 | 0 | 0 | 0 | 0 | 0 | 0 |
| 1557 | pstC | 1 | 0 | 0 | 0 | 0 | 0 | 0 | 0 | 0 | 0 |
| 1558 | moeA | 0 | 0 | 0 | 0 | 0 | 0 | 0 | 0 | 0 | 0 |
| 1559 | ydeO | 0 | 0 | 0 | 0 | 0 | 0 | 0 | 0 | 0 | 0 |
| 1560 | ydeI | 0 | 0 | 0 | 0 | 0 | 0 | 0 | 0 | 0 | 0 |
| 1561 | ydeJ | 0 | 0 | 0 | 0 | 0 | 0 | 0 | 0 | 0 | 0 |
| 1562 | ydeA | 0 | 0 | 0 | 0 | 0 | 0 | 0 | 0 | 0 | 0 |
| 1563 | yhdX | 0 | 0 | 0 | 0 | 0 | 0 | 0 | 0 | 0 | 0 |
| 1564 | yaeQ | 0 | 0 | 0 | 0 | 0 | 0 | 0 | 0 | 0 | 0 |
| 1565 | yfiF | 0 | 0 | 0 | 0 | 0 | 0 | 0 | 0 | 0 | 0 |
| 1566 | feaB | 0 | 0 | 0 | 0 | 0 | 0 | 0 | 0 | 0 | 0 |
| 1567 | yfiE | 0 | 0 | 0 | 0 | 0 | 0 | 0 | 0 | 0 | 0 |
| 1568 | yfiC | 0 | 0 | 0 | 0 | 0 | 0 | 0 | 0 | 0 | 0 |
| 1569 | nagE | 0 | 0 | 0 | 0 | 0 | 0 | 0 | 0 | 0 | 0 |
| 1570 | nagB | 0 | 0 | 0 | 0 | 0 | 0 | 0 | 0 | 0 | 0 |
| 1571 | nagA | 0 | 0 | 0 | 0 | 0 | 0 | 0 | 0 | 0 | 0 |
| 1572 | feaR | 0 | 0 | 0 | 0 | 0 | 0 | 0 | 0 | 0 | 0 |
| 1573 | yaeH | 0 | 0 | 0 | 0 | 0 | 0 | 0 | 0 | 0 | 0 |
| 1574 | uidC | 0 | 0 | 0 | 0 | 0 | 0 | 0 | 0 | 0 | 0 |
| 1575 | uidA | 41 | 0 | 35 | 0 | 60 | 0 | 32 | 0 | 66 | 0 |
| 1576 | aes | 0 | 0 | 0 | 0 | 0 | 0 | 0 | 0 | 0 | 0 |
| 1577 | aer | 0 | 0 | 0 | 0 | 0 | 0 | 0 | 0 | 0 | 0 |
| 1578 | dps | 0 | 0 | 0 | 0 | 0 | 0 | 0 | 0 | 0 | 0 |
| 1579 | b0725 | 0 | 0 | 0 | 0 | 0 | 0 | 0 | 0 | 0 | 0 |
| 1580 | aroP | 0 | 0 | 0 | 0 | 0 | 0 | 0 | 0 | 0 | 0 |
| 1581 | tyrP | 0 | 0 | 0 | 0 | 0 | 0 | 0 | 0 | 0 | 0 |
| 1582 | yfgA | 0 | 0 | 0 | 0 | 0 | 0 | 0 | 0 | 0 | 0 |
| 1583 | yfgB | 0 | 0 | 0 | 0 | 0 | 0 | 0 | 0 | 0 | 0 |

| | | | | | | | | | | | |
|---|---|---|---|---|---|---|---|---|---|---|---|
| 1584 | yfgH | 0 | 0 | 1 | 0 | 0 | 0 | 0 | 0 | 0 | 0 |
| 1585 | yheO | 0 | 0 | 0 | 0 | 0 | 0 | 0 | 0 | 0 | 0 |
| 1586 | tyrB | 0 | 0 | 0 | 0 | 0 | 0 | 0 | 0 | 0 | 0 |
| 1587 | yheS | 0 | 0 | 0 | 0 | 0 | 0 | 0 | 0 | 0 | 0 |
| 1588 | aroF | 0 | 0 | 0 | 0 | 0 | 0 | 0 | 0 | 0 | 0 |
| 1589 | aroH | 0 | 0 | 0 | 0 | 0 | 0 | 0 | 0 | 0 | 0 |
| 1590 | aroK | 0 | 0 | 0 | 0 | 0 | 0 | 0 | 0 | 0 | 0 |
| 1591 | kefF | 0 | 1 | 0 | 0 | 0 | 0 | 0 | 0 | 0 | 0 |
| 1592 | kefG | 0 | 0 | 0 | 0 | 0 | 0 | 0 | 0 | 0 | 0 |
| 1593 | uvrD | 0 | 0 | 0 | 0 | 0 | 0 | 0 | 0 | 0 | 0 |
| 1594 | uvrC | 0 | 0 | 0 | 0 | 0 | 0 | 0 | 0 | 0 | 0 |
| 1595 | b4283 | 0 | 0 | 0 | 0 | 0 | 0 | 0 | 0 | 0 | 0 |
| 1596 | yidI | 0 | 0 | 0 | 0 | 0 | 0 | 0 | 0 | 0 | 0 |
| 1597 | ycfQ | 0 | 0 | 0 | 0 | 0 | 0 | 0 | 0 | 0 | 0 |
| 1598 | rsuA | 0 | 0 | 0 | 0 | 0 | 0 | 0 | 0 | 0 | 0 |
| 1599 | aegA | 0 | 0 | 0 | 0 | 0 | 0 | 0 | 0 | 0 | 0 |
| 1600 | yniC | 0 | 0 | 0 | 0 | 0 | 0 | 0 | 0 | 0 | 0 |
| 1601 | yniA | 0 | 0 | 0 | 0 | 0 | 0 | 0 | 0 | 0 | 0 |
| 1602 | caiC | 0 | 0 | 0 | 0 | 0 | 0 | 0 | 0 | 0 | 0 |
| 1603 | rumA | 9 | 0 | 5 | 1 | 30 | 0 | 12 | 0 | 0 | 0 |
| 1604 | ttdA | 0 | 0 | 0 | 0 | 0 | 0 | 0 | 0 | 0 | 0 |
| 1605 | rpe | 0 | 0 | 0 | 0 | 0 | 0 | 0 | 0 | 0 | 0 |
| 1606 | rph | 9 | 0 | 1 | 0 | 19 | 0 | 7 | 0 | 8 | 0 |
| 1607 | nhoA | 0 | 0 | 0 | 0 | 0 | 0 | 0 | 0 | 0 | 0 |
| 1608 | rhlE | 0 | 0 | 0 | 0 | 0 | 0 | 0 | 0 | 0 | 0 |
| 1609 | rhlB | 0 | 0 | 0 | 0 | 0 | 0 | 0 | 0 | 0 | 0 |
| 1610 | rpmI | 0 | 0 | 0 | 0 | 0 | 0 | 0 | 0 | 0 | 0 |
| 1611 | agaR | 0 | 0 | 0 | 0 | 0 | 0 | 0 | 0 | 0 | 0 |
| 1612 | rpmB | 0 | 0 | 0 | 0 | 0 | 0 | 0 | 0 | 0 | 0 |
| 1613 | rpmE | 0 | 0 | 0 | 0 | 0 | 0 | 0 | 0 | 0 | 0 |
| 1614 | lgt | 0 | 0 | 0 | 0 | 0 | 0 | 0 | 0 | 0 | 0 |
| 1615 | yjgH | 0 | 0 | 0 | 0 | 0 | 0 | 0 | 0 | 0 | 0 |
| 1616 | yjgJ | 0 | 0 | 0 | 0 | 0 | 0 | 0 | 0 | 0 | 0 |
| 1617 | yddA | 0 | 0 | 0 | 0 | 0 | 0 | 0 | 0 | 0 | 0 |
| 1618 | yjgA | 0 | 0 | 0 | 0 | 0 | 0 | 0 | 0 | 0 | 0 |
| 1619 | yjgR | 0 | 0 | 0 | 0 | 0 | 0 | 0 | 0 | 0 | 0 |
| 1620 | fabR | 4 | 3022 | 767 | 3455 | 9 | 1777 | 27 | 325 | 3 | 194 |
| 1621 | fabZ | 0 | 0 | 0 | 2 | 0 | 0 | 0 | 0 | 0 | 0 |
| 1622 | rfbB | 0 | 0 | 0 | 0 | 0 | 0 | 0 | 0 | 0 | 0 |
| 1623 | rfbA | 0 | 0 | 0 | 0 | 0 | 0 | 0 | 0 | 0 | 0 |
| 1624 | cpsG | 0 | 0 | 0 | 0 | 0 | 0 | 0 | 0 | 0 | 0 |
| 1625 | ygaV | 0 | 0 | 0 | 0 | 0 | 0 | 0 | 0 | 0 | 0 |
| 1626 | fkpA | 0 | 0 | 0 | 0 | 0 | 0 | 0 | 0 | 0 | 0 |
| 1627 | dam | 0 | 0 | 0 | 0 | 0 | 0 | 0 | 0 | 0 | 0 |
| 1628 | ackA | 0 | 0 | 0 | 0 | 0 | 0 | 0 | 0 | 0 | 0 |
| 1629 | setA | 0 | 0 | 0 | 0 | 0 | 0 | 0 | 0 | 0 | 0 |
| 1630 | setB | 0 | 0 | 0 | 0 | 0 | 0 | 0 | 0 | 0 | 0 |
| 1631 | mglB | 0 | 0 | 0 | 0 | 0 | 0 | 0 | 0 | 0 | 0 |
| 1632 | fhlA | 0 | 0 | 0 | 0 | 0 | 0 | 0 | 0 | 0 | 0 |
| 1633 | iscS | 0 | 0 | 0 | 0 | 0 | 0 | 0 | 0 | 0 | 0 |
| 1634 | iscR | 0 | 0 | 0 | 0 | 0 | 0 | 0 | 0 | 0 | 0 |
| 1635 | dusA | 0 | 0 | 0 | 0 | 0 | 0 | 0 | 0 | 0 | 0 |
| 1636 | dusB | 0 | 0 | 0 | 0 | 0 | 0 | 0 | 0 | 0 | 0 |
| 1637 | dusC | 0 | 0 | 0 | 0 | 0 | 0 | 0 | 0 | 0 | 0 |
| 1638 | nfnB | 0 | 0 | 0 | 0 | 0 | 0 | 0 | 0 | 0 | 0 |
| 1639 | b2639 | 0 | 0 | 0 | 0 | 0 | 0 | 0 | 0 | 0 | 0 |
| 1640 | ycfD | 1 | 0 | 0 | 0 | 1 | 0 | 2 | 0 | 0 | 0 |
| 1641 | mfd | 0 | 0 | 0 | 0 | 0 | 0 | 0 | 0 | 0 | 0 |
| 1642 | modF | 0 | 0 | 0 | 0 | 0 | 0 | 0 | 0 | 0 | 0 |
| 1643 | gutM | 0 | 0 | 0 | 0 | 0 | 0 | 0 | 0 | 0 | 0 |
| 1644 | gmk | 0 | 0 | 0 | 0 | 0 | 0 | 0 | 0 | 0 | 0 |
| 1645 | skp | 0 | 0 | 0 | 0 | 0 | 0 | 0 | 0 | 0 | 0 |
| 1646 | sufI | 0 | 0 | 0 | 0 | 0 | 0 | 0 | 0 | 0 | 0 |
| 1647 | add | 0 | 0 | 0 | 0 | 0 | 0 | 0 | 0 | 0 | 0 |
| 1648 | ada | 0 | 0 | 0 | 0 | 0 | 0 | 0 | 0 | 0 | 0 |
| 1649 | gmr | 0 | 0 | 0 | 0 | 0 | 0 | 0 | 0 | 0 | 0 |

| | | | | | | | | | | | |
|---|---|---|---|---|---|---|---|---|---|---|---|
| 1650 | adk | 0 | 0 | 0 | 0 | 0 | 0 | 0 | 0 | 0 | 0 |
| 1651 | eutN | 0 | 0 | 0 | 0 | 0 | 0 | 0 | 0 | 0 | 0 |
| 1652 | ppdD | 0 | 0 | 0 | 0 | 0 | 0 | 0 | 0 | 0 | 0 |
| 1653 | otsB | 0 | 0 | 0 | 0 | 0 | 0 | 0 | 0 | 0 | 0 |
| 1654 | yfjG | 0 | 0 | 0 | 0 | 0 | 0 | 0 | 0 | 0 | 0 |
| 1655 | pagB | 0 | 0 | 0 | 0 | 0 | 0 | 0 | 0 | 0 | 0 |
| 1656 | nadC | 0 | 0 | 0 | 0 | 0 | 0 | 0 | 0 | 0 | 0 |
| 1657 | b0663 | 0 | 0 | 1 | 0 | 0 | 0 | 0 | 0 | 0 | 0 |
| 1658 | yadG | 0 | 0 | 0 | 0 | 0 | 0 | 0 | 0 | 0 | 0 |
| 1659 | rstA | 0 | 0 | 0 | 0 | 0 | 0 | 0 | 0 | 0 | 0 |
| 1660 | grpE | 0 | 0 | 0 | 0 | 0 | 0 | 0 | 0 | 0 | 0 |
| 1661 | murA | 0 | 0 | 0 | 0 | 0 | 0 | 0 | 0 | 0 | 0 |
| 1662 | murC | 0 | 0 | 0 | 0 | 0 | 0 | 0 | 0 | 0 | 0 |
| 1663 | molR2 | 0 | 0 | 0 | 0 | 0 | 0 | 0 | 0 | 0 | 0 |
| 1664 | dcm | 0 | 0 | 0 | 0 | 0 | 0 | 0 | 0 | 0 | 0 |
| 1665 | yddW | 0 | 0 | 0 | 0 | 0 | 0 | 0 | 0 | 0 | 0 |
| 1666 | dcd | 0 | 0 | 0 | 0 | 0 | 0 | 0 | 0 | 0 | 0 |
| 1667 | yqaD | 0 | 0 | 0 | 0 | 0 | 0 | 0 | 0 | 0 | 0 |
| 1668 | nohA | 0 | 0 | 0 | 0 | 0 | 0 | 0 | 0 | 0 | 0 |
| 1669 | gudP | 0 | 0 | 0 | 0 | 0 | 0 | 0 | 0 | 0 | 0 |
| 1670 | yddM | 0 | 0 | 0 | 0 | 0 | 0 | 0 | 0 | 0 | 0 |
| 1671 | yddL | 0 | 0 | 0 | 0 | 0 | 0 | 0 | 0 | 0 | 0 |
| 1672 | yddH | 0 | 0 | 0 | 0 | 0 | 0 | 0 | 0 | 0 | 0 |
| 1673 | yddE | 0 | 0 | 0 | 0 | 0 | 0 | 0 | 0 | 0 | 0 |
| 1674 | dcp | 0 | 0 | 0 | 0 | 0 | 0 | 0 | 0 | 0 | 0 |
| 1675 | b1403 | 0 | 0 | 0 | 0 | 0 | 0 | 0 | 0 | 0 | 0 |
| 1676 | yabP | 0 | 0 | 0 | 0 | 0 | 0 | 0 | 0 | 0 | 0 |
| 1677 | yfhH | 0 | 0 | 0 | 0 | 0 | 0 | 0 | 0 | 0 | 0 |
| 1678 | ndk | 0 | 0 | 0 | 0 | 0 | 0 | 0 | 0 | 0 | 0 |
| 1679 | yfhM | 0 | 0 | 0 | 0 | 0 | 0 | 0 | 0 | 0 | 0 |
| 1680 | yfhL | 0 | 0 | 0 | 0 | 0 | 0 | 0 | 0 | 0 | 0 |
| 1681 | yfhB | 0 | 0 | 0 | 0 | 0 | 0 | 0 | 0 | 0 | 0 |
| 1682 | yfhG | 0 | 0 | 0 | 0 | 0 | 0 | 0 | 0 | 0 | 0 |
| 1683 | yabI | 0 | 0 | 0 | 0 | 0 | 0 | 0 | 0 | 0 | 0 |
| 1684 | yfhR | 0 | 0 | 0 | 0 | 0 | 0 | 0 | 0 | 0 | 0 |
| 1685 | yabN | 0 | 0 | 0 | 0 | 0 | 0 | 0 | 0 | 0 | 0 |
| 1686 | cvrA | 0 | 0 | 0 | 0 | 0 | 0 | 0 | 0 | 0 | 0 |
| 1687 | yaaU | 87 | 0 | 21 | 1 | 149 | 0 | 102 | 0 | 13 | 0 |
| 1688 | mazE | 0 | 0 | 0 | 0 | 0 | 0 | 0 | 0 | 0 | 0 |
| 1689 | mazG | 0 | 0 | 0 | 0 | 0 | 0 | 0 | 0 | 0 | 0 |
| 1690 | appA | 0 | 0 | 0 | 0 | 0 | 0 | 0 | 0 | 0 | 0 |
| 1691 | yjfK | 0 | 0 | 0 | 0 | 0 | 0 | 0 | 0 | 0 | 0 |
| 1692 | yjfI | 0 | 0 | 0 | 0 | 0 | 0 | 0 | 0 | 0 | 0 |
| 1693 | mraZ | 0 | 0 | 0 | 0 | 0 | 0 | 0 | 0 | 0 | 0 |
| 1694 | ydaQ | 0 | 0 | 0 | 0 | 0 | 0 | 0 | 0 | 0 | 0 |
| 1695 | hdhA | 0 | 0 | 0 | 0 | 0 | 0 | 0 | 0 | 0 | 0 |
| 1696 | yjfY | 158 | 1 | 53 | 0 | 297 | 1 | 78 | 0 | 139 | 0 |
| 1697 | yjfQ | 0 | 0 | 0 | 0 | 0 | 0 | 0 | 1 | 0 | 0 |
| 1698 | yhdA | 0 | 0 | 0 | 0 | 0 | 0 | 0 | 0 | 0 | 0 |
| 1699 | fliY | 0 | 0 | 0 | 0 | 0 | 0 | 0 | 0 | 0 | 0 |
| 1700 | yhdJ | 0 | 0 | 0 | 0 | 0 | 0 | 0 | 0 | 0 | 0 |
| 1701 | yhdH | 0 | 0 | 0 | 0 | 0 | 0 | 0 | 0 | 0 | 0 |
| 1702 | yhdN | 0 | 0 | 0 | 0 | 0 | 0 | 0 | 0 | 0 | 0 |
| 1703 | yffH | 0 | 0 | 0 | 1 | 0 | 0 | 0 | 0 | 0 | 0 |
| 1704 | fliL | 0 | 0 | 0 | 0 | 0 | 0 | 0 | 0 | 0 | 0 |
| 1705 | yffS | 0 | 0 | 0 | 0 | 0 | 0 | 0 | 0 | 0 | 0 |
| 1706 | fliE | 0 | 0 | 0 | 0 | 0 | 0 | 0 | 0 | 0 | 0 |
| 1707 | yhdY | 1 | 0 | 0 | 0 | 0 | 0 | 0 | 0 | 0 | 0 |
| 1708 | fliF | 0 | 0 | 0 | 0 | 0 | 0 | 0 | 0 | 0 | 0 |
| 1709 | fliA | 0 | 0 | 0 | 0 | 0 | 0 | 0 | 0 | 0 | 0 |
| 1710 | fliC | 0 | 0 | 0 | 0 | 0 | 0 | 0 | 0 | 0 | 0 |
| 1711 | yaiA | 0 | 0 | 0 | 0 | 0 | 0 | 0 | 0 | 0 | 0 |
| 1712 | rplL | 0 | 0 | 0 | 0 | 0 | 0 | 0 | 0 | 0 | 0 |
| 1713 | srmB | 0 | 0 | 0 | 0 | 0 | 0 | 0 | 0 | 0 | 0 |
| 1714 | mglA | 0 | 0 | 0 | 0 | 0 | 0 | 0 | 0 | 0 | 0 |
| 1715 | yohJ | 0 | 0 | 0 | 0 | 0 | 0 | 0 | 0 | 0 | 0 |

| | | | | | | | | | | | |
|---|---|---|---|---|---|---|---|---|---|---|---|
| 1716 | yohL | 0 | 0 | 0 | 0 | 0 | 0 | 0 | 0 | 0 | 0 |
| 1717 | yohM | 0 | 0 | 0 | 0 | 0 | 0 | 0 | 1 | 0 | 0 |
| 1718 | pdxY | 0 | 0 | 0 | 0 | 0 | 0 | 0 | 0 | 0 | 0 |
| 1719 | yohC | 0 | 0 | 0 | 0 | 0 | 0 | 0 | 0 | 0 | 0 |
| 1720 | yohD | 0 | 0 | 0 | 0 | 0 | 0 | 0 | 0 | 0 | 0 |
| 1721 | yaiB | 0 | 0 | 0 | 0 | 0 | 0 | 0 | 0 | 0 | 0 |
| 1722 | yeaE | 1 | 0 | 0 | 0 | 0 | 0 | 0 | 0 | 0 | 0 |
| 1723 | yqaC | 0 | 0 | 0 | 0 | 0 | 0 | 0 | 0 | 0 | 0 |
| 1724 | yqaB | 0 | 0 | 0 | 0 | 0 | 0 | 0 | 0 | 0 | 0 |
| 1725 | yqaE | 0 | 0 | 0 | 0 | 0 | 0 | 0 | 0 | 0 | 0 |
| 1726 | nrdB | 0 | 0 | 0 | 0 | 0 | 0 | 0 | 0 | 0 | 0 |
| 1727 | nrdD | 0 | 0 | 0 | 0 | 0 | 0 | 0 | 0 | 0 | 0 |
| 1728 | rhsE | 0 | 0 | 0 | 0 | 0 | 0 | 0 | 0 | 0 | 0 |
| 1729 | rhsD | 0 | 0 | 0 | 0 | 0 | 0 | 0 | 0 | 0 | 0 |
| 1730 | rhsB | 0 | 0 | 0 | 0 | 0 | 0 | 0 | 0 | 0 | 0 |
| 1731 | rhsA | 0 | 0 | 0 | 0 | 0 | 0 | 0 | 0 | 0 | 0 |
| 1732 | pdxH | 0 | 0 | 0 | 0 | 0 | 0 | 0 | 0 | 0 | 0 |
| 1733 | pdxK | 0 | 0 | 0 | 0 | 0 | 0 | 0 | 0 | 0 | 0 |
| 1734 | pdxB | 0 | 0 | 0 | 0 | 0 | 0 | 0 | 0 | 0 | 0 |
| 1735 | nrdG | 0 | 0 | 0 | 0 | 0 | 0 | 0 | 0 | 0 | 0 |
| 1736 | nrdH | 0 | 0 | 0 | 0 | 0 | 0 | 0 | 0 | 0 | 0 |
| 1737 | yghU | 0 | 0 | 0 | 0 | 0 | 0 | 0 | 0 | 0 | 0 |
| 1738 | mdoB | 0 | 0 | 0 | 4 | 0 | 0 | 0 | 2 | 0 | 0 |
| 1739 | negCtrl | 0 | 0 | 0 | 0 | 0 | 0 | 0 | 0 | 0 | 0 |

Data Set S2 - MG1655 library 7 day experiment
Odds Ratios

| | Promoter | gavage | M3-day1 | M4-day1 | M5-day1 | M3-day7 | M4-day7 | M5-day7 | #(OR>.02 & p<.05) / #non-N/A | AVG OR |
|---|---|---|---|---|---|---|---|---|---|---|
| 64 | ydiL | 0.0000 | 0.7479 | 0.9117 | 0.4953 | N/A | N/A | N/A | 1 | 0.7183 |
| 88 | ydjL | 0.0000 | 0.6555 | 0.7266 | 0.3133 | 0.1251 | N/A | 1.6286 | 1 | 0.6898 |
| 8 | gcvA | 0.0061 | 0.4132 | 0.1093 | 0.9384 | 0.1053 | N/A | 0.4845 | 1 | 0.4102 |
| 37 | fucR | 0.0000 | 0.0583 | 0.1965 | 0.0465 | 0.4388 | N/A | N/A | 1 | 0.1850 |
| 69 | ubiG | 0.0011 | 0.0617 | 0.0123 | 0.3473 | 0.0144 | N/A | 0.1302 | 0.6 | 0.1132 |
| 5 | gatY | 0.0003 | 0.0498 | 0.0034 | 0.1944 | 0.1036 | N/A | 0.0005 | 0.6 | 0.0704 |
| 71 | galS | 0.0021 | 0.1423 | 0.0000 | 0.6656 | N/A | N/A | 0.0000 | 0.5 | 0.2020 |
| 65 | dinG | 0.0000 | 0.0000 | 0.0724 | N/A | N/A | N/A | N/A | 0.5 | 0.0362 |
| 66 | guaC | 0.0000 | 0.1185 | 0.0000 | 0.0000 | 0.0000 | N/A | 0.2336 | 0.4 | 0.0704 |
| 93 | yhbQ | 0.0001 | 0.0013 | 0.0002 | 0.0259 | 0.0478 | N/A | 0.0803 | 0.4 | 0.0311 |
| 30 | rof | 0.0003 | 0.0197 | 0.0001 | 0.6460 | N/A | N/A | 0.0000 | 0.25 | 0.1664 |
| 63 | ybiA | 0.0000 | 0.0198 | 0.0653 | 0.0037 | N/A | N/A | 0.0000 | 0.25 | 0.0222 |
| 6 | fadB | 0.0007 | 0.0280 | 0.0036 | 0.0373 | N/A | N/A | 0.0000 | 0.25 | 0.0172 |
| 10 | pgsA | 0.0003 | 0.0132 | 0.0001 | 0.2185 | 0.0000 | N/A | 0.0002 | 0.2 | 0.0464 |
| 32 | cysH | 0.0000 | 0.0028 | 0.0002 | 0.1272 | 0.0007 | N/A | 0.0023 | 0.2 | 0.0266 |
| 73 | yjcO | 0.0000 | 0.0013 | 0.0100 | 0.0083 | 0.1102 | N/A | 0.0019 | 0.2 | 0.0264 |
| 68 | ubiF | 0.0002 | 0.0094 | 0.0027 | 0.0796 | 0.0111 | N/A | 0.0031 | 0.2 | 0.0212 |
| 24 | cynR | 0.0001 | 0.0063 | 0.0006 | 0.0482 | 0.0022 | N/A | 0.0023 | 0.2 | 0.0119 |
| 52 | pepQ | 0.0002 | 0.0528 | 0.0053 | N/A | N/A | N/A | N/A | 0 | 0.0291 |
| 83 | edd | 0.0004 | 0.0317 | 0.0048 | N/A | N/A | N/A | N/A | 0 | 0.0183 |
| 87 | mutS | 0.0002 | 0.0028 | 0.0129 | 0.0000 | N/A | N/A | 0.0371 | 0 | 0.0132 |
| 36 | ydcF | 0.0000 | 0.0006 | 0.0214 | 0.0135 | 0.0138 | N/A | 0.0019 | 0 | 0.0102 |
| 29 | idnT | 0.0000 | 0.0000 | 0.0000 | 0.0283 | N/A | N/A | N/A | 0 | 0.0094 |
| 13 | glyQ | 0.0000 | 0.0159 | 0.0007 | N/A | N/A | N/A | N/A | 0 | 0.0083 |
| 81 | b3044 | 0.0000 | 0.0226 | 0.0013 | N/A | N/A | N/A | 0.0000 | 0 | 0.0080 |
| 82 | pabC | 0.0000 | 0.0004 | 0.0001 | 0.0296 | N/A | N/A | 0.0003 | 0 | 0.0076 |
| 75 | yicC | 0.0001 | 0.0000 | 0.0000 | 0.0283 | N/A | N/A | 0.0000 | 0 | 0.0071 |
| 22 | yijO | 0.0000 | 0.0000 | 0.0000 | 0.0252 | 0.0000 | N/A | N/A | 0 | 0.0063 |
| 4 | b0671 | 0.0000 | 0.0006 | 0.0002 | 0.0280 | 0.0003 | N/A | 0.0015 | 0 | 0.0061 |
| 57 | yegT | 0.0000 | 0.0014 | 0.0043 | 0.0105 | N/A | N/A | 0.0000 | 0 | 0.0040 |
| 11 | dcuS | 0.0000 | 0.0017 | 0.0000 | 0.0120 | N/A | N/A | 0.0000 | 0 | 0.0034 |
| 76 | ybhC | 0.0000 | 0.0088 | 0.0011 | N/A | N/A | N/A | 0.0000 | 0 | 0.0033 |
| 18 | yjdI | 0.0000 | 0.0019 | 0.0000 | 0.0110 | N/A | N/A | 0.0000 | 0 | 0.0032 |
| 55 | fumB | 0.0000 | 0.0000 | 0.0000 | 0.0044 | 0.0058 | N/A | 0.0000 | 0 | 0.0021 |
| 7 | gcvT | 0.0001 | 0.0051 | 0.0001 | 0.0000 | N/A | N/A | N/A | 0 | 0.0017 |
| 38 | rrlG | 0.0000 | 0.0000 | 0.0001 | 0.0084 | 0.0000 | N/A | 0.0000 | 0 | 0.0017 |
| 21 | ygjG | 0.0000 | 0.0026 | 0.0000 | 0.0046 | 0.0005 | N/A | 0.0001 | 0 | 0.0016 |
| 40 | tktB | 0.0000 | 0.0004 | 0.0000 | 0.0016 | 0.0050 | N/A | 0.0000 | 0 | 0.0014 |
| 12 | ybbB | 0.0000 | 0.0003 | 0.0000 | 0.0050 | N/A | N/A | 0.0002 | 0 | 0.0014 |
| 17 | yjdF | 0.0000 | 0.0017 | 0.0000 | 0.0035 | 0.0000 | N/A | N/A | 0 | 0.0013 |
| 49 | grxB | 0.0000 | 0.0019 | 0.0000 | 0.0000 | 0.0044 | N/A | 0.0000 | 0 | 0.0013 |
| 27 | phnQ | 0.0000 | 0.0003 | 0.0000 | 0.0052 | 0.0003 | N/A | 0.0001 | 0 | 0.0012 |
| 59 | cueR | 0.0000 | 0.0004 | 0.0000 | 0.0047 | 0.0000 | N/A | 0.0008 | 0 | 0.0012 |
| 60 | ybjK | 0.0000 | 0.0007 | 0.0001 | 0.0032 | 0.0007 | N/A | N/A | 0 | 0.0012 |
| 45 | argZ_argQ | 0.0001 | 0.0000 | 0.0001 | 0.0043 | N/A | N/A | 0.0002 | 0 | 0.0011 |
| 25 | nudC | 0.0000 | 0.0004 | 0.0000 | 0.0000 | 0.0035 | N/A | 0.0014 | 0 | 0.0011 |
| 102 | negCtrl | 0.0000 | 0.0027 | 0.0001 | 0.0000 | 0.0012 | N/A | 0.0000 | 0 | 0.0008 |
| 92 | yhbW | 0.0000 | 0.0003 | 0.0000 | 0.0030 | 0.0000 | N/A | 0.0000 | 0 | 0.0006 |

Data Set S2 - MG1655 library 7 day experiment
Odds Ratios

| | | | | | | | | | | |
|---|---|---|---|---|---|---|---|---|---|---|
| 43 | yeaU | 0.0000 | 0.0024 | 0.0000 | 0.0000 | 0.0000 | N/A | N/A | 0 | 0.0006 |
| 86 | fabF | 0.0000 | 0.0022 | 0.0000 | 0.0000 | 0.0000 | N/A | 0.0008 | 0 | 0.0006 |
| 101 | yjfY | 0.0000 | 0.0004 | 0.0000 | 0.0000 | 0.0024 | N/A | 0.0000 | 0 | 0.0006 |
| 23 | yfcC | 0.0000 | 0.0004 | 0.0000 | 0.0017 | N/A | N/A | 0.0000 | 0 | 0.0005 |
| 72 | hrpB | 0.0000 | 0.0003 | 0.0000 | 0.0009 | 0.0011 | N/A | 0.0003 | 0 | 0.0005 |
| 100 | yaaU | 0.0000 | 0.0004 | 0.0000 | 0.0018 | 0.0000 | N/A | 0.0000 | 0 | 0.0004 |
| 95 | uidA | 0.0000 | 0.0016 | 0.0001 | 0.0000 | N/A | N/A | 0.0000 | 0 | 0.0004 |
| 16 | yhhT | 0.0000 | 0.0004 | 0.0000 | 0.0000 | 0.0013 | N/A | 0.0002 | 0 | 0.0004 |
| 33 | cysQ | 0.0000 | 0.0013 | 0.0000 | 0.0000 | N/A | N/A | 0.0000 | 0 | 0.0003 |
| 94 | pgi | 0.0000 | 0.0008 | 0.0000 | 0.0000 | 0.0000 | N/A | 0.0007 | 0 | 0.0003 |
| 97 | rph | 0.0000 | 0.0011 | 0.0000 | 0.0000 | N/A | N/A | 0.0000 | 0 | 0.0003 |
| 67 | yceF | 0.0000 | 0.0007 | 0.0001 | 0.0000 | N/A | N/A | N/A | 0 | 0.0003 |
| 51 | rluF | 0.0000 | 0.0008 | 0.0000 | 0.0000 | N/A | N/A | N/A | 0 | 0.0003 |
| 89 | yfjN | 0.0000 | 0.0006 | 0.0000 | 0.0000 | 0.0006 | N/A | 0.0000 | 0 | 0.0002 |
| 42 | ybgE | 0.0000 | 0.0005 | 0.0001 | 0.0000 | N/A | N/A | 0.0002 | 0 | 0.0002 |
| 15 | pdhR | 0.0000 | 0.0010 | 0.0000 | 0.0000 | 0.0000 | N/A | 0.0000 | 0 | 0.0002 |
| 79 | mreC | 0.0001 | 0.0005 | 0.0000 | 0.0000 | 0.0003 | N/A | 0.0000 | 0 | 0.0002 |
| 14 | gst | 0.0000 | 0.0006 | 0.0000 | 0.0000 | 0.0000 | N/A | 0.0000 | 0 | 0.0001 |
| 74 | barA | 0.0000 | 0.0005 | 0.0000 | 0.0000 | N/A | N/A | 0.0000 | 0 | 0.0001 |
| 77 | mobA | 0.0000 | 0.0000 | 0.0000 | 0.0000 | N/A | N/A | 0.0005 | 0 | 0.0001 |
| 44 | nupX | 0.0000 | 0.0003 | 0.0000 | 0.0000 | 0.0000 | N/A | 0.0003 | 0 | 0.0001 |
| 48 | yiiP | 0.0000 | 0.0003 | 0.0000 | 0.0000 | 0.0003 | N/A | 0.0000 | 0 | 0.0001 |
| 50 | ynbA | 0.0000 | 0.0004 | 0.0001 | 0.0000 | N/A | N/A | 0.0000 | 0 | 0.0001 |
| 35 | mdtO | 0.0000 | 0.0004 | 0.0000 | 0.0000 | N/A | N/A | 0.0000 | 0 | 0.0001 |
| 20 | yebF | 0.0000 | 0.0004 | 0.0000 | 0.0000 | N/A | N/A | 0.0000 | 0 | 0.0001 |
| 84 | yjaB | 0.0000 | 0.0004 | 0.0000 | 0.0000 | 0.0000 | N/A | 0.0000 | 0 | 0.0001 |
| 31 | yfbJ | 0.0000 | 0.0002 | 0.0000 | 0.0000 | 0.0000 | N/A | 0.0000 | 0 | 0.0000 |
| 91 | metK | 0.0000 | 0.0000 | 0.0001 | 0.0000 | N/A | N/A | N/A | 0 | 0.0000 |
| 62 | yqgA | 0.0000 | 0.0000 | 0.0001 | 0.0000 | N/A | N/A | 0.0000 | 0 | 0.0000 |
| 2 | speC | 0.0000 | 0.0000 | 0.0000 | 0.0000 | N/A | N/A | 0.0000 | 0 | 0.0000 |
| 3 | eco | 0.0000 | 0.0000 | 0.0000 | 0.0000 | N/A | N/A | 0.0000 | 0 | 0.0000 |
| 9 | aniC | 0.0000 | 0.0000 | 0.0000 | 0.0000 | N/A | N/A | 0.0000 | 0 | 0.0000 |
| 28 | msbB | 0.0000 | 0.0000 | 0.0000 | 0.0000 | 0.0000 | N/A | 0.0000 | 0 | 0.0000 |
| 34 | yihD | 0.0000 | 0.0000 | 0.0000 | 0.0000 | N/A | N/A | 0.0000 | 0 | 0.0000 |
| 39 | ycaO | 0.0000 | 0.0000 | 0.0000 | N/A | N/A | N/A | N/A | 0 | 0.0000 |
| 41 | yqgD | 0.0000 | 0.0000 | 0.0000 | 0.0000 | N/A | N/A | N/A | 0 | 0.0000 |
| 46 | argA_amiC | 0.0000 | 0.0000 | 0.0000 | 0.0000 | N/A | N/A | 0.0000 | 0 | 0.0000 |
| 47 | argD | 0.0000 | 0.0000 | 0.0000 | 0.0000 | N/A | N/A | N/A | 0 | 0.0000 |
| 53 | yidK | 0.0000 | 0.0000 | 0.0000 | 0.0000 | N/A | N/A | 0.0000 | 0 | 0.0000 |
| 54 | yidL | 0.0000 | 0.0000 | 0.0000 | 0.0000 | N/A | N/A | 0.0000 | 0 | 0.0000 |
| 56 | ynaI | 0.0000 | 0.0000 | 0.0000 | 0.0000 | N/A | N/A | N/A | 0 | 0.0000 |
| 58 | rihC | 0.0000 | 0.0000 | 0.0000 | 0.0000 | N/A | N/A | N/A | 0 | 0.0000 |
| 70 | bglA | 0.0000 | 0.0000 | 0.0000 | 0.0000 | 0.0000 | N/A | 0.0000 | 0 | 0.0000 |
| 90 | yfjR | 0.0000 | 0.0000 | 0.0000 | 0.0000 | 0.0000 | N/A | N/A | 0 | 0.0000 |
| 96 | rumA | 0.0000 | 0.0000 | 0.0000 | 0.0000 | N/A | N/A | 0.0000 | 0 | 0.0000 |
| 99 | ycfD | 0.0000 | 0.0000 | 0.0000 | N/A | N/A | N/A | N/A | 0 | 0.0000 |
| 19 | flgH | 0.0001 | 0.0000 | 0.0000 | 0.0000 | 0.0000 | N/A | 0.0000 | 0 | 0.0000 |
| 26 | ygeF | 0.0000 | N/A | N/A | N/A | N/A | N/A | N/A | | #DIV/0! |
| 85 | sirA | 0.0000 | N/A | N/A | N/A | N/A | N/A | N/A | | #DIV/0! |
| 78 | ydhR | 0.0219 | 0.4341 | 0.0383 | 1.0803 | 0.0088 | N/A | 0.0002 | 0.6 | 0.3123 |
| 80 | ychH | 0.2976 | 0.4938 | 0.5021 | 0.6612 | 0.4913 | N/A | 0.6053 | 1 | 0.5507 |

Data Set S2 - MG1655 library 7 day experiment
Odds Ratios

| | | | | | | | | | | |
|---|---|---|---|---|---|---|---|---|---|---|
| 1 | speE | 0.5199 | 1.1906 | 1.6953 | 0.9328 | N/A | N/A | 1.2642 | 1 | 1.2707 |
| 98 | fabR | 1.0000 | 1.0000 | 1.0000 | 1.0000 | 1.0000 | N/A | 1.0000 | 1 | 1.0000 |
| 61 | yahN | N/A | N/A | N/A | N/A | N/A | N/A | N/A | | #DIV/0! |

### Data Set S2 - MG1655 library 7 day experiment
#### Odds Ratios & p-Values

| | Promoter | gavage | gavage_pValu | M3-day1 | M3-day1_pVa | M4-day1 | M4-day1_pVa | M5-day1 | M5-day1_pVa | M3-day7 | M3-day7_pVa | M5-day7 | M5-day7_pValue |
|---|---|---|---|---|---|---|---|---|---|---|---|---|---|
| 1 | speE | 0.51991154 | 3.49E-23 | 1.19056678 | 0 | 1.69532402 | 2.24E-52 | 0.93280591 | 0 | N/A | 1 | 1.26424416 | 2.45E-176 |
| 2 | speC | 0 | 1 | 0 | 1 | 0 | 1 | 0 | 1 | N/A | 1 | 0 | 1 |
| 3 | eco | 0 | 1 | 0 | 1 | 0 | 1 | 0 | 1 | N/A | 1 | 0 | 1 |
| 4 | b0671 | 0 | 1 | 0.00061594 | 1 | 0.00021283 | 1 | 0.02801819 | 0.13243473 | 0.00030473 | 1 | 0.00145298 | 1 |
| 5 | gatY | 0.00028467 | 1 | 0.04981331 | 1.38E-18 | 0.00337081 | 1 | 0.19444878 | 9.36E-68 | 0.10363147 | 0.00120965 | 0.00052712 | 1 |
| 6 | fadB | 0.00067675 | 1 | 0.02798414 | 0.04026411 | 0.00360248 | 1 | 0.03725491 | 0.07639405 | N/A | 1 | 0 | 1 |
| 7 | gcvT | 6.10E-05 | 1 | 0.00511591 | 0.9997474 | 0.00010284 | 1 | 0 | 1 | N/A | 1 | N/A | 1 |
| 8 | gcvA | 0.0061274 | 1 | 0.41319472 | 1.35E-256 | 0.1093439 | 7.22E-24 | 0.93844722 | 0 | 0.10530492 | 2.59E-13 | 0.48451502 | 1.06E-233 |
| 9 | aniC | 0 | 1 | 0 | 1 | 0 | 1 | 0 | 1 | N/A | 1 | 0 | 1 |
| 10 | pgsA | 0.0003234 | 1 | 0.01323372 | 0.99999562 | 9.23E-05 | 1 | 0.218536 | 7.79E-298 | 0 | 1 | 0.00017509 | 1 |
| 11 | dcuS | 0 | 1 | 0.00169999 | 0.99999999 | 0 | 1 | 0.01204069 | 0.84368874 | N/A | 1 | 0 | 1 |
| 12 | ybbB | 0 | 1 | 0.00028913 | 1 | 0 | 1 | 0.00501863 | 0.99664823 | N/A | 1 | 0.00024449 | 1 |
| 13 | glyQ | 0 | 1 | 0.01585225 | 0.72378041 | 0.00068237 | 0.99999998 | N/A | 1 | N/A | 1 | N/A | 1 |
| 14 | gst | 0 | 1 | 0.00064755 | 1 | 0 | 1 | 0 | 1 | 0 | 1 | 0 | 1 |
| 15 | pdhR | 0 | 1 | 0.00098736 | 1 | 0 | 1 | 0 | 1 | 0 | 1 | 0 | 1 |
| 16 | yhhT | 0 | 1 | 0.00041068 | 1 | 0 | 1 | 0 | 1 | 0.00131046 | 0.99999185 | 0.00018327 | 1 |
| 17 | yjdF | 0 | 1 | 0.00170843 | 1 | 0 | 1 | 0.00349545 | 0.99635692 | 0 | 1 | N/A | 1 |
| 18 | yjdI | 0 | 1 | 0.00193902 | 0.9999999 | 0 | 1 | 0.0109711 | 0.83903571 | N/A | 1 | 0 | 1 |
| 19 | flgH | 6.29E-05 | 1 | 0 | 1 | 0 | 1 | 0 | 1 | 0 | 1 | 0 | 1 |
| 20 | yebF | 0 | 1 | 0.00036136 | 1 | 0 | 1 | 0 | 1 | N/A | 1 | 0 | 1 |
| 21 | ygjG | 0 | 1 | 0.00264135 | 0.99999984 | 0 | 1 | 0.00458693 | 0.98663429 | 0.00052773 | 1 | 0.00012751 | 1 |
| 22 | yijO | 0 | 1 | 0 | 1 | 0 | 1 | 0.02516797 | 0.55611429 | 0 | 1 | N/A | 1 |
| 23 | yfcC | 0 | 1 | 0.00041923 | 1 | 0 | 1 | 0.00174916 | 0.99998782 | N/A | 1 | 0 | 1 |
| 24 | cynR | 9.39E-05 | 1 | 0.00625094 | 1 | 0.00063331 | 1 | 0.0482391 | 1.15E-103 | 0.00220548 | 1 | 0.00225448 | 1 |
| 25 | nudC | 0 | 1 | 0.00038412 | 1 | 0 | 1 | 0 | 1 | 0.00351748 | 0.99226912 | 0.00138956 | 0.9999978 |
| 26 | ygeF | 0 | 1 | N/A | 1 | N/A | 1 | N/A | 1 | N/A | 1 | N/A | 1 |
| 27 | phnQ | 0 | 1 | 0.00026757 | 1 | 0 | 1 | 0.00521561 | 0.99995814 | 0.00029728 | 1 | 0.00014564 | 1 |
| 28 | msbB | 0 | 1 | 0 | 1 | 0 | 1 | 0 | 1 | 0 | 1 | 0 | 1 |
| 29 | idnT | 0 | 1 | 0 | 1 | 0 | 1 | 0.02832969 | 0.51577082 | N/A | 1 | N/A | 1 |
| 30 | rof | 0.00025374 | 1 | 0.01973068 | 0.55850286 | 6.92E-05 | 1 | 0.64599502 | 1.75E-90 | N/A | 1 | 0 | 1 |
| 31 | yfbJ | 0 | 1 | 0.00023761 | 1 | 0 | 1 | 0 | 1 | 0 | 1 | 0 | 1 |
| 32 | cysH | 0 | 1 | 0.00281253 | 1 | 0.00016073 | 1 | 0.12718795 | 2.63E-12 | 0.00065503 | 1 | 0.00226738 | 1 |
| 33 | cysQ | 0 | 1 | 0.00130706 | 1 | 0 | 1 | 0 | 1 | N/A | 1 | 0 | 1 |
| 34 | yihD | 0 | 1 | 0 | 1 | 0 | 1 | 0 | 1 | N/A | 1 | 0 | 1 |
| 35 | mdtO | 0 | 1 | 0.00037536 | 1 | 0 | 1 | 0 | 1 | N/A | 1 | 0 | 1 |
| 36 | ydcF | 0 | 1 | 0.00060809 | 1 | 0.02140027 | 0.3273113 | 0.01350604 | 0.98685728 | 0.01376885 | 0.89581479 | 0.00188588 | 1 |
| 37 | fucR | 0 | 1 | 0.05834275 | 1.00E-20 | 0.19648011 | 7.02E-35 | 0.0464672 | 0.00112385 | 0.43877523 | 2.08E-32 | N/A | 1 |
| 38 | rrlG | 0 | 1 | 0 | 1 | 8.18E-05 | 1 | 0.0084489 | 0.94846731 | 0 | 1 | 0 | 1 |
| 39 | ycaO | 0 | 1 | 0 | 1 | 0 | 1 | N/A | 1 | N/A | 1 | N/A | 1 |
| 40 | tktB | 0 | 1 | 0.00042922 | 1 | 0 | 1 | 0.00162026 | 0.999995 | 0.00501852 | 0.97198353 | 0 | 1 |
| 41 | yqgD | 0 | 1 | 0 | 1 | 0 | 1 | 0 | 1 | N/A | 0.59931175 | N/A | 1 |
| 42 | ybgE | 0 | 1 | 0.00054086 | 1 | 6.38E-05 | 1 | 0 | 1 | N/A | 1 | 0.00020662 | 1 |
| 43 | yeaU | 0 | 1 | 0.00242608 | 1 | 0 | 1 | 0 | 1 | 0 | 1 | N/A | 1 |
| 44 | nupX | 0 | 1 | 0.00034911 | 1 | 0 | 1 | 0 | 1 | 0 | 1 | 0.00025091 | 1 |
| 45 | argZ_argQ | 6.10E-05 | 1 | 0 | 1 | 6.33E-05 | 1 | 0.00426258 | 0.99027397 | N/A | 1 | 0.00023185 | 1 |
| 46 | argA_amiC | 0 | 1 | 0 | 1 | 0 | 1 | 0 | 1 | N/A | 1 | 0 | 1 |
| 47 | argD | 0 | 1 | 0 | 1 | 0 | 1 | 0 | 1 | N/A | 1 | N/A | 1 |
| 48 | yiiP | 0 | 1 | 0.00025735 | 1 | 0 | 1 | 0 | 1 | 0.00033395 | 1 | 0 | 1 |
| 49 | grxB | 0 | 1 | 0.00190909 | 1 | 0 | 1 | 0 | 1 | 0.00439383 | 0.98181449 | 0 | 1 |
| 50 | ynbA | 0 | 1 | 0.00038502 | 1 | 6.61E-05 | 1 | 0 | 1 | N/A | 0.3681694 | 0 | 1 |
| 51 | rluF | 0 | 1 | 0.00079673 | 1 | 0 | 1 | 0 | 1 | N/A | 1 | N/A | 1 |
| 52 | pepQ | 0.00024235 | 1 | 0.05282211 | 0.22446456 | 0.00534904 | 0.99480982 | N/A | 1 | N/A | 1 | N/A | 1 |
| 53 | yidK | 0 | 1 | 0 | 1 | 0 | 1 | 0 | 1 | N/A | 1 | 0 | 1 |
| 54 | yidL | 0 | 1 | 0 | 1 | 0 | 1 | 0 | 1 | N/A | 1 | 0 | 1 |
| 55 | fumB | 0 | 1 | 0 | 1 | 0 | 1 | 0.00442538 | 0.99983737 | 0.00584885 | 0.95670513 | 0 | 1 |
| 56 | ynaI | 0 | 1 | 0 | 1 | 0 | 1 | 0 | 1 | N/A | 1 | N/A | 1 |
| 57 | yegT | 0 | 1 | 0.0014315 | 1 | 0.00428227 | 1 | 0.01046439 | 0.85243347 | N/A | 1 | 0 | 1 |
| 58 | rihC | 0 | 1 | 0 | 1 | 0 | 1 | 0 | 1 | N/A | 1 | N/A | 1 |
| 59 | cueR | 0 | 1 | 0.00037281 | 1 | 0 | 1 | 0.00468401 | 0.98542359 | 0 | 1 | 0.00082234 | 1 |
| 60 | ybjK | 0 | 1 | 0.00066721 | 1 | 6.81E-05 | 1 | 0.00319022 | 0.99806926 | 0.0006987 | 1 | N/A | 1 |
| 61 | yahN | N/A | 1 | N/A | 0.06501067 | N/A | 6.84E-11 | N/A | 1 | N/A | 1 | N/A | 1 |
| 62 | yqgA | 0 | 1 | 0 | 1 | 9.06E-05 | 1 | 0 | 1 | N/A | 1 | 0 | 1 |
| 63 | ybiA | 0 | 1 | 0.01976343 | 0.55572381 | 0.06525766 | 3.63E-13 | 0.00369127 | 0.99513749 | N/A | 0.00115874 | 0 | 1 |
| 64 | ydiL | 0 | 1 | 0.74789627 | 6.41E-298 | 0.91169137 | 1.76E-71 | 0.49534115 | 1.91E-38 | N/A | 0.00023054 | N/A | 1 |
| 65 | dinG | 0 | 1 | 0 | 1 | 0.07240954 | 5.78E-05 | N/A | 1 | N/A | 1 | N/A | 1 |
| 66 | guaC | 0 | 1 | 0.1184639 | 1.58E-92 | 0 | 1 | 0 | 1 | 0 | 1 | 0.23364452 | 2.49E-110 |
| 67 | yceF | 0 | 1 | 0.00073042 | 1 | 7.54E-05 | 1 | 0 | 1 | N/A | 1 | N/A | 1 |
| 68 | ubiF | 0.00015378 | 1 | 0.00935554 | 1 | 0.00271468 | 1 | 0.07956947 | 0 | 0.01105121 | 0.99995941 | 0.00314811 | 1 |
| 69 | ubiG | 0.00107216 | 1 | 0.06165452 | 1.30E-55 | 0.01227243 | 0.99997473 | 0.34729731 | 0 | 0.01441965 | 0.98238315 | 0.13023222 | 1.05E-107 |
| 70 | bglA | 0 | 1 | 0 | 1 | 0 | 1 | 0 | 1 | 0 | 1 | 0 | 1 |
| 71 | galS | 0.0021321 | 1 | 0.14225181 | 1.88E-79 | 0 | 1 | 0.66556208 | 0 | N/A | 7.54E-05 | 0 | 1 |
| 72 | hrpB | 0 | 1 | 0.00029708 | 1 | 0 | 1 | 0.00086758 | 1 | 0.0011047 | 0.99999877 | 0.0003207 | 1 |
| 73 | yjcO | 0 | 1 | 0.00134276 | 1 | 0.01002275 | 1 | 0.00832692 | 1 | 0.11024284 | 8.39E-16 | 0.00194649 | 1 |
| 74 | barA | 0 | 1 | 0.00049819 | 1 | 0 | 1 | 0 | 1 | N/A | 1 | 0 | 1 |
| 75 | yicC | 6.10E-05 | 1 | 0 | 1 | 0 | 1 | 0.02832969 | 0.51577082 | N/A | 1 | 0 | 1 |
| 76 | ybhC | 0 | 1 | 0.00881021 | 0.89525267 | 0.00106248 | 1 | N/A | 1 | N/A | 1 | 0 | 1 |
| 77 | mobA | 0 | 1 | 0 | 1 | 0 | 1 | 0 | 1 | N/A | 1 | 0.00049446 | 1 |
| 78 | ydhR | 0.02194768 | 0.30921387 | 0.43409611 | 8.47E-263 | 0.0382584 | 6.24E-06 | 1.08032193 | 0 | 0.0087665 | 0.98361707 | 0.00017767 | 1 |
| 79 | mreC | 6.34E-05 | 1 | 0.00050277 | 1 | 0 | 1 | 0 | 1 | 0.00033982 | 1 | 0 | 1 |
| 80 | ychH | 0.29756392 | 4.54E-30 | 0.49376051 | 1.10E-283 | 0.5020663 | 1.31E-58 | 0.66124343 | 0 | 0.4912677 | 1.27E-37 | 0.60527747 | 1.29E-256 |
| 81 | b3044 | 0 | 1 | 0.02262746 | 0.61407297 | 0.00133777 | 0.99993164 | N/A | 1 | N/A | 1 | 0 | 1 |
| 82 | pabC | 0 | 1 | 0.000427 | 1 | 6.36E-05 | 1 | 0.02956129 | 0.23183988 | N/A | 1 | 0.00031961 | 1 |
| 83 | edd | 0.00035634 | 0.99999995 | 0.03170609 | 0.51050058 | 0.00483394 | 0.98623654 | N/A | 0.0563817 | N/A | 0.6032597 | N/A | 0.29084781 |

### Data Set S2 - MG1655 library 7 day experiment
#### Odds Ratios & p-Values

| | | | | | | | | | | | | | |
|---|---|---|---|---|---|---|---|---|---|---|---|---|---|
| 84 | yjaB | 0 | 1 | 0.00042261 | 1 | 0 | 1 | 0 | 1 | 0 | 1 | 0 | 1 |
| 85 | sirA | 0 | 1 | N/A | 1 | N/A | 1 | N/A | 1 | N/A | 1 | N/A | 0.17801785 |
| 86 | fabF | 0 | 1 | 0.00223215 | 1 | 0 | 1 | 0 | 1 | 0 | 1 | 0.00079416 | 1 |
| 87 | mutS | 0.00015252 | 1 | 0.00281302 | 0.99999999 | 0.01292398 | 0.9988222 | 0 | 1 | N/A | 1 | 0.03708203 | 0.11253669 |
| 88 | ydjL | 0 | 1 | 0.65549026 | 5.53E-293 | 0.72663692 | 8.06E-67 | 0.31332918 | 7.17E-88 | 0.12507044 | 5.26E-06 | 1.62858289 | 4.32E-287 |
| 89 | yfjN | 0 | 1 | 0.0006264 | 1 | 0 | 1 | 0 | 1 | 0.00061954 | 1 | 0 | 1 |
| 90 | yfjR | 0 | 1 | 0 | 1 | 0 | 1 | 0 | 1 | 0 | 1 | N/A | 1 |
| 91 | metK | 0 | 1 | 0 | 1 | 0.00011443 | 1 | 0 | 1 | N/A | 1 | N/A | 1 |
| 92 | yhbW | 0 | 1 | 0.00028964 | 1 | 0 | 1 | 0.00295506 | 0.99881762 | 0 | 1 | 0 | 1 |
| 93 | yhbQ | 9.34E-05 | 1 | 0.0013232 | 1 | 0.00016345 | 1 | 0.0259385 | 0.09229543 | 0.04784784 | 2.75E-05 | 0.08028006 | 2.44E-61 |
| 94 | pgi | 0 | 1 | 0.00084915 | 1 | 0 | 1 | 0 | 1 | 0 | 1 | 0.00074324 | 1 |
| 95 | uidA | 0 | 1 | 0.00159727 | 1 | 7.03E-05 | 1 | 0 | 1 | N/A | 1 | 0 | 1 |
| 96 | rumA | 0 | 1 | 0 | 1 | 0 | 1 | 0 | 1 | N/A | 1 | 0 | 1 |
| 97 | rph | 0 | 1 | 0.00111785 | 0.99999994 | 0 | 1 | 0 | 1 | N/A | 1 | 0 | 1 |
| 98 | fabR | 1 | 6.18E-43 | 1 | 1.670291854 | 1 | 3.43E-66 | 1 | 0 | 1 | 1.82E-42 | 1 | 4.23E-266 |
| 99 | ycfD | 0 | 1 | 0 | 1 | 0 | 1 | N/A | 0.08336795 | N/A | 1 | N/A | 1 |
| 100 | yaaU | 0 | 1 | 0.0003642 | 1 | 0 | 1 | 0.00183844 | 0.99999999 | 0 | 1 | 0 | 1 |
| 101 | yjfY | 0 | 1 | 0.0004062 | 1 | 0 | 1 | 0 | 1 | 0.00237715 | 0.99904332 | 0 | 1 |
| 102 | negCtrl | 0 | 1 | 0.00265012 | 0.9992408 | 0.00014602 | 1 | 0 | 1 | 0.00120434 | 1 | 0 | 1 |

### Data Set S2 - MG1655 library 7 day experiment
#### Aligned Reads

| | Promoter | gavage-ALL_S2 | gavage-ON_S1 | M3-day1-ALL_ | M3-day1-ON_ | M3-day7-ALL_ | M3-day7-ON_ | M4-day1-ALL_ | M4-day1-ON_ | M5-day1-ALL_ | M5-day1-ON_ | M5-day7-ALL_ | M5-day7-ON_S11 |
|---|---|---|---|---|---|---|---|---|---|---|---|---|---|
| 1 | xapA | 0 | 0 | 0 | 0 | 0 | 0 | 0 | 0 | 0 | 0 | 0 | 0 |
| 2 | abrB | 0 | 0 | 0 | 0 | 0 | 0 | 0 | 0 | 0 | 0 | 0 | 0 |
| 3 | speF | 0 | 0 | 0 | 0 | 0 | 0 | 0 | 0 | 0 | 0 | 0 | 0 |
| 4 | xseA | 0 | 0 | 0 | 0 | 0 | 0 | 0 | 0 | 0 | 0 | 0 | 1 |
| 5 | xseB | 0 | 0 | 0 | 0 | 0 | 0 | 0 | 0 | 0 | 0 | 0 | 0 |
| 6 | speE | 8 | 1603 | 256 | 1928 | 0 | 0 | 7 | 1000 | 1280 | 1757 | 35 | 479 |
| 7 | speB | 0 | 0 | 0 | 0 | 1 | 0 | 0 | 0 | 0 | 0 | 0 | 0 |
| 8 | speC | 94 | 0 | 72 | 0 | 1 | 0 | 107 | 0 | 35 | 0 | 13 | 0 |
| 9 | ybcL | 0 | 0 | 0 | 0 | 0 | 0 | 0 | 0 | 0 | 0 | 0 | 0 |
| 10 | ytfE | 0 | 0 | 0 | 0 | 0 | 0 | 0 | 1 | 0 | 0 | 0 | 0 |
| 11 | ybcK | 0 | 0 | 0 | 0 | 0 | 0 | 0 | 0 | 0 | 0 | 0 | 0 |
| 12 | frdA | 0 | 0 | 0 | 0 | 0 | 0 | 0 | 0 | 0 | 0 | 0 | 0 |
| 13 | ytfF | 0 | 0 | 0 | 0 | 0 | 0 | 0 | 0 | 0 | 0 | 0 | 0 |
| 14 | ybcY | 0 | 0 | 0 | 0 | 0 | 0 | 0 | 0 | 0 | 0 | 0 | 0 |
| 15 | pgpB | 0 | 0 | 0 | 0 | 0 | 0 | 0 | 0 | 0 | 0 | 0 | 0 |
| 16 | ybcV | 0 | 0 | 0 | 0 | 0 | 0 | 0 | 0 | 0 | 0 | 0 | 0 |
| 17 | ybcW | 0 | 0 | 0 | 0 | 0 | 0 | 0 | 0 | 0 | 0 | 0 | 0 |
| 18 | kbaZ | 0 | 0 | 0 | 0 | 0 | 0 | 0 | 0 | 0 | 0 | 0 | 0 |
| 19 | ucpA | 0 | 0 | 0 | 0 | 0 | 0 | 0 | 0 | 0 | 0 | 0 | 0 |
| 20 | cmk | 0 | 0 | 0 | 0 | 0 | 0 | 0 | 0 | 0 | 0 | 0 | 0 |
| 21 | cmr | 0 | 0 | 0 | 0 | 0 | 0 | 0 | 0 | 0 | 0 | 0 | 0 |
| 22 | upp | 0 | 0 | 0 | 0 | 0 | 0 | 0 | 0 | 0 | 0 | 0 | 0 |
| 23 | ydgR | 0 | 0 | 0 | 0 | 0 | 0 | 0 | 1 | 0 | 0 | 0 | 0 |
| 24 | ydgK | 0 | 0 | 0 | 0 | 0 | 0 | 0 | 0 | 0 | 0 | 0 | 0 |
| 25 | ydgH | 0 | 0 | 0 | 0 | 0 | 0 | 0 | 0 | 0 | 0 | 0 | 0 |
| 26 | ydgI | 0 | 0 | 0 | 0 | 0 | 0 | 0 | 0 | 0 | 0 | 0 | 0 |
| 27 | ydgL | 0 | 0 | 1 | 0 | 0 | 0 | 0 | 0 | 0 | 0 | 0 | 0 |
| 28 | tolC | 0 | 0 | 0 | 0 | 0 | 0 | 0 | 0 | 0 | 0 | 0 | 0 |
| 29 | tolB | 0 | 0 | 0 | 0 | 0 | 0 | 0 | 1 | 0 | 0 | 0 | 0 |
| 30 | tolA | 0 | 0 | 0 | 0 | 0 | 0 | 0 | 0 | 0 | 0 | 0 | 0 |
| 31 | ydgA | 0 | 0 | 0 | 0 | 0 | 0 | 0 | 0 | 0 | 0 | 0 | 0 |
| 32 | ydgD | 0 | 0 | 0 | 0 | 0 | 0 | 0 | 0 | 0 | 0 | 0 | 0 |
| 33 | entC | 0 | 0 | 0 | 0 | 0 | 0 | 0 | 0 | 0 | 0 | 0 | 0 |
| 34 | entD | 0 | 1 | 0 | 0 | 0 | 0 | 0 | 0 | 0 | 0 | 0 | 0 |
| 35 | entF | 0 | 0 | 0 | 0 | 1 | 0 | 0 | 0 | 0 | 0 | 0 | 0 |
| 36 | eco | 124 | 0 | 74 | 0 | 2 | 0 | 130 | 0 | 9 | 0 | 216 | 0 |
| 37 | entS | 0 | 0 | 0 | 0 | 0 | 0 | 0 | 0 | 0 | 0 | 0 | 0 |
| 38 | yacG | 0 | 0 | 0 | 0 | 0 | 0 | 0 | 0 | 0 | 0 | 0 | 0 |
| 39 | fecA | 0 | 0 | 0 | 0 | 0 | 0 | 0 | 0 | 0 | 0 | 0 | 0 |
| 40 | yacH | 0 | 0 | 0 | 0 | 0 | 0 | 0 | 0 | 0 | 0 | 0 | 0 |
| 41 | yhbO | 0 | 0 | 0 | 0 | 0 | 0 | 0 | 0 | 0 | 0 | 0 | 0 |
| 42 | fecI | 0 | 0 | 0 | 0 | 0 | 0 | 0 | 0 | 0 | 0 | 0 | 0 |
| 43 | rcsA | 0 | 0 | 0 | 0 | 0 | 1 | 0 | 0 | 0 | 0 | 0 | 0 |
| 44 | b2394 | 0 | 0 | 1 | 0 | 0 | 0 | 0 | 0 | 0 | 0 | 0 | 0 |
| 45 | rcsC | 0 | 0 | 0 | 0 | 0 | 0 | 0 | 0 | 0 | 0 | 0 | 0 |
| 46 | rcsF | 0 | 0 | 0 | 0 | 0 | 0 | 0 | 0 | 0 | 0 | 0 | 0 |
| 47 | uspA | 0 | 0 | 0 | 0 | 0 | 0 | 0 | 0 | 0 | 0 | 0 | 0 |
| 48 | uspB | 0 | 0 | 0 | 0 | 0 | 0 | 0 | 0 | 0 | 0 | 0 | 0 |
| 49 | uspE | 0 | 0 | 0 | 0 | 0 | 0 | 0 | 0 | 0 | 0 | 0 | 0 |
| 50 | uspF | 0 | 0 | 0 | 0 | 0 | 0 | 0 | 0 | 0 | 0 | 0 | 0 |
| 51 | uspG | 0 | 0 | 0 | 0 | 0 | 0 | 0 | 0 | 0 | 0 | 0 | 0 |
| 52 | gltX | 0 | 0 | 0 | 0 | 0 | 0 | 0 | 0 | 0 | 0 | 0 | 0 |
| 53 | yifM2 | 0 | 0 | 0 | 0 | 0 | 0 | 0 | 0 | 0 | 0 | 0 | 0 |
| 54 | gltJ | 0 | 0 | 1 | 0 | 0 | 0 | 0 | 0 | 1 | 0 | 0 | 0 |
| 55 | cspI | 0 | 0 | 0 | 0 | 0 | 0 | 0 | 0 | 0 | 0 | 0 | 0 |
| 56 | gltA | 0 | 0 | 0 | 0 | 0 | 0 | 0 | 0 | 0 | 0 | 0 | 0 |
| 57 | cspG | 0 | 0 | 0 | 0 | 0 | 0 | 0 | 0 | 0 | 1 | 0 | 0 |
| 58 | cspD | 0 | 0 | 0 | 0 | 0 | 0 | 0 | 0 | 0 | 0 | 0 | 0 |
| 59 | gltB | 0 | 0 | 0 | 0 | 0 | 0 | 0 | 0 | 0 | 0 | 0 | 0 |
| 60 | cspB | 0 | 0 | 0 | 0 | 0 | 0 | 0 | 0 | 0 | 0 | 0 | 0 |
| 61 | cspA | 0 | 0 | 0 | 0 | 0 | 0 | 0 | 0 | 0 | 0 | 0 | 0 |
| 62 | yhgN | 0 | 0 | 0 | 0 | 0 | 0 | 0 | 0 | 0 | 0 | 0 | 0 |
| 63 | yfeC | 0 | 0 | 0 | 0 | 0 | 0 | 0 | 0 | 0 | 0 | 0 | 0 |
| 64 | maeB | 0 | 0 | 0 | 0 | 0 | 0 | 0 | 0 | 0 | 0 | 0 | 0 |
| 65 | yhgF | 0 | 0 | 0 | 0 | 0 | 0 | 0 | 0 | 0 | 0 | 0 | 0 |
| 66 | yfeK | 0 | 0 | 0 | 0 | 0 | 0 | 0 | 0 | 0 | 0 | 0 | 0 |
| 67 | yhgE | 0 | 0 | 0 | 0 | 0 | 0 | 0 | 0 | 0 | 0 | 0 | 0 |
| 68 | yfeN | 0 | 0 | 0 | 0 | 0 | 0 | 0 | 0 | 0 | 0 | 0 | 0 |
| 69 | yhgA | 0 | 0 | 0 | 0 | 0 | 0 | 0 | 0 | 0 | 0 | 0 | 0 |
| 70 | yfeY | 0 | 0 | 0 | 0 | 0 | 0 | 0 | 0 | 0 | 0 | 0 | 0 |
| 71 | b0671 | 1089 | 8 | 774 | 3 | 35814 | 293 | 820 | 16 | 364 | 15 | 258 | 4 |
| 72 | b0298 | 0 | 0 | 0 | 0 | 1 | 0 | 0 | 0 | 0 | 0 | 0 | 0 |
| 73 | yjiX | 0 | 0 | 0 | 0 | 0 | 0 | 0 | 0 | 0 | 0 | 0 | 0 |
| 74 | rssA | 0 | 0 | 0 | 0 | 0 | 0 | 0 | 0 | 0 | 0 | 0 | 0 |
| 75 | rssB | 0 | 0 | 0 | 0 | 0 | 0 | 0 | 0 | 0 | 0 | 0 | 0 |
| 76 | ddlA | 0 | 0 | 0 | 0 | 0 | 0 | 0 | 0 | 0 | 0 | 0 | 0 |
| 77 | gatY | 1221 | 134 | 699 | 220 | 5 | 15 | 960 | 272 | 430 | 123 | 175 | 1 |
| 78 | yjiW | 0 | 0 | 0 | 0 | 0 | 0 | 0 | 0 | 0 | 0 | 0 | 0 |
| 79 | ytjB | 0 | 0 | 0 | 0 | 0 | 0 | 0 | 0 | 0 | 0 | 0 | 0 |
| 80 | ygiF | 0 | 0 | 0 | 0 | 0 | 0 | 0 | 0 | 0 | 0 | 0 | 0 |
| 81 | ygiD | 0 | 0 | 0 | 0 | 0 | 1 | 0 | 0 | 0 | 0 | 0 | 0 |
| 82 | ygiH | 0 | 0 | 0 | 0 | 0 | 0 | 0 | 0 | 0 | 0 | 0 | 0 |
| 83 | eno | 0 | 0 | 0 | 0 | 0 | 0 | 0 | 1 | 0 | 0 | 0 | 0 |

| | | | | | | | | | | | | | |
|---|---|---|---|---|---|---|---|---|---|---|---|---|---|
| 84 | ygiS | 0 | 0 | 0 | 0 | 0 | 0 | 0 | 0 | 0 | 0 | 0 | 0 |
| 85 | ygiR | 0 | 0 | 0 | 0 | 0 | 0 | 0 | 0 | 0 | 1 | 0 | 0 |
| 86 | ygiP | 0 | 0 | 0 | 0 | 0 | 0 | 0 | 0 | 0 | 0 | 0 | 0 |
| 87 | ygiW | 0 | 0 | 0 | 0 | 0 | 0 | 0 | 0 | 0 | 0 | 0 | 0 |
| 88 | ygiU | 0 | 0 | 0 | 0 | 0 | 0 | 0 | 0 | 0 | 0 | 0 | 0 |
| 89 | ygiZ | 0 | 0 | 0 | 0 | 0 | 0 | 0 | 0 | 0 | 0 | 0 | 0 |
| 90 | smtA | 0 | 0 | 0 | 0 | 0 | 0 | 0 | 0 | 0 | 0 | 0 | 0 |
| 91 | b0582 | 0 | 0 | 0 | 0 | 0 | 0 | 0 | 0 | 0 | 0 | 0 | 0 |
| 92 | amiA | 0 | 0 | 0 | 0 | 0 | 0 | 0 | 0 | 0 | 0 | 0 | 0 |
| 93 | yhiD | 0 | 0 | 0 | 0 | 0 | 0 | 0 | 0 | 0 | 0 | 0 | 0 |
| 94 | yjiA | 0 | 0 | 0 | 0 | 0 | 0 | 0 | 0 | 0 | 0 | 0 | 0 |
| 95 | hslJ | 0 | 0 | 0 | 0 | 0 | 0 | 0 | 0 | 0 | 0 | 0 | 0 |
| 96 | yhiI | 0 | 0 | 0 | 0 | 0 | 0 | 0 | 0 | 0 | 0 | 0 | 0 |
| 97 | yhiJ | 0 | 0 | 0 | 0 | 0 | 0 | 0 | 0 | 0 | 0 | 0 | 0 |
| 98 | adiA | 0 | 0 | 0 | 0 | 0 | 0 | 0 | 0 | 0 | 0 | 0 | 0 |
| 99 | yhiP | 0 | 0 | 0 | 0 | 0 | 0 | 0 | 0 | 0 | 0 | 0 | 0 |
| 100 | hslV | 0 | 0 | 0 | 0 | 0 | 0 | 0 | 0 | 0 | 0 | 0 | 0 |
| 101 | arcB | 0 | 1 | 0 | 0 | 0 | 0 | 0 | 0 | 0 | 0 | 0 | 0 |
| 102 | yjeN | 0 | 0 | 0 | 0 | 0 | 0 | 0 | 0 | 0 | 0 | 0 | 0 |
| 103 | yjeK | 0 | 0 | 0 | 0 | 0 | 0 | 0 | 0 | 0 | 0 | 0 | 0 |
| 104 | yjeI | 0 | 0 | 0 | 0 | 0 | 0 | 0 | 0 | 0 | 0 | 0 | 0 |
| 105 | fadB | 256 | 64 | 238 | 42 | 1 | 0 | 264 | 80 | 146 | 8 | 30 | 0 |
| 106 | fdrA | 0 | 0 | 0 | 0 | 0 | 0 | 0 | 0 | 0 | 0 | 0 | 0 |
| 107 | fadE | 0 | 0 | 0 | 0 | 0 | 0 | 0 | 0 | 0 | 0 | 0 | 0 |
| 108 | ypeA | 0 | 0 | 0 | 0 | 0 | 0 | 0 | 0 | 0 | 0 | 0 | 0 |
| 109 | creA | 0 | 0 | 0 | 0 | 0 | 0 | 0 | 0 | 0 | 0 | 0 | 0 |
| 110 | creB | 0 | 1 | 1 | 0 | 0 | 0 | 0 | 0 | 0 | 0 | 0 | 0 |
| 111 | inaA | 0 | 0 | 0 | 0 | 0 | 0 | 0 | 0 | 0 | 0 | 0 | 0 |
| 112 | ompT | 0 | 0 | 0 | 0 | 0 | 0 | 0 | 0 | 0 | 0 | 0 | 0 |
| 113 | ompW | 0 | 0 | 0 | 0 | 0 | 0 | 0 | 0 | 0 | 0 | 0 | 0 |
| 114 | ompR | 0 | 0 | 0 | 0 | 0 | 0 | 0 | 0 | 0 | 0 | 0 | 0 |
| 115 | fiu | 0 | 1 | 0 | 0 | 0 | 0 | 0 | 0 | 0 | 0 | 0 | 0 |
| 116 | cld | 0 | 0 | 0 | 0 | 0 | 0 | 0 | 0 | 0 | 0 | 0 | 0 |
| 117 | ompX | 0 | 0 | 0 | 0 | 0 | 0 | 0 | 0 | 0 | 0 | 0 | 0 |
| 118 | yggE | 0 | 0 | 0 | 0 | 0 | 0 | 0 | 0 | 0 | 0 | 0 | 0 |
| 119 | ompG | 0 | 1 | 1 | 0 | 0 | 0 | 0 | 0 | 0 | 0 | 0 | 0 |
| 120 | ompA | 0 | 0 | 0 | 0 | 0 | 0 | 0 | 0 | 0 | 0 | 0 | 0 |
| 121 | ompC | 0 | 0 | 0 | 0 | 0 | 0 | 0 | 0 | 0 | 0 | 0 | 1 |
| 122 | ompN | 0 | 0 | 0 | 0 | 0 | 0 | 0 | 0 | 0 | 0 | 0 | 0 |
| 123 | htrL | 0 | 0 | 0 | 0 | 0 | 0 | 0 | 0 | 0 | 0 | 0 | 0 |
| 124 | cls | 0 | 0 | 0 | 0 | 0 | 0 | 0 | 0 | 0 | 0 | 0 | 0 |
| 125 | gcvP | 0 | 0 | 0 | 0 | 0 | 0 | 0 | 0 | 0 | 0 | 0 | 0 |
| 126 | gcvT | 230 | 3 | 124 | 4 | 2 | 0 | 121 | 1 | 31 | 0 | 0 | 0 |
| 127 | tnaC | 0 | 0 | 0 | 0 | 0 | 0 | 0 | 0 | 0 | 0 | 0 | 0 |
| 128 | srlR | 0 | 0 | 0 | 0 | 0 | 0 | 0 | 1 | 0 | 0 | 0 | 0 |
| 129 | gcvA | 2562 | 6041 | 4104 | 10726 | 116 | 355 | 3405 | 31370 | 7651 | 10566 | 10486 | 54994 |
| 130 | ftsN | 0 | 0 | 1 | 0 | 0 | 0 | 0 | 0 | 1 | 0 | 1 | 0 |
| 131 | ftsK | 0 | 0 | 0 | 0 | 0 | 0 | 0 | 0 | 0 | 0 | 0 | 0 |
| 132 | aniC | 109 | 0 | 60 | 0 | 0 | 0 | 65 | 0 | 19 | 0 | 53 | 0 |
| 133 | ftsH | 0 | 0 | 0 | 0 | 0 | 0 | 0 | 0 | 0 | 0 | 0 | 0 |
| 134 | yieE | 0 | 0 | 0 | 0 | 0 | 0 | 0 | 0 | 0 | 0 | 0 | 0 |
| 135 | ftsQ | 0 | 0 | 0 | 0 | 0 | 0 | 0 | 0 | 0 | 0 | 0 | 0 |
| 136 | ftsZ | 0 | 0 | 0 | 0 | 0 | 0 | 0 | 0 | 0 | 0 | 0 | 0 |
| 137 | ftsY | 0 | 0 | 0 | 0 | 0 | 0 | 0 | 0 | 0 | 0 | 0 | 0 |
| 138 | hemA | 0 | 0 | 0 | 0 | 0 | 0 | 0 | 0 | 0 | 0 | 0 | 0 |
| 139 | yrfF | 0 | 0 | 1 | 0 | 0 | 0 | 0 | 0 | 0 | 0 | 0 | 0 |
| 140 | pgsA | 2832 | 346 | 2414 | 202 | 28 | 0 | 2486 | 12 | 1832 | 589 | 1215 | 2 |
| 141 | fepB | 0 | 0 | 1 | 0 | 0 | 0 | 0 | 0 | 0 | 0 | 0 | 0 |
| 142 | fepA | 0 | 0 | 0 | 0 | 0 | 0 | 0 | 0 | 0 | 0 | 0 | 0 |
| 143 | fepD | 0 | 0 | 0 | 0 | 0 | 0 | 0 | 0 | 0 | 0 | 0 | 0 |
| 144 | yiiR | 0 | 0 | 0 | 0 | 0 | 0 | 0 | 0 | 0 | 0 | 0 | 0 |
| 145 | ligA | 0 | 0 | 0 | 0 | 0 | 0 | 0 | 0 | 0 | 0 | 0 | 0 |
| 146 | ligB | 0 | 0 | 0 | 0 | 0 | 0 | 0 | 0 | 0 | 0 | 0 | 0 |
| 147 | aidB | 0 | 0 | 0 | 0 | 0 | 0 | 0 | 0 | 0 | 0 | 0 | 0 |
| 148 | ligT | 0 | 0 | 0 | 0 | 0 | 0 | 0 | 0 | 0 | 0 | 0 | 0 |
| 149 | smf | 0 | 0 | 0 | 0 | 0 | 0 | 0 | 0 | 0 | 0 | 0 | 0 |
| 150 | prfB | 0 | 0 | 0 | 0 | 0 | 0 | 0 | 0 | 0 | 0 | 0 | 0 |
| 151 | prfC | 0 | 0 | 0 | 0 | 0 | 0 | 0 | 0 | 0 | 0 | 0 | 0 |
| 152 | dcuA | 0 | 0 | 0 | 0 | 0 | 0 | 0 | 0 | 0 | 0 | 0 | 0 |
| 153 | sodB | 0 | 0 | 0 | 0 | 0 | 0 | 0 | 0 | 0 | 0 | 0 | 0 |
| 154 | sodC | 0 | 0 | 0 | 0 | 0 | 0 | 0 | 0 | 0 | 0 | 0 | 1 |
| 155 | sodA | 0 | 0 | 0 | 0 | 0 | 0 | 0 | 0 | 0 | 0 | 0 | 0 |
| 156 | erfK | 0 | 0 | 0 | 0 | 0 | 0 | 0 | 0 | 0 | 0 | 0 | 0 |
| 157 | dcuS | 203 | 2 | 187 | 2 | 3 | 0 | 153 | 0 | 113 | 2 | 164 | 0 |
| 158 | ybbL | 0 | 0 | 0 | 0 | 0 | 0 | 0 | 0 | 0 | 0 | 0 | 0 |
| 159 | oxyR | 0 | 0 | 0 | 0 | 0 | 0 | 0 | 0 | 0 | 0 | 0 | 0 |
| 160 | umuD | 0 | 0 | 0 | 0 | 0 | 0 | 0 | 0 | 0 | 0 | 0 | 0 |
| 161 | ybbC | 0 | 0 | 0 | 0 | 0 | 0 | 0 | 0 | 0 | 0 | 0 | 0 |
| 162 | ybbB | 662 | 1 | 573 | 1 | 3 | 0 | 574 | 2 | 271 | 2 | 410 | 1 |
| 163 | yphG | 0 | 0 | 0 | 0 | 0 | 0 | 0 | 0 | 0 | 0 | 0 | 0 |
| 164 | ybdG | 0 | 0 | 0 | 0 | 0 | 0 | 0 | 0 | 0 | 0 | 0 | 0 |
| 165 | fixC | 0 | 0 | 0 | 0 | 0 | 0 | 0 | 0 | 0 | 0 | 0 | 0 |
| 166 | glyQ | 26 | 0 | 20 | 2 | 1 | 0 | 18 | 1 | 1 | 0 | 0 | 0 |
| 167 | glyU | 0 | 0 | 0 | 0 | 0 | 0 | 0 | 0 | 0 | 0 | 0 | 0 |

Data Set S2 - MG1655 library 7 day experiment
Aligned Reads

| | | | | | | | | | | | | | |
|---|---|---|---|---|---|---|---|---|---|---|---|---|---|
| 168 | aqpZ | 0 | 0 | 0 | 0 | 0 | 0 | 0 | 0 | 0 | 0 | 0 | 0 |
| 169 | glyA | 0 | 0 | 0 | 0 | 0 | 0 | 0 | 0 | 0 | 0 | 0 | 0 |
| 170 | trkA | 0 | 0 | 0 | 0 | 0 | 0 | 0 | 0 | 0 | 0 | 0 | 0 |
| 171 | ynfM | 0 | 0 | 0 | 0 | 0 | 0 | 0 | 0 | 0 | 0 | 0 | 0 |
| 172 | ynfL | 0 | 0 | 0 | 0 | 0 | 0 | 0 | 0 | 0 | 0 | 0 | 0 |
| 173 | ynfK | 0 | 0 | 0 | 0 | 0 | 0 | 0 | 0 | 0 | 0 | 1 | 0 |
| 174 | ynfE | 0 | 0 | 0 | 0 | 0 | 0 | 0 | 0 | 0 | 0 | 0 | 0 |
| 175 | ynfC | 0 | 0 | 0 | 0 | 0 | 0 | 0 | 0 | 0 | 0 | 0 | 0 |
| 176 | ynfB | 0 | 1 | 0 | 1 | 2 | 0 | 0 | 0 | 1 | 0 | 0 | 0 |
| 177 | ynfA | 0 | 0 | 0 | 0 | 1 | 0 | 0 | 0 | 1 | 0 | 0 | 0 |
| 178 | treA | 0 | 0 | 0 | 0 | 1 | 0 | 0 | 0 | 0 | 0 | 0 | 0 |
| 179 | qseB | 0 | 0 | 1 | 0 | 0 | 0 | 0 | 0 | 0 | 0 | 0 | 0 |
| 180 | treB | 0 | 0 | 0 | 0 | 0 | 0 | 0 | 0 | 0 | 0 | 0 | 0 |
| 181 | treF | 0 | 0 | 0 | 0 | 0 | 0 | 1 | 0 | 0 | 0 | 0 | 0 |
| 182 | pgmI | 0 | 0 | 0 | 0 | 0 | 0 | 1 | 0 | 0 | 0 | 0 | 0 |
| 183 | gst | 588 | 0 | 487 | 2 | 9 | 0 | 595 | 1 | 151 | 0 | 795 | 0 |
| 184 | ydfE | 0 | 0 | 0 | 0 | 0 | 0 | 0 | 0 | 0 | 0 | 0 | 0 |
| 185 | ydfA | 0 | 0 | 0 | 0 | 0 | 0 | 0 | 0 | 0 | 0 | 0 | 0 |
| 186 | chbA | 0 | 0 | 0 | 0 | 0 | 0 | 0 | 0 | 0 | 0 | 0 | 0 |
| 187 | ydfO | 0 | 0 | 0 | 0 | 0 | 0 | 0 | 0 | 0 | 0 | 0 | 0 |
| 188 | ydfI | 0 | 0 | 0 | 0 | 0 | 0 | 0 | 0 | 0 | 0 | 0 | 0 |
| 189 | ydfH | 0 | 0 | 0 | 0 | 0 | 0 | 0 | 0 | 0 | 0 | 0 | 0 |
| 190 | ydfV | 0 | 0 | 0 | 0 | 0 | 0 | 0 | 0 | 0 | 0 | 0 | 0 |
| 191 | ydfZ | 0 | 0 | 0 | 0 | 0 | 1 | 0 | 0 | 0 | 0 | 0 | 0 |
| 192 | serW | 0 | 0 | 0 | 0 | 0 | 0 | 0 | 0 | 1 | 0 | 0 | 0 |
| 193 | pdhR | 582 | 1 | 483 | 3 | 36 | 0 | 376 | 0 | 360 | 0 | 635 | 0 |
| 194 | ydgC | 0 | 0 | 1 | 0 | 0 | 0 | 0 | 0 | 0 | 0 | 0 | 0 |
| 195 | pqiA | 0 | 0 | 0 | 0 | 0 | 0 | 0 | 0 | 0 | 0 | 0 | 0 |
| 196 | yhhF | 0 | 0 | 0 | 0 | 0 | 0 | 0 | 0 | 0 | 0 | 0 | 0 |
| 197 | flgB | 0 | 0 | 0 | 0 | 0 | 0 | 0 | 0 | 0 | 0 | 0 | 0 |
| 198 | yhhN | 0 | 0 | 0 | 0 | 0 | 0 | 0 | 1 | 0 | 0 | 0 | 0 |
| 199 | yhhM | 0 | 0 | 0 | 0 | 0 | 0 | 0 | 0 | 0 | 0 | 0 | 0 |
| 200 | minC | 0 | 0 | 0 | 0 | 0 | 0 | 0 | 0 | 0 | 0 | 0 | 0 |
| 201 | flgA | 0 | 0 | 0 | 0 | 0 | 0 | 0 | 0 | 0 | 1 | 0 | 0 |
| 202 | yhhW | 0 | 0 | 0 | 0 | 0 | 0 | 0 | 0 | 0 | 0 | 0 | 0 |
| 203 | yhhT | 992 | 4 | 730 | 2 | 27 | 1 | 877 | 0 | 301 | 0 | 545 | 1 |
| 204 | adhE | 0 | 0 | 0 | 0 | 0 | 0 | 0 | 0 | 0 | 0 | 0 | 0 |
| 205 | yjdF | 618 | 3 | 373 | 4 | 23 | 0 | 476 | 0 | 193 | 1 | 0 | 0 |
| 206 | yjdI | 246 | 0 | 164 | 2 | 0 | 0 | 170 | 0 | 62 | 1 | 172 | 0 |
| 207 | slyB | 0 | 0 | 0 | 0 | 0 | 0 | 0 | 0 | 0 | 0 | 0 | 0 |
| 208 | yjdK | 0 | 0 | 0 | 0 | 0 | 0 | 0 | 0 | 0 | 0 | 0 | 0 |
| 209 | yhhZ | 0 | 0 | 0 | 0 | 0 | 0 | 0 | 0 | 0 | 0 | 0 | 0 |
| 210 | yhhY | 0 | 0 | 0 | 0 | 0 | 0 | 0 | 0 | 0 | 0 | 0 | 0 |
| 211 | yfdL | 0 | 0 | 0 | 0 | 0 | 0 | 0 | 0 | 0 | 0 | 0 | 0 |
| 212 | yhfK | 0 | 0 | 0 | 0 | 0 | 0 | 1 | 0 | 1 | 0 | 0 | 0 |
| 213 | yhfL | 0 | 0 | 0 | 0 | 0 | 0 | 0 | 0 | 0 | 0 | 1 | 0 |
| 214 | yhfA | 0 | 0 | 0 | 0 | 0 | 0 | 0 | 0 | 0 | 0 | 0 | 0 |
| 215 | yfdG | 0 | 0 | 0 | 0 | 0 | 0 | 0 | 0 | 0 | 0 | 0 | 0 |
| 216 | sgcC | 0 | 0 | 0 | 0 | 0 | 0 | 0 | 0 | 0 | 0 | 0 | 0 |
| 217 | yfdC | 0 | 0 | 0 | 0 | 0 | 0 | 0 | 0 | 0 | 0 | 0 | 0 |
| 218 | yhfX | 0 | 0 | 0 | 0 | 0 | 0 | 0 | 1 | 0 | 0 | 0 | 0 |
| 219 | yhfZ | 0 | 0 | 0 | 0 | 0 | 0 | 0 | 0 | 0 | 0 | 0 | 0 |
| 220 | yfdZ | 0 | 0 | 0 | 0 | 0 | 0 | 0 | 0 | 0 | 0 | 0 | 0 |
| 221 | yfdU | 0 | 0 | 0 | 0 | 0 | 0 | 0 | 0 | 0 | 0 | 0 | 0 |
| 222 | yhfP | 0 | 0 | 0 | 0 | 0 | 0 | 0 | 0 | 0 | 0 | 0 | 0 |
| 223 | yfdV | 0 | 0 | 0 | 0 | 0 | 0 | 0 | 0 | 0 | 0 | 0 | 0 |
| 224 | mcrB | 0 | 0 | 0 | 0 | 0 | 0 | 0 | 0 | 0 | 0 | 0 | 0 |
| 225 | gsk | 0 | 0 | 0 | 0 | 0 | 0 | 0 | 0 | 0 | 0 | 0 | 0 |
| 226 | mcrA | 0 | 0 | 0 | 0 | 0 | 0 | 0 | 0 | 0 | 0 | 0 | 0 |
| 227 | flgH | 841 | 9 | 543 | 0 | 5 | 0 | 701 | 0 | 391 | 0 | 146 | 0 |
| 228 | gloA | 0 | 0 | 1 | 0 | 0 | 0 | 0 | 0 | 0 | 0 | 0 | 0 |
| 229 | lpp | 0 | 0 | 0 | 0 | 0 | 0 | 0 | 0 | 0 | 0 | 0 | 0 |
| 230 | lpd | 0 | 0 | 0 | 0 | 0 | 2 | 0 | 0 | 0 | 0 | 0 | 0 |
| 231 | arpA | 0 | 0 | 0 | 0 | 0 | 0 | 0 | 0 | 0 | 0 | 0 | 0 |
| 232 | aldH | 0 | 0 | 0 | 0 | 0 | 0 | 0 | 0 | 0 | 0 | 0 | 0 |
| 233 | menD | 0 | 0 | 0 | 0 | 0 | 0 | 0 | 0 | 0 | 0 | 0 | 0 |
| 234 | sieB | 0 | 0 | 0 | 0 | 0 | 0 | 0 | 0 | 0 | 0 | 0 | 0 |
| 235 | menF | 0 | 0 | 0 | 0 | 0 | 0 | 0 | 1 | 1 | 0 | 0 | 2 |
| 236 | menG | 0 | 0 | 0 | 0 | 0 | 0 | 0 | 0 | 0 | 0 | 0 | 0 |
| 237 | aldB | 0 | 0 | 0 | 0 | 0 | 0 | 0 | 0 | 0 | 0 | 0 | 0 |
| 238 | aldA | 0 | 0 | 0 | 0 | 0 | 0 | 0 | 0 | 0 | 0 | 0 | 0 |
| 239 | sbcB | 0 | 0 | 0 | 0 | 1 | 0 | 0 | 0 | 0 | 0 | 0 | 0 |
| 240 | priA | 0 | 0 | 0 | 0 | 0 | 0 | 0 | 0 | 0 | 0 | 0 | 0 |
| 241 | priC | 0 | 0 | 0 | 0 | 0 | 0 | 0 | 0 | 0 | 0 | 0 | 0 |
| 242 | yojI | 0 | 0 | 0 | 0 | 0 | 0 | 0 | 0 | 0 | 0 | 0 | 0 |
| 243 | yebU | 0 | 0 | 0 | 0 | 0 | 0 | 0 | 0 | 0 | 0 | 0 | 0 |
| 244 | yebQ | 0 | 0 | 0 | 0 | 0 | 0 | 0 | 0 | 0 | 0 | 0 | 0 |
| 245 | yebR | 0 | 0 | 0 | 0 | 0 | 0 | 0 | 0 | 0 | 0 | 0 | 0 |
| 246 | yebS | 0 | 0 | 0 | 0 | 0 | 0 | 0 | 0 | 0 | 0 | 0 | 0 |
| 247 | yebO | 0 | 0 | 0 | 0 | 0 | 0 | 0 | 0 | 0 | 0 | 0 | 0 |
| 248 | yebE | 0 | 0 | 0 | 0 | 0 | 0 | 0 | 0 | 0 | 0 | 0 | 0 |
| 249 | yebF | 729 | 4 | 459 | 1 | 1 | 0 | 438 | 0 | 415 | 0 | 504 | 0 |
| 250 | yebG | 0 | 0 | 0 | 0 | 0 | 0 | 0 | 0 | 0 | 0 | 0 | 0 |
| 251 | yebA | 0 | 0 | 0 | 0 | 0 | 0 | 0 | 0 | 0 | 0 | 0 | 0 |

| | | | | | | | | | | | | | |
|---|---|---|---|---|---|---|---|---|---|---|---|---|---|
| 252 | yebB | 0 | 0 | 0 | 0 | 0 | 0 | 0 | 0 | 0 | 0 | 0 | 0 |
| 253 | ygjN | 0 | 0 | 1 | 1 | 0 | 0 | 0 | 0 | 0 | 0 | 0 | 0 |
| 254 | ygjO | 1 | 0 | 0 | 0 | 0 | 0 | 0 | 0 | 0 | 0 | 0 | 0 |
| 255 | ygjJ | 0 | 0 | 0 | 0 | 0 | 0 | 0 | 1 | 0 | 1 | 0 | 0 |
| 256 | ygjH | 0 | 0 | 0 | 0 | 0 | 0 | 0 | 0 | 0 | 0 | 0 | 0 |
| 257 | ygjI | 0 | 0 | 0 | 0 | 0 | 0 | 0 | 0 | 0 | 0 | 0 | 0 |
| 258 | ygjG | 356 | 0 | 181 | 3 | 343 | 5 | 242 | 1 | 148 | 1 | 782 | 1 |
| 259 | ygjD | 0 | 0 | 1 | 0 | 0 | 0 | 0 | 0 | 0 | 0 | 1 | 0 |
| 260 | bax | 0 | 0 | 0 | 0 | 0 | 0 | 0 | 0 | 0 | 0 | 0 | 0 |
| 261 | ygjV | 0 | 0 | 0 | 0 | 0 | 0 | 1 | 0 | 0 | 0 | 0 | 0 |
| 262 | yacL | 0 | 0 | 0 | 0 | 0 | 0 | 1 | 0 | 0 | 0 | 0 | 0 |
| 263 | ygjR | 0 | 0 | 1 | 0 | 0 | 0 | 0 | 0 | 0 | 0 | 0 | 0 |
| 264 | ygjQ | 0 | 0 | 0 | 0 | 0 | 0 | 0 | 0 | 0 | 0 | 0 | 0 |
| 265 | ecfG | 0 | 0 | 0 | 0 | 0 | 0 | 0 | 0 | 0 | 0 | 0 | 0 |
| 266 | ecfD | 0 | 0 | 0 | 0 | 0 | 0 | 0 | 0 | 0 | 0 | 0 | 0 |
| 267 | ecfE | 0 | 0 | 0 | 0 | 0 | 0 | 0 | 0 | 0 | 0 | 0 | 0 |
| 268 | ecfK | 0 | 0 | 0 | 0 | 0 | 0 | 0 | 0 | 0 | 0 | 0 | 0 |
| 269 | ecfI | 0 | 0 | 0 | 0 | 0 | 0 | 0 | 0 | 0 | 0 | 0 | 0 |
| 270 | dctA | 0 | 0 | 0 | 1 | 0 | 0 | 0 | 0 | 0 | 0 | 0 | 0 |
| 271 | pmrD | 0 | 0 | 0 | 0 | 0 | 0 | 0 | 0 | 0 | 0 | 0 | 0 |
| 272 | ybaK | 0 | 0 | 0 | 0 | 0 | 0 | 0 | 0 | 0 | 0 | 0 | 0 |
| 273 | ybaN | 0 | 0 | 0 | 0 | 0 | 0 | 0 | 0 | 0 | 0 | 0 | 0 |
| 274 | ybaO | 0 | 0 | 0 | 0 | 0 | 0 | 1 | 0 | 0 | 0 | 0 | 0 |
| 275 | ybaL | 0 | 0 | 0 | 0 | 0 | 0 | 0 | 0 | 0 | 0 | 0 | 0 |
| 276 | ybaB | 1 | 0 | 0 | 0 | 0 | 0 | 0 | 0 | 0 | 0 | 0 | 0 |
| 277 | ybaA | 0 | 0 | 0 | 0 | 0 | 0 | 0 | 0 | 0 | 0 | 0 | 0 |
| 278 | ybaD | 0 | 0 | 0 | 0 | 0 | 0 | 0 | 0 | 0 | 0 | 0 | 0 |
| 279 | ybaE | 0 | 0 | 0 | 0 | 0 | 0 | 0 | 0 | 0 | 0 | 0 | 0 |
| 280 | mug | 0 | 0 | 0 | 0 | 0 | 0 | 0 | 0 | 0 | 0 | 0 | 0 |
| 281 | ybaY | 0 | 0 | 0 | 0 | 0 | 0 | 0 | 0 | 0 | 0 | 0 | 0 |
| 282 | yzgL | 0 | 0 | 0 | 0 | 0 | 0 | 0 | 0 | 0 | 0 | 0 | 0 |
| 283 | ybaP | 0 | 0 | 0 | 0 | 0 | 0 | 0 | 0 | 0 | 0 | 0 | 0 |
| 284 | ybaQ | 0 | 0 | 0 | 0 | 0 | 0 | 0 | 0 | 0 | 0 | 0 | 0 |
| 285 | ybaV | 0 | 0 | 0 | 0 | 0 | 0 | 0 | 0 | 0 | 0 | 0 | 0 |
| 286 | ybaT | 0 | 0 | 0 | 1 | 0 | 0 | 0 | 0 | 0 | 0 | 0 | 0 |
| 287 | yijF | 0 | 0 | 0 | 0 | 0 | 0 | 0 | 0 | 0 | 0 | 0 | 0 |
| 288 | ygdL | 0 | 0 | 0 | 0 | 0 | 0 | 0 | 0 | 0 | 0 | 0 | 0 |
| 289 | ygdI_csdA | 0 | 0 | 0 | 0 | 0 | 0 | 0 | 0 | 1 | 0 | 0 | 1 |
| 290 | yijO | 222 | 0 | 156 | 0 | 18 | 0 | 218 | 0 | 27 | 1 | 0 | 0 |
| 291 | ygdQ | 0 | 0 | 0 | 0 | 0 | 0 | 0 | 0 | 0 | 0 | 0 | 0 |
| 292 | ygdR | 0 | 0 | 0 | 0 | 0 | 0 | 0 | 0 | 0 | 0 | 0 | 0 |
| 293 | pitA | 0 | 0 | 0 | 0 | 0 | 0 | 0 | 0 | 0 | 0 | 0 | 0 |
| 294 | b2974 | 0 | 0 | 0 | 0 | 0 | 0 | 0 | 0 | 0 | 0 | 0 | 0 |
| 295 | pitB | 0 | 0 | 0 | 0 | 0 | 0 | 0 | 0 | 0 | 0 | 0 | 0 |
| 296 | ung | 0 | 0 | 0 | 0 | 0 | 0 | 0 | 0 | 0 | 0 | 0 | 0 |
| 297 | rimL | 0 | 0 | 0 | 0 | 0 | 0 | 0 | 0 | 0 | 1 | 0 | 0 |
| 298 | rimJ | 0 | 0 | 0 | 0 | 0 | 0 | 0 | 0 | 0 | 0 | 0 | 0 |
| 299 | insA2 | 0 | 0 | 0 | 0 | 0 | 0 | 0 | 0 | 0 | 0 | 0 | 0 |
| 300 | tonB | 0 | 0 | 0 | 0 | 0 | 0 | 0 | 0 | 0 | 0 | 0 | 0 |
| 301 | ydaG | 0 | 0 | 0 | 0 | 0 | 0 | 0 | 0 | 0 | 0 | 0 | 0 |
| 302 | crcA | 0 | 0 | 0 | 0 | 0 | 0 | 0 | 0 | 0 | 0 | 0 | 1 |
| 303 | yaaW | 0 | 0 | 0 | 0 | 0 | 0 | 0 | 0 | 0 | 0 | 0 | 0 |
| 304 | ydaO | 0 | 0 | 0 | 0 | 0 | 0 | 0 | 0 | 0 | 0 | 0 | 0 |
| 305 | yaaI | 0 | 0 | 0 | 0 | 0 | 0 | 0 | 0 | 0 | 0 | 0 | 0 |
| 306 | frr | 0 | 0 | 0 | 0 | 0 | 0 | 0 | 1 | 0 | 0 | 0 | 0 |
| 307 | yaaJ | 0 | 0 | 0 | 0 | 0 | 0 | 0 | 0 | 0 | 0 | 0 | 0 |
| 308 | yaaA | 0 | 0 | 0 | 0 | 0 | 0 | 0 | 0 | 0 | 0 | 0 | 0 |
| 309 | lctP | 0 | 0 | 0 | 0 | 0 | 0 | 0 | 0 | 0 | 0 | 0 | 0 |
| 310 | znuC | 0 | 0 | 0 | 0 | 0 | 0 | 0 | 0 | 0 | 0 | 0 | 0 |
| 311 | znuA | 0 | 0 | 0 | 0 | 0 | 0 | 0 | 0 | 0 | 0 | 0 | 0 |
| 312 | slt | 0 | 0 | 0 | 0 | 1 | 0 | 0 | 0 | 0 | 0 | 0 | 0 |
| 313 | ais | 0 | 0 | 0 | 0 | 0 | 0 | 0 | 0 | 0 | 0 | 0 | 0 |
| 314 | hepA | 1 | 0 | 1 | 0 | 0 | 0 | 0 | 0 | 0 | 0 | 0 | 0 |
| 315 | gnd | 0 | 0 | 0 | 0 | 0 | 0 | 0 | 0 | 0 | 0 | 0 | 0 |
| 316 | lepA | 0 | 0 | 0 | 0 | 0 | 0 | 0 | 0 | 0 | 0 | 0 | 0 |
| 317 | hupA | 0 | 0 | 0 | 0 | 0 | 0 | 0 | 1 | 0 | 0 | 0 | 0 |
| 318 | yfcI | 0 | 0 | 0 | 0 | 0 | 0 | 0 | 0 | 0 | 0 | 0 | 0 |
| 319 | yfcJ | 0 | 0 | 0 | 0 | 0 | 0 | 0 | 0 | 0 | 0 | 0 | 0 |
| 320 | hupB | 0 | 0 | 0 | 0 | 0 | 0 | 0 | 0 | 0 | 0 | 0 | 1 |
| 321 | yfcC | 887 | 4 | 717 | 2 | 4 | 0 | 508 | 1 | 391 | 1 | 249 | 0 |
| 322 | yfcD | 0 | 0 | 0 | 0 | 0 | 0 | 0 | 0 | 0 | 0 | 0 | 0 |
| 323 | yfcE | 0 | 0 | 0 | 0 | 0 | 0 | 0 | 0 | 0 | 0 | 0 | 0 |
| 324 | yfcG | 0 | 0 | 0 | 0 | 0 | 0 | 0 | 0 | 0 | 0 | 0 | 0 |
| 325 | yfcU | 0 | 0 | 0 | 0 | 0 | 0 | 0 | 0 | 0 | 0 | 0 | 0 |
| 326 | yfcV | 0 | 0 | 0 | 0 | 0 | 0 | 0 | 0 | 0 | 0 | 0 | 0 |
| 327 | glnK | 0 | 0 | 0 | 0 | 0 | 0 | 0 | 0 | 0 | 0 | 0 | 0 |
| 328 | glnH | 0 | 0 | 1 | 0 | 0 | 0 | 1 | 0 | 0 | 0 | 0 | 0 |
| 329 | glnL | 0 | 0 | 0 | 0 | 0 | 0 | 0 | 0 | 0 | 0 | 0 | 0 |
| 330 | glnB | 0 | 0 | 0 | 0 | 0 | 0 | 0 | 0 | 0 | 0 | 0 | 0 |
| 331 | glnA | 0 | 0 | 0 | 0 | 0 | 0 | 0 | 0 | 0 | 0 | 0 | 0 |
| 332 | glnW | 0 | 0 | 0 | 0 | 0 | 0 | 0 | 0 | 0 | 0 | 0 | 0 |
| 333 | nudE | 1 | 0 | 0 | 0 | 1 | 0 | 0 | 0 | 0 | 0 | 0 | 0 |
| 334 | glnU | 0 | 0 | 1 | 0 | 2 | 0 | 0 | 0 | 1 | 0 | 0 | 0 |
| 335 | cynT | 2 | 0 | 8 | 0 | 0 | 0 | 6 | 0 | 2 | 0 | 2 | 0 |

### Data Set S2 - MG1655 library 7 day experiment
Aligned Reads

| | | | | | | | | | | | | | |
|---|---|---|---|---|---|---|---|---|---|---|---|---|---|
| 336 | cynR | 42939 | 1563 | 43271 | 1717 | 14250 | 914 | 37005 | 1922 | 24873 | 1765 | 27677 | 679 |
| 337 | rne | 0 | 0 | 1 | 0 | 0 | 0 | 1 | 0 | 0 | 0 | 1 | 0 |
| 338 | glxK | 0 | 0 | 0 | 0 | 0 | 0 | 0 | 1 | 1 | 0 | 0 | 0 |
| 339 | kil | 0 | 0 | 0 | 0 | 0 | 0 | 0 | 0 | 0 | 0 | 0 | 0 |
| 340 | gssA | 0 | 0 | 0 | 0 | 0 | 0 | 0 | 0 | 0 | 0 | 0 | 0 |
| 341 | yneJ | 0 | 0 | 0 | 0 | 0 | 0 | 0 | 0 | 0 | 0 | 0 | 0 |
| 342 | yneK | 0 | 0 | 0 | 0 | 0 | 0 | 0 | 0 | 0 | 0 | 0 | 0 |
| 343 | yneH | 0 | 0 | 1 | 0 | 0 | 0 | 0 | 0 | 0 | 0 | 0 | 0 |
| 344 | yneI | 0 | 0 | 0 | 0 | 0 | 0 | 0 | 0 | 0 | 0 | 0 | 0 |
| 345 | amyA | 0 | 0 | 0 | 1 | 0 | 0 | 0 | 0 | 0 | 0 | 0 | 0 |
| 346 | yneE | 0 | 0 | 0 | 0 | 0 | 0 | 0 | 0 | 0 | 0 | 0 | 0 |
| 347 | yecM | 0 | 0 | 0 | 0 | 0 | 0 | 0 | 0 | 0 | 0 | 0 | 1 |
| 348 | yfeA | 0 | 0 | 0 | 0 | 0 | 0 | 0 | 0 | 0 | 0 | 0 | 0 |
| 349 | yecH | 0 | 0 | 0 | 0 | 0 | 0 | 0 | 0 | 0 | 0 | 0 | 0 |
| 350 | yecF | 0 | 0 | 0 | 0 | 0 | 0 | 0 | 0 | 0 | 0 | 0 | 0 |
| 351 | fadI | 0 | 0 | 0 | 0 | 1 | 0 | 1 | 0 | 0 | 0 | 0 | 0 |
| 352 | yjjU | 0 | 0 | 0 | 0 | 0 | 0 | 0 | 2 | 0 | 0 | 0 | 0 |
| 353 | ychJ | 0 | 0 | 1 | 0 | 0 | 0 | 0 | 0 | 0 | 0 | 0 | 0 |
| 354 | rnk | 0 | 0 | 0 | 0 | 0 | 0 | 0 | 0 | 0 | 0 | 0 | 0 |
| 355 | astC | 0 | 0 | 0 | 0 | 0 | 0 | 0 | 0 | 0 | 0 | 0 | 0 |
| 356 | nudH | 0 | 0 | 0 | 0 | 0 | 0 | 0 | 0 | 0 | 0 | 0 | 0 |
| 357 | nudG | 0 | 0 | 0 | 0 | 0 | 0 | 0 | 0 | 0 | 0 | 0 | 0 |
| 358 | nudF | 0 | 0 | 0 | 0 | 0 | 0 | 0 | 0 | 0 | 0 | 0 | 0 |
| 359 | rna | 0 | 1 | 0 | 0 | 0 | 0 | 0 | 0 | 0 | 0 | 0 | 0 |
| 360 | yliG | 0 | 0 | 0 | 0 | 0 | 0 | 0 | 0 | 0 | 0 | 0 | 0 |
| 361 | nudC | 848 | 2 | 432 | 1 | 10 | 1 | 436 | 0 | 334 | 0 | 67 | 1 |
| 362 | nudB | 0 | 0 | 0 | 0 | 0 | 0 | 1 | 0 | 0 | 0 | 0 | 0 |
| 363 | gadB | 0 | 0 | 0 | 0 | 0 | 0 | 0 | 0 | 0 | 0 | 0 | 0 |
| 364 | rnd | 1 | 0 | 0 | 0 | 0 | 0 | 0 | 0 | 0 | 0 | 0 | 0 |
| 365 | gadX | 0 | 0 | 0 | 0 | 0 | 0 | 0 | 1 | 0 | 0 | 1 | 0 |
| 366 | gadW | 0 | 0 | 0 | 0 | 0 | 0 | 0 | 0 | 0 | 0 | 0 | 0 |
| 367 | rnt | 0 | 0 | 0 | 0 | 0 | 0 | 0 | 0 | 0 | 0 | 0 | 0 |
| 368 | stpA | 0 | 0 | 0 | 0 | 0 | 0 | 0 | 0 | 0 | 0 | 0 | 0 |
| 369 | araB | 0 | 0 | 0 | 0 | 0 | 0 | 0 | 0 | 0 | 0 | 0 | 0 |
| 370 | araC | 0 | 0 | 0 | 0 | 0 | 0 | 0 | 0 | 0 | 0 | 0 | 0 |
| 371 | araF | 0 | 0 | 0 | 0 | 0 | 0 | 0 | 0 | 0 | 0 | 0 | 0 |
| 372 | araD | 0 | 0 | 0 | 0 | 0 | 0 | 0 | 0 | 0 | 0 | 0 | 0 |
| 373 | araE | 2 | 0 | 0 | 0 | 0 | 0 | 0 | 0 | 0 | 0 | 0 | 0 |
| 374 | araJ | 0 | 0 | 0 | 0 | 0 | 0 | 0 | 0 | 1 | 0 | 0 | 0 |
| 375 | ygeG | 0 | 0 | 0 | 0 | 0 | 0 | 0 | 0 | 0 | 0 | 0 | 0 |
| 376 | ygeF | 5 | 0 | 2 | 0 | 0 | 0 | 4 | 0 | 0 | 0 | 0 | 0 |
| 377 | ygeA | 0 | 0 | 0 | 0 | 0 | 0 | 0 | 0 | 0 | 0 | 0 | 0 |
| 378 | ygeI | 0 | 0 | 0 | 0 | 0 | 0 | 0 | 0 | 0 | 0 | 0 | 0 |
| 379 | cirA | 0 | 0 | 0 | 0 | 0 | 0 | 0 | 0 | 0 | 0 | 0 | 0 |
| 380 | ygeR | 0 | 0 | 0 | 0 | 0 | 0 | 0 | 0 | 0 | 1 | 0 | 0 |
| 381 | ygeQ | 0 | 0 | 0 | 0 | 0 | 0 | 0 | 0 | 0 | 0 | 0 | 0 |
| 382 | fruL | 0 | 0 | 0 | 0 | 0 | 0 | 0 | 0 | 0 | 0 | 0 | 0 |
| 383 | fruB | 0 | 0 | 0 | 0 | 0 | 0 | 0 | 0 | 0 | 0 | 0 | 0 |
| 384 | ygeY | 0 | 0 | 0 | 0 | 0 | 0 | 0 | 0 | 0 | 0 | 0 | 0 |
| 385 | ygeX | 0 | 0 | 0 | 0 | 0 | 0 | 0 | 0 | 0 | 0 | 0 | 0 |
| 386 | rffC | 0 | 0 | 0 | 0 | 0 | 0 | 0 | 0 | 0 | 0 | 0 | 0 |
| 387 | ypfG | 0 | 0 | 1 | 0 | 0 | 0 | 0 | 0 | 0 | 0 | 0 | 0 |
| 388 | ypfH | 0 | 0 | 0 | 0 | 0 | 0 | 0 | 0 | 0 | 0 | 1 | 0 |
| 389 | ypfJ | 0 | 0 | 0 | 0 | 0 | 0 | 0 | 0 | 0 | 0 | 0 | 0 |
| 390 | deaD | 0 | 0 | 0 | 0 | 0 | 0 | 0 | 0 | 0 | 0 | 0 | 0 |
| 391 | icd | 0 | 0 | 0 | 0 | 0 | 0 | 0 | 0 | 0 | 0 | 0 | 0 |
| 392 | borD | 0 | 0 | 0 | 0 | 0 | 0 | 0 | 1 | 0 | 0 | 0 | 0 |
| 393 | hdeD | 0 | 0 | 0 | 0 | 0 | 0 | 0 | 0 | 0 | 0 | 0 | 0 |
| 394 | hdeB | 0 | 0 | 0 | 0 | 0 | 0 | 0 | 0 | 0 | 0 | 0 | 0 |
| 395 | hdeA | 0 | 0 | 0 | 0 | 0 | 0 | 0 | 0 | 0 | 0 | 0 | 0 |
| 396 | ribA | 0 | 0 | 0 | 0 | 0 | 0 | 0 | 0 | 0 | 0 | 0 | 0 |
| 397 | ribC | 0 | 0 | 0 | 0 | 0 | 0 | 0 | 0 | 0 | 0 | 0 | 0 |
| 398 | ribB | 0 | 0 | 0 | 0 | 0 | 0 | 0 | 0 | 0 | 0 | 0 | 0 |
| 399 | ribE | 0 | 0 | 0 | 0 | 0 | 0 | 0 | 0 | 0 | 0 | 0 | 0 |
| 400 | htpX | 0 | 0 | 0 | 0 | 0 | 0 | 0 | 0 | 0 | 0 | 0 | 0 |
| 401 | yqjI | 0 | 0 | 0 | 0 | 0 | 0 | 0 | 0 | 0 | 0 | 0 | 0 |
| 402 | corA | 0 | 0 | 0 | 0 | 0 | 0 | 0 | 0 | 1 | 0 | 0 | 0 |
| 403 | yqjC | 0 | 0 | 0 | 1 | 0 | 0 | 0 | 0 | 0 | 0 | 0 | 0 |
| 404 | yqjF | 0 | 0 | 0 | 0 | 0 | 0 | 0 | 0 | 0 | 0 | 0 | 0 |
| 405 | yqjG | 0 | 0 | 0 | 0 | 0 | 0 | 0 | 0 | 0 | 0 | 0 | 0 |
| 406 | pheS | 0 | 0 | 0 | 0 | 0 | 0 | 0 | 0 | 0 | 0 | 0 | 0 |
| 407 | pheP | 0 | 0 | 0 | 0 | 0 | 0 | 0 | 0 | 0 | 0 | 0 | 0 |
| 408 | ileX | 0 | 0 | 0 | 0 | 0 | 0 | 0 | 0 | 0 | 0 | 0 | 0 |
| 409 | htpG | 0 | 0 | 0 | 0 | 0 | 0 | 1 | 0 | 0 | 0 | 0 | 0 |
| 410 | ppa | 0 | 0 | 0 | 0 | 0 | 0 | 0 | 0 | 0 | 0 | 0 | 0 |
| 411 | dkgB | 0 | 0 | 0 | 0 | 0 | 0 | 0 | 0 | 0 | 0 | 0 | 0 |
| 412 | dkgA | 0 | 0 | 0 | 0 | 0 | 0 | 0 | 0 | 0 | 0 | 0 | 0 |
| 413 | relB | 0 | 0 | 0 | 0 | 0 | 0 | 0 | 0 | 0 | 0 | 0 | 0 |
| 414 | relA | 0 | 0 | 0 | 0 | 0 | 0 | 0 | 0 | 0 | 0 | 0 | 0 |
| 415 | phnQ | 1896 | 4 | 1294 | 2 | 1881 | 16 | 1278 | 1 | 652 | 5 | 2378 | 3 |
| 416 | pykF | 1 | 0 | 0 | 0 | 0 | 0 | 0 | 0 | 0 | 0 | 0 | 0 |
| 417 | sfmA | 0 | 0 | 0 | 0 | 0 | 0 | 0 | 0 | 0 | 0 | 0 | 0 |
| 418 | sfmC | 0 | 0 | 0 | 0 | 0 | 0 | 0 | 0 | 0 | 0 | 0 | 0 |
| 419 | b1997_b1998 | 0 | 0 | 0 | 0 | 0 | 0 | 0 | 0 | 0 | 0 | 0 | 0 |

| | | | | | | | | | | | | | |
|---|---|---|---|---|---|---|---|---|---|---|---|---|---|
| 420 | pphA | 0 | 0 | 0 | 0 | 0 | 0 | 0 | 0 | 0 | 0 | 0 | 0 |
| 421 | mtr | 0 | 0 | 0 | 0 | 0 | 0 | 0 | 0 | 0 | 0 | 0 | 0 |
| 422 | thrU | 0 | 0 | 0 | 0 | 0 | 0 | 0 | 0 | 0 | 0 | 0 | 1 |
| 423 | msbB | 387 | 2 | 292 | 0 | 12 | 0 | 299 | 0 | 184 | 0 | 342 | 0 |
| 424 | dacA | 0 | 0 | 0 | 1 | 0 | 0 | 1 | 0 | 0 | 0 | 0 | 0 |
| 425 | dacB | 0 | 0 | 0 | 0 | 0 | 0 | 0 | 0 | 0 | 0 | 0 | 0 |
| 426 | tesB | 0 | 0 | 0 | 0 | 0 | 0 | 0 | 0 | 0 | 0 | 0 | 0 |
| 427 | yigI | 0 | 0 | 0 | 0 | 0 | 0 | 0 | 0 | 0 | 0 | 0 | 0 |
| 428 | cyoA | 0 | 0 | 0 | 0 | 0 | 0 | 0 | 0 | 0 | 0 | 0 | 0 |
| 429 | dgkA | 0 | 0 | 0 | 0 | 0 | 0 | 0 | 0 | 0 | 0 | 0 | 0 |
| 430 | shiA | 0 | 0 | 0 | 0 | 0 | 0 | 0 | 0 | 0 | 1 | 0 | 0 |
| 431 | ybfG | 0 | 0 | 0 | 0 | 0 | 0 | 0 | 0 | 0 | 0 | 0 | 0 |
| 432 | tig | 0 | 0 | 0 | 0 | 1 | 0 | 0 | 0 | 0 | 0 | 0 | 0 |
| 433 | ykgD | 0 | 0 | 0 | 0 | 0 | 0 | 0 | 0 | 0 | 0 | 0 | 0 |
| 434 | sdaA | 0 | 0 | 0 | 0 | 0 | 0 | 0 | 0 | 0 | 0 | 0 | 0 |
| 435 | nmpC | 0 | 0 | 1 | 0 | 0 | 0 | 0 | 0 | 0 | 0 | 0 | 0 |
| 436 | idnT | 101 | 0 | 53 | 0 | 0 | 0 | 33 | 0 | 24 | 1 | 0 | 0 |
| 437 | idnR | 0 | 0 | 0 | 0 | 0 | 0 | 0 | 0 | 0 | 0 | 0 | 0 |
| 438 | sbmA | 0 | 0 | 0 | 0 | 0 | 0 | 0 | 0 | 0 | 0 | 0 | 0 |
| 439 | sbmC | 0 | 0 | 0 | 0 | 0 | 0 | 0 | 0 | 0 | 0 | 0 | 0 |
| 440 | idnK | 0 | 0 | 0 | 0 | 0 | 0 | 0 | 0 | 0 | 0 | 0 | 0 |
| 441 | mltC | 0 | 0 | 0 | 0 | 0 | 0 | 0 | 0 | 0 | 0 | 1 | 0 |
| 442 | mltD | 0 | 0 | 0 | 0 | 0 | 0 | 0 | 0 | 0 | 0 | 0 | 0 |
| 443 | pppA | 0 | 0 | 0 | 0 | 0 | 0 | 0 | 0 | 0 | 0 | 0 | 0 |
| 444 | xthA | 0 | 0 | 0 | 0 | 0 | 0 | 0 | 0 | 0 | 0 | 0 | 0 |
| 445 | fsr | 0 | 0 | 0 | 0 | 0 | 0 | 0 | 0 | 0 | 0 | 0 | 0 |
| 446 | rhaT | 0 | 0 | 0 | 0 | 0 | 0 | 0 | 0 | 0 | 0 | 0 | 0 |
| 447 | aspA | 0 | 0 | 0 | 0 | 0 | 0 | 0 | 0 | 0 | 0 | 0 | 0 |
| 448 | rhaS | 0 | 0 | 0 | 0 | 0 | 0 | 0 | 0 | 0 | 0 | 0 | 0 |
| 449 | aspC | 0 | 0 | 0 | 0 | 0 | 0 | 0 | 0 | 0 | 0 | 0 | 0 |
| 450 | rob | 0 | 0 | 0 | 0 | 0 | 0 | 0 | 0 | 0 | 0 | 0 | 0 |
| 451 | grxA | 0 | 0 | 0 | 0 | 0 | 0 | 0 | 0 | 0 | 0 | 0 | 0 |
| 452 | rof | 357 | 38 | 233 | 29 | 2 | 0 | 369 | 2 | 101 | 96 | 121 | 0 |
| 453 | aspU | 0 | 0 | 0 | 0 | 0 | 0 | 0 | 0 | 0 | 1 | 0 | 0 |
| 454 | rhaB | 0 | 0 | 0 | 0 | 0 | 0 | 0 | 0 | 0 | 0 | 0 | 0 |
| 455 | yhjK | 0 | 0 | 0 | 0 | 0 | 0 | 0 | 0 | 0 | 0 | 0 | 0 |
| 456 | yhjJ | 0 | 0 | 0 | 0 | 0 | 0 | 0 | 0 | 0 | 0 | 0 | 0 |
| 457 | yjjV | 0 | 0 | 0 | 0 | 0 | 0 | 0 | 0 | 1 | 0 | 0 | 0 |
| 458 | yhjG | 0 | 0 | 0 | 0 | 0 | 0 | 0 | 0 | 0 | 0 | 0 | 0 |
| 459 | dsdC_dsdX | 0 | 0 | 0 | 1 | 0 | 0 | 0 | 0 | 0 | 0 | 0 | 0 |
| 460 | yhjC | 0 | 0 | 0 | 0 | 0 | 0 | 0 | 0 | 0 | 0 | 0 | 0 |
| 461 | yhjB | 0 | 0 | 0 | 0 | 0 | 0 | 0 | 0 | 0 | 0 | 0 | 0 |
| 462 | yjjL | 0 | 0 | 0 | 0 | 0 | 0 | 0 | 0 | 0 | 0 | 0 | 0 |
| 463 | yhjY | 0 | 0 | 0 | 0 | 0 | 0 | 0 | 0 | 0 | 0 | 0 | 0 |
| 464 | pldA | 0 | 0 | 0 | 0 | 0 | 0 | 0 | 0 | 0 | 0 | 0 | 0 |
| 465 | yjjI | 0 | 0 | 0 | 0 | 0 | 0 | 0 | 0 | 0 | 0 | 0 | 0 |
| 466 | yhjR | 0 | 0 | 0 | 0 | 0 | 0 | 0 | 0 | 0 | 0 | 0 | 0 |
| 467 | yfbJ | 1065 | 1 | 697 | 1 | 13 | 0 | 985 | 0 | 380 | 0 | 155 | 0 |
| 468 | yfbM | 0 | 0 | 0 | 0 | 0 | 0 | 1 | 0 | 0 | 0 | 0 | 0 |
| 469 | yfbE | 0 | 0 | 0 | 0 | 0 | 0 | 0 | 0 | 0 | 0 | 0 | 0 |
| 470 | apbE | 0 | 0 | 0 | 0 | 0 | 1 | 0 | 0 | 0 | 0 | 0 | 0 |
| 471 | yfbS | 0 | 0 | 0 | 0 | 0 | 0 | 0 | 0 | 0 | 0 | 0 | 0 |
| 472 | yfbR | 0 | 0 | 0 | 0 | 0 | 0 | 0 | 0 | 0 | 2 | 0 | 0 |
| 473 | yfbV | 0 | 0 | 0 | 0 | 0 | 0 | 0 | 0 | 0 | 0 | 0 | 0 |
| 474 | yfbU | 0 | 0 | 0 | 0 | 0 | 0 | 0 | 0 | 0 | 0 | 0 | 0 |
| 475 | glmU | 0 | 0 | 0 | 1 | 0 | 0 | 0 | 0 | 0 | 0 | 0 | 0 |
| 476 | eutB | 0 | 0 | 0 | 0 | 0 | 0 | 0 | 0 | 0 | 0 | 0 | 0 |
| 477 | cyaA_hemC | 0 | 0 | 0 | 0 | 0 | 0 | 0 | 0 | 1 | 0 | 0 | 0 |
| 478 | der | 0 | 0 | 0 | 0 | 0 | 0 | 0 | 0 | 0 | 0 | 0 | 0 |
| 479 | hdfR | 0 | 0 | 0 | 0 | 0 | 0 | 0 | 0 | 0 | 0 | 0 | 0 |
| 480 | hscC | 0 | 0 | 0 | 0 | 0 | 0 | 0 | 0 | 0 | 0 | 0 | 0 |
| 481 | acpP | 0 | 0 | 0 | 0 | 0 | 0 | 0 | 0 | 0 | 0 | 0 | 0 |
| 482 | seqA | 0 | 0 | 0 | 0 | 0 | 0 | 0 | 0 | 0 | 0 | 0 | 0 |
| 483 | nupG | 0 | 0 | 0 | 1 | 0 | 0 | 0 | 0 | 0 | 0 | 0 | 0 |
| 484 | pmbA | 0 | 0 | 0 | 0 | 0 | 0 | 0 | 0 | 0 | 0 | 0 | 0 |
| 485 | yodB | 0 | 0 | 0 | 0 | 0 | 0 | 0 | 0 | 0 | 0 | 0 | 0 |
| 486 | nlpC | 0 | 0 | 0 | 0 | 0 | 0 | 0 | 0 | 0 | 0 | 0 | 0 |
| 487 | nlpA | 0 | 0 | 0 | 0 | 0 | 0 | 0 | 0 | 1 | 0 | 0 | 0 |
| 488 | cysH | 506 | 5 | 391 | 7 | 336 | 6 | 507 | 7 | 139 | 26 | 203 | 5 |
| 489 | nlpD | 0 | 0 | 0 | 0 | 0 | 0 | 0 | 0 | 0 | 0 | 0 | 0 |
| 490 | cysJ | 0 | 0 | 0 | 0 | 0 | 0 | 0 | 0 | 0 | 0 | 0 | 0 |
| 491 | cysD | 0 | 0 | 0 | 0 | 0 | 0 | 0 | 0 | 0 | 0 | 0 | 0 |
| 492 | nlpI | 0 | 0 | 0 | 0 | 0 | 0 | 0 | 0 | 0 | 0 | 0 | 0 |
| 493 | cysB | 0 | 0 | 0 | 0 | 0 | 0 | 0 | 0 | 0 | 1 | 0 | 0 |
| 494 | uxuR | 0 | 0 | 0 | 0 | 0 | 1 | 0 | 0 | 1 | 0 | 0 | 0 |
| 495 | cysZ | 0 | 0 | 0 | 0 | 0 | 0 | 0 | 0 | 0 | 0 | 0 | 0 |
| 496 | cysT | 0 | 0 | 0 | 0 | 0 | 0 | 0 | 0 | 0 | 0 | 0 | 0 |
| 497 | cysW | 0 | 0 | 0 | 0 | 0 | 0 | 0 | 0 | 1 | 0 | 0 | 0 |
| 498 | cysQ | 522 | 0 | 365 | 3 | 0 | 0 | 397 | 0 | 252 | 0 | 470 | 0 |
| 499 | cysP | 0 | 0 | 0 | 0 | 0 | 0 | 0 | 0 | 0 | 0 | 0 | 0 |
| 500 | cysS | 0 | 0 | 0 | 0 | 0 | 0 | 0 | 0 | 0 | 0 | 0 | 0 |
| 501 | def | 0 | 0 | 0 | 0 | 0 | 0 | 0 | 0 | 0 | 0 | 0 | 0 |
| 502 | perR | 0 | 0 | 0 | 0 | 0 | 0 | 0 | 0 | 0 | 0 | 0 | 0 |
| 503 | ppiD | 0 | 0 | 0 | 0 | 0 | 1 | 0 | 0 | 0 | 0 | 0 | 0 |

Data Set S2 - MG1655 library 7 day experiment
Aligned Reads

| | | | | | | | | | | | | | |
|---|---|---|---|---|---|---|---|---|---|---|---|---|---|
| 504 | ppiC | 0 | 0 | 0 | 0 | 0 | 0 | 0 | 0 | 0 | 0 | 0 | 0 |
| 505 | ppiB | 0 | 0 | 0 | 0 | 0 | 0 | 0 | 0 | 0 | 0 | 0 | 0 |
| 506 | ppiA | 0 | 0 | 0 | 0 | 0 | 0 | 0 | 0 | 0 | 0 | 0 | 0 |
| 507 | ybgQ | 0 | 0 | 0 | 0 | 0 | 0 | 0 | 0 | 0 | 0 | 0 | 0 |
| 508 | ybgS | 0 | 0 | 0 | 0 | 0 | 0 | 0 | 0 | 0 | 0 | 0 | 0 |
| 509 | yifB | 0 | 0 | 0 | 0 | 0 | 0 | 0 | 0 | 0 | 0 | 0 | 0 |
| 510 | mscS | 0 | 0 | 0 | 0 | 0 | 0 | 1 | 0 | 0 | 0 | 0 | 0 |
| 511 | ykfG | 0 | 0 | 0 | 0 | 0 | 0 | 0 | 0 | 0 | 0 | 0 | 0 |
| 512 | ybgC | 0 | 0 | 1 | 0 | 0 | 0 | 0 | 0 | 0 | 0 | 0 | 0 |
| 513 | ybgD | 0 | 0 | 0 | 0 | 0 | 0 | 0 | 0 | 0 | 0 | 0 | 0 |
| 514 | mscL | 0 | 0 | 0 | 0 | 0 | 0 | 0 | 0 | 0 | 0 | 0 | 0 |
| 515 | ykfA | 0 | 0 | 0 | 0 | 0 | 0 | 0 | 0 | 0 | 0 | 0 | 0 |
| 516 | ybgH | 0 | 0 | 0 | 0 | 0 | 0 | 0 | 0 | 0 | 0 | 0 | 0 |
| 517 | ybgI | 0 | 0 | 0 | 0 | 0 | 0 | 0 | 0 | 0 | 0 | 0 | 0 |
| 518 | ykfJ | 0 | 0 | 0 | 0 | 0 | 0 | 0 | 0 | 0 | 0 | 0 | 0 |
| 519 | ykfI | 0 | 0 | 0 | 0 | 0 | 0 | 0 | 0 | 0 | 0 | 0 | 0 |
| 520 | yihN | 0 | 0 | 0 | 0 | 0 | 0 | 0 | 0 | 0 | 0 | 0 | 0 |
| 521 | bolA | 0 | 0 | 0 | 0 | 0 | 0 | 0 | 0 | 0 | 0 | 0 | 0 |
| 522 | yihL | 0 | 1 | 0 | 0 | 0 | 0 | 0 | 0 | 0 | 0 | 0 | 0 |
| 523 | ygfF | 0 | 0 | 1 | 0 | 0 | 0 | 0 | 0 | 0 | 0 | 0 | 0 |
| 524 | yihD | 133 | 0 | 103 | 0 | 2 | 0 | 129 | 0 | 109 | 0 | 216 | 0 |
| 525 | ygfI | 0 | 0 | 0 | 0 | 0 | 0 | 0 | 0 | 0 | 0 | 0 | 1 |
| 526 | mdtG | 0 | 0 | 0 | 0 | 0 | 0 | 0 | 0 | 0 | 0 | 0 | 0 |
| 527 | yihX | 0 | 0 | 0 | 0 | 0 | 0 | 0 | 0 | 0 | 0 | 0 | 0 |
| 528 | ygfU | 0 | 0 | 0 | 0 | 0 | 0 | 0 | 0 | 0 | 0 | 0 | 0 |
| 529 | mdtK | 0 | 0 | 0 | 0 | 0 | 0 | 0 | 0 | 0 | 0 | 0 | 0 |
| 530 | focA | 0 | 0 | 0 | 0 | 0 | 0 | 0 | 0 | 0 | 0 | 0 | 0 |
| 531 | yihU | 0 | 0 | 0 | 0 | 0 | 0 | 0 | 0 | 0 | 0 | 0 | 0 |
| 532 | mdtO | 696 | 1 | 442 | 1 | 3 | 0 | 529 | 0 | 248 | 0 | 276 | 0 |
| 533 | mdtN | 0 | 0 | 1 | 0 | 0 | 1 | 0 | 0 | 0 | 0 | 0 | 0 |
| 534 | hycB | 0 | 0 | 0 | 0 | 0 | 0 | 0 | 0 | 0 | 1 | 0 | 0 |
| 535 | flk | 0 | 0 | 0 | 0 | 0 | 1 | 0 | 0 | 0 | 0 | 0 | 0 |
| 536 | spy | 0 | 0 | 0 | 0 | 0 | 0 | 0 | 0 | 0 | 0 | 0 | 0 |
| 537 | pinH | 0 | 0 | 0 | 0 | 0 | 0 | 0 | 0 | 0 | 0 | 0 | 0 |
| 538 | purT | 0 | 0 | 0 | 0 | 0 | 0 | 0 | 0 | 0 | 0 | 0 | 0 |
| 539 | mpaA | 0 | 0 | 0 | 0 | 0 | 0 | 0 | 0 | 0 | 0 | 0 | 0 |
| 540 | spr | 0 | 0 | 0 | 0 | 0 | 0 | 0 | 0 | 0 | 0 | 0 | 0 |
| 541 | dxr | 0 | 0 | 0 | 0 | 0 | 0 | 0 | 0 | 0 | 0 | 0 | 0 |
| 542 | tilS | 0 | 0 | 0 | 0 | 0 | 0 | 0 | 0 | 0 | 0 | 0 | 0 |
| 543 | trxC | 0 | 0 | 0 | 0 | 0 | 0 | 0 | 0 | 0 | 0 | 0 | 0 |
| 544 | trxA | 0 | 0 | 0 | 0 | 0 | 0 | 0 | 0 | 0 | 0 | 0 | 0 |
| 545 | pheM | 0 | 0 | 0 | 0 | 0 | 0 | 0 | 1 | 0 | 0 | 0 | 0 |
| 546 | pinR | 0 | 0 | 0 | 0 | 0 | 0 | 0 | 0 | 0 | 0 | 0 | 0 |
| 547 | ylaB | 0 | 0 | 0 | 0 | 0 | 1 | 0 | 0 | 0 | 0 | 0 | 0 |
| 548 | ylaC | 0 | 0 | 0 | 0 | 0 | 0 | 0 | 0 | 0 | 0 | 0 | 0 |
| 549 | purM | 0 | 0 | 0 | 0 | 0 | 0 | 0 | 0 | 0 | 0 | 0 | 0 |
| 550 | ydcL | 0 | 0 | 0 | 0 | 0 | 0 | 0 | 0 | 0 | 0 | 0 | 0 |
| 551 | ydcM | 1 | 1 | 0 | 1 | 0 | 0 | 0 | 1 | 0 | 1 | 0 | 0 |
| 552 | ydcJ | 0 | 0 | 0 | 0 | 0 | 0 | 0 | 0 | 0 | 0 | 0 | 0 |
| 553 | ydcK | 0 | 0 | 0 | 0 | 0 | 0 | 0 | 0 | 0 | 0 | 0 | 0 |
| 554 | ydcI | 0 | 0 | 0 | 0 | 0 | 0 | 0 | 0 | 0 | 0 | 0 | 0 |
| 555 | ydcF | 4865 | 9 | 3214 | 12 | 38 | 15 | 4294 | 7733 | 1409 | 28 | 3536 | 72 |
| 556 | ydcG | 0 | 0 | 0 | 0 | 0 | 0 | 0 | 0 | 0 | 0 | 0 | 0 |
| 557 | ydcA | 0 | 0 | 0 | 0 | 0 | 0 | 0 | 0 | 0 | 0 | 0 | 0 |
| 558 | fucR | 481 | 1 | 467 | 172 | 73 | 932 | 315 | 5214 | 278 | 19 | 1 | 0 |
| 559 | ydcX | 0 | 0 | 1 | 0 | 0 | 0 | 0 | 0 | 0 | 0 | 0 | 0 |
| 560 | fpr | 0 | 0 | 0 | 0 | 0 | 0 | 0 | 0 | 0 | 0 | 0 | 0 |
| 561 | chiA | 0 | 0 | 0 | 0 | 0 | 0 | 0 | 0 | 0 | 0 | 0 | 0 |
| 562 | ydcR | 1 | 0 | 0 | 0 | 0 | 0 | 0 | 0 | 0 | 0 | 0 | 0 |
| 563 | ydcS | 0 | 0 | 0 | 0 | 0 | 0 | 0 | 0 | 0 | 0 | 0 | 0 |
| 564 | map | 0 | 0 | 1 | 0 | 0 | 0 | 0 | 0 | 0 | 0 | 0 | 0 |
| 565 | rrlG | 514 | 0 | 303 | 0 | 12 | 0 | 313 | 2 | 161 | 2 | 65 | 0 |
| 566 | cdaR | 0 | 0 | 0 | 0 | 0 | 0 | 0 | 0 | 0 | 0 | 0 | 0 |
| 567 | cpdA | 0 | 0 | 0 | 0 | 0 | 0 | 0 | 0 | 0 | 0 | 0 | 0 |
| 568 | cpdB | 2 | 0 | 0 | 0 | 0 | 0 | 0 | 0 | 0 | 0 | 0 | 0 |
| 569 | talA | 0 | 0 | 0 | 0 | 0 | 0 | 0 | 0 | 0 | 0 | 0 | 0 |
| 570 | talB | 0 | 0 | 0 | 0 | 0 | 0 | 0 | 0 | 0 | 0 | 0 | 0 |
| 571 | yncA | 0 | 0 | 0 | 0 | 0 | 0 | 0 | 0 | 0 | 0 | 0 | 0 |
| 572 | yncB | 0 | 0 | 0 | 0 | 0 | 0 | 0 | 0 | 0 | 0 | 0 | 0 |
| 573 | yncC | 0 | 0 | 0 | 0 | 0 | 0 | 0 | 0 | 0 | 0 | 0 | 0 |
| 574 | yncD | 0 | 0 | 0 | 0 | 0 | 0 | 0 | 0 | 0 | 0 | 0 | 0 |
| 575 | yncE | 0 | 0 | 0 | 0 | 0 | 0 | 0 | 0 | 0 | 0 | 0 | 0 |
| 576 | uvrY | 0 | 0 | 0 | 0 | 0 | 0 | 0 | 0 | 0 | 0 | 0 | 0 |
| 577 | yncG | 0 | 0 | 0 | 0 | 0 | 0 | 0 | 0 | 0 | 0 | 0 | 0 |
| 578 | yncJ | 0 | 0 | 0 | 0 | 0 | 0 | 0 | 0 | 0 | 0 | 0 | 0 |
| 579 | pncB | 0 | 0 | 0 | 0 | 0 | 0 | 0 | 0 | 0 | 0 | 0 | 0 |
| 580 | yfaY | 0 | 0 | 0 | 1 | 0 | 0 | 0 | 0 | 0 | 0 | 0 | 0 |
| 581 | torT | 0 | 0 | 0 | 0 | 0 | 0 | 0 | 0 | 0 | 0 | 0 | 0 |
| 582 | yfaD | 2 | 0 | 0 | 0 | 0 | 0 | 0 | 0 | 0 | 0 | 0 | 0 |
| 583 | torS | 0 | 0 | 0 | 0 | 0 | 0 | 0 | 0 | 0 | 0 | 0 | 0 |
| 584 | torR | 0 | 0 | 0 | 0 | 0 | 0 | 0 | 0 | 0 | 0 | 0 | 0 |
| 585 | yfaL | 0 | 0 | 0 | 0 | 0 | 0 | 0 | 0 | 0 | 0 | 0 | 0 |
| 586 | ygiL | 0 | 0 | 0 | 0 | 0 | 0 | 0 | 0 | 0 | 0 | 0 | 0 |
| 587 | torY | 0 | 0 | 0 | 0 | 0 | 0 | 0 | 0 | 0 | 1 | 0 | 0 |

| | | | | | | | | | | | | | |
|---|---|---|---|---|---|---|---|---|---|---|---|---|---|
| 588 | uvrA | 0 | 0 | 0 | 0 | 0 | 0 | 0 | 0 | 0 | 0 | 0 | 0 |
| 589 | ycaO | 10 | 0 | 5 | 0 | 0 | 0 | 17 | 0 | 4 | 0 | 0 | 0 |
| 590 | ycaN | 0 | 0 | 0 | 0 | 0 | 0 | 0 | 0 | 0 | 0 | 0 | 0 |
| 591 | ycaL | 0 | 0 | 0 | 0 | 0 | 0 | 0 | 0 | 0 | 0 | 0 | 0 |
| 592 | ycaK | 0 | 0 | 0 | 0 | 0 | 0 | 0 | 0 | 0 | 0 | 0 | 0 |
| 593 | cutA | 0 | 0 | 0 | 0 | 0 | 0 | 0 | 0 | 0 | 0 | 0 | 0 |
| 594 | ycaI | 0 | 0 | 0 | 0 | 0 | 0 | 0 | 0 | 0 | 0 | 0 | 0 |
| 595 | ycaD | 0 | 0 | 0 | 0 | 0 | 0 | 0 | 0 | 0 | 0 | 0 | 0 |
| 596 | ycaC | 0 | 0 | 0 | 0 | 0 | 0 | 0 | 0 | 0 | 0 | 0 | 0 |
| 597 | cybC | 0 | 0 | 2 | 0 | 0 | 0 | 0 | 0 | 0 | 0 | 0 | 0 |
| 598 | tktB | 755 | 2 | 387 | 1 | 7 | 1 | 342 | 0 | 422 | 1 | 204 | 0 |
| 599 | ssb | 1 | 0 | 0 | 0 | 0 | 0 | 0 | 0 | 0 | 0 | 0 | 0 |
| 600 | ycaP | 0 | 0 | 0 | 0 | 0 | 0 | 0 | 0 | 0 | 0 | 0 | 0 |
| 601 | b0540 | 0 | 0 | 0 | 0 | 0 | 0 | 0 | 0 | 0 | 0 | 0 | 0 |
| 602 | yqgD | 139 | 0 | 122 | 0 | 1 | 1 | 132 | 0 | 61 | 0 | 0 | 0 |
| 603 | lexA | 0 | 0 | 0 | 0 | 0 | 0 | 0 | 0 | 0 | 0 | 0 | 0 |
| 604 | cobU | 0 | 0 | 0 | 0 | 0 | 0 | 0 | 0 | 0 | 1 | 0 | 0 |
| 605 | ybgE | 486 | 2 | 308 | 1 | 2 | 0 | 377 | 1 | 149 | 0 | 484 | 1 |
| 606 | ymcE | 0 | 0 | 0 | 0 | 0 | 0 | 0 | 0 | 0 | 0 | 0 | 0 |
| 607 | ymcC | 0 | 0 | 0 | 0 | 0 | 0 | 0 | 0 | 0 | 0 | 0 | 0 |
| 608 | b0309 | 0 | 0 | 0 | 0 | 0 | 0 | 0 | 0 | 0 | 0 | 0 | 0 |
| 609 | ilvY | 0 | 0 | 0 | 0 | 0 | 0 | 0 | 0 | 0 | 0 | 0 | 0 |
| 610 | cytR | 0 | 0 | 0 | 0 | 0 | 0 | 0 | 0 | 0 | 0 | 0 | 0 |
| 611 | nupC | 0 | 0 | 0 | 0 | 0 | 0 | 0 | 0 | 0 | 0 | 0 | 0 |
| 612 | hcp | 0 | 0 | 0 | 0 | 0 | 1 | 0 | 0 | 0 | 0 | 0 | 0 |
| 613 | ytfQ | 0 | 0 | 0 | 0 | 0 | 0 | 0 | 0 | 0 | 0 | 0 | 0 |
| 614 | ytfR | 0 | 0 | 0 | 0 | 0 | 0 | 0 | 0 | 0 | 0 | 0 | 0 |
| 615 | yeaM | 0 | 0 | 0 | 0 | 0 | 0 | 0 | 0 | 0 | 0 | 0 | 0 |
| 616 | yeaL | 0 | 0 | 0 | 0 | 0 | 0 | 0 | 0 | 0 | 0 | 0 | 0 |
| 617 | rhtB | 0 | 0 | 0 | 0 | 0 | 0 | 0 | 0 | 0 | 0 | 0 | 0 |
| 618 | yeaH | 0 | 0 | 0 | 0 | 0 | 0 | 0 | 0 | 0 | 0 | 0 | 0 |
| 619 | yeaU | 405 | 1 | 263 | 4 | 6 | 0 | 311 | 0 | 151 | 0 | 2 | 0 |
| 620 | yeaT | 1 | 0 | 0 | 0 | 0 | 0 | 0 | 0 | 0 | 0 | 0 | 0 |
| 621 | yeaV | 0 | 0 | 0 | 0 | 0 | 0 | 0 | 0 | 0 | 0 | 0 | 0 |
| 622 | yeaP | 0 | 0 | 0 | 0 | 0 | 0 | 0 | 0 | 0 | 0 | 0 | 0 |
| 623 | yeaS | 0 | 0 | 0 | 0 | 0 | 0 | 1 | 1 | 0 | 0 | 0 | 0 |
| 624 | ytfB | 0 | 0 | 0 | 1 | 0 | 0 | 0 | 0 | 1 | 0 | 0 | 0 |
| 625 | ytfM | 0 | 0 | 0 | 0 | 0 | 0 | 0 | 0 | 0 | 0 | 0 | 0 |
| 626 | ytfL | 0 | 0 | 0 | 0 | 0 | 0 | 0 | 0 | 0 | 0 | 0 | 0 |
| 627 | nupX | 610 | 2 | 475 | 1 | 8 | 0 | 355 | 0 | 305 | 0 | 680 | 2 |
| 628 | ytfI | 0 | 0 | 0 | 0 | 0 | 0 | 0 | 0 | 0 | 0 | 0 | 0 |
| 629 | yeaX | 0 | 0 | 0 | 0 | 0 | 0 | 0 | 0 | 0 | 0 | 0 | 0 |
| 630 | blc | 0 | 0 | 0 | 0 | 0 | 0 | 0 | 0 | 0 | 0 | 0 | 0 |
| 631 | yeaZ | 0 | 0 | 0 | 0 | 0 | 0 | 0 | 0 | 0 | 0 | 0 | 0 |
| 632 | argZ_argQ | 338 | 4 | 242 | 0 | 0 | 0 | 195 | 1 | 159 | 1 | 432 | 1 |
| 633 | yjiY | 0 | 0 | 0 | 0 | 0 | 0 | 0 | 0 | 0 | 0 | 0 | 0 |
| 634 | ilvL | 0 | 0 | 0 | 0 | 0 | 0 | 0 | 0 | 0 | 0 | 0 | 0 |
| 635 | argP | 0 | 0 | 1 | 0 | 0 | 0 | 0 | 0 | 0 | 0 | 0 | 0 |
| 636 | intB | 0 | 0 | 0 | 0 | 1 | 0 | 0 | 0 | 0 | 0 | 0 | 0 |
| 637 | argU | 0 | 0 | 0 | 0 | 0 | 0 | 0 | 0 | 0 | 0 | 0 | 0 |
| 638 | yjiT | 0 | 0 | 0 | 1 | 0 | 0 | 0 | 0 | 0 | 0 | 0 | 0 |
| 639 | argW | 0 | 0 | 0 | 0 | 0 | 0 | 0 | 0 | 0 | 0 | 0 | 0 |
| 640 | yjiJ | 0 | 0 | 0 | 0 | 0 | 0 | 0 | 0 | 0 | 0 | 0 | 0 |
| 641 | argI | 0 | 0 | 0 | 0 | 0 | 0 | 0 | 0 | 0 | 0 | 0 | 0 |
| 642 | yjiH | 0 | 0 | 0 | 0 | 0 | 0 | 0 | 0 | 0 | 0 | 0 | 0 |
| 643 | yjiM | 0 | 0 | 0 | 0 | 0 | 0 | 0 | 0 | 0 | 0 | 0 | 0 |
| 644 | argA_amiC | 260 | 1 | 199 | 0 | 3 | 0 | 334 | 0 | 84 | 0 | 148 | 0 |
| 645 | argC | 0 | 0 | 0 | 1 | 0 | 0 | 0 | 0 | 0 | 0 | 0 | 0 |
| 646 | argD | 641 | 4 | 487 | 0 | 4 | 0 | 669 | 0 | 274 | 0 | 0 | 0 |
| 647 | argE | 0 | 0 | 0 | 0 | 0 | 0 | 0 | 0 | 0 | 0 | 0 | 0 |
| 648 | argF | 0 | 0 | 0 | 0 | 0 | 0 | 0 | 0 | 0 | 0 | 0 | 0 |
| 649 | argG | 0 | 0 | 0 | 0 | 0 | 0 | 0 | 1 | 0 | 0 | 0 | 0 |
| 650 | yggM | 0 | 0 | 0 | 0 | 0 | 0 | 0 | 0 | 0 | 0 | 0 | 0 |
| 651 | tyrS | 0 | 0 | 0 | 0 | 0 | 0 | 0 | 0 | 0 | 0 | 0 | 0 |
| 652 | yggH | 0 | 0 | 0 | 0 | 0 | 0 | 0 | 0 | 0 | 0 | 0 | 0 |
| 653 | htrA | 0 | 0 | 0 | 0 | 0 | 0 | 0 | 0 | 0 | 0 | 0 | 0 |
| 654 | iaaA | 0 | 0 | 0 | 0 | 0 | 0 | 0 | 0 | 0 | 0 | 0 | 0 |
| 655 | yggD | 0 | 0 | 0 | 0 | 0 | 0 | 0 | 0 | 0 | 0 | 0 | 0 |
| 656 | citT | 0 | 1 | 0 | 1 | 0 | 0 | 0 | 0 | 0 | 0 | 0 | 0 |
| 657 | ysgA | 0 | 0 | 0 | 0 | 0 | 0 | 0 | 0 | 0 | 0 | 0 | 0 |
| 658 | sstT | 0 | 0 | 0 | 0 | 0 | 0 | 0 | 0 | 0 | 0 | 0 | 0 |
| 659 | yiiF | 0 | 0 | 0 | 0 | 0 | 0 | 0 | 0 | 0 | 0 | 0 | 0 |
| 660 | yiiQ | 0 | 0 | 0 | 0 | 0 | 0 | 0 | 0 | 0 | 0 | 0 | 0 |
| 661 | yiiP | 2106 | 4 | 1708 | 3 | 298 | 3 | 1839 | 1 | 1046 | 0 | 1457 | 0 |
| 662 | citC | 0 | 0 | 0 | 0 | 0 | 0 | 0 | 0 | 0 | 0 | 0 | 0 |
| 663 | yiiU | 0 | 0 | 0 | 1 | 0 | 0 | 0 | 0 | 0 | 0 | 0 | 0 |
| 664 | tyrV | 0 | 0 | 0 | 0 | 0 | 0 | 1 | 0 | 0 | 0 | 0 | 0 |
| 665 | degQ | 0 | 0 | 1 | 0 | 0 | 0 | 1 | 0 | 0 | 1 | 0 | 0 |
| 666 | yphH | 0 | 0 | 0 | 0 | 0 | 0 | 0 | 0 | 0 | 0 | 0 | 0 |
| 667 | hybO | 0 | 0 | 0 | 0 | 0 | 0 | 0 | 0 | 0 | 0 | 0 | 0 |
| 668 | yphF | 0 | 0 | 0 | 0 | 0 | 0 | 0 | 0 | 0 | 0 | 0 | 0 |
| 669 | accB | 0 | 0 | 1 | 0 | 0 | 1 | 0 | 1 | 0 | 0 | 0 | 0 |
| 670 | iap | 0 | 0 | 0 | 0 | 0 | 0 | 0 | 0 | 0 | 0 | 0 | 0 |
| 671 | smpA | 0 | 0 | 0 | 0 | 0 | 0 | 0 | 0 | 0 | 0 | 0 | 0 |

| | | | | | | | | | | | | | |
|---|---|---|---|---|---|---|---|---|---|---|---|---|---|
| 672 | smpB | 0 | 0 | 0 | 0 | 0 | 0 | 0 | 0 | 0 | 0 | 0 | 0 |
| 673 | serU | 0 | 0 | 0 | 0 | 0 | 0 | 0 | 0 | 0 | 0 | 0 | 0 |
| 674 | pnp | 0 | 0 | 0 | 0 | 0 | 0 | 0 | 0 | 0 | 0 | 0 | 0 |
| 675 | serB | 0 | 0 | 0 | 0 | 0 | 0 | 0 | 0 | 0 | 0 | 0 | 0 |
| 676 | serC | 0 | 0 | 0 | 0 | 1 | 0 | 1 | 0 | 0 | 0 | 0 | 0 |
| 677 | serA | 0 | 0 | 0 | 0 | 0 | 0 | 0 | 0 | 0 | 0 | 0 | 0 |
| 678 | alaS | 0 | 0 | 0 | 0 | 0 | 0 | 0 | 0 | 0 | 0 | 0 | 0 |
| 679 | yqhH | 0 | 0 | 0 | 0 | 0 | 0 | 0 | 0 | 0 | 0 | 0 | 0 |
| 680 | rrmA | 0 | 0 | 0 | 0 | 0 | 0 | 0 | 0 | 0 | 0 | 0 | 0 |
| 681 | aroL | 0 | 1 | 1 | 0 | 0 | 0 | 0 | 0 | 0 | 0 | 0 | 0 |
| 682 | yjeB | 0 | 0 | 0 | 0 | 0 | 0 | 0 | 0 | 0 | 0 | 0 | 0 |
| 683 | sulA | 0 | 0 | 0 | 0 | 0 | 0 | 0 | 0 | 0 | 0 | 0 | 0 |
| 684 | gyrB | 0 | 0 | 1 | 0 | 0 | 0 | 0 | 1 | 0 | 0 | 0 | 0 |
| 685 | yhiR | 0 | 0 | 0 | 0 | 0 | 0 | 0 | 0 | 0 | 0 | 1 | 0 |
| 686 | sohB | 0 | 0 | 0 | 0 | 0 | 0 | 0 | 0 | 0 | 0 | 0 | 0 |
| 687 | sohA | 0 | 0 | 0 | 0 | 0 | 0 | 0 | 0 | 0 | 0 | 0 | 0 |
| 688 | brnQ | 0 | 0 | 0 | 0 | 0 | 0 | 0 | 0 | 1 | 0 | 0 | 0 |
| 689 | betT | 0 | 0 | 0 | 0 | 0 | 0 | 0 | 0 | 0 | 0 | 0 | 0 |
| 690 | mrp | 0 | 0 | 0 | 0 | 0 | 0 | 0 | 0 | 0 | 0 | 0 | 0 |
| 691 | mrr | 0 | 0 | 0 | 0 | 0 | 0 | 0 | 0 | 0 | 0 | 0 | 0 |
| 692 | ykgC | 0 | 0 | 0 | 0 | 0 | 0 | 0 | 0 | 0 | 0 | 0 | 0 |
| 693 | ykgE | 0 | 0 | 0 | 0 | 0 | 0 | 0 | 0 | 0 | 0 | 0 | 0 |
| 694 | ybfF | 0 | 0 | 0 | 0 | 0 | 0 | 0 | 0 | 0 | 0 | 0 | 0 |
| 695 | ybfE | 0 | 1 | 0 | 0 | 0 | 0 | 0 | 0 | 0 | 0 | 0 | 0 |
| 696 | ykgF | 0 | 0 | 0 | 0 | 0 | 0 | 0 | 0 | 0 | 0 | 0 | 0 |
| 697 | ykgI | 0 | 0 | 0 | 0 | 0 | 0 | 0 | 0 | 0 | 0 | 0 | 0 |
| 698 | ykgH | 0 | 0 | 0 | 0 | 0 | 0 | 0 | 0 | 0 | 0 | 0 | 0 |
| 699 | ykgJ | 0 | 0 | 0 | 0 | 0 | 0 | 0 | 0 | 1 | 0 | 0 | 0 |
| 700 | ykgM | 0 | 0 | 0 | 0 | 0 | 0 | 0 | 0 | 0 | 0 | 0 | 0 |
| 701 | grxC | 0 | 0 | 0 | 0 | 0 | 0 | 1 | 0 | 0 | 0 | 0 | 0 |
| 702 | grxB | 680 | 0 | 418 | 5 | 8 | 1 | 704 | 1 | 362 | 0 | 832 | 0 |
| 703 | dicB | 0 | 0 | 0 | 0 | 0 | 0 | 0 | 0 | 0 | 0 | 0 | 0 |
| 704 | elaD | 0 | 0 | 0 | 1 | 0 | 0 | 0 | 0 | 0 | 0 | 0 | 0 |
| 705 | elaA | 0 | 0 | 0 | 0 | 0 | 0 | 0 | 0 | 0 | 0 | 0 | 0 |
| 706 | elaB | 0 | 0 | 0 | 0 | 0 | 0 | 0 | 0 | 0 | 0 | 0 | 0 |
| 707 | hcaT | 0 | 0 | 0 | 0 | 0 | 0 | 0 | 0 | 0 | 0 | 0 | 0 |
| 708 | osmC | 0 | 0 | 0 | 0 | 0 | 0 | 0 | 0 | 0 | 0 | 0 | 0 |
| 709 | osmB | 0 | 0 | 0 | 0 | 0 | 0 | 0 | 1 | 0 | 0 | 0 | 0 |
| 710 | osmE | 0 | 0 | 0 | 0 | 0 | 0 | 0 | 0 | 0 | 0 | 0 | 0 |
| 711 | hcaR | 0 | 0 | 0 | 0 | 0 | 0 | 0 | 0 | 0 | 0 | 0 | 0 |
| 712 | leuA | 0 | 0 | 0 | 0 | 0 | 0 | 0 | 0 | 0 | 0 | 0 | 0 |
| 713 | asnV | 0 | 0 | 0 | 0 | 0 | 0 | 0 | 0 | 0 | 0 | 0 | 0 |
| 714 | asnS | 0 | 0 | 0 | 0 | 0 | 0 | 0 | 0 | 0 | 0 | 0 | 0 |
| 715 | leuS | 0 | 0 | 0 | 0 | 0 | 0 | 0 | 0 | 0 | 0 | 0 | 0 |
| 716 | asnC | 0 | 0 | 0 | 0 | 0 | 0 | 0 | 0 | 0 | 0 | 0 | 0 |
| 717 | asnA | 0 | 0 | 0 | 0 | 0 | 0 | 0 | 0 | 0 | 0 | 0 | 0 |
| 718 | ydbH | 0 | 0 | 0 | 0 | 0 | 0 | 0 | 0 | 0 | 0 | 0 | 0 |
| 719 | ydbC | 0 | 0 | 0 | 0 | 0 | 0 | 0 | 0 | 0 | 0 | 1 | 0 |
| 720 | ydbD | 0 | 0 | 0 | 0 | 0 | 0 | 0 | 0 | 0 | 0 | 0 | 0 |
| 721 | dksA | 0 | 0 | 0 | 0 | 0 | 0 | 0 | 0 | 0 | 0 | 0 | 0 |
| 722 | b1228 | 0 | 0 | 0 | 0 | 0 | 0 | 0 | 0 | 0 | 0 | 0 | 0 |
| 723 | hisL | 0 | 0 | 1 | 0 | 0 | 0 | 0 | 0 | 0 | 0 | 0 | 0 |
| 724 | hisJ | 0 | 0 | 0 | 0 | 0 | 0 | 0 | 0 | 0 | 1 | 0 | 0 |
| 725 | rmf | 0 | 0 | 0 | 0 | 0 | 0 | 0 | 0 | 0 | 0 | 0 | 0 |
| 726 | hisQ | 0 | 0 | 0 | 0 | 0 | 0 | 0 | 0 | 0 | 0 | 0 | 0 |
| 727 | hisS | 0 | 0 | 0 | 0 | 0 | 0 | 0 | 0 | 0 | 0 | 0 | 0 |
| 728 | ypdA | 0 | 0 | 0 | 0 | 0 | 0 | 0 | 0 | 0 | 0 | 0 | 0 |
| 729 | fadL | 0 | 0 | 0 | 0 | 0 | 0 | 0 | 0 | 0 | 0 | 0 | 0 |
| 730 | ynbA | 660 | 4 | 431 | 1 | 0 | 1 | 386 | 2 | 234 | 0 | 847 | 0 |
| 731 | manX | 0 | 0 | 0 | 0 | 0 | 0 | 0 | 0 | 0 | 0 | 0 | 0 |
| 732 | manY | 0 | 0 | 0 | 0 | 0 | 0 | 0 | 0 | 0 | 0 | 0 | 0 |
| 733 | yajR | 0 | 0 | 1 | 0 | 0 | 0 | 0 | 0 | 1 | 0 | 0 | 0 |
| 734 | yibK | 0 | 0 | 0 | 0 | 0 | 0 | 0 | 0 | 0 | 0 | 0 | 0 |
| 735 | yajO | 0 | 0 | 0 | 0 | 0 | 0 | 0 | 0 | 0 | 0 | 0 | 0 |
| 736 | yajB | 0 | 0 | 0 | 0 | 0 | 0 | 0 | 0 | 0 | 0 | 0 | 0 |
| 737 | yajC | 2 | 0 | 0 | 0 | 0 | 0 | 2 | 0 | 1 | 0 | 0 | 0 |
| 738 | b0374 | 0 | 0 | 0 | 0 | 0 | 0 | 0 | 0 | 0 | 0 | 0 | 0 |
| 739 | yajG | 0 | 0 | 0 | 0 | 0 | 0 | 0 | 0 | 0 | 0 | 0 | 0 |
| 740 | ycbK | 0 | 0 | 0 | 0 | 0 | 0 | 0 | 0 | 0 | 0 | 0 | 0 |
| 741 | ycbB | 0 | 0 | 0 | 0 | 0 | 0 | 0 | 0 | 0 | 0 | 0 | 0 |
| 742 | ycbC | 0 | 0 | 0 | 0 | 0 | 0 | 0 | 0 | 0 | 0 | 0 | 0 |
| 743 | lpxC | 0 | 0 | 0 | 0 | 0 | 0 | 0 | 0 | 0 | 0 | 0 | 0 |
| 744 | ycbZ | 1 | 0 | 1 | 0 | 0 | 0 | 0 | 0 | 0 | 0 | 0 | 0 |
| 745 | csiE | 0 | 0 | 0 | 1 | 0 | 0 | 0 | 0 | 0 | 0 | 0 | 0 |
| 746 | csiD | 0 | 0 | 0 | 0 | 0 | 0 | 0 | 0 | 0 | 0 | 0 | 0 |
| 747 | ycbR | 0 | 0 | 0 | 0 | 0 | 0 | 0 | 0 | 0 | 0 | 0 | 0 |
| 748 | ycbW | 0 | 0 | 0 | 0 | 0 | 0 | 0 | 0 | 0 | 0 | 0 | 0 |
| 749 | glcB | 0 | 0 | 0 | 0 | 0 | 0 | 0 | 0 | 0 | 1 | 0 | 0 |
| 750 | glcC | 0 | 0 | 0 | 0 | 0 | 0 | 0 | 0 | 0 | 0 | 0 | 0 |
| 751 | artP | 0 | 0 | 0 | 0 | 0 | 0 | 0 | 0 | 0 | 0 | 0 | 0 |
| 752 | marC_marR | 0 | 0 | 0 | 0 | 0 | 0 | 0 | 0 | 0 | 0 | 0 | 0 |
| 753 | rluE | 0 | 0 | 0 | 0 | 0 | 0 | 0 | 0 | 0 | 0 | 0 | 0 |
| 754 | rluD | 0 | 0 | 0 | 0 | 0 | 0 | 0 | 0 | 0 | 0 | 0 | 0 |
| 755 | rluF | 247 | 1 | 196 | 1 | 1 | 0 | 172 | 0 | 107 | 0 | 0 | 0 |

Data Set S2 - MG1655 library 7 day experiment
Aligned Reads

| | | | | | | | | | | | | | |
|---|---|---|---|---|---|---|---|---|---|---|---|---|---|
| 756 | rluC | 0 | 0 | 0 | 0 | 0 | 0 | 0 | 0 | 0 | 0 | 0 | 0 |
| 757 | rluB | 0 | 0 | 0 | 0 | 0 | 0 | 0 | 0 | 0 | 0 | 0 | 0 |
| 758 | b0373 | 0 | 0 | 0 | 0 | 0 | 0 | 0 | 0 | 0 | 0 | 0 | 0 |
| 759 | helD | 0 | 0 | 0 | 0 | 0 | 0 | 0 | 0 | 0 | 0 | 0 | 0 |
| 760 | yqiC | 0 | 0 | 0 | 0 | 0 | 0 | 0 | 0 | 0 | 0 | 0 | 0 |
| 761 | polB | 0 | 0 | 0 | 0 | 0 | 0 | 0 | 0 | 0 | 0 | 0 | 0 |
| 762 | polA | 0 | 0 | 0 | 0 | 0 | 0 | 0 | 1 | 0 | 0 | 0 | 0 |
| 763 | yqiJ | 0 | 0 | 0 | 0 | 0 | 0 | 0 | 0 | 0 | 0 | 0 | 0 |
| 764 | ecpD | 0 | 0 | 0 | 0 | 0 | 0 | 0 | 0 | 0 | 0 | 0 | 0 |
| 765 | yjhF | 0 | 0 | 0 | 0 | 0 | 0 | 0 | 0 | 0 | 0 | 0 | 0 |
| 766 | yjhI | 0 | 0 | 0 | 0 | 0 | 0 | 0 | 0 | 0 | 0 | 0 | 0 |
| 767 | yjhU | 0 | 0 | 0 | 0 | 0 | 0 | 0 | 0 | 0 | 0 | 0 | 0 |
| 768 | yjhS | 0 | 0 | 0 | 0 | 0 | 0 | 0 | 0 | 0 | 0 | 0 | 0 |
| 769 | hda | 0 | 0 | 0 | 0 | 0 | 0 | 0 | 0 | 0 | 0 | 0 | 0 |
| 770 | pepN | 0 | 0 | 0 | 0 | 0 | 0 | 0 | 0 | 0 | 0 | 0 | 0 |
| 771 | pepB | 0 | 0 | 0 | 0 | 0 | 0 | 0 | 0 | 0 | 0 | 0 | 0 |
| 772 | pepE | 0 | 0 | 0 | 0 | 0 | 0 | 0 | 0 | 0 | 0 | 0 | 0 |
| 773 | pepD | 0 | 0 | 0 | 0 | 0 | 0 | 0 | 0 | 0 | 0 | 0 | 0 |
| 774 | mqo | 0 | 1 | 0 | 0 | 0 | 0 | 0 | 0 | 0 | 0 | 0 | 1 |
| 775 | pepQ | 11 | 1 | 6 | 2 | 0 | 0 | 9 | 4 | 1 | 0 | 0 | 0 |
| 776 | pepT | 0 | 0 | 0 | 0 | 0 | 0 | 0 | 0 | 0 | 0 | 0 | 0 |
| 777 | yidC | 0 | 0 | 0 | 0 | 0 | 0 | 0 | 0 | 0 | 0 | 0 | 0 |
| 778 | ybeZ | 0 | 0 | 0 | 0 | 0 | 1 | 0 | 0 | 0 | 0 | 0 | 0 |
| 779 | yidE | 0 | 0 | 0 | 0 | 0 | 0 | 0 | 0 | 0 | 1 | 0 | 0 |
| 780 | yidK | 111 | 0 | 74 | 0 | 1 | 0 | 98 | 0 | 14 | 0 | 20 | 0 |
| 781 | yidH | 0 | 0 | 0 | 0 | 0 | 0 | 0 | 0 | 0 | 0 | 0 | 0 |
| 782 | ybeU | 0 | 0 | 0 | 0 | 0 | 0 | 0 | 0 | 0 | 0 | 0 | 0 |
| 783 | ybeR | 0 | 0 | 0 | 0 | 0 | 0 | 0 | 0 | 0 | 0 | 0 | 0 |
| 784 | yidL | 164 | 0 | 81 | 0 | 1 | 0 | 133 | 0 | 41 | 0 | 30 | 0 |
| 785 | ybeQ | 0 | 0 | 0 | 0 | 0 | 0 | 0 | 0 | 0 | 0 | 0 | 0 |
| 786 | yidR | 0 | 0 | 0 | 0 | 0 | 0 | 0 | 0 | 0 | 0 | 0 | 0 |
| 787 | yidS | 0 | 0 | 0 | 0 | 0 | 0 | 0 | 0 | 0 | 0 | 0 | 0 |
| 788 | ybeL | 0 | 0 | 0 | 0 | 0 | 0 | 0 | 0 | 0 | 0 | 0 | 0 |
| 789 | ybeF | 0 | 0 | 0 | 0 | 0 | 0 | 0 | 0 | 0 | 0 | 0 | 0 |
| 790 | yidX | 0 | 0 | 0 | 0 | 0 | 0 | 0 | 0 | 0 | 0 | 0 | 0 |
| 791 | ybeB | 0 | 0 | 0 | 0 | 0 | 0 | 0 | 0 | 0 | 0 | 0 | 0 |
| 792 | chaB | 1 | 0 | 0 | 0 | 0 | 0 | 0 | 0 | 0 | 0 | 0 | 2 |
| 793 | cvpA | 0 | 0 | 0 | 0 | 0 | 0 | 0 | 1 | 0 | 0 | 0 | 0 |
| 794 | ycgX | 0 | 0 | 0 | 0 | 0 | 0 | 0 | 0 | 0 | 0 | 0 | 0 |
| 795 | kdgR | 0 | 0 | 0 | 0 | 0 | 0 | 0 | 0 | 0 | 0 | 0 | 0 |
| 796 | lpxP | 0 | 0 | 0 | 0 | 0 | 0 | 0 | 0 | 0 | 0 | 0 | 0 |
| 797 | ssuE | 0 | 0 | 0 | 0 | 0 | 0 | 0 | 0 | 0 | 0 | 0 | 0 |
| 798 | hyaA | 0 | 0 | 0 | 0 | 0 | 0 | 0 | 0 | 0 | 0 | 0 | 0 |
| 799 | yaaH | 0 | 0 | 0 | 0 | 0 | 0 | 0 | 0 | 0 | 0 | 0 | 0 |
| 800 | fimD | 1 | 0 | 0 | 0 | 0 | 0 | 0 | 0 | 0 | 0 | 0 | 0 |
| 801 | cca | 0 | 0 | 0 | 0 | 0 | 0 | 0 | 0 | 0 | 0 | 0 | 0 |
| 802 | ylcE | 0 | 0 | 0 | 0 | 0 | 0 | 0 | 0 | 0 | 0 | 0 | 0 |
| 803 | fumB | 1601 | 4 | 1003 | 0 | 6 | 1 | 1247 | 1 | 461 | 3 | 527 | 0 |
| 804 | ycgL | 0 | 0 | 0 | 0 | 0 | 0 | 0 | 0 | 0 | 0 | 0 | 0 |
| 805 | rrnB | 0 | 0 | 0 | 0 | 0 | 0 | 0 | 0 | 0 | 0 | 0 | 0 |
| 806 | rrnC | 0 | 0 | 0 | 0 | 0 | 0 | 0 | 0 | 0 | 0 | 0 | 0 |
| 807 | rrnA | 0 | 0 | 0 | 0 | 0 | 0 | 0 | 0 | 0 | 0 | 0 | 0 |
| 808 | rrnG | 0 | 0 | 1 | 0 | 0 | 0 | 1 | 0 | 0 | 0 | 1 | 0 |
| 809 | rrnD | 0 | 0 | 0 | 0 | 0 | 0 | 0 | 0 | 0 | 0 | 0 | 0 |
| 810 | amn | 0 | 0 | 0 | 0 | 0 | 0 | 0 | 0 | 0 | 0 | 0 | 0 |
| 811 | rrnH | 0 | 0 | 0 | 0 | 0 | 0 | 0 | 0 | 0 | 0 | 0 | 0 |
| 812 | psd | 0 | 0 | 0 | 0 | 0 | 0 | 0 | 0 | 0 | 0 | 0 | 0 |
| 813 | b1142 | 0 | 0 | 0 | 0 | 0 | 0 | 0 | 0 | 0 | 0 | 0 | 0 |
| 814 | mog | 0 | 0 | 0 | 0 | 0 | 0 | 0 | 0 | 0 | 0 | 0 | 0 |
| 815 | ycgE | 0 | 0 | 0 | 0 | 0 | 0 | 0 | 0 | 0 | 0 | 0 | 0 |
| 816 | wrbA | 0 | 0 | 0 | 0 | 0 | 0 | 0 | 0 | 0 | 0 | 0 | 0 |
| 817 | fldA | 0 | 0 | 0 | 1 | 0 | 0 | 0 | 0 | 0 | 0 | 0 | 0 |
| 818 | ynaI | 52 | 0 | 36 | 0 | 0 | 0 | 38 | 0 | 5 | 0 | 2 | 0 |
| 819 | b1367 | 0 | 0 | 0 | 0 | 0 | 0 | 0 | 0 | 0 | 0 | 0 | 0 |
| 820 | ftsJ | 0 | 0 | 0 | 0 | 0 | 0 | 0 | 0 | 0 | 0 | 0 | 0 |
| 821 | yccR | 0 | 0 | 0 | 0 | 0 | 0 | 0 | 0 | 0 | 0 | 0 | 0 |
| 822 | b1331 | 0 | 0 | 0 | 0 | 0 | 0 | 0 | 0 | 0 | 0 | 0 | 0 |
| 823 | yccE | 0 | 0 | 0 | 0 | 0 | 0 | 0 | 0 | 0 | 0 | 0 | 0 |
| 824 | yccF | 0 | 0 | 0 | 0 | 0 | 0 | 0 | 0 | 0 | 0 | 0 | 0 |
| 825 | yccA | 0 | 0 | 0 | 0 | 0 | 0 | 0 | 0 | 0 | 0 | 0 | 0 |
| 826 | b1966 | 0 | 0 | 0 | 0 | 0 | 0 | 0 | 0 | 0 | 0 | 0 | 0 |
| 827 | b1788 | 1 | 0 | 9 | 0 | 0 | 0 | 3 | 0 | 0 | 0 | 0 | 0 |
| 828 | yccM | 0 | 0 | 0 | 0 | 0 | 0 | 0 | 0 | 0 | 0 | 0 | 0 |
| 829 | cydD | 0 | 0 | 0 | 0 | 0 | 0 | 0 | 0 | 0 | 0 | 0 | 0 |
| 830 | ushA | 0 | 0 | 0 | 0 | 0 | 0 | 0 | 0 | 0 | 0 | 0 | 0 |
| 831 | rmuC | 1 | 0 | 0 | 0 | 0 | 1 | 0 | 0 | 1 | 0 | 0 | 0 |
| 832 | dpiB | 0 | 0 | 0 | 0 | 0 | 0 | 0 | 0 | 0 | 0 | 1 | 0 |
| 833 | dapA | 0 | 0 | 0 | 0 | 0 | 0 | 0 | 0 | 0 | 0 | 0 | 0 |
| 834 | mngR | 0 | 0 | 0 | 0 | 0 | 0 | 0 | 0 | 0 | 0 | 0 | 0 |
| 835 | lolB | 0 | 0 | 0 | 0 | 0 | 0 | 0 | 0 | 0 | 0 | 0 | 0 |
| 836 | b0360 | 0 | 0 | 0 | 0 | 0 | 0 | 0 | 0 | 0 | 0 | 0 | 0 |
| 837 | rnhA | 0 | 0 | 0 | 0 | 0 | 0 | 0 | 0 | 0 | 0 | 0 | 0 |
| 838 | yegE | 0 | 0 | 0 | 0 | 0 | 0 | 0 | 0 | 0 | 0 | 0 | 0 |
| 839 | garD | 0 | 0 | 0 | 0 | 0 | 0 | 0 | 0 | 0 | 0 | 0 | 0 |

| | | | | | | | | | | | | | |
|---|---|---|---|---|---|---|---|---|---|---|---|---|---|
| 840 | ybiU | 0 | 0 | 0 | 0 | 0 | 0 | 0 | 0 | 0 | 0 | 0 | 0 |
| 841 | lon | 0 | 0 | 0 | 0 | 0 | 0 | 0 | 0 | 0 | 0 | 0 | 0 |
| 842 | nrfA | 0 | 0 | 0 | 0 | 0 | 0 | 0 | 0 | 0 | 0 | 0 | 0 |
| 843 | yegP | 0 | 0 | 0 | 0 | 0 | 0 | 0 | 0 | 0 | 0 | 0 | 0 |
| 844 | yegW | 0 | 0 | 0 | 0 | 0 | 0 | 0 | 0 | 0 | 0 | 0 | 0 |
| 845 | yegT | 326 | 1 | 222 | 2 | 3 | 0 | 290 | 104 | 65 | 1 | 161 | 0 |
| 846 | ygaD | 0 | 0 | 0 | 0 | 0 | 0 | 0 | 0 | 0 | 0 | 0 | 0 |
| 847 | ygaZ | 0 | 0 | 0 | 0 | 0 | 0 | 0 | 0 | 0 | 0 | 0 | 0 |
| 848 | zntA | 0 | 0 | 0 | 0 | 0 | 0 | 0 | 0 | 0 | 0 | 0 | 0 |
| 849 | soxR | 0 | 0 | 0 | 0 | 0 | 0 | 0 | 0 | 0 | 0 | 0 | 0 |
| 850 | soxS | 0 | 0 | 0 | 0 | 0 | 0 | 0 | 0 | 0 | 0 | 0 | 0 |
| 851 | ygaQ | 0 | 0 | 0 | 0 | 0 | 0 | 0 | 0 | 0 | 0 | 0 | 0 |
| 852 | aceK | 0 | 0 | 0 | 0 | 0 | 0 | 0 | 0 | 0 | 0 | 0 | 0 |
| 853 | ssrA | 0 | 0 | 0 | 0 | 0 | 0 | 0 | 1 | 0 | 0 | 0 | 0 |
| 854 | aceB | 0 | 0 | 1 | 0 | 0 | 0 | 0 | 0 | 0 | 0 | 0 | 0 |
| 855 | fryB | 0 | 0 | 0 | 0 | 0 | 0 | 0 | 0 | 0 | 0 | 0 | 0 |
| 856 | ypjD | 0 | 0 | 0 | 0 | 0 | 0 | 0 | 0 | 0 | 0 | 0 | 0 |
| 857 | ypjA | 0 | 0 | 0 | 0 | 0 | 0 | 0 | 0 | 0 | 0 | 0 | 0 |
| 858 | rihC | 326 | 1 | 276 | 0 | 0 | 0 | 291 | 0 | 173 | 0 | 0 | 0 |
| 859 | rihA | 0 | 0 | 0 | 0 | 0 | 0 | 0 | 0 | 0 | 0 | 0 | 0 |
| 860 | hsdM | 0 | 0 | 0 | 0 | 0 | 0 | 0 | 0 | 0 | 0 | 0 | 0 |
| 861 | lysA | 0 | 0 | 0 | 0 | 0 | 0 | 0 | 0 | 0 | 0 | 0 | 0 |
| 862 | hsdR | 0 | 0 | 0 | 0 | 0 | 0 | 0 | 0 | 0 | 0 | 0 | 0 |
| 863 | lysR | 0 | 0 | 0 | 0 | 0 | 0 | 0 | 0 | 0 | 0 | 0 | 0 |
| 864 | lysU | 0 | 0 | 0 | 0 | 0 | 0 | 0 | 0 | 1 | 0 | 0 | 1 |
| 865 | eaeH | 0 | 0 | 0 | 0 | 0 | 0 | 0 | 0 | 0 | 0 | 0 | 0 |
| 866 | ansA | 0 | 1 | 0 | 0 | 0 | 0 | 0 | 0 | 0 | 0 | 0 | 0 |
| 867 | ansB | 0 | 0 | 0 | 0 | 0 | 0 | 0 | 0 | 0 | 0 | 0 | 0 |
| 868 | mgsA | 0 | 0 | 0 | 0 | 0 | 0 | 0 | 0 | 0 | 0 | 0 | 0 |
| 869 | yqfA | 0 | 0 | 0 | 1 | 0 | 0 | 0 | 0 | 0 | 0 | 0 | 0 |
| 870 | norR | 0 | 0 | 0 | 0 | 0 | 0 | 0 | 0 | 0 | 0 | 0 | 0 |
| 871 | norV | 0 | 1 | 0 | 1 | 0 | 0 | 0 | 0 | 1 | 0 | 1 | 0 |
| 872 | pth | 0 | 18 | 9 | 4 | 0 | 2 | 1 | 2 | 4 | 9 | 0 | 0 |
| 873 | lacA | 0 | 3 | 1 | 0 | 0 | 0 | 0 | 0 | 1 | 1 | 0 | 0 |
| 874 | lacI | 0 | 0 | 0 | 0 | 0 | 0 | 0 | 0 | 0 | 0 | 0 | 0 |
| 875 | lacY | 0 | 0 | 0 | 0 | 0 | 0 | 0 | 0 | 0 | 0 | 0 | 0 |
| 876 | lacZ | 0 | 0 | 0 | 0 | 0 | 0 | 0 | 0 | 0 | 0 | 0 | 0 |
| 877 | atpI | 0 | 0 | 0 | 0 | 0 | 0 | 0 | 0 | 0 | 0 | 0 | 0 |
| 878 | sanA | 0 | 0 | 0 | 0 | 0 | 0 | 0 | 0 | 0 | 0 | 1 | 0 |
| 879 | pfkB | 0 | 0 | 0 | 0 | 0 | 0 | 0 | 0 | 0 | 0 | 0 | 0 |
| 880 | pfkA | 0 | 0 | 0 | 1 | 0 | 0 | 0 | 0 | 0 | 0 | 0 | 0 |
| 881 | hemH | 0 | 0 | 1 | 0 | 0 | 0 | 0 | 0 | 0 | 0 | 0 | 0 |
| 882 | hemN | 0 | 0 | 0 | 0 | 0 | 0 | 0 | 0 | 0 | 0 | 0 | 0 |
| 883 | yieH | 0 | 0 | 0 | 0 | 0 | 0 | 0 | 0 | 0 | 0 | 0 | 0 |
| 884 | hemL | 1 | 0 | 0 | 0 | 0 | 0 | 0 | 0 | 0 | 0 | 0 | 0 |
| 885 | rhaD | 0 | 0 | 0 | 0 | 0 | 0 | 0 | 0 | 0 | 0 | 1 | 0 |
| 886 | hemB | 0 | 0 | 0 | 0 | 0 | 0 | 0 | 0 | 0 | 0 | 0 | 0 |
| 887 | yieG | 0 | 0 | 0 | 0 | 0 | 0 | 0 | 0 | 0 | 0 | 0 | 0 |
| 888 | ybdR | 0 | 0 | 0 | 0 | 0 | 0 | 0 | 0 | 0 | 0 | 0 | 0 |
| 889 | hemG | 0 | 0 | 0 | 0 | 0 | 0 | 0 | 0 | 0 | 0 | 0 | 0 |
| 890 | ybdH | 0 | 0 | 0 | 0 | 0 | 0 | 0 | 0 | 0 | 0 | 0 | 0 |
| 891 | ybdK | 0 | 0 | 0 | 0 | 0 | 0 | 0 | 0 | 0 | 0 | 0 | 0 |
| 892 | ybdL | 0 | 0 | 0 | 0 | 0 | 0 | 0 | 0 | 0 | 0 | 0 | 0 |
| 893 | ybdN | 0 | 0 | 0 | 0 | 0 | 0 | 0 | 0 | 0 | 0 | 0 | 0 |
| 894 | ykiA | 0 | 0 | 0 | 1 | 0 | 0 | 0 | 0 | 0 | 0 | 0 | 0 |
| 895 | ybdF | 0 | 0 | 0 | 0 | 0 | 0 | 0 | 0 | 0 | 0 | 0 | 0 |
| 896 | cueR | 430 | 2 | 445 | 1 | 9 | 0 | 477 | 0 | 145 | 1 | 113 | 1 |
| 897 | frlA | 0 | 0 | 0 | 0 | 0 | 0 | 0 | 0 | 0 | 0 | 1 | 0 |
| 898 | nanA | 0 | 0 | 0 | 0 | 0 | 0 | 0 | 0 | 0 | 0 | 0 | 0 |
| 899 | cueO | 0 | 0 | 0 | 0 | 0 | 0 | 0 | 0 | 0 | 0 | 0 | 0 |
| 900 | nanR | 0 | 0 | 0 | 0 | 0 | 0 | 0 | 0 | 0 | 0 | 0 | 0 |
| 901 | nanT | 0 | 0 | 0 | 0 | 0 | 0 | 0 | 0 | 0 | 0 | 0 | 0 |
| 902 | trmU | 0 | 0 | 0 | 0 | 0 | 0 | 0 | 0 | 0 | 0 | 1 | 0 |
| 903 | trmA | 0 | 0 | 0 | 0 | 0 | 0 | 0 | 0 | 0 | 0 | 0 | 0 |
| 904 | trmE | 0 | 0 | 1 | 0 | 0 | 0 | 0 | 0 | 0 | 0 | 0 | 0 |
| 905 | dgoR | 0 | 0 | 0 | 0 | 0 | 0 | 0 | 0 | 0 | 0 | 0 | 0 |
| 906 | ggt | 0 | 0 | 0 | 0 | 0 | 0 | 0 | 0 | 0 | 0 | 0 | 0 |
| 907 | ybjS | 0 | 0 | 0 | 0 | 0 | 0 | 0 | 0 | 0 | 0 | 0 | 0 |
| 908 | ybjQ | 0 | 0 | 0 | 0 | 0 | 0 | 0 | 0 | 0 | 0 | 0 | 0 |
| 909 | ybjP | 0 | 0 | 0 | 0 | 0 | 0 | 0 | 0 | 0 | 0 | 0 | 0 |
| 910 | ybjG | 0 | 0 | 0 | 0 | 0 | 0 | 0 | 0 | 0 | 0 | 0 | 0 |
| 911 | ybjE | 0 | 0 | 0 | 0 | 0 | 0 | 0 | 0 | 0 | 0 | 0 | 0 |
| 912 | ybjC | 0 | 0 | 0 | 0 | 0 | 0 | 0 | 0 | 0 | 0 | 0 | 0 |
| 913 | ybjO | 0 | 0 | 0 | 0 | 0 | 0 | 0 | 0 | 0 | 0 | 0 | 0 |
| 914 | ybjN | 0 | 0 | 0 | 0 | 0 | 0 | 0 | 1 | 0 | 1 | 0 | 0 |
| 915 | ybjM | 0 | 0 | 0 | 0 | 0 | 0 | 0 | 0 | 0 | 0 | 0 | 0 |
| 916 | ybjL | 0 | 0 | 0 | 0 | 0 | 0 | 0 | 0 | 0 | 0 | 0 | 0 |
| 917 | ybjK | 572 | 2 | 474 | 2 | 2713 | 55 | 375 | 2 | 215 | 1 | 0 | 0 |
| 918 | ybjJ | 1 | 0 | 1 | 0 | 0 | 0 | 0 | 0 | 0 | 0 | 0 | 0 |
| 919 | ybjH | 0 | 0 | 0 | 0 | 0 | 1 | 0 | 0 | 0 | 0 | 0 | 0 |
| 920 | ylbE | 0 | 0 | 0 | 0 | 0 | 0 | 0 | 0 | 0 | 0 | 0 | 0 |
| 921 | yjjY | 0 | 0 | 0 | 0 | 0 | 0 | 0 | 1 | 0 | 0 | 0 | 0 |
| 922 | ylbG | 0 | 0 | 0 | 0 | 0 | 0 | 0 | 0 | 0 | 0 | 0 | 0 |
| 923 | ylbF | 0 | 0 | 0 | 0 | 0 | 0 | 0 | 0 | 0 | 0 | 0 | 0 |

| | | | | | | | | | | | | | |
|---|---|---|---|---|---|---|---|---|---|---|---|---|---|
| 924 | fklB | 0 | 0 | 0 | 0 | 0 | 0 | 0 | 0 | 0 | 0 | 0 | 0 |
| 925 | valV | 0 | 0 | 0 | 0 | 0 | 0 | 0 | 0 | 0 | 0 | 0 | 0 |
| 926 | valT | 0 | 0 | 0 | 0 | 1 | 0 | 1 | 0 | 0 | 0 | 0 | 0 |
| 927 | zraS | 0 | 0 | 0 | 0 | 0 | 0 | 0 | 0 | 0 | 0 | 0 | 0 |
| 928 | zraP | 0 | 0 | 0 | 0 | 0 | 0 | 0 | 0 | 0 | 0 | 0 | 0 |
| 929 | pntA | 0 | 0 | 0 | 0 | 0 | 0 | 0 | 0 | 0 | 0 | 0 | 0 |
| 930 | aphA | 0 | 0 | 0 | 0 | 0 | 0 | 0 | 0 | 0 | 0 | 0 | 0 |
| 931 | yjjK | 0 | 0 | 0 | 0 | 0 | 0 | 0 | 0 | 0 | 0 | 0 | 0 |
| 932 | yjjJ | 0 | 0 | 0 | 0 | 0 | 0 | 0 | 0 | 0 | 0 | 0 | 0 |
| 933 | tpiA | 0 | 0 | 0 | 0 | 0 | 0 | 0 | 0 | 0 | 0 | 0 | 0 |
| 934 | pldB | 0 | 0 | 0 | 0 | 0 | 0 | 0 | 0 | 0 | 0 | 0 | 0 |
| 935 | malX | 0 | 0 | 0 | 0 | 0 | 0 | 0 | 0 | 0 | 0 | 0 | 0 |
| 936 | malS | 0 | 0 | 0 | 0 | 0 | 0 | 0 | 0 | 0 | 0 | 0 | 0 |
| 937 | malI | 0 | 0 | 0 | 0 | 0 | 0 | 0 | 0 | 0 | 0 | 0 | 0 |
| 938 | malF | 0 | 0 | 0 | 0 | 0 | 0 | 0 | 0 | 0 | 0 | 0 | 0 |
| 939 | malE | 0 | 0 | 0 | 0 | 0 | 0 | 0 | 1 | 0 | 0 | 0 | 1 |
| 940 | crcB | 0 | 0 | 0 | 0 | 0 | 1 | 0 | 0 | 0 | 0 | 0 | 0 |
| 941 | yahD | 0 | 0 | 0 | 0 | 0 | 0 | 0 | 0 | 0 | 0 | 0 | 0 |
| 942 | yahB | 0 | 0 | 0 | 0 | 0 | 0 | 0 | 0 | 0 | 0 | 0 | 0 |
| 943 | yahC | 0 | 0 | 0 | 0 | 0 | 0 | 0 | 0 | 0 | 0 | 0 | 0 |
| 944 | yahN | 1 | 0 | 2 | 2 | 0 | 0 | 3 | 81 | 0 | 0 | 0 | 0 |
| 945 | yahO | 0 | 0 | 0 | 0 | 0 | 0 | 0 | 0 | 0 | 0 | 0 | 0 |
| 946 | ycdZ | 0 | 0 | 0 | 0 | 0 | 0 | 0 | 0 | 0 | 0 | 0 | 0 |
| 947 | yahM | 0 | 0 | 0 | 0 | 0 | 0 | 0 | 0 | 0 | 0 | 0 | 0 |
| 948 | yahK | 0 | 0 | 0 | 0 | 0 | 0 | 0 | 0 | 0 | 0 | 0 | 0 |
| 949 | yahI | 0 | 0 | 0 | 0 | 0 | 0 | 0 | 0 | 0 | 0 | 0 | 0 |
| 950 | ycdC | 0 | 0 | 0 | 0 | 0 | 0 | 0 | 0 | 0 | 0 | 0 | 0 |
| 951 | ibpB | 0 | 0 | 0 | 1 | 0 | 0 | 0 | 0 | 0 | 0 | 0 | 0 |
| 952 | eutR | 0 | 0 | 0 | 0 | 0 | 0 | 0 | 1 | 0 | 0 | 0 | 0 |
| 953 | eutS | 0 | 0 | 0 | 0 | 0 | 0 | 0 | 1 | 0 | 0 | 0 | 0 |
| 954 | csgC | 0 | 0 | 0 | 0 | 0 | 0 | 0 | 0 | 0 | 0 | 0 | 0 |
| 955 | ycdM | 0 | 0 | 0 | 0 | 0 | 0 | 0 | 0 | 0 | 0 | 0 | 0 |
| 956 | ycdN | 0 | 0 | 0 | 0 | 1 | 0 | 0 | 0 | 0 | 1 | 0 | 0 |
| 957 | ycdO | 0 | 0 | 0 | 0 | 0 | 0 | 0 | 0 | 0 | 0 | 0 | 0 |
| 958 | dsrA | 0 | 0 | 0 | 0 | 0 | 0 | 0 | 0 | 0 | 0 | 0 | 0 |
| 959 | sseA | 0 | 0 | 0 | 0 | 0 | 0 | 0 | 0 | 0 | 0 | 0 | 0 |
| 960 | sseB | 0 | 0 | 0 | 0 | 0 | 0 | 0 | 0 | 0 | 0 | 0 | 0 |
| 961 | mntR | 0 | 1 | 0 | 0 | 0 | 0 | 0 | 0 | 0 | 0 | 0 | 0 |
| 962 | yqgA | 153 | 0 | 91 | 0 | 0 | 0 | 137 | 1 | 25 | 0 | 21 | 0 |
| 963 | secA | 0 | 0 | 0 | 0 | 0 | 0 | 1 | 0 | 0 | 0 | 0 | 0 |
| 964 | secB | 0 | 0 | 0 | 0 | 0 | 0 | 0 | 0 | 1 | 0 | 0 | 0 |
| 965 | secE | 0 | 0 | 0 | 0 | 0 | 0 | 0 | 0 | 0 | 0 | 0 | 0 |
| 966 | secM | 0 | 0 | 0 | 0 | 0 | 0 | 0 | 0 | 0 | 0 | 0 | 0 |
| 967 | ymfI | 0 | 0 | 0 | 0 | 0 | 0 | 0 | 0 | 0 | 0 | 0 | 0 |
| 968 | ilvI | 0 | 0 | 0 | 0 | 0 | 0 | 0 | 0 | 0 | 0 | 0 | 0 |
| 969 | coaE | 0 | 0 | 0 | 0 | 0 | 0 | 0 | 0 | 0 | 0 | 0 | 0 |
| 970 | ilvD | 0 | 0 | 0 | 0 | 0 | 0 | 0 | 0 | 0 | 1 | 0 | 0 |
| 971 | ymfE | 0 | 0 | 0 | 0 | 0 | 0 | 0 | 0 | 0 | 0 | 0 | 0 |
| 972 | ilvC | 0 | 0 | 0 | 0 | 0 | 0 | 0 | 0 | 0 | 0 | 0 | 0 |
| 973 | hfq | 0 | 0 | 0 | 0 | 0 | 0 | 0 | 0 | 0 | 0 | 0 | 0 |
| 974 | ego | 0 | 0 | 0 | 0 | 0 | 0 | 0 | 0 | 0 | 0 | 0 | 0 |
| 975 | yedX | 0 | 0 | 0 | 0 | 0 | 0 | 0 | 0 | 0 | 0 | 0 | 0 |
| 976 | yedY | 0 | 0 | 0 | 0 | 0 | 0 | 0 | 0 | 0 | 0 | 0 | 0 |
| 977 | yedW | 0 | 0 | 0 | 0 | 0 | 0 | 0 | 0 | 0 | 0 | 0 | 0 |
| 978 | ybeT | 0 | 0 | 0 | 0 | 0 | 0 | 0 | 0 | 0 | 0 | 0 | 0 |
| 979 | yedP | 0 | 0 | 0 | 0 | 0 | 0 | 0 | 0 | 0 | 0 | 0 | 0 |
| 980 | yedQ | 0 | 0 | 0 | 0 | 0 | 0 | 0 | 0 | 0 | 0 | 0 | 0 |
| 981 | allD | 0 | 0 | 0 | 0 | 0 | 0 | 0 | 0 | 0 | 0 | 0 | 0 |
| 982 | allA | 1 | 0 | 0 | 0 | 0 | 0 | 0 | 0 | 0 | 0 | 0 | 0 |
| 983 | poxB | 0 | 0 | 0 | 0 | 0 | 0 | 0 | 0 | 0 | 0 | 0 | 0 |
| 984 | emrR | 0 | 0 | 0 | 0 | 0 | 0 | 0 | 0 | 0 | 0 | 0 | 1 |
| 985 | ugpA | 0 | 0 | 0 | 0 | 0 | 0 | 0 | 0 | 0 | 0 | 0 | 0 |
| 986 | ugpB | 0 | 0 | 0 | 0 | 0 | 0 | 0 | 0 | 0 | 0 | 0 | 0 |
| 987 | allS | 0 | 0 | 0 | 0 | 0 | 0 | 0 | 0 | 0 | 0 | 0 | 0 |
| 988 | allR | 0 | 0 | 0 | 0 | 0 | 0 | 0 | 1 | 0 | 0 | 0 | 0 |
| 989 | emrA | 0 | 0 | 0 | 0 | 0 | 0 | 0 | 0 | 0 | 0 | 0 | 0 |
| 990 | emrE | 0 | 0 | 0 | 0 | 0 | 0 | 0 | 0 | 0 | 0 | 0 | 0 |
| 991 | pcnB | 0 | 0 | 0 | 0 | 0 | 0 | 0 | 0 | 0 | 0 | 0 | 0 |
| 992 | yibL | 0 | 0 | 1 | 0 | 0 | 0 | 0 | 0 | 0 | 0 | 0 | 0 |
| 993 | yibF | 0 | 0 | 0 | 0 | 0 | 0 | 0 | 0 | 0 | 0 | 0 | 0 |
| 994 | panB | 0 | 0 | 0 | 0 | 0 | 0 | 0 | 0 | 0 | 0 | 0 | 0 |
| 995 | panE | 0 | 0 | 0 | 1 | 0 | 0 | 0 | 0 | 0 | 0 | 0 | 0 |
| 996 | panD | 0 | 0 | 0 | 0 | 0 | 0 | 0 | 0 | 0 | 0 | 0 | 0 |
| 997 | yejG | 0 | 0 | 0 | 0 | 0 | 0 | 0 | 0 | 0 | 0 | 1 | 0 |
| 998 | yejA | 0 | 0 | 0 | 0 | 0 | 0 | 0 | 0 | 0 | 0 | 0 | 0 |
| 999 | yejL | 0 | 0 | 0 | 0 | 0 | 0 | 0 | 0 | 0 | 0 | 0 | 0 |
| 1000 | yejO | 0 | 0 | 0 | 0 | 0 | 0 | 0 | 0 | 0 | 0 | 0 | 0 |
| 1001 | yejH | 0 | 0 | 1 | 0 | 0 | 0 | 0 | 0 | 0 | 0 | 0 | 0 |
| 1002 | rbsD | 0 | 0 | 0 | 0 | 0 | 0 | 0 | 0 | 0 | 0 | 0 | 0 |
| 1003 | yejK | 0 | 0 | 0 | 0 | 0 | 0 | 0 | 0 | 0 | 0 | 0 | 0 |
| 1004 | lsrR | 0 | 0 | 0 | 0 | 0 | 0 | 0 | 0 | 0 | 0 | 0 | 0 |
| 1005 | ydiF | 0 | 0 | 0 | 0 | 0 | 0 | 0 | 1 | 0 | 0 | 0 | 0 |
| 1006 | ygbA | 0 | 0 | 0 | 0 | 0 | 0 | 0 | 0 | 0 | 0 | 0 | 0 |
| 1007 | ygbN | 0 | 0 | 0 | 0 | 0 | 0 | 0 | 0 | 0 | 0 | 0 | 0 |

| | | | | | | | | | | | | | |
|---|---|---|---|---|---|---|---|---|---|---|---|---|---|
| 1008 | ygbJ | 0 | 0 | 0 | 0 | 0 | 0 | 0 | 0 | 0 | 0 | 0 | 0 |
| 1009 | ygbI | 0 | 0 | 0 | 0 | 0 | 0 | 0 | 0 | 0 | 0 | 0 | 0 |
| 1010 | adrA | 0 | 0 | 0 | 0 | 0 | 0 | 0 | 0 | 0 | 1 | 0 | 0 |
| 1011 | dedA | 0 | 0 | 1 | 0 | 0 | 0 | 0 | 0 | 0 | 0 | 9 | 0 |
| 1012 | lolC | 0 | 0 | 0 | 0 | 0 | 0 | 0 | 0 | 0 | 0 | 0 | 0 |
| 1013 | ybiS | 0 | 1 | 0 | 0 | 0 | 0 | 0 | 0 | 0 | 0 | 0 | 0 |
| 1014 | lolA | 0 | 0 | 1 | 0 | 0 | 0 | 0 | 0 | 0 | 0 | 0 | 0 |
| 1015 | ybiV | 0 | 0 | 0 | 0 | 0 | 0 | 0 | 0 | 1 | 0 | 0 | 0 |
| 1016 | dapF | 0 | 0 | 0 | 0 | 0 | 0 | 0 | 0 | 0 | 1 | 0 | 0 |
| 1017 | ybiA | 396 | 0 | 369 | 46 | 0 | 7 | 424 | 2331 | 183 | 1 | 80 | 0 |
| 1018 | ybiF | 0 | 0 | 0 | 0 | 0 | 0 | 0 | 0 | 0 | 0 | 0 | 0 |
| 1019 | ybiJ | 1 | 0 | 1 | 0 | 0 | 0 | 0 | 0 | 0 | 0 | 0 | 0 |
| 1020 | ybiH | 0 | 0 | 0 | 0 | 0 | 0 | 0 | 0 | 0 | 0 | 0 | 0 |
| 1021 | ybiN | 0 | 0 | 0 | 0 | 0 | 0 | 0 | 0 | 0 | 0 | 0 | 0 |
| 1022 | ybiO | 0 | 0 | 0 | 1 | 0 | 0 | 0 | 0 | 0 | 0 | 0 | 0 |
| 1023 | ybiM | 0 | 0 | 0 | 0 | 0 | 0 | 0 | 0 | 0 | 0 | 0 | 0 |
| 1024 | nac | 0 | 0 | 0 | 0 | 0 | 0 | 0 | 0 | 0 | 0 | 0 | 0 |
| 1025 | b2680 | 1 | 0 | 0 | 0 | 0 | 0 | 0 | 0 | 0 | 0 | 0 | 0 |
| 1026 | asr | 0 | 0 | 0 | 0 | 0 | 0 | 0 | 0 | 0 | 0 | 0 | 0 |
| 1027 | oppA | 0 | 0 | 0 | 0 | 0 | 0 | 0 | 0 | 0 | 0 | 0 | 0 |
| 1028 | oppB | 0 | 0 | 0 | 0 | 0 | 0 | 0 | 0 | 0 | 0 | 0 | 0 |
| 1029 | asd | 0 | 0 | 0 | 0 | 0 | 0 | 0 | 0 | 0 | 0 | 0 | 0 |
| 1030 | yaiE | 0 | 0 | 0 | 0 | 0 | 0 | 0 | 0 | 0 | 0 | 0 | 0 |
| 1031 | dinB | 0 | 0 | 0 | 0 | 0 | 0 | 0 | 0 | 0 | 0 | 0 | 0 |
| 1032 | recA | 0 | 0 | 0 | 0 | 0 | 0 | 0 | 0 | 0 | 0 | 0 | 0 |
| 1033 | recE | 0 | 0 | 0 | 0 | 0 | 0 | 0 | 0 | 0 | 0 | 0 | 0 |
| 1034 | recX | 0 | 0 | 0 | 0 | 0 | 0 | 0 | 0 | 0 | 1 | 0 | 1 |
| 1035 | ispB | 0 | 0 | 0 | 0 | 0 | 0 | 0 | 0 | 0 | 0 | 0 | 0 |
| 1036 | hscB | 0 | 0 | 0 | 0 | 0 | 0 | 0 | 0 | 0 | 0 | 0 | 0 |
| 1037 | ydiL | 684 | 0 | 501 | 2370 | 1 | 11 | 350 | 26887 | 59 | 43 | 1 | 0 |
| 1038 | ydiM | 0 | 0 | 0 | 0 | 0 | 0 | 0 | 0 | 0 | 0 | 0 | 0 |
| 1039 | tatE | 0 | 0 | 0 | 0 | 0 | 0 | 0 | 0 | 0 | 0 | 0 | 0 |
| 1040 | pspF | 0 | 0 | 0 | 0 | 0 | 0 | 0 | 0 | 0 | 0 | 0 | 0 |
| 1041 | pspE | 0 | 0 | 0 | 0 | 0 | 0 | 0 | 0 | 1 | 0 | 0 | 0 |
| 1042 | pspB | 0 | 0 | 0 | 0 | 0 | 0 | 0 | 0 | 0 | 0 | 0 | 0 |
| 1043 | moaA | 0 | 0 | 0 | 0 | 0 | 0 | 0 | 0 | 0 | 0 | 0 | 0 |
| 1044 | dinJ | 0 | 0 | 0 | 0 | 0 | 0 | 0 | 0 | 0 | 0 | 0 | 0 |
| 1045 | ydiY | 0 | 0 | 0 | 0 | 0 | 0 | 0 | 0 | 0 | 0 | 0 | 0 |
| 1046 | ydiP | 0 | 0 | 0 | 0 | 0 | 0 | 0 | 0 | 0 | 0 | 0 | 0 |
| 1047 | ydiQ | 0 | 0 | 0 | 0 | 0 | 0 | 0 | 0 | 0 | 0 | 0 | 0 |
| 1048 | ydiS | 0 | 0 | 0 | 0 | 0 | 0 | 0 | 0 | 0 | 0 | 0 | 0 |
| 1049 | ydiU | 0 | 0 | 0 | 0 | 0 | 0 | 0 | 1 | 0 | 0 | 0 | 0 |
| 1050 | ydiV | 0 | 0 | 0 | 0 | 0 | 0 | 0 | 0 | 0 | 0 | 0 | 0 |
| 1051 | dinG | 10 | 0 | 7 | 0 | 0 | 0 | 11 | 67 | 3 | 0 | 0 | 0 |
| 1052 | yceP | 0 | 0 | 0 | 1 | 0 | 0 | 0 | 0 | 0 | 0 | 0 | 0 |
| 1053 | guaC | 675 | 5 | 1031 | 772 | 6 | 0 | 743 | 2 | 375 | 0 | 216 | 546 |
| 1054 | guaB | 0 | 0 | 0 | 0 | 0 | 0 | 0 | 0 | 0 | 0 | 0 | 0 |
| 1055 | guaA | 0 | 0 | 0 | 0 | 0 | 0 | 0 | 0 | 0 | 0 | 0 | 0 |
| 1056 | cadA | 0 | 0 | 0 | 0 | 0 | 0 | 0 | 0 | 0 | 0 | 0 | 0 |
| 1057 | yaiI | 0 | 0 | 0 | 0 | 0 | 0 | 0 | 0 | 0 | 0 | 0 | 0 |
| 1058 | yaiM | 0 | 0 | 0 | 0 | 0 | 1 | 0 | 0 | 0 | 0 | 0 | 0 |
| 1059 | dfp | 0 | 0 | 0 | 0 | 0 | 0 | 0 | 0 | 0 | 0 | 0 | 0 |
| 1060 | yaiN | 0 | 0 | 0 | 0 | 0 | 0 | 0 | 0 | 0 | 0 | 0 | 0 |
| 1061 | yaiP | 0 | 0 | 1 | 0 | 0 | 0 | 0 | 0 | 0 | 0 | 0 | 0 |
| 1062 | cadB | 0 | 0 | 0 | 1 | 0 | 0 | 0 | 1 | 0 | 0 | 0 | 0 |
| 1063 | yceF | 247 | 0 | 212 | 1 | 4 | 0 | 322 | 1 | 96 | 0 | 0 | 0 |
| 1064 | yaiW | 0 | 0 | 0 | 0 | 0 | 0 | 0 | 0 | 0 | 0 | 0 | 0 |
| 1065 | gpt | 0 | 0 | 0 | 0 | 0 | 0 | 0 | 0 | 0 | 0 | 0 | 0 |
| 1066 | yaiX | 0 | 0 | 0 | 0 | 0 | 0 | 0 | 0 | 0 | 0 | 0 | 0 |
| 1067 | yaiZ | 0 | 0 | 0 | 0 | 0 | 0 | 0 | 0 | 0 | 0 | 0 | 0 |
| 1068 | gpp | 0 | 0 | 0 | 0 | 0 | 1 | 0 | 0 | 0 | 0 | 0 | 0 |
| 1069 | gntT | 0 | 0 | 0 | 0 | 0 | 0 | 0 | 0 | 0 | 0 | 0 | 0 |
| 1070 | ubiX | 0 | 0 | 0 | 0 | 0 | 0 | 0 | 0 | 0 | 0 | 0 | 0 |
| 1071 | b0501 | 0 | 0 | 0 | 0 | 0 | 0 | 0 | 0 | 0 | 0 | 0 | 0 |
| 1072 | yceK | 0 | 0 | 0 | 0 | 0 | 0 | 0 | 0 | 0 | 0 | 0 | 0 |
| 1073 | greA | 0 | 0 | 0 | 0 | 0 | 0 | 0 | 0 | 0 | 0 | 0 | 0 |
| 1074 | greB | 0 | 0 | 0 | 0 | 0 | 0 | 0 | 0 | 0 | 0 | 0 | 0 |
| 1075 | gntR | 1 | 0 | 1 | 0 | 0 | 0 | 0 | 0 | 1 | 0 | 1 | 0 |
| 1076 | ubiF | 163276 | 8509 | 152783 | 9044 | 132871 | 42827 | 150216 | 34330 | 74946 | 8775 | 194248 | 6619 |
| 1077 | ubiG | 48673 | 19788 | 52371 | 20430 | 109756 | 46138 | 45535 | 47075 | 31617 | 16158 | 37900 | 53416 |
| 1078 | ubiE | 0 | 1 | 0 | 0 | 0 | 0 | 2 | 0 | 0 | 1 | 1 | 0 |
| 1079 | ubiC | 0 | 0 | 0 | 0 | 0 | 0 | 0 | 0 | 0 | 0 | 0 | 0 |
| 1080 | gntP | 0 | 0 | 0 | 0 | 0 | 0 | 0 | 0 | 0 | 0 | 0 | 0 |
| 1081 | parC | 0 | 0 | 0 | 0 | 0 | 0 | 0 | 1 | 0 | 0 | 0 | 0 |
| 1082 | bglA | 840 | 4 | 564 | 0 | 5 | 0 | 634 | 2 | 358 | 0 | 295 | 0 |
| 1083 | ascG | 0 | 0 | 0 | 0 | 0 | 0 | 0 | 0 | 0 | 1 | 0 | 0 |
| 1084 | b0165 | 0 | 0 | 0 | 0 | 0 | 0 | 0 | 0 | 0 | 0 | 0 | 0 |
| 1085 | selD | 0 | 0 | 0 | 0 | 0 | 0 | 0 | 0 | 0 | 0 | 0 | 0 |
| 1086 | alsR | 0 | 0 | 0 | 0 | 0 | 0 | 0 | 0 | 0 | 0 | 0 | 0 |
| 1087 | selA | 0 | 0 | 0 | 0 | 0 | 0 | 0 | 1 | 0 | 0 | 0 | 0 |
| 1088 | selC | 0 | 0 | 0 | 0 | 0 | 0 | 0 | 0 | 0 | 0 | 0 | 0 |
| 1089 | alsI | 0 | 0 | 0 | 0 | 0 | 0 | 0 | 0 | 0 | 0 | 0 | 0 |
| 1090 | alsB | 0 | 0 | 0 | 0 | 0 | 0 | 0 | 0 | 0 | 0 | 0 | 0 |
| 1091 | ymgE | 0 | 0 | 0 | 0 | 0 | 0 | 0 | 0 | 0 | 0 | 1 | 0 |

| | | | | | | | | | | | | | |
|---|---|---|---|---|---|---|---|---|---|---|---|---|---|
| 1092 | alsA | 0 | 0 | 0 | 0 | 0 | 0 | 0 | 0 | 0 | 0 | 0 | 0 |
| 1093 | yeeI | 0 | 0 | 0 | 0 | 0 | 0 | 0 | 0 | 0 | 0 | 0 | 0 |
| 1094 | galU | 0 | 0 | 0 | 0 | 0 | 0 | 0 | 0 | 0 | 0 | 0 | 0 |
| 1095 | yeeJ | 0 | 0 | 0 | 0 | 0 | 0 | 0 | 0 | 0 | 0 | 0 | 0 |
| 1096 | galP | 0 | 0 | 0 | 0 | 0 | 0 | 0 | 0 | 0 | 0 | 0 | 0 |
| 1097 | galR | 0 | 0 | 0 | 0 | 0 | 0 | 0 | 0 | 0 | 0 | 0 | 0 |
| 1098 | galS | 506 | 415 | 376 | 338 | 0 | 10 | 498 | 0 | 725 | 710 | 151 | 0 |
| 1099 | yeeA | 0 | 0 | 0 | 1 | 0 | 0 | 0 | 0 | 0 | 0 | 0 | 0 |
| 1100 | yeeE | 0 | 0 | 0 | 0 | 0 | 0 | 0 | 0 | 0 | 0 | 0 | 0 |
| 1101 | lit | 0 | 0 | 0 | 0 | 0 | 0 | 0 | 0 | 0 | 0 | 0 | 0 |
| 1102 | galE | 0 | 0 | 0 | 0 | 0 | 0 | 0 | 0 | 0 | 0 | 0 | 0 |
| 1103 | yeeR | 0 | 0 | 0 | 1 | 0 | 0 | 0 | 0 | 0 | 0 | 0 | 0 |
| 1104 | yeeU | 1 | 0 | 0 | 0 | 0 | 0 | 0 | 1 | 0 | 0 | 0 | 0 |
| 1105 | yeeV | 0 | 0 | 0 | 0 | 0 | 0 | 0 | 0 | 0 | 0 | 0 | 0 |
| 1106 | hipB | 0 | 0 | 0 | 0 | 0 | 0 | 0 | 0 | 0 | 0 | 0 | 0 |
| 1107 | rho | 0 | 0 | 0 | 0 | 0 | 0 | 0 | 0 | 0 | 0 | 1 | 0 |
| 1108 | msrA | 0 | 0 | 0 | 0 | 0 | 0 | 0 | 0 | 0 | 0 | 0 | 0 |
| 1109 | hofM | 0 | 0 | 0 | 0 | 0 | 0 | 0 | 0 | 0 | 0 | 0 | 0 |
| 1110 | yhaH | 0 | 0 | 0 | 0 | 0 | 0 | 0 | 0 | 0 | 0 | 0 | 0 |
| 1111 | yhaJ | 0 | 0 | 0 | 0 | 0 | 0 | 0 | 0 | 0 | 0 | 0 | 0 |
| 1112 | yhaK | 0 | 0 | 0 | 0 | 0 | 0 | 0 | 0 | 0 | 0 | 0 | 0 |
| 1113 | cdd | 0 | 0 | 0 | 0 | 0 | 0 | 0 | 1 | 0 | 0 | 1 | 0 |
| 1114 | yoaB | 0 | 0 | 0 | 0 | 0 | 0 | 0 | 0 | 0 | 0 | 0 | 0 |
| 1115 | yoaA | 0 | 0 | 0 | 0 | 0 | 0 | 0 | 0 | 0 | 0 | 0 | 0 |
| 1116 | prpB | 0 | 0 | 0 | 0 | 0 | 0 | 0 | 0 | 0 | 0 | 0 | 0 |
| 1117 | prpC | 0 | 0 | 0 | 0 | 0 | 0 | 0 | 0 | 0 | 0 | 0 | 0 |
| 1118 | yoaE | 0 | 0 | 0 | 0 | 0 | 0 | 0 | 0 | 0 | 0 | 0 | 0 |
| 1119 | yoaD | 0 | 0 | 0 | 0 | 0 | 0 | 0 | 0 | 0 | 0 | 0 | 0 |
| 1120 | yoaH | 0 | 0 | 0 | 1 | 0 | 0 | 0 | 0 | 0 | 0 | 0 | 0 |
| 1121 | ugd | 0 | 0 | 0 | 0 | 0 | 0 | 0 | 0 | 0 | 0 | 0 | 0 |
| 1122 | ygcB | 0 | 0 | 0 | 0 | 0 | 0 | 0 | 0 | 0 | 0 | 0 | 0 |
| 1123 | prpR | 0 | 0 | 1 | 0 | 0 | 0 | 0 | 0 | 0 | 0 | 0 | 0 |
| 1124 | ygcF | 0 | 0 | 0 | 0 | 0 | 0 | 0 | 0 | 0 | 0 | 0 | 0 |
| 1125 | clpX | 0 | 0 | 0 | 0 | 0 | 0 | 0 | 1 | 0 | 0 | 0 | 0 |
| 1126 | stfE | 0 | 0 | 0 | 0 | 0 | 0 | 0 | 0 | 0 | 0 | 0 | 0 |
| 1127 | bfd | 0 | 0 | 0 | 0 | 0 | 0 | 0 | 0 | 0 | 0 | 0 | 0 |
| 1128 | clpP | 0 | 0 | 0 | 0 | 0 | 0 | 0 | 0 | 0 | 0 | 0 | 1 |
| 1129 | clpS | 0 | 0 | 0 | 0 | 0 | 0 | 0 | 0 | 0 | 0 | 0 | 0 |
| 1130 | gshA | 0 | 0 | 0 | 0 | 0 | 0 | 0 | 0 | 0 | 0 | 0 | 0 |
| 1131 | sspA | 0 | 0 | 0 | 0 | 0 | 0 | 0 | 0 | 0 | 0 | 0 | 0 |
| 1132 | stfR | 0 | 0 | 0 | 0 | 0 | 0 | 0 | 0 | 0 | 0 | 0 | 0 |
| 1133 | hyfR | 0 | 0 | 0 | 0 | 0 | 0 | 0 | 0 | 0 | 0 | 0 | 0 |
| 1134 | clpB | 0 | 0 | 0 | 0 | 0 | 0 | 0 | 0 | 0 | 0 | 0 | 0 |
| 1135 | sthA | 0 | 1 | 0 | 0 | 0 | 0 | 0 | 0 | 1 | 0 | 0 | 0 |
| 1136 | plsX | 0 | 0 | 0 | 0 | 0 | 0 | 0 | 0 | 0 | 0 | 0 | 0 |
| 1137 | hrpB | 3320 | 17 | 2684 | 5 | 32 | 1 | 2761 | 2 | 1579 | 2 | 1458 | 5 |
| 1138 | hrpA | 1 | 1 | 0 | 0 | 0 | 0 | 0 | 0 | 0 | 0 | 1 | 0 |
| 1139 | plsC | 0 | 0 | 0 | 0 | 0 | 0 | 0 | 1 | 0 | 0 | 0 | 0 |
| 1140 | plsB | 0 | 0 | 0 | 0 | 0 | 0 | 0 | 0 | 0 | 0 | 0 | 0 |
| 1141 | rpiA | 0 | 0 | 0 | 0 | 0 | 0 | 0 | 0 | 0 | 0 | 0 | 0 |
| 1142 | pck | 0 | 0 | 0 | 0 | 0 | 0 | 0 | 0 | 0 | 0 | 0 | 0 |
| 1143 | ldhA | 0 | 0 | 0 | 0 | 0 | 0 | 0 | 0 | 0 | 0 | 0 | 0 |
| 1144 | yjcD | 0 | 0 | 0 | 0 | 0 | 0 | 0 | 0 | 0 | 0 | 0 | 0 |
| 1145 | yjcE | 0 | 0 | 0 | 0 | 0 | 0 | 0 | 0 | 0 | 0 | 0 | 0 |
| 1146 | crp | 0 | 0 | 0 | 0 | 0 | 0 | 0 | 0 | 0 | 0 | 0 | 0 |
| 1147 | fsaA | 0 | 0 | 0 | 0 | 0 | 0 | 0 | 0 | 0 | 0 | 0 | 0 |
| 1148 | fkpB | 0 | 0 | 1 | 0 | 0 | 0 | 0 | 0 | 1 | 0 | 0 | 0 |
| 1149 | yjcO | 32316 | 260 | 29060 | 249 | 2425 | 7781 | 29072 | 24542 | 14190 | 174 | 13569 | 289 |
| 1150 | yjcH | 1 | 0 | 2 | 0 | 0 | 0 | 2 | 1 | 1 | 0 | 1 | 0 |
| 1151 | rfe | 0 | 0 | 0 | 0 | 1 | 0 | 0 | 0 | 0 | 0 | 0 | 0 |
| 1152 | aaeR | 0 | 0 | 0 | 0 | 0 | 0 | 0 | 0 | 0 | 0 | 0 | 0 |
| 1153 | crl | 0 | 0 | 0 | 0 | 0 | 0 | 0 | 0 | 0 | 0 | 0 | 0 |
| 1154 | agn43 | 0 | 0 | 0 | 0 | 0 | 0 | 0 | 0 | 0 | 0 | 0 | 0 |
| 1155 | uxuA | 0 | 0 | 0 | 0 | 0 | 0 | 0 | 0 | 0 | 0 | 0 | 0 |
| 1156 | cysK | 0 | 0 | 0 | 0 | 0 | 0 | 0 | 0 | 0 | 0 | 0 | 0 |
| 1157 | barA | 346 | 0 | 334 | 1 | 3 | 0 | 258 | 0 | 166 | 0 | 26 | 0 |
| 1158 | cbpA | 0 | 0 | 0 | 0 | 0 | 0 | 0 | 0 | 0 | 0 | 0 | 0 |
| 1159 | imp | 0 | 0 | 0 | 0 | 0 | 0 | 0 | 0 | 0 | 0 | 0 | 0 |
| 1160 | mhpR | 0 | 0 | 0 | 0 | 0 | 0 | 0 | 0 | 0 | 0 | 0 | 0 |
| 1161 | yicH | 0 | 0 | 0 | 0 | 0 | 0 | 1 | 0 | 0 | 0 | 0 | 0 |
| 1162 | yicC | 84 | 1 | 66 | 0 | 3 | 0 | 107 | 0 | 24 | 1 | 47 | 0 |
| 1163 | solA | 0 | 0 | 0 | 1 | 0 | 0 | 0 | 0 | 0 | 0 | 0 | 0 |
| 1164 | fdhF | 0 | 0 | 0 | 0 | 0 | 0 | 0 | 1 | 0 | 0 | 0 | 0 |
| 1165 | fdhD | 0 | 0 | 0 | 0 | 0 | 0 | 0 | 0 | 0 | 0 | 0 | 0 |
| 1166 | tgt | 0 | 0 | 0 | 0 | 0 | 0 | 0 | 0 | 0 | 0 | 0 | 0 |
| 1167 | napF | 0 | 0 | 0 | 0 | 0 | 0 | 0 | 0 | 1 | 0 | 0 | 0 |
| 1168 | sppA | 0 | 0 | 0 | 0 | 0 | 0 | 0 | 0 | 0 | 0 | 0 | 0 |
| 1169 | ybhQ | 0 | 0 | 0 | 0 | 0 | 0 | 1 | 0 | 0 | 0 | 0 | 0 |
| 1170 | ybhM | 0 | 0 | 0 | 1 | 0 | 0 | 0 | 0 | 0 | 0 | 0 | 0 |
| 1171 | ybhL | 0 | 0 | 0 | 0 | 0 | 0 | 0 | 0 | 0 | 0 | 0 | 0 |
| 1172 | ybhI | 0 | 0 | 0 | 0 | 0 | 0 | 0 | 0 | 0 | 0 | 0 | 0 |
| 1173 | ybhK | 0 | 0 | 0 | 0 | 0 | 0 | 0 | 0 | 0 | 0 | 0 | 0 |
| 1174 | ybhJ | 0 | 0 | 0 | 1 | 0 | 0 | 0 | 0 | 0 | 0 | 0 | 0 |
| 1175 | ybhE | 0 | 0 | 0 | 0 | 0 | 0 | 0 | 0 | 0 | 0 | 0 | 0 |

| | | | | | | | | | | | | | |
|---|---|---|---|---|---|---|---|---|---|---|---|---|---|
| 1176 | ybhD | 0 | 0 | 0 | 0 | 0 | 0 | 0 | 0 | 0 | 0 | 0 | 0 |
| 1177 | ybhA | 0 | 0 | 0 | 0 | 0 | 0 | 0 | 0 | 0 | 0 | 0 | 0 |
| 1178 | ybhC | 35 | 0 | 18 | 1 | 0 | 0 | 34 | 3 | 0 | 0 | 32 | 0 |
| 1179 | mlc | 0 | 0 | 0 | 0 | 0 | 0 | 0 | 0 | 0 | 0 | 0 | 0 |
| 1180 | fur | 0 | 0 | 0 | 0 | 0 | 0 | 0 | 0 | 0 | 0 | 0 | 0 |
| 1181 | tdcB | 0 | 0 | 0 | 0 | 0 | 0 | 0 | 0 | 0 | 0 | 1 | 0 |
| 1182 | b1172 | 0 | 0 | 0 | 0 | 0 | 0 | 0 | 0 | 0 | 0 | 0 | 0 |
| 1183 | glgB | 0 | 0 | 0 | 0 | 0 | 0 | 0 | 0 | 0 | 0 | 0 | 0 |
| 1184 | glgS | 0 | 0 | 0 | 0 | 0 | 0 | 0 | 0 | 0 | 0 | 0 | 0 |
| 1185 | motA | 0 | 0 | 0 | 0 | 0 | 0 | 0 | 0 | 0 | 0 | 0 | 0 |
| 1186 | ptrB | 0 | 0 | 2 | 0 | 0 | 0 | 0 | 0 | 0 | 1 | 0 | 0 |
| 1187 | dsbG | 0 | 0 | 0 | 0 | 0 | 0 | 0 | 0 | 0 | 0 | 0 | 0 |
| 1188 | ampH | 0 | 0 | 0 | 0 | 0 | 0 | 0 | 0 | 0 | 0 | 0 | 0 |
| 1189 | ydhH | 0 | 0 | 0 | 0 | 0 | 0 | 0 | 0 | 0 | 0 | 0 | 0 |
| 1190 | ydhO | 0 | 0 | 0 | 0 | 0 | 0 | 0 | 0 | 0 | 0 | 0 | 0 |
| 1191 | syd | 0 | 0 | 0 | 0 | 0 | 0 | 0 | 0 | 0 | 0 | 0 | 0 |
| 1192 | ydhB | 0 | 0 | 0 | 0 | 0 | 0 | 0 | 0 | 0 | 0 | 0 | 0 |
| 1193 | mobA | 252 | 2 | 210 | 0 | 2 | 0 | 214 | 0 | 201 | 0 | 376 | 2 |
| 1194 | ampD | 0 | 0 | 0 | 0 | 0 | 0 | 0 | 0 | 0 | 0 | 0 | 0 |
| 1195 | ydhD | 0 | 0 | 0 | 0 | 0 | 0 | 0 | 0 | 0 | 0 | 0 | 0 |
| 1196 | ydhZ | 0 | 0 | 0 | 0 | 0 | 0 | 0 | 0 | 0 | 0 | 0 | 0 |
| 1197 | dgt | 0 | 0 | 0 | 0 | 0 | 0 | 1 | 0 | 0 | 0 | 0 | 0 |
| 1198 | ydhS | 0 | 1 | 0 | 0 | 0 | 0 | 0 | 0 | 0 | 0 | 0 | 0 |
| 1199 | ydhR | 5759 | 48713 | 4193 | 11513 | 28 | 7 | 4409 | 14212 | 12568 | 19980 | 562 | 1 |
| 1200 | ydhQ | 0 | 1 | 0 | 0 | 0 | 0 | 1 | 0 | 0 | 0 | 0 | 0 |
| 1201 | ydhP | 0 | 0 | 0 | 0 | 0 | 0 | 0 | 0 | 0 | 0 | 0 | 0 |
| 1202 | yafL | 0 | 0 | 0 | 0 | 0 | 0 | 0 | 0 | 0 | 0 | 0 | 0 |
| 1203 | yafM | 0 | 1 | 0 | 0 | 0 | 0 | 0 | 0 | 0 | 0 | 0 | 0 |
| 1204 | yafN | 0 | 0 | 0 | 0 | 0 | 0 | 0 | 0 | 0 | 0 | 0 | 0 |
| 1205 | zitB | 0 | 0 | 0 | 0 | 0 | 0 | 0 | 0 | 0 | 0 | 0 | 0 |
| 1206 | yafJ | 0 | 0 | 0 | 0 | 0 | 0 | 0 | 0 | 0 | 0 | 0 | 0 |
| 1207 | yafK | 1 | 0 | 0 | 0 | 0 | 0 | 0 | 0 | 0 | 0 | 0 | 0 |
| 1208 | yafD | 0 | 0 | 0 | 0 | 0 | 0 | 0 | 0 | 0 | 0 | 0 | 0 |
| 1209 | ycfT | 0 | 0 | 0 | 0 | 0 | 0 | 0 | 0 | 0 | 0 | 0 | 0 |
| 1210 | mipA | 1 | 0 | 0 | 0 | 0 | 0 | 0 | 1 | 0 | 0 | 0 | 0 |
| 1211 | ycfR | 0 | 0 | 0 | 0 | 0 | 0 | 0 | 0 | 0 | 0 | 0 | 0 |
| 1212 | ycfS | 0 | 0 | 0 | 0 | 0 | 0 | 0 | 0 | 0 | 0 | 0 | 0 |
| 1213 | slyX | 0 | 0 | 0 | 0 | 0 | 0 | 0 | 0 | 0 | 0 | 0 | 0 |
| 1214 | yafC | 0 | 0 | 0 | 0 | 0 | 0 | 0 | 0 | 0 | 0 | 0 | 0 |
| 1215 | efp | 0 | 0 | 0 | 0 | 0 | 0 | 0 | 0 | 0 | 0 | 0 | 0 |
| 1216 | slyD | 0 | 0 | 0 | 0 | 0 | 0 | 0 | 0 | 0 | 0 | 0 | 0 |
| 1217 | ycfJ | 0 | 0 | 0 | 0 | 0 | 0 | 0 | 0 | 0 | 0 | 0 | 0 |
| 1218 | yafZ | 0 | 0 | 0 | 0 | 0 | 0 | 0 | 0 | 0 | 0 | 0 | 0 |
| 1219 | slyA | 0 | 0 | 0 | 0 | 0 | 0 | 0 | 0 | 0 | 0 | 0 | 0 |
| 1220 | yafT | 0 | 0 | 0 | 0 | 0 | 0 | 0 | 0 | 0 | 0 | 0 | 0 |
| 1221 | yafU | 0 | 0 | 0 | 0 | 0 | 0 | 0 | 0 | 0 | 0 | 1 | 0 |
| 1222 | yafV | 0 | 0 | 0 | 0 | 0 | 0 | 0 | 1 | 0 | 0 | 0 | 0 |
| 1223 | uxaC | 0 | 0 | 0 | 0 | 0 | 0 | 0 | 0 | 0 | 0 | 0 | 0 |
| 1224 | flgM | 0 | 0 | 0 | 0 | 0 | 1 | 1 | 0 | 0 | 0 | 0 | 0 |
| 1225 | yafS | 0 | 0 | 1 | 0 | 0 | 0 | 0 | 0 | 0 | 0 | 0 | 0 |
| 1226 | sscR | 0 | 0 | 0 | 0 | 0 | 0 | 0 | 0 | 0 | 0 | 0 | 0 |
| 1227 | dhaM | 0 | 0 | 0 | 0 | 0 | 0 | 0 | 0 | 0 | 0 | 0 | 0 |
| 1228 | dhaR | 0 | 0 | 0 | 0 | 0 | 0 | 0 | 0 | 0 | 1 | 0 | 0 |
| 1229 | yjbF | 1 | 0 | 0 | 0 | 0 | 0 | 0 | 0 | 1 | 0 | 0 | 0 |
| 1230 | yjbE | 0 | 0 | 0 | 0 | 1 | 0 | 0 | 0 | 0 | 0 | 0 | 0 |
| 1231 | yjbB | 0 | 0 | 0 | 0 | 0 | 0 | 0 | 0 | 0 | 0 | 0 | 0 |
| 1232 | yeeX | 0 | 0 | 0 | 0 | 0 | 0 | 0 | 0 | 0 | 0 | 0 | 0 |
| 1233 | mreC | 475 | 6 | 331 | 1 | 978 | 10 | 373 | 0 | 174 | 0 | 179 | 0 |
| 1234 | yjbJ | 0 | 0 | 0 | 0 | 0 | 0 | 0 | 0 | 0 | 0 | 0 | 0 |
| 1235 | yjbI | 0 | 0 | 0 | 0 | 0 | 0 | 0 | 0 | 0 | 0 | 0 | 0 |
| 1236 | yjdB | 0 | 0 | 0 | 0 | 0 | 0 | 0 | 0 | 0 | 0 | 0 | 0 |
| 1237 | yjbQ | 0 | 0 | 0 | 0 | 0 | 0 | 0 | 0 | 0 | 0 | 0 | 0 |
| 1238 | deoR | 1 | 0 | 0 | 0 | 0 | 0 | 0 | 0 | 1 | 0 | 0 | 0 |
| 1239 | yhhQ | 0 | 0 | 0 | 0 | 0 | 0 | 0 | 0 | 0 | 0 | 0 | 0 |
| 1240 | ychF | 0 | 0 | 0 | 0 | 0 | 0 | 0 | 0 | 0 | 0 | 0 | 0 |
| 1241 | ptsA | 0 | 0 | 0 | 0 | 0 | 0 | 0 | 0 | 0 | 0 | 0 | 0 |
| 1242 | ychM | 0 | 3 | 1 | 2 | 0 | 0 | 0 | 1 | 1 | 0 | 0 | 0 |
| 1243 | ychH | 1410 | 161702 | 47778 | 149215 | 7413 | 106010 | 2906 | 122936 | 130660 | 127138 | 12014 | 78713 |
| 1244 | ptsG | 0 | 4 | 2 | 3 | 0 | 1 | 1 | 1 | 2 | 3 | 1 | 0 |
| 1245 | ymdB | 0 | 0 | 0 | 0 | 0 | 0 | 0 | 0 | 0 | 0 | 0 | 0 |
| 1246 | dppA | 0 | 0 | 0 | 0 | 0 | 0 | 0 | 1 | 0 | 0 | 0 | 0 |
| 1247 | dppB | 0 | 0 | 0 | 0 | 0 | 0 | 0 | 0 | 0 | 0 | 0 | 0 |
| 1248 | uhpT | 0 | 0 | 0 | 0 | 0 | 0 | 0 | 0 | 0 | 0 | 0 | 0 |
| 1249 | fhuF | 0 | 0 | 0 | 0 | 0 | 0 | 0 | 0 | 0 | 0 | 0 | 0 |
| 1250 | fhuA | 0 | 0 | 0 | 0 | 0 | 0 | 0 | 0 | 0 | 0 | 1 | 0 |
| 1251 | fhuC | 0 | 0 | 0 | 0 | 0 | 0 | 0 | 0 | 0 | 1 | 0 | 0 |
| 1252 | uhpA | 0 | 0 | 0 | 0 | 0 | 0 | 0 | 1 | 0 | 0 | 0 | 0 |
| 1253 | nuoM | 0 | 0 | 0 | 0 | 0 | 0 | 0 | 0 | 0 | 0 | 0 | 0 |
| 1254 | proX | 0 | 0 | 0 | 0 | 0 | 0 | 0 | 0 | 1 | 0 | 0 | 0 |
| 1255 | proS | 0 | 0 | 0 | 0 | 0 | 0 | 0 | 0 | 0 | 0 | 0 | 0 |
| 1256 | proQ | 0 | 0 | 0 | 0 | 0 | 0 | 0 | 1 | 0 | 0 | 0 | 0 |
| 1257 | proL | 0 | 0 | 0 | 0 | 0 | 0 | 0 | 0 | 0 | 0 | 0 | 0 |
| 1258 | phnC | 0 | 0 | 0 | 0 | 0 | 0 | 0 | 0 | 0 | 0 | 0 | 0 |
| 1259 | luxS | 0 | 0 | 0 | 0 | 0 | 0 | 0 | 0 | 0 | 0 | 0 | 0 |

| | | | | | | | | | | | | | |
|---|---|---|---|---|---|---|---|---|---|---|---|---|---|
| 1260 | phnL | 0 | 0 | 0 | 0 | 0 | 0 | 1 | 0 | 0 | 0 | 0 | 0 |
| 1261 | proB | 0 | 0 | 1 | 0 | 0 | 0 | 0 | 1 | 0 | 0 | 0 | 0 |
| 1262 | lipA | 0 | 0 | 0 | 0 | 0 | 0 | 0 | 0 | 0 | 0 | 0 | 0 |
| 1263 | lipB | 0 | 0 | 0 | 0 | 0 | 0 | 0 | 0 | 0 | 0 | 0 | 0 |
| 1264 | bacA | 0 | 0 | 0 | 0 | 0 | 0 | 0 | 0 | 0 | 0 | 0 | 0 |
| 1265 | somA_macA | 0 | 0 | 0 | 0 | 0 | 0 | 0 | 0 | 0 | 0 | 0 | 0 |
| 1266 | bcsE | 0 | 0 | 0 | 0 | 0 | 0 | 0 | 0 | 0 | 0 | 0 | 0 |
| 1267 | b3044 | 6 | 0 | 7 | 1 | 0 | 0 | 9 | 1 | 0 | 0 | 5 | 0 |
| 1268 | yqeG | 0 | 0 | 0 | 0 | 0 | 0 | 0 | 0 | 0 | 0 | 0 | 0 |
| 1269 | yehD | 0 | 0 | 0 | 0 | 0 | 0 | 0 | 0 | 0 | 0 | 0 | 0 |
| 1270 | yifE | 0 | 0 | 0 | 0 | 0 | 0 | 0 | 0 | 0 | 0 | 0 | 0 |
| 1271 | yobF | 0 | 0 | 0 | 0 | 0 | 0 | 0 | 0 | 0 | 0 | 0 | 0 |
| 1272 | b2641 | 0 | 0 | 0 | 0 | 0 | 0 | 0 | 0 | 0 | 0 | 0 | 0 |
| 1273 | yqeI | 0 | 0 | 0 | 0 | 0 | 0 | 0 | 0 | 0 | 0 | 0 | 0 |
| 1274 | yqeH | 0 | 0 | 0 | 0 | 0 | 0 | 0 | 0 | 0 | 0 | 0 | 0 |
| 1275 | yqeK | 0 | 0 | 0 | 0 | 0 | 0 | 0 | 0 | 0 | 0 | 0 | 0 |
| 1276 | yehS | 0 | 0 | 0 | 0 | 0 | 0 | 0 | 0 | 0 | 0 | 0 | 0 |
| 1277 | yhfG | 0 | 0 | 0 | 0 | 0 | 0 | 0 | 0 | 0 | 0 | 0 | 0 |
| 1278 | yehZ | 0 | 1 | 0 | 0 | 0 | 0 | 0 | 0 | 0 | 0 | 0 | 0 |
| 1279 | bisC | 0 | 0 | 0 | 0 | 0 | 0 | 0 | 0 | 0 | 0 | 0 | 0 |
| 1280 | pbpG | 0 | 0 | 0 | 0 | 0 | 0 | 0 | 0 | 0 | 0 | 0 | 0 |
| 1281 | idi | 0 | 0 | 0 | 1 | 0 | 0 | 0 | 0 | 0 | 0 | 0 | 0 |
| 1282 | bioB | 0 | 0 | 0 | 0 | 0 | 0 | 0 | 0 | 0 | 0 | 1 | 0 |
| 1283 | b0332 | 0 | 0 | 0 | 0 | 0 | 0 | 0 | 0 | 0 | 0 | 0 | 0 |
| 1284 | pabC | 517 | 2 | 389 | 1 | 4 | 0 | 401 | 2 | 138 | 6 | 266 | 1 |
| 1285 | yeeO | 0 | 0 | 0 | 0 | 0 | 0 | 0 | 0 | 0 | 0 | 0 | 0 |
| 1286 | intF | 0 | 0 | 0 | 0 | 0 | 1 | 0 | 0 | 0 | 0 | 0 | 0 |
| 1287 | intA | 0 | 0 | 0 | 0 | 0 | 0 | 0 | 0 | 0 | 0 | 0 | 0 |
| 1288 | ybgA | 0 | 0 | 0 | 1 | 0 | 0 | 0 | 0 | 0 | 0 | 0 | 0 |
| 1289 | yfdQ | 0 | 0 | 0 | 0 | 0 | 0 | 0 | 0 | 0 | 0 | 0 | 0 |
| 1290 | intS | 0 | 0 | 0 | 0 | 0 | 0 | 0 | 0 | 0 | 0 | 0 | 0 |
| 1291 | intZ | 0 | 0 | 0 | 0 | 0 | 0 | 0 | 0 | 0 | 0 | 0 | 0 |
| 1292 | infA | 0 | 0 | 0 | 0 | 0 | 0 | 0 | 0 | 0 | 0 | 0 | 0 |
| 1293 | rlmB | 0 | 0 | 0 | 1 | 0 | 0 | 0 | 0 | 0 | 0 | 0 | 0 |
| 1294 | ihfB | 0 | 0 | 0 | 0 | 0 | 0 | 0 | 0 | 0 | 0 | 0 | 0 |
| 1295 | atoS | 0 | 0 | 0 | 0 | 0 | 0 | 0 | 0 | 0 | 0 | 0 | 0 |
| 1296 | yhcG | 0 | 0 | 0 | 0 | 0 | 0 | 0 | 0 | 0 | 0 | 0 | 0 |
| 1297 | exuR | 0 | 0 | 0 | 0 | 0 | 0 | 0 | 0 | 0 | 0 | 0 | 0 |
| 1298 | tufB | 0 | 0 | 0 | 1 | 0 | 1 | 0 | 0 | 0 | 0 | 1 | 0 |
| 1299 | tufA | 0 | 0 | 0 | 0 | 0 | 0 | 0 | 0 | 0 | 0 | 0 | 0 |
| 1300 | ycgY | 0 | 0 | 0 | 0 | 0 | 0 | 0 | 0 | 0 | 0 | 0 | 0 |
| 1301 | chaC | 0 | 0 | 0 | 0 | 0 | 0 | 0 | 0 | 0 | 0 | 0 | 0 |
| 1302 | yagM | 0 | 0 | 0 | 0 | 0 | 0 | 0 | 0 | 0 | 0 | 0 | 0 |
| 1303 | yagI | 0 | 0 | 0 | 0 | 0 | 0 | 0 | 0 | 0 | 0 | 0 | 0 |
| 1304 | yagG | 0 | 0 | 0 | 0 | 0 | 0 | 0 | 0 | 0 | 0 | 0 | 0 |
| 1305 | yagE | 0 | 0 | 0 | 0 | 0 | 0 | 0 | 0 | 0 | 0 | 0 | 0 |
| 1306 | ycgR | 0 | 0 | 0 | 0 | 0 | 0 | 0 | 0 | 0 | 0 | 0 | 0 |
| 1307 | yagB | 0 | 0 | 0 | 0 | 0 | 0 | 0 | 0 | 0 | 0 | 0 | 0 |
| 1308 | ycgI | 0 | 0 | 0 | 0 | 0 | 0 | 0 | 0 | 0 | 0 | 0 | 0 |
| 1309 | ycgH | 0 | 0 | 0 | 0 | 0 | 0 | 0 | 0 | 0 | 0 | 0 | 0 |
| 1310 | ycgK | 0 | 0 | 0 | 0 | 0 | 0 | 0 | 0 | 0 | 0 | 0 | 0 |
| 1311 | ycgJ | 0 | 0 | 0 | 0 | 0 | 0 | 0 | 0 | 0 | 0 | 0 | 0 |
| 1312 | ycgM | 0 | 0 | 0 | 0 | 0 | 0 | 0 | 0 | 0 | 0 | 0 | 0 |
| 1313 | yagZ | 0 | 0 | 0 | 0 | 0 | 0 | 0 | 0 | 1 | 0 | 0 | 0 |
| 1314 | yagY | 0 | 0 | 0 | 0 | 0 | 0 | 0 | 0 | 1 | 1 | 0 | 0 |
| 1315 | ycgN | 0 | 0 | 0 | 0 | 0 | 0 | 0 | 0 | 0 | 0 | 0 | 0 |
| 1316 | yagU | 0 | 0 | 0 | 0 | 1 | 0 | 1 | 0 | 0 | 0 | 0 | 0 |
| 1317 | yagT | 0 | 0 | 0 | 0 | 0 | 0 | 0 | 0 | 0 | 0 | 0 | 0 |
| 1318 | fldB | 0 | 1 | 0 | 1 | 0 | 1 | 0 | 0 | 2 | 0 | 0 | 0 |
| 1319 | ycgF | 0 | 0 | 0 | 0 | 0 | 0 | 0 | 0 | 0 | 0 | 0 | 0 |
| 1320 | yhbP | 0 | 0 | 1 | 0 | 0 | 0 | 0 | 0 | 0 | 0 | 0 | 0 |
| 1321 | pyrH | 0 | 0 | 0 | 0 | 0 | 0 | 0 | 0 | 0 | 0 | 0 | 0 |
| 1322 | pyrG | 0 | 0 | 0 | 0 | 0 | 0 | 0 | 0 | 0 | 0 | 0 | 0 |
| 1323 | pyrC | 0 | 0 | 0 | 0 | 0 | 0 | 0 | 0 | 0 | 0 | 0 | 0 |
| 1324 | pyrB | 0 | 0 | 0 | 0 | 0 | 0 | 0 | 0 | 0 | 0 | 0 | 0 |
| 1325 | glpX | 0 | 0 | 0 | 0 | 0 | 0 | 0 | 0 | 0 | 0 | 0 | 0 |
| 1326 | glpE | 0 | 0 | 0 | 0 | 0 | 0 | 0 | 0 | 0 | 0 | 0 | 0 |
| 1327 | glpD | 0 | 0 | 0 | 0 | 0 | 0 | 0 | 0 | 1 | 0 | 0 | 0 |
| 1328 | glpA | 0 | 0 | 0 | 0 | 0 | 0 | 0 | 0 | 0 | 0 | 0 | 0 |
| 1329 | yciW | 0 | 0 | 0 | 0 | 0 | 0 | 0 | 0 | 0 | 0 | 0 | 0 |
| 1330 | fxsA | 0 | 0 | 0 | 0 | 0 | 0 | 0 | 0 | 0 | 0 | 0 | 0 |
| 1331 | yciT | 0 | 0 | 0 | 0 | 0 | 0 | 0 | 0 | 0 | 0 | 0 | 0 |
| 1332 | yciS | 0 | 0 | 0 | 0 | 0 | 0 | 0 | 0 | 0 | 0 | 0 | 0 |
| 1333 | edd | 7 | 1 | 5 | 1 | 3 | 2 | 5 | 2 | 1 | 1 | 4 | 2 |
| 1334 | yciG | 0 | 0 | 0 | 0 | 0 | 0 | 0 | 0 | 0 | 0 | 0 | 0 |
| 1335 | b1437 | 0 | 0 | 0 | 0 | 0 | 0 | 0 | 0 | 0 | 0 | 0 | 0 |
| 1336 | yciC | 0 | 0 | 0 | 0 | 0 | 0 | 0 | 0 | 0 | 0 | 0 | 0 |
| 1337 | yciA | 0 | 0 | 0 | 0 | 0 | 0 | 0 | 0 | 0 | 0 | 0 | 0 |
| 1338 | phoH | 0 | 1 | 0 | 1 | 0 | 0 | 0 | 0 | 0 | 1 | 0 | 0 |
| 1339 | ygcU | 0 | 0 | 0 | 0 | 0 | 0 | 0 | 0 | 0 | 0 | 0 | 1 |
| 1340 | yciK | 0 | 0 | 0 | 0 | 0 | 0 | 0 | 0 | 0 | 0 | 0 | 0 |
| 1341 | yciI | 0 | 0 | 0 | 0 | 0 | 0 | 0 | 0 | 0 | 0 | 0 | 0 |
| 1342 | yoaF | 0 | 0 | 0 | 0 | 0 | 0 | 0 | 0 | 0 | 0 | 0 | 0 |
| 1343 | mdtH | 0 | 0 | 0 | 0 | 0 | 0 | 0 | 0 | 0 | 0 | 0 | 0 |

### Data Set S2 - MG1655 library 7 day experiment
#### Aligned Reads

| | | | | | | | | | | | | | |
|---|---|---|---|---|---|---|---|---|---|---|---|---|---|
| 1344 | codB | 0 | 0 | 0 | 0 | 0 | 0 | 0 | 0 | 0 | 0 | 0 | 0 |
| 1345 | ygcW | 0 | 0 | 0 | 0 | 0 | 0 | 0 | 0 | 0 | 0 | 0 | 0 |
| 1346 | yihR | 0 | 0 | 0 | 0 | 0 | 0 | 0 | 0 | 0 | 0 | 0 | 0 |
| 1347 | zupT | 0 | 0 | 0 | 0 | 0 | 0 | 0 | 0 | 0 | 0 | 0 | 0 |
| 1348 | katE | 0 | 0 | 0 | 0 | 0 | 0 | 0 | 0 | 0 | 0 | 0 | 0 |
| 1349 | katG | 0 | 0 | 0 | 0 | 0 | 0 | 0 | 0 | 0 | 0 | 0 | 0 |
| 1350 | racR | 0 | 0 | 0 | 0 | 0 | 0 | 0 | 0 | 0 | 0 | 0 | 0 |
| 1351 | acnB | 0 | 0 | 0 | 0 | 0 | 0 | 0 | 0 | 0 | 0 | 0 | 0 |
| 1352 | kch | 0 | 0 | 0 | 0 | 0 | 0 | 0 | 0 | 0 | 0 | 0 | 0 |
| 1353 | b0259 | 0 | 0 | 0 | 0 | 0 | 0 | 0 | 0 | 0 | 0 | 0 | 0 |
| 1354 | racC | 0 | 0 | 0 | 0 | 0 | 0 | 0 | 0 | 0 | 0 | 0 | 0 |
| 1355 | b0257 | 0 | 0 | 0 | 0 | 0 | 0 | 0 | 0 | 0 | 0 | 0 | 0 |
| 1356 | b0255 | 0 | 0 | 0 | 0 | 0 | 0 | 0 | 0 | 0 | 0 | 0 | 0 |
| 1357 | yhcO | 0 | 0 | 0 | 0 | 0 | 0 | 0 | 0 | 0 | 0 | 0 | 0 |
| 1358 | yhcM | 0 | 0 | 0 | 0 | 0 | 0 | 0 | 0 | 0 | 0 | 0 | 0 |
| 1359 | yhcB | 0 | 0 | 0 | 0 | 0 | 0 | 0 | 0 | 0 | 0 | 0 | 0 |
| 1360 | yhcA | 0 | 0 | 0 | 0 | 0 | 0 | 0 | 0 | 0 | 0 | 0 | 0 |
| 1361 | yhjX | 0 | 0 | 0 | 0 | 0 | 0 | 0 | 0 | 0 | 0 | 0 | 0 |
| 1362 | yeiM | 0 | 0 | 1 | 1 | 0 | 0 | 0 | 0 | 1 | 0 | 0 | 0 |
| 1363 | yeiE | 0 | 0 | 0 | 0 | 0 | 0 | 0 | 0 | 0 | 0 | 0 | 0 |
| 1364 | yeiG | 0 | 0 | 0 | 0 | 0 | 0 | 0 | 0 | 0 | 0 | 0 | 0 |
| 1365 | sapA | 0 | 0 | 0 | 0 | 0 | 0 | 0 | 0 | 0 | 0 | 0 | 0 |
| 1366 | yeiC | 0 | 0 | 0 | 0 | 0 | 0 | 0 | 0 | 0 | 0 | 0 | 0 |
| 1367 | nhaA | 0 | 0 | 0 | 0 | 0 | 0 | 0 | 0 | 0 | 0 | 0 | 0 |
| 1368 | yeiT | 0 | 0 | 0 | 0 | 0 | 0 | 0 | 0 | 0 | 0 | 0 | 0 |
| 1369 | yeiP | 0 | 0 | 0 | 0 | 0 | 0 | 0 | 0 | 0 | 0 | 0 | 0 |
| 1370 | nfrB | 0 | 0 | 0 | 0 | 0 | 0 | 0 | 0 | 0 | 1 | 0 | 0 |
| 1371 | folX | 0 | 0 | 0 | 0 | 0 | 0 | 0 | 0 | 0 | 0 | 0 | 0 |
| 1372 | folE | 0 | 0 | 0 | 0 | 0 | 0 | 0 | 0 | 0 | 0 | 0 | 0 |
| 1373 | folD | 0 | 0 | 0 | 0 | 0 | 0 | 0 | 0 | 0 | 0 | 0 | 0 |
| 1374 | folA | 0 | 0 | 0 | 0 | 0 | 0 | 0 | 0 | 0 | 0 | 0 | 0 |
| 1375 | folC | 0 | 0 | 0 | 0 | 1 | 0 | 0 | 0 | 0 | 0 | 1 | 0 |
| 1376 | folB | 0 | 0 | 0 | 0 | 0 | 0 | 0 | 0 | 0 | 0 | 0 | 0 |
| 1377 | hydN | 0 | 0 | 0 | 0 | 0 | 0 | 0 | 0 | 0 | 0 | 0 | 0 |
| 1378 | rpoS | 0 | 0 | 0 | 0 | 0 | 0 | 0 | 0 | 0 | 0 | 0 | 0 |
| 1379 | kdtA | 0 | 0 | 0 | 0 | 0 | 0 | 0 | 0 | 0 | 0 | 0 | 0 |
| 1380 | rpoE | 0 | 0 | 0 | 0 | 0 | 0 | 0 | 0 | 0 | 0 | 0 | 0 |
| 1381 | rpoD | 1 | 0 | 0 | 1 | 0 | 0 | 0 | 0 | 0 | 0 | 1 | 0 |
| 1382 | rpoH | 0 | 0 | 0 | 0 | 0 | 0 | 0 | 0 | 0 | 0 | 0 | 0 |
| 1383 | yjaB | 537 | 2 | 393 | 1 | 11 | 0 | 530 | 1 | 238 | 0 | 82 | 0 |
| 1384 | yjaH | 0 | 0 | 0 | 0 | 0 | 0 | 0 | 0 | 0 | 0 | 0 | 0 |
| 1385 | nohB | 0 | 0 | 0 | 0 | 0 | 0 | 0 | 0 | 0 | 0 | 0 | 0 |
| 1386 | rpsG | 0 | 0 | 0 | 0 | 0 | 0 | 0 | 0 | 0 | 0 | 0 | 0 |
| 1387 | rpsB | 0 | 0 | 0 | 0 | 0 | 0 | 0 | 0 | 0 | 0 | 0 | 0 |
| 1388 | rpsA | 0 | 0 | 0 | 0 | 0 | 0 | 0 | 0 | 0 | 0 | 0 | 0 |
| 1389 | fdnG | 0 | 0 | 0 | 0 | 0 | 0 | 0 | 0 | 0 | 0 | 0 | 0 |
| 1390 | rpsO | 0 | 0 | 1 | 0 | 0 | 0 | 0 | 0 | 0 | 0 | 0 | 0 |
| 1391 | rpsM | 0 | 0 | 0 | 0 | 0 | 0 | 0 | 0 | 0 | 0 | 0 | 0 |
| 1392 | rpsL | 0 | 0 | 0 | 0 | 0 | 0 | 0 | 0 | 1 | 0 | 0 | 0 |
| 1393 | rpsJ | 0 | 0 | 0 | 0 | 0 | 0 | 0 | 0 | 0 | 0 | 0 | 0 |
| 1394 | apt | 1 | 0 | 0 | 0 | 1 | 0 | 0 | 0 | 0 | 0 | 1 | 0 |
| 1395 | rpsU | 0 | 0 | 0 | 0 | 0 | 0 | 0 | 0 | 0 | 0 | 0 | 0 |
| 1396 | rpsT | 0 | 0 | 0 | 0 | 0 | 0 | 0 | 0 | 0 | 0 | 0 | 0 |
| 1397 | rpsP | 0 | 0 | 0 | 0 | 0 | 0 | 0 | 0 | 0 | 0 | 0 | 0 |
| 1398 | deoA | 0 | 0 | 0 | 0 | 0 | 0 | 0 | 0 | 0 | 0 | 0 | 0 |
| 1399 | deoB | 0 | 0 | 0 | 0 | 0 | 0 | 0 | 0 | 0 | 0 | 0 | 0 |
| 1400 | deoC | 0 | 0 | 0 | 0 | 0 | 0 | 0 | 0 | 0 | 0 | 0 | 0 |
| 1401 | deoD | 0 | 0 | 0 | 0 | 0 | 0 | 0 | 0 | 0 | 0 | 0 | 0 |
| 1402 | usg | 0 | 0 | 0 | 0 | 0 | 0 | 0 | 0 | 0 | 0 | 0 | 0 |
| 1403 | sirA | 6 | 0 | 1 | 0 | 0 | 0 | 3 | 0 | 1 | 0 | 0 | 1 |
| 1404 | fes | 0 | 0 | 0 | 0 | 0 | 0 | 0 | 0 | 0 | 0 | 0 | 0 |
| 1405 | xasA | 0 | 0 | 0 | 0 | 0 | 0 | 0 | 0 | 0 | 0 | 0 | 0 |
| 1406 | pheL | 0 | 0 | 0 | 0 | 0 | 0 | 0 | 0 | 0 | 0 | 0 | 0 |
| 1407 | phoP | 0 | 0 | 0 | 0 | 0 | 0 | 0 | 0 | 0 | 0 | 0 | 0 |
| 1408 | phoR | 0 | 0 | 0 | 0 | 0 | 0 | 0 | 0 | 0 | 0 | 0 | 0 |
| 1409 | ndh | 0 | 0 | 0 | 0 | 0 | 0 | 0 | 0 | 0 | 0 | 0 | 1 |
| 1410 | yfhK | 0 | 0 | 0 | 0 | 0 | 0 | 0 | 0 | 1 | 0 | 0 | 0 |
| 1411 | phoA | 0 | 0 | 0 | 0 | 0 | 0 | 0 | 0 | 0 | 0 | 0 | 0 |
| 1412 | thiI | 0 | 0 | 0 | 0 | 0 | 0 | 0 | 0 | 0 | 0 | 0 | 0 |
| 1413 | phoB | 0 | 0 | 0 | 0 | 0 | 0 | 0 | 0 | 0 | 0 | 0 | 0 |
| 1414 | pinE | 0 | 0 | 0 | 0 | 0 | 0 | 0 | 0 | 0 | 0 | 0 | 0 |
| 1415 | thiM | 0 | 0 | 0 | 0 | 0 | 0 | 0 | 0 | 0 | 0 | 0 | 0 |
| 1416 | thiL | 0 | 0 | 0 | 0 | 0 | 0 | 0 | 0 | 0 | 0 | 0 | 0 |
| 1417 | thiC | 0 | 0 | 0 | 0 | 0 | 0 | 0 | 0 | 0 | 1 | 0 | 0 |
| 1418 | prlC | 0 | 0 | 0 | 0 | 0 | 0 | 0 | 0 | 0 | 0 | 0 | 0 |
| 1419 | thiG | 0 | 0 | 0 | 0 | 0 | 0 | 0 | 0 | 0 | 0 | 0 | 0 |
| 1420 | yiaU | 0 | 0 | 0 | 0 | 0 | 0 | 0 | 0 | 0 | 0 | 0 | 0 |
| 1421 | yiaT | 0 | 0 | 0 | 0 | 0 | 0 | 0 | 0 | 0 | 0 | 0 | 0 |
| 1422 | yiaW | 0 | 0 | 0 | 0 | 0 | 0 | 0 | 0 | 0 | 0 | 0 | 0 |
| 1423 | fabF | 670 | 4 | 430 | 6 | 10 | 0 | 652 | 1 | 188 | 0 | 117 | 1 |
| 1424 | yiaD | 0 | 0 | 0 | 0 | 0 | 0 | 0 | 0 | 0 | 0 | 0 | 0 |
| 1425 | yiaG | 0 | 0 | 0 | 0 | 0 | 0 | 0 | 0 | 0 | 0 | 0 | 0 |
| 1426 | yiaF | 0 | 0 | 0 | 0 | 0 | 0 | 0 | 0 | 0 | 0 | 0 | 0 |
| 1427 | yiaJ | 0 | 0 | 0 | 0 | 0 | 0 | 0 | 0 | 0 | 0 | 0 | 0 |

| | | | | | | | | | | | | | |
|---|---|---|---|---|---|---|---|---|---|---|---|---|---|
| 1428 | yqjH | 0 | 0 | 0 | 0 | 0 | 0 | 0 | 0 | 0 | 0 | 0 | 0 |
| 1429 | narX | 0 | 0 | 0 | 0 | 0 | 0 | 0 | 0 | 0 | 0 | 0 | 0 |
| 1430 | tar | 0 | 0 | 0 | 0 | 0 | 0 | 0 | 0 | 0 | 0 | 0 | 0 |
| 1431 | narZ | 0 | 0 | 0 | 0 | 0 | 0 | 0 | 0 | 0 | 0 | 0 | 0 |
| 1432 | narU | 0 | 0 | 0 | 1 | 0 | 0 | 0 | 0 | 0 | 0 | 0 | 0 |
| 1433 | cbdA | 0 | 0 | 0 | 0 | 0 | 0 | 0 | 0 | 0 | 0 | 0 | 0 |
| 1434 | narQ | 0 | 0 | 0 | 0 | 0 | 0 | 0 | 0 | 0 | 0 | 0 | 0 |
| 1435 | tam | 0 | 0 | 0 | 0 | 0 | 0 | 0 | 0 | 0 | 0 | 0 | 0 |
| 1436 | gcl | 0 | 0 | 0 | 0 | 0 | 0 | 1 | 0 | 0 | 0 | 0 | 0 |
| 1437 | yneF | 0 | 0 | 0 | 0 | 0 | 0 | 0 | 0 | 0 | 0 | 0 | 0 |
| 1438 | paaX | 0 | 0 | 0 | 0 | 0 | 0 | 0 | 0 | 0 | 0 | 0 | 0 |
| 1439 | yrbL | 0 | 0 | 0 | 0 | 0 | 0 | 0 | 0 | 0 | 0 | 0 | 0 |
| 1440 | yrbG | 0 | 0 | 0 | 0 | 0 | 0 | 0 | 0 | 0 | 0 | 0 | 0 |
| 1441 | yrbF | 0 | 0 | 0 | 0 | 0 | 0 | 0 | 0 | 0 | 0 | 0 | 0 |
| 1442 | yrbA | 0 | 0 | 0 | 0 | 0 | 0 | 0 | 0 | 0 | 0 | 0 | 0 |
| 1443 | kbl | 0 | 0 | 0 | 0 | 0 | 0 | 0 | 0 | 0 | 0 | 0 | 0 |
| 1444 | dmsA | 0 | 0 | 0 | 0 | 0 | 0 | 0 | 0 | 1 | 0 | 0 | 0 |
| 1445 | ruvA | 0 | 0 | 0 | 0 | 0 | 0 | 0 | 0 | 0 | 0 | 0 | 0 |
| 1446 | tsf | 0 | 0 | 0 | 0 | 0 | 0 | 0 | 0 | 0 | 0 | 0 | 0 |
| 1447 | mutH | 0 | 0 | 0 | 0 | 0 | 0 | 0 | 0 | 0 | 0 | 0 | 0 |
| 1448 | ynjF | 0 | 0 | 0 | 0 | 0 | 0 | 0 | 0 | 0 | 0 | 0 | 0 |
| 1449 | mutM | 0 | 0 | 0 | 0 | 0 | 1 | 0 | 0 | 0 | 0 | 0 | 0 |
| 1450 | mutS | 256 | 13 | 227 | 4 | 3 | 0 | 320 | 347 | 113 | 0 | 20 | 8 |
| 1451 | mutT | 0 | 0 | 0 | 0 | 0 | 0 | 0 | 0 | 0 | 0 | 0 | 0 |
| 1452 | mutY | 0 | 0 | 0 | 0 | 0 | 0 | 0 | 0 | 0 | 0 | 0 | 0 |
| 1453 | ydjX | 0 | 0 | 0 | 0 | 0 | 0 | 0 | 0 | 0 | 0 | 0 | 0 |
| 1454 | ydjA | 0 | 0 | 0 | 0 | 0 | 0 | 0 | 0 | 0 | 0 | 0 | 0 |
| 1455 | ydjE | 0 | 0 | 0 | 0 | 0 | 0 | 0 | 0 | 0 | 0 | 0 | 0 |
| 1456 | b1420 | 0 | 0 | 0 | 0 | 0 | 0 | 1 | 0 | 0 | 0 | 0 | 0 |
| 1457 | ydjF | 0 | 0 | 0 | 0 | 0 | 0 | 0 | 0 | 0 | 0 | 0 | 0 |
| 1458 | ydjL | 580 | 0 | 767 | 3180 | 8 | 29 | 671 | 41082 | 269 | 124 | 119 | 2098 |
| 1459 | ydjN | 0 | 0 | 0 | 0 | 0 | 0 | 0 | 0 | 0 | 0 | 0 | 0 |
| 1460 | nadK | 0 | 0 | 0 | 0 | 0 | 0 | 0 | 0 | 0 | 0 | 0 | 0 |
| 1461 | yadK | 0 | 0 | 0 | 0 | 0 | 0 | 0 | 0 | 0 | 0 | 0 | 0 |
| 1462 | cusR | 0 | 0 | 0 | 0 | 0 | 0 | 0 | 0 | 0 | 0 | 0 | 0 |
| 1463 | yadN | 0 | 0 | 0 | 0 | 0 | 0 | 0 | 0 | 0 | 0 | 0 | 0 |
| 1464 | nfo | 0 | 0 | 0 | 0 | 0 | 0 | 0 | 0 | 0 | 0 | 0 | 0 |
| 1465 | yadC | 0 | 0 | 0 | 0 | 0 | 0 | 0 | 0 | 0 | 0 | 0 | 0 |
| 1466 | yfjI | 0 | 0 | 0 | 0 | 0 | 0 | 0 | 0 | 0 | 0 | 0 | 0 |
| 1467 | yadF | 0 | 0 | 0 | 0 | 0 | 0 | 0 | 0 | 0 | 0 | 0 | 0 |
| 1468 | yfjN | 1547 | 11 | 1272 | 5 | 444 | 8 | 1142 | 2 | 889 | 0 | 897 | 0 |
| 1469 | yfjM | 0 | 0 | 0 | 0 | 0 | 0 | 0 | 0 | 0 | 0 | 0 | 0 |
| 1470 | yfjL | 0 | 0 | 0 | 0 | 0 | 0 | 0 | 0 | 0 | 0 | 0 | 0 |
| 1471 | yfjR | 798 | 0 | 495 | 0 | 17 | 0 | 723 | 0 | 174 | 0 | 1 | 0 |
| 1472 | cusC | 1 | 0 | 0 | 0 | 0 | 0 | 0 | 0 | 0 | 0 | 0 | 0 |
| 1473 | yfjP | 0 | 0 | 0 | 0 | 0 | 0 | 0 | 0 | 0 | 0 | 0 | 0 |
| 1474 | yfjW | 0 | 0 | 0 | 0 | 0 | 0 | 0 | 0 | 0 | 0 | 0 | 0 |
| 1475 | yadR | 0 | 1 | 0 | 0 | 0 | 0 | 0 | 0 | 0 | 0 | 0 | 0 |
| 1476 | yadS | 0 | 0 | 0 | 0 | 0 | 0 | 0 | 0 | 0 | 0 | 0 | 0 |
| 1477 | psiE | 0 | 0 | 0 | 0 | 0 | 0 | 1 | 1 | 0 | 0 | 0 | 0 |
| 1478 | psiF | 0 | 0 | 0 | 0 | 0 | 0 | 0 | 0 | 0 | 0 | 0 | 0 |
| 1479 | rplY | 0 | 0 | 1 | 0 | 0 | 0 | 0 | 0 | 0 | 0 | 0 | 0 |
| 1480 | rplT | 0 | 0 | 0 | 0 | 0 | 0 | 0 | 0 | 0 | 0 | 0 | 0 |
| 1481 | hlyE | 0 | 0 | 0 | 0 | 0 | 0 | 0 | 0 | 0 | 0 | 0 | 0 |
| 1482 | rplN | 0 | 0 | 1 | 0 | 0 | 0 | 1 | 0 | 0 | 0 | 0 | 0 |
| 1483 | kdsB | 0 | 0 | 0 | 0 | 1 | 0 | 0 | 0 | 0 | 0 | 0 | 0 |
| 1484 | metZ | 0 | 0 | 0 | 0 | 0 | 0 | 0 | 0 | 0 | 0 | 0 | 0 |
| 1485 | metB | 0 | 0 | 0 | 0 | 0 | 0 | 0 | 0 | 0 | 0 | 0 | 0 |
| 1486 | metC | 0 | 0 | 0 | 0 | 0 | 1 | 0 | 0 | 0 | 0 | 0 | 0 |
| 1487 | mrcB | 0 | 0 | 0 | 0 | 0 | 0 | 0 | 0 | 0 | 0 | 1 | 0 |
| 1488 | metA | 1 | 0 | 0 | 0 | 0 | 0 | 0 | 0 | 0 | 0 | 0 | 0 |
| 1489 | metF | 0 | 0 | 0 | 0 | 0 | 0 | 0 | 0 | 0 | 0 | 0 | 0 |
| 1490 | metG | 0 | 0 | 0 | 0 | 0 | 0 | 0 | 0 | 0 | 0 | 0 | 0 |
| 1491 | metJ | 0 | 0 | 0 | 0 | 0 | 0 | 0 | 0 | 0 | 0 | 0 | 0 |
| 1492 | metK | 110 | 0 | 109 | 0 | 1 | 0 | 109 | 1 | 47 | 0 | 1 | 0 |
| 1493 | yhbE | 0 | 1 | 0 | 0 | 0 | 0 | 0 | 0 | 0 | 0 | 0 | 0 |
| 1494 | ycjW | 0 | 0 | 0 | 0 | 0 | 0 | 0 | 0 | 0 | 0 | 0 | 0 |
| 1495 | yhbC | 0 | 0 | 0 | 0 | 0 | 0 | 0 | 0 | 0 | 0 | 1 | 0 |
| 1496 | ycjZ | 0 | 0 | 0 | 0 | 0 | 0 | 0 | 0 | 0 | 0 | 0 | 0 |
| 1497 | ycjX | 0 | 0 | 0 | 0 | 0 | 0 | 0 | 0 | 0 | 0 | 0 | 0 |
| 1498 | ycjY | 0 | 0 | 0 | 0 | 0 | 0 | 0 | 0 | 0 | 0 | 0 | 0 |
| 1499 | yhbU | 0 | 0 | 0 | 0 | 0 | 0 | 0 | 0 | 0 | 0 | 0 | 0 |
| 1500 | yhbT | 0 | 0 | 0 | 0 | 0 | 0 | 0 | 0 | 0 | 0 | 0 | 0 |
| 1501 | yhbW | 826 | 4 | 572 | 1 | 5 | 0 | 697 | 0 | 232 | 1 | 37 | 0 |
| 1502 | yhbQ | 1478 | 37 | 1344 | 11 | 157 | 218 | 1073 | 17 | 865 | 33 | 2330 | 2024 |
| 1503 | ycjG | 0 | 0 | 0 | 0 | 0 | 0 | 0 | 0 | 0 | 0 | 0 | 1 |
| 1504 | ycjK | 0 | 0 | 0 | 0 | 0 | 0 | 0 | 0 | 0 | 0 | 0 | 0 |
| 1505 | yhbX | 0 | 0 | 0 | 0 | 0 | 0 | 0 | 0 | 0 | 0 | 0 | 0 |
| 1506 | ycjL | 0 | 0 | 0 | 0 | 0 | 0 | 0 | 0 | 0 | 0 | 0 | 0 |
| 1507 | ycjM | 0 | 0 | 0 | 0 | 0 | 0 | 0 | 0 | 0 | 0 | 0 | 0 |
| 1508 | ymjA | 0 | 0 | 0 | 0 | 0 | 0 | 0 | 0 | 0 | 0 | 0 | 0 |
| 1509 | rhoL | 0 | 0 | 0 | 0 | 0 | 0 | 0 | 0 | 0 | 0 | 0 | 0 |
| 1510 | nrfE | 0 | 0 | 0 | 0 | 0 | 0 | 0 | 0 | 0 | 0 | 0 | 0 |
| 1511 | pgk | 0 | 0 | 0 | 1 | 0 | 0 | 0 | 0 | 1 | 0 | 0 | 0 |

Data Set S2 - MG1655 library 7 day experiment
Aligned Reads

| | | | | | | | | | | | | | |
|---|---|---|---|---|---|---|---|---|---|---|---|---|---|
| 1512 | pgi | 184 | 0 | 185 | 1 | 14 | 0 | 94 | 0 | 71 | 0 | 125 | 1 |
| 1513 | potF | 0 | 0 | 0 | 0 | 0 | 0 | 0 | 0 | 0 | 0 | 0 | 0 |
| 1514 | potA | 1 | 0 | 0 | 0 | 0 | 0 | 0 | 0 | 0 | 0 | 0 | 0 |
| 1515 | mviN | 0 | 0 | 0 | 0 | 0 | 0 | 0 | 1 | 0 | 0 | 0 | 0 |
| 1516 | gatR1 | 0 | 0 | 0 | 0 | 0 | 0 | 0 | 0 | 0 | 0 | 0 | 0 |
| 1517 | pflA | 0 | 0 | 0 | 0 | 0 | 0 | 0 | 0 | 0 | 0 | 0 | 0 |
| 1518 | ahpF | 1 | 1 | 2 | 3 | 1 | 1 | 0 | 4 | 1 | 3 | 0 | 1 |
| 1519 | ahpC | 0 | 0 | 0 | 0 | 0 | 0 | 0 | 0 | 0 | 0 | 0 | 0 |
| 1520 | udp | 0 | 0 | 0 | 0 | 0 | 0 | 0 | 0 | 0 | 0 | 0 | 0 |
| 1521 | yqcE | 0 | 0 | 0 | 0 | 0 | 0 | 0 | 0 | 0 | 0 | 0 | 0 |
| 1522 | yqcD | 0 | 0 | 0 | 0 | 0 | 0 | 0 | 0 | 0 | 0 | 0 | 0 |
| 1523 | rsd | 1 | 0 | 0 | 0 | 0 | 0 | 0 | 0 | 0 | 0 | 0 | 0 |
| 1524 | udk | 0 | 0 | 0 | 0 | 0 | 0 | 0 | 0 | 0 | 0 | 0 | 0 |
| 1525 | xylB | 0 | 0 | 0 | 0 | 0 | 0 | 0 | 0 | 0 | 0 | 0 | 0 |
| 1526 | xylE | 0 | 0 | 0 | 0 | 0 | 0 | 0 | 0 | 0 | 0 | 0 | 0 |
| 1527 | xylG | 0 | 0 | 0 | 0 | 0 | 0 | 0 | 0 | 0 | 0 | 0 | 0 |
| 1528 | xylF | 0 | 0 | 0 | 0 | 0 | 0 | 0 | 0 | 0 | 0 | 0 | 0 |
| 1529 | dnaQ | 0 | 0 | 0 | 0 | 0 | 0 | 0 | 0 | 0 | 0 | 0 | 0 |
| 1530 | dnaX | 0 | 0 | 1 | 0 | 0 | 0 | 0 | 0 | 1 | 0 | 0 | 0 |
| 1531 | rdoA | 0 | 0 | 1 | 0 | 0 | 0 | 0 | 0 | 0 | 0 | 0 | 0 |
| 1532 | dnaK | 0 | 0 | 0 | 0 | 0 | 0 | 0 | 0 | 0 | 0 | 0 | 0 |
| 1533 | dnaJ | 0 | 0 | 0 | 0 | 0 | 0 | 0 | 0 | 0 | 0 | 0 | 0 |
| 1534 | sdhC | 0 | 0 | 0 | 0 | 0 | 0 | 0 | 0 | 0 | 0 | 0 | 0 |
| 1535 | trpL | 0 | 0 | 0 | 0 | 0 | 0 | 0 | 0 | 0 | 0 | 0 | 0 |
| 1536 | yraR | 0 | 0 | 0 | 0 | 0 | 0 | 0 | 0 | 0 | 0 | 0 | 0 |
| 1537 | yraQ | 0 | 0 | 1 | 0 | 0 | 0 | 0 | 0 | 0 | 0 | 0 | 0 |
| 1538 | vacJ | 0 | 0 | 0 | 0 | 0 | 0 | 0 | 0 | 0 | 0 | 0 | 0 |
| 1539 | yliE | 0 | 0 | 0 | 0 | 0 | 0 | 0 | 0 | 0 | 0 | 0 | 0 |
| 1540 | yraL | 0 | 0 | 0 | 0 | 0 | 0 | 0 | 0 | 0 | 0 | 0 | 0 |
| 1541 | yraM | 0 | 0 | 0 | 0 | 0 | 0 | 0 | 1 | 0 | 0 | 0 | 0 |
| 1542 | trpR | 0 | 0 | 0 | 0 | 0 | 0 | 0 | 0 | 0 | 0 | 0 | 0 |
| 1543 | yliI | 0 | 0 | 0 | 0 | 0 | 0 | 0 | 0 | 0 | 0 | 0 | 0 |
| 1544 | fdoG | 0 | 0 | 0 | 0 | 0 | 0 | 0 | 0 | 0 | 0 | 0 | 0 |
| 1545 | rfaQ | 0 | 0 | 0 | 0 | 0 | 0 | 0 | 0 | 0 | 0 | 0 | 0 |
| 1546 | rfaZ | 0 | 0 | 0 | 0 | 0 | 0 | 0 | 0 | 0 | 0 | 0 | 0 |
| 1547 | rfaD | 0 | 0 | 0 | 0 | 0 | 0 | 0 | 0 | 0 | 0 | 0 | 0 |
| 1548 | rfaH | 0 | 0 | 0 | 0 | 0 | 0 | 0 | 0 | 0 | 0 | 0 | 0 |
| 1549 | pstS | 0 | 0 | 0 | 2 | 0 | 0 | 1 | 2 | 0 | 0 | 0 | 0 |
| 1550 | evgA | 0 | 0 | 0 | 0 | 0 | 0 | 0 | 0 | 0 | 0 | 0 | 0 |
| 1551 | envY | 0 | 0 | 0 | 0 | 0 | 0 | 0 | 0 | 0 | 0 | 0 | 0 |
| 1552 | ydeV | 0 | 0 | 0 | 0 | 0 | 0 | 0 | 0 | 0 | 0 | 0 | 0 |
| 1553 | ydeP | 0 | 0 | 0 | 0 | 0 | 0 | 0 | 0 | 0 | 0 | 0 | 0 |
| 1554 | ydeQ | 0 | 0 | 0 | 0 | 0 | 0 | 0 | 0 | 0 | 0 | 0 | 0 |
| 1555 | btuC | 0 | 0 | 0 | 0 | 0 | 0 | 0 | 0 | 0 | 0 | 0 | 0 |
| 1556 | btuB | 0 | 0 | 0 | 0 | 0 | 0 | 0 | 0 | 0 | 0 | 0 | 0 |
| 1557 | pstC | 0 | 0 | 0 | 0 | 1 | 0 | 0 | 0 | 1 | 0 | 0 | 0 |
| 1558 | moeA | 0 | 0 | 0 | 0 | 0 | 0 | 0 | 0 | 0 | 0 | 0 | 0 |
| 1559 | ydeO | 0 | 0 | 0 | 0 | 0 | 0 | 0 | 0 | 0 | 0 | 0 | 0 |
| 1560 | ydeI | 0 | 0 | 0 | 0 | 0 | 0 | 0 | 0 | 0 | 0 | 0 | 0 |
| 1561 | ydeJ | 0 | 0 | 0 | 0 | 0 | 0 | 0 | 0 | 0 | 0 | 0 | 0 |
| 1562 | ydeA | 0 | 0 | 0 | 0 | 0 | 0 | 0 | 0 | 0 | 0 | 0 | 0 |
| 1563 | yhdX | 0 | 0 | 0 | 0 | 0 | 0 | 0 | 0 | 0 | 0 | 0 | 0 |
| 1564 | yaeQ | 0 | 0 | 0 | 0 | 0 | 0 | 0 | 0 | 0 | 0 | 0 | 0 |
| 1565 | yfiF | 0 | 0 | 0 | 0 | 0 | 0 | 0 | 0 | 0 | 0 | 0 | 0 |
| 1566 | feaB | 0 | 0 | 0 | 0 | 0 | 0 | 0 | 0 | 0 | 0 | 0 | 0 |
| 1567 | yfiE | 0 | 0 | 0 | 0 | 0 | 0 | 0 | 0 | 0 | 0 | 0 | 0 |
| 1568 | yfiC | 0 | 0 | 0 | 0 | 0 | 0 | 0 | 0 | 0 | 0 | 0 | 0 |
| 1569 | nagE | 0 | 0 | 0 | 1 | 0 | 0 | 0 | 0 | 1 | 0 | 0 | 0 |
| 1570 | nagB | 0 | 0 | 0 | 0 | 0 | 0 | 0 | 0 | 0 | 0 | 0 | 0 |
| 1571 | nagA | 0 | 0 | 0 | 0 | 0 | 0 | 0 | 0 | 0 | 0 | 0 | 0 |
| 1572 | feaR | 0 | 0 | 0 | 0 | 0 | 0 | 0 | 0 | 0 | 0 | 0 | 0 |
| 1573 | yaeH | 0 | 0 | 0 | 0 | 0 | 0 | 0 | 0 | 0 | 0 | 0 | 0 |
| 1574 | uidC | 0 | 0 | 0 | 0 | 0 | 0 | 0 | 0 | 0 | 0 | 0 | 0 |
| 1575 | uidA | 378 | 0 | 199 | 2 | 0 | 0 | 344 | 1 | 30 | 0 | 41 | 0 |
| 1576 | aes | 0 | 0 | 0 | 0 | 0 | 0 | 0 | 0 | 0 | 0 | 0 | 0 |
| 1577 | aer | 0 | 0 | 0 | 0 | 0 | 0 | 0 | 0 | 0 | 0 | 0 | 0 |
| 1578 | dps | 0 | 0 | 1 | 0 | 0 | 0 | 0 | 0 | 0 | 0 | 0 | 0 |
| 1579 | b0725 | 0 | 0 | 0 | 1 | 0 | 0 | 0 | 0 | 0 | 0 | 0 | 0 |
| 1580 | aroP | 0 | 0 | 0 | 0 | 0 | 0 | 0 | 0 | 0 | 0 | 0 | 0 |
| 1581 | tyrP | 0 | 0 | 0 | 0 | 0 | 0 | 0 | 0 | 0 | 0 | 0 | 0 |
| 1582 | yfgA | 0 | 0 | 0 | 0 | 0 | 0 | 0 | 0 | 0 | 0 | 0 | 0 |
| 1583 | yfgB | 0 | 0 | 0 | 0 | 0 | 0 | 0 | 0 | 0 | 0 | 0 | 0 |
| 1584 | yfgH | 0 | 0 | 0 | 0 | 0 | 0 | 0 | 0 | 0 | 1 | 0 | 0 |
| 1585 | yheO | 0 | 0 | 0 | 0 | 0 | 0 | 0 | 0 | 0 | 0 | 0 | 0 |
| 1586 | tyrB | 0 | 0 | 0 | 0 | 0 | 0 | 0 | 0 | 0 | 0 | 0 | 0 |
| 1587 | yheS | 0 | 0 | 0 | 0 | 0 | 0 | 0 | 0 | 0 | 0 | 0 | 0 |
| 1588 | aroF | 0 | 0 | 0 | 0 | 0 | 0 | 0 | 0 | 0 | 0 | 0 | 0 |
| 1589 | aroH | 0 | 0 | 0 | 0 | 0 | 0 | 0 | 0 | 0 | 0 | 0 | 0 |
| 1590 | aroK | 0 | 0 | 0 | 0 | 0 | 0 | 0 | 0 | 0 | 0 | 0 | 0 |
| 1591 | kefF | 0 | 0 | 1 | 0 | 0 | 0 | 0 | 0 | 0 | 0 | 0 | 0 |
| 1592 | kefG | 0 | 0 | 0 | 0 | 0 | 0 | 0 | 0 | 0 | 0 | 0 | 0 |
| 1593 | uvrD | 0 | 0 | 0 | 0 | 0 | 0 | 0 | 0 | 0 | 0 | 0 | 0 |
| 1594 | uvrC | 0 | 0 | 0 | 0 | 0 | 0 | 0 | 0 | 0 | 0 | 0 | 0 |
| 1595 | b4283 | 0 | 0 | 0 | 0 | 0 | 0 | 0 | 0 | 0 | 0 | 0 | 0 |

| | | | | | | | | | | | | | |
|---|---|---|---|---|---|---|---|---|---|---|---|---|---|
| 1596 | yidI | 0 | 0 | 0 | 0 | 0 | 0 | 0 | 0 | 0 | 0 | 0 | 0 |
| 1597 | ycfQ | 0 | 0 | 0 | 0 | 0 | 0 | 0 | 0 | 0 | 0 | 0 | 0 |
| 1598 | rsuA | 0 | 0 | 0 | 0 | 0 | 0 | 0 | 0 | 0 | 0 | 0 | 0 |
| 1599 | aegA | 0 | 0 | 0 | 0 | 0 | 0 | 0 | 0 | 0 | 0 | 0 | 0 |
| 1600 | yniC | 0 | 0 | 0 | 0 | 0 | 0 | 0 | 0 | 0 | 0 | 0 | 0 |
| 1601 | yniA | 0 | 0 | 0 | 0 | 0 | 0 | 0 | 0 | 0 | 0 | 0 | 0 |
| 1602 | caiC | 0 | 0 | 0 | 0 | 0 | 0 | 0 | 0 | 0 | 0 | 0 | 0 |
| 1603 | rumA | 416 | 0 | 355 | 0 | 4 | 0 | 297 | 0 | 237 | 0 | 29 | 0 |
| 1604 | ttdA | 0 | 0 | 0 | 0 | 0 | 0 | 0 | 0 | 0 | 0 | 0 | 0 |
| 1605 | rpe | 0 | 0 | 1 | 0 | 0 | 0 | 0 | 0 | 0 | 0 | 0 | 0 |
| 1606 | rph | 141 | 1 | 142 | 1 | 4 | 0 | 122 | 0 | 69 | 0 | 69 | 0 |
| 1607 | nhoA | 0 | 0 | 0 | 0 | 0 | 0 | 0 | 0 | 0 | 0 | 0 | 0 |
| 1608 | rhlE | 0 | 0 | 0 | 0 | 0 | 0 | 0 | 0 | 0 | 0 | 0 | 0 |
| 1609 | rhlB | 0 | 0 | 0 | 0 | 0 | 0 | 0 | 0 | 0 | 0 | 0 | 0 |
| 1610 | rpmI | 0 | 0 | 0 | 0 | 0 | 0 | 0 | 0 | 0 | 0 | 0 | 0 |
| 1611 | agaR | 0 | 0 | 0 | 0 | 0 | 0 | 0 | 0 | 0 | 0 | 0 | 0 |
| 1612 | rpmB | 0 | 0 | 0 | 0 | 0 | 0 | 0 | 0 | 1 | 0 | 0 | 0 |
| 1613 | rpmE | 0 | 0 | 0 | 0 | 0 | 0 | 0 | 0 | 0 | 0 | 0 | 0 |
| 1614 | lgt | 0 | 0 | 0 | 0 | 0 | 0 | 0 | 0 | 0 | 0 | 0 | 0 |
| 1615 | yjgH | 0 | 0 | 0 | 0 | 0 | 0 | 0 | 0 | 0 | 0 | 0 | 0 |
| 1616 | yjgJ | 0 | 0 | 0 | 0 | 0 | 0 | 0 | 0 | 0 | 0 | 0 | 0 |
| 1617 | yddA | 0 | 0 | 0 | 0 | 0 | 0 | 0 | 0 | 0 | 0 | 0 | 0 |
| 1618 | yjgA | 0 | 0 | 0 | 0 | 0 | 0 | 0 | 0 | 0 | 0 | 0 | 0 |
| 1619 | yjgR | 0 | 0 | 0 | 0 | 0 | 0 | 0 | 0 | 0 | 0 | 0 | 0 |
| 1620 | fabR | 37 | 14260 | 298 | 1885 | 37 | 1077 | 58 | 4887 | 1775 | 2612 | 245 | 2652 |
| 1621 | fabZ | 0 | 0 | 0 | 0 | 0 | 0 | 0 | 0 | 0 | 0 | 0 | 0 |
| 1622 | rfbB | 0 | 0 | 0 | 0 | 0 | 0 | 0 | 0 | 0 | 0 | 0 | 0 |
| 1623 | rfbA | 0 | 0 | 0 | 0 | 0 | 0 | 0 | 0 | 0 | 0 | 0 | 0 |
| 1624 | cpsG | 0 | 0 | 1 | 0 | 0 | 0 | 0 | 0 | 0 | 0 | 0 | 0 |
| 1625 | ygaV | 0 | 0 | 0 | 0 | 0 | 0 | 0 | 0 | 0 | 0 | 0 | 0 |
| 1626 | fkpA | 0 | 0 | 0 | 0 | 0 | 0 | 0 | 0 | 0 | 0 | 0 | 0 |
| 1627 | dam | 0 | 0 | 0 | 0 | 0 | 0 | 0 | 0 | 0 | 0 | 0 | 0 |
| 1628 | ackA | 0 | 0 | 0 | 0 | 0 | 0 | 0 | 0 | 0 | 0 | 0 | 0 |
| 1629 | setA | 0 | 0 | 0 | 0 | 0 | 0 | 0 | 0 | 0 | 0 | 0 | 0 |
| 1630 | setB | 0 | 0 | 0 | 0 | 0 | 0 | 0 | 0 | 0 | 0 | 0 | 0 |
| 1631 | mglB | 0 | 0 | 0 | 0 | 0 | 0 | 0 | 0 | 0 | 0 | 0 | 0 |
| 1632 | fhlA | 0 | 0 | 0 | 0 | 0 | 1 | 0 | 0 | 0 | 0 | 0 | 0 |
| 1633 | iscS | 0 | 0 | 0 | 0 | 0 | 0 | 0 | 0 | 0 | 1 | 0 | 0 |
| 1634 | iscR | 0 | 0 | 0 | 0 | 0 | 0 | 0 | 0 | 0 | 0 | 0 | 0 |
| 1635 | dusA | 0 | 0 | 0 | 0 | 0 | 0 | 0 | 0 | 0 | 0 | 0 | 0 |
| 1636 | dusB | 0 | 0 | 0 | 0 | 0 | 0 | 0 | 0 | 0 | 0 | 0 | 0 |
| 1637 | dusC | 0 | 0 | 0 | 0 | 0 | 0 | 0 | 0 | 0 | 0 | 0 | 0 |
| 1638 | nfnB | 0 | 0 | 0 | 0 | 0 | 0 | 0 | 0 | 0 | 0 | 0 | 0 |
| 1639 | b2639 | 0 | 0 | 0 | 0 | 0 | 0 | 0 | 0 | 0 | 0 | 0 | 0 |
| 1640 | ycfD | 25 | 0 | 14 | 0 | 3 | 0 | 25 | 0 | 2 | 1 | 0 | 0 |
| 1641 | mfd | 0 | 0 | 0 | 0 | 0 | 0 | 0 | 0 | 0 | 0 | 0 | 0 |
| 1642 | modF | 0 | 0 | 0 | 0 | 0 | 0 | 0 | 0 | 0 | 0 | 0 | 0 |
| 1643 | gutM | 0 | 0 | 0 | 0 | 0 | 0 | 0 | 0 | 0 | 0 | 0 | 0 |
| 1644 | gmk | 0 | 0 | 0 | 0 | 0 | 0 | 0 | 0 | 0 | 0 | 0 | 0 |
| 1645 | skp | 0 | 0 | 0 | 0 | 0 | 0 | 0 | 0 | 0 | 0 | 0 | 0 |
| 1646 | sufI | 0 | 0 | 0 | 0 | 0 | 1 | 0 | 0 | 0 | 0 | 0 | 0 |
| 1647 | add | 0 | 0 | 0 | 0 | 0 | 0 | 0 | 0 | 0 | 0 | 0 | 0 |
| 1648 | ada | 0 | 0 | 0 | 0 | 0 | 0 | 0 | 0 | 0 | 0 | 0 | 0 |
| 1649 | gmr | 0 | 0 | 0 | 0 | 0 | 0 | 0 | 0 | 0 | 0 | 0 | 0 |
| 1650 | adk | 0 | 0 | 0 | 0 | 0 | 0 | 0 | 0 | 0 | 0 | 0 | 0 |
| 1651 | eutN | 1 | 0 | 0 | 0 | 0 | 0 | 0 | 0 | 0 | 0 | 0 | 0 |
| 1652 | ppdD | 0 | 0 | 0 | 0 | 0 | 0 | 0 | 0 | 0 | 0 | 0 | 0 |
| 1653 | otsB | 1 | 0 | 0 | 0 | 0 | 0 | 0 | 0 | 1 | 0 | 0 | 0 |
| 1654 | yfjG | 0 | 0 | 0 | 0 | 0 | 0 | 0 | 0 | 0 | 0 | 0 | 0 |
| 1655 | pagB | 0 | 0 | 0 | 0 | 0 | 0 | 0 | 0 | 0 | 0 | 0 | 0 |
| 1656 | nadC | 0 | 0 | 0 | 0 | 0 | 0 | 0 | 0 | 0 | 0 | 0 | 0 |
| 1657 | b0663 | 0 | 0 | 0 | 0 | 0 | 0 | 0 | 0 | 0 | 0 | 0 | 0 |
| 1658 | yadG | 0 | 0 | 0 | 0 | 0 | 0 | 0 | 0 | 0 | 0 | 0 | 0 |
| 1659 | rstA | 0 | 0 | 0 | 0 | 0 | 0 | 0 | 0 | 0 | 0 | 0 | 0 |
| 1660 | grpE | 0 | 0 | 0 | 0 | 0 | 0 | 0 | 0 | 0 | 0 | 0 | 0 |
| 1661 | murA | 0 | 0 | 0 | 0 | 0 | 0 | 0 | 0 | 0 | 0 | 0 | 0 |
| 1662 | murC | 0 | 0 | 0 | 0 | 0 | 0 | 0 | 0 | 0 | 0 | 0 | 0 |
| 1663 | molR2 | 0 | 0 | 0 | 0 | 0 | 0 | 0 | 0 | 0 | 0 | 0 | 0 |
| 1664 | dcm | 0 | 0 | 0 | 0 | 0 | 0 | 0 | 1 | 0 | 0 | 0 | 0 |
| 1665 | yddW | 0 | 0 | 0 | 0 | 0 | 0 | 0 | 0 | 0 | 0 | 0 | 0 |
| 1666 | dcd | 0 | 0 | 0 | 0 | 0 | 0 | 0 | 0 | 0 | 0 | 0 | 0 |
| 1667 | yqaD | 0 | 0 | 0 | 0 | 0 | 0 | 0 | 0 | 0 | 0 | 0 | 0 |
| 1668 | nohA | 0 | 0 | 0 | 0 | 0 | 0 | 0 | 0 | 0 | 0 | 0 | 0 |
| 1669 | gudP | 0 | 0 | 0 | 0 | 0 | 0 | 0 | 0 | 0 | 1 | 0 | 0 |
| 1670 | yddM | 0 | 0 | 0 | 0 | 0 | 0 | 0 | 0 | 0 | 0 | 0 | 0 |
| 1671 | yddL | 0 | 0 | 0 | 0 | 0 | 0 | 0 | 0 | 0 | 0 | 0 | 0 |
| 1672 | yddH | 0 | 0 | 0 | 0 | 0 | 0 | 0 | 0 | 0 | 0 | 0 | 0 |
| 1673 | yddE | 0 | 0 | 0 | 0 | 0 | 0 | 0 | 0 | 0 | 0 | 0 | 0 |
| 1674 | dcp | 0 | 0 | 0 | 0 | 0 | 0 | 0 | 0 | 0 | 0 | 0 | 0 |
| 1675 | b1403 | 0 | 0 | 0 | 0 | 0 | 0 | 0 | 0 | 0 | 0 | 0 | 0 |
| 1676 | yabP | 0 | 0 | 0 | 0 | 0 | 0 | 0 | 0 | 0 | 0 | 0 | 0 |
| 1677 | yfhH | 0 | 0 | 0 | 0 | 0 | 0 | 0 | 0 | 0 | 0 | 0 | 0 |
| 1678 | ndk | 0 | 0 | 0 | 0 | 0 | 0 | 0 | 0 | 0 | 0 | 0 | 0 |
| 1679 | yfhM | 0 | 0 | 0 | 0 | 0 | 0 | 0 | 0 | 0 | 0 | 0 | 0 |

| | | | | | | | | | | | | | |
|---|---|---|---|---|---|---|---|---|---|---|---|---|---|
| 1680 | yfhL | 0 | 0 | 0 | 0 | 0 | 0 | 0 | 0 | 0 | 0 | 0 | 0 |
| 1681 | yfhB | 0 | 0 | 0 | 0 | 0 | 0 | 0 | 0 | 0 | 0 | 0 | 0 |
| 1682 | yfhG | 0 | 0 | 0 | 0 | 0 | 0 | 0 | 0 | 0 | 0 | 0 | 0 |
| 1683 | yabI | 0 | 0 | 0 | 0 | 0 | 0 | 0 | 0 | 0 | 0 | 0 | 0 |
| 1684 | yfhR | 0 | 0 | 0 | 1 | 0 | 0 | 0 | 0 | 0 | 0 | 0 | 0 |
| 1685 | yabN | 0 | 0 | 0 | 0 | 0 | 0 | 0 | 0 | 0 | 0 | 0 | 0 |
| 1686 | cvrA | 0 | 0 | 0 | 0 | 0 | 0 | 0 | 0 | 0 | 0 | 0 | 0 |
| 1687 | yaaU | 1677 | 3 | 1267 | 3 | 43 | 0 | 1109 | 1 | 746 | 2 | 2170 | 0 |
| 1688 | mazE | 1 | 0 | 0 | 0 | 0 | 0 | 0 | 0 | 0 | 0 | 0 | 0 |
| 1689 | mazG | 0 | 0 | 0 | 0 | 0 | 0 | 0 | 0 | 0 | 0 | 0 | 0 |
| 1690 | appA | 0 | 0 | 0 | 0 | 0 | 0 | 0 | 0 | 0 | 0 | 0 | 0 |
| 1691 | yjfK | 0 | 0 | 0 | 0 | 0 | 0 | 0 | 0 | 0 | 0 | 0 | 0 |
| 1692 | yjfI | 0 | 0 | 0 | 0 | 0 | 0 | 0 | 0 | 0 | 0 | 0 | 0 |
| 1693 | mraZ | 0 | 0 | 0 | 0 | 0 | 0 | 1 | 0 | 0 | 0 | 0 | 0 |
| 1694 | ydaQ | 0 | 0 | 0 | 0 | 0 | 0 | 0 | 0 | 0 | 0 | 0 | 0 |
| 1695 | hdhA | 0 | 0 | 0 | 0 | 0 | 0 | 0 | 0 | 0 | 1 | 0 | 0 |
| 1696 | yjfY | 1190 | 1 | 737 | 2 | 15 | 1 | 1390 | 1 | 341 | 0 | 93 | 0 |
| 1697 | yjfQ | 0 | 0 | 0 | 0 | 0 | 0 | 0 | 0 | 0 | 0 | 0 | 0 |
| 1698 | yhdA | 0 | 0 | 0 | 0 | 0 | 0 | 0 | 0 | 0 | 0 | 0 | 0 |
| 1699 | fliY | 0 | 0 | 0 | 0 | 0 | 0 | 0 | 0 | 0 | 0 | 0 | 0 |
| 1700 | yhdJ | 0 | 0 | 0 | 0 | 0 | 0 | 0 | 0 | 0 | 0 | 0 | 0 |
| 1701 | yhdH | 0 | 0 | 0 | 0 | 0 | 0 | 0 | 0 | 0 | 0 | 0 | 0 |
| 1702 | yhdN | 0 | 0 | 0 | 0 | 0 | 0 | 0 | 0 | 0 | 0 | 0 | 0 |
| 1703 | yffH | 0 | 0 | 0 | 0 | 0 | 0 | 0 | 0 | 0 | 0 | 0 | 0 |
| 1704 | fliL | 0 | 0 | 0 | 0 | 0 | 0 | 0 | 0 | 1 | 0 | 1 | 0 |
| 1705 | yffS | 0 | 0 | 0 | 0 | 0 | 0 | 0 | 0 | 0 | 0 | 0 | 0 |
| 1706 | fliE | 0 | 0 | 0 | 0 | 0 | 0 | 0 | 0 | 0 | 0 | 0 | 0 |
| 1707 | yhdY | 0 | 0 | 0 | 0 | 0 | 0 | 0 | 0 | 0 | 0 | 0 | 0 |
| 1708 | fliF | 0 | 0 | 0 | 0 | 0 | 0 | 0 | 0 | 0 | 0 | 0 | 0 |
| 1709 | fliA | 0 | 0 | 0 | 0 | 0 | 0 | 0 | 0 | 0 | 1 | 0 | 1 |
| 1710 | fliC | 0 | 0 | 0 | 0 | 0 | 0 | 0 | 0 | 0 | 0 | 0 | 0 |
| 1711 | yaiA | 0 | 0 | 0 | 0 | 0 | 0 | 0 | 0 | 0 | 0 | 0 | 0 |
| 1712 | rplL | 0 | 0 | 0 | 0 | 0 | 0 | 1 | 0 | 0 | 0 | 0 | 0 |
| 1713 | srmB | 0 | 0 | 0 | 0 | 0 | 0 | 0 | 0 | 0 | 0 | 0 | 0 |
| 1714 | mglA | 0 | 0 | 0 | 0 | 0 | 1 | 0 | 0 | 0 | 0 | 0 | 0 |
| 1715 | yohJ | 0 | 0 | 0 | 0 | 0 | 0 | 0 | 0 | 0 | 0 | 0 | 0 |
| 1716 | yohL | 0 | 0 | 0 | 0 | 0 | 0 | 0 | 0 | 0 | 0 | 0 | 0 |
| 1717 | yohM | 0 | 0 | 0 | 0 | 0 | 0 | 0 | 0 | 0 | 0 | 0 | 0 |
| 1718 | pdxY | 0 | 0 | 0 | 0 | 0 | 0 | 0 | 0 | 0 | 0 | 0 | 0 |
| 1719 | yohC | 0 | 0 | 0 | 0 | 0 | 0 | 0 | 0 | 0 | 0 | 0 | 0 |
| 1720 | yohD | 0 | 0 | 0 | 0 | 0 | 0 | 0 | 0 | 0 | 0 | 0 | 0 |
| 1721 | yaiB | 0 | 0 | 0 | 0 | 0 | 0 | 0 | 0 | 0 | 0 | 0 | 0 |
| 1722 | yeaE | 0 | 0 | 0 | 0 | 0 | 0 | 0 | 0 | 0 | 0 | 0 | 0 |
| 1723 | yqaC | 0 | 0 | 0 | 0 | 0 | 0 | 0 | 0 | 0 | 0 | 0 | 0 |
| 1724 | yqaB | 1 | 0 | 0 | 0 | 0 | 0 | 1 | 0 | 1 | 0 | 2 | 0 |
| 1725 | yqaE | 0 | 0 | 0 | 0 | 0 | 0 | 0 | 0 | 0 | 0 | 0 | 0 |
| 1726 | nrdB | 0 | 0 | 1 | 0 | 0 | 0 | 0 | 1 | 1 | 0 | 0 | 0 |
| 1727 | nrdD | 0 | 0 | 0 | 0 | 0 | 0 | 0 | 0 | 0 | 0 | 0 | 0 |
| 1728 | rhsE | 0 | 0 | 0 | 0 | 0 | 0 | 0 | 0 | 0 | 0 | 0 | 0 |
| 1729 | rhsD | 0 | 0 | 0 | 0 | 0 | 0 | 0 | 0 | 0 | 0 | 0 | 0 |
| 1730 | rhsB | 0 | 0 | 0 | 0 | 0 | 0 | 0 | 0 | 0 | 0 | 0 | 0 |
| 1731 | rhsA | 0 | 0 | 0 | 0 | 0 | 0 | 0 | 0 | 0 | 0 | 0 | 0 |
| 1732 | pdxH | 0 | 0 | 0 | 0 | 0 | 0 | 0 | 0 | 0 | 0 | 0 | 0 |
| 1733 | pdxK | 0 | 0 | 0 | 0 | 0 | 0 | 0 | 0 | 0 | 0 | 0 | 0 |
| 1734 | pdxB | 0 | 0 | 0 | 0 | 0 | 0 | 0 | 0 | 0 | 0 | 0 | 0 |
| 1735 | nrdG | 0 | 0 | 0 | 0 | 0 | 0 | 0 | 0 | 0 | 0 | 0 | 0 |
| 1736 | nrdH | 0 | 0 | 0 | 0 | 0 | 0 | 0 | 0 | 1 | 0 | 0 | 0 |
| 1737 | yghU | 0 | 0 | 0 | 0 | 0 | 0 | 0 | 0 | 0 | 0 | 0 | 0 |
| 1738 | mdoB | 1 | 1 | 0 | 2 | 2 | 1 | 0 | 1 | 0 | 0 | 1 | 2 |
| 1739 | negCtrl | 33 | 0 | 60 | 1 | 59 | 2 | 86 | 1 | 66 | 0 | 40 | 0 |

### Data Set S3 - Nissle 1917 library 5 day experiment (-DSS and +DSS)
### Odds Ratios

| | Day | 0 | 1 | | | | | | | |
|---|---|---|---|---|---|---|---|---|---|---|
| | | | -DSS | | | | +DSS | | | |
| | Promoter | GV-11-13 | M1-11-14 | M2-11-14 | M3-11-14 | M4-11-14 | M5-11-14 | M6-11-14 | M7-11-14 | M8-11-14 |
| 41 | torC23cI | 0.0051 | 0.1065 | 0.3867 | 0.1589 | 0.2054 | 0.5833 | 0.4655 | 0.1525 | 0.0808 |
| 55 | ynfE10cI | 0.0000 | 0.1426 | 0.0851 | 0.1026 | 0.2166 | 0.2615 | 0.5247 | 0.2128 | 0.1278 |
| 56 | ynfE15cI | 0.0001 | 0.1101 | 0.0565 | 0.0774 | 0.1844 | 0.2575 | 0.5539 | 0.2032 | 0.1247 |
| 39 | torC15cI | 0.0036 | 0.0848 | 0.1217 | 0.1122 | 0.2226 | 0.1793 | 0.4238 | 0.1575 | 0.0986 |
| 37 | torC5cI | 0.0025 | 0.0873 | 0.1108 | 0.1041 | 0.2047 | 0.1830 | 0.4300 | 0.1586 | 0.0970 |
| 42 | yeaRWTcI | 0.0003 | 0.0302 | 0.0169 | 0.0540 | 0.0704 | 0.0260 | 0.0671 | 0.0213 | 0.0173 |
| 58 | ynfE23cI | 0.0000 | 0.0528 | 0.0123 | 0.0718 | 0.0501 | 0.2078 | 0.3934 | 0.1567 | 0.1143 |
| 38 | torC10cI | 0.0049 | 0.0776 | 0.1143 | 0.2044 | 0.3071 | 0.2345 | 0.3556 | 0.1448 | 0.0903 |
| 44 | yeaR10cI | 0.0000 | 0.0277 | 0.0024 | 0.0188 | 0.0331 | 0.0201 | 0.0409 | 0.0181 | 0.0089 |
| 45 | yeaR15cI | 0.0000 | 0.0162 | 0.0068 | 0.0230 | 0.0404 | 0.0157 | 0.0270 | 0.0133 | 0.0091 |
| 40 | torC17cI | 0.0132 | 0.0370 | 0.0604 | 0.0336 | 0.0671 | N/A | N/A | N/A | N/A |
| 47 | yeaR23cI | 0.0000 | 0.0000 | N/A | N/A | 0.1274 | 0.0108 | N/A | N/A | N/A |
| 13 | hycAWTcI | 0.0001 | 0.0141 | 0.0046 | 0.0037 | 0.0059 | 0.0143 | 0.0469 | 0.0121 | 0.0073 |
| 16 | hycA15cI | 0.0000 | 0.0037 | 0.0014 | 0.0019 | 0.0019 | 0.0023 | 0.0136 | 0.0046 | 0.0026 |
| 32 | narK10cI | 0.0004 | N/A | N/A | 0.0067 | 0.0000 | N/A | N/A | N/A | 0.0000 |
| 57 | ynfE17cI | 0.0001 | 0.0055 | 0.0007 | 0.0091 | 0.0025 | 0.1103 | 0.1408 | 0.0602 | 0.0779 |
| 8 | fdnG5cI | 0.0001 | 0.0002 | 0.0033 | 0.0002 | 0.0059 | 0.0006 | 0.0000 | 0.0004 | 0.0006 |
| 30 | narKWTcI | 0.0000 | 0.0008 | 0.0003 | 0.0013 | 0.0003 | 0.0006 | 0.0037 | 0.0007 | 0.0007 |
| 54 | ynfE5cI | 0.0000 | 0.0006 | 0.0008 | 0.0012 | 0.0014 | 0.0018 | 0.0000 | 0.0007 | 0.0009 |
| 49 | yedY10cI | 0.0000 | 0.0005 | 0.0006 | 0.0000 | 0.0012 | 0.0006 | 0.0000 | 0.0007 | 0.0006 |
| 50 | yedY15cI | 0.0000 | 0.0003 | 0.0004 | 0.0006 | 0.0003 | 0.0005 | 0.0016 | 0.0006 | 0.0003 |
| 17 | hycA17cI | 0.0000 | 0.0000 | 0.0010 | 0.0010 | 0.0006 | 0.0004 | 0.0000 | 0.0005 | 0.0000 |
| 35 | narK23cI | 0.0000 | 0.0008 | 0.0000 | 0.0011 | 0.0005 | 0.0065 | 0.0082 | 0.0030 | 0.0036 |
| 51 | yedY17cI | 0.0001 | 0.0001 | 0.0000 | 0.0007 | 0.0005 | 0.0003 | 0.0047 | 0.0006 | 0.0002 |
| 14 | hycA5cI | 0.0000 | 0.0000 | 0.0000 | 0.0003 | 0.0004 | 0.0009 | 0.0013 | 0.0000 | 0.0004 |
| 19 | moaAWTcI | 0.0001 | 0.0012 | 0.0000 | 0.0004 | 0.0008 | 0.0003 | 0.0000 | 0.0000 | 0.0002 |
| 52 | yedY23cI | 0.0001 | 0.0001 | 0.0003 | 0.0006 | 0.0010 | 0.0003 | 0.0011 | 0.0002 | 0.0002 |
| 53 | ynfEWTcI | 0.0000 | 0.0002 | 0.0006 | 0.0006 | 0.0002 | 0.0014 | 0.0021 | 0.0005 | 0.0007 |
| 31 | narK5cI | 0.0001 | 0.0002 | 0.0000 | 0.0002 | 0.0000 | 0.0002 | 0.0009 | 0.0000 | 0.0003 |
| 10 | fdnG15cI | 0.0000 | 0.0000 | N/A | 0.0000 | 0.0000 | 0.0000 | N/A | N/A | 0.0043 |
| 12 | fdnG23cI | 0.0000 | 0.0000 | N/A | 0.0000 | 0.0000 | 0.0000 | N/A | N/A | 0.0000 |
| 34 | narK17cI | 0.0000 | N/A | N/A | N/A | 0.0000 | N/A | N/A | N/A | 0.0086 |
| 18 | hycA23cI | 0.0000 | 0.0000 | N/A | 0.0000 | 0.0000 | 0.0000 | 0.0000 | 0.0000 | 0.0051 |
| 15 | hycA10cI | 0.0000 | N/A | N/A | N/A | 0.0000 | N/A | N/A | N/A | 0.0000 |
| 48 | yedY5cI | 0.0000 | 0.0000 | N/A | 0.0000 | 0.0000 | 0.0000 | N/A | N/A | 0.0000 |
| 33 | narK15cI | 0.0002 | 0.0000 | N/A | 0.0000 | 0.0000 | 0.0000 | 0.0000 | 0.0150 | 0.0029 |
| 11 | fdnG17cI | 0.0000 | N/A | N/A | N/A | N/A | N/A | N/A | N/A | N/A |
| 46 | yeaR17cI | 0.0000 | N/A | N/A | N/A | N/A | N/A | N/A | N/A | N/A |
| 43 | yeaR5cI | 0.0000 | N/A | N/A | N/A | N/A | N/A | N/A | N/A | N/A |
| 9 | fdnG10cI | 0.0008 | N/A | N/A | N/A | N/A | N/A | N/A | N/A | 0.0000 |
| 36 | narG23cI | 0.0225 | N/A | N/A | N/A | 0.0531 | N/A | N/A | N/A | N/A |
| 1 | cydAWTcI | 0.0261 | 0.1433 | 0.1602 | 0.1479 | 0.2100 | 0.2438 | 0.5780 | 0.2555 | 0.1417 |
| 2 | cydA5cI | 0.0297 | 0.0207 | N/A | N/A | 0.0000 | 0.1935 | N/A | N/A | 0.0771 |
| 26 | napF10cI | 0.0349 | 0.1551 | 0.1303 | 0.0967 | 0.2493 | 0.5373 | 0.5331 | 0.1345 | 0.2804 |
| 28 | napF17cI | 0.0502 | 0.0431 | N/A | 0.0806 | 0.1911 | 0.2269 | 0.0000 | N/A | 0.1363 |
| 7 | fdnGWTcI | 0.0554 | 0.1411 | 0.1817 | 0.1465 | 0.2286 | 0.2726 | 0.4587 | 0.2340 | 0.1153 |
| 27 | napF15cI | 0.0658 | 0.1219 | 0.1781 | 0.1418 | 0.3089 | 0.2432 | 0.6704 | 0.2103 | 0.1413 |
| 25 | napF5cI | 0.0723 | 0.1277 | 0.1699 | 0.1411 | 0.3200 | 0.2468 | 0.5672 | 0.2125 | 0.1470 |
| 29 | napF23cI | 0.0726 | 0.1320 | 0.1585 | 0.1401 | 0.2908 | 0.2363 | 0.5424 | 0.2087 | 0.1347 |
| 4 | cydA15cI | 0.0752 | 0.2068 | 0.2712 | 0.1244 | 0.1638 | 1.2295 | 0.5331 | 0.1345 | 0.1367 |
| 20 | moaA5cI | 0.0832 | 0.1871 | 0.1832 | 0.2096 | 0.2463 | 0.3247 | 0.7026 | 0.2789 | 0.1586 |
| 22 | moaA15cI | 0.0842 | 0.1576 | 0.1985 | 0.1905 | 0.2730 | 0.2679 | 0.6160 | 0.2535 | 0.1603 |
| 21 | moaA10cI | 0.0848 | 0.1756 | 0.1961 | 0.2088 | 0.3110 | 0.3150 | 0.6366 | 0.2643 | 0.1529 |
| 5 | cydA17cI | 0.0875 | 0.1435 | 0.1801 | 0.1688 | 0.3140 | 0.2905 | 0.6690 | 0.2334 | 0.1766 |
| 6 | cydA23cI | 0.0882 | 0.1451 | 0.1822 | 0.1483 | 0.2868 | 0.2702 | 0.6046 | 0.2487 | 0.1704 |
| 24 | moaA23cI | 0.0893 | 0.1790 | 0.2181 | 0.2238 | 0.2819 | 0.3023 | 0.6439 | 0.2614 | 0.1661 |
| 3 | cydA10cI | 0.0956 | 0.1635 | 0.1889 | 0.1689 | 0.3072 | 0.3380 | 0.6060 | 0.2614 | 0.1751 |
| 23 | moaA17cI | 0.1678 | 0.3691 | 0.3508 | 0.4588 | 0.5138 | 0.6293 | 1.6569 | 0.6470 | 0.2671 |
| 59 | fabR | 1.0000 | 1.0000 | 1.0000 | 1.0000 | 1.0000 | 1.0000 | 1.0000 | 1.0000 | 1.0000 |

| | Day | 2 | | | | | | | | 4 | | | | | | | |
|---|---|---|---|---|---|---|---|---|---|---|---|---|---|---|---|---|---|
| | | -DSS | | | | +DSS | | | | -DSS | | | | +DSS | | | |
| | Promoter | M1-11-15 | M2-11-15 | M3-11-15 | M4-11-15 | M5-11-15 | M6-11-15 | M7-11-15 | M8-11-15 | M1-11-17 | M2-11-17 | M3-11-17 | M4-11-17 | M5-11-17 | M6-11-17 | M7-11-17 | |
| 41 | torC23cI | 0.1532 | 0.1545 | 0.2154 | 0.1014 | 0.0699 | 0.1590 | 0.0917 | 0.1091 | 0.0975 | 0.1803 | 0.1773 | 0.1254 | 0.1661 | 0.1727 | 0.0470 | N/A |
| 55 | ynfE10cI | 0.1658 | 0.1601 | 0.1576 | 0.1278 | 0.1252 | 0.1607 | 0.1086 | 0.1824 | 0.1054 | 0.2393 | 0.1623 | 0.1705 | 0.2274 | 0.1880 | 0.1957 | N/A |
| 56 | ynfE15cI | 0.1533 | 0.1250 | 0.1412 | 0.1297 | 0.1144 | 0.1653 | 0.1086 | 0.1614 | 0.1237 | 0.1403 | 0.1892 | 0.1225 | 0.2121 | 0.2305 | 0.1250 | N/A |
| 39 | torC15cI | 0.1618 | 0.1368 | 0.1440 | 0.0928 | 0.0684 | 0.1121 | 0.0778 | 0.1160 | 0.0500 | 0.0771 | 0.1010 | 0.0918 | 0.0992 | 0.1050 | 0.0423 | N/A |
| 37 | torC5cI | 0.1513 | 0.1409 | 0.1351 | 0.0910 | 0.0649 | 0.1119 | 0.0731 | 0.1168 | 0.0513 | 0.0761 | 0.0989 | 0.0864 | 0.0891 | 0.0947 | 0.0376 | N/A |
| 42 | yeaRWTcI | 0.0566 | 0.0317 | 0.1027 | 0.0907 | 0.0711 | 0.0361 | 0.0228 | 0.0321 | 0.0602 | 0.0744 | 0.1325 | 0.0663 | 0.1271 | 0.1705 | 0.1199 | N/A |
| 58 | ynfE23cI | 0.0795 | 0.0439 | 0.0971 | 0.0553 | 0.0966 | 0.1329 | 0.0763 | 0.1403 | 0.0568 | 0.0681 | 0.1034 | 0.0572 | 0.1787 | 0.1905 | 0.1006 | N/A |
| 38 | torC10cI | 0.2223 | 0.1354 | 0.1550 | 0.1086 | 0.0511 | 0.1820 | 0.1077 | 0.3074 | 0.0329 | 0.0550 | 0.1248 | 0.1405 | 0.3053 | 0.1652 | 0.0166 | N/A |
| 44 | yeaR10cI | 0.0797 | 0.0204 | 0.0658 | 0.0556 | 0.0431 | 0.0230 | 0.0227 | 0.0205 | 0.0454 | 0.0602 | 0.0750 | 0.0655 | 0.0731 | 0.1179 | 0.0758 | N/A |
| 45 | yeaR15cI | 0.0471 | 0.0156 | 0.0294 | 0.1505 | 0.0821 | 0.0246 | 0.0132 | 0.0161 | 0.0371 | 0.0567 | 0.0814 | 0.0754 | 0.0890 | 0.1292 | 0.0784 | N/A |
| 40 | torC17cI | 0.1993 | 0.0488 | N/A | 0.0931 | 0.0500 | 0.2274 | 0.1205 | 0.1279 | 0.0782 | N/A | N/A | 0.0644 | 0.1069 | 0.0221 | 0.0106 | N/A |
| 47 | yeaR23cI | 0.0000 | N/A | N/A | N/A | 0.0530 | 0.0149 | N/A | 0.0747 | N/A | N/A | N/A | N/A | N/A | N/A | 0.0927 | N/A |
| 13 | hycAWTcI | 0.0295 | 0.0138 | 0.0080 | 0.0209 | 0.0053 | 0.0152 | 0.0095 | 0.0078 | 0.0101 | 0.0221 | 0.0206 | 0.0161 | 0.0208 | 0.0054 | 0.0148 | N/A |
| 16 | hycA15cI | 0.0043 | 0.0000 | 0.0022 | 0.0136 | 0.0020 | 0.0055 | 0.0025 | 0.0030 | 0.0053 | 0.0024 | 0.0023 | 0.0077 | 0.0064 | 0.0053 | 0.0021 | N/A |
| 32 | narK10cI | N/A | N/A | N/A | N/A | 0.0000 | N/A | N/A | 0.0000 | N/A | N/A | N/A | N/A | N/A | N/A | N/A | N/A |
| 57 | ynfE17cI | 0.0025 | 0.0000 | 0.0057 | 0.0050 | 0.0557 | 0.0727 | 0.0402 | 0.1016 | 0.0000 | 0.0000 | 0.0056 | 0.0000 | 0.0771 | 0.0872 | 0.0409 | N/A |
| 8 | fdnG5cI | 0.0022 | 0.0043 | 0.0000 | 0.0121 | 0.0008 | 0.0005 | 0.0013 | 0.0007 | 0.0000 | 0.0042 | 0.0020 | 0.0040 | 0.0025 | 0.0068 | 0.0030 | N/A |
| 30 | narKWTcI | 0.0000 | 0.0043 | 0.0036 | 0.0006 | 0.0002 | 0.0007 | 0.0004 | 0.0008 | 0.0016 | 0.0030 | 0.0025 | 0.0000 | 0.0029 | 0.0008 | 0.0037 | N/A |
| 54 | ynfE5cI | 0.0018 | 0.0000 | 0.0000 | 0.0011 | 0.0008 | 0.0022 | 0.0002 | 0.0011 | 0.0000 | 0.0000 | 0.0000 | 0.0083 | 0.0023 | 0.0015 | 0.0008 | N/A |
| 49 | yedY10cI | 0.0000 | 0.0026 | 0.0000 | 0.0040 | 0.0001 | 0.0007 | 0.0005 | 0.0004 | 0.0000 | 0.0000 | 0.0000 | 0.0030 | 0.0009 | 0.0007 | 0.0013 | N/A |
| 50 | yedY15cI | 0.0019 | 0.0000 | 0.0000 | 0.0023 | 0.0002 | 0.0009 | 0.0001 | 0.0002 | 0.0000 | 0.0010 | 0.0024 | 0.0015 | 0.0004 | 0.0000 | 0.0017 | N/A |
| 17 | hycA17cI | 0.0000 | 0.0000 | 0.0000 | 0.0000 | 0.0002 | 0.0000 | 0.0000 | 0.0002 | 0.0000 | 0.0000 | 0.0025 | 0.0043 | 0.0003 | 0.0008 | 0.0005 | N/A |
| 35 | narK23cI | 0.0000 | 0.0000 | 0.0027 | 0.0000 | 0.0027 | 0.0055 | 0.0019 | 0.0027 | 0.0021 | 0.0000 | 0.0000 | 0.0000 | 0.0025 | 0.0034 | 0.0060 | N/A |
| 51 | yedY17cI | 0.0000 | 0.0000 | 0.0015 | 0.0012 | 0.0002 | 0.0010 | 0.0005 | 0.0002 | 0.0000 | 0.0017 | 0.0018 | 0.0000 | 0.0004 | 0.0005 | 0.0021 | N/A |
| 14 | hycA5cI | 0.0000 | 0.0000 | 0.0000 | 0.0000 | 0.0001 | 0.0011 | 0.0000 | 0.0000 | 0.0023 | 0.0000 | 0.0000 | 0.0000 | 0.0002 | 0.0000 | 0.0014 | N/A |
| 19 | moaAWTcI | 0.0000 | 0.0000 | 0.0000 | 0.0000 | 0.0001 | 0.0012 | 0.0002 | 0.0002 | 0.0011 | 0.0014 | 0.0012 | 0.0000 | 0.0008 | 0.0009 | 0.0003 | N/A |
| 52 | yedY23cI | 0.0004 | 0.0000 | 0.0000 | 0.0000 | 0.0001 | 0.0005 | 0.0001 | 0.0002 | 0.0004 | 0.0010 | 0.0005 | 0.0007 | 0.0006 | 0.0009 | 0.0016 | N/A |
| 53 | ynfEWTcI | 0.0000 | 0.0004 | 0.0003 | 0.0002 | 0.0005 | 0.0011 | 0.0005 | 0.0006 | 0.0006 | 0.0003 | 0.0003 | 0.0003 | 0.0007 | 0.0011 | 0.0018 | N/A |
| 31 | narK5cI | 0.0000 | 0.0000 | 0.0013 | 0.0000 | 0.0002 | 0.0003 | 0.0000 | 0.0003 | 0.0000 | 0.0018 | 0.0000 | 0.0000 | 0.0006 | 0.0004 | 0.0008 | N/A |
| 10 | fdnG15cI | N/A | 0.0000 | N/A | N/A | 0.0191 | 0.0000 | N/A | N/A | N/A | N/A | N/A | N/A | N/A | 0.0294 | 0.0000 | N/A |
| 12 | fdnG23cI | N/A | N/A | N/A | N/A | 0.0000 | N/A | N/A | N/A | N/A | N/A | N/A | N/A | N/A | N/A | N/A | N/A |
| 34 | narK17cI | N/A | N/A | N/A | N/A | 0.0000 | N/A | N/A | 0.0000 | N/A | N/A | N/A | N/A | N/A | N/A | N/A | N/A |
| 18 | hycA23cI | 0.0000 | N/A | N/A | N/A | 0.0019 | 0.0000 | 0.0000 | 0.0150 | N/A | N/A | N/A | N/A | 0.0000 | 0.0000 | 0.0000 | N/A |
| 15 | hycA10cI | N/A | N/A | N/A | N/A | 0.0000 | N/A | N/A | 0.0000 | N/A | N/A | N/A | N/A | N/A | N/A | N/A | N/A |
| 48 | yedY5cI | N/A | N/A | N/A | N/A | 0.0000 | 0.0000 | 0.0000 | 0.0000 | N/A | N/A | N/A | N/A | N/A | N/A | 0.0000 | N/A |
| 33 | narK15cI | 0.0000 | 0.0000 | N/A | N/A | 0.0099 | 0.0000 | 0.0000 | 0.0062 | N/A | N/A | N/A | N/A | 0.0000 | 0.0000 | 0.0000 | N/A |
| 11 | fdnG17cI | N/A | N/A | N/A | N/A | N/A | N/A | N/A | N/A | N/A | N/A | N/A | N/A | N/A | N/A | N/A | N/A |
| 46 | yeaR17cI | N/A | N/A | N/A | N/A | 0.0294 | N/A | N/A | N/A | N/A | N/A | N/A | N/A | N/A | N/A | N/A | N/A |
| 43 | yeaR5cI | N/A | N/A | N/A | N/A | 0.0273 | N/A | N/A | N/A | N/A | N/A | N/A | N/A | N/A | N/A | N/A | N/A |
| 9 | fdnG10cI | N/A | N/A | N/A | N/A | 0.0000 | N/A | N/A | N/A | N/A | N/A | N/A | N/A | N/A | N/A | N/A | N/A |
| 36 | narG23cI | N/A | N/A | N/A | N/A | N/A | N/A | N/A | N/A | N/A | N/A | N/A | N/A | N/A | N/A | N/A | N/A |
| 1 | cydAWTcI | 0.2349 | 0.1625 | 0.1751 | 0.2068 | 0.1295 | 0.1792 | 0.1163 | 0.1475 | 0.1631 | 0.2778 | 0.3582 | 0.1877 | 0.2620 | 0.1992 | 0.3189 | N/A |
| 2 | cydA5cI | N/A | N/A | N/A | N/A | 0.0191 | N/A | N/A | N/A | N/A | N/A | N/A | N/A | N/A | N/A | N/A | N/A |
| 26 | napF10cI | 0.1744 | N/A | N/A | 0.1628 | 0.1385 | 0.3176 | 0.1271 | 0.1657 | N/A | N/A | N/A | N/A | N/A | 0.2921 | 0.1541 | N/A |
| 28 | napF17cI | 0.0399 | N/A | N/A | N/A | 0.0868 | 0.0487 | N/A | 0.0640 | N/A | N/A | N/A | N/A | N/A | N/A | 0.0167 | N/A |
| 7 | fdnGWTcI | 0.2178 | 0.1722 | 0.1515 | 0.1619 | 0.1114 | 0.1676 | 0.1028 | 0.1483 | 0.1288 | 0.2084 | 0.2278 | 0.1837 | 0.2088 | 0.1959 | 0.2424 | N/A |
| 27 | napF15cI | 0.1970 | 0.2454 | 0.2280 | 0.0989 | 0.1094 | 0.1249 | 0.0971 | 0.1344 | 0.0747 | 0.1239 | 0.1451 | 0.1115 | 0.2024 | 0.2083 | 0.1622 | N/A |
| 25 | napF5cI | 0.1824 | 0.2111 | 0.1884 | 0.0984 | 0.1181 | 0.1468 | 0.1026 | 0.1644 | 0.0715 | 0.1203 | 0.2036 | 0.1210 | 0.1956 | 0.1810 | 0.2006 | N/A |
| 29 | napF23cI | 0.1872 | 0.1916 | 0.1627 | 0.1135 | 0.1237 | 0.1506 | 0.1149 | 0.1568 | 0.1064 | 0.1846 | 0.2153 | 0.1401 | 0.2106 | 0.2231 | 0.2540 | N/A |
| 4 | cydA15cI | 0.1495 | N/A | N/A | N/A | 0.1799 | 0.2090 | 0.1038 | 0.1438 | N/A | N/A | N/A | N/A | 0.3103 | 0.0660 | 0.1052 | N/A |
| 20 | moaA5cI | 0.1910 | 0.2653 | 0.2244 | 0.1898 | 0.1566 | 0.2118 | 0.1306 | 0.1766 | 0.1844 | 0.2301 | 0.3138 | 0.2035 | 0.2356 | 0.2704 | 0.2456 | N/A |
| 22 | moaA15cI | 0.2226 | 0.2330 | 0.2117 | 0.1853 | 0.1573 | 0.1963 | 0.1332 | 0.1810 | 0.1693 | 0.2584 | 0.2808 | 0.2210 | 0.2460 | 0.2509 | 0.3415 | N/A |
| 21 | moaA10cI | 0.1993 | 0.2476 | 0.2252 | 0.2128 | 0.1391 | 0.2023 | 0.1324 | 0.1710 | 0.1565 | 0.4059 | 0.2071 | 0.2869 | 0.2301 | 0.2651 | 0.3170 | N/A |
| 5 | cydA17cI | 0.3092 | 0.2271 | 0.1235 | 0.2281 | 0.1763 | 0.2049 | 0.1332 | 0.2256 | 0.1405 | 0.2218 | 0.2153 | 0.3444 | 0.2628 | 0.2307 | 0.3570 | N/A |
| 6 | cydA23cI | 0.2436 | 0.1460 | 0.1953 | 0.1812 | 0.1520 | 0.2047 | 0.1341 | 0.2163 | 0.1395 | 0.2938 | 0.2241 | 0.2070 | 0.2467 | 0.2474 | 0.3452 | N/A |
| 24 | moaA23cI | 0.2496 | 0.2120 | 0.2379 | 0.2132 | 0.1530 | 0.1865 | 0.1217 | 0.1993 | 0.1679 | 0.2715 | 0.3379 | 0.2041 | 0.2992 | 0.2424 | 0.2805 | N/A |
| 3 | cydA10cI | 0.2830 | 0.1961 | 0.1392 | 0.1867 | 0.1578 | 0.2253 | 0.1522 | 0.2342 | 0.1460 | 0.3066 | 0.4104 | 0.3202 | 0.2704 | 0.2307 | 0.3515 | N/A |
| 23 | moaA17cI | 0.3431 | N/A | N/A | 0.5962 | 0.2463 | 0.5207 | 0.2283 | 0.2601 | 0.2140 | N/A | N/A | 0.2937 | 0.4119 | 0.2847 | 0.4545 | N/A |
| 59 | fabR | 1.0000 | 1.0000 | 1.0000 | 1.0000 | 1.0000 | 1.0000 | 1.0000 | 1.0000 | 1.0000 | 1.0000 | 1.0000 | 1.0000 | 1.0000 | 1.0000 | 1.0000 | N/A |

| | Day | 5 | | | | | | | | | | | | |
|---|---|---|---|---|---|---|---|---|---|---|---|---|---|---|
| | | -DSS | | | | +DSS | | | | (-DSS) | | (+DSS) | | (+DSS)-(-DSS) |
| | Promoter | M1-11-18 | M2-11-18 | M3-11-18 | M4-11-18 | M5-11-18 | M6-11-18 | M7-11-18 | M8-11-18 | #(OR>.02 & p<.05) / #non-N/A | AVG OR (-DSS) | #(OR>.02 & p<.05) / #non-N/A | AVG OR (+DSS) | #(OR>.02 & p<.05) / #non-N/A |
| 41 | torC23cI | 0.0634 | 0.0433 | 0.0553 | 0.0381 | 0.1127 | 0.0760 | 0.0402 | 0.1190 | 1 | 0.1414 | 1 | 0.1630 | 0 |
| 55 | ynfE10cI | 0.0774 | 0.0926 | 0.0591 | 0.0722 | 0.1789 | 0.1068 | 0.0627 | 0.1175 | 1 | 0.1336 | 1 | 0.1854 | 0 |
| 56 | ynfE15cI | 0.0548 | 0.0590 | 0.0609 | 0.0843 | 0.1411 | 0.1078 | 0.0608 | 0.1269 | 1 | 0.1133 | 1 | 0.1795 | 0 |
| 39 | torC15cI | 0.0265 | 0.0327 | 0.0270 | 0.0318 | 0.1105 | 0.0645 | 0.0370 | 0.1104 | 1 | 0.0947 | 1 | 0.1202 | 0 |
| 37 | torC5cI | 0.0253 | 0.0308 | 0.0305 | 0.0268 | 0.0958 | 0.0514 | 0.0320 | 0.1032 | 1 | 0.0907 | 1 | 0.1159 | 0 |
| 42 | yeaRWTcI | 0.0378 | 0.0305 | 0.0300 | 0.0342 | 0.1349 | 0.0955 | 0.0368 | 0.1076 | 0.9375 | 0.0574 | 0.8 | 0.0724 | -0.1375 |
| 58 | ynfE23cI | 0.0266 | 0.0289 | 0.0349 | 0.0323 | 0.1156 | 0.0881 | 0.0442 | 0.1048 | 0.9375 | 0.0544 | 1 | 0.1427 | 0.0625 |
| 38 | torC10cI | 0.0171 | 0.0375 | 0.0351 | 0.0254 | 0.0803 | 0.0837 | 0.0453 | 0.1646 | 0.6875 | 0.1121 | 0.933333 | 0.1556 | 0.24583333 |
| 44 | yeaR10cI | 0.0166 | 0.0225 | 0.0045 | 0.0238 | 0.0754 | 0.0823 | 0.0202 | 0.0693 | 0.5625 | 0.0386 | 0.533333 | 0.0474 | -0.02916667 |
| 45 | yeaR15cI | 0.0151 | 0.0240 | 0.0161 | 0.0605 | 0.0409 | 0.0662 | 0.0183 | 0.0684 | 0.5 | 0.0435 | 0.466667 | 0.0461 | -0.03333333 |
| 40 | torC17cI | 0.0214 | 0.0187 | 0.0085 | N/A | 0.0371 | 0.0628 | 0.0125 | 0.1610 | 0.333333 | 0.0609 | 0.636364 | 0.0853 | 0.3030303 |
| 47 | yeaR23cI | N/A | N/A | N/A | N/A | 0.0220 | N/A | 0.0146 | 0.0895 | 0 | 0.0425 | 0.5 | 0.0465 | 0.5 |
| 13 | hycAWTcI | 0.0049 | 0.0031 | 0.0073 | 0.0036 | 0.0151 | 0.0054 | 0.0062 | 0.0217 | 0 | 0.0118 | 0.066667 | 0.0139 | 0.06666667 |
| 16 | hycA15cI | 0.0000 | 0.0017 | 0.0054 | 0.0016 | 0.0023 | 0.0019 | 0.0002 | 0.0040 | 0 | 0.0035 | 0 | 0.0039 | 0 |
| 32 | narK10cI | N/A | N/A | N/A | N/A | N/A | N/A | 0.0000 | 0.0000 | 0 | 0.0034 | 0 | 0.0000 | 0 |
| 57 | ynfE17cI | 0.0013 | 0.0043 | 0.0048 | 0.0000 | 0.0381 | 0.0381 | 0.0136 | 0.0423 | 0 | 0.0029 | 0.933333 | 0.0664 | 0.93333333 |
| 8 | fdnG5cI | 0.0000 | 0.0000 | 0.0014 | 0.0057 | 0.0027 | 0.0035 | 0.0006 | 0.0037 | 0 | 0.0028 | 0 | 0.0018 | 0 |
| 30 | narKWTcI | 0.0006 | 0.0015 | 0.0003 | 0.0000 | 0.0018 | 0.0002 | 0.0005 | 0.0030 | 0 | 0.0013 | 0 | 0.0014 | 0 |
| 54 | ynfE5cI | 0.0000 | 0.0009 | 0.0025 | 0.0000 | 0.0021 | 0.0003 | 0.0003 | 0.0003 | 0 | 0.0012 | 0 | 0.0010 | 0 |
| 49 | yedY10cI | 0.0000 | 0.0000 | 0.0000 | 0.0000 | 0.0000 | 0.0000 | 0.0002 | 0.0001 | 0 | 0.0007 | 0 | 0.0004 | 0 |
| 50 | yedY15cI | 0.0000 | 0.0003 | 0.0000 | 0.0000 | 0.0002 | 0.0000 | 0.0000 | 0.0002 | 0 | 0.0007 | 0 | 0.0005 | 0 |
| 17 | hycA17cI | 0.0000 | 0.0000 | 0.0000 | 0.0000 | 0.0005 | 0.0004 | 0.0000 | 0.0002 | 0 | 0.0006 | 0 | 0.0003 | 0 |
| 35 | narK23cI | 0.0013 | 0.0000 | 0.0000 | 0.0000 | 0.0005 | 0.0016 | 0.0008 | 0.0023 | 0 | 0.0005 | 0 | 0.0034 | 0 |
| 51 | yedY17cI | 0.0007 | 0.0000 | 0.0000 | 0.0000 | 0.0000 | 0.0000 | 0.0000 | 0.0000 | 0 | 0.0005 | 0 | 0.0007 | 0 |
| 14 | hycA5cI | 0.0010 | 0.0000 | 0.0015 | 0.0015 | 0.0004 | 0.0000 | 0.0000 | 0.0002 | 0 | 0.0004 | 0 | 0.0004 | 0 |
| 19 | moaAWTcI | 0.0000 | 0.0000 | 0.0006 | 0.0000 | 0.0006 | 0.0000 | 0.0003 | 0.0003 | 0 | 0.0004 | 0 | 0.0004 | 0 |
| 52 | yedY23cI | 0.0000 | 0.0000 | 0.0004 | 0.0003 | 0.0002 | 0.0003 | 0.0000 | 0.0002 | 0 | 0.0003 | 0 | 0.0004 | 0 |
| 53 | ynfEWTcI | 0.0003 | 0.0000 | 0.0001 | 0.0003 | 0.0003 | 0.0002 | 0.0001 | 0.0005 | 0 | 0.0003 | 0 | 0.0008 | 0 |
| 31 | narK5cI | 0.0000 | 0.0000 | 0.0000 | 0.0000 | 0.0003 | 0.0004 | 0.0002 | 0.0000 | 0 | 0.0002 | 0 | 0.0003 | 0 |
| 10 | fdnG15cI | N/A | N/A | N/A | N/A | 0.0000 | 0.0000 | 0.0000 | 0.0067 | 0 | 0.0000 | 0 | 0.0060 | 0 |
| 12 | fdnG23cI | N/A | N/A | N/A | N/A | N/A | N/A | N/A | 0.0112 | 0 | 0.0000 | 0 | 0.0028 | 0 |
| 34 | narK17cI | N/A | N/A | N/A | N/A | N/A | N/A | 0.0000 | 0.0000 | 0 | 0.0000 | 0 | 0.0017 | 0 |
| 18 | hycA23cI | N/A | 0.0000 | N/A | N/A | 0.0000 | 0.0000 | 0.0000 | 0.0000 | 0 | 0.0000 | 0 | 0.0015 | 0 |
| 15 | hycA10cI | N/A | N/A | N/A | N/A | N/A | N/A | 0.0000 | 0.0000 | 0 | 0.0000 | 0 | 0.0000 | 0 |
| 48 | yedY5cI | N/A | N/A | N/A | N/A | 0.0000 | N/A | 0.0000 | 0.0000 | 0 | 0.0000 | 0 | 0.0000 | 0 |
| 33 | narK15cI | N/A | N/A | 0.0000 | N/A | 0.0000 | 0.0000 | 0.0000 | 0.0000 | 0 | 0.0000 | 0 | 0.0023 | 0 |
| 11 | fdnG17cI | N/A | N/A | N/A | N/A | N/A | N/A | N/A | N/A | | #DIV/0! | | #DIV/0! | |
| 46 | yeaR17cI | N/A | N/A | N/A | N/A | N/A | N/A | N/A | N/A | | #DIV/0! | 0 | 0.0294 | |
| 43 | yeaR5cI | N/A | N/A | N/A | N/A | N/A | N/A | N/A | N/A | | #DIV/0! | 0 | 0.0273 | |
| 9 | fdnG10cI | N/A | N/A | N/A | N/A | N/A | N/A | N/A | N/A | | #DIV/0! | 0 | 0.0000 | |
| 36 | narG23cI | N/A | N/A | N/A | N/A | N/A | N/A | N/A | N/A | 0 | 0.0531 | | #DIV/0! | |
| 1 | cydAWTcI | 0.0796 | 0.0985 | 0.0921 | 0.0659 | 0.1654 | 0.1129 | 0.1079 | 0.1412 | 1 | 0.1727 | 1 | 0.2066 | 0 |
| 2 | cydA5cI | N/A | N/A | N/A | N/A | N/A | N/A | N/A | N/A | 0 | 0.0104 | 0.666667 | 0.0966 | 0.66666667 |
| 26 | napF10cI | N/A | N/A | N/A | N/A | 0.0803 | N/A | 0.0466 | 0.0533 | 1 | 0.1614 | 0.923077 | 0.2200 | -0.07692308 |
| 28 | napF17cI | N/A | N/A | N/A | N/A | 0.0241 | N/A | 0.0146 | 0.0671 | 0.5 | 0.0887 | 0.5 | 0.0685 | 0 |
| 7 | fdnGWTcI | 0.0932 | 0.0765 | 0.0720 | 0.1038 | 0.1773 | 0.1204 | 0.0891 | 0.1354 | 1 | 0.1560 | 1 | 0.1853 | 0 |
| 27 | napF15cI | 0.0587 | 0.0518 | 0.0502 | 0.0414 | 0.1512 | 0.0939 | 0.0640 | 0.1153 | 1 | 0.1361 | 1 | 0.1819 | 0 |
| 25 | napF5cI | 0.0766 | 0.0582 | 0.0773 | 0.0634 | 0.1404 | 0.0895 | 0.0743 | 0.1379 | 1 | 0.1394 | 1 | 0.1817 | 0 |
| 29 | napF23cI | 0.0760 | 0.0708 | 0.0708 | 0.0705 | 0.1603 | 0.1112 | 0.0766 | 0.1259 | 1 | 0.1444 | 1 | 0.1886 | 0 |
| 4 | cydA15cI | N/A | N/A | N/A | 0.0286 | 0.1003 | N/A | 0.0477 | 0.2430 | 0.833333 | 0.1574 | 0.928571 | 0.2531 | 0.0952381 |
| 20 | moaA5cI | 0.0816 | 0.1093 | 0.0801 | 0.1004 | 0.1411 | 0.1214 | 0.0935 | 0.1498 | 1 | 0.1875 | 1 | 0.2265 | 0 |
| 22 | moaA15cI | 0.1199 | 0.1168 | 0.1069 | 0.1133 | 0.1927 | 0.1351 | 0.0931 | 0.1402 | 1 | 0.1912 | 1 | 0.2243 | 0 |
| 21 | moaA10cI | 0.1688 | 0.1408 | 0.0945 | 0.1475 | 0.1889 | 0.1516 | 0.0890 | 0.1296 | 1 | 0.2115 | 1 | 0.2257 | 0 |
| 5 | cydA17cI | 0.0930 | 0.1563 | 0.1003 | 0.1629 | 0.2486 | 0.1433 | 0.1106 | 0.1703 | 1 | 0.1956 | 1 | 0.2422 | 0 |
| 6 | cydA23cI | 0.1087 | 0.0998 | 0.1260 | 0.0993 | 0.1704 | 0.1371 | 0.0995 | 0.1695 | 1 | 0.1767 | 1 | 0.2278 | 0 |
| 24 | moaA23cI | 0.1079 | 0.1050 | 0.0879 | 0.1013 | 0.1988 | 0.1296 | 0.0799 | 0.1358 | 1 | 0.1999 | 1 | 0.2267 | 0 |
| 3 | cydA10cI | 0.1492 | 0.1216 | 0.0989 | 0.1480 | 0.2177 | 0.1689 | 0.1008 | 0.2236 | 1 | 0.2084 | 1 | 0.2476 | 0 |
| 23 | moaA17cI | N/A | N/A | N/A | N/A | 0.2605 | 0.4151 | 0.1642 | 0.2737 | 1 | 0.3924 | 1 | 0.4480 | 0 |
| 59 | fabR | 1.0000 | 1.0000 | 1.0000 | 1.0000 | 1.0000 | 1.0000 | 1.0000 | 1.0000 | 1 | 1.0000 | 1 | 1.0000 | 0 |

### Data Set S3 - Nissle 1917 library 5 day experiment (-DSS and +DSS)
### Odds Ratios & p-Values

| | Promoter | GV-11-13 | GV-11-13_pV | M1-11-14 | M1-11-14_pV | M2-11-14 | M2-11-14_pV | M3-11-14 | M3-11-14_pV | M4-11-14 | M4-11-14_pV | M5-11-14 | M5-11-14_pV | M6-11-14 | M6-11-14_pV |
|---|---|---|---|---|---|---|---|---|---|---|---|---|---|---|---|
| 1 | cydAWTcI | 0.02611864 | 0.00010198 | 0.14325589 | 9.58E-106 | 0.16018244 | 1.00E-52 | 0.14794419 | 4.39E-112 | 0.21003061 | 2.47E-108 | 0.2438183 | 5.41E-242 | 0.57795888 | 9.27E-182 |
| 2 | cydA5cI | 0.02972825 | 0.05458934 | 0.02071492 | 0.65384882 | N/A | 1 | N/A | 0.00420199 | 0 | 1 | 0.19350189 | 5.69E-05 | N/A | 1 |
| 3 | cydA10cI | 0.09557549 | 4.27E-93 | 0.16346819 | 4.33E-129 | 0.18893079 | 2.11E-72 | 0.16885408 | 3.36E-131 | 0.30715866 | 5.90E-217 | 0.33800926 | 2.359706931 | 0.60597163 | 6.50E-221 |
| 4 | cydA15cI | 0.07517839 | 2.56E-26 | 0.20676575 | 7.24E-13 | 0.27116082 | 2.22E-08 | 0.12439273 | 2.75E-10 | 0.16384434 | 9.01E-10 | 1.22945783 | 1.01E-67 | 0.53313298 | 6.17E-09 |
| 5 | cydA17cI | 0.08745497 | 9.53E-86 | 0.14353695 | 1.26E-121 | 0.18009556 | 3.90E-90 | 0.16884798 | 1.71E-135 | 0.31396476 | 1.86E-242 | 0.29047706 | 3.38E-301 | 0.66898977 | 1.85E-254 |
| 6 | cydA23cI | 0.08821836 | 3.22E-86 | 0.14513978 | 1.18E-127 | 0.18221881 | 4.04E-83 | 0.14827537 | 1.85E-124 | 0.28676215 | 1.30E-219 | 0.2701662 | 4.97E-287 | 0.60462299 | 3.24E-234 |
| 7 | fdnGWTcI | 0.05543458 | 2.41E-43 | 0.14113278 | 2.97E-107 | 0.18166346 | 5.36E-79 | 0.14650985 | 7.33E-122 | 0.22859203 | 4.41E-165 | 0.27262677 | 7.29E-270 | 0.45872467 | 7.90E-178 |
| 8 | fdnG5cI | 0.00011609 | 1 | 0.00022371 | 1 | 0.00328029 | 1 | 0.00020267 | 1 | 0.00593207 | 0.99999945 | 0.00060191 | 1 | 0 | 1 |
| 9 | fdnG10cI | 0.0008409 | 0.99999982 | N/A | 1 | N/A | 1 | N/A | 1 | N/A | 1 | N/A | 1 | N/A | 1 |
| 10 | fdnG15cI | 0 | 1 | 0 | 1 | N/A | 1 | 0 | 1 | 0 | 1 | 0 | 1 | N/A | 1 |
| 11 | fdnG17cI | 0 | 1 | N/A | 1 | N/A | 1 | N/A | 1 | N/A | 1 | N/A | 1 | N/A | 1 |
| 12 | fdnG23cI | 0 | 1 | 0 | 1 | N/A | 1 | 0 | 1 | 0 | 1 | 0 | 1 | N/A | 1 |
| 13 | hycAWTcI | 9.25E-05 | 1 | 0.01413287 | 0.99741362 | 0.00459624 | 0.99999996 | 0.00371973 | 1 | 0.00591138 | 1 | 0.01426681 | 0.99966749 | 0.04693447 | 3.52E-06 |
| 14 | hycA5cI | 0 | 1 | 0 | 1 | 0 | 1 | 0.00026926 | 1 | 0.00042504 | 1 | 0.000869 | 1 | 0.00131608 | 0.99999965 |
| 15 | hycA10cI | 0 | 1 | N/A | 1 | N/A | 1 | N/A | 0.76055239 | 0 | 1 | N/A | 1 | N/A | 1 |
| 16 | hycA15cI | 0 | 1 | 0.00366225 | 1 | 0.00137439 | 1 | 0.00192583 | 1 | 0.00188621 | 1 | 0.00226506 | 1 | 0.01360797 | 0.91287109 |
| 17 | hycA17cI | 0 | 1 | 0 | 1 | 0.00100931 | 0.99999998 | 0.00096879 | 1 | 0.00060955 | 1 | 0.00036468 | 1 | 0 | 1 |
| 18 | hycA23cI | 0 | 1 | 0 | 1 | N/A | 1 | 0 | 1 | 0 | 1 | 0 | 1 | 0 | 1 |
| 19 | moaAWTcI | 6.10E-05 | 1 | 0.00115262 | 1 | 0 | 1 | 0.0003755 | 1 | 0.00081681 | 1 | 0.00034272 | 1 | 0 | 1 |
| 20 | moaA5cI | 0.08322278 | 4.69E-76 | 0.18714196 | 3.10E-121 | 0.18316417 | 1.66E-60 | 0.20962408 | 1.51E-184 | 0.24630356 | 5.13E-116 | 0.3246898 | 1.36E-276 | 0.70263171 | 4.29E-182 |
| 21 | moaA10cI | 0.08478248 | 6.61E-82 | 0.17558914 | 1.86E-135 | 0.19609925 | 1.01E-79 | 0.20879822 | 5.39E-161 | 0.31101286 | 3.06E-215 | 0.31500268 | 1.27E-305 | 0.63664516 | 4.95E-227 |
| 22 | moaA15cI | 0.08417127 | 2.00E-85 | 0.15764532 | 9.48E-166 | 0.19851291 | 4.93E-124 | 0.19049517 | 6.24E-196 | 0.27303394 | 1.22E-290 | 0.26787421 | 2.070869702 | 0.61602782 | 7.13E-297 |
| 23 | moaA17cI | 0.16783317 | 1.41E-125 | 0.3690933 | 2.19E-70 | 0.35081836 | 2.72E-28 | 0.45882553 | 3.15E-62 | 0.51380861 | 2.90E-69 | 0.62933326 | 3.79E-205 | 1.6569488 | 1.59E-74 |
| 24 | moaA23cI | 0.08925777 | 1.74E-88 | 0.17896962 | 4.94E-159 | 0.2180974 | 6.78E-121 | 0.22376412 | 1.53E-185 | 0.28191635 | 4.31E-254 | 0.302275 | 1.964817019 | 0.6438796 | 2.39E-267 |
| 25 | napF5cI | 0.07226241 | 4.45E-64 | 0.12769792 | 5.39E-104 | 0.16988033 | 4.91E-92 | 0.1410612 | 1.31E-100 | 0.3199861 | 1.54E-236 | 0.2468086 | 5.32E-238 | 0.56724966 | 3.93E-223 |
| 26 | napF10cI | 0.0348697 | 0.00347389 | 0.15510957 | 7.29E-10 | 0.13028094 | 0.00336797 | 0.0966619 | 9.11E-05 | 0.2492815 | 5.77E-12 | 0.53728732 | 1.53E-21 | 0.53313298 | 6.17E-09 |
| 27 | napF15cI | 0.06584283 | 9.48E-54 | 0.12194299 | 3.19E-117 | 0.17811351 | 1.30E-110 | 0.14177367 | 2.09E-109 | 0.30891935 | 1.51E-263 | 0.24317591 | 1.00E-217 | 0.67041625 | 1.46E-257 |
| 28 | napF17cI | 0.05017459 | 0.00010935 | 0.04314666 | 0.12876756 | N/A | 0.0034901 | 0.08056347 | 0.01027295 | 0.19113794 | 0.00021058 | 0.22691608 | 1.08E-09 | 0 | 1 |
| 29 | napF23cI | 0.07260189 | 2.27E-69 | 0.13203628 | 2.31E-156 | 0.15851147 | 6.02E-120 | 0.14007609 | 2.53E-177 | 0.29076223 | 0 | 0.23627085 | 1.21E-294 | 0.54235221 | 8.20E-306 |
| 30 | narKWTcI | 0 | 1 | 0.00082099 | 1 | 0.00031604 | 1 | 0.00132794 | 1 | 0.00027106 | 1 | 0.00059624 | 1 | 0.00374008 | 0.99999981 |
| 31 | narK5cI | 7.45E-05 | 1 | 0.00024843 | 1 | 0 | 1 | 0.00021044 | 1 | 0 | 1 | 0.00019284 | 1 | 0.00088025 | 1 |
| 32 | narK10cI | 0.00040059 | 1 | N/A | 1 | N/A | 1 | 0.00672209 | 0.94242553 | 0 | 1 | N/A | 1 | N/A | 1 |
| 33 | narK15cI | 0.00024341 | 1 | 0 | 1 | N/A | 1 | 0 | 1 | 0 | 1 | 0 | 1 | 0 | 1 |
| 34 | narK17cI | 0 | 1 | N/A | 1 | N/A | 1 | N/A | 1 | 0 | 1 | N/A | 0.49473471 | N/A | 1 |
| 35 | narK23cI | 0 | 1 | 0.0008092 | 1 | 0 | 1 | 0.00109092 | 1 | 0.00047201 | 1 | 0.00648942 | 1 | 0.00821725 | 0.99517441 |
| 36 | narG23cI | 0.02246996 | 0.46921851 | N/A | 1 | N/A | 1 | N/A | 0.57605633 | 0.05310601 | 0.34735358 | N/A | 1 | N/A | 0.00026213 |
| 37 | torC5cI | 0.00251581 | 1 | 0.08728916 | 2.67E-101 | 0.11075239 | 2.57E-86 | 0.10412822 | 5.17E-132 | 0.20465534 | 1.92E-264 | 0.1830282 | 5.45E-242 | 0.43004373 | 1.16E-270 |
| 38 | torC10cI | 0.00488617 | 0.99984788 | 0.07759176 | 4.81E-05 | 0.11427003 | 3.63E-07 | 0.20439595 | 5.65E-16 | 0.3070886 | 1.08E-18 | 0.23450717 | 5.12E-12 | 0.35558645 | 2.92E-12 |
| 39 | torC15cI | 0.00360089 | 1 | 0.08476826 | 9.83E-102 | 0.12169368 | 4.46E-97 | 0.11219928 | 2.25E-146 | 0.22262187 | 3.75E-290 | 0.17928935 | 1.68E-243 | 0.42377529 | 4.32E-273 |
| 40 | torC17cI | 0.01322673 | 0.86810972 | 0.03697327 | 0.18441044 | 0.06035551 | 0.05473392 | 0.03358732 | 0.2558193 | 0.06710311 | 0.04429147 | N/A | 5.16E-10 | N/A | 8.62E-07 |
| 41 | torC23cI | 0.00505965 | 0.99999979 | 0.10647853 | 4.88E-34 | 0.38674113 | 1.67E-93 | 0.15893216 | 3.76E-41 | 0.20537343 | 1.98E-58 | 0.58330448 | 5.81E-45 | 0.46548198 | 1.54E-43 |
| 42 | yeaRWTcI | 0.00028361 | 1 | 0.03019744 | 1.36E-07 | 0.01686229 | 0.90592881 | 0.05396763 | 2.65E-45 | 0.07043307 | 1.43E-56 | 0.02601158 | 4.75E-05 | 0.06712826 | 8.54E-25 |
| 43 | yeaR5cI | 0 | 1 | N/A | 1 | N/A | 1 | N/A | 1 | N/A | 1 | N/A | 1 | N/A | 1 |
| 44 | yeaR10cI | 0 | 1 | 0.02769241 | 0.01066599 | 0.00243766 | 0.99999551 | 0.01883518 | 0.67369021 | 0.03312558 | 0.00677999 | 0.02012536 | 0.50281028 | 0.04092612 | 0.0119216 |
| 45 | yeaR15cI | 0 | 1 | 0.01616115 | 0.93386669 | 0.00679787 | 0.99902387 | 0.02304669 | 0.12812618 | 0.040395 | 2.79E-06 | 0.01573888 | 0.9772073 | 0.02695588 | 0.19164187 |
| 46 | yeaR17cI | 0 | 1 | N/A | 1 | N/A | 1 | N/A | 1 | N/A | 1 | N/A | 1 | N/A | 1 |
| 47 | yeaR23cI | 0 | 1 | 0 | 1 | N/A | 1 | N/A | 0.46331146 | 0.12744413 | 0.06032928 | 0.01075913 | 0.8466653 | N/A | 1 |
| 48 | yedY5cI | 0 | 1 | 0 | 1 | N/A | 1 | 0 | 1 | 0 | 1 | 0 | 1 | N/A | 1 |
| 49 | yedY10cI | 0 | 1 | 0.00050062 | 1 | 0.00059384 | 1 | 0 | 1 | 0.00117596 | 1 | 0.00063895 | 1 | 0 | 1 |
| 50 | yedY15cI | 0 | 1 | 0.00030207 | 1 | 0.00035184 | 1 | 0.0006417 | 1 | 0.00028755 | 1 | 0.00049297 | 1 | 0.00157116 | 1 |
| 51 | yedY17cI | 7.12E-05 | 1 | 0.0001372 | 1 | 0 | 1 | 0.00073404 | 1 | 0.00045802 | 1 | 0.0003359 | 1 | 0.00472516 | 0.99992957 |
| 52 | yedY23cI | 6.14E-05 | 1 | 0.00010024 | 1 | 0.00029946 | 1 | 0.00056824 | 1 | 0.00096064 | 1 | 0.00031995 | 1 | 0.00110103 | 1 |
| 53 | ynfEWTcI | 0 | 1 | 0.00023769 | 1 | 0.00056015 | 1 | 0.00060638 | 1 | 0.00020976 | 1 | 0.00142802 | 1 | 0.00205192 | 1 |
| 54 | ynfE5cI | 0 | 1 | 0.0006001 | 1 | 0.00080118 | 1 | 0.00124997 | 1 | 0.00141817 | 1 | 0.0018365 | 1 | 0 | 1 |
| 55 | ynfE10cI | 0 | 1 | 0.14256497 | 1.79E-137 | 0.08505218 | 6.89E-44 | 0.10263284 | 2.98E-121 | 0.21664563 | 6.15E-209 | 0.261518 | 1.11E-302 | 0.52466817 | 1.16E-257 |
| 56 | ynfE15cI | 0.00011592 | 1 | 0.1101403 | 2.54E-97 | 0.05646924 | 1.56E-17 | 0.07739425 | 8.18E-82 | 0.18442082 | 1.07E-160 | 0.25746387 | 1.13E-286 | 0.55391784 | 2.43E-242 |
| 57 | ynfE17cI | 9.04E-05 | 1 | 0.0055374 | 1 | 0.00070285 | 1 | 0.00908241 | 0.99999989 | 0.00249447 | 1 | 0.11028514 | 3.50E-97 | 0.14082542 | 3.89E-31 |
| 58 | ynfE23cI | 0 | 1 | 0.05280615 | 3.47E-39 | 0.01230556 | 0.99994372 | 0.07175273 | 1.87E-66 | 0.05011911 | 1.10E-28 | 0.20777361 | 2.49E-261 | 0.39337802 | 2.93E-224 |
| 59 | fabR | 1 | 0 | 1 | 0 | 1 | 5.65E-256 | 1 | 0 | 1 | 0 | 1 | 0 | 1 | 3.680747348 |

| | Promoter | M7-11-14 | M7-11-14_pV | M8-11-14 | M8-11-14_pV | M1-11-15 | M1-11-15_pV | M2-11-15 | M2-11-15_pV | M3-11-15 | M3-11-15_pV | M4-11-15 | M4-11-15_pV |
|---|---|---|---|---|---|---|---|---|---|---|---|---|---|
| 1 | cydAWTcI | 0.25553942 | 9.74E-122 | 0.14167871 | 3.54E-91 | 0.23493431 | 2.82E-57 | 0.16248538 | 6.90E-18 | 0.17513656 | 3.24E-28 | 0.20678564 | 2.44E-25 |
| 2 | cydA5cI | N/A | 1 | 0.07706727 | 0.00226621 | N/A | 1 | N/A | 1 | N/A | 1 | N/A | 1 |
| 3 | cydA10cI | 0.26144786 | 6.37E-142 | 0.17508978 | 1.83E-117 | 0.28304382 | 6.51E-69 | 0.19614656 | 5.60E-22 | 0.13915484 | 1.21E-18 | 0.18672981 | 6.16E-38 |
| 4 | cydA15cI | 0.13446013 | 3.52E-05 | 0.13669687 | 9.37E-21 | 0.14948948 | 0.00092226 | N/A | 0.00520899 | N/A | 0.00100312 | N/A | 8.86E-08 |
| 5 | cydA17cI | 0.2333891 | 6.24E-141 | 0.17664168 | 2.24E-120 | 0.30919118 | 1.04E-76 | 0.22708479 | 4.52E-31 | 0.12353929 | 6.85E-23 | 0.2280804 | 1.55E-53 |
| 6 | cydA23cI | 0.24869904 | 1.81E-142 | 0.17042292 | 1.49E-116 | 0.24355191 | 7.94E-83 | 0.14597348 | 4.33E-20 | 0.1952943 | 2.37E-34 | 0.18120901 | 1.02E-49 |
| 7 | fdnGWTcI | 0.23400908 | 1.38E-127 | 0.11526616 | 8.98E-78 | 0.21780669 | 3.16E-182 | 0.17216391 | 1.97E-86 | 0.15151317 | 3.03E-76 | 0.16191676 | 1.18E-72 |
| 8 | fdnG5cI | 0.00039752 | 1 | 0.00063471 | 1 | 0.00219352 | 0.99984138 | 0.00427537 | 0.98937642 | 0 | 1 | 0.0120668 | 0.9305969 |
| 9 | fdnG10cI | N/A | 1 | 0 | 1 | N/A | 1 | N/A | 1 | N/A | 1 | N/A | 1 |
| 10 | fdnG15cI | N/A | 1 | 0.00428991 | 0.98722622 | N/A | 1 | 0 | 1 | N/A | 1 | N/A | 1 |
| 11 | fdnG17cI | N/A | 1 | N/A | 1 | N/A | 1 | N/A | 1 | N/A | 1 | N/A | 1 |
| 12 | fdnG23cI | N/A | 1 | 0 | 1 | N/A | 1 | N/A | 1 | N/A | 1 | N/A | 1 |
| 13 | hycAWTcI | 0.01214129 | 0.99910935 | 0.00727268 | 1 | 0.02945094 | 0.08941373 | 0.01376089 | 0.87243223 | 0.00797857 | 0.99569451 | 0.02087746 | 0.47668003 |
| 14 | hycA5cI | 0 | 1 | 0.00037787 | 1 | 0 | 1 | 0 | 1 | 0 | 1 | 0 | 1 |
| 15 | hycA10cI | N/A | 1 | 0 | 1 | N/A | 1 | N/A | 1 | N/A | 1 | N/A | 0.15585275 |
| 16 | hycA15cI | 0.00464499 | 1 | 0.00263119 | 1 | 0.0043486 | 0.99864579 | 0 | 1 | 0.00223574 | 0.99978503 | 0.01356147 | 0.89645718 |
| 17 | hycA17cI | 0.00048449 | 1 | 0 | 1 | 0 | 1 | 0 | 1 | 0 | 1 | 0 | 1 |
| 18 | hycA23cI | 0 | 1 | 0.0051472 | 0.97468382 | 0 | 1 | N/A | 1 | N/A | 1 | N/A | 1 |
| 19 | moaAWTcI | 0 | 1 | 0.00023939 | 1 | 0 | 1 | 0 | 1 | 0 | 1 | 0 | 1 |
| 20 | moaA5cI | 0.27892063 | 2.09E-126 | 0.15856028 | 3.46E-101 | 0.19098706 | 4.71E-33 | 0.26531839 | 2.92E-27 | 0.22437416 | 1.07E-51 | 0.1898395 | 2.85E-38 |
| 21 | moaA10cI | 0.26433844 | 2.45E-143 | 0.15289661 | 7.78E-108 | 0.19928156 | 5.78E-67 | 0.24757095 | 2.90E-57 | 0.22522333 | 1.72E-47 | 0.21275785 | 3.87E-115 |
| 22 | moaA15cI | 0.2535347 | 9.71E-168 | 0.16032867 | 5.12E-120 | 0.22263774 | 1.88E-250 | 0.23301393 | 5.08E-192 | 0.21166334 | 3.66E-116 | 0.18527187 | 5.60E-113 |
| 23 | moaA17cI | 0.64695717 | 5.31E-64 | 0.26707444 | 8.47E-100 | 0.3431371 | 2.24E-20 | N/A | 7.72E-10 | N/A | 1.61E-05 | 0.59619214 | 1.30E-20 |
| 24 | moaA23cI | 0.26138981 | 8.78E-156 | 0.16608308 | 6.18E-118 | 0.24960473 | 1.42E-198 | 0.21199607 | 3.14E-155 | 0.23791837 | 3.77E-93 | 0.21322931 | 2.17E-104 |
| 25 | napF5cI | 0.21252156 | 1.49E-116 | 0.14702541 | 5.56E-99 | 0.18242212 | 2.23E-119 | 0.21109762 | 6.54E-107 | 0.18835627 | 1.49E-64 | 0.09839528 | 3.13E-35 |
| 26 | napF10cI | 0.13446013 | 3.52E-05 | 0.28039323 | 6.46E-22 | 0.17439197 | 0.00017893 | N/A | 3.27E-06 | N/A | 0.01496775 | 0.16278869 | 6.76E-07 |
| 27 | napF15cI | 0.21032564 | 4.01E-113 | 0.14127905 | 7.53E-92 | 0.19699902 | 4.04E-267 | 0.24540553 | 4.43E-240 | 0.22804786 | 2.82E-128 | 0.09890325 | 2.36E-58 |
| 28 | napF17cI | N/A | 0.05127275 | 0.13626471 | 3.86E-08 | 0.03988262 | 0.30067087 | N/A | 0.41181028 | N/A | 1 | N/A | 0.39828403 |
| 29 | napF23cI | 0.20872525 | 3.15E-158 | 0.13465011 | 1.13E-105 | 0.18719639 | 0 | 0.1915909 | 4.62E-277 | 0.16273954 | 2.22E-155 | 0.11346361 | 1.62E-97 |
| 30 | narKWTcI | 0.00065795 | 1 | 0.00068922 | 1 | 0 | 1 | 0.00428126 | 0.99984617 | 0.00362291 | 0.99997699 | 0.00059929 | 1 |
| 31 | narK5cI | 0 | 1 | 0.00028575 | 1 | 0 | 1 | 0 | 1 | 0.00131948 | 0.99999923 | 0 | 1 |
| 32 | narK10cI | N/A | 1 | 0 | 1 | N/A | 1 | N/A | 1 | N/A | 1 | N/A | 1 |
| 33 | narK15cI | 0.0149587 | 0.74997633 | 0.00285399 | 0.9983431 | 0 | 1 | 0 | 1 | N/A | 1 | N/A | 1 |
| 34 | narK17cI | N/A | 1 | 0.00856616 | 0.90006933 | N/A | 1 | N/A | 1 | N/A | 1 | N/A | 1 |
| 35 | narK23cI | 0.00298512 | 1 | 0.00357145 | 1 | 0 | 1 | 0 | 1 | 0.00274436 | 0.99901829 | 0 | 1 |
| 36 | narG23cI | N/A | 1 | N/A | 0.60290169 | N/A | 0.17419459 | N/A | 0.10074546 | N/A | 1 | N/A | 1 |
| 37 | torC5cI | 0.15855825 | 2.60E-128 | 0.09699063 | 1.64E-76 | 0.15128049 | 2.210123616 | 0.14093068 | 8.94E-206 | 0.13510982 | 1.03E-128 | 0.09100765 | 4.09E-75 |
| 38 | torC10cI | 0.14479901 | 1.79E-06 | 0.09033104 | 1.53E-08 | 0.22227736 | 3.60E-16 | 0.13542071 | 1.12E-10 | 0.15504377 | 3.39E-07 | 0.10855551 | 7.47E-06 |
| 39 | torC15cI | 0.15753989 | 2.94E-130 | 0.09856721 | 1.03E-78 | 0.161848 | 0 | 0.13677532 | 6.42E-216 | 0.14395018 | 2.21E-142 | 0.09278758 | 2.05E-80 |
| 40 | torC17cI | N/A | 0.0015009 | N/A | 6.79E-05 | 0.19929092 | 0.0001037 | 0.04879506 | 0.16117809 | N/A | 0.00110845 | 0.09306605 | 0.00937399 |
| 41 | torC23cI | 0.152522 | 3.91E-22 | 0.08075668 | 3.54E-16 | 0.15320425 | 9.67E-87 | 0.154508 | 1.33E-83 | 0.21542252 | 5.25E-57 | 0.10136034 | 1.70E-35 |
| 42 | yeaRWTcI | 0.02134321 | 0.27898044 | 0.01730886 | 0.96561864 | 0.05663842 | 1.40E-15 | 0.03166934 | 0.01081227 | 0.10266834 | 5.36E-39 | 0.09071684 | 1.67E-36 |
| 43 | yeaR5cI | N/A | 1 | N/A | 1 | N/A | 1 | N/A | 1 | N/A | 1 | N/A | 1 |
| 44 | yeaR10cI | 0.01812829 | 0.69366853 | 0.00892173 | 0.99999974 | 0.07974619 | 1.10E-06 | 0.02043831 | 0.56168228 | 0.06582286 | 0.00045557 | 0.05557414 | 0.00029057 |
| 45 | yeaR15cI | 0.01329166 | 0.98119622 | 0.00911496 | 1 | 0.04705617 | 0.00100025 | 0.0155553 | 0.75359429 | 0.02943763 | 0.14149969 | 0.15052113 | 6.27E-16 |
| 46 | yeaR17cI | N/A | 1 | N/A | 1 | N/A | 1 | N/A | 1 | N/A | 1 | N/A | 1 |
| 47 | yeaR23cI | N/A | 1 | N/A | 0.68462444 | 0 | 1 | N/A | 1 | N/A | 1 | N/A | 0.06545463 |
| 48 | yedY5cI | N/A | 1 | 0 | 1 | N/A | 1 | N/A | 1 | N/A | 1 | N/A | 1 |
| 49 | yedY10cI | 0.00067136 | 1 | 0.00062152 | 1 | 0 | 1 | 0.00256004 | 0.99943621 | 0 | 1 | 0.00397955 | 0.99990031 |
| 50 | yedY15cI | 0.00064128 | 1 | 0.0002503 | 1 | 0.00192872 | 0.99999994 | 0 | 1 | 0 | 1 | 0.00225571 | 0.99999999 |
| 51 | yedY17cI | 0.00064564 | 1 | 0.00016688 | 1 | 0 | 1 | 0 | 1 | 0.00145164 | 0.99999733 | 0.00119796 | 0.99999974 |
| 52 | yedY23cI | 0.00023709 | 1 | 0.00024391 | 1 | 0.00036638 | 1 | 0 | 1 | 0 | 1 | 0 | 1 |
| 53 | ynfEWTcI | 0.00051782 | 1 | 0.00065165 | 1 | 0 | 1 | 0.00037399 | 1 | 0.00028671 | 1 | 0.00019718 | 1 |
| 54 | ynfE5cI | 0.00065031 | 1 | 0.00093625 | 1 | 0.00176606 | 0.9999801 | 0 | 1 | 0 | 1 | 0.00112367 | 0.9999999 |
| 55 | ynfE10cI | 0.21281262 | 1.18E-143 | 0.12782345 | 5.82E-99 | 0.16579162 | 6.53E-77 | 0.16013643 | 3.17E-44 | 0.15763713 | 4.77E-73 | 0.12781213 | 2.58E-53 |
| 56 | ynfE15cI | 0.20318921 | 5.07E-129 | 0.12466556 | 3.67E-94 | 0.15325204 | 1.05E-82 | 0.12499801 | 7.56E-34 | 0.14120854 | 2.43E-58 | 0.1296868 | 1.65E-50 |
| 57 | ynfE17cI | 0.06021475 | 7.49E-18 | 0.07785782 | 8.12E-45 | 0.00246474 | 0.99959167 | 0 | 1 | 0.00568487 | 0.99089061 | 0.00502008 | 0.99894595 |
| 58 | ynfE23cI | 0.15673241 | 4.02E-114 | 0.11431258 | 1.10E-89 | 0.07945905 | 9.40E-38 | 0.04387711 | 2.92E-06 | 0.09712633 | 2.84E-34 | 0.05532128 | 4.50E-15 |
| 59 | fabR | 1 | 2.82E-283 | 1 | 3.51E-262 | 1 | 0 | 1 | 0 | 1 | 0 | 1 | 2.88E-275 |

| | Promoter | M5-11-15 | M5-11-15_pV | M6-11-15 | M6-11-15_pV | M7-11-15 | M7-11-15_pV | M8-11-15 | M8-11-15_pV | M1-11-17 | M1-11-17_pV | M2-11-17 | M2-11-17_pV | M3-11-17 | M3-11-17_pV |
|---|---|---|---|---|---|---|---|---|---|---|---|---|---|---|---|
| 1 | cydAWTcI | 0.12949299 | 2.68E-125 | 0.17918823 | 1.69E-80 | 0.11625717 | 1.15E-45 | 0.14745238 | 5.73E-59 | 0.16313808 | 1.09E-17 | 0.27776234 | 2.13E-22 | 0.35823063 | 1.10E-28 |
| 2 | cydA5cI | 0.01912499 | 0.64129046 | N/A | 0.08510666 | N/A | 0.78445623 | N/A | 0.02558144 | N/A | 1 | N/A | 1 | N/A | 1 |
| 3 | cydA10cI | 0.15775971 | 1.02E-173 | 0.22530339 | 4.29E-115 | 0.15215655 | 1.41E-66 | 0.23417938 | 1.81E-86 | 0.1459904 | 2.32E-16 | 0.306625 | 1.40E-22 | 0.41038519 | 2.59E-28 |
| 4 | cydA15cI | 0.17993123 | 1.85E-19 | 0.20903155 | 1.87E-12 | 0.10380596 | 2.76E-07 | 0.14384477 | 2.83E-09 | N/A | 4.11E-05 | N/A | 0.09212005 | N/A | 0.0008396 |
| 5 | cydA17cI | 0.17634506 | 1.72E-193 | 0.20490486 | 2.36E-113 | 0.13322354 | 4.34E-63 | 0.22563569 | 1.90E-87 | 0.14050655 | 7.45E-17 | 0.22181519 | 8.13E-23 | 0.21534304 | 9.08E-25 |
| 6 | cydA23cI | 0.15199624 | 7.86E-169 | 0.20472978 | 8.19E-113 | 0.13412042 | 1.10E-61 | 0.21631588 | 7.40E-84 | 0.13945648 | 8.06E-19 | 0.29384381 | 9.12E-29 | 0.22411961 | 1.16E-19 |
| 7 | fdnGWTcI | 0.11137628 | 1.08E-120 | 0.16763628 | 3.89E-98 | 0.10276955 | 2.41E-59 | 0.14830838 | 2.03E-64 | 0.12883575 | 8.43E-41 | 0.20839941 | 1.58E-45 | 0.22784941 | 2.30E-43 |
| 8 | fdnG5cI | 0.00075235 | 1 | 0.00051747 | 1 | 0.00125936 | 1 | 0.00074637 | 1 | 0 | 1 | 0.00415663 | 0.9986444 | 0.00203174 | 0.99986605 |
| 9 | fdnG10cI | 0 | 1 | N/A | 1 | N/A | 1 | N/A | 1 | N/A | 1 | N/A | 1 | N/A | 1 |
| 10 | fdnG15cI | 0.01912472 | 0.68047388 | 0 | 1 | N/A | 1 | N/A | 1 | N/A | 1 | N/A | 1 | N/A | 1 |
| 11 | fdnG17cI | N/A | 1 | N/A | 1 | N/A | 1 | N/A | 1 | N/A | 1 | N/A | 1 | N/A | 1 |
| 12 | fdnG23cI | 0 | 1 | N/A | 1 | N/A | 1 | N/A | 0.61336791 | N/A | 1 | N/A | 1 | N/A | 1 |
| 13 | hycAWTcI | 0.00526628 | 1 | 0.01524741 | 0.9608144 | 0.00946591 | 0.99999988 | 0.00776492 | 1 | 0.01014072 | 0.98787007 | 0.02211454 | 0.44146095 | 0.02064141 | 0.51283959 |
| 14 | hycA5cI | 0.00010343 | 1 | 0.00113043 | 1 | 0 | 1 | 0 | 1 | 0.00233565 | 0.99999511 | 0 | 1 | 0 | 1 |
| 15 | hycA10cI | 0 | 1 | N/A | 1 | N/A | 1 | 0 | 1 | N/A | 1 | N/A | 1 | N/A | 1 |
| 16 | hycA15cI | 0.0019937 | 1 | 0.00552831 | 0.99999992 | 0.00246593 | 1 | 0.00304655 | 1 | 0.00533154 | 0.9979022 | 0.00244871 | 0.99948782 | 0.00226015 | 0.99969035 |
| 17 | hycA17cI | 0.00015047 | 1 | 0 | 1 | 0 | 1 | 0.00019876 | 1 | 0 | 1 | 0 | 1 | 0.00248561 | 0.99927899 |
| 18 | hycA23cI | 0.00188368 | 0.99994479 | 0 | 1 | 0 | 1 | 0.01496025 | 0.76447874 | N/A | 1 | N/A | 1 | N/A | 1 |
| 19 | moaAWTcI | 8.16E-05 | 1 | 0.00118698 | 1 | 0.00015807 | 1 | 0.00018071 | 1 | 0.00111393 | 0.99999985 | 0.00139172 | 0.99999737 | 0.00123444 | 0.99999926 |
| 20 | moaA5cI | 0.15662807 | 1.23E-153 | 0.21175045 | 2.78E-99 | 0.13064569 | 1.49E-55 | 0.17663266 | 2.14E-68 | 0.18441285 | 1.16E-21 | 0.23007734 | 8.08E-22 | 0.31384407 | 1.13E-36 |
| 21 | moaA10cI | 0.13909154 | 2.29E-170 | 0.20225916 | 3.36E-126 | 0.13237602 | 3.16E-66 | 0.17103336 | 1.22E-74 | 0.15646433 | 1.41E-29 | 0.40591078 | 4.28E-51 | 0.20713688 | 5.97E-28 |
| 22 | moaA15cI | 0.15730146 | 1.86E-213 | 0.19626744 | 3.28E-150 | 0.13315219 | 7.22E-77 | 0.18103492 | 1.77E-83 | 0.16927105 | 2.34E-51 | 0.2584358 | 3.58E-56 | 0.28081013 | 1.16E-50 |
| 23 | moaA17cI | 0.24632027 | 1.02E-89 | 0.52065893 | 1.33E-44 | 0.2282501 | 9.47E-42 | 0.26012597 | 1.33E-54 | 0.2140018 | 1.18E-07 | N/A | 3.17E-06 | N/A | 7.83E-13 |
| 24 | moaA23cI | 0.15299718 | 2.24E-200 | 0.18649438 | 6.36E-132 | 0.12167922 | 7.82E-66 | 0.19928598 | 7.23E-85 | 0.16788406 | 1.06E-48 | 0.27148959 | 6.09E-55 | 0.33794333 | 1.12E-50 |
| 25 | napF5cI | 0.11814921 | 3.24E-117 | 0.14680497 | 5.62E-88 | 0.10255515 | 7.72E-48 | 0.16441122 | 1.72E-64 | 0.0714648 | 5.49E-14 | 0.12034548 | 8.23E-19 | 0.2036205 | 5.15E-26 |
| 26 | napF10cI | 0.13845086 | 9.99E-11 | 0.31763234 | 1.83E-11 | 0.12706533 | 2.55E-05 | 0.16570214 | 8.30E-08 | N/A | 0.21572105 | N/A | 0.0506681 | N/A | 0.00361234 |
| 27 | napF15cI | 0.10943594 | 3.00E-95 | 0.12491043 | 1.73E-79 | 0.09712452 | 1.67E-45 | 0.13443531 | 9.41E-50 | 0.07470349 | 4.36E-24 | 0.12388506 | 2.87E-31 | 0.14506421 | 8.54E-33 |
| 28 | napF17cI | 0.08683049 | 0.00139051 | 0.04871749 | 0.1216734 | N/A | 0.00168324 | 0.0640309 | 0.0402273 | N/A | 0.03133874 | N/A | 1 | N/A | 1 |
| 29 | napF23cI | 0.12371882 | 5.88E-180 | 0.15062581 | 1.57E-136 | 0.11486531 | 1.23E-71 | 0.15682654 | 6.87E-77 | 0.1063551 | 9.37E-42 | 0.18462483 | 2.45E-51 | 0.21527723 | 4.08E-50 |
| 30 | narKWTcI | 0.00021189 | 1 | 0.00066436 | 1 | 0.00042393 | 1 | 0.00079013 | 1 | 0.00157627 | 0.99999999 | 0.00304375 | 0.99999537 | 0.00250897 | 0.99998697 |
| 31 | narK5cI | 0.00017965 | 1 | 0.0003024 | 1 | 0 | 1 | 0.00032431 | 1 | 0 | 1 | 0.00182481 | 0.99995373 | 0 | 1 |
| 32 | narK10cI | 0 | 1 | N/A | 1 | N/A | 1 | 0 | 1 | N/A | 1 | N/A | 1 | N/A | 1 |
| 33 | narK15cI | 0.00988239 | 0.9321618 | 0 | 1 | 0 | 1 | 0.00623827 | 0.95360182 | N/A | 1 | N/A | 1 | N/A | 1 |
| 34 | narK17cI | 0 | 1 | N/A | 1 | N/A | 1 | 0 | 1 | N/A | 1 | N/A | 1 | N/A | 1 |
| 35 | narK23cI | 0.00266978 | 1 | 0.00547216 | 0.99999947 | 0.00189942 | 1 | 0.00270823 | 1 | 0.0021142 | 0.99982277 | 0 | 1 | 0 | 1 |
| 36 | narG23cI | N/A | 0.07979857 | N/A | 1 | N/A | 0.28727279 | N/A | 0.50991576 | N/A | 1 | N/A | 1 | N/A | 1 |
| 37 | torC5cI | 0.06486705 | 1.47E-84 | 0.11186437 | 7.55E-104 | 0.07305278 | 1.00E-43 | 0.11676055 | 3.17E-59 | 0.05129233 | 2.32E-16 | 0.07614053 | 6.20E-23 | 0.09892535 | 4.72E-27 |
| 38 | torC10cI | 0.0510784 | 0.0005544 | 0.18196505 | 3.12E-15 | 0.10773757 | 4.50E-08 | 0.30737742 | 6.51E-15 | 0.03291414 | 0.19029436 | 0.05503252 | 0.04706298 | 0.12478862 | 0.00028163 |
| 39 | torC15cI | 0.06836256 | 1.87E-92 | 0.11210206 | 1.59E-106 | 0.07776045 | 4.06E-48 | 0.11600458 | 9.41E-60 | 0.04995883 | 3.02E-16 | 0.07714659 | 4.68E-24 | 0.10098437 | 1.89E-28 |
| 40 | torC17cI | 0.05003639 | 0.0158795 | 0.22740394 | 2.85E-09 | 0.12046062 | 0.00018854 | 0.12789062 | 0.00011088 | 0.078206 | 0.03196447 | N/A | 0.05824903 | N/A | 0.26619996 |
| 41 | torC23cI | 0.06986927 | 1.76E-14 | 0.15901125 | 8.23E-62 | 0.09168439 | 1.09E-26 | 0.10912687 | 5.66E-18 | 0.09751033 | 1.52E-24 | 0.18025565 | 2.58E-32 | 0.17730052 | 1.18E-29 |
| 42 | yeaRWTcI | 0.07112702 | 1.25E-91 | 0.03605305 | 3.02E-10 | 0.02284155 | 0.09488323 | 0.03213481 | 1.07E-06 | 0.06023689 | 8.39E-13 | 0.07435992 | 4.73E-15 | 0.13253197 | 1.41E-23 |
| 43 | yeaR5cI | 0.02730744 | 0.47920626 | N/A | 1 | N/A | 1 | N/A | 1 | N/A | 1 | N/A | 1 | N/A | 1 |
| 44 | yeaR10cI | 0.0431447 | 5.14E-14 | 0.02303951 | 0.31317014 | 0.02270691 | 0.26727952 | 0.0205016 | 0.46953865 | 0.04536892 | 0.01581112 | 0.06015055 | 0.00152403 | 0.07495557 | 4.57E-05 |
| 45 | yeaR15cI | 0.08205772 | 1.21E-106 | 0.0245765 | 0.08828441 | 0.01320582 | 0.99396993 | 0.0160867 | 0.94166229 | 0.0371219 | 0.03333558 | 0.05669629 | 0.00076341 | 0.08141177 | 1.36E-05 |
| 46 | yeaR17cI | 0.02940775 | 0.33902705 | N/A | 1 | N/A | 1 | N/A | 0.28729659 | N/A | 1 | N/A | 1 | N/A | 1 |
| 47 | yeaR23cI | 0.0530477 | 0.00085843 | 0.01488818 | 0.75146867 | N/A | 0.70726261 | 0.07467254 | 0.01672588 | N/A | 0.17643567 | N/A | 1 | N/A | 0.02841158 |
| 48 | yedY5cI | 0 | 1 | 0 | 1 | 0 | 1 | 0 | 1 | N/A | 1 | N/A | 1 | N/A | 1 |
| 49 | yedY10cI | 9.50E-05 | 1 | 0.00066313 | 1 | 0.00047113 | 1 | 0.00037353 | 1 | 0 | 1 | 0 | 1 | 0 | 1 |
| 50 | yedY15cI | 0.00024173 | 1 | 0.00087122 | 1 | 0.00012377 | 1 | 0.00023464 | 1 | 0 | 1 | 0.00100254 | 0.99999997 | 0.00238459 | 0.99999282 |
| 51 | yedY17cI | 0.00020057 | 1 | 0.00096288 | 1 | 0.00046207 | 1 | 0.00021809 | 1 | 0 | 1 | 0.00166037 | 0.99998157 | 0.00181211 | 0.99995054 |
| 52 | yedY23cI | 9.98E-05 | 1 | 0.00048498 | 1 | 0.00014327 | 1 | 0.00019137 | 1 | 0.00041802 | 1 | 0.001008 | 1 | 0.00047408 | 1 |
| 53 | ynfEWTcI | 0.00050861 | 1 | 0.00106808 | 1 | 0.00049481 | 1 | 0.0005991 | 1 | 0.00061227 | 1 | 0.00026329 | 1 | 0.00025997 | 1 |
| 54 | ynfE5cI | 0.00078132 | 1 | 0.00216805 | 1 | 0.00019178 | 1 | 0.00110179 | 1 | 0 | 1 | 0 | 1 | 0 | 1 |
| 55 | ynfE10cI | 0.12524418 | 6.77E-171 | 0.16067637 | 6.85E-120 | 0.10860653 | 3.79E-61 | 0.18238382 | 1.68E-82 | 0.10541947 | 1.20E-24 | 0.23930558 | 2.13E-43 | 0.16226724 | 2.09E-31 |
| 56 | ynfE15cI | 0.11441853 | 6.59E-146 | 0.16529518 | 4.24E-113 | 0.1085903 | 1.54E-57 | 0.16136588 | 1.38E-72 | 0.12366042 | 4.89E-30 | 0.14032176 | 4.56E-29 | 0.18921364 | 9.65E-35 |
| 57 | ynfE17cI | 0.05569757 | 2.25E-33 | 0.07271118 | 1.55E-22 | 0.04022356 | 1.12E-07 | 0.10155511 | 1.24E-39 | 0 | 1 | 0 | 1 | 0.00558447 | 0.99029801 |
| 58 | ynfE23cI | 0.09657643 | 3.97E-132 | 0.13292493 | 3.50E-104 | 0.07633299 | 4.03E-42 | 0.1403436 | 4.68E-68 | 0.05683335 | 2.75E-12 | 0.06810932 | 1.12E-13 | 0.10338916 | 7.91E-21 |
| 59 | fabR | 1 | 0 | 1 | 1.23E-305 | 1 | 9.78E-200 | 1 | 5.65E-168 | 1 | 1.59E-121 | 1 | 1.38E-96 | 1 | 9.99E-86 |

| | Promoter | M4-11-17 | M4-11-17_pV | M5-11-17 | M5-11-17_pV | M6-11-17 | M6-11-17_pV | M7-11-17 | M7-11-17_pV | M1-11-18 | M1-11-18_pV | M2-11-18 | M2-11-18_pV |
|---|---|---|---|---|---|---|---|---|---|---|---|---|---|
| 1 | cydAWTcI | 0.18766645 | 4.02E-19 | 0.26196217 | 6.42E-36 | 0.1992354 | 3.02E-36 | 0.31894811 | 1.52E-45 | 0.07961655 | 1.37E-05 | 0.09850603 | 8.18E-10 |
| 2 | cydA5cI | N/A | 1 | N/A | 0.45698754 | N/A | 0.10289667 | N/A | 1 | N/A | 1 | N/A | 1 |
| 3 | cydA10cI | 0.32022024 | 2.01E-27 | 0.27044299 | 6.63E-38 | 0.2306838 | 1.12E-43 | 0.35145494 | 1.18E-49 | 0.14917903 | 1.04E-09 | 0.12162401 | 6.57E-14 |
| 4 | cydA15cI | N/A | 0.37596833 | 0.31025604 | 1.42E-12 | 0.06603497 | 0.0224762 | 0.10521436 | 0.00053658 | N/A | 0.10239987 | N/A | 0.02218989 |
| 5 | cydA17cI | 0.34440031 | 2.60E-33 | 0.26275455 | 7.95E-38 | 0.23070156 | 5.94E-46 | 0.35700825 | 1.18E-50 | 0.09302204 | 1.92E-09 | 0.15628454 | 3.78E-17 |
| 6 | cydA23cI | 0.20697375 | 4.00E-24 | 0.24668292 | 2.11E-36 | 0.24740397 | 4.70E-48 | 0.34519953 | 7.78E-50 | 0.10871655 | 4.94E-11 | 0.09981554 | 1.91E-14 |
| 7 | fdnGWTcI | 0.18367134 | 8.32E-44 | 0.20884656 | 3.71E-34 | 0.19591372 | 8.52E-45 | 0.24244341 | 3.42E-45 | 0.0931608 | 6.01E-39 | 0.07648998 | 2.00E-35 |
| 8 | fdnG5cI | 0.00400617 | 0.99112246 | 0.0024932 | 1 | 0.00675729 | 0.99983016 | 0.00302586 | 1 | 0 | 1 | 0 | 1 |
| 9 | fdnG10cI | N/A | 1 | N/A | 1 | N/A | 0.10289667 | N/A | 1 | N/A | 1 | N/A | 1 |
| 10 | fdnG15cI | N/A | 1 | N/A | 1 | 0.02942572 | 0.53093112 | 0 | 1 | N/A | 1 | N/A | 1 |
| 11 | fdnG17cI | N/A | 1 | N/A | 1 | N/A | 1 | N/A | 1 | N/A | 1 | N/A | 1 |
| 12 | fdnG23cI | N/A | 1 | N/A | 0.03456879 | N/A | 0.41809491 | N/A | 1 | N/A | 1 | N/A | 1 |
| 13 | hycAWTcI | 0.0161351 | 0.78555666 | 0.02082836 | 0.43677444 | 0.00543814 | 0.99999955 | 0.01477372 | 0.95567316 | 0.00488785 | 0.99957803 | 0.00307056 | 1 |
| 14 | hycA5cI | 0 | 1 | 0.00021673 | 1 | 0 | 1 | 0.00141753 | 1 | 0.00096539 | 0.99999997 | 0 | 1 |
| 15 | hycA10cI | N/A | 1 | N/A | 1 | N/A | 1 | N/A | 1 | N/A | 1 | N/A | 1 |
| 16 | hycA15cI | 0.00768477 | 0.96058276 | 0.00638715 | 0.99999992 | 0.00530881 | 0.99999155 | 0.00205195 | 1 | 0 | 1 | 0.00171251 | 1 |
| 17 | hycA17cI | 0.00433922 | 0.98755419 | 0.00028939 | 1 | 0.00080197 | 1 | 0.00046136 | 1 | 0 | 1 | 0 | 1 |
| 18 | hycA23cI | N/A | 1 | 0 | 1 | 0 | 1 | 0 | 1 | N/A | 1 | 0 | 1 |
| 19 | moaAWTcI | 0 | 1 | 0.00083827 | 1 | 0.00089602 | 1 | 0.00026941 | 1 | 0 | 1 | 0 | 1 |
| 20 | moaA5cI | 0.20346159 | 4.94E-15 | 0.23556373 | 2.41E-34 | 0.2703829 | 1.97E-45 | 0.24558174 | 5.42E-39 | 0.08163636 | 5.40E-11 | 0.10932639 | 5.29E-20 |
| 21 | moaA10cI | 0.28687104 | 1.97E-44 | 0.23009486 | 2.15E-35 | 0.26512196 | 7.04E-51 | 0.31701754 | 1.35E-48 | 0.16883469 | 6.83E-25 | 0.14077595 | 1.04E-39 |
| 22 | moaA15cI | 0.22100986 | 8.00E-52 | 0.24603721 | 3.09E-38 | 0.25092503 | 1.57E-54 | 0.34145806 | 2.57E-53 | 0.11987443 | 5.62E-50 | 0.11677819 | 3.09E-65 |
| 23 | moaA17cI | 0.29370376 | 3.42E-07 | 0.41192146 | 5.74E-31 | 0.28471841 | 6.28E-20 | 0.45448349 | 3.09E-31 | N/A | 0.00104349 | N/A | 0.05519805 |
| 24 | moaA23cI | 0.20410315 | 9.94E-47 | 0.29916578 | 9.97E-42 | 0.24243826 | 2.23E-51 | 0.28052034 | 1.04E-46 | 0.10786097 | 2.47E-40 | 0.10495747 | 9.20E-55 |
| 25 | napF5cI | 0.1209547 | 9.86E-21 | 0.19562358 | 5.91E-31 | 0.18101405 | 3.89E-37 | 0.20057941 | 1.30E-33 | 0.07663036 | 1.93E-15 | 0.05824518 | 4.09E-11 |
| 26 | napF10cI | N/A | 0.37596833 | N/A | 2.45E-12 | 0.29211438 | 9.26E-07 | 0.15409998 | 0.0003932 | N/A | 0.03288747 | N/A | 0.02218989 |
| 27 | napF15cI | 0.11147542 | 3.63E-30 | 0.20240932 | 1.97E-31 | 0.20827008 | 2.55E-41 | 0.16223419 | 1.56E-27 | 0.05867059 | 1.89E-23 | 0.05175835 | 9.35E-21 |
| 28 | napF17cI | N/A | 1 | N/A | 0.00093334 | N/A | 0.00111686 | 0.01666306 | 0.72053817 | N/A | 0.68452602 | N/A | 0.56831161 |
| 29 | napF23cI | 0.14007002 | 1.59E-43 | 0.21057628 | 1.66E-35 | 0.22311225 | 3.19E-53 | 0.25400705 | 1.21E-46 | 0.07602769 | 9.38E-52 | 0.07083673 | 3.70E-59 |
| 30 | narKWTcI | 0 | 1 | 0.00286184 | 1 | 0.0007891 | 1 | 0.00372949 | 1 | 0.00055958 | 1 | 0.00145892 | 1 |
| 31 | narK5cI | 0 | 1 | 0.00063706 | 1 | 0.00038469 | 1 | 0.00081163 | 1 | 0 | 1 | 0 | 1 |
| 32 | narK10cI | N/A | 1 | N/A | 1 | N/A | 1 | N/A | 1 | N/A | 1 | N/A | 1 |
| 33 | narK15cI | N/A | 1 | 0 | 1 | 0 | 1 | 0 | 1 | N/A | 1 | N/A | 1 |
| 34 | narK17cI | N/A | 1 | N/A | 1 | N/A | 1 | N/A | 1 | N/A | 1 | N/A | 1 |
| 35 | narK23cI | 0 | 1 | 0.00251089 | 1 | 0.00335849 | 0.99999984 | 0.00603915 | 0.99999925 | 0.00125034 | 0.99999869 | 0 | 1 |
| 36 | narG23cI | N/A | 1 | N/A | 0.090601 | N/A | 0.10289667 | N/A | 0.38169492 | N/A | 0.10239987 | N/A | 1 |
| 37 | torC5cI | 0.08635064 | 4.09E-27 | 0.08913771 | 3.44E-18 | 0.09466848 | 5.53E-27 | 0.03755514 | 1.09E-05 | 0.02529706 | 0.00259915 | 0.03078621 | 6.30E-09 |
| 38 | torC10cI | 0.14046083 | 2.21E-06 | 0.30533614 | 1.75E-13 | 0.16522339 | 9.41E-08 | 0.01664976 | 0.70539652 | 0.01709421 | 0.70465467 | 0.03746742 | 0.1064956 |
| 39 | torC15cI | 0.09184174 | 6.79E-30 | 0.09924109 | 2.81E-20 | 0.10499443 | 2.88E-30 | 0.04231021 | 1.78E-07 | 0.02654448 | 0.00012186 | 0.03270394 | 1.54E-12 |
| 40 | torC17cI | 0.06437248 | 0.17919234 | 0.10685464 | 0.00888125 | 0.02205631 | 0.62166984 | 0.01059054 | 0.8510179 | 0.02136164 | 0.5876287 | 0.01872624 | 0.64818422 |
| 41 | torC23cI | 0.12536024 | 1.21E-24 | 0.16609674 | 4.93E-22 | 0.17271797 | 3.16E-22 | 0.04697108 | 0.00086755 | 0.06341149 | 1.74E-16 | 0.04332026 | 1.41E-09 |
| 42 | yeaRWTcI | 0.0663459 | 2.18E-12 | 0.12708111 | 1.34E-24 | 0.17050512 | 2.36E-42 | 0.11989614 | 5.22E-27 | 0.03781875 | 1.13E-05 | 0.03046623 | 0.00022783 |
| 43 | yeaR5cI | N/A | 1 | N/A | 0.55641149 | N/A | 1 | N/A | 1 | N/A | 1 | N/A | 1 |
| 44 | yeaR10cI | 0.06552568 | 0.00288536 | 0.07311158 | 2.21E-11 | 0.11786375 | 2.38E-19 | 0.07577672 | 5.14E-11 | 0.01661934 | 0.7297615 | 0.02249651 | 0.42101036 |
| 45 | yeaR15cI | 0.07535416 | 0.00019783 | 0.08898322 | 3.89E-17 | 0.12918838 | 8.97E-28 | 0.07844865 | 1.59E-14 | 0.0150696 | 0.83824298 | 0.02402672 | 0.28476541 |
| 46 | yeaR17cI | N/A | 1 | N/A | 1 | N/A | 1 | N/A | 1 | N/A | 1 | N/A | 1 |
| 47 | yeaR23cI | N/A | 1 | N/A | 8.35E-11 | N/A | 6.33E-07 | 0.09266554 | 0.03418885 | N/A | 0.3196116 | N/A | 1 |
| 48 | yedY5cI | N/A | 1 | N/A | 1 | N/A | 1 | 0 | 1 | N/A | 1 | N/A | 1 |
| 49 | yedY10cI | 0.00300629 | 0.99813742 | 0.00087892 | 1 | 0.00065398 | 1 | 0.00125576 | 1 | 0 | 1 | 0 | 1 |
| 50 | yedY15cI | 0.0015425 | 0.99999235 | 0.00040961 | 1 | 0 | 1 | 0.00174116 | 1 | 0 | 1 | 0.00032493 | 1 |
| 51 | yedY17cI | 0 | 1 | 0.00043587 | 1 | 0.00048824 | 1 | 0.00212881 | 1 | 0.00066411 | 1 | 0 | 1 |
| 52 | yedY23cI | 0.00066319 | 1 | 0.00058338 | 1 | 0.00085485 | 1 | 0.001585 | 1 | 0 | 1 | 0 | 1 |
| 53 | ynfEWTcI | 0.00030897 | 1 | 0.00073103 | 1 | 0.0010784 | 1 | 0.00183429 | 1 | 0.00034596 | 1 | 0 | 1 |
| 54 | ynfE5cI | 0.00827551 | 0.94780357 | 0.00225022 | 1 | 0.00146757 | 1 | 0.00082489 | 1 | 0 | 1 | 0.00086752 | 0.99999999 |
| 55 | ynfE10cI | 0.17048852 | 7.57E-31 | 0.22742876 | 3.23E-36 | 0.18797529 | 1.93E-44 | 0.19572123 | 7.87E-38 | 0.07736925 | 1.04E-16 | 0.09256277 | 1.32E-25 |
| 56 | ynfE15cI | 0.12247188 | 1.14E-26 | 0.21207148 | 4.05E-34 | 0.23054227 | 1.63E-48 | 0.12503965 | 6.21E-26 | 0.05484043 | 4.16E-11 | 0.05901393 | 4.93E-16 |
| 57 | ynfE17cI | 0 | 1 | 0.07712243 | 8.71E-13 | 0.08718496 | 2.79E-14 | 0.04086061 | 0.00037464 | 0.00131129 | 0.99999604 | 0.00428855 | 0.99988772 |
| 58 | ynfE23cI | 0.05724924 | 2.74E-10 | 0.17869616 | 2.51E-31 | 0.19054714 | 5.51E-45 | 0.10064056 | 1.75E-22 | 0.02661323 | 0.02907161 | 0.02886099 | 0.00146846 |
| 59 | fabR | 1 | 1.15E-99 | 1 | 3.56E-61 | 1 | 1.21E-88 | 1 | 2.18E-72 | 1 | 1.95E-220 | 1 | 5.91E-277 |

| | Promoter | M3-11-18 | M3-11-18_pV | M4-11-18 | M4-11-18_pV | M5-11-18 | M5-11-18_pV | M6-11-18 | M6-11-18_pV | M7-11-18 | M7-11-18_pV | M8-11-18 | M8-11-18_pValue |
|---|---|---|---|---|---|---|---|---|---|---|---|---|---|
| 1 | cydAWTcI | 0.09206008 | 3.17E-11 | 0.06587152 | 6.36E-06 | 0.16538015 | 9.40E-40 | 0.11286178 | 9.61E-36 | 0.10789489 | 4.08E-31 | 0.14117111 | 3.80E-74 |
| 2 | cydA5cI | N/A | 0.28219258 | N/A | 1 | N/A | 1 | N/A | 1 | N/A | 1 | N/A | 0.0953587 |
| 3 | cydA10cI | 0.09892315 | 7.91E-08 | 0.14802316 | 3.14E-20 | 0.21773508 | 6.63E-55 | 0.16886411 | 1.54E-52 | 0.10079577 | 1.38E-32 | 0.22358548 | 1.10E-116 |
| 4 | cydA15cI | N/A | 0.73385121 | 0.02856913 | 0.4390469 | 0.10034764 | 0.00051137 | N/A | 1.86E-06 | 0.04770783 | 0.08286799 | 0.24298148 | 6.58E-14 |
| 5 | cydA17cI | 0.1003198 | 3.88E-13 | 0.16293078 | 2.56E-19 | 0.24858656 | 1.48E-65 | 0.14329282 | 5.28E-52 | 0.11064883 | 1.42E-39 | 0.17029551 | 6.78E-100 |
| 6 | cydA23cI | 0.12600832 | 4.33E-12 | 0.09927302 | 4.35E-14 | 0.17038847 | 4.42E-50 | 0.13709051 | 5.38E-51 | 0.09949555 | 8.76E-36 | 0.16946687 | 3.45E-102 |
| 7 | fdnGWTcI | 0.07200338 | 2.33E-26 | 0.10376339 | 1.78E-42 | 0.17732667 | 2.60E-66 | 0.1204432 | 6.44E-59 | 0.08905154 | 3.23E-43 | 0.13537888 | 3.09E-88 |
| 8 | fdnG5cI | 0.00139368 | 1 | 0.00572069 | 0.99535377 | 0.00268909 | 0.99999999 | 0.00345181 | 1 | 0.00061417 | 1 | 0.00368269 | 1 |
| 9 | fdnG10cI | N/A | 1 | N/A | 1 | N/A | 1 | N/A | 1 | N/A | 1 | N/A | 1 |
| 10 | fdnG15cI | N/A | 1 | N/A | 1 | 0 | 1 | 0 | 1 | 0 | 1 | 0.00672146 | 0.94452164 |
| 11 | fdnG17cI | N/A | 1 | N/A | 1 | N/A | 1 | N/A | 1 | N/A | 1 | N/A | 1 |
| 12 | fdnG23cI | N/A | 1 | N/A | 1 | N/A | 1 | N/A | 1 | N/A | 1 | 0.0111922 | 0.83647974 |
| 13 | hycAWTcI | 0.0072966 | 0.99979257 | 0.00356188 | 1 | 0.0150971 | 0.95500812 | 0.00541409 | 1 | 0.006169 | 1 | 0.02170116 | 0.23847648 |
| 14 | hycA5cI | 0.00152932 | 0.99999999 | 0.00151538 | 0.99998405 | 0.00038687 | 1 | 0 | 1 | 0 | 1 | 0.00015366 | 1 |
| 15 | hycA10cI | N/A | 1 | N/A | 1 | N/A | 1 | N/A | 1 | 0 | 1 | 0 | 1 |
| 16 | hycA15cI | 0.00537984 | 0.99989535 | 0.00162881 | 0.99996856 | 0.00231344 | 1 | 0.00187346 | 1 | 0.00024192 | 1 | 0.00404362 | 1 |
| 17 | hycA17cI | 0 | 1 | 0 | 1 | 0.00053837 | 1 | 0.00039439 | 1 | 0 | 1 | 0.00019976 | 1 |
| 18 | hycA23cI | N/A | 1 | N/A | 0.6512702 | 0 | 1 | 0 | 1 | 0 | 1 | 0 | 1 |
| 19 | moaAWTcI | 0.00055238 | 1 | 0 | 1 | 0.00058299 | 1 | 0 | 1 | 0.00030308 | 1 | 0.00025867 | 1 |
| 20 | moaA5cI | 0.08008893 | 2.66E-15 | 0.10036176 | 6.14E-12 | 0.14111397 | 1.34E-36 | 0.12137158 | 2.64E-37 | 0.09347814 | 9.24E-27 | 0.14983657 | 4.29E-75 |
| 21 | moaA10cI | 0.09449861 | 4.07E-18 | 0.147495 | 1.20E-44 | 0.18887153 | 3.47E-63 | 0.15163831 | 1.40E-59 | 0.08899179 | 2.51E-34 | 0.12955983 | 7.61E-84 |
| 22 | moaA15cI | 0.1069009 | 7.70E-46 | 0.11329396 | 2.12E-56 | 0.19274133 | 1.23E-78 | 0.13506654 | 2.32E-72 | 0.09310134 | 4.49E-43 | 0.14018736 | 4.14E-106 |
| 23 | moaA17cI | N/A | 0.00277752 | N/A | 9.15E-05 | 0.26054088 | 2.22E-13 | 0.41510823 | 4.83E-23 | 0.16424204 | 7.00E-12 | 0.27370648 | 4.90E-23 |
| 24 | moaA23cI | 0.08789228 | 8.16E-29 | 0.10130198 | 2.95E-42 | 0.19876197 | 1.77E-72 | 0.12959717 | 5.79E-61 | 0.07988624 | 7.48E-33 | 0.13582055 | 5.78E-91 |
| 25 | napF5cI | 0.07726666 | 1.47E-14 | 0.06343468 | 1.97E-09 | 0.14041303 | 5.35E-42 | 0.08948174 | 9.79E-32 | 0.07433061 | 2.89E-21 | 0.13793987 | 2.28E-73 |
| 26 | napF10cI | N/A | 0.14091244 | N/A | 0.15893053 | 0.08032812 | 0.04347337 | N/A | 2.26E-07 | 0.04664788 | 0.05305127 | 0.05331459 | 0.02095962 |
| 27 | napF15cI | 0.05019086 | 3.76E-15 | 0.04144884 | 2.79E-09 | 0.15119968 | 7.73E-46 | 0.09389469 | 7.50E-36 | 0.06403935 | 5.60E-16 | 0.11528453 | 1.55E-56 |
| 28 | napF17cI | N/A | 0.48453574 | N/A | 1 | 0.02414542 | 0.6026342 | N/A | 0.04682319 | 0.01460124 | 0.76366446 | 0.06714387 | 0.02754372 |
| 29 | napF23cI | 0.07081637 | 1.11E-46 | 0.07052238 | 5.72E-50 | 0.16025802 | 9.01E-77 | 0.11122164 | 6.16E-70 | 0.07656021 | 1.61E-37 | 0.12590821 | 2.42E-105 |
| 30 | narKWTcI | 0.00033509 | 1 | 0 | 1 | 0.00179553 | 1 | 0.00018391 | 1 | 0.00053208 | 1 | 0.00295157 | 1 |
| 31 | narK5cI | 0 | 1 | 0 | 1 | 0.00032742 | 1 | 0.00043041 | 1 | 0.00019832 | 1 | 0 | 1 |
| 32 | narK10cI | N/A | 1 | N/A | 1 | N/A | 1 | N/A | 1 | 0 | 1 | 0 | 1 |
| 33 | narK15cI | 0 | 1 | N/A | 1 | 0 | 1 | 0 | 1 | 0 | 1 | 0 | 1 |
| 34 | narK17cI | N/A | 1 | N/A | 1 | N/A | 1 | N/A | 1 | 0 | 1 | 0 | 1 |
| 35 | narK23cI | 0 | 1 | 0 | 1 | 0.00045358 | 1 | 0.00156716 | 1 | 0.00076781 | 1 | 0.00232233 | 1 |
| 36 | narG23cI | N/A | 1 | N/A | 1 | N/A | 1 | N/A | 1 | N/A | 0.18685509 | N/A | 0.00462139 |
| 37 | torC5cI | 0.03047279 | 6.08E-07 | 0.02676983 | 0.00026521 | 0.0958378 | 5.75E-49 | 0.05136898 | 9.27E-26 | 0.03200219 | 8.87E-07 | 0.10321597 | 5.94E-86 |
| 38 | torC10cI | 0.03507734 | 0.11445145 | 0.02539691 | 0.36838181 | 0.0803185 | 0.00061412 | 0.08367682 | 0.00024512 | 0.04534463 | 0.01581096 | 0.16459655 | 4.23E-14 |
| 39 | torC15cI | 0.0269711 | 6.28E-05 | 0.03179758 | 8.08E-10 | 0.11052575 | 4.32E-57 | 0.06451141 | 3.77E-38 | 0.03696033 | 1.25E-10 | 0.11036408 | 2.97E-94 |
| 40 | torC17cI | 0.00851424 | 0.90094807 | N/A | 0.18851768 | 0.03711857 | 0.21847295 | 0.06275612 | 0.03351365 | 0.01253456 | 0.80747959 | 0.16095161 | 0.00035853 |
| 41 | torC23cI | 0.05531646 | 4.90E-11 | 0.03806347 | 0.00012071 | 0.11271024 | 1.47E-19 | 0.07601049 | 7.20E-18 | 0.04022707 | 0.00023437 | 0.11898802 | 4.60E-29 |
| 42 | yeaRWTcI | 0.03001599 | 0.00245376 | 0.03417581 | 0.0002232 | 0.13488293 | 2.63E-61 | 0.09545717 | 1.91E-52 | 0.03683771 | 6.71E-09 | 0.10755524 | 6.36E-87 |
| 43 | yeaR5cI | N/A | 1 | N/A | 1 | N/A | 1 | N/A | 1 | N/A | 1 | N/A | 1 |
| 44 | yeaR10cI | 0.00451011 | 0.99933558 | 0.02381098 | 0.40730365 | 0.07539555 | 1.21E-12 | 0.08229113 | 1.18E-16 | 0.02023778 | 0.51178103 | 0.06932532 | 5.97E-25 |
| 45 | yeaR15cI | 0.01606507 | 0.78339129 | 0.06051514 | 0.00013792 | 0.04092813 | 2.33E-09 | 0.0662446 | 1.50E-19 | 0.0183032 | 0.73466223 | 0.06837425 | 1.68E-33 |
| 46 | yeaR17cI | N/A | 1 | N/A | 1 | N/A | 1 | N/A | 1 | N/A | 1 | N/A | 0.3044819 |
| 47 | yeaR23cI | N/A | 1 | N/A | 1 | 0.02195019 | 0.57197417 | N/A | 0.05881358 | 0.01460124 | 0.76366446 | 0.08947006 | 0.00356333 |
| 48 | yedY5cI | N/A | 1 | N/A | 1 | 0 | 1 | N/A | 1 | 0 | 1 | 0 | 1 |
| 49 | yedY10cI | 0 | 1 | 0 | 1 | 0 | 1 | 0 | 1 | 0.00019952 | 1 | 0.00014418 | 1 |
| 50 | yedY15cI | 0 | 1 | 0 | 1 | 0.00020575 | 1 | 0 | 1 | 0 | 1 | 0.00018028 | 1 |
| 51 | yedY17cI | 0 | 1 | 0 | 1 | 0 | 1 | 0 | 1 | 0 | 1 | 0 | 1 |
| 52 | yedY23cI | 0.00035951 | 1 | 0.00033448 | 1 | 0.00018103 | 1 | 0.00028583 | 1 | 0 | 1 | 0.0002197 | 1 |
| 53 | ynfEWTcI | 0.00012693 | 1 | 0.00028609 | 1 | 0.00031156 | 1 | 0.00021718 | 1 | 0.00012776 | 1 | 0.00047721 | 1 |
| 54 | ynfE5cI | 0.00247351 | 0.99999971 | 0 | 1 | 0.00207354 | 1 | 0.00029478 | 1 | 0.00026091 | 1 | 0.00030222 | 1 |
| 55 | ynfE10cI | 0.05911205 | 5.91E-15 | 0.07218421 | 3.09E-16 | 0.17886179 | 2.14E-70 | 0.1068148 | 5.86E-55 | 0.06271609 | 4.62E-24 | 0.11754052 | 2.58E-87 |
| 56 | ynfE15cI | 0.06090616 | 6.40E-17 | 0.08430634 | 1.08E-19 | 0.14111842 | 1.52E-51 | 0.107795 | 1.23E-48 | 0.06080064 | 2.13E-19 | 0.12691626 | 2.65E-85 |
| 57 | ynfE17cI | 0.00482202 | 0.99992063 | 0 | 1 | 0.03809192 | 0.00054139 | 0.03810451 | 0.00010004 | 0.01363315 | 0.9846565 | 0.04227931 | 1.42E-09 |
| 58 | ynfE23cI | 0.0348867 | 1.87E-05 | 0.03234164 | 0.00069183 | 0.11557645 | 1.58E-50 | 0.08810033 | 1.71E-44 | 0.04424911 | 4.92E-13 | 0.10480044 | 7.79E-79 |
| 59 | fabR | 1 | 1.10E-218 | 1 | 2.14E-235 | 1 | 3.83E-164 | 1 | 2.10E-201 | 1 | 1.47E-158 | 1 | 3.25E-278 |

### Data Set S3 - Nissle 1917 library 5 day experiment (-DSS and +DSS)
#### Aligned Reads

| | Promoter | GV-11-13-ALL | GV-11-13-ON | M1-11-14-ALL | M1-11-14-ON | M1-11-15-ALL | M1-11-15-ON | M1-11-17-ALL | M1-11-17-ON | M1-11-18-ALL | M1-11-18-ON |
|---|---|---|---|---|---|---|---|---|---|---|---|
| 1 | cydAWTcI | 1067 | 2110 | 407 | 564 | 95 | 112 | 31 | 54 | 15 | 28 |
| 2 | cydA5cI | 28 | 63 | 5 | 1 | 0 | 0 | 0 | 0 | 0 | 0 |
| 3 | cydA10cI | 1736 | 12571 | 466 | 737 | 88 | 125 | 34 | 53 | 10 | 35 |
| 4 | cydA15cI | 81 | 461 | 14 | 28 | 8 | 6 | 1 | 7 | 0 | 2 |
| 5 | cydA17cI | 2145 | 14207 | 584 | 811 | 87 | 135 | 38 | 57 | 22 | 48 |
| 6 | cydA23cI | 1999 | 13359 | 658 | 924 | 135 | 165 | 45 | 67 | 20 | 51 |
| 7 | fdnGWTcI | 1196 | 5020 | 438 | 598 | 419 | 458 | 326 | 449 | 183 | 400 |
| 8 | fdnG5cI | 873 | 4 | 488 | 1 | 91 | 1 | 56 | 0 | 31 | 0 |
| 9 | fdnG10cI | 16 | 1 | 1 | 0 | 0 | 0 | 0 | 0 | 1 | 0 |
| 10 | fdnG15cI | 18 | 0 | 9 | 0 | 2 | 0 | 2 | 0 | 0 | 0 |
| 11 | fdnG17cI | 6 | 0 | 1 | 0 | 0 | 0 | 0 | 0 | 0 | 0 |
| 12 | fdnG23cI | 15 | 0 | 6 | 0 | 1 | 0 | 1 | 0 | 0 | 0 |
| 13 | hycAWTcI | 1413 | 5 | 587 | 80 | 122 | 18 | 93 | 10 | 35 | 4 |
| 14 | hycA5cI | 1116 | 1 | 637 | 0 | 118 | 0 | 80 | 2 | 45 | 1 |
| 15 | hycA10cI | 22 | 0 | 3 | 0 | 0 | 0 | 0 | 0 | 0 | 0 |
| 16 | hycA15cI | 973 | 2 | 509 | 18 | 92 | 2 | 53 | 3 | 35 | 0 |
| 17 | hycA17cI | 834 | 1 | 416 | 0 | 87 | 0 | 61 | 0 | 37 | 0 |
| 18 | hycA23cI | 33 | 0 | 15 | 0 | 10 | 0 | 1 | 0 | 1 | 0 |
| 19 | moaAWTcI | 1175 | 3 | 635 | 7 | 166 | 0 | 85 | 1 | 49 | 0 |
| 20 | moaA5cI | 1049 | 6611 | 285 | 516 | 72 | 69 | 33 | 65 | 35 | 67 |
| 21 | moaA10cI | 1812 | 11639 | 435 | 739 | 146 | 146 | 73 | 122 | 26 | 103 |
| 22 | moaA15cI | 5559 | 35446 | 1836 | 2801 | 656 | 733 | 300 | 543 | 155 | 436 |
| 23 | moaA17cI | 340 | 4322 | 42 | 150 | 18 | 31 | 7 | 16 | 2 | 9 |
| 24 | moaA23cI | 2513 | 16989 | 712 | 1233 | 372 | 466 | 234 | 420 | 130 | 329 |
| 25 | napFWTcI | 0 | 0 | 0 | 1 | 0 | 1 | 0 | 1 | 1 | 1 |
| 26 | napF5cI | 1110 | 6074 | 544 | 672 | 331 | 303 | 139 | 106 | 64 | 115 |
| 27 | napF10cI | 36 | 95 | 16 | 24 | 8 | 7 | 3 | 2 | 0 | 3 |
| 28 | napF15cI | 817 | 4074 | 957 | 1129 | 974 | 963 | 556 | 444 | 311 | 428 |
| 29 | napF17cI | 20 | 76 | 12 | 5 | 10 | 2 | 0 | 2 | 2 | 1 |
| 30 | napF23cI | 2549 | 14018 | 9563 | 12221 | 15569 | 14632 | 11798 | 13428 | 6210 | 11089 |
| 31 | narKWTcI | 1679 | 7 | 1149 | 9 | 257 | 0 | 121 | 2 | 77 | 1 |
| 32 | narK5cI | 1307 | 4 | 912 | 2 | 166 | 0 | 80 | 0 | 53 | 0 |
| 33 | narK10cI | 31 | 1 | 3 | 0 | 0 | 0 | 0 | 0 | 0 | 0 |
| 34 | narK15cI | 56 | 1 | 27 | 0 | 11 | 0 | 1 | 0 | 4 | 0 |
| 35 | narK17cI | 19 | 0 | 4 | 0 | 0 | 0 | 0 | 0 | 0 | 0 |
| 36 | narK23cI | 881 | 1 | 514 | 4 | 116 | 0 | 45 | 1 | 35 | 1 |
| 37 | narGWTcI | 0 | 4 | 0 | 0 | 0 | 0 | 0 | 0 | 0 | 0 |
| 38 | narG5cI | 1 | 0 | 0 | 0 | 0 | 0 | 0 | 0 | 0 | 0 |
| 39 | narG10cI | 1 | 0 | 0 | 0 | 0 | 0 | 0 | 0 | 0 | 0 |
| 40 | narG15cI | 0 | 0 | 0 | 0 | 0 | 0 | 0 | 0 | 0 | 0 |
| 41 | narG17cI | 0 | 0 | 0 | 0 | 0 | 0 | 0 | 0 | 0 | 0 |
| 42 | narG23cI | 10 | 17 | 1 | 0 | 1 | 1 | 0 | 0 | 0 | 2 |
| 43 | torCWTcI | 0 | 0 | 0 | 0 | 0 | 0 | 0 | 0 | 0 | 0 |
| 44 | torC5cI | 1841 | 351 | 5018 | 4238 | 5955 | 4522 | 3409 | 1871 | 1993 | 1183 |
| 45 | torC10cI | 19 | 7 | 24 | 18 | 26 | 29 | 20 | 7 | 15 | 6 |
| 46 | torC15cI | 1856 | 505 | 11831 | 9706 | 23649 | 19217 | 21505 | 11498 | 14663 | 9139 |
| 47 | torC17cI | 9 | 9 | 14 | 5 | 7 | 7 | 6 | 5 | 6 | 3 |
| 48 | torC23cI | 42 | 16 | 135 | 139 | 307 | 236 | 170 | 177 | 117 | 174 |
| 49 | yeaRWTcI | 2797 | 55 | 1480 | 432 | 352 | 100 | 213 | 137 | 132 | 117 |
| 50 | yeaR5cI | 7 | 0 | 0 | 0 | 0 | 0 | 0 | 0 | 0 | 0 |
| 51 | yeaR10cI | 439 | 1 | 314 | 84 | 55 | 22 | 29 | 14 | 18 | 7 |
| 52 | yeaR15cI | 701 | 0 | 372 | 58 | 89 | 21 | 43 | 17 | 34 | 12 |
| 53 | yeaR17cI | 7 | 0 | 1 | 0 | 0 | 0 | 0 | 0 | 0 | 0 |
| 54 | yeaR23cI | 17 | 0 | 5 | 0 | 5 | 0 | 0 | 1 | 0 | 1 |
| 55 | yedYWTcI | 1 | 1 | 0 | 0 | 0 | 0 | 0 | 0 | 0 | 0 |
| 56 | yedY5cI | 33 | 0 | 7 | 0 | 0 | 0 | 0 | 0 | 1 | 0 |
| 57 | yedY10cI | 1197 | 2 | 622 | 3 | 133 | 0 | 79 | 0 | 38 | 0 |
| 58 | yedY15cI | 1844 | 4 | 972 | 3 | 207 | 2 | 105 | 0 | 61 | 0 |
| 59 | yedY17cI | 1148 | 6 | 792 | 1 | 200 | 0 | 96 | 0 | 65 | 1 |
| 60 | yedY23cI | 3537 | 15 | 2277 | 2 | 555 | 1 | 226 | 1 | 188 | 0 |
| 61 | ynfEWTcI | 8630 | 19 | 4884 | 11 | 984 | 0 | 468 | 3 | 375 | 3 |
| 62 | ynfE5cI | 1246 | 1 | 519 | 3 | 113 | 1 | 56 | 0 | 41 | 0 |
| 63 | ynfE10cI | 4794 | 15 | 954 | 1316 | 232 | 193 | 135 | 152 | 70 | 127 |
| 64 | ynfE15cI | 3250 | 24 | 748 | 797 | 290 | 223 | 134 | 177 | 98 | 126 |
| 65 | ynfE17cI | 1093 | 4 | 486 | 26 | 81 | 1 | 56 | 0 | 32 | 1 |
| 66 | ynfE23cI | 4991 | 13 | 1619 | 827 | 434 | 173 | 249 | 151 | 165 | 103 |
| 67 | fabR | 293 | 22194 | 362 | 3504 | 737 | 3700 | 111 | 1188 | 199 | 4673 |

| | Promoter | M2-11-14-ALL | M2-11-14-ON | M2-11-15-ALL | M2-11-15-ON | M2-11-17-ALL | M2-11-17-ON | M2-11-18-ALL | M2-11-18-ON | M3-11-14-ALL | M3-11-14-ON | M3-11-15-ALL | M3-11-15-ON | M3-11-17-ALL | M3-11-17-ON | M3-11-18-ALL | M3-11-18-ON |
|---|---|---|---|---|---|---|---|---|---|---|---|---|---|---|---|---|---|
| 1 | cydAWTcI | 128 | 189 | 44 | 40 | 23 | 50 | 19 | 50 | 289 | 708 | 57 | 69 | 20 | 72 | 31 | 56 |
| 2 | cydA5cI | 0 | 0 | 0 | 0 | 0 | 0 | 0 | 0 | 2 | 6 | 0 | 0 | 0 | 0 | 0 | 1 |
| 3 | cydA10cI | 155 | 270 | 41 | 45 | 20 | 48 | 20 | 65 | 299 | 836 | 52 | 50 | 16 | 66 | 17 | 33 |
| 4 | cydA15cI | 6 | 15 | 4 | 4 | 2 | 2 | 1 | 5 | 17 | 35 | 1 | 4 | 0 | 4 | 3 | 1 |
| 5 | cydA17cI | 280 | 465 | 48 | 61 | 34 | 59 | 17 | 71 | 324 | 906 | 82 | 70 | 37 | 80 | 32 | 63 |
| 6 | cydA23cI | 222 | 373 | 60 | 49 | 30 | 69 | 30 | 80 | 369 | 906 | 60 | 81 | 24 | 54 | 19 | 47 |
| 7 | fdnGWTcI | 200 | 335 | 246 | 237 | 193 | 315 | 207 | 423 | 361 | 876 | 273 | 286 | 155 | 355 | 201 | 284 |
| 8 | fdnG5cI | 199 | 6 | 42 | 1 | 62 | 2 | 40 | 0 | 327 | 1 | 56 | 0 | 50 | 1 | 75 | 2 |
| 9 | fdnG10cI | 0 | 0 | 0 | 0 | 0 | 0 | 0 | 0 | 2 | 0 | 1 | 0 | 0 | 0 | 0 | 0 |
| 10 | fdnG15cI | 2 | 0 | 5 | 0 | 0 | 0 | 1 | 0 | 6 | 0 | 0 | 0 | 2 | 0 | 1 | 0 |
| 11 | fdnG17cI | 0 | 0 | 0 | 0 | 0 | 0 | 0 | 0 | 1 | 0 | 0 | 0 | 0 | 0 | 0 | 0 |
| 12 | fdnG23cI | 0 | 0 | 1 | 0 | 0 | 0 | 1 | 0 | 7 | 0 | 3 | 0 | 0 | 0 | 0 | 0 |
| 13 | hycAWTcI | 213 | 9 | 91 | 7 | 64 | 11 | 110 | 9 | 683 | 42 | 109 | 6 | 68 | 14 | 77 | 11 |
| 14 | hycA5cI | 178 | 0 | 51 | 0 | 64 | 0 | 56 | 0 | 426 | 2 | 91 | 0 | 52 | 0 | 66 | 2 |
| 15 | hycA10cI | 0 | 0 | 1 | 0 | 0 | 0 | 1 | 0 | 4 | 1 | 0 | 0 | 0 | 0 | 0 | 0 |
| 16 | hycA15cI | 159 | 2 | 49 | 0 | 53 | 1 | 66 | 3 | 377 | 12 | 65 | 1 | 45 | 1 | 57 | 6 |
| 17 | hycA17cI | 108 | 1 | 44 | 0 | 53 | 0 | 44 | 0 | 380 | 6 | 55 | 0 | 40 | 1 | 53 | 0 |
| 18 | hycA23cI | 2 | 0 | 4 | 0 | 1 | 0 | 8 | 0 | 20 | 0 | 1 | 0 | 0 | 0 | 2 | 0 |
| 19 | moaAWTcI | 171 | 0 | 111 | 0 | 93 | 1 | 109 | 0 | 485 | 3 | 81 | 0 | 82 | 1 | 93 | 1 |
| 20 | moaA5cI | 122 | 206 | 33 | 49 | 30 | 54 | 38 | 111 | 464 | 1611 | 78 | 121 | 39 | 123 | 63 | 99 |
| 21 | moaA10cI | 172 | 311 | 83 | 115 | 50 | 159 | 59 | 222 | 299 | 1034 | 70 | 109 | 50 | 104 | 55 | 102 |
| 22 | moaA15cI | 782 | 1432 | 470 | 613 | 280 | 567 | 223 | 696 | 927 | 2925 | 289 | 423 | 170 | 480 | 192 | 403 |
| 23 | moaA17cI | 17 | 55 | 2 | 11 | 3 | 9 | 2 | 5 | 20 | 152 | 4 | 8 | 1 | 18 | 1 | 6 |
| 24 | moaA23cI | 436 | 877 | 386 | 458 | 189 | 402 | 200 | 561 | 388 | 1438 | 155 | 255 | 88 | 299 | 131 | 226 |
| 25 | napFWTcI | 0 | 1 | 2 | 0 | 0 | 0 | 1 | 0 | 0 | 1 | 1 | 0 | 0 | 1 | 0 | 0 |
| 26 | napF5cI | 353 | 553 | 226 | 267 | 84 | 79 | 72 | 112 | 259 | 605 | 139 | 181 | 45 | 92 | 64 | 97 |
| 27 | napF10cI | 5 | 6 | 2 | 7 | 1 | 2 | 1 | 5 | 10 | 16 | 2 | 3 | 1 | 4 | 2 | 3 |
| 28 | napF15cI | 644 | 1058 | 725 | 996 | 299 | 290 | 298 | 412 | 305 | 716 | 305 | 481 | 249 | 363 | 260 | 256 |
| 29 | napF17cI | 3 | 5 | 4 | 1 | 1 | 0 | 3 | 2 | 6 | 8 | 3 | 0 | 0 | 0 | 1 | 1 |
| 30 | napF23cI | 11378 | 16642 | 15048 | 16142 | 8507 | 12317 | 7397 | 14014 | 5714 | 13259 | 7269 | 8184 | 4875 | 10560 | 5359 | 7456 |
| 31 | narKWTcI | 654 | 2 | 126 | 3 | 127 | 3 | 181 | 7 | 729 | 16 | 121 | 3 | 80 | 2 | 153 | 1 |
| 32 | narK5cI | 248 | 0 | 72 | 0 | 71 | 1 | 64 | 0 | 656 | 2 | 110 | 1 | 76 | 0 | 77 | 0 |
| 33 | narK10cI | 1 | 0 | 0 | 0 | 0 | 0 | 0 | 0 | 9 | 1 | 1 | 0 | 0 | 0 | 0 | 0 |
| 34 | narK15cI | 3 | 0 | 6 | 0 | 3 | 0 | 3 | 0 | 17 | 0 | 3 | 0 | 2 | 0 | 5 | 0 |
| 35 | narK17cI | 0 | 0 | 0 | 0 | 0 | 0 | 0 | 0 | 4 | 0 | 0 | 0 | 0 | 0 | 2 | 0 |
| 36 | narK23cI | 163 | 0 | 65 | 0 | 70 | 0 | 66 | 0 | 338 | 6 | 53 | 1 | 47 | 0 | 62 | 0 |
| 37 | narGWTcI | 0 | 0 | 0 | 0 | 0 | 0 | 0 | 0 | 0 | 0 | 0 | 0 | 0 | 0 | 0 | 0 |
| 38 | narG5cI | 0 | 0 | 0 | 0 | 0 | 0 | 0 | 0 | 0 | 0 | 0 | 0 | 0 | 0 | 0 | 0 |
| 39 | narG10cI | 0 | 0 | 0 | 0 | 0 | 0 | 0 | 0 | 0 | 0 | 0 | 0 | 0 | 0 | 0 | 0 |
| 40 | narG15cI | 0 | 0 | 0 | 0 | 0 | 0 | 0 | 0 | 0 | 0 | 0 | 0 | 0 | 0 | 0 | 0 |
| 41 | narG17cI | 0 | 0 | 0 | 0 | 0 | 0 | 0 | 0 | 0 | 0 | 0 | 0 | 0 | 0 | 0 | 0 |
| 42 | narG23cI | 0 | 0 | 0 | 1 | 0 | 0 | 0 | 0 | 2 | 1 | 0 | 0 | 0 | 0 | 0 | 0 |
| 43 | torCWTcI | 0 | 0 | 0 | 0 | 0 | 0 | 0 | 0 | 0 | 0 | 0 | 0 | 0 | 0 | 0 | 0 |
| 44 | torC5cI | 3703 | 3783 | 4199 | 3313 | 2499 | 1492 | 2082 | 1713 | 3276 | 5652 | 3107 | 2904 | 1608 | 1600 | 1725 | 1032 |
| 45 | torC10cI | 19 | 20 | 33 | 25 | 14 | 6 | 11 | 11 | 13 | 44 | 14 | 15 | 8 | 10 | 16 | 11 |
| 46 | torC15cI | 12222 | 13724 | 21011 | 16088 | 14122 | 8544 | 14352 | 12550 | 5969 | 11094 | 10257 | 10215 | 10472 | 10642 | 13533 | 7172 |
| 47 | torC17cI | 9 | 5 | 11 | 3 | 4 | 3 | 8 | 4 | 9 | 5 | 3 | 5 | 4 | 2 | 6 | 1 |
| 48 | torC23cI | 67 | 239 | 288 | 249 | 95 | 134 | 153 | 177 | 57 | 150 | 94 | 140 | 82 | 146 | 105 | 114 |
| 49 | yeaRWTcI | 522 | 81 | 209 | 37 | 222 | 129 | 235 | 191 | 1288 | 1151 | 234 | 166 | 100 | 133 | 192 | 113 |
| 50 | yeaR5cI | 0 | 0 | 0 | 0 | 0 | 0 | 0 | 0 | 4 | 0 | 0 | 0 | 0 | 0 | 0 | 0 |
| 51 | yeaR10cI | 90 | 2 | 35 | 4 | 32 | 15 | 25 | 15 | 180 | 56 | 33 | 15 | 28 | 21 | 34 | 3 |
| 52 | yeaR15cI | 96 | 6 | 46 | 4 | 43 | 19 | 39 | 25 | 320 | 122 | 64 | 13 | 27 | 22 | 35 | 11 |
| 53 | yeaR17cI | 0 | 0 | 0 | 0 | 0 | 0 | 1 | 0 | 0 | 0 | 0 | 0 | 0 | 0 | 0 | 0 |
| 54 | yeaR23cI | 0 | 0 | 1 | 0 | 0 | 0 | 0 | 0 | 4 | 2 | 0 | 0 | 0 | 2 | 0 | 0 |
| 55 | yedYWTcI | 0 | 1 | 0 | 0 | 0 | 0 | 0 | 0 | 0 | 0 | 0 | 0 | 0 | 0 | 0 | 0 |
| 56 | yedY5cI | 0 | 0 | 1 | 0 | 0 | 0 | 1 | 0 | 9 | 0 | 1 | 0 | 0 | 0 | 0 | 0 |
| 57 | yedY10cI | 182 | 1 | 70 | 1 | 75 | 0 | 80 | 0 | 437 | 0 | 79 | 0 | 65 | 0 | 74 | 0 |
| 58 | yedY15cI | 333 | 1 | 112 | 0 | 129 | 1 | 116 | 1 | 668 | 7 | 107 | 0 | 84 | 2 | 102 | 0 |
| 59 | yedY17cI | 234 | 0 | 66 | 0 | 78 | 1 | 78 | 0 | 416 | 5 | 100 | 1 | 56 | 1 | 97 | 0 |
| 60 | yedY23cI | 684 | 2 | 261 | 0 | 258 | 2 | 294 | 0 | 1533 | 14 | 262 | 0 | 213 | 1 | 286 | 2 |
| 61 | ynfEWTcI | 1380 | 7 | 492 | 1 | 463 | 1 | 545 | 0 | 2962 | 30 | 529 | 1 | 378 | 1 | 462 | 1 |
| 62 | ynfE5cI | 136 | 1 | 64 | 0 | 65 | 0 | 44 | 1 | 396 | 8 | 76 | 0 | 46 | 0 | 62 | 3 |
| 63 | ynfE10cI | 463 | 363 | 125 | 112 | 103 | 193 | 74 | 183 | 1635 | 2779 | 234 | 255 | 127 | 207 | 138 | 160 |
| 64 | ynfE15cI | 325 | 169 | 143 | 100 | 143 | 157 | 118 | 186 | 1255 | 1609 | 209 | 204 | 111 | 211 | 154 | 184 |
| 65 | ynfE17cI | 155 | 1 | 43 | 0 | 57 | 0 | 35 | 4 | 306 | 46 | 51 | 2 | 36 | 2 | 53 | 5 |
| 66 | ynfE23cI | 752 | 85 | 220 | 54 | 248 | 132 | 226 | 174 | 877 | 1042 | 222 | 149 | 159 | 165 | 209 | 143 |
| 67 | fabR | 237 | 2187 | 497 | 2783 | 89 | 698 | 250 | 6685 | 390 | 6461 | 310 | 2145 | 78 | 785 | 198 | 3891 |

| | Promoter | M4-11-14-ALL | M4-11-14-ON | M4-11-15-ALL | M4-11-15-ON | M4-11-17-ALL | M4-11-17-ON | M4-11-18-ALL | M4-11-18-ON | M5-11-14-ALL | M5-11-14-ON | M5-11-15-ALL | M5-11-15-ON | M5-11-17-ALL | M5-11-17-ON | M5-11-18-ALL | M5-11-18-ON |
|---|---|---|---|---|---|---|---|---|---|---|---|---|---|---|---|---|---|
| 1 | cydAWTcI | 373 | 246 | 32 | 61 | 41 | 48 | 26 | 36 | 632 | 1434 | 561 | 761 | 149 | 441 | 104 | 143 |
| 2 | cydA5cI | 8 | 0 | 0 | 0 | 0 | 0 | 0 | 0 | 5 | 9 | 10 | 2 | 2 | 1 | 1 | 0 |
| 3 | cydA10cI | 538 | 519 | 61 | 105 | 28 | 56 | 26 | 81 | 803 | 2526 | 772 | 1276 | 214 | 654 | 106 | 192 |
| 4 | cydA15cI | 35 | 18 | 2 | 11 | 3 | 1 | 5 | 3 | 9 | 103 | 26 | 49 | 8 | 28 | 12 | 10 |
| 5 | cydA17cI | 651 | 642 | 68 | 143 | 33 | 71 | 21 | 72 | 1022 | 2763 | 781 | 1443 | 262 | 778 | 117 | 242 |
| 6 | cydA23cI | 635 | 572 | 91 | 152 | 48 | 62 | 34 | 71 | 1019 | 2562 | 795 | 1266 | 259 | 722 | 151 | 214 |
| 7 | fdnGWTcI | 592 | 425 | 205 | 306 | 264 | 303 | 158 | 345 | 702 | 1781 | 822 | 959 | 538 | 1270 | 307 | 453 |
| 8 | fdnG5cI | 590 | 11 | 63 | 7 | 40 | 1 | 25 | 3 | 717 | 4 | 638 | 5 | 324 | 9 | 179 | 4 |
| 9 | fdnG10cI | 1 | 0 | 0 | 0 | 0 | 0 | 0 | 0 | 1 | 0 | 5 | 0 | 0 | 0 | 1 | 0 |
| 10 | fdnG15cI | 8 | 0 | 0 | 0 | 2 | 0 | 0 | 0 | 6 | 0 | 5 | 1 | 3 | 0 | 6 | 0 |
| 11 | fdnG17cI | 2 | 0 | 0 | 0 | 0 | 0 | 0 | 0 | 3 | 0 | 3 | 0 | 0 | 0 | 4 | 0 |
| 12 | fdnG23cI | 12 | 0 | 0 | 0 | 0 | 0 | 2 | 0 | 5 | 0 | 19 | 0 | 0 | 2 | 2 | 0 |
| 13 | hycAWTcI | 863 | 16 | 99 | 19 | 110 | 11 | 174 | 13 | 1032 | 137 | 1285 | 71 | 512 | 120 | 375 | 47 |
| 14 | hycA5cI | 764 | 1 | 73 | 0 | 63 | 0 | 31 | 1 | 872 | 7 | 963 | 1 | 418 | 1 | 297 | 1 |
| 15 | hycA10cI | 7 | 0 | 0 | 1 | 0 | 0 | 0 | 0 | 2 | 0 | 20 | 0 | 2 | 0 | 2 | 0 |
| 16 | hycA15cI | 677 | 4 | 72 | 9 | 42 | 2 | 29 | 1 | 711 | 15 | 766 | 16 | 378 | 27 | 260 | 5 |
| 17 | hycA17cI | 533 | 1 | 59 | 0 | 37 | 1 | 27 | 0 | 637 | 2 | 664 | 1 | 313 | 1 | 223 | 1 |
| 18 | hycA23cI | 27 | 0 | 3 | 0 | 3 | 0 | 2 | 1 | 13 | 0 | 51 | 1 | 11 | 0 | 10 | 0 |
| 19 | moaAWTcI | 809 | 2 | 116 | 0 | 86 | 0 | 48 | 0 | 886 | 3 | 1219 | 1 | 536 | 5 | 416 | 2 |
| 20 | moaA5cI | 309 | 239 | 60 | 105 | 26 | 33 | 27 | 57 | 464 | 1402 | 543 | 891 | 183 | 487 | 127 | 149 |
| 21 | moaA10cI | 517 | 505 | 317 | 622 | 77 | 138 | 77 | 239 | 805 | 2360 | 1183 | 1724 | 323 | 840 | 210 | 330 |
| 22 | moaA15cI | 1954 | 1676 | 460 | 786 | 304 | 420 | 237 | 565 | 2791 | 6959 | 2784 | 4589 | 875 | 2434 | 546 | 876 |
| 23 | moaA17cI | 57 | 92 | 6 | 33 | 6 | 11 | 3 | 12 | 77 | 451 | 93 | 240 | 23 | 107 | 12 | 26 |
| 24 | moaA23cI | 961 | 851 | 238 | 468 | 236 | 301 | 167 | 356 | 1199 | 3373 | 1877 | 3009 | 432 | 1461 | 295 | 488 |
| 25 | napFWTcI | 0 | 1 | 1 | 0 | 3 | 1 | 2 | 0 | 1 | 3 | 1 | 0 | 1 | 1 | 0 | 0 |
| 26 | napF5cI | 591 | 594 | 193 | 175 | 114 | 86 | 54 | 72 | 573 | 1316 | 627 | 776 | 224 | 495 | 167 | 195 |
| 27 | napF10cI | 23 | 18 | 10 | 15 | 3 | 1 | 2 | 3 | 7 | 35 | 20 | 29 | 4 | 23 | 6 | 4 |
| 28 | napF15cI | 844 | 819 | 498 | 454 | 448 | 312 | 218 | 190 | 449 | 1016 | 499 | 572 | 206 | 471 | 167 | 210 |
| 29 | napF17cI | 10 | 6 | 2 | 1 | 1 | 0 | 2 | 0 | 9 | 19 | 11 | 10 | 2 | 6 | 5 | 1 |
| 30 | napF23cI | 13139 | 12004 | 9856 | 10318 | 11130 | 9755 | 6033 | 8957 | 3311 | 7280 | 4527 | 5870 | 3670 | 8740 | 3127 | 4173 |
| 31 | narKWTcI | 1199 | 1 | 180 | 1 | 153 | 0 | 86 | 0 | 1453 | 8 | 1611 | 4 | 687 | 22 | 470 | 7 |
| 32 | narK5cI | 1049 | 0 | 100 | 0 | 58 | 0 | 43 | 0 | 1196 | 2 | 1149 | 2 | 552 | 4 | 342 | 1 |
| 33 | narK10cI | 9 | 0 | 0 | 0 | 0 | 0 | 0 | 0 | 3 | 0 | 25 | 0 | 1 | 0 | 3 | 0 |
| 34 | narK15cI | 28 | 0 | 3 | 0 | 3 | 0 | 0 | 0 | 9 | 0 | 29 | 3 | 10 | 0 | 15 | 0 |
| 35 | narK17cI | 5 | 0 | 0 | 0 | 0 | 0 | 0 | 0 | 3 | 1 | 11 | 0 | 2 | 0 | 4 | 0 |
| 36 | narK23cI | 688 | 1 | 47 | 0 | 40 | 0 | 35 | 0 | 729 | 44 | 752 | 21 | 393 | 11 | 259 | 1 |
| 37 | narGWTcI | 0 | 0 | 0 | 0 | 0 | 0 | 0 | 0 | 0 | 0 | 0 | 0 | 0 | 0 | 0 | 0 |
| 38 | narG5cI | 0 | 0 | 0 | 0 | 0 | 0 | 0 | 0 | 0 | 0 | 0 | 0 | 0 | 0 | 0 | 0 |
| 39 | narG10cI | 0 | 0 | 0 | 0 | 0 | 0 | 0 | 0 | 0 | 0 | 0 | 0 | 0 | 0 | 0 | 0 |
| 40 | narG15cI | 0 | 0 | 0 | 0 | 0 | 0 | 0 | 0 | 0 | 0 | 0 | 0 | 0 | 0 | 0 | 0 |
| 41 | narG17cI | 0 | 0 | 0 | 0 | 0 | 0 | 0 | 0 | 0 | 0 | 0 | 0 | 0 | 0 | 0 | 0 |
| 42 | narG23cI | 6 | 1 | 0 | 0 | 0 | 0 | 0 | 0 | 2 | 0 | 1 | 2 | 1 | 2 | 0 | 0 |
| 43 | torCWTcI | 0 | 0 | 0 | 0 | 0 | 0 | 0 | 0 | 0 | 0 | 0 | 0 | 0 | 0 | 0 | 0 |
| 44 | torC5cI | 5843 | 3757 | 3485 | 2926 | 2607 | 1408 | 1682 | 947 | 2471 | 4209 | 4851 | 3297 | 3204 | 3229 | 3393 | 2707 |
| 45 | torC10cI | 28 | 27 | 16 | 16 | 16 | 14 | 15 | 8 | 11 | 24 | 43 | 23 | 9 | 31 | 18 | 12 |
| 46 | torC15cI | 13251 | 9269 | 13329 | 11411 | 14151 | 8131 | 10076 | 6747 | 3259 | 5438 | 5418 | 3881 | 5350 | 6003 | 5274 | 4854 |
| 47 | torC17cI | 19 | 4 | 7 | 6 | 5 | 2 | 4 | 4 | 3 | 15 | 21 | 11 | 5 | 6 | 13 | 4 |
| 48 | torC23cI | 183 | 118 | 182 | 170 | 147 | 115 | 95 | 76 | 14 | 76 | 108 | 79 | 71 | 133 | 79 | 74 |
| 49 | yeaRWTcI | 1918 | 424 | 250 | 209 | 249 | 103 | 142 | 102 | 2609 | 631 | 3122 | 2326 | 1237 | 1777 | 799 | 897 |
| 50 | yeaR5cI | 3 | 0 | 0 | 0 | 0 | 0 | 0 | 0 | 1 | 0 | 7 | 2 | 3 | 1 | 3 | 0 |
| 51 | yeaR10cI | 327 | 34 | 43 | 22 | 27 | 11 | 18 | 9 | 331 | 62 | 434 | 196 | 176 | 145 | 107 | 67 |
| 52 | yeaR15cI | 505 | 64 | 31 | 43 | 32 | 15 | 22 | 28 | 575 | 84 | 2333 | 2006 | 588 | 591 | 638 | 217 |
| 53 | yeaR17cI | 2 | 0 | 0 | 0 | 0 | 0 | 0 | 0 | 0 | 0 | 13 | 4 | 3 | 0 | 3 | 0 |
| 54 | yeaR23cI | 5 | 2 | 1 | 2 | 1 | 0 | 0 | 0 | 10 | 1 | 36 | 20 | 3 | 19 | 11 | 2 |
| 55 | yedYWTcI | 0 | 0 | 0 | 0 | 0 | 0 | 0 | 0 | 0 | 0 | 0 | 0 | 0 | 0 | 0 | 0 |
| 56 | yedY5cI | 9 | 0 | 1 | 0 | 0 | 0 | 0 | 0 | 5 | 0 | 31 | 0 | 3 | 0 | 12 | 0 |
| 57 | yedY10cI | 814 | 3 | 82 | 3 | 54 | 1 | 44 | 0 | 845 | 5 | 1048 | 1 | 409 | 4 | 335 | 0 |
| 58 | yedY15cI | 1130 | 1 | 146 | 3 | 105 | 1 | 63 | 0 | 1315 | 6 | 1452 | 4 | 690 | 3 | 647 | 1 |
| 59 | yedY17cI | 709 | 1 | 91 | 1 | 66 | 0 | 38 | 0 | 901 | 3 | 1030 | 2 | 422 | 2 | 398 | 0 |
| 60 | yedY23cI | 2331 | 7 | 295 | 0 | 244 | 1 | 143 | 1 | 2707 | 8 | 3239 | 3 | 1468 | 10 | 1219 | 2 |
| 61 | ynfEWTcI | 5044 | 3 | 588 | 1 | 500 | 1 | 335 | 2 | 5858 | 78 | 6584 | 36 | 2689 | 22 | 1979 | 5 |
| 62 | ynfE5cI | 675 | 3 | 97 | 1 | 39 | 2 | 23 | 0 | 751 | 13 | 862 | 7 | 359 | 9 | 230 | 4 |
| 63 | ynfE10cI | 1111 | 756 | 202 | 238 | 109 | 116 | 83 | 126 | 1929 | 4695 | 2248 | 2950 | 641 | 1648 | 393 | 585 |
| 64 | ynfE15cI | 903 | 523 | 179 | 214 | 191 | 146 | 75 | 133 | 1270 | 3043 | 1484 | 1779 | 421 | 1009 | 276 | 324 |
| 65 | ynfE17cI | 512 | 4 | 65 | 3 | 34 | 0 | 24 | 0 | 537 | 551 | 631 | 368 | 215 | 187 | 155 | 49 |
| 66 | ynfE23cI | 2090 | 329 | 265 | 135 | 297 | 106 | 153 | 104 | 2080 | 4022 | 2539 | 2569 | 848 | 1713 | 601 | 578 |
| 67 | fabR | 492 | 1546 | 255 | 2353 | 93 | 582 | 213 | 4486 | 471 | 4384 | 458 | 4800 | 55 | 622 | 152 | 1266 |

| | Promoter | M6-11-14-ALL | M6-11-14-ON | M6-11-15-ALL | M6-11-15-ON | M6-11-17-ALL | M6-11-17-ON | M6-11-18-ALL | M6-11-18-ON | M7-11-14-ALL | M7-11-14-ON | M7-11-15-ALL | M7-11-15-ON | M7-11-17-ALL | M7-11-17-ON | M7-11-18-ALL | M7-11-18-ON |
|---|---|---|---|---|---|---|---|---|---|---|---|---|---|---|---|---|---|
| 1 | cydAWTcI | 241 | 294 | 209 | 280 | 138 | 157 | 120 | 216 | 246 | 468 | 174 | 364 | 131 | 363 | 152 | 188 |
| 2 | cydA5cI | 1 | 0 | 2 | 2 | 0 | 1 | 0 | 0 | 1 | 0 | 4 | 1 | 3 | 0 | 4 | 0 |
| 3 | cydA10cI | 326 | 417 | 257 | 433 | 173 | 228 | 98 | 264 | 374 | 728 | 222 | 608 | 166 | 507 | 206 | 238 |
| 4 | cydA15cI | 8 | 9 | 16 | 25 | 16 | 6 | 1 | 11 | 11 | 11 | 15 | 28 | 11 | 10 | 11 | 6 |
| 5 | cydA17cI | 393 | 555 | 310 | 475 | 223 | 294 | 140 | 320 | 530 | 921 | 294 | 705 | 185 | 574 | 253 | 321 |
| 6 | cydA23cI | 380 | 485 | 307 | 470 | 215 | 304 | 150 | 328 | 445 | 824 | 263 | 635 | 189 | 567 | 277 | 316 |
| 7 | fdnGWTcI | 345 | 334 | 363 | 455 | 450 | 504 | 347 | 667 | 353 | 615 | 955 | 1767 | 1720 | 3626 | 2052 | 2097 |
| 8 | fdnG5cI | 343 | 0 | 275 | 1 | 235 | 9 | 164 | 9 | 359 | 1 | 264 | 6 | 231 | 6 | 287 | 2 |
| 9 | fdnG10cI | 0 | 0 | 1 | 0 | 0 | 1 | 0 | 0 | 0 | 0 | 2 | 0 | 1 | 0 | 2 | 0 |
| 10 | fdnG15cI | 3 | 0 | 8 | 0 | 6 | 1 | 7 | 0 | 2 | 0 | 2 | 0 | 8 | 0 | 5 | 0 |
| 11 | fdnG17cI | 1 | 0 | 0 | 0 | 1 | 0 | 0 | 0 | 1 | 0 | 2 | 0 | 3 | 0 | 1 | 0 |
| 12 | fdnG23cI | 3 | 0 | 3 | 0 | 4 | 1 | 1 | 0 | 2 | 0 | 1 | 0 | 2 | 0 | 4 | 0 |
| 13 | hycAWTcI | 445 | 44 | 422 | 48 | 389 | 12 | 302 | 26 | 477 | 43 | 377 | 64 | 447 | 57 | 567 | 40 |
| 14 | hycA5cI | 369 | 1 | 357 | 3 | 343 | 0 | 222 | 0 | 388 | 0 | 322 | 0 | 329 | 4 | 681 | 0 |
| 15 | hycA10cI | 1 | 0 | 4 | 0 | 4 | 0 | 1 | 0 | 2 | 0 | 1 | 0 | 3 | 0 | 8 | 0 |
| 16 | hycA15cI | 349 | 10 | 291 | 12 | 266 | 8 | 168 | 5 | 319 | 11 | 314 | 14 | 284 | 5 | 327 | 1 |
| 17 | hycA17cI | 268 | 0 | 259 | 0 | 221 | 1 | 160 | 1 | 263 | 1 | 203 | 0 | 254 | 1 | 334 | 0 |
| 18 | hycA23cI | 5 | 0 | 9 | 0 | 12 | 0 | 7 | 0 | 6 | 0 | 10 | 0 | 14 | 0 | 14 | 0 |
| 19 | moaAWTcI | 419 | 0 | 454 | 4 | 398 | 2 | 287 | 0 | 446 | 0 | 406 | 1 | 435 | 1 | 581 | 2 |
| 20 | moaA5cI | 178 | 264 | 216 | 342 | 123 | 190 | 109 | 211 | 222 | 461 | 205 | 482 | 143 | 305 | 168 | 180 |
| 21 | moaA10cI | 317 | 426 | 437 | 661 | 227 | 344 | 167 | 404 | 376 | 740 | 374 | 891 | 241 | 664 | 388 | 396 |
| 22 | moaA15cI | 1138 | 1480 | 1239 | 1819 | 687 | 986 | 559 | 1205 | 1261 | 2381 | 1134 | 2719 | 674 | 2001 | 1038 | 1109 |
| 23 | moaA17cI | 20 | 70 | 19 | 74 | 24 | 39 | 8 | 53 | 22 | 106 | 37 | 152 | 19 | 75 | 17 | 32 |
| 24 | moaA23cI | 515 | 700 | 694 | 968 | 404 | 560 | 295 | 610 | 580 | 1129 | 572 | 1253 | 365 | 890 | 587 | 538 |
| 25 | napFWTcI | 1 | 1 | 1 | 1 | 1 | 1 | 1 | 2 | 1 | 1 | 0 | 1 | 1 | 0 | 0 | 0 |
| 26 | napF5cI | 380 | 455 | 399 | 438 | 206 | 213 | 178 | 254 | 347 | 549 | 304 | 561 | 143 | 249 | 216 | 184 |
| 27 | napF10cI | 8 | 9 | 8 | 19 | 6 | 10 | 2 | 14 | 11 | 11 | 7 | 16 | 6 | 8 | 15 | 8 |
| 28 | napF15cI | 407 | 576 | 499 | 466 | 195 | 232 | 199 | 298 | 327 | 512 | 321 | 561 | 130 | 183 | 203 | 149 |
| 29 | napF17cI | 5 | 0 | 11 | 4 | 4 | 5 | 1 | 3 | 4 | 3 | 4 | 9 | 7 | 1 | 6 | 1 |
| 30 | napF23cI | 6506 | 7450 | 8375 | 9438 | 4601 | 5874 | 4173 | 7410 | 5079 | 7896 | 4528 | 9365 | 2624 | 5796 | 4550 | 3999 |
| 31 | narKWTcI | 637 | 5 | 607 | 3 | 452 | 2 | 389 | 1 | 616 | 3 | 534 | 4 | 561 | 18 | 841 | 5 |
| 32 | narK5cI | 552 | 1 | 467 | 1 | 459 | 1 | 294 | 2 | 571 | 0 | 358 | 0 | 444 | 3 | 498 | 1 |
| 33 | narK10cI | 2 | 0 | 3 | 0 | 4 | 0 | 2 | 0 | 2 | 0 | 4 | 0 | 3 | 0 | 6 | 0 |
| 34 | narK15cI | 7 | 0 | 20 | 0 | 13 | 0 | 6 | 0 | 9 | 1 | 11 | 0 | 13 | 0 | 19 | 0 |
| 35 | narK17cI | 1 | 0 | 1 | 0 | 4 | 0 | 1 | 0 | 3 | 0 | 3 | 0 | 3 | 0 | 5 | 0 |
| 36 | narK23cI | 346 | 6 | 245 | 10 | 263 | 5 | 159 | 4 | 361 | 8 | 235 | 8 | 308 | 16 | 346 | 3 |
| 37 | narGWTcI | 1 | 0 | 0 | 0 | 0 | 0 | 0 | 0 | 0 | 1 | 0 | 0 | 0 | 0 | 1 | 0 |
| 38 | narG5cI | 0 | 0 | 0 | 0 | 0 | 0 | 0 | 0 | 0 | 0 | 0 | 0 | 0 | 0 | 0 | 0 |
| 39 | narG10cI | 0 | 0 | 0 | 0 | 0 | 0 | 0 | 0 | 0 | 0 | 0 | 0 | 0 | 0 | 0 | 0 |
| 40 | narG15cI | 0 | 0 | 0 | 0 | 0 | 0 | 0 | 0 | 0 | 0 | 0 | 0 | 0 | 0 | 0 | 0 |
| 41 | narG17cI | 0 | 0 | 0 | 0 | 0 | 0 | 0 | 0 | 0 | 0 | 0 | 0 | 0 | 0 | 0 | 0 |
| 42 | narG23cI | 1 | 3 | 1 | 0 | 0 | 1 | 2 | 0 | 3 | 0 | 2 | 2 | 2 | 1 | 0 | 1 |
| 43 | torCWTcI | 0 | 0 | 0 | 0 | 0 | 0 | 1 | 0 | 0 | 0 | 0 | 0 | 0 | 0 | 0 | 0 |
| 44 | torC5cI | 4421 | 4014 | 4970 | 4160 | 3254 | 1762 | 3030 | 2485 | 3357 | 3965 | 3129 | 4116 | 2294 | 748 | 3442 | 1263 |
| 45 | torC10cI | 20 | 15 | 25 | 34 | 17 | 16 | 12 | 16 | 13 | 14 | 16 | 31 | 14 | 2 | 27 | 14 |
| 46 | torC15cI | 8075 | 7225 | 10102 | 8472 | 6770 | 4066 | 6350 | 6540 | 5948 | 6979 | 5571 | 7800 | 4303 | 1583 | 6299 | 2673 |
| 47 | torC17cI | 4 | 6 | 10 | 17 | 8 | 1 | 7 | 7 | 3 | 5 | 6 | 13 | 11 | 1 | 7 | 1 |
| 48 | torC23cI | 56 | 55 | 175 | 208 | 67 | 66 | 104 | 126 | 52 | 59 | 114 | 188 | 74 | 30 | 113 | 52 |
| 49 | yeaRWTcI | 1067 | 151 | 1061 | 286 | 801 | 781 | 686 | 1045 | 1108 | 176 | 982 | 403 | 958 | 998 | 1183 | 500 |
| 50 | yeaR5cI | 1 | 0 | 0 | 0 | 2 | 0 | 1 | 0 | 1 | 0 | 0 | 0 | 1 | 0 | 1 | 0 |
| 51 | yeaR10cI | 174 | 15 | 128 | 22 | 125 | 84 | 77 | 101 | 156 | 21 | 130 | 53 | 125 | 82 | 160 | 37 |
| 52 | yeaR15cI | 229 | 13 | 387 | 71 | 240 | 177 | 180 | 190 | 264 | 26 | 207 | 49 | 247 | 168 | 320 | 67 |
| 53 | yeaR17cI | 0 | 0 | 3 | 0 | 0 | 0 | 1 | 0 | 0 | 0 | 0 | 0 | 0 | 0 | 2 | 0 |
| 54 | yeaR23cI | 0 | 0 | 9 | 1 | 2 | 8 | 0 | 2 | 2 | 0 | 3 | 1 | 5 | 4 | 6 | 1 |
| 55 | yedYWTcI | 0 | 0 | 0 | 0 | 1 | 0 | 0 | 0 | 0 | 0 | 0 | 0 | 0 | 0 | 0 | 0 |
| 56 | yedY5cI | 2 | 0 | 8 | 0 | 3 | 0 | 2 | 0 | 1 | 0 | 5 | 0 | 10 | 0 | 8 | 0 |
| 57 | yedY10cI | 394 | 0 | 405 | 2 | 271 | 1 | 236 | 0 | 402 | 2 | 359 | 3 | 371 | 4 | 495 | 1 |
| 58 | yedY15cI | 605 | 2 | 774 | 5 | 585 | 0 | 466 | 0 | 632 | 3 | 516 | 1 | 535 | 8 | 788 | 0 |
| 59 | yedY17cI | 403 | 4 | 419 | 3 | 363 | 1 | 291 | 0 | 418 | 2 | 366 | 3 | 383 | 7 | 502 | 0 |
| 60 | yedY23cI | 1296 | 3 | 1390 | 5 | 1016 | 5 | 891 | 4 | 1264 | 2 | 1121 | 3 | 1101 | 15 | 1639 | 0 |
| 61 | ynfEWTcI | 2521 | 11 | 2114 | 17 | 1927 | 12 | 1473 | 5 | 2664 | 10 | 2051 | 18 | 2016 | 32 | 2795 | 4 |
| 62 | ynfE5cI | 375 | 0 | 312 | 5 | 243 | 2 | 206 | 1 | 415 | 2 | 258 | 1 | 287 | 2 | 379 | 1 |
| 63 | ynfE10cI | 781 | 865 | 828 | 995 | 575 | 618 | 450 | 767 | 903 | 1431 | 758 | 1482 | 458 | 779 | 720 | 518 |
| 64 | ynfE15cI | 514 | 601 | 576 | 712 | 324 | 427 | 275 | 473 | 589 | 891 | 508 | 993 | 350 | 380 | 396 | 276 |
| 65 | ynfE17cI | 256 | 76 | 243 | 132 | 159 | 79 | 122 | 74 | 306 | 137 | 213 | 154 | 167 | 59 | 257 | 40 |
| 66 | ynfE23cI | 996 | 827 | 1019 | 1013 | 592 | 645 | 486 | 683 | 1012 | 1181 | 976 | 1341 | 684 | 598 | 877 | 445 |
| 67 | fabR | 332 | 701 | 286 | 2140 | 83 | 475 | 183 | 2922 | 265 | 1974 | 181 | 3260 | 66 | 574 | 145 | 1665 |

| | Promoter | M8-11-14-ALL | M8-11-14-ON | M8-11-15-ALL | M8-11-15-ON | M8-11-18-ALL | M8-11-18-ON_S56 |
|---|---|---|---|---|---|---|---|
| 1 | cydAWTcI | 528 | 972 | 283 | 560 | 282 | 396 |
| 2 | cydA5cI | 11 | 11 | 4 | 5 | 4 | 3 |
| 3 | cydA10cI | 817 | 1859 | 292 | 918 | 272 | 605 |
| 4 | cydA15cI | 40 | 71 | 14 | 27 | 12 | 29 |
| 5 | cydA17cI | 957 | 2197 | 375 | 1136 | 366 | 620 |
| 6 | cydA23cI | 917 | 2031 | 344 | 999 | 401 | 676 |
| 7 | fdnGWTcI | 667 | 999 | 433 | 862 | 528 | 711 |
| 8 | fdnG5cI | 615 | 5 | 408 | 4 | 520 | 19 |
| 9 | fdnG10cI | 8 | 0 | 1 | 0 | 4 | 0 |
| 10 | fdnG15cI | 18 | 1 | 4 | 0 | 15 | 1 |
| 11 | fdnG17cI | 4 | 0 | 4 | 0 | 2 | 0 |
| 12 | fdnG23cI | 5 | 0 | 3 | 1 | 9 | 1 |
| 13 | hycAWTcI | 1065 | 100 | 770 | 80 | 784 | 169 |
| 14 | hycA5cI | 823 | 4 | 636 | 0 | 701 | 1 |
| 15 | hycA10cI | 13 | 0 | 9 | 0 | 9 | 0 |
| 16 | hycA15cI | 733 | 25 | 516 | 21 | 598 | 24 |
| 17 | hycA17cI | 621 | 0 | 428 | 1 | 540 | 1 |
| 18 | hycA23cI | 15 | 1 | 15 | 3 | 15 | 0 |
| 19 | moaAWTcI | 952 | 3 | 775 | 2 | 881 | 2 |
| 20 | moaA5cI | 530 | 1092 | 267 | 633 | 249 | 371 |
| 21 | moaA10cI | 1002 | 1991 | 537 | 1233 | 520 | 670 |
| 22 | moaA15cI | 2728 | 5685 | 1339 | 3255 | 1188 | 1657 |
| 23 | moaA17cI | 119 | 413 | 57 | 199 | 18 | 49 |
| 24 | moaA23cI | 1280 | 2763 | 623 | 1667 | 587 | 793 |
| 25 | napFWTcI | 1 | 0 | 0 | 0 | 0 | 0 |
| 26 | napF5cI | 682 | 1303 | 271 | 598 | 293 | 402 |
| 27 | napF10cI | 14 | 51 | 9 | 20 | 17 | 9 |
| 28 | napF15cI | 554 | 1017 | 214 | 386 | 273 | 313 |
| 29 | napF17cI | 13 | 23 | 7 | 6 | 9 | 6 |
| 30 | napF23cI | 3639 | 6369 | 2404 | 5063 | 3812 | 4776 |
| 31 | narKWTcI | 1400 | 12 | 936 | 10 | 1092 | 32 |
| 32 | narK5cI | 1088 | 4 | 699 | 3 | 829 | 0 |
| 33 | narK10cI | 17 | 0 | 14 | 0 | 7 | 0 |
| 34 | narK15cI | 27 | 1 | 12 | 1 | 18 | 0 |
| 35 | narK17cI | 9 | 1 | 5 | 0 | 7 | 0 |
| 36 | narK23cI | 670 | 31 | 442 | 16 | 521 | 12 |
| 37 | narGWTcI | 1 | 0 | 0 | 0 | 1 | 0 |
| 38 | narG5cI | 0 | 0 | 0 | 0 | 0 | 0 |
| 39 | narG10cI | 0 | 0 | 0 | 0 | 0 | 1 |
| 40 | narG15cI | 0 | 0 | 0 | 0 | 0 | 0 |
| 41 | narG17cI | 0 | 0 | 0 | 0 | 0 | 0 |
| 42 | narG23cI | 3 | 1 | 2 | 1 | 0 | 3 |
| 43 | torCWTcI | 0 | 0 | 0 | 0 | 0 | 0 |
| 44 | torC5cI | 2617 | 3298 | 1584 | 2483 | 2873 | 2951 |
| 45 | torC10cI | 29 | 34 | 8 | 33 | 22 | 36 |
| 46 | torC15cI | 3189 | 4085 | 2150 | 3349 | 4815 | 5288 |
| 47 | torC17cI | 4 | 10 | 7 | 12 | 5 | 8 |
| 48 | torC23cI | 83 | 87 | 54 | 79 | 93 | 110 |
| 49 | yeaRWTcI | 2456 | 552 | 1661 | 716 | 1952 | 2089 |
| 50 | yeaR5cI | 2 | 0 | 0 | 0 | 2 | 0 |
| 51 | yeaR10cI | 363 | 42 | 226 | 62 | 270 | 186 |
| 52 | yeaR15cI | 549 | 65 | 390 | 84 | 484 | 329 |
| 53 | yeaR17cI | 2 | 0 | 3 | 2 | 1 | 1 |
| 54 | yeaR23cI | 4 | 1 | 7 | 7 | 9 | 8 |
| 55 | yedYWTcI | 0 | 0 | 0 | 0 | 0 | 0 |
| 56 | yedY5cI | 19 | 0 | 16 | 0 | 12 | 0 |
| 57 | yedY10cI | 756 | 6 | 607 | 3 | 747 | 1 |
| 58 | yedY15cI | 1242 | 4 | 966 | 3 | 1269 | 2 |
| 59 | yedY17cI | 789 | 2 | 670 | 2 | 800 | 0 |
| 60 | yedY23cI | 2589 | 8 | 2001 | 5 | 2785 | 6 |
| 61 | ynfEWTcI | 5666 | 49 | 3750 | 31 | 4084 | 20 |
| 62 | ynfE5cI | 800 | 10 | 469 | 7 | 628 | 2 |
| 63 | ynfE10cI | 2278 | 3784 | 977 | 2393 | 992 | 1160 |
| 64 | ynfE15cI | 1543 | 2500 | 631 | 1367 | 602 | 760 |
| 65 | ynfE17cI | 539 | 545 | 287 | 391 | 331 | 139 |
| 66 | ynfE23cI | 2404 | 3571 | 1118 | 2107 | 1120 | 1168 |
| 67 | fabR | 240 | 3120 | 153 | 2055 | 257 | 2558 |

```
>xapA
GAGGCAATTCCGACGTCTAAGAAACCATTATTATCATGACATTAACCTATAAAAATAGGCGTATCACGAGGCCCTTTCGTCTTCACCTC
GATAGATCTCACGCCGATATCTGCAAAACCGCCTTTAAATTCTGCATGAGATACAACAGGTATCGCTAATATAGAGGAAAGTGTCAGAG
TTAAAAGATGTTTTTTCATAGCATAATTCCCTATGCCGATCGTTATTAGTAGATTGCTTAATAAAGTGGTTTGCTAAAAACGACATATT
GTTAATGCAAAGGAATTAATATCGCCAATACAGTTTTTACCGTGGATTAATAGAGAAAAGATATAAATCCATCGATATCGAATCGCTAT
TGAATGCGTGAGAAAAGTGTATTGGTAAGAGCCGGGACAACCTCGCTATAAAAAGGCGTTAGATTCCACCCTACAGAAAAAGGATATGT
ATGTCTCAGGTTCAATTTTCTCATAACCCGCTGTTTTGCATAGATATTATCAAGACTTATAAACCTGATTTCACGCCACGCTCGAGAGA
TCCTCTAGATTTAAGAAGGAGATATACATATG
>abrB
GAGGCAATTCCGACGTCTAAGAAACCATTATTATCATGACATTAACCTATAAAAATAGGCGTATCACGAGGCCCTTTCGTCTTCACCTC
GATGGATCCTTCGTCCATTCCAGGCACCCCCATCGAGGCTGGCCCCTTCGATACAGTGGTGCTTTTTAAGATTAACTATAACTGATTTA
CCATGATGCCCTTTGCCGCCGGAAAGGGCATCTCTTTCAGAAAATTGCTGATTTTCCCCTTTTCCTTGATTGCTGTTATTTTTTCCTGG
CCCAGCTTTCTTTATGCTGTATCGATACACCAAAGAGAAAATAATGAGGGAGCGAAGGATGCCAGTTTTGCAGTGGGGAATGTTCTCGA
GAGATCCTCTAGATTTAAGAAGGAGATATACATATG
>speF
GAGGCAATTCCGACGTCTAAGAAACCATTATTATCATGACATTAACCTATAAAAATAGGCGTATCACGAGGCCCTTTCGTCTTCACCTC
GATGGATCCTTCAACCGTATTTCCTGGATAGAATATCTTCACCTTCATTCACATCAGGAAAGGTAAATTAAATGGAAAATAACAGCCGC
ACTATGCCCCATATAAGGCGGACAACTCATATTATGAAGTTTGCTCATCGCAATAGCTTCGACTTTCACTTCTTCAATGCCCGTTAGTC
TACCGACTAAGGGCACTTCAGCGTACAGGTCTTCCTGACTCTCTGTATTACAGGCATATGCCTGTATTCCGCAGTGCTCATTCTCTGTT
AACCCTTGTATTATCAGCCACCTGACTGGTAAGGGTTTCTGCATTCTGAAAAAAGCAATCTACCTGATTTATTTATCAGCGGGGTAACT
TTGCTTTTTTCCGGCACGATCGATTTCTCATTCGAGAAATTGAGGACCTGCTATTACCTAAAATAAAGAGATGAAAAATGTCAAAATTA
AAAATTGCGGTTAGTGATTCTTGCCCGGACTGTTTTACCACGCAGCGAGAATGTATCTACATTAATGAAAGTCGTAATATCGATGTGGC
GGCAATAGTTTTATCGCTCAACGATGTTACATGCGGAAAACTCGATGAAATCGATGCCACGGGTTATGGCATCCCGGTATTTATTGCTA
CTGAAAATCAAGAACGTGTACCCGCAGAGTATTTGCCCCGTATTTCGGGTGTCTTTGAGAATTGCGAATCGCGACGAGAATTTTATGGT
CGCCAGTTAGAAACCGCCTCGAGAGATCCTCTAGATTTAAGAAGGAGATATACATATG
>xseA
GAGGCAATTCCGACGTCTAAGAAACCATTATTATCATGACATTAACCTATAAAAATAGGCGTATCACGAGGCCCTTTCGTCTTCACCTC
GAGTCGGCAGAACGGTAGAGTGAGCAGGAACGAGGAGAACGTCGTCAAACGTCAGAGCTTCTTTAGCGATACGTAGCATGGGCAATATC
TCGACCAGAGTGGTTAATAAATATTGCCGCGGCATTATACAGAGCGTAACCGATTGCATCTACCCCTTTTTGCAAAAAATGCTTGCTAT
CCCCGAAGGGCGGGTTACTATCGACTGAATAACCTGCTGATTTAGAATTTGATCTCGCTCACATGTTACCTTCTCAATCCCCTGCAATT
TTTACCGTTAGTCGCCTGAATCAAACGGTTCGTCTGCTGCTTGAGCATGAGATGGGACAGGTTTGGATCAGCGGGGATCCCGCTCTAGA
TTTAAGAAGGAGATATACATATG
>xseB
GAGGCAATTCCGACGTCTAAGAAACCATTATTATCATGACATTAACCTATAAAAATAGGCGTATCACGAGGCCCTTTCGTCTTCACCTC
GATGGATCCGCGCACAGATTGGCTTTTGATGGTGATTTCCGGGAACAATTTAATGATAAACTTCATGGCGGCAATGGTTCGTTGGCAAG
CCTTAAGCGACTTGTATAGGGAAAAATACAGCAGCCCACACCTGCGGCTGCATCCAGGCGCGGAAGTATACCACTAACATCGCTTTGCT
GTGCACATCACCTTACCATTGCGCGTTATTTGCTATTTGCCCTGAGTCCGTTACCATGACGGGGCGAAAAATATTGAGAGTCAGACATT
CATTATGCCGAAGAAAAATGAGGCGCCCGCCAGCTTTGAAAAGGCGCTGAGCGAGCTGGAACAGATTGTAACCCGTCTGGAAAGTGGCG
ACCTGCTCGAGAGATCCTCTAGATTTAAGAAGGAGATATACATATG
>speE
GAGGCAATTCCGACGTCTAAGAAACCATTATTATCATGACATTAACCTATAAAAATAGGCGTATCACGAGGCCCTTTCGTCTTCACCTC
GATGGATCCGCATTCCCGTCGCTAAAAAATGCAAAGCCCTGGCCCGCGATTCCTTAAGCCTGCTTGCCTACGTCAAATAATCCCTGATA
CCTTTTTGTAGAAATATTGACCGTGCATCCGCGGTCAATGTTAGCTATTATGTTGCGCCCTTTTTTTACGGGTGTTAACAAAGGAGGTA
TCAACCCATGGCCGAAAAAAAACAGTGGCATGAAACGCTACACGACCAGTTTGGGCAGTACTTTGCGGTAGATAACGTTCTGTATCATG
AAAAGACCGATCACCAGGACTCGAGAGATCCTCTAGATTTAAGAAGGAGATATACATATG
>speB
GAGGCAATTCCGACGTCTAAGAAACCATTATTATCATGACATTAACCTATAAAAATAGGCGTATCACGAGGCCCTTTCGTCTTCACCTC
GATGGATCCCGAGGCAGGTTTGTACGGTTATACTTATCTTGAAGATGAGTAAGTCCTGTGTTACTTGAATCCGCTTAATTTAGCGGTGA
TAATCCGCCACAATTTATTGTGACAAATCCAACCCTTCCTCGTCGGGCCTAACGACGCGGAAGGGTTTTTTTATATCGACTTTGTAATA
GGAGTCCATCCATGAGCACCTTAGGTCATCAATACGATAACTCACTGGTTTCCAATGCCTTTGGTTTTTTACGCCTGCCGATGAACTTC
CAGCCGTCTCGAGAGATCCTCTAGATTTAAGAAGGAGATATACATATG
>speC
GAGGCAATTCCGACGTCTAAGAAACCATTATTATCATGACATTAACCTATAAAAATAGGCGTATCACGAGGCCCTTTCGTCTTCACCTC
GATGGATCCTAAAAGGGCCGATGACCACAAGAGTTCCTGTTAGCTGTATTGTTTGATTTTCTTATTATGACCGTTTGGTGCCGTCGATG
GCTTGTTTGTTCGCAAAGTCCTGGCTTGCACGCTTTAGCGAAAGGTGCCATGATTGCGCGAATTTTCTCCTCTCTGTACGGAGTTTGCC
CGATGCACGCCATCTCCTTACATTCTCTCGCTTATCGCCGTTTCGCGCGAAACGTTTCCCTTTCTATGTTACTGCTCATGCGGTGAAGT
TAACGCACGCTCACTGCAGGACAACAGTAAAATCAGAGCGTTTCTGCTTTTACTGATGTCTGGCGGTCGGAGCTGGTGACCAGTTTGAC
CCATATCTCATGGGGCAGGGTTTTCCACCTTGTCCGGTATTCTTACTTCCCCGAAACGGGTTTGCGCTTATGAAATCAATGAATATTGC
CGCCAGTAGTGAACTCTCGAGAGATCCTCTAGATTTAAGAAGGAGATATACATATG
>ybcL
GAGGCAATTCCGACGTCTAAGAAACCATTATTATCATGACATTAACCTATAAAAATAGGCGTATCACGAGGCCCTTTCGTCTTCACCTC
GAGTGAAAGTAATATCCGCCCAGCAGGCGATAAGTGCTCTCAAATATATGGTTGATGGTGAGATTTATTTCTAAATAATGATCTCGGAT
TTTAAGTTATGCTATGGTGATAAAGTGCAAGACAGAATTAATTATCTTTGACGAAACTTAATGGGTAATTACTTTGTTTGCTCCCACAA
```

GCGAGTTTTGTACGGCTGTATTGGGGTAGTAAATGAGCTATACAATCTTAATCATTTGTTAGGTGAGAACTCTTGGTCGCAGATTCAAA
TACTGAAAATACGTGACAAATTATTATGAGCAAAATGGTGTATGTCACGTATTTTGAATGGTAGGTTAAAAAATAACACCGACTTTCGT
AGGTGTTACTAATAATAAAGCAGAGTTTTTAGATAGTATCAATGTGCTTTGTGTATATTGTGGCAAATAATTGGGTTGGGGGTACAATT
GTGATTGCTTTTGCATGAACATTGCGCCTTTATGCATAATGAGATAAAGGAATATCAAATAAAATAACGATAGGTCATAACAAAGAGGT
TTTTATGAAAACACTTATCGTTTCAACTGTATTGGCATTCATAACATTTTCTGCGCAGGCTGGATCCTCTAGATTTAAGAAGGAGATAT
ACATATG
>ytfE
GAGGCAATTCCGACGTCTAAGAAACCATTATTATCATGACATTAACCTATAAAAATAGGCGTATCACGAGGCCCTTTCGTCTTCACCTC
GATGGATCCATCGCGGTCAGAGCAAAACCAGAAAAACCTTTAACTGAATCTGTCTCAGAAAGTTGACACGCTGGCAGTGAGTTAAATAA
GCCTCTGCTACGTAAGGGTTATAGCTTTTGCCTTAAAGATGCATTTAAAATACATCTTATCTTATTAAGAATGAGGTATCAGCTATGGC
TTATCGCGACCAACCTTTAGGTGAACTGGCGCTCTCTATTCCTCGCGCTTCAGCTCTGTTTCGTAAATATGATATGGATTACTGCTGTG
GCGGCTCGAGAGATCCTCTAGATTTAAGAAGGAGATATACATATG
>ybcK
GAGGCAATTCCGACGTCTAAGAAACCATTATTATCATGACATTAACCTATAAAAATAGGCGTATCACGAGGCCCTTTCGTCTTCACCTC
GAGCTGCCAGCCATTATAGGCATGATGTTGATTTGTGCCGGTGTGTTGATTATTAATTTATTGTCACGAAGCACACCACATTAAAATAA
TTTGTTTCTAAACGACTAAAATATGGAGGCTCTTATATTTATATGAGCCTCGTTTTATGCTTTTTGTTAATGTCTTTATTTTTTATGTA
TTCTTTTGTGCTTTCAAGATTATGGCGTAAGAAAATTGCAATACGATTATTGTTGTATATTCAAGATAATGTGACCTTAATTGTCTTTT
TAAATAAAAAATAAACAAAAATTATATCCCACCACTAAGGTTTATAAAAGCATACGTTAGCAGGTGTCACCATGAAAAAAGCCATAGCA
TATATGCGATTTTCATCACCAGGTCAGATGTCTGGCGGATCCTCTAGATTTAAGAAGGAGATATACATATG
>frdA
GAGGCAATTCCGACGTCTAAGAAACCATTATTATCATGACATTAACCTATAAAAATAGGCGTATCACGAGGCCCTTTCGTCTTCACCTC
GATGGATCCTAGGAATGGATGCGCTCGGCTGCCAGGATGCCGTTTCGCTCATAGTTAAATCTCCAGTTTTTGACAAGGGCACGAAGTCT
ACTCGCAACGCGACGGCGAGACAAATTTTACGCAGGAATCAAACAGCGGTGGGCAGTGACTAAAAAAAGCACGATCTGATGGTTTAGTA
ATTAAATTAATCATCTTCAGTGATAATTTAGCCCTCTTGCGCACTAAAAAAATCGATCTCGTCAAATTTCAGACTTATCCATCAGACTA
TACTGTTGTACCTATAAAGGAGCAGTGGAATAGCGTTCGCAGACCGTAACTTTCAGGTACTTACCCTGAAGTACGTGGCTGTGGGATAA
AAACAATCTGGAGGAATGTCGTGCAAACCTTTCAAGCCGATCTTGCCATTGTAGGCGCCGGTGGCGCGGGATTACGTGCTGCAATTGCT
GCCGCGCAGGCAAATCCGAATGCAACTCGAGAGATCCTCTAGATTTAAGAAGGAGATATACATATG
>ytfF
GAGGCAATTCCGACGTCTAAGAAACCATTATTATCATGACATTAACCTATAAAAATAGGCGTATCACGAGGCCCTTTCGTCTTCACCTC
GATGGATCCGCAAATTGATTGGCCACCCAACGACAACGTTAGCCGAAAGCGTAAGCCATCTTTTTAATGTTAATAACTAGTTAATTAAA
GTGGCATCCTCCCGCATCCTCTCTGATAATGACGGGATGCCGGGAGCAATCATGTCTGCTTCCTGAACTTTCTTCTGACAGATCAATGG
ATGCCAGTAATGATTAGCGGCGTGCTGTACGCCCTGTTAGCAGGGTTGATGTGGGGGCTTATTTTTGTCGGGCCGTTGATCGTGCCGGA
ATACCCTCGAGAGATCCTCTAGATTTAAGAAGGAGATATACATATG
>ybcY
GAGGCAATTCCGACGTCTAAGAAACCATTATTATCATGACATTAACCTATAAAAATAGGCGTATCACGAGGCCCTTTCGTCTTCACCTC
GATGGATCCTGACGGTCTTCCAGCCATTCGGCTCCTGCTGTATTGAAGCATACCAGGCTATTTCAATATCGCTATGCTGCGGCAGCATT
TAACCCCTTGTAATTCATTGCCATAATTGATTTAATTCACAAATAAAACTATAACATGGTGAAATTAATAAAAAAACACAGATGATGGG
GCTAGAATTTACACACCACTTACCCTAAAGCTTTATGACTGGTGGGTTTTGGGAGTATCAAATCGGCTTGCATGGGGATGTCCTACAAA
GGAACACCTTCTTCCACACTTTCTGGAACATTTAGGTAACAACCATCTGGATATTGGTGTTGGAACTGGGTTTTACCTTACTCACGTAC
CTGAGAGTAGTCTGATATCTTTAATGGATTTGAACGAAGCTAGCCTGAACGCGGCATCTACAAGGGCTGGGGAATCAAAAATTAAACAT
AAAATTAGCCATGATGTTTTTGAACCTTATCCCGCCTCGAGAGATCCTCTAGATTTAAGAAGGAGATATACATATG
>pgpB
GAGGCAATTCCGACGTCTAAGAAACCATTATTATCATGACATTAACCTATAAAAATAGGCGTATCACGAGGCCCTTTCGTCTTCACCTC
GAGGTGTCCGGTTGCCAGTTCTTCAAATCCCACCATCAGGAAATCGCCCCATGGGGTTGGCAGTTTGGCTTCTGCCACACGTTTAAGCT
GCATGAAATTCTCCAGATAATGCTGGTTCTGTATTGGCTTATTTTGCCATAACGAGAAGGGGTTCACCTAATCACGACGCGTCGATCGT
TCACGGAATGGCACAAATCTGTCAATTTTTCCTGGAACTGGCGTTTTCAGTTATGATTGTGGGACTTATCAAAAAGGAGAGGCCATGCG
TTCGATTGCCAGACGTACCGCAGTGGGAGCTGCACTATTGCTTGTCATGCCAGTAGCCGTATGGATTTCTGGCTGGCGTTGGGATCCTC
TAGATTTAAGAAGGAGATATACATATG
>ybcV
GAGGCAATTCCGACGTCTAAGAAACCATTATTATCATGACATTAACCTATAAAAATAGGCGTATCACGAGGCCCTTTCGTCTTCACCTC
GATGGATCCCCACGGAGGGATCATCAGCAGATTGTTCTTTATTCATTTTGTCGCTCCATGCGCTTGCTCTTCATCTAGCGGTTAAAATA
TTACTTCAAATCTTTCTGTATGAAGGTTTGAGCACGTTGGCCTTACATACATCTGTCGGTTGTATTTCCCTCCAGAATGCCAGCAGGAC
CGCACTTTGTTACGCAACCAATACTATTAATTGAAAACATTCCTAATATTTGACATAAATCATCAACAAAACACAAAGAGGTCAGACCA
GATTGAAGCGATAAAAACGATAATGCAAACTACGCGCCCTCGTATCACATGGAAGGTTTTACCAATGGCTCAGGTTGCCATTTTTAAAG
AAATATTCGATCAAGTGCGAAAAGATTCTCGAGAGATCCTCTAGATTTAAGAAGGAGATATACATATG
>ybcW
GAGGCAATTCCGACGTCTAAGAAACCATTATTATCATGACATTAACCTATAAAAATAGGCGTATCACGAGGCCCTTTCGTCTTCACCTC
GAGAATCTTTTCGCACTTGATCGAATATTTCTTTAAAAATGGCAACCTGAGCCATTGGTAAAACCTTCCATGTGATACGAGGGCGCGTA
GTTTGCATTATCGTTTTTATCGCTTCAATCTGGTCTGACCTCTTTGTGTTTTGTTGATGATTTATGTCAAATATTAGGAATGTTTTCAA
TTAATAGTATTGGTTGCGTAACAAAGTGCGGTCCTGCTGGCATTCTGGAGGGAAATACAACCGACAGATGTATGTAAGGCCAACGTGCT
CAAACCTTCATACAGAAAGATTTGAAGTAATATTTTAACCGCTAGATGAAGAGCAAGCGCATGGAGCGACAAAATGAATAAAGAACAAT
CTGCTGATGATCCCTCCGTGGGGATCCTCTAGATTTAAGAAGGAGATATACATATG
>kbaZ

```
GAGGCAATTCCGACGTCTAAGAAACCATTATTATCATGACATTAACCTATAAAAATAGGCGTATCACGAGGCCCTTTCGTCTTCACCTC
GAGACACTCCCTTGCTGTCGCAGACGCTGAATGATCTGTTCTCGTCGCTCGCTGGTGCCAGTCACTCGCTTCTCACCTGAAGCGTCGGT
ATTACTCATAGTAAGTCCTTTCGTAAAACTTTCGTTTCATTTCGTTTTGCCTATTAACGCCTTTCTATTAAGCAAATGCAAGCCCACCT
TGCCCATTGACGCAAGCTACTCTCGTTTCAGTGACTTTCATTATGTTTCTTTTGTGAATCAGATCAGAAAACCATTATCTTTCGTTTTA
TTTTTATCTCACCATGACGCAGTATCAACTGAAACAAAACGAAAGATTAATATCGCAGTAATCTGAACTGGAGAGGAAAGTGAAACATC
TGACAGAAATGGTGAGACAGCACAAAGCGGGCAAAACAAATGGAATTTATGCCGTTTGTTCCGCACATCCGCTGGTGCTGGAAGCTGCA
ATGGATCCTCTAGATTTAAGAAGGAGATATACATATG
>ucpA
GAGGCAATTCCGACGTCTAAGAAACCATTATTATCATGACATTAACCTATAAAAATAGGCGTATCACGAGGCCCTTTCGTCTTCACCTC
GATGGATCCCAGCGCAGCAGTGAGCTAACACAACGCCTGAAATGATGCATAAAGCAGCGACTGGATTGAGATTTTCCTGAATTAGTGAG
CTGATCCGCAGCAATATTTTGTTTATCCTGTATTTTCAGAGGGAATGGAGTGTAACGCTCTGTATTAACAAGGAGAGCATTAAAATGGG
TAAACTCACGGGCAAGACAGCCTCGAGAGATCCTCTAGATTTAAGAAGGAGATATACATATG
>cmk
GAGGCAATTCCGACGTCTAAGAAACCATTATTATCATGACATTAACCTATAAAAATAGGCGTATCACGAGGCCCTTTCGTCTTCACCTC
GAGCGCACCCAGTCCATGTTTGACTCTCACCCACCATCAACAGAGCGTGCGCAACACATCCGTGATCGTATCGCCTCTGGTAAGTAAAT
CATTGTCATCTTTCGGGCTGGTCTTCTGCCAGCCCGCTATAATTGCGCAATAAATCCCCATCTGAATACAGACAAAACTGGTTTTTGCA
CACAACGTTAACGATTTGTGGCGTCGGCGCGTATAATGCGCGCGGTTATGTTAACGGTACGCCTGTTTTAAGGAGATAAAGATGACGGC
AATTGCCCCGGTTATTACCAGGATCCTCTAGATTTAAGAAGGAGATATACATATG
>cmr
GAGGCAATTCCGACGTCTAAGAAACCATTATTATCATGACATTAACCTATAAAAATAGGCGTATCACGAGGCCCTTTCGTCTTCACCTC
GAGCGAGATCATCCACGGAGCCGAGTCTGGCGTCGCGTTAATAAGAGAGAATAGAGAGAGATTCAAATTTTCCAGCATAACTTCCCGAC
GCAAAGTGATTAAAAGGGAGCCAATACAGGCAAGTCGTTGAGAATAAAGTGCAGGTTAAACTGGGTAAAGCGGCATCGTCTTATTTCCC
TCAAGCGGCCTGTTTACGGTGGGTGATTGTAACGGGCATAGGTTAAATAAAACTTAAAGAAAGCGTAGCTATACTCGTAATAATGTAAG
AATGTGCTTAACCGTGGTTTCAGCTACAAAATTCGCTTTCTCGTTAGCTGCGCTTTTATTAAACTCTGCGCGATTATTATTGGCGAAGA
AATTGCATGCAAAATAAATTAGCTTCCGGTGCCAGGCTTGGACGTCAGGCGTTACTTTTCCCTCTCTGTCTGGTGCTGGATCCTCTAGA
TTTAAGAAGGAGATATACATATG
>upp
GAGGCAATTCCGACGTCTAAGAAACCATTATTATCATGACATTAACCTATAAAAATAGGCGTATCACGAGGCCCTTTCGTCTTCACCTC
GATGGATCCCAGAGCATTACCCGCGTCAATATCAACACCGGCATCTTTGTAGCTAAGAGAGGTTTTATCGGTCACTGCTTGGGTCCCCA
CGCGTTACTTGCGGTAGAAAAATAAAATTCGGCGCAATTCTAACAGGGAAAGCAAACGTTTGCGAGACTGCTTTACACAACCTTTTTGC
ACGTCTTTTCCCCAGGCGCGCGGCGAAAGAAGACTTGTGCCAGGGTAAAGGTTAGTTTTCGGATGGAATAATCTTCTTTCATAACCATC
TGAATATAAAATAACTTTATCTCAAACCGTTATCATTTTGACTAAAGTCAACGAAAAGAATATTGCCGCCTTGAAGAAAGGAGGTATAA
TCCGTCGATTTTTTTTGTGGCTGCCCCTCAAAGGAGAAAGAGTATGAAGATCGTGGAAGTCAAACACCCACTCGTCAAACACAAGCTGG
GACTGATGCGTGAGCAACTCGAGAGATCCTCTAGATTTAAGAAGGAGATATACATATG
>ydgR
GAGGCAATTCCGACGTCTAAGAAACCATTATTATCATGACATTAACCTATAAAAATAGGCGTATCACGAGGCCCTTTCGTCTTCACCTC
GAGGCACGGGCGTTATACCTGCATTGCCCGCAAGCCCCGCTGTGGCTCTTGTATTATTGAAGATCTTTGTGAATACAAAGAGAAAGTTG
ACATCTGAAGAAAAGGGGTAACACCGATTACCCCATTGATAACCTTTCTTTATCCTCTTTTAAAACATCTTTTTAACGTCAATGATGCC
ATTGCTTAGCGTTATCATCAGGTAATCCGTTTGCGGATAACCAAAAATGCAGGTTAATTGTTTTTTTAATAGCGAAATTTTCTATTCAT
TCGTGATCAAGATCACGATGCTGTCGCAATGTAAAAACATTGCAAGAATCAGGTTAAAATCATTTAAATTTACACACGCAACAAATATT
GACCTACAAAACATTACACTGGCTATTTTTCAGAAACTGGACTATCTCACTAGTAAAAACGCTAAATAGCAGAACATATCGCCGTTAAG
CCATCACCCCCCTCTCAGTGCAGTGAAAAAATCTGCCGTTACGTTTTTTGAAAAATTTAACGCTGGATAACATTTCCCGGAATGGTTGA
ATTCCCCGCCTCAGTTATATGTAACAGATTATTACAAAGGACTTGTCTGAAAGTGCAAGATAGTGAACATTACCTGCCGTTTCCCCTCC
CACTATAACAATTGCGCGTATGTTTTTTATACATAACGCGAGAAAGCACCCCCGTTAATATGGGATGTAAAAAAAGAGGTAAAAGTGTC
CACTGCAAACCAAAAACCAACTGAAAGCGTCAGTTTGAACGCTTTCAAACAACCGAAGGCGTTGGATCCTCTAGATTTAAGAAGGAGAT
ATACATATG
>ydgK
GAGGCAATTCCGACGTCTAAGAAACCATTATTATCATGACATTAACCTATAAAAATAGGCGTATCACGAGGCCCTTTCGTCTTCACCTC
GAGGCAGAGCTGGTTTCTGGCGGGCGTTTGTTTGACCTCGGCCAGGTACCGAAGTCCGTCTGGCACTATGTCCAATAAAGAAAGTAGCG
ATCGGTTATAAAAGCCTTATAATAACGCCCCTAAAATTAGATTTGCCTGTTTGCGTCGCTATTCTGGAGAACCTATGACCACCACAACG
CCACAACGTATTGGAGGCTGGTTGCTTGGCCCGTTGGCCTGGCTTTTAGTCGGGATCCTCTAGATTTAAGAAGGAGATATACATATG
>ydgH
GAGGCAATTCCGACGTCTAAGAAACCATTATTATCATGACATTAACCTATAAAAATAGGCGTATCACGAGGCCCTTTCGTCTTCACCTC
GAGGCGACGGTAAAACCCAGTTTCAGCAGCTGTTCCACTGTTTTTGGCGTTGCTGCAACACGGGTTTCATTGGTTAACCGTTCTCTTGG
TATGCCAATTCGCATGATATTCCCTTCCATCGGTTTTATTGATGATGGTTTGCCTGTGTCAGGAGCCACACAAGCTGCTCATGTACGAG
CTAAATGTTACTCCGTTAAAATAAATTAGTAACAAACGCCCTATAACGTACTGAAAATTATGCCTGTGATCTAGCGCCAAAAATCAGTA
TTTCGGCGTGAACTCGCAAAATATTAACGATTCAGCCGTGATAGTGGGATAAACACCTTAGAACGCCGGATAAAGACTGATAATTGTCT
TCGACGGTCGGGTAAAACGAGACATCGCCCCGGCACGAATCACTACTTAACATTAAATTAACTTATACAATTCAGTTGCTTCAGTAGTA
ATGATGCTGATACGGCTGTTTTTTAAGCATAGACGGTCATTTGAGCAGGATTAAAATTGGCTTAAGGAATGTGATATGAAAAATGACGC
AGACAGTTACACCGTTTAAATGCAATAATCAGCCACGTTTCTCGTTAATAACAATACCAGTACCTGGTTTGCGCAAGGCGAAGGATTAT
TTTTATGAAGCTTAAGAACACCCTCCTGGCGTCGGCACTGCTTTCTGCTATGGCATTCTCCGTTAACGCAGCAACAGAACTGACACCGG
AGCAAGCGGATCCTCTAGATTTAAGAAGGAGATATACATATG
>ydgI
```

```
GAGGCAATTCCGACGTCTAAGAAACCATTATTATCATGACATTAACCTATAAAAATAGGCGTATCACGAGGCCCTTTCGTCTTCACCTC
GAGCCGCGAAGAAAGGTGCTAAGTATTACCACATCACCCGCCAGTGGCAGGAACGTGGTAATAACCTGACCGTCAGCGCAGATCTGTAT
AAATAACTATTCAGTTACTCCTATAAGGCGGCTTGATGCCGCCTTTCGCATTTTTAGTCACAAATTTCCCCGCGACCATTGCATGCAGA
CACATCTCTTCGTAAAATCCCGCGCTTCAGTGGCAAACTCCATTTTTATGCGTTTTGCCCTAATATTTATTCTTCTTATCACGTTTTAA
TCACTGGATATCGATGGAAAAGAAACTGGGACTGAGCGCACTCACCGCGCTGGTATTAAGCTCAATGCTGGGCGCGGGTGTTTTCAGTC
TGCCGCAGGATCCTCTAGATTTAAGAAGGAGATATACATATG
>ydgL
GAGGCAATTCCGACGTCTAAGAAACCATTATTATCATGACATTAACCTATAAAAATAGGCGTATCACGAGGCCCTTTCGTCTTCACCTC
GAGGTTGGCAACCGCATTGATCGTCCCCTATTTCAAACGCTCGTCGCGGGTAAAAGCGACGTTTGTGAATCCGTAATAACCTTACAGTT
AACCTGTTGTCGCCTGCTCTGGATTAACGGATAATAGGCGGCTTTTTTATTTCAGGCCGAAAAATGACTGACTACCTGTTACTGTTTGT
CGGAACTGTACTGGTCAATAACTTTGTACTGGTCAAGTTTCTCGGTCTCTGTCCGTTTATGGGGGTTTCCGGATCCTCTAGATTTAAGA
AGGAGATATACATATG
>tolC
GAGGCAATTCCGACGTCTAAGAAACCATTATTATCATGACATTAACCTATAAAAATAGGCGTATCACGAGGCCCTTTCGTCTTCACCTC
GAGAAGCCGCGATAAAGTGTTTCTCGTGCAATAATTTCTACATCGTTTTTGCCAAATGTAACGGGCAGGTTGTCTGGCTTAAGCATTGT
TAATGTCCTGGCACTAATAGTGAATTAAATGTGAATTTCAGCGACGTTTGACTGCCGTTTGAGCAGTCATGTGTTAAATTGAGGCACAT
TAACGCCCTATGGCACGTAACGCCAACCTTTTGCGGTAGCGGCTTCTGCTAGAATCCGCAATAATTTTACAGTTTGATCGCGCTAAATA
CTGCTTCACCACAAGGAATGCAAATGAAGAAATTGCTCCCCATTCTTATCGGCCTGAGCCTTTCTGGGTTCAGTTCGTTGAGCCAGGCC
GAGAACCTGATGGGATCCTCTAGATTTAAGAAGGAGATATACATATG
>tolB
GAGGCAATTCCGACGTCTAAGAAACCATTATTATCATGACATTAACCTATAAAAATAGGCGTATCACGAGGCCCTTTCGTCTTCACCTC
GAGTTGCGAAGATCCCGAAACCACCAAGCCAGGCAGTATATGAAGTGTTCAAAAACGCGCCATTGGACTTCAAACCGTAATCGCGATGT
TGACTGTTCGGACGGTCAACATCAGGCACCGGTTGCCACGGGGTTCTGGTAGTTTTGTGTATTTTAGTTTGTTAACATTCTGCTAAATT
ATCGTGGGCCATCGGTCCAGATAAGGGAGATATGATGAAGCAGGCATTACGAGTAGCATTTGGTTTTCTCATACTGTGGGCATCAGTTC
TGCATGCTGAAGTCCGCATTGTGATCGACAGCGGGATCCTCTAGATTTAAGAAGGAGATATACATATG
>tolA
GAGGCAATTCCGACGTCTAAGAAACCATTATTATCATGACATTAACCTATAAAAATAGGCGTATCACGAGGCCCTTTCGTCTTCACCTC
GAGGCGCAAAAGATGTGCCTTACGATGAAATAATTAAAGCACTGAACTTGTTACATAGTGCGGGTGTGAAATCGGTTGGTTTAATGACG
CAGCCTATCTAAACATCTGCGTTTCCCTTGCTTGAAAGAGAGCGGGTAACAGGCGAACAGTTTTTGGAAACCGAGAGTGTCAAAGGCAA
CCGAACAAAACGACAAGCTCAAGCGGGCGATAATTATTTCAGCAGTGCTGCATGTCATCTTATTTGCGGCGCTGGATCCTCTAGATTTA
AGAAGGAGATATACATATG
>ydgA
GAGGCAATTCCGACGTCTAAGAAACCATTATTATCATGACATTAACCTATAAAAATAGGCGTATCACGAGGCCCTTTCGTCTTCACCTC
GAGTTATTGCCGCCAACGAATCACCGGTGACTGTCAAAGGCCACGGCCGTTTAGCGCGTGTTTACAACAAGCTGTAAGAGCTTACTGAA
AAAATTAACATCTCTTGCTAAGCTTATTAAAGGCTTATAACACCTTCAGGCGGCCAGTCCGCCTGATTTCATTTTATGGATAATCATTA
TGAATAAATCGCTGGTAGCGGTAGGCGTCATTGTTGCGCTAGGCGTAGTCTGGACAGGCGGCGCATGGTATACAGGCGGATCCTCTAGA
TTTAAGAAGGAGATATACATATG
>ydgD
GAGGCAATTCCGACGTCTAAGAAACCATTATTATCATGACATTAACCTATAAAAATAGGCGTATCACGAGGCCCTTTCGTCTTCACCTC
GAGAACAAAAAGCGCAGGCAGCGAAGAAACACGCCAAGAAACACAGCCATCAGCAACCGGCAAAACCTGCTGCACAACCCGCAGCGTAA
GTTTTCAACAGTAATGCTGGCGCGCCCCCTCGCGCCTGAAAATTACGGTGCTAAGCGGGTAACGTTTAGCACCGCCTTTAGCGGAGGGT
AGTATGGTGGGCCGTTATCGCTTTGAGTTCATTCTGATCATCCTTATTTTATGCGCACTGATTACCGCCCGTTTTTATCTTTCCTGATT
GTAGTTATCTGATTTTACTCCCACTTTCATCCCGTCCCGTCTATAGTATTTATGAGGGTTTGCTTTTAATAATCATAATTACCCACCAG
AGTGTGATATGCGTACAACCATTGCTGTAGTGTTGGGTGCAATTAGTTTGACGTCTGCTTTTGTGTTTGCAGATAAACCAGACGTTGCC
AGATCGGCAAACGATGAGGTGGATCCTCTAGATTTAAGAAGGAGATATACATATG
>entC
GAGGCAATTCCGACGTCTAAGAAACCATTATTATCATGACATTAACCTATAAAAATAGGCGTATCACGAGGCCCTTTCGTCTTCACCTC
GAGAGTCAGCGGCCTGAACTGCGGCTATTCCTGAAAGCAAAAGTCCTGTTAATAGAAGGGCGTTGCGGTAGAGCGGGGCGAGTCTCACA
AATCAGCTTCCTGTTATTAATAAGGTTAAGGGCGTAATGACAAATTCGACAAAGCGCACAATCCGTCCCCTCGCCCCTTTGGGGAGAGG
GTTAGGGTGAGGGGAACAGCCAGCACTGGTGCGAACATTAACCCTCACCCCAGCCCTCACCCTGGAAGGGAGAGGGGGCAGAACGGCGC
AGGACATCACATTGCGCTTATGCGAATCCATCAATAATGCTTCTCATTTTCATTGTAACCACAACCAGATGCAACCCCGAGTTGCAGAT
TGCGTTACCTCAAGAGTTGACATAGTGCGCGTTTGCTTTTAGGTTAGCGACCGAAAATATAAATGATAATCATTATTAAAGCCTTTATC
ATTTTGTGGAGGATGATATGGATACGTCACTGGCTGAGGAAGTACAGCAGACCATGGCAACACTTGCGCCCAATCGCTTTTTCTTTATG
TCGCCGTACCGCGGATCCTCTAGATTTAAGAAGGAGATATACATATG
>entD
GAGGCAATTCCGACGTCTAAGAAACCATTATTATCATGACATTAACCTATAAAAATAGGCGTATCACGAGGCCCTTTCGTCTTCACCTC
GATGGATCCCCGGGACGTACGTGGTATATGAGCGTAAACACCCACTTCTGATGCTAACGTCAGATTGTTGACAAAGTGCGCGTCGTTCA
TGCCGGATGCGGCGTGAACGCCTTATCCGGCCTACAAAAATCTTGCCAATTCAATATATTGCAGGACCATGTAGGCCTGATAAGCGTAG
CGCATCAGGCAGTTTTGCGTTTGTCATCAGTCTCGAATATGGTCGATATGAAAACTACGCATACCTCCCTCCCCTTTGCCGGACATACG
CTGCATTTTGTTGAGTTCGATCCGGCGAATTTTTGTGAGCCTCGAGAGATCCTCTAGATTTAAGAAGGAGATATACATATG
>entF
GAGGCAATTCCGACGTCTAAGAAACCATTATTATCATGACATTAACCTATAAAAATAGGCGTATCACGAGGCCCTTTCGTCTTCACCTC
GAGCATGATGCGCTTTGTTGGCGCGGTGGCTTGATGCAGGGGCTAATCGACCTCTGGCAACCACTTTTCCATGACAGGAGTTGAATATG
```

GCATTCAGTAATCCCTTCGATGATCCGCAGGGAGCGTTTTACATATTGCGCAATGCGCAGGGGCAATTCAGTCTGTGGCCGCAACAATG
CGTCTTACCGGCAGGCTGGGACATTGTGTGTCAGCCGCAGTCACAGGCGTCCTGCCAGCAGTGGCTGGAAGCCCACTGGCGTACTCTGA
CACCGACGAATTTTACCCAGTTGCAGGAGGCACAATGAGCCAGCATTTACCTTTGGTCGCCGCACAGCCCGGCATCTGGATGGCAGAAA
AACTGTCAGAATTACCCTCCGCCTGGAGGGATCCTCTAGATTTAAGAAGGAGATATACATATG
>eco
GAGGCAATTCCGACGTCTAAGAAACCATTATTATCATGACATTAACCTATAAAAATAGGCGTATCACGAGGCCCTTTCGTCTTCACCTC
GAGATTCATCACCGGACCAGGGCGGACGGATACCGTTACTGGCTTTGCGCCAGCGACCAGTGAGTATGCCCCGACGGGATGCATCAATC
TTCACATTGACCTTCCATCATTAACGCGCTCTGAAAATTGAGAGCGACCAAATAAACCGCATAATTAATAAGCCATTTTTATAGCCGCT
AAGATATTAAAGGATGTGTCAAAGATGCATACCCCGATCGGGGTAAAACCTGTAGCAGGATCAAAAGAGTGGCGGGAAGCGTGGCAAAA
ACGGGCTTTTGCTCACATTTCAAATGGTTATAAATATATTTATATAGCGATTGATTCACCAGAGATATTTCTGCTGGTTTGCTCTCTCA
TTAGAATTTAACACTAAAAGAGCAGGTAAAATTGTCTGAATGTTCTTTAAGTTATTCATAAAGCAAATTAATAAATCTGATGAATATGT
TAACCTTCAGCGACATCATCGGTGAAAACCTATAAATGAAGAAGGAAAGCAAAAAAATGAAGACCATTCTACCTGCAGTATTGTTTGCC
GCTTTCGCTACCACTTCCGCGGATCCTCTAGATTTAAGAAGGAGATATACATATG
>entS
GAGGCAATTCCGACGTCTAAGAAACCATTATTATCATGACATTAACCTATAAAAATAGGCGTATCACGAGGCCCTTTCGTCTTCACCTC
GAGATTTTGCCCCAATGAGCAGGCTTAATGCCGTCGCGATAATCAGTAATAACAGCAATCCGGGCACGGCAATGGCGCGTGTCACGGCA
ACAGAACCAGACATACGAACTTCCATGATAATGAAATTAATTATCGTTATCGATCTTATTTGGATATGTTAGCATGTGCAGCCTAAGAA
TAGGTATTTAAAATATTTGATGGCAAGGCATTGTAATGAATAAACAATCCTGGCTGCTTAACCTCAGCCTGTTGAAAACGCACCCGGCG
TTTCGCGCAGTATTCCGGATCCTCTAGATTTAAGAAGGAGATATACATATG
>yacG
GAGGCAATTCCGACGTCTAAGAAACCATTATTATCATGACATTAACCTATAAAAATAGGCGTATCACGAGGCCCTTTCGTCTTCACCTC
GATGGATCCGCCATTCGTTTTATGCCGCTGGACACTGAAAACGGACAGGTACCGGAACGTCTGGATTTCGAACTGGCCTGTTGCTAAGG
AGTAAAAATGTCAGAAACTATTACGGTGAATTGCCCAACCTGCGGGAAAACGGTGGTGTGGGGTGAAATCAGCCCGTTTCGGCCATTTT
GCTCCAAACGCTCGAGAGATCCTCTAGATTTAAGAAGGAGATATACATATG
>fecA
GAGGCAATTCCGACGTCTAAGAAACCATTATTATCATGACATTAACCTATAAAAATAGGCGTATCACGAGGCCCTTTCGTCTTCACCTC
GATGGATCCACCGATGCGATCCTGAACGTTATCGCGCAAACGCTTCCCGTTAAAATTCAGTCTATTACGCGGTACTGGATAAACATTTC
ACCACTGTAAGGAAAATAATTCTTATTTCGATTGTCCTTTTTACCCTTCTCGTTCGACTCATAGCTGAACACAACAAAAATGATGATGG
GGAAGGTATGACGCCGTTACGCGTTTTTCGTAAAACAACACCTTTGGTTAACACCATTCGCCTGAGCCTGCTGCCGCTGGCCGGTCTCT
CGTTTTCCGCTTTTGCTGCCTCGAGAGATCCTCTAGATTTAAGAAGGAGATATACATATG
>yacH
GAGGCAATTCCGACGTCTAAGAAACCATTATTATCATGACATTAACCTATAAAAATAGGCGTATCACGAGGCCCTTTCGTCTTCACCTC
GATGGATCCGGCCATTTGGTTTGCATCCAGGGGTTTGGGCGCAATCCCCTCAGCGGCACGCTCAGCTACGTGCTTACGGTATTCTTCTA
GCACGACGGTTCTCCTCGCTCTCATTGTCATAGTGCGGCAGGCGATTCTCTTCACGCTCCTGTGAGACAGCAGTTTGTAGGGTAAATGC
CCAGTACCGCAACGGGCAGAATAGCAGGATTTTGACGAGGTGTTAATCTGTTTACAAAAAAGCAACATAAAAAAGTGGCTGAATCGTTA
AGGATGGTCTAGAGATTGTTTCCCTCCGTAAAAACTTCGTTAATTCGCATGGTGATAATCACCTTTCTCAACGAATCCCAAGATTAGTC
AAAATTTAAACTACCGCCTCTTTATACTCGGATTCACAGCACCTGCGGGTGGCAGTTCGCCGACCATTGCGATTTCCTTGAGATCCGAA
TTATGAAAATGACTTTGCCGTTTAAACCCCATGTGCTGGCGCTAATTTGCAGTGCCGGGCTTTGTGCCGCCTCTACCTCGAGAGATCCT
CTAGATTTAAGAAGGAGATATACATATG
>yhbO
GAGGCAATTCCGACGTCTAAGAAACCATTATTATCATGACATTAACCTATAAAAATAGGCGTATCACGAGGCCCTTTCGTCTTCACCTC
GAGCGTTGCGCCTGTAATCAGTACCTGACTCATCTTCGCTCCTTTACAGAATTGTCTGCCTTGCGCTCCACGGCTCATGCACCAGGCTT
AATAGACCATGAGGTAATTATCCCCGATTGTGGGAAATTCGCCTCATCCAATGCAACAACGCGGAGGAAGCATGAGTAAGAAAATTGCC
GTTTTAATCACTGATGGGATCCTCTAGATTTAAGAAGGAGATATACATATG
>fecI
GAGGCAATTCCGACGTCTAAGAAACCATTATTATCATGACATTAACCTATAAAAATAGGCGTATCACGAGGCCCTTTCGTCTTCACCTC
GATGGATCCACCACGCCTTCAGTAGCGGAGCAGGAAGGACATCTGATGGAAATGGAAGCCACGCAAGCACCTTAAAATCACCATCATAC
ACTAAATCAGTAAGTTGGCAGCATTACCGAAATCCCCATCCTTTAGGGCAGTGAGGATGTCAATAATAGCAGACCACATCATGAGTCAT
TCTGACCGACACATGCCAGAAGCAGAGGCTCCATCCCATTTTATACCTACCTTATAACACTTAGAAAAACAACATGTTAAAAATGTCTA
TTGGAAACAATTTTATTTCCAATTGTAATGATAACCATTCTCATATTAATATGACTACGTGATAATTAACTTTTGATGCACTCCGCATG
TCTGACCGCGCCACTACCACAGCTTCCTTAACGTTCGAGTCGCTTTATGGCACACATCACGGCTGGTTGACTCGAGAGATCCTCTAGAT
TTAAGAAGGAGATATACATATG
>rcsA
GAGGCAATTCCGACGTCTAAGAAACCATTATTATCATGACATTAACCTATAAAAATAGGCGTATCACGAGGCCCTTTCGTCTTCACCTC
GAGTGCACCTTTTTGCGAACATTTATTCAGTGAAATTTTTAATTTGCTGGCTGATATTATTAGTGAATTGCCATTAATATAATTCCGTA
ACGTTTATCATGTTATCCTAAGGATTATCCGAAAAATAATACCTACGAACATCTTCCAGGATACTCCTGCAGCGAAATATTTGTTTTAA
GCTCACTCACATATCGCAACATTTACTTTACTTTAAGACAATTCCAGGCAAATTATACAACACTTTACGGGATAGTAAGTCCGCCTGAA
AAATCGCGAGAGTGGCGCATTAGGTGACCCATGTTGTTCCGTTTAGTCATGATGAAATATTCAGGTAAGGGGAATTATCGTTACGCATT
GAGTGAGGGTATGCCATGTCAACGATTATTATGGATTTATGTAGTTACACCCGACTAGGTTTAACCGGGTATCTGTTGAGTAGAGGGGT
TAAAAAAAGAGAAATCAACGACATTGAAACCGTTGATGACCTTGCCATAGCTTGTGATTCACAGCGCCCTTCAGGGATCCTCTAGATTT
AAGAAGGAGATATACATATG
>b2394

Data Set S4 - MG1655 library reference file.fasta

GAGGCAATTCCGACGTCTAAGAAACCATTATTATCATGACATTAACCTATAAAAATAGGCGTATCACGAGGCCCTTTCGTCTTCACCTC
GAGACGTGGTTTCTCGCTTCGGTCTGAAGCTGGTTTACGGCTCTACCCTGGTGAGTGTGCTGTCTGCGTCAATCGCAGCACTGGTGCTG
TAAGACCATACATAAAAAAGCCGGGGATAATTCCCATAAGCGCTAACTTAAGGGTTGTGGTATTACGCCTGATATGATTTAACGTGCCG
ATGAATTACTCTCACGATAACTGGTCAGCAATTCTGGCCCATATTGGTAAGCCCGAAGAACTGGATACTTCGGCACGTAATGCCGGATC
CTCTAGATTTAAGAAGGAGATATACATATG
>rcsC
GAGGCAATTCCGACGTCTAAGAAACCATTATTATCATGACATTAACCTATAAAAATAGGCGTATCACGAGGCCCTTTCGTCTTCACCTC
GATGGATCCCGTAAGCGGCGTGGATAAATCCACTTCATATAATGCATGCTGAAATCCACTAGTCTTGTCCGGTATATGACGATTATCAG
AGGTTAAGGTGATGATTTCTCGGCGGTGTATCATATTCCAGAGAAGAGAGAACATTGCGGTAACACGCTTTTACCGCTACCTTAACCAC
ACTCCATCGGTCACCTGAGGCGGAGCTTCGCCCCTTTGAAATACCTTGCTTCTTTTCGTACAACCCTGAAAGCCTCGCGCTACATGTTC
AGAGCATTGGCGTTAGTGCTCTGGCTGTTGATTGCTTTTTCATCCGTTTTTACATCGTTAATGCGTTACATCAGCGAGAATCGGAAAT
TCGCTCGAGAGATCCTCTAGATTTAAGAAGGAGATATACATATG
>rcsF
GAGGCAATTCCGACGTCTAAGAAACCATTATTATCATGACATTAACCTATAAAAATAGGCGTATCACGAGGCCCTTTCGTCTTCACCTC
GATGGATCCACGCCGAGAACGTGAAGAAATTCGTCCAGGCTTATCAGTCTGACGAAGTTTACGAAGCAGCAAACAAAGTGTTTAACGGC
GGAGCTGTTAAAGGCTGGTAATTTTTAGGCTGTTTCCACAATTTGTAATATCATTCAGGACGGGCGCTTGCCCGTCTTGTCATTTTTAC
AAGCTCCTGATTCAATATTGACGTTTTGATCATACATTGAGGAAATACTATGCGTGCTTTACCGATCTGTTTAGTAGCACTCATGCTAA
GCGGCTGTTCCATGTTAAGCAGATCCCCTGTCGAACCCCTCGAGAGATCCTCTAGATTTAAGAAGGAGATATACATATG
>uspA
GAGGCAATTCCGACGTCTAAGAAACCATTATTATCATGACATTAACCTATAAAAATAGGCGTATCACGAGGCCCTTTCGTCTTCACCTC
GAGCGCGCGTAGTGATGAGAAATAGCGCGCCATGTTAACAATGCAAACGACACATAAAGCCCAAAATAATGCGACGGTGCTTATCATAC
CTCCTCCCCGGCGACCTGCCCGCGGAGTTCCACCCCGGGGCTACCGCTCCCGATACGCTGCCAATCAGTTAACACCAGGTCCTGGAGAA
ACCGCTTTTGTGGTGACCAACATACGAGCGGCTCTATAGATAGTGTAGGAGATCAGGTTGTTTTTTTTCCAGAAGGTTAACCACTATCA
ATATATTCATGTCGAAAATTTGTTTATCTAACGAGTAAGCAAGGCGGATTGACGGATCATCCGGGTCGCTATAAGGTAAGGATGGTCTT
AACACTGAATCTTTACGGCTGGGTTAGCCCCGCGCACGTAGTTCGCAGGACGCGGGTGACGTAACGGCACAAGAAACGCTAGCTGGCCA
GTCATCGACAACTTTATGGAAGGAGTAACACTATGGCTTATAAACACATTCTCATCGCGGTCGACCTCTCCCCGGAAAGCAAAGTTCTG
GTAGAGAAAGCAGTCTCTATGGCTCGCCCCTACAAGGATCCTCTAGATTTAAGAAGGAGATATACATATG
>uspB
GAGGCAATTCCGACGTCTAAGAAACCATTATTATCATGACATTAACCTATAAAAATAGGCGTATCACGAGGCCCTTTCGTCTTCACCTC
GATGGATCCTTGTAGGGGCGAGCCATAGAGACTGCTTTCTCTACCAGAACTTTGCTTTCCGGGGAGAGGTCGACCGCGATGAGAATGTG
TTTATAAGCCATAGTGTTACTCCTTCCATAAAGTTGTCGATGACTGGCCAGCTAGCGTTTCTTGTGCCGTTACGTCACCCGCGTCCTGC
GAACTACGTGCGCGGGGCTAACCCAGCCGTAAAGATTCAGTGTTAAGACCATCCTTACCTTATAGCGACCCGGATGATCCGTCAATCCG
CCTTGCTTACTCGTTAGATAAACAAATTTTCGACATGAATATATTGATAGTGGTTAACCTTCTGGAAAAAAAACAACCTGATCTCCTAC
ACTATCTATAGAGCCGCTCGTATGTTGGTCACCACAAAAGCGGTTTCTCCAGGACCTGGTGTTAACTGATTGGCAGCGTATCGGGAGCG
GTAGCCCCGGGGTGGAACTCCGCGGGCAGGTCGCCGGGGAGGAGGTATGATAAGCACCGTCGCATTATTTTGGGCTTTATGTGTCGTTT
GCATTGTTAACATGGCGCGCTATTTCTCATCACTACGCGCGCTCGAGAGATCCTCTAGATTTAAGAAGGAGATATACATATG
>uspE
GAGGCAATTCCGACGTCTAAGAAACCATTATTATCATGACATTAACCTATAAAAATAGGCGTATCACGAGGCCCTTTCGTCTTCACCTC
GATGGATCCAAATAACGATGCGCTGGCCCAGCTTGCTGGTCATACGCGTAACGTTGCCTGATTTTTCCGCATAACTCACTATCCTTCTG
TCATATCATTAAATTTTTCTGATTTATTGATCTGGCAGAAGGTTCATCACTGTTTCATTCACCAGATATGGGTTAATCTTTTAATTACA
AACTGCGTTGACAGTTGTTGTAAGGAGACCCTGTATGGCTATGTATCAGAACATGCTCGTTGTTATCGATCCTAACCAGGACGACCAAC
CAGCATTGCGGCGAGCTGTTTACTCGAGAGATCCTCTAGATTTAAGAAGGAGATATACATATG
>uspF
GAGGCAATTCCGACGTCTAAGAAACCATTATTATCATGACATTAACCTATAAAAATAGGCGTATCACGAGGCCCTTTCGTCTTCACCTC
GATGGATCCCAGCTTCTACGCTGCCAATGGCATCTCTACCGATGATATTGTCGCTTTAGGTCTGGTTTATCAGTTCTAAATCCTCCTGC
CCGCTGTTATGGCGGGCTTTTTCTGCTTATTCTTCCTCCTTTGATCTAAATTAAAAATGTGAACTCCGTCATTACACAAAAAGTGTCAT
CTGGCGTTACACTTTATGCGGATACTAAAACAGGAGGTTTTATGAACAGAACGATTCTTGTCCCTATCGACTCGAGAGATCCTCTAGAT
TTAAGAAGGAGATATACATATG
>uspG
GAGGCAATTCCGACGTCTAAGAAACCATTATTATCATGACATTAACCTATAAAAATAGGCGTATCACGAGGCCCTTTCGTCTTCACCTC
GATGGATCCATCTGCCTGTTCAACGCCCGGATCGGGAACATTTTCTACCTGAACGTGATGTGGGCCGTGATAAGTCAATGCTTTCATGC
AACCTCCAGAGACAGTTCAAAAAAGAGTCAGTCTTTAAGGGTAGCTATCACAGCGGGAAAACGGGCGGTACCACAGGTTTTGGGAGAAT
TCCTGGTGTTCAGTGAGAATATATTCATCACAGAAACAGTTCGTAAGGAAAAAGAACCAGACAAATATCAGGGATAAATATTATAATCA
TGGCGAATGATTTGTTCATGATTAACAGGGAGAAAGGGTTATGTATAAGACAATCATTATGCCAGTTGATGTATTTGAAATGGAATTGA
GCGACAAAGCTGTTCGCCACGCTGAATTCCTCGAGAGATCCTCTAGATTTAAGAAGGAGATATACATATG
>gltX
GAGGCAATTCCGACGTCTAAGAAACCATTATTATCATGACATTAACCTATAAAAATAGGCGTATCACGAGGCCCTTTCGTCTTCACCTC
GATGGATCCAGGGAGGTGCTCTCCCAGCTGAGCTAATCACCCCCGCTGTGTGGAGTCGCATTATAGGGAGAGTTCAAAATGAGTCAACG
CATTTTCTAAAGAAATTGTTCGTTCGTCGTAAATTTAAGCAAGATGATCGCAAAACAGACCGTGTTGCGCAATTTGTCAACGAAAACAA
TAATGCGTAAGGTAGAAACCCGAACTACATTGAGGAATCAGGCGGGAGTGATAGAATATCGCCCACTTAATTTTTCCAGGATTTGCCGG
TTGTCGGCATCTTTCTAAACGTAAGGCCATTTCATGAAAATCAAAACTCGCTTCGCGCCAAGCCCAACAGGCTATCTGCACGTTGGCGG
CGCGCGTACTGCTCTTTACTCCTGGCTTTTTGCACGTCTCGAGAGATCCTCTAGATTTAAGAAGGAGATATACATATG
>yifM2

GAGGCAATTCCGACGTCTAAGAAACCATTATTATCATGACATTAACCTATAAAAATAGGCGTATCACGAGGCCCTTTCGTCTTCACCTC
GAGAGTTGGCGTCGGTGGATAAAAACACCATTGCCTTCTTTAGCCCTAACTATCTACAAGGCTGGCAGCGGGCGTTGGCGATTGCCGCC
AGGGAGGTCGCATGAGTCTGCTGCAATTCAGTGGCCTGTTTGTTGTCTGGCTGCTCTGCACGCTGTTTATTGCCACGCTGACCTGGTTT
GAGTTTCGCCGTGGATCCTCTAGATTTAAGAAGGAGATATACATATG
>gltJ
GAGGCAATTCCGACGTCTAAGAAACCATTATTATCATGACATTAACCTATAAAAATAGGCGTATCACGAGGCCCTTTCGTCTTCACCTC
GATGGATCCAATTCCGCCGAAAAACCTGAACATGAATTTCGAACTGTCAGACGAAATGAAAGCACTGTTCAAAGAACCGAATGACAAGG
CACTGAACTAATTACAAGAACCAGGGGCGGAAATTCCAGCCCTCTCGATTGTTACGTAGCACGGACAGACTATACGCCTGATGGTCGTT
CCCCATCGGGCCTGAAAACCGCAATACGCTGGGTAACAATCTTCGAGGGTAGCAGTTAACGCTGCTACCCTCTTTTTTCTGGAGTAGAT
TTATGTCTATAGACTGGAACTGGGGTATTTTTTTACAACAAGCCCCGTTCGGCAACACCACCTATCTTGGTTGGATCTGGAGTGGCTCG
AGAGATCCTCTAGATTTAAGAAGGAGATATACATATG
>cspI
GAGGCAATTCCGACGTCTAAGAAACCATTATTATCATGACATTAACCTATAAAAATAGGCGTATCACGAGGCCCTTTCGTCTTCACCTC
GATGGATCCGACGAACGCTACATCGCAGGAGTGACTTGTCAACAGATCTATGAATCTTTAAGAGATAAAAAGCATCAAATGTAGGGTAA
TATTAAATCGGAACATTTACATCGCGGAATGTAAAATTTAAATAAAAAGGACTCTTCCATGAGCCAAAATTCCTGAAATCTTAAGGGTA
AGATAAAAGGTCTTAATCAGAATGACACGTTTTATTAATAAATAAAGCTATTCTTTCATTGCTGTGTTTTTCTTTACAAAAGTAATCCT
TGCTATGGGTGGTTAATCATGCGTTAATGGTGTTCTGGTTTGTTACAAATTTATCTGAAGCAGTCATTGTTATAATTTTATTATTTGTA
CCTCTTGAGATTTCCTTGTTGGTTTTTCTCTCTGATATTTTTTTTCGGACCATTCTGCCCAAGGGCTAATTTCTTCAAAAGGTAATAAT
TATGTCTAACAAAATGACTGGTTTAGTGAAATGGTTTAACCCTGAAAAAGGTTTTGGTTTCATCACGCCGAAACTCGAGAGATCCTCTA
GATTTAAGAAGGAGATATACATATG
>gltA
GAGGCAATTCCGACGTCTAAGAAACCATTATTATCATGACATTAACCTATAAAAATAGGCGTATCACGAGGCCCTTTCGTCTTCACCTC
GATGGATCCCGATGGAGAATGGACGCTATCGCCGTGATGGGGAACCGGATGGTCTGTAGGTCCAGATTAACAGGTCTTTGTTTTTTCAC
ATTTCTTATCATGAATAACGCCCACATGCTGTTCTTATTATTCCCTGGGGACTACGGGCACAGAGGTTAACTTTCTGTTACCTGGAGAC
GTCGGGATTTCCTTCCTCCGGTCTGCTTGCGGGTCAGACAGCGTCCTTTCTATAACTGCGCGTCATGCAAAACACTGCTTCCAGATGCG
AAAACGACACGTTACAACGCTGGGTGGCTCGGGATTGCAGGGTGTTCCGGAGACCTGGCGGCAGTATAGGCTGTTCACAAAATCATTAC
AATTAACCTACATATAGTTTGTCGGGTTTTATCCTGAACAGTGATCCAGGTCACGATAACAACATTTATTTAATTTTTAATCATCTAAT
TTGACAATCATTCAACAAAGTTGTTACAAACATTACCAGGAAAAGCATATAATGCGTAAAAGTTATGAAGTCGGTATTTCACCTAAGAT
TAACTTATGTAACAGTGTGGAAGTATTGACCAATTCATTCGGGACAGTTATTAGTGGTAGACAAGTTTAATAATTCGGATTGCTAAGTA
CTTGATTCGCCATTTATTCGTCATCAATGGATCCTTTACCTGCAAGCGCCCAGAGCTCTGTACCCAGGTTTTCCCCTCTTTCACAGAGC
GGCGAGCCAAATAAAAAACGGGTAAAGCCAGGTTGATGTGCGAAGGCAAATTTAAGTTCCGGCAGTCTTACGCAATAAGGCGCTAAGGA
GACCTTAAATGGCTGATACAAAAGCAAAACTCACCCTCAACGGGGATACAGCTGTTGAACTGGATGTGCTGAAAGGCAGTCGAGAGATC
CTCTAGATTTAAGAAGGAGATATACATATG
>cspG
GAGGCAATTCCGACGTCTAAGAAACCATTATTATCATGACATTAACCTATAAAAATAGGCGTATCACGAGGCCCTTTCGTCTTCACCTC
GAGCGTCGGAGGGGATAATGAATCCTTTGCCGCTTTTGCGATCAAAGGTTTTGACAATTCCTGTCATTTTACGGGACAAACAAATTCCT
TACTGAAAATACTGCGCTGCACTATACGGGGTTAATAAAATAAAGCCAGCGATATTTAAGACCGCCGGACGGCTAAAATAAAATTTGCT
TAATCTCAATTATCATGCGTTAATAGCTGCGTCGGTTTGAAAGACAGACAGCATACAAAGTAGTTTACTAAAGCAGTTCTCATTATCAG
GCATTATCCCCTTCTTTTGAGTCTCTCTCCTGAACACTAAGTAGTTTCTGTATTAAAGCCCTGTTTGCCGAAAGGCCCAAAATGAAGGA
AGTAAAATATGTCTAATAAAATGACTGGTTTAGTAAAATGGTTTAACGCAGATAAAGGTTTTGGCTTTATCACTCCTGATGATGGCAGC
AAGGATCCTCTAGATTTAAGAAGGAGATATACATATG
>cspD
GAGGCAATTCCGACGTCTAAGAAACCATTATTATCATGACATTAACCTATAAAAATAGGCGTATCACGAGGCCCTTTCGTCTTCACCTC
GATGGATCCAGTCCAGCCAGTCGTTCGTTTTACCCATTGTCAGTTATCATCTTCGGTTACGGTTATCGGCAGAACGCCCTGCCGCTGAC
AAGAGCTTATTTGCCTGACCAATCTACTCCACATGATAGATAACTATCATCTATCGTTGGCATCAGCGACATCTGTCACATTCCTGTCA
ATAGCGTTAACTGCTTCAAATTTTTGATTCATACCTGCCCCTTGTCCCCTGTCAAATGCTTGACGGCTCGCCCTAATTCTCTAAATTGT
ATTTCTAGAGTTGGCGAGGTTTTGAACAGCCCCCTCTCTGACCCCGGTTTATTCCATCTTACTTGTATAAGATTTGCGAAGGATGTCGA
AGCATGGAAAAGGGTACTGTTAAGTGGTTCAACAATGCCAAAGGGTTTGGTTTCATCTGCCCTGAAGGCGGCGGCGAAGATATTTTCCT
CGAGAGATCCTCTAGATTTAAGAAGGAGATATACATATG
>gltB
GAGGCAATTCCGACGTCTAAGAAACCATTATTATCATGACATTAACCTATAAAAATAGGCGTATCACGAGGCCCTTTCGTCTTCACCTC
GAGAGCTAAAACCGCCGTGCAGCGTCAGCTTATGCACCTTTTGCCCATAACGACGGGTGAGATCACCACCAAACATATTGACTAATTTC
TGTAACTGCATAATCTGATAGACCGCGCCTTGAAAAGAGGCCAAAGCCTGCCATTTTTAGCTCAATTCGGCGATGACCTGGATCAATCG
TCCCGGCCTGCTTTTATCAACTGCATAATCAATCAAAATTACCGAAATTTCATGCATAATCACATAAATCACTTTTGCTTATCTTGTGT
CAGATTTTTTTATCTCCTGATGGATTTTAGGCAAAAACAGTAGCATGAAACGTCATTACCAATTAAGGCAGTATAAAATGCTGGTTTTG
TCGTCAGTTCAAGGCAGGATAAGGGTTAACACACCTTTATGACAGTCAGGAATTGACTGTTTCTCTAACGACTTCCCTTTTAGCCTTAA
AGATAAAATCCATTTTAATTTCAGTCATTTAATAAAGAATTTTGCGCTAAAGCACATTTCTGTACCAATAAGCTTGCCATTTGACCTGT
ATCAGCTTTCCCGATAAGTTGGAAATCCGCTGGAAGCTTTCTGGATGAGCAGCCTGCTCATCATATTTATGCAGTAATTGAGATCCCCT
CTTCACCGTATTAACCGATGCGAAAAGGACAACAAGGGGGCGAATGCGAGGCGCGCGTATGACACGCAAACCCCGTCGCCACGCTCTTT
CTGTGCCCGTGCGCAGCGGTTCGGAAGTGGGGTTCCCGCAGAGCCTGGGGGAGGTTCACGATAGGATCCTCTAGATTTAAGAAGGAGAT
ATACATATG
>cspB

```
GAGGCAATTCCGACGTCTAAGAAACCATTATTATCATGACATTAACCTATAAAAATAGGCGTATCACGAGGCCCTTTCGTCTTCACCTC
GATGGATCCTGATAAGACCCTTGCCGCTTTTGCCGTCAAAGGTTTTGACAATTCCTGTCATTTTACGGGACAAAAAAATTCCTTAATAC
TGATAACTTGGCGCACTATACACACGTTCCTGAAGAAAGCTATAGTTTTTTGATGGGGTTGAAGATGGCTGGATGTCTAAAATAAACAT
TGCTTCATATGTTCAACTATGCGTTAATGATTGCGTCGGTTTGAAGAACAGACGATATACGAAGTAGTTTACTAAAGCAGTTCTCATTT
CAGGTGTTATTCACTTATTCCTTCTTTGAGTCTCTCCAATTAAGTACGAAGTCGTTTCTGTTATGCAAACCATTTATGCCGAAAGGCTC
AAGTTAAGGAATGTAGAATGTCAAATAAAATGACTGGTTTAGTAAAATGGTTTAACGCTGATAAAGGTTTCGGCTTTATTTCTCCTGTT
GCTCGAGAGATCCTCTAGATTTAAGAAGGAGATATACATATG
>cspA
GAGGCAATTCCGACGTCTAAGAAACCATTATTATCATGACATTAACCTATAAAAATAGGCGTATCACGAGGCCCTTTCGTCTTCACCTC
GAGAATCCAGACGCGTGAAGCCTTCAAGTGCCGAACTAAAATTGATGCGTTTGATTCAAGCCAACCCGGCATTAAGTAAGCAGTTGATG
GAATAGACTTTTATCCACTTTATTGCTGTTTACGGTCCTGATGACAGGACCGTTTTCCAACCGATTAATCATAAATATGAAAAATAATT
GTTGCATCACCCGCCAATGCGTGGCTTAATGCACATCAACGGTTTGACGTACAGACCATTAAAGCAGTGTAGTAAGGCAAGTCCCTTCA
AGAGTTATCGTTGATACCCCTCGTAGTGCACATTCCTTTAACGCTTCAAAATCTGTAAAGCACGCCATATCGCCGAAAGGCACACTTAA
TTATTAAAGGTAATACACTATGTCCGGTAAAATGACTGGTATCGTAAAATGGTTCAACGCTGACAAAGGCTTCGGCTTCATCACTCCTG
ACGATGGCTCGGATCCTCTAGATTTAAGAAGGAGATATACATATG
>yhgN
GAGGCAATTCCGACGTCTAAGAAACCATTATTATCATGACATTAACCTATAAAAATAGGCGTATCACGAGGCCCTTTCGTCTTCACCTC
GAGGTCGCGCTCTTCAACCATGCGTTGCATGAGAACGGAGCCGACCATACCGCGCCAGCCGATAAAACCAACATTTTTCATAAGCGTTT
TTTTCCTGCAAAGATGTGTGCTGTATAAATGTGCCGGTCTCCTCTTGGCACATCTTTCACCATACAAAAAGCAGCCAAAGTCGCAAGTG
AAATTAATCAATGATAGCGAAGCCATCAGTAATGCGACTTATCCTGCTTTGTTAGCACGCAGAAAGTCCGCGGCAATTATCAGGGAATT
TGAGTTATGAATGAAATCATTTCTGCAGCAGTTTTATTGATCCTGATTATGGATCCGCTCGGAAGGATCCTCTAGATTTAAGAAGGAGA
TATACATATG
>yfeC
GAGGCAATTCCGACGTCTAAGAAACCATTATTATCATGACATTAACCTATAAAAATAGGCGTATCACGAGGCCCTTTCGTCTTCACCTC
GAGTTTGGTGGAGCTAAGCGGGATCGAACCGCTGACCTCTTGCATGCCATGCAAGCGCTCTCCCAGCTGAGCTATAGCCCCACGATGCG
TTTACGTACCAAGTTTGCTGGGTGCAAAATTTGGTGGAGCTAAGCGGGATCGAACCGCTGACCTCTTGCATGCCATGCAAGCGCTCTCC
CAGCTGAGCTATAGCCCCGTCACGTAAAGCTTGTCGAGTTGACGGGCGGCATCATATGAATTCCGCCCGAATGTGTCAACGGCAAATTG
CAACGTGTAGTTTCAATCGCTGAAAAATCAGGCAAATGAACAATTTTGGAAAACCGCTCGCATTCAGTAGTTATTCATGTCACGGTTTC
CTGTAAAGTGGTGTTATAAAATGAACTACTAATAGACCCACATACATTCAGGGAATTGTTATGTTCAAGGAGAGGATGACACCAGATGA
ACTTGCCAGACTGACCGGCTATAGCCGCCAGACCATTAATAAATGGGTACGCAAGGAAGGCTGGACAAGGATCCTCTAGATTTAAGAAG
GAGATATACATATG
>maeB
GAGGCAATTCCGACGTCTAAGAAACCATTATTATCATGACATTAACCTATAAAAATAGGCGTATCACGAGGCCCTTTCGTCTTCACCTC
GATGGATCCAATATCGCCGCTGTCTGCCACGACAGTGGTGAACTGTTTGATGCCGTCTAACTCGTTCATAGGAAATACTCCTTGAAAAG
TAAAGTGTTAGATGAGTGCGTTAATTCACACTTCTGAGAAATTTCGCTAAACGCATCAAAAAAGCATAGCAGACAGGCATGGTATTGCT
GGATTAAGCAGGTAACATCAGTGTTATAGGATTATTACCAAAACATTATATTAATGGTGAGAGTTTGGACTTGCTCAAAGTCTGTAGAC
TCCGGCAGGGTAATAATGTGCGCCACGTTGTGGGCAGGGGCTGTTGCCCACACACTTTATTTGTGAACGTTACGTGAAAGGAACAACCA
AATGGATGACCAGTTAAAACAAAGTGCACTTGATTTCCATGAATTTCCAGTTCCAGGGAAAATCCAGGTTTCTCCAACCAAGCCTCTGG
CAACACAGCCTCGAGAGATCCTCTAGATTTAAGAAGGAGATATACATATG
>yhgF
GAGGCAATTCCGACGTCTAAGAAACCATTATTATCATGACATTAACCTATAAAAATAGGCGTATCACGAGGCCCTTTCGTCTTCACCTC
GAGCTGGCGGTGGTGAATACCCCTGCCGGGGAAGCGAGCTGGTATGTTAATGCTATCGAGTACGTGAAACCGTAAGGAAGAGTCTTAAC
CTCCTGCCGATGGCTGGCATTTTTGCCAGCCAGTCCGTATAACTATCCCCTGATTTTTGATCCGAAAAGATGAACTCAAACCATGATGA
ATGATTCGTTCTGCCGCATTATTGCGGGTGAAATTCAGGCGCGCCCGGAACAGGTTGACGCTGCCGTTCGCCTGCTTGACGAAGGGAAT
ACCGTGCCGTTTATCGCACGTTATCGTAAGGAAATCACCGGCGGTCTGGATGACACGCAGCTGCGTAATCTGGAAACGCGTCTGAGCTA
TCTGCGCGAGCTGGATCCTCTAGATTTAAGAAGGAGATATACATATG
>yfeK
GAGGCAATTCCGACGTCTAAGAAACCATTATTATCATGACATTAACCTATAAAAATAGGCGTATCACGAGGCCCTTTCGTCTTCACCTC
GAGCCGTAAACCACCTGCGACTGCACGGCGACGATATCCGCCTGCAGTGCCCTACTCTTATCGTTAAACAACAACAAACTACTCATTTA
ATTTTTTCTCCTTGCCGATGATCCTCATCGTAATCCAACCGAAACTTTACCTGATTCTGGCAGTCAAATCGGCTATCACAAAACAAGGA
TAAGGTAATTCAATGAAGAAAATCATTTGTCTGGTCATTACACTACTAATGACACTCCCCGTTTACGCGAAGTTAACTGCCCATGAAGA
AGCCCGCATCAGGATCCTCTAGATTTAAGAAGGAGATATACATATG
>yhgE
GAGGCAATTCCGACGTCTAAGAAACCATTATTATCATGACATTAACCTATAAAAATAGGCGTATCACGAGGCCCTTTCGTCTTCACCTC
GATGGATCCGAGTTCTTGCGGGGTCAAACCATTGTTAACGCGCATTTCACTGCTCCTTAGCCAATATGTATTGCCTGAATAGTAAAGTC
TTTTTGGGGGTGTTAACCGCGACAAGGCTCATAGATTTACGTATCTGGAGAAATTCATATTTGACAAAAAACTCCGTGATTCCTGTCAC
GAAACGGTTGCTATTATCGCAGAAATAGCATTTTAGGTGTGGAAAATGTTCTCAAAAGGTTAAATCTTGGTTTATATGGCGGGAATATT
GTGAGTGGAATCGCATTCCTGAATGAAATTGATGCAAATTCAACCGGTAAAGTATCGATTCACCACGCTAACTTTTTTTGATTATTTTC
TGAAAAAAGCAAAATACTACTGCCCGGATGCGATAATTAACCCACCTTTTGATTCTGGATAACCAACATGGATAACGTTGAACTTTCAC
CCGCAACACGCTGGGGGATGATTGCTACCGGATTACTTCAGGGTCTGGTTTGCTACCTGCTGATCGCCTGGGTCGAGAGATCCTCTAGA
TTTAAGAAGGAGATATACATATG
>yfeN
```

GAGGCAATTCCGACGTCTAAGAAACCATTATTATCATGACATTAACCTATAAAAATAGGCGTATCACGAGGCCCTTTCGTCTTCACCTC
GAGCGTGGCGTGAAATCAGGTTTATAAGTCTTGATAATATCTATGCAAAACAGCGGGTTATGAGAAAATTGAACCTGAGACATACATAT
CCTTTTTCTGTAGGGTGGAATCTAACGCCTTTTTATAGCGAGGTTGTCCCGGCTCTTACCAATACACTTTTCTCACGCATTCAATAGCG
ATTCGATATCGATGGATTTATATCTTTTCTCTATTAATCCACGGTAAAAACTGTATTGGCGATATTAATTCCTTTGCATTAACAATATG
TCGTTTTTAGCAAACCACTTTATTAAGCAATCTACTAATAACGATCGGCATAGGGAATTATGCTATGAAAAAACATCTTTTAACTCTGA
CACTTTCCTCTATATTAGCGATACCTGTTGTATCTCATGCAGAATTTAAAGGCGGTTTTGCAGATATCGGCGTGGGATCCCGGGATCCT
CTAGATTTAAGAAGGAGATATACATATG
>yhgA
GAGGCAATTCCGACGTCTAAGAAACCATTATTATCATGACATTAACCTATAAAAATAGGCGTATCACGAGGCCCTTTCGTCTTCACCTC
GAGGAAGAACCTGACGGCTGCCTCTCTGGCAGTTGTAAAAGCTGCCCGGAAGGAAAAGCCTGTCTGCGCGAGTGGTGGGCGCTGCGTTA
ACCTTACTCCATCGCCTGTTTTGGAAAGCCCGGTATGCGTCTGCATCCGGGCTTTTTTGCGTGCGGCTTTCCATAAAAATGCAACTCTT
GCAGCACGGCGTAAGTTCCTTTGAAAGCATCTCGCAGGGATGAAAACTCGCTAATACACAGGTGTGGAGTGGCGCGTAGAGTCGCGGCA
TTCAAACAACAGGTGAAGGAACGCCATGAGCAAAAAGCAGAGTTCCACCCCACACGATGCGCTGTTCAAACTCTTTTTACGCCAACCGG
ACACGGCTCGTGATTTGGATCCTCTAGATTTAAGAAGGAGATATACATATG
>yfeY
GAGGCAATTCCGACGTCTAAGAAACCATTATTATCATGACATTAACCTATAAAAATAGGCGTATCACGAGGCCCTTTCGTCTTCACCTC
GATGGATCCGGCTGCTGACCAATCGACGTTTACGCGCTTGTTTTTATGAAGTAAAAGAATAACGGCACTTTTTGGTGAATTTGCACTCC
AAGCAACGTTATTGAATAACCAAAGGAAGTGACATGAAATCGCTGCGTTTAATGTTATGCGCTATGCCATTGATGCTGACCGGCTGTTC
CACGATGTCGTCAGTTAACTGGTCTGCCGCTAACCCGTGGAACTGCTCGAGAGATCCTCTAGATTTAAGAAGGAGATATACATATG
>b0671
GAGGCAATTCCGACGTCTAAGAAACCATTATTATCATGACATTAACCTATAAAAATAGGCGTATCACGAGGCCCTTTCGTCTTCACCTC
GATGGATCCGCGAGTTCAAGTCTCGCCTCCCGCACCATTCACCAGAAAGCGTTGTACGGATGGGGTATCGCCAAGCGGTAAGGCACCGG
TTTTTGATACCGGCATTCCCTGGTTCGAATCCAGGTACCCCAGCCATCTTCTTCGAGTAAGCGGTTCACCGCCTCGAGAGATCCTCTAG
ATTTAAGAAGGAGATATACATATG
>b0298
GAGGCAATTCCGACGTCTAAGAAACCATTATTATCATGACATTAACCTATAAAAATAGGCGTATCACGAGGCCCTTTCGTCTTCACCTC
GAGCGATGAAGTCGGGCTGTTTGGTAAAGATAAGCGCCAGAAAGACCCGCATGCTATTTCTGCCGAGGTGACCTATACGCCAGTGCCTC
TTACCCAGCAATAGTGGACACGCGGCTAAGTGAGTAAACTCTCAGTCAGAGGTGACTCACATGACAAAAACAGTATCAACCAGTAAAAA
ACCCCGTAAACAGCATTCGCCTGAATTTCGCAGTGAAGCCCTGAAGCTTGCCTCGAGCGGAGATCCTCTAGATTTAAGAAGGAGATATA
CATATG
>yjiX
GAGGCAATTCCGACGTCTAAGAAACCATTATTATCATGACATTAACCTATAAAAATAGGCGTATCACGAGGCCCTTTCGTCTTCACCTC
GATGGATCCGGTGCGTAACAGCGACAAACGTACTGACAAAGAAACACCGTACGTTCCAATCCCGGAAGGCGGCGTGAAGATCTCTTCGC
ACCACTAACCGTGTTTAGCCCCGCTTCGGCGGGGCTTTGTTCTTATCAGAGTGAACTATGTTTGGTAACTTAGGACAGGCAAAAAAATA
TCTCGGCCAGGCGGCGAAGATGTTGATTGGCATTCCAGACTCGAGAGATCCTCTAGATTTAAGAAGGAGATATACATATG
>rssA
GAGGCAATTCCGACGTCTAAGAAACCATTATTATCATGACATTAACCTATAAAAATAGGCGTATCACGAGGCCCTTTCGTCTTCACCTC
GAGGGTGCAACCTTTTCACCAGACACATAAGGGTGGCAACATAGGCTATACTCGACAGCACTACCACAGGGACAAAGCTGAGACACAAA
TAATCTCCCTGGAAACAATAACGGCGTATTAACCGCCTGAGTAGCACTATGTTAACCGAGCAGTAGCGATGTGGCTACGATTGCATTCC
AGGGGAATCTTGCGGGAATAATGAGAAAGATAAAAATAGGGCTGGCGCTGGATCCTCTAGATTTAAGAAGGAGATATACATATG
>rssB
GAGGCAATTCCGACGTCTAAGAAACCATTATTATCATGACATTAACCTATAAAAATAGGCGTATCACGAGGCCCTTTCGTCTTCACCTC
GAGCGCTTGATTTCCATCGCGCGCACGCTGCCATTGCGGCCGGACAGCTGGCAGTGGAAAGGAAAATGGACGAACTTTTGCCGTTGGTA
CGCACCAACATTTGACCAGAATTTTTATCTACACTTAAGTTAATTCTGACAGGCGCAGGTGGCAATAGCATGCCACTATTGAGTAAAGC
CAGTCAGGGGAGAGAACATGACGCAGCCATTGGTCGGAAAACAGATTCTCATTGTTGAAGATGAGCAGGTATTTCGCTCGCTTCTGGAT
TCATGGTTTTCCTCATTGGGAGCGGGATCCTCTAGATTTAAGAAGGAGATATACATATG
>ddlA
GAGGCAATTCCGACGTCTAAGAAACCATTATTATCATGACATTAACCTATAAAAATAGGCGTATCACGAGGCCCTTTCGTCTTCACCTC
GATGGATCCGCGCACACAGTTCTTTCGACTCTTCTTCTTTTTGGGCAAGCTTAAATAACAACTCAGCAATGAGATTTTTCATGTCTGTA
TTTCCTTATCCAAAGTATGGAGAAGTTAACATTATGTTTCACAACTTCATAGTGTCTGGAAGTATTATTTTTATGACAAAGGAAATACT
TTGCGCAAATATTTTTTGTTTGATTACCAGCGAAACGTTTCTCTTATTTAAATAATTACCCACACAAAATATAGGCTTTTAATTATCAA
CCTTTATTATAAATAAAAGTTATGAAATATATTCTCTTCGTTAACTTATAATTAAGAGAAAAAAACTCACCGCATGGCCATTTAAGCCA
GGAATAAATTAGCTTCCCGTTCATCATTTTTGTGATCTTGCGCACATCTTTGAGATAACGCACTTTCTTCACCAGTACGTTGCACGCTT
TGTCAGCAGAAGATTGCAACAATCTTTCAGAAGGCCGCGAATCCCCCGTTTTGTCGTGGTAATGTATGCACTTTGTGTTAGACGGGATA
GGTTTTTAAGATGGAAAAACTGCGGGTAGGAATCGTTTTTGGTGGTAAATCAGCGGAACATGAAGTGTCTCTGCAATCGGCAACTCGAG
AGATCCTCTAGATTTAAGAAGGAGATATACATATG
>gatY
GAGGCAATTCCGACGTCTAAGAAACCATTATTATCATGACATTAACCTATAAAAATAGGCGTATCACGAGGCCCTTTCGTCTTCACCTC
GATGGATCCGAAATCGATGGCTGACGGCGTGAAACTGATTAACGCCGTGCAGGACGTTTATCTCGATAGCAAAATTACTATCGCCTGAT
ACGCCTTATACGAACAACACCACCCCGTAGGACGGATAAGGCGTTTACGCTGCATCCGGCAACCTATGCCTGATGCGACGCTGAAGCGT
CTTATCATGCCTACATAGCACTGCCACGTATGTTTACACCGCATCCGGCATAAAAACACGCGCACTTTGCTACGGCTTCCCTATCGGGA
GGCCGTTTTTTTGCCTTTCACTCCTCGAATAATTTTCATATTGTCGTTTTTGTGATCGTTATCTCGATATTTAAAAACAAATAATTTCA

Data Set S4 - MG1655 library reference file.fasta

```
TTATATTTTGAAATCGAAAACAAACGACAGGATATGAAAATGTACGTGGTATCGACAAAGCAGATGCTGAACAACGCACAGCGCGGCGG
TTATGCGGTTCCGGCATCTCGAGAGATCCTCTAGATTTAAGAAGGAGATATACATATG
>yjiW
GAGGCAATTCCGACGTCTAAGAAACCATTATTATCATGACATTAACCTATAAAAATAGGCGTATCACGAGGCCCTTTCGTCTTCACCTC
GATGGATCCGCAGCTAGCGGGGGTAAAAAAGCCTCACGTAAAAAATCCTGAACATTATTTTCTGGCGCACCTTTCCGGTGCGCTTTTTA
TTATTTCACGCCAATCATAACCCACATAAATATATTTAAATCATTCCAGAAATTGCCCATTTTATTCTATTTTTAGCTGGACTTTCCCC
ATATTTACTGATGATATATACAGGTATTTAGCGCGGTGCGGATGTGCGCCAACACACCCGCATCGCTAATCACAATCACTATTCCTGGA
GAATAGCAGTTATGACTGACACGCATTCTATTGCACAACCGTTCGAAGCAGCTCGAGAGATCCTCTAGATTTAAGAAGGAGATATACAT
ATG
>ytjB
GAGGCAATTCCGACGTCTAAGAAACCATTATTATCATGACATTAACCTATAAAAATAGGCGTATCACGAGGCCCTTTCGTCTTCACCTC
GATGGATCCCCCGGCCATAAAGAGACATCTTCAGGCAGGTCGCACCAGGTAATGTTAGGCATTAAGGCTCCTGTAAAATCGTTCGAAGC
AGGGAAAATAACGCATGAGGCTACCTTGTATCCATTGCTTCTGGCAACATTAAGTCTCAAATTTTCAAAGGGTGGAAGATGGCTCGCAC
AAAACTGAAATTCCGGCTGCATCGGGCAGTGATTGTCCTGTTCTGTCTTGCCTTGTTAGTGGCGCTGATGCAGCTCGAGAGATCCTCTA
GATTTAAGAAGGAGATATACATATG
>ygiF
GAGGCAATTCCGACGTCTAAGAAACCATTATTATCATGACATTAACCTATAAAAATAGGCGTATCACGAGGCCCTTTCGTCTTCACCTC
GATGGATCCCGGAAACATAGCGCGTTTCTTCAGCGTGTGAGACGGCAGTCGCGCTAAGTGCGAGTAAAGTTAATCCGATCAGGCGTAAT
TTTGGCATCAGGCTGTCGTTATTGTCATGGAAAGTGGAACGATAGTAGTGGCATCAGTGTCGCTACGCAAAGCATTCGACATCAATCGG
AATCCTCTGTGTCAGGTCTTGCCACTGCTTCAAACTTTTACGCCCGTCAAATTGCCCGTTGCATGGCAATTTGGCGCAAAATACTATCT
ACTGACAAAAAAGATCGCATTAAGTTCGCCGTTCATCGTCACTTATGTGACATAACCGATAAGTAAAGGCCATGGCTCAGGAAATCGAA
TTAAAGTTTATTGTTAATCACAGTGCCGTTGAGGCGTTGCGTGACCATCTCAATACGCTGGGCGCTCGAGAGATCCTCTAGATTTAAGA
AGGAGATATACATATG
>ygiD
GAGGCAATTCCGACGTCTAAGAAACCATTATTATCATGACATTAACCTATAAAAATAGGCGTATCACGAGGCCCTTTCGTCTTCACCTC
GATGGATCCCAATAAACGTGGCTGCCCCCGCCAGTATGGTCAGAATGAGAGGTACTGACATCGACAACTCCTTATAAATCTTCACCTTC
AGGCAGATCGTCAAAATAACTGATGATCATCATCATTGTTATCCGGGTTCTTACTCCGCCGAACAAGCGTATTGTGAGGATATCAAAAT
GACACCTTTAGTTAAGGATATCATCATGTCTTCAACACGTATGCCAGCATTGTTTTTAGGTCACGGTAGTCCGATGAACGTGCTGGAAG
CTCGAGAGATCCTCTAGATTTAAGAAGGAGATATACATATG
>ygiH
GAGGCAATTCCGACGTCTAAGAAACCATTATTATCATGACATTAACCTATAAAAATAGGCGTATCACGAGGCCCTTTCGTCTTCACCTC
GAGTTCGATGGTCTGTTCCCAGTCGTAAACACCAATAGTGGTGATTACCGAAAGTTGCTCTATAAATACAATATCCATCACGTCCTGCC
TGCTTTTTGGCTAACCGGATACCACTTCCGGCGAAATGTGCGTATTATCCACAGATTCATCGTTGAACACGAATTTTCAAAACGGAACA
GCTTATGAGTGCAATCGCGCCTGGAATGATCCTCATCGCGTACCTCTGCGGCTCCATTTCCAGTGCCATTCTGGTTTGCCGGATCCTCT
AGATTTAAGAAGGAGATATACATATG
>eno
GAGGCAATTCCGACGTCTAAGAAACCATTATTATCATGACATTAACCTATAAAAATAGGCGTATCACGAGGCCCTTTCGTCTTCACCTC
GATGGATCCGCCAGCGAGTTCCAGAAACGTCAGGCGAAGTAAGTAAAAAAGTTAGAGCGGCAACGCGTACCCTGGGTACGCGTTGTTTG
TCTGGAGTTTCAGTTTAACTAGTGACTTGAGGAAAACCTAATGTCCAAAATCGTAAAAATCATCGGTCGTGAAATCATCGACTCCCGTG
GTAACCCGACTGTTGAAGCCGAAGTACATCTGGAGGGTGGTTTCGTCGGTCTCGAGAGATCCTCTAGATTTAAGAAGGAGATATACATA
TG
>ygiS
GAGGCAATTCCGACGTCTAAGAAACCATTATTATCATGACATTAACCTATAAAAATAGGCGTATCACGAGGCCCTTTCGTCTTCACCTC
GATGGATCCGCAATGCCCAACCTCATCCTTCCACAATCAAACTTTTACGTGTTCTGGATAAGCATCCAGAACTATTGAATGAAATCCGT
TAATTAATGGAAAAGCGGGGGCAACTCGCTTTTTTCTTTACCGGAGTGTGAGCCAAATCACTCGCGATAAGTTTGAGCGAACAATGCCA
CAGGCGTATTGTTGTTGGGCTTATAATTTCTCCGGTAAGGAATACCATGTATACGCGAAATTTATTATGGCTGGTCAGCCTGGTAAGTG
CGGCTCCTCTCTACGCTGCTGACGTTCCCCTCGAGAGATCCTCTAGATTTAAGAAGGAGATATACATATG
>ygiR
GAGGCAATTCCGACGTCTAAGAAACCATTATTATCATGACATTAACCTATAAAAATAGGCGTATCACGAGGCCCTTTCGTCTTCACCTC
GATGGATCCGCTCGATTGGGCAACTGTTGGTCAATCCGGTACCGTAATCTTCATACTCGCCCGAGGAGGGCGAGTATGGAATACCTTTC
GATTTCTCTTTCGTTCCGGCTTATAATCCCCGGCCTTTGATTATTTTTTTATCATTTCTGGAAGCAAAATGAGCTCTATCTCCCTGATC
CAACCGGATCGCGACCTGTTCTCCTGGCCGCAGTACTGGGCCGCCTGTTTTGGACCGGCACCGTTTTTGCCGATGTCTGTCGAGAGATC
CTCTAGATTTAAGAAGGAGATATACATATG
>ygiP
GAGGCAATTCCGACGTCTAAGAAACCATTATTATCATGACATTAACCTATAAAAATAGGCGTATCACGAGGCCCTTTCGTCTTCACCTC
GATGGATCCCGGGTAGAAATCATGGCGGTAAAGTTAGCGACAATCTCTGTCAACTTATTCACTGCCTGTTGCTTATTACTTTCGCTCAT
CATGGCAACTCCAGTTTTTGTATTTGTCTGGAAAAATTCACCGCCACAAACTACCACTAAAAATGTGGTTTTATATTTTTTGCCACCGT
GAATATGTAATTAACAGTTCGTGAATCAACGCGAAAATAAGAATGAATATTGCGCCAGAGCAGGGAATGGCAATGTTTTGCGAAATACA
CTGCGCTTATCACAACTGCGGATAATTAACGATGCTGAATAGCTGGCCTTTAGCCAAAGACTTGCAGGTACTGGTAGAAATCGTTCACA
GCGGGAGTTTTAGCGCTCGAGAGATCCTCTAGATTTAAGAAGGAGATATACATATG
>ygiW
GAGGCAATTCCGACGTCTAAGAAACCATTATTATCATGACATTAACCTATAAAAATAGGCGTATCACGAGGCCCTTTCGTCTTCACCTC
GATGGATCCTAAGGCCCGTTTTGATGCCGTCGCCAATCAGCATGTCATCTTCTATCAGTAAAATTCGCATTTTTTCATCCCTGCGATAA
```

```
CCGTATTTGCTGCCAGTAAACCGTGTTGTAACGCGCTCTGTACAGCACTAAATTTTATTTTTTTCCCTTAAGAAGTTGTTAAGGACTAT
CTTGTTAAATGCTCGGAAACAGACATTAAAGGGAGTAATAAACATGAAAAAATTCGCAGCAGTAATCGCAGTAATGGCCCTGTGCAGCG
CACCGGTGATGGCAGCAGAGCAGGGCGGTTTTTCTCTCGAGAGATCCTCTAGATTTAAGAAGGAGATATACATATG
>ygiU
GAGGCAATTCCGACGTCTAAGAAACCATTATTATCATGACATTAACCTATAAAAATAGGCGTATCACGAGGCCCTTTCGTCTTCACCTC
GATGGATCCTGACAGAGCAGCGACTGGAAACGGATGTTTATGTGCCGTTGGCGTGATGCCTGACTCCAGCTTCCCTTATAAATTCCTGC
TTCAAACTGGCGCTACTAATGCGCCACTTTCTTATATGGTCAATCACTTTCCTTAAAGAATACGTTTTGTGTGGTCACTATCTCCGTAC
ATCTAACTAACCTTTTAGGTGCTTTTTCATTGCTGTAATTAACCTTTTAGGTTATAACTAAAGTAACAGGGAGGCGGGGGTTATGGAAA
AACGCACACCACATACACGTTTGAGTCAGGTTAAAAAACTTGTCAATGCCGGGCAAGTTCGTACAACACTCGAGCGGGCAAGTGAGATC
CTCTAGATTTAAGAAGGAGATATACATATG
>ygiZ
GAGGCAATTCCGACGTCTAAGAAACCATTATTATCATGACATTAACCTATAAAAATAGGCGTATCACGAGGCCCTTTCGTCTTCACCTC
GATGGATCCTGACTCACCGATGCAGGATAAATATTTTCTGAAAATGACAATACTGATAGCCATTCCTCATAATTCTTACTCGCACCGGA
ATAATTTACACGTTTGGTTCGGTTTTTTTGGGGATTGAAAATATGGCTTTGAATAAGTAAAAGATGAAAGTATCAACGTGTCATGTTGA
TAATTATAAATGGATTTTTATGAGCGTACTGATATATGTTAAAGCAGAAAATTAAAACAATATTTGAAGCATTGCTATATATAATGCTT
ACTTATTGGTTAATAGATTCTTTTTTTGCTTTCAATAAGTATGACTGGATGCTTGAGTCGGGAGGAAATATTTGCAGTATACCTTCCGT
TTCAGGCGAAGATCGTATATTGCAAGCAATGATTGCAGCATTTTTTCTTCTTACGCCACTTATTATTCTTATCTTAAGAAAACTCTTTA
TGCGGGAAATGTTCGAGTTTTGGGTATATGTATTCTCGCTCGGTATTTGTCTGGCTCGAGAGATCCTCTAGATTTAAGAAGGAGATATA
CATATG
>smtA
GAGGCAATTCCGACGTCTAAGAAACCATTATTATCATGACATTAACCTATAAAAATAGGCGTATCACGAGGCCCTTTCGTCTTCACGTC
GACGGCTAAACCACAGCAGCGCCAGGCCAGCGCCAATAATCAACAACATCAGCGGAAGTGGCAGCAACATATTGCCAATCACTTTTTTC
AGTGTAAAAAGCATCCTTTCTGGTTCCTTTTTTAACCATATAGCAAGGGATCGCCAGCGATATTACACCAGACGGGTTCATTCTCGCCG
TGGCTGTGACAAAATAGCGTCTTTATCAGAAAACGTCTTAGCCAACCGGTGGAGAAAGCAATGCAGGATCGCAATTTTGATGATATTGC
GGAAAAGTTTTCCCGTAACATTTACGGCACCACCAAAGGGCAGCTTCGACAGGCTAGGATCCTCTAGATTTAAGAAGGAGATATACATA
TG
>b0582
GAGGCAATTCCGACGTCTAAGAAACCATTATTATCATGACATTAACCTATAAAAATAGGCGTATCACGAGGCCCTTTCGTCTTCACCTC
GAGATAGCCCGGCGGATTAACCACCTGCATTTCCAGTTCAATACCGAGGGTAAAAGGTTCAGAAACATGAAAATCGGGTAATGGCATAG
GTTTCTCTTATGTTGGCGTTTTCTATTCAGTATAGAAGTCGGAGCGGCTGGGCGAGATGCGGAAGTTCTGGAATGTTTCTTTTTTTTGG
TGATGGTGAACTGATGGTGCCTGAAGCAATTTGGCTACTTTTGCAATGTGACAAGTTATGGCAGTGCTGACTGGTGGCGAAGAATTTTG
ACGATTGAGTCATGCAGAAAAAAACGGGTTCAGCTTCAGTTGATTAAATGAGGAGTGAGAAGTCCGAAACAGGACTCACTGTATAAATA
AACAGCTATTTTGTTGAGGAAGGGTAAGATAACGGCGGGTGCCTGAAGCTTTCCGGTTTCAGGTTTACTCTGAGGTCTGGAAAGATGAA
GCCCCAGGAGATATTTCTATCAACCCTGGGGCTGCCACTCCAAACCCGAACAATTTGGATGGTAGTCCCTTCTTCGCATGGAGGCAATA
TAAACATGCTGACGAAATATGCCCTTGCGGCAGTCATAGTGCTGTGTTTAACGGTGCTGGGATTTACGCTTCTGGTCGGAGACTCGCTG
TGTGAGTTTACGGTGAAGGAACGTAATATTGAGTTTAAGGCTGTTCTCGCTTACGAACCGAAGAAGTAGCCGTTGTGCGGGGAGTAATC
CCATAAGCGCTAACTTAAGGGTTGTGGTATTACGCCTGATATGATTTAACGTGCCGATGAATTACTCTCACGATAACTGGTCAGCAATT
CTGGCCCATATTGGTAAGCCCGAAGAACTGGATACTTCGGCACGTAATGCCGGATCCTCTAGATTTAAGAAGGAGATATACATATG
>amiA
GAGGCAATTCCGACGTCTAAGAAACCATTATTATCATGACATTAACCTATAAAAATAGGCGTATCACGAGGCCCTTTCGTCTTCACCTC
GAGTTGCGGGTTAAAAGGCTGATTATGGCGTGAACGGTCGAATTAGCCAATATCTGACGAAAATCGGTTGAAAAAGTGGCATAATGGGG
AGTTGTCAACTATTGAAATGAAAAGTAAAACAATTCTCAACAGCAAACCGTCGTAACGGATTACGCGATACGATATAACATCTGGAACT
TTATTATTACAACTCAGGCCGTATGAGCACTTTTAAACCACTAAAAACACTCACTTCGCGCCGCCAGGTGCTGAAAGCCGGTTTGGCTG
CCCTGACGTTGTCAGGAATGTCGCAAGCCATCGGATCCTCTAGATTTAAGAAGGAGATATACATATG
>yhiD
GAGGCAATTCCGACGTCTAAGAAACCATTATTATCATGACATTAACCTATAAAAATAGGCGTATCACGAGGCCCTTTCGTCTTCACCTC
GATGGATCCAAGAAAAACCCGCAGAAAAATTTGTATACCTTCAAAAATCAAGCATCTAATGACTTGCCGAATTAATGAGGTGCAAGTAA
AAAGGAGTAGCAAGTTGAGCCATCTTGCTGCTCCTTTTTGCATTTTTATATGACAGCAGAATTTATTATACGTCTTATACTCGCGGCAA
TTGCCTGTGGCGCTATTGGCATGGAAAGGCAAATGCTCGAGAGATCCTCTAGATTTAAGAAGGAGATATACATATG
>yjiA
GAGGCAATTCCGACGTCTAAGAAACCATTATTATCATGACATTAACCTATAAAAATAGGCGTATCACGAGGCCCTTTCGTCTTCACCTC
GATGGATCCATCCCGACAAGCCGTACATGAGCTATGAAGAATTCTTCCGCGAACGCCAGAACGCGCGCTACGGCGGCGATGGTAAAGGC
GGTATGCGCTGTTGTTAATGGAGAGAGCATGAACCCGATTGCAGTTACCCTACTCACCGGTTTTTTAGGCGCAGGAAAAACCACCCTGC
TGCGCCATATTCTTAACGAACAGCACGGCTACAAGATTGCCGTGATTGAAAACGAATTCGGCGAAGTCTCTGTTGATGATCAACTGATT
GGCGACCGCGCCACGCAGATCAAAACGCTGACCAACGGCTGCATCTGTCTCGAGAGATCCTCTAGATTTAAGAAGGAGATATACATATG
>hslJ
GAGGCAATTCCGACGTCTAAGAAACCATTATTATCATGACATTAACCTATAAAAATAGGCGTATCACGAGGCCCTTTCGTCTTCACCTC
GATGGATCCAAAAAGGCGAAACCTGCCCGAACGAACTGGTTTAATCTTGCCGCTCCCCTGCATTCCAGGGGAGCTGATTCAGATAATCC
CCAATGACCTTTCATCCTCTATTCTTAAAATAGTCCTGAGTCAGAAACTGTAATTGAGAACCACAATGAAGAAAGTAGCCGCGTTTGTT
GCGCTAAGCCTGCTGATGGCGGGATGTGTAAGTAATGACAAAATTGCTGTTACGCCAGAACAGCTACAGCCTCGAGAGATCCTCTAGAT
TTAAGAAGGAGATATACATATG
>yhiI
```

GAGGCAATTCCGACGTCTAAGAAACCATTATTATCATGACATTAACCTATAAAAATAGGCGTATCACGAGGCCCTTTCGTCTTCACCTC
GATGGATCCCCTTTGATTAGAAGGCGCACAATGTTACAGCTCTGAACCATAGACAACATGAATATTAATGTTACTATCTGGTTGAGACT
GTTGATAAAACGTAAAAAGGAATGCTCTGTTCCTGCAACTGAACAGGCGTGAAACGCTATCCAACAGGATGGATACCGTTTTGTCAGCC
AGTTTTATAGATCACACTTATGAGTATGCGTGTTTCATTTACGGCCAGGAATGCTGATGATGACAACATTACTACCCGTTTTCACTAAG
CCTTCCCCGTTGGCGCTCAATGCTCTGCGCGCTGGTCGTATTTGCCGTTTCCTTCTTATCCCGGATGGAAGAATCCGCTGAACGTCGTT
CGGCGGCAGTTGTAACACGCGTCGTTTGCTTACAGGGATTATGGCGGGTCGATTATGGATAAGAGTAAGCGCCATCTGGCGTGGTGGGT
TGTCGGGTTACTGGCGGTGGCGGCTATCGTGGCGTGGTGCTCGAGAGATCCTCTAGATTTAAGAAGGAGATATACATATG
>yhiJ
GAGGCAATTCCGACGTCTAAGAAACCATTATTATCATGACATTAACCTATAAAAATAGGCGTATCACGAGGCCCTTTCGTCTTCACCTC
GATGGATCCTTCCCTGAAGGAAGCACGTGATAAAAATAATAAAATAATTTATTCCTCAGGGCTTAGTTGTTTTCAACTACATTAAAACT
CCATGAATACGGTGGTATGTCTTTTTAATCCTGCCAGTATTTCATAGGTTTTCTTAATCGATTGTAGAGGCATGTTCTGATGACGCCTC
TACAATCTTTAGGGACTTTCTCTGCTATGAGATAAGAAATTTCATATTATGCTTTGAGGCTGAATGATTTGTGCAAAATATAATTATTT
TCCCATCGAACATGAGGTGTATGTCATAAGATAAAATGTTAACTATGTATTCCTCGCGGTAAATAATATGAGGTAAAATATGAAAATTG
GAACTGTGGCAGGCACTAACGACAGTACAACGACAATCGCAACAAATGATATGGTGCAAGAGCATGTTACCCTCGAGAGATCCTCTAGA
TTTAAGAAGGAGATATACATATG
>adiA
GAGGCAATTCCGACGTCTAAGAAACCATTATTATCATGACATTAACCTATAAAAATAGGCGTATCACGAGGCCCTTTCGTCTTCACCTC
GATGGATCCTACAGCACGTTCGGCAAATATGTCGGCATGTCGCCGCAACAATACCGCAAACTTAGCCAACAGCGCCGCCAGACGTTTCC
CGGCTAAAGACGATATCAGTATCAGCCAAAAAAATAGTTTAGCCGTCGATAATAAGCAGTGAAAATAATTTAAAATAATCACATAACGT
ATTGTTTATAAAACTAAATATTTCAAAAATGTTTGTTTTTCACGCGCTTTACAGCCCGAAAAGGCCGGAAGATACTTGCCCGCAACGAA
GATTCCTTCATAACCGGGTAAGCAATGATGAAAGTATTAATTGTTGAAAGCGAGTTTCTCCATCAAGACACCTGGGTCGGTAACGCCGT
TGAGCGTCTGGCAGATGCTCGAGAGATCCTCTAGATTTAAGAAGGAGATATACATATG
>yhiP
GAGGCAATTCCGACGTCTAAGAAACCATTATTATCATGACATTAACCTATAAAAATAGGCGTATCACGAGGCCCTTTCGTCTTCACCTC
GAGTGATGTCTTCCGCACGTCAGCTGATCAACACCGTTCACGTTGATATGCTGATTGTTCCGCTGCGCGACGAAGAAGAATAATCTTCC
CTCTACGACGTGTTCCTGAACGCCCGCATATGCGGGCGTTTTGCTTTTTGGCGCGCCTTGTTACCTGATCAGCGTAAACACCTTATCTG
GCCTACGGTCTGCGTACGCAATCAAAATCCCCAGCCAATACAACATTTAACACCATCATATTTTCCATCATTAGTGTGATCATCTGGTT
ATTTTCTGTTGTAATAGTGTATTAATCTATTCACCGCATCAATATTAAGAATCTCTGACAGATGTAAACTTTTTCGCGCGTTATCCCTT
ACGCGTTCATACTTTTCAGGATGGTATTGGAAGGTTAATAAATATGAATACAACAACACCCATGGGGATGCTGCAGCAACCTCGCCCAT
TTTTCATGATCTTTTTTGTCGAGTTATGGGAGCGATTCGGCTACTGGATCCTCTAGATTTAAGAAGGAGATATACATATG
>hslV
GAGGCAATTCCGACGTCTAAGAAACCATTATTATCATGACATTAACCTATAAAAATAGGCGTATCACGAGGCCCTTTCGTCTTCACCTC
GATGGATCCAACGCAGACAGCACCCTCAATCGGTTGAAGATGGCGGGTCATACAAACTGCATTCGGCTCGCCGCCGGGGGTTGAAACCC
TCAAAATCCCCCCCATCTATAATTGCATTATGCCCCGTACTTTTGTACGGGGTTTGTACTCTGTATTCGTAACCAAGGGGTCAGCTCGT
GACAACTATAGTAAGCGTACGCCGTAACGGCCATGTGGTCATCGCTGGTGATGGTCAGGCCACGTTGGGCAATACCGTCTCGAGAGATC
CTCTAGATTTAAGAAGGAGATATACATATG
>arcB
GAGGCAATTCCGACGTCTAAGAAACCATTATTATCATGACATTAACCTATAAAAATAGGCGTATCACGAGGCCCTTTCGTCTTCACCTC
GATGGATCCGGTCGAGCTGGATCGCTATCTGAATGAACATGGCGTACAGGGATCGGCGCTGGGACGTCCGTGGCTACCTCCAACGGAGT
AGGTCGTTGAGGGGAATTCCGCATTTCTCACACAATTTATAACGTAACTGTCAGAATTGGGTATTATTGGGGCAGGTTGTCGTGAAGGA
ATTCCCTAATGAAGCAAATTCGTCTGCTGGCGCAGTATTATGTTGACCTGATGATGAAGTTAGGTCTGGTGCGCTTCTCAATGTTGCCT
CGAGAGATCCTCTAGATTTAAGAAGGAGATATACATATG
>yjeN
GAGGCAATTCCGACGTCTAAGAAACCATTATTATCATGACATTAACCTATAAAAATAGGCGTATCACGAGGCCCTTTCGTCTTCACCTC
GAGGCGGCCCTGTCTTCTTCTCGCTGTTAGCGATGGCGATTTACCAGAACTATTGCAGCAGAGTGGCAAAAAATCCGCAGTGGGCGGTG
GAATAACCCGATGCCCTATCGTCTACCTGACGATAGGGCGTAAAGACAGGAAATGTTATGGATGACACCAGCCGGGACCCGGCAATAAC
CGAAGATGAAATCAGGGCGCTGCAATTTTCTGCCGGGGATGTGGATCCTCTAGATTTAAGAAGGAGATATACATATG
>yjeK
GAGGCAATTCCGACGTCTAAGAAACCATTATTATCATGACATTAACCTATAAAAATAGGCGTATCACGAGGCCCTTTCGTCTTCACCTC
GATGGATCCACTCGCTTCAACCGCGTAAGGTTCGCCGTCTAACATGATTTTAAGACCAGCACGAAAATCGTTGCTATAGTACGTTGCCA
TAAGGCCCTCTGAAATTTGTTAATTGGTAGCTAAGCCACAAAATGGCGCATATTGTAACCCTAAATACCCCATCCAGAGAAGATTGGTT
AACGCAACTTGCCGATGTTGTGACCGATGTCGACAGATCCTCTAGATTTAAGAAGGAGATATACATATG
>yjeI
GAGGCAATTCCGACGTCTAAGAAACCATTATTATCATGACATTAACCTATAAAAATAGGCGTATCACGAGGCCCTTTCGTCTTCACCTC
GAGGCATGGTTACCGACCTGCCGAAAAACGATGCAGCTGACTTAGGCGCTGCTGGCGGTATGGGCGGCATGGGTGGCATGGGCGGCATG
ATGTAATTGCCCTGCACCTCGCAGAAATAAACAAACCCCCGGGCAGAAATGTCTGGGGGTTTTTCTTTTGGTCATCTTTCTAGTATAAG
ATTCAGACACGGACGACGCGAGTGGCGTCCAGCTCATTGATTATGGGGAATAACATGCACGTAAAATACTTAGCAGGGATTGTCGGTGC
CGCGCTACTGATGGCGGGTTGTAGCGGATCCTCTAGATTTAAGAAGGAGATATACATATG
>fadB
GAGGCAATTCCGACGTCTAAGAAACCATTATTATCATGACATTAACCTATAAAAATAGGCGTATCACGAGGCCCTTTCGTCTTCACCTC
GATGGATCCCATCGCGAGTCCGTTCTTGTAAGGTAGCTATATGATTTTTATAGAGCGAGGCCAGTGATTCCATTTTTTACCCTTCTGTT
TTTTTGACCTTAAGTCTCCGCATCTTAGCACATCGTTCATCCAGAGCGTGATTTCTGCCGAGCGTGATCAGATCGGCATTTCTTTAATC
TTTTGTTTGCATATTTTTAACACAAAATACACACTTCGACTCATCTGGTACGACCAGATCACCTTGCGGATTCAGGAGACTGACATGCT

```
TTACAAAGGCGACACCCTGTACCTTGACTGGCTGGAAGATGGCATTGCCGAACTGGTATTTGATGCCCCAGGTTCAGTCTCGAGAGATC
CTCTAGATTTAAGAAGGAGATATACATATG
>fdrA
GAGGCAATTCCGACGTCTAAGAAACCATTATTATCATGACATTAACCTATAAAAATAGGCGTATCACGAGGCCCTTTCGTCTTCACCTC
GAGGCAGCGTGCTCACGTTTTAACCCAGCCTGGCAGAGTTTATTCTCAATTAGCTGGTGGAGTGTTTCCCGACTGATTTTCATCTGTCT
TCCTTTTTAACGACGGTGTGAAGCATGACTGCAATTAACATACAGGGAAAATATCTGGATTATGTGATCCAGACAGGCAAAAAAATATA
GTTAGAATTTATTTGATAATCCGCTCACTTTTAACCTGATTTTTAAAACAACAACGCTTATTAAAAAATAATGAGTAATAGCCTGGTGG
TTATTTGAATTCTTTTGTTAATAATTCCTGTGTGATATTCATCACCTTATTTACTCGTTGTCATCGATACCGTAATCGCCACATTAACA
CTGCTCGTGCAATTGCCATGGGTGCAATTTTTAAGGAGTTGTTATGATCCACGCCTTTATTAAAAAAGGGTGTTTTCAGGATTCGGTCA
GTTTAATGATTATTTCACGAAAACTCAGCGAATCAGGATCCTCTAGATTTAAGAAGGAGATATACATATG
>fadE
GAGGCAATTCCGACGTCTAAGAAACCATTATTATCATGACATTAACCTATAAAAATAGGCGTATCACGAGGCCCTTTCGTCTTCACCTC
GATGGATCCCCGCTTCGTTCAGTTCGTTACGAATAAGATCCTGGTACATGAGGATATCCTTCAGCATAAATGTAATAGACAAAATGCAG
TGTACCGGATACCGCCAAAAGCGAGAAGTACGGGCAGGTGCTATGACCAGGACTTTTTGACCTGAAGTGCGGATAAAAACAGCAACAAT
GTGAGCTTTGTTGTAATTATATTGTAAACATATTGCTAAATGTTTTTACATCCACTACAACCATATCATCACAAGTGGTCAGACCTCCT
ACAAGTAAGGGGCTTTTCGTTATGATGATTTTGAGTATTCTCGCTACGGTTGTCCTGCTCTCGAGAGATCCTCTAGATTTAAGAAGGAG
ATATACATATG
>ypeA
GAGGCAATTCCGACGTCTAAGAAACCATTATTATCATGACATTAACCTATAAAAATAGGCGTATCACGAGGCCCTTTCGTCTTCACCTC
GATGGATCCGGCGCGAAGTGAGTGTTTTTAGTGGTTTAAAAGTGCTCATACGGCCTGAGTTGTAATAATAAAGTTCCAGATGTTATATC
GTATCGCGTAATCCGTTACGACGGTTTGCTGTTGAGAATTGTTTTACTTTTCATTTCAATAGTTGACAACTCCCCATTATGCCACTTTT
TCAACCGATTTTCGTCAGATATTGGCTAATTCGACCGTTCACGCCATAATCAGCCTTTTAACCCGCAACTCGAGAGATCCTCTAGATTT
AAGAAGGAGATATACATATG
>creA
GAGGCAATTCCGACGTCTAAGAAACCATTATTATCATGACATTAACCTATAAAAATAGGCGTATCACGAGGCCCTTTCGTCTTCACCTC
GAGCTGCTTTCGCCGCTACATTGTCGAGCGACAGGGGCTGATCCAGATGACCTTCCAGCCAGATTAAAAGGTCGCGAATAATGCCGGCC
TGATCCATAAAATATCCTCATCCTTTCAACAACGAGCACCTGACATCAGGTAATTGGATAATAGCATTTTTTGCTGTTTTAGCATTCAG
TGTTTTTTTCTTAGTAGAGTATGTTTTAGGGCTTGACGGAAATAAAAGTATTGAGATTTTGTTCTTAATCAATATGTTATTTACCGTGA
CGAACTAATTGCTCGTGTAATAGATAAAAATGGTAACAATATGAAATACAAGCATTTGATCCTGTCTTTAAGCCTGATAATGCTGGGGC
CATTGGCTCATGCAGAAGAGATTGGTTCGGTCGACACCGGATCCTCTAGATTTAAGAAGGAGATATACATATG
>creB
GAGGCAATTCCGACGTCTAAGAAACCATTATTATCATGACATTAACCTATAAAAATAGGCGTATCACGAGGCCCTTTCGTCTTCACCTC
GAGCAACGCGCTCGCTTATCTGGCTTACTCCGACAAAGTTGTAGAAGGTTCGCCGAAAAACGCGATTAGCGCGGTTCCTGTCATGCCGT
GGCGGCAATAACAGAGGCGATTTATGCAACGGGAAACGGTCTGGTTAGTGGAAGATGAGCAAGGGATAGCCGACACGCTGGTGGATCCT
CTAGATTTAAGAAGGAGATATACATATG
>inaA
GAGGCAATTCCGACGTCTAAGAAACCATTATTATCATGACATTAACCTATAAAAATAGGCGTATCACGAGGCCCTTTCGTCTTCACCTC
GATGGATCCAAGGCGATGCCTAATCCCTGTCGAATAAAATTCATCGCTGTATCCGCCTGATTAAATTCACATTTTATCTGCTGGGCGCA
CTATTAGTGGTAGTGGCTGGTGGAATCTATGCGCTGCAAATGAAGGGTAAAAGCGGTGTCAAGCTAGTAGTGGCAAAAGAAATTCATTA
ATACGACACGTTTCATTAAGATTTTCCTCAGGTAACCACCTATAGTCATTGTCGTTAATAAAAATGACAGGCGATAGGGTATGGCAGTT
TCAGCAAAGTACGACGAATTTAATCACTGGTGGGCAACAGAGGGCGACTGGGTTGAAGAGCCTCGAGAGATCCTCTAGATTTAAGAAGG
AGATATACATATG
>ompT
GAGGCAATTCCGACGTCTAAGAAACCATTATTATCATGACATTAACCTATAAAAATAGGCGTATCACGAGGCCCTTTCGTCTTCACCTC
GATGGATCCTCTCTGCGCTCACACGTCCCACTACCTCAAAACATGTAAAGCCTTGCAAGCCATTGCGAGGCCTTATGTGTCTCAGTTTT
GTCCCTCTTTTTTGTACTAAAAAACATAGTAATTGAGGATAAACCTCATGCTATTTTCGCTTATATGCCTCTAAAGGCATGGCACTTAA
ATAGATAAAAGCACCACAAAAGCATAAAAAAACCACACAGTAAAACCGAAATATGAAACAATAACAGATAATTAAACCAAAAACAGATA
GCGCATTGTGATAATCATTCAATACTAAACAAAATATAAACAGTGGAGCAATATGTAATTGACTCATTAAGTTAGATATAAAAAATACA
TATTCAATCATTAAAACGATTGAATGGAGAACTTTTATGCGGGCGAAACTTCTGGGAATAGTCCTGACAACCCCTATTGCGATCAGCTC
TTTTGCTTCTACCGAGACTTTATCGTTTACTCCTGACAACATAAATGCGGACATTAGTCTTGGAACTCTGAGCGGAAAAACAAAAGAGC
GTGTTTATCTAGCCGAAGAAGGAGGCCGAAAAGTCAGTCAACTCGACTGGAAATTCAATAACGCTGCAATTATTAAAGGTGCAATTAAT
TGGGATTTGATGCCCCAGATATCTATCGGGGCTGCTGGCTGGACAACTCTCGAGAGATCCTCTAGATTTAAGAAGGAGATATACATATG
>ompW
GAGGCAATTCCGACGTCTAAGAAACCATTATTATCATGACATTAACCTATAAAAATAGGCGTATCACGAGGCCCTTTCGTCTTCACCTC
GAGCGCACATAGCAACGATACCAACAGAATGGTCATAAATTGATTACGGAAGAAATTTCCGGTGTCACGGTATACGGACTGCGCCGTGA
TAGACATGCACTCTCCTTGAGTTTTATGCAGGTGTTAATTAGCGGGCAATTGTACCCTGGTTAAGCCTCGGGTGGCAGCATCAGGTCTT
GTATGGAAAAGCATATCTTTGTAAAGCGGAGGTAATTGCTGGCCTTCAGGCATGGGCAAATCGCGGCAAGTGGCGACCGAGTTAATATT
TGCGTAGCGAAAATATTTAAAAATTGATTTAAATCACATTAACCAGGATTCTCAATGCAACTTCTAAATTAATCCAGATCAATAAAGGG
TGAATTATCATATGTAATGTGATCTATGTAGGATCATTTGTTACTCCAATGTAGGTATATTCGTCACGTTTTTATAACCATAACGACGG
AGCGGATATGAAAAAGTTAACAGTGGCGGCTTTGGCAGTAACAACTCTTCTCTCTGGCAGTGCCTTTGCGCATGAAGCAGGCGAATTGG
ATCCCGGGATCCTCTAGATTTAAGAAGGAGATATACATATG
>ompR
```

GAGGCAATTCCGACGTCTAAGAAACCATTATTATCATGACATTAACCTATAAAAATAGGCGTATCACGAGGCCCTTTCGTCTTCACCTC
GATGGATCCTAACCAGGGGCGTTTTCATCTCGTTGATTCCCTTTGTCTGTTTGATAATGCGCACATTGGGTATAACGTGATCATATCAA
CAGAATCAATAATGTTTCGCCGAATAAATTGTATACTTAAGCTGCTGTTTAATATGCTTTGTAACAATTTAGGCTGAAATTCATACCAG
ATTTAGCTGGTGACGAACGTGAGCTTTTTTAAGAATACACGCTTACAAATTGTTGCGAACCTTTGGGAGTACAAACAATGCAAGAGAAC
TACAAGATTCTGGTGGTCGATGACGACATGCGCCTGCGTGCGCTGCTGGAACGTTATCTCACCGAACAAGGCTTCCAGGTTCGACTCGA
GAGATCCTCTAGATTTAAGAAGGAGATATACATATG
>fiu
GAGGCAATTCCGACGTCTAAGAAACCATTATTATCATGACATTAACCTATAAAAATAGGCGTATCACGAGGCCCTTTCGTCTTCACCTC
GATGGATCCGGCGGAAATGACCAGATGTTCGGTACTGCAACCATCTACAAATAACCCCACCAACGTAACTGCATTCGCCTCTGGATGCA
GTTTTCGTCTGTTTTGTAATCAAAAAACATCAAATATGAACTCAATGTAAATAAATGTATTTCTTTTTCGCGCAATGGGTGATAGAAAA
TCGCTCCAAGTGATAATGCTTATCAAAATTATTATCACTTTCACGAGCACTATCACGGGATTAACAGTGGCATCGCATCCGCAGAGAGG
CTTTCTCGTGGCAGTGAAAATTTCAACATATAAGAAAAAGTCACCTGCAAAATGGAAAACAATCGCAATTTCCCTGCCAGACAATTTCA
TTCGCTCACGTTCTTTGCCGGTCTTTGTATTGGCCTCGAGAGATCCTCTAGATTTAAGAAGGAGATATACATATG
>cld
GAGGCAATTCCGACGTCTAAGAAACCATTATTATCATGACATTAACCTATAAAAATAGGCGTATCACGAGGCCCTTTCGTCTTCACCTC
GATGGATCCACACCCGCGATCTCTTTGGCAGCGACTAACATCTTGTTATCAGGGCTATTTACGCCCTGATTGTCTTTTGTTACTTACAC
AACAATTCATTATTTTTATCACTTATCCTATAGCATTCACGAGGATTATCGCTAAACTATGCGGACTTGGAAATTTCCGTCAGTTAGGG
TAATGATGAGAGTAGAAAATAATAATGTTTCTGGGCAAAACCATGACCCTCGAGAGATCCTCTAGATTTAAGAAGGAGATATACATATG
>ompX
GAGGCAATTCCGACGTCTAAGAAACCATTATTATCATGACATTAACCTATAAAAATAGGCGTATCACGAGGCCCTTTCGTCTTCACCTC
GAGTAACGAGGCTCCACCCTGAATAGACGCCATGGCAACGAGCAATATGACTATTGGTAACCAGACCGGCATTTTACGTAATGAACCAG
GCATCCTTTCTCCCACAAATATCTAGACTTAAGTAAAGCGTGGAGTGTACTGGATATACCCAATGCTGGTTGAGCATTTGTTGAAAAAA
TTTTCCCCCGTTTTGACTAAAATGCGCCAGGATTGATGGAATCATTAGTCTGGTGATTAGGAATAATCTGGATGAATGACAGGGAAAAC
ATGCGTAATACTTACGCAGTTCTCTGAAAAAGTGATTTAAATTTAGATGGATAGCGGTGTATGGAAACGTTCTGTTACATGAAATGGCC
CGTTAGACATCACAAATCGCGAAGAGTTTCCCATTAATTTTTGATATATTTAAAACTTAGGACTTATTTGAATCACATTTGAGGTGGTT
ATGAAAAAAATTGCATGTCTTTCAGCACTGGCCGCAGTTCTGGCTTTCACCGCAGGTACTTCCGTAGCTGCGACTTCTACTGTAACTGG
CGGTTACGCACAGGATCCTCTAGATTTAAGAAGGAGATATACATATG
>yggE
GAGGCAATTCCGACGTCTAAGAAACCATTATTATCATGACATTAACCTATAAAAATAGGCGTATCACGAGGCCCTTTCGTCTTCACCTC
GATGGATCCAGCTGGCGAGAGACGGTATTGCTCATGCACAAGCCTTGTTCAGTTAGGCGCTATCTGATGGAAAAATAAAACAGAGGCGC
TAAGCTTGCCTCCAGAGGTCCTGAATTTTGGGCAAGTCAATTAACACGGAGAGACTAACGTGAAGTTCAAAGTTATCGCCCTGGCGGCA
TTAATGGGTATTAGCGGGATGGCAGCGCAGGCTAACGAATTGCCGCTCGAGAGATCCTCTAGATTTAAGAAGGAGATATACATATG
>ompG
GAGGCAATTCCGACGTCTAAGAAACCATTATTATCATGACATTAACCTATAAAAATAGGCGTATCACGAGGCCCTTTCGTCTTCACCTC
GAGGCCAAAATTAGCGTGGCAGAATTAACCGGTGCGGAATTTATGCTCTACACCCACGGTTGGGGGCACGAGTTAGTGGTCCGTGCTGGT
GCGTTAAATGATTATCATGCAGGAGAAAATATCACTATTCATTTTGATATGACGAAATGTCATTTCTTTGATGCAGAAACGGAAATAGC
AATTCGCTAAATACAGGGGGAAGGCATTCCCCCAGGATAATACAAGGAACAATAATGAAAAAGTTATTACCCTGTACCGCACTGGTGAT
GTGTGCGGGAATGGCCTGCGCACAGGCCGAGGAAAGGAACGACTGGCAGGATCCTCTAGATTTAAGAAGGAGATATACATATG
>ompA
GAGGCAATTCCGACGTCTAAGAAACCATTATTATCATGACATTAACCTATAAAAATAGGCGTATCACGAGGCCCTTTCGTCTTCACCTC
GATGGATCCATCCTCTCACGCCACGAGACAACTTTCCGGGCTAAAAATTCACTCTAATTTGTATCATTAAGTAAATTTAGGATTAATCC
TGGAACTTTTTTTGTCGCCCAGCCAATGCTTTCAGTCGTGACTAATTTTCCTTGCGGAGGCTTGTCTGAAGCGGTTTCCGCGATTCTCT
TCTGTAAATTGTCGCTGACAAAAAAGATTAAACATACCTTATACAAGACTTTTTTTTCATATGCCTGACGGAGTTCACACTTGTAAGTT
TTCAACTACGTTGTAGACTTTACATCGCCAGGGGTGCTCGGCATAAGCCGAAGATATCGGTAGAGTTAATATTGAGCAGATCCCCCGGT
GAAGGATTTAACCGTGTTATCTCGTTGGAGATATTCATGGCGTATTTTGGATGATAACGAGGCGCAAAAAATGAAAAAGACAGCTATCG
CGATTGCAGTGGCACTGGCTGGTTTCGCTACCGTAGCGCAGGCCGCTCCGAAAGATACTCGAGAGATCCTCTAGATTTAAGAAGGAGAT
ATACATATG
>ompC
GAGGCAATTCCGACGTCTAAGAAACCATTATTATCATGACATTAACCTATAAAAATAGGCGTATCACGAGGCCCTTTCGTCTTCACCTC
GATGGATCCAATGCTCCCCGGTAGGAGTGAAAAGCGGGTCGTGGCCGTTGTCTCTTTCTGACGCATTCGTGTTTATGACCTGTTAAAAC
TTCGCGAAGTGATAGGGCAGTATAAAGGGTACAGTGTGAAATTCCAGACTCTTACCTGATTGATGATTCTCTATTCTCCCGTGATATTT
ATCATCCTGCATTGGGGGAAATAGATGTACAAGCGCCATTTTGCTGTACAAATAACCTACAAAAAGCCCAACAAAAATAATTGCAAATA
AACAAAGATTGCTGGAAATTATGCGGATGTTAATTATTTGTGAAATAGTTAACAAGCGTTATAGTTTTTCTGTGGTAGCACAGAATAAT
GAAAAGTGTGTAAAGAAGGGTAAAAAAAACCGAATGCGAGGCATCCGGTTGAAATAGGGGTAAACAGACATTCAGAAATGAATGACGGT
AATAAATAAAGTTAATGATGATAGCGGGAGTTATTCTAGTTGCGAGTGAAGGTTTTGTTTTGACATTCAGTGCTGTCAAATACTTAAGA
ATAAGTTATTGATTTTAACCTTGAATTATTATTGCTTGATGTTAGGTGCTTATTTCGCCATTCCGCAATAATCTTAAAAAGTTCCCTTG
CATTTACATTTTGAAACATCTATAGCGATAAATGAAACATCTTAAAAGTTTTAGTATCATATTCGTGTTGGATTATTCTGCATTTTTGG
GGAGAATGGACTTGCCGACTGATTAATGAGGGTTAATCAGTATGCAGTGGCATAAAAAAGCAAATAAAGGCATATAACAGAGGGTTAAT
AACATGAAAGTTAAAGTACTGTCCCTCCTGGTCCCAGCTCTGCTGGTAGCAGGCGCAGCAAACGCTGCTGAAGTTTCTCGAGAGATCCT
CTAGATTTAAGAAGGAGATATACATATG
>ompN
GAGGCAATTCCGACGTCTAAGAAACCATTATTATCATGACATTAACCTATAAAAATAGGCGTATCACGAGGCCCTTTCGTCTTCACCTC
GATGGATCCCACAAATGGCCGGAAAAGCGGAAAAAAGCAACACCGATTAAATGCTCTGGATAAGGATTATCCAATTCTAAAAAAAAAGC

CCGGACGACTGTTCGGGCTTGTCTTTTTATATGTTGGAAAATCAGTGGCAATGCAATGGCCCAACAGAAAATAAAATCTGTGACAATAA
AGGCATATAACCCGCGCAGAATAACGTATAGCAAATTAAAAGTATAATTTTTATTTTGTATGTATTATCTTCGATACATTCACCAGACT
TATTATCATTTCAGAGGAATTATTTGATTAAGGTTTTACTTAAGGCGTAACAAATGATATTTATCGGCTAACTGAACTTCTCCTTTAGG
ATGTTTTCACTCCCCGAAATGGGGATAAAAGGCAAATAAAATAACTAAAGGATTTATTCAATGAAAAGCAAAGTACTGGCACTTTTAAT
TCCTGCCCTGCTCGCCGCAGGTGCTGCACATGCAGCCGAAGTTTCTCGAGAGATCCTCTAGATTTAAGAAGGAGATATACATATG
>htrL
GAGGCAATTCCGACGTCTAAGAAACCATTATTATCATGACATTAACCTATAAAAATAGGCGTATCACGAGGCCCTTTCGTCTTCACCTC
GATGGATCCTAACGATGTTGCTGCCGATAAAGCCCGCGCCGCCGGTAACGATGATCATAACTGTAACCTTCGAATTATGGAGTCAGAGA
CAATCTCAGACACGAATGTCTGTAATCATACCATCACATCCGAATCCCTTCAGCCCTTTACATAGAGGTGCAAGGATGCTAATTGCAAC
AAACTAGTTTGGGTTTTAGCTAGTCCTTCATGGCGGATGTTGGTTTTGCATGTGAATAGTGCTTTAGTATCAATATTCCGATGTATGAT
AAATAACGGAAGCATTTGAATGGATAATATACAGTATACATTCTATTAATATTTATGATATAATTCTTTTTTTATATTAATGGTCTTAT
TAGTAATGAAATCATCCACCACAATTATCACTGCATATTTTGACATTGGTAGAGGGGATTGGACTGCTCGAGAGATCCTCTAGATTTAA
GAAGGAGATATACATATG
>cls
GAGGCAATTCCGACGTCTAAGAAACCATTATTATCATGACATTAACCTATAAAAATAGGCGTATCACGAGGCCCTTTCGTCTTCACCTC
GATGGATCCAAAGAATCGGCGCAAAAATAGAGTTTAAAGACGGAAGAGTAAACTTATTGGGCACCGCAATGGTGCCCAGTACATTTAGA
TATTGTAGATCGCAAACAATAACGAGTTACGCTGTTTGTTTAGAATGCACTTCCTGATGCTGTGGATACGCATATTTCGGCGCGATTGC
CCCTCAAGCCAACGCGATTTACGTCGGCTAGCCTGACGCTGCATGCGCCAGCGCCCCACTTCCGTTCTACTCCGCTTCATGTTAACTAC
TCTATGCAATAACAGAATGGTCATTATACCCCCACCCCCGACGCTGACCAGTGCTTTTCCCGCGTTTTTATTTGCCAGATGAATCCATA
TGCGTAAACTCATAACAATGCGCTTTCAAAAGGATTTCTAATCTATGACAACCGTTTATACGTTGGTGAGTTGGTTGGCCATTCTGGGA
TACTGGTTGCTCATTGCAGGCGTAACTTTACGCATTCTAATGAAACGACGCGCACTCGAGAGATCCTCTAGATTTAAGAAGGAGATATA
CATATG
>gcvP
GAGGCAATTCCGACGTCTAAGAAACCATTATTATCATGACATTAACCTATAAAAATAGGCGTATCACGAGGCCCTTTCGTCTTCACCTC
GATGGATCCCAGGCGGCTGGATCTTTAAAATCAAAGCCAGCGATGAAAGCGAACTGGAATCACTGCTGGATGCGACCGCATACGAAGCA
TTGTTAGAAGACGAGTAACGGCTTTATTCCTCTTCTGCGGGAGAGGATCAGGGTGAGGAAAATTTATGCCTCACCCTCACTCTCTTCGT
AAGGAGAGAGGTTCACAATTCACTGCACGTTTCAGGAACCATCGCTCATGACACAGACGTTAAGCCAGCTTGAAAACAGCGGCGCTTTT
ATTGAACGCCATATCGGACCGGACGCCGCGCAACAGCAAGAAATGCTGAATGCCGTTGGCTCGAGAGATCCTCTAGATTTAAGAAGGAG
ATATACATATG
>gcvT
GAGGCAATTCCGACGTCTAAGAAACCATTATTATCATGACATTAACCTATAAAAATAGGCGTATCACGAGGCCCTTTCGTCTTCACCTC
GATGGATCCCTTATCCGCCAGGCAATGGGATTAAACGATTTGCCTGAATGGCTGCGTTAAAAATTTCTCCTCTGTTGTTTATTTGATAC
CCATCACACTTTCATCTCCCGGTTTTTTCGCCGGGAGATTTTCCTCATTTGAAATAAACTAATTTCACCTCCGTTTTCGCATTATATTT
TCTAATGCCATTATTTTTTTGATTTAGTGTTTTTTGACATTTTTTTAGCTCTTAATATTGTCTTATTCAAATTGACTTTCTCATCACATC
ATCTTTGTATAGAAACTGGTGTATTTTTTTGGTTTTTTATTCTGTCGCGATTTTTTGCATTTTTTAACCATAAGCTAATGTGATGATCAAT
TTTACCTTATGGTTAACAGTCTGTTTCGGTGGTAAGTTCAGGCAAAAGAGAACGATTGCGTTGGGGACCGGGAGTGGCTCCGATGCTGG
GTTTCGTGGTGATAATTTCACCATGAAAAAGTTGTCAGCCCCGCTTATTCAATGAGGACAAGATGGCACAACAGACTCCTTTGTACGAA
CAACACACGCTTTGCGGCGCTCGCATGGTGGATTTCCACGGCTGGATGATCTCGAGAGATCCTCTAGATTTAAGAAGGAGATATACATA
TG
>tnaC
GAGGCAATTCCGACGTCTAAGAAACCATTATTATCATGACATTAACCTATAAAAATAGGCGTATCACGAGGCCCTTTCGTCTTCACCTC
GAGTTAAGCGAAATCACCGGGGAATTTACTTCAGACGACCTGCTGGGGCGGATTTTCTCCAGCTTCTGTATTGGTAAGTAACCGCGCTT
ACGAAGCCGCATTCTGACTGTCAGATGCGGCTTCGCTTCATTGTTACCACTCCTGTTATTCCTCAACCCTTTTTTTAAACATTAAAATT
CTTACGTAATTTATAATCTTTAAAAAAAGCATTTAATATTGCTCCCCGAACGATTGTGATTCGATTCACATTTAAACAATTTCAGAATA
GACAAAAACTCTGAGTGTAATAATGTAGCCTCGTGTCTTGCGAGGATAAGTGCATTATGAATATCTTACATATATGTGTGACCTCAAAA
TGGTTCAATATTGACAACAAAATTGTCGATCACCGCCGGATCCTCTAGATTTAAGAAGGAGATATACATATG
>srlR
GAGGCAATTCCGACGTCTAAGAAACCATTATTATCATGACATTAACCTATAAAAATAGGCGTATCACGAGGCCCTTTCGTCTTCACCTC
GAGGGTATGCATGCGGGTGATTTACAGCCCGATGTGATCTTTCCCCATGATCCACTATCACAGAATGCTCTATCATTGGCGCTTAAACT
GAAACGTGGATAATTTCGTTGTGAATGTTACTTGCTTGCGAAGTTATCATTTTGAAACCTAAATCAGGTAATCACGCCCATGAAACCTC
GTCAGCGTCAGGCCGCCATTCTGGAGTATCTGCAAGAGCAGGGTAAATGCTCGGTTGAAGAATTGGCGGGATCCTCTAGATTTAAGAAG
GAGATATACATATG
>gcvA
GAGGCAATTCCGACGTCTAAGAAACCATTATTATCATGACATTAACCTATAAAAATAGGCGTATCACGAGGCCCTTTCGTCTTCACCTC
GATGGATCCGCTAATGGCAACAAACAGCAGATCAACCGTACTGACGTGAAAGAGATGGTCGAACTGGATCAGTAATTCGCGATCGCAAG
GTAAAAAAAAGCACCGCAATTAGGCGGTGCTACATTAATCACTATGGACAGACAGGGTAAATGTACAGGAAGTGAAAAAAGGTAGCTTT
GCTACCATGGTCTGAATCGCAGACCAATTGCAAACACAACAACACAACATCACAACCGTAAGCCAAAAGTTCACCAGAACACGCATTCC
GATAAAACTTTTCGTTCCGGCTCAGGAAGTGCCGCCACTATAGGTATTTGCTGGTAGAAGCTCAACGGACAATTTATAATGGCTCAGAT
TAAAAAAACTAATAGGTTACATAGTGTGATCTAATTGTTAAATTCATTTAACATCAAAGTTTAATAGCCATGTCTAAACGATTACCACC
GCTAAATGCCTTACGAGTTTTTGATGCCGCAGCACGCCATTTAAGTTTCACTCGCGCAGCCTCGAGAGATCCTCTAGATTTAAGAAGGA
GATATACATATG
>ftsN

Data Set S4 - MG1655 library reference file.fasta
GAGGCAATTCCGACGTCTAAGAAACCATTATTATCATGACATTAACCTATAAAAATAGGCGTATCACGAGGCCCTTTCGTCTTCACCTC
GATGGATCCGTTGGCAGTGGCTCTCGTTTAATGGACTGCGAACTTATCATCCGGGGATCAACACGCGCGTTACCTTAAAGTAAACCTTT
AAGACTGCCGTGACTGGTCAAAGGCCCGCCGCTTAAGTAACATGGCGGGCTGACGAACGAATAAATACAGCGAAACGATAGTGGCACAA
CGAGATTATGTACGCCGCAGCCAACCGGCACCTTCGCGGCGAAAAAAGAGCACCTCACGGAAAAAGCAACGAAATCTGCCTGCCTCGAG
AGATCCTCTAGATTTAAGAAGGAGATATACATATG
>ftsK
GAGGCAATTCCGACGTCTAAGAAACCATTATTATCATGACATTAACCTATAAAAATAGGCGTATCACGAGGCCCTTTCGTCTTCACCTC
GAGGCGTCAATGACACACGGACATACGTTGTTATGGAAGAAGTCAAGCAGAGTAATCGTCTGGTTATTAAGACGCGCTAACACGGAACA
GGTGCAAAATCGGCGTATTTTGATTACACTCCTGTTAATCCATACAGCAACAGTACTGGGGTAACCTGGTACTGTTGTCCGTTTTAGCA
TCGGGCAGGAAAAGCCTGTAACCTGGAGAGCCTTTCTTGAGCCAGGAATACATTGAAGACAAAGAAGTCACATTGACAAAGTTAAGTAG
CGGCCGCCGCCTTCTGGAGGCGTTGCTGATGGATCCTCTAGATTTAAGAAGGAGATATACATATG
>aniC
GAGGCAATTCCGACGTCTAAGAAACCATTATTATCATGACATTAACCTATAAAAATAGGCGTATCACGAGGCCCTTTCGTCTTCACCTC
GATTGATCACTACGGTATGACGCCGCTGCATTATGTTAGTCAGCACAGAGAACGCACTGTCGCCTGATTTTTAACCTTAACGAAGAGCT
ATATTAATAACGGCATCAGCGATAACCCGGTCGATAATAATTCAACTATCGAATGCAGGCGTATGATATGACGTAATTTATTGTCACGA
AGCTCGCCTTCGCAGGAGTTTAATTATGTCTTCGGATGCTGATGCTCACAAAGTGGGCTTAATCCCCGTCACCCTGATGGTGTCGGGGA
ATATTATGGGGCTCGAGAGATCCTCTAGATTTAAGAAGGAGATATACATATG
>ftsH
GAGGCAATTCCGACGTCTAAGAAACCATTATTATCATGACATTAACCTATAAAAATAGGCGTATCACGAGGCCCTTTCGTCTTCACCTC
GATGGATCCACGTTCGCGGGAAGTGTATATTGTAGCGACCGGGCGTAAACCCTAACCGGGAGATTTCAGACGAAAGTTTGAAAGATGCT
GGATATAGAGTATCCTGACGCTGTTTTTAACACAGTTGTAATAAGAGGTTAATCCCTTGAGTGACATGGCGAAAAACCTAATACTCTGG
CTGGTCATTGCCGTTGTGCTGATGTCAGTATTCCAGAGCTTTGGGCCCTCGAGAGATCCTCTAGATTTAAGAAGGAGATATACATATG
>yieE
GAGGCAATTCCGACGTCTAAGAAACCATTATTATCATGACATTAACCTATAAAAATAGGCGTATCACGAGGCCCTTTCGTCTTCACCTC
GAGCGGAACAGCCATAATCCGAAGATCGTCTGGTTACGGGAAACCATTAAAAACCTTTACGCGTCGATGGCATAACCGAATCGTATGAA
ACGCGACCCAATTTCACAATAAAATGTAAAAAAGTTGTAATAAGCTTGTCTGAATCGAACTTTTAGCCGCTTTAGTCTGTCCATCATTC
CAGTAAATGATTACTCTTGTATTCATAATGGACCATTAAGCATGGAGCGAAAAATGGCGACTCACTTTGCCCGAGGGATTTTAACGGAA
GGACATCTGATTTCTGTTCGTCTCCCCTCCCAGTGTCAGGATCCTCTAGATTTAAGAAGGAGATATACATATG
>ftsQ
GAGGCAATTCCGACGTCTAAGAAACCATTATTATCATGACATTAACCTATAAAAATAGGCGTATCACGAGGCCCTTTCGTCTTCACCTC
GAGCGGGTATGACCAGCCACAGCCTGGTGCCGATGGCGGCACGTCAGGCAGGTATGAGCTTCTCGCAGTTGGTAGTACGAATTCTGGAA
CTGGCGGACTAATATGTCGCAGGCTGCTCTGAACACGCGAAACAGCGAAGAAGAGGTTTCTTCTCGCCGCAATAATGGAACGCGTCTGG
CGGGGATCCTTTTCCTGCTGACCGAGATCTTCTAGATTTAAGAAGGAGATATACATATG
>ftsZ
GAGGCAATTCCGACGTCTAAGAAACCATTATTATCATGACATTAACCTATAAAAATAGGCGTATCACGAGGCCCTTTCGTCTTCACCTC
GAGTAAACATCACCTGGCGGCAGGCATTGTATTAACCGGTGGCGCAGCGCAGATCGAAGGTCTTGCAGCCTGTGCTCAGCGCGTGTTTC
ATACGCAAGTGCGTATCGGCGCGCCGCTGAACATTACCGGTTTAACGGATTATGCTCAGGAGCCGTATTATTCGACGGCGGTGGGATTG
CTTCACTGGATCCTCTAGATTTAAGAAGGAGATATACATATG
>ftsY
GAGGCAATTCCGACGTCTAAGAAACCATTATTATCATGACATTAACCTATAAAAATAGGCGTATCACGAGGCCCTTTCGTCTTCACCTC
GATGGATCCGGAACCGGGAGTTTACGGCCTCGCCACTGCCCGCCAATAATGCGGATTTGGCCGCTGCCGGAATGATTCGGTTTTTTCAT
GATAAATTGCTCAATCCGCCAGATAACACATACTTGCAGGCGGTGATGTGAATTAAGTTAAGTGATAGACTATTTCATCATTTTTTTAG
CTGCTATGTACATAGCGTTAACGCTGTGCCATGAAGCAACAGCGAGGAGTGTAGTCGCAAATGGCGAAAGAAAAAAAACGTGGCTTTTT
TTCCTGGCTGGGCTTTGGTCAAAAAGAGCAGACCCCGGAAAAAGCTCGAGAGATCCTCTAGATTTAAGAAGGAGATATACATATG
>hemA
GAGGCAATTCCGACGTCTAAGAAACCATTATTATCATGACATTAACCTATAAAAATAGGCGTATCACGAGGCCCTTTCGTCTTCACCTC
GAGACGGAACAGGCAGTGAGCACAAGAGCAGCCAGCGGTAGCAGGCGGATAAGACGAAAATCGGGCAGGGGCATAGTGATGACAAGTCC
TTGAGATACGTTGCAGTTATAACCCTTAATGCTAGCGTTACCGTCCGCTATCGTCTATGTTCAAGTTGTCTTAATTGCCAGAATCTAAC
GGCTTTCGGCAATTACTCCAAAAGGGGGCGCTCTCTTTTATTGATCTTACGCATCCTGTATGATGCAAGCAGACTAACCCTATCAACGT
TGGTATTATTTCCCGCAGACATGACCCTTTTAGCACTCGGTATCAACCATAAAACGGCACCTGTATCGCTGCGAGAACGTGTATCGTTT
TCGCCGGGGATCCTCTAGATTTAAGAAGGAGATATACATATG
>yrfF
GAGGCAATTCCGACGTCTAAGAAACCATTATTATCATGACATTAACCTATAAAAATAGGCGTATCACGAGGCCCTTTCGTCTTCACCTC
GAGAGATCCACGCTCTCGACGGTAAACAGTCGGGAACGGGCTACAGTTTCAACATTCAGAATGGTGGGTTTTTGTAATGATTTGCTCAT
CGTGGGATCTATGCTGTGAAATCAGCGGTTATTGTGCGATATCGGACACGCTTTCGGCAATGTGAATTGCATGTTATTTACATTTATGT
AACTTAATAAATAATCGTCCTCAAATCAAATTAAAAGTCAATAGGTTGAAATAACTCCAGGAATTTGCTGATATTCCGCCTTCAGAGGG
TTTGCTATGATCAGCGGTTACTGTGATGTGCTTAATGATGCTCAAGTTAAACTCCACGCTTGCCGATAGCCAACCGCAGAATCATGTAT
TGTGTCCGGTGCGACTGACCACGCCTGACAGACTAAGTAAGATGGGGAAAGCATGAGCACCATTGTGATTTTTTTAGCTGCTTTGCTGG
CCTGCTCACTACTTGCGGGATGGCTGATAAAAGTGCGATCCAGACGGGGATCCTCTAGATTTAAGAAGGAGATATACATATG
>pgsA
GAGGCAATTCCGACGTCTAAGAAACCATTATTATCATGACATTAACCTATAAAAATAGGCGTATCACGAGGCCCTTTCGTCTTCACCTC
GATGGATCCTGCAAAAGTGCCGGGTATTTCGCAAGGTCTGGCAGAAAAGATCTTCTGGTCGTTGAAACATTGATGTCTCTGTAGCAACA

```
TAGGGGTAATCTTACTGACAACAGATAGTTACCCGTCATTATGCAATTTAATATCCCTACGTTGCTTACACTGTTCCGTGTCATCCTTA
TCCCATTCTTTGTATTGGTCTTTTATCTGCCTGTCACCTGGTCGCTCGAGAGATCCTCTAGATTTAAGAAGGAGATATACATATG
>fepB
GAGGCAATTCCGACGTCTAAGAAACCATTATTATCATGACATTAACCTATAAAAATAGGCGTATCACGAGGCCCTTTCGTCTTCACCTC
GATAGATCTTGTTGCCATGGTCTGCTGTACTTCCTCAGCCAGTGACGTATCCATATCATCCTCCACAAAATGATAAAGGCTTTAATAAT
GATTATCATTTATATTTTCGGTCGCTAACCTAAAAGCAAACGCGCACTATGTCAACTCTTGAGGTAACGCAATCTGCAACTCGGGGTTG
CATCTGGTTGTGGTTACAATGAAAATGAGAAGCATTATTGATGGATTCGCATAAGCGCAATGTGATGTCCTGCGCCGTTCTGCCCCCTC
TCCCTTCCAGGGTGAGGGCTGGGGTGAGGGTTAATGTTCGCACCAGTGCTGGCTGTTCCCCTCACCCTAACCCTCTCCCCAAAGGGGCG
AGGGGACGGATTGTGCGCTTTGTCGAATTTGTCATTACGCCCTTAACCTTATTAATAACAGGAAGCTGATTTGTGAGACTCGCCCCGCT
CTACCGCAACGCCCTTCTATTAACAGGACTTTTGCTTTCAGGAATAGCCGCAGTTCAGGCCGCTGACTCTCGAGAGATCCTCTAGATTT
AAGAAGGAGATATACATATG
>fepA
GAGGCAATTCCGACGTCTAAGAAACCATTATTATCATGACATTAACCTATAAAAATAGGCGTATCACGAGGCCCTTTCGTCTTCACCTC
GATGGATCCCGCGCTTTATCGTCGAGTATTCTTCAGAACCTTGGGGATCACGCCACCAGAAAGTGACCTCAAACATTTCGTCATTCAGA
CGCTGCCATTCCGGGCCATGTTTCGACTGCCACCAGCTCTCACTTCCTACTTTTAACGCCGTCACACCATAACCCCATGTTTACTGTGC
AATTTTTCATTGATTGCAGAAATATATTGATAATATTATTGATAACTATTTGCATTTGCAATAGCGTAATGGCGCGCCGTGGGAAGCGC
GGACATTAATTAACCAACTGCACTGCGTGTCTTTCAGGATCAAAGGTTTTCGCGGTAGCGGGATGCGTCGTGTTGATGACGACCATGCC
CGACAGTTGCAATTCGTGGCAAAAATGCAGGAATAAAACAATGAACAAGAAGATTCATTCCCTGGCCTTGTTGGTCAATCTGGGGATTT
ATGGGGTAGCGCAGGCACAAGAGCCGACCCTCGAGAGATCCTCTAGATTTAAGAAGGAGATATACATATG
>fepD
GAGGCAATTCCGACGTCTAAGAAACCATTATTATCATGACATTAACCTATAAAAATAGGCGTATCACGAGGCCCTTTCGTCTTCACCTC
GATGGATCCGGGTGCGTTTTCAACAGGCTGAGGTTAAGCAGCCAGGATTGTTTATTCATTACAATGCCTTGCCATCAAATATTTTAAAT
ACCTATTCTTAGGCTGCACATGCTAACATATCCAAATAAGATCGATAACGATAATTAATTTCATTATCATGGAAGTTCGTATGTCTGGT
TCTGTTGCCGTGACACGCGCCATTGCCGTGCCCGGATTGCTGTTATTACTGATTATCGCGACGGCATTAAGCCTGCTCATTGGGGCAAA
ATCTCGAGAGATCCTCTAGATTTAAGAAGGAGATATACATATG
>yiiR
GAGGCAATTCCGACGTCTAAGAAACCATTATTATCATGACATTAACCTATAAAAATAGGCGTATCACGAGGCCCTTTCGTCTTCACCTC
GAGGGTCGTTGCCGTATCTGGTGTTTGTGCATGCGCCTCTGTTGTAACGGTTAATGCAGAACATAAGAGAAAAAACAGCGTACACCCTG
GCTTCATTGACAGTACCTTTTGTTATGACTGGCTCCGATTAAAACGATATCTGTCGCCCTTGTCAAAAGGGTAGAATCCTGGAAAAGAC
AACCATCAAGGAACTTACATGACCATACAGCAATGGTTATTCTCATTTAAAGGGCGTATTGGACGCCGTGATTTCTGGATTTGGATAGG
CCTGTGGTTCGGGATCCTCTAGATTTAAGAAGGAGATATACATATG
>ligA
GAGGCAATTCCGACGTCTAAGAAACCATTATTATCATGACATTAACCTATAAAAATAGGCGTATCACGAGGCCCTTTCGTCTTCACCTC
GATGGATCCAGTCAAAGACGCCAACGCCTGATACACTTAAGGCAAATTAACTCCTCTTCGAACCCCCGCTTGTCGGGGGTTTTTAGCAT
TGATGGTGCGATATGGAATCAATCGAACAACAACTGACAGAACTGCGAACGACGCTTCGCCATCATGAATATCTTTATCATGTGATGGA
TGCGCCTCGAGCGGATCATGTGATGGATGCGCCTCGAGAGATCCTCTAGATTTAAGAAGGAGATATACATATG
>ligB
GAGGCAATTCCGACGTCTAAGAAACCATTATTATCATGACATTAACCTATAAAAATAGGCGTATCACGAGGCCCTTTCGTCTTCACCTC
GATGGATCCTCAGGCTGGATTTACCCGCGCCACTGGGGGCAGAAACAATATAAAGCGTGCCTTGAGCCATGAGTATCTTTCGTATGTGA
TTAGCGAAATAAAGCCTACATACGAGCTTATTATACACGGCGCGGCCATGTGACGTAGCCCTTGCTCACACTTTTTGCGCAAAGTTGTT
GCGTTTTGCGTAGCTGCTTCCGCATTTGCCTGTCATTCACTGCAATAAAGACGATTTTCAGGAAGCGTCCTCGCAGACTCCGTTTTTGC
CACTCGGCTTTGCTTTCATCACGTTGTATATCTGCCTGACTATCTACAGCGGAGGTGATGATGAAAGTATGGATGGCGATATTAATAGG
TATCTTGTGCTGGCAATCATCTGTGTGGGCGGTCTCTCGAGAGATCCTCTAGATTTAAGAAGGAGATATACATATG
>aidB
GAGGCAATTCCGACGTCTAAGAAACCATTATTATCATGACATTAACCTATAAAAATAGGCGTATCACGAGGCCCTTTCGTCTTCACCTC
GAGGATGATGAAGCGTGCGGAATGGGGATCCGTGAAGATAACACACTGATCACCAAAGACACCTCACGTTTCGTTCCGCATTACATTGC
TGGATAAGAATGTTTTAGCAATCTCTTTCTGTCATGAATCCATGGCAGTGACCATACTAATGGTGACTGCCATTGATGGAGGGAGACAC
AGTGCACTGGCAAACTCACACCGTTTTTAATCAACCTATACCATTAAATAACAGCAACTTATACCTGTCTGATGGCGCGCTCTGCGAAG
CGGTAACGCGTGAAGGGCACCAAAAGGCGCTGGAAAGCCTTCGTGCACTCGAGAGATCCTCTAGATTTAAGAAGGAGATATACATATG
>ligT
GAGGCAATTCCGACGTCTAAGAAACCATTATTATCATGACATTAACCTATAAAAATAGGCGTATCACGAGGCCCTTTCGTCTTCACCTC
GATGGATCCTCTTCGCGCCACATTGTAGCATCGCGGTAATTCATAACCGAGTGCCTCACATGTCTGAACCGCAACGTCTGTTCTTTGCT
ATCGACTTACCTGCAGAAATCCGCGAACAGATTATCCATTGGCGCGCTCGAGAGATCCTCTAGATTTAAGAAGGAGATATACATATG
>smf
GAGGCAATTCCGACGTCTAAGAAACCATTATTATCATGACATTAACCTATAAAAATAGGCGTATCACGAGGCCCTTTCGTCTTCACCTC
GATGGATCCCGCTCGTCCGGAATATGTAACACTTGCAAAACTGACATAAATCTCCAGAGATGTGTTCAGGAGTTAGAAAGATTATTTCT
TCTATTCTAGACAAATCCCCCTCTGATTGACAGCATCACTGACCAATCGCAAAGATTGCTAAGGCTGCTTATGGCAGGGAGATAAGGAT
GGTCGATACAGAAATTTGGCTGCGTTTAATGAGTATCAGCAGCTTGTACGGCGATGATATGGTCCGTATCGCTCACTGGGTCTCGAGAG
ATCCTCTAGATTTAAGAAGGAGATATACATATG
>prfB
GAGGCAATTCCGACGTCTAAGAAACCATTATTATCATGACATTAACCTATAAAAATAGGCGTATCACGAGGCCCTTTCGTCTTCACCTC
GATGGATCCGCGAAGTGCAACTGGCTTATAAGCTCGATATCAACGAGTTTCGCGGCAACCGCAGCCTGCAAATTATCATCGACAATATC
TGGCCAATTTAGCGTCATCTTCTCTATAAAAAAGAGCGTGGATTGGGTACAATCCCGCTCTTATCACCGCATTTTGACTAGCTCAATAA
```

AAGAAATCAGACCATGTTTGAAATTAATCCGGTAAATAATCGCATTCAGGACCTCACGGAACGCTCCGACGTTCTTAGGGGGTCTCGAG
AGATCCTCTAGATTTAAGAAGGAGATATACATATG
>prfC
GAGGCAATTCCGACGTCTAAGAAACCATTATTATCATGACATTAACCTATAAAAATAGGCGTATCACGAGGCCCTTTCGTCTTCACCTC
GAGAGCAACCAGAAGGCATCGCGCCCACCTGGACCGTTTCTTCGTTGCACGAACTGGAGCAGCTCCTGTGTAAACACTGATTGCCTCCC
CCCCGTTGATGGGTAAAATAGCCGCAATTTTTCGTTTTCAACAAGCGCGGCGCGATGCCGCTTACTCAAGAAGAAAGAATTATGACGTT
GTCTCCTTATTTGCAAGAGGTGGCGAAGCGCCGCACTTTTGCCATTATTTCTCACCCGGACGCGGATCCTCTAGATTTAAGAAGGAGAT
ATACATATG
>dcuA
GAGGCAATTCCGACGTCTAAGAAACCATTATTATCATGACATTAACCTATAAAAATAGGCGTATCACGAGGCCCTTTCGTCTTCACCTC
GATGGATCCATGCACCCGGCTTACAAAGCAAAACGCTATACTGATGAAAGCGAACAGTAATCGTACAGGGTAGTACAAATAAAAAAGGC
ACGTCAGATGACGTGCCTTTTTTCTTGTGAGCAGTAACTTAAAAATAACAATCTAATATCAACTTGTTAAAAAACAAGGAAGGCTAATA
TGCTAGTTGTAGAACTCATCATAGTTTTGCTGGCGATCTTCTTGGGCGCCAGATTGGGGGGAATAGGTATTGGTTTTGCAGGCGGCTCG
AGAGATCCTCTAGATTTAAGAAGGAGATATACATATG
>sodB
GAGGCAATTCCGACGTCTAAGAAACCATTATTATCATGACATTAACCTATAAAAATAGGCGTATCACGAGGCCCTTTCGTCTTCACCTC
GAGGCACAGGTCAGGAAATTCAAATCACTTCTCTCAGTGAAGACTACTGGCAGCGCCACTATGTTGGCGCTCGTCGGGTAATGACCCCA
AAAACACTTCGCTAAAACTTTACCCTGTTGTTACGGCAACAGGGTAAGTTCATCTTTTGTCTCACCTTTTAATTTGCTACCCTATCCAT
ACGCACAATAAGGCTATTGTACGTATGCAAATTAATAATAAAGGAGAGTAGCAATGTCATTCGAATTACCTGCACTACCATATGCTAAA
GATGCTCTGGCACCGCACATTTCTGCGGAAACCATCGAGTATCACTACGGCAAGCACCAGGATCCTCTAGATTTAAGAAGGAGATATAC
ATATG
>sodC
GAGGCAATTCCGACGTCTAAGAAACCATTATTATCATGACATTAACCTATAAAAATAGGCGTATCACGAGGCCCTTTCGTCTTCACCTC
GATGGATCCATCCGTAAAGCGGCACTGGGGTACGACGTACCGTAAGCACTTTTAGGAATAGCCGCCGTTCAAAAATGTGTCACTGGTTT
ACACTTATTCAGGAATGCACAATGAACGGAGGTCCTATGAAACGTTTTAGTCTGGCTATTCTGGCGCTGGTTGTTGCAACCGGCGCACA
AGCTGCCAGTGAAAAAGTCGAGATGAACCTCGTCACGTCGCACTCGAGAGATCCTCTAGATTTAAGAAGGAGATATACATATG
>sodA
GAGGCAATTCCGACGTCTAAGAAACCATTATTATCATGACATTAACCTATAAAAATAGGCGTATCACGAGGCCCTTTCGTCTTCACCTC
GAGCGGAGCGTAAAAACAGGCTGCACTGGCCGCGCCGATCAAATGCCAAAATATCCCCATCGTAATCGCGTTACTCATCTTCTTATCCT
CATCATTTTTCGTCGCGTCACATCTCCGACGAGATGAGTGTAAAAATCGTGCTGTCGATTAACCTTTCGCCTGTTGCCGCCGTTGTCGA
TTTACTGGCAATCACGGCATTAAGTGGGTGATTTGCTTCACATCTCGGGCATTTTCCTGCAAAACCATACCCTTACGAAAAGTACGGCA
TTGATAATCATTTTCAATATCATTTAATTAACTATAATGAACCAACTGCTTACGCGGCATTAACAATCGGCCGCCCGACAATACTGGAG
ATGAATATGAGCTATACCCTGCCATCCCTGCCGTATGCTTACGATGCCCTGGAACCGCACTTCGATAAGCAGACCATGGAAATCCACCA
CACCAAACACCAGGATCCCGCTCGAGCGGAGCGTAAAAACAGGCACCACACCAAACACCAGGATCCTCTAGATTTAAGAAGGAGATATA
CATATG
>erfK
GAGGCAATTCCGACGTCTAAGAAACCATTATTATCATGACATTAACCTATAAAAATAGGCGTATCACGAGGCCCTTTCGTCTTCACCTC
GATGGATCCCCATCATCGAAGCTGCCTGTGCGATATACAACAACATGGGCGAACTTGCTGCCAGTAATATTGTTCTACCGGGGAATACG
ACTTCTGATTTGAACAGTTAAAAACGAAAAGTCTGATAAAAGCTATACTTAACGGATAGCTTTCGCGACATAGGAAAGGGACATGATGC
GTCGTGTAAATATTCTTTGCTCATTTGCTCTGCTTTTTGCCAGCCATACTAGCCTGGCGGTAACTTATCCATTACCTCCAGAGGGTAGC
CGTTTCTCGAGAGATCCTCTAGATTTAAGAAGGAGATATACATATG
>dcuS
GAGGCAATTCCGACGTCTAAGAAACCATTATTATCATGACATTAACCTATAAAAATAGGCGTATCACGAGGCCCTTTCGTCTTCACCTC
GATGGATCCCGTCCAGTAGCGCCTGATCCATGACCTTCCCTCTCTCATCAGTATGATTATCGCTATAAGCATAGCCCCGGAATCTTCAT
ACAGCACTGGCAGTTTCCGTTGCCAGAGTCTCTTCCCCTATATAGCCAATCATTCTTACTCCCTTTGAATTACCCGCCTCATCAGAGAT
AATGCTTAAGAAATTTGTCACACAAGGAAGCTGATGAGACATTCATTGCCCTACCGCATGTTACGCAAACGTCCGATGAAATTGAGTAC
CACAGTGATCTTAATGGTCAGTGCGGTACTGTTCTCGGTGCTCGAGAGATCCTCTAGATTTAAGAAGGAGATATACATATG
>ybbL
GAGGCAATTCCGACGTCTAAGAAACCATTATTATCATGACATTAACCTATAAAAATAGGCGTATCACGAGGCCCTTTCGTCTTCACCTC
GAGCCCTGCGGTACGATTTTGACACCCGCGCCGACAATGACCAGCGCGACAAAAATGAGAATCGGGATAAAGATAAGCATCGGAAAAAC
CTCCTGTTGTACCGTCCATAATCAGCAAAATTGCTGCTTGATTAAACAAATTATACCTGATTACTGAAAGAGAGTTCCCCCTTATTCCT
GCGAAGGATAAACTGTTTTTAGTAAAAATCAGAAAAAGGGAACAGCGATGCAGGAAAATAGTCCTTTGCTTCAGCTACAAAACGTAGGA
TATCTGGCGGGTGATGCGAAGATTCTTAATAACATCAATTTTTCGCTGCGTGGGATCCTCTAGATTTAAGAAGGAGATATACATATG
>oxyR
GAGGCAATTCCGACGTCTAAGAAACCATTATTATCATGACATTAACCTATAAAAATAGGCGTATCACGAGGCCCTTTCGTCTTCACCTC
GAGAATCGTGCCTCGACAAGCGTGCGGCAAAAGGCGGCGTCTCACCGCAGCAGGTGGCGCAGGCGATTGCTTTTGCGCAGGCTCGGTTA
GGGTAAGAACATTTATATGTATAAATTTGAGCCTGGCTTATCGCCGGGCTTTTTTATGGCAAAAAAAAGCGGATCCTGGAGATCCGCAA
AAGTTCACGTTGGCTTTAGTTATTCGAGTTGAGAAACTCTCGAAACGGGCAGTGACTTCAAGGGTTAAAAGAGGTGCCGCTCCGTTTCT
GTGAGCAATTATCAGTCAGAATGCTTGATAGGGATAATCGTTCATTGCTATTCTACCTATCGCCATGAACTATCGTGGCGATGGAGGAT
GGATAATGAATATTCGTGATCTTGAGTACCTGGTGGCATTGGCTAGATCCTCTAGATTTAAGAAGGAGATATACATATG
>umuD
GAGGCAATTCCGACGTCTAAGAAACCATTATTATCATGACATTAACCTATAAAAATAGGCGTATCACGAGGCCCTTTCGTCTTCACCTC
GAGTCCATCTGCGGTTTCGATTGCGTTTTTAACTACTTCTACCGTTTTATCTGCAACGATTTCAGTCATAATCATTCGCCTCTTTAAAT

ATATAAATTGTAATGAAACTCCTGTTTTACAACTATTAATAAATTTTACTTCATCTAATTCATAGTTAGCCGGGCGGGATGCGTCAATG
TCTTTATTTCTATTAATATGATAAATATCAAACAATGTTTAATGTCATTATGGCGAATGCTTCTATTCTATTTTTTAGCCGGGTGATAT
TTTTCATTTCTGCTGGATGAGCGTCGTCGCCAGAAGGCCACGTGAGCACAAGATAAGAGAACGAAAAATCAGCAGCCTATGCAGCGACA
AATATTGATAGCCTGAATCAGTATTGATCTGCTGGCAAGAACAGACTACTGTATATAAAAACAGTATAACTTCAGGCAGATTATTATGT
TGTTTATCAAGCCTGCGGATCTCCGCGAAATTGTGACTTTTCCGCTATTTAGCGATCTTGTTCAGTGTGGCTTTCCTTCACCGGCAGCA
GATTGGATCCTCTAGATTTAAGAAGGAGATATACATATG
>ybbC
GAGGCAATTCCGACGTCTAAGAAACCATTATTATCATGACATTAACCTATAAAAATAGGCGTATCACGAGGCCCTTTCGTCTTCACCTC
GAGGATGGACGATTTGAGGCGCTTAATAATAAAGGAGAATATATTTTTATCAGAAATAGTGTCCCGGGATTGAGTTCAGTATGCATACC
GTATCATGATTAATTTTAGTGCTTTTATTAGTGGGGCCTATAAGGAGATTCAATGAAATATAGTTCAATATTTTCGATGCTTTCATTTT
TTATACTATTTGCCTGTAATGAGACAGCTGGGATCCTCTAGATTTAAGAAGGAGATATACATATG
>ybbB
GAGGCAATTCCGACGTCTAAGAAACCATTATTATCATGACATTAACCTATAAAAATAGGCGTATCACGAGGCCCTTTCGTCTTCACCTC
GATGGATCCCGGCAAAGCGGTAGAAGATATTGTGACCTTGTTTACCCAGCGCAGGCCGGAAATCAGCGGATTTTTAGAAATTTTCGGCA
ACCCACGCAGTTAAAAATCAAGTTTCCTTGCGCGTTGGCGCGTAAAATAGCGCGCAAGCTGATTTGAGGTTCACTTTTAACTATGCAAG
AGAGACACACGGAACAGGACTATCGTGCCCTGCTGATTGCTGATACGCCCATTATTGATGTTCGCGCCCCTATCGAGTTTGAGCACGGC
CTCGAGAGATCCTCTAGATTTAAGAAGGAGATATACATATG
>yphG
GAGGCAATTCCGACGTCTAAGAAACCATTATTATCATGACATTAACCTATAAAAATAGGCGTATCACGAGGCCCTTTCGTCTTCACCTC
GATGGATCCTGGTGGCGAATCTGTTGATTATTAATGCAGGCTCTCATCTTCACTAAACCTTGTTTAGTTAATTGCGTGATTATTCCGCG
AAAACATTGCCAGATTGTTCAATACACTGCCACAAATCTTTTAATTCAGTATGTCTTGTTAATATTGAGGGCACCATGACTCCAGCTCG
AGAGATCCTCTAGATTTAAGAAGGAGATATACATATG
>ybdG
GAGGCAATTCCGACGTCTAAGAAACCATTATTATCATGACATTAACCTATAAAAATAGGCGTATCACGAGGCCCTTTCGTCTTCACCTC
GATGGATCCTCTGGTGGTTGTTCCGGTAGGTCATCACAGCGTTGAAGATTTTAACGCTACGCTGCCGAAATCTCGTCTGCCGCAAAACA
TCACCTTAACCGAAGTGTAATTCTCTCTTGCCGGGCATCTGCCCGGCTATTTCCTCTCAGATTCTCCTGATTTGCATAACCCTGTTTCA
GCCGTCATCATAGGCTGCTGTTGTATAAAGGAGACGTTATGCAGGATTTAATATCCCAGGTTGAAGATTTAGCGGGTATTGAGATCGAT
CACACCACCTCGATGGTGATGATTTTCGGCTCGAGAGATCCTCTAGATTTAAGAAGGAGATATACATATG
>fixC
GAGGCAATTCCGACGTCTAAGAAACCATTATTATCATGACATTAACCTATAAAAATAGGCGTATCACGAGGCCCTTTCGTCTTCACGTC
GACTACGCGGATTACGGCATTGTTGGCGACGCCGTGAAGATCCTTCCGGCGCTGACCGCAGCTTTAGCGCGTTGATCCACTCTGGCAGG
GCTGCATTTTGGCCCTGCCGCTGACAGGGAGCTCTTATGTCCGAAGATATCTTTGACGCCATCATCGTCGGTGCAGGGCTTGCCGGTTC
GGTTGCCGCACTGGTGCTCGCCCGCGAAGATGCGCAAGTGTTGGATCCTCTAGATTTAAGAAGGAGATATACATATG
>glyQ
GAGGCAATTCCGACGTCTAAGAAACCATTATTATCATGACATTAACCTATAAAAATAGGCGTATCACGAGGCCCTTTCGTCTTCACCTC
GATGGATCCACACGCTATCCCTCGCAGGTTATCAATCCAGTAAATTTTGGGCTGCATCTGTGTCTCTGTATCTGGTTAAAAAAAGTCTG
ACCGATAAATCATTTGGTTGGCGCACTGGAATAATCTGAGTTTTATCACTACAGCTTATAGAGGCTTAAGGAAATTCGTAAGATATCAG
CCACTATACCGATATAAATAATAAGACTCACCTGCAAACCAGACGGTAATTTAATGATGATGAACGCTTTCTTTCCGGCAATGGCGCTT
ATGGTGCTAGTGGGTTGTTCTATACCGTCACCCGTGCAGAAAGCACAACGGGTAAAGGTTGATCCTCTGCGTTCGTTGAATATGGAAGC
GTTATGCAAGGATCAGGCGGCAAAACGTTATAACACCGGCGAGCAAAAAATCGACGTCACCGCCTTCGAACAGTTCCAGGGAAGCTATG
AAATGCGCGGTTATACCTTCCGTAAAGAGCAGTTTGTCTGTTCTTTTGACGCGGATGGCCATTTTTTGCATCTTTCCATGCGTTAAGCC
CTGCTTTTTCCCGTTTCGTACTGTATATCTTCCATCCAGCGGGTATACTGATCCCTTCCTTTAAATCCACACGTATCCAGCACGAAATA
ATATGCAAAAGTTTGATACCAGGACCTTCCAGGGCTTGATCCTGACCTTACAGGATTACTGGGCTCGCCAGCTCGAGAGATCCTCTAGA
TTTAAGAAGGAGATATACATATG
>glyU
GAGGCAATTCCGACGTCTAAGAAACCATTATTATCATGACATTAACCTATAAAAATAGGCGTATCACGAGGCCCTTTCGTCTTCACCTC
GATGGATCCCGGCAATTGATCCGCTACGTTACTTGCCGCCTCAGGGCAGCAAGCCAAAATGCTGATGGCGAATTAATCAGCAGTCAGCA
GCGTGGCACTTGCTAAGGAGAGCGTAAGGTTTATAATGCCTTACGCATCTCGAAGCGGGCGTAGTTCAATGGTAGAACGAGAGCTTCCC
AAGCTCTATACGAGGGTTCGATTCCCTTCGCCCGCTCCAATTTATCTTCGCCGTAATACTTCCCCACTCGAGAGATCCTCTAGATTTAA
GAAGGAGATATACATATG
>aqpZ
GAGGCAATTCCGACGTCTAAGAAACCATTATTATCATGACATTAACCTATAAAAATAGGCGTATCACGAGGCCCTTTCGTCTTCACCTC
GATGGATCCAACGGTTGATACCGCGAAAACCCACAATTTCAACGCGCTCAAGAATCATATCCCCTCCAGAAATATGTGACTTATCCTTT
ACGCTGGCAGTATAACGGGAAAGGCCTTGCTAATACGTGATGACATCACATTTAAGCATGGCAAATGTCTGAAAAATGAGCGAATAATC
GCGTTATACTCCCTCTTGCTGATAAGTCTGCAAAAACAACAAAATCAAATAATTAGCATGATAAATTATTTTTTATTACGCCTAAAGGA
TTAAATCTTTTGCGTGCACTACACTGTCAATCAGACAAATGACTATATGGGGCAACCCGCTGTTGCCCTAAATCAAATTTACTGGATTT
ATTTATCTGGTGAATGACTCGCCTTCGTTTTAAAATAGCCGCGCATTATTTCCGTCGTCACTTTTTTACGACGACCCTATAAAACGAC
CATATTTTTCACAGGGTCAATTTTTAATTGTGGTGGATATGTTCAGAAAATTAGCAGCTGAATGTTTTGGTACTTTCTGGCTTGTTTTT
GGTGGCTGTGGTAGTGCTGTACTGGCCGCAGGCTTCCCGGAATTAGGCTCGAGAGATCCTCTAGATTTAAGAAGGAGATATACATATG
>glyA
GAGGCAATTCCGACGTCTAAGAAACCATTATTATCATGACATTAACCTATAAAAATAGGCGTATCACGAGGCCCTTTCGTCTTCACCTC
GATGGATCCAGCGATGGTTTGAGCGTCAAGCATATGGTCTTCCTTTTTTTGCATCTTAATTGATGTATCTCAAATGCATCTTATAAAAA
ATAGCCCTGCAATGTAAATGGTTCTTTGGTGTTTTTCAGAAAGAATGTGATGAAGTGAAAAATTTGCATCACAAACCTGAAAAGAAATC

CGTTTCCGGTTGCAAGCTCTTTATTCTCCAAAGCCTTGCGTAGCCTGAAGGTAATCGTTTGCGTAAATTCCTTTGTCAAGACCTGTTAT
CGCACAATGATTCGGTTATACTGTTCGCCGTTGTCCAACAGGACCGCCTATAAAGGCCAAAAATTTTATTGTTAGCTGAGTCAGGAGAT
GCGGATGTTAAAGCGTGAAATGAACATTGCCGATTATGATGCCGAACTGTGGCAGGCTATGGAGCAGGAAAAAGTACGTCAGGAAGAGC
ACATCGAACTGATCGCCTCGAGAGATCCTCTAGATTTAAGAAGGAGATATACATATG
>trkA
GAGGCAATTCCGACGTCTAAGAAACCATTATTATCATGACATTAACCTATAAAAATAGGCGTATCACGAGGCCCTTTCGTCTTCACCTC
GAGTGCGAAACAGGAACACCAGAGCAACCGGGTAAACAAAATCTACCTGGTGCCGAAGAGGGCGACGGCTTCTTTTACGCTAAGCTAAT
CAAAAAGTGATGAGATAACGGGTCGCGACTGATGAAAATTATCATTCTGGGTGCCGGCCAGGTTGGCGGCACACTGGCGGAAAACCTGG
ATCCTCTAGATTTAAGAAGGAGATATACATATG
>ynfM
GAGGCAATTCCGACGTCTAAGAAACCATTATTATCATGACATTAACCTATAAAAATAGGCGTATCACGAGGCCCTTTCGTCTTCACCTC
GAGTGCGAAATATTCAGGCGGGCAGCGGCGCGCCCGAAATGCAGCTCTTCCGCAACAGCAACAAAGTAACGCAGATGACGAAGTTCAAT
ATTCATATTTAAAACATCTTATTTGAGATTATTAATATATTAGACAGAACAATTCGATTTTCCTACCCTATGTATAAGCCTGATCTACA
GGCATATTTAGCAAGGATTTCAAGTGAGCCGTACTACAACTGTTGATGGCGCTCCGGCAAGCGACACTGACAAGCAAAGCATTTCTCAG
CCAAATCAATTTATTAAACGCGGTACGCCGGGATCCTCTAGATTTAAGAAGGAGATATACATATG
>ynfL
GAGGCAATTCCGACGTCTAAGAAACCATTATTATCATGACATTAACCTATAAAAATAGGCGTATCACGAGGCCCTTTCGTCTTCACCTC
GATGGATCCTTGTCAGTGTCGCTTGCCGGAGCGCCATCAACAGTTGTAGTACGGCTCACTTGAAATCCTTGCTAAATATGCCTGTAGAT
CAGGCTTATACATAGGGTAGGAAAATCGAATTGTTCTGTCTAATATATTAATAATCTCAAATAAGATGTTTTAAATATGAATATTGAAC
TTCGTCATCTGCGTTACTTTGTTGCTGTTGCGGAAGAGCTGCATTTCGGGCGCGCCGCTGCCCGCCTGAATATTTCGCACTCGAGAGAT
CCTCTAGATTTAAGAAGGAGATATACATATG
>ynfK
GAGGCAATTCCGACGTCTAAGAAACCATTATTATCATGACATTAACCTATAAAAATAGGCGTATCACGAGGCCCTTTCGTCTTCACCTC
GATGGATCCCTAACCAGGGCACGATGGCAGGCGCTGCACTGGTAAAAGACGCGATGTATAACGGTTCTTTGTTGATTCGTCTGTTGCAG
GGTTAACATTTTTTAACTGTTCTACCAAAATTTGCGCTATCTCAATTTGGGCCAGGAAAGCATAACTTAGACTTTCAAGGTTAATTATT
TTCCTGGTTTATATTTGTGAAGCATAACGGTGGAGTTAGTGATGCTGAAGCGTTTCTTTATTACCGGTACAGACACTTCTGTAGGGAAA
ACGGTGGTTTCCCGCGCATTGCTACAAGCGTTAGCCTCCCACTCGAGAGATCCTCTAGATTTAAGAAGGAGATATACATATG
>ynfE
GAGGCAATTCCGACGTCTAAGAAACCATTATTATCATGACATTAACCTATAAAAATAGGCGTATCACGAGGCCCTTTCGTCTTCACCTC
GAGACCACCAGCGGTAACGTCTCTGCTCCCGCGCAGTCTAGCCAGGATGGCGCGCCTGCCGAACCGCAGTAACACATTATCGACTGAAC
GCCGGATATGACAAATCCGGCGATTTGAACATACAACATAATCCCACCTTATTACTCATACCCTTCTATTGATATGGATTAATAATTCT
TAACCCAAAATGGGTAGACTCCCTCTATTGTTAGCGCGCTAAATATTCAATATATAAACTTTTATATAACGATAAAGAACAGGGAGTGA
GTTATGTCCAAAAATGAACGAATGGTGGGCATCAGCCGCAGAACACTCGTTAAATCTACCGCGATAGGTTCTCTGGCGCTGGGGATCCT
CTAGATTTAAGAAGGAGATATACATATG
>ynfC
GAGGCAATTCCGACGTCTAAGAAACCATTATTATCATGACATTAACCTATAAAAATAGGCGTATCACGAGGCCCTTTCGTCTTCACCTC
GATGGATCCGGCACAGCAGGTTGAGAGTTTCATCATAAGCTCCTTACGCGAAGATAACTTCTTTAAGCATAGCATTTAACGTGTAAAGT
ACTGTATTTGCTACTATGATTGAGAATCATCTCTACTCTCTGGTGACTGTTGTGAAATACAAATTACTACCATGCTTACTCGCGATATT
CCTCACAGGATGTGACCGCACAGCTCGAGAGATCCTCTAGATTTAAGAAGGAGATATACATATG
>ynfB
GAGGCAATTCCGACGTCTAAGAAACCATTATTATCATGACATTAACCTATAAAAATAGGCGTATCACGAGGCCCTTTCGTCTTCACCTC
GAGACCACAACCAGGGCAGAAAGCATCCAATAATTTCACACAGCGCAGTAGCAAAAAATAGTAACGTTGTTTTAATCATCTTTGTCTCT
TATTGACATCATGTATAGTTATAGGGCGACATAATATCATCAATATAAACACCCTCCTGGTACGTTTTGCGTCCGCAGTGGATGGTGTA
GAATCACCTTTAATCATTCATACAGGGAATGAATTATGAAAATCACTCTCAGCAAACGAATCGGCCTGCTCGCTATTCTGCTGCCTTGC
GCACTGGCATTGAGCACAGGATCCATCGAGGTGAAAACGAAAGGGCCTCGTGATACGCCTATTTTTATAGGTTAATGTCATGATAATAA
TGGTTTCTTAGACGCCGCTATGGATCCTCTAGATTTAAGAAGGAGATATACATATG
>ynfA
GAGGCAATTCCGACGTCTAAGAAACCATTATTATCATGACATTAACCTATAAAAATAGGCGTATCACGAGGCCCTTTCGTCTTCACCTC
GATGGATCCGCAGAATAGCGAGCAGGCCGATTCGTTTGCTGAGAGTGATTTTCATAATTCATTCCCTGTATGAATGATTAAAGGTGATT
CTACACCATCCACTGCGGACGCAAAACGTACCAGGAGGGTGTTTATATTGATGATATTATGTCGCCCTATAACTATACATGATGTCAAT
AAGAGACAAAGATGATTAAAACAACGTTACTATTTTTTGCTACTGCGCTGTGTGAAATTATTGGATGCTTTCTGCCCTGGTTGTGGTCT
CGAGAGATCCTCTAGATTTAAGAAGGAGATATACATATG
>treA
GAGGCAATTCCGACGTCTAAGAAACCATTATTATCATGACATTAACCTATAAAAATAGGCGTATCACGAGGCCCTTTCGTCTTCACCTC
GATGGATCCAAACGCTAACGCTGGACGTTAAAACGCAACGTTTCAACCGTCAGGGTTAAGCCACTGTTCGGATGGCATCGTTCTGATGT
CATCCGGCAATTTACTCAGCACTAATGGTTTTTAACTACCTGTTATATTTTTCACCAATTTATGATGATTTTTCTTCGCTAATCACGAA
CCCAACGCCGCTGGCTGGCTAATAAACAGACGTTATTTGATCACTCTATTTCGCCGTTATTTATCCCTTTAATTCCCTTTTCTAAAATG
CCTGACAGTTCGCAGAATGAGATTTCGATCATGCAGCTAGTGCGATCCTGAACTAAGGTTTTCTGATACTTGAATACCGTTTTATTCGG
TTTCGCCAAAGGAGAATGATTGATGAAATCCCCCGCACCTTCTCGCCCGCAAAAAATGGCGTTAATTCCAGCCTGTATCTTTTTGTGTT
TCGCTGCGCTACTCGAGAGATCCTCTAGATTTAAGAAGGAGATATACATATG
>qseB
GAGGCAATTCCGACGTCTAAGAAACCATTATTATCATGACATTAACCTATAAAAATAGGCGTATCACGAGGCCCTTTCGTCTTCACCTC
GAGGAAAAACCGCCCTGCTCTGCTGCCATCACCGGTGCGCTGCACAGGGCCATTACTGCGATTACTGCTGCGAATTTTTCATGTTTAT

TACTCCCTTTAATGTCTGTTTCCGAGCATTTAACAAGATAGTCCTTAACAACTTCTTAAGGGAAAAAAATAAAATTTAGTGCTGTACAG
AGCGCGTTACAACACGGTTTACTGGCAGCAAATACGGTTATCGCAGGGATGAAAAAATGCGAATTTTACTGATAGAAGATGACATGCTG
ATTGGCGACGGCATCAAAACGGGCCTTAGTAAAATGGGTTTTAGCGTCGACTGGGATCCTCTAGATTTAAGAAGGAGATATACATATG
>treB
GAGGCAATTCCGACGTCTAAGAAACCATTATTATCATGACATTAACCTATAAAAATAGGCGTATCACGAGGCCCTTTCGTCTTCACCTC
GATGGATCCAGCGAACCGCAACAAATCATCATCCCCGCCACCCTGTCCTGATCGTTTCCTGAACGATAAATTGTGATCTTCGCTGCGTT
TCGGGAACGTTCCCGTTTTTAAATTTTTCCGCGCAATATATTCTGCAGCCAACCAAAAATGTCATCTGCCATGGGGCTTTATGATGAGC
AAAATAAACCAAACGGATATCGATCGGTTGATTGAACTGGTCGGCGGGCGCGGCAATATTGCGACGGTGAGCCACTGTACTCGAGAGAT
CCTCTAGATTTAAGAAGGAGATATACATATG
>treF
GAGGCAATTCCGACGTCTAAGAAACCATTATTATCATGACATTAACCTATAAAAATAGGCGTATCACGAGGCCCTTTCGTCTTCACCTC
GAGGCACGGCCTCAAGGTTAAAAGATTTATATCCAAGCTTAATGTCGTAATCCATCAACTGTTTCGCGCCAGGAATATAATAATAGGCG
GGTAATTCTGCCGAAGGCGTGTGGCAATAGTCGCATCCTTTTTCGCGGAGAAAGCCTAAAACCTTATTATTTTCACTGACAGCAGATGC
CTGAACATCGGCCTGTTTACTGCGTTTATTATCGTGGTACCACACATAACCAGATAACCCTAAATAGCAAATCGCGACGCCAGCCAGGC
CGATCGCGGTAATACGTGAGACCATTTTCATTATTCTTCCCTCACGGTGGTGAGTTATTGTGACCCAGAAAATTTTCCAGAATGCTGCA
AACGTCTCTGCGAGACAAAGCCATCATAAAAAACAGGGAGATACGCTTTTTGATAGGAATCAATCAATTACATGAATATAAACTATCAA
TAAGATAGCCTGTGCCTATTACCCAGATCAGGAATGGCATATCATGATTGCGGCGAGAGCAGGATTGAATGTTCTGAAAATGAAAAACA
ACACCAAATCGGGGATCCTCTAGATTTAAGAAGGAGATATACATATG
>pgmI
GAGGCAATTCCGACGTCTAAGAAACCATTATTATCATGACATTAACCTATAAAAATAGGCGTATCACGAGGCCCTTTCGTCTTCACCTC
GAGCACCAGTAACGCGATCCAGGCGATACTCAGTATGGGATGACGGCCAACAAATTGCATAATTTCTTGCATGGGGGGTAACAACTCCC
GACGTAGTGATTAAAAAAACCAGGAAAGGAGTATACCTGCGCGGTGCGGCAAATACAGCCAGCGCGTTAACTGGAATGCAATTTTGCGG
GGCGCGACGAAAAAAAAGCGTAAATCTGGCATCGGCTTGCCGCGAGCGGTCGTATATTTTGATCTTCAGAGGCTATTTTATCGATTCAG
CTGTAGTAAAATTACGCAAATTTTGACTCTTGAGTATGAGGTTGTCGCAATGTTGGTTTCTAAAAAACCTATGGTACTGGTGATTCTGG
ATGGCTATGGCTATCGCGAAGAACAGCAGGATAACGCCAGGGGATCCTCTAGATTTAAGAAGGAGATATACATATG
>gst
GAGGCAATTCCGACGTCTAAGAAACCATTATTATCATGACATTAACCTATAAAAATAGGCGTATCACGAGGCCCTTTCGTCTTCACCTC
GAGACTGATGCTGCTGACCGCGCCGAAACTGCACCGCATGACGCAGGATGACGCTGCAGACAAAGCGGCGAAAGCAGCCGTAGCGTAAA
ATTCAGGGAAACTCTTTTACAAGCCGCTAACTTTTCGTTAGCGGCTTTTTTTTTGTTCAGCAACGCACTACCATACTTTAAACCACAGA
CAAAAGGAGTTACCGATGAAATTGTTCTACAAACCGGGTGCCTGCTCTCTCGCTTCCCATATCACCCTGCGTGAGAGCGGAAAGGATGG
ATCCCGGGATCCTCTAGATTTAAGAAGGAGATATACATATG
>ydfE
GAGGCAATTCCGACGTCTAAGAAACCATTATTATCATGACATTAACCTATAAAAATAGGCGTATCACGAGGCCCTTTCGTCTTCACCTC
GAGTGCAGCAACCGAACAGAAAATTCCCGGTAACTGTTACCCGGTCGATAAAGTTATTCACCAGGATAATATCGAAATCCCGGCAGGTC
TTTAAAACAGTTCCGTAATAAATATCCGGTTTCATTCTTATATGCCAGCAATGGCAGGGATTTGTTCATCCTTAAATCTGTCATGAGGT
TAAAACAAATGAGTAAAGTCTTTATTTGCGCCGCTATTCCTGACGAACTGGCAACAAGGGAAGAAGGCGCTGTGGCTGTAGCCACAGCC
ATTGAAGCGGATCCTCTAGATTTAAGAAGGAGATATACATATG
>ydfA
GAGGCAATTCCGACGTCTAAGAAACCATTATTATCATGACATTAACCTATAAAAATAGGCGTATCACGAGGCCCTTTCGTCTTCACCTC
GAGACAGGATACCCAGCTCGCCGAAATGCGAGCTCGAGTAAAAAACTTCAATAAACTCTTTGAAGAATTACTAAAAGCCCGTCAGCGGA
CAAATAAAAGATAACATCATCAATGAGTTATCTTTTACCACATCAATTATGTTAGCTATAGCATACAAAATCACTTGACCGATATGTTA
GTCATGGCTAATCTTGTTTGCATCAACACACCGCACGGTGTTCTCAGCAAACAGTTCCGCTACCCCAGCGTTAAGGGGAAATGAGGTCA
GCATGGATACTATCGATCTTGGCAACAACGAATCTCTGGTGTACGGCGTGTTTCCAAACCAGGACGGCACGTTCACCGCAATGACGGAT
CCTCTAGATTTAAGAAGGAGATATACATATG
>chbA
GAGGCAATTCCGACGTCTAAGAAACCATTATTATCATGACATTAACCTATAAAAATAGGCGTATCACGAGGCCCTTTCGTCTTCACCTC
GATGGATCCCCCTTTGTTGTGGTGGCTAACAAAGCACAAAATGCGATTGATAAAGAAGAGAGCGAAGAAGATATCGCTAACGCCCTGAA
ATTTTAATCGATGCCGGTACGGGTAATCGTGCCGGTAAGAAAAGAGAGGAACGATGTATGATGGATCTCGATAACATTCCCGATACGCA
AACGGAAGCTGAAGAGCTGGAAGAAGTGGTGATGGGGCTGCTCGAGAGATCCTCTAGATTTAAGAAGGAGATATACATATG
>ydfO
GAGGCAATTCCGACGTCTAAGAAACCATTATTATCATGACATTAACCTATAAAAATAGGCGTATCACGAGGCCCTTTCGTCTTCACCTC
GAGGCGTAAGTCGATGGCGTTCGTACTCAATAGTCCCTGGCTGGAGATCTGTCTCGCTGGCCTGCCGCAGTTCTTCAACTTCCCGGCGC
AGCTTTTCGTTCTCAATTTCAGCATCCCTTTCGGCATACCATCTTATAACGGCGGCAGAGTCATAAAGCACCTCATTACCCTTGCCACC
GCCTCGCAGAACGGGCATTCCCTGTTCCTGCCAGTTCTGAATGGTACGGATACTCGCACCGAAAATGTCAGCCAGCTGCTTTTTGTTGA
CTTCCATTGTTCATTCCACGGCCAAAAACAGAGAAAGGAAACGACAGAGGCCCAAAAGCTCGTTTTCAGCACCTGTCGTTTCCTTTCTT
TTCAGGGGGTATTTTAAATAAAAACATTAAGTTACGACGAAGAAGAACGGAAATGCCTTAAACCGGAAAATTTTCATAAATAGCGAAAA
CCCGCGAGGTCGCCGCCCCGTAACCTGTCGGATCACCGGAAAGGACCCGTAAAGTGATAATGATTATCATCTACATATCACAACGTGCG
TGGAGGCCATCAAACCACGTCAAATAATCAATTATGACGCAGGTATCGTATTAATTGATCTGCATCAACTTAACGTAAAAACAACTTCA
GACAATACAAATCAGCGACACTGAATACGGGGCAACCTCATGTCAACGAAGAACAGAACCCGCAGAACAACAACCCGCAACATCCGCTT
TCCTAACCAAATGATTGAACAAATTAACATCGCTCTTGAGCAAAAAGGGTCCGGGAATGGATCCTCTAGATTTAAGAAGGAGATATACA
TATG
>ydfI

GAGGCAATTCCGACGTCTAAGAAACCATTATTATCATGACATTAACCTATAAAAATAGGCGTATCACGAGGCCCTTTCGTCTTCACCTC
GATGGATCCACGCTGCGGAAGATCAACCGCGTGTTGTAAGAGCTGCGCAACCTTCCAGAAGTCTGTAAACCCTTAATCCCTTCTCTTAC
CGGAGAGGGGATTTTTATTCATATAACAAAACATATAGCTTGCCATATTTATATTTAAGGAATTGTTATGGGAAATAATTTGTTATCAG
CAAAAGCGACACTCCCTGTTTATGATCTTAATAACCTGGCTCCAAGAATTGTTCATTTAGGCTTTGGTGCATTTCACCGCTCGAGAGAT
CCTCTAGATTTAAGAAGGAGATATACATATG
>ydfH
GAGGCAATTCCGACGTCTAAGAAACCATTATTATCATGACATTAACCTATAAAAATAGGCGTATCACGAGGCCCTTTCGTCTTCACCTC
GAGCTGGTGGGTGTCAACGCTGCCTGCTCACGTCAATATCAATACCCTGGAAATGATGCCGGTTACCCAAAGCTATGCCGGACTGAATG
TCCACCGTCAGTAATTTTTATACCCGGCGTAACTGCCGGGTTATTGCTTGTCACAAAAAAGTGGTAGACTCATGCAGTTAACTCACTCA
CAAGCAAGAACGAATGACCGTCGAAACGCAACTTAATCCCACACAGCCTGTCAATCAGCAGATTTATCGTATTCTTCGTCGCGACATTG
TCCATTGCCTGATTGCTCCGGATCCTCTAGATTTAAGAAGGAGATATACATATG
>ydfV
GAGGCAATTCCGACGTCTAAGAAACCATTATTATCATGACATTAACCTATAAAAATAGGCGTATCACGAGGCCCTTTCGTCTTCACCTC
GAGGCGCTTCAGAAGGAGTTACACCCATTTTTTCAAGCGCGGCGTAAGAACGCGCTTTAAGTTCATCGTCAATACGCAGGTTAATGCTA
CCCATGTCTTACACCTCTTGTAATTACAAATGTCATTACAAGTATCGCACTACAACATGCTTAGGGCAAGTCACGAAGGAAGTCAGAAA
GTAGTCGTAAGAACGGTGATCACTGTCCGCTTTGTGCGGATCCTCTAGATTTAAGAAGGAGATATACATATG
>ydfZ
GAGGCAATTCCGACGTCTAAGAAACCATTATTATCATGACATTAACCTATAAAAATAGGCGTATCACGAGGCCCTTTCGTCTTCACCTC
GAGAGAACGTGCAATGACGCAACATTTGCAGGAAATCAGCGAATCCGTGCGCCAGATCCGCCAGGAAAACAGCGACTGGTTTAGCGAAG
AGTAATTCATTTCCTCTCATCCCATCCGGGGTGAGAGTCTTTTCCCCCGCCTTATGGCTCATGCATGCATCAAAAAAGATGTGAGCTTG
ATCAAAAACAAAAAATATTTCACTCGACAGGAGTATTTATATTGCGCCCGTTACGTGGGCTTCGACTGTAAATCAGAAAGGAGAAAACA
CCTATGACGACCTACGATCGTAACCGTAACGCAATCACCACTGGCAGCCGTGTTATGGTTAGCGGCACCGGTCACACTGGCAAGATCGG
ATCCTCTAGATTTAAGAAGGAGATATACATATG
>serW
GAGGCAATTCCGACGTCTAAGAAACCATTATTATCATGACATTAACCTATAAAAATAGGCGTATCACGAGGCCCTTTCGTCTTCACCTC
GATGGATCCCTGACCCCGTACGACCTGAGCAAAGGCCGCATTGTCTTCCGTAGTCGCTGATTGTTTTACCGCCTGATGGGCGAAGAGAA
AGAACGAGTAAAAGGTCGGTTTAACCGGCCTTTTTATTTTGTGATATGTATGAAGTACTTTGGAAGTATAAGTCCGTAACTTGTCTCGA
TGTAGATATACTGCACAACTTGTTTACAGTATGAGCAAACAATAATTATCATTAAAATTCTTCTTGACCGTTTTACTCCAACTCCCTAT
AGTAGCGCCCCGTTGCCACCCATGAGGTTTGGTAGCAAAACAATTTGGTGAGGTGTCCGAGTGGCTGAAGGAGCACGCCTGGAAAGTGT
GTATACGGCAACGTATCGGGGGTTCGAATCCCTCGAGAGATCCTCTAGATTTAAGAAGGAGATATACATATG
>pdhR
GAGGCAATTCCGACGTCTAAGAAACCATTATTATCATGACATTAACCTATAAAAATAGGCGTATCACGAGGCCCTTTCGTCTTCACCTC
GAGTAACCCGGTCCCTATCGCGCCACCCAGCGCGATAAGCTGAATATGGCGGTTTTTAAGGCCGCGCTTTAGCTGCTCGCCGTGCTGTT
GACCTTCCATCATGAAACCTCGTGCGGTGGTTGTTTTTTTGATCTACGCAGTGATGCGTGTGTAAGTTTGCAATTCCGTTTGTTGTATT
AATTTGTTTACATCAAAGAAGTTTGAATTGTTACAAAAAGACTTCCGTCAGATCAAGAATAATGGTATGCGGCAGCGAATGCACCCGCT
TTATGCATGGTTGAAGATGAGTTGCTTAAAAAGAAACCGTTTGTAAAGCTCAGCCTCAACCCCTCTCAATATGTAGAATGAATTTAAAT
TCGTTTTAATTGAATTAAAAATCACAAAATTGGTAAGTGAATCGGTTCAATTCGGATTTTTATAGTTTAATAATCGTTAAAAAACTCCT
TTCCTACGTAAAGTCTACATTTGTGCATAGTTACAACTTTGAAACGTTATATATGTCAAGTTGTTAAAATGTGCACAGTTTCATGATTT
CAATCAAAACCTGTATGGACATAAGGTGAATACTTTGTTACTTTAGCGTCACAGACATGAAATTGGTAAGACCAATTGACTTCGGCAAG
TGGCTTAAGACAGGAACTCATGGCCTACAGCAAAATCCGCCAACGGATCCTCTAGATTTAAGAAGGAGATATACATATG
>ydgC
GAGGCAATTCCGACGTCTAAGAAACCATTATTATCATGACATTAACCTATAAAAATAGGCGTATCACGAGGCCCTTTCGTCTTCACCTC
GATGGATCCTACGCGGCAATCAGTGAACCGACTTCCGCATCATCTTCCACAAATACGATAGTGTTCATAACATTCACGGTAGATATAAA
AACGTCACAATACACCGCGCTGTTTTTACTTACTATCAATCGTTTCTAAACAAGTTTTCATCAGGTATTCTACTGATGACTTGTTATGG
AAGTGTTAAGGTAAAAAGATGGGGCTGGTAATCAAAGCCGCGCTCGGTGCGCTGGTAGTGCTGTTGATTGGTGTTTTAGCAAAAACGAA
AAATTATTATATCGCCGGGCTGATCTCGAGAGATCCTCTAGATTTAAGAAGGAGATATACATATG
>pqiA
GAGGCAATTCCGACGTCTAAGAAACCATTATTATCATGACATTAACCTATAAAAATAGGCGTATCACGAGGCCCTTTCGTCTTCACCTC
GAGGCAGAGCAGGAGCTGGAGCAAGCCTTTGAACGCTGGGAGTATCTTGAAGCGTTAAAAAATGGTGGCTGATCGCAAAATAGTCGATT
TAAACAGAGACAGCTGACAACGGTAAAATTGCCTGATGAACTGGGTTTATCAGGCATGAAAACCAACACTTTGTTAGTCATCTGTAAAA
TAGCGCATCATTAAGGAGTACCAATGTGCGAACATCATCATGCCGCGAAGCACATCCTGTGCTCGCAGTGTGACATGCTGGTGGCGTTA
CCGCGCCTTGAGCATGGTCAGAAAGCGGCATGTGGATCCTCTAGATTTAAGAAGGAGATATACATATG
>yhhF
GAGGCAATTCCGACGTCTAAGAAACCATTATTATCATGACATTAACCTATAAAAATAGGCGTATCACGAGGCCCTTTCGTCTTCACCTC
GAGCTTTTTCCGGGGTCTGCTCTTTTTGACCAAAGCCCAGCCAGGAAAAAAAGCCACGTTTTTTTTCTTTCGCCATTTGCGACTACACT
CCTCGCTGTTGCTTCATGGCACAGCGTTAACGCTATGTACATAGCAGCTAAAAAAATGATGAAATAGTCTATCACTTAACTTAATTCAC
ATCACCGCCTGCAAGTATGTGTTATCTGGCGGATTGAGCAATTTATCATGAAAAAACCGAATCATTCCGGCAGCGGCCAAATCCGCATT
ATTGGCGGGCAGTGGCGAGGCCGTAAACTCCCGGTTCCGGATCCTCTAGATTTAAGAAGGAGATATACATATG
>flgB
GAGGCAATTCCGACGTCTAAGAAACCATTATTATCATGACATTAACCTATAAAAATAGGCGTATCACGAGGCCCTTTCGTCTTCACCTC
GAGTGTGCAATTGCGATGTGAGATTGCTCGCCGTACTTAACGGACTGAACAGTATCGCGATGATCGCCACGCTACGTTTTATTATCAGC
ATTTTCGCCCCCAGCCATTTCTACAACGTGAATTGTACCTGTCCGCAATGACCATCAACGGCATAAATAGCGACCCATTTTGCGTTTAT
TCCGCCGATAACGCGCGCGTAAAGGCATTTAAGCTGATGGCAGAATTTTGATACCTGCGGAGGAGATATGCTCGATAAGCTCGACGCCG

```
CCTTACGTTTTCAACAAGAGGCGCTCAATCTGCGCGCCCAGCGTCAGGAAGTGCTGGGGATCCTCTAGATTTAAGAAGGAGATATACAT
ATG
>yhhN
GAGGCAATTCCGACGTCTAAGAAACCATTATTATCATGACATTAACCTATAAAAATAGGCGTATCACGAGGCCCTTTCGTCTTCACCTC
GAGCCGTCGCTGTTGCATAAAACGAAACGATGCGGCGACGACAATTAAGCCAATGATAACAATAAAGAAAAGAGGTGGTTTGCTCATCT
TTATCCCTCATCGGAAAATGCGGAAATAAGCATACCCTGCCAGTTATGGTGTTGTCATCCGTCCACCCTCGCCACTAAACTGGAAGCAA
GACCGTAGGCATTCCGCTTACGAAAAAATAACGAATTCAAGGAACTAAGATGCTTTGGTCGTTTATCGCTGTCTGTCTTTCCGCATGGC
TATCTGTGGATGCATCGTATCGTGGGCCAACCTGGGATCCTCTAGATTTAAGAAGGAGATATACATATG
>yhhM
GAGGCAATTCCGACGTCTAAGAAACCATTATTATCATGACATTAACCTATAAAAATAGGCGTATCACGAGGCCCTTTCGTCTTCACCTC
GATGGATCCAGCCATGCGGAAAGACAGACAGCGATAAACGACCAAAGCATCTTAGTTCCTTGAATTCGTTATTTTTTCGTAAGCGGAAT
GCCTACGGTCTTGCTTCCAGTTTAGTGGCGAGGGTGGACGGATGACAACACCATAACTGGCAGGGTATGCTTATTTCCGCATTTTCCGA
TGAGGGATAAAGATGAGCAAACCACCTCTTTTCTTTATTGTTATCATTGGCTTAATTGTCGTCGCCGCATCGTTTCGTTTTATGCAACA
GCGACGGCTCGAGAGATCCTCTAGATTTAAGAAGGAGATATACATATG
>minC
GAGGCAATTCCGACGTCTAAGAAACCATTATTATCATGACATTAACCTATAAAAATAGGCGTATCACGAGGCCCTTTCGTCTTCACCTC
GATAGATCTCCCCACACCTGACAATCCGATATAAAAAATACGCATCATATTCATAGCACATCCTGATAATCCAACCGTACAACACTTAT
GCTCATTTCATCATCAATCCGTAATCAAAACCGCTGACCTGGACGAGTTACGATGACCTGACAGAAGTATCAGACGTCTTCTGCGGTTT
CTATGATAGAGCAATTCTGTGGCGGATATTAAGATAACGGGTATCTGTTCTTTAATGATGCCGCAAAAAATGGAAAGTTAAGAAAGCCT
GGCGTGGTGAAAAAATACACTTGCCTCAATACTGGATATTTAAACAGGTCTGGAGATGCTGTCACGATTAGGGTGCGGAAAAAGGAAGC
GTAAATAAGAACAATCAAAAGTACGACGGCAATGGGTTGATTGACAAGGGTATTTTTTAAGCTATGAATCAGCGCCATTTATCACAGAA
TAGACTTTTACTCTGAATAAATGGGAGGGTGACTTGCCTCAATATAATCCAGACTATAACATGCCTTATAGTCTTCGGAACATCATCGC
GCGCTGGCGATGATTAATAGCTAATTGAGTAAGGCCAGGATGTCAAACACGCCAATCGAGCTTAAAGGCAGTAGCTTCACTTTATCTGT
GGTTCATCTGCATGAGGCAGAACCTAAGGTTATCCATCAGGCGCTGGCTCGAGAGATCCTCTAGATTTAAGAAGGAGATATACATATG
>flgA
GAGGCAATTCCGACGTCTAAGAAACCATTATTATCATGACATTAACCTATAAAAATAGGCGTATCACGAGGCCCTTTCGTCTTCACCTC
GATGGATCCCCAGCACTTCCTGACGCTGGGCGCGCAGATTGAGCGCCTCTTGTTGAAAACGTAAGGCGGCGTCGAGCTTATCGAGCATA
TCTCCTCCGCAGGTATCAAAATTCTGCCATCAGCTTAAATGCCTTTACGCGCGCGTTATCGGCGGAATAAACGCAAAATGGGTCGCTAT
TTATGCCGTTGATGGTCATTGCGGACAGGTACAATTCACGTTGTAGAAATGGCTGGGGGCGAAAATGCTGATAATAAAACGTAGCGTGG
CGATCATCGCGATACTGTTCAGTCCGTTAAGTACGGCGAGCAATCTCACATCGCAATTGCACACTCGAGAGATCCTCTAGATTTAAGAA
GGAGATATACATATG
>yhhW
GAGGCAATTCCGACGTCTAAGAAACCATTATTATCATGACATTAACCTATAAAAATAGGCGTATCACGAGGCCCTTTCGTCTTCACCTC
GATGGATCCGAGCAAGCCTCTCCCTCCACAGTAACTCTCGCTAAGTAAGTTTGATGGCCCCTCGAATAGTTCAATTTTTTTGAACAGAG
GGGTCAATTTTCACCCTCTATCATCCCAGGCAGATCGGGTCCACACTAAGCCCATCGAAATCATTCAGGGGGCGAATACAAATGATCTA
CTTACGCAAAGCAAATGAACGTGGTCATGCAAATCATGGCTGGCTGGACTCCTGGCATACTTTCTCTTTTGCCAACTATTACGATCCGA
CTCGAGAGATCCTCTAGATTTAAGAAGGAGATATACATATG
>yhhT
GAGGCAATTCCGACGTCTAAGAAACCATTATTATCATGACATTAACCTATAAAAATAGGCGTATCACGAGGCCCTTTCGTCTTCACCTC
GAGCAAATTCAGGCGCAATCCGTTTAGCGCGGGTTCGGCTACGGGTTCGGGCATGAGGATCACCACATTTTTACAACAGTGTTTCATTT
TTACCACGTGCTGACGTGAAAATCAGCAGTAAGAATCAGAATATTGCTGGCGTGACTCCCGCTACACTTAATACAAAAAGTCACAAGGA
AGCCCCAATGGAAACCGGATCCTCTAGATTTAAGAAGGAGATATACATATG
>adhE
GAGGCAATTCCGACGTCTAAGAAACCATTATTATCATGACATTAACCTATAAAAATAGGCGTATCACGAGGCCCTTTCGTCTTCACCTC
GATGGATCCAGCGCAAATAACCCGATGAAAAATTTGAAGTAAACGGGAAAATCAAAAAAGGTCTGAATCACGGTTAGCTCCGAAGCAAA
AGCCGGATAATGTTAGCCATAAATAAGGTTGAAAAGACGCGCTGACAATACGCCTTTTGACAGCATTTTTCACCTCCTAACTACTTAAA
ATTGCTATCATTCGTTATTGTTATCTAGTTGTGCAAAACATGCTAATGTAGCCACCAAATCATACTACAATTTATTAACTGTTAGCTAT
AATGGCGAAAAGCGATGCTGAAAGGTGTCAGCTTTGCAAAAATTTGATTTGGATCACGTAATCAGTACCCAGAAGTGAGTAATCTTGCT
TACGCCACCTGGAAGTGACGCATTAGAGATAATAACTCTAATGTTTAAACTCTTTTAGTAAATCACAGTGAGTGTGAGCGCGAGTAAGC
TTTTGATTTTCATAGGTTAAGCAAATCATCACCGCACTGACTATACTCTCGTATTCGAGCAGATGATTTACTAAAAAAGTTTAACATTA
TCAGGAGAGCATTATGGCTGTTACTAATGTCGCTGAACTTAACGCACTCGTAGAGCGTGTAAAAAAAGCCCAGCGTGAATATGCCTCGA
GAGATCCTCTAGATTTAAGAAGGAGATATACATATG
>yjdF
GAGGCAATTCCGACGTCTAAGAAACCATTATTATCATGACATTAACCTATAAAAATAGGCGTATCACGAGGCCCTTTCGTCTTCACCTC
GATGGATCCGAAGATTTCCCGGCGTTTATCCTGGTCGATGACAAAGGTAACGACTTCTTCCAGCAAATCGTCAACAAACAGTGCGCGAA
CTGCACTAAGTAACCTCTTCGGCCCAGCGCCTGGCAGCATGCTGCCAGGTGATCCCCCTGGCCACCTCTTTTGCGATTGTAATTTCACG
CTTGCTGGTGAATAGTCAGTATTTTCCCTGATTTGCGAACTCACCATGACCCGCACACTCAAGCCGTTAATTCTTAACACCAGCGCACT
GACGCTAACGTTAATCCTGATTTATACCGGCATTTCGGCCCATGACAAACTCTCGAGAGATCCTCTAGATTTAAGAAGGAGATATACAT
ATG
>yjdI
GAGGCAATTCCGACGTCTAAGAAACCATTATTATCATGACATTAACCTATAAAAATAGGCGTATCACGAGGCCCTTTCGTCTTCACCTC
GAGCACCGAGAACAGTACCGCACTGACCATTAAGATCACTGTGGTACTCAATTTCATCGGACGTTTGCGTAACATGCGGTAGGGCAATG
AATGTCTCATCAGCTTCCTTGTGTGACAAATTTCTTAAGCATTATCTCTGATGAGGCGGGTAATTCAAAGGGAGTAAGAATGATTGGCT
```

ATATAGGGGAAGAGACTCTGGCAACGGAAACTGCCAGTGCTGTATGAAGATTCCGGGGCTATGCTTATAGCGATAATCATACTGATGAG
AGAGGGAAGGTCATGGATCAGGCGCTACTGGACGGGGGTTATCGCTGTTATACCGGCGAAAAGATCGATGTCTATTTCAACACTGCGAT
ATGTCAGCATTCTGGCGGATCCTCTAGATTTAAGAAGGAGATATACATATG
>slyB
GAGGCAATTCCGACGTCTAAGAAACCATTATTATCATGACATTAACCTATAAAAATAGGCGTATCACGAGGCCCTTTCGTCTTCACCTC
GAGCGACCCGGTGTTCATCAATTGTCGCCAACACAACATCAACACCATCAAGGCTGGTGCCTGACATAACGCCAATAAAGCGGCCCGAT
TTCATAGTTCATCCTTTTTCAATCTGACGTTTGCGCACCACTCAAACATAAACTTTTCGTGAATACCATGCGGAATGACCGATTTTTAC
CGTTGGTAGTAAAACATTATCTTCAAATCAATAATCATCATGAATGTTTTGTTTATAATTGGTTGATCCTACTTTCATTATGATTTGCT
CATATTTGGTAGAACATGTAACCATGGATTCACATATGCCATATACTTTGACCATGAGGGATGCTTGCGTGGCGTTTCATGGTGAACAG
GAGATTTTTCAATGATTAAACGCGTATTGGTTGTTTCAATGGTAGGTCTGTCTCTTGTCGGTTGTGTTAATAACGACACCCTGTCAGGG
GATGTTTATACCGCGGATCCTCTAGATTTAAGAAGGAGATATACATATG
>yjdK
GAGGCAATTCCGACGTCTAAGAAACCATTATTATCATGACATTAACCTATAAAAATAGGCGTATCACGAGGCCCTTTCGTCTTCACCTC
GAGAAATACGCGAAGGCCACAATAAATTTTACATTAATGACAAACAAGGCAAGCAAATCGCTGAAATTGTCTTTGTGCCGACCGGAGAG
AATTTAGCGATTATCGAACATACCGATGTCGATGAAAGCCTGAAAGGGCAAGGGATTGGTAAACAGCTGGTTGCGAAAGTCGTGGAAAA
AATGCGTCGGGAAAAACGAAAAATTATCCCATTATGCCCATTTGCGAAACATGAATTTGATAAAACGCGGGAGTATGATGATATTCGCA
GTTGATGGGAGAGTACAGAGTCACGATATTTTTCATTCTCTCCGCGATGTGATGCAGGAGAGCATCTGAAGGGTAGGGGGATGCACGGA
TCCTCTAGATTTAAGAAGGAGATATACATATG
>yhhZ
GAGGCAATTCCGACGTCTAAGAAACCATTATTATCATGACATTAACCTATAAAAATAGGCGTATCACGAGGCCCTTTCGTCTTCACCTC
GAGCGGCTTTGAAATTGAAGGGACTGGTAAGAAGTACGCATTGCGTAATGGTGAATATGTCGATGCATATTATATGGCGCGGGTGAAGT
AAGATAGTGCCCTTTTTCTGAGATGGAAAAAGGGTGTCATTCAAAATCGACATACCTTCCTTTAAGGTATTTATTTGCCCAATACATAT
ATTGAGATTTAACTCATAATTACTTCCTAAAGTGTAATATTTTATTTTTAATATATACGCCTACAATTTCCTGGAGTAAATAAATAAC
AATTAACAAGCATAATATTGCCATTGATAAAATAGCATGCCATAAAAGGACTTTTCAGGGATGAGTAATATTGTTTACCTGACAGTAAC
GGGAGAACAACAAGGAAGCATCTCCGCAGGTTGTGGGACTTCGGATCCTCTAGATTTAAGAAGGAGATATACATATG
>yhhY
GAGGCAATTCCGACGTCTAAGAAACCATTATTATCATGACATTAACCTATAAAAATAGGCGTATCACGAGGCCCTTTCGTCTTCACCTC
GAGATGCCAGCTATCCTTGCGGTTAAGTACATACGGCAGATGGTAACGGGTGGTGCTTTTGCCGAAGCCAATAAAGGCGCAGTTGATGA
CCATGATTTCGTCCTTTTTAAGGTGGTTATTTACACCTTAGCGCAAAGCGGACGTGGTTCCTACTGGAGTGCGCATAACGAACACAAGC
ACTCCCGTGGATAAATTGAGAACGAAAGATCAAAAAAAAAGCCAGCACCCGGCTGGCTAAGTAATACTGGAAGCAATGTGAGCAATGTC
GTGCTTTCAGGTTCTCCGCGAGGGTCTTCCTGATCGCGAGACAATAATAATCATTCTCATTCGCACTTGTCCAACACTTTTTGCAAAAA
AATGCATTTGACTCGCATTTGAAAGTCAATGATGTTGAAAGGGACATTTACCCCAAAGAGGACAAAGGAATGAGTGAGATAGTAATACG
CCACGCAGAAACACGGGATTACGAGGCCATCAGGCAGATTCACGCCCAGCCGGAGGTGTATTGCAGGATCCTCTAGATTTAAGAAGGAG
ATATACATATG
>yfdL
GAGGCAATTCCGACGTCTAAGAAACCATTATTATCATGACATTAACCTATAAAAATAGGCGTATCACGAGGCCCTTTCGTCTTCACCTC
GATGGATCCTGGTAATCCCCCGTACCCGTGGTAACCCCCAGACCGGCACGCCTGCCACCGATCTGGATGATGACTACTTTGACATGTTG
CAGGAGGAGCTTTGCAGCGTTGTGGAGGCCTCCGGTGCCAGCCTGGAGAAGGGGCGGCATGACCAGCTGCTTACCGCGCTTCGTGCGCT
GCTGTTCTCGAGAGATCCTCTAGATTTAAGAAGGAGATATACATATG
>yhfK
GAGGCAATTCCGACGTCTAAGAAACCATTATTATCATGACATTAACCTATAAAAATAGGCGTATCACGAGGCCCTTTCGTCTTCACCTC
GAGCTCGTGAAACCGTGGGACGCATTCTGAAGATGCTGGAAGATCAGAACCTGATCTCCGCACACGGTAAAACCATCGTCGTTTACGGC
ACTCGTTAATCCCGTCGGAGTGGCGCGTTACCTGGTAGCGCGCCATTTTGTTTCCCCCGATGTGGCGCAGACTGATTTATCACCCCGAT
ATCAACTATGCACTTCGACAAACGCTGGTGCTATGTTTGCCCGTGGCCGTTGGGTTAATGGGATCCTCTAGATTTAAGAAGGAGATATA
CATATG
>yhfL
GAGGCAATTCCGACGTCTAAGAAACCATTATTATCATGACATTAACCTATAAAAATAGGCGTATCACGAGGCCCTTTCGTCTTCACCTC
GAGAGAACTGGCGCAGCAAATGAACAGTCCATCGCTAATTATTATTGGTCGGGTTGTTGGCCTGCGCGATAAACTGAACTGGTTCTCCA
ACCATTAATTTAACCCGGGCCAGAGAATTCTGGCCTTCTTAATGACTCTTTTTATATATTCAGCAAATAAAACATAGCCCCTTAATAAA
TATTATCTGCTAATGGGTTATATTCACCGTTACAGCGACTTACAAAATTAAGTCATTCGCTCACTGACGCCAATTAAATATTCCTGTAT
CGTTCTTAACGAGCAGTCTGCTTATATTCATATAATCAATGAATATTAATTAATATATAATACATAGGGAATGTAAATGAACAAATTTA
TTAAAGTTGCACTGGTAGGTGCAGGATCCTCTAGATTTAAGAAGGAGATATACATATG
>yhfA
GAGGCAATTCCGACGTCTAAGAAACCATTATTATCATGACATTAACCTATAAAAATAGGCGTATCACGAGGCCCTTTCGTCTTCACCTC
GATGGATCCAAGCGTGCTCTTGGATGGGTACTTATGAATGTGGCAATGAGACAAGAACCATTCGAGAGTCGGGTCTGTTTGCGGTTTGC
CAAGCACCATGCGCGGTTATCCTCTGTTATAAGCTTTCTCCAGAGCCAGATAACGCCGCTGTCTCTGGATTGCCGAAATATGCTTCCCG
CTACCTGGGAAGGGGCTATCAACTGTACTGCACGGTAATGTGACGTCCTTTGCATACATGCAGTACATCAATGTATTACTGTAGCATCC
TGACTGTTTTAGCATAGCTTTCGCTTTGTGTCTCCTGGTGTCTCGCTTCAGCATGACCCAGGTCGCCTTCCGTTGCGCGATTTGGTTAG
TACGCGTACTCTGTCAGGAAAATTGACGCAGTGGAGTAGCAAAAATGCAAGCGCGAGTGAAGTGGGTCGAAGGGTTAACTTTTCTGGGC
GAATCCGCCTCTGGTCATCAGATTTTAATGGACGGCAACTCAGGCTCGAGAGATCCTCTAGATTTAAGAAGGAGATATACATATG
>yfdG
GAGGCAATTCCGACGTCTAAGAAACCATTATTATCATGACATTAACCTATAAAAATAGGCGTATCACGAGGCCCTTTCGTCTTCACCTC
GAGTGCGTGGGATTTACAACCATGCTCAGTATCTCGATAAACGCAGAGAAATGATGCAATGGTGGGCGGACTGGCTTGATGAGAAGGTG

GAGTGAGCGACCTTAACAACTATCGAATAGCACAAAGTCTTGCAATCCAGTGCAAAGCTTTGTGTGTATAAGTTTTGTCTCATCAACCA
CAGCAAGTATCGATCGATTAAGACTTGGATGATAGACTTCATTCCTTTGATTATTAGCTGATAGAAGAAATGTTAAAGCTATTTGCAAA
GTACACCTCTATTGGTGTGCTGAACACCCTTATACACTGGGTGGTTTTTGGTGGGATCCCGGGATCCTCTAGATTTAAGAAGGAGATAT
ACATATG
>sgcC
GAGGCAATTCCGACGTCTAAGAAACCATTATTATCATGACATTAACCTATAAAAATAGGCGTATCACGAGGCCCTTTCGTCTTCACCTC
GATAGATCTGCACATTTCCCCGTTGAGCCTGATTCAGGCACTACACAAGAGGCACATCCATTATGAAAAAGATCCTTGTGGCATGCGGT
ACCGGCATGTCGACTTCAACCATGATTGCACACAAGCTGCAAGAGTTCCTGACTGAACAGGGCATTTCAGCCACTACCGCCCAATGCTG
TCTGAATGAGATCCCTCTTAACTGTAACGGTATGGACCTTATTGTAACCTCCATGCGAACCAACAGTGATTACGGCATCCCTACGCTTA
ATGGCGCCGCCCTGCTCACAGGGATCAACGACGACGCGTTAAAACAACAAATCAAGGCTCTGTTAACTCAATAAAAGGAGGCATTATGT
TTGATTACATCCTGTCTCTCGGCGGTACCGTTTTCGTCCCTATCATCATGATCGTGATTGGCCTTATCTTCCGCACTCGAGAGATCCTC
TAGATTTAAGAAGGAGATATACATATG
>yfdC
GAGGCAATTCCGACGTCTAAGAAACCATTATTATCATGACATTAACCTATAAAAATAGGCGTATCACGAGGCCCTTTCGTCTTCACCTC
GAGCAGAACGCCCTTGCTGATCTGTACCGGAACTCGCACACCCCACCAGAAGCGTAGTTCCCAGAGCAAGCGCCGACAGGCGAAGCTTC
ATAAATGTCTCCCTGTTTTTTTATGGCTTATGCAGTTTGCCATCCATGACGGAACGATACCGTATCCGCCTGTTTAGGTGTGGGCGATT
GTAACAGCACGTCAACTGATGTCCAGACGCCCTGATTTGCTGATTTGATCATAGCCTGGTAATCGCCGCCCTGTAGGCTACTTGATTCT
ATAGAAACAGAAAAAGGCAAACGCAGCCTTTTCTACATTTTCAGAGTAACTCCCGTCAGTTGCGAGCAAAAAAGCCGCTACGCTTTAGC
TATACGTGCTAATTCAAGAGAAGAGACCATGGACAACGACAAAATTGATCAACACAGCGACGAAATTGAAGTTGAGAGCGAAGAAAAAG
AGCGCGGCGGATCCTCTAGATTTAAGAAGGAGATATACATATG
>yhfX
GAGGCAATTCCGACGTCTAAGAAACCATTATTATCATGACATTAACCTATAAAAATAGGCGTATCACGAGGCCCTTTCGTCTTCACCTC
GATGGATCCGCTGGCCAACCTTTATGGATTATGGATGGCTGCTAACGAAGAGGTTTGAAGCTCCCGACAAATCCACGTCGCACAACCTA
AATATTCAAAAAAATGAATATTTAATTCAAGGAACAGAAACGAGGCAACAAGATGTTTGTAGAAGCATTGAAACGCCAGAACCCGGCGC
TGATTTCCGCCGCACTAAGCCTGTGGCAGCAGGGCAAGATCGCCCCGGACAGCTGGGTGATCGACTCGAGAGATCCTCTAGATTTAAGA
AGGAGATATACATATG
>yhfZ
GAGGCAATTCCGACGTCTAAGAAACCATTATTATCATGACATTAACCTATAAAAATAGGCGTATCACGAGGCCCTTTCGTCTTCACCTC
GATGGATCCAAAGCGGTGTACGAAGCGATTGGTTTTGTGGCGAAGCCGTAATTCACTATTGCTGGCAAATTGCGCTTTGTTCATGCCGG
ATGCGGCGTGAACGCCTTATCCGGCCTACAAAGTAATGCACATATTGCAAGGAAATGTAGGCCTGATAAGCGTAGCGCATCAGGCAATT
TTGCTTTTATCATCACTCTGAAACCGGGGAAACCCGGTTTTTTTATCCTCATTTGCAATAATCCGAAAAAATGTGAAGTGCCTCGCCGT
TTCCACATCTCGCTGTTGCGCCTTATCTTCAAATTCCTGAAGATATAAATATTCAATTTACTGAAAATTTATTATGCGTCGAACGTTTA
TCAAAAAAGAAGGCGTCGTCATCACGACGCTGGCCCGTTATTTGTTGGGTGAAAAGTGCGGTAATCGATTGAAAACCATAGATGAGCTG
GCAAATGAATGCCGTTCATCCGTTGGCCTGACGCAGGCCGCGTTGAAAACGCTGGAATCAAGCGGAGCGATACGGATTGAACGCCGTGG
GCGCAATGGCAGTTATCTGGTCGAGATGGATAACAAAGCATTGCTGACTCATGTGGATATCAACAACGTGGTATGTGCAATGCCCTTGC
CCTATACCCGTCTCGAGAGATCCTCTAGATTTAAGAAGGAGATATACATATG
>yfdZ
GAGGCAATTCCGACGTCTAAGAAACCATTATTATCATGACATTAACCTATAAAAATAGGCGTATCACGAGGCCCTTTCGTCTTCACCTC
GATGGATCCACAGGCGGATACGGATGAGAAAGAACAGGCAGATAAGCATTAACGCTGCCCGATCGAAGACCGCCAGCAGCATGTTGAAT
ATTTCGTGCACGGGTGACTCTGAATAAGGATTAAACCTCTATGATAGGTAACCTGAAGGCTGATGACCAGCAGGCCGTTTTTGAGGAAA
AACCACAATTTTAAAAGGGGATTGATAGATAAATGGTGTATTAGCAATGCATCTCTTGATAATGCACTTCTGGTTAGTAAATTTATTGA
AATTATTATGTTAATAAAATGAGTGGTAATTTTCTTGTTAGCGCAGATAAATTTCCTGTTTTTAAAGCAGAGTTCCTCATTTAACGTCA
TTAATAAAATGTTCTAAAACGCTGGACAGACAGAAATTAATCAGGCTATTTTCAAAACAGCCACCATCGTGGCAGCGTCGCTCGGACGG
TCCGGGCGCTAACGTTAATCTGAGGAAATTATGGCTGACACTCGCCCTGAACGTCGCTTTACGCGCATTGATCGTCTCCCGCCCTACGT
TTTTAACATCACCGCTGAACTGAAAATGGCTGCGCCTCGAGGCGCAGCCATTTTCAGTTCCGCTCGAGAGATCCTCTAGATTTAAGAAG
GAGATATACATATG
>yfdU
GAGGCAATTCCGACGTCTAAGAAACCATTATTATCATGACATTAACCTATAAAAATAGGCGTATCACGAGGCCCTTTCGTCTTCACCTC
GATGGATCCTGTGACGGATATGGCACGCCATGCCCAGGCGGAAGGCATTCGTTATATTGGTTTTCGTCATGAGCAGTCGGCAGGCTATG
CCGCTGCGGCAAGCGGTTTTCTTACCCAAAAACCGGGGATCTGCCTGACAGTTTCTGCGCCAGGATTCCTCAATGGTTTGACCGCATTG
GCCAACGCAACGGTAAATCTCGAGAGATCCTCTAGATTTAAGAAGGAGATATACATATG
>yhfP
GAGGCAATTCCGACGTCTAAGAAACCATTATTATCATGACATTAACCTATAAAAATAGGCGTATCACGAGGCCCTTTCGTCTTCACCTC
GAGGGCGATGGATATGGCAAAAGAGATGAACGCGGGTTATACGCTGATTTCCGCGGCCCACGCGGGCTATCTCACGCCACCTAATGTTA
TCTGGGGACGGCTGGCAGAGAACCTAAGCGAACTGTGTGAGTACGCGGAAAACATCGGCATGGATCTAATCCTCGAACCGTTAACGCCG
TATGAATCGAACGTCGTGTGTAATGCTAATGATGTGCTTCATGCGCTGGGATCCTCTAGATTTAAGAAGGAGATATACATATG
>yfdV
GAGGCAATTCCGACGTCTAAGAAACCATTATTATCATGACATTAACCTATAAAAATAGGCGTATCACGAGGCCCTTTCGTCTTCACCTC
GATGGATCCCCCAAAACAAGTCGCTGGTAATTAAATACTGACATCGTCTCCTCTTGCCAACTCATTATGCGATGGTGAGGGGACCCTTA
TCGGAAATACTATTATGCTAACATTTTTTATTGGCGATTTATTGCCTATTATCGTAATCATGCTGTTGGGTTATTTTAGCGGCAGACGA
GAAACATCTCGAGAGATCCTCTAGATTTAAGAAGGAGATATACATATG
>mcrB

GAGGCAATTCCGACGTCTAAGAAACCATTATTATCATGACATTAACCTATAAAAATAGGCGTATCACGAGGCCCTTTCGTCTTCACCTC
GATGGATCCACAAAAGCAGGTGCAGGCGTTTATTGGAGTGATTGCCGGTAAACAGAAAGTCGCGTAACAGCGTTTACTCCCGGTTAACC
ACCGGGAGCCTTCCACTGACTCAATAGAAACTTTCCCCCTCAGTAAATATTTACCAGTCTGATTTTGCAGTAAAAATCTATTGTTTCAG
TACGTTGCGAAAGCGATAATAGAGGCTTAGCAATGAGGAAGGCATATCTTATGGAATCTATTCAACCCTGGATTGAAAAATTTATTAAG
CAAGCACAGCAACAACGTTCGCCTCGAGAGATCCTCTAGATTTAAGAAGGAGATATACATATG
>gsk
GAGGCAATTCCGACGTCTAAGAAACCATTATTATCATGACATTAACCTATAAAAATAGGCGTATCACGAGGCCCTTTCGTCTTCACCTC
GAGCGGTAAATCCGGCTGAAGAGTATTCACAACGGTCTTCATTTCAGCAGAAATAAGGTCCAGAACAGGTAGTTTGTTTTCCGGCTTCA
TACAAAAACTCCTTTCAAATTACGTCATTGTAAGGAACCACTGCCATGAAAATGCGATCCCGCCTGCTGATATTGAAACTGGCTGCGTC
TCGCGCGCTCCCGTCAGATTGTGTTAACATTCGCCGCTCAGTTAACCACCCGTAAAAACAACCATGAAATTTCCCGGTAAACGTAAATC
CAAACATTACTTCCCCGTAAACGCACGCGATCCGCTGCTTCAGCAATTCCAGCCAGAAAACGAAACCAGCGCTGGGATCCTCTAGATTT
AAGAAGGAGATATACATATG
>mcrA
GAGGCAATTCCGACGTCTAAGAAACCATTATTATCATGACATTAACCTATAAAAATAGGCGTATCACGAGGCCCTTTCGTCTTCACCTC
GAGCAGGAACTCCTCGCCAGAAGGTGGCGATTATCTATGATGTTGGTGTGTCAACTTTGTATAAGAGGTTTCCTGCAGGGGATAAATAA
AGTTAAAGACACTTTGTGTACAAAAGAAAGTAAAACAACAGCAACTTGTTGCAATTTTATCAATAAAAGTAGTATTGTCGTGAAAAATT
GATTAAAGATTAATATTATGCATGTTTTTGATAATAATGGAATTGAACTGAAAGCTGAGTGTTCGATAGGTGAAGAGGATGGTGTTTAT
GGTCTAATCCTTGAGTCGTGGGGGGGATCCTCTAGATTTAAGAAGGAGATATACATATG
>flgH
GAGGCAATTCCGACGTCTAAGAAACCATTATTATCATGACATTAACCTATAAAAATAGGCGTATCACGAGGCCCTTTCGTCTTCACCTC
GAGGCAACGCGCTTACGAAATCAACAGTAAAGCGGTGTCCACCACCGATCAGATGCTGCAAAAACTGACGCAACTCTAAGGCTTAACCG
GTGGCAGGTTCACCGGTTTACTGATTTTTGAAGATGATAGCCATGCAAAAAAACGCTGCGCATACTTATGCCATTTCCAGCTTGTTGGT
GCTTTCACTAACCGGCTGCGCCTGGATACCCTGGATCCTCTAGATTTAAGAAGGAGATATACATATG
>gloA
GAGGCAATTCCGACGTCTAAGAAACCATTATTATCATGACATTAACCTATAAAAATAGGCGTATCACGAGGCCCTTTCGTCTTCACCTC
GAGCTTAACCCACAGCGTGCCGAAAGTTTCTACGGTGGCGGCGCGGAAGGCTATACCGATTACCCGACGTTGTAATCCAACATTGCGAG
CGGCGTAAAGCCGCCGCTATACTAAAACAACATTTTGAATCTGTTAGCCATTTTGAGGATAAAAAGATGCGTCTTCTTCATACCATGCT
GCGCGTTGGCGATTTGCAACGCTCCATCGATTTTTATACCAAAGTGCTGGGCATGAAACTGCTGGATCCTCTAGATTTAAGAAGGAGAT
ATACATATG
>lpp
GAGGCAATTCCGACGTCTAAGAAACCATTATTATCATGACATTAACCTATAAAAATAGGCGTATCACGAGGCCCTTTCGTCTTCACCTC
GAGTGGCACACAAAGGTGACGTTGTAGTTATGGTTTCTGGTGCACTGGTACCGAGCGGCACTACTAACACCGCATCTGTTCACGTCCTG
TAATATTGCTTTTGTGAATTAATTTGTATATCGAAGCGCCCTGATGGGCGCTTTTTTTATTTAATCGATAACCAGAAGCAATAAAAAAT
CAAATCGGATTTCACTATATAATCTCACTTTATCTAAGATGAATCCGATGGAAGCATCCTGTTTTCTCTCAATTTTTTTATCTAAAACC
CAGCGTTCGATGCTTCTTTGAGCGAACGATCAAAAATAAGTGCCTTCCCATCAAAAAAATATTCTCAACATAAAAAACTTTGTGTAATA
CTTGTAACGCTACATGGAGATTAACTCAATCTAGAGGGTATTAATAATGAAAGCTACTAAACTGGTACTGGGCGCGGTAATCCTGGGTT
CTACTCTGCTGGCAGGTTGCTCCAGCAACGCTAAAATCGAGGATCCTCTAGATTTAAGAAGGAGATATACATATG
>lpd
GAGGCAATTCCGACGTCTAAGAAACCATTATTATCATGACATTAACCTATAAAAATAGGCGTATCACGAGGCCCTTTCGTCTTCACCTC
GAGATCGACGGTGCTGATGGTGCCCGTTTCATTACCATCATTAACAACACGCTGTCTGACATTCGCCGTCTGGTGATGTAAGTAAAAGA
GCCGGCCCAACGGCCGGCTTTTTTCTGGTAATCTCATGAATGTATTGAGGTTATTAGCGAATAGACAAATCGGTTGCCGTTTGTTGTTT
AAAAATTGTTAACAATTTTGTAAAATACCGACGGATAGAACGACCCGGTGGTGGTTAGGGTATTACTTCACATACCCTATGGATTTCTG
GGTGCAGCAAGGTAGCAAGCGCCAGAATCCCCAGGAGCTTACATAAGTAAGTGACTGGGGTGAGGGCGTGAAGCTAACGCCGCTGCGGC
CTGAAAGACGACGGGTATGACCGCCGGAGATAAATATATAGAGGTCATGATGAGTACTGAAATCAAAACTCAGGTCGTGGTACTTGGGG
CAGGCCCCGCAGGTTACTCCGCTGCCTTCCGTTGCGCTGATTTGGATCCTCTAGATTTAAGAAGGAGATATACATATG
>arpA
GAGGCAATTCCGACGTCTAAGAAACCATTATTATCATGACATTAACCTATAAAAATAGGCGTATCACGAGGCCCTTTCGTCTTCACCTC
GATGGATCCTGATTAAAGCGGCGAAGGAAGTGACGCTGGCGTACGGTGGAATGCGCTGACTTTTTCTGGCGGGCAGAGGCAATATTCTG
CCCATCATACCTGAGTGGCAATAGAATAAGGGTGTCTGTTAATCGCATTGACGCCAAAATAACTTAATGTCATACACTTCACACACTTT
TATTAACTCAGCATTATTTTTAAACATCAAACCACTTAATTATTAACAAAATCGCAAGATCTCATTCCACCGAAACTTTTACGGCTGCA
TTATGAGATTGAGCGATTTATGATATTTCTTTAGTTGTAATCATATTAAATTTTTTCACTGTTCTGATAGTTAAAATTCAAGACATCAA
TAAACAATGAGATATTTAAATGATTACTCGTATTCCTCGTAGTTCTTTCTCTGCAAATATTAATAATACAGCCCAGACAAATGAACACC
CTCGAGAGATCCTCTAGATTTAAGAAGGAGATATACATATG
>aldH
GAGGCAATTCCGACGTCTAAGAAACCATTATTATCATGACATTAACCTATAAAAATAGGCGTATCACGAGGCCCTTTCGTCTTCACCTC
GAGCATCCCGCACAGTTTCAGTAATACGTCGGCAGGTATTTGCCGAATTATCAGCGCCCATACGCCCACCACGTTTTAATCTTTTTGTT
CTGTAAGCCGGGTAAGCGCAATGCGCCCGGCAATCTATACACAAAATCATTCAAGTTGCATCAAGGCGGCAAGTGAGCGAATCCCGATG
AGCTTACTCAGGTAAGTGATTCGGGGGAGCGAACGCAGCCAAGGCAGAGGCGGCTTGAAGGATGAAGTGTATATAAAGAATGTCGCCAG
AAATAAACGGGCATACGGCCCGGGGATCTCTGCGCCCTGACGTTCACAAACTGCATATATCTGATAGACGTGAAACAGGAGTCATAATG
AATTTTCATCATCTGGCTTACTGGCAGGATAAAGCGTTAAGTCTCGCCATTGAAAACCGCTTATTTATTAACGGTGAATATACTGCTGC
GGCGGGATCCTCTAGATTTAAGAAGGAGATATACATATG
>menD

Data Set S4 - MG1655 library reference file.fasta

```
GAGGCAATTCCGACGTCTAAGAAACCATTATTATCATGACATTAACCTATAAAAATAGGCGTATCACGAGGCCCTTTCGTCTTCACCTC
GATGGATCCAATCGACAACAAAGCGGCAGGGCTGCGTACTTTATTACAAATGGAATAGTAATGAGTCGCATCATTACCGATTCATATCA
ATAATCTATTTTTGTAGCTCCTTATACTTAGCCGCAATATTGATACCGGACAAACTCATGTCAGTAAGCGCATTTAACCGACGCTGGGC
GGCGGTCATTCTGGAAGCATTAACGCGTCACGGCGTCAGACACATCTGTATCGCCCCAGGCTCGAGAGATCCTCTAGATTTAAGAAGGA
GATATACATATG
>sieB
GAGGCAATTCCGACGTCTAAGAAACCATTATTATCATGACATTAACCTATAAAAATAGGCGTATCACGAGGCCCTTTCGTCTTCACCTC
GAGTCGCCAGGAGTCACACACTGACGTGGTATTTCATCAGTTCCGAAGTGATGTGCAATCATAATTATCTCCATTTTTACAAATGAACT
TTGTTGATGCGGTGTCTGGTGCCTCCAGGTGACTGCAACCAGTTAACAATTACAGTCGGCTTTCCCACCCAAACCAATAAGGACTAACA
TGACTTTTAACTGTGCCACGTGCGCTTAGCCGCATTCACCGCATCACAAAATTCACTTTAAAAAGGGCGGACATCAGCCGAACTTCAAG
AAAAAAACTGATGCCGCCAGGACTACACACAGCAATGTCGTTATTTACAACCGGAGGCGCACTCCCACCATTTAAATTTAACAGACAAG
ACCGACTCTTTATGGATATCGGAAATGCGCCTTCGTGTTGTGCCCGGTTTTATTTCACCACCTCCGGGCTTCGGTGGTCTCGGCTAGGA
TCCTCTAGATTTAAGAAGGAGATATACATATG
>menF
GAGGCAATTCCGACGTCTAAGAAACCATTATTATCATGACATTAACCTATAAAAATAGGCGTATCACGAGGCCCTTTCGTCTTCACCTC
GATGGATCCCCTGGCAAGGAATTGGTGTGGGCGCGGCCGTTGGGCTGGTACTAGGACTGTTGCTGGCACGCCGTTAAAACCTCCCTACA
CCGGGGTACTGCATTTTTTTTGTAGTACCCCGTATAATGTGAGGCTTTTAACAGGGAGAGGTCCGCGTGCAATCACTTACTACGGCGCT
GGAAAATCTACTGCGCCATTTGTCGCAAGAGATTCCGGCGACACCCGGCATTCGGGTTATCGATATTCCTTTCCCTCTCAAAGACGCTT
TTGATGCCTTGAGCTGGCTGGCCAGTCAGCAAACATACCCGCAATTCTACTGGCAACAACGTAATGGTGATGAAGAAGCTGTCGTCCTG
GGCGCGATTACCCGTTTTACGTCGTTGGACCAGGCACAACGTTTTCTTCGCCAGCACCCGGAACACGCCGACTTACGCATTTGGGGGCT
GAATGCTTTTGACCCGTCGCAGGGCAATTTACCTCGAGAGATCCTCTAGATTTAAGAAGGAGATATACATATG
>menG
GAGGCAATTCCGACGTCTAAGAAACCATTATTATCATGACATTAACCTATAAAAATAGGCGTATCACGAGGCCCTTTCGTCTTCACCTC
GATGGATCCTCCTAAGCCAGTGGGCAGCATAACTGACAAATATCAATTAACAATTGATGATTTTGCCAACAGCCCACATAGCGCGATAT
ACTGAAAATTCTCGCAGCAACTGAATGTTAAGCCTATGAAATACGATACTTCCGAGCTTTGTGACATCTATCAAGAAGATGTTAACGTC
GTGGAACCGCTGTTCTCCAACTTTGGCGGCTCGAGAGATCCTCTAGATTTAAGAAGGAGATATACATATG
>aldB
GAGGCAATTCCGACGTCTAAGAAACCATTATTATCATGACATTAACCTATAAAAATAGGCGTATCACGAGGCCCTTTCGTCTTCACCTC
GATGGATCCGGCGACTAATGCCCTGAAAGATGCCTGTGGCTTTACTAACCCGATCCAGGCAACTCACGAAGAAATTGTGGCGATTTATC
GCGCAGCGATGTAATCATCATTTCCACAACGGCTGGCAAATTGTTAGCCGCTTTTTCAACTATCTCTGTAACCCTTGCCCGTAAATTCG
TGATAGCTGTCGTAAAGCTGTTACCGACTGGCGAAGATTTCGCCAGTCACGTCTACCCTTGTTATACCTCACACCGCAAGGAGCTCGAG
AGATCCTCTAGATTTAAGAAGGAGATATACATATG
>aldA
GAGGCAATTCCGACGTCTAAGAAACCATTATTATCATGACATTAACCTATAAAAATAGGCGTATCACGAGGCCCTTTCGTCTTCACCTC
GAGTTCCGGCAGAAGTCATCCATGCATGGCAAACGCTGAAACATGATGCGGTGCTCATCGAGGCGATGGAAAGTCGCTCGTTACGTTAA
AAATTGCCCGTTTGTGAACCACTTGTTTGCAAACGGGCATGACTCCTGACTTTTATTTCTGCCTTTTATTCCTTTTACACTTGTTTTTA
TGAAGCCCTTCACAGAATTGTCCTTTCACGATTCCGTCTCTCTGATGATTGATGTTAATTAACAATGTATTCACCGAAAACAAACATAT
AAATCACAGGAGTCGCCCATGTCAGTACCCGTTCAACATCCTATGTATATCGATGGACAGTTTGTTACCTGGCGTGGAGACGCATGGAT
TGAGGATCCTCTAGATTTAAGAAGGAGATATACATATG
>sbcB
GAGGCAATTCCGACGTCTAAGAAACCATTATTATCATGACATTAACCTATAAAAATAGGCGTATCACGAGGCCCTTTCGTCTTCACCTC
GAGAAGGAATGTTTTCCGCCGCAAAACCAGACGATAAGTTAAAAACGAACAAAGAAGCAGCAATAATAAGACGGCGTTTCAACAGCAAA
CCCTCAGGAGTTTCAAATAGCTGTTCTTTTTACGGAAATACTTATGAACTGGCTGGAATAAAGTGCAAGAAAATGTGACTACCCTCTCA
TTTTTATCTGACATGATCTGTTGCCACTCGCTGCCAAATTGTGGCGCTAAAGCTGATTAGCACGGTGATATTTGATACTCTGGCAGACA
GCAGAAATAACGGATTTAACCTAATGATGAATGACGGTAAGCAACAATCTACCTTTTTGTTTCACGATTACGAAACCTTTGGCACGCAC
CCCGCGTTAGATCGCCCTGCACGGATCCCCGGCGGAAAACATTCCTTCTCGAGAAGGAATGTTTTCCGCCGAGCGGCGTTAGATCGCCC
TGCACGGATCCTCTAGATTTAAGAAGGAGATATACATATG
>priA
GAGGCAATTCCGACGTCTAAGAAACCATTATTATCATGACATTAACCTATAAAAATAGGCGTATCACGAGGCCCTTTCGTCTTCACCTC
GATGGATCCAACGGTGGAGCGGATTTTCATTACGTTACCGCAAGAGCAGCTAGCAGTAATTTCTTCGTATTTCGGGTGAATATCTTTTT
TCATGGGAAAACCTCGGTTTAAGGCCGCGTCGCTCTTCCAGCCCTAACGCCAGACACCACGCGATGTTAAAAGTATAGCTTCAATACGA
TCATTTCGTACGAAGGCGCGAAATCATACAGAAATTAACCAGCGTATGCAAACTGATCCGCACTCTTCTACGGCAATGTGTATACTAAC
CCACCGAATTTCAAGTCAGGATGATGCTATGCCCGTTGCCCACGTTGCCTTGCCCGTTCCGCTTCCTCGTACCTTTGACTATCTGCTGC
CAGAAGGCATGACGGTTCTCGAGAACCGTCATGCCTTCTGGATCCAACGGTGGCGGCCGCTCGAGAGATCCTCTAGATTTAAGAAGGAG
ATATACATATG
>priC
GAGGCAATTCCGACGTCTAAGAAACCATTATTATCATGACATTAACCTATAAAAATAGGCGTATCACGAGGCCCTTTCGTCTTCACCTC
GATGGATCCCGGTAATACCACGCCCAGCGTACCCAGCACTACCGCCAGCCAGCCAATGATGATTAAAATGATTCGTTGCATATTGAGTG
TTGTCAGCTTACAGAGTGGCTACTTTAGCATAACAATTATCATTTTCATTGAGGTCTTATCGTGAAAACCGCCCTGCTGCTGGAAAAAC
TGGAAGGACAGCTCGCTACGCTGCGTCAGCGTTGTGCCCCGGTGTCACAGTTCGCCACGCTACTCGAGAGATCCTCTAGATTTAAGAAG
GAGATATACATATG
>yojI
```

GAGGCAATTCCGACGTCTAAGAAACCATTATTATCATGACATTAACCTATAAAAATAGGCGTATCACGAGGCCCTTTCGTCTTCACCTC
GATGGATCCTGCCGCTACAACCTGACATTCCGTCAGGCAGGTAAAAAGAATAAAAATAAGAATTATTATTGCTGTGCGCGCGAAGATG
TTTAAACTGCGGGCTGTAATTCATTGTCCGGGTTTTCTGCATGGAACTTCTTGTACTTGTCTGGCGGCAGTATCGCTGGCCATTTATCA
GTGTGATGGCGCTAAGCCTCCTCGAGAGATCCTCTAGATTTAAGAAGGAGATATACATATG
>yebU
GAGGCAATTCCGACGTCTAAGAAACCATTATTATCATGACATTAACCTATAAAAATAGGCGTATCACGAGGCCCTTTCGTCTTCACCTC
GAGAAGCCCAGGAAGGCAAACACTTCCTGTTGCAGGAAAGTGAACCGAAAGAGTGGCGTGAATGGGGAACTGCGCTTCCCAAATAATGC
CCACTGCTCCGGCGTGCCTGCGCCGGAGCGTTTATGCTAAACTGCGCGCCTGTTTTTTTGCCAGTGGTACATGCTCGTGGCCCAACACA
CCGTTTATTTCCCGGACGCCTTTCTGACACAAATGCGCGAAGCGATGCCTTCGACGCTCTCGGATCCTCTAGATTTAAGAAGGAGATAT
ACATATG
>yebQ
GAGGCAATTCCGACGTCTAAGAAACCATTATTATCATGACATTAACCTATAAAAATAGGCGTATCACGAGGCCCTTTCGTCTTCACCTC
GAGCTCTTCACCCAGCGCCTGCAAAATGCCAAAAACTTTTAGCACGGAAGATACAGAATCAGGCTGTTTATCCAGATCTGCGTTAGCCA
TTTATCACCTCATTGCGAGTGTTTTATAAAAATCAGAACTGTTTTTTATTATAATTTCGCACCAGGGTGGTCGCAATCCATCTTTTGCC
GGTTAGTTACAATTCTGCGACATCCACCGTGAATATCAGTGCTAGAATCATACCCCTGTTGATTATTCACCAGGATCCCGGGATCCTCT
AGATTTAAGAAGGAGATATACATATG
>yebR
GAGGCAATTCCGACGTCTAAGAAACCATTATTATCATGACATTAACCTATAAAAATAGGCGTATCACGAGGCCCTTTCGTCTTCACCTC
GATGGATCCCGGGCAACGTTGGTAATCACCACGCGGCAGTTCCTCGCCGATTGCCCTCACTGTTATCTTTTTTGTCGGCGTAATTTGTG
GTGTGTTAAGAGCCATGACGTGCCAAATGTAATAAGAATTAACATTAATCTTAACCCATGATGGGAATGATCTTGAGCATTAACGCATT
TAATGCTTATTTTAATAGGCTGGCAGATCCAGGTAAGACAACCAAGTGATTATATAATGAACAAAACAGAATTTTACGCGGCTCGAGAG
ATCCTCTAGATTTAAGAAGGAGATATACATATG
>yebS
GAGGCAATTCCGACGTCTAAGAAACCATTATTATCATGACATTAACCTATAAAAATAGGCGTATCACGAGGCCCTTTCGTCTTCACCTC
GAGCCGCGTAAAATTCTGTTTTGTTCATTATATAATCACTTGGTTGTCTTACCTGGATCTGCCAGCCTATTAAAATAAGCATTAAATGC
GTTAATGCTCAAGATCATTCCCATCATGGGTTAAGATTAATGTTAATTCTTATTACATTTGGCACGTCATGGCTCTTAACACACCACAA
ATTACGCCGACAAAAAAGATAACAGTGAGGGCAATCGGCGAGGAACTGCCGCGTGGTGATTACCAACGTTGCCCGGGATCCTCTAGATT
TAAGAAGGAGATATACATATG
>yebO
GAGGCAATTCCGACGTCTAAGAAACCATTATTATCATGACATTAACCTATAAAAATAGGCGTATCACGAGGCCCTTTCGTCTTCACCTC
GATGGATCCTGGGCGCAGGTATTCAACATGATGTGCGATCAGGATGTACAATTTTTCAGCGGAATTTGTGCCATTAACCAGTTTATCCC
GTGGTGATATCATTTTTTCAGCACTAGATTTTCTTCCTCTCCAGCAGTGGTAGAATAGCGCCTCTACTCAGAGGTGGTGAAATGAACGA
AGTTGTAAATTCAGGCGTGATGAACATTGCGTCTTTGGTTGTATCGGTGGTGGTTCTTCTTATCGGGCTCATCTTGTGGTCTCGAGAGA
TCCTCTAGATTTAAGAAGGAGATATACATATG
>yebE
GAGGCAATTCCGACGTCTAAGAAACCATTATTATCATGACATTAACCTATAAAAATAGGCGTATCACGAGGCCCTTTCGTCTTCACCTC
GATGGATCCAGTACCGCTAACCGTGCGTGGTAAAAGTGCCGATATTCATTACCAGGTCAGCGTGGACTGCAAAGCGGGAATGGCGGAAT
ATCAGCGGCGTTAATGGCTTCTGATAATTCGATTGACTGTTGGCGGATGCGGCGCGAGTGCCTTGTCCGCCCCAGAACGCACCACAATC
GTAGGCCTGAGAAGACGCAACAGCGTCGCATCGGGCAATCAGCATCAAACAATGCCACTCAAAACGACAAACTACTTTCACCTAAAATA
CCCTTATCACCTGATATACAGTAAATACCTTACCTGGCGCGAATTTCTTTGAATAACGGCCTACCGCCCCGCTTATTTGCGCATCTTTT
GTCTCGATATACATTTTCTGACATCCTCATCCGCTATGCTACAGGCACTAAAAATGGTTATTGAGGAGCAAATATGGCTAACTGGTTAA
ATCAACTGCAATCCCTGCTTGGGCAAAGCAGTTCTTCTACCTCCTCGTCTGCGGCTCGAGAGATCCTCTAGATTTAAGAAGGAGATATA
CATATG
>yebF
GAGGCAATTCCGACGTCTAAGAAACCATTATTATCATGACATTAACCTATAAAAATAGGCGTATCACGAGGCCCTTTCGTCTTCACCTC
GATGGATCCGTCGTCGGTGCTGAATCCGAAGAGGAAGATGCCTCACACGCGGCCTGAAGCAACTTGCGCTCCTGACAATTTGTACCATG
CTTTCTCCTGAATCATTGTGCTATGGAGAAAAACATGAAAAAAAGAGGGGCGTTTTTAGGGCTGTTGTTGGTTTCTGCCTGCGCATCAG
TTTTCGCTGCCACTCGAGAGATCCTCTAGATTTAAGAAGGAGATATACATATG
>yebG
GAGGCAATTCCGACGTCTAAGAAACCATTATTATCATGACATTAACCTATAAAAATAGGCGTATCACGAGGCCCTTTCGTCTTCACCTC
GATGGATCCCCAGTTCACCGGAGCCTAATAACATCACGCGAGTTGCTGCCGGACGCAGCGCAGTGCCTAATAACGTCATAACGAATATC
CCCTGATTAATTCCGCGCGCAGTATAAACGAAAACGTTTGCGTGTGTCTTTATTGCTGATGTTGATTTCTCAACCGAAAAGAAATATAC
TGTATAAAATCACAGTTATTATGAGAGGCTGCAAATGGCGGTTGAAGTTAAATACGTAGTCATTCGTGAGGGCGAAGAGAAAATGTCGT
TTACCAGCAAAAAGGAAGCCGACTCGAGAGATCCTCTAGATTTAAGAAGGAGATATACATATG
>yebA
GAGGCAATTCCGACGTCTAAGAAACCATTATTATCATGACATTAACCTATAAAAATAGGCGTATCACGAGGCCCTTTCGTCTTCACCTC
GATGGATCCACCAGTATGCGAGCTGCCTGAAAGGAGATTAATGAGGAAGTGATTACGTGCAACAGATAGCCCGCTCTGTCGCCCTGGCG
TTTAATAATTTACCGCGACCACACCGCGTTATGTTGGGGTCGCTCACCGTTCTTACTCTGGCCGTCGCTGTCTGGCGGCCTTATGTTTA
TCACCGTGATGCCACGCCAATTGTCAAAACTCGAGAGATCCTCTAGATTTAAGAAGGAGATATACATATG
>yebB
GAGGCAATTCCGACGTCTAAGAAACCATTATTATCATGACATTAACCTATAAAAATAGGCGTATCACGAGGCCCTTTCGTCTTCACCTC
GAGCTTCATAGCCTACGCCGCCCACTTCAATTAACACCAGCGGGGGTTGTTTTTCAATGATGATGCCTCTGAGTCTGCCTATCACATGA
CGCTCCTGCGTAATGAATCAAAGATAATGCTGTATGATAAAAAAATGCTGGATAGATATCCAGCGAAGGATGAAGAAAACTTGCGAGGT

GTCTCGATGAACTGAAAAATGGCACAGTATAATTTATTCTACAGGTTATATTGGAAGCAAATATTTTAATATTACATATTCAGTGAAGA
AATGCGTAATAAAAATATACATTGCGCCTCCTGAAAGGATCCTCTAGATTTAAGAAGGAGATATACATATG
>ygjN
GAGGCAATTCCGACGTCTAAGAAACCATTATTATCATGACATTAACCTATAAAAATAGGCGTATCACGAGGCCCTTTCGTCTTCACCTC
GATGGATCCTCGGCAACTGCACCACTATTGCGACGAATAATAAATTTGTGGTACTGAAAGCGGTGAAGCTGGGACGTCGTCGGTAAGAT
TGGGAGTTACCTGCCGGATGCGGCGTAAACGCCTTATCCGGCCTACGTTCAGCACGTTACTCCAGGCCGGATAAGACGCGGCAAGCGTC
GCATCCGGCAATGTCTGCCCAGTTCTGGAATCGCGCTTCCATAAAACAAAAAATCCCTGTCACCATCGTCTACATTCTCTTGTTTAGCG
TTTTTCTACGTTTATTCTTCCGTCACACAGATAAATTCATCCGTTGCCAATCTGGCAACGGATGTTATCATATTGCCAATTTGTCAACG
GACGTGATGAAGAATGCACCTGATAACTCAAAAAGCATTGAAAGATGCTGCGGAAAAATACCCGCAACATAAAACGGAGTTGGTGGCTC
TGCTCGAGAGATCCTCTAGATTTAAGAAGGAGATATACATATG
>ygjO
GAGGCAATTCCGACGTCTAAGAAACCATTATTATCATGACATTAACCTATAAAAATAGGCGTATCACGAGGCCCTTTCGTCTTCACCTC
GATGGATCCATTGCTCATAACCGCCCACGAAAAAGAGAAAAAGGGTATACTCACGCACCCTTTTCAGGGGCAATGTCGAAATTTTATCA
TTCAGGAGGGCCGATGAGCCACTTAGACAACGGTTTCCGTTCACTGACACTACAACGTTTTCCGGCGACGGATGACGTTAACTCGAGCG
GGCGACGGATGACGTTAACTCGAGAGATCCTCTAGATTTAAGAAGGAGATATACATATG
>ygjJ
GAGGCAATTCCGACGTCTAAGAAACCATTATTATCATGACATTAACCTATAAAAATAGGCGTATCACGAGGCCCTTTCGTCTTCACCTC
GAGGCTTTGCGTGGTGGAAATACAGTAAATATATAAAGGGATTAACGGCTGAAGAGCGCCATATTGAAGCGACGCCAGCCAGCAATGTT
GATTAAGCAGAATAATAAAAAATGAGCAGGCATTTTCCCTCTTCCTGGATGAGGAGAGGGAAAATAGTTTCTGCCTTTATATTTTTATA
ATTACAACGATAAAAGGCTGTACTTTTTCTTTAGCTCATGGATTAACACAATGAAATTAATCACTGCACCATGCAGAGCATTACTTGCT
CTGCCGTTTTGCTACGCCTTTTCTGCGGCAGGAGAAGAAGCACGGGATCCTCTAGATTTAAGAAGGAGATATACATATG
>ygjH
GAGGCAATTCCGACGTCTAAGAAACCATTATTATCATGACATTAACCTATAAAAATAGGCGTATCACGAGGCCCTTTCGTCTTCACCTC
GATGGATCCACTGTCGCCAGGGATACGCCAGCTTCGATTGCGATGTCTTTTAGTGTTGCCATGTTTACCTTTTTTACTTTCCCTGATGT
GCGCCCCCATTGTAGAGGAAGTGCTTATCGGGTCACCTGTTTTCAGTAAATAAAGCGCGTTTCTTTGACGCTGTAACGGTAACACAGCG
GCTCACGAATGTGATCACGCCTGCATTCTCCATTTTTAGCCTACAGGATTGAGAAAAACTTTTGCTATAGTTTAGTAAAATTTTACCCT
AAGGGAGGAAGCAATGGAAACCGTGGCTTACGCTGATTTTGCACGTCTGGAAATGCGCGTCGGAAAGATTGTGGAAGTGAAACGCCATG
AAAACGCCGACAAGCCTCGAGAGATCCTCTAGATTTAAGAAGGAGATATACATATG
>ygjI
GAGGCAATTCCGACGTCTAAGAAACCATTATTATCATGACATTAACCTATAAAAATAGGCGTATCACGAGGCCCTTTCGTCTTCACCTC
GAGTCTGCAATAACGCGGTCAAAAAAGTGGTTCTCAAAGTCACCATCGAAGATGGTTATTTCCATAACAAATAACAACTACGGCGGCAA
AAGGAGTTTGCCGCCACCGCTACCCTACTCATTTTCGGAGATGTGTTATGTCTGATACCAAACGTAATACAATCGGCAAATTCGGCTTG
CTCTCGCTGACTTTTGCCGCCGTTTAGGATCCTCTAGATTTAAGAAGGAGATATACATATG
>ygjG
GAGGCAATTCCGACGTCTAAGAAACCATTATTATCATGACATTAACCTATAAAAATAGGCGTATCACGAGGCCCTTTCGTCTTCACCTC
GAGTGCAGATCGGTAGTGGACATCAGAGTGGTATCGTCCGCCAGCGGGGTATTTTGCTGGGTGACATACGGATGAGAAGACATGTTCGC
GTCCTGTGCAGGTTATATGGTTGTTAACTTCTTGTCAGAGTTTATGTCGGCCCCGCTGCGGTTATCTTTAACCGATTAATTTGATTTAG
ATCGCAATTTGCGATTTAAACACAAATCTAATTCCTTGATTTAAAATACTTTCACTCTGTTACTATACGAAAACGTTAATTATCTTGCC
CAAAAATCAGGCAATTATTGCCCTGAAAACGTGCATTTGCGCAGCAATCATCAAATCCATACCCGACAAAAACCGTGCAAAATAACAAC
AAATGTTAACAGATAGCATTAAATATTGCACAAATGATAACCGAATTTGTGTTTATCCCGATTTTCGCGATCGCAGCCGGAGTGGCGCA
ATCCCTGCAATACTTAAATCGGTATCATGTGATACGCGAGCCTCCGGATCCTCTAGATTTAAGAAGGAGATATACATATG
>ygjD
GAGGCAATTCCGACGTCTAAGAAACCATTATTATCATGACATTAACCTATAAAAATAGGCGTATCACGAGGCCCTTTCGTCTTCACCTC
GATGGATCCCGCCAGAACACCTGCTTTTTCGCAGGAACGCTTGAAGCGACGCAGAGCTACGTCGAACGGCTCGTTTTCACGTACTTTAA
TTACCGGCATGTGCCTCTCACCTTTGATTAATTCGGTTTGCCGCTGGCATCAACGCCAGCTTATTTCAAAATGGTGCGGAATTTTACTG
CAATTGCTGCTGCTTTGTAAAGCACCGCGGCCTTTTTTGAAAGGGACTTTTATAAGGGTGAGGAGTATACACGAGGCTTTCTCCAGGGG
CGAACAAAGTTTTACATCAACCCGCATTGGTCCTACACTGCGCGGTAATAAAGCGAGGTAAAACAAGTCATGCGTGTACTGGGTATTGA
AACTTCCTGCGATGAAACCGGCATCGCCATTTACGACGATGAAAAAGGTTTGCTCGAGAGATCCTCTAGATTTAAGAAGGAGATATACA
TATG
>bax
GAGGCAATTCCGACGTCTAAGAAACCATTATTATCATGACATTAACCTATAAAAATAGGCGTATCACGAGGCCCTTTCGTCTTCACCTC
GATGGATCCAACCCCGGAGAAGTCCAGCTGGCGGCAACGGCGAAGCCAGGAAGGAGTGTCAGAAAACAGGCGGCGAGTTTCATGGCGGA
TGAGTCCTTATTGCTGCGATTTCAGACAGTTTGCCAGCGATAAGCCAAACAAAACTCATCCCTGAGCGGTAAGTCCACAGGATGAGAAG
CAAGGGTGAGCGATCTCGCGCAAAAAACGGCTGAATTTTGCGATAACCCCCACATTTTCTGCGATTTAGCGCCAATCTGAATCGTTAAC
ACGTGACATAGTTTCAGATTTGGACTATTCCTTGGGGTGCTCTTGACAGCTATTTTTGATATGATTTGAGATTCAACTCTCAAATTTGT
GAAAAATATAAGGTGTTGGAATGATTAAATCCGACCAGGAGACCTGATGATTTTGACTCCCATACGACGATATGGGGCGATGATTCTTA
TGTTACTCACTCTGGTGTTTTCGAGTGAGGTGTTAGCGAAGACGCACACAACACTCGAGAGATCCTCTAGATTTAAGAAGGAGATATAC
ATATG
>ygjV
GAGGCAATTCCGACGTCTAAGAAACCATTATTATCATGACATTAACCTATAAAAATAGGCGTATCACGAGGCCCTTTCGTCTTCACCTC
GATGGATCCAACGACTTCCGCGAACTGGCGATCTTCAAAAGCGGCGTAACGCTATAAAGTCGAAATGAGTTGTTAATAAAAAACGGCGT
TTCATAATGTGAAGCGCCGTTTGCTTTTCCATACCTGCTAAGGATTTATCATGACCGCGTATTGGCTGGCCCAGGGCGTGGGTGTCATC
GCCTTTCTGATTGGTATCACAACATTTTTCAATCGTGACGAACGTCGCCTCGAGAGATCCTCTAGATTTAAGAAGGAGATATACATATG

Data Set S4 - MG1655 library reference file.fasta

```
>yacL
GAGGCAATTCCGACGTCTAAGAAACCATTATTATCATGACATTAACCTATAAAAATAGGCGTATCACGAGGCCCTTTCGTCTTCACCTC
GAGACAGCCGTTGATACTTACCGTTATCTGAACTTCAACCAGCTTTCTCAGTACACCGAGAAAGCCGATGGGGTGATTTTCCAGACTGC
GGTTTAAAAAGTCAGCGCACGCGCTGCGCATAAACGACACAATGCCCGGTGAATGAGATTCCCGGGCATTTTTTTATTTCTAAACCATC
GCCGTTCCGCTGTTTTTCTCCGGTAAGGCTGCGATAATTACATCAATGGCGCAATGCGATTTCGGTGCATTGCCGGGAGCAGAGGAACA
CACTATGGATTACGAATTTCTGCGCGATATTACCGGAGTGGTAAAGGTGCGGGATCCTCTAGATTTAAGAAGGAGATATACATATG
>ygjR
GAGGCAATTCCGACGTCTAAGAAACCATTATTATCATGACATTAACCTATAAAAATAGGCGTATCACGAGGCCCTTTCGTCTTCACCTC
GAGTTGACGCCAAAATCCTCCATCGCCAGCCGAAATATCTCGGGCCTTCGGTCATGATTGGGCCGTTTAGTGAGCATGGCTGTCCGGCG
CAGAAATAATGCGTATCTGCGCACGTCGAAGATGAAAAAGGCGTGCTACATTGACGACAGAATCCCTTTATGGAGTATCCACGCGTTAT
GATACGTTTCGCTGTGATTGGTACGAACTGGATCACTCGCCAGTTCGTCGAGGCCGCCCATGAGAGCGGTAAAGGATCCTCTAGATTTA
AGAAGGAGATATACATATG
>ygjQ
GAGGCAATTCCGACGTCTAAGAAACCATTATTATCATGACATTAACCTATAAAAATAGGCGTATCACGAGGCCCTTTCGTCTTCACCTC
GAGGGAACCCCAGTACCACCAGCTTGAGTTCGACACCCGACTGTGGCTAACGCAGTTGTCGCTTGGGCAGAACAAAATCTGAAAATGAT
GTCAGGCGTTATCTTTTCTTTCTTTAGCGCCTAAAATCGACCTCCCCCCTTTCGTTAAGGATAACGACCATGCTGCGCGCATTTGCCCG
CCTTCTTCTCCGTATTTGTTTTTCTCGCCGGACTCTAAAGATTGCCTGCTTACTGTTACTTGTGGCAGGAGCGACGGATCCTCTAGATT
TAAGAAGGAGATATACATATG
>ecfG
GAGGCAATTCCGACGTCTAAGAAACCATTATTATCATGACATTAACCTATAAAAATAGGCGTATCACGAGGCCCTTTCGTCTTCACCTC
GAGCGCCCAGCGTATTGAGATGGTCACGCAACGCCTCAACGGCACTGTGATTAACAATAAACTTTAATTCGATTTCCTGAGCCATGGCC
TTTACTTATCGGTTATGTCACATAAGTGACGATGAACGGCGAACTTAATGCGATCTTTTTTGTCAGTAGATAGTATTTTGCGCCAAATT
GCCATGCAACGGGCAATTTGACGGGCGTAAAAGTTTGAAGCAGTGGCAAGACCTGACACAGAGGATTCCGATTGATGTCGAATGCTTTG
CGTAGCGACACTGATGCCACTACTATCGTTCCACTTTCCATGACAATAACGACAGCCTGATGCCAAAATTACGCCTGATCGGATTAACT
TTACTCGCACTTAGCGCGACTGCCGTCTCACACGCTGAAGAAACGCGCTATGTTTCCGGGATCCTCTAGATTTAAGAAGGAGATATACA
TATG
>ecfD
GAGGCAATTCCGACGTCTAAGAAACCATTATTATCATGACATTAACCTATAAAAATAGGCGTATCACGAGGCCCTTTCGTCTTCACCTC
GAGCGGCCAAAGCCTGATCTAAGCGTTGACCGAGTTGGTTTTCGGACACCGTTGCAGTGAGCTGTACTCGTTGTGCCATATATACTGCT
TCTTCGTTTATCGTTGGGGTTTTACGGCTTTGCCGTTTAATATAGTGTGCTATTGTAGCTGGTCTTAACCGGGAGCAGGAACAGAGAAT
CTCCCGTATTACATTTTGAGGAAAGTCAAAACGTCATGACGCGCATGAAATATCTGGTGGCAGCCGCCACACTAAGCCTGTTTTTGGCG
GGTTGCTCGGGGTCAAAGGAAGGGATCCCGGGGATCCTCTAGATTTAAGAAGGAGATATACATATG
>ecfE
GAGGCAATTCCGACGTCTAAGAAACCATTATTATCATGACATTAACCTATAAAAATAGGCGTATCACGAGGCCCTTTCGTCTTCACCTC
GAGAATTCCAGGACACGGTGGTATTTTAGATCGTATTGATAGCCTGACGGCTGCGGTACCGGTCTTTGCTTGCTTGTTGTTACTGGTAT
TCAGGACGCTTTAACGGAAGGTAATATGCTGAGTTTTCTCTGGGATTTGGCTTCGTTCATCGTTGCACTGGGTGGGATCCTCTAGATTT
AAGAAGGAGATATACATATG
>ecfK
GAGGCAATTCCGACGTCTAAGAAACCATTATTATCATGACATTAACCTATAAAAATAGGCGTATCACGAGGCCCTTTCGTCTTCACCTC
GAGTATCCGAGCGGGTTCAAGACTTTTGTTATCGCATTGGCTCGATTCTGCTGGTGCTGTTAATGGGGCTTGCACTTTTCAATGATTTC
TCTCGGTTATGAGAGTTAGTTAGGAAGAACGCATAATAACGATGGCGATGAAAAAGTTGCTCATAGCGTCGCTGCTGTTTAGCAGGATC
CTCTAGATTTAAGAAGGAGATATACATATG
>ecfI
GAGGCAATTCCGACGTCTAAGAAACCATTATTATCATGACATTAACCTATAAAAATAGGCGTATCACGAGGCCCTTTCGTCTTCACCTC
GAGCCGCCATTACTCTGGCTCTGGTTGTTTTCTAAGTGGTTAAACAAACGGTCAAATCCAATAGCAGAACGGTAAAGCGGGGATAAATC
AAAGTTACGCATAATCAATAGCTCCTGAAATCAGCGAGAATGTAAGACCTTCCACAATGGACAGGTCAGGTAGCCAGAACACCCATCAG
GCGCGTCCTCATCGGCTACGATGTAAAAATGGGTCTGGAAATGAACTTTTCAAGCCTTATCAGCGGACTTTTTTTTGCAGTTTATGGTC
TATTGCATAGACTGAGGGGGCAGCACGAAACGTTAAAGTGCGATAGCCGCCACAGTGCGACGTAATGATGATGTTATTTTCAGCAAGGG
ATCGCTATAACTCATCAGGCCAACACCCACGATCAACAGATGAAGGATCCTCTAGATTTAAGAAGGAGATATACATATG
>dctA
GAGGCAATTCCGACGTCTAAGAAACCATTATTATCATGACATTAACCTATAAAAATAGGCGTATCACGAGGCCCTTTCGTCTTCACCTC
GATGGATCCTCGCCCACTCCCTATTGAAATCTTCGAAGAGAGTTACCTGGAAGAAAAGTAGCTACCCCAAACTGATTACAAAACTTTAA
AAAGTGCTGGTTTGTGCGAGCCAGCTCAAACTTTTTAACCTTTTTGTTTCAATTATGATCCAGGTACATTTCTGTGATGTTGTCTGGGT
GTTATTTTAAGGCCGCAGGTACCCCATAACCTTACAAGACCTGTGGTTTTACTAAAGGACACCCTATGAAAACCTCTCTGTTTAAAAGC
CTTTACTTTCAGGTCCTGACAGCGATAGCCATTGGTATTCTCCTTGGCCATTTCTATCCTGAAATAGGCGAGCAACTCGAGAGATCCTC
TAGATTTAAGAAGGAGATATACATATG
>pmrD
GAGGCAATTCCGACGTCTAAGAAACCATTATTATCATGACATTAACCTATAAAAATAGGCGTATCACGAGGCCCTTTCGTCTTCACCTC
GATGGATCCAGCTGAAAAACGGCGGTATTAAAATTTCACGTCAGGCGCTAAAAGAGTGGGTGCAACGTCAGCAATAATTAACCCTCCGT
TTCACTTCTCATTTCTGCCATTCCATTGCCGTTGATAAAGAGTTTATCTGTTAATCTGTAAATTGATGTGAAAACTCTTAGCAAACAGG
ATAATGCAATGGAATGGCTGGTCAAAAAATCGTGCTGCAATAAACAAGACTCGAGAGATCCTCTAGATTTAAGAAGGAGATATACATAT
G
>ybaK
```

GAGGCAATTCCGACGTCTAAGAAACCATTATTATCATGACATTAACCTATAAAAATAGGCGTATCACGAGGCCCTTTCGTCTTCACCTC
GATGGATCCGGGATAAATTACGCGCCATGCCGCCGGGGCGATATGTGGTCGCGGTCGGTGCACTACACCTGTATGGAGAAGGGAATTTG
CCGCAAATGTTGCGCTAAAAAAATGGCCAATATCGCTATTGGCCCGTCAAAGAGGAATTTCATTTTTTTATTATTATGCCGTCACTTTA
AGCGACGGTGTAACTCGATTGTCGCTCAATCTCACCATCCTGCCAATACTCTTAGGCAAGATGACTCTTTTTTTTGGACCGCCATCAGG
CGTATGCTTGCTTCGTTTTTTTGTTTAAGGCAGGAAAATTACCATGACACCCGCAGTTAAATTACTCGAAAAAAACAAGATTTCGTTTC
AAATCCATACCTACGAGCACGATCCGCTCGAGAGATCCTCTAGATTTAAGAAGGAGATATACATATG
>ybaN
GAGGCAATTCCGACGTCTAAGAAACCATTATTATCATGACATTAACCTATAAAAATAGGCGTATCACGAGGCCCTTTCGTCTTCACCTC
GAGTAGCGTGGCGAACTGTGACACCGGGGCACAACGCTGACGCAGCGTAGCGAGCTGTCCTTCCAGTTTTTCCAGCAGCAGGGCGGTTT
TCACGATAAGACCTCAATGAAAATGATAATTGTTATGCTAAAGTAGCCACTCTGTAAGCTGACAACACTCAATATGCAACGAATCATTT
TAATCATCATTGGCTGGCTGGCGGTAGTGCTGGGTACGCTGGGCGTGGTATTACCGGGATCCTCTAGATTTAAGAAGGAGATATACATA
TG
>ybaO
GAGGCAATTCCGACGTCTAAGAAACCATTATTATCATGACATTAACCTATAAAAATAGGCGTATCACGAGGCCCTTTCGTCTTCACGTC
GACCATTGCCGAAATCAGGCTGTCTCTCACTATTTGACGCACTGGCTGGACTATCCACATCTACCTTATTCCCCCGAATAACGAGATCC
CTTCCAGCACCGGGCAATTGCCCGGTTTTTTTTGCGTTGAATTTGTCATTTTGTGCCGTGGTGTTTAAACCGCACAGAATAAATTGTCG
TGATTTCACCTTTAAAATAAAATTAAAAGAGAAAAAAATTCTCTGTGGAAGGGCTATGGGATCCTCTAGATTTAAGAAGGAGATATACA
TATG
>ybaL
GAGGCAATTCCGACGTCTAAGAAACCATTATTATCATGACATTAACCTATAAAAATAGGCGTATCACGAGGCCCTTTCGTCTTCACCTC
GATGGATCCTCCTGCCACTATTGGGGATGTTGACCATATTCCTGCCTGATAACCGGCATAAAGACTGATTTCATTGCAGCCAAAGGCAA
ACTTTGGCTGCATCGTTTTACAGTCGCCATAAGCCTTTCCTCTGTTAAACCGCCTTCTGTTTTCCGATCCGTGCTTTTTTTGCAGCTAA
TTCCATTTTTCTGCAAAATTCAATAATTATTCATGAACATCATCAATGAAATTGTCATAAACTATTACCTGTAATTTTGGTTTTCCCGG
CCAAAAATGGACAATAACCACCAGGAAAAGGAGACGGAATGCATCACGCCACCCCGCTTATCACCACCATTGTTGGCGGCCTTGTGCTC
GCCTTTATCCTCGGCATGCTGGCTCGAGAGATCCTCTAGATTTAAGAAGGAGATATACATATG
>ybaB
GAGGCAATTCCGACGTCTAAGAAACCATTATTATCATGACATTAACCTATAAAAATAGGCGTATCACGAGGCCCTTTCGTCTTCACCTC
GAGGCGAGTCCATTATTGCGGATAATAATATTCAGACCCTGCGTCGGTTCTTCGATGCGGAGCTGGATGAAGAAAGTATCCGCCCCATT
TGATCGTAAGCACAGCTTACGTTCGTCATCCTTAACGTGATTGAGAGAGAAACCTATGTTTGGTAAAGGCGGTCTGGGTAACCTGATGA
AGCAAGCCCAGCAGATGCAAGAAAAAATGCAGAAAATGCAGGAAGAGATCGCGGGATCCTCTAGATTTAAGAAGGAGATATACATATG
>ybaA
GAGGCAATTCCGACGTCTAAGAAACCATTATTATCATGACATTAACCTATAAAAATAGGCGTATCACGAGGCCCTTTCGTCTTCACCTC
GAGCCCGCAACGCTACTCTGTTTACCAGGTCAGGTCCGGAAGGAAGCAGCCAAGGCAGATGACGCGTGTGCCGGGATGTAGCTGGCAGG
GCCCCCACCCATTTCTGCCTCCCACCGTTTCGTCAAAAAATCCCAACATGGCTAAACTTTAACCACAACTGACGTCGCAAGAATTGTCT
GGCTGCGCAGTACGCTTCGGAGGTATGTCTGATGAAGTATGTTGATGGTTTTGTGGTTGCCGTTCCTGGATCCTCTAGATTTAAGAAGG
AGATATACATATG
>ybaD
GAGGCAATTCCGACGTCTAAGAAACCATTATTATCATGACATTAACCTATAAAAATAGGCGTATCACGAGGCCCTTTCGTCTTCACCTC
GAGTTCCCAGCAGGAGCAGTCGAAAAACGTTAGTATTCATAGGTCGTCCCTTTCTCGTCGCAATTAGCCTAATGGTAGAGGGTAAAAGC
AGCGAAAGCATCGATCCTTTATCTCAAAAGCGTTGCGCCTTTGTTGTATCGTCAGTTCAGGGTAAAATAGATTTCCGTTAACCACCTGG
TCAGGACGCCGTATGCATTGCCCATTCTGTTTCGCCGTGGACACTAAGGTAATTGACTCTCGTCTCGTGGGCGAGGGTTCATCCGTACG
GGATCCTCTAGATTTAAGAAGGAGATATACATATG
>ybaE
GAGGCAATTCCGACGTCTAAGAAACCATTATTATCATGACATTAACCTATAAAAATAGGCGTATCACGAGGCCCTTTCGTCTTCACCTC
GATGGATCCCTGCCAGACGAGCCATTTCTTTCTCCACTCTGTGACACGGTTAATGGTGATTGAGTATCACCTGATATATACGGAAGTAA
TAGTGAATAGTTAAATAATATTGTTCCGGGTTTATATGCGATTGCTCAACCGTCTTAACCAGTATCAACGTCTGTGGCAACCTTCCGCC
GGAAAGCCGCAAACCGTCACCGTCAGCGAACCTCGAGAGATCCTCTAGATTTAAGAAGGAGATATACATATG
>mug
GAGGCAATTCCGACGTCTAAGAAACCATTATTATCATGACATTAACCTATAAAAATAGGCGTATCACGAGGCCCTTTCGTCTTCACCTC
GATGGATCCGCTGGACTTGAACCAGCGACCAAGCGATTATGAGTCGCCTGCTCTAACCACTGAGCTAAGGGGCCATGGTAGCGGATTAT
AAAGTAACTCCGTGTCGCAATCCAGCCATTACCGCGCGCCTGCTGTTTTTATAAACAATGTATTTTCAATCCGTTATACTTTTCCTGGT
GATGTTAGAAAGGAGTAAATATGGTTGAGGATATTTTGGCTCCAGGGTTACGGGTCGTGTTTTGCGGTATCAACCCTGGGCTTTCATCC
GCCGGGACTGGTTTTCCCTTCTCGAGAGATCCTCTAGATTTAAGAAGGAGATATACATATG
>ybaY
GAGGCAATTCCGACGTCTAAGAAACCATTATTATCATGACATTAACCTATAAAAATAGGCGTATCACGAGGCCCTTTCGTCTTCACCTC
GAGCGCGAAAGAGTCCTTCCTCAATTTTTTCCAGATTTAACAATGTCAGTAAATTTTTTAGCGCCTGACTCATATAACTCTCCAGTAAC
AAAGCTGCCGCAGCAAGCCAAAGTGAGTTGAGTATAACGCAAATTTGCTACTGGTCCGATGGGTGCAATGGTCTGAATTACGGGCTAAT
TACAGGCAGAAATGCGTGATGTGTGCCACACTTGTTGACGTTACTATTTTGTTAACCACTCTTCCGGCGAGGAAAGTTAGCCCGCTGGT
GCATTGATAATAAGGAGAAATGAATGAAACTCGTGCACATGGCCAGTGGTTTAGCGGTTGCGATTGCGTTGGCGGCTTGCGCAGATAAA
AGCGCGGATATTCAGACGGGATCCTCTAGATTTAAGAAGGAGATATACATATG
>yzgL
GAGGCAATTCCGACGTCTAAGAAACCATTATTATCATGACATTAACCTATAAAAATAGGCGTATCACGAGGCCCTTTCGTCTTCACCTC
GATGGATCCCGATGCTGAACATTGCCAATATGGGCTACTTCTCTTCTGACCGTACTATCAAAGAGTACGCCGATCATATCTGGCATATC

Data Set S4 - MG1655 library reference file.fasta

GATCCGGTGAGATTGTAAGTTCACCAATAAATAGAACGGGGCCAAAGGGTCCCGTTTTTTTCCGCCATCATTTCGGAAAAGTGTCCAGA
AGGGGCAAGGGGACTCGGGGAGTATAATTTCACCATTCAAAAAGAATGGAAAGAATAATGCAAAACAGAAAATGGATTTTGACCTCGCT
GGTAATGACTTTTTTCGGCATCCCCATACTCGAGAGATCCTCTAGATTTAAGAAGGAGATATACATATG
>ybaP
GAGGCAATTCCGACGTCTAAGAAACCATTATTATCATGACATTAACCTATAAAAATAGGCGTATCACGAGGCCCTTTCGTCTTCACCTC
GATGGATCCTATCTCCGGGTGTCGTCGGTTTTCTTGTTGCCTGTTTCATGGTTTTTAACTGTTGCTTCCTTGTGAATAGCGATGCAAGA
ACATACTGAATCATACAAACACATTAAACAAGTCTCCAACTCCATTTGCGCCATCCCTCACCGCGCGCTAGCATGATATTTCACAAAGG
GAGGTCGTATGGATCTGTTGTACCGGGTAAAAACGCTTTGGGCCGCGCTGCGCGGTAATCATTACACCTGGCCTGCCCTCGAGAGATCC
TCTAGATTTAAGAAGGAGATATACATATG
>ybaQ
GAGGCAATTCCGACGTCTAAGAAACCATTATTATCATGACATTAACCTATAAAAATAGGCGTATCACGAGGCCCTTTCGTCTTCACCTC
GAGGGCAGGCCAGGTGTAATGATTACCGCGCAGCGCGGCCCAAAGCGTTTTTACCCGGTACAACAGATCCATACGACCTCCCTTTGTGA
AATATCATGCTAGCGCGCGGTGAGGGATGGCGCAAATGGAGTTGGAGACTTGTTTAATGTGTTTGTATGATTCAGTATGTTCTTGCATC
GCTATTCACAAGGAAGCAACAGTTAAAAACCATGAAACAGGCAACAAGAAAACCGACGACACCCGGAGATAGGATCCCGGGATCCTCTA
GATTTAAGAAGGAGATATACATATG
>ybaV
GAGGCAATTCCGACGTCTAAGAAACCATTATTATCATGACATTAACCTATAAAAATAGGCGTATCACGAGGCCCTTTCGTCTTCACCTC
GAGCACCCAGAACAACGCACAAATCGTCTTTGAAGCTCTGATGAGTAACCTGCGTAAAGAGGCGAAAATCAAAATTGGCGATGCGCTGG
AACAGCAATAATCCTGAAGCCGCCTCGCAAAAAAATGCGTTGCAGCTGTAACAAGAAAAGGTCGCTTTCGCGGCCTTTTCCATTTCTGA
ACATTGCCATTTGTTTACTGTTTTCACTGCCGTTAAGGTGATTCCACTGTTAACAAACAAGGAGAAAACAGTATGAAACACGGAATTAA
AGCACTGCTCATTACCCTGTCCCTGGCCTGTGCCGGAATGTCTCGGATCCTCTAGATTTAAGAAGGAGATATACATATG
>ybaT
GAGGCAATTCCGACGTCTAAGAAACCATTATTATCATGACATTAACCTATAAAAATAGGCGTATCACGAGGCCCTTTCGTCTTCACCTC
GAGTGGACGAAGATGGCAACAGTGTTCGCGGTCAAAAAATGGTGGCATCGGTCGCTAAGCAACTCGGCTATAACGTGTTTAAGGGCTGA
TCATGATGAACACGGAAGGTAATAACGGTAACAAACCTCTCGGTCTATGGAACGTCGTTTCCATCGGCATTGGATCCTCTAGATTTAAG
AAGGAGATATACATATG
>yijF
GAGGCAATTCCGACGTCTAAGAAACCATTATTATCATGACATTAACCTATAAAAATAGGCGTATCACGAGGCCCTTTCGTCTTCACCTC
GATAGATCTGCTCTGCTGGTAGCCGACCAGTACGGTCAGCGTTTCCTGCAAGAGTGGGAATAACCTACTCCAAACTCCCGGCTTGTCCG
GGAGTTTGAACGCAAAATTGCCTGATGCGCTACGCTTATCAGGCCTACGCAATCTCTGCAATATATTGAATTTGCGTGCTTTTGTAGGC
CGGATAAGGCGTTCACTCCGCATCCGGCACTCTGTGCCAGTAATCTGAAACTCCCACGTTGTTCAGGAATTTTTTATCCGCTTCTGGAC
ATTTTCTCTACAGAACAATCGTTAGCCCGGAAGGTCGAATCTGTGACTCTTATTTTCACGGACGAACCAGATGAAAGCTTCACTGGCGC
TGCTCAGCCTGCTAACGGCATTTACCAGCCACTCCTTAAAATCTCCCGCCGCTCGAGAGATCCTCTAGATTTAAGAAGGAGATATACAT
ATG
>ygdL
GAGGCAATTCCGACGTCTAAGAAACCATTATTATCATGACATTAACCTATAAAAATAGGCGTATCACGAGGCCCTTTCGTCTTCACCTC
GATGGATCCATGGACGTGTCTGGGTGCTGAAAACCGCCCCGGGCGCAGGTAACGTCTTTAGCGGCTGATGTGATATTCTGTATGCCGTT
TTTACGGATATGACAGGGTGAGGGGTAACCTCACATCCTGATTATTGACTAAGTGCGCGTTGTTCATGCCGGATGCGGCGTGAACGCCT
TATCCGGCCTACAAATCATGCAAATCCAAAATATTGCAGGAGTTATGTAGGCCTGATAAGCGTAGCGCATCAGGCATTTATGCGTTTAG
GGTGAGGGATAACCTCGCCCTTTTTAATTCTGAGGTGCTATGTCTGTGGTAATTAGTGATGCATGGCGTCAGCGTTTTGGTGGCACAGC
GCGTCTGTATGGTGAAAAAGCGTTGCACTCGAGAGATCCTCTAGATTTAAGAAGGAGATATACATATG
>ygdI_csdA
GAGGCAATTCCGACGTCTAAGAAACCATTATTATCATGACATTAACCTATAAAAATAGGCGTATCACGAGGCCCTTTCGTCTTCACCTC
GATGGATCCTCAGGTTTAAGCGCGGTCGCGGCGCTGTCGAGATAGACGCCCGCATCCTGTAGTGCGGGAAACTGGGCGCGAAACTGCGC
GGGATTAAAAACGTTCATGGTACTCCTCGGCTTGATGCACCGATCGTGACGCAAAATTCACTTTGATACAAGGAGAGTGCTAACCATCA
GAATTATCTGGATATCCTGGCGAAGCCGCGTAAGTAACATATGCTGAATAGTGTTAGGCGTTAATTTAGCCTGTTAATCAATACTATT
AAGATTTATTAATCGCTTTAAGGAGAAGATGATGAAAAAGACTGCCGCAATTATTTCTGCCTGTATGCTGACTTTTGCCCTGAGCGCCT
GTTCCGGTTCGAACTATGTGATGCACACCAATGACGGCTCGAGAGATCCTCTAGATTTAAGAAGGAGATATACATATG
>yijO
GAGGCAATTCCGACGTCTAAGAAACCATTATTATCATGACATTAACCTATAAAAATAGGCGTATCACGAGGCCCTTTCGTCTTCACCTC
GATGGATCCACTGCCCTATGGCGATCAGGTGGGTAATCAGTAATTATCAGCCAGGCGCTATTTGCCGGATGCGACGCTTGACGCGTCTT
ATCCGGCCTACAGATCGCGGCGGTAGGCTGGATAAGGCGTTTACGCCACATCCGGCAATACAACTGCCGCAATGTTTACATCGAACTCC
CCTCAGGCCACACTTTTCAGGTGGCCTTTTTTAATAATGCCAGTTACTGTTTTCCCGTCTACATACCTAAGGGAATAACATGTATCACG
ACGTCAGCTATCTGCTCTCCCGCCTGATCAACGGCCCGCTGTCCCTGCGCCAGATTTACCTCGAGAGATCCTCTAGATTTAAGAAGGAG
ATATACATATG
>ygdQ
GAGGCAATTCCGACGTCTAAGAAACCATTATTATCATGACATTAACCTATAAAAATAGGCGTATCACGAGGCCCTTTCGTCTTCACCTC
GAGCCACGCGGCTTTTATTTGAAGAAGAATTTCACCAGTGCACTACTGGCCCGTCATTTTCTGATCCAGTAGCCATCGCTTTGACCTGC
CGCTTTCCGGGCATATAATTACCGCTTCATTTTTTTGGCAGGGCTTTTTAGATGTTATTTGCATGGATAACCGATCCTAACGCCTGGCT
TGCGCTCGGTACGCTGACGCTGCTGGAGATCGTGGATCCTCTAGATTTAAGAAGGAGATATACATATG
>ygdR
GAGGCAATTCCGACGTCTAAGAAACCATTATTATCATGACATTAACCTATAAAAATAGGCGTATCACGAGGCCCTTTCGTCTTCACCTC
GAGTCGACATCCACGTACCGAAAGGTTACATCTACTTCGCGATGTTCTTCTCTATTGCGGTTGAAAGCCTCAACTTGATTCGCAACAAA

```
AAGAATCCGCTCTGATACTCCGTACGCTCTCCTGCGGGAGGGCGTACTCACTCCTTTTACCCTCACAGTTTAAGATTTTCCTGCTTTCA
AGATATATAACGTCGGTTTATAAACAGACTATTATCATAGGTAGGCCAAATGAGGATAACCGGATGAAAAAATGGGCAGTAATAATTTC
TGCAGTCGGACTGGCGTTTGCTGTTTCCGGGTGTTCCAGTGATGGATCCTCTAGATTTAAGAAGGAGATATACATATG
>pitA
GAGGCAATTCCGACGTCTAAGAAACCATTATTATCATGACATTAACCTATAAAAATAGGCGTATCACGAGGCCCTTTCGTCTTCACCTC
GAGATCGATCAGCAGAACCCGGCGTCCTGCCTGACCTGCCAGCGCAGAACAGAACATACCCGCCGCACCAGCGCCTATAATAATGGCAT
CAAACCTTTCCACGTTGCGCTCCTCTTAGAAAAAACGGGCGTGAATTGTAAAGATTCCTCAGTGGTCGCACCAGCATCAATATTACTAA
AAGGAAGTATTTGCCTGAATTATATAAGATAATTATTTTTTGAGTGAAATCCATACAGGGGGCAAATCAAAAAAAGTCTATATTTCACT
TTGCCCGCGCCGCGAAAGTCACTGATAATGCGCCGCGTTCATGTCCTCAAAATGGCGTAACGTCCTATGCTACATTTGTTTGCTGGCCT
GGATTTGCATACCGGGCTGTTATTATTGCTTGCACTGGCTTTTGTGCGGATCCTCTAGATTTAAGAAGGAGATATACATATG
>b2974
GAGGCAATTCCGACGTCTAAGAAACCATTATTATCATGACATTAACCTATAAAAATAGGCGTATCACGAGGCCCTTTCGTCTTCACCTC
GATGGATCCTGCTGCAGGCGTATGTGTTTACCGGGATGTTAGTCTCGTAAATCAATAAAGCCGGATGATCGTTAATATCATCCGGCTTC
ATTTACATTATTCTGTCTTATTAATTGAAAATCATTTCTCGCTTTATTATCCATGCAGCAATAATGTTTTGCGTTCAGATAACCTGCTA
AATAATTAAGAGCGTGAGAACATCTATTTTTTTATCACGCAAAACAAGTCTGTATTAATTATGTTCGCTTGATGAGCAAACACCGTTCT
ATCTGTTGCTTTTGCAATCTTAATCGTTAAAACTACCAACAAATTTATATAGTTACGTTGTGAAATGTTGTTAATTTTTTGTTTGTTGA
AATTTCGTAAAAATAGCCAGATAGACATCGTATAGGGGTTATTCTTATAAATTTCACCTGTTTTCTATCAATGATGTCGTTTTCTTAAG
AATGGAGGAAGTTATTAGAGAAGATAAAGCATAAAACTATTTCTCCCAGTTACGAATTTTTTAACATTGTTTTGTCACTTGCGTTATTA
ATGAATAAGAAATTTAAATATAAGAAATCGCTTTTAGCGGCTATTTTGAGCGCAACCCTGTTAGCCGGTTGTGCTCGAGAGATCCTCTA
GATTTAAGAAGGAGATATACATATG
>pitB
GAGGCAATTCCGACGTCTAAGAAACCATTATTATCATGACATTAACCTATAAAAATAGGCGTATCACGAGGCCCTTTCGTCTTCACCTC
GATGGATCCGTTTGCGCGGTGATGAATCACTGGTCATCAAAAAAGAGAGTGATATTGAACCGTTAATTGTGGTGAAAAAGTAATACATG
TTGTAGCAACCCCGTACTGATTATGAAGAATAATCAGTACGGGGATTAATCAAAATATTCAATATTTATATATTCCAATATTTATTCAT
TTCATATGTGAATATATTAACCAGTGGAATACCTGTGTTTTGATTATTTCTAAAGGTTTTGAATGAATGCTTATTGTCTGATACACAAG
AAATAACACTCTTTTTATCGTTAAAAAATGATATTTCACTTTGCCCATGCCGTTAAAGTCACTGATAATGCGTCCGTTCGTAAATTCAA
AATGGCGTAATCTAATATATGCTAAATTTATTTGTTGGCCTTGATATATACACAGGGCTTTTGTTATTGCTTGCTCTGGCATTTGTGTT
GCTCGAGAGATCCTCTAGATTTAAGAAGGAGATATACATATG
>ung
GAGGCAATTCCGACGTCTAAGAAACCATTATTATCATGACATTAACCTATAAAAATAGGCGTATCACGAGGCCCTTTCGTCTTCACCTC
GAGTTCGCCTTTTTCGCTGTCCAGCAGCCAGAAAGAGTTCAGCAGATCGTCGTTAGCGGCTTTAGTAATCTGGATACCTGTAATCATGT
GATGCCTCCCCGGCAAAATTATTTGATTTGTTCAGCCTGTCGCGGCCAATTGGTAAAACCATTGTTGCTTGAGTGTATATATACTCCTC
AAACACCCTTGAATCTTTGATTTAAATCAATAAAAACCACACATCAAGTATGGTCGCAAATGGATTTTATTGTTTTACATCAACTTATG
CGGGTGTGAAATTTTACCAATTTACATTTTTTTGCACTCGTTTAAGTCTAAAAAATGAGCATGATTTTGTTCTGTAGAAAGAAGCAGTT
AAGCTAGGCGGATTGAAGATTCGCAGGAGAGCGAGATGGCTAACGAATTAACCTGGCATGACGTGCTGGCTGAAGAGAAGCAGCAACCC
TATTTTCTTAATACCCTTCAGACCGTCGCGGATCCTCTAGATTTAAGAAGGAGATATACATATG
>rimL
GAGGCAATTCCGACGTCTAAGAAACCATTATTATCATGACATTAACCTATAAAAATAGGCGTATCACGAGGCCCTTTCGTCTTCACCTC
GAGACAATGCGGAAGTGGTCCGCATGAAAAAACAAAAGCTACAGTTAAAAGATGAGATGCTCAAAATCCTGCAGCAGGAGAGCGTCAAA
GAGGTGTAAACTTTCCTAAGCCGCCAGCCAGGCGGCTTTTTTAACAACTGCATGGATTGACTGGAGATAAGATGACTGAAACGATAAAA
GTAAGCGAATCACTTGAATTACATGCTGTTGCAGAAAATCACGTCAAACCTCTTTATCAGTTAATCTGTAAAAATAAAACCTGGTTACA
GCAGTCGCTAAACTGGCCGCAGTTTGTTCAAAGTGAAGAGGACACGCGAAAAACGGTGCAGGGTAATGTGATGTTGCATCAACGCGGCT
ATGCCAAAATGTGGATCCTCTAGATTTAAGAAGGAGATATACATATG
>rimJ
GAGGCAATTCCGACGTCTAAGAAACCATTATTATCATGACATTAACCTATAAAAATAGGCGTATCACGAGGCCCTTTCGTCTTCACCTC
GAGGCACGACCAGCATATTATCGATGAGCAGGAAATATTTACCCAGGTTCCTCGCCTGCGACACGCGGGACATTTCCCCTCCCGGGAAA
TAAAAGATGAGCACTTTCTATTCTGCATTGCCGCTAAGACTTTTCCCATCACTTCGGGGACAATATTTTTTTATCAAAACGTCGCTTTG
ATAGAGAGTTTTTATCAAATATGTGAATAATGCAGAAAATGGCATTTTGGACTTTTCACAGGGTCTGGTTGCGCAGGTATAGTAATACT
TACAGCGTATTAAAGACGTTACGGGAAGGAGTAGGTATAGAATGTTTGGCTATCGCAGTAACGTGCCAAAAGTGCGCTTAACCACAGAC
CGACTGGTCGTGCGTCTGGTGGATCCTCTAGATTTAAGAAGGAGATATACATATG
>insA2
GAGGCAATTCCGACGTCTAAGAAACCATTATTATCATGACATTAACCTATAAAAATAGGCGTATCACGAGGCCCTTTCGTCTTCACCTC
GATGGATCCATCGCCACGGGTTTAACAGACACCTCCGAGTCATTTAAAATGGCTTAAAGAGAGGTGCCCATGAGCGGTAAGCGTTATCC
CGAAGAGTTTAAAACTGAAGCAGTCAAACAGGTTGTTGATCTCGGTTATTCTGTTGCCAGCGTTGCAACACGTCTCGATATCACCACCC
ACAGCCTTTATGCCTGGATAAAGAAGTACGGTCCGGATTCTTCCACTAATAAAGAACAGTCAGATGCTCAGGCCGAGATCCGCCGTCTC
CAGAAAGAGCTGAAACGGGTTACCGACGAACGGGACATATTAAAAAAAGCTGCTGTAGATTCAATCTGTCAATGCAACACCCCTTTCAA
TTATCTCTTTCGGTGTTTTGAACTTCAGTGTCTTTCTCGGTCTGTTGTTTAGCTGAGCAGCAACCAGATCTAGTTCATGTTGAGTATAT
TGGGCAAGACATGTCTTTTTAGGAAAGTACTGCCGAATTAGCCCATTTGTGTTCTCGGTGATGCTGCCAACTTACTGATTTAGTGTATG
ATGGTGTTTTTGAGGTGCTCCAGTGGCTTCTGTTTCTATCAGCTGTCCCTCCTGTTCAGCTACTGACGGGGTGGTGCGTAACGGCAAAA
GCACTGCCGGACATCCTCGAGAGATCCTCTAGATTTAAGAAGGAGATATACATATG
>tonB
GAGGCAATTCCGACGTCTAAGAAACCATTATTATCATGACATTAACCTATAAAAATAGGCGTATCACGAGGCCCTTTCGTCTTCACCTC
GAGCGGGCACAACTCCTTGTTCGGGAAAGTTGTAAAGTACGTTATTTGAAAGGGCGAAGATCTGCAACGGAAAGATGATGTCTTTGTTA
```

```
AGGCCATGCATAAAGTAAGGGTAATTACGCCAAAAATGACATTTTCACTGATCCTGATCGTCTTGCCTTATTGAATATGATTGCTATTT
GCATTTAAAATCGAGACCTGGTTTTTCTACTGAAATGATTATGACTTCAATGACCCTTGATTTACCTCGCCGCTTCCCCTGGCCGACGT
TACTTTCGGTCTGCATTCATGGTGCTGTTGTGGCGGATCCTCTAGATTTAAGAAGGAGATATACATATG
>ydaG
GAGGCAATTCCGACGTCTAAGAAACCATTATTATCATGACATTAACCTATAAAAATAGGCGTATCACGAGGCCCTTTCGTCTTCACCTC
GATGGATCCCGGAAAAGCCAGAACATCACAAAAACTGGAACTTAAACTTATATGGTCTGACGGAAAACTCCTGGATTCCGTTATTTAAC
CCCCCCATCACTTTCTGCTGTCGCCATCACCTATTAGGTTACGCTCAAAACATTAGGCATAGCCTATTGACAATCAATTAGGCATTACC
TATAGTTCCAGCATACCACCCACCCCGCCCCACAGAACGCCGGGCAATACTTCGAGTTACCAGGCAGTGGTAAGGGGTTAAGTAGCCAG
CCCGAGGCGTATGAACATGACGGCGGGATTCAAATTTTGCAGTGCAGCAGTTAGTTCCGCCACCCGGCGTTAAGGGGAGAGATAAGATG
GTGCATTACGAAGTAGTTCAGTATTTGATGGATTGTTGCGGCTCGAGAGATCCTCTAGATTTAAGAAGGAGATATACATATG
>crcA
GAGGCAATTCCGACGTCTAAGAAACCATTATTATCATGACATTAACCTATAAAAATAGGCGTATCACGAGGCCCTTTCGTCTTCACCTC
GAGACACACCAGTGGCGGAATACCCTTTAATGATGTAGCGAGCTACACCCACAATAACCACAACCCCAATAAGGAGCTCAATGAATGTC
AGCATAATTTTTCCTGTCTCCAGGCCCCAAAGTAAATAATAAAAAATTCCTAAAGCTTAAGGAAAAAATATGCCCAATAAATTGGCGAT
GAATGCTGATTAAAATCAAGAAAAACTGCCATTAAGACATTGAAGTTGCTGTTTTTATACATAGATAACAAAACGCATATTTAGCGGTT
GGCAATTATCACATAATAATGTGTGCAAATTGATGGTGGATTGATGCAAATTTGTTAATTGCAATGGTGTTAGCTGCGGCTGTGCGCTC
AAAAAATAATCTAATATGAGCATAGGTTGACGATATATATTATATACTTCGTTAAGATGATTGTTGTATCTCGTTAAAAAATAAAATAA
TTTTTCCTTGATTGCATTTTGTCATCAAAAAAGACTTGGTTTTTCTTTTTTGACTATTCCCATCGCAGAAAACGACGCATCATCTTTAA
TCGATGCGCGGAAATATTTAACTTGAACAAGCGGAAATAAATAGAGCAGCTATTCAGATTATTCTTTATGTTGGGTCTATTAAGGTTAT
GTTAATTGTAGCTTTGCTATGCTAGTAGTAGATTTTTGATAAATGTTTTATGGTCACAAATGAACGTGAGTAAATATGTCGCTATCTTT
TCCTTTGTTTTTATTCAGTTAATCAGCGTTGGTAAAGTTTTTGCTAACGCAGATGAGTGGATGACAGGATCCTCTAGATTTAAGAAGGA
GATATACATATG
>yaaW
GAGGCAATTCCGACGTCTAAGAAACCATTATTATCATGACATTAACCTATAAAAATAGGCGTATCACGAGGCCCTTTCGTCTTCACCTC
GATGGATCCGGATATGGCCACGCTGAAAATTATCGGCGACGACTAACGCCAGAATATAGAAGCCACAAAAAATGAATGTTAATTACCTG
AATGATTCAGATCTGGATTTTCTTCAGCATTGTAGTGAGGAACAGTTGGCAAATTTCGCCCTCGAGAGATCCTCTAGATTTAAGAAGGA
GATATACATATG
>ydaO
GAGGCAATTCCGACGTCTAAGAAACCATTATTATCATGACATTAACCTATAAAAATAGGCGTATCACGAGGCCCTTTCGTCTTCACCTC
GATGGATCCTGAAACTGCGCAGATGGTGTATGAAATTTACGGTATGTGGATTGATGACATGAACGACGAACAGATAGCCATGTTGAATG
CGCGGTTATCGTAGTTGCAAAGTTTGCCCCCAATTTGCCCCATTTAGTACCAGAGAACTGAAATAATGCAAGAAAATCAACAAATTACA
AAGAAAGAACAATACAACCTGAACAAATTACAAAAACGTCTGCGTCGTAACGTGGGCGAAGCCATTGCTGACTCTCGAGAGATCCTCTA
GATTTAAGAAGGAGATATACATATG
>yaaI
GAGGCAATTCCGACGTCTAAGAAACCATTATTATCATGACATTAACCTATAAAAATAGGCGTATCACGAGGCCCTTTCGTCTTCACCTC
GATGGATCCGGTGCCATCCATAATCGCTACACAAGAGTTGGTAGTACCCAGGTCGATACCAATTATTTTACCCATCTAAACGTCTCCAC
TATATATTCGGTCATCATGTGGTTGTGAGTCTGTAATAGGGGCGAAACGTCTGGTTTCAACTGCCCAATTTCATTTTTTTGCAGACTCA
CTCACTGCGGTTGACTACTAAATGGGGTCGTAAACCACGTCATCAAGGGGGAGATGCAAAAATTTTTTGTGCTTTTGCGCAGAAATTTC
GCGATATGGCAGGACAAGACGTAAAATTGACAGCAATTTTATTCATATAAAGAATGAATCGCAGCCAAGGCATTCATCAAAAAATTGTA
ATAAAAAGAAAAGATTACGTGCCTGAATCTTCTCTTTATCAGCAGTAAACTAGTGGGTATTCATCCCCCTACCTCTTCCCACTAAGAGA
ATCCTTATGAAATCCGTTTTTACGATTTCCGCCAGCCTGGCGATTAGCCTGATGCTGTGCTGCACGGCGCAGGCAAACGACCATAAACT
CGAGAGATCCTCTAGATTTAAGAAGGAGATATACATATG
>frr
GAGGCAATTCCGACGTCTAAGAAACCATTATTATCATGACATTAACCTATAAAAATAGGCGTATCACGAGGCCCTTTCGTCTTCACCTC
GAGTATGAACAAACCGGGTGCGCTGCGCCGTGTGGTAATGGGTGAAAAAGAAGGGACTTTAATCACGGAATAATTCCCGTGATGGATAA
ATAAGGGTAAGATTCCGCGTAAGTATCGCGGGGGCGTAAGTCTGGTTATAAGGCGTTATTGTTGCAGGCAGTTTGGTCACGGCCAGCGC
GCAGCAACCGGAGCGTACAAAAGTACGTGAGGATGGCGAGCACTGCCCGGGGCCAAAATGGCAAATAAAATAGCCTAATAATCCAGACG
ATTACCCGTAATATGTTTAATCAGGGCTATACTTAGCACACTTCCACTGTGTGTGACTGTCTGGTCTGACTGAGACAAGTTTTCAAGGA
TTCGTAACGTGATTAGCGATATCAGAAAAGATGCTGAAGTACGCATGGACAAATGCGTAGAAGCGTTCAAAACCCAAATCAGCAAAATA
CGCACGGGTCGGATCCTCTAGATTTAAGAAGGAGATATACATATG
>yaaJ
GAGGCAATTCCGACGTCTAAGAAACCATTATTATCATGACATTAACCTATAAAAATAGGCGTATCACGAGGCCCTTTCGTCTTCACCTC
GATGGATCCGGCATCCTGCGGTTGATACAGCTTCATTGCCGCGATGTCCCCAGTGTCGGCCACTACGGTGGTGTACTGACGAAGGGAGG
TCAATTTGTCCGTCATGATAGTATTTCTCTTTAAACAGCTTGTTAGGGGGATGTAACCGGTCTGCCCTGATGATATCACGACGCAACGT
CAGCGCAACCGCCGACAAGACGGTTATCGCCAGGCGAGACTGTTTCGGATTTCTGACACTGCGTTGATATCGCCCGCCATTTTTATACA
AAACCTCATGTATGCTACGCAGAAGTTATCAAGTACCTCGTAGCGTATATACTTCTTAAACAATTGGTAACGTTTACACAGGAAAGTCA
TCGCGACCGGCAATAAGAGGGATATGCATGCCAGATTTTTTCTCCTTCATTAACAGCGTCCTTTGGGGATCGGTAATGATTTACCTGCT
CTTCGGCGCTCGAGAGATCCTCTAGATTTAAGAAGGAGATATACATATG
>yaaA
GAGGCAATTCCGACGTCTAAGAAACCATTATTATCATGACATTAACCTATAAAAATAGGCGTATCACGAGGCCCTTTCGTCTTCACCTC
GATGGATCCTATCGGTCGCCAGCTTTCTCCGGACGCGTGGGATGATGTTTCGCAGGAGTAATCACAACTATCGATCAACTCATTCTCAT
TTTTTGCTAAAGTCGGCATAAATTTCCTGCAAGGACTGGATATGCTGATTCTTATTTCACCTGCGAAAACGCTTGATTACCAAAGCCCG
TTGACCACCACGCGCTATACGCTCGAGAGATCCTCTAGATTTAAGAAGGAGATATACATATG
```

Data Set S4 - MG1655 library reference file.fasta

```
>lctP
GAGGCAATTCCGACGTCTAAGAAACCATTATTATCATGACATTAACCTATAAAAATAGGCGTATCACGAGGCCCTTTCGTCTTCACCTC
GAGAAAGTGTTCGCGGACTGGTCGTTGTGCACCCAATGACCGCACTGGGCCGCGAAATGGGCCTGCAGGAGATGACTGGGTTTTCAAAG
ACCGCGTTTTAAGAACACAGTATCTACAGGGTGATTCTGCACATTCCTATAGGCCGAGTAAGGTGTTCACGCCGCATCCGGCAAGATAA
GGCGCTCTGGATCAACAACCTAAGGGCAATTCTCTGATGAGGATTGCCCTTTTCTTTACCAGACATCTCCCCCCACAAGAATTGGCCCT
ACCAATTCTTCGCTTATCTGACCTCTGGTTCACAATTTCCCAATTAAAACTCACATCAATGTTGCCAATACATAACATTTAGTTAACCA
TTCATTGTCATTATCCCTACACAACACAATTGGCAGTGCCACTTTTACACAACGTGTGACAAGGAGATGAGCAACAGACTCATTACACG
ATGTGCGTGGACTCCAGGAGACCTGCAATGAATCTCTGGCAACAAAACTACGATCCCGCCGGGAATATCTGGCTTTCCAGTCTGATAGC
ATCGCTTCCCATCCGGATCCTCTAGATTTAAGAAGGAGATATACATATG
>znuC
GAGGCAATTCCGACGTCTAAGAAACCATTATTATCATGACATTAACCTATAAAAATAGGCGTATCACGAGGCCCTTTCGTCTTCACCTC
GAGATGCTGCGAAAAGAAGCGTTTTTTTATGTAACATAATGCGACCAATAATCGTAATGAATATGAGAAGTGTGATATTATAACATTTC
ATGACTACTGCAAGACTAAAATTAACATGACAAGTCTGGTTTCCCTGGAAAATGTCTCGGTTTCTTTTGGCCAACGCCGCGTCCTCTCT
GATGTGTCGCTGGAACTTAAACCTGGGGATCCTCTAGATTTAAGAAGGAGATATACATATG
>znuA
GAGGCAATTCCGACGTCTAAGAAACCATTATTATCATGACATTAACCTATAAAAATAGGCGTATCACGAGGCCCTTTCGTCTTCACCTC
GATGGATCCACATCAGAGAGGACGCGGCGTTGGCCAAAAGAAACCGAGACATTTTCCAGGGAAACCAGACTTGTCATGTTAATTTTAGT
CTTGCAGTAGTCATGAAATGTTATAATATCACACTTCTCATATTCATTACGATTATTGGTCGCATTATGTTACATAAAAAAACGCTTCT
TTTCGCAGCATCTCGAGAGATCCTCTAGATTTAAGAAGGAGATATACATATG
>slt
GAGGCAATTCCGACGTCTAAGAAACCATTATTATCATGACATTAACCTATAAAAATAGGCGTATCACGAGGCCCTTTCGTCTTCACCTC
GAGGCCCCAGGGAAGAAACTCAGAGAGATGTTTTTCAAAATATGACGTTTCGGCGGAACAACTTTGCCGACACGATGCATGGTATAAAC
GAATTGAGCCACGTTGGACTTCGCCTCTATGTTTATCGTGATAATGAGTTTTCAAAGGCGAAGTGTAGCCTTTTTCCCCGCCTAATCCC
AGCCGGTCGATCACCCTCGCAGTAAAAGTAAAAAAGTGTCCGTAACGTGGCGTAAACGGCAATGACTGGTTAGCATAAATCTATTACGC
GGCATGACGCTGCATTGATGTATTTACACTTAGAGGATGCGCTTGTGGAAAAAGCCAAACAAGTTACCTGGCGGCTGTTGGCTGCCGGT
GTCTGTCTGCTGACGGGGATCCTCTAGATTTAAGAAGGAGATATACATATG
>ais
GAGGCAATTCCGACGTCTAAGAAACCATTATTATCATGACATTAACCTATAAAAATAGGCGTATCACGAGGCCCTTTCGTCTTCACCTC
GATGGATCCGTCGCGAAAAAGGCAAAAATTCTGACATTGCTTTTCCTTCCGCCATTGAATACCTGTCCACTTATATTTGCTATAAAGAG
TGTTGTGTATATTTTGCCATTTGGAGCGAAATTTTAAGGATAGAATATTAACTTAACCTTAAGAAACTAATATTAGACGTAAATATTGA
AATTTTTATATTTTTTCTTATTTAGGCTTTGCATTTGGCAAAATTTTGAGGCATTTTGCCGACATCGTAGGATTTTTAATATTACACCA
ACTGCGAATTATCGCCAGAAATGTAGCTCAATTTCACGGTAATTGTCTGGTTGCGCTTGTCTATAGGTGGAGTTTACGTGTTAGCTTTT
TGCCGCTCTTCGTTGAAGTCAAAAAAATATATCATCATTTTACTGGCGCTCGCTGCAATTGCCGGACTGGGTACTCCTCGAGAGATCCT
CTAGATTTAAGAAGGAGATATACATATG
>hepA
GAGGCAATTCCGACGTCTAAGAAACCATTATTATCATGACATTAACCTATAAAAATAGGCGTATCACGAGGCCCTTTCGTCTTCACCTC
GATGGATCCTGACCCGCCAGCTACAACCCGTGGCGGAGGGAATACTCCCTTTTATTGAGGATAATTTTGCTACACTTATGACCGGGCAA
CTTGGGCTATTTTGAGCAAAAAAAAGAGTTCGCCAGATACCATTTTGATGCGTGACGAATGCTTTGCCATCCAGTACCATAGCGCCCTT
TCCATTCCTGGACCTGAATAACACCACTACCTCATAAGCACGGTAGCGGGTGGTTATTGCCTGCAATTAAAGATATAGAGCCGAACACA
TATGCCTTTTACACTTGGTCAACGCTGGATCAGCGATACAGAAAGCGAATTGGGACTTGGAACCGTTGTCGCCTCGAGAGATCCTCTAG
ATTTAAGAAGGAGATATACATATG
>gnd
GAGGCAATTCCGACGTCTAAGAAACCATTATTATCATGACATTAACCTATAAAAATAGGCGTATCACGAGGCCCTTTCGTCTTCACCTC
GATGGATCCGGGGTTGCTGAAAAACGATAACCAACTGGCGATGTTATTCACGCTGGCCAACCTGTTTCGGGCGGACCAAATGATACGTC
AGTGGGAGAGATCTCACTAAAAACTGGGGATAACGCCTTAAATGGCGAAGAAACGGTCTAAATAGGCTGATTCAAGGCATTTACGGGAG
AAAAAATCGGCTCAAACATGAAGAAATGAAATGACTGAGTCAGCCGAGAAGAATTTCCCCGCTTATTCGCACCTTCCTTAATAAAACAA
AAATTCCTAAAGAAAGTATCTATTCTGATACGGTTGTTGATTGGTGCGCAGGATCATTTATGCTGGTACGTTTTTCAGATTTTGTGCGT
GTAAATGGCTTCGATCAAGGTTACTTTATGTACTGTGAAGATATTGACCTGTGCTTGAGGCTTAGCCTGGCTGGTGTCAGACTTCATTA
TGTTCCCGCTTTTCATGCGATACATTATGCTCATCATGACAATCGAAGTTTTTTTTCAAAAGCCTTCAGATGGCACTTAAAAAGTACTT
TTAGATATTTAGCCAGAAAACGTATTTTATCAAATCGCAACTTTGATCGAATTTCATCAGTTTTTCACCCGTAATATAAAGCCGTAAGC
ATATAAGCATGGATAAGCTATTTATACTTTAATAAGTACTTTGTATACTTATTTGCGAACATTCCAGGCCGCGAGCATTCAGCGCGGTG
ATCACACCTGACAGGAGTATGTAATGTCCAAGCAACAGATCGGCGTAGTCGGTATGGCAGTGATGGGACGCAACCTTGCGCTCAACATC
GAAAGCCGTGGTTCTCGAGAGATCCTCTAGATTTAAGAAGGAGATATACATATG
>lepA
GAGGCAATTCCGACGTCTAAGAAACCATTATTATCATGACATTAACCTATAAAAATAGGCGTATCACGAGGCCCTTTCGTCTTCACCTC
GATGGATCCGGCCTTGTGCGATTTGAAACATCTTCGGAAGACGCGAGCCAGTGATTTCACTGGCTTTGTTGCATTTTGCATAACAAATG
CGAGGACGTTTCCAGATACCACAATCCATACGGTATGAACATTTCCTCGCCTCAATGTTGTAGTGTAGAATGCGGCGTTTCTATTAATA
CAGACGTTAAGCTCAGAACAGCGACTTCTAAAAGCCTGGTTAACCGGGCATTAAGGCACAATAATCATACTTTATGAAGAATATACGTA
ACTTTTCGATCATAGCTCACATTGACCACGGTAAATCGACGCTGTCTGACCGTACTCGAGAGATCCTCTAGATTTAAGAAGGAGATATA
CATATG
>hupA
GAGGCAATTCCGACGTCTAAGAAACCATTATTATCATGACATTAACCTATAAAAATAGGCGTATCACGAGGCCCTTTCGTCTTCACCTC
GAGGCTGAGTGCGAAGAACGCGATATCGAGCTGATAAAAGGCCTTAGGGCAGACCTGCGTGAGGCGGGTGAGAGCAATATTGGTATAAT
```

Data Set S4 - MG1655 library reference file.fasta
TTTTCAGCAATAAGACCAGAAAACGTGATTTAACGCCTGATTTGTCGTACCTGGAGTCTTCCCTTTCGCCCCCCGTCTGGTCTACATTT
GGGGGGCGAAAAAAAGTGGCTATCGGTGCGTGTATGCAGGAGAGTGCTATTCTGGCATTTCCGTCGCACTCGATGCTTAGCAAGCGATA
AACACATTGTAAGGATAACTTATGAACAAGACTCAACTGATTGATGTAATTGCAGAGAAAGCAGAACTGTCCAAAACCCAGGCTAAAGC
TGCTCTGGATCCTCTAGATTTAAGAAGGAGATATACATATG
>yfcI
GAGGCAATTCCGACGTCTAAGAAACCATTATTATCATGACATTAACCTATAAAAATAGGCGTATCACGAGGCCCTTTCGTCTTCACCTC
GATGGATCCCGTCTGCAACGGTTCCTTAAGGGATCGCTGAAATAATTTCCTCTCCGGTGGCTAACGCTGCCGGAGTATTTTCATGGGGA
ATTTGCGAGTCAGCTTCCGCTGAAATGTAAAAGTTACAGTCCGCTAAAAATACTTCTGAATTCGCTGTCCAAAGATAAAAACTGCCTGC
TGCATTGGGTGTTAATCGTGGTTATTGTCGCGTCATCTTAAATTTGAGGGGTGACGGAACGCCATGACAATATCGACAACTTCCACGCC
GCATGATGCGGTATTTAAATCTTTTTTACGCCATCCAGACACCGCTCGAGAGATCCTCTAGATTTAAGAAGGAGATATACATATG
>yfcJ
GAGGCAATTCCGACGTCTAAGAAACCATTATTATCATGACATTAACCTATAAAAATAGGCGTATCACGAGGCCCTTTCGTCTTCACCTC
GATGGATCCTAACAGCTTCGGCTTCGGCGGCACCAACGCCACGCTGGTAATGCGCAAGCTGAAAGATTAATTCGCAGTAGGTCGGAGTA
GACGCGCCAGCGTCGCATCCGACGTTACGCGCCAATGCGGCCTCCGGCACTAACGCAAAAGGGAACCTGATGGTTCCCTTTTTCACATC
ATTGACAATCGCCGCCAGTTCCAGGCAAACTTCCCGCTTTGTCGATTTCCTTCTGAAAAGACGTACGCGTTAAATCCTGCCAACGCACC
GTAACCCTGAAACCAGAGAGATGAGACGGGGATACTCCTCGCCTTGCGCTGCATTCTGGAGTAATGCATGACTGCTGTAAGCCAAACCG
AAACACGATCTTCTGCCAATTTTTCGCTCTTCCGCATCGCTTTTGCGGTTTTTCCTCGAGAGATCCTCTAGATTTAAGAAGGAGATATA
CATATG
>hupB
GAGGCAATTCCGACGTCTAAGAAACCATTATTATCATGACATTAACCTATAAAAATAGGCGTATCACGAGGCCCTTTCGTCTTCACCTC
GAGTGTGAAGCGCATTGAGGAAGTTCTGACTCTGGCGCTGCAAAATGAACCGTCTGGTATGCAGGTTGTGACTGCAAAATAGTGACCTC
GCGCAAAATGCACTAATAAAAACAGGGCTGGCAGGCTAATTCGGGCTTGCCAGCCTTTTTTTGTCTCGCTAAGTTAGATGGCGGATCGG
GCTTGCCCTTATTAAGGGGTGTTGTAAGGGGATGGCTGGCCTGATATAACTGCTGCGCGTTCGTACCTTGAAGGATTCAAGTGCGATAT
AAATTATAAAGAGGAAGAGAAGAGTGAATAAATCTCAATTGATCGACAAGATTGCTGCAGGGGCTGATATCTCTAAAGCTGCGGCTGGC
CGTGCGTTAGATGGGATCCTCTAGATTTAAGAAGGAGATATACATATG
>yfcC
GAGGCAATTCCGACGTCTAAGAAACCATTATTATCATGACATTAACCTATAAAAATAGGCGTATCACGAGGCCCTTTCGTCTTCACCTC
GAGTAACGACCTGTCCCGTGGCGCACTGGTTGACGATATCGTCTACACCATCGCGCTGACTGCGATTCAGTCTGCACAGCAGCAGTAAT
CTCGTCATCATCCGCAGCTTTGCGCTGCGGATATCTGAACCGGAAATAATCACTATTTCCGGTTTTTTATTCTCTTAATTTGCATTAAT
CCTTTCTGATTATCTTGCTTAACTGCGCTGCATCAATGAATTGCGCCATCCCACTTTGCATACTTACCACTTTGTTTTGTGCAAGGGAA
TATTTGCGCTATGTCCGCAATCACTGAATCCAAACCAACAAGAAGATGGGCAATGCCCGATACGTTGGTGGGATCCTCTAGATTTAAGA
AGGAGATATACATATG
>yfcD
GAGGCAATTCCGACGTCTAAGAAACCATTATTATCATGACATTAACCTATAAAAATAGGCGTATCACGAGGCCCTTTCGTCTTCACCTC
GATGGATCCAATCCGGCGAGTTATGGCATGCTGGATAATGACGTACTTAGTGTAATCGCACTCAATGATCAAAGTATCATTGCGCAGGT
CGCGATTAATCCGTAATTTACCCACAACTCAAAATGCGCCGTAAGAGCGCCAGAAAAAGAAGGTTTCCTGATAATGGAACAGCGTCGTT
TGGCAAGTACTGAATGGGTGGATATTGTCAATGAAGAGAACGAAGTCATTGCACAAGCCAGCCGGGAACAAATGCGGGCACAGTCTCGA
GAGATCCTCTAGATTTAAGAAGGAGATATACATATG
>yfcE
GAGGCAATTCCGACGTCTAAGAAACCATTATTATCATGACATTAACCTATAAAAATAGGCGTATCACGAGGCCCTTTCGTCTTCACCTC
GATGGATCCGCGTCTGTCCAGCGTTTTATTGCACTTTCGGCGAAGCAATCTGGCTGATAGCAACGACAGAATCCAGTATCATAGCTTAC
GTTTTTTCGACGAGGAGTGAGTGATGAAACTGATGTTTGCATCGGACATTCATGGGTCGTTACCGGCGACGGAACGTGTTCTGGAGTTG
TTTGCCCAAAGCGCTCGAGAGATCCTCTAGATTTAAGAAGGAGATATACATATG
>yfcG
GAGGCAATTCCGACGTCTAAGAAACCATTATTATCATGACATTAACCTATAAAAATAGGCGTATCACGAGGCCCTTTCGTCTTCACCTC
GAGACCCAGGCGGATAACACATAAGGGGAGAAAAAGTGGGCATCTGACCAAAGCGTGATAGCGGGTTTACTCATTATGTCCTCGGCAAT
AAAGGGGGCTGGCTATCAAACTATAGAGCCTTTTTGTCACTGTCACGTGACGAATACTCACGTAAGTCGCCGCGCTTTCTATACTTGTG
GTGTTCAGATGATTAAACAACCGGAGCTGCAATGATCGATCTCTATTTCGCCCCGACACCCAATGGTCACAAAATTACGCTGTTTCTCG
AAGAAGCAGAGCTGGGGATCCTCTAGATTTAAGAAGGAGATATACATATG
>yfcU
GAGGCAATTCCGACGTCTAAGAAACCATTATTATCATGACATTAACCTATAAAAATAGGCGTATCACGAGGCCCTTTCGTCTTCACCTC
GATGGATCCTTAAAGCCTGGCTGGTCGGCGCAGCTGATGCGCCAGATCTGGGTAATTTCGAAGCAAATACCACCTTCCAGATTACTTAC
CTGTAAGGCATACCCCCATAGCGACACTGCTATGGGGTATTTAAATGGATTTTGACCGAACAAGAATAAAAACGGATAAAAACGTGTAT
GCCTGACCATTCTCTTTTTCGATTACGGATTCTTCCGTGGTGCATTGCGCTGGCAATGTCAGGGAGTTATAGCAGTGTCTGGGCTGAAG
ACGACACTCGAGAGATCCTCTAGATTTAAGAAGGAGATATACATATG
>yfcV
GAGGCAATTCCGACGTCTAAGAAACCATTATTATCATGACATTAACCTATAAAAATAGGCGTATCACGAGGCCCTTTCGTCTTCACCTC
GATGGATCCCAACTGGCAGATGAGCCCGTGTAATCTGAAGATGGCAAAAGCTATCTGATGAGTTCCAAATTATGCCCAATGAGGTAGAT
TCATTGGGCATAATTGCTCATCAGGCAGCGGATATCTGTCAACCTCACTACTATCGAAGACTCTCCCCATCTATAAATAAAACACCTTT
AACTTTCATTCGGAAAGTATTCTTCTGCGCGTATCTTGATTTTGCCCAAAGTTACATGTTGAAATGAATAATGTCTGTTTGACACTCAT
TATCATTTATTAATACAGATAATGGAGAAAATAGAGTTGAATAATCAGGCAGGACGTAATTGAAACTTGATTAAATTCTGGCTATAGTG
TTTATATATTATATCGACAAAAGCGGGTTAATGCCAATCGCATTATATGTTCATTGTTTGTTTTTTAAGCTATTGGCATTCATTGTTGA
TATGCTTCCTGTTTTTTCTAAATATATATTTCGTGTTTTTTTAGAAGATCCTTAAGATAATCTTTCCCCTATCAATATATTAATTAAAT

CTACATTTACTCATTAGTAAAATTAAATTGAATCAGGGGAGTGAAAACATCACTATTCGACCCCACACCAACGAATACATTTTTTACGA
CCTGTTTTTTTGGATCGCATGCCGTATTCAATGGCTGGGTATAAACATTGGCTATTGACTGAATCAATGTATTGCAATGTTTATATTTT
AATTATTTATAGTTTAAGGACAAATAATGAGTAAGTTTGTTAAAACAGCTATTGCTGCGGCAATGGTGATGGGCGCGTTCACCTCTACG
GCAACAATCGCTGCGGGTAACTCGAGAGATCCTCTAGATTTAAGAAGGAGATATACATATG
>glnK
GAGGCAATTCCGACGTCTAAGAAACCATTATTATCATGACATTAACCTATAAAAATAGGCGTATCACGAGGCCCTTTCGTCTTCACCTC
GAGAGGGACGCTACTGGCAGATGTATCAACTGCAACTTGCGGGCGAAGAGCTGGCAGCCAGCGTGCGTGAAGAGGAATCATTGAGCGCC
TGAATAGCGCAATATTTCATCGTTGGTGCAAAAATGTAACGCACTGTGCACTGTCATAGTGCGTTTTCATTTTCAAACTTCTTAACTTC
CTGCTCTCTTTCTCGTTTTTCATTTCTGGCACACCGCTTGCAATACCTTCTTCGTGTAGCAGAACCATTACCGAATTCTGACCGGAGGG
GATCTATGAAGCTGGTGACCGTGATAATCAAACCATTCAAGCTGGAAGACGTTCGTGAAGCGTTATCTTCCATTGGTATTCAGGGCCTG
ACCGTCACCGAAGTGGATCCCGGATCCTCTAGATTTAAGAAGGAGATATACATATG
>glnH
GAGGCAATTCCGACGTCTAAGAAACCATTATTATCATGACATTAACCTATAAAAATAGGCGTATCACGAGGCCCTTTCGTCTTCACCTC
GATGGATCCGATTGGCGAAGCGAAAGATGACGACACCGCAGATATCCTGACCGCCGCGTCTCGCGACCTGGATAAATTCCTGTGGTTTA
TCGAGTCTAACATCGAATAAATCCATCGCTGATGGTGCAGAACTTTAGTACCCGATAAAAGCGGCTTCCTGACAGGAGGCCGTTTTGTT
TTGCAGCCCACCTCAACGCACTTATTTAGTGCATCCATCTGCTATCTCCAGCTGATTAAGTAAATTTTTTGTATCCACATCATCACACA
ATCGTTACATAAAGATTGTTTTTTCATCAGGTTTTACGCTAAATAATCACTGTGTTGAGTGCACAATTTTAGCGCACCAGATTGGTGCC
CCAGAATGGTGCATCTTCAGGGTATTGCCCTATAAATCGTGCATCACGTTTTTGCCGCATCTCGAAAAATCAAGGAGTTGCAAAACTGG
CACGATTTTTTCATATATGTGAATGTCACGCAGGGGATCGTCCCGTGGATAGAAAAAAGGAAATGCTATGAAGTCTGTATTAAAAGTTT
CACTGGCTGCACTGACCCTGGCTTTTGCGGTTTCTTCTCATGCCGCGGATAAAAAATTAGTTGTCGCGACGGATACCCTCGAGAGATCC
TCTAGATTTAAGAAGGAGATATACATATG
>glnL
GAGGCAATTCCGACGTCTAAGAAACCATTATTATCATGACATTAACCTATAAAAATAGGCGTATCACGAGGCCCTTTCGTCTTCACCTC
GATGGATCCAAGAAGATGACCGCGTGCGTATGACTCCGCATCCGGTAGAGTTTGAGCTGTACTACAGCGTCTAAGTGTTTTAGTTGCCG
TGGAAACTTTTCGCCTGTCTCTGGCAGGCCTGGGATCGGTGGCAAGCACATCACGCCGGATGCGACGCAAATGCGTCTTATCCGGCCTA
CACGGTGATGATGTGGTAGGCCGGAGCAGGTGAGTCGCTCTCCAACGTGAAGTTTGTCAGCTATCTGTAGCCCATCTCTGCATGGGCTT
TTTCTCCGTCAATTCTCTGATGCTTCGCGCTTTTTATCCGTAAAAAGCTATAATGCACTAAAATGGTGCAACCTGTTCAGGAGACTGC
TTTATGGCAACAGGCACGCAGCCCGATGCTGGGCAGATCCTCAACTCGCTGATTAACAGTATTTTGTTAATCGATGACAACCTGGCGAT
CCATTACGCCAACCTCGAGAGATCCTCTAGATTTAAGAAGGAGATATACATATG
>glnB
GAGGCAATTCCGACGTCTAAGAAACCATTATTATCATGACATTAACCTATAAAAATAGGCGTATCACGAGGCCCTTTCGTCTTCACCTC
GATGGATCCAACTGCTTTCCCGACACGAGCTGGATGCAAACGATTTCAAGGAATGAATTGGCGTTATGTGTTACGTTTAGCAGATCAAA
AGACAGGCGACCTTTTCAAGGAATAGCATGAAAAAGATTGATGCGATTATAAAACCCTTCAAGCTGGACGATGTCCGCGAAGCACTGGC
CGAAGTCGGTATTACCGGCATGACGGTGACCGAAGTGACTCGAGAGATCCTCTAGATTTAAGAAGGAGATATACATATG
>glnA
GAGGCAATTCCGACGTCTAAGAAACCATTATTATCATGACATTAACCTATAAAAATAGGCGTATCACGAGGCCCTTTCGTCTTCACCTC
GATGGATCCATGGTCTACGTGCGCGATGATGGCGATATTACGCAATTTTTCGATCACAACTTTGCCTCAGGCATTAGAAATAGCGCGTT
ATTGTACACGGATTAATCGCACTACAAAACAGGATCACAAACATCCTCCGCAAACAAGTATTGCAGAGTCCCTTTGTGATCGCTTTCAC
GGAGCATAAAAAGGGTTATCCAAAGGTCATTGCACCAACATGGTGCTTAATGTTTCCATTGAAGCACTATATTGGTGCAACATTCACAT
CGTGGTGCAGCCCTTTTGCACGATGGTGCGCATGATAACGCCTTTTAGGGGCAATTTAAAAGTTGGCACAGATTTCGCTTTATCTTTTT
TACGGCGACACGGCCAAAATAATTGCAGATTTCGTTACCACGACGACCATGACCAATCCAGGAGAGTTAAAGTATGTCCGCTGAACACG
TACTGACGATGCTGAACGAGCACGAAGTGAAGTTTGTTGATTTGCGCTTCACCGATCTCGAGAGATCCTCTAGATTTAAGAAGGAGATA
TACATATG
>glnW
GAGGCAATTCCGACGTCTAAGAAACCATTATTATCATGACATTAACCTATAAAAATAGGCGTATCACGAGGCCCTTTCGTCTTCACCTC
GATGGATCCTCGAATCCAGGTACCCCAGCCATCGAAGAAACAATCTGGCTACGTAGCTCAGTTGGTTAGAGCACATCACTCATAATGAT
GGGGTCACAGGTTCGAATCCCGTCGTAGCCACCAAATTCTGAATGTATCGAATATGTTCGGCAAATTCAAAACCAACTCGAGAGATCCT
CTAGATTTAAGAAGGAGATATACATATG
>nudE
GAGGCAATTCCGACGTCTAAGAAACCATTATTATCATGACATTAACCTATAAAAATAGGCGTATCACGAGGCCCTTTCGTCTTCACCTC
GATGGATCCAGGCCAGCAAAGCAGCTAAAAAAATCACAATGGTGCTCATGCTTTCCCCATCTTACTTAGTCTGTCAGGCGTGGTCAGTC
GCACCGGACACAATACATGATTCTGCGGTTGGCTATCGGCAAGCGTGGAGTTTAACTTGAGCATCATTAAGCACATCACAGTAACCGCT
GATCATAGCAAACCCTCTGAAGGCGGAATATCAGCAAATTCCTGGAGTTATTTCAACCTATTGACTTTTAATTTGATTTGAGGACGATT
ATTTATTAAGTTACATAAATGTAAATAACATGCAATTCACATTGCCGAAAGCGTGTCCGATATCGCACAATAACCGCTGATTTCACAGC
ATAGATCCCACGATGAGCAAATCATTACAAAAACCCACCATTCTGAATGTTGAAACTGTAGCCCGTTCCCGACTGTTTACCGTCGAGAG
CGTGGATCTCTCGAGAGATCCTCTAGATTTAAGAAGGAGATATACATATG
>glnU
GAGGCAATTCCGACGTCTAAGAAACCATTATTATCATGACATTAACCTATAAAAATAGGCGTATCACGAGGCCCTTTCGTCTTCACCTC
GATGGATCCTATCGCCAAGCGGTAAGGCACCGGTTTTTGATACCGGCATTCCCTGGTTCGAATCCAGGTACCCCAGCCATCTTCTTCGA
GTAAGCGGTTCACCGCCTCGAGAGATCCTCTAGATTTAAGAAGGAGATATACATATG
>cynT
GAGGCAATTCCGACGTCTAAGAAACCATTATTATCATGACATTAACCTATAAAAATAGGCGTATCACGAGGCCCTTTCGTCTTCACGTC
GACGGTTGGGAGACGTGCAACGCACTGGCGGCACGGGTGAAGCTGCCATGTTCAGCCACGGCAAGAAAATAATTGATATGTCGAGAGAG

```
CATTCGCAACCTATAAGTAAATCCAATGGAACTCATCATAAATGAGACTTTTACCTTATGACAATCGGCGAGTAGTCTGCCTCTCATTC
CAGAGACAGACAGAGGTTAACGGTGAAAGAGATTATTGATGGATTCCTTAAATTCCAGCGCGGGATCCTCTAGATTTAAGAAGGAGATA
TACATATG
>cynR
GAGGCAATTCCGACGTCTAAGAAACCATTATTATCATGACATTAACCTATAAAAATAGGCGTATCACGAGGCCCTTTCGTCTTCACCTC
GATGGATCCCGCGCTGGAATTTAAGGAATCCATCAATAATCTCTTTCACCGTTAACCTCTGTCTGTCTCTGGAATGAGAGGCAGACTAC
TCGCCGATTGTCATAAGGTAAAAGTCTCATTTATGATGAGTTCCATTGGATTTACTTATAGGTTGCGAATGCTCTCTCGACATATCAAT
TATTTTCTTGCCGTGGCTGAACATGGCAGCTTCACCCGTGCCGCCAGTGCGTTGCACGTCTCCCAACCCTCGAGAGATCCTCTAGATTT
AAGAAGGAGATATACATATG
>rne
GAGGCAATTCCGACGTCTAAGAAACCATTATTATCATGACATTAACCTATAAAAATAGGCGTATCACGAGGCCCTTTCGTCTTCACCTC
GATGGATCCCTCTGCGCCTCTTTGAGCTGACGATAACCGTGAGGTTGGCGACGCGACTAGACACGAGGCCATCGGTTCACACCCGGAAA
GGCGTTACTTTGCCCGCAGCTTAGTCGTCAATGTAAGAATAATGAGTAAGTTACGATGAAAAGAATGTTAATCAACGCAACTCAGCAGG
AAGAGTTGCGCGTTGCCCTTGTAGATGGGCAGCGTCTGTATGACTCGAGAGATCCTCTAGATTTAAGAAGGAGATATACATATG
>glxK
GAGGCAATTCCGACGTCTAAGAAACCATTATTATCATGACATTAACCTATAAAAATAGGCGTATCACGAGGCCCTTTCGTCTTCACCTC
GAGACGCTTCGTCCTCTGCTCAGTAACGGCTTATTGGTCGGGATTTTACTGGCTGTTCTTATGGATAACCTTATTCCGTGGGAACGCAT
CGAATAATTTGTTGAAAAAGGATTGATAATGAAGATTGTCATTGCGCCAGACTCTTTTAAAGAGAGCTTAAGTGCAGAAAAATGTTGTC
AGGCAATTAAAGCCGGGTTTTCGACCCTCTTGGATCCTCTAGATTTAAGAAGGAGATATACATATG
>kil
GAGGCAATTCCGACGTCTAAGAAACCATTATTATCATGACATTAACCTATAAAAATAGGCGTATCACGAGGCCCTTTCGTCTTCACCTC
GATGGATCCTAGCCGAGACCACCGAAGCCCGGAGGTGGTGAAATAAAACCGGGCACAACACGAAGGCGCATTTCCGATATCCATAAAGA
GTCGGTCTTGTCTGTTAAATTTAAATGGTGGGAGTGCGCCTCCGGTTGTAAATAACGACATTGCTGTGTGTAGTCCTGGCGGCATCAGT
TTTTTTCTTGAAGTTCGGCTGATGTCCGCCCTTTTTAAAGTGAATTTTGTGATGCGGTGAATGCGGCTAAGCGCACGTGGCACAGTTAA
AAGTCATGTTAGTCCTTATTGGTTTGGGTGGGAAAGCCGACTGTAATTGTTAACTGGTTGCAGTCACCTGGAGGCACCAGACACCGCAT
CAACAAAGTTCATTTGTAAAAATGGAGATAATTATGATTGCACATCACTTCGGAACTGATGAAATACCACGTCAGTGTGTGACTCCTGG
CGACTCGAGAGATCCTCTAGATTTAAGAAGGAGATATACATATG
>gssA
GAGGCAATTCCGACGTCTAAGAAACCATTATTATCATGACATTAACCTATAAAAATAGGCGTATCACGAGGCCCTTTCGTCTTCACCTC
GATGGATCCGACTTTCGCGGGCTGATAAGTATTGTCTGTCATAGTGAGTTGCCTTCTGAGTGGTAGTGTTGAAGCAGTGTAGCAGGTCA
CTTGCTGTACAGAATGTGAATTTTTTAACCCTTTGCGCCAGCCGCTGCGATGAGGTGTATGATTAGTCTGACCTCTGTCCGGTTAATGT
CGCTGAGCAGGTAATCAGAGCACAACAATTACGCCAACTGCTTAGGTTGTATTGTTAAAGGTAAAGTGATGAGCAAAGGAACGACCAGC
CAGGATGCCCCGTTCGGGACATTATTGGGCTACGCCCCAGGTGGGCTCGAGAGATCCTCTAGATTTAAGAAGGAGATATACATATG
>yneJ
GAGGCAATTCCGACGTCTAAGAAACCATTATTATCATGACATTAACCTATAAAAATAGGCGTATCACGAGGCCCTTTCGTCTTCACCTC
GAGCCGTGGCAGGATTTATCGAAATTGCATGAGTTGCCGGAGTAATGGTCATCGGGGTATCTCCTTTATGAGTCATGGTATGAAGATAC
GCAGATTTACTCTTGCTTTAAAATGAATAATATTAAGCCACTTATTCACGAATCGAGAATGCTATGGATCTGACCCAACTGGAGATGTT
CAACGCGGTTGCCGAAGCTGGCAGCATAACCCAGGCTGCAGCAAAAGTGGATCCTCTAGATTTAAGAAGGAGATATACATATG
>yneK
GAGGCAATTCCGACGTCTAAGAAACCATTATTATCATGACATTAACCTATAAAAATAGGCGTATCACGAGGCCCTTTCGTCTTCACCTC
GAGCGTCGTGGTGCGAAAACACGTCCGCTCGAGGCATTTATTCAACTGCTGGATGTGCCTGACTCGGCAAGACAGGGATATCAATGAGC
TATTTTTGATAGTTCTTGCGGTTAATATGCTCTATATAGTGATGTTCCGATGACTTATGACTATATGGGGCAAATATGGTTACGCCAGT
AAGCATCAGCAGGATCCTCTAGATTTAAGAAGGAGATATACATATG
>yneH
GAGGCAATTCCGACGTCTAAGAAACCATTATTATCATGACATTAACCTATAAAAATAGGCGTATCACGAGGCCCTTTCGTCTTCACCTC
GATGGATCCCCCATTTCGGCTTACACGAATTCTGTAATATCCAGACGGTGTGGAAAGACCGGATCTGACGCGACATATTCAGCTCTGAT
ATACTCGCAGGTCTTTTCAGACCTGCGGTCCAGGAGTAGAAAGTGGCAGTCGCCATGGATAATGCAATTTTAGAAAACATCTTGCGGCA
AGTGCGGCCGCTCATTGGTCAGGGTAAAGTCGCGGACTCGAGAGATCCTCTAGATTTAAGAAGGAGATATACATATG
>yneI
GAGGCAATTCCGACGTCTAAGAAACCATTATTATCATGACATTAACCTATAAAAATAGGCGTATCACGAGGCCCTTTCGTCTTCACCTC
GATGGATCCACTTTTGCTGCAGCCTGGGTTATGCTGCCAGCTTCGGCAACCGCGTTGAACATCTCCAGTTGGGTCAGATCCATAGCATT
CTCGATTCGTGAATAAGTGGCTTAATATTATTCATTTTAAAGCAAGAGTAAATCTGCGTATCTTCATACCATGACTCATAAAGGAGATA
CCCCGATGACCATTACTCCGGCAACTCATGCAATTTCGATAAATCCTGCCACGGCTCGAGAGATCCTCTAGATTTAAGAAGGAGATATA
CATATG
>amyA
GAGGCAATTCCGACGTCTAAGAAACCATTATTATCATGACATTAACCTATAAAAATAGGCGTATCACGAGGCCCTTTCGTCTTCACCTC
GAGCGGTGCAAAAATCGGTGTTAAGTGCCTATGGCGATCAGGGCGGCTTTGTGCTGGCTCCGCAGGATAACCTCTTTTGAATCTGAATG
AGTCGATGGCTCGCGAATAATCCGATTACGGCTACGCTTCTAATGTTCCCCTTGAATGGAGTCGAAGAATGCGTAATCCCACGCTGTTA
CAATGTTTTCACTGGTATTACCCGGAAGGCGGTAAGCTCTGGCCTGAGGATCCTCTAGATTTAAGAAGGAGATATACATATG
>yneE
GAGGCAATTCCGACGTCTAAGAAACCATTATTATCATGACATTAACCTATAAAAATAGGCGTATCACGAGGCCCTTTCGTCTTCACCTC
GATGGATCCGACGCAAGTGCCTGGTCTGGTCGAGCAGGTTGCTAATGACCTGGATGCGATTCTCGAAAAAGGGATGCGCGAAGCTGTAC
GTCCGTTGTGCTAATAACACCGTTACTTCTGAACCCGGTTTCGACCGGGTTTCTTATTTCACCATATTAGTTACAACATCCTATATGCC
```

TTTATCATTAACAGGTGCCACATGCAGCGCCTGCTAATGTTGTAAGCTTGTAGGTAATTTGTAAGTAGCGTCTTGAGAATGTGATCGTA
ATCACGTTTAATAGGGCGACTTTTCCACGCCTGAAGATAACCATGATTGTTCGTCCACAACAACACTGGCTGCGCCGTATTTTTGTCTC
GAGAGATCCTCTAGATTTAAGAAGGAGATATACATATG
>yecM
GAGGCAATTCCGACGTCTAAGAAACCATTATTATCATGACATTAACCTATAAAAATAGGCGTATCACGAGGCCCTTTCGTCTTCACCTC
GATGGATCCTCATGGCCTGACGGACTTTTTCTGAGAGAAGAGCCTGAATATTCACCGGAATACCTTACATTGATGAAGCGAGCAAAAAT
TGCCCGCACCAGTTAAAGAAATTAGGGCGGGAGTATACTGCAAATGGCTGTTTTCGTCAGCAACGTGCGCGTGGCGTATTGCCGTTTTC
ACTATTGCATCAGAAGCGGCGTGAAATCAGCGGTTGGTTGGCACAGGGCAACAGGGTAAATTAACGCTTTTACTGAAAGACGAAGAGTG
ATTATGGCGAACTGGCAATCTATCGACGAGCTGCAGGATATTGCATCGGATTTACCGCGATTTATCCACGCATTAGATGAACTTTCCCG
CCGTCTGCTCGAGAGATCCTCTAGATTTAAGAAGGAGATATACATATG
>yfeA
GAGGCAATTCCGACGTCTAAGAAACCATTATTATCATGACATTAACCTATAAAAATAGGCGTATCACGAGGCCCTTTCGTCTTCACCTC
GATGGATCCATCCCGCTTAGCTCCACCAAATTTCCAACCCTCGCTGCAAAGCGGGGGTTTTTTGTCTCTGCTTTTTGCCGCTTTTGTAA
TACAGTCTACGTCCGGGTTAGTGCCGCCTGGTGAAAGCATCATTGGATGAAAAATCGGCAACAGGCTGGCCCCCTGTTTGCTTCGCGAT
GCGAATAAACTTATTATTTGTGTGCCTGAAAACCCCCTCGAGAGATCCTCTAGATTTAAGAAGGAGATATACATATG
>yecH
GAGGCAATTCCGACGTCTAAGAAACCATTATTATCATGACATTAACCTATAAAAATAGGCGTATCACGAGGCCCTTTCGTCTTCACCTC
GATGGATCCTTCCCGCCACGATAAAAACACTTCCCAGGGTTCTGTTTTTCACGCTTTCTTCTGTCCTGACGATCTTTATGAGTGATATC
TGCGGCGCAGGTTAAGGCATAACGGCTGCTTCGTCAAACCGACGTTACATATGGTGTAAATATAAATGTACAATAAAAAGCATTCAGTC
CTGCGCTGGCGCAAAGTCTTCATAAGCGGCAATCGCTACGCTAGGCTTTTTTTGAGGAGGGACGATGGACTCTATTCACGGTCATGAAG
TGTTAAATATGATGATTGAATCAGGCGAGCAATATACGCATGCCAGTCTGGACTCGAGAGATCCTCTAGATTTAAGAAGGAGATATACA
TATG
>yecF
GAGGCAATTCCGACGTCTAAGAAACCATTATTATCATGACATTAACCTATAAAAATAGGCGTATCACGAGGCCCTTTCGTCTTCACCTC
GAGGATGCCTCACCGACGACTTTTATACCCTTTATATCTTCCAGAATGCGTCGTATCCCTGCGCGCACCAGTTCGTGGTCATCAACAAG
TAGAACGTTGATCAAAGGAATATCTCCAGAAATAGGGATAACGCTACTGATAGTTAGTCATTCGTATATTAGCGGTTTTTATTGCAACT
TTGAAACGTTAAAAATGGTGGGTATTCGATTTTTCTCTCGTTTTTGTCAATTCGATTGTCCACAGTTATGGTCACGCCCGTCACCAACG
GCTTACCGGATCGTTTCACAGTAAGCCCTACGATGAAATGTGACAAAAATTGACTTTATTCAGCAAAAATGAAAATCAGCCCGCTGAAT
ATGTAACATTAATTAACCCGAACAATACGGGCAAAAAACATTCATGATAAAAATATTTATTGGTCATTATATTAACGTATTTTATAGCA
CTGCCGATATCACGCTCAAAAAACAACCACTGCTATTTTTAGCAAAGCTTATGGTATACTCCGCCGCCTTAACATTTTTCACCGCAAAT
TTTCATTGCAACATGACGAGGAAAATAAATGAGTACGCCTGATTTTTCTACTGCCGAGAATAATCAAGAACTGGCAAATGAAGTCTCCT
GCCTGAAAGCGATGCTGGATCCTCTAGATTTAAGAAGGAGATATACATATG
>fadI
GAGGCAATTCCGACGTCTAAGAAACCATTATTATCATGACATTAACCTATAAAAATAGGCGTATCACGAGGCCCTTTCGTCTTCACCTC
GAGAAAAGCCGTCGCCTGACGGGCAAAAGGCGTACGTAAACCGCTAACAATGGCGATACGATCGCCCTGGCGGGTAACCAGCGGTAAAA
CCTGACCCATAACACTCCCCTGTAAAAAAATAAATAAAGTGGTCTGACCTGATCATAGTCTTAACCATTTTTTTACATTTAGCCAAGTG
GAGAAAAGGGAAAGTGGGAGCTATGACACAGAGAGAAAAGAAGAAGAGAAAAGAAAACGCCCCTGCCAGCAACTGACAGGGGCGTACGT
ACCCGAGAGGAATTAACGCAGACCCAGCTGGAAAAGGATCCTCTAGATTTAAGAAGGAGATATACATATG
>yjjU
GAGGCAATTCCGACGTCTAAGAAACCATTATTATCATGACATTAACCTATAAAAATAGGCGTATCACGAGGCCCTTTCGTCTTCACCTC
GAGCCGTCGATTCTCAGGCACAAAGTGACCGTGCTGAAAGTATCGCCAAAGCGGTAGATGGTGTGAAAAGCGTTAAAAATGATCTGAAA
ACTAAGTAATTCGTCGTAATTCGTCCTCCCGAACTTTGTCGGGAGGCGCGATGTGCACCACACTAAAAATATCGCATTGAGTGATTTTC
ACGCTCATATCAAGCGGTGACATTAACTATGGTAAAGGAGACGCTTATGTTTCGTTGGGGCATCATATTTCTGGTTATCGCGTTAATCG
CCGCCGCACTTGGGTTTGGTGGTCTGGCCGGTACCGCTGCAGGCGCAGCTAAAATTGTCTTTGTCGTCGGGATTATTCTGTTCCTGGTG
AGTTTGTTCATGGGCCGAAAACGACCCTAGATTTCAGACCATTAAGAAAATTTTATCCAAAGCCAGTCCAGCGGACTGGCTTTTGCGGT
TTTAGCGAATAATAAATTGCGTTACTTTGTCATTCTTTCACAACAAAAACAGGAAGGCAGAGGTGGGGCAGCGAATACCTGTAACGCTT
GGTAATATTGCGCCGTTGTCGCTAAGGCCGTTCGGATCCTCTAGATTTAAGAAGGAGATATACATATG
>ychJ
GAGGCAATTCCGACGTCTAAGAAACCATTATTATCATGACATTAACCTATAAAAATAGGCGTATCACGAGGCCCTTTCGTCTTCACCTC
GATGGATCCAGCGCCAGCCCTATTTTTATCTTTCTCATTATTCCCGCAAGATTCCCCTGGAATGCAATCGTAGCCACATCGCTACTGCT
CGGTTAACATAGTGCTACTCAGGCGGTTAATACGCCGTTATTGTTTCCAGGGAGATTATTTGTGTCTCAGCTTTGTCCCTGTGGTAGTG
CTGTCGAGTATAGCCTATGTTGCCACCCTTATGTGTCTGGTGAAAAGGTTGCACCCTCGAGAGATCCTCTAGATTTAAGAAGGAGATAT
ACATATG
>rnk
GAGGCAATTCCGACGTCTAAGAAACCATTATTATCATGACATTAACCTATAAAAATAGGCGTATCACGAGGCCCTTTCGTCTTCACCTC
GATGGATCCCTTGCCACAACCTCACCCAGGTAACTGTGGCAAAACCTTTGTGATTGATAAAGCGGGTTATTAATCATCTGATTTACAAA
ACACGCTAATTCGTCTTCCATAAACCAAGGCGATTAAGATTTATACATAATGCGCTGCATTGAGGAATTCCCTGAATGTGACGCAAATC
ACTTCAAAGAGTGAGCAACAGTCAGTATCATCATGTGAGTTAAACCCTCGCCGCCTGACGGTGAGGGTTTTCTTTTGGGTATGTTTATC
TGCGACACTCGCAGCACCGACGACATGGAGTAAAAATGTCCAGACCAACTATCATCATTAACGACCTGGATGCCGAACGCATCGATATT
CTGCTGGAGCAACCCGCCTATGCTGGTTTGCCAATCGCTCGAGAGATCCTCTAGATTTAAGAAGGAGATATACATATG
>astC
GAGGCAATTCCGACGTCTAAGAAACCATTATTATCATGACATTAACCTATAAAAATAGGCGTATCACGAGGCCCTTTCGTCTTCACCTC
GATGGATCCCGATGGCTTCAAGCTGGTGAGGTCTGGCGCGCAGGCCGTTGATATTAAAAGAGACAAATTTCATAGTCGCTGCCATTTAT

Data Set S4 - MG1655 library reference file.fasta

```
TCAGATAAAGAGTGCGTGGATGGTAGCAGAATTTCGCGTTGCTTACCGCCTGTTGTTAAGTGATTTGATCGATGGTTGTTAATGATGTC
AACGATGGCGCAAAAAATGCCCGCTTTTGGTGCGCGCTGCGTCAGAATGGCGCAGTAATTTCCAGTAAATTTCGTCAATGATCACAATT
CCAGAATTATATTTTACTTCTGCATAACATTGCGTTTTTATTCTTTTTATTCACCCACATCGCAAACGTATTCACTTTATATGCACTT
TAAATGCATATGGTTTGGTTATAACTCCTTGATTTCCAGTTAGCCTCCGCCGTTTATGCACTTTTATCACTGGCTGGCACGAACCCTGC
AATCTACATTTACAGCGCAAACATTACTTATTATTAACATATAAATAACGAATTATTTACTGTAGAGGTCGCTATGTCTCAGCCAATTA
CGCGTGAAAACTTTGATGAATGGATGATACCTGTTTACGCTCCGGCACCCTTTATACCGGTACGTGGCGAAGGTTCCTCGAGAGATCCT
CTAGATTTAAGAAGGAGATATACATATG
>nudH
GAGGCAATTCCGACGTCTAAGAAACCATTATTATCATGACATTAACCTATAAAAATAGGCGTATCACGAGGCCCTTTCGTCTTCACCTC
GATGGATCCCGGGAGGAGAGAGCAGTGGGCGAGGTTGGGACATGTCATGATACCTTGATTAAAAAACAAACAGCAATTATGTGCCGAAT
ATTCGCAGTGTAACTTTTTCCACATGCGTTGTTCAATTCTCTGATTTATATTGCTTTTTAGGTTGCCGGTAATCGAGTTGCGCAACACT
GCAACGGGTTTTCCAAATCTGGTCACTGAAAATAAACAGGATCTTACACCATGTTATCCACAGAAAGCTGGGATAACTGTGAAAAACCT
CCACTACTGTTTCCATTTACAGCCTTGACGTGCGACGAAACTCCAGTTTTTAGTGGTTCAGTGCTTAACTTAGTGGCACATTCTGTGGA
TAAATACGGCATTGGTTGATCTTTCGCCAACCCGTAAATGAGAGGTGTGACCTGGGTCACGAATTACTCTTTTTCATTACTAAAATTAC
ATGTAATGCATTGACTGATAATGAATATTTTAAAATGTCTATATCGGGCTTATTCAGAATGCTCCGGGTTTTCCCGTAGTAATTCCGTA
GTTCTTCACAACTCTATCCACAGAAAAGGTGAATAAAAACGGCTATCCACCCCTTCCTCTGTTTATAACTCTGCACATAACTGTGAGTT
ATTCAATTGTTATTAGTGTGTAATGCCCTGCAGAGAGTGGTTTACCGTCTTCCTGGAGTATGAAACAATCATTCGTATATAAAGCTTTA
TTTTGAGGTAGTCCGGTGATTGATGACGATGGCTACCGCCCAAACGTAGGTATCGTGATTTGTAATCGCCAGGGGCAGGTAATGTGGCT
CGAGAGATCCTCTAGATTTAAGAAGGAGATATACATATG
>nudG
GAGGCAATTCCGACGTCTAAGAAACCATTATTATCATGACATTAACCTATAAAAATAGGCGTATCACGAGGCCCTTTCGTCTTCACCTC
GAGTATCAAGAACCCGCACGCACTGATGCAACAACGGTTTAATCCGGGGATGAAGATGGCGGTCTAGCACAGGCACTCCTTAAATATAA
AGCCTTTCTGATTGAGCAACAGTGCGGATATTATGGCATTTTTCGCTTATCTGCCCGTGTGTAATTTATGAAAATGATTGAAGTTGTTG
CCGCCATCATTGAACGTGATGGCAAAATTTTACTCGCGCAACGCCCCGCCCAGAGCGATCAGGCGGGATTATGGGATCCTCTAGATTTA
AGAAGGAGATATACATATG
>nudF
GAGGCAATTCCGACGTCTAAGAAACCATTATTATCATGACATTAACCTATAAAAATAGGCGTATCACGAGGCCCTTTCGTCTTCACCTC
GATGGATCCAGAAAGGCTCAGGCCGATAAGAATGGGGAGCAATTTCTTCATTTGCATTCCTTGTGGTGAAGCAGTATTTAGCGCGATCA
AACTGTAAAATTATTGCGGATTCTAGCAGAAGCCGCTACCGCAAAAGGTTGGCGTTACGTGCCATAGGGCGTTAATGTGCCTCAATTTA
ACACATGACTGCTCAAACGGCAGTCAAACGTCGCTGAAATTCACATTTAATTCACTATTAGTGCCAGGACATTAACAATGCTTAAGCCA
GACAACCTGCCCGTTACATTTGGCAAAAACGATGTAGAAATTATTGCACGAGAAACACTTTATCGCGGCTTCTCGAGAGATCCTCTAGA
TTTAAGAAGGAGATATACATATG
>rna
GAGGCAATTCCGACGTCTAAGAAACCATTATTATCATGACATTAACCTATAAAAATAGGCGTATCACGAGGCCCTTTCGTCTTCACCTC
GATGGATCCATTACTGGCGTCTTGGCGCAATCTTCGGGGTGATTTACATCTCTATGTTGCTGTTGGTTGGCTGGCCGATTCTCGCCATG
TGGAACTAATACGTTCGCCAAACGCTGAATAAATCATCACGCCCGCCAGGTAAGCCCACCTGGCGGGCTTTTTTATGATTTAATAGATA
GTATTCAGTGTATGTATGAGTTCCACACCCATTATGAAAGCATTCTGGCGTAACGCCGCGTTGCTCGCGGTTTCTCTGCTTCCCTTCTC
TTCTGCCAACGCCCTCGAGAGATCCTCTAGATTTAAGAAGGAGATATACATATG
>yliG
GAGGCAATTCCGACGTCTAAGAAACCATTATTATCATGACATTAACCTATAAAAATAGGCGTATCACGAGGCCCTTTCGTCTTCACCTC
GATGGATCCAACCGGTTCAGCAGATCGATTCGCTGTCTGTCAACGAACATACTTCGATCCTCCTCTTCCCAGCCTTGTTGCAATTAAAT
AGTATGGCTATTTTTTGAGCAAAATGCTGTCCAGGATCAATTCTTCACGCCGAATTAACAAAATTTACATTTCCTTCGCCTCCGCCAGC
GAATGAGATTGCGATACAAATTTTATCCAGAACTCTGTAGAATCCCTGCCCTGACCTTTCCTGTAACTGAATATTTTTTAAGCTATGAG
CAAAGTAACTCCCCAGCCGAAAATCGGCTTTGTTTCCCTTGGCTGTCCGAAAAACCTTGTTGATTCAGAGCGTATTCTCACCGAACTCC
GCACTCTCGAGAGATCCTCTAGATTTAAGAAGGAGATATACATATG
>nudC
GAGGCAATTCCGACGTCTAAGAAACCATTATTATCATGACATTAACCTATAAAAATAGGCGTATCACGAGGCCCTTTCGTCTTCACCTC
GAGAGCCACGAGCAGATGCTTACGTACATGTAGCCAGCGATCAACCAGTTTGTTACTTCCTCTGACGCGCTCCGTCAGGTTATCGAGCT
GGTTAAGCATGATTGACTCCGCAAGTTTGTATTCAAAAACTGCTCAGTGAGAAATGTAAAAACCATGTTAAACATGCCAGTGATGCAAA
GGTAGTGCAAGAGCTATGGATCGTATAATTGAAAAATTAGATCACGGCTGGTGGGTCGTCAGCCATGAACAAAAATTATGGTTGCCGAA
GGGAGGATCCTCTAGATTTAAGAAGGAGATATACATATG
>nudB
GAGGCAATTCCGACGTCTAAGAAACCATTATTATCATGACATTAACCTATAAAAATAGGCGTATCACGAGGCCCTTTCGTCTTCACCTC
GATGGATCCCGAGCTTTGCTAACCCGACTGCACTGGCTGAGCTGAGCATTCAGGTTGTGAAGAAGGCTGAGAATAACTGATATGACTCA
AATACACGAAATCATTCGCGTTGCATCGAGGCGGCAACTGAGTGAACTCCCATGAGCATAGATAACTATGTGAATGGGATGAGCGAAGG
CAGTCAACGAAGAGGCAGCGTGAAGGATAAAGTGTATAAGCGTCCCGTTTCGATCTTAGTGGTCATCTACGCACAAGATACGAAACGGG
TGCTGATGCTCGAGAGATCCTCTAGATTTAAGAAGGAGATATACATATG
>gadB
GAGGCAATTCCGACGTCTAAGAAACCATTATTATCATGACATTAACCTATAAAAATAGGCGTATCACGAGGCCCTTTCGTCTTCACCTC
GATGGATCCACCGGAGTATTATTGCCAAAATAATAACAGCCCCGTCAACACATCGTTGGCGGGGATTTTAGCAATATTCGCTATTTTTA
TGTAATAATTTTATAAATGCGTTCAAAATAATAATCAAGTACTAATAGTGATATTTTAAGGTCTGATTTTTACGTGATAATTCAGGAGA
CACAGAATGCGCATAAAAATAACAGCATAAAACACCTTACCACCACCCAAGAATTTCATATTGTATTGTTTTTCAATGAAAAAATATTA
TTCGCGTAATATCTCACGATAAATAACATTAGGATTTTGTTATTTAAACACGAGTCCTTTGCACTTGCTTACTTTATCGATAAATCCTA
```

```
CTTTTTTAATGCGATCCAATCATTTTAAGGAGTTTAAAATGGATAAGAAGCAAGTAACGGATTTAAGGTCGGAACTACTCGATTCACGT
TTTGGTGCGAAGTCTATTTCCACTATCGCAGAATCAAAACGTTTTCCGCTGCACGAAATGCGCGACGATGTCGCATTCCAGATTATCAA
TGACGAATTATATCTTGATGGCAACGCTCGTCAGAACCTGGCCACTTTCTGCCAGACCTGGGACGACGAAAATGCTCGAGAGATCCTCT
AGATTTAAGAAGGAGATATACATATG
>rnd
GAGGCAATTCCGACGTCTAAGAAACCATTATTATCATGACATTAACCTATAAAAATAGGCGTATCACGAGGCCCTTTCGTCTTCACCTC
GATGGATCCGCGGCAAAGTGGACAATAAAGCCTGAGCGTTAAGTCAGTCGTCAGACGCCGGTTAATCCGGCGTTTTTTTGACGCCCAC
TAAAGAGAAAACAATTTGAATTACCAAATGATTACCACGGACGATGCGCTGGCTTCTTTGTGTGAAGCCGTCCGTGCCTTTCCGGCGAT
AGCTCGAGAGATCCTCTAGATTTAAGAAGGAGATATACATATG
>gadX
GAGGCAATTCCGACGTCTAAGAAACCATTATTATCATGACATTAACCTATAAAAATAGGCGTATCACGAGGCCCTTTCGTCTTCACCTC
GATGGATCCATTGCCCAGCAGAACAGCTTTAAACACACCTGATAACATAACGTTGTAAAAACCGAATGCCCAGCCTTTAAAAAAACAGC
TGGGCATTCGGTTGCTTATTAATCGCAATAATATATTGGCTGTCTATTCATCGTGTTGATAAGATATATGTTATATGAATGTTTATTAA
ATGTAGCGATTACCTTTAATGTCACGAAAAGGGATTATTTGCTTACTATTAATTTCCCTGTGTTCTTAAACACGTTTTTGTATTATAAA
TCCAGTCATCCTGCCCGTAGAACATCACACATTATCATCCTGTTCTCCCGCTTCGTTTAAATTTATTTATCAATCAATTTGACTTAAGA
GGGCGGCGTGCTACATTAATAAACAGTAATATGTTTATGTAATATTAAGTCAACTATGCAATCACTACATGGGAATTGTCTAATTGCGT
ATGCAAGACATAAATATATTCTCACCATGGTTAATGGTGAATATCGCTATTTTAATGGCGGTGACCTCTCGAGAGATCCTCTAGATTTA
AGAAGGAGATATACATATG
>gadW
GAGGCAATTCCGACGTCTAAGAAACCATTATTATCATGACATTAACCTATAAAAATAGGCGTATCACGAGGCCCTTTCGTCTTCACCTC
GATAGATCTGGCACGGGAAACTTTGTGCTCTCAGTAAGTTAAATATAACTTTTACTGGAAATAAGATCAGCCATTTTTTTATAAACATA
AGCTATACGCTGTGCGAAAATATATTCGTGCTGCATTTACTTATTATCAATTAACTGTTATGCAAAACTACTTTGTGGATAAATTTTGG
TCCTACCAAATCTGGCAGTTTTTGCGCTAAGAAACAGTCTGGCATCATTTCATTAGTATACTGAAATTGAAATAATCGCAGTATGAAAT
ATAAGGGATAATCATGACTCATGTCTGCTCGGTGATCCTCATTCGTCGTTCATTCGATATTTATCATGAACAGCAAAAAATATCGCTGC
ATAACGAGAGTATTCTGCTGCTGGAGAAAAATTTGGCAGACGATTTTGCGTTTTGTTCACCGGATACGCGACGACTGGATATCGATGAG
CTGACAGTTTGCCATTACTTACAAAATATTCGTCAGCTACCACGCAATTTAGGGTTACACAGCAAAGACCGTTTGTTAATTAACCAGTC
ACCCCCCATGCCGCTGGTGACGGCGATTTTTCTCGAGAGATCCTCTAGATTTAAGAAGGAGATATACATATG
>rnt
GAGGCAATTCCGACGTCTAAGAAACCATTATTATCATGACATTAACCTATAAAAATAGGCGTATCACGAGGCCCTTTCGTCTTCACCTC
GAGGCGTTTGTGGAAGATCCGGACGGTTACAAAATTGAGTTAATCGAAGAGAAAGACGCCGGTCGCGGTCTGGGCAACTAATCTCCTGC
CGGGCGTGAACTCATCGCGCCCGCATCTTTACTGCATCGACAAGTAATATTTGTCATAATGCGCGCTGCAATTTATCCGTATTAAGAGA
ATCAGATGTCCGATAACGCTCAACTTACCGGTCTGTGCGACCGTTTTCGTGGTTTTTATCCTGTTGTGATCGATGTTGAAACAGCCGGA
TTTAACGCGGATCCTCTAGATTTAAGAAGGAGATATACATATG
>stpA
GAGGCAATTCCGACGTCTAAGAAACCATTATTATCATGACATTAACCTATAAAAATAGGCGTATCACGAGGCCCTTTCGTCTTCACCTC
GATGGATCCGCAACTGCATGACGCAAGCGTGACTGCGGTGAGAACATGCTAATGCTCCTTAATAAAAGCGTAATGGGATGTTAATGGAG
ATGAAAGCAGTGGATCTCATTGCCATCTGCATTTTCGCATAGCGTATATCGCTGGCTATGACTAATCAATTAGTGAAAAAATTTTTATTT
AGTCAGTTTTTGTAGATAAAATATTCCGTAACGGCTCGTTTTTTGTACATTGTGGATTTTTACATGCCGATATGATTTATTTTTAATTA
AATCAATTAGATATGGCCTTACCTTTAATATCACTAATGAAAGTTTGGGGGTGAGGTAACGCTATAAGCGTAGAGAATTCGGTGTATGT
AGTAGCTGGGTCATTTTCAGCATTTTTTATGCCAGACAATGTACATTAACAGCAGAAAGCACCAGTGATGGATGTTTTCTTTTATTCTG
TTATATTGCATTTTTATTCACTTTGATTTTGTCAGGCTTGCGGAATTAGCGAGCAGAGAGCGCCGCTCTGTTCACCAATAATCTAAATT
ACTCATCAAAATGGCTAATATATAATCTTGAAATTATTCTGAGAGTTTCAGAAAATAACAGCTGAAATAATCTCGCGCAGGACTGTAAA
TAGATTAAATTTTGTGGAAATATAATAAGTGATCGCTTACACTACGCGACGAAATACTTTTTTTGTTTTGGCGTTAAAAGGTTTTCTTT
ATTATGTCCGTAATGTTACAAAGTTTAAATAACATTCGCACCCTCCGTGCGATGGCTCGCGAATTCTCCATTCTCGAGAGATCCTCTAG
ATTTAAGAAGGAGATATACATATG
>araB
GAGGCAATTCCGACGTCTAAGAAACCATTATTATCATGACATTAACCTATAAAAATAGGCGTATCACGAGGCCCTTTCGTCTTCACCTC
GATAGATCTGCCTCAATCGGCGTTAAACCCGCCACCAGATGGGCGTTAAACGAGTATCCCGGCAGCAGGGGATCATTTTGCGCTTCAGC
CATACTTTTCATACTCCCACCATTCAGAGAAGAAACCAATTGTCCATATTGCATCAGACATTGCCGTCACTGCGTCTTTTACTGGCTCT
TCTCGCTAACCCAACCGGTAACCCCGCTTATTAAAAGCATTCTGTAACAAAGCGGGACCAAAGCCATGACAAAAACGCGTAACAAAAGT
GTCTATAATCACGGCAGAAAAGTCCACATTGATTATTTGCACGGCGTCACACTTTGCTATGCCATAGCATTTTTATCCATAAGATTAGC
GGATCCTACCTGACGCTTTTTATCGCAACTCTCTACTGTTTCTCCATACCCGTTTTTTTGGATGGAGTGAAACGATGGCGATTGCAATT
GGCCTCGATTTTGGCAGTGATTCTGTGCGAGCTTTGGCGGTGGACTGCGCTACCGGTGAAGCTCGAGAGATCCTCTAGATTTAAGAAGG
AGATATACATATG
>araC
GAGGCAATTCCGACGTCTAAGAAACCATTATTATCATGACATTAACCTATAAAAATAGGCGTATCACGAGGCCCTTTCGTCTTCACCTC
GAGCTTCACCGGTAGCGCAGTCCACCGCCAAAGCTCGCACAGAATCACTGCCAAAATCGAGGCCAATTGCAATCGCCATCGTTTCACTC
CATCCAAAAAAAACGGGTATGGAGAAACAGTAGAGAGTTGCGATAAAAAGCGTCAGGTAGGATCCGCTAATCTTATGGATAAAAATGCTA
TGGCATAGCAAAGTGTGACGCCGTGCAAATAATCAATGTGGACTTTTCTGCCGTGATTATAGACACTTTTGTTACGCGTTTTTGTCATG
GCTTTGGTCCCGCTTTGTTACAGAATGCTTTTAATAAGCGGGGTTACCGGTTGGGTTAGCGAGAAGAGCCAGTAAAAGACGCAGTGACG
GCAATGTCTGATGCAATATGGACAATTGGTTTCTTCTCTGAATGGTGGGAGTATGAAAAGTATGGCTGAAGCGCAAAATGATCCCCTGC
TGCCGGGATACTCGTTTAACGCCCATCTGGTGGCGGGTTTAACGCCGATTGAGGCAGATCCTCTAGATTTAAGAAGGAGATATACATAT
G
```

Data Set S4 - MG1655 library reference file.fasta

```
>araF
GAGGCAATTCCGACGTCTAAGAAACCATTATTATCATGACATTAACCTATAAAAATAGGCGTATCACGAGGCCCTTTCGTCTTCACCTC
GATGGATCCGAAGCATTCCAGCGGTTGCCATACCTAATATCCTTATATCCAGAAGTGGAGAGGTGCAAGATAAATCAAAACACCATAAT
TACTGTTTGCAGAATAATTCCTTTCTGCCGGTCCGAAATGTGATCGTGGAGTCAATTCTGACGATGATCTGAAATAAGAATAACGTCTG
AACGGGTATAAGAAAATGCTTAAATCATCAAGTTTATATATTTTTACATTTCATTGATGAAATCAATGTAACTGCAATGAATTTTAATG
ATAGTGCTTTTTGTATCTTGTTGATATTAATCAATGAATTTGAATTCATGATGAAGTGATGATTTTATAAAACGTTTTTTCATTTTTGC
GAGATGGCTCTCATTATACGTGTTCTGTTAATTAATGAGCACAGTGATAATTAATGACGAAATGAATGACGTGCATTTTCCACATCTTT
GAGTTGCGGTTATTACACCATTTCAAAAACTCAACGCCAGGTAATGCGGCCTATTGACTGGTTAAAAAGAAGACATCCCGCATGGGTAC
CAAAGACAACAAGGATTTCCAGGCTAATCTTATGGATTAATCTGCTGTGCATTCGACAATTTGTCTGACAAATTGGCTTTCCCTTATGT
CTTTTCCCGCTAAATTTATGCACGTTCTCACTGTAATTCTGCGATGTGATATTGCTCTCCTATGGAGAATTAATTTCTCGCTAAAACTA
TGTCAACACAGTCACTTATCTTTTAGTTAAAAGGTAATGCTTTGTTTTCCGATTAATTTAACGAATGTCATTCGTTTTTGCCCTACACA
AAACGACACTAAAGCTGGAGAGAACCATGCACAAATTTACTAAAGCCCTGGCAGCCATTGGTCTGGCAGCCGTTATGTCACAATCCGCT
ATGGCGGAGAACCTGAAGCTCGCTCGAGAGATCCTCTAGATTTAAGAAGGAGATATACATATG
>araD
GAGGCAATTCCGACGTCTAAGAAACCATTATTATCATGACATTAACCTATAAAAATAGGCGTATCACGAGGCCCTTTCGTCTTCACCTC
GATGGATCCCGTTTAAAGACGCGCTGCGCTGGAACGAAGTGTATTACGGGTTTCGTCGCTAAGTAGCCGCATCCGGTATGTAACGCCTG
ATGCGACGCTGACGCGTCTTATCTGGCCTACACGCTGCGATTTTGTAGGCCGGATAAGCAAAGCGCATCCGGCATTCAACGCCTGATGC
GACGCTGGCGCGTCTTATCAGGCCTACGCGCTGCGATTTTGTAGGCCGGATAAGCAAAGCGCATCCGGCATTCAACGCCTGATGCGACG
CTGGCGCGTCTTATCAGGCCTACACGCTGCGATTTTGTAGGCCGGATAAGCAAAGCGCATCCGGCACGAAGGAGTCAACATGTTAGAAG
ATCTCAAACGCCAGGTATTAGAAGCCAACCTGGCGCTGCCAAAACACAACCTGGTCACGCTCACATGGCTCGAGAGATCCTCTAGATTT
AAGAAGGAGATATACATATG
>araE
GAGGCAATTCCGACGTCTAAGAAACCATTATTATCATGACATTAACCTATAAAAATAGGCGTATCACGAGGCCCTTTCGTCTTCACCTC
GATGGATCCCTGATGGGGCCGATAGTGTTCCTTGCCTCCAGCGCTTCAGATTATGTGAATGGTTATACCATTGCCGTGGATGGCGGTTG
GCTGGCGCGTTAATTCATTCTTCTTACTTTTATGACCCTGCCGCATGGCAGGGTTTTTTATACCTGTAGATCATCATAATCCATATCAT
GGTTATGAAATAATCCATATTAATTATCAATTAATGAACTTTATGAATTTTATCTGCTGTAAAATTAGGTGGTTAATAATAATCTCAAT
AATTCAACTTAATTTGAAAATTGGAATATCCATCACATAACGACATGTCGCAGCAATTTAATCCATATTTATGCTGTTTCCGACCTGAC
ACCTGCGTGAGTTGTTCACGTATTTTTTCACTATGTCTTACTCTCTGCTGGCAGGAAAAAATGGTTACTATCAATACGGAATCTGCTTT
AACGCCACGTTCTTTGCGGGATACGCGGCGTATGAATATGTTTGTTTCGGTAGCTGCTGCGCTCGAGAGATCCTCTAGATTTAAGAAGG
AGATATACATATG
>araJ
GAGGCAATTCCGACGTCTAAGAAACCATTATTATCATGACATTAACCTATAAAAATAGGCGTATCACGAGGCCCTTTCGTCTTCACCTC
GATGGATCCGCGTATTCCGGTGCAGATCAAAGTGAAAAAGATCAACGGCCTGGGCTACAGCAAACTGGAAAGTACGTTTGCAGTGAAAT
AACTATTCAGCAGGATAATGAATACAGAGGGGCGAATTATCTCTTGGCCTTGCTGGTCGTTATCCTGCAAGCTATCACTTTATTGGCTA
CGGTGATTGGTAGCCGTTCTGGTGGTTGTGATGGTGGTATGAAAAAAGTCATTTTATCTTTGGCTCTGGGCACGTTTGGTTTGGGGATG
GCCGAATTTGGCATTATGGGCGTGCTCACGGAGCTGGCGCATAACGTAGGCTCGAGAGATCCTCTAGATTTAAGAAGGAGATATACATA
TG
>ygeG
GAGGCAATTCCGACGTCTAAGAAACCATTATTATCATGACATTAACCTATAAAAATAGGCGTATCACGAGGCCCTTTCGTCTTCACCTC
GAGCCGCGTCACCAGACATAATAAAGAAAAAAACAAACGACATTCTACCCTTCTCAAAAATCCCCTTTTTAATTTTCATTGAAAAAATG
GCAATTGATACTATGAATCCAGGTCATACACATTAAGGCATTTATGTAATGTCGTAATTAAGATAACTAATAAGGTGAATATTAGTAAA
ACAACTAATGATGTTATTAGTTTGTAGTGAACAGTACTTTTACCAATAATGAAAAATATACCATAAGCAATGCAAGCTTAAAACTCATA
ATTTAAAAAAGATAAATATAAAAATATCAATGAGTCAATCAATAATAATTTGGCATCACGAGACACATCACAGAGGAATATTATGAGCA
CAGAAACAATTGAAATATTCAATAATAGTGATGAATGGGCAAATCAACTAAAACACGCATTATCGAAAGGAGAAAATCTGGCAGGATCC
TCTAGATTTAAGAAGGAGATATACATATG
>ygeF
GAGGCAATTCCGACGTCTAAGAAACCATTATTATCATGACATTAACCTATAAAAATAGGCGTATCACGAGGCCCTTTCGTCTTCACCTC
GAGTGGTTTCTTTTTCGAGTCCGAATTTACAATATCATGCATATAGACCATTTCATGAATCTGCGAAAATTCAATGTCCATTGTATACC
TCACATTTTTACCGTGACTCGATGTTACTGTTCAATAATCACCTTCCATCAATACTAAAATTAATACCCCTAATGTGCCGATAACAAAT
ATAGTCATTCTACGTAACGTCTCCATAAGGTGATATTTGACATTATCAGAAGCTGCGAATTCGGATTTTGCTCTAATCAAATGAGAGAA
ATGTAAAATGAAGCCACGAAATATTAATAATAGCCTACCACTGCAACCAGGATCCTCTAGATTTAAGAAGGAGATATACATATG
>ygeA
GAGGCAATTCCGACGTCTAAGAAACCATTATTATCATGACATTAACCTATAAAAATAGGCGTATCACGAGGCCCTTTCGTCTTCACCTC
GATGGATCCATGGCAGGCGAGAAGTTGAGAAATATCGGCGTCTGATTTCACGGGCCGGATGTGCTGTACATCCGGCCCTTTTTTCGTTA
ATAGAGATTGGGCACTTGGCCGTTGAGGCGTTTGTCTCGTTCCTTATTCAGCCTTGTTGCGGTAACACACATCAGGAGAGAGGAATGAA
AACAATTGGTTTGCTGGGAGGAATGAGCTGGGAATCCACCATTCCTTACTATCGTTTGATAAATGAAGGCATTAAACAGCGGCTTGGTG
GCTCGAGAGATCCTCTAGATTTAAGAAGGAGATATACATATG
>ygeI
GAGGCAATTCCGACGTCTAAGAAACCATTATTATCATGACATTAACCTATAAAAATAGGCGTATCACGAGGCCCTTTCGTCTTCACCTC
GAGTGAGAGTGAAAGCCATCGAGTTATAATTGACAACATTTTGAAATTAAAGCAGCTGACGAGAATTTGTATGCGATAAATTTTTATTG
ACGCAATTTTATTATTTAGATGTATGCAACTTGTTATTTAAACAACAGAGATAAAAATATCTAAGATATTCACCTTATTGCAAGATATT
TAAAATGCTCTAGAGTTAAAAGTATGATGTAAAGACCATTGGCCTGGACAATTAAAAAAGGGAGCTTTATGACTAATCCAATCGGTATC
```

AATAATTTATCTCAAAGTTCAAATATAGCGAATGCAACGGGTGATGAAGTCGGGATCCCGGGATCCTCTAGATTTAAGAAGGAGATATA
CATATG
>cirA
GAGGCAATTCCGACGTCTAAGAAACCATTATTATCATGACATTAACCTATAAAAATAGGCGTATCACGAGGCCCTTTCGTCTTCACCTC
GATGGATCCGAAGTTCCCGCAGAACGATAAGAAATAAGTTTCCTCCCTTCCTTGCTAAGCCCTCTCAACCGAGAGGGCTTTTTCAATTC
CATTTCCCTGACAAATCATGCGGATATAAAATTTAACATTTGGATTGATAATTGTTATCGTTTGCATTATCGTTACGCCGCAATCAAAA
AAGGCTGACAAATCAGAGGCTGTTCCGGCTTTCTGGGATGATCACCTGCATAAAAAATAAGTCCACCGCGATGCTGCCGTACGCAAGGG
GACGTGAAGAAGATGTGAGCGATAACCCATTTTATTTTCGTAGTTACCTCATGGAGATATGGAATGTTTAGGTTGAACCCTTTCGTACG
GGTCGGGCTGTGTTTGTCCGCTATTTCTTGTGCATGGCCTGTGTTAGCGGTCGATGATGATGGCGAAACGATGGCTCGAGAGATCCTCT
AGATTTAAGAAGGAGATATACATATG
>ygeR
GAGGCAATTCCGACGTCTAAGAAACCATTATTATCATGACATTAACCTATAAAAATAGGCGTATCACGAGGCCCTTTCGTCTTCACCTC
GATGGATCCCAGTATATCGTGCCCGCCCGGTGACCTTAGCAATGGCATCGACGCGCATGCATGATTCACCCGTAGCGGTTGCTTCCCGC
GCTTCCATAGAAATCCCCTCGATTTGAACATTTGGTTCTTATGGCAAAGATTTAGTCTAAGCAGGCAAGATTTACGCCAGAAACGACGA
TAAGCGTTAGGGGAGGTAGGAAGTGTGATGACGTTAAAAAAACACAGATATGATACGAAACAGGACTCTGAATAGGGGCAGTGATACGT
ATCAATATGAGTGGCGATAAAATCAATGTGATAAATTGGGATTATTACGCATCAGGATGGGAAATAGGGTAAAAAGTCAGGAAGTTCGA
TACCTCTACAGCGAATATTTTGCTGATGATACTTCCCTTTGCCAAATGAAATTAACCTCTTTCTCTATTACCACGTTTTTCCAGAAGCA
AGAGATTGGGTATCAAGAGGCTGGCTGCTATGATAAGGCGTCTTGTTTTTTAAGCGAGGAAAGATTTTGAGTGCGGGACGCCTGAATAA
AAAATCTCTGGGTATCGTGATGTTGTTATCGGTTGGACTGCTTTTGGCTCGAGAGATCCTCTAGATTTAAGAAGGAGATATACATATG
>ygeQ
GAGGCAATTCCGACGTCTAAGAAACCATTATTATCATGACATTAACCTATAAAAATAGGCGTATCACGAGGCCCTTTCGTCTTCACCTC
GATGGATCCAGGGTTCGATTCCCTTCGCCCGCTCCAATTTATCTTCGCCGTAATACTTCCCCATTTTGTTGTAAATAAAAAGCGTTTAC
GCAATGCAATCCTTTTGTAACAACCCAGGTAAAACACCTTATTTTCAAGTTGTTAAATAAGTGGTTATCTACGCTTTTAACAAACGATT
GATGGAATATTTTATGAAAGGGAAATCTGCATTAACTCTTCTGCTCGCTGGTATATTTTCGTGCGGCACATCTCGAGAGATCCTCTAGA
TTTAAGAAGGAGATATACATATG
>fruL
GAGGCAATTCCGACGTCTAAGAAACCATTATTATCATGACATTAACCTATAAAAATAGGCGTATCACGAGGCCCTTTCGTCTTCACCTC
GAGAGCATCGATTCGCGATGTGGCGAAAATTGTGGAGGTTGCTCGCTCTGGTGTGGTCGGACTTTCGCGCGGCGATAAAATAATGCGTT
GAGAATGATCTCAATGCGCAATTTACAGCCCAACATGTCACGTTGGGCTTTTTTTGCGAAATCAGTGGGAACCTGGAATAAAAGCAGTT
GCCGCAGTTGGATCCTCTAGATTTAAGAAGGAGATATACATATG
>fruB
GAGGCAATTCCGACGTCTAAGAAACCATTATTATCATGACATTAACCTATAAAAATAGGCGTATCACGAGGCCCTTTCGTCTTCACCTC
GATGGATCCGTCAAACGATTTCGCGCTGGAGACTGCGGGGGAGTTATGCATGCTGGTCGATCACTGGAAGAGGAAGACGCGGAGTTTAC
GCTGCTTGTCAGAAAATAAACAGACAGCGAAATCAAAAGCGCCACGAATAGTCAAAATGCCATTTCACTGTTGCTGATTGGCTGCTGAC
GCTGCTGAAATGAAATTGCTGATAACGGATTTTCCCATCAGCAATTAGGAAAAATGGCAAAAAATTGTGCAGCACATCAAACTTTTGCT
CATAACTTTACGGCTTTCCTTGCGTGCTGAAACAGAAGTATTATGCTTTCTTGAAACGTTTCAGCGCGATCTTGTCTTTAACCCTAAGC
CAGGTTGGCGCTTTTTTTCTCATAGAGGCTGAATCGTTTCAATTCAGCAAGAGAGGAGAACTATGTTCCAGTTATCCGTACAGGACATC
CATCCGGGCGAAAAGGCCGGAGACAAAGAAGAGGCGATTCGCCTCGAGAGATCCTCTAGATTTAAGAAGGAGATATACATATG
>ygeY
GAGGCAATTCCGACGTCTAAGAAACCATTATTATCATGACATTAACCTATAAAAATAGGCGTATCACGAGGCCCTTTCGTCTTCACCTC
GAGAGCACTGAAGGCGACACCGACGTGAAGCACTACCGCGAAGTTGTCTGGGAAGGCAAACACGCTGTAGCACCTTAATCACCCTATTG
GAACTGCTCCCTGCAAAACGGGGAGTAAAAAATCTGGAGAAAAATAATGGCTAAGAATATTCCATTCAAACTGATTCTTGAAAAAGCAA
AAGATTACCAGGCGGATATGACTCGGGATCCTCTAGATTTAAGAAGGAGATATACATATG
>ygeX
GAGGCAATTCCGACGTCTAAGAAACCATTATTATCATGACATTAACCTATAAAAATAGGCGTATCACGAGGCCCTTTCGTCTTCACCTC
GAGAGAAGCCAGCTGGAAGCCTTATATCATCGCCGCGATGATCCTGTCCCGTAAATACGCCAAACCAGGTGCACTGCTCGAGCAACTGC
TGAAAGAAGCGCAAGAACGCGTGAAATAACCCCTCGGGCCGATCAGCTGGTCGGCCCTCTATTTCCAGAGGCCAAAAGGATAGGATATG
TCCGTTTTCTCATTGAAGATTGATATCGCCGATAACAAATTTTTCAACGGCGAAACATCACCGCTCTGGATCCTCTAGATTTAAGAAGG
AGATATACATATG
>rffC
GAGGCAATTCCGACGTCTAAGAAACCATTATTATCATGACATTAACCTATAAAAATAGGCGTATCACGAGGCCCTTTCGTCTTCACCTC
GAGGACGAGGGTGTGAAGCGTGCTGCCAGTTCATTAGCGAAAACTGGCTACGGCCAATATCTGCTGGAGTTACTTCGTGCCCGTCCGCG
CCAGTATTGAGCCACTAACCTGGGAAAACGCCTTCTTTGGTGTTAACAGCGCCATCGTGCGCATTACGTCTGAAGCACCGCTCCTGACG
CCAGACGCGTTAGCGCCGTGGTCACGGGTGCAGGCCAAAATTGCGGCATCAAATACGGGTGAACTGGATGCCCTGCAACAGCTGGGATT
CTCCCTGGTGGATCCTCTAGATTTAAGAAGGAGATATACATATG
>ypfG
GAGGCAATTCCGACGTCTAAGAAACCATTATTATCATGACATTAACCTATAAAAATAGGCGTATCACGAGGCCCTTTCGTCTTCACCTC
GATGGATCCCAAAACCGGCGAGATACGTGACGGTAAGACGGTGTTATTGCTTAACTATTTGCAAACGTCACATTTAATGGACTGAAAAA
TAACAATAATATTTCGTTGTTTATTATTGGCTCAATCCGACAAATCCGATTGAGCCGCGCTACCTGCGCAACGAAGATACGACTTGTGC
TGTTTGTTTGAACTTCTGGGGTCGTACCGTCCATGCGCTATCGCATTTTCCTTCTCTTTTTTTTCGCTTTGTTGCCGACGTCTTTGGTG
TGGGCGGCACCAGCGCAACGGGCGTTTTCCGACTGGCAGGTCCTCGAGAGATCCTCTAGATTTAAGAAGGAGATATACATATG
>ypfH

Data Set S4 - MG1655 library reference file.fasta
GAGGCAATTCCGACGTCTAAGAAACCATTATTATCATGACATTAACCTATAAAAATAGGCGTATCACGAGGCCCTTTCGTCTTCACCTC
GATGGATCCAGGCGCTGTTCGCACTTAATGATGTTCGCACTGAGCGTCTGCGCGATCGCATAACGCAATGGCAATTATTTCACTGACTC
CTTCAATTTTATCTCATGGTCATTGTGAGTCAGCGGCGTAAAATTAATGACATCAGTTAAGGAGTTCGCCATGAAACATGACCATTTTG
TTGTTCAAAGCCCGGATAAACCTGCGCAACCTCGAGAGATCCTCTAGATTTAAGAAGGAGATATACATATG
>ypfJ
GAGGCAATTCCGACGTCTAAGAAACCATTATTATCATGACATTAACCTATAAAAATAGGCGTATCACGAGGCCCTTTCGTCTTCACCTC
GATGGATCCTCGAAGCCTATGAAGCCGTCGCCCGCCGCCTGGGTGTACAGCTGGACTGATTTTTCTGTTCATCATCTTGCCGTGCAGCT
GGCACGGCAAGACAACCGCTCTCGGAATATATGCATTGAACGCTCATTCCTCCGCCATTTCATCCCGCTTTCCTGTGGTAATCTTCCTC
TGAACCACCTACGATCATTTCTATAATGAATATATTGTTGAGGTATCTATGCGTTGGCAAGGGCGACGTGAAAGTGACAATGTTGAAGA
CAGGCGCAACAGCTCTGGTGGTCCATCTATGGGCGCTCGAGAGATCCTCTAGATTTAAGAAGGAGATATACATATG
>deaD
GAGGCAATTCCGACGTCTAAGAAACCATTATTATCATGACATTAACCTATAAAAATAGGCGTATCACGAGGCCCTTTCGTCTTCACCTC
GATGGATCCATGACCTGGCAGAATCGGACCAGCAATAGCTGACGTACACATCAGCCCGTAATCTTTTTGATTGCCATCACCTTAACGG
GTGAGGGCGTTGTTGTTCGTTAATACACCTACTTTGAGCCGGTTCACACTTTTCAATGAAAATTGCTGATCAATTTCATGATGAGTTAT
GTAGACTGGCCGCCATTAATTTTGAGGCACACGTACTACATGGCTGAATTCGAAACCACTTTTGCAGATCTGGGCCTGAAGGCTCCTCG
AGAGATCCTCTAGATTTAAGAAGGAGATATACATATG
>icd
GAGGCAATTCCGACGTCTAAGAAACCATTATTATCATGACATTAACCTATAAAAATAGGCGTATCACGAGGCCCTTTCGTCTTCACCTC
GAGTGGGCTGGTTTTCAGGTTTACGCCTGGTAGAACGTTGCGAGCTGAATCGCTTAACCTGGTGATTTCTAAAAGAAGTTTTTTGCATG
GTATTTTCAGAGATTATGAATTGCCGCATTATAGCCTAATAACGCGCATCTTTCATGACGGCAAACAATAGGGTAGTATTGACAAGCCA
ATTACAAATCATTAACAAAAAATTGCTCTAAAGCATCCGTATCGCAGGACGCAAACGCATATGCAACGTGGTGGCAGACGAGCAAACCA
GTAGCGCTCGAAGGAGAGGTGAATGGAAAGTAAAGTAGTTGTTCCGGCACAAGGCAAGAAGATCACCCTGCAAAACGGCAAACTGGATC
CTCTAGATTTAAGAAGGAGATATACATATG
>borD
GAGGCAATTCCGACGTCTAAGAAACCATTATTATCATGACATTAACCTATAAAAATAGGCGTATCACGAGGCCCTTTCGTCTTCACCTC
GATGGATCCATTCATCGCACCTGGCATACATTAATAAATATTAACAATATGAAATTTCAACTCATTGTTTAGGGTTTGTTTAATTTTCT
ACACATACGATTCTGCGAACTTCAAAAAGCATCGGGAATAACACCATGAAAAAAATGCTACTCGCTACTGCGCTGGCCCTGCTTATTAC
AGGATGTGCTCAACAGACATTTACTGTTCAAAACAAACAGACAGCAGTAGCACCAAAGGAAACCATCACCCATCATTTCTTCGTTTCTG
GAATTGGGCAGAAGAAAACTGTCGATGCAGCTAAAATTTGTGGCGGCGCAGAAAATGTTGTTAAAACAGAAACCCAGCAAACATTCGTA
AATGGATTGCTCGGTTTTATTACTTTAGGCATTTATACTCCGCTGGAAGCGCGTGTGTATTGCCTCGAGAGATCCTCTAGATTTAAGAA
GGAGATATACATATG
>hdeD
GAGGCAATTCCGACGTCTAAGAAACCATTATTATCATGACATTAACCTATAAAAATAGGCGTATCACGAGGCCCTTTCGTCTTCACCTC
GAGAGCTGCTTTTTGCGCATCCGCTGCATTGCTCACAACTGGCAGAAGAAGCAGACCACCAAGAATAACGCCTAATACTTTTTTCATCG
TAATATCCTCAACTATAAAGTGAAAGAGCCGTCACGAATCAATTTCGACACTGAGGTTATAACCTGGTTTTCTGTATATGTCATGTTGA
TGGAAAATATCAAAATCAGATATTTTTATTTCAATACAATGAGTTACAGATGCATCAGATACTGCAATTAGGAAATTTTTATTAAATCG
ACTGCATTCTTAGACGCGTTTTTGGCATAGATTGATAGCAGGGGATTTTCTTCTTAATTTTATAGGGTGGTTCTATGTTATATATAGAT
AAGGCAACAATTTTGAAGTTTGATCTGGAGATGCTTAAAAAACATCGCAGAGCAATCCAGTTTATTGCCGTGCTGCTGTTTAGGATCCT
CTAGATTTAAGAAGGAGATATACATATG
>hdeB
GAGGCAATTCCGACGTCTAAGAAACCATTATTATCATGACATTAACCTATAAAAATAGGCGTATCACGAGGCCCTTTCGTCTTCACCTC
GATAGATCTCCCAGCTATCGTTCAGGCTTGTACTCAGGATAAACAAGCCAACTTTAAAGATAAAGTTAAAGGCGAATGGGACAAAATTA
AGAAAGATATGTAATTCCGGGAATGCGTTACATCGTACTTCCTTGCATATTGAACAGGCCGGAATATCTTCTTTAAAAGCAGCTATTCC
TCCTGTTCATATATAATCTCTATATTGAATGGGTTACAAAATGAATATTTCATCTCTCCGTAAAGCGTTTATTTTTATGGGCGCTGTAG
CGGCTTTGTCACTGGTGAACGCACAATCTGCGCTCGAGAGATCCTCTAGATTTAAGAAGGAGATATACATATG
>hdeA
GAGGCAATTCCGACGTCTAAGAAACCATTATTATCATGACATTAACCTATAAAAATAGGCGTATCACGAGGCCCTTTCGTCTTCACCTC
GATAGATCTTAAACAGCAGCACGGCAATAAACTGGATTGCTCTGCGATGTTTTTTAAGCATCTCCAGATCAAACTTCAAAATTGTTGCC
TTATCTATATATAACATAGAACCACCCTATAAAATTAAGAAGAAAATCCCCTGCTATCAATCTATGCCAAAAACGCGTCTAAGAATGCA
GTCGATTTAATAAAAATTTCCTAATTGCAGTATCTGATGCATCTGTAACTCATTGTATTGAAATAAAAATATCTGATTTTGATATTTTC
CATCAACATGACATATACAGAAAACCAGGTTATAACCTCAGTGTCGAAATTGATTCGTGACGGCTCTTTCACTTTATAGTTGAGGATAT
TACGATGAAAAAAGTATTAGGCGTTATTCTTGGTGGTCTGCTTCTTCTGCCAGTTGTGAGCAATGCAGCGGATGCGCAAAAAGCAGCTC
TCGAGAGATCCTCTAGATTTAAGAAGGAGATATACATATG
>ribA
GAGGCAATTCCGACGTCTAAGAAACCATTATTATCATGACATTAACCTATAAAAATAGGCGTATCACGAGGCCCTTTCGTCTTCACCTC
GATGGATCCTCCCACTGCGGTACGTCTGGCAATCGAACGCATGGCCTCTCCTTTTTGATAAGTCCCACAATCATAACTGAAAACGCCAG
TTCCAGGAAAAATTGACAGATTTGTGCCATTCCGTGAACGATCGACGCGTCGTGATTAGGTGAACCCCTTCTCGTTATGGCAAAATAAG
CCAATACAGAACCAGCATTATCTGGAGAATTTCATGCAGCTTAAACGTGTGGCAGAAGCCAAACTGCCAACCCCATGGGGCGATTTCCT
GATGGTGGGATTTGAAGAACTGGCAACCGGACACCTCGAGAGATCCTCTAGATTTAAGAAGGAGATATACATATG
>ribC
GAGGCAATTCCGACGTCTAAGAAACCATTATTATCATGACATTAACCTATAAAAATAGGCGTATCACGAGGCCCTTTCGTCTTCACCTC
GATGGATCCTCCAGCAGGCTGTCATTGATAAAGTTGTTGAAACGCGCGATGGTGATGGCGACGCGAGCGTCCGGGGTAGCAACGTTAGC
TTCAATAATGTTCATTTTCTTCCTTCGGGTTCGAGTATGGCCCCGCAGGGGGGCGGATTTTAGCATAATATTTCGTGCGCTGCTTCCCT

TTCGAGCCGGGAGATCATGCACCCACTAAATGCAGGCAAACATCCGGGCCTACATGACGTATCTCTTTGAATTTAAATTGGGGGGCGTC
GGCTAATTTCTCAAGCCCTGGCAGCTCGAGAGATCCTCTAGATTTAAGAAGGAGATATACATATG
>ribB
GAGGCAATTCCGACGTCTAAGAAACCATTATTATCATGACATTAACCTATAAAAATAGGCGTATCACGAGGCCCTTTCGTCTTCACCTC
GATGGATCCGCGAGCGATTTGCTCAATTTTTTTCGGGTCAATCATTGTAGTTTCCCTGTACTGATAGGTGTTGAGTGCCATTGTAGTGC
GATAAGGGTAAGTCATAAACCAGAATTATGTGAAGCTATGCGTTGCTGCCGCTAATCATTAGCGTTATAGTGAATCCGCTTATTCTCAG
GGCGGGGCGAAATTCCCCACCGGCGGTAAATCAACTCAGTTGAAAGCCCGCGAGCGCTTTGGGTGCGAACTCAAAGGACAGCAGATCCG
GTGTAATTCCGGGGCCGACGGTTAGAGTCCGGATGGGAGAGAGTAACGATTCTGTCGGGCATGGACCCGCTCACGTTATTTTGGCTATA
TGCCGCCACTCCTAAGACTGCCCTGATTCTGGTAACCATAATTTTAGTGAGGTTTTTTTACCATGAATCAGACGCTACTTTCCTCTTTT
GGTACGCCTTTCGAACGTGTTGAAAATGCACTGGCTGCGCTGCGTGAAGGACGCGGTGTAATGGTGCTCTCGAGAGATCCTCTAGATTT
AAGAAGGAGATATACATATG
>ribE
GAGGCAATTCCGACGTCTAAGAAACCATTATTATCATGACATTAACCTATAAAAATAGGCGTATCACGAGGCCCTTTCGTCTTCACCTC
GATGGATCCATTTGCGCGAGAATCACCGGGATTGCTAATGCTAATAACAGACGCGCTTCACTGATATACTTCTGCACGTGAACACCTTT
TATTTGTAGTTATATGAGAGACTAAAAAGCCGCCTGAATGGGCAGCAAAAAGAAGAAGCAGGGGAAATTCAGTCTATTGTAGCGAGGAT
TTATTAATTCTCCAGTGAAAAAATAGCCATTCTTACCCCTTCACTGGTAAGGGCGTTTTCCACCTGCTATTGTGCTGAACAGAATGTCT
TAACTGATTTCAGGAGTTGTAAGTATGTTTACGGGGATTGTACAGGGCACCGCAAAACTGGTGTCGATTGACGAGAAACCAAATTTTCG
TACGCATGTGGTGGAGTTACCCGACCCTCGAGAGATCCTCTAGATTTAAGAAGGAGATATACATATG
>htpX
GAGGCAATTCCGACGTCTAAGAAACCATTATTATCATGACATTAACCTATAAAAATAGGCGTATCACGAGGCCCTTTCGTCTTCACCTC
GATGGATCCTTGAAAAAGCCAGACCCGCGGAACAACCCGCTCCCGTCAAGTAATATCAATCAGGCACAAGAAATTGTGCCTGATTTTTT
AACAGCGACAAGATGCCGTAAATCAGATGCTACAAAATGTAAAGTTGTGTCTTTCTGGTGACTTACGCACTATCCAGACTTGAAAATAG
TCGCGTAACCCATACGATGTGGGTATCGCATATTGCGTTTTGTTAAACTGAGGTAAAAAGAAAATTATGATGCGAATCGCGCTCTTCCT
GCTAACGAACCTGGCCGTAATGGTCGTTTTCGGGCTGGTACTGAGCCTCGAGAGATCCTCTAGATTTAAGAAGGAGATATACATATG
>yqjI
GAGGCAATTCCGACGTCTAAGAAACCATTATTATCATGACATTAACCTATAAAAATAGGCGTATCACGAGGCCCTTTCGTCTTCACCTC
GAGCGCTCAACGCGTAACACAGTCAGTTCACGGAAGCGCAGATCATTGCGAACGCGCTGCGGGTAGCGGGGGGTGTTATTCATTGTTAT
CGCCTTCGTGATGGTAATCAGATATATCTAAATAAAACTCGCAAATGATAATGATTGTTAATCATGATAAATGCAAGCGATTTGTAGAA
CTGATATGTCTATAGTCTGATAAGACGAACCGCCTCTTCTCAGGCATCATTACTCAACGCCGGATGCGGCGTGAACGCCTTATCCGGCC
TACGTGTGAGATGAGTCCACGTTTAAAAATCATTTTTACACTTGCAAGAACGCTCATATCAGATATAAATTAGATATATCTAATTAAGC
AAAAGGAGGCTGATATGAGCCATCATCACGAAGGGTGTTGTAAACATGAAGGCCAGCCACGCCATGAGGGCTGCTGCAAAGGTGAGGGA
TCCTCTAGATTTAAGAAGGAGATATACATATG
>corA
GAGGCAATTCCGACGTCTAAGAAACCATTATTATCATGACATTAACCTATAAAAATAGGCGTATCACGAGGCCCTTTCGTCTTCACCTC
GAGGGCAACAGCAAACAAGGGAAGTAAGGTGAGGGCGAGAAAAATCCGTTTGAGATTCAAGGTGATCATTCCTTTACCAATGAGTAGCT
GATGCGCCATTATAGGTCCTGGATGTGGGATTTTTTTATCCTGTTAGCGACCTTGACGAGTACCAAAAAGCGCGAAGTTCAACTATTGT
TCTGTGGTGTTCTGTTGCGTGTTGACGGCAAAATTTTGCTGGCGTAACATGCGCGCACGATCACTCTAAGAGGACATTCGCCTTGGACA
CACCCAGTAGATACTGGCTCACTATCCTGTCATCCAGGATCAACTCCTAAGGCTATCCCTTTTTGCTGATAGCCTTAGCGGTTGTCAGC
GACCTCAATTTTTCCCGTCGCGCTGAGTCAGGCTGTTTAATGGTCTGAAACCCAATTTGTTTCTGTGTGCCCACCGAACTGTCCGATAT
TTTAAGCATTGGGAGTCCCGGTCATGCTGAGCGCATTTCAACTGGAAAATAACCGACTGACCCGGCTGGAAGTCGAAGAGTCACAACCC
CTTGTAAATGCAGGATCCTCTAGATTTAAGAAGGAGATATACATATG
>yqjC
GAGGCAATTCCGACGTCTAAGAAACCATTATTATCATGACATTAACCTATAAAAATAGGCGTATCACGAGGCCCTTTCGTCTTCACCTC
GAGTGCGCAGCAGGACAATAACAGTCAGGCTATGCAGTGGCTGACCCGGTTACGGGATAACTCTCATCGCTTCGGATAACTTCCTGTTA
TCCGAAACATATCACTCACTTTGGTGATTTCACCGTAACTGTCTATGATTAATGAGGCGGTGGTTAACACTACCGCTTCGTTGTCTGGA
CAATCGTTCCTTTGTAATAGGTCCGAATCACAATGGAAGGTTCAAGAATGAAATACCGCATCGCTTTAGCTGTTTCTCTCTTTGCTCTT
AGTGCCGGTAGTTATGCCACTACCCTGTGTCAGGGGGATCCTCTAGATTTAAGAAGGAGATATACATATG
>yqjF
GAGGCAATTCCGACGTCTAAGAAACCATTATTATCATGACATTAACCTATAAAAATAGGCGTATCACGAGGCCCTTTCGTCTTCACCTC
GAGCGGTTTTGGCGTATGGAGCGCCTGGCGTCTGGTTAAAACGACCCTCAAGCAGCAACAGCTTCGCGGTTAACTTCCCTCTGGCCGGA
GCCATTCCGGCCTTATCCCTCAAATTTTTTGAAGATTTTTGACAGTTTTCCTTGCTAACAATCATCATTCACCACGTTTATGATTCTCT
CCATCGACAGCAACGACGCTAATACCGCGCCATTGCACAAAAAAACAATCAGCAGCCTGAGTGGCGGATCCTCTAGATTTAAGAAGGAG
ATATACATATG
>yqjG
GAGGCAATTCCGACGTCTAAGAAACCATTATTATCATGACATTAACCTATAAAAATAGGCGTATCACGAGGCCCTTTCGTCTTCACCTC
GAGCCTGACAATTTCTGGCGGATTCCTGCTGCTGGCAATTACCGGTCCGGGCGCGTATAGCATCGACCGCCTGCTGAATAAAAAGTGGT
AAGCACGCGCTATACTTAACGATAAAAAACGAGGAGGAAGCTCCTCGTTTTTGCTATTGGAGGAGAGAAAAATGGGTCAACTGATTGAC
GGCGTCTGGCATGACACCTGGTACGATACCAAATCTACCGGCGGTAAATTTCAACGTTCAGCTTCCGCATTTGGATCCTCTAGATTTAA
GAAGGAGATATACATATG
>pheS
GAGGCAATTCCGACGTCTAAGAAACCATTATTATCATGACATTAACCTATAAAAATAGGCGTATCACGAGGCCCTTTCGTCTTCACCTC
GATGGATCCCGCCGTAGCCTTTTCAATAAAGGTCTTTTGAAGAGTACCAAAAGGTAACGCAAGCAATGAATGCTGCTATTTTCCGCTTC
TTTTTTTACTTTAGCACCTGAATCCAGGAGGCTAGCGCGTGAGAAGAGAAACGGAAAACAGCGCCTGAAAGCCTCCCAGTGGAGGCTTT

TTTTGTATGCGCGTTTTGAAAAATAAAAAGCCTGTCTCATCAGGTGTAATTGTCGCAGCCAGACTGGACACGGCCAGCGCGGAGAAACC
GGAGCGTATTTAAGTACGTGAGAATTTCGAGCACAGCCCGGGACCAAAATGGCAAGTAAAATAGCCTGATGGGATAGGCTCTAAGTCCA
ACGAACCAGTGTCACCACTGACACAATGAGGAAAACCATGTCACATCTCGCAGAACTGGTTGCCAGTGCGAAGGCGGCCATTAGCCAGG
CGTCAGATGTTGCCGCGTTACTCGAGAGATCCTCTAGATTTAAGAAGGAGATATACATATG
>pheP
GAGGCAATTCCGACGTCTAAGAAACCATTATTATCATGACATTAACCTATAAAAATAGGCGTATCACGAGGCCCTTTCGTCTTCACCTC
GAGGATCACCGCACCTTTGCTGTCGCTGTTTATTATCCCGGCGGCGTATAAGCTGATGTGGCTGCACCGACATCGGGTACGGAAATAAA
AGCAGGATACCCCGTTTAACCGTGTGGATTGTGTCTTGCGACGATGGGCACTAAATGTTAAAAGGTGCCCCTCAACAAAAAAGACACAC
AGGGGAAAGGCGTGAAAAACGCGTCAACCGTATCGGAAGATACTGCGTCGAATCAAGAGCCGACGCTTCATCGCGGATTAGGATCCTCT
AGATTTAAGAAGGAGATATACATATG
>ileX
GAGGCAATTCCGACGTCTAAGAAACCATTATTATCATGACATTAACCTATAAAAATAGGCGTATCACGAGGCCCTTTCGTCTTCACCTC
GAGAAGGGAAAACCAGTCCCGGCGGATGAAAGCCCAGGGTTGATACCGCAAAACACGACCCGTAACCCTGGAGCCAAAATATCCTCAAC
CATATTTACTCCTTTCTAACATCACCAGGAAAAGTATAACGGATTGAAAATACATTGTTTATAAAAACAGCAGGCGCGCGGTAATGGCT
GGATTGCGACACGGAGTTACTTTATAATCCGCTACCATGGCCCCTTAGCTCAGTGGTTAGAGCAGGCGACTCATAATCGCTTGGTCGCT
GGTTCAAGTCCAGCAGGGGGATCCCGGGATCCTCTAGATTTAAGAAGGAGATATACATATG
>htpG
GAGGCAATTCCGACGTCTAAGAAACCATTATTATCATGACATTAACCTATAAAAATAGGCGTATCACGAGGCCCTTTCGTCTTCACGTC
GACGGCGAGCTGGAAATGGTCGACGGCACCACGTTGTCACACTCCCTTGCCGGGCGTCATAAGATTCGTTTTTAAGCAAACGAGAGCAG
GATCACCTGCTCTCGCTTGAAATTATTCTCCCTTGTCCCCATCTCTCCCACATCCTGTTTTTAACCTTAAAATGGCATTATTGAGGTAG
ACCTACATGAAAGGACAAGAAACTCGTGGTTTTCAGTCAGAAGTGAAACAGCTTCTGCACCTGATGATCCAGGATCCTCTAGATTTAAG
AAGGAGATATACATATG
>ppa
GAGGCAATTCCGACGTCTAAGAAACCATTATTATCATGACATTAACCTATAAAAATAGGCGTATCACGAGGCCCTTTCGTCTTCACCTC
GATGGATCCGACATGGCTGCCGAGACTGCAGAGACTATAAGTAAGCGTTTCCACATAGGGTTGTCCTCGTCGGGGTAGTTATTATTGGT
TAAAAGATGTTCGCGAAAAAACTATAGACAATTCGTTATGTAACAGATTGCGTTACATCACACTTCAAAAAAGTAAATAAAACGTTAAT
CACAAGTTTGTAATCGCTTTCATCTCACTATGAAAAATGCGGCTACGGTTATGGATTTTCCTGCTCTGTATACCGTCTTAAAACTGGTG
AAAAAGGAAAATGAAGACGAAAACAAGCGAAGACATTCGGCGCGAGTTGGCTATAATACTCGGCACTTGTTTGCCACATATTTTTAAAG
GAAACAGACATGAGCTTACTCAACGTCCCTGCGGGTAAAGATCTGCCGGAAGACATCTACGTTGTTATTGAGATCCCGGCTAACGCTCG
AGAGATCCTCTAGATTTAAGAAGGAGATATACATATG
>dkgB
GAGGCAATTCCGACGTCTAAGAAACCATTATTATCATGACATTAACCTATAAAAATAGGCGTATCACGAGGCCCTTTCGTCTTCACCTC
GAGTCGGTGGAGCGGTAGTTCAGTCGGTTAGAATACCTGCCTGTCACGCAGGGGGTCGCGGGTTCGAGTCCCGTCCGTTCCGCCACTTA
TTAAGAAGCCTCGAGTTAACGCTCGAGGTTTTTTTTCGTCTGTATATCTATTATTGCCAGAATCGCAAAAATCCTCTGCATTTTACGCT
CTTTTTCCTCAACAGTCTGAAGCCCATAATCACCTCAGTTAACGAAAATAGCATTAAAAGAGGCATATTATGGCTATCCCTGCATTTGG
TTTAGGTACTTTCCGTCTGAAAGACGACGTTGTTATTTCATCTGTGATAACGGCGCTTGAGGATCCTCTAGATTTAAGAAGGAGATATA
CATATG
>dkgA
GAGGCAATTCCGACGTCTAAGAAACCATTATTATCATGACATTAACCTATAAAAATAGGCGTATCACGAGGCCCTTTCGTCTTCACCTC
GAGACCCAACTGGGCGAAAATCATGACATTACGTTGGATGTCAGCCGCCGTATATACGAAGCCGCCCGCTAAGCTTTTTACGCCTCAAA
CTTTCGTTTTCGGGCATTTCGTCCAGACTTAAGTTCACAACACCTCACCGGAGCCTGCTCCGGTGAGTTCATATAAAGGAGGAACGTAT
GGCTAATCCAACCGTTATTAAGCTACAGGATGGCAATGTCATGCCCCAGCTGGGACTGGGCGTCTGGCAAGCAAGTAATGAGGAAGTAA
TCACCGCCATTCAAAAAGCGTTAGAAGTGGGTTATCGCTCGATTGATACCGCCGCGGCCTACAAGAACGAAGAAGGTGTCGGCAAAGCC
CTGAAAAATGCCTCAGGATCCTCTAGATTTAAGAAGGAGATATACATATG
>relB
GAGGCAATTCCGACGTCTAAGAAACCATTATTATCATGACATTAACCTATAAAAATAGGCGTATCACGAGGCCCTTTCGTCTTCACCTC
GATGGATCCTTCCTTCGTGACTTGCCCTAAGCATGTTGTAGTGCGATACTTGTAATGACATTTGTAATTACAAGAGGTGTAAGACATGG
GTAGCATTAACCTGCGTATTGACGATGAACTTAAAGCGCGTTCTTACGCCGCGCTTGAAAAAATGGGTGTAACTCCTTCTGAAGCGCTT
CGTCTCATGCTCGAGGTCGAGAGATCCTCTAGATTTAAGAAGGAGATATACATATG
>relA
GAGGCAATTCCGACGTCTAAGAAACCATTATTATCATGACATTAACCTATAAAAATAGGCGTATCACGAGGCCCTTTCGTCTTCACCTC
GATGGATCCCCCACACACGGGACATCTGGAATCGATGGTACTTTTCTCGCGCGTTAAATAGTTGCGATTTGCCGATTTCGGCAGGTCTG
GTCCCTAAAGGAGAGGACGATGGTTGCGGTAAGAAGTGCACATATCAATAAGGCTGGTGAATTTGATCCGGAAAAATGGATCGCAAGTC
TGGGCTCGAGAGATCCTCTAGATTTAAGAAGGAGATATACATATG
>phnQ
GAGGCAATTCCGACGTCTAAGAAACCATTATTATCATGACATTAACCTATAAAAATAGGCGTATCACGAGGCCCTTTCGTCTTCACCTC
GATGGATCCCAGTTTGATGCGTGGCTGATGGAAAACGCACTACCGTCAGGGTTTGAGGTGGGGTTTGATGGGATGGAGATTGGGGTGGC
GTGATGAGAGGGAATGTGCGCGCTGGCCCCCTCACCCTAACCCTCTCCCCAGAGGGGCGAGGGGACCGATTGTGCTCGATATTGAATAT
TGCGCTCGTTTTCTCCCTCTCCCCATTGGGGTGAGGGGCGATGCCTGCTCCATACCCAACCTCATCGCCCATACTCATCTTCCATTCTC
CGCTCTTCATCCTCCAGTTGCCGACCTCGAGAGATCCTCTAGATTTAAGAAGGAGATATACATATG
>pykF
GAGGCAATTCCGACGTCTAAGAAACCATTATTATCATGACATTAACCTATAAAAATAGGCGTATCACGAGGCCCTTTCGTCTTCACCTC
GAGTGGTTCCTGCCCCAGATCGCGCATTTCCAGCACAACTTTACCGACAACTCTGCACATTTCTCGGTACAGTTCATCCTCTTTTGTCC

Data Set S4 - MG1655 library reference file.fasta

```
GATTTCCCATTCTTTATACCTATTTATCATGCCAACTATCAGCATATATCAATCTAACCAATTAAACAAAAATCAAACAAAATCAGACA
AATAACGCGATAAATTATTTTAACTGTAGCAATTGAGCGATGATATATTTATACACCGGATGAACTTTCACTTATCCTCACACTGACAA
CTTCGGCACCAGACGTTGCGCAAACAGTGAAGTTTTTGCGTAACCTTTTCCCTGGAACGTTAAATCTTTGATAACAATTTATTGTCTAA
CAAGTTGTATATTTTTTGAAACGCTGTTTTTGTTTTCCTTTTGGATTAATTTCAGCGTATAATGCGCGCCAATTGACTCTTGAATGGTT
TCAGCACTTTGGACTGTAGAACTCAACGACTCAAAAACAGGCACTCACGTTGGGCTGAGACACAAGCACACATTCCTCTGCACGCTTTT
TCGATGTCACCTATCCTTAGAGCGAGGCACCACCACTTTCGTAATACCGGATTCGCTTTCCGGCAGTGCGCCCAGAAAGCAAGTTTCTC
CCATCCTTCTCAACTTAAAGACTAAGACTGTCATGAAAAAGACCAAAATTGTTTGCACCATCGGACCGAAAACGGATCCTCTAGATTTA
AGAAGGAGATATACATATG
>sfmA
GAGGCAATTCCGACGTCTAAGAAACCATTATTATCATGACATTAACCTATAAAAATAGGCGTATCACGAGGCCCTTTCGTCTTCACCTC
GAGACGCGCCTGAACTTTTTGAGCAACTTCAGAGCGCACCTGCTGCGCAATCGTTTTACCGTCAATAATCTTTGCTGCCATCAGAGAGA
GGATTCCATCTGTTACGTAGATCGAAGGGGATGCGCCTATTTTGTCAGAAGCGGGGCGCGCTGTCAGGTTTCGTTTCAGATTTATCGCG
TGAAGCGACCTCTTGCGAAGGTGAGGCGCACCGTCGCTGAGACTGAAAGCTTCATTTTTCGTCCATGATGGCGTTGTAAATCTGGAACT
GATTTATTTCCTTGTCTAAGGATTAAGATAATTTAAGAAATACCTGACAATATAAAAAGAATTTTCAGCCTGGTAATTTACCGCTTCAG
GTCTATATTTGTGTTGAATATATTTTGCGCGGAAGTATTCATCTAACGGGGCTCTCTATTTTTTAGAATAGAGTGCATATTTTCAATTA
AGACATTCTTAGAGGATAAAAAGGAATTTACTACTATCAGTGTCTTAAATAAAGTAATCGGTTATATACGGATGTGGAGTCGATAAATG
AGATTGAAGGAATATATATGAAATTAAGATTTATTTCGTCTGCGCTGGCTGCCGCACTATTCGCCGCTACGGGTAGTTAGGATCCTCTA
GATTTAAGAAGGAGATATACATATG
>sfmC
GAGGCAATTCCGACGTCTAAGAAACCATTATTATCATGACATTAACCTATAAAAATAGGCGTATCACGAGGCCCTTTCGTCTTCACCTC
GAGGCACGTTATAAAGGCACCGGTACAAGTGCATCAGCAGGGCAAGCAAATGCTGACGCGACTTTTATTATGAGATATGAATAATCAAA
ACCACGTTGTTTTGAATTATATATCACGTCTTATAACAAAGTAATGTACCGGTTGTCTGAAGCGGTATGGTGGCAATGTAAATCGAAAT
CATGTTCACTTTGTATCATGCCGCTTTATTAAATGAAAAGGGAATGATGTGTTGTAAGAAACCAAAGCAATCATTTCTTTATATTCCTT
ATTTTTGCCGTCAGGAATACACAAGGCGTATTAACTATGATGACTAAAATAAAGTTATTGATGCTCATTATATTTTATTTAATCATTTC
GGCCAGCGCCCATGCTGCCGGAGGGATCGCATTAGGTGCCAGGATCCTCTAGATTTAAGAAGGAGATATACATATG
>b1997_b1998
GAGGCAATTCCGACGTCTAAGAAACCATTATTATCATGACATTAACCTATAAAAATAGGCGTATCACGAGGCCCTTTCGTCTTCACCTC
GATGGATCCGACTGGCCCCCTGAATCTCCAGACAACCAATATCACTTAAATAAGTGATAGTCTTAATACTAGTTTTTAGACTAGTCATT
GGAGAACAGATGATTGATGTCTTAGGGCCGGAGAAACGCAGACGGCGTACCACACAAGATCCTCTAGATTTAAGAAGGAGATATACATA
TG
>pphA
GAGGCAATTCCGACGTCTAAGAAACCATTATTATCATGACATTAACCTATAAAAATAGGCGTATCACGAGGCCCTTTCGTCTTCACCTC
GATGGATCCTAACAATCCCGGCGAAAAAATGAGTCTGGCACCGTTTATAGAACGAGCTGAAAAACTCTGTTAATTACCTAAAATAGCCT
TTTGATTTCCAATAAAAAAACCGCCTCAGTTCTTTCACCAGAACGGGCGGTTTTTAACATTTCAGCTGATGACCACCACGCTTTTTATT
GACCATTTTGCACGCAAACTGGAAAACCTGGCGTCGTCATCTATTCTTAAAGGGCAAGGCAACTAAGCCTGCATTAATGCCAACTTTTA
GCGCACGGCTCTCTCCCAAGAGCCATTTCCCTGGACCGAATACAGGAATCGTGTTCGGTCTCTTTTTATCTGTTAAAAGCCAGAAGCAT
TTCCTTCGCTGACTTTATAGTCAACCATAACACACACTCTACTGTCTGAGTCCAGCGTTTTTTAACATTCTTGTTAAGATTATGTGATC
TTTAGCGCGGGAGGAAAATATTGATGAAACAGCCTGCGCCCGTTTATCAGAGAATTGCGGGTCATCAATGGCGACATATCTGGCTTTCT
GGCGATCTCGAGAGATCCTCTAGATTTAAGAAGGAGATATACATATG
>mtr
GAGGCAATTCCGACGTCTAAGAAACCATTATTATCATGACATTAACCTATAAAAATAGGCGTATCACGAGGCCCTTTCGTCTTCACCTC
GATGGATCCTACCGGTCGTCGTCGTTTCGGTGGTGATGCGTAATCATCGCTGAACAGCGAACACAATCTGTAAAATAATATATACAGCC
CCGATTTTTACCATCGGGGCTTTTTTTCTGTCTTTTGTACTCGTGTACTGGTACAGTGCAATGCATAACAACGCAGTCGCACTATTTTT
CACTGGAGAGAAGCCCTCATGGCAACACTAACCACCACCCAAACGTCACCGTCGCTGCTTGGCGGCGTGGTGATTATCGGCGGCACCAT
TATTGGCGCAGGGATGTTCTCGAGAGATCCTCTAGATTTAAGAAGGAGATATACATATG
>thrU
GAGGCAATTCCGACGTCTAAGAAACCATTATTATCATGACATTAACCTATAAAAATAGGCGTATCACGAGGCCCTTTCGTCTTCACCTC
GAGTCCGTTCGGCTGTTTCTTCATCGTGTCGCATAAAATGTGACCAATAAAACAAATTATGCAATTTTTTAGTTGCATGAACTCGCATG
TCTCCATAGAATGCGCGCTACTTGATGCCGACTTAGCTCAGTAGGTAGAGCAACTGACTTGTAATCAGTAGGTCACCAGTTCGATTCCG
GTAGTCGGCACCATCAAGTCCGGATCCTCTAGATTTAAGAAGGAGATATACATATG
>msbB
GAGGCAATTCCGACGTCTAAGAAACCATTATTATCATGACATTAACCTATAAAAATAGGCGTATCACGAGGCCCTTTCGTCTTCACCTC
GATGGATCCCCGCAGCTACGGTTTGATTAATTAACATCCATTCGCAGCCGGTACGCAGTCAGTACCGGCTTTTTTTATTTGGTGCGGGG
CAAGTTGCGCCGCTACACTATCACCAGATTGATTTTTGCCTTATCCGAAACTGGAAAAGCATGGAAACGAAAAAAAAATAATAGCGAATA
CATTCCTGAGTTTGATAAATCCTTTCGCCACCCGCGCTACTGGGGAGCATGGCTGCTCGAGAGATCCTCTAGATTTAAGAAGGAGATAT
ACATATG
>dacA
GAGGCAATTCCGACGTCTAAGAAACCATTATTATCATGACATTAACCTATAAAAATAGGCGTATCACGAGGCCCTTTCGTCTTCACCTC
GATGGATCCGCCGAAGCCAGTACCTTGCAGCAACGTTTGCAAACCGAAGCCCAATTACAGTCATTTATTACTACCGCGCAGTAGCGTAA
AGCAGGCATCTGATGTGTCAATTTCTGTAAATGACGTTAACAAAGTCACGCGGAAAGTCAGATGCCTGCCGGTAGTGCATTTGCTATAG
TAGGGCACTTTTTTAATTCCATCACGGATGTCGTAGTTCAGACCATGAATACCATTTTTTCCGCTCGTATCATGAAGCGCCTGGCGCTC
ACCACGGCTCTTTGCACAGCCTTTATCTCTGCTGCACATGCCGATGACCTGACTCGAGAGATCCTCTAGATTTAAGAAGGAGATATACA
TATG
```

Data Set S4 - MG1655 library reference file.fasta

```
>dacB
GAGGCAATTCCGACGTCTAAGAAACCATTATTATCATGACATTAACCTATAAAAATAGGCGTATCACGAGGCCCTTTCGTCTTCACCTC
GAGAATGATTTCAGGACGGCGCACAGATTTCAGAAAATCCAGCTCTTCGCGTAATTTTTCAGCGCCGCGTAAGGTCATCGGAATAGCTT
GCATTTGTTATACCTCTTGAATATTCCTGATAGGGCAAGTCTTCACCCACCCCGGCTGTTCGGCCTGCCCGGCATAACTCCTTACCGGG
ACCAGAAGCAAAAAAATACCGACCCGGGTACAAGTCCCAGGTCAGCTACAATTCACATTTTGATAGTCATTTTACCCTGAAGTTCCCGA
AGGGTCATCGTTTACTTTATAGGGCGTTGCGCCGTAGTATGACGGCTCGATTCCAGGTTGTTAGCGCGAGATTATGCGATTTTCCAGAT
TTATCATCGGATTGACCAGCTGTATAGCGTTCAGTGTTCAGGCCGCAAATGTTGATGAGTACATTACTCAACTCCCCGCTGGGGATCCT
CTAGATTTAAGAAGGAGATATACATATG
>tesB
GAGGCAATTCCGACGTCTAAGAAACCATTATTATCATGACATTAACCTATAAAAATAGGCGTATCACGAGGCCCTTTCGTCTTCACCTC
GATGGATCCCGTCTGAATATCCGCGCTTTTATCTGCGCAAGCCGCCAACGCAATCGCAACCGCTAAACCACTGGCCATGTGCACGAGTT
TCATTCATTTCTCCTTATTATCAATGCACCAGCGGGCTAACTTTCCTCGCCGGAAGAGTGGTTAACAAAATAGTAACGTCAACAAGTGT
GGCACACATCACGCATTTCTGCCTGTAATTAGCCCGTAATTCAGACCATTGCACCCATCGGACCAGTAGCAAATTTGCGTTATACTCAA
CTCACTTTGGCTTGCTGCGGCAGCTTTGTTACTGGAGAGTTATATGAGTCAGGCGCTAAAAAATTTACTGACATTGTTAAATCTGGAAA
AAATTGAGGAAGGACTCTTTCGCGCTCGAGAGATCCTCTAGATTTAAGAAGGAGATATACATATG
>yigI
GAGGCAATTCCGACGTCTAAGAAACCATTATTATCATGACATTAACCTATAAAAATAGGCGTATCACGAGGCCCTTTCGTCTTCACCTC
GATGGATCCGGCAACATAAACACCGGCAACAACCAGCCCTGCAGAGTCCGCATCGGTCGTGTTCTCCGTAACGAATTATTGATAAGGTG
AATATTTTTCTGCGGGCTATTTTACATACTTAACTCCATAGCCGTTAGTTATCCCTTCCTAAAGTAGAAACCAACAACAAAGAAGCATA
AAATCAACATTTCATTAACCAATGAAGTGTAAGGATGATATGTCTGCCGTACTGACCGCTGAACAAGCCCTGAAATTAGTGGGTGAGAT
GTTTGTTTATCACATGCCATTTAACCGCCTCGAGAGATCCTCTAGATTTAAGAAGGAGATATACATATG
>cyoA
GAGGCAATTCCGACGTCTAAGAAACCATTATTATCATGACATTAACCTATAAAAATAGGCGTATCACGAGGCCCTTTCGTCTTCACCTC
GATGGATCCGCGCTACGAAAAACGCATCTGACGTAATCTGTAAATATTATTTAGCTGTAGTAATCACTCGCCGTAATTATTACGGCTAA
ATAAACATCAGTATCGCTTATTAATATTTGTTCTTTTGTGCGGCTTAGCGTTTGGAAATTATTGGCGCCATTTATAAATTCTATTTTTC
TTACACGATTCAGCTAATGAGTCTTTATTTTCTCATCACCCAGTTGTCACTCTAATGATAATTATTTGTTAAATAATTGTTTTATTTCA
CATTGGTTATACCAATTGCCCGCCCAGACTCCTTCAGCACTCCCCTTTTGTTATAACGCCCCTTTTGCAACAGCTTCTTAAAATCAACCT
GATATGTTTTGCAACATATGTGACCTGGCAGCCAAATCCAAGTAACAGGAATTTAATCATGTTTACAGTAATGTAACCTTCCCGTAAAA
TGCCCACACACTTTAAACGCCACCAGATCCCGTGGAATTGAGGTCGTTAAATGAGACTCAGGAAATACAATAAAAGTTTGGGATGGTTG
TCATTATTTGCAGGCACTGTATTGCTCAGTGGCTGTAATTCTGCGCTGTTAGATCCCAACTCGAGAGATCCTCTAGATTTAAGAAGGAG
ATATACATATG
>dgkA
GAGGCAATTCCGACGTCTAAGAAACCATTATTATCATGACATTAACCTATAAAAATAGGCGTATCACGAGGCCCTTTCGTCTTCACCTC
GAGCCAGCCGGACATAAACGATGTAAAGCCTCTGGTTAATAATGCAAATGCGCGGCAAGGATAGCAGAAAGTCATGGGAAATTCTGTGG
TATCCGCTCATGTTTCGCGCGGCGCTACGCAAACCCGAATCATCGGATTTAACGGTACACTGATATTGACGCTCATAATGTAAAAAGGT
TCTTTCAATGGCCAATAATACCACTGGATTCACCCGAATTATCAAAGCTGCTGGCTATTCCTGGAAAGGTTTACGCGCTGCGGATCCTC
TAGATTTAAGAAGGAGATATACATATG
>shiA
GAGGCAATTCCGACGTCTAAGAAACCATTATTATCATGACATTAACCTATAAAAATAGGCGTATCACGAGGCCCTTTCGTCTTCACCTC
GAGTTGCGTGATCCTGCATAGTCATTGTGACTAATGTTAATTTAGTCTGTTCAAGTTTAAGCGCATTAAAGACTTCTAAATTAATTGTC
GACGTTACAATTAAAAGATGCTTAATTTTATGCAATTCAAGCGCCCGAATAACAGGAAAGATGGCCATAGCATCGCCAATCTGATCGGG
AATATGGATGACAACAAAGTCTGTTTTTTCAATATTGAAATTATAAGCTTTATAATCGTAGTAACTAAATGCAATACGTCTCAACAATG
ATGCTAAAAACATACCTAACCTCGCCTCCCTACTGGTTATAATGCAATGCAGTCTATCAGACTCATCAGGGTGCCATTTTGTGCATATG
CGGACTTTTATGTTTCATATCTCTAACCTGTGGGTCCTCTGCTTAATCCTTAAACAACACCAGCAACTCCTGCGCTTTCATCTTCCGGA
TCCTCTAGATTTAAGAAGGAGATATACATATG
>ybfG
GAGGCAATTCCGACGTCTAAGAAACCATTATTATCATGACATTAACCTATAAAAATAGGCGTATCACGAGGCCCTTTCGTCTTCACCTC
GATGGATCCGGGTGCGCTTGTCTATGGCTCCGGAGGTCTGCTGATTGATAATGGTTAGTTACGCATAGCAGGATCTGGTCACCCCCGGT
TGCCACGAGATCCAGTAAGTTGGACACAACGCCCTGAATTCGCAGGCGTGCGGGCTTTACCCATCGCTGACGATGTCGCAGGGGTAATT
TTTGCTCTTAATGGTGGCGATCTTGGGGAAGATCTCGAGAGATCCTCTAGATTTAAGAAGGAGATATACATATG
>tig
GAGGCAATTCCGACGTCTAAGAAACCATTATTATCATGACATTAACCTATAAAAATAGGCGTATCACGAGGCCCTTTCGTCTTCACCTC
GAGAGGAGTGGGAAGGGTTGCAGGACACCGTCTTTGCCTCTCCTCCCTGTCGTGGAGCAGGAAGCATCGCGTAAAAACGCATTTGCAAC
TGTCGGCGCTTTTCCAGTATGTTGCTAAAGATTTTATGAAAAACGGCCTGCGGGCCGTTTTGTTTTGTCTGGATTTTGCGCTTTTTGCC
CAGCATTCAGACGAAAATTGCCCGGGAATTGTGAAAAAAATACGCGACAGCGCGCAATAACCGTTCTCGACTCATAAAAGTGATGCCGCT
ATAATGCCGCGTCCTATTTGAATGCTTTCGGGATGATTCTGGTAACAGGGAATGTGATTGATTATAAGAACATCCCGGTTCCGCGAAGC
CAACAACCTGTGCTTGCGGGGTAAGAGTTGACCGAGCACTGTGATTTTTTGAGGTAACAAGATGCAAGTTTCAGTTGAAACCACTCAAG
GCCTTGGCCGCCGTGTAACGATTACTATCGCTGCTGACAGCATCGAGACCGCTGGATCCCGGGGATCCTCTAGATTTAAGAAGGAGATA
TACATATG
>ykgD
GAGGCAATTCCGACGTCTAAGAAACCATTATTATCATGACATTAACCTATAAAAATAGGCGTATCACGAGGCCCTTTCGTCTTCACCTC
GAGTGTTTTTCCAGCCTTGCCAAAACCAATAATCACTGCCTGATATTTATTCATGATGATTTCCTGTCGAAACGCGTTCATTTGACATC
ATGACTATAGACGCAAAAGAGCAAATAACATTTCTTCACAACGCTTGATATTTTGTCTTATCGTCTGATATTTTCTTTTTCTTAATCAG
```

AATCACAAAACAAAAATCATGATTAACTATTTGATGGTATTAGATTAGTTATTTACTAAGATTGTTGGTGTTTGTAATCAAAAACCACT
CAGGAGTCTGATATGGATGCCCTTAGCCGTTTGTTGATGCTTAACGCTCCACAAGGAACGATCGATAAGAATTGCGTGTTAGGAAGTGA
CTGGCAGCTTCCACAGGATCCTCTAGATTTAAGAAGGAGATATACATATG
>sdaA
GAGGCAATTCCGACGTCTAAGAAACCATTATTATCATGACATTAACCTATAAAAATAGGCGTATCACGAGGCCCTTTCGTCTTCACCTC
GAGCATCGGGTATGGCTGTCCTGGTACGAACAGTATTTTGTATGGGGAATGACCGCAGGCATAATTCGTGAGCTGGCGCTGCAAATTGG
TGTGAAACCCTGACTATACTTATCTTTACATCTACAAAACACTACTTGAGACAATCATCGCAATATTAGTTAAATCGCGGTTTTTGATT
AGTTTAATTCATGTGAATAGTTAAGCCAGTCGCCGCGTTCCCTCTTACACTATGCGCTGTTATTAGTTCGTTACTGGAAGTCCAGTCAC
CTTGTCAGGAGTATTATCGTGATTAGTCTATTCGACATGTTTAAGGTGGGGATTGGTCCCTCATCTTCCCATACCGTAGGGCCTATGAA
GGCAGGTGGATCCTCTAGATTTAAGAAGGAGATATACATATG
>nmpC
GAGGCAATTCCGACGTCTAAGAAACCATTATTATCATGACATTAACCTATAAAAATAGGCGTATCACGAGGCCCTTTCGTCTTCACCTC
GATGGATCCAGGCGACACCCGTTGTTAACTTATCCATGGATTTCATAACCCCACCTCGCAGACAAAGCGGGTGTAAATTGAGGGAATAC
TACGAAACGTAACAGACTCGGAGTCAGTGAATAACTCAGGTATTGGGTTATCAGCTAATATCGAGACTCAAAAAATGGAAAAACCCGCT
CGACGGCGGGTTTAAGCTGTGTGACGAAGTAACCACTCTTAACAGCATAACCAATTTTTTACGTACGTAAACCACTAAATGATATTTGA
GAGAATGCTACCGAGTATTGAAAACACCACTACAAATACATAAGCAAATCTCAACAAATAACCAAAAAATAATTTCCAGTGTTATTTTT
AGCCGGTTTAAATTGAACCTTCAAATTATAGAGCACTTATAAATAACAGCCGTTAATATAAATTGGCTAATAGATTTATTTTTATTCAG
CCAAGAGCCATGAATAGGATTCGATAGAAAAAAGTTCAGATAAAAATAGAGATCTACTTCACAAATCAAACGAGAAACCAAAACTTACA
TCTTGAAATAATCACATTGATTAGATGAATATTTATCGCGCAGTGACATCATTTTTTAATAATAGTTCAAAAAAAGGGCTCACGATGAA
AAAATTAACAGTGGCAATTTCTGCTGCTCGAGAGATCCTCTAGATTTAAGAAGGAGATATACATATG
>idnT
GAGGCAATTCCGACGTCTAAGAAACCATTATTATCATGACATTAACCTATAAAAATAGGCGTATCACGAGGCCCTTTCGTCTTCACCTC
GATGGATCCTCGTAAACGGCCACCTGTTGTTTGTTGATGGCGGCATGTTAGTGGCTGTTTAACAGCCCCCCTACGTGCAGCGCGAAAGC
GCTGCTTTTCTGGCACTAATAAGAGATAAAGACTATGCCATTAATCATTATTGCGGCAGGCGTCGCGCTGCTTCTTATCCTGATGATCG
GCTTTAAAGTTAACGGCTTTATTGCCCTCGTTCTGGTAGCTGCCGTCTCGAGAGATCCTCTAGATTTAAGAAGGAGATATACATATG
>idnR
GAGGCAATTCCGACGTCTAAGAAACCATTATTATCATGACATTAACCTATAAAAATAGGCGTATCACGAGGCCCTTTCGTCTTCACCTC
GATGGATCCACCTGGACGGTGATGGAAACCCCTTATTTCTATTATGGGTTTGCTGGGCGTGTTAGCCATTAACGCCGTATTGCACTGAAA
GAATAAAATATTGTGATGTAGTTTCGGGGGAAGATTGAAGATTCCCCCGACGTTGGGAGAAGTTATGCGCAATCACAGAATTTCTTTAC
AGGATATCGCTACGCTGGCTGGCGTAACAAAAATGACCGTGAGTCGTTATATCCGCTCGCCCTCGAGAGATCCTCTAGATTTAAGAAGG
AGATATACATATG
>sbmA
GAGGCAATTCCGACGTCTAAGAAACCATTATTATCATGACATTAACCTATAAAAATAGGCGTATCACGAGGCCCTTTCGTCTTCACCTC
GAGGAGGCAAACTCCGGTTCAGTGATCGGCTGCGCCGCATGGACGGAGGTCAATGACGCCGCACACAGCACGGCAGAAAAAAGCAGACT
ACGTTTCAACGGTGGTGTCCATAGATTGAATGAAGGTAACTGAGCGTATTTATACTACTCATTGATAGATTGCAAAGGGCGAATTAGTG
CAATAATAGTGTTAAAAACGTAACGGCGCGTAAACACGATCTTTTTCCTGCTCACCAATGCCTGCATTTGTGATCCTGCGCGTGTTAAT
CCGCTTATGTAAACAATCTTTGGTTTGTGAAATCCGTCGAAACAATTCTTATGGTCAGGCTGGCGAACTAAGCGCCTTGCTATGGGTCA
CAATGGGCGGTTTCATTTGGCTGACGCTTTGTAGCGGTCATGCGGTTAATACACAGCAGTTGTTAAAACGATAAGAAGTTAGCAGGAGT
GCATATGTTTAAGTCTTTTTTCCCAAAGCCGGGAACGTTTTTTCTCTCGGCCTTCGTTTGGGCATTGATTGCCGTTATCTTCTGGCAAG
CCGGGGATCCTCTAGATTTAAGAAGGAGATATACATATG
>sbmC
GAGGCAATTCCGACGTCTAAGAAACCATTATTATCATGACATTAACCTATAAAAATAGGCGTATCACGAGGCCCTTTCGTCTTCACCTC
GATGGATCCTGATAAACAGGTGGCGCACTGGCCGCTGGTTACCCTGGAATCTGTCGGGGAAGGCAGCATGTTTTCTCGCCTGAGTGATT
ATTTCCACCATAAGGCCTGACCTTTCTTTTGCAGCAGACTGGCAGGAGTGCGAGTCTGCTCGCATAATCAACACTCATTCCTTGTGGTT
TTAATTTTGCAACTATACTGTATATAAAAACAGTATCAATGGAGGCGTCATGAACTACGAGATTAAGCAGGAAGAGAAACGTACCGTTG
CAGGTTTCCATCTCGTTGGCCCGTGGGAACAGACGGTAAACTCGAGAGATCCTCTAGATTTAAGAAGGAGATATACATATG
>idnK
GAGGCAATTCCGACGTCTAAGAAACCATTATTATCATGACATTAACCTATAAAAATAGGCGTATCACGAGGCCCTTTCGTCTTCACCTC
GAGCCAATCTATCGTCTGCTCGGTAACGGCAACAGTTTTCTTGCCCGCAACAACGCAGGACTGTGTTTTCACTTGCATAAATTCCTCAC
TGGTCAGGTAGACACCTCGGAAGCATTTAAGCGGTTTTAACTGTCATTATTTGTGATGAAGATCACGTCAGAAAATTGTTACATTACTA
TGTTACGCATAACGTGATGTGCCTTGTAATTCTTATCAGTAGAAATAAAAAAACGTGAAATTATTATGCCGCCAGGCGTAGTATCGCAG
CAGGTAAGATGATTCAGGAGATTTTAAATGGCGGGTGAAAGCTTTATTTTGATGGGCGTTTCAGGGAGTGGTAAAACATTAATTGGTAG
CAAGGTTGCCGCGTTATTATCTGCGGATCCTCTAGATTTAAGAAGGAGATATACATATG
>mltC
GAGGCAATTCCGACGTCTAAGAAACCATTATTATCATGACATTAACCTATAAAAATAGGCGTATCACGAGGCCCTTTCGTCTTCACCTC
GAGGCTGCTTGAGCAGGAGATGGTCAACTTCCTGTTCGAGGGTAAAGAGGTGCATATCGAGGGCTATACGCCGGAAGATAAAAAATAAA
AACAGTGCCGGAGCACGCCTCCGGCAACTTGCATAAAAACAAACACAACACGCACCCGGAATGATGAAAAAATATCTCGCGCTGGCTTT
GATTGCGCCGTTGCTCATCTCCTGTTCGACGACCAAAAAAGGCGATACCTATAACGAAGCCTGGGTCAAAGAGGATCCTCTAGATTTAA
GAAGGAGATATACATATG
>mltD
GAGGCAATTCCGACGTCTAAGAAACCATTATTATCATGACATTAACCTATAAAAATAGGCGTATCACGAGGCCCTTTCGTCTTCACCTC
GATGGATCCGCGTTTTGCATGGTTAAGGTCAAAGAAAGATAGGTTCTGATAAAACTTTCTTGTCATCGGCTCCGTTCGCCGTTATGATC
GGTCGTCTTTTAAGCAACTATTGACACACACATGAAGGCAAAAGCGATATTACTCGCCTCTGTCCTGCTCGTGGGTTGCCAGAGTACCG

```
GCAACGTTCAACAGCACGCACAGAGCCTTTCTGCAGCTGGTCAAGGGGAAGCAGCAAAGTTTACAAGTCAGGCACGATGGATGGACGAT
GGGACGTCCTCGAGAGATCCTCTAGATTTAAGAAGGAGATATACATATG
>pppA
GAGGCAATTCCGACGTCTAAGAAACCATTATTATCATGACATTAACCTATAAAAATAGGCGTATCACGAGGCCCTTTCGTCTTCACCTC
GATGGATCCACAGGGACCGGAAACCATTAATCAGGTTACCGAGCATAAGATGTCTGCCGAGTAAGCCCGTAGGCCGGATAAGGCGCTCG
CGCCGCATCCGGCATGAAATAAGGCGCACCGGGTTAACACATTTGCCCGATGCGCCTGCTTATCGGGCCTACAGACGGGGAAAGCCGCG
ATTATTTGCGCTGGCAGACACCCGTAGGACGGCTCGAGAGATCCTCTAGATTTAAGAAGGAGATATACATATG
>xthA
GAGGCAATTCCGACGTCTAAGAAACCATTATTATCATGACATTAACCTATAAAAATAGGCGTATCACGAGGCCCTTTCGTCTTCACCTC
GAGGAACCTTCGCCACGTACCGGTATAAAGGGTGCCGGAGCGTAAACAGGTATCATCCATTCATCAAAGTTTTCACGCGTAATTGGCTG
AGACATAGCGACCTCTACAGTAAATAATTCGTTATTTATATGTTAATAATAAGTAATGTTTGCGCTGTAAATGTAGATTGCAGGGTTCG
TGCCAGCCAGTGATAAAAGTGCATAAACGGCGGAGGCTAACTGGAAATCAAGGAGTTATAACCAAACCATATGCATTTAAAGTGCATAT
AAAGTGAATACGTTTGCGATGTGGGTGAATAAAAAGAATAAAAAACGCAATGTTATGCAGAAGTAAAATATAATTCTGGAATTGTGATC
ATTGACGAAATTTACTGGAAATTACTGCGCCATTCTGACGCAGCGCGCACCAAAAGCGGGCATTTTTTGCGCCATCGTTGACATCATTA
ACAACCATCGATCAAATCACTTAACAACAGGCGGTAAGCAACGCGAAATTCTGCTACCATCCACGCACTCTTTATCTGAATAAATGGCA
GCGACTATGAAATTTGTCTCTTTTAATATCAACGGCCTGCGCGCCAGACCTCACCAGCTTGAAGCCATCGTCGAAAAGCACCGGATCCT
CTAGATTTAAGAAGGAGATATACATATG
>fsr
GAGGCAATTCCGACGTCTAAGAAACCATTATTATCATGACATTAACCTATAAAAATAGGCGTATCACGAGGCCCTTTCGTCTTCACCTC
GATGGATCCTTAACAGCGCTAACGCCACGCCCCGCTGCAATAATTTCATACTTCTCTCCCTGACCTGATTTCAACGTGTTGAAAAATAG
TATGACGCGACATCATAGACAAGTCTGCGTCCGGTGAAGATAAATCTGCATCCAGATTAGAAATTCTGTATTCGGCGTTGAGACTTCCT
TCAGACGACGCATACAGGTATGTTAGTCTGGTAATAATCACAAATAAGATCATTCTTTACATCGCAACCCTGAAAAAGTGACTTTCCTA
TGGCAATGAGTGAACAACCCCAGCCTGTGGCGGGCGCGGCTGCGTCAACGACCAAGGCCCGAACATCGTTTCTCGAGAGATCCTCTAGA
TTTAAGAAGGAGATATACATATG
>rhaT
GAGGCAATTCCGACGTCTAAGAAACCATTATTATCATGACATTAACCTATAAAAATAGGCGTATCACGAGGCCCTTTCGTCTTCACCTC
GATGGATCCTGGTGTTTGGTGTGGTGGATTTCCATGGTCTGCTTATCGAAGTGCGGTTCCAGGGCATCGTAAGCATACGGCAGGGATGG
CAGGGTATAGCTCATATTCATCTCCAGTATTGTCGGGCGGCCGATTGTTAATGCCGCGTAAGCAGTTGGTTCATTATAGTTAATTAAAT
GATATTGAAAATGATTATCAATGCCGTACTTTTCGTAAGGGTATGGTTTTGCAGGAAAATGCCCGAGATGTGAAGCAAATCACCCACTT
AATGCCGTGATTGCCAGTAAATCGACAACGGCGGCAACAGGCGAAAGGTTAATCGACAGCACGATTTTTACACTCATCTCGTCGGAGAT
GTGACGCGACGAAAAATGATGAGGATAAGAAGATGAGTAACGCGATTACGATGGGGATATTTTGGCATTTGATCGGCGCGGCCAGTGCA
GCCTGTTTTTACGCTCCGCTCGAGAGATCCTCTAGATTTAAGAAGGAGATATACATATG
>aspA
GAGGCAATTCCGACGTCTAAGAAACCATTATTATCATGACATTAACCTATAAAAATAGGCGTATCACGAGGCCCTTTCGTCTTCACCTC
GATGGATCCTTGCTGCATCAGCACGAAATTCTTAAAGCCCTGGTTACGTACCAGTGACATACCGATAACTGACGTGAATATAACCAGCA
CGAGGGTCAGCAATACCCCCAATACATGGGCAACCTGAATAAAGATTGAAATCTCAATATAGACATAAAGGAAAATGGCAATAAAAGGT
AACCAGCGCAAAGGTTTCTCCTGTAATAGCAGCCGGTTAACCCCGGCTACCTGAATGGGTTGCGAATCGCGTTTAGCTTATATTGTGGT
CATTAGCAAAATTTCAAGATGTTTGCGCAACTATTTTTGGTAGTAATCCCAAAGCGGTGATCTATTTCACAAATTAATAATTAAGGGGT
AAAAACCGACACTTAAAGTGATCCAGATTACGGTAGAAATCCTCAAGCAGCATATGATCTCGGGTATTCGGTCGATGCAGGGGATAATC
GTCGGTCGAAAAACATTCGAAACCACATATATTCTGTGTGTTTAAAGCAAATCATTGGCAGCTTGAAAAAGAAGGTTCACATGTCAAAC
AACATTCGTATCGAAGAAGATCTGTTGGGTACCAGGGAAGCTCGAGAGATCCTCTAGATTTAAGAAGGAGATATACATATG
>rhaS
GAGGCAATTCCGACGTCTAAGAAACCATTATTATCATGACATTAACCTATAAAAATAGGCGTATCACGAGGCCCTTTCGTCTTCACCTC
GAGCATTCACGCTCGTAACGCGCCAGCATCACGCGCCCACTGGATGCGCCGAGATCGACGGCGACACAATTGCGAAAGGTCATAATGTG
ATCCTGCTGAATTTCATTACGACCAGTCTAAAAAGCGCCTGAATTCGCGACCTTCTCGTTACTGACAGGAAAATGGGCCATTGGCAACC
AGGGAAAGATGAACGTGATGATGTTCACAATTTGCTGAATTGTGGTGATGTGATGCTCACCGCATTTCCTGAAAATTCACGCTGTATCT
TGAAAAATCGACGTTTTTTACGTGGTTTTCCGTCGAAAATTTAAGGTAAGAACCTGACCTCGTGATTACTATTTCGCCGTGTTGACGAC
ATCAGGAGGCCAGTATGACCGTATTACATAGTGTGGATTTTTTTCCGTCTGGTAACGCGTCCGTGGCGATAGAACGGATCCTCTAGATT
TAAGAAGGAGATATACATATG
>aspC
GAGGCAATTCCGACGTCTAAGAAACCATTATTATCATGACATTAACCTATAAAAATAGGCGTATCACGAGGCCCTTTCGTCTTCACCTC
GATGGATCCCAGACGACACCGTTGCTGTGGGTATCGTTTACCAGTTCTAATAGCACACCTCTTTGTTAAATGCCGAAAAAACAGGACTT
TGGTCCTGTTTTTTTTATACCTTCCAGAGCAATCTCACGTCTTGCAAAAACAGCCTGCGTTTTCATCAGTAATAGTTGGAATTTTGTAA
ATCTCCCGTTACCCTGATAGCGGACTTCCCTTCTGTAACCATAATGGAACCTCGTCATGTTTGAGAACATTACCGCCGCTCCTGCCGAC
CCGATTCTGGGCCTGGCCGATCTGTTTCGTGCCGATGACTCGAGAGATCCTCTAGATTTAAGAAGGAGATATACATATG
>rob
GAGGCAATTCCGACGTCTAAGAAACCATTATTATCATGACATTAACCTATAAAAATAGGCGTATCACGAGGCCCTTTCGTCTTCACCTC
GATGGATCCATGGCCCCAGCATTATCAGGCTTAAAGACAGGATCAAATGCTTGTATTTCATATTGTTACCATTTTTATCTATTACACGA
GCAATTAGTTCGTCACGGTAAATAACATATTGATTAAGAACAAAATCTCAATACTTTTATTTCCGTCAAGCCCTAAAACATACTCTACT
AAGAAAAAAACACTGAATGCTAAAACAGCAAAAAATGCTATTATCCAATTACCTGATGTCAGGTGCTCGTTGTTGAAAGGATGAGGATA
TTTTATGGATCAGGCCGGCATTATTCGCGACCTTTTAATCTGGCTGGAAGGTCATCTGGATCAGCCCCTGTCGCTCGACAATGTAGCGG
CGAAAGCAGCTCGAGAGATCCTCTAGATTTAAGAAGGAGATATACATATG
>grxA
```

GAGGCAATTCCGACGTCTAAGAAACCATTATTATCATGACATTAACCTATAAAAATAGGCGTATCACGAGGCCCTTTCGTCTTCACCTC
GATGGATCCGAGTCGCTTACCGACAGCAGCGCCACCGCCAGTAGCATCATTCCGATAAATTCCAGTATCAACACGCCTTTAGGCAATTT
ACCGATCGCGCGCATACGCTTCCCTCTGCAAAGTGAGCCTTCAGTCTAAAACTTTTCACTGTATTGTGTTTAACAGTTATAGCTTTTAG
CAATTAATGCAACAGGTTAAACCTACTTTCAGCGAATACATTTTAGCGTGATCATTACAGGCATAAATCTATGAGGAGAGAAATAATGC
AAACCGTTATTTTTGGTCGTTCGGGTTGCCCTTACTGTGTGCGTGCAAAAGATCTGGCTGAGAAATTGAGCAATGAACGCGATGCTCGA
GAGATCCTCTAGATTTAAGAAGGAGATATACATATG
>rof
GAGGCAATTCCGACGTCTAAGAAACCATTATTATCATGACATTAACCTATAAAAATAGGCGTATCACGAGGCCCTTTCGTCTTCACCTC
GATGGATCCCCAGCGTCAGAGAGGCATCGAGAAACTGGTTGCGGTCGAGATCGGCCACATTAACCGTCGCTTTATAAATTGTCGCTTTA
AGCGCCATAAGAAAACTCCTTTCGAATCAGGAGGCCATTGTAACGGAATACGTGCAGTTGCGGCGCTGATATTGCAGAAATGAGTGTGG
TATAGTCACCTTGCGAAACAGAGTTTATCTTCGTAGGCTTAGACTTGCATCCACGGTTAAGTCAGAGTGCTGACAGGAGGGCATGTGGA
AAAATATTGTGAGTTAATACGCAAGCGGTACGCGGAAATCGCCAGCGGAGACTTAGGATACGTTCCGGACGCGCTGGGCTGCGTTTTGA
AAGTGCTGAATGAAATGGCTGCCGACGACGCCCTTTCTGAGGCTGTCAGGGAAAAAGCAGCATACGCTGCGGCAAACTTACTGGTGAGC
GATTATGTCAATGAATGATACGTATCAACCAATCAATTGTGATGATTACGATAATCTTGAGCTCGCCTGCCAGCATCATTTAAGTCGAG
AGATCCTCTAGATTTAAGAAGGAGATATACATATG
>aspU
GAGGCAATTCCGACGTCTAAGAAACCATTATTATCATGACATTAACCTATAAAAATAGGCGTATCACGAGGCCCTTTCGTCTTCACCTC
GAGAAGTGAAACGCCGTAGCGCCGATGGTAGTGTGGGGTCTCCCCATGCGAGAGTAGGGAACTGCCAGGCATCAAATTAAGCAGTAAGC
CGGTCATAAAACCGGTGGTTGTAAAAGAATTCGGTGGAGCGGTAGTTCAGTCGGTTAGAATACCTGCCTGTCACGCAGGGGGTCGCGGG
TTCGAGTCCCGTCCGTTCCGCCACTTATTGGATCCTCTAGATTTAAGAAGGAGATATACATATG
>rhaB
GAGGCAATTCCGACGTCTAAGAAACCATTATTATCATGACATTAACCTATAAAAATAGGCGTATCACGAGGCCCTTTCGTCTTCACCTC
GATGGATCCGTTCTATCGCCACGGACGCGTTACCAGACGGAAAAAAATCCACACTATGTAATACGGTCATACTGGCCTCCTGATGTCGT
CAACACGGCGAAATAGTAATCACGAGGTCAGGTTCTTACCTTAAATTTTCGACGGAAAACCACGTAAAAAACGTCGATTTTTCAAGATA
CAGCGTGAATTTTCAGGAAATGCGGTGAGCATCACATCACCACAATTCAGCAAATTGTGAACATCATCACGTTCATCTTTCCCTGGTTG
CCAATGGCCCATTTTCCTGTCAGTAACGAGAAGGTCGCGAATTCAGGCGCTTTTTAGACTGGTCGTAATGAAATTCAGCAGGATCACAT
TATGACCTTTCGCAATTGTGTCGCCGTCGATCTCGGCGCATCCAGTGGGCGCGTGATGCTGGCGCGTTACGAGCGTGAATGCTCGAGAG
ATCCTCTAGATTTAAGAAGGAGATATACATATG
>yhjK
GAGGCAATTCCGACGTCTAAGAAACCATTATTATCATGACATTAACCTATAAAAATAGGCGTATCACGAGGCCCTTTCGTCTTCACCTC
GATGGATCCGACATGGATTTACCGCCGCAGCCGCTGATACCTTACGCCGACTGGTAAGTTTTCAGATAGCGCCTCTCTTAATGCCGCTG
CGATCGGGTATACTCGGGCGGCAATCTGGGATTTCCGGGGGGAGACAATTTGCGCGTAAGTCGCTCGTTAACAATCAAGCAGATGGCAA
TGGTGGCAGCCGTTGTCCTGGTGTTCGTTTTTATTTTTTGCACCGTTTTGCTGTTCCATCTGGTCCAGCAGACTCGAGAGATCCTCTAG
ATTTAAGAAGGAGATATACATATG
>yhjJ
GAGGCAATTCCGACGTCTAAGAAACCATTATTATCATGACATTAACCTATAAAAATAGGCGTATCACGAGGCCCTTTCGTCTTCACCTC
GATGGATCCATGGCAAAACGCACGAATTATCCTCTTAATCTCATCACTTATGCCCGCACTCCCTTCCCCGAGTGCGGGTGTTTGCGCAT
AATTGACCCCTGTGCCTGAACTTACCACCTGGGTGAGTTCAGGTTTATTTCACTTCTTCCGTGACATTTTCATGTTCTTGCGGTCTAAC
ACGAAGTGTTTTTAACGTCATTTAGACTGACCGACAACGGGCAGTTTTTTTATTACCAGGATAGTTGATCAGGGGTTCACATGCAGGGC
ACAAAAATTCGACTTTTAGCGGGCGGTTTGCTGATGATGGCCACTGCTGGCTATGTGCAGGCAGACTCGAGAGATCCTCTAGATTTAAG
AAGGAGATATACATATG
>yjjV
GAGGCAATTCCGACGTCTAAGAAACCATTATTATCATGACATTAACCTATAAAAATAGGCGTATCACGAGGCCCTTTCGTCTTCACCTC
GAGGCCAGTCGCCAACGATACGCTGGTTGCCGAAGTGAGCGACGCTCCGCAGGCGAACGACCCGACATTTAACAATGAGGATCTGGCTT
GATTTGCCGTTTTATCGACACCCACTGCCATTTTGATTTCCCGCCGTTTAGTGGCGATGAAGAGGCCAGCCTGCAACGCGCGGCACAAG
CGGGCGTAGGCAAGATCATTGTTCCGGCAACAGAGGCGGAAAATTTTGCCCGTGTGTTGGCATTAGCGGAAAATTATCAACCGCTGTAT
GCCGGGATCCTCTAGATTTAAGAAGGAGATATACATATG
>yhjG
GAGGCAATTCCGACGTCTAAGAAACCATTATTATCATGACATTAACCTATAAAAATAGGCGTATCACGAGGCCCTTTCGTCTTCACCTC
GATGGATCCCGCACAAGGCTGGTTTCTTTCACGCCCGGCACCGATAGAAACGCTGAATACGGCGGTTCTGGCGCTATAAGCTGCCTCAT
TTTCCGCCTGGCTGGACTATCTTTAGGACTGGCACAGGAAGAATATGAGGAAGCGAACGATGAGCAAGGCAGGCAAAATAACCGCTGCG
ATTTCAGGGGCTTTCTTGTTGTTGATTGTCGTGGCGCTCGAGAGATCCTCTAGATTTAAGAAGGAGATATACATATG
>dsdC_dsdX
GAGGCAATTCCGACGTCTAAGAAACCATTATTATCATGACATTAACCTATAAAAATAGGCGTATCACGAGGCCCTTTCGTCTTCACCTC
GAGCGAAGGACTGATGCCTGGCAGCCACTTCAAAAGTATGCATTTTTGATAATTGCCAGCCGTTAAGCAGCCGATTTCTTATTTCACGA
AGGGGTTCCATAATCACCTCATTTTTCGCTTAAGTGTAAAAAAATAGCGGCAAAATTTCAGCTATGAGATGAGCTAAAGTGAACCATAT
CTCAATTCACCTTCATTTTTAGATGTAAATCACTCCATTGATGCAATTTACCTCATGTGAAAGGCAAATTTTATCGTTTGTCAGCCTGC
GTTGTTTTTTTGTCCAATATCATCAGGTTAATCACAGGGGAAGGTGAGATATGCACTCTCAAATCTGGGTTGTGAGCACGCTGCTTATC
AGCATCGTGTTAATTGTACTGACCATCGTGAAGTTCAAATTCCACCCGTTTCTGGCGGATCCTCTAGATTTAAGAAGGAGATATACATA
TG
>yhjC
GAGGCAATTCCGACGTCTAAGAAACCATTATTATCATGACATTAACCTATAAAAATAGGCGTATCACGAGGCCCTTTCGTCTTCACCTC
GAGTGACTGGCCCCCCTGAATACTCACTCCTGGAATACGCTGCTGTAAACTGATTTTCATTCCATGAATAAATATTGACTGCCTGTCAAA

CATGACTATTTGCATAACTGAATCTCCACCTGAATACGTTAAAAAGACTTAAGTAGTGGAAGGGTATTACCCGCGAGAAAAAATAAGAA
TTCGCCATTTGGCGGTGGCCATTCTACAGAGATGACGTGTAGAAAATAGTTACCGATATAAATAGTTACAGCTAAACGCCTGAAATTAC
ATGTCGAGGGCACTATTTAAAACAATTTTGAGGATTTCCTTATATTGGTGGTTAGTACGCATGCAATTAAAAATGAAATTCCGCGACCA
CAAGCCAAAATAACAAACGGCAAGGAGACAAAAATAAGCACAAATAGCCAACACGTCCTCTGTTCACTTTAAAGGGAATCGCTGAAAAA
TACGCTCTGTTTAAGGGGATTCACCTTTCTCAGAAAGCTATTCCGCCCTTTTCCTGCTGAGAAATCGCCACATTCGGCATGACAACATT
GTGAAACCCGGCATTAGATGTTAGAAAAAACAATAAACAATGCGATATGCGCGTTACTGTTTCGCTGTGAACAACAATCGGTCAAAGAA
ATGGATAAAATTCACGCAATGCAGTTGTTGGATCCTCTAGATTTAAGAAGGAGATATACATATG
>yhjB
GAGGCAATTCCGACGTCTAAGAAACCATTATTATCATGACATTAACCTATAAAAATAGGCGTATCACGAGGCCCTTTCGTCTTCACCTC
GATGGATCCACGCGCATATCGCATTGTTTATTGTTTTTTCTAACATCTAATGCCGGGTTTCACAATGTTGTCATGCCGAATGTGGCGAT
TTCTCAGCAGGAAAAGGGCGGAATAGCTTTCTGAGAAAGGTGAATCCCCTTAAACAGAGCGTATTTTTCAGCGATTCCCTTTAAAGTGA
ACAGAGGACGTGTTGGCTATTTGTGCTTATTTTTGTCTCCTTGCCGTTTGTTATTTTGGCTTGTGGTCGCGGAATTTCATTTTTAATTG
CATGCGTACTAACCACCAATATAAGGAAATCCTCAAAATTGTTTTAAATAGTGCCCTCGACATGTAATTTCAGGCGTTTAGCTGTAACT
ATTTATATCGGTAACTATTTTCTACACGTCATCTCTGTAGAATGGCCACCGCCAAATGGCGAATTCTTATTTTTTCTCGCGGGTAATAC
CCTTCCACTACTTAAGTCTTTTTAACGTATTCAGGTGGAGATTCAGTTATGCAAATAGTCATGTTTGACAGGCAGTCAATATTTATTCA
TGGAATGAAAATCAGTTTACAGCAGCGTATTCCAGGAGTGAGTATTCAGGGGGCCAGTCACTCGAGAGATCCTCTAGATTTAAGAAGGA
GATATACATATG
>yjjL
GAGGCAATTCCGACGTCTAAGAAACCATTATTATCATGACATTAACCTATAAAAATAGGCGTATCACGAGGCCCTTTCGTCTTCACCTC
GATGGATCCTGACCTTGATGCCACCCTTGCGCTCCGTAACCATTTGAATTCAGCTAAACAATCAATGATTCGATCCATTAATGAGAACA
CGCGTTATGCTCATTAAGTACGGATTAAAAACAGCAAACTATTAGCGGGTGCACGGTTTTATCAAAAGATAAATACCGGGAAATTCCCG
CTTACCTATGCTCACAATCAGTCGACTAAAAATGTTTTTGTGAGCGGTAGTAAAGTCCTAAAACTTTAACCTGCCGTTAATGTCAACCT
TGCGTAATGCACAATTACGTAAACAATTAAAATTATAAATACAGGAGTCCGGCGTGGAAAAAGAAAATATCACCATCGATCCGCGTTCT
TCATTTACTCCATCTTCGTCGGCAGATATTCCCGTGCCACCACTCGAGAGATCCTCTAGATTTAAGAAGGAGATATACATATG
>yhjY
GAGGCAATTCCGACGTCTAAGAAACCATTATTATCATGACATTAACCTATAAAAATAGGCGTATCACGAGGCCCTTTCGTCTTCACCTC
GATGGATCCTGTCAGTTTCAGGCACGCCCCACTCATTATCATGGTAGGCAATATAAAGCGGGTCCTGACTCACCCAGCCGCAACGTTCC
ATACTTTCCCTCGCTATGTACCCGATCGTAAAAATTTCAGCCGCTTTTCTTGACGCATCGGTGAAAAAGTCAATATTTAATACAGGGCT
ACTATGCCCGATATCCTTCATACAATGACAATAATATTGCCGAATTCGGCTCTTCCCTGGAGTCGTGTTAATGATCATAAAAAAAAGCG
GTGGTCGCTGGCAGCTAAGCCTGCTGGCGAGCGTGGTAATCAGTGCCTTTTTTCTCAACACAGCTTACGCCTGGCCTCGAGAGATCCTC
TAGATTTAAGAAGGAGATATACATATG
>pldA
GAGGCAATTCCGACGTCTAAGAAACCATTATTATCATGACATTAACCTATAAAAATAGGCGTATCACGAGGCCCTTTCGTCTTCACCTC
GAGGCGGTTAAATGGCATGTGATAAACAAACATCTCACCCACTAATTTCAGGGCTTGTTCAGCGGTCAGTACGGCAGACATATCATCCT
TACACTTCATTGGTTAATGAAATGTTGATTTTATGCTTCTTTGTTGTTGGTTTCTACTTTAGGAAGGGATAACTAACGGCTATGGAGTT
AAGTATGTAAAATAGCCCGCAGAAAAATATTCACCTTATCAATAATTCGTTACGGAGAACACGACCGATGCGGACTCTGCAGGGCTGGT
TGTTGCCGGTGTTTATGTTGCCTATGGCAGTATATGCACAAGAGGCAACGGTGAGGATCCTCTAGATTTAAGAAGGAGATATACATATG
>yjjI
GAGGCAATTCCGACGTCTAAGAAACCATTATTATCATGACATTAACCTATAAAAATAGGCGTATCACGAGGCCCTTTCGTCTTCACCTC
GATGGATCCCGGTGTCGTCGTCATTCAGGGTGGTCAGGTCCATCAATTTCAGTGCACGCAGGCTGCTTGCTTTCAGATCAGTCATTTCA
TTCTCCTGGCTTGTCGCCAATAAAATTCACCTTGCGAGTTTGTTAGTATTCTAACATCTACTCCGCAACACACTTCGATACACATCACA
ATTAAGGAAATCTACTTACAAGTTTGCATCACTGTAATGCGATCTGGTTCAAATAATTCACTTTCAAATGAATGCGTCAGTGGTGGCAA
ACGCATCAGGATCAAAGTGAACATCACGAAACTTCTTACAATGGCGCATCACCTTCGGCATAAAAGGAAAAGAAGATGCCCACTTCTCA
TGAAAATGCACTGCAACAACGTTGCCAGCAAATTGTCACCAGCCCAGTGCTTAGCCCGGAGCAGAAGCGCCATTTTCTGGCACTGGCTC
GAGAGATCCTCTAGATTTAAGAAGGAGATATACATATG
>yhjR
GAGGCAATTCCGACGTCTAAGAAACCATTATTATCATGACATTAACCTATAAAAATAGGCGTATCACGAGGCCCTTTCGTCTTCACCTC
GATGGATCCCAGGGTCCACAATGTCCCTCATGTCACCATTTTGTCTGATTATCAGTTTACTAGCGAAAGCTAAGAAATAAACCTACCAT
TGAAATTAAAACTTATCAGATTTAGCATGTGAATCAATTTAGAAGGTCACAAGACGTGTGATTTTAGTCTTTTCGTTTTTAACTTTTTC
TGTTGCGTAACGGAAAGTCAAAAAGTGAGCAAATTCCCGTCTCGCCGTAAACAATAAACCGGCGAAATGCTCACGGTTAGAATTGTCTC
ATGAACGGTACGGTTATTTCATAGGGATCAAGCAAAATGAATAACAATGAACCAGATACTCTGCCTGATCCCGCGATAGGCTATATCTT
CCAGAATGATATTGTGGCGTTAAAGCAGGCCTCGAGAGATCCTCTAGATTTAAGAAGGAGATATACATATG
>yfbJ
GAGGCAATTCCGACGTCTAAGAAACCATTATTATCATGACATTAACCTATAAAAATAGGCGTATCACGAGGCCCTTTCGTCTTCACCTC
GAGAACAGTCTCGCCATTCCGCCCGCAGATGCCATCGATCGTCAGGAGCGTCTGGTGCTGATTCAGTATCGTCCCAAATGATCTGGCTA
ACATTAGTCTTTGCCAGCTTGCTTAGCGTTGCCGGGCAGTTGTGTCAGAAACAGGCAACCTGCTTTGTGGCGATAAACAAACGGCGCAA
ACATATCGTGCTGTGGCTGGGACTGGCGCTGGCTTGTCTTGGTCGGATCCTCTAGATTTAAGAAGGAGATATACATATG
>yfbM
GAGGCAATTCCGACGTCTAAGAAACCATTATTATCATGACATTAACCTATAAAAATAGGCGTATCACGAGGCCCTTTCGTCTTCACCTC
GAGCCACTTGCCCGGTGATACCGCAGACAGATTGAATTATCAGAAAATGGCTCAGGTAGTGGATGGTGTTATAACTTTGTTATACAACA
GTAAATAATAAAAATTTCTCGAGGCGGGTCCAGAAAAAGCGACATGCACATTTTATTTTCCCTCATCTGTTTTAAGATGCGAAACATGA
AATTCAACGGAGGAAAAATAAATGGGAATGATTGGCTATTTTGCGGATCCTCTAGATTTAAGAAGGAGATATACATATG
>yfbE

GAGGCAATTCCGACGTCTAAGAAACCATTATTATCATGACATTAACCTATAAAAATAGGCGTATCACGAGGCCCTTTCGTCTTCACCTC
GAGGAGTACCCAGTCCGGCAATTGCAGCGAGCGCCAGTAAAATGATGATATATTTTTTTGACTTCAACGAAGAGCGGCAAAAAGCTAAC
ACGTAAACTCCACCTATAGACAAGCGCAACCAGACAATTACCGTGAAATTGAGCTACATTTCTGGCGATAATTCGCAGTTGGTGTAATA
TTAAAAATCCTACGATGTCGGCAAAATGCCTCAAAATTTTGCCAAATGCAAAGCCTAAATAAGAAAAAATATAAAAATTTCAATATTTA
CGTCTAATATTAGTTTCTTAAGGTTAAGTTAATATTCTATCCTTAAAATTTCGCTCCAAATGGCAAAATATACACAACACTCTTTATAG
CAAATATAAGTGGACAGGTATTCAATGGCGGAAGGAAAAGCAATGTCAGAATTTTTGCCTTTTTCGCGACGGATCCTCTAGATTTAAGA
AGGAGATATACATATG
>apbE
GAGGCAATTCCGACGTCTAAGAAACCATTATTATCATGACATTAACCTATAAAAATAGGCGTATCACGAGGCCCTTTCGTCTTCACCTC
GATGGATCCTCAACCTGCTGGACGACAACCAGTTCACTCGTGACGCTGGCATCAACACTGATAACATCGTAGCTCTGGGTCTGGTTTAC
CAGTTCTAATCTCGATTGATATCGAACAAAGGGCCTGCGGGCCCTTTTTTCATTGTTTTCAGCGTACAAACTCAGTTTTTTGGTGTACT
CTTGCGACCGTTCGCATGAGGATAATCACGTATGGAAATAAGCTTTACCCGCGTGGCACTGCTGGCTGCCGCGCTCTTCTTTGTTGGTT
GCGATCAAAAACCACAACCCGCCTCGAGAGATCCTCTAGATTTAAGAAGGAGATATACATATG
>yfbS
GAGGCAATTCCGACGTCTAAGAAACCATTATTATCATGACATTAACCTATAAAAATAGGCGTATCACGAGGCCCTTTCGTCTTCACCTC
GATGGATCCCGCCTGAATGAGGTCGATTTGGTCCTCCACAGTCTGGAGCAAATTACTGTGACCAAACAGCCAAATGGCGATGTTATTAT
TCAGTGAAACACCTGATCATGATTTAGCCCCGTCTCGTCGGGGCTTTTTTATGGCAGAATCAAGTCATCCCCCTCAATTAACAAGGATA
AGTTGTGAACGGTGAATTGATTTGGGTTCTTTCATTACTGGCCGTTGCCATCGTCTTGTTTGCGACGGGCAGAGTGCGTATGGATGCGG
TCTCGAGAGATCCTCTAGATTTAAGAAGGAGATATACATATG
>yfbR
GAGGCAATTCCGACGTCTAAGAAACCATTATTATCATGACATTAACCTATAAAAATAGGCGTATCACGAGGCCCTTTCGTCTTCACCTC
GAGTTCCGCATTGTCACGCTACCGCGTGTCGATGATATCGAGCTGTCTTTGAGCAAGTTCGCGCGTTTCCTTTCTGGTTATCATCAGCT
GTAATCTTAATTTCACTGCCGGAGATTGCATCCGGCAGCGTTATCCCGCCACAATGACCTGATGATGTCATCATACGTAAGGTCACTAT
GAAACAGAGCCATTTTTTTGCCCATCTCTCCCGCCTGAAACTCATTAACCGCTGGCCGCTCATGCGCAACGTGCGGACGGAAAATGTGT
CGGATCCTCTAGATTTAAGAAGGAGATATACATATG
>yfbV
GAGGCAATTCCGACGTCTAAGAAACCATTATTATCATGACATTAACCTATAAAAATAGGCGTATCACGAGGCCCTTTCGTCTTCACCTC
GATGGATCCTGCATCGATGATGGCAAATTTCAGTGAAGAACTACCGCAGTTCAGAACCAGTACTAACTTACTCGACATGGAAGTACCTA
TAATTGATACGTGGCTAAAAAAACGTCAGGGAGCCATAGAGCGTAGCGCATGATGACACCGACATTTATGATTAACATCATGCACTGCA
AATTTTTTTTGCATGATGGAAGAAATATCTTGAGTCTATGCCATTTTGCGTAATTTTCAGCCAATGGCATAATGTATATGAATTTGACG
ACTCAATGAATATGTACTACTTAGGCCTTCAGGCTGCTAAAGGATACCTGATAGTGGGTATGGAAGACAAAATGTTTTGAACGTTGTCC
CGCTGAGTTGTGAATTTGCACAAATTTTTGAATCTTTATATGTGAAGTTGAGGTGAAGCATGTCAACGCCGGATAATCGTTCTGTTAAT
TTTTTTAGCTTGTTTCGCCGGGGACAGCATTACTCAAAGACGTGGCCGCTCGAGAGATCCTCTAGATTTAAGAAGGAGATATACATATG
>yfbU
GAGGCAATTCCGACGTCTAAGAAACCATTATTATCATGACATTAACCTATAAAAATAGGCGTATCACGAGGCCCTTTCGTCTTCACCTC
GATGGATCCTTAAGCGCGCCTTCAAACAACTGGATAAAACTTTCCTTGATGATTTGTAATTGACCCCTGATTACGCATATAAAAGAAGG
CATACTCAGCCTTCTTTTTTTTCACCGTAGTAGAGTGATACAGGAGTCAACAATGGAAATGACCAACGCGCAACGTCTTATTTTGTCTA
ATCAGTACAAGATGATGACTATGCTCGATCCGGCACTCGAGAGATCCTCTAGATTTAAGAAGGAGATATACATATG
>glmU
GAGGCAATTCCGACGTCTAAGAAACCATTATTATCATGACATTAACCTATAAAAATAGGCGTATCACGAGGCCCTTTCGTCTTCACCTC
GATGGATCCTGCGCGTTATCGAGTTGACCAAAAAAGCGATGTAACACCGGCTTGAAAAGCACAAAAGCCAGTCTGGAAACAGGCTGGCT
TTTTTTTGCGCGTGTGACCCGTCCTGAATAGCGTTCACATAGATCCTGCTGATATAAAACCCCCCTGTTTTCCTGTTTATTCATTGATC
GAAATAAGAGCAAAAACATCCACCTGACGCTTAAATTAAGGTACTGCCTTAATTTTCTGCAGACAAAAGGCGTGACGATGGTCGAAAAT
GGCGCTTTCGTCAGCGGGGATAATCCGTTATTGAACAATTTATCCTCTGTCCATTTCACGATGAAAAAAATGTAGTTTTTTCAAGGTGA
AGCGGTTTAAATTCGTTCTCAAATTACAGTCAGGACGCGTATGTTGAATAATGCTATGAGCGTAGTGATCCTTGCCGCAGGCAAAGGCA
CGCGCATGTATTCCGATCTTCCGAAAGTGCTGCATACCCTTGCCGGGAAAGCTCGAGAGATCCTCTAGATTTAAGAAGGAGATATACAT
ATG
>eutB
GAGGCAATTCCGACGTCTAAGAAACCATTATTATCATGACATTAACCTATAAAAATAGGCGTATCACGAGGCCCTTTCGTCTTCACCTC
GATGGATCCGCGTGATGAATAACGGGCGTGTTTAGGCAACCACCTTACACGACAAGGGCTGACGATGAAACGTGATTGTTAGAATGCTG
TGAAAAATGCCAATCATCGATTCTTCCTGAGGGAACGACTTATGAAACTAAAGACCACATTGTTCGGCAATGCTCGAGAGATCCTCTAG
ATTTAAGAAGGAGATATACATATG
>cyaA_hemC
GAGGCAATTCCGACGTCTAAGAAACCATTATTATCATGACATTAACCTATAAAAATAGGCGTATCACGAGGCCCTTTCGTCTTCACCTC
GAGCCAGAGTGCAAGTGGGCTTTGGCGTGTGGCAATTCTTAAAACATTGTCTAACATGCTTGTTACCGTCATTATCATCCGTGGTCCAT
CCTAACATCCTTGCCAGAGTGATGTCAGTGTTGTGGTGAAACGTAGACGCCTGCGCAAACCGTAAAATGAGGTCTGGCAGTGGATCCTG
ACAGGCGTTTCACGCCGTTGTAATAAGGAATTTACAGAGAATAAACGGTGCTACACTTGTATGTAGCGCATCTTTCTTTACGGTCAATC
AGCAAGGTGTTAAATTGATCACGTTTTAGACCATTTTTTCGTCGTGAAACTAAAAAAACCAGGCGCGAAAAGTGGTAACGGTTACCTTT
GACATACGAAATATCCCGAATGCCGCGTGTTACCGTTGATGTTGGCGGAATCACAGTCATGACGGGTAGCAAATCAGGCGATACGTCTT
GTACCTCTATATTGAGACTCTGAAACAGAGACTGGATGCCATAAATCAATTGCGTGTGGATCGCGCGCTTGCTGCTATGGGGCCTGCAT
TCCAACAGGTAGATCCTCTAGATTTAAGAAGGAGATATACATATG
>der

```
GAGGCAATTCCGACGTCTAAGAAACCATTATTATCATGACATTAACCTATAAAAATAGGCGTATCACGAGGCCCTTTCGTCTTCACCTC
GATGGATCCACGGCAAACTGCTGATCCAGGCAAAAGACGGAACCGTGTACTCTATTACACGTTAATCGTCTCTGTCGTTCACTTTGAAA
ACGGCTCCTGGACAGGGGCCGTTTTCCTGTTTTTAACAACGACGCGAATATAGCGTGCGTTGTCTGATGATTTATAAAAATGAGGCTTT
AAACATGGTACCTGTGGTCGCGCTTGTCGGGCGCCCTAACGTAGGAAAATCCTCGAGAGATCCTCTAGATTTAAGAAGGAGATATACAT
ATG
>hdfR
GAGGCAATTCCGACGTCTAAGAAACCATTATTATCATGACATTAACCTATAAAAATAGGCGTATCACGAGGCCCTTTCGTCTTCACCTC
GATGGATCCTTGTGCCTCTTTGATGGTGAAATCACCATGACGAGAGAATCCACGTGGATAATGTTTATTGTCGAAATATCGATTAGTCG
TCGTAAAGCTTTCCGCCATCCTGCACGCTCCTAATTCTTTGACCGAGCTAGTTATGGCGCGGAGTATTAGTTACGCTTGACAGAGTGTA
AAACAAAACATTTAAATCATAACGACAAATAATTTTGCGGAGAGCACTGTGGATACGGAATTGTTAAAAACTTTCCTGGAAGTTAGCCG
AACGCGTCACTTTGGTCGAGCGGCTGAATCGCTCTATCTCGAGAGATCCTCTAGATTTAAGAAGGAGATATACATATG
>hscC
GAGGCAATTCCGACGTCTAAGAAACCATTATTATCATGACATTAACCTATAAAAATAGGCGTATCACGAGGCCCTTTCGTCTTCACCTC
GATGGATCCAACCGAATGCCACCGTAATGGTCGATGTTGATCGTCAGGGCTTTGTTGATTTACTGGCTGATCGTCTGAAATTTTACGCT
TAACACACGTCATTGATTAAGATCACCACGCGTTACCTGGGTGGTCTTAATCTGTTTATCTCTTTTTCTTTCAAGATATAAATGCAATG
GATAATGCAGAACTCGCCATTGGTATCGATCTCGGTACTACCAATAGTTTAATTGCCGTCTGGAAAGACGGTGCCGCGCAATTAATTCC
TCGAGAGATCCTCTAGATTTAAGAAGGAGATATACATATG
>acpP
GAGGCAATTCCGACGTCTAAGAAACCATTATTATCATGACATTAACCTATAAAAATAGGCGTATCACGAGGCCCTTTCGTCTTCACCTC
GAGGGTTGCATTCCTGGCATCCGACGAAGCAGCTTACATCACGGGTGAAACTTTGCATGTGAACGGCGGGATGTACATGGTCTGACCGC
GATTTGCACAAAATGCTCATGTTGCGCGCAGTCTGCGTGGTTATGAGTAATAATTAGTGCAAAATGATTTGCGTTATTGGGGGGTAAGG
CCTCAAAATAACGTAAAATCGTGGTAAGACCTGCCGGGATTTAGTTGCAAATTTTTCAACATTTTATACACTACGAAAACCATCGCGAA
AGCGAGTTTTGATAGGAAATTTAAGAGTATGAGCACTATCGAAGAACGCGTTAAGAAAATTATCGGCGAACAGCTGGGCGTTAAGCAGG
AAGAAGTTACCAACAATGCTTCTTTCGTTGAAGACCTGGGGGATCCTCTAGATTTAAGAAGGAGATATACATATG
>seqA
GAGGCAATTCCGACGTCTAAGAAACCATTATTATCATGACATTAACCTATAAAAATAGGCGTATCACGAGGCCCTTTCGTCTTCACCTC
GAGGCCAAACAGACCGTGGACAAGAACGATGGGAGAATTATTGTGCTGGTTTTGTGCAGTTTGCGCGCGGATATTCAATTTCATGGCAA
AGTTCTTTTTTTCGCGTTGTCGGGTTAGGGTATTATGTTGACCATTGTGCCACAGGGCTGCAACAAATAAGGTTTATTCCGAGTTTTTC
TGCAAGCCAGGCTTGACGCTATCCGCTGCCGGGATTTATTCATATACTCCTGGCGACTTGTATTCAGCTAAGACACTGCACTGGATTAA
GATGAAAACGATTGAAGTTGATGATGAACTCTACAGCTATATTGCCAGCCACACTAAGCATATCGGCGAGAGCGCATCCGACATTTTAC
GGCGTGGATCCTCTAGATTTAAGAAGGAGATATACATATG
>nupG
GAGGCAATTCCGACGTCTAAGAAACCATTATTATCATGACATTAACCTATAAAAATAGGCGTATCACGAGGCCCTTTCGTCTTCACCTC
GAGATCAGACGCTGACGACCCGCCATCCCTCTGCGGAATCTCGCCGTTATCTTTATAAAGTGAATACCGCGCAAAAATCCTACCGCCGC
CGATAATTCCATTAACCGCCCCTGACGATGCTCAGGGGCAAAAATGTTATCCACATCACAATTTCGTTTTGCAAATTGGGAATGTTTGC
AATTATTTGCCACAGGTAACAAAAAACCAGTCCGCGAAGTTGATAGAATCCCATCATCTCGCACGGTCAAATGTGCTTTTTCAAACACT
CATCCGCATCACGATGTGAGGAAATTAACATGAATCTTAAGCTGCAGCTGAAAATCCTCTCTTTTCTGCAGTTCTGTCTGTGGGGAAGC
TGGCTGACGACCCTCGGCTCCTATATGTTTGTTACCCTGAAGTTTGACGGTGCTTCTATTGGCGGGATCCTCTAGATTTAAGAAGGAGA
TATACATATG
>pmbA
GAGGCAATTCCGACGTCTAAGAAACCATTATTATCATGACATTAACCTATAAAAATAGGCGTATCACGAGGCCCTTTCGTCTTCACCTC
GAGGCGCAGCATCTTACCGATGAGCTGCAACTGGCGGCGGCGACCTTCCATCTTAATACGCTGGGCGAGCTCAATAGCCGCGCGCAGAT
CCGCATCTAACGGGATCTTATCCAGCGCGTTTTTCCCCAGATCAACAATTTCCGCGCCAAGGCGTTTTAGCTCCTCGGCATCACGTTTA
ATTTCACTTTTACTGACCCAGATAATTTCATCGTCTTCGTCTTCGATGTCATCACCGGGAACGTCGTCGAGCCAGTAGATCTTCTAGAT
TTAAGAAGGAGATATACATATG
>yodB
GAGGCAATTCCGACGTCTAAGAAACCATTATTATCATGACATTAACCTATAAAAATAGGCGTATCACGAGGCCCTTTCGTCTTCACCTC
GAGATGGAAAACTGGCCAACGTATTATCCATATCAGTTGAGTAGCGAAGAAGTGGTCGAGGAAATGATGTCTCATTGAGTATTCTCATG
ATAACGCCTCGATGCCGCTTTAGTAAGTTATCATAACTGCCACTGGTCCATCCACAAACGCCACTGAACGCAAGCTAGCTACAGACACG
CTCATCACTATGACGTGTCTGTATATTAATAAGCTAACCCGCATTGAGTTAACCAATAACGGATTCCATACACAATACGGCCAACAGAG
AAAGATACCTGTGCTCACGCCATTGCTTATATTGGCTGGTTACAATATGCACTATCAATTTTTTAAATAACAAATTATTAAAACCACAT
CTGTATAAGGAATTTTTAAGGTTCGTGGGATAGCTTGACTGTGAAAATCACAGGAGCTACAAAAATGAACCGATTCTCAAAAACTCAAA
TTTATTTACATTGGATAACGCTGCTTTTCGTTGCGGATCCTCTAGATTTAAGAAGGAGATATACATATG
>nlpC
GAGGCAATTCCGACGTCTAAGAAACCATTATTATCATGACATTAACCTATAAAAATAGGCGTATCACGAGGCCCTTTCGTCTTCACCTC
GATGGATCCGAATTTTCGCCGTCTGGATATCGAAGGTCACAGAATGCTGATTTCGACCATCTGATTTCATTAGTATCTTGAAAATCGGG
TAATTACGACGCTAAATTAATGCAAGAATAAAAAACAGAGGAACCGTACGGAATGCGTTTCTGCCTTATTTTGATCACAGCACTGCTTC
TGGCCGGGTGTAGCCACCATAAAGCACCGCCGCCAAATGCCAGACTTTCTCGAGAGATCCTCTAGATTTAAGAAGGAGATATACATATG
>nlpA
GAGGCAATTCCGACGTCTAAGAAACCATTATTATCATGACATTAACCTATAAAAATAGGCGTATCACGAGGCCCTTTCGTCTTCACCTC
GATGGATCCATTGGTCTGACTTCGCCGTTGATGTTGTCCGGCACATTGATGTTGCTGACTGCATTGTTGGTTACTGCAAAGGTGAAAAT
GAAGAAATCCTGATATCAGGTAAAGAAGCGTTGGTCAGTAGTTGTTGCTGTTCTTACATATCCGGGCATTCTACCTTACCCAACGCTTC
CCAATAATTGCTCTGATTATTACGTTCCATCGCCACGATGGCCTCGCGTATATGTTGTTTATTGTTTTCATCTGACAACATTAACTTTT
```

CCCACGCTTCATCTGATAATGACGCCTGTGGCGTGCAAATTTTACGTAAATCTTGTAATACCGTGGTTTTCTCTTTTGCGGACTGCAGT
GAGCTAAAAGGCGACTCGGCGTATACGATGCCTGGCATAGCGAAAAAAGTGAAAATAAGCAGTAGTGTCGTTGGTTTCATGGATTCAAC
CTTTTACAAATGTGTATCGAATGAAGCATAAGTGTAGTTTGCTTTCATCGCTCTCATCATGCTGTACGAAACGCGCTTGCGTGGTCAGT
AAGAAGTGCCAGACTTTATATTCCACTTTTATTCCTTTTTATTCTTATCGATAGCGTTTCGTTTTTTAAACCGCAGCGACCTTACCGCT
ATAGTCAGGTAATCATTAATAAAAGGATAAAAAAATGAAACTGACAACACATCATCTACGGACAGGGGCCGCATTATTGCTGGCCGGAA
TTCTGCTGGCAGGTTGCGACCAGAGTAGCAGCGCTCGAGAGATCCTCTAGATTTAAGAAGGAGATATACATATG
>cysH
GAGGCAATTCCGACGTCTAAGAAACCATTATTATCATGACATTAACCTATAAAAATAGGCGTATCACGAGGCCCTTTCGTCTTCACCTC
GATGGATCCCGCGTGATTTGTGGGATTAACCATCAGCCCGGTCTTGTAGGCCTGATAAGAACGCGTGAGCGTCGCATCAGGCAAGGCAA
ACAGTGAGGAATCTATGTCCAAACTCGATCTAAACGCCCTGAACGAACTGCCGAAGGTAGATCGCATTCTGGCGCTGGCGGAAACTAAC
GCCGAACCTCGAGAGATCCTCTAGATTTAAGAAGGAGATATACATATG
>nlpD
GAGGCAATTCCGACGTCTAAGAAACCATTATTATCATGACATTAACCTATAAAAATAGGCGTATCACGAGGCCCTTTCGTCTTCACCTC
GATGGATCCTCGGGGAGGCGAATTTATTATCGATACCGTGGAGGCCGTGCGCTTTGTCCCTTTAGTGAAGGGTGAGCTGGCTTAAAACG
TGAGGAAATACCTGGATTTTTCCTGGTTATTTTGCCGCAGGTCAGCGTATCGTGAACATCTTTTCCAGTGTTCAGTAGGGTGCCTTGCA
CGGTAATTATGTCACTGGTTATTAACCAATTTTTCCTGGGGGATAAATGAGCGCGGGAAGCCCAAAATTCACCGTTCGCCGCATTGCGG
CTTTGTCACTGGTTTCGCTATGGCTGGCAGGCTGTCTCGAGAGATCCTCTAGATTTAAGAAGGAGATATACATATG
>cysJ
GAGGCAATTCCGACGTCTAAGAAACCATTATTATCATGACATTAACCTATAAAAATAGGCGTATCACGAGGCCCTTTCGTCTTCACCTC
GATGGATCCGCGACCACATTTATGCCCTTCCGGGACGTGTGGTAAGCGGTGAGCGGCTTCGAAGGTGAAATCTTTAAATAACGTGGTGG
ACATCATAATTTCTCTACAGGACGATAAAACCGCCGTAGAGTACCGGAAAGGCTGTTTTTTCGCATTATCTAAAACGACTTTGTTGCGC
AAAATCGCTGATTTATCTTAATGATTGGCTAAATTCATTTGTTTTTCATTAGGTTGGTTAATCTATTTTGTTGTTAAAGACTATTGCTA
AAACAGGTTAGTCGATTTGGTTATTAGTTATCGCTATCCCGTCTTTAATCCACACCGTTTGCCCCGTTAACCTTACCTTCTCTTCTGTT
TTATGGGCGCTGACAGGGCGCAGAAACAGCTTTGCTTACTGGAACATAACGACGCATGACGACACAGGTCCCACCTTCCGCGTTGCTTC
CGTTGAACCCGGAGCAACTGGCACGCCTTCAGGCGGCCACGACCGATTTAACTCCCACCCAGCTTGCCTCGAGAGATCCTCTAGATTTA
AGAAGGAGATATACATATG
>cysD
GAGGCAATTCCGACGTCTAAGAAACCATTATTATCATGACATTAACCTATAAAAATAGGCGTATCACGAGGCCCTTTCGTCTTCACCTC
GATGGATCCCGCCAGGTTTAGGTGACGAGGCGTGTACGGGGAGAATAAAGCATACGCCGAGCGCCAGGGCAGCGGTACGGTGGCGCAAT
GCGGAAAACATAGTGAGTCCTTAAATACCATGCAAATTTTTTTACCGCCATAGTATGAAACTGCCGCTGCGCTAAAACAATTTCAAATC
TTCCTAAACGCCCGAAATCCGGTGCCTTAAGCACTTTTTGATATTAGCTTTGCCAAATCGTTATTCCGTTAAGGAACTACTCATTCTAA
TTGGTAATTTCATTCGTTCTCTTACGCTCCCTATAGTCGAAACATCTGATGGCAAGAAAATAGCGGTATTGCAAAGGAACGGTTATGGA
TCAAATACGACTTACTCACCTGCGGCAACTGGAGGCGGAAAGCATCCACATTATTCGCGAGGTGGCGGCAGAATTCTCAAACTCGAGAG
ATCCTCTAGATTTAAGAAGGAGATATACATATG
>nlpI
GAGGCAATTCCGACGTCTAAGAAACCATTATTATCATGACATTAACCTATAAAAATAGGCGTATCACGAGGCCCTTTCGTCTTCACCTC
GATGGATCCCTGCTGAACAGGGCGAGTAAGGTTGCCATTTGCCCTCCGCTGCGGCGGGGGGCTTTTAACCGGGCAGGACGCCTTGTTAG
CAACCGGGAACAGGACGTTCATTCAACCGTGGTCTTCGGGAGTGGGAAATGAAGCCTTTTTTGCGCTGGTGTTTCGTTGCGACAGCACT
TACGCTTGCAGGATGCAGTAATACTTCCTGGCGTAAAAGTGAAGTCCTCGCGGTACCCTCGAGAGATCCTCTAGATTTAAGAAGGAGAT
ATACATATG
>cysB
GAGGCAATTCCGACGTCTAAGAAACCATTATTATCATGACATTAACCTATAAAAATAGGCGTATCACGAGGCCCTTTCGTCTTCACCTC
GAGTCGGAAAAAGACGGAAAGGCGACTGGCTGGTCAGCATTTTATGTTGATGGCAAATGGGTTGAAGGAAAAAAATAACCTTTAATTCT
GTCAGGTTTTTATAAACAAAGGGTCGCGAAAGCGGCCCTTTTTTATTGCATATTATTTTTTCTTCACACCTATACACTAAGGCTATAAA
TGATATAGTGGTTATAGTTAGCACCTTTTTTATTATTAAATCGTATTAGTCACCCGCCAGGTGTGACGAAAAAACGATGTTCTGATGGC
GTCTAAGTGGATGGTTTAACATGAAATTACAACAACTTCGCTATATTGTTGAGGTGGTCAATCATAACCTGAATGTCTCATCAACAGCG
GAAGGACTTTACACATCACAACCCGGGATGGATCCTCTAGATTTAAGAAGGAGATATACATATG
>uxuR
GAGGCAATTCCGACGTCTAAGAAACCATTATTATCATGACATTAACCTATAAAAATAGGCGTATCACGAGGCCCTTTCGTCTTCACCTC
GAGACTTTGTTGGCGCAGTGACGGCGGCATATCAGCAGCTGTGCGAACGCGGTGCGCGCGAGTGTGTGGCTGCGCTGTAACTAACTGAT
TACCCTACAGACTTACTGGTCAATCAAACTGATATTTGGTTGACCAGTTTTCGTTTTTTTGCCCACCTGTACGTGCCAACTTCCAGTGC
TAATGGTATAGTTTGAGATTAACGGGGGCCGTAAAATTGCCCGTTGTAGGCCGGATAAGGCGTTCACGCCGCATCCGGCAAAAATTTGA
TTAACCGCACCTAACGGACACAACACCATGAAATCTGCCACCTCTGCGCAAAGACCTTACCAGGAAGTCGGGGCGATGATCCGCGATCT
GATCATAAAGACGCCGTACAATCCTGGGGATCCTCTAGATTTAAGAAGGAGATATACATATG
>cysZ
GAGGCAATTCCGACGTCTAAGAAACCATTATTATCATGACATTAACCTATAAAAATAGGCGTATCACGAGGCCCTTTCGTCTTCACCTC
GAGACGGCTGGTCCAGAAACCATGTACCAGTAAAGCGATTATGGCGATCGCGCCAACAATGATTAATATCAGACGCAAATCCTGCATCA
TTATATTCTCTGTTGTTCTAACACCTTGCCACCACGGCAAACATTTACTCACTAAGAGTATTTGCCGATTACCTCAAGTGCAAGTGCAC
TATTAACTTTCACAGCACAAAGATAGATGAAATCGTGCTTTTTGCTGTTTTTTCGAACATATCCTAACTGTCCATTGCGCAATTACCCG
GTAAAATACGCAGAATTTTCCTGGGATTGGTCAAAAGGAGCTCATCCTGATTATGGTTTCATCATTCACATCTGCCCCACGCAGCGGTT
TTTACTATTTTGCGCAAGGCTGGAAGCTCGTCTCGCAACGGATCCCGGATCCTCTAGATTTAAGAAGGAGATATACATATG
>cysT

```
GAGGCAATTCCGACGTCTAAGAAACCATTATTATCATGACATTAACCTATAAAAATAGGCGTATCACGAGGCCCTTTCGTCTTCACCTC
GATGGATCCTTCGAGTCTCGTTTCCCGCTCCAGTTTAAAAGACATCGGCGTCAAGCGGATGTCTGGCTGAAAGGCCTGAAGAATTTGGC
GCGTTAACAAAGCGGTTATGTAGCGGATTGCAAATCCGTCTAGTCCGGTTCGACTCCGGAACGCGCCTCCACTTTCTTCCCGAGCCCTC
GAGAGATCCTCTAGATTTAAGAAGGAGATATACATATG
>cysW
GAGGCAATTCCGACGTCTAAGAAACCATTATTATCATGACATTAACCTATAAAAATAGGCGTATCACGAGGCCCTTTCGTCTTCACCTC
GATGGATCCGTCGGCGTGTGGTAGGTCATTAATGGCGGAAGTTACCCAATTGAAGCGTTATGACGCGCGCCCGATTAACTGGGGCAAAT
GGTTTCTGATTGGCATCGGGATGCTGGTTTCGGCGTTCATCCTGCTGGTGCCGATGATTTACATCTTCGTGCAGGCATTCAGCAAGGGG
CTGATGCCGGTTTTACAGAATCTGGCCGATCCGGACATGCTGCACGCCATCTGGCTGACGGTGATGATCGCGCTGATTGCCGTACCGGT
AAACCTGGTGTTCGGCATTCTGCTGGCCTGGCTGGTGACGCGCTTTAACTTCCCTGGACGCCAGTTACTGCTGACGCTACTGGACATTC
CGTTTGCCGTATCGCCGGTGGTTGCCGGTCTGGTGTATTTGCTGTTCTACGGCTCTAACGGCCCGCTCGGCGGTTGGCTCGACGAGCAT
AACCTGCAAATTATGTTCTCCTGGCCGGGAATGGTGCTGGTCACCATCTTCGTGACGTGTCCGTTTGTGGTGCGCGAACTGGTGCCGGT
GATGTCTCGAGAGATCCTCTAGATTTAAGAAGGAGATATACATATG
>cysQ
GAGGCAATTCCGACGTCTAAGAAACCATTATTATCATGACATTAACCTATAAAAATAGGCGTATCACGAGGCCCTTTCGTCTTCACCTC
GAGTGATACGCAGATCGACCGTCGCTGCATTCACACTGGCGGCAATCAGCGTGGCCAGGAGCGTTGCGCTAAACTTAATCATCAGGGAC
ATCCTTTTATCATCGGGAATACGAAAGAAAAGGGAGAATAAACGTCTTACTTATAGAACAGTGAAGAATGCCACAATTTTACGCTTTGA
AAATGATGACACTATCACAGTTGGCGCATTCATTAACGATAGGGTATAAGTAAAACAATAAGTTAACACCGCTCACAGAGACGAGGTGG
AGAAATGTTAGATCAAGTATGCCAGCTTGCACGGAATGCAGGCGATGCCATTATGCAGGTCTACGACGGGACGAAGGATCCTCTAGATT
TAAGAAGGAGATATACATATG
>cysP
GAGGCAATTCCGACGTCTAAGAAACCATTATTATCATGACATTAACCTATAAAAATAGGCGTATCACGAGGCCCTTTCGTCTTCACCTC
GATGGATCCTCCTCGCATCGGATGAATCCAGCTATTTAACCGGTACACAGAATGTGATTGATGGCGGCAGCACACTGCCGGAGACGGTT
AGCGTCGGTATCTGATTCACCTCTGTTTCCTCCCTGCATTTGTGGGGAGGATTTCGTCTTGAACTAAGTTCACCAGGCTATTTTATTTG
TCATTTTGGCCCCGGGCAGTGCTCGAAATCCTCACGTACTATGTGTACGCTCCGGTTTCTCCGCGCTGTTCGTGTCCAAACTGACTGCA
ACAATTACGCCTGTTGAACCAAGTTCTTATTCCCTTTTCAACTTCCAAATCACCAAACGGTATATAAAACCGTTACTCCTTTCACGTCC
GTTATAAATATGATGGCTATTAGAAAGTCATTAAATTTATAAGGGTGCGCAATGGCCGTTAACTTACTGAAAAAGAACTCACTCGCGCT
GGTCGCTTCTCTGCTGCTGGCGGGCCATGTACAGGCAACGGAACTGCTGACTCGAGAGATCCTCTAGATTTAAGAAGGAGATATACATA
TG
>cysS
GAGGCAATTCCGACGTCTAAGAAACCATTATTATCATGACATTAACCTATAAAAATAGGCGTATCACGAGGCCCTTTCGTCTTCACCTC
GAGGCGGCAGTAGTCCAGGAAGTTTTTAACTGTTTCAGGTGCTTTATCGTCAAAAGTTTTGATGACAATATCGCCGTGATTGGTGTGGA
AAGTAACCATTTTTGCATCCTGTTCCGTTTGATTGGTGCTTCAACCCAGTTCGGGTCATATATAGGGTGGTGTTATAGCATAACCGCAC
GATCGGATCATCACGCAATGTATGCTGATTCGCGCGGGAAATATGGGTATTATACGCAACTCAATTACCCACACATGTCTAAACGGAAT
CTTCGATGCTAAAAATCTTCAATACTCTGACACGCCAAAAAGAGGAATTTAAGCCTATTCACGCCGGGGAAGTCGGCATGTACGGGATC
CTCTAGATTTAAGAAGGAGATATACATATG
>def
GAGGCAATTCCGACGTCTAAGAAACCATTATTATCATGACATTAACCTATAAAAATAGGCGTATCACGAGGCCCTTTCGTCTTCACCTC
GAGACCCAGTGAGCGATACGGACCATATCATCGCCGTACAAGCTGCTGATACTCATTAAACGCAGCCAAATTTCTGTATCGACCATCCT
TATCTCCCTGCCATAAGCAGCCTTAGCAATCTTTGCGATTGGTCAGTGATGCTGTCAATCAGAGGGGGATTTGTCTAGAATAGAAGAAA
TAATCTTTCTAACTCCTGAACACATCTCTGGAGATTTATGTCAGTTTTGCAAGTGTTACATATTCCGGACGAGCGGCTTCGCAAAGTTG
CTAAACCGGTAGAAGAAGTGAATGCAGAAATTCAGCGTATCGTCGATGAGGATCCCGGGATCCTCTAGATTTAAGAAGGAGATATACAT
ATG
>perR
GAGGCAATTCCGACGTCTAAGAAACCATTATTATCATGACATTAACCTATAAAAATAGGCGTATCACGAGGCCCTTTCGTCTTCACCTC
GATGGATCCCGACCAGTTGAGCGGATTCGCGTTTAAACTCTGCGCTGAAATTTCTTTTTTTCATTATGACACCTGTGTTGTTCTGAGGT
GAGCATATCACCTCTGTTCAGGTGGCCAAATTCAGTAAACCACTTCATCCCTAGTGAAAAGGATTCATGAATGAAGCTCTTAGCAAAAG
CACCTCTAAATCTGTTACGCGCCTTTGAAGCGGCTGGTCGTACCGGAGCTTTTGCCTCGAGAGATCCTCTAGATTTAAGAAGGAGATAT
ACATATG
>ppiD
GAGGCAATTCCGACGTCTAAGAAACCATTATTATCATGACATTAACCTATAAAAATAGGCGTATCACGAGGCCCTTTCGTCTTCACCTC
GAGCCGCAGACCGGTAAAGAGATCACCATCGCTGCTGCTAAAGTACCGAGCTTCCGTGCAGGTAAAGCACTGAAAGACGCGGTAAACTA
AGCGTTGTCCCCAGTGGGGATGTGACGAAGTTCAAGGGCGCATCTACTGATGTGCCTTTTTTATTTGTATTCGGTGACTTTCTGCGTCT
TGTGGGCTGACAATTGCCCCCGTTTCTTGTCACAATAGGCCTTTGCGCGCATCGATACGTTGCGTGAGGTACACAGTCATCTACAGCGG
AGTGTTGTTACACCATGATGGACAGCTTACGCACGGCTGGATCCTCTAGATTTAAGAAGGAGATATACATATG
>ppiC
GAGGCAATTCCGACGTCTAAGAAACCATTATTATCATGACATTAACCTATAAAAATAGGCGTATCACGAGGCCCTTTCGTCTTCACCTC
GATGGATCCACAGAGGGCGGTCATCAAGTCAATCAATGCCATATCGCTCATAAATTGACAATAAAATAACCATCCCCGATAATGACTTT
GTAGCCCATAGAGGCCCTAAGTTTTCTCCCCTTGAGGGACCTGCGAAGCCCGGATTTTCCGGGCTTTTGCCATATATGTACCAGCCACT
CCCTGACTGACAATTACGTTAGCATCTGCGGATCATGCTAAAATCGCCGCCTGACAATTATTCACTCAAGGCACGATCATGGCAAAAAC
AGCAGCAGCACTGCATATCCTTGTAAAAGAAGAGAAACTGGCTCTGGATCTTCTCGAGCAGATTAAGAACGGGGCCGGTCGAGAGATCC
TCTAGATTTAAGAAGGAGATATACATATG
>ppiB
```

GAGGCAATTCCGACGTCTAAGAAACCATTATTATCATGACATTAACCTATAAAAATAGGCGTATCACGAGGCCCTTTCGTCTTCACCTC
GATGGATCCCGTACATGCCGACTTCCCCGGCGTGAATAGGCTTAAATTCCTCTTTTTGGCGTGTCAGAGTATTGAAGATTTTTAGCATC
GAAGATTCCGTTTAGACATGTGTGGGTAATTGAGTTGCGTATAATACCCATATTTCCCGCGCGAATCAGCATACATTGCGTGATGATCC
GATCGTGCGGTTATGCTATAACACCACCCTATATATGACCCGAACTGGGTTGAAGCACCAATCAAACGGAACAGGATGCAAAAATGGTT
ACTTTCCACACCAATCACGGCGATATTGTCATCAAAACTTTTGACGATAAAGCACCTGAAACAGTTAAAAACTTCCTGGACTACTGCCG
CCTCGAGAGATCCTCTAGATTTAAGAAGGAGATATACATATG
>ppiA
GAGGCAATTCCGACGTCTAAGAAACCATTATTATCATGACATTAACCTATAAAAATAGGCGTATCACGAGGCCCTTTCGTCTTCACCTC
GATGGATCCCAACGCACCAGTCAGTGCGTAGGAGAGAAAGCTAATCCATGTGAGCTTGATGCGATTGCTGTTAGTCATGATTGGCCTGC
GTTCAAAAATAAAATGGCATAGCGGGATATGCCGCGAGCGGGCGATTTTAGGTGATTTTGTGATCTGTTTAAATGTTTTATTGCAATCG
GTTGCTAAATTGCATTTTAAGAGGTGATTTTGATCACGGAATAAAAAGTGATCGTCAGGTTACATATATTTCAGATACGTAAAATTAGG
TAAAGGGATGGCCTTGTTCTTGAAGGCTATTTAGAATCTCTTCACTTGCTTTTTTTCTGCTCTGTTTGTTAAGGAAATCTCATGTTCAA
ATCGACCCTGGCGGCGATGGCTGCTGTTTTCGCTCTTTCTGCTCTTTCTCCCGCAGCAATGCTCGAGAGATCCTCTAGATTTAAGAAGG
AGATATACATATG
>ybgQ
GAGGCAATTCCGACGTCTAAGAAACCATTATTATCATGACATTAACCTATAAAAATAGGCGTATCACGAGGCCCTTTCGTCTTCACCTC
GATGGATCCCAGTCGATGTCACGGCAGGTGAAGTAACCGCTAACGCAACCTACGTGCTGGATTATAAATAAGTATTATTTTGGCGGCAT
CATCGCCGCCTTTCATAAAATATTTATCAGGTATGGACACCGTGAATATTTATCGACTCTCTTTTGTATCCTGCCTGGTCATGGCGATG
CCTTGCGCAATGGCGGTCGAATTCAACTCGAGAGATCCTCTAGATTTAAGAAGGAGATATACATATG
>ybgS
GAGGCAATTCCGACGTCTAAGAAACCATTATTATCATGACATTAACCTATAAAAATAGGCGTATCACGAGGCCCTTTCGTCTTCACCTC
GATGGATCCTTTTCCAGCAATGCGACAGGAGGAAGTAACTCTTTGATTTCTTTGATGCGTAAATCGTCGTTCTGATAATTCATGTCTGT
TCCAGTGTTGCCATACTTATCTTAGTGAATGCAATACTTCCAATCTATATCTTCCGTCAGAATGTGTAAACGGGGTTTTACACTATGAA
CGGATTAATCCTGGAATTAGCAAGAAAAACGCCAGATTGTCGCGAAAAACGAGATCTCTCCTACAATTTCTAACTGTAACTCCTTTGTT
TGTTAATTATTTCAAGATTCTCTGCTGCGTTTCATAACCTGGCTGAAATCTTAAACCAATGCCTTATATTCACCTGCAAATGCACTGTT
GGAAGAGGTTATCCGACATAACGACCATAACAGGAGCATCCTATGAAAATGACAAAACTGGCCACATTATTTCTGACTGCCACTCTAAG
CCTTGCCAGCGGTGCCGCACTGGCTGCCGATAGCGGAGCGCAAACTCTCGAGAGATCCTCTAGATTTAAGAAGGAGATATACATATG
>yifB
GAGGCAATTCCGACGTCTAAGAAACCATTATTATCATGACATTAACCTATAAAAATAGGCGTATCACGAGGCCCTTTCGTCTTCACCTC
GATGGATCCCCGCACGGTGGGATAATAATCACCACCACGCTAATCACGACCAGGCTAATCACTCGTAGAAGGGCTGTCATTTTGTCTTT
TCTTGCATCTTGTTCGAAGGAATGCCTAAAGAGTTACCATAGGTTTTGTAAATAGTACAAGATATTTTTTGGCCGATTTTTATGATGCG
TTTAATAATTTCAATATAAATTGTTGTTTTTTAAAAATAAAATAGTGAGTGAAAAAAATTCAATTTCAGTATGAATGTTGTTCCCTTCC
TCGTAGTTCATCCTTTGTCCGCTGCATTTTTGCGAGCATCATTCCACCGCCAGAACAACTCAACTTCACTTGCACAAATAAACAGGAAT
CAGCCACTAAAATTCACGAATTAGCGGGCTACAGCAGCGATATCTTATGCACATAATGGTGACTCAAGGAGGGCTTATGTCACTGTCAA
TTGTTCATACCCGCGCAGCCCTGGGAGTAAATGCGCCCCCGATCACTGTTGAGCTCGAGAGATCCTCTAGATTTAAGAAGGAGATATAC
ATATG
>mscS
GAGGCAATTCCGACGTCTAAGAAACCATTATTATCATGACATTAACCTATAAAAATAGGCGTATCACGAGGCCCTTTCGTCTTCACCTC
GATGGATCCTGCCGGTCAGACTTCGATGATCGCTCGTCTGGAGAAAGCATTCCAGGAACTGAACGCGATCGACGTTCTGTAAGATATTC
CTTTCTGCTTATCTCAAGGCCCGCTCTGCGGGTCTTTTTTTCGCCAAAAGCAAAATTGCCTGATGCGCTACGCTTATCAGGCCTACACG
TATCCTGCAATATATTGAATTTGCAGGTTTTTGTAGGCCGGATAAGGCGTTCACGCCGCATCCGGCATGACAAAGCGCACATTGATATC
TGTCTGAGCCCCGCTCTGCGGGGATTTTTTCGCCAAAAGCAAGGTGATTCAGATGAGAAATCTGTGATCTATTTGGCAAAATTATGCTT
TATTGTTTACCCTTGTCAGACTGCCCGTCATAAGGCGGCGGAGTGTATTTCTCCATTTTGAGTCAGTTGAAAAGGAATATTGAATGGAA
GATTTGAATGTTGTCGATAGCATAAACGGCGCGGGAAGCTGGCTGGTAGCTAACCAGGCGCTGCTGCTAAGTTATGCAGTAAACATCGT
GGCGGCCTCGAGAGATCCTCTAGATTTAAGAAGGAGATATACATATG
>ykfG
GAGGCAATTCCGACGTCTAAGAAACCATTATTATCATGACATTAACCTATAAAAATAGGCGTATCACGAGGCCCTTTCGTCTTCACCTC
GATGGATCCACATCCAGAGGCTCACGCCATTCTGCGTATTATCGACTGACCGAAGGAGCAACAGATGAAACAGCTTTCCTTTTTACCCG
GCGAGATGACGCCACAGGACCGGCGTCTCATTCAGCGGGCGCTCAGGGCTCTGGACCTCGAGAGATCCTCTAGATTTAAGAAGGAGATA
TACATATG
>ybgC
GAGGCAATTCCGACGTCTAAGAAACCATTATTATCATGACATTAACCTATAAAAATAGGCGTATCACGAGGCCCTTTCGTCTTCACCTC
GAGCGCAGAAGGTTCTTTGGCAAGGGATTTTTTGCCCATTGCTTGCCGATATTGTTCTCATTGTCGGGCTGATTTTCTTCTTCTTTTAA
ATCAGAATTCTCTTTAAAAAATTATGGGCCGCTCCAGGCCCATAAATTTTTACGCTCCCTTAACTTGCCCTCATTCCCAAACCTCAATC
GCGCGCGTATAGTAGCAGCGTTTAAAAGCTCTAACTTTTGTTGCATTACCGGGATGTAAAGTGAATACAACGCTGTTTCGATGGCCGGT
TCGCGTCTACTATGAAGATACCGATGCCGGTGGTGTGGTGTACCACGCGGATCCTCTAGATTTAAGAAGGAGATATACATATG
>ybgD
GAGGCAATTCCGACGTCTAAGAAACCATTATTATCATGACATTAACCTATAAAAATAGGCGTATCACGAGGCCCTTTCGTCTTCACCTC
GATGGATCCGCGAAATGCACAGTGACGGTATGAAGATTGCCCGTCCGCGTCAGCTGTATACAGGATATGAAAAACGCGACTTTAAAAGC
GATATCAAGCGTTAATGGTTGATTGCTAAGTTGTAAATATTTTAACCCGCCGTTCATATGGCGGGTTGATTTTTATATGCCTAAACACA
AAAAATTGTAAAAATAAAATCCATTAACAGACCTATATAGATATTTAAAAAGAATAGAACAGCTCAAATTATCAGCAACCCAATACTTT
CAATTAAAAACTTCATGGTAGTCGCATTTATAACCCTATGAAAATGACGTCTATCTATACCCCCCTATATTTTATTCATCATACAACAA
ATTCATGATACCAATAATTTAGTTTTGCATTTAATAAAACTAACAATATTTTTAAGCAAAACTAAAAACTAGCAATAATCAAATACGAT

ATTCTGGCGTAGCTATACCCCTATTCTATATCCTTAAAGGACTCTGTTATGTTTAAAGGACAAAAAACATTGGCCGCACTGGCCGTATC
TCTGCTGTTCACTGCACCTGTTTATGCTGCTGATGAAGGTTCTGGCCTCGAGAGATCCTCTAGATTTAAGAAGGAGATATACATATG
>mscL
GAGGCAATTCCGACGTCTAAGAAACCATTATTATCATGACATTAACCTATAAAAATAGGCGTATCACGAGGCCCTTTCGTCTTCACCTC
GAGCATTGAGCAAGGCGATCACGTAATTATGTTCCTCACAGATAAAAAGTTTATTACCGACGTCGAAAGACTCTTCCAGCCAAGCCCTT
TCTTCTTGTAATTAATAAGGCGTCTAATGACGCCTTATTATTTCCCTTTGATTATCAAGGATTAATTAAATTCATTCCTGGCAGGAAAA
TGGCTTAACATTTGTTAGACTTATGGTTGTCGGCTTCATAGGGAGAATAACATGAGCATTATTAAAGAATTTCGCGAATTTGCGATGCG
CGGGAACGTGGTGGATTTGGCGGTGGGTGTCATTATCGGTGCGGCATTGGATCCCGGGATCCTCTAGATTTAAGAAGGAGATATACATA
TG
>ykfA
GAGGCAATTCCGACGTCTAAGAAACCATTATTATCATGACATTAACCTATAAAAATAGGCGTATCACGAGGCCCTTTCGTCTTCACCTC
GATGGATCCATTTGGTTTTCCCCCAGCACCAACACATGCATTCTGCGCTTGATGTGTTCAAAACCTGGCTACTGAATGAACTCAATCTG
GGTAAAATTCGTTGATCATGTTCACATTAATCGGGGCTCTGAATTCAAAGTAACTTTATTACTAAGCATTATTTTGTGAAGGCTCTCAC
TTTTGCTCGCGGCTCCGAAACTTTGCTGGGTTGAGATTTTAAGGTGTATCACGGCGGCTCATACTCTCAATAAATCCCTGTTAGTAAAT
GCTGCTATTATAACTCATTCATTGCCTGAGCCAGCAGTCGCGCAGCTGCCTCAAGCGCAGAGGATTGAAAAAATCTTCAGATATATATC
ATCACTCTGAACATACGCAATTAACGGCGGAATCTCACGATTCTACCGTTTTTTATTATTTTTCAGAGGAGTAGTGATGAATAATTCTG
AAGGTTTGAAGTCGTTTCAGCAATCGCTTGCTGATCTGCCGCAATGGGTATCAGAGCGGCTTCTGCAGCAAATTCTCGAGAGATCCTCT
AGATTTAAGAAGGAGATATACATATG
>ybgH
GAGGCAATTCCGACGTCTAAGAAACCATTATTATCATGACATTAACCTATAAAAATAGGCGTATCACGAGGCCCTTTCGTCTTCACCTC
GATGGATCCTCTTTGCCTTCCACCTGCAAACCATTCGGCGCGTAATCACTAATCGCCGCGCTGTTCAGTTTTTCGTTAATCAGTTGTTC
CAGTTCGGTGTTTTTCATCTTCGCGCCTTCCGTATTCTGATAGCTACACCGTTAACATAGCGTTACCTCGCCAGTTCGTCGATAGCCAA
TCTTTTGACATATTAAGAAACAAACCCGACGCCAAAAGACGCAAATTGTTTATATCATCACGCTTCAGCTAATGTATATTTATGCAATC
AGACAGCATATTGCATAAATAATCAGATCTGTATCAAAGTTTTAACTTTAAGGAAACACAAGCACGCCGCGCAACTATTACGTGACGTT
GCCGTCAATTGTGGAATTATTATGAATAAACACGCATCACAGCCGCGCGCTATTTACTATGTCGTTGCGCTGCAAATCTGGGAATATTT
TAGCTTTTACGGCATGCGCTCGAGAGATCCTCTAGATTTAAGAAGGAGATATACATATG
>ybgI
GAGGCAATTCCGACGTCTAAGAAACCATTATTATCATGACATTAACCTATAAAAATAGGCGTATCACGAGGCCCTTTCGTCTTCACCTC
GAGCGCATGCCGTAAAAGCTAAAATATTCCCAGATTTGCAGCGCAACGACATAGTAAATAGCGCGCGGCTGTGATGCGTGTTTATTCAT
AATAATTCCACAATTGACGGCAACGTCACGTAATAGTTGCGCGGCGTGCTTGTGTTTCCTTAAAGTTAAAACTTTGATACAGATCTGAT
TATTTATGCAATATGCTGTCTGATTGCATAAATATACATTAGCTGAAGCGTGATGATATAAACAATTTGCGTCTTTTGGCGTCGGGTTT
GTTTCTTAATATGTCAAAAGATTGGCTATCGACGAACTGGCGAGGTAACGCTATGTTAACGGTGTAGCTATCAGAATACGGAAGGCGCG
AAGATGAAAAACACCGAACTGGAACAACTGATTAACGAAAAACTGAACAGCGCGGCGATTAGTGATTACGCGCCGAATGGTTTGCAGGT
GGAAGGCAAAGAGGATCCTCTAGATTTAAGAAGGAGATATACATATG
>ykfJ
GAGGCAATTCCGACGTCTAAGAAACCATTATTATCATGACATTAACCTATAAAAATAGGCGTATCACGAGGCCCTTTCGTCTTCACCTC
GAGAGTTAAGCAGCAGCGCGTTGAATGCCGGGGAGCGTGGTTTACTAATTTTTATATGCGATATAAACCGCAACATTAAATCCAGCTTG
CAATGAAAATAACGCCCGCCTGGTATGTGCGCGTTTTCTGATTCCACAAACTGCAAGGAGGTAAATCATGACAAATCCTTTATCAATGA
CTCTTTGCAGACCTTTCCAGGATTAATTCTTTTTTTCTTGCCCTGGATTCGTCTGCCATTTCCTGATTTTTATATTTATATACTCTAAA
TAATTCGAGTTGCAGGAAGGCGACAAGCGAGTGAATCGCCAGGAGCTTACATAAGTAAGTGACTGGGGTGAACGAACGCAGTCGCAGTA
CATGCAACTTGAAGTATGACGAGTATAAATGGAATGGTACATGGGCAAATATATTCGTCCCTTATCCGATGCGGATCCTCTAGATTTAA
GAAGGAGATATACATATG
>ykfI
GAGGCAATTCCGACGTCTAAGAAACCATTATTATCATGACATTAACCTATAAAAATAGGCGTATCACGAGGCCCTTTCGTCTTCACCTC
GATGGATCCTTAACCTGCGAAGCCGACACCCTTGGCAGTTGCGGCTACGTATACATCGCCATTTACCCCACTCAGCGTTAATTACCTAC
ACGAGAGCAAACATGAAAACTTTACCTGCAATAACTCAGCGGGCGGTGAAGCCCTGCCTGTCACCCGTGGCTGTCTGGCAAATCTCGAG
AGATCCTCTAGATTTAAGAAGGAGATATACATATG
>yihN
GAGGCAATTCCGACGTCTAAGAAACCATTATTATCATGACATTAACCTATAAAAATAGGCGTATCACGAGGCCCTTTCGTCTTCACCTC
GAGCGCACCTTATCTGCCTGGGTGATGATGAGCCGCAGGTGACGGCATATGGCCTGGAAGAAGAGGTTGATTACTATGCGCCGGCGTTC
CGCTTTGAAGACGAAGATGATAATCCGTGGATCCCTTATCGCCAGATGAGTGAAACACCATTGCCAGAAAATCATTTACTGGATGCGCG
GTTACGTAAAGAAAAAGAAGATGCAATTAATCAGATAAATCATGTGCGTAACGTACTACAACAAATCAAACAAGAGGCAAACCATCTTC
TGAACCGGATCCTCTAGATTTAAGAAGGAGATATACATATG
>bolA
GAGGCAATTCCGACGTCTAAGAAACCATTATTATCATGACATTAACCTATAAAAATAGGCGTATCACGAGGCCCTTTCGTCTTCACCTC
GAGTGGTAAAAAATTCCCGCCATCATAACATTGCCAACGGCGAGGGGAAGTGGGTAAGGCATGTAAATTCATCATGTTGACGAAATAATC
GCCCCTGGTAAAAGAAACACTGATGCGAGGCCTGTGTTTCAATCTTTAAATCAGTAAACTTCATACGCTTGACGGAAAAACCAGGACGA
AACCTAAATATTTGTTGTTAAGCTGCAATGGAAACGGTAAAAGCGGCTAGTATTTAAAGGGATGGATGACATCTCAGCGTTGTCGGAGG
AGATATTTCATGATGATACGTGAGCGGATAGAAGAAAAATTAAGGGCGGCGTTCGGATCCTCTAGATTTAAGAAGGAGATATACATATG
>yihL
GAGGCAATTCCGACGTCTAAGAAACCATTATTATCATGACATTAACCTATAAAAATAGGCGTATCACGAGGCCCTTTCGTCTTCACCTC
GAGCGCCTATCCGCATGACTCTGGAACAAGCTCTGGAGTTCATCGATGATGACGAACTGGTAGAAGTGACTCCGACCTCTATCCGTATT
CGTAAACGTCACCTGACGGAAAACGATCGTCGCCGCGCCAACCGCGCACCGAAAGACGATTAATTTCGTTTTTAGTCATAAAAAACCT

Data Set S4 - MG1655 library reference file.fasta
GCCAGCGATGGCAGGTTTTTTTTTGCCTGTCGTGAGCGCACACCAATGCGAGTATCTCCCTTATAAGTCTGTGGTTACGTCAATAGAGA
GCTTTGTCACAATTATCTGCAAAGTCATACGCCGTTAATTGCTTTCTTTTTTGGCGTAAGCGTAAGATGCTTCATCTGGTTTAAACCAA
AAGGATTAAACAATGGCGGAGAATCAATCCACCGTAGAAAATGCAAAAGAGAAACTGGATCGGTGGTTGAAAGATGGCATCACCACGCC
GGGTGGAAAACTCCCTTGGATCCCGGGGATCCTCTAGATTTAAGAAGGAGATATACATATG
>ygfF
GAGGCAATTCCGACGTCTAAGAAACCATTATTATCATGACATTAACCTATAAAAATAGGCGTATCACGAGGCCCTTTCGTCTTCACCTC
GATGGATCCCTGCTCCTGCGTACCGATTAGCGAATACCAGTAATTCACTGATTCGACTATTTTCTAAAGGCGCTTCGGCGCCTTTTTAG
TCAGATGACAAAGTACAAAAGTGCTCAGACAGTCCCCTCGCCCCTTTGGGGAGAGGGTTAGGGTGAGGGGAACAGGCCGGCACTGGCGC
GAATATTTACCCTCACCCCGGCCCTCTCCCTGAAAGGGCGAGGGGGAAAAGCGTGCCAACATTGAAGATTGAGCCAGTTTGTTAGCAAT
CTCAAAGATACGTCAACGAATTAATTTTTCTCGGAAAAACAAATGGCTATAGCACTTGTGACTGGTGGCAGTCGCGGCATCGGGCGGGC
AACTGCATTACTGTTGGCGCAAGAAGGCTCGAGAGATCCTCTAGATTTAAGAAGGAGATATACATATG
>yihD
GAGGCAATTCCGACGTCTAAGAAACCATTATTATCATGACATTAACCTATAAAAATAGGCGTATCACGAGGCCCTTTCGTCTTCACCTC
GAGTATCTACGCCGCCCATTCGTCTGGCTTTACCGCCTGCCAGCACAACGCCTGTTATCGTCGTCATCAGATTCACCGATATCGCCTCT
TTTATTGTGGGATTGACCCTGCTAACGTGTCTGTCTCAAGAGTAAGGAGCATCACTATGAAATGTAAACGTCTGAATGAAGTTATTGAA
CTCCTCCAGCCAGCCTGGCAAAAAGAGCCAGACCTTAACCTGCTGCAATTTTTGCAGGATCCTCTAGATTTAAGAAGGAGATATACATA
TG
>ygfI
GAGGCAATTCCGACGTCTAAGAAACCATTATTATCATGACATTAACCTATAAAAATAGGCGTATCACGAGGCCCTTTCGTCTTCACCTC
GATGGATCCGAGCAGGCCGCTATTCAGTTTGATGATCAGGTCGATGTGGTCTTCCAGTTAGAACCTGTGGATCAACAACCCGCTAAAAC
ACCTGCAGCACAATAATATTTGTAGCCCGACGTATTCGTCGGGCTAATGCCTTTACCCGATATTGCCTCCTTTTTCTGTAATTTTCTGT
AAAACCATGACGTAATCACCTATCTTTGCTCGCGGTTTCATCAAGGATAGGCTGACTGAAACGGTTAAGCGGACAAGGATGCGGGGGAG
GGGGATGGACATTTTTATTTCGAAAAAGATGCGCAATTTCATTTTATTAGCGCAAACCAATAATATTGCCCGGGCGGCAGAAAAACTCG
AGAGATCCTCTAGATTTAAGAAGGAGATATACATATG
>mdtG
GAGGCAATTCCGACGTCTAAGAAACCATTATTATCATGACATTAACCTATAAAAATAGGCGTATCACGAGGCCCTTTCGTCTTCACCTC
GATGGATCCTTTAAAACACGCCCGGAAGGCGTTCCTTCACGCTATTAAATCTCCCATGCCGGATGCTTCAGAATGGCATCCGGCATTAC
CACAGCAAATCCCCCTGATTTAGCGATAAAAGCTCTCTGGATTGCGCCCCCTGGAAGTCGGGCGCATAATTAGTGTGCTTATCTTTTCT
TCTTATGTTCACCGCGCCTGGCGCACCAACAGCGGATTGCTATGTCACCCTGTGAAAATGACACCCCTATAAACTGGAAACGAAACCTG
ATCGTCGCCTGGCTAGGCTGTTTTCTTACCGGTGCCGCCTTCAGTCTGGTACTCGAGAGATCCTCTAGATTTAAGAAGGAGATATACAT
ATG
>yihX
GAGGCAATTCCGACGTCTAAGAAACCATTATTATCATGACATTAACCTATAAAAATAGGCGTATCACGAGGCCCTTTCGTCTTCACCTC
GAGGCGCTCTTTACCGACGAGCTGCCGCCCGCTGCGCTAAAATCACGCTTACAAGACAGCCAAATTGAAATCATTCTTCCCCAGGAAGA
CGCGTAGATTTTTGTGACCGTTAACCTGGCTTCATACCTTGCCACATAGCCAAACCCATCCTTTCCCGCTACAGTTAATTTCTTGTGGC
GCGAAAGGAGGCAAAAATGCTCTATATCTTTGATTTAGGTAATGTGATTGTCGATATCGACTTTAACCGTGTGCTGGGAGCGGATCCTC
TAGATTTAAGAAGGAGATATACATATG
>ygfU
GAGGCAATTCCGACGTCTAAGAAACCATTATTATCATGACATTAACCTATAAAAATAGGCGTATCACGAGGCCCTTTCGTCTTCACCTC
GAGCTTGATTGTGTGCCACCGCACAGGCAATTTCACAAGCATGACAGCCTATACATTCCGCAGCTTCAGCAGCAATAAACTTATTCATT
TGCATCCCTTTCATTTAATGAGTTATGTCTTCTCAAATTTCGCGATGCAATAAACGGGCATATATTAGGTAGCATGACCGTTTTAATTA
TAATTTTGCGACTGAGTTCAAATTATTGCCCCTCAATCTATGCCGCTTTCATTATTTCAACATTGCCGATCAAAAGAGCGCTATCCAGA
TAAAAACGATTAAATGCGAGAGTGCGACATGCCAGAATGATTAATGAAATATCAATATGAGAAAATTACAGATAATTCCATTGTCAGGT
TCTCTTCTCATTTTCGTGGTTTAACCACGATTATGTGATTGCCTCCGCAATTCCTGTCTCTAACTCCCCTTCCTCGCAAAAACTGGCAC
TCCACGAGCATGTGTTTAGACAGTTTCATTAACGTAAACGGTTGCTTTTTACTCTGGCGGGCGAAAGGAGAAACACTGATGAGCGCCAT
AGATTCCCAGGATCCTCTAGATTTAAGAAGGAGATATACATATG
>mdtK
GAGGCAATTCCGACGTCTAAGAAACCATTATTATCATGACATTAACCTATAAAAATAGGCGTATCACGAGGCCCTTTCGTCTTCACCTC
GAGGGTCGGGTAACTCCACCACATGCGTACGAAAATTTGGTTTCTCGTCAATCGACACCAGTTTTGCGGTGCCCTGTACAATCCCCGTA
AACATACTTACAACTCCTGAAATCAGTTAAGACATTCTGTTCAGCACAATAGCAGGTGGAAAACGCCCTTACCAGTGAAGGGGTAAGAA
TGGCTATTTTTTCACTGGAGAATTAATAAATCCTCGCTACAATAGACTGAATTTCCCCTGCTTCTTCTTTTTGCTGCCCATTCAGGCGG
CTTTTTAGTCTCTCATATAACTACAAATAAAAGGTGTTCACGTGCAGAAGTATATCAGTGAAGCGCGTCTGTTATTAGCATTAGCAATC
CCGGTGATTCTCGCGCAAATGGATCCCGGGATCCTCTAGATTTAAGAAGGAGATATACATATG
>focA
GAGGCAATTCCGACGTCTAAGAAACCATTATTATCATGACATTAACCTATAAAAATAGGCGTATCACGAGGCCCTTTCGTCTTCACCTC
GATAGATCTGCGTTTGCTGCACATCAGTCGTTGTTGAAGGCCTACGAAAAGCTGCAGCGCGCCAAAGCAGCATTCTGGGCAAAATAAAA
TCAAATAGCCTACGCAATGTAGGCTTAATGATTAGTCTGAGTTATATTACGGGGCGTTTTTTTAATGCCCCGCTTTACATATATTTGCA
TTAATAAAATAATTGTAATTATAAGGTTAAATATCGGTAATTTGTATTTAATAAATACGATCGATATTGTTACTTTATTCGCCTGATGC
TCCCTTTTAATTAACTGTTTTAGCGGAGGATGCGGAAAAAATTCAACTCATTTGTTAATTTTTAAAATTTATTTTTATTTGGATAATCA
AATATTTACTCCGTATTTGCATAAAAACCATGCGAGTTACGGGCCTATAAGCCAGGCGAGATATGATCTATATCAATTTCTCATCTATA
ATGCTTTGTTAGTATCTCGTCGCCGACTTAATAAAGAGAGAGTTAGTGTGAAAGCTGACAACCCTTTTGATCTTTTACTTCCTGCTGCA
ATGGCCAAAGTGGCCGAAGAGGCGGGTGTCTACTCGAGAGATCCTCTAGATTTAAGAAGGAGATATACATATG
>yihU

Data Set S4 - MG1655 library reference file.fasta

```
GAGGCAATTCCGACGTCTAAGAAACCATTATTATCATGACATTAACCTATAAAAATAGGCGTATCACGAGGCCCTTTCGTCTTCACCTC
GATGGATCCATGCGATCCATCACGGTTATACCTACACAAGCAACACGAATCATGGTCATTCCTTAAACATTTTGGGATGCGTTCATTTT
AATTTCACCAATTGTTTAAAAAGTGACGGCTGTCAATTTTTTGACGATAAATTACAAATACGATCAAAAACAGACAAGCAAAGCATGTC
ATAAATGACAAAAAGTGACATAACTGTATTCAGGAGAGGGTTATGGCAGCAATCGCGTTTATCGGTTTAGGACAAATGGGTTCGCCAAT
GGCGAGCAATTTATTGCAGCAAGGGCATCAACTTCGCGTCTCTCGAGAGATCCTCTAGATTTAAGAAGGAGATATACATATG
>mdtO
GAGGCAATTCCGACGTCTAAGAAACCATTATTATCATGACATTAACCTATAAAAATAGGCGTATCACGAGGCCCTTTCGTCTTCACCTC
GATGGATCCTCGCTAATCTTGAGCCGCAATAATGAGCGCGCTCAACTCCCTGCCATTACCGGTGGTCAGGCTGCTGGCGTTCTTTCATG
AAGAGTTAAGCGAGCGGCGACCAGGTCGCGTGCCGCAGACCGTGCAACTCTGGGTAGGCTGCCTGCTGGTGATTCTGATCTCGATGACC
TTTGAGATCCCTTTTGTGGCGTTATCGCTGGCAGTGCTGTTTTACGGTATTCAGTCGAACGCGTTTTACACCAAATTTGTCGCGACTCG
AGAGATCCTCTAGATTTAAGAAGGAGATATACATATG
>mdtN
GAGGCAATTCCGACGTCTAAGAAACCATTATTATCATGACATTAACCTATAAAAATAGGCGTATCACGAGGCCCTTTCGTCTTCACCTC
GATGGATCCGCGCGAAACTGGAAGAAATTATCGCCTGTCTGGATAATTTCGATTTGTGGGTGAATATCGTAACCCCAAATTAATGTCAC
TCGCTTTTGGGGGCGCTGAAGCCCCCAAAAGTATTTATTTTAAAAATATATACAGCCAATTAATAAATAAAATATATAAGTTATGAATG
CGGTGGCTGAATAAAGCTATTTTTATTCATCAAGAAAATAGCCATACTGACGCCTTCAATGGATATTGATATTCACTACAGATATTGTT
CGCAGCAGACAGGATGTATTTAATATGCCAACAGTTCTCTCTCGCATGGCGATGCAACTCAAAAAAACAGCCTGGATAATTCCCGTCTT
CATGGTTTCGGGATGCTCATTATCTCCGGCAATCCCGGTGATCGGCGCTTATTATCCCAGCTGGTTTTTCTGCGCCATTGCCAGCCTTA
TTTTGACGCTCATCACGAGGCGAGTTATTCAGCGGGCAAATATCAATCTGGCATTTGTCGGAATTATTTATACCGCCCTTTTTGCTCTC
TACGCCATGCTGTTCTGGCTGGCATTTTTCTAATCTATTTGAGACGCGCTCATGGAAAGTACGCCGAAAAAAGCTCCTCGCAGTAAATT
CCCTGCTCTGTTAGTGGTTGCGTTGGCGCTGGTTGCCCTTGTTTTCGTTATCTGGCGCGTCTCGAGAGATCCTCTAGATTTAAGAAGGA
GATATACATATG
>hycB
GAGGCAATTCCGACGTCTAAGAAACCATTATTATCATGACATTAACCTATAAAAATAGGCGTATCACGAGGCCCTTTCGTCTTCACCTC
GATGGATCCAATCGCTGGCGTTTACGACAGCTTATACACCGCTATTGAAAACAATGTGCCGGTGAATTTAAGCCAACTGGTAAAGGGAC
AAAGCCCGGCAGCATGAGCTGAGGCTTTGCCCGTTTTGCAGGCGTTACGCCTGTTTGGGGATGGGCGTGTCGATGAGTGTCGAAAATGA
CATTTCATCGGCATGTTTTCGTCAAAAATGACAATCACCTGAGGAATGCCTGGTGAATCGTTTTGTAATTGCTGACTCCACGCTCTGTA
TCGGCTGCCACACTTGTGAGGCCGCCTGTTCAGAGACGCATCGCCTCGAGAGATCCTCTAGATTTAAGAAGGAGATATACATATG
>flk
GAGGCAATTCCGACGTCTAAGAAACCATTATTATCATGACATTAACCTATAAAAATAGGCGTATCACGAGGCCCTTTCGTCTTCACCTC
GAGCGGTCACCTCACCCAAACGGCTAAATAAGTCGCGGGCATAAGGCATATTTTCATCAACAAGGATTTTCACGTTTGTGTTACCTGTA
TGAGACGAGAGTTAACCGGACAAGTGTGCCATAATCTCGCGGCCAGGCATACTTGCGAAGATTTCAGGTATAAGGATACGTAATGATAC
AACCTATTTCCGGCCCTCCTCCTGGGCAACCACCAGGTCAGGGAGATAATCTGCCGTCTGGCACGGGCAATCAGCCTTTAGGATCCTCT
AGATTTAAGAAGGAGATATACATATG
>spy
GAGGCAATTCCGACGTCTAAGAAACCATTATTATCATGACATTAACCTATAAAAATAGGCGTATCACGAGGCCCTTTCGTCTTCACCTC
GATGGATCCGCGGGATTAATGCTCGAAAAAATAAGCTAATACTTACCCGCAGAAATCATTCTGCGGGTATTAAGTTGCTTTCTTATAAA
TTAATACAGATTAATCCTGGTTATTTGCTTTATTTATCACCAGTCATCCGGTATAGTTCTTCATAATCTCTGCAAAATCATCGTGTTGT
AATATTCTCTCATCACTCTCCATCAAATTTTCTTTTTTTCTCCATAATTGGCGCAAAAGTGTTTTTTACACTTTCATTGTTTTACCGTT
GCTCTGATTAATTGACGCTAAAGTCAGTAAAGTTAATCTCGTCAACACGGCACGCTACTTAAGAAAGCCGTAATAAATAACTGAAAGGA
AGGATATAGAATATGCGTAAATTAACTGCACTGTTTGTTGCCTCTACCCTGGCTCTTGGCGCGGCTAACCTGGCCCATGCCGCAGACAC
CACTACCCTCGAGAGATCCTCTAGATTTAAGAAGGAGATATACATATG
>pinH
GAGGCAATTCCGACGTCTAAGAAACCATTATTATCATGACATTAACCTATAAAAATAGGCGTATCACGAGGCCCTTTCGTCTTCACCTC
GATGGATCCCCTCTGAGGATGATCCCGTTGAGAATAGTCCTGTTGTGACTAGTCCCGTTGTATCAAGTTCTAAAAGCAGTTTTCAATGA
TTAATAAATGGCAATTTTTACAAAGAGATAATAACGTAGATTGAAACAATATGTATTATTGATTGAGTAAACGGGATTATTCTTCACAC
TACATCCATTTTAGTGTCACTACCAAAGAAAAAGGGGCTACGTTTTCATGTAACCCCTTGATTTATTTGGTGGGCTGGCAGTGACTGAA
TTGGTGCTGTAATTGGTTGAAATATTTCATCTAATGTCTGGTTCAACTTTCCTTAGCTGCCTGAAAGTTTTTTGATCTTACCCAGCAAT
AACAGCGTTACGGTGCACCACGCCTGACTGAAGAACTGCGTGCTCAGGGTTACCACTTCAACGTAAAACCGTGGCAGGAGGTCCGGACT
GAAAAAACTGGTCAGGACATCATCGGCAGGGAACGCACTGCTAGTCTGTAAACTGAACCGACTGGGGCGCAGTATGTGGCATCTTGTCG
TGCTGTTGGAGGAGCTGTGCGAACGTGGTATTAACTTTCGTGCTCTGGCCCAATCTATATTTGCCCAACAATGGGGCTCGAGAGATCCT
CTAGATTTAAGAAGGAGATATACATATG
>purT
GAGGCAATTCCGACGTCTAAGAAACCATTATTATCATGACATTAACCTATAAAAATAGGCGTATCACGAGGCCCTTTCGTCTTCACCTC
GAGTCGGCTTCCTTTTTGCTGGTAAACGACATTTTCTCTTCGCCCTCACGAATGACTACGTATTTAACTTCAACCGCCATTTGCAGCCT
CTCATAATAACTGTGATTTTATACAGTATATTTCTTTTCGGTTGAGAAATCAACATCAGCAATAAAGACACACGCAAACGTTTTCGTTT
ATACTGCGCGCGGAATTAATCAGGGGATATTCGTTATGACGTTATTAGGCACTGCGCTGCGTCCGGCAGCAACTCGCGTGATGTTATTA
GGCTCCGGTGAACTGGGGATCCTCTAGATTTAAGAAGGAGATATACATATG
>mpaA
GAGGCAATTCCGACGTCTAAGAAACCATTATTATCATGACATTAACCTATAAAAATAGGCGTATCACGAGGCCCTTTCGTCTTCACCTC
GATGGATCCACGGCAAAGCCTATGTGGCTGAAGCAATTTCAGTATTAGCTCCGTTCTTTGAGGAAACTCTGTAATGAAACATATTCAAA
TCCGCAATAGTGATATGGACTGGCATATTGCTGCCAATAATCTGGGATAATTTTCATGAAAAATGTACTGATTCTTGGTGCTGGCGGTC
AGATAGCCCGCCATGTGATTAATCAACTGGCAGATAAGCAAACCATTAAACAAACTTTATTTGCCCGTCAGCCAGCAAAAATCCACAAA
```

```
CCCTATCCAACTAATAAAATGCAAACAACTTCCGGCAAAAAAGTGATACAAGATCGATGACATAGCGTTGAAGAAAAGGATTTGATAAT
CCGCAAAATTTGCACAACACTGCCTGTAATACTTCCTGAACCAGAGACGATCATGACCGTAACCCGCCCACCTCGAGAGATCCTCTAGA
TTTAAGAAGGAGATATACATATG
>spr
GAGGCAATTCCGACGTCTAAGAAACCATTATTATCATGACATTAACCTATAAAAATAGGCGTATCACGAGGCCCTTTCGTCTTCACCTC
GAGGGTGTTGCTGACGCCATTAAGTGATCGATTAATCACCTTTTTTGACAAATCACTACCAGGAAAAAACAAACATTTCCAAAACAAAT
AACTCACAGTAATTAACATCATCAGGGTTATTTTTATAGTGAGGATAATCCTGATGATGCGCACCGTGCTTTCATCTATCGAACGCAAA
AATCATTCTCTAAGTAAATGAATGGATTGCATGCGTTTCACTCAATTGTACTTTAATTGACCAACCCCGCTTATTAACTTTCTGTATCA
CTTTTTCTTATAAAAAATCATGTAAAACCGCTCGCCAAGACCGCACCAATCGGGTAATCTCGAACTCGTTTTGCCTCGGCGGTAGATTA
TCCTCACAGCATATAATTTTGTGCGTTAGTCCACAGATTTGGCCTTAAGGAATTGTTTCAACATGCCCAGGTAATTAGTCTCGTGTCGC
TTGGCATTTTTTTATAACGATATTTGTCGTTAAGGACTTCAAGGGAAAACAAACAACATGGTCAAATCTCAACCGATTTTGAGATATAT
CTTGCGCGGGATTCCCGCGATTGCAGTAGCGGTTCTGCTTTCTGCAGGATCCTCTAGATTTAAGAAGGAGATATACATATG
>dxr
GAGGCAATTCCGACGTCTAAGAAACCATTATTATCATGACATTAACCTATAAAAATAGGCGTATCACGAGGCCCTTTCGTCTTCACCTC
GAGGCCGTTCTCAGGACGATGTACAGAAACTGACTGATGCTGCAATCAAGAAAATTGAAGCGGCGCTGGCAGACAAAGAAGCAGAACTG
ATGCAGTTCTGATTTCTTGAACGACAAAAACGCCGCTCAGTAGATCCTTGCGGATCGGCTGGCGGCGTTTTGCTTTTTATTCTGTCTCA
ACTCTGGATGTTTCATGAAGCAACTCACCATTCTGGGCTCGACCGGCTCGATTGGTTGCAGCACGCTGGACGTGGTGCGCCATAATCCC
GAACACGGATCCTCTAGATTTAAGAAGGAGATATACATATG
>tilS
GAGGCAATTCCGACGTCTAAGAAACCATTATTATCATGACATTAACCTATAAAAATAGGCGTATCACGAGGCCCTTTCGTCTTCACCTC
GAGCCATCCGTGTCGATCCATACACGCAAAAACGCTTCACCTTCTTTAACGATCCGGACGGGCTGCCGTTGGAACTGTATGAGCAGTAA
GGCTTGTCATCGCCGCATTTGCCCGGTAACGTGCCGGGCATTGCTACTGTAAAATCGCACCATCATGACACTCACGCTCAATAGACAAC
TTCTCACCTCACGCCAGATTCTGGTGGCCTTTAGCGGCGGGCTTGACTCCACCGTTCTGCTGCGGATCCTCTAGATTTAAGAAGGAGAT
ATACATATG
>trxC
GAGGCAATTCCGACGTCTAAGAAACCATTATTATCATGACATTAACCTATAAAAATAGGCGTATCACGAGGCCCTTTCGTCTTCACCTC
GAGGTACGCGGGTTGTGGGTACGTTTATCAGAATCATCATCACTGCGGACATACATCACTTTGACCTTGCCGCTTTTACCTTTCATTTC
ATCGTTCATGCTTTTCTCCACCAGCGCTGCGCGAAGCGCGCAGATTACCCGAAGTCCGCGCGGTTCGCCATGATTTCGTACCAAAGCCT
GCGACTATCATACCTATTGAATAAAACAGATTGTTGTCTGGAACAATGTCCCCGATAATATGTAACATATTAGAAACATACCGGCGTCG
TTGCCGATAAGTCTCCTTACTCATCCCGAGGTTAGTTATGAATACCGTTTGTACCCATTGTCAGGCCATCAATCGCATTCCCGACGATC
GGATCGAAGATGCGGCAAAAGGATCCTCTAGATTTAAGAAGGAGATATACATATG
>trxA
GAGGCAATTCCGACGTCTAAGAAACCATTATTATCATGACATTAACCTATAAAAATAGGCGTATCACGAGGCCCTTTCGTCTTCACCTC
GAGGGGCTTCTACAACCTTCGGATGCAGGGCGAAGTCGGAAAACTTCTGTTCTGTTAAATGTGTTTTGCTCATAGTGTGGTAGAATATC
AGCTTACTATTGCTTTACGAAAGCGTATCCGGTGAAATAAAGTCAACCTTTAGTTGGTTAATGTTACACCAACAACGAAACCAACACGC
CAGGCTTATTCCTGTGGAGTTATATATGAGCGATAAAATTATTCACCTGACTGACGACAGTTTTGACACGGAGGATCCCGGATCCTCTA
GATTTAAGAAGGAGATATACATATG
>pheM
GAGGCAATTCCGACGTCTAAGAAACCATTATTATCATGACATTAACCTATAAAAATAGGCGTATCACGAGGCCCTTTCGTCTTCACCTC
GATGGATCCCGACCGTAAGATCCTGGCTGATATCGCAGTATTCGACAAAGTAGCGTTCACCGCTCTGGTTGAAAAAGCGAAAGCAGCTC
TGGCATAAGCCAGTTGAAAGAGGGAGCTAGTCTCCCTCTTTTCGTTTCAACGCCATCAAAACATTGACTTTTATCGCCGTAGCCTTTTC
AATAAAGGTCTTTTGAAGAGTACCAAAAGGTAACGCAAGCAATGAATGCTGCTATTTTCCGCTTCTTTTTTTACTTTAGCACCTGAATC
CAGGAGGCTAGCGCGTGAGAAGAGAAACGGAAAACAGCGCCTGAAACTCGAGAGATCCTCTAGATTTAAGAAGGAGATATACATATG
>pinR
GAGGCAATTCCGACGTCTAAGAAACCATTATTATCATGACATTAACCTATAAAAATAGGCGTATCACGAGGCCCTTTCGTCTTCACCTC
GATGGATCCAAGAAAACATCATGGCTGGATAACAAAGAATTATTACAAACAGCGATATCAGTTCTTAAAGACAACCAAAATTTAAATGG
TAAAATGTAATATAATAAACTTACTTTTTTATCATTTTTCCACTTTAACAACATTTTGCTCCACTTTTCCACGACCAAACAACTTGAAA
TCTGGTTAAAATAACACGCAACACTATTCTTCTTCCTTGAGTCCGCCCGGAACTCGAAAAACAAACCGAGTTAAAGCCATTTTTCACAA
AATCGATTTTGGGTCTCACCAAAATTACGGGGTTGCATACGCATTCGTTTATTTTCGAACGTGTACATACAAATATGCACAAAAATAAT
CATAATTATTTTCTGAGATGCATTATGATATGAACACCAATTTCGTATAGAGTCTCACTATGTCTCGAATTTTTGCTTACTGTCGGATA
TCAACGCTGGATCAGACCACCGAAAATCAACGCCGGGAAATCGAAAGTGCACTCGAGAGATCCTCTAGATTTAAGAAGGAGATATACAT
ATG
>ylaB
GAGGCAATTCCGACGTCTAAGAAACCATTATTATCATGACATTAACCTATAAAAATAGGCGTATCACGAGGCCCTTTCGTCTTCACCTC
GATGGATCCGCGCCGAATCAACATCTTAAGTTAGGGTTACATACCAGGCGTAAAGCTCTGCGCCTGGTCAAATGACAATGATCGTTTCC
ACCCATCACTTCATGAAATACCAGCTCTACCTCCTTATCTCCAGCCAGCCTTTTTCCACAATCAGATATACTTTCCCTACACTGTGTTA
ATAAGGATATGCTGGTGAGAACACGACATCTGGTCGGCCTTATTTCGGGAGTACTGATTCTTTCAGTATTGCTGCCTGTCGGCTTAAGC
ATCTGGCCTCGAGAGATCCTCTAGATTTAAGAAGGAGATATACATATG
>ylaC
GAGGCAATTCCGACGTCTAAGAAACCATTATTATCATGACATTAACCTATAAAAATAGGCGTATCACGAGGCCCTTTCGTCTTCACCTC
GATGGATCCGCGGTAATCCAGCCAGAATAATTAAAAAATTGTAATCGGTTTTTCGCAACTGTTATGCAAAATTGTGGTAGATCTGTTAC
TTCCCCTCTACTATTCCCACGTTAAAATAGGGTGTTCCCTGGAAAGTTGCAGATACCACGAAGGCAAACGATGACCGAAATACAACGCC
TGCTGACCGAAACGATCTCGAGAGATCCTCTAGATTTAAGAAGGAGATATACATATG
```

Data Set S4 - MG1655 library reference file.fasta

```
>purM
GAGGCAATTCCGACGTCTAAGAAACCATTATTATCATGACATTAACCTATAAAAATAGGCGTATCACGAGGCCCTTTCGTCTTCACCTC
GAGTTGCTCACGCATCAGTCCCAGCTTGTGTTTGACGAGTGGGTGTTTGACTTCCACGATCTTCATACTCTTTCTCCTTTGAGGGGCAG
CCACAAAAAAAATCGACGGATTATACCTCCTTTCTTCAAGGCGGCAATATTCTTTTCGTTGACTTTAGTCAAAATGATAACGGTTTGAG
ATAAAGTTATTTTATATTCAGATGGTTATGAAAGAAGATTATTCCATCCGAAAACTAACCTTTACCCTGGCACAAGTCTTCTTTCGCCG
CGCGCCTGGGGAAAAGACGTGCAAAAAGGTTGTGTAAAGCAGTCTCGCAAACGTTTGCTTTCCCTGTTAGAATTGCGCCGAATTTTATT
TTTCTACCGCAAGTAACGCGTGGGGACCCAAGCAGTGACCGATAAAACCTCTCTTAGCTACAAAGATGCCGGTGTTGATATTGACGCGG
GTAATGCTCTGGTTGGATCCTCTAGATTTAAGAAGGAGATATACATATG
>ydcL
GAGGCAATTCCGACGTCTAAGAAACCATTATTATCATGACATTAACCTATAAAAATAGGCGTATCACGAGGCCCTTTCGTCTTCACCTC
GAGAATGAAGACGTCGGCGAGCTGCACCGCACCGACGCCAACGGTAATCGTATTAAACTGCGTTTCGCCACGATGCTGGCACGTAAAAA
ATGACCCGGTAAGCACAAAACGCGTGAAAATTCCCCACGCTGAGATGATTTACTGTTCTTCTTTTCGGTAAGCATATTTTTTATCGAAG
GGATGTGAAATTAATCACAGTAGTCGAAGTTTTTAGCAGCTTAACTTACTGAAATTTAAGTACTGATGATTGACTTAGCCCCTTTTTCG
GCATTGACTATGTCGTCTGAAAAGGGGCTGAAAAATTTATTTTCACCAACACTTTTTTTGCCACAACACGAAGCGGCGCTTTTTGCTAT
AACTTAGAAAGTAATATAATCATCTCAGGAAACTATTCATGCGTACCACATCATTTGCGAAAGTTGCAGCTTTATGCGGCTTATTGGCT
CTGTCTGGTTGTGCATCTAAAATCACCCAGCCGGATCCTCTAGATTTAAGAAGGAGATATACATATG
>ydcM
GAGGCAATTCCGACGTCTAAGAAACCATTATTATCATGACATTAACCTATAAAAATAGGCGTATCACGAGGCCCTTTCGTCTTCACCTC
GAGGTACGCCAATGGCTTTCGAAAATATTAAACAAGTTATTGATGACATGGCGACCGATGCCACCATGTTTGACGTTAACAAAAAGTAG
TTCCAGACGCGCCATCGTTAGATGGCGCTTTTTATCCGGTGCGCCGTAAAACCCCATCCTTCAGGGCGGGGATATAAGGCGCGGTTTTC
CACCTGACCAGGTGTTTGCTGTTGTTCAATGTATTGGCGGATGACGGATATTGGCGCACCTCCGCAGCTACTGGCAAAATAGCCAGGAC
TCCACAAAACGCCTTTGTAGTAATACCTGAATGCAATATCTGGTCGATCTCGTCGCAGTCACCTACCCGATACGCCTTTGAGGCTGTTT
ACCAGACTGGATATCGCCAGTTTGGGAGGATAGTTAATTAACAAATGGACGTGATCTGGTTCGCCATCCATTTCAACCAGTTCAGCTTC
AAAATCAGCACATACATTTGAAAAGTAAGTGCGTAGTTTTTCTGTCGCGTCATGGTCAAAAATCTGGCGTCGGTATCTGGTGACAAAGA
GCAGGTGAACATGCATCAGGAAAACACAATGCCTTCCACGCCGGATATCGGTTTCTTTTTTCACAGACCAAAGTATCATTTGACCTGTG
AAACGATTACAGGCATTTAAATTCCAGTTAAGACCCGGTGGTCAACAGGAGTGTGAAATGAAGCGCTTCGGGATCCTCTAGATTTAAGA
AGGAGATATACATATG
>ydcJ
GAGGCAATTCCGACGTCTAAGAAACCATTATTATCATGACATTAACCTATAAAAATAGGCGTATCACGAGGCCCTTTCGTCTTCACCTC
GAGCAGCTGCTCCAGTTCATTCAATGTCTTAGAGAGCGCAGGTTGACTCAAATTAAGGGTTTCAGCCGCGCGCCCCAAAGTTCCTTGTT
GTGCGACAGCTACGAATGTATGAAGGTGGCGCAAACGGATGCGCTGACTAAACAGACTATTTTTTTCCATAAGCGATGTTAAAAACGAA
GCGGTGTCGCTGACAAGTGGGATCCTCTAGATTTAAGAAGGAGATATACATATG
>ydcK
GAGGCAATTCCGACGTCTAAGAAACCATTATTATCATGACATTAACCTATAAAAATAGGCGTATCACGAGGCCCTTTCGTCTTCACCTC
GATGGATCCTCCCTATCGTCCCCAACACAATACCAAAGTAGCCTGCCGGCAAATTGAGCACTTTATCGCTCTGCATCTTCTTACTCATT
TTGTTTAATTTATAAAGTATATTTGAAATGCATTTTATGGAATTTTCTCGAGCATAGCCAGAGCCGCAGAATTTGCTACGGTTATAGTA
AGTCACCTGTGGGAAAACGCTATGCGCAAATATCGTCTAAGTGAAGAACAGCGAGCCTTTAGTTATCAGGAAGATGGCACTAAAAAAAA
TGTGTTATTACGGCAGATTATCGCCAGTCGAGAGATCCTCTAGATTTAAGAAGGAGATATACATATG
>ydcI
GAGGCAATTCCGACGTCTAAGAAACCATTATTATCATGACATTAACCTATAAAAATAGGCGTATCACGAGGCCCTTTCGTCTTCACCTC
GATGGATCCCACTTGTCAGCGACACCGCTTCGTTTTTAACATCGCTTATGGAAAAAAATAGTCTGTTTAGTCAGCGCATCCGTTTGCGC
CACCTTCATACATTCGTAGCTGTCGCACAACAAGGAACTTTGGGGCGCGCGGCTGAAACCCTTAATTTGAGTCAACCTGCGCTCTCTAA
GACATTGAATGAACTGGAGCAGCTGCTCGAGAGATCCTCTAGATTTAAGAAGGAGATATACATATG
>ydcF
GAGGCAATTCCGACGTCTAAGAAACCATTATTATCATGACATTAACCTATAAAAATAGGCGTATCACGAGGCCCTTTCGTCTTCACCTC
GAGTCGCTCAACAACTTGGTACGCCTTATCCGATTTCAGATAAGCGTATTTTGCAGGCGATGGAGCAGATTAGCGGTTAACCCTGCTAT
TTGCCTGATAAAGAAAAAACCCGGTAAGCATTTAGCGCCGGGTTTTTTATTAATTCTAAAACGGTAAGGGTAAAAATTCAGGAATTCAG
AAAAATACAATTCTCTGCTGCAAGATGAATAATGTTTATCTACAGCATTTCCTTAAAAGATATGTCAGGCTTGCGGAGTGGCGGTTAAG
GACATACGATTTCCTCCTTTCAGAGTGCTCCGCTTCTCACTATTATCTCACGCAGTATTCTTAAGGGAACGATAAGGAGGAACCATGAA
CATTACCCCGTTTCCGACGCTTTCGCCGGCAACTATAGATGCCATAAATGTTATCGGACAGTGGCTGGCGCAGGATGATTTCTCCGGGA
TCCTCTAGATTTAAGAAGGAGATATACATATG
>ydcG
GAGGCAATTCCGACGTCTAAGAAACCATTATTATCATGACATTAACCTATAAAAATAGGCGTATCACGAGGCCCTTTCGTCTTCACCTC
GAGTTGGGTTGTCCGGTTTTGGATGAGTTCCAGCTTTACCAGGAAGCGGAAGAACGCAGTAAACGTCGCTGTGGTTTGCTTTAAAATCT
GACCATCCGCCTTTGCAAAAATTTGCCTGATTTTACAAACGAATCAGGCTCATCCCATCGACATAAAAAAAATGCCGATTTATGCATAT
TCTCTCAGTTCAACAATTGGATTATTAATAAATATTGTCTAGAGTGAGCGGTCATAAATAAGCACTTTCTTGCCGCTGAAAACGACCAG
CGCGGGACCATTCACAACACCAGAAGGACTCACTTTCAGGTATGGATCGTAGACGATTTATTAAAGGTTCAATGGCTATGGCCGCCGTG
TGCGGTACCAGCGGCATTGCTTCTCTTTTTTCTCAGGCGGCATTCGGATCCTCTAGATTTAAGAAGGAGATATACATATG
>ydcA
GAGGCAATTCCGACGTCTAAGAAACCATTATTATCATGACATTAACCTATAAAAATAGGCGTATCACGAGGCCCTTTCGTCTTCACCTC
GAGCTGCACTGGCACACCACTATTTCTGGAAGGACAACACACTTCTACGCATGATGCCGCGTAAACGTTCCTGAAGGATATTTAAAGAA
AACGCCTGTACTAAAACCGACCCGTGGTACAGGCGAAGAATACGGGTCTACATCGGAAGCGCCTATTATATTTATTTGTATGATAAATA
AAACCATAATCCTTGCCCATACGTCCATCTGGCTTATTTTTAATCAATTCACCCGATCTTTGATCTCATCAACGGTATCAAAATAAACC
```

```
AGCGTATAACGTTCAAATCGAACATAAGATTCGATAAACCATGGAGGTTATATGAAAAAACTGGCACTTATTTTGTTTATGGGAACGCT
TGTTTCCTTTTATGCCGATGCCGGGCGCAAACCCTGTTCTGGTGGATCCTCTAGATTTAAGAAGGAGATATACATATG
>fucR
GAGGCAATTCCGACGTCTAAGAAACCATTATTATCATGACATTAACCTATAAAAATAGGCGTATCACGAGGCCCTTTCGTCTTCACCTC
GAGTGCGTTTTATGAACGGGCGCAAAAAGCCTTTGCGATCGTTATCACAGGCGAACGAGCGAAGTACGGGAATATTCTTTTAAAAAAAG
GGGTAACACCGTAATCTCATACCGGTACGCCCGCATGACGCGGGCGGTTATCGAATGATGGGGTGAAAAATATGAAAGCGGCACGCCAG
CAAGCGATAGTCGACCTGCTGCTGAACCATACCAGCCTGACCACGGAAGCTGGATCCTCTAGATTTAAGAAGGAGATATACATATG
>ydcX
GAGGCAATTCCGACGTCTAAGAAACCATTATTATCATGACATTAACCTATAAAAATAGGCGTATCACGAGGCCCTTTCGTCTTCACCTC
GAGCACGGTGGGCAGAAACTTTCTGGTTACGGCAAGGATATGTCACTTTATGGGCTGGAGGATTACACCGTCGTCCGCCACGTCATGGT
TAAACATTAAGGATAATATTGCAGATCGTAAGAGTATGAGATGATCTTGCGTACTGTCAGAGCACTGATTTCTCTGACAGTACGTGCTG
CGTGCAAAATCACTACATCAAATAAACCAGCCAAATCTCATAGAGATAAGAGTAAACAAACCAACAGCAGCAAAGAAATTTATCAGTAC
TACAGTTCTACTGGAAACATTCATTTTTGCCACCTTTTATCATTGCCCCTATGTGTATAGTACGGGTTAAGAAAATCCGCAATAATACT
CATTCTGAAGATGGATAAGGGCAAGTTGCTGTTTGATGATTTTTTCTCAAACTGGTGAAAAAACTTGATGCACGTCAAAAAATGACGCA
TATTTGCGCGCGTTTTATTCATCTGGCTGGACGCGGATCCTCTAGATTTAAGAAGGAGATATACATATG
>fpr
GAGGCAATTCCGACGTCTAAGAAACCATTATTATCATGACATTAACCTATAAAAATAGGCGTATCACGAGGCCCTTTCGTCTTCACCTC
GATGGATCCTCTGGATCGCAAAGACCCGGAAATGCAGGTGCACATCCTCTGATTGATTTGATCGATTGAGCCTTCCAGTCCTTCGGGAC
TGGAATTTTTTTGTTCGGAGAACGAAGATAAGGCAAGTCAATCAAAACAGGAGAAAAACATGGCTGATTGGGTAACAGGCAAAGTCACT
AAAGTGCAGAACTGGACCGACGCCCTGTTTAGTCTCACCGTTCACGCCCTCGAGAGATCCTCTAGATTTAAGAAGGAGATATACATATG
>chiA
GAGGCAATTCCGACGTCTAAGAAACCATTATTATCATGACATTAACCTATAAAAATAGGCGTATCACGAGGCCCTTTCGTCTTCACCTC
GATGGATCCGTCTGCGTTTCGCAATCCGTGAAGGCGGCCGTACCGTTGGCGCGGGCGTTGTTGCTAAAGTTCTGGGCTAATTACACGTT
AATTAGTTTTGAATTGAAAAGGGCGCTTCGGCGCCCTTTTTGCATTTGTTGACGGCATATCTTATTCGTACTGTCAGAACGAATAAATT
ATTCATATTTAAAATATCAATTCAATTTAACATTTTAAATATGAATGTTTCTGTTTTTAAAATGCTTCATTTGTCAGGCTTTTTTCTTA
TTTTTAATGCCGTAATTTAATTCATATTGAAGGGTTCTCGTAAAACGTAAATAATTCCTGCGTAGGACTTTTGTTTTGCAGTTTTTACG
TCACAAGGGATTATAATGAAATTAAATATATTTACTAAATCTATGATTGGTATGGGGCTGGTGTGTTCCGCTCTGCCAGCATTGGCAAT
GGAAGCATGGAACTCGAGAGATCCTCTAGATTTAAGAAGGAGATATACATATG
>ydcR
GAGGCAATTCCGACGTCTAAGAAACCATTATTATCATGACATTAACCTATAAAAATAGGCGTATCACGAGGCCCTTTCGTCTTCACCTC
GAGTTGCATCATGCGACAAAAATCGACGCCGTCCAGCTCGCGGCAAAGGCGCTTGGCAAAGAGTTATCGCTGGTGATGGTTTAATTACA
GTTAACGAAAAGTTGTCATTTTTAACAACTGATATAGACTGCCGAATCATCTGCACATAATTACGATTCGATAATGAAAAAATACCAGC
AGCTTGCAGAACAATTACGCGAGCAGATTGCGTCGGGTATCTGGCAACCCGGCGATCGTTTGCCTTCGTTGCGGATCCTCTAGATTTAA
GAAGGAGATATACATATG
>ydcS
GAGGCAATTCCGACGTCTAAGAAACCATTATTATCATGACATTAACCTATAAAAATAGGCGTATCACGAGGCCCTTTCGTCTTCACCTC
GAGTAATACCGCGTGGCAGTGGGGAGAGCGTGAAGAACAGGCGGTAAAACAATTAGGCAAACTTATTCAAGAACGGCTGTAATAGCGTT
TAATTTAATTCCTCTTAGATTGGGTAATATGAATTTCGAATAGCAGTCATATTTCCTAACTCCTTGACTATACTCCAGAAGATAACCTT
ACAGACGGCATAATGCGCGGTAGCTCACAACCTGAATAAATTTTCTCAGGGGCGAAGGTGTGCCTGCAAGCCGCCGTCTATGGTTAAAC
AAGGAGATATTTTTACGGCACGGCGGCTGAACAATTAATTACGACAGGAGTAAGACCTTATGAGCAAGACATTTGCCCGCAGCAGCCTG
TGTGCGCTCAGCATGACAATAATGACCGCTCACGCCGCCGAACCGCCTACCAATTGGATCCTCTAGATTTAAGAAGGAGATATACATAT
G
>map
GAGGCAATTCCGACGTCTAAGAAACCATTATTATCATGACATTAACCTATAAAAATAGGCGTATCACGAGGCCCTTTCGTCTTCACCTC
GATGGATCCCACCGAAGATGAACGGCTTCATTTTCGGGTTCCAGTAACGGGTCTGGTGACCGAAGTGAACACCAGCCTTGAGCATGTCG
CGCATGGAAACAGTTGCCATGATTAAAACCTCTATATAAAAGTTGGGGTTATGCCTCCACGTATCCCATATTACCGACCCCAAAGGGCA
CCCCGGAATATGTGCCGATACGTGTGTGTTGTTACACAAAGTGAGATTGTCGCTTCCGTCCAGTCTGTGTATACGAAATGGATCGGAAG
TCCGGCGCGCTTTATACCACAAATACGTCGTGGACACCAATAATTGTTGGCGCTGTGTACAGCATCAGACGTCGAATTTTCTATTATAG
AAAACCTTCAGTGGCACGTTTGGCGAAATTCAGAATGATTCTCAATTTGCCCGGGTGTGATACCATTGACGGCACTTACATATATATTG
TCGGTATCACCGACGCTGATGGACAGAATTAATGGCTATCTCAATCAAGACCCCAGAAGATATCGAAAAAATGCGCGTCGCTGGCCGAC
TGGCTGCCGAAGTGCTGGAGATCTCGAGAGATCCTCTAGATTTAAGAAGGAGATATACATATG
>rrlG
GAGGCAATTCCGACGTCTAAGAAACCATTATTATCATGACATTAACCTATAAAAATAGGCGTATCACGAGGCCCTTTCGTCTTCACCTC
GATGGATCCCCCTTCGTCTAGAGGCCCAGGACACCGCCCTTTCACGGCGGTAACAGGGGTTCGAATCCCCTAGGGGACGCCACTTGCTG
GTTTGTGAGTGAAAGTCGCCGACCTCAATATCTCAAAACAGACTGTTAAGTCTTGTTTGATATATTTGCTCTTTAAAAATCTGGATCAA
GCTGAAAATTGAAACACTGAACAATGAAAGTTGTTCGTGAGTCTCTCAAATTTTCGCAACTCTGAAGTGAAACATCTTCGGGTTGTGAG
GTTAAGCGACTAAGCGTACACGGTGGATGCCCTGGCAGTCAGAGGCGATGAAGGACGTGCTAATCTGCGATAAGCGTCGGTCTCGAGAG
ATCCTCTAGATTTAAGAAGGAGATATACATATG
>cdaR
GAGGCAATTCCGACGTCTAAGAAACCATTATTATCATGACATTAACCTATAAAAATAGGCGTATCACGAGGCCCTTTCGTCTTCACCTC
GAGTTCTCGACAGCAAACCGTCTGTGCTGGCACTCAACATTCAGCGCGGCGACAGCACCATCTACCTGTTAATGCAGTAATCTCCCTCA
ACCCCTTCCTGAAAACGGGAAGGGGTTCTCCTTACAATCTGTGAACTTCACCACAACTCCATACATCTTCATCATCCTTTAGGCATTTG
```

```
CACAATGCCGTACGTTACGTACTTCCTTATGCTAAGCCGTGCATAACGGAGGACTTATGGCTGGCTGGCATCTTGATGGATCCTCTAGA
TTTAAGAAGGAGATATACATATG
>cpdA
GAGGCAATTCCGACGTCTAAGAAACCATTATTATCATGACATTAACCTATAAAAATAGGCGTATCACGAGGCCCTTTCGTCTTCACCTC
GATGGATCCGCGGACTGGTTGCGATACTGTTTAGCACATGGAGCGATGGCGATTCCGGTTTATTAGCGTCGTGAAACCTAAGGACACCA
TTTGGAAAGCCTGTTAACCCTTCCTCTGGCTGGTGAGGCCAGAGTCAGGATTTTACAAATTACCGACACTCACCTGTTTGCACAAAAGC
ACGAAGCCCCTCGAGAGATCCTCTAGATTTAAGAAGGAGATATACATATG
>cpdB
GAGGCAATTCCGACGTCTAAGAAACCATTATTATCATGACATTAACCTATAAAAATAGGCGTATCACGAGGCCCTTTCGTCTTCACCTC
GATGGATCCTTCGTCCCGTCGTAGACCTGCATAATGGCATCGCCTGCATTCCGTGCAAGCTGGCATACTTGATCTAACATTTCTCCACC
TCGTCTCTGTGAGCGGTGTTAACTTATTGTTTTACTTATACCCTATCGTTAATGAATGCGCCAACTGTGATAGTGTCATCATTTTCAAA
GCGTAAAATTGTGGCATTCTTCACTGTTCTATAAGTAAGACGTTTATTCTCCCTTTTCTTTCGTATTCCCGATGATAAAAGGATGTCCC
TGATGATTAAGTTTAGCGCAACGCTCCTGGCCACGCTGATTGCCGCCAGTGTGAATGCAGCGACGGTCGATCTGCGTATCACTCGAGAG
ATCCTCTAGATTTAAGAAGGAGATATACATATG
>talA
GAGGCAATTCCGACGTCTAAGAAACCATTATTATCATGACATTAACCTATAAAAATAGGCGTATCACGAGGCCCTTTCGTCTTCACCTC
GAGGCTGTGTTGCCAGAGGCTTGGTTGGAGAAACCTGGATTTTCCCTGGAACTGGAAATTCATGGAAATCAAGTGCACTTTGTTTTAAC
TGGTCATCCATTTGGTTGTTCCTTTCACGTAACGTTCACAAATAAAGTGTGTGGGCAACAGCCCCTGCCCACAACGTGGCGCACATTAT
TACCCTGCCGGAGTCTACAGACTTTGAGCAAGTCCAAACTCTCACCATTAATATAATGTTTTGGTAATAATCCTATAACACTGATGTTA
CCTGCTTAATCCAGCAATACCATGCCTGTCTGCTATGCTTTTTTGATGCGTTTAGCGAAATTTCTCAGAAGTGTGAATTAACGCACTCA
TCTAACACTTTACTTTTCAAGGAGTATTTCCTATGAACGAGTTAGACGGCATCAAACAGTTCACCACTGTCGTGGCAGACAGCGGCGAT
ATTGGATCCTCTAGATTTAAGAAGGAGATATACATATG
>talB
GAGGCAATTCCGACGTCTAAGAAACCATTATTATCATGACATTAACCTATAAAAATAGGCGTATCACGAGGCCCTTTCGTCTTCACCTC
GAGCGCCGAAGAGCAGGTAAATCATTACCGATCCCCAAAGGACGCTGTTAATGAAGGAGAAAAAATCTGGCATGCATATCCCTCTTATT
GCCGGTCGCGATGACTTTCCTGTGTAAACGTTACCAATTGTTTAAGAAGTATATACGCTACGAGGTACTTGATAACTTCTGCGTAGCAT
ACATGAGGTTTTGTATAAAAATGGCGGGCGATATCAACGCAGTGTCAGAAATCCGAAACAGTCTCGCCTGGCGATAACCGTCTTGTCGG
CGGTTGCGCTGACGTTGCGTCGTGATATCATCAGGGCAGACCGGTTACATCCCCCTAACAAGCTGTTTAAAGAGAAATACTATCATGAC
GGACAAATTGACCTCCCTTCGTCAGTACACCACCGTAGTGGCCGACACTGGGGACATCGCGGCAATGAAGCTGTATCAACCGCAGGATG
CCGGATCCTCTAGATTTAAGAAGGAGATATACATATG
>yncA
GAGGCAATTCCGACGTCTAAGAAACCATTATTATCATGACATTAACCTATAAAAATAGGCGTATCACGAGGCCCTTTCGTCTTCACCTC
GATGGATCCTTTTGTTTGCAGCCGTCACATCCCCTCTGCAACACGCAATAATTTCATCGAGGAATCCGTTGCTTCGGGCTGTTTTCGAT
AGTATTGATAACATCATCATCATTATTTTATTGAGGCCAGACATGTCCATCCGTTTTGCCCGCAAAGCCGACTGTGCTGCCATTGCGGA
AATTTATAACCACGCCGTGTTGTATACGGCGGCTATCTGGAATGACTCGAGAGATCCTCTAGATTTAAGAAGGAGATATACATATG
>yncB
GAGGCAATTCCGACGTCTAAGAAACCATTATTATCATGACATTAACCTATAAAAATAGGCGTATCACGAGGCCCTTTCGTCTTCACCTC
GAGTCATTCCAGATAGCCGCCGTATACAACACGGCGTGGTTATAAATTTCCGCAATGGCAGCACAGTCGGCTTTGCGGGCAAAACGGAT
GGACATGTCTGGCCTCAATAAAATAATGATGATGATGTTATCAATACTATCGAAAACAGCCCGAAGCAACGGATTCCTCGATGAAATTA
TTGCGTGTTGCAGAGGGGATGTGACGGCTGCAAACAAAACCAATCCCGTGAGCCTGCTACTATTGATGGATTCATGCCAAAAGCCGGAG
AGCATTATGGGGCAGATCCCGGGATCCTCTAGATTTAAGAAGGAGATATACATATG
>yncC
GAGGCAATTCCGACGTCTAAGAAACCATTATTATCATGACATTAACCTATAAAAATAGGCGTATCACGAGGCCCTTTCGTCTTCACCTC
GAGACGTTTATCGGCCTGCTGAAGGGTAAAAACTTCGGCAAAGTGGTGATCCGCGTGGCGGGTGATGATTAAAGTTTCAACGGCGGCGT
AAGCCGCCATAAAAATCCAGGCAAAAAGCTTATTTTCAGCTTTAATTAACTAACCACATCATTGTTCTGCTGCTAATAACAACAAAGGT
GAGTGGTTTATGATTATTCACTTTAATACACCAGGTGAATTCCTTCTGCCATGCAGGCAGGGTTGGACAGAAAACACGCGAAGGAATTA
AAATGCCTGGAACGGGAAAAATGGATCCTCTAGATTTAAGAAGGAGATATACATATG
>yncD
GAGGCAATTCCGACGTCTAAGAAACCATTATTATCATGACATTAACCTATAAAAATAGGCGTATCACGAGGCCCTTTCGTCTTCACCTC
GATGGATCCGGCGCGATGAAAACAGATGACGTAAATGCATGACGACTCCCTTTGATGAACCGATATTTTTGCTGCTTCACAATGCTTCG
CTAAGCGTAATCATCGTTACGCTCTTGTTATAAATGGGAATGGTAATCATTATTTTCTCAAATGTGGAGCATTTCTTGCCGTTACGACT
ACAATTTTCTCAATTACTTAACGTATTGTGCCGTTTCATTTTCCATAAATTTACAATTTCATTACCATGACGGCGTTTTCCCATTCTTA
ACCTGGCTTTTTTCATGAAGATTTTTTCCGTCCGACAGACCGTTTTGCCCGCACTGCTTGTCCTTTCCCCCGTTGTTTTTGCCGCTGAT
GAACAGACTCGAGAGATCCTCTAGATTTAAGAAGGAGATATACATATG
>yncE
GAGGCAATTCCGACGTCTAAGAAACCATTATTATCATGACATTAACCTATAAAAATAGGCGTATCACGAGGCCCTTTCGTCTTCACCTC
GAGTCATCAGCGGCAAAAACAACGGGGGAAAGGACAAGCAGTGCGGGCAAAACGGTCTGTCGGACGGAAAAAATCTTCATGAAAAAAGC
CAGGTTAAGAATGGGAAAACGCCGTCATGGTAATGAAATTGTAAATTTATGGAAAATGAAACGGCACAATACGTTAAGTAATTGAGAAA
ATTGTAGTCGTAACGGCAAGAAATGCTCCACATTTGAGAAAATAATGATTACCATTCCCATTTATAACAAGAGCGTAACGATGATTACG
CTTAGCGAAGCATTGTGAAGCAGCAAAAATATCGGTTCATCAAAGGGAGTCGTCATGCATTTACGTCATCTGTTTTCATCGCGCCTGCG
TGGTTCATTACTGTTAGGTTCATTGCTTGTTGTTTCATCATTCAGTACGCAGGCCGGGATCCTCTAGATTTAAGAAGGAGATATACATA
TG
>uvrY
```

GAGGCAATTCCGACGTCTAAGAAACCATTATTATCATGACATTAACCTATAAAAATAGGCGTATCACGAGGCCCTTTCGTCTTCACCTC
GATGGATCCAGCATCGCTTTCAGGCAGGAGACTTCATTTGCCAGTTCTTGATTATTCTCGGCAGTAGAAAAATCAGGCGTACTCATTTA
TTTTCCTCGTCATGTTGCAATGAAAATTTGCGGTGAAAAATGTTAAGGCGGCGGAGTATACCATAAGCTTTGCTAAAAATAGCAGTGGT
TGTTTTTTGAGCGTGATATCGGCAGTGCTATAAAATACGTTAATATAATGACCAATAAATATTTTTATCATGAATGTTTTTTGCCCGTA
TTGTTCGGGTTAATTAATGTTACATATTCAGCGGGCTGATTTTCATTTTTGCTGAATAAAGTCAATTTTTGTCACATTTCATCGTAGGG
CTTACTGTGAAACGATCCGGTAAGCCGTTGGTGACGGGCGTGACCATAACTGTGGACAATCGAATTGACAAAAACGAGAGAAAAATCGA
ATACCCACCATTTTTAACGTTTCAAAGTTGCAATAAAAACCGCTAATATACGAATGACTAACTATCAGTAGCGTTATCCCTATTTCTGG
AGATATTCCTTTGATCAACGTTCTACTTGTTGATGACCACGAACTGGTGCGCGCAGGGATACGACGCATTCTGGAAGATATAAAGGGTA
TAAAAGTCGTCGGTGAGGCATCCTCGAGAGATCCTCTAGATTTAAGAAGGAGATATACATATG
>yncG
GAGGCAATTCCGACGTCTAAGAAACCATTATTATCATGACATTAACCTATAAAAATAGGCGTATCACGAGGCCCTTTCGTCTTCACCTC
GAGCGGCGTGTTGATCTGAAGTGTCGGTGTCGTGTTTACTCATTGCTCTCCCTGATTGCTTTAATGAAAAAGTCATATAAGTTGCCATG
AACAATGTTTATTCATCCTGGCGATGATAGTTATCACCATTATTCTTTATGGGTATAAAACTATCGGCGCAGCATCCTACCCGTTATCG
ATAAACGATGCAAAACATCCCCTTACAATCCTGAAGGGGATTAATACAACTGACGAAAAAATGACAAATCCTTTTGCTGGTTAACCTGT
GTACTGTCCTACACTTAATCTTTAAAAGATTGTGAGGGGCATATGATTAAGGTGTATGGCGTACCCGGATGGGGCTCTACAATCAGTGA
GCTGATGCTAACACTGGCTGATATTCCTTATCAGTTCGTTGATGTCAGTGGGGATCCTCTAGATTTAAGAAGGAGATATACATATG
>yncJ
GAGGCAATTCCGACGTCTAAGAAACCATTATTATCATGACATTAACCTATAAAAATAGGCGTATCACGAGGCCCTTTCGTCTTCACCTC
GATGGATCCTTGCTGCCATTCGCTACATCGACGCCCTGAGATTCGAGCCAACGTCTGAACTCGCTTTGTTTCACATCGCCTCCCTGCCG
CTGAGTATATCAACAAAGTAAACATTAGTGTTTACCTCCGTGTGTAATTTGCTCGAAAATAGCACTAAATCGCCGCAACGGCAGAATGC
AATGCAAAATCAGGAAGGAAGATTCCAGAATTTCTTTGCGCTAATTTGCAAAACTGAATAATAACCTCAAGAAACGATCGGCATCATAG
CGGCAAAAGGTAAGAAAAAGGTGGCAGGATGTTTACGAAGGCGTTATCGGTTGTCTTATTAACGTGTGCTCTGTTTTCAGGACAACTCA
TGGCAGGGCTCGAGAGATCCTCTAGATTTAAGAAGGAGATATACATATG
>pncB
GAGGCAATTCCGACGTCTAAGAAACCATTATTATCATGACATTAACCTATAAAAATAGGCGTATCACGAGGCCCTTTCGTCTTCACCTC
GATGGATCCTTGGCTTGTGGCTGTTGAGTCATAAATAACCTTTAGCATCTTTTATAGAGTCTGGTGTTCAGTCTATTCCTGTTGCGTAA
ATCGCGCTATGCAGAATCTTCATCTTTTCAGGTACAAACGCCTTTATTGCTACATTTTTATAACATACACCGCGTAATGCCATCGACCA
GAAAGGTGGCATATGGTGTGATCGGGGTTCAATAAATTGCGAAACAAGGTATACTCCAGCAGTTCCTGAAGATGTTTATTGTACTAAAC
GCTCCTGTACGAGGACGCTACTGCGCACCTATGACACAATTCGCTTCTCCTGTTCTGCACTCGTTGCTGGATACAGATGCTTATAAGTT
GCATATGCAGCAAGCCGTGTTTCTCGAGAGATCCTCTAGATTTAAGAAGGAGATATACATATG
>yfaY
GAGGCAATTCCGACGTCTAAGAAACCATTATTATCATGACATTAACCTATAAAAATAGGCGTATCACGAGGCCCTTTCGTCTTCACCTC
GATGGATCCGGCGGGTTATCGCTACCTGAATCTGTCGGGTAAAGACGGTAACCGCGACAACGCCGTGGCTGACGGCCCGTATGTTGGGG
TTAACGCCAGTTTCTGATCCCTTCCGGCGCGGCATTCTGTCGCGCCATTCAGTTTCTTACTACGCCTTATCTCCCTTTGCGTTATAGTG
ATTCGATAACAAAAGCAGGAGATAACCATGTTAAAAGTGGAAATGTTATCCACCGGGGATGAAGTGTTACACGGGCAAATCGTTGACAC
TAACGCTGCCTGGCCTCGAGAGATCCTCTAGATTTAAGAAGGAGATATACATATG
>torT
GAGGCAATTCCGACGTCTAAGAAACCATTATTATCATGACATTAACCTATAAAAATAGGCGTATCACGAGGCCCTTTCGTCTTCACCTC
GAGGGCTGATAAAGCGCAGGTTGTACCATCCCACCAGGGTACTGGTCAGGGTTAACAGCGCCATCAGGGCAAAGCCCATCCAGAGTCTT
CGGGTCAGGGTTAAATTCACGGTCGGTGCACTTTAGGTGAAAAAGGTTGAGTCGCAAAGCGGAATGCATCTAGCATAAAGCCTTATTAT
TGATGAGGCTATCATGCGCGTACTGCTATTTTTACTTCTTTCCCTTTTCATGTTGCCGGCATTTTCGGCTGATAACCTGTTGCGCTGGC
GGATCCTCTAGATTTAAGAAGGAGATATACATATG
>yfaD
GAGGCAATTCCGACGTCTAAGAAACCATTATTATCATGACATTAACCTATAAAAATAGGCGTATCACGAGGCCCTTTCGTCTTCACCTC
GAGGTCACCGACTGCGAAACCTGTAAATGGCAGATTGAGATGTCCACAAGTCTTCGCTGCGAACATCCGATTACGCTACTGGCCCAGGC
GCTGGCTTAAACTCCTTTCTGATGCCCGGTAAGCATGTGGTTACCGGGCATTTTTGCGTACACGATTCCGTGCCCAATGTATGCGTTGC
AACGCAGTGAAAATTCCTCTGAAAACGTCTCGCAAAGGCTGAAACTGGCAGATGTCAAAGGCCTGGGATAACCGTAATGTCGCGTCATC
ATAAATATCAGGTGACGGACAACCATGACCGAATCAACAACCTCCTCCCCGCATGATGCGGTATTTAAAACCTTTATGTTCACACCCGA
AACCGCACGGGATTTTCTCGAAGGATCCTCTAGATTTAAGAAGGAGATATACATATG
>torS
GAGGCAATTCCGACGTCTAAGAAACCATTATTATCATGACATTAACCTATAAAAATAGGCGTATCACGAGGCCCTTTCGTCTTCACCTC
GATGGATCCGCCAGCGCAACAGGTTATCAGCCGAAAATGCCGGCAACATGAAAAGGGAAAGAAGTAAAAATAGCAGTACGCGCATGATA
GCCTCATCAATAATAAGGCTTTATGCTAGATGCATTCCGCTTTGCGACTCAACCTTTTTCACCTAAAGTGCACCGACCGTGAATTTAAC
CCTGACCCGAAGACTCTGGATGGGCTTTGCCCTGATGGCGCTGTTAACCCTGACCAGTACCCTGGTGGGATGGTACAACCTGCGCTTTA
TCAGCCCTCGAGAGATCCTCTAGATTTAAGAAGGAGATATACATATG
>torR
GAGGCAATTCCGACGTCTAAGAAACCATTATTATCATGACATTAACCTATAAAAATAGGCGTATCACGAGGCCCTTTCGTCTTCACCTC
GATGGATCCCGTAGCGCGTTCCAGAGTTTCCGCATAATAACCCCTGTAGAATTATGGTTTAGTGAAGCGATCTTAATGAGCAAATATGA
ACAGCGGCACTGGTCAGGATGAACGGCTTACGGCAGAATATGAACAGATATGAACAGAATGAGTAAAACCCTCTGATGCCACATCACAT
TGTTATTGTTGAAGATGAGCCGGTTACCCAGGCGCGATTACAATCCTACTTCACTCAGGAGGGGTATACCGTTTCCGTTACGGCCTCGA
GAGATCCTCTAGATTTAAGAAGGAGATATACATATG
>yfaL

Data Set S4 - MG1655 library reference file.fasta
GAGGCAATTCCGACGTCTAAGAAACCATTATTATCATGACATTAACCTATAAAAATAGGCGTATCACGAGGCCCTTTCGTCTTCACCTC
GATGGATCCTCCAGAACGCGATGGATTTTGTCGAGATTGATGCGCTCTGTGCTACCGTCGCGCTTTGTCACCAGCAGATTCTGATTCAT
GTATGTCGTACCTGTTTTTGGAATTCTTTCCGCAATAGTTTCATGCAGACTACCTGTAGTACGGGAAGCTGCTTAGAACACTATATATA
GGGGGTATGTTTGGGATTCACTGCAAGATAGTGTGAAAATGACCCTCTTGCAAGTGCATAACTTTGTGGATAACTCAGGAAGGAAAAAG
TGGCTTTCGCGCACCTTAGGTCAGACAAGGTGTCCGGGAAAGTCAATGGGAAGAGAAAAATTTGTTAAAAATAACTGTTCGGCGATTTC
TTGAACGGCATAATAGGTTAGATAAATTGATATAGATGGCTTATTAATCTTTTTGCTAATGATTGCAAAAATTGATCTAAGCACGAGTC
ATCACATAATCTGATGTATCAGTTGTGGTCAATATATAAGCGAATGTAAGAGATTGTTGATGGCGAAATGGTTGTTTTCTATTCGGTTA
TTGTTAAATGCTAAGCCATTTTGTTAATTTACCTTATATTATAAGAAAATTATTCATGACTTTAATTTAGTTTAAATATAACTGATATC
CCTGAAGGTTATTAACTTGAATTATGACGCACTGATATTATTCATCAAATAATAACAAAATAGCCATTGCACCGGGTTGAACAATTTAC
AAAAGAAAGATATTCCATATCGTATAATGCGATTAAATACGCCGTCTTATAGAAAATGATGGTTAGATTAATATGCGGATTATCTTTCT
ACGCAAGGAGTATTTATCTTTACTCCCGTCAATGATTGCATCTCTTTTCTCTGCTAACGGTGTCGCGGCGGTCACTGATTCATCTCGAG
AGATCCTCTAGATTTAAGAAGGAGATATACATATG
>ygiL
GAGGCAATTCCGACGTCTAAGAAACCATTATTATCATGACATTAACCTATAAAAATAGGCGTATCACGAGGCCCTTTCGTCTTCACCTC
GAGAAAAACTGGCGCTGCTTGAACAAAGAATCAGCGAACTGGAAAATCGCAGTACTGAGATCAAAAAGCAGCCAGATCCAGAGACTCTC
CCCCCAACGCTGTAAACATTCGTCTTTAATGCACTGTAAATTATGAATTAATCCTCTAATTTATAGTGCTTTCTATTACCTTATCGCGT
AATCCCATTCAGAAAACCATCACAAAAATGAATGAGTACAATTGAAAGTATAATTGCAATGTATTTTTTAAGTTTTACTTAATCGATAT
TTAGATAAACTAATTTTATAAAAAACACAAAATAAAATATCATATTTCAATGATTAGTAGTCCTTAAATAACAGTTACTAATCATTCTC
TTTTCTACTTATCTTTATATAAAGGAATATTATGTCCGCTTTTAAAAAGTCACTTCTTGTTGCTGGCGTAGCAATGATTCTTTCCAATA
ACGTGTTTGCTGATGAAGGTGGATCCTCTAGATTTAAGAAGGAGATATACATATG
>torY
GAGGCAATTCCGACGTCTAAGAAACCATTATTATCATGACATTAACCTATAAAAATAGGCGTATCACGAGGCCCTTTCGTCTTCACCTC
GATGGATCCTGAACACGCGGACGAGTATTCGCGTTATATCGTAGACGGGGCGGCGGTTGCTGAAATGAAAGGAATCATTGAACGCCATC
AGGCCAAATGATTTTTACCGTTGCATCATGTCGCCCAATATGATGCTTGCTCGTACCAGGCCCCTGCAATTTCAACAGGGGCCTTTTTT
TATCCCTGAACAGTATAAAAAACGAACGATAACCGTGATCTGTTGAGCGGGTGACAGTGCGCATAGCGTTGTGCTAAAAATATTGTATA
TATTCACATTAATTATGGGATTTAAATTACTAAAACTGATAAATATATATTCTAAATAGCAACTGGGTTATTCCTTAGCAATTAATGAT
TACATTGTAATAAATCATATTCTTTATCGATTGTTTCAGGCAGTGTTGTGCCTAATTATGCAAGCGGTTAATTCGTTGTATATTTAATT
ATACAATGATTTCGGTGTCCAGTAATTTAATTAGAGGAATCTATGCGAGGGAAAAAACGCATTGGGTTATTGTTTTTGCTGATAGCGGT
TGTGGTTGGTGGCGGCGGGTTATTGCTGGCGCAAAAAGTCTCGAGAGATCCTCTAGATTTAAGAAGGAGATATACATATG
>uvrA
GAGGCAATTCCGACGTCTAAGAAACCATTATTATCATGACATTAACCTATAAAAATAGGCGTATCACGAGGCCCTTTCGTCTTCACCTC
GATGGATCCAGCGTACTTCCGGGTCCTGACCCAGATTACCAACGAGAATAACCTTGTTTACGCCTCTGCTGGCCATGTTCGTGTCTCCT
GAAAAAAATCGTTCTGAATAAGTGTAAACGCGCGATTGTACCATTACCAATAGCGCTTTTACTATGTTGTGACCTCGGTTCCGGGAAAC
AAACCTGGCCAGACATTGTTACACAACACTCCGGGTAATGCATTCCAATACTGTATATTCATTCAGGTCAATTTGTGTCATAATTAACC
GTTTGTGATCGCCGGTAGCACCATGCCACCGGGCAAAAAAGCGTTTAATCCGGGAAAGGTGAATGGATAAGATCGAAGTTCGGGGCGCC
CGCACCCATAATCTCAAAAACATCAACCTCGTTATCCCCCGCGACAAGCTCATTGTCTCGAGAGATCCTCTAGATTTAAGAAGGAGATA
TACATATG
>ycaO
GAGGCAATTCCGACGTCTAAGAAACCATTATTATCATGACATTAACCTATAAAAATAGGCGTATCACGAGGCCCTTTCGTCTTCACCTC
GATGGATCCGCCATACGGTGGAGATCGAACGCCTTCATTTTTATGCTCTTAACGTGGTTTATCCGCTAAAAGCATAGTCAACAGTTCAT
CGGCATGGCGGCGGTGATAGCGCCTTTGAGATAAAACCCGTATTATTGCGCGCTTTCCGTACGACTAAAGTGATTTTCATGACGCAAAC
ATTTATCCCCGGCAAAGATGCCGCTCTGGAAGATTCCATCGCTCGCTTCCAGCAAAAACTCGAGAGATCCTCTAGATTTAAGAAGGAGA
TATACATATG
>ycaN
GAGGCAATTCCGACGTCTAAGAAACCATTATTATCATGACATTAACCTATAAAAATAGGCGTATCACGAGGCCCTTTCGTCTTCACCTC
GATGGATCCATCATGACGAGGATGGGCCCATACCAAATAAATACGTTCAGACTGCATAATTTTACCTCCACGTTAAGTTCTGGAGCGCA
CTTTACGCAGGCGGCATAATCCTGTAAACGTGATGATTCAGGATGAGTTAATGAAGGATTTTCACTAATGCGGATGAATATGTCTGACT
TTGCCACTTTCTTTGCCGTGGCCCGTAATCAAAGCTTTCGTGCAGCGGGCGATGAGTTAGGCTCTCGAGAGATCCTCTAGATTTAAGAA
GGAGATATACATATG
>ycaL
GAGGCAATTCCGACGTCTAAGAAACCATTATTATCATGACATTAACCTATAAAAATAGGCGTATCACGAGGCCCTTTCGTCTTCACCTC
GAGTGATCCCAAATGCACGGCCAAAACATTTCCGGATTATTTCGAGCAGCTGGCGCGGATTAGCCAGGCAGCCTGAATGAACAACGGGC
AATAAATAGCCAAATCTTTCTTTATCAAAACGTCGGCACATTGTCGGCGTTTTTTTTCGGACCTTGTGAGTCATTTTGATTAATGGTAG
CGTCGCTTGTCAATGTAAGTTGTTGATACATAATATTTATATATGATTAATCAACGGATGATTCACATGAAGAATACTAAATTACTGCT
GGCGATTGCGACCTCTGCAGCGGATCCTCTAGATTTAAGAAGGAGATATACATATG
>ycaK
GAGGCAATTCCGACGTCTAAGAAACCATTATTATCATGACATTAACCTATAAAAATAGGCGTATCACGAGGCCCTTTCGTCTTCACCTC
GAGCCTAACTCATCGCCCGCTGCACGAAAGCTTTGATTACGGGCCACGGCAAAGAAAGTGGCAAAGTCAGACATATTCATCCGCATTAG
TGAAAATCCTTCATTAACTCATCCTGAATCATCACGTTTACAGGATTATGCCGCCTGCGTAAAGTGCGCTCCAGAACTTAACGTGGAGG
TAAAATTATGCAGTCTGAACGTATTTATTTGGTATGGGCCCATCCTCGTCATGATTCATTGACCGCACATATTGCTGATGCGATCCATC
GGATCCTCTAGATTTAAGAAGGAGATATACATATG
>cutA

GAGGCAATTCCGACGTCTAAGAAACCATTATTATCATGACATTAACCTATAAAAATAGGCGTATCACGAGGCCCTTTCGTCTTCACCTC
GATGGATCCCATCCGTTCTTCATCCCGGGTACTCTGGGTGTTGCCCTGGCCGTTTGCTTCGGCTTCGTGCTGGGTAGCTTCATGCTGTA
ATGATCAATCGCGGGGCGTTCACGCCCCGCTTTCTTTCCCGCCGACTAACATCCTTCCCCCGTCCGTTGTATAGTGACCTCTCTCTTGC
GGTTCCATCTGTTCTTGCGAGGTGTTTATGCTTGATGAAAAAAGTTCGAATACCGCGTCTGTCGTGGTGCTATGTACGGCACCAGATGA
AGCGACAGCCCAGGATTTAGCCGCCAAACTCGAGAGATCCTCTAGATTTAAGAAGGAGATATACATATG
>ycaI
GAGGCAATTCCGACGTCTAAGAAACCATTATTATCATGACATTAACCTATAAAAATAGGCGTATCACGAGGCCCTTTCGTCTTCACCTC
GAGTCCGAAGACTGGCGATAAAGTAGAACTGGAAGGAAAATACGTTCCTCACTTTAAACCTGGTAAAGAACTGCGCGATCGCGCCAATA
TTTACGGTTAAGTTTTTTACTCAAACTTGAACGAGAGAAAAGCACCTGTCGGGTGCTTTTTTCATTTCTCTAATCTGGAACTGGAAGCT
GCCTCGCAGAGTTTTGAACAGTTTTCACCCTTTCGTTAAATTCTTCTGAATATGCCTCGGGGAACGCAAAATTCCCACCAGACAACCGC
TCAACAAAGTTGCACACTTTCCATAAACAGAGAGGGGTGCAGGATCCTCTAGATTTAAGAAGGAGATATACATATG
>ycaD
GAGGCAATTCCGACGTCTAAGAAACCATTATTATCATGACATTAACCTATAAAAATAGGCGTATCACGAGGCCCTTTCGTCTTCACCTC
GAGCGGGTTCGATATCCCGTACAAGGGAAAGTAAACCAGCCTGGTGATCAACAAGCAAAACGGCAGCATCATTTTTATCAAGACGAACA
TACGGTTTGGTCATCCTCTTCTCCTTTCGAGAGTCAGTGGTACGTCTGAAATCGATCAGACGCGATGCATTGCTCTGAAAGCATAGACG
GGAAATATGAGTTTGCTGTGACCATGAAATTTTTCGACTGAACGAAGCGTATGAAATTTGTGTTAGTTCAATAAAAACAATCAGATGAG
ACTAATCATCTCTTCAGCATTAGCGTTATGAATGAGTAATATAAATCGATAATACCGCCGCTGGGATATCGCGTATTTTCACCCATTGA
CAATGTTTTTGGCGGTGGCATGATGCGCATGAAATTTGAACTTCCTCACGGTTTTAATTCATGTCCACGTATACCCAGCCTGTCATGCT
TTTGCTGTCTGGCCTGCTTTTGTTGACTCTGGCGATTACGGTGTTAAATACACTCGTGCCGCTTTGGGATCCTCTAGATTTAAGAAGGA
GATATACATATG
>ycaC
GAGGCAATTCCGACGTCTAAGAAACCATTATTATCATGACATTAACCTATAAAAATAGGCGTATCACGAGGCCCTTTCGTCTTCACCTC
GATGGATCCAAAGCATGACAGGCTGGGTATACGTGGACATGAATTAAAACCGTGAGGAAGTTCAAATTTCATGCGCATCATGCCACCGC
CAAAAACATTGTCAATGGGTGAAAATACGCGATATCCCAGCGGCGGTATTATCGATTTATATTACTCATTCATAACGCTAATGCTGAAG
AGATGATTAGTCTCATCTGATTGTTTTTATTGAACTAACACAAATTTCATACGCTTCGTTCAGTCGAAAAATTTCATGGTCACAGCAAA
CTCATATTTCCCGTCTATGCTTTCAGAGCAATGCATCGCGTCTGATCGATTTCAGACGTACCACTGACTCTCGAAAGGAGAAGAGGATG
ACCAAACCGTATGTTCGTCTTGATAAAAATGATGCTGCCGTTTTGCTTGTTGATCACCAGGCTGGTTTACTTTCCCTTGTACGGGATAT
CGAACCCGCTCGAGAGATCCTCTAGATTTAAGAAGGAGATATACATATG
>cybC
GAGGCAATTCCGACGTCTAAGAAACCATTATTATCATGACATTAACCTATAAAAATAGGCGTATCACGAGGCCCTTTCGTCTTCACCTC
GAGGCGCAATATTGTCACCGTCGGTAACGATATTGAAACACGCAGTAATATACAGTGTGGTTCTGTGCTGTTGCCGGAGATGAAAATCG
CCGGACAGTAAAAAAACTGGCGCGACCTGCCGCGCTATAAACTAAAAAATTACACAAAATCATTCGCACTGCATCGAGACGGAAAGTGA
ATGAGGGCCGGGGAGCGTACTCGCAGTACGTGACCCGGTCGAATGAGCGTAGTCGTCGATGAGGCAGTGTGAAGGATGAAGTGTAAATA
AAAAAGGAACTATTCTTGCAGTCTCCTCGTTGGTATTCAGTTCGGCGTCATTTGCTGCTGATCTCGAAGACAATATGGAAACCCTCAAC
GACAATTTAAAAGTGGTCGAAAAAGCCGATAACGCGGCGCAAGTCAAAGACGCGTGGATCCTCTAGATTTAAGAAGGAGATATACATAT
G
>tktB
GAGGCAATTCCGACGTCTAAGAAACCATTATTATCATGACATTAACCTATAAAAATAGGCGTATCACGAGGCCCTTTCGTCTTCACGTC
GACATGCGATGGCGGTAGAAAAACTGTCTGAAGGCATTCGTCTGTTCGCCGTTGATCAACGCAAACTGGAAGATCTTCTTGCCGCCAAA
CTATAAACCAGCCACGGAGTGTTATATGTCCCGAAAAGACCTTGCCAATGCGATTCGCGCACTCAGTATGGATGCGGTACAAAAAGCCA
ACTCTGGTCATCCCGGGGATCCCGGGATCCTCTAGATTTAAGAAGGAGATATACATATG
>ssb
GAGGCAATTCCGACGTCTAAGAAACCATTATTATCATGACATTAACCTATAAAAATAGGCGTATCACGAGGCCCTTTCGTCTTCACCTC
GAGACAATGAGCTTGTCGCGGGGGATAACGAGGTTGATGTTTTTGAGATTATGGGTGCGGGCGCCCCGAACTTCGATCTTATCCATTCA
CCTTTCCCGGATTAAACGCTTTTTTGCCCGGTGGCATGGTGCTACCGGCGATCACAAACGGTTAATTATGACACAAATTGACCTGAATG
AATATACAGTATTGGAATGCATTACCCGGAGTGTTGTGTAACAATGTCTGGCCAGGTTTGTTTCCCGGAACCGAGGTCACAACATAGTA
AAAGCGCTATTGGTAATGGTACAATCGCGCGTTTACACTTATTCAGAACGATTTTTTTCAGGAGACACGAACATGGCCAGCAGAGGCGT
AAACAAGGTTATTCTCGTTGGTAATCTGGGTCAGGACCCGGAAGTACGCTGGATCCTCTAGATTTAAGAAGGAGATATACATATG
>ycaP
GAGGCAATTCCGACGTCTAAGAAACCATTATTATCATGACATTAACCTATAAAAATAGGCGTATCACGAGGCCCTTTCGTCTTCACCTC
GAGTTTTTGCTGGAAGCGAGCGATGGAATCTTCCAGAGCGGCATCTTTGCCGGGGATAAATGTTTGCGTCATGAAAATCACTTTAGTCG
TACGGAAAGCGCGCAATAATACGGGTTTTATCTCAAAGGCGCTATCACCGCCGCCATGCCGATGAACTGTTGACTATGCTTTTAGCGGA
TAAACCACGTTAAGAGCATAAAAATGAAGGCGTTCGATCTCCACCGTATGGCATTTGATAAAGTGCCTTTTGATTTCCTTGGCGAAGTT
GCGGATCCTCTAGATTTAAGAAGGAGATATACATATG
>b0540
GAGGCAATTCCGACGTCTAAGAAACCATTATTATCATGACATTAACCTATAAAAATAGGCGTATCACGAGGCCCTTTCGTCTTCACCTC
GAGATCTTTTCGCGTCACCCACATGCTGTACTGCACCTGGGCCATGTAAGCCGATTTTATTGCCTCGAAACCACCGAGCCGGAATTTCA
TGAAATCCCGGGAGGTACGAGTATTGCCGGAAGCGTGGCCTGTATCCGGATGCAGAGTCTTATCCGTGGAAATCGAACGCGCATTACTG
GTTGGTTACCAACTTGTACCAGAACATGCGGGCCAATGCGCTGGCTGACGCGGAATTACGGCGCAAGGCTGCCGATGAACTGACCTGTA
TGACAGCGCGAATTAACCGTGGTGAGACGATACCTGAACCAGTAAAACAACTTCCTGTTATGGGCGGTAGACCTCTAAATCGTGCACAG
GCTCTGGCGAAGATCGCAGAAATTAAAGCTAAGTTCGGACTGAAAGGAGCAAGTGTATGACGGGCAAAGAGGCAATTATTCATTACCTG
GGGACGCATAAGAGCTTCTGTGCACAGGACGTTGCCGCGGTAACAGGCGCAACCGTAATCTGATCTTACCCAGCAATAGTGGACACGCG

GCTAAGTGAGTAAACTCTCAGTCAGAGGTGACTCACATGACAAAAACAGTATCAACCAGTAAAAAACCCCGTAAACAGCATTCGCCTGA
ATTTCGCAGTGAAGCCCTGAAGCTTGCCTCGAGCGGAGATCCTCTAGATTTAAGAAGGAGATATACATATG
>yqgD
GAGGCAATTCCGACGTCTAAGAAACCATTATTATCATGACATTAACCTATAAAAATAGGCGTATCACGAGGCCCTTTCGTCTTCACCTC
GATGGATCCTGCCCTTCAGAGACGGACTCGGACGTAAAAAGGTGTTTTGCCATATTTAATATCACCTAAAGAGAATTTGGTTAGCTCAA
ACTGTTGTGTGGATTTTCTGTGGTAGCGGATCCTACCACGACTCTGCAGGTTAAAAACACTGGCAGTCTGAGTGTTAATCGGTATGGAT
GGATTAACATCTGGATGGCCTCGAGAGATCCTCTAGATTTAAGAAGGAGATATACATATG
>lexA
GAGGCAATTCCGACGTCTAAGAAACCATTATTATCATGACATTAACCTATAAAAATAGGCGTATCACGAGGCCCTTTCGTCTTCACCTC
GAGACGCGCAAAAGATATGGGATCCGCTGCGGTGCTGATTGCCATTATCGTCGCCGTGATTACCTGGTGCATTCTGTTATGGTCGCATT
TTGGATAACCCTTCCAGAATTCGATAAATCTCTGGTTTATTGTGCAGTTTATGGTTCCAAAATCGCCTTTTGCTGTATATACTCACAGC
ATAACTGTATATACACCCAGGGGGCGGAATGAAAGCGTTAACGGCCAGGCAACAAGAGGTGTTTGATCTCATCCGTGATCACATCAGCC
AGACAGGTATGCCGAGATCCTCTAGATTTAAGAAGGAGATATACATATG
>cobU
GAGGCAATTCCGACGTCTAAGAAACCATTATTATCATGACATTAACCTATAAAAATAGGCGTATCACGAGGCCCTTTCGTCTTCACCTC
GATGGATCCGGGGTTGCTGAAAAACGATAACCAACTGGCGATGTTATTCACGCTGGCCAACCTGTTTCGGGCGGACCAAATGATACGTC
AGTGGGAGAGATCTCACTAAAAACTGGGGATAACGCCTTAAATGGCGAAGAAACGGTCTAAATAGGCTGATTCAAGGCATTTACGGGAG
AAAAAATCGGCTCAAACATGAAGAAATGAAATGACTGAGTCAGCCGAGAAGAATTTCCCCACTTATTCGCACCTTCCCTAGTGCATAGT
CGCCACCATTCCGGATAATTCTATTACGTTGTTTTACAAGTTCTTTTACAGCGCAGATCTGTCGCTGACGTTGAATCAATAGCCTGGAT
TTCAGAACACAATGCCCGTTTGCTGGTTGTTCCGGCAGTAGCATTAATTCATTATCTGAATGAAATGTGTCAATCCACAGAAACCGCTG
TCAATGACGCATTTTGTCGGACGTTAACACGAGAAAGTCTGGCTATTCCGTAATGCCATCTTATTTCGTAATGGATTGAGAATGCTGGA
TTTTTAATTATATTACGCGCAGTATTAATCGCGTAAATGTTTTGCGGTCAGCAACGGTGGGGGATCCATTGTTCTGTTCAATGAATACC
ATCATACATTTGAATCAGGAGTCGGTATGATGATTTTGGTGACGGGGGGCGCACGGAGCGGGAAGAGTCGCCACGCAGAGGCGCTTATT
GGGGACTCCTCGAGAGATCCTCTAGATTTAAGAAGGAGATATACATATG
>ybgE
GAGGCAATTCCGACGTCTAAGAAACCATTATTATCATGACATTAACCTATAAAAATAGGCGTATCACGAGGCCCTTTCGTCTTCACCTC
GAGTGCTCTACACCGCCTGGTGTTACTGGAAAATGTTCGGTCGTATCACCAAAGAAGATATTGAACGTAACACCCACTCTCTGTACTAA
GTAAGGAGCTAAAAATGTGGTATTTCGCATGGATTCTGGGAACGCTTCTTGCCTGTTCGTTTGGGGTAATCACCGCGCTGGCGCTTGAA
CACGTCGAATCAGGCAAAGCCGGTCAAGAAGACATCTGATGAGTAAGATTATCGCGACTTTGTATGCGGTAATGGACAAGCGCCCCCTG
CGGGCGCTTTCCTTCGTGATGGCGCTTCTGGATCCTCTAGATTTAAGAAGGAGATATACATATG
>ymcE
GAGGCAATTCCGACGTCTAAGAAACCATTATTATCATGACATTAACCTATAAAAATAGGCGTATCACGAGGCCCTTTCGTCTTCACCTC
GAGCAATGAATTCCGCACGCTGAACGAAAATCAGAAAGTTGAATTTTCTATTGAGCAGGGGCAACGTGGCCCCGCGGCAGCGAACGTTG
TTACGCTCTAAGGTTGCCATTATTACTCAACATCTCCATTTCCGCTGTCCATGTTGTCATGGTTCACAGTACCGCACATCGGCATTCGA
TGTGACGGAGCGAAACCCTTTGGGCGCTAAGTGTATTTTTTGTAAATCGACGATGATCACCTTTGATAACGTCGCGCTGCAAATACGCA
CTGACCATGCGCCGCTGGATTTCACAAAATAATATCAGGCTCCCTCGTGGAGCCTTTTTGGATCCTCTAGATTTAAGAAGGAGATATAC
ATATG
>ymcC
GAGGCAATTCCGACGTCTAAGAAACCATTATTATCATGACATTAACCTATAAAAATAGGCGTATCACGAGGCCCTTTCGTCTTCACCTC
GATGGATCCCGTTGGCAGTTGTTGCAGCAAATGACAGTGGCGATGGCGGTTCTCATAACACCTCAACCACCACGTCCACTACACGCTAA
TACTGCTTTAACGCGTCGAAATTAAAGTTAAATGATACCCGGTTGCGATATTCGTACCGGGCTTTAGAGATTTGCATTTTTTTAATTAA
CCTTTCAAGGATTAAAGGTGCGCCCTCTTATTTTATCGATTTTCGCACTATTTCTTGCGGGATGTACGCACAGCCAGCAAAGTATGGTC
GATACATTTCGCGCCAGCCTTTTCGATAACTCGAGAGATCCTCTAGATTTAAGAAGGAGATATACATATG
>b0309
GAGGCAATTCCGACGTCTAAGAAACCATTATTATCATGACATTAACCTATAAAAATAGGCGTATCACGAGGCCCTTTCGTCTTCACCTC
GATGGATCCGCCTGGCTCTGGGATTTTTTCTAAGCATTGTAATTTCTGTAATGTTTTGCCTTGTCAAAAAAATGGTAGATGAATACCAA
CAAAACTCTGGGCAGTAATAAACAACATTCTGATAGAGAGTGCCTGTATAAAGCTGAAGTGACAATGACCGGGAAGGGAACCCGTTTTC
AGGAATGGATGCCTGTCTTTTTCACTGTAAATATCCTGGCATTGGTAATACCAGGATATTTACAGAAGAGCAGCATAACCATGACGGAA
CGGGCTCGAGAGATCCTCTAGATTTAAGAAGGAGATATACATATG
>ilvY
GAGGCAATTCCGACGTCTAAGAAACCATTATTATCATGACATTAACCTATAAAAATAGGCGTATCACGAGGCCCTTTCGTCTTCACCTC
GATGGATCCGGCCCATAAAGCGACATTTGCCCAGCTGTGCCAGCTGCTGGCGCAGATTCAGTGTATTGAAGTAGTTAGCCATGGTGATT
CCTCGTGATGTTGTGCTTCTTATTGTTCGGTTCGCGTTTGCGAGATTGAATTCACTATATGACAGGAAATTTATTGCGGAAATTGATAT
ATTCACAACGTCACATTGCAATTTTTGCAACGTCAACATCGAGGGCTGTCCCTGTGGATTTACGCGATCTGAAAACCTTCCTGCATCTG
GCGGAAAGCCGCCATTTTGGCCGCAGCGCGCGGGCGATGCACGTTAGCCCATCCACCTCGAGAGATCCTCTAGATTTAAGAAGGAGATA
TACATATG
>cytR
GAGGCAATTCCGACGTCTAAGAAACCATTATTATCATGACATTAACCTATAAAAATAGGCGTATCACGAGGCCCTTTCGTCTTCACCTC
GATGGATCCTGCTGGATGTTGATCCGATTGAGGGTTAAACCGCTCACGATGCGAGGCGGATCGAAAAATTCAATATTCATCACACTTTT
CATGAAAATTCTGTAACCGTTTTCACGCGCTATCTGCTAAAAATGTTGCCGATGTGAAGTAAACATGGATGTAGTACGCCTGACGTGCC
AGGCGAGGAGTGAGTGTGAAAGCGAAGAAGCAGGAAACTGCCGCGACCATGAAAGACGTTGCCCTCAAGGCAAAAGTCTCTACAGCGAC
CGTCTCCCGAGCATTAACTCGAGAGATCCTCTAGATTTAAGAAGGAGATATACATATG
>nupC

```
GAGGCAATTCCGACGTCTAAGAAACCATTATTATCATGACATTAACCTATAAAAATAGGCGTATCACGAGGCCCTTTCGTCTTCACCTC
GAGCGCAATGAACGCAGGTCCCATTAATGCGAGCCTCATCTTGCGCGCCGCCCGTCCGCTGCTACTCTCAACGCGATAGTTCGTCATCT
TGTGCCTCTAAAACATAGCCTTTGCTATGTTTCATGCTATGCCAAACGAGAATGATTATCAAATTCATTTAAATGGATTGTGGTGATTT
CTCTGATAGACCAGGATTATGACTACGAAAAGATTGATGACTTGAATGTTATGGTTATGTTTAATGTTAGCACATTTACATAACTTTCA
GCTTCCATACACAACATAGCAGAAATGTATGACAGATCACTATTTTTGAAGCCTGTCACAGGACGTCATTATAGTGTGTGTCAGATCTC
GTTTTCCTTAACCATGTTACATAGAATGTGCACGGAAATTTAACCTGCCTCATATTTGGAGCAAATATGGACCGCGTCCTTCATTTTGT
ACTGGCACTTGCCGTTGTTGCGATTCTCGCACTGCTGGTAAGCAGGGATCCTCTAGATTTAAGAAGGAGATATACATATG
>hcp
GAGGCAATTCCGACGTCTAAGAAACCATTATTATCATGACATTAACCTATAAAAATAGGCGTATCACGAGGCCCTTTCGTCTTCACCTC
GATGGATCCTCGCCTTTTTCTCTGCGTAATACCTCTCTGGCGGTAGATCCCTGCCGCCAAAATTGCGCTAAATCAATCTCCCTTAAAGT
TGCATGAAAAATCCCTTTTATCCCCGCGTTAAGCGTCTTAACCTTAAACATGTATATTAAATATAACTTTAAAAGGTGTGATCATGTTT
TGTGTGCAATGTGAACAAACTATCCGTACTCCGGCAGGAAACGGCTGCTCATACGCGCAGGGGATGTGTGGTAAAACGGCGGAAACCTC
GAGAGATCCTCTAGATTTAAGAAGGAGATATACATATG
>ytfQ
GAGGCAATTCCGACGTCTAAGAAACCATTATTATCATGACATTAACCTATAAAAATAGGCGTATCACGAGGCCCTTTCGTCTTCACCTC
GAGCGTTAGCCGGGATCTCAATAACAACGTAGATGTCTTCCGGCAGATCTTTACCCGCAGGGACGTTGAGTAAGCTCATGTCTGTTTCC
TTTAAAAATATGTGGCAAACAAGTGCCGAGTATTATAGCCAACTCGCGCCGAATGTCTTCGCTTGTTTTCGTCTTCATTTTCCTTTTTC
ACCAGTTTTAAGACGGTATACAGAGCAGGAAAATCCATAACCGTAGCCGCATTTTTCATAGTGAGATGAAAGCGATTACAAACTTGTGA
TTAACGTTTTATTTACTTTTTTGAAGTGTGATGTAACGCAATCTGTTACATAACGAATTGTCTATAGTTTTTTCGCGAACATCTTTTAA
CCAATAATAACTACCCCGACGAGGACAACCCTATGTGGAAACGCTTACTTATAGTCTCTGCAGTCTCGGCAGCCATGTCGTCTATGGCG
TTGGCCGCTCCATTAACCGTTGGGATCCTCTAGATTTAAGAAGGAGATATACATATG
>ytfR
GAGGCAATTCCGACGTCTAAGAAACCATTATTATCATGACATTAACCTATAAAAATAGGCGTATCACGAGGCCCTTTCGTCTTCACCTC
GAGGGCACCATGCCTGAAAAGCTGACGTTAACCAAATCCACCCTTTACCTGCCTGATACCGCAAAAGAAGAATTAGAGAAGAAGAAAAA
TATGGGGTATTGAGGGTTGCTATGCCTGATGCCGATTCGTAGGCCGGATAAGGCGCTCGCGCCGCATCCGGCGATGGTGCACTGAAGCC
TGATGCGACGCTTACCGCGTCTTATCATGCCTACTGGGAGCACGCTTTACACCGGGGGAAACCATGACGACCGACCAACACCAGGAGAT
CCTCCGCACCGAAGGATTAAGTAAATTTTTCCCCGCCGTCAAAGCGTTAGACAGGATCCTCTAGATTTAAGAAGGAGATATACATATG
>yeaM
GAGGCAATTCCGACGTCTAAGAAACCATTATTATCATGACATTAACCTATAAAAATAGGCGTATCACGAGGCCCTTTCGTCTTCACCTC
GATGGATCCTCATCAGAATGCCAGCGATAAGGACAATCCTGTTTTTGCCGCTTAATGAAGTTGAACAGGTCATATGCTTCTCATTGTTA
CAAACGAGGCTGTACCTTACCGTTTTTATTGTTATGTTGAATCAAGCTAAAATGACAAATTATCGCTAAGTTCGGACAAAATGATGCAC
AGGTTGAATCTCAATGGCTATGAACCCGACCGACACCATGAAGCCGCGGTCGCATTTTGTATTCACGCCGGAACGGATGAGTTACTCGA
GAGATCCTCTAGATTTAAGAAGGAGATATACATATG
>yeaL
GAGGCAATTCCGACGTCTAAGAAACCATTATTATCATGACATTAACCTATAAAAATAGGCGTATCACGAGGCCCTTTCGTCTTCACCTC
GAGCTGGTTGCCGACCCTCTACTCTTCGAACGATTTGATGAAATCGCCTTCAATGCAGGCATGCTGGATAAATCCGTTATTTGAAAAC
CGCGGATTATTTGCGCATTGCGCAACCAGAACTGGTCAATTTCCGCCGCACTGCGTAACTAGCCGGTCCGTTCGATAAGAAGAACGGAC
AAAACCAGTACAACAGCAATGGCAAAAAACGATGACGTGATAATCAGTGTCTCGACAAACATTTGATCGTTCATAGCATGCCCCAGATC
TTCTAGATTTAAGAAGGAGATATACATATG
>rhtB
GAGGCAATTCCGACGTCTAAGAAACCATTATTATCATGACATTAACCTATAAAAATAGGCGTATCACGAGGCCCTTTCGTCTTCACCTC
GATGGATCCACGCTGACGCCAGAAATCAGTCAGCGGTCCCATGGTAAAAGCAGCAAACGCGTTTTCTCTTGTTTCCCAGTCTTTTTGCT
GCTGAAACATCGGGTAATCTGCCTCTTAAACCACGTAAAATCGTTTTTTTTAGCGTGCCTGACACAACGCTGCGACAGTAGCGTATTGT
GGCACAAAAATAGACACACCGGGAGTTCATCATGACCTTAGAATGGTGGTTTGCCTACCTGCTGACATCGATCATTTTAAGCCTGTCGC
CAGGCTCTGGTGCAATCAACACTATGACCACCTCGCTCAACCACGGTTATCGCGGCGCGGTGGCGTCTATTGCTGGGCTTCAGACCGGA
CTGGCGATTCATATTGTGCTGGTTGGCGTGGGGTTGGGGACGCTATTTTCCCGCTCAGTGATTGCGTTTGAAGTGTTGAAGTGGGCAGG
CGCGGCTTACTTGATTTGGCTGGGAATCCAGCAGTGGCGCGCCGCTGGTGCAATTGACCTCTCGAGAGATCCTCTAGATTTAAGAAGGA
GATATACATATG
>yeaH
GAGGCAATTCCGACGTCTAAGAAACCATTATTATCATGACATTAACCTATAAAAATAGGCGTATCACGAGGCCCTTTCGTCTTCACCTC
GAGCGACGACTTTGTCGACCGTATGATGGAAAAAGGCTACACCCGTAAACAGGTGCGTTTACTGTGCGAATGGTATTTGCGCGTACGTA
AATCGTCTTAACAACCCTGGCCCGGTACACATGTTACCGGGCCTACAACGACAGCGAACCGTGGGCCTGAGAAGCGGCAACACAGGCGT
AGCATACAGTTGGCAAATGTAGTACGGGGGGCATATGACCTGGTTTATTGACCGGCGTCTGAACGGCAAAAACAAAAGCATGGTGAATC
GCCAGCGTTTTGGATCCTCTAGATTTAAGAAGGAGATATACATATG
>yeaU
GAGGCAATTCCGACGTCTAAGAAACCATTATTATCATGACATTAACCTATAAAAATAGGCGTATCACGAGGCCCTTTCGTCTTCACCTC
GAGGCGAGCCACCAGCATAAAGACGCGCAAATCATTCAGCAGCGGTAAATTATTCATGATTCGTGTTTTATGTTTCACCAGTTACGGGG
ATTAATTCCTTTTCAGTCAATTATAGGATGGTGATGTTGTCAATTTTGATGGTCAGGAAGTGAGAACCCAATGATGAAAACGATGCGTA
TTGCTGCGATCCCGGGAGACGGGATTGGCAAAGAAGTCCTTCCTGAAGGGATTCGCGTGTTACAGGCTGCCGCTGGATCCTCTAGATTT
AAGAAGGAGATATACATATG
>yeaT
GAGGCAATTCCGACGTCTAAGAAACCATTATTATCATGACATTAACCTATAAAAATAGGCGTATCACGAGGCCCTTTCGTCTTCACCTC
GATGGATCCAGCGGCAGCCTGTAACACGCGAATCCCTTCAGGAAGGACTTCTTTGCCAATCCCGTCTCCCGGGATCGCAGCAATACGCA
```

TCGTTTTCATCATTGGGTTCTCACTTCCTGACCATCAAAATTGACAACATCACCATCCTATAATTGACTGAAAAGGAATTAATCCCCGT
AACTGGTGAAACATAAAACACGAATCATGAATAATTTACCGCTGCTGAATGATTTGCGCGTCTTTATGCTGGTGGCTCGCCTCGAGCGG
GATCCAGCGGCAGCCTGTAACACTATGCTGGTGGCTCGCCTCGAGAGATCCTCTAGATTTAAGAAGGAGATATACATATG
>yeaV
GAGGCAATTCCGACGTCTAAGAAACCATTATTATCATGACATTAACCTATAAAAATAGGCGTATCACGAGGCCCTTTCGTCTTCACCTC
GAGAAGTGATTGCTCACGGGCCGAAAACACCTGATATGAAAGGCAATGCCACCACGCCACAGGTTGCCGACGCGATTTGCAAAATTATT
TTGCGTTAAGGTCAAACCAGTTTATTTGAACCGCGTCACTGACGCGGTTTTTTTATTCGTTCTTTGCAGTAAATAACCTGCGTCATTTC
ACCTTTTATTGTTTCCGTTTCGTGTTTTATGGCTTTCCGTATTCTTAATTGTTTAATTTATGTAACATGCAAATTTTGTTACGCGTACG
TTAGGTTCCGCCGTACAGGTATTGGTTTTGCTGGCAATGGGACTGGTGATTTATTTAGCCACCAGTAAATACGGCAATATTCGTCTTGG
CGAAGGAAAACCGGAATGGATCCTCTAGATTTAAGAAGGAGATATACATATG
>yeaP
GAGGCAATTCCGACGTCTAAGAAACCATTATTATCATGACATTAACCTATAAAAATAGGCGTATCACGAGGCCCTTTCGTCTTCACCTC
GAGTGGCTGAGAAGGGGTTGAACAACCAGCCAGGACCAGCAAGCAAGGCAGAACAAAACTGATAATTTTCATTTATTGATCTCACATAT
TTATCCAAGATTAGAGTATCGCGGTATCGTTTTGTTTTGCAGCACTATTTTTATTACATTCACTCAAAACATATTACGTCTTGTTTCAT
CTTTGTTGATGATGTTTTATCATGCCTGCAAAGGATCCTCTAGATTTAAGAAGGAGATATACATATG
>yeaS
GAGGCAATTCCGACGTCTAAGAAACCATTATTATCATGACATTAACCTATAAAAATAGGCGTATCACGAGGCCCTTTCGTCTTCACCTC
GATGGATCCGTCAGCGAAAGTGCGGATAACGGTAGAATTTTTACGCCAGTATTTTGCCGAGCACTACCCGAATTTTTCACTGGAGCATG
CCTGATTAATGATTCAATTATCGGGTTGATATCAGGTTAAAACCTGATTTTCTCCTTTCTAAGCCGCTACAGATTGGTTAGCATATTCA
CCTTTAATCGCGCATGATCGAAAGATAATTAAAGAGGTTAATGTGTTCGCTGAATACGGGGTTCTGAATTACTGGACCTATCTGGTTGG
GGCCATTTTTATTGTGTTGGTGCCAGGGCTCGAGAGATCCTCTAGATTTAAGAAGGAGATATACATATG
>ytfB
GAGGCAATTCCGACGTCTAAGAAACCATTATTATCATGACATTAACCTATAAAAATAGGCGTATCACGAGGCCCTTTCGTCTTCACCTC
GATGGATCCACTTTGTCGCGGGAATGATGACAAATCCCAGGGAAAAAGAAACAATAGCAATATCCCGTATGAATATAATAAAGGCGGGT
TTACCACGGACATCATAGCGTTAGCTTATTCATAGGCTATGATTGAGGAACAAGACGCGGAGCAGGAGGAAACCATGCCCGGGCGCTTT
GAACTAAAACCAACCCTGGAGAAAGTCTGGCACGCGCCGGACAATTTTCGCTTCATGGACCCCTCGAGAGATCCTCTAGATTTAAGAAG
GAGATATACATATG
>ytfM
GAGGCAATTCCGACGTCTAAGAAACCATTATTATCATGACATTAACCTATAAAAATAGGCGTATCACGAGGCCCTTTCGTCTTCACCTC
GAGGCATCGGGGTGTTACGTCCAGGCAGGGCATCGGCGGGGGAAACCAGATGCTTTTTATCAAATAAACTCATGGTGTCGCTCTCCCGA
AAATCGGTCATTGGGGTTAAGGTTGTAACAAGAGGCGTATTTGCACACAATAACCACCGTGAATAGGTCTAAAGTAAAACATAAGAAAT
ATTTGGGGTTTAGTCTGCTTTTTAATCCATATTACTGGATTTTTGTTAAGCCGTTTAACGGCGTTCCAGGGGCAGGAAAAAAGGATATT
CAGGAGAAAATGTGCGCTATATCCGACAGTTATGCTGTGTAAGCTTACTCTGCTTAAGCGGATCTGCCGTCGCCGCGAACGTCCGTCTA
CAGGTCGAGGGGTTATCGGGGATCCTCTAGATTTAAGAAGGAGATATACATATG
>ytfL
GAGGCAATTCCGACGTCTAAGAAACCATTATTATCATGACATTAACCTATAAAAATAGGCGTATCACGAGGCCCTTTCGTCTTCACCTC
GATGGATCCACTGTGGAATTGGCGGAATTGGCGTCTGTCTGCCGCCGGAAGCATAGCGTTACGGGTACAAATGTAGTTTGTTGATAAAG
TGCTCTTTGTCTATGCCTGATGCGGCGTAAACGACTTACCAGGCCTACAAAATCGTCCAAATTCAATATATTGCAAGGACTGCGTAGGC
CTGATCGGCATAGCGTATCAGGCAGTTTTGCATTTTTCAGCTGTCTGAATTTTAACCCTCTGGCGGCTTTACAGTACCTTACGCTATAC
TAGCCACTGAAAATGCCGGTTCACTTTCTTCGAATCGGCTTTCAATGTGTATTTCACACAAATTAATCAACTTCCCCTTCCGAGGATCT
GGCCTGAAAGGTCGGATAAGATATGTTAAACAGTATTTTAGTCATACTCTGCTTGATCGCTGTAAGTGCGTTCTTCTCGATGTCCGAGA
TCTCGCTTGCCGCCTCACGCAAAATCACTCGAGAGATCCTCTAGATTTAAGAAGGAGATATACATATG
>nupX
GAGGCAATTCCGACGTCTAAGAAACCATTATTATCATGACATTAACCTATAAAAATAGGCGTATCACGAGGCCCTTTCGTCTTCACCTC
GATGGATCCCCGCCAATACCAAAGTCGGCATCACTATTGATACAGACTGGTTCTGGGGATTAGTCGAAGAGTGCGTGCGCGGCTACATC
AAAACCCATTAAGTGTTGATAACAAGCCGGGCCGGTATCGCAAGATGCAGGCCTGGCTATTTCTGATGGGCAGAAAGCCGTCGCCCAAT
ATGACATGAAGAGATAATCACTATGGATGTCATGAGAAGTGTTCTGGGAATGGTGGTATTGCTGACGATTGCGTTTTTACTGTCAGTAA
ACAAGAAGAAGATCAGCCTGCGTACCGCTCGAGAGATCCTCTAGATTTAAGAAGGAGATATACATATG
>ytfI
GAGGCAATTCCGACGTCTAAGAAACCATTATTATCATGACATTAACCTATAAAAATAGGCGTATCACGAGGCCCTTTCGTCTTCACCTC
GAGGTTCACGACTGGCAGGGTAAACCGCTGGATTACACTCCGCGTGAGTCGTTCCTGAATCCGGGGTTCAGAGTGTCTATTTACTAAAT
TCAGATGGCAGAAACAGTGTATTTCCTGATTCTGCCATCCTGATTTCTCCCAACCTAAAAAGTTATAAATAAAAAGAGATTGTATTTAA
AGTGCAAAAATTCAATTGCTAATAAGTTACATTTTAATAATGAGCGTTTTTTGATAGTTTACTTCTATAGTGAGATATTTAATGGCGAC
ATAAAGTAACCAAATAAAATAAGGTTGTCATATGTTACCCAGGATCAGACACAATAATTTTATTGGTGCGGTGGAGTTATTTGTAAAGT
CTTCGTATACAAAAACACATTCAAACAATTTTTTTAACAACATCCATCATGCATTTAAGAAAAAAGATTGGATTTCGAATTATGATAGT
CTGTTAACTTTGAGGGAGTTCTTTCGTTGCGCGGATCCTCTAGATTTAAGAAGGAGATATACATATG
>yeaX
GAGGCAATTCCGACGTCTAAGAAACCATTATTATCATGACATTAACCTATAAAAATAGGCGTATCACGAGGCCCTTTCGTCTTCACCTC
GAGGGCCGACAGTAGCGGTAGTGGCATTTCCGAACATGGTATCGCCCATTTCCATAATCTGCTGGCGCAGGTGTTTAAGGACTAATGAC
ATCGGCGGCGGTATTTTCCGCCGCTGGGCTGATTTTTGATGGAGTACAGCAATGTCAGACTATCAAATGTTTGAAGTACAGGTGAGCCA
GGTTGAACCCCTTACCGAACAGGTGAAACGCTGGATCCTCTAGATTTAAGAAGGAGATATACATATG
>blc

```
GAGGCAATTCCGACGTCTAAGAAACCATTATTATCATGACATTAACCTATAAAAATAGGCGTATCACGAGGCCCTTTCGTCTTCACCTC
GATGGATCCGGGTATCGTGATGCTGGCAAACAAAAACTATCCCAATCCAGCGAGAGTCGACGCCGCCTGGCAGATTCTTAACGCTCTAC
AGTAAAATTCCATCGGGTCCGAATTTTCGGACCTTTTCTCCGCTTTTCCTTGCTGTCATCTACACTTAGAAAAAAACCAGTAAGGAAAC
ATTTATGCGCCTGCTCCCTCTCGTTGCCGCAGCGACAGCTGCATTTCTGGTCGTTGCCTGCAGTTCTCCTACGCCGCCGCGTGGCGTGA
CCGTAGTACTCGAGAGATCCTCTAGATTTAAGAAGGAGATATACATATG
>yeaZ
GAGGCAATTCCGACGTCTAAGAAACCATTATTATCATGACATTAACCTATAAAAATAGGCGTATCACGAGGCCCTTTCGTCTTCACCTC
GATGGATCCGTGCGGTTCGTTTCCTTGCGATACCATCCTCCAGGTAATTTGTTACGGAGTATGGTATGATGCGCGCCTTTTTATTGATC
TAAGCACGAAGACCCATGCGAATTCTGGCTATCGATACCGCGACAGAGGCCTGCTCTGTCGCCCTGTGGAACGACGGTACTGTCAACGC
TCATTTTGAGCTTTGCCCTCGCTCGAGAGATCCTCTAGATTTAAGAAGGAGATATACATATG
>argZ_argQ
GAGGCAATTCCGACGTCTAAGAAACCATTATTATCATGACATTAACCTATAAAAATAGGCGTATCACGAGGCCCTTTCGTCTTCACCTC
GATGGATCCCGATGCATCCGTAGCTCAGCTGGATAGAGTACTCGGCTACGAACCGAGCGGTCGGAGGTTCGAATCCTCCCGGATGCACC
ATATTCCTCGAGAGATCCTCTAGATTTAAGAAGGAGATATACATATG
>yjiY
GAGGCAATTCCGACGTCTAAGAAACCATTATTATCATGACATTAACCTATAAAAATAGGCGTATCACGAGGCCCTTTCGTCTTCACCTC
GATGGATCCCAAAAACGGCCAAAACCAGCAGTAAGCTGGTCACAATTTTGATACGTTTTAACATGGTTTCTCTTTCCTGTGGACCGATG
CAAGATTTTCGGCCTGGAAAGGAAAAACTTTATGAATATCCCGCGGAGATTACGCCGAGTGAATTTTATTCACACTCTGAATTTAAAAA
GCCATTATTTACATAAATTATTCACAATATAAATATGTGATATGAATCACATATTTATCGTCACTTAAACGACGCCTTTGCCGCTCAAC
CGCAAAACTGACCGCTTACATCCCTAAAATAACCACTCAGTTATTTACCTTACTTTACGCGCGCGTAACTCTGGCAACATCACTACAGG
ATAGCGGTCAATTTACCTCCTCAAACGCAACGCAAACCTAGAACGGCTTCGGCCAACTATTAATCAATACATGCCAGGTTTTACTATGG
ATACTAAAAAGATATTCAAGCACATACCCTGGGTGATTCTCGGAATCATCGGTGCATTCTGCCTCCTCGAGAGATCCTCTAGATTTAAG
AAGGAGATATACATATG
>ilvL
GAGGCAATTCCGACGTCTAAGAAACCATTATTATCATGACATTAACCTATAAAAATAGGCGTATCACGAGGCCCTTTCGTCTTCACCTC
GAGCTCAACAGTGATCGGGGGCGCATTTACTCCCAGGGCTGCGCGGGTATGAACAATTGACAGTGACATAAGCCCTCCTTGAGTCACCA
TTATGTGCATAAGATATCGCTGCTGTAGCCCGCTAATTCGTGAATTTTAGTGGCTGATTCCTGTTTATTTGTGCAAGTGAAGTTGAGTT
GTTCTGGCGGTGGAATGATGCTCGCAAAAATGCAGCGGACAAAGGATGAACTACGAGGAAGGGAACAACATTCATACTGAAATTGAATT
TTTTTCACTCACTATTTTATTTTTAAAAAACAACAATTTATATTGAAATTATTAAACGCATCATAAAAATCGGCCAAAAAATATCTTGT
ACTATTTACAAAACCTATGGTAACTCTTTAGGCATTCCTTCGAACAAGATGCAAGAAAAGACAAAATGACAGCCCTTCTACGAGTGATT
AGCCTGGTCGTGATTAGCGTGGTGGTGATTATTATCCCACCGTGCGGGGATCCTCTAGATTTAAGAAGGAGATATACATATG
>argP
GAGGCAATTCCGACGTCTAAGAAACCATTATTATCATGACATTAACCTATAAAAATAGGCGTATCACGAGGCCCTTTCGTCTTCACCTC
GAGAGTTCACCGTTCGCAAGCCCCTGTTTACAGGCAAATTGAGGCAATAACGCCACGCCATTCAACACAACCAGGGCGCGAACTCGCTG
AGCGAAATGCATTAGCGCAAATCCCTGATCTTTTAAGCTGCGCTTTTAACCACGGATACAAATGTTACCCGCCGACGGCTTCGGTATAT
GCAACCTGACACAAAATTGTGTCATAGTGCAGGAAAAAGCATTTACCAGGAGCAGACAACAGCAATGAAACGCCCGGACTACAGAACAT
TACAGGCACTGGATGCGGTGATACGTGAACGAGGATTTGAGCGCGGATCCGCGCTCAAATCCTCCGGGATCCCGAACGAGGATTTGAGC
GCGGATCCCGAACGAGGATTTGAGCGCGGAACGAGGATTTGAGCGCGGATCCTCTAGATTTAAGAAGGAGATATACATATG
>intB
GAGGCAATTCCGACGTCTAAGAAACCATTATTATCATGACATTAACCTATAAAAATAGGCGTATCACGAGGCCCTTTCGTCTTCACCTC
GAGCTTCAGTGCCGAAGTGGCGAAATCGGTAGACGCAGTTGATTCAAAATCAACCGTAGAAATACGTGCCGGTTCGAGTCCGGCCTTCG
GCACCAAAAGTATGTAAATAGACCTCAACTGAGGTCTTTTTTTATGCCTGAAATCCAGTGTTTATCTATCTTTCCCGCTATATTAACTC
TCTCAAGGTCAACCGATATCAACGTACATCTACCAACATATGTTGGTACAGATGATGGTATTTCCGGTTCGATAATGCTTGTACCAACA
GGGAGGGAATACGATGGCATTAACAGATATCAAAGTCAGAGCAGCCAAGCCAACGGATAAGCAATATTAGCTGACTGATGGTGGCGGTA
TGCATCTGCTTGTCCATCCAAATGGTTCTAAGTACTGGCGTTTGCAGTACCGTTGGATCCTCTAGATTTAAGAAGGAGATATACATATG
>argU
GAGGCAATTCCGACGTCTAAGAAACCATTATTATCATGACATTAACCTATAAAAATAGGCGTATCACGAGGCCCTTTCGTCTTCACCTC
GAGTCGCTCTGCCAGCATAAAAGCATTCTGATTTCGATGATAAAAATAACACTTTCACTGTGCTCTGGATTTGTTTGATCCTTTTCAGG
AAGGTAAAACCGTCTGTTCCGGGCAAGTCTATATCCATAATGATTAAATCAACAGGACGGGTTCGGAGATAATCGATGGTTATGCGATA
ATCATCCGTTTTCAGGACAATCTGCAATTCACTGTTTTTTTGCAACAGAACTTCAATAGACATTCTGATGATAGGATGAGTATCCATAA
TGATCACCGACGTTGGTTTCATAGTTACCAGTCTCATAGGAGCGGACAATTTTCCGTTAGGTCGGGAAATTGTACTTTGATACATGAAA
ATACGGGTTTTCTTGATTCAGACGCGCAGCGGTGTGCGTTTGTTTGCCGCTATGGATCCTCTAGATTTAAGAAGGAGATATACATATG
>yjiT
GAGGCAATTCCGACGTCTAAGAAACCATTATTATCATGACATTAACCTATAAAAATAGGCGTATCACGAGGCCCTTTCGTCTTCACCTC
GAGCGACGCGTGTTACAGCAGATGAGTGATGAGCGGTTGAAGGATATCGGGTTACGCAGGGAGGATGTGGAGTGAGGGGGGATATAGAT
TTATATATAATAAAACGTTTTTATGTTTTTAAATTAAGTTATAAAAATTTTCCCGAGACAATTCATCAATAGGTATGGAGTGTACAGGA
ATATCTTCTTCACTAACCTCTTTAAACTCATGTCGATAACGGTCAAAAAACGCACAGTGATTACTTGTGTTTTTTGACTGACTTTACAA
CTCTCATTATTCGCTATTGTGCAGTTTCTCTAATTGTTTTATACCCTGGAAAGTTAAATGTCAGCTACTGAATACTTTTTGATTGTTTG
AGATTTATTTTCATTTGAAATTATAAAATCAGGTGATAAATGAGTTGTGATTAATAGATGGTTGATATCATTTTTATCTAAAATTGATT
TATAGTATCGACCTGAAAAAATAGTTGTTGCCGCCTGAGTAACTATACAATATTCTGAAAGGTTTTCTTTCAAATTAGAAATGTTGTGG
GGTTTTGTTTTTGTATCTTTTATCTCTAAGGAGCTTGCTTTGGGTCAGGATCCTCTAGATTTAAGAAGGAGATATACATATG
>argW
```

```
GAGGCAATTCCGACGTCTAAGAAACCATTATTATCATGACATTAACCTATAAAAATAGGCGTATCACGAGGCCCTTTCGTCTTCACCTC
GAGAAAGCAGAAGCACGCCAAAAAGCAGAACGTGCGGAAAACATTAAGAAAAATTATAAAAACCCGGCATAAATGGCGAGGGTTTAAGC
AATCGAGCGGCAGCGTACTTACCCCGCACTCCATTAGCGGGTATACTCATGCCGCATTGTCCTCTTAGTTAAATGGATATAACGAGCCC
CTCCTAAGGGCTAATTGCAGGTTCGATTCCTGCAGGGGACACCATTTATCAGTTCGCTCCCATCCGTAGGATCCTCTAGATTTAAGAAG
GAGATATACATATG
>yjiJ
GAGGCAATTCCGACGTCTAAGAAACCATTATTATCATGACATTAACCTATAAAAATAGGCGTATCACGAGGCCCTTTCGTCTTCACCTC
GATGGATCCGTTGCGATGGATGCCTCTGGTAATATTTATATCGTTAGCGAACCAAATCGTTTTTATCGCTTTACCCCTCAATCATCGCA
TTAACAGGCTGGCATGCTTCACCTTTTTGTTATTAACTTGTTAACAACAAAAATGACAAACAAGGTTAAACATGCCTTCGTCCACGCAT
CCCGTAGAACGCTTTTCTTTCAGCACCGCGCTTTTCGGGATGCTGGTTCTGACCTTAGGTATGGGTTTAGGCCGCTTCCTCGAGAGATC
CTCTAGATTTAAGAAGGAGATATACATATG
>argI
GAGGCAATTCCGACGTCTAAGAAACCATTATTATCATGACATTAACCTATAAAAATAGGCGTATCACGAGGCCCTTTCGTCTTCACCTC
GATGGATCCCCAGTTGTTCCGGGTTTGCCATGGCTCATTCCTCTTAATGTCCTGCCGATACTCTTATTGTCACACACCACTGACATTGC
CTCCACCTTTGTGACAAAGATTTATGCTTTAGACTTGCAAATGAATAATCATCCATATAAATTGAATTTTAATTCATTGAGGCGTTAGC
CACAGGAGGGATCTATGTCCGGGTTTTATCATAAGCATTTCCTGAAATTACTCGATTTCACGCCAGCTGAACTCAACAGCCTGCTGCAG
TTAGCCGCGAAGCTGAACTCGAGAGATCCTCTAGATTTAAGAAGGAGATATACATATG
>yjiH
GAGGCAATTCCGACGTCTAAGAAACCATTATTATCATGACATTAACCTATAAAAATAGGCGTATCACGAGGCCCTTTCGTCTTCACCTC
GATGGATCCCTGCAACTCAATCCTGCGCAGAGCCTTTTCCACACAGCGTAATTTTTCCGCTATGCAAAAAACGCATAGCCAGAAACATA
CTTGCATTTTCCCCATACCCACATGGCTGATAAGGTTTTTCAGTTCCAGGAATTTTCCTGGCTCAAGCAACACACAAACACAAAAAAAC
AATGGGGATTGTTATGACACAACAAGGGGATGCTGTAGCCGGGGAGCTGGCGACAGAAAAAGTGGGCATTAAAGGGTATCTGGCCTCGA
GAGATCCTCTAGATTTAAGAAGGAGATATACATATG
>yjiM
GAGGCAATTCCGACGTCTAAGAAACCATTATTATCATGACATTAACCTATAAAAATAGGCGTATCACGAGGCCCTTTCGTCTTCACCTC
GATGGATCCGTTGGCGGTTGTATTGGGCTAATTTTATATTTGCTGTCGCAGCTCCCGGCCTTGTTCCCCCTCGGCAATTTTTAGAAATC
CATGAAAAACTAAGTGGCTAACCGCGAGAGAGATCAAATAAACCATACTAAAAGAATGGTAATAATGTGGCCTCTTTTCATTATCTCCC
GTGGTACGGGGAAGGAAAATCATGTCACTTGTCACCGATCTACCCGCCATTTTCGATCAGTTCTCTGAAGCTCGCCAGACAGGCTTTCT
CTCGAGAGATCCTCTAGATTTAAGAAGGAGATATACATATG
>argA_amiC
GAGGCAATTCCGACGTCTAAGAAACCATTATTATCATGACATTAACCTATAAAAATAGGCGTATCACGAGGCCCTTTCGTCTTCACCTC
GATGGATCCTTTTCCCCGGTGGGTATTGATATAGGGAACCGAATGGCGGAATCCCTCGACCAACTCGGTTTTACGTTCCTTTACCACGG
CACACCTCTTTGCATGATTATTCGAAATTAGTGTATTTTTATTCTGTTTTTTCGCAGGGTGCAAGTGTAAATTTTATGCGAGAGCGACT
TTTTTATCAGTAATGCCGTGAATATAAAAAGAATGTTTACCGTTTATTAGATGACAGATTATGCGTCTTTCGCTAAAGTTTCCGGTCAA
ATTAGTCGTTTACTTGTTACACAGCTTAGATTTCTTTTGCTCGGGAGAGGCATGTCAGGACTCGAGAGATCCTCTAGATTTAAGAAGGA
GATATACATATG
>argC
GAGGCAATTCCGACGTCTAAGAAACCATTATTATCATGACATTAACCTATAAAAATAGGCGTATCACGAGGCCCTTTCGTCTTCACCTC
GAGCTTCCGTGGCGCTTATTGAAGGTGTGGCAATCAGAGCGCGGTAAATCTCGATAAATGGCGGTAATTTGTTTTTCATTGTTGACACA
CCTCTGGTCATGATAGTATCAATATTCATGCAGTATTTATGAATAAAAATACACTAACGTTGAGCGTAATAAAACCCACCAGCCGTAAG
GTGAATGTTTTACGTTTAACCTGGCAACCAGACATAAGAAGGTGAATAGCCCCGATGTTGAATACGCTGATTGTGGGTGCCAGCGGCTA
CGCTGGCGCAGAGCTAGTGACCTATGTAAATCGCCATCCGCATATGGGATCCTCTAGATTTAAGAAGGAGATATACATATG
>argD
GAGGCAATTCCGACGTCTAAGAAACCATTATTATCATGACATTAACCTATAAAAATAGGCGTATCACGAGGCCCTTTCGTCTTCACCTC
GATGGATCCGCCAGTGGGATCTGGAAGGTGTGCAGTTCCATCCAGAAAGTATTCTTAGCGAACAAGGACATCAACTGCTGGCTAATTTC
CTGCATCGCTGATTTCTGATTGCCATTTAGTGATTTTTTATGCATATTTTGTGGTTATAATTTCACATTTGTTTATGCGTAACAGGGTG
ATCATGAGATGGCAATTGAACAAACAGCAATTACACGCGCGACTTTCGATGAAGTGATCCTGCCGATTTATGCTCCGCTCGAGAGATCC
TCTAGATTTAAGAAGGAGATATACATATG
>argE
GAGGCAATTCCGACGTCTAAGAAACCATTATTATCATGACATTAACCTATAAAAATAGGCGTATCACGAGGCCCTTTCGTCTTCACCTC
GATGGATCCGGTCACTAGCTCTGCGCCAGCGTAGCCGCTGGCACCCACAATCAGCGTATTCAACATCGGGGCTATTCACCTTCTTATGT
CTGGTTGCCAGGTTAAACGTAAAACATTCACCTTACGGCTGGTGGGTTTTATTACGCTCAACGTTAGTGTATTTTTATTCATAAATACT
GCATGAATATTGATACTATCATGACCAGAGGTGTGTCAACAATGAAAAACAAATTACCGCCATTTATCGAGATTTACCGCGCTCTGATT
GCCACACCTTCAATAAGCGCCACGGAAGCTCGAGAGATCCTCTAGATTTAAGAAGGAGATATACATATG
>argF
GAGGCAATTCCGACGTCTAAGAAACCATTATTATCATGACATTAACCTATAAAAATAGGCGTATCACGAGGCCCTTTCGTCTTCACCTC
GATGGATCCTGACAAAGTCATCGGGCATTATCTGAACATAAAACACTATCAATAAGTTGGAGTCATTACCCGGTAACAGATATATCGCC
TCCATCACCGCTGAAGTAATGAAGTAAGGCAGCAAGCCTCGCCATGTTCTCCGCCATTTTAGAAGCATAATCCTTAAAATCATAAAGAA
GGCCAATCATTCTCATTTCTGACTCGACCTGGTTGTAGAATTCGATCCAATGTCTTTCTGCTTCTGCAGAGAATCGGAGGCAGATACGA
TTATTTTCACACACGGACGGGTTTGCCTCCACCTTTGTAAGAAAGAATTGTGAAATGGGGTTGCAAATGAATAATTACACATATAAAGT
GAATTTTAATTCAATAAGTGGCGTTCGCCATGCGAGGATAAAATGTCCGATTTATACAAAAAACACTTTCTGAAACTGCTCGACTTTAC
CCCTGCACAGTTCACTTCTCTGCTGACCCTTGCCCTCGAGAGATCCTCTAGATTTAAGAAGGAGATATACATATG
>argG
```

GAGGCAATTCCGACGTCTAAGAAACCATTATTATCATGACATTAACCTATAAAAATAGGCGTATCACGAGGCCCTTTCGTCTTCACCTC
GATGGATCCCCGCCAGAAAAAGCGATACCAATACGTTGACCTACCGGGAGATGCTTGAGAATCGTCGTCATAAAATAACACCCTGCTTA
ATTAACTGATGATGAGCCTGGATTTCCGCTCTCACTGAATTTTTATGCAAAATAAATGAGTTTTCATTTAATCATCTTTTATCGGAGAC
AGGAAGAGTTTAGTGTGTTTTTTGTAAAATAATGCGCTTAAGGGAGAGCAGGAGAAGGCAAAAGTATTCAACAAATGAAAGTGAACTGG
ATATTCATTCACATGATTAGCAATAAACGTTGACAAAATGTGGCGTGGATCACTATAATGCCTGCAGATTTTACGTCCCGTCTCGGTAC
ACCAAATCCCAGCAGTATTTGCATTTTTTACCCAAAACGAGTAGAATTTGCCACGTTTCAGGCGCGGGGTGGAGCAGCCTGGTAGCTCG
TCGGGCTCATAACCCGAAGATCGTCGGTTCAAATCCGGCCCCCGCAACCACTTTCCCTTCTCGAGAGATCCTCTAGATTTAAGAAGGAG
ATATACATATG
>yggM
GAGGCAATTCCGACGTCTAAGAAACCATTATTATCATGACATTAACCTATAAAAATAGGCGTATCACGAGGCCCTTTCGTCTTCACCTC
GATGGATCCACCAAAGATCCGCAGCAGATCCAGCAGATCTTCAATCAGTACTAATCGCCTCGCCCCGGTATCGTGCCGGGGCTTTTTCA
CTTCAGACTCACGTCCATTGCCAATTTTAATTACCCTAATGATAATCACCGGAATAAATTATTCCGCGTGAGGTTTTTCGGGTGAAAAA
GCAATGGATTGTTGGCACGGCACTGCTTATGTTGATGACTGGTAATGCCTGGGCAGATGGCGAACCGCCAACTGAAAACTCGAGAGATC
CTCTAGATTTAAGAAGGAGATATACATATG
>tyrS
GAGGCAATTCCGACGTCTAAGAAACCATTATTATCATGACATTAACCTATAAAAATAGGCGTATCACGAGGCCCTTTCGTCTTCACCTC
GATGGATCCCCTGCATGACCGCTTTTTGTACCAGCGTGAAAATGATGCGTGGAAGATTGATCGTCTTGCACCCTGAAAAGATGCAAAAA
TCTTGCTTTAATCGCTGGTACTCCTGATTCTGGCACTTTATTCTATGTCTCTTTCGCATCTGGCGAAAAGTCGTGTACCGGCAAAGGTG
CAGTCGTTATATACATGGAGATTTTGATGGCAAGCAGTAACTTGATTAAACAATTGCAAGAGCGGGGGCTGGTAGCCCAGGTGACGGAC
GAGGAAGCGTTAGCAGAGCGCTCGAGAGATCCTCTAGATTTAAGAAGGAGATATACATATG
>yggH
GAGGCAATTCCGACGTCTAAGAAACCATTATTATCATGACATTAACCTATAAAAATAGGCGTATCACGAGGCCCTTTCGTCTTCACCTC
GATGGATCCCCAGAACCTGGGCTGAAAATTGCGACGCTTGCATGGTCACCGAATTCACTGTTGTTGGGGGCAAGATTGCAGCACAGCGA
CGGCAGGGTGTAAACCGGAACTTTCCGCAGCCACGGCCTTATCATGCTTGCATCCGGCAACTAACTTTGGATAATGCCCGTTTTCAGAA
CACTTTCACAAGCGACTAAACCTTTATGAAAAACGACGTCATTTCACCGGAATTTGATGAAAACGGCCGCCCACTGCGCCGTATCCGTA
GTTTTGTGCGCCCTCGAGAGATCCTCTAGATTTAAGAAGGAGATATACATATG
>htrA
GAGGCAATTCCGACGTCTAAGAAACCATTATTATCATGACATTAACCTATAAAAATAGGCGTATCACGAGGCCCTTTCGTCTTCACCTC
GAGTATTACCGTTGCCGCCTGCTGCAGGATTATATCAGCGGTATGACCGACCTCTATGCGTGGGATGAATACCGACGTCTGATGGCCGT
AGAACAATAACCAGGCTTTTGTAAAGACGAACAATAAATTTTTACCTTTTGCAGAAACTTTAGTTCGGAACTTCAGGCTATAAAACGAA
TCTGAAGAACACAGCAATTTTGCGTTATCTGTTAATCGAGACTGAAATACATGAAAAAAACCACATTAGCACTGAGTGCACTGGCTCTG
AGTTTAGGTTTGGCGTTATCTCCGCTCTCTGCAACGGCGGCTGAGACTTCGGATCCCGCTCGAGTATTACCGTTGCCGCCTGACGGCGG
CTGAGACTTCGGATCCTCTAGATTTAAGAAGGAGATATACATATG
>iaaA
GAGGCAATTCCGACGTCTAAGAAACCATTATTATCATGACATTAACCTATAAAAATAGGCGTATCACGAGGCCCTTTCGTCTTCACCTC
GAGGTTTCCTGGGCGGTCAGTGGGGTGACGCGAGAAAGCATCTCATTAAGCGCGGTGTCGAGCGACATCAATCCGGTGGTAAATTCCAT
GAAAACACTCCTGCGGAGGCAAAATCGAATTTGCCTATTATGTCAGAAAAACGCCACAGACTGTATGCCACCTCGGGCGTAGCGCTGGG
TCCTGCCTTTACATGCCATATCCATCTTTCTATATTCAAAAATTGAATGAGTAATTCATAAAAATTCTGATATTTATAGCAAAAGTGGC
GAACCACCCTTAATGGACGAATACTATGGGCAAAGCAGTCATTGCAATTCATGGTGGCGCAGGTGCAATTAGCCGCGCGCAGATGAGTC
TGCAACAGGAATTACGCTACATCGAGGCGGGATCCCGGATCCTCTAGATTTAAGAAGGAGATATACATATG
>yggD
GAGGCAATTCCGACGTCTAAGAAACCATTATTATCATGACATTAACCTATAAAAATAGGCGTATCACGAGGCCCTTTCGTCTTCACCTC
GATGGATCCCGGTATCCAACAGACGGCGAATGGGGTGCGGACGCAGACATTACTGATCGGCGGCGCGGACCAAACGTGTAATATAATAG
ACTCCCTGCATTGATGGTGATCTATCGGAAGACCAATGGCGACGCTGACAGAAGATGATGTGCTTGAGCAACTGGATGCACAGGACAAT
TTATTTTCATTTATGAAAACTGCGCATACCATTTTGCTCCAGGGGCTCGAGAGATCCTCTAGATTTAAGAAGGAGATATACATATG
>citT
GAGGCAATTCCGACGTCTAAGAAACCATTATTATCATGACATTAACCTATAAAAATAGGCGTATCACGAGGCCCTTTCGTCTTCACCTC
GATGGATCCATCTCCGGCAGTTCGACAGGGAGTGTATCGAACGAAATCTCAGTCCAGGCGGCAGTGCTGACCTACTGATCCTTACCTGG
TTTTTAGCACAGATTTAATTATTTAAGCACTTGATAAATTTGGAAATATTAATTTTCGGAGAACCCGTATGTCTTTAGCAAAAGATAAT
ATATGGAAACTATTGGCCCCACTGGTGGTGATGGGTGTCATGTTTCTTATCCCTGTCCCCGACTCGAGAGATCCTCTAGATTTAAGAAG
GAGATATACATATG
>ysgA
GAGGCAATTCCGACGTCTAAGAAACCATTATTATCATGACATTAACCTATAAAAATAGGCGTATCACGAGGCCCTTTCGTCTTCACCTC
GATGGATCCGGACGATGGCAAGCGTAGCCCCTTGTAAATCGTTTTTAGTGAGGCCGAGATGAAAAACATCAGACTTGGACATATACAAC
TCCTCTGTGAATCGGTTTAGTCAGAAGAAGCAAAAAGACACTTTACCGAAGGGTTTAACATTTTTTCGTGATACTCATCACCATGACGC
AAATGCGTTGCATAAATTTACCCACCAAAAGTGATGCAAATCACATAAATCACCCTTCATTCTGCAGGACTGCATAAAAGATGAGCCGA
TTCAGGCAATGTGATTTGTTGCATGCTGACTATAGTTAACATTGCGCAACATTTTTATAAGGGTACGGAGAATACCATGGCAACAACAC
AACAATCTGGATTTGCACCTGCTGCATCGCCTCTCGCTTCGACCATCGTTCAGACCCCGGACGACGCGATTGTGGCGGGCTTCACCTCT
ATCCCTTCACAAGGGGATAACATGCCTGCCTCGAGAGATCCTCTAGATTTAAGAAGGAGATATACATATG
>sstT
GAGGCAATTCCGACGTCTAAGAAACCATTATTATCATGACATTAACCTATAAAAATAGGCGTATCACGAGGCCCTTTCGTCTTCACCTC
GAGTGGTGTTTGGCATTCTGGTGATGACGTTTATTATCAACGCCTGGGTGAATTATCGGCATGATAAGCAGCGGGGTGGATAATTTTTA
ATCTGCCTAAGCCGTGTACCCTGTCATTAACATGAGCACCGTTTCTCCCTCTCCCTTCCAGGGAGAGGGTCGGGGTGAGGGTAATTTT

TCGCACCGATGCTGGCCTGTTCCCCTCACCCTAACCCTCTCCCCAAACGGGGCGAGGGGACTGACCGAGTCCTTTTTTGATGTTGTCAT
CAGTCTGGAAGCCGCACGTTGGCTTTATTTTTATGTCAAAGAAATGTAACCATTAAGTTTCAAAATATGACCTCTCTTTAAAATCCAGC
ATTTTTCGCTTCCCGAAGCTGTAACTTTCCTTATACTCGACCTTGCAAACACTTTGTTACATCCTGAAAGATGCGTCGACAGAACGCAC
CAGGGATGTGCGACAACACAATGAAAGGATCGAAAAATGACTACGCAACGTTCACCGGGGCTATTCCGGCGTCTGGCTCATGGCAGCCT
GGTAAAACAAATCCTGGTCGGCCTGGATCCTCTAGATTTAAGAAGGAGATATACATATG
>yiiF
GAGGCAATTCCGACGTCTAAGAAACCATTATTATCATGACATTAACCTATAAAAATAGGCGTATCACGAGGCCCTTTCGTCTTCACCTC
GAGAAAGCAAAGGGAGGCAGGCAAAAGGGGGAGGTAGTTGGTGTTGATGATCAGTGTAAGGAGCATAAGGAATGACAAAAATTTTTCAT
TCTTGAATATAAAAAACAGATGCCCTTATTCTGGTATTAATACAAGGCTGTTTTACTTGAACTTATAATAACTGCAACTGTTACATCAT
ATCTGGAAAACGCCTCGCAAAAATAAAAAATGATGCGTAAATGAGCTTTCTTTAAAGATTCGGCAAGGGTACTTTTCTCTCCATAGATT
CAAAAATGATATTGGCGAGATATTTATGAACTCTCTGGCAGGTATAGATATGGGCAGAATTTTACTCGATTTATCGAATGAGGTGATTA
AGCAACTTGATGATCTCGAGGTGCAGCGTAATCTTGGATCCTCTAGATTTAAGAAGGAGATATACATATG
>yiiQ
GAGGCAATTCCGACGTCTAAGAAACCATTATTATCATGACATTAACCTATAAAAATAGGCGTATCACGAGGCCCTTTCGTCTTCACCTC
GATGGATCCAGAAATCACGGCGTCCAATACGCCCTTTAAATGAGAATAACCATTGCTGTATGGTCATGTAAGTTCCTTGATGGTTGTCT
TTTCCAGGATTCTACCCTTTTGACAAGGGCGACAGATATCGTTTTAATCGGAGCCAGTCATAACAAAAGGTACTGTCAATGAAGCCAGG
GTGTACGCTGTTTTTTCTCTTATGTTCTGCATTAACCGTTACAACAGAGGCGCATGCACAAACACCAGATACGGCAACGACCCTCGAGA
GATCCTCTAGATTTAAGAAGGAGATATACATATG
>yiiP
GAGGCAATTCCGACGTCTAAGAAACCATTATTATCATGACATTAACCTATAAAAATAGGCGTATCACGAGGCCCTTTCGTCTTCACCTC
GAGGAGCAGTTGCGTGACGTGACGCAATGGCAAAAAAGTTCATCGTTGAAGCTATTGAGTAGTAGCAACTCACGTTCCCAGTAGTAAAC
CCTGTTTTCCTTGCCATAGACACCATCCCTGTCTTCCCCCACATGCTGTGGGGGTTTTTTTTATCCTCAATTTGCCTGCTGCTTAATGC
ATTGCAGATGATTTGCTTCCGTTATACTAGCGTCAGTTGATAGCGGGAGTATTTATGAATCAATCTTATGGACGGCTGGTCAGTCGGGC
GGCGATTGCTGCGACGGCGATGGCTTCGCTGCTATTGCTGGATCCTCTAGATTTAAGAAGGAGATATACATATG
>citC
GAGGCAATTCCGACGTCTAAGAAACCATTATTATCATGACATTAACCTATAAAAATAGGCGTATCACGAGGCCCTTTCGTCTTCACCTC
GATGGATCCCAGCGGAAATGCCAGTCTTTGGAAAAATGCAAACTGTTTATTCTCGTTAAGCTGCAACATTGTTTTCCTTGCGGGTTGTC
CCGAGCCTCTTTGACGGATGTAATTAAAATGTTGAATCAAAATACATAGATATTAATTATGATCGTGGCAAATTTTATTCATTAAATCA
ATTAAATCACTTAAACCTGCGAGTCATTTCGTAATTACATTAATTAAAAAACTTAAAATCATTAAATAAATAAAAAACCACTAACTCTA
TGTGAAATAAATCAAAATTTCACGCCGAAATACTCCTTAGGATGTATAGCGAAAGGAGAAAAAGATATACCTCGATCACCCCCTTTCTC
CCAAGTGAAAATAAAAGGTTATCAGTTTGCAACATTGAACAACATTCGTTGCAAATCGATAACAACATGCACCTTCAGGACTCGAGAGA
TCCTCTAGATTTAAGAAGGAGATATACATATG
>yiiU
GAGGCAATTCCGACGTCTAAGAAACCATTATTATCATGACATTAACCTATAAAAATAGGCGTATCACGAGGCCCTTTCGTCTTCACCTC
GAGTAGTGCTGCAACGCAACCCACACCGAAGAAAATCAACAACCCGGTACCGAGGAATTCAGCAATGCACTGGCCTTTCAAGGTTGATG
TTTGACTCATAATCGGATCCTGAAGAGTTAATGTTTGTTGTATGCGTGAAAGTCACGGACCTCCACGATGCTTGTAGGCATGCTGTAAA
CTTATCGTTAACGAGCAAAAACGAGAAATATCGAACTTAAAATGTGTGTGCCTCGTCATAAAATGAGCGTTATCGCGCCATTTATTCCA
TCTGATTGTACATGATGCATGTTTGTTAGATCATTCGCGCAACAAATTTATTAACATTTTCGATTGATGTGGATTATGTCACCAGGAGA
AGGATGAAAAGTGTGGCAAACCGTAATCTGCAAGGCGTGCCGCTGGACAGCGATGGCGCGGCTTCATACAATCGGAGCTAACTAAAGTG
CGCTCGTATTTATTAAGGCGTCACCGGTAATCGGGACGAGGATTTTTATCCATCAACGCCTTGCAATTCAGGAGAGGTATGACAATGTC
ATTAGAAGTGTTTGAGAAACTGGAAGCAAAAGTACAGCAGGCGAGGATCCTCTAGATTTAAGAAGGAGATATACATATG
>tyrV
GAGGCAATTCCGACGTCTAAGAAACCATTATTATCATGACATTAACCTATAAAAATAGGCGTATCACGAGGCCCTTTCGTCTTCACCTC
GATGGATCCGGCCAAAGGGAGCAGACTGTAAATCTGCCGTCATCGACTTCGAAGGTTCGAATCCTTCCCCCACCACCTCGAGAGATCCT
CTAGATTTAAGAAGGAGATATACATATG
>degQ
GAGGCAATTCCGACGTCTAAGAAACCATTATTATCATGACATTAACCTATAAAAATAGGCGTATCACGAGGCCCTTTCGTCTTCACCTC
GAGAAGCCAGCAACGATCAGGCACCGGTGCAGATGCCTCGCGACTATTCTGAAGGCGCATCCGGCCTGCTGCGTACTGGCGCGAAGCGC
GACTAATTTATTTTTCGGGCGCAGCCATTGCGCCCTCCTCTTCTCTCCCTCCCCGACTATCATTTAATCTGGTGTCTCATTGTTAGCCG
TCTGAAAATTCAATAACATCAAACTGTTTTGAATCTCTTTTCTTATCATTCAGGTACGAGAGCAGGAATAATGAAAAAACAAACCCAGC
TGTTGAGTGCATTAGCGTTAAGTGTCGGGTTAACTCTCTCGGCGTCATTTCAGGGGATCCTCTAGATTTAAGAAGGAGATATACATATG
>yphH
GAGGCAATTCCGACGTCTAAGAAACCATTATTATCATGACATTAACCTATAAAAATAGGCGTATCACGAGGCCCTTTCGTCTTCACGTC
GACGGAGTCATGGTGCCCTCAATATTAACAAGACATACTGAATTAAAAGATTTGTGGCAGTGTATTGAACAATCTGGCAATGTTTTCGC
GGAATAATCACGCAATTAACTAAACAAGGTTTAGTGAAGATGAGAGCCTGCATTAATAATCAACAGATTCGCCACCATAACAAATGCGT
GATTCTGGAACTGCTGTACCGGCAAAAGCGCGCCAATAAATCAGGATCCTCTAGATTTAAGAAGGAGATATACATATG
>hybO
GAGGCAATTCCGACGTCTAAGAAACCATTATTATCATGACATTAACCTATAAAAATAGGCGTATCACGAGGCCCTTTCGTCTTCACCTC
GATGGATCCTTTATGGCCGGTTATCGCCTCGAAAATTGATCAAACATACGTATTATCTTGCTTTAATTAATTACACTAATGCTTCTTCC
CTTCGTTTTAGCGCCCCGCCGCAGTATCATGATATCGATAACCATAATAAATGTGTGGTAAATGGCGCATCGATCGCATTATTGATTTT
GCGATTGAGGCAAAATATATGCCAGGTCTTCGCAACGGAATAACTATAAATGACTGGAGATAACACCCTCATCCATTCTCACGGCATTA
ACCGTCGTGATTTCATGAAGCTTTGTGCAGCATTAGCCGCCACCATGGGGTTAAGTAGCAAAGCCGCTGCAGAGATGGCCGAATCGGTT

ACTAACCCGCAGCGTCCGCCAGTTATCTGGATTGGCGCGCAGGAGTGCACCGGTTGTACGGAATCTCTGCTTCGTGCAACGCTCGAGAG
ATCCTCTAGATTTAAGAAGGAGATATACATATG
>yphF
GAGGCAATTCCGACGTCTAAGAAACCATTATTATCATGACATTAACCTATAAAAATAGGCGTATCACGAGGCCCTTTCGTCTTCACCTC
GATGGATCCGCGGCATTATTCACCACCGTGATGCCTTTTATTTTTAACCGCGTTCACTAAACCCTATTAACTGTAATATTTGAGCGGCA
CGCGCTGCCGCTCATCACAAAATAATTAGAATAAGCCGTTGTTAATAAAAACCCTACAGGAGGCTCTATGCCTACAAAAATGAGAACAA
CCCGTAATCTATTATTAATGGCTACGCTATTAGGTAGCGCACTATTTGCCAGGGCTGCGGAAAAAGAACTCGAGAGATCCTCTAGATTT
AAGAAGGAGATATACATATG
>accB
GAGGCAATTCCGACGTCTAAGAAACCATTATTATCATGACATTAACCTATAAAAATAGGCGTATCACGAGGCCCTTTCGTCTTCACCTC
GAGACGGGAAAAGCGGTTAGCCGGACAGCTTTCGTCGAACACCTTGGTCAGGAAGTTATACAACATCATCCAAACTGGAAAGTCATGAT
TTCGACTGACCACAAATTGATGCGCATTGATACTCCACTAAACAGCTATTATTGATACGCCTCCGTCGCCTGTTAGGTTTATGTTGCTT
TGCCTGGGCGACGCTACGCTTAGCCCCTTACTTATTTCTGGTACCATGGGGTGAATAATCTGATTTTGTTTGACTACAAATTAATCACT
CGAACCTATTTAATGCTGAGCATTGTCAATCGGTTAATTTTGCGTGCTTTAGCATTCACATCTATCCAGACGATGCAGTGAAAATTGGG
TAATCCCCAGCAACCGCTGCGTAATGTCGTCTATCTTGTCGCGATCCTGGCATCCCTACATTATTTGTGGTCTGTGAAGATTATCTCAT
TGCAGCCCCTCATCTTCGCAGGGCTGGCTTTTCAGCTTTTCACCTTACGTTATAAGAAGTTCCGTCGATGATGGCGCTAATTTCGTGAA
TTGTGCGGCTTGTTGCAAATTACACGGTGTTGAAGGTTATTTACATGTTAGCTGTTGATTATCTTCCCTGATAAGACCAGTATTTAGCT
GCCAATTGCTACGAAATCGTTATAATGTGCGACCTCGTCCTCCCTGACGCAGTTTTTGCGCTGCGGAAAAGGTGACATTGGCGCAACGA
AGGTATATTTTGTTTTTTGCCGGAGGATAGCAGCAGATCGCTGCACAATGTCCGTCAAGTCTAACATTGACACTCTGGGGCAAAATAGA
CCGGCGTCCCGGCCTGCTGGAATTTATCGCTATGCATACAGCTGTCGGGGCATACGCTTTACAGACGGCGGTGAAACGCCTGTCACAAT
CACACTAAACAAAGAGTACGGAACCCACTCATGGATATTCGTAAGATTAAAAAACTGATCGAGCTGGTTGAAGAATCAGGCATCTCCGA
ACTGGAAATTTCTGAAGGCGAAGAGTCAGTACGCAGGATCCTCTAGATTTAAGAAGGAGATATACATATG
>iap
GAGGCAATTCCGACGTCTAAGAAACCATTATTATCATGACATTAACCTATAAAAATAGGCGTATCACGAGGCCCTTTCGTCTTCACCTC
GAGTTTGAGAATTCTGCCGCCACCTCGCGAATAATGTGGATGCTTTCCGCCTCCAGTTGCCGCAGGTGAGTAAGTCGTATTTGATCCAT
AACCGTTCCTTTGCAATACCGCTATTTTCTTGCCATCAGATGTTTCGACTATAGGGAGCGTAAGAGAACGAATGAAATTACCAATTAGA
ATGAGTAGTTCCTTAACGGAATAACGATTTGGCAAAGCTAATATCAAAAAGTGCTTAAGGCACCGGATTTCGGGCGTTTAGGAAGATTT
GAAATTGTTTTAGCGCAGCGGCAGTTTCATACTATGGCGGTAAAAAAATTTGCATGGTATTTAAGGACTCACTATGTTTTCCGCATTGC
GCCACCGTACCGCTGCCCTGGCGCTCGGCGTATGCTTTATTCTCCCCGTACACGCCTCGTCACCTAAACCTGGCGGGATCCTCTAGATT
TAAGAAGGAGATATACATATG
>smpA
GAGGCAATTCCGACGTCTAAGAAACCATTATTATCATGACATTAACCTATAAAAATAGGCGTATCACGAGGCCCTTTCGTCTTCACCTC
GAGGCGGTTACAAGAGCTGGCGCGCCTGCTTGGTGGCAGTGAAGTCACACGTAATACACTGGCGAATGCGAAAGAACTGCTTGCAGCGT
AAACTTTTTTCCTGCTTCACGGTCAGAGTAAACAGCAAAACGCCGTAAGACCGGAAAGCAAAAGGTTTTAAAGTGATGAAAGGTCTATT
ATCATCGGCATATTACAGATGAGCCACGTACTGCTCGGGCCCGAAAAGGAATCAAATCACTATGCGCTGTAAAACGCTGACTGCTGCAG
CAGCAGTACTATTGATGTTGACCGCAGGCTGTTCCACTCTGGAGCGAGTGGTTTACCGTCCTGACATCAACCAGGGGAACTATCTGACC
GCTAACGACGTATCCAAAATACGTGTTGGCATGACGCAACAACAAGTTGCGTACGCATTGGGTACACCGCTGATGTCCGATCCATTTGG
TACGAATACCTGGTTCTATGTCTTCCGCCAGCAACGGATCCTCTAGATTTAAGAAGGAGATATACATATG
>smpB
GAGGCAATTCCGACGTCTAAGAAACCATTATTATCATGACATTAACCTATAAAAATAGGCGTATCACGAGGCCCTTTCGTCTTCACCTC
GAGCGCGCTGTAGGGTACCAGTGCGGTCCGGCTAATCTGAGGCATAACAATTTCCATCAACAAAAATCCAACAAATAATATCATTTATC
CTGCTGTAAAAAAAACGCTATCCCGGCGCTGGGTAACATCGGGTTCATGCTAAGATAGAGCCTTGTCCCCCGCAGGATTGATATGGGGT
GTTTTCGATTTCAGATTACCGATGATTCACGACGCTTATGACGAAGAAAAAAGCACATAAACCTGGTTCAGCGACCATCGCGCTTAACA
AGCGCGCCCGTCACGAATACTTGGATCCTCTAGATTTAAGAAGGAGATATACATATG
>serU
GAGGCAATTCCGACGTCTAAGAAACCATTATTATCATGACATTAACCTATAAAAATAGGCGTATCACGAGGCCCTTTCGTCTTCACCTC
GATGGATCCCAGGGAAGGGCAGTTTGATGTGCTGATTCTTGTACTTTCCAGGGCCACTTAATCATCGTTTTGCTCGCAAACTCGTCACT
TGAACAAAATTGCACGGACAGGGACTGTTAAAATGCCAAATTTCCTGGCATCATGGCAACCATCTGAACGGAGAGATGCCGGAGCGGCT
GAACGGACCGGTCTCGAAAACCGGAGTAGGGGCAACTCTACCGGGGGTTCAAATCCCCCTCTCTCCGCCACTTTCTCGAGAGATCCTCT
AGATTTAAGAAGGAGATATACATATG
>pnp
GAGGCAATTCCGACGTCTAAGAAACCATTATTATCATGACATTAACCTATAAAAATAGGCGTATCACGAGGCCCTTTCGTCTTCACCTC
GATGGATCCCGCATGGTTTCTCAGCGTCGTAAACTGCTCGACTACCTGAAACGTAAAGACGTAGCACGTTACACCCAGCTCATCGAGCG
CCTGGGTCTGCGTCGCTAATTCTTGCGAGTTTCAGAAAAGGGGGCCTGAGTGGCCCCTTTTTTCAAGCTGACGGCAGCAATTCACTGGA
AACTAATGTATTGTTGCTATGAATGATCTTCCGTTGCAGAGGTTCGCGCGGCTAATGAGAGGCTTTACCCACATAGAGCTGGGTTAGGG
TTGTCATTAGTCGCGAGGATGCGCAGAAGATCGGGTATTAACACCAGTGCCGTAAGGTACTGTCTAAGAAAGAGAAAGGATATTACATT
GCTTAATCCGATCGTTCGCTCGAGAGATCCTCTAGATTTAAGAAGGAGATATACATATG
>serB
GAGGCAATTCCGACGTCTAAGAAACCATTATTATCATGACATTAACCTATAAAAATAGGCGTATCACGAGGCCCTTTCGTCTTCACCTC
GAGCTGCATCAGCGCCACTAACAAGGCAAGACAGAACAGGACAATCACTGCCCGATGCAGCCGGAATTTCAGTTTTGTGCGAGCCATCT
TCCACCCTTTGAAAATTTGAGACTTAATGTTGCCAGAAGCAATGGATACAAGGTAGCCTCATGCGTTATTTTCCCTGCTTCGAACGATT
TTACAGGAGCCTTAATGCCTAACATTACCTGGTGCGACCTGCCTGAAGATGTCTCTTTATGGCCGGGTCTGCCTCTTTCATTAAGTGGT
GATGAAGTGATGCCAGGATCCTCTAGATTTAAGAAGGAGATATACATATG

Data Set S4 - MG1655 library reference file.fasta
>serC
GAGGCAATTCCGACGTCTAAGAAACCATTATTATCATGACATTAACCTATAAAAATAGGCGTATCACGAGGCCCTTTCGTCTTCACCTC
GAGTCATATGAAAGCGGGGGAAAAACAATTATGTCCGCGCTGTGCAAATCCAGAATGGACGAAGGCAAGTCGGGCAAAACGGGTGACCT
GACAGTAAAAACATCGGCTTTTTGCTAATAATCCGAGAGATTCTTTTGTGTGATGCAAGCCACATTTTTGCCCTCAACGGTTTTACTCA
TTGCGATGTGTGTCACTGAATGATAAAACCGATAGCCACAGGAATAATGTATTACCTGTGGTCGCAATCGATTGACCGCGGGTTAATAG
CAACGCAACGTGGTGAGGGGAAATGGCTCAAATCTTCAATTTTAGTTCTGGTCCGGCAATGCTACCGGCAGAGGTGCTTAAACAGGCTC
AACAGGAACTGCGCGACTGGAACGGTCTTGGGGATCCTCTAGATTTAAGAAGGAGATATACATATG
>serA
GAGGCAATTCCGACGTCTAAGAAACCATTATTATCATGACATTAACCTATAAAAATAGGCGTATCACGAGGCCCTTTCGTCTTCACCTC
GATGGATCCATTGGCACACCTGACGGTGTCAAAACCATTGTGAAATGATCTGACGGGGGAACCTCCCCCGTTAAAAAAATTCTCTTCAT
TAAATTTGGTGACATGTGTCACGCTTTTACCAGGCAATTGTCGATTGCTCTAAATAAATCCTCTAAACCAGCATATTCATCCAAGAATT
ACCTTTGCGTGATATTTCCTCAACATCGCGACGCAAACGTTCATATTGCCGCAATATTATTTTTGATATGTTGAAAGGCGGATGCAAA
TCCGCACACAACATTTCAAAAGACAGGATTGGGTAAATGGCAAAGGTATCGCTGGAGAAAGACAAGATTAAGTTTCTGCTGGTAGAAGG
CGTGCACCAAAAGGCGCTGGAAAGCCTTCGTGCACTCGAGAGATCCTCTAGATTTAAGAAGGAGATATACATATG
>alaS
GAGGCAATTCCGACGTCTAAGAAACCATTATTATCATGACATTAACCTATAAAAATAGGCGTATCACGAGGCCCTTTCGTCTTCACCTC
GATGGATCCGCGAAATTTTGCCGACTGAACGCATACGGTATTTTACTTCCCAGTCAAGAAAACTTATCTTATTCCCACTTTTCAGTTAC
CAGCCCGGCGGTTAAGACACGCTGGAGCTGGTGGCGATATTTCGTTAGCTTGATTTCAGGATAATTATGAGCAAGAGCACCGCTGAGAT
CCGTCAGGCGTTTCTCGACTTTTTCCATAGTAAGGGACATCAGGTAGTTGCCAGCAGCTCCCTGGTACCCCATAACTCGAGAGATCCTC
TAGATTTAAGAAGGAGATATACATATG
>yqhH
GAGGCAATTCCGACGTCTAAGAAACCATTATTATCATGACATTAACCTATAAAAATAGGCGTATCACGAGGCCCTTTCGTCTTCACCTC
GAGTGATTTTATCCGCCCGAACAGCGGCAGCCTACTTAAGCAGATTGAGGCAAAAACTGCCGCCAGATTAAAGCAATGAGCTGAATTAA
ATAACAATTAGCCGGAACAATAAATAAAAGGGAACACTATATGAAAACGATTTTCACCGTGGGAGCTGTTGTTCTGGCAACCTGCTTGC
TCAGTGGCTGCGTCAATGAGCAAAAGGTCAATCAGCTGGCGGGATCCTCTAGATTTAAGAAGGAGATATACATATG
>rrmA
GAGGCAATTCCGACGTCTAAGAAACCATTATTATCATGACATTAACCTATAAAAATAGGCGTATCACGAGGCCCTTTCGTCTTCACCTC
GATGGATCCTTCAGGACGGCCAGAAAGGTCCGGCAGCTGTTAACGTAACAGCTATCTGATCGAATCCACTGATCTGAAGTGTGAATACG
CTTCAATCTCGCTATAAAGCCTCGTCGAATGCGAGGCTTTTTACTATGCTTTATCTTCGCTCCTGGCGTTCGGATATTTGCCCGCCGCG
TGATTCGCGTTACACTTGCGGCCTTTAGTATCCTGCCGGAGTTGTCATGTCTTTTTCCTGTCCACTTTGCCATCAGCCTCTTTCGCGTG
AAAAAAACAGCTATATCTGTCCCCAGCGACATCAGTTCTCGAGAGATCCTCTAGATTTAAGAAGGAGATATACATATG
>aroL
GAGGCAATTCCGACGTCTAAGAAACCATTATTATCATGACATTAACCTATAAAAATAGGCGTATCACGAGGCCCTTTCGTCTTCACCTC
GAGGGCGGACCAGATAGCCTTTCACAACGTGACCGCCAGGCCTTTGCCGCGGAGCTGGAGAAGTGGTGGCTGGAAGTGCAACGTAGTCG
TGGCTAAATGTAATTTATTATTTACACTTCATTCTTGAATATTTATTGGTATAGTAAGGGGTGTATTGAGATTTTCACTTTAAGTGGAA
TTTTTTCTTTACAATCGAAATTGTACTAGTTTGATGGTATGATCGCTATTCTCATGACACCGGCTTTCGCCGCATTGCGACCTATTGGG
GAAAACCCACGATGACACAACCTCTTTTTCTGATCGGGCCTCGGGGCTGTGGTAAAACAACGGTCGGAATGGCCCTTGCCGATTCGCTT
AACCGGATCCTCTAGATTTAAGAAGGAGATATACATATG
>yjeB
GAGGCAATTCCGACGTCTAAGAAACCATTATTATCATGACATTAACCTATAAAAATAGGCGTATCACGAGGCCCTTTCGTCTTCACCTC
GAGGCTGACTGGTGTGCCGATCGATATCATCTCTACCGGTCCGGATCGTACTGAAACCATGATTCTGCGCGACCCGTTCGACGCGTAAT
TCTGGTACGCCTGGCAGATATTTTGCCTGCCGGGCGAACAGTGTGATACATTGCTGTGTCGGGTAAGCCATTACGCTATCCGACACAGT
GTTAAATCCTCGCTTTTTTCCTTCCCCGAACTGAAATAAATTAGCGACACAGCTTGTGGCTGGTTTATCATCAATATAAATGTATTTTT
TCCCGATTTCCCTTTTGAGGTTGATGTGCAGTTAACGAGTTTCACTGATTACGGATTACGTGCGCTGATCTACATGGCGTCATTGCCAG
AAGGGCGGATGACCAGTAGGATCCTCTAGATTTAAGAAGGAGATATACATATG
>sulA
GAGGCAATTCCGACGTCTAAGAAACCATTATTATCATGACATTAACCTATAAAAATAGGCGTATCACGAGGCCCTTTCGTCTTCACCTC
GATGGATCCCCCAACGTTGCCAGGTATTCTTGTGATTTATAGATCCGCTTATAGGAGAGGCTTTTCATAAAATTCCTTTTAAAATCATA
ACATAAAAGAATGATTCACATTAACGGATCCGTTAACTACGAAAATAGGCAACTTATTCTTAAGGGGCAAGATTAATTTATGTTTTCCC
GTCACCAACGACAAAATTTGCGAGGCTCTTTCCGAAAATAGGGTTGATCTTTGTTGTCACTGGATGTACTGTACATCCATACAGTAACT
CACAGGGGCTGGATTGATTATGTACACTTCAGGCTATGCACATCGTTCTTCGTCGTTCTCATCCGCAGCAAGTAAAATTGCGCGTGTCT
CTACGGAAAACACTACAGCCGGGCTCTCGAGAGATCCTCTAGATTTAAGAAGGAGATATACATATG
>gyrB
GAGGCAATTCCGACGTCTAAGAAACCATTATTATCATGACATTAACCTATAAAAATAGGCGTATCACGAGGCCCTTTCGTCTTCACCTC
GATGGATCCTTGTCAGCGCGATCAGTGCTGAACACGTTATAGACATGTCGGACGAAAATTCGAAGATGTTTACCGTGGAAAAGGGTAAA
ATAACGGATTAACCCAAGTATAAATGAGCGAGAAACGTTGATGTCGAATTCTTATGACTCCTCCAGTATCAAAGTCCTGAAAGGGCTGG
ATGCGGTGCGTAAGCGCCCGGGTATGTACTCGAGAGATCCTCTAGATTTAAGAAGGAGATATACATATG
>yhiR
GAGGCAATTCCGACGTCTAAGAAACCATTATTATCATGACATTAACCTATAAAAATAGGCGTATCACGAGGCCCTTTCGTCTTCACCTC
GAGGCCTTAGTCACGGCTGGAACGACATGTTCCGGGAGAATTTTAGAAAACGGAGGCAATTCAAAGGGAGTCAGTAACGGATTCGTCAT
TCGCGCAGTCCTGGTTAAAGAGGTTAAGGAAGCGCTCAACAGGCGCTTTACATAATGTGTGTAGCATGGGGTTAAGTGTAGTGAATTTC
AATGAGAAACGTTACGCTTTCGCGGCGGCGGCCTCTTTTCGGTATACTGTCCTGATACGCTTTTGTGCGCCCCGAATACGGGCCGATTT

TTACTTACCGGAACACCTTTACCCATGCTCAGTTATCGCCACAGCTTTCACGCTGGCAACCACGCCGACGTCCTTAAACATACCGTTCA
GAGCCTGATCATCGAGTCGCTGGGATCCCGCCTGATCATCGAGTCGCTGGGATCCTCTAGATTTAAGAAGGAGATATACATATG
>sohB
GAGGCAATTCCGACGTCTAAGAAACCATTATTATCATGACATTAACCTATAAAAATAGGCGTATCACGAGGCCCTTTCGTCTTCACCTC
GAGATACGTCATCGCGGCTTCACGACCAATACCATCGCTGGCTCCCGTCACCAGGATAATGCGATCATTGAGTAAATCTTGTTTTGGCT
GGTAATGCATGGCTACTCCTCAACGACGTTGTCTGTCGCGCTGCCAGTGTACTTTATGACTTTGGGGCTTTATGCCTGAAACAGAAGGG
CATTTCAATCAGCCTGGGGCGATAACACCGCAAAATGGATACTTTGTCATACTTTCGCTGCAATAACATCTCTGCGAGACGGCTTAACA
TGCCTGTTGTAAACTGTGAGCCAAAGCGTTGTTTAACCAAGGTGGGGACTCGTGGAATTGTTGTCTGAATATGGTTTGTTTTTGGCGAA
AATCGTTACCGTTGTGCTAGCGATTGCGGCGATTGCCGCCATTATTGTCAGGATCCTCTAGATTTAAGAAGGAGATATACATATG
>sohA
GAGGCAATTCCGACGTCTAAGAAACCATTATTATCATGACATTAACCTATAAAAATAGGCGTATCACGAGGCCCTTTCGTCTTCACCTC
GAGGCCAGCGGGAAGAAGAAAACCTTCTCGGATCAATGGGGGCTGCATAACCAGCTGGCGGTGTTTAACCCGGCACCGGTGACCTGATT
TCTTTTGGAACGTCCCTCGCAAAACATGGCCTTAGTGCCATGTTTTTATTGTTTAAAGCCCCCACGTCCATTAATAATGCATTTGCATT
ACCTTAGTTCAAGCTTATAATTTGAGCAGAAAACAGGCTGTAAAAGGACAGTGAATCATGCCCGCTAATGCTCGCTCTCACGCTGTACT
GACCACTGAATCAAAGGTCACGATACGCGGACAAACAACTATCCCCGCGGGATCCTCTAGATTTAAGAAGGAGATATACATATG
>brnQ
GAGGCAATTCCGACGTCTAAGAAACCATTATTATCATGACATTAACCTATAAAAATAGGCGTATCACGAGGCCCTTTCGTCTTCACCTC
GAGTCACGAAAGTCGCCTGAATATTGAGAGTACAGTAGGAAAAGGAACACGTTTCAGTTTTGTTATCCCGGAACGTTTAATTGCCAAAA
ACAGCGATTAATCCGCCTTTGTCATCTTTTATTGCCATAAGCCAGTCGATGCTGGCTTATTTTCTTTGCAGTCAAAATACGGGCGTTAG
ATTTTACAACGATTGGTGATTTTTTGTTCGCATGATTAGCCATGTCTTTTTCACGGAAATAGTGTTTTATACTGGTTGGTGATTTCTTA
TCGCTATATACCTCTGGTTTTTAGATCCCTCCTTGCTTTAAAACGTTATAAGCGTTTAAATTGCGCTTCAGGTGCTGTCATACTGACTG
CATTAACGCGGTAAATCGAAAAACTATTCTTCGCCGCGCCTGGTTGGGAGTATTTCCCGCTAAAATTGTTTAAATATACCGCTGTATCA
TCCCCAGGGATTGGCACAAAAATTTAACGTTACAACACCACATCCACAGGCAGTATGATTTATGACCCATCAATTAAGATCGCGCGATA
TCATCGCTCTGGGCTTTATGACATTTGCGTTGTTCGTCGGCGCAGGTAACATTGGATCCTCTAGATTTAAGAAGGAGATATACATATG
>betT
GAGGCAATTCCGACGTCTAAGAAACCATTATTATCATGACATTAACCTATAAAAATAGGCGTATCACGAGGCCCTTTCGTCTTCACCTC
GAGATCGTGCATGCCCACTTCATTTATTGCTTCCAGTGTGGCGTCGATCAGTTGTCTGCGCCGGATCGACTGCATCCCCAATTTGGGCA
TTTTCGCCACTCCATTCATCAGCGGTGTTTATCTATTAAAGCGGTTATTGATTGGACGTTCAATATAAAATGTGTCTTAATTGTTACGA
ATTTGATTTTAAATAGTAACAATAACAGTGGGGATACTGGATGACAGACCTTTCACACAGCAGGGAAAAGGACAAAATCAATCCGGTGG
TGTTTTACACCTCCGCCGGACTGATTTTGTTGGATCCCGGGATCCTCTAGATTTAAGAAGGAGATATACATATG
>mrp
GAGGCAATTCCGACGTCTAAGAAACCATTATTATCATGACATTAACCTATAAAAATAGGCGTATCACGAGGCCCTTTCGTCTTCACCTC
GATGGATCCTATGGCCGAGGTGGATTGAGCCGTTAGCGTACGGCAGTGCGCACGTCACCAGAATTTTCTTCGCGACTTGAGTCATAGTA
GGCATTACTTCTTAATAGTGAAAAGGGGTTAGATAGTACCAAATGGGAAAATGTTAAGTAAGTACTTCTCTATATTCTTCAGGTAAACT
ATCAGCTGATGGTTTATCTATACAAAAAAGGAGTCGGGATGAACGAACAATCCCAGGCCAAATCACCAGAAGCCTTGCGCTCGAGAGAT
CCTCTAGATTTAAGAAGGAGATATACATATG
>mrr
GAGGCAATTCCGACGTCTAAGAAACCATTATTATCATGACATTAACCTATAAAAATAGGCGTATCACGAGGCCCTTTCGTCTTCACCTC
GAGCGCCCTTCAGGAATTCAAAATTGGATTTATTCATCATTGTTATTAATCCATTGCTGTGCGGGCCTGTCCAAATATTTAAGGCCCAT
AACATCTCATCTTAGCTTTCTGTACCTTTCCGGGCAATGACCACGGTCACAGCAACTGACTCATTTCTAACGTGTTCGTCTATTTTTGT
AGTGCTATAGTAGCCGAAAAACATCTACCTGATTCTGCAAGGATGTACTATGACGGTTCCTACCTATGACAAATTTATTGAACCTGTTC
TGCGTTATCTGGCAACAAAACCGGAAGGTGCAGGATCCTCTAGATTTAAGAAGGAGATATACATATG
>ykgC
GAGGCAATTCCGACGTCTAAGAAACCATTATTATCATGACATTAACCTATAAAAATAGGCGTATCACGAGGCCCTTTCGTCTTCACCTC
GATGGATCCTGTGGAAGCTGCCAGTCACTTCCTAACACGCAATTCTTATCGATCGTTCCTTGTGGAGCGTTAAGCATCAACAAACGGCT
AAGGGCATCCATATCAGACTCCTGAGTGGTTTTTGATTACAAACACCAACAATCTTAGTAAATAACTAATCTAATACCATCAAATAGTT
AATCATGATTTTTGTTTTGTGATTCTGATTAAGAAAAAGAAAATATCAGACGATAAGACAAAATATCAAGCGTTGTGAAGAAATGTTAT
TTGCTCTTTTGCGTCTATAGTCATGATGTCAAATGAACGCGTTTCGACAGGAAATCATCATGAATAAATATCAGGCAGTGATTATTGGT
TTTGGCAAGGCTGGAAAAACACTCGAGAGATCCTCTAGATTTAAGAAGGAGATATACATATG
>ykgE
GAGGCAATTCCGACGTCTAAGAAACCATTATTATCATGACATTAACCTATAAAAATAGGCGTATCACGAGGCCCTTTCGTCTTCACCTC
GAGCACAAGGCGTTTGTCCGCGAGTTTGGTTGTACCCCGGGAGAATATCGGGAAAGGGTCAGACAGCTTGCACCCTGAATAAAACCGCC
AGAAATCAGGGCAAAGATAATCCGCATTCCGGGAGTTGTGAGCCTTTCAACTATTTCTATTTCCAACGGTGGTTTGGGCTTTATATATT
TTTTCTGATGGACTCTAGCTCAGTTTTTAAATAAAAATGCAAACTAAATTGCTTCAATTAAATAAAATCCCGACTATTACTTGATACGT
GATTATTGTCGGTGATTTTTTTGTACACCATCAGTGTATATCTCAATAACCCCTGAATAAGTAGCTCTGAATAGGTATAAGGGATGTAG
CCATTTTTAAATGGATTTCTTATGCCAAAAATGATCGTCGATTTCCCCATAAAATGTGAGCGATGCCGAAAGAAATAAAATTAGTTATT
GCATTTGACGTTTGGATGAAAGATTATCATTTGTCATACAAATGAGGGGTGGTATGTTGCTAGCCAATTAAAAAAGAACGCCATATTTA
TTGATGATTGATCCCCCGGTGGGGGGGAAACATCGGGCATAAATGGGCATGAAGTAATGGAGTATTAGTTGTGAATGTCAATTTCTTTG
TCACCTGTATTGGTGACGCCCTGAAATCAAGAATGGCACGAGACTCCGTGCTGCTACTGGAAAAACTCGGCTGTCGCGGATCCTCTAGA
TTTAAGAAGGAGATATACATATG
>ybfF
GAGGCAATTCCGACGTCTAAGAAACCATTATTATCATGACATTAACCTATAAAAATAGGCGTATCACGAGGCCCTTTCGTCTTCACCTC
GATGGATCCACGCCGTAAAATGTCGGATGCGCTCTCGCCGATATGCTTAGTGTGGCTGGCAATATAGCTGTAGAGTTCATCATCAACTT

```
CAATCGTTTTCATCTTAATCCAGTGCAGTGTCTTAGCTGAATACAAGTCGCCAGGAGTATATGAATAAATCCCGGCAGCGGATAGCGTC
AAGCCTGGCTTGCAGAAAAACTCGGAATAAACCTTATTTGTTGCAGCCCTGTGGCACAATGGTCAACATAATACCCTAACCCGACAACG
CGAAAAAAAGAACTTTGCCATGAAATTGAATATCCGCGCGCAAACTGCACAAAACCAGCACAATAATTCTCCCATCGTTCTTGTCCACG
GTCTGTTTGGCCTCGAGAGATCCTCTAGATTTAAGAAGGAGATATACATATG
>ybfE
GAGGCAATTCCGACGTCTAAGAAACCATTATTATCATGACATTAACCTATAAAAATAGGCGTATCACGAGGCCCTTTCGTCTTCACCTC
GATGGATCCATTACGCGCTATCCGTCGCTATCTCAATGATTAACGCAACTGATTAAAAACCCAGCGTCCGCGTCGCTTTCCGCGCGTGG
GCGTTGGCGTGACTCATCGCGCTGATGTATTATGGCGCGCTATCCATCCGGGCAGAGGCCTGGCTGATTGTTTCCCCCGAAGTCACCAA
GATCATGGCCAAAGAACAAACGGACCGTACGACACTCGAGAGATCCTCTAGATTTAAGAAGGAGATATACATATG
>ykgF
GAGGCAATTCCGACGTCTAAGAAACCATTATTATCATGACATTAACCTATAAAAATAGGCGTATCACGAGGCCCTTTCGTCTTCACCTC
GAGAGTGGGCGATTACAACGGGAAGGGCAGAAAGTCAAAGTGATGCATATTGCTGAAGTGTTGATGAGCCGCTGAGGATATAAAGATGT
CGATCAAAACCAGTAATACAGATTTTAAGACACGCATCCGTCAGCAAATTGAAGATCCGATCATGCGGGATCCCGGATCCTCTAGATTT
AAGAAGGAGATATACATATG
>ykgI
GAGGCAATTCCGACGTCTAAGAAACCATTATTATCATGACATTAACCTATAAAAATAGGCGTATCACGAGGCCCTTTCGTCTTCACCTC
GATGGATCCCGGGCTGCCTTATAGCATATTACGCGATCAGATATTTACTCATCCGTCGATGAGCGAATCACTCAATGATCTATTTTCAT
TAGTCAAATAAACTCAAAATCAGACGCCAGAACAAATATTCTGGCGTCTCAGAGAAAAGAATCTTATTAATTCTCTGTTACTCTATAAC
TTCTTAATCACTTCATTGATGGTATTTTATATGTTTAAAAAATCTGTTTTATTTGCAACACTATTATCTGGCGTTATGGCATTTTCCAC
CAATGCCTCGAGAGATCCTCTAGATTTAAGAAGGAGATATACATATG
>ykgH
GAGGCAATTCCGACGTCTAAGAAACCATTATTATCATGACATTAACCTATAAAAATAGGCGTATCACGAGGCCCTTTCGTCTTCACCTC
GATGGATCCGCGGGCTTTTTGCTGATATTTAACCCGAACATTCCTGATACAGACTATTAATACAACCTTATCTGCGACAGATTTTATTC
TCTGGATATATTATGCGTGAACAAATCAAACAGGATATCGATCTGATTGAGATTTTATTTTATCTGAAGAAAAAGATTCGTGTTATCCT
TTTTATTATGGCTATATGTATGGCTATGGTGCTGTTGTTTCTGTATATCAATAAAGACAATATAAAAGTGATTTACAGCCTAAAAATAA
ACCAGACAACGCCAGGTATACTTGTTAGCTGTGATAGCAATAATAATTTTGCCTGTCAGACTACAATGACTGAAGATGTTATTCAGCGA
ATTACTACATTTTTTCACACCAGCCCAGATGCTCGAGAGATCCTCTAGATTTAAGAAGGAGATATACATATG
>ykgJ
GAGGCAATTCCGACGTCTAAGAAACCATTATTATCATGACATTAACCTATAAAAATAGGCGTATCACGAGGCCCTTTCGTCTTCACCTC
GATGGATCCTCAGCCGGGGAATCACTATCTGGTCTATAACGACAAATTCATTAAGATTAGCGATTCCTGTCCGGCTAAGCCCCCTTCGG
CGGACTAAGACTTATCCATCGGTCGAGGGAAGAATTCCACCTCAGAGCCCCAAAAATCGTTTAAATGATGTGGAAGCGATCGTCAATAT
CACCTGTAACGATGGGCTTTCTCATATGCACACCATAGTGTTCAGGAGACACGGAAAGGTATCAGGCAGCCTGCTAACGAGTGGTTTAA
TCACGAATTAGTACGTAAAATCGGTAACGGCTGGAAATCATTCAATACTCACACTATCGAAAGTTCCCCAGCCAACCGCGGTACGTTCT
TACATACGATGTACCGCTGTTCTCTTTACGATTTTTAGCTGTACTGGTGAATTATGAGCAATCTGAATCCATGCATGACGTGTGGTGCC
TGTTGTGCATTTTTCCGCGTCTCTTTTTACTGGGCCGAAGCCGACGATGCTGGCGGTACTCTCGAGAGATCCTCTAGATTTAAGAAGGA
GATATACATATG
>ykgM
GAGGCAATTCCGACGTCTAAGAAACCATTATTATCATGACATTAACCTATAAAAATAGGCGTATCACGAGGCCCTTTCGTCTTCACCTC
GATGGATCCAGATATTTGCCCACGCCACGCAGCGGGCCAGAACTGAATAACGAAATCGTGGTTGTTTATGACCTGTTTTATAATGTGAC
ATTGAGATAATTTCCTGATGAACGAATTAACGAACATAGGCAGACACAAATAATGGCGTCTGCTTGTCAGAAAAACCGGGGATTTCCCG
ATTTCCTTTTCCGCAGAAAAGGAAATACCTTTATGTAAGAAAGTGCACCGCCACCTAATATATCCAGTGCCTATAGCCACTCATCAGGT
GGTTGGGCGCGGGATTATATTTTTATTTATGGGGGAAAATTTAGCTTTTTGTGCAAGTCATTAAATAGATACCTAAGTTATATTTCTTA
CTCTGCGGACAGTGAATAATTCACTATTCAGAGTGAATAACAATGACCTGCCTTTAATTAAAACTATTGGAAATACGTCGAAATAGGTA
TATATGCCAGGATAAACCAATATATTTTTTATGGATATTCATTTTCCTTCATCGGACTGTAAATTGCGGAGATATCGAACTTGCACTAC
CCATCGCAACACTCAAACAGATTTATGAATTTCGTTCATAACCCCTGACAAATTCTGCCGTCTAATCGAGCTTTTCCCTTTACGTTATG
CTTCTCATGAACGATAACACAACTTGTTCATGAATTAACCATTCCGGATAAACTATGGGATCAAATTCAGGCATCGAAAGGAACTGGCT
GGTTTCCGTTAATGGATTAAATATCTATTATTCTCCCGAGAGTAAAACTATCAAATCAATGAGTAACCGATACAGCCTCACAAATTCTT
TCAGATAATTAACCCGTCACTGCGGCACGTTCTCTCATAAGGCGTGCCTGCAGTGTAATCCAGTATGTATCCGGCATAGTTGACGGTTT
AATAGAACTCAATTCCTCTCCCAGGTCACCAACAACGATATCTTGTATAACACCCCAGGAGATACGACATTATGATGAGATGCGATATG
CTCCTCAGCACCAATCAACGCCCGGACTGGGCCAGTCTTGAGATCAGACTCATTCCCGGAACTGACTTCAGGCGACATATATCAATGAC
GTTGGTGCAATAAAAGATAGTTGATGTAATACATCTATTCAAATTTGTGATACATGTCAAAAAATTGGTTGACCAAACTCGTTATTTAT
ATAAGGGCACTTACGAAGTGCACTCTTTTTTAAAGCGAGGAAGTACCAATGAAAGAGAATAAAGTACAGCAAATCAGTCATAAACTGAT
TAATATCGTTGTTTTTGTCGCAATTGTAGAATACGCCTATTTATTTCTCCATTTCTATTAATAACGGAAATAAACTGTTCACTTCAGTG
ATATTTAAAATATGCATCCTCTCCCTTTTTTGTAAGTAATTATTATATCCGTGGGAGAGGAATACACGCTGTCAGGTAATCAATCATAC
TGCGATAAATTATCGGCCAGTAAAGTGGAGATAACCTCCATTCTCGAAAAATCCATACTCTCAGCAAAACCATCATCAATCACTCATCC
AGGCGTTTATGGGAGCGTCACCAATGGCTGCTAACAATGCCAGAGTTCCCCGTTGCGAAAATTCCACATCCACAAAGAGTCACAGGGAT
TGAGTGTTGAAATGATCCGGATGAGCATGTATCTTTATGGTTATGTTATAACATAACAGGTAAAAATGATGAAGCCCAATATCCATCCT
GAGTATCGTACTGTGGTGTTCCACGACACCAGTGTTGATGAGTACTTTAAAATCGGCTCGACTATCAAAACAGCTCGAGAGATCCTCTA
GATTTAAGAAGGAGATATACATATG
>grxC
GAGGCAATTCCGACGTCTAAGAAACCATTATTATCATGACATTAACCTATAAAAATAGGCGTATCACGAGGCCCTTTCGTCTTCACCTC
GATGGATCCAAACTTGCCTTTGGTGCGCGGCAAATAATTTACCTGTCGTCTTTCATGATGCAGGCCGTTAGCTGCAATTAGGCCCCCAG
CGACTTATTGTTATAAGCTTCTGGGGACACTCCATTTTGCTGCCGCCCTGCAACATGAAATTTCAGGGTAAAAAAACAGGAGCTAATTT
```

ATGGCCAATGTTGAAATCTATACCAAAGAAACCTGCCCGTATTGCCATCGTGCAAAAGCACTGCTGAGCAGCAAGGGCGTGAGTTTCTC
GAGAGATCCTCTAGATTTAAGAAGGAGATATACATATG
>grxB
GAGGCAATTCCGACGTCTAAGAAACCATTATTATCATGACATTAACCTATAAAAATAGGCGTATCACGAGGCCCTTTCGTCTTCACCTC
GATGGATCCTGGGTTGGCAGTTTAGCCAGAAACGCGCCGCGCGTCGTTTGCTTGAACGCGACGCCTGATTTGATACTCCCCCCTGACTC
CTTCATACTGAAAGTGAACTGACAGCCAAATGGAGGAGTCACGTGAAGCTATACATTTACGATCACTGCCCTTACTGCCTCAAAGCCCG
CATGATTTTCGGCCTGAAAAATATCCCCTCGAGAGATCCTCTAGATTTAAGAAGGAGATATACATATG
>dicB
GAGGCAATTCCGACGTCTAAGAAACCATTATTATCATGACATTAACCTATAAAAATAGGCGTATCACGAGGCCCTTTCGTCTTCACCTC
GAGAAGTCAGACCATAACGCGAAAGCGCATGGCGAGGTAGCTGGTTCATAGATAGCCTGTCGTTAAATTTTCGTCGACCGTGCGCTTCC
GGTTGTGGCAACCCGCGAAATGGCGCGGCGGTAAGTATGGCGGGGTTATTCCTTCCCCGTTGAGGACACCGGGTTGTCAGGTTGACCAT
ACGCTTAAGTGACAACCCCGCTGCAACGCCCTCTGTTATCAATTTTCTGGTGACGTTTGGCGGTATCAGTTTTACTCCGTGACTGCTCT
GCCGCCCTTTTTAAAGTGAATTTTGTGATGTGGTGAATGCGGAGATCTTCTAGATTTAAGAAGGAGATATACATATG
>elaD
GAGGCAATTCCGACGTCTAAGAAACCATTATTATCATGACATTAACCTATAAAAATAGGCGTATCACGAGGCCCTTTCGTCTTCACCTC
GAGCGCGCTATGATGACAAAGGTTGTCAGCACCTGTTACGTGAATGCAGGTCAATTTTCCCGGCGACTGAACTGGCGAATGATTTCACC
GTGTTTAACGTTTAACTCTGCTTATTATTAAAAACAGGGCGAAACTTGCCCTGTTATCGCAACCCGCGCTACATATATCCACCCACACA
GGCATATTGAGCGATTTTTGATATTTATTCATCTCCTTACGCACTATATTCAGAGCAATAAGTTGCCTCTTTTCATCTTGATTCAGCAC
AACCTTTGCAACTAAAAGGAGCCCATGATGGTTACAGTTGTCAGCAATTATTGTCAATTATCTCAAACGCAACTCAGTCAGACATTTGC
AGAAAAATTTACTGTGACCGAGGAATTACTGCAGGGATCCTCTAGATTTAAGAAGGAGATATACATATG
>elaA
GAGGCAATTCCGACGTCTAAGAAACCATTATTATCATGACATTAACCTATAAAAATAGGCGTATCACGAGGCCCTTTCGTCTTCACCTC
GATGGATCCTATTGCCGTGACATTGCGCGTGCGGGTTGGCACACCGGCTGAAGTACCTAAAAAAATTAATTCCATGAGTTCATCACGTT
TCAAAAAATATTCCGGCTAGTATAAGGCATACATCTAAAAGGAGAGCATTATGATTGAATGGCAAGATCTGCATCACTCTGAATTATCG
GTTTCCCAGCTTTATGCCTTATTACAACTGCGTTGCGCGGTGTTTGTTGTCGAGCCTCGAGAGATCCTCTAGATTTAAGAAGGAGATAT
ACATATG
>elaB
GAGGCAATTCCGACGTCTAAGAAACCATTATTATCATGACATTAACCTATAAAAATAGGCGTATCACGAGGCCCTTTCGTCTTCACCTC
GATGGATCCGATGGCACGCGAGGTAATTCAGGCGTAATCAACAACCCTTGTCTATAGTTAGTGACAGGTTTTACGCAAATGGAGAACGA
GAATGTCTAATCAGTTTGGTGATACACGTATCGATGACGACCTGACGCTGCTTAGTGAAACACTGGAAGAGGTGCTCCGCTCCTCGAGA
GATCCTCTAGATTTAAGAAGGAGATATACATATG
>hcaT
GAGGCAATTCCGACGTCTAAGAAACCATTATTATCATGACATTAACCTATAAAAATAGGCGTATCACGAGGCCCTTTCGTCTTCACCTC
GATGGATCCAAAGGGGAGATGAAGCCGGCATTACGCGATTTCATCGCCATTGTGCAGGAACGTTTGGCAAGCGTAACGGCATAATTACG
CATTATTTTTGCAACATGCGTCGCGGATCTCAGTTTTTTAGTGCTTGTGAACTGGCGTTACAGACATCGGGAAGCAGGCTCGAATATTC
TTATTTTTCAACTTGCACCAACATTCGATTAACCCCAGACTAACGCCTCCTGACGGGAGGGACTCATGGTTTTGCAATCCACGCGCTGG
TTGGCGCTCGGCTATTTCACATACTTTTTTAGTTACGGCATTTTTCTACCTTTCTGGAGCGTCTGGCCTCGAGAGATCCTCTAGATTTA
AGAAGGAGATATACATATG
>osmC
GAGGCAATTCCGACGTCTAAGAAACCATTATTATCATGACATTAACCTATAAAAATAGGCGTATCACGAGGCCCTTTCGTCTTCACCTC
GAGTGCTGTTGAATTTTCTGCCTGATAATAAGTAAACATAGTGATTCTCCGTGTCTGTGTATTTATGGTGTCTGCTACGGATCGCAGAT
TTATAAAGCACATTCAGCATGGCAAATATTTGCCGCTTCGTTGTTAAGATTAGTCCTGGTTGATGATTTTTATATTTTAACACCATGAT
ATTCATAGGGATTGTGATTGGTATGATCCGATTAATATTGATACAATATCTTTTGGGTTATATATTCCCGGTAATCTATTGTGGGAATT
TAATTTAAGTGCAGAAGTAATATTTTCGCCGGATTTTATTCGGAATATCCTGCTTATCCTCGTGCTGTTTCTCACGTAGTCTATAATTT
CCTTTTTAAGCCCACAGGAGAGCAACAATGACAATCCATAAGAAAGGTCAGGCACACTGGGAAGGCGATATCAAACGCGGGAAGGGAAC
AGTATCCACCGAGAGTGGCGTGCTGAACCGGATCCTCTAGATTTAAGAAGGAGATATACATATG
>osmB
GAGGCAATTCCGACGTCTAAGAAACCATTATTATCATGACATTAACCTATAAAAATAGGCGTATCACGAGGCCCTTTCGTCTTCACCTC
GATGGATCCATTCGAAAATACGCGCCAGCGATCTGATGCATCTTGAGCACAGTAAACTCACCATTCACGTCGTTGACATTTAACCATTT
CCCTTCTCTACGGATGATTTGCAGTTTGGCAAATCATCCGCTCTAAGATGATTCCTGGTTGATAATTAAGACTATTTACCTGTTATTAA
CACTCTCAAGATATAAAATTATTATCAGCGATATAACAGGAAGTCATTATCACCTGCGTGATATAACCCTGCGCGCGAGCAGATTTCAC
GGAATAATTTCACCAGACTTATTCTTAGCTATTATAGTTATAGAGAGCTTACTTCCGTGAATCATAAATTCAGGAGAGAGTATTATGTT
TGTAACGAGCAAAAAAATGACCGCGGCTGTTCTGGCAATTACTTTGGCAATGTCTCTGAGTGCCTGTTCTAACTGGTCTAAACGGGACC
GCAACCTCGAGAGATCCTCTAGATTTAAGAAGGAGATATACATATG
>osmE
GAGGCAATTCCGACGTCTAAGAAACCATTATTATCATGACATTAACCTATAAAAATAGGCGTATCACGAGGCCCTTTCGTCTTCACCTC
GATGGATCCGCCCAGCGCCTTTATTATTTGTTGTTGCAATGTCATTGAACCCCTCCAGACAGAAAAGACAAAACTTCCTTAAAGCTAAC
CCGTTGCTACTGAAACGACAAGCGACAGCGGGCTGAATGGTGGTTAAAACGTCATACAGGAATCCTCTGAAAGCGGAGCCGTTTCGTTC
ACGGGCTTGAAAAAGCGCCCAATGTATTCCAGGCTTATCTAACACGCTGATAAACAAGAGGACGGAATATGAACAAGAATATGGCAGGA
ATTCTGAGTGCAGCGGCGGTATTAACCATGCTGGCGGGTTGTACGGCTTATCTCGAGCGGTCGAGAGATCCTCTAGATTTAAGAAGGAG
ATATACATATG
>hcaR

GAGGCAATTCCGACGTCTAAGAAACCATTATTATCATGACATTAACCTATAAAAATAGGCGTATCACGAGGCCCTTTCGTCTTCACCTC
GATGGATCCCCGACCATTTTGGGTGTCAATCAGTTGGTAAATGTTCAAATCTGAGGGTGTGGTCATATTTTTTAATCCTGAAAGATAAG
ATGTATTGACCTTCTATTTTTGTGCCATAAGGATGTTGCTGGTTGGTTGCTAATATGTGAAATAGTTAATTTCTACCAGCTGAGACGTT
TTTTCGATAACAAGGGGAAGGTGATGGAACTACGGCATTTACGCTATTTCGTCGCAGTGGCGCAGGCACTGAACTTTACCCGTGCGGCG
GAAAAACTGCATACTCGAGAGATCCTCTAGATTTAAGAAGGAGATATACATATG
>leuA
GAGGCAATTCCGACGTCTAAGAAACCATTATTATCATGACATTAACCTATAAAAATAGGCGTATCACGAGGCCCTTTCGTCTTCACCTC
GATGGATCCAGACGAGTGAGCGGCATCCAGCATTAAGCCAGCACGCAGTCAAACAAAAAACCCGCGCCATTGCGCGGGTTTTTTTATGC
CCGAAGCGAGGCGCTCTAAAAGAGACAAGGACCCAAACCATGAGCCAGCAAGTCATTATTTTCGATACCACATTGCGCGACGGTGAACA
GGCGTTACAGGCAAGCTTGAGTGTGAAAGAAAAACTGCAAATTGCGCTGCTCGAGAGATCCTCTAGATTTAAGAAGGAGATATACATAT
G
>asnV
GAGGCAATTCCGACGTCTAAGAAACCATTATTATCATGACATTAACCTATAAAAATAGGCGTATCACGAGGCCCTTTCGTCTTCACCTC
GAGCGCTGGTTGTGCGATATGCAATACTTCAGCAGCCTGGGTCAGGCTACCAATATCTACAATTTTTACGAAGTATTTCAGGCGTCTGA
AGTTCATGTTGCCTCCGGTTTTTAAGAATCGGCCCAAGTGCCGCCATTACTTACAACCAGATTGCAAGATGCTTGCCAGTTTTATTTTG
GTGTTGATGTACAAGCTAACCAACTGTCAAATAAGAGATTATGATAGATTCGTCATTTGCTCCTTTAATCAGCTGTCGCGTTCCCCTGC
CCTATAAAAGGAGGGTATGCACCACGATGGTTCATTACCCAATAAGATTGAAAGCTCACCACTTTGTTGAAATTGACAGCAAACAAACA
AAAAAATGCATTTCACCCTTTGACATCACCATGCACTGCCATTAATATGCGCCCCGTTCACACGATTCCTCTGTAGTTCAGTCGGTAGA
ACGGCGGACTGTTAATCCGTATGTCACTGGTTCGAGTCCAGTCAGAGGAGCCAAAGGATCCTCTAGATTTAAGAAGGAGATATACATAT
G
>asnS
GAGGCAATTCCGACGTCTAAGAAACCATTATTATCATGACATTAACCTATAAAAATAGGCGTATCACGAGGCCCTTTCGTCTTCACCTC
GATGGATCCTAAAGCGTTTGTTCGGGCGCTGCGCAAAGCGTTCGACCTTCCGCATATTAAAAAAGCCAGTTAATATCATCAGGGAGCTA
ATCGGCTCCCTTTTTTTACCTTTAATTCCGAAATCTTTCGCTGCATTTGCGAATTCTGCTTGTCTGATTGCAGATGCCAGGTAACATAG
GTATCCCCCCATTTACGGGTGGACAAGTGTTTATTTTTTCCGACTATTAACAGAGAGAATATTATGAGCGTTGTGCCTGTAGCCGACGT
ACTCCAGGGCCGTGTAGCCGTTGACAGCGAAGTCACCGTGCGCGGATGGGTACGTACCCGCCGAGATTCAAACTCGAGAGATCCTCTAG
ATTTAAGAAGGAGATATACATATG
>leuS
GAGGCAATTCCGACGTCTAAGAAACCATTATTATCATGACATTAACCTATAAAAATAGGCGTATCACGAGGCCCTTTCGTCTTCACCTC
GATGGATCCAGGCGTTCGCTCAGTGACGCAACCAGTTCACGGTAATATTGAGCAACCTTGTTCATCGTTTCATCTCCCGGGTCGTTAAC
TGACTATAATAATAGACGTTCTTTGTTCCTTGCTTTGCTAATACGCCCACAAACTCTGTAAATTAATGTATGAATAATCCGGGCATCAT
TTCAACGTCGAGTGCGCGAAAGGCTGTTTTGACGCGGACGTTTGGGCTATGCTATGCGGATCTGAAAAACCACCTCAACGCTACATTTG
TAGCCGTATTGAAAACAGGACCACTGGCTGCCATGCAAGAGCAATACCGCCCGGAAGAGATAGAATCCAAAGTACAGCTTCATTGGGAT
GAGAAGCGCACATCTCGAGAGATCCTCTAGATTTAAGAAGGAGATATACATATG
>asnC
GAGGCAATTCCGACGTCTAAGAAACCATTATTATCATGACATTAACCTATAAAAATAGGCGTATCACGAGGCCCTTTCGTCTTCACCTC
GATGGATCCAGCCCCAGACGTTCTTCCAGTTGACGAGAAAAGTGAGATTTCACGAAGCTAATTTGACGTTGTTTGGCAATGTAAGCGGT
TTTCATTTTTTATACTCCTGCGTCCTGTTGCTTATGATTAAGCAACAAAAACCGACACACATGCAATAATCATTCAATAAAAAGCCTAA
CTCACTTTTGATTCATTAAAAAAGAAGGCTAAAATAGAATGAATCATCAATCCGCATAAGAAAATCCTATGGAAAATTATCTGATCGAC
AATCTGGACCGTGGCATCCTGGAAGCATTAATGGGCAATGCGCGCACCGCTTACGCCGAACTGGCGAAACAATTTGGCCTCGAGAGATC
CTCTAGATTTAAGAAGGAGATATACATATG
>asnA
GAGGCAATTCCGACGTCTAAGAAACCATTATTATCATGACATTAACCTATAAAAATAGGCGTATCACGAGGCCCTTTCGTCTTCACCTC
GAGGCCAAATTGTTTCGCCAGTTCGGCGTAAGCGGTGCGCGCATTGCCCATTAATGCTTCCAGGATGCCACGGTCCAGATTGTCGATCA
GATAATTTTCCATAGGATTTTCTTATGCGGATTGATGATTCATTCTATTTTAGCCTTCTTTTTTAATGAATCAAAAGTGAGTTAGGCTT
TTTATTGAATGATTATTGCATGTGTGTCGGTTTTTGTTGCTTAATCATAAGCAACAGGACGCAGGAGTATAAAAAATGAAAACCGCTTA
CATTGCCAAACAACGTCAAATTAGCTTCGTGAAATCTCACTTTTCTCGTCAACTGGAAGAACGTCTGGGGCTGGATCCTCTAGATTTAA
GAAGGAGATATACATATG
>ydbH
GAGGCAATTCCGACGTCTAAGAAACCATTATTATCATGACATTAACCTATAAAAATAGGCGTATCACGAGGCCCTTTCGTCTTCACCTC
GAGCCAGCTCAAAGCCAAAGGACTCGTTCACCTGTTGCAGGTACTTCTTGTCGTACTGTTTTGTGCTATAAACGGCGAGTTTCATAAGA
CTTTCTCCAGTGATGTTGAATCACATTTAAGCTACTAAAAATATTTTACAAAATTTCAAATTTAATTGAAAGCTATGGCGATATTGAAA
AATTCATCAACAACTATGCTTAGTGTAGGCGCAACCTTCAACTGAACGGTTAAACATGCCACAATACCCGTATTGAATGCTTAATTTTT
CGCTAAATCAGGATATTAACTACCCATGCTGGGTAAATATAAAGCCGTTCTCGCGCTGTTATTACTGATTATTCTTGTGCCGTTGACGC
TGCTGATGACGGGATCCTCTAGATTTAAGAAGGAGATATACATATG
>ydbC
GAGGCAATTCCGACGTCTAAGAAACCATTATTATCATGACATTAACCTATAAAAATAGGCGTATCACGAGGCCCTTTCGTCTTCACCTC
GAGCGCTGAATCTGGATGCGTACAACTGGAAAGAGGATGGCATCAGCGATAAAGGCGTGATGCTGAACTTCAAGAAAACGTTCTAATTT
TTAGCATGTGATCCCTAAACCGCAACGCTGATACAGGTTGCGGTTTTTTTATTGCCGGATGTGGTACGTGACGCGTTTTGTTTTGTGTC
TTTCAGGACAATAGAGCAACTCATCCAGTAATCTTGTTTACACCTTCGCATTATTTATCTCTTTTCGTTTCTATACTGATTTTTCTTAA
TCCGTTTTATTACAGGGCAGGGTGCGATGAGCAGCAATACATTTACTCTCGGTACAAAATCCGTTAACCGTCTTGGTTATGGCGCGATG
GATCCTCTAGATTTAAGAAGGAGATATACATATG
>ydbD

GAGGCAATTCCGACGTCTAAGAAACCATTATTATCATGACATTAACCTATAAAAATAGGCGTATCACGAGGCCCTTTCGTCTTCACCTC
GAGGTCTTCGGTTGCGCATTTACGGGAGAATATGGCTGCTGAAAAATTGCATCTTTCTGAGGAAGTGTTGTCTACGTTGGATGGTATTT
CGCGAGAATAACGAATATACAAAAGGGAAAGATGCATTTCCCTTTTTTTCTTTTTAATGGCATGGAGTGCATATGTTGAAGGCCAGGA
ATTGCGGCTGGATACGTTTACTGCCCCTGTTTATGCTGAGTCTACCAGTACAGGCGGAACTTCGATAGGATCCTCTAGATTTAAGAAGG
AGATATACATATG
>dksA
GAGGCAATTCCGACGTCTAAGAAACCATTATTATCATGACATTAACCTATAAAAATAGGCGTATCACGAGGCCCTTTCGTCTTCACCTC
GATGGATCCTCTGGCTTACAAAGCGGAAATTTCTGCTGAAGGCATGGCTCTTAAAAAATCACTGCCGGTTACATTGTAGTAAAGTAAGT
AACTGGTTAATTTACATTCTGGTCGCGTGCGCAAATACGCTTTTCCTCACACAGTTGTCAAGTGTTACGTTTAGATAATTGCTATCCGG
AAAAGCATCTGCTATTTATAGCGGCCTCATTTTTCCCCCGAACATGGGGATCGATAGTGCGTGTTAAGGAGAAGCAACATGCAAGAAGG
GCAAAACCGTAAAACATCGTCCCTGAGTATTCTCGCCATCGCTGGGGTGGAACCATATCAGGAGAAGCCGGGCGAAGAGTATCTCGAGA
GATCCTCTAGATTTAAGAAGGAGATATACATATG
>b1228
GAGGCAATTCCGACGTCTAAGAAACCATTATTATCATGACATTAACCTATAAAAATAGGCGTATCACGAGGCCCTTTCGTCTTCACCTC
GAGTTCCTGCTGTTCCCGTTCTCGCGTCTGATACACATCTGGAGCGTACCGGTGGAGTATCTGACACGTAAGTACCAGCTGGTGCGCGC
TCGTCACTAAGCGAATTTTAGTTCACATAGACCCTGCTTCGGCGGGGTTTTTTTATGGGCACGGTGCGGGGTGAGTTGTCGGATGCGCT
TCGCTTATCCGACCTACAGGGGAGGATATTGTAGGCCCGGTAAACGTGGTGCCATCGGGCAAAATTAGCTCGGAGTAACAGGTTTTGAT
TATTTAAGGAAGCGATGGTGGTGGGGGAAGGATTACTCAGCGCTGCGCGCTTCGCCCTTCGGGTCGTTGCCTGCGGCAACGCTCTCTCG
CTGGCGCTCAAGTCGAACCTTGGTCGGGATCCTCTAGATTTAAGAAGGAGATATACATATG
>hisL
GAGGCAATTCCGACGTCTAAGAAACCATTATTATCATGACATTAACCTATAAAAATAGGCGTATCACGAGGCCCTTTCGTCTTCACCTC
GAGGCTTTCATCATTGTTGCCGACAAATTCTGACGCGCTTCGCTGTAGCTAATTGTACGCATGTCAATCTCCTCTTTTGTACAGTTCAT
TGTACAATGATGAGCGTTAATTAACTATTTATTAATTAGTTTGTAGATCAAGGTATTGTCAGTGAGACGAAAATCCAGGCTTCGCTATT
TTTGGTGCCATCAGCTAAGAGGACAGTCCTCTTAGCCCCCTCCTTTCCCCGCTCATTCATTAAACAAATCCATTGCCATAAAATATATA
AAAAAGCCCTTGCTTTCTAACGTGAAAGTGGTTTAGGTTAAAAGACATCAGTTGAATAAACATTCACAGAGACTTTTATGACACGCGTT
CAATTTAAACACCACCATCATCACCATCATCCTGACTAGTCTTTCAGGCGATGTGTGCTGGAAGAGGATCCTCTAGATTTAAGAAGGAG
ATATACATATG
>hisJ
GAGGCAATTCCGACGTCTAAGAAACCATTATTATCATGACATTAACCTATAAAAATAGGCGTATCACGAGGCCCTTTCGTCTTCACCTC
GATGGATCCTTCAATAAGGCGCTTGGCGAGCTGCGTCAGGACGGCACCTACGACAAGATGGCGAAAAAGTATTTCGACTTTAATGTCTA
CGGTGACTGATACGTCGCTGGGAAGCTGTACCTGATGGAATGATCATCATGGTGCACGCCAGGTTTGTTGCACTATCGTGGTGCATTGA
AATGCATACTTAAGCATTTTTAATGAAAAATAATACGTCTAACGGGGCGGGATATTTTGCCTTGATGGTCAATTTTATGGCACGATAAG
TGTAACAAACCTGTAAATATTCCCTATAAAAAGACTGTCAGTTGAGGACATTATGAAAAAACTGGTGCTATCGCTCTCTCTGGTTCTGG
CCTTCTCCAGCGCAACTGCGGCGTTTGCTGCGATTCCGCAAAACATCCCTCGAGAGATCCTCTAGATTTAAGAAGGAGATATACATATG
>rmf
GAGGCAATTCCGACGTCTAAGAAACCATTATTATCATGACATTAACCTATAAAAATAGGCGTATCACGAGGCCCTTTCGTCTTCACCTC
GAGTAAAGTGCTGGCCGGTGTCTGGAGTCAGGAAGCCGCTTCTATTGCACAAGAGATAAAGCGTCTACCTTAATTATAAAGATTTGTAA
ATATAACCGTCTCCGGTATGTTGCCTGAGGCGGTTTTTTTGTCTCTAACGTGCGGAAAAATTTGTTCCTCTTCACATTTTTTGTACAAC
CGACATGCCCGTGTAGCTCACAAATATGACAGTGGCGTGAATTTTGCGCATTGACGGCAGTTATGATTCGCGGTATTGCTTAACTGTGA
TTGCACATTTAGTAATCACTGTTTTCTTTTCCACCAGAAACCAGTATGAGGGAAACGAGGCATGAAGAGACAAAAACGAGATCGCCTGG
AACGGGCACATCAACGTGGTTATCAGGCCGGCATCGCCGGACGCTCAAAAGAAATGGGATCCTCTAGATTTAAGAAGGAGATATACATA
TG
>hisQ
GAGGCAATTCCGACGTCTAAGAAACCATTATTATCATGACATTAACCTATAAAAATAGGCGTATCACGAGGCCCTTTCGTCTTCACCTC
GATGGATCCCGCGAAGCACTGAACAAAGCCTTTGCCGAAATGCGCGCTGACGGTACTTACGAGAAATTAGCGAAAAAGTACTTCGATTT
TGATGTTTATGGTGGCTAATCGCCATCGGTGAAAGAAAGCCCGCTCCCTTCGGGTAACCGGAGGGAGACAAGAGGCGGTAATTCACCAC
ACACGACAGGACAGGCAGCATGTTGTATGGGTTTTCAGGTGTTATTTTACAGGGTGCGCTCGTCACGCTGGAGCTGGCTATCAGCTCTG
TAGTGCTCGCTGTAATCATCGGCTCGAGAGATCCTCTAGATTTAAGAAGGAGATATACATATG
>hisS
GAGGCAATTCCGACGTCTAAGAAACCATTATTATCATGACATTAACCTATAAAAATAGGCGTATCACGAGGCCCTTTCGTCTTCACCTC
GATGGATCCAGTCAGCTGGACGAAGCGCGTCGAATTGACGTTCAGCAGGTTGAAAAATAATAACGTGATGGGAAGCGCCTCGCTTCCCG
TGTATGATTGAACCCGCATGGCTCCCGAAACATTGAGGGAAGCGTTGAGGGTTCATTTTTATATTCAGAAAGAGAATAAACGTGGCAAA
AAACATTCAAGCCATTCGCGGCATGAACGATTACCTGCCTGGCGAAACGGCCATCTGGCAGCGCATTGAAGGCACACTGCTCGAGAGAT
CCTCTAGATTTAAGAAGGAGATATACATATG
>ypdA
GAGGCAATTCCGACGTCTAAGAAACCATTATTATCATGACATTAACCTATAAAAATAGGCGTATCACGAGGCCCTTTCGTCTTCACCTC
GAGGCGCAGCCATTTTCAGTTCAGCGGTGATGTTAAAAACGTAGGGCGGGAGACGATCAATGCGCGTAAAGCGACGTTCAGGGCGAGTG
TCAGCCATAATTTCCTCAGATTAACGTTAGCGCCCGGACCGTCCGAGCGACGCTGCCACGATGGTGGCTGTTTTGAAAATAGCCTGATT
AATTTCTGTCTGTCCAGCGTTTTAGAACATTTTATTAATGACGTTAAATGAGGAACTCTGCTTTAAAAACAGGAAATTTATCTGCGCTA
ACAAGAAAATTACCACTCATTTTATTAACATAATAATTTCAATAAATTTACTAACCAGAAGTGCATTATCAAGAGATGCATTGCTAATA
CACCATTTATCTATCAATCCCCTTTTAAAATTGTGGTTTTTCCTCAAAAACGGCCTGCTGGTCATCAGCCTTCAGGTTACCTATCATAG
AGGTTTAATCCTTATTCAGAGTCACCCGTGCACGAAATATTCAACATGCTGCTGGCGGTCTTCGATCGGGCAGCGTTAATGCTTATCTG
CCTGTTCTTTCTCATCCGTATCCGCCTGTGGATCCTCTAGATTTAAGAAGGAGATATACATATG

Data Set S4 - MG1655 library reference file.fasta

```
>fadL
GAGGCAATTCCGACGTCTAAGAAACCATTATTATCATGACATTAACCTATAAAAATAGGCGTATCACGAGGCCCTTTCGTCTTCACCTC
GAGCCATACAGCAGCAAACCGGGGTTTCATCAGCACTACATTTACTCATCGTTGATTTCCTCTGTATGTGCACCCAAGGTGCCAGATAA
ACGTTGTGGATATTTTACGCTTCCGGAAAGTGCTGCTCCAGTTGTTAATTCTGCAAAATCGGATAAGTGACCGAAATCACACTTAAAAA
TGATCTAAAACAAAATTCACCCGAATCCATGAGTGCGCCACCTCCAAATTTTGCCAGCTGGATCGCGTTTCTTAGATCATATTTGAAAA
AAGATAGAAACATACTTGCAACATTCCAGCTGGTCCGACCTATACTCTCGCCACTGGTCTGATTTCTAAGATGTACCTCAGACCCTACA
CTTCGCGCTCCTGTTACAGCACGTAACATAGTTTGTATAAAAATAAATCATTGAGGTTATGGTCATGAGCCAGAAAACCCTGTTTACAA
AGTCTGCTCTCGCAGTCGCAGTGGCACTTATCTCCACCCAGGCCTGGGATCCTCTAGATTTAAGAAGGAGATATACATATG
>ynbA
GAGGCAATTCCGACGTCTAAGAAACCATTATTATCATGACATTAACCTATAAAAATAGGCGTATCACGAGGCCCTTTCGTCTTCACCTC
GAGTGGATTTAGTCCGCCAGCTGGCGCTTAACGGTAAGACGCGCGCAATACGGGAACAAGCGATGAGAAAACTTGATGAGTTGTATTGA
TCCACAACGTAATGTTTTTTAACTATCTGATTAATTGGGGATAATCATTCCTGACAGTGAGTCCCCAATACCTTGATATATTCTGAATT
TTTAATGAAACGGCGTGTTGCGATATCTCCGTCAGGGGAATTGATGCACCATAGCGCAAACCGAATTATCAAGGATTGATAATGACGCT
CTACCAGATAAAACCGCTCTTTCAGTCGCTGTTAAGGCCGACGATGTTTTGGCTTGGATCCTCTAGATTTAAGAAGGAGATATACATAT
G
>manX
GAGGCAATTCCGACGTCTAAGAAACCATTATTATCATGACATTAACCTATAAAAATAGGCGTATCACGAGGCCCTTTCGTCTTCACCTC
GAGATGGCGATGAAGACCAGGTTATCGATACCCAGCACAATTTCGAGAACAACAAGCGTGAGTAGCCCCGCCCAAATTGAGGGGTCCAT
TAAGAATTCCATGACAAGCTCCTGCTTAAGGAATAGCTATTCGACGCCAGAAATAATGCAGGCGTAACGACAAAATGCAAACGAAAGGT
GCGGCATAGAGTGCCAGAAAGGCAGGCGTTAAAAGGCCTGATGCTGAAATGACGTCGGTGACGATCCATACTGCGGGCTACTGCCCTAT
ACTCCATGGTTGTTAAACGGGAGTTAAACATATCAGAGACGCCTCTGATTTGGCAAAGATTTACCTTCCTTTGCAAACGAATGTGACAA
GGATATTTTACCTTTCGAAATTTCTGCTAATCGAAAGTTAAATTACGGATCTTCATCACATAAAATAATTTTTTCGATATCTAAAATAA
ATCGCGAAACGCAGGGGTTTTTGGTTGTAGCCCTTATCTGAATCGATTCGATTGTGGACGACGATTCAAAAATACATCTGGCACGTTGA
GGTGTTAACGATAATAAAGGAGGTAGCAAGTGACCATTGCTATTGTTATAGGCACACATGGTTGGGCTGCAGAGCAGTTGCTTAAAACG
GCAGAAATGCTGTTAGGCGAGCAGGAAAACGGGATCCTCTAGATTTAAGAAGGAGATATACATATG
>manY
GAGGCAATTCCGACGTCTAAGAAACCATTATTATCATGACATTAACCTATAAAAATAGGCGTATCACGAGGCCCTTTCGTCTTCACCTC
GAGCGCGGTATTGAGCTGGAAGTCCGTAAGGTTTCCACCGATCCGAAACTGAAAATGATGGATCTGATCAGCAAAATCGATAAGTAACG
TATTGTGTTGATTATCACTCAGTTTTCACACTTAAGTCTTACGTAAACAGGAGAAGTACAATGGAGATTACCACTCTTCAAATTGTGCT
GGTATTTATCGTAGCCTGTATCGCAGGTATGGGATCAATCCTCGATGAATTTCAGTTTCACCGTCCGCTGGATCCTCTAGATTTAAGAA
GGAGATATACATATG
>yajR
GAGGCAATTCCGACGTCTAAGAAACCATTATTATCATGACATTAACCTATAAAAATAGGCGTATCACGAGGCCCTTTCGTCTTCACCTC
GATGGATCCGGACTCGCATACGCTGCTGGCTGCTGTGTGGTAACAAAACCTCTCTATTAAAAAGGTGCTACGGCACCTTTTTTCTTAGC
ATTAGAAACATATCCCTCTCGAAATATTTACTAAAAAATCCGCATGTTTACCCCATTCGTTTGCCGCTTTACACTAGTCGCGAATTTAA
AACAGAGGTGGTAATGAACGATTATAAAATGACGCCAGGTGAGAGGCGCGCGACCTGGGGTTTAGGGACCGTATTCTCGTTGCGCCTCG
AGAGATCCTCTAGATTTAAGAAGGAGATATACATATG
>yibK
GAGGCAATTCCGACGTCTAAGAAACCATTATTATCATGACATTAACCTATAAAAATAGGCGTATCACGAGGCCCTTTCGTCTTCACCTC
GAGAAGATTCGCTGGTGCAGGGGCTGGGTAAAGAGTTGCCTGCGGCACTGGCTCCCATGGCGAAAGGGAATGCGGCATAGTCGTTTGCG
CCCCCTCACCCTAACCCTCTCCCTCAGGGAGAGGGGACCGTTCGGCGCTGTATGTACTCCCTCACTCTGAAACGACACCGCACTCTTTT
TTTCTCCCTCGCCCCTCCGGGGAGAGGGCCGGGGTGAGGGGAAAAGGCCGCACTGCTCACCAATTTTCTGCTATCCTGCCCCCGCACTA
AGGGGGCAGTATGCTAAACATCGTACTTTACGAACCAGAAATTCCGCCAAATACTGGCAACATCATCCGTCTTTGCGCTGGATCCTCTA
GATTTAAGAAGGAGATATACATATG
>yajO
GAGGCAATTCCGACGTCTAAGAAACCATTATTATCATGACATTAACCTATAAAAATAGGCGTATCACGAGGCCCTTTCGTCTTCACCTC
GATGGATCCCCGCAAGGAACTCAGGAAGAAATGCGCGCCGAACTCGGCCTCGATGCCGCTGGTATGGAAGCCAAAATCAAGGCCTGGCT
GGCATAATCCCTACTCCACTCCTGCTATGCTTAAGAAATTATTCATAGACTCTAAATAATTCGAGTTGCAGGAAGGCGGCAAACGAGTG
AAGCCCCAGGAGCTTACATAAGTAAGTGACTGGGGTGAACGAATGCAGCCGCAGCACATGCAACTTGAAGTATGACGAGTATAGCAGGA
GTGGCAGCATGCAATCTCGAGAGATCCTCTAGATTTAAGAAGGAGATATACATATG
>yajB
GAGGCAATTCCGACGTCTAAGAAACCATTATTATCATGACATTAACCTATAAAAATAGGCGTATCACGAGGCCCTTTCGTCTTCACCTC
GATGGATCCTTCGGGCAATTCAAAGGAGAAATCGGTAACGCGCATGACACTGACTCAGACTAAAATAAGAGGCGGGTAGTCTAGTGCCG
GGGCGCTCTCCCTGCAACCTTTACCCCCTTTTGGATAGAATATAATCGATGAATTTTTTAGCTCACCTGCATTTAGCCCATCTCGCGGA
AAGCTCGCTTTCCGGCAATTTACTGGCTGATTTCGTACGCGGAAATCCCTCGAGAGATCCTCTAGATTTAAGAAGGAGATATACATATG
>yajC
GAGGCAATTCCGACGTCTAAGAAACCATTATTATCATGACATTAACCTATAAAAATAGGCGTATCACGAGGCCCTTTCGTCTTCACCTC
GAGCGGGTTTACGCAAGGCTATTGAAGAGGGTAAATTAGAGAGCTTCGTAACTGATTTTTACCAGCGTCAGGGGCGAGAAGTACCACCT
TTGAACGTTGATTAATATTAATAATGAGGGAAATTTAATGAGCTTTTTTATTTCTGATGCGGTAGCGGGGATCCTCTAGATTTAAGAAG
GAGATATACATATG
>b0374
GAGGCAATTCCGACGTCTAAGAAACCATTATTATCATGACATTAACCTATAAAAATAGGCGTATCACGAGGCCCTTTCGTCTTCACCTC
GAGTCAGCAAGCTTCAGGGCTTCACTGCGAAATTCAGGCGAATGCTGTTTACGGGGTTTTTTACTGGTTGATACTGTTTTTGTCATGTG
```

AGTCACCTCTGACTGAGAGTTTACTCACTTAGCCGCGTGTCCACTATTGCTGGGTAAGATCAGATTACAACCTGTACGCCAATACTATC
ACCGTAACTAACGGCGGTGTATTGGATGTGAACGTTGATCAGTTCGATACTGAAGCTTTCCGTACTGACAAACTGGAACTGACCAGCGG
GGATCCTCTAGATTTAAGAAGGAGATATACATATG
>yajG
GAGGCAATTCCGACGTCTAAGAAACCATTATTATCATGACATTAACCTATAAAAATAGGCGTATCACGAGGCCCTTTCGTCTTCACCTC
GATGGATCCGAACGCCGCCCTTAATTTTTCTTCTATCCGCTCACGTATCATCATGAAATATCTCCTCCGACAACGCTGAGATGTCATCC
ATCCCTTTAAATACTAGCCGCTTTTACCGTTTCCATTGCAGCTTAACAACAAATATTTAGGTTTCGTCCTGGTTTTTCCGTCAAGCGTA
TGAAGTTTACTGATTTAAAGATTGAAACACAGGCCTCGCATCAGTGTTTCTTTTACCAGGGGCGATTATTTCGTCAACATGATGAATTT
ACATGCCTTACCCACTTCCCCTCGCCGTTGGCAATGTTATGATGGCGGGAATTTTTACCACTCGAGAGATCCTCTAGATTTAAGAAGGA
GATATACATATG
>ycbK
GAGGCAATTCCGACGTCTAAGAAACCATTATTATCATGACATTAACCTATAAAAATAGGCGTATCACGAGGCCCTTTCGTCTTCACCTC
GAGGATGGTCGTACCCAGTATCGTACAGATATTTACAATTATGATCTGCCTGCGCGATCCAGCTCGCAAATCGTATCGAAAGCGGAACA
ATTAATCAGGTAAATGAAGTAGTTCGGGGAAATCAGTTGTCGTAATAATTAGCATGATTGGGGGCGATTCTCTGCAGCCCCCGTCACTG
CTGGGGTTGAGTCATCTTGACGTCTGCTTTACGGGCGGTTAAGGTGCCTCTTGTGCGCCAGAAGTGCATATAAACGATAACATTGACCT
GTAGACTTGATTATCATGGACAAATTCGACGCTAATCGCCGCAAATTGCTGGCGCTTGGTGGCGTTGCACTCGGTGCCGCCATCCTGCC
GACCCCTGCGTTTGCAACACTGGATCCTCTAGATTTAAGAAGGAGATATACATATG
>ycbB
GAGGCAATTCCGACGTCTAAGAAACCATTATTATCATGACATTAACCTATAAAAATAGGCGTATCACGAGGCCCTTTCGTCTTCACCTC
GAGACACGTTCATGTCGTCGGCCTGCGAGGATTTGCGCCGCAACTCCCTGAAACGCTTCCAGGAACTGACGAAGCGCCTTCTCAGGCGA
GTTAAAATTAAGGCGGCAGCAATGCCGCCTTTTCTTTTTCCGAAAACTCCGTTTCTGCACTAAAAAAGTGGCACATTACGGCGCGCTTT
TCTTTAAACTTCTTTACATTAGGTTATGTAAAAACGTATCGGCGTTTATATACTGAAGATAAGCCTGATGAGTAACAGGCTTGCTCGTC
ATACTTTCGTGAGTATTGGCGTTGTACAGGCAAGTCGTAAAATAACAGCCTGGCTATTCAGAGTATGATAAAAACAGGGGGCAAGGGAT
GTTGCTTAATATGATGTGTGGTCGTCAGCTGTCGGCAATCAGTTTGTGCCTGGCCGTAACATTCGCTGGATCCTCTAGATTTAAGAAGG
AGATATACATATG
>ycbC
GAGGCAATTCCGACGTCTAAGAAACCATTATTATCATGACATTAACCTATAAAAATAGGCGTATCACGAGGCCCTTTCGTCTTCACCTC
GATGGATCCTAGCCTGTCGAAGCTGCCCTTTGGTGGTGCCGTAAATGTTACGGGAAAACTTTTCCGCAATATCATCAAAATTGCGATCC
TGCATTGCTTTCTCCACCGGTTGGCTAAGACGTTTTCTGATAAAGACGCTATTTTGTCACAGCCACGGCGAGAATGAACCCGTCTGGTG
TAATATCGCTGGCGATCCCTTGCTATATGGTTAAAAAAGGAACCAGAAAGGATGCTTTTTACACTGAAAAAAGTGATTGGCAATATGTT
GCTGCCACTTCCGCTGATGTTGTTGATTATTGGCGCTGGCCTGGCGCTGCTGTGGTTTAGCCCTCGAGAGATCCTCTAGATTTAAGAAG
GAGATATACATATG
>lpxC
GAGGCAATTCCGACGTCTAAGAAACCATTATTATCATGACATTAACCTATAAAAATAGGCGTATCACGAGGCCCTTTCGTCTTCACCTC
GAGAGCCGGTTGCTAAAGTCGTGAATGACAATGCGCCGCAAACTGCGAAAGAGCCGGATTATCTGGATATCCCAGCATTCCTGCGTAAG
CAAGCTGATTAAGAATTGACTGGAATTTGGGTTTCGAGGCTCTTTGTGCTAAACTGGCCCGCCGAATGTATAGTACACTTCGGTTGGAT
AGGTAATTTGGCGAGATAATACGATGATCAAACAAAGGACACTTAAACGTATCGTTCAGGCGACGGGTGTCGGTTTACATACCGGCAAG
AAAGTCACCCTGACGTTACGCCCTGGATCCTCTAGATTTAAGAAGGAGATATACATATG
>ycbZ
GAGGCAATTCCGACGTCTAAGAAACCATTATTATCATGACATTAACCTATAAAAATAGGCGTATCACGAGGCCCTTTCGTCTTCACCTC
GATGGATCCCCATTTCCAACCGCTTTCAAGATTTTCAAGTTGTTGATATTTCATAGCGATCTCAATGTTACCGTGTAACTCTTTACAGA
ATATCAGCTTTTTAGGCCAGTGAAGAAAAGAATCTCCATCCTGTGAGCTTTAACGCCCATGCCAGGAGTATTGTCGGTGCTTTCTACGT
GTGCTGTAGTGCCGGTTACGGTATAATCGCGGCTTTGACAACAGACTAAAAAACATCAACTTTGACCATTACGAAACTTGCATGGCGTG
ACCTGGTTCCTGATACCGATAGCTATCAGGAAATATTTGCTCAGCCACATTTGACTCGAGAGATCCTCTAGATTTAAGAAGGAGATATA
CATATG
>csiE
GAGGCAATTCCGACGTCTAAGAAACCATTATTATCATGACATTAACCTATAAAAATAGGCGTATCACGAGGCCCTTTCGTCTTCACCTC
GAGATCGTCACGGCGATGTTTATCGCGAACAGATGGTGGACTTTATCCTTAGCGCGTTGAATCCGCAGAACTAACCCATGATCGCTAGC
ACGATAATCATTCACAAAACCACCTTAAGACATGCTAATCCACTGGTCAGAACAGTTTAAGATGAGAAAAATTCTGTGACGCTTGCCAA
CATTTCTGATGATTAGCATTCCCTTCGCCATTTCCTTGAGCAAACTTTAGCTATTCTTATCAATTATGCTTATGGGAGATCTACAGATG
ATGCCTACGCTTGCTCCACCATCTGTCCTTTCGGCTCCCCAGCGCCGCTGCCAGATCTTGCTGACGCTCTTTCAGCGGATCCTCTAGAT
TTAAGAAGGAGATATACATATG
>csiD
GAGGCAATTCCGACGTCTAAGAAACCATTATTATCATGACATTAACCTATAAAAATAGGCGTATCACGAGGCCCTTTCGTCTTCACCTC
GAGCGCTTAGAGGCGGAACTCTAAATATCAATATTCCTGGGCGTAGCGCTCTTCTTTTAGGGAAAACAGGAGAACCGCCGAACTATCTC
TATTTATAATTTAATGTTATATCTGCCCCGATAAAACGGGGCAGATAATATGTTTAGTTTACTAACGGTCATTTTGCAGTGAAGCCATT
TACTGTTTTTTATCGACCAGATAATCTGTTCTCTAATGTTAACTCCCCCTAACCTGTTGCTTTAGTTATTCATTTCCTGTCTCACTTTG
CCTTAATACCCTACGTTAAATGTTACTAATTTGTTGCTTTTGATCACAATAAGAAAACAATATGTCGCTTTTGTGCGCATTTTTCAGAA
ATGTAGATATTTTTAGATTATGGCTACGAAATGAGCATCGCCATGTCACCCTACAGGATCCTCTAGATTTAAGAAGGAGATATACATAT
G
>ycbR
GAGGCAATTCCGACGTCTAAGAAACCATTATTATCATGACATTAACCTATAAAAATAGGCGTATCACGAGGCCCTTTCGTCTTCACCTC
GAGTATCGCTACGGGGAAAGCGACTTCAGGTAATGTGAATGCGGTAACAAATTTCCATATTAACTATTATTAATAGAACTCATTAATTG

TTTTATTAATTAGTACCCCTCCAGTGTTCTGGAGGGGATATTCATATTTTTTAAGAGTGACTATTTATGAAAACTTGCATAACAAAGGG
AATTGTGACCGTAAGTTTAACGGCAATATTACTCAGCTGCTCATCCGCATGGGCGGCAGGTAAAGGCGGGATTGGATCCTCTAGATTTA
AGAAGGAGATATACATATG
>ycbW
GAGGCAATTCCGACGTCTAAGAAACCATTATTATCATGACATTAACCTATAAAAATAGGCGTATCACGAGGCCCTTTCGTCTTCACCTC
GAGAAAAGATTGCTGCGGGTGCCTCACTGGTGCAAATTTATTCTGGTTTTATTTTTAAAGGTCCGCCGCTGATTAAAGAAATCGTTACC
CATATCTAATTATTTCTTCGAATTCCTCTTTATAAGATAATCAGGGCTTTATTTTCAGCCCTGGTTGTTTTATATTCATCGCTGTTGCT
TATTTAGACATTTTGTACTTTTATTGTTGAGGTTATTAAGCGAAGCGACAATGGATTGTGTTGCTGCGGTTTTATAGGGTAGGGGAGAG
GCAGATGCGAATTAAACCAGACGATAACTGGCGTTGGTATTACGATGAAGAGCACGATCGGGATCCTCTAGATTTAAGAAGGAGATATA
CATATG
>glcB
GAGGCAATTCCGACGTCTAAGAAACCATTATTATCATGACATTAACCTATAAAAATAGGCGTATCACGAGGCCCTTTCGTCTTCACCTC
GATGGATCCGGCAAATTATTGGTGCCGTGGGCGTTTCTGGTTTAACCGGAGCACAGGATGCGCAGGTCGCGAAAGCGGCAGCAGCGGTG
TTGGCGAAATAAGCGAAAACGAGGAGATAAACAATGAGTCAAACCATAACCCAGAGCCGTTTACGCATTGACGCCAATTTTAAACGTTT
TGTGGATGAAGAAGTTTTACCGGGAACAGGGCTGGACGCTCTCGAGAGATCCTCTAGATTTAAGAAGGAGATATACATATG
>glcC
GAGGCAATTCCGACGTCTAAGAAACCATTATTATCATGACATTAACCTATAAAAATAGGCGTATCACGAGGCCCTTTCGTCTTCACCTC
GAGTCACGCAGTGCCATCAGTACCGATGTGCGGTCGACATCGGGTAAAGCGCCATCAAGACGCTCTTCGTACAAGATGCTCATGAGTAG
GCTTCGCTTTGTTGTGTTGTGTGGCAGCTGATTTTTGCGCGCTGCTTCTGTGAACAGTTATTAAGCGGGCTTTTCGTTTTCGTCTATCT
CTTTAGCTACCGGTCAGACCATTTTTTTTCCAGCTCTGTGACCTTGTCTTGGTTAACTCAATGTTAAATTGATGTAACATAATCACTTA
CGTGATGTGCGTGTTTTGCGAGTTAAGAACAGAAAAATTGGTCCTACCTGTGCACGAGGTCCGGGAATGAAAGATGAACGTCGCCCTAT
TTGCGAAGTGGTTGCAGAGAGTATCGAACGGTTAATTATCGACGGCGTACTGAAGGTCCGTCAGGATCCTCTAGATTTAAGAAGGAGAT
ATACATATG
>artP
GAGGCAATTCCGACGTCTAAGAAACCATTATTATCATGACATTAACCTATAAAAATAGGCGTATCACGAGGCCCTTTCGTCTTCACCTC
GATGGATCCAAGGCCAGTGTTGGGTCATCGACGATGTGCGTTACTTAGGCGGTAGCGTTCATGCCACGGCGGGTACGCTCCGTCAGTCC
ATCGAGAATCGCTAACGCCTTGATCAACCAATAAACCCCGTAAATTGTCCGGCATTTCGTGAGGAATGCCGACATTTATGCTCGCCGAC
CACCGCCCCCGTTATTTTGTGCTATGTTTATTGAATAATGCGCTTTGCTTTTAACTTTTAAAGCAGAAATATTGCATAATTATTCTGTC
AAAGGTACTATCTTCGACCTCAATAGCTATCAGACTGCCAGTATACGAGTGTCAATGAGTATTCAATTAAACGGCATTAATTGCTTCTA
CGGCGCGCATCAGGCGCTGTTCGATATCACGCTGGATTGCCCCTCGAGAGATCCTCTAGATTTAAGAAGGAGATATACATATG
>marC_marR
GAGGCAATTCCGACGTCTAAGAAACCATTATTATCATGACATTAACCTATAAAAATAGGCGTATCACGAGGCCCTTTCGTCTTCACCTC
GAGGGCCAAGAAATAACGCGACAGTTGTTAATGGGTTAGCTAACGGCAGCAACACCACCAGCCCCAGGCCAATTGCTTTAAACAAATCT
AACATTGGTGGTTGTTATCCTGTGTATCTGGGTTATCAGCGAAAAGTATAAGGGGTAAACAAGGATAAAGTGTCACTCTTTAGCTAGCC
TTGCATCGCATTGAACAAAACTTGAACCGATTTAGCAAAACGTGGCATCGGTCAATTCATTCATTTGACTTATACTTGCCTGGGCAATA
TTATCCCCTGCAACTAATTACTTGCCAGGGCAACTAATGTGAAAAGTACCAGCGATCTGTTCAATGAAATTATTCCATTGGGTCGCTTA
ATCCATATGGTTAATCAGAAGAAAGATCGCCTGCTTAACGAGTATCTGTCTCCGCTGGATATTACCGCGGCACAGGATCCTCTAGATTT
AAGAAGGAGATATACATATG
>rluE
GAGGCAATTCCGACGTCTAAGAAACCATTATTATCATGACATTAACCTATAAAAATAGGCGTATCACGAGGCCCTTTCGTCTTCACCTC
GATGGATCCAGTTTGCCGTTTTGCAGGGTGATCTTCTTGCCTTGTGCCGGAACAACTACTTTACTTTCCATTCACCTCTCCTTCGAGCG
CTACTGGTTTGCTCGTCTGCCACCACGTTGCATATGCGTTTGCGTCCTGCGATACGGATGCTTTAGAGCAATTTTTTGTTAATGATTTG
TAATTGGCTTGTCAATACTACCCTATTGTTTGCCGTCATGAAAGATGCGCGTTATTAGGCTATAATGCGGCAATTCATAATCTCTGAAA
ATACCATGCAAAAAACTTCTTTTAGAAATCACCAGGTTAAGCGATTCAGCTCGCAACGTTCTACCAGGCGTAAACCTGAAAACCAGCCC
ACTCGAGAGATCCTCTAGATTTAAGAAGGAGATATACATATG
>rluD
GAGGCAATTCCGACGTCTAAGAAACCATTATTATCATGACATTAACCTATAAAAATAGGCGTATCACGAGGCCCTTTCGTCTTCACCTC
GATGGATCCACAGGCTTAGTGTGGCGGCTGCCACCAGATATTTCATGCGCGTCATGACGTTTTGACTTTCCTCAAAATGTAATACGGGA
GATTCTCTGTTCCTGCTCCCGGTTAAGACCAGCTACAATAGCACACTATATTAAACGGCAAAGCCGTAAAACCCCAACGATAAACGAAG
AAGCAGTATATATGGCACAACGAGTACAGCTCACTGCAACGGTGTCCGAAAACCAACTCGGTCAACGCTTAGATCAGGCTTTGGCCGCT
CGAGAGATCCTCTAGATTTAAGAAGGAGATATACATATG
>rluF
GAGGCAATTCCGACGTCTAAGAAACCATTATTATCATGACATTAACCTATAAAAATAGGCGTATCACGAGGCCCTTTCGTCTTCACCTC
GAGGCGACCCTGCAACTGTTCAGCAATTAGCGGCAGTGCATGTTCCAGCCAGGCTTTACCCGGCAGCGTCGAGTTACTCAATAAAAGCA
GTTCCATTACTTCTCCGGTTAGTAAGAAATCACGGGAGGCATGCTACCACTGCGAATTTAACCACTTACTGACATGGCTCACGATGTCG
CACATTGCTTCTGGAAAGTGATTTGAGCAGGTAACGAATTCAAGCCAATGGAAATAACAAATAGTTCGCTCATGACTATTTTTTATCAA
CCCAAAGGTACGGTATATTCCGTTTTTTACAGGAGAATTTATGCTGCCCGACTCATCAGTCCGTTTAAATAAATACATCAGCGAAAGCG
GAATTTGCTCACGCCGCGAAGCGGATCGCTATATCGAGCAAGGCAATGTGGGATCCTCTAGATTTAAGAAGGAGATATACATATG
>rluC
GAGGCAATTCCGACGTCTAAGAAACCATTATTATCATGACATTAACCTATAAAAATAGGCGTATCACGAGGCCCTTTCGTCTTCACCTC
GAGCGGAAACGGCCTCATTATTCCACTGCAAGCCTTGTTATAGCAAGATGACTTTTACCATTTATCACCCGCTTACTCACAGTTTTTTC
ACTTCTTGCTGGTGATTGGTTTAATAACCATCAATAAAGTGATCACGCAGTGAAGTAATAAGGTGCAAAAGTAAATATAAGCATAGAAA

```
AATGAGTGGCGCGAATCCTGTTGGCTAATTAGAATCGCCAACCATGAAAACAGAGACTCCATCCGTAAAAATTGTTGCTATCACCGCTG
ACGAAGCGGGGCAACGTATCGATAGGATCCTCTAGATTTAAGAAGGAGATATACATATG
>rluB
GAGGCAATTCCGACGTCTAAGAAACCATTATTATCATGACATTAACCTATAAAAATAGGCGTATCACGAGGCCCTTTCGTCTTCACCTC
GAGTAAGTTGCGGGGCCAATACCAGTTTGTTGGGGCGAAGTTATTCCGGTGGTGGTGCTGGCGGCGGCGCGGGTGGCGGAGGCGGTGGT
GGCTGGTAATTAAGCTGATGTTAATCGAAACAAGCTTTAACCATAAAGATAGTGTAGTATGTTGCGCCTCAAAGCCAGGCCGATAAACG
TCGAGTCGTTTACTTAAGGCCTGAAGAGTTCAAACAGACTTTACTATCAGTGAGTCAGATGTAAGTTCTGACTCACTGTTGCACAAAAG
CATATCAGACGCCTGGGAAGGCGACACCAAAAGGAAGCTCTATGAGCGAAAAGCTACAGAAAGTGCTGGCGCGTGCCGGCCACGGTTCT
CGCCGTGAAATCGAATCTATTATTGAAGCCGGTCGTGTGAGTGGATCCTCTAGATTTAAGAAGGAGATATACATATG
>b0373
GAGGCAATTCCGACGTCTAAGAAACCATTATTATCATGACATTAACCTATAAAAATAGGCGTATCACGAGGCCCTTTCGTCTTCACCTC
GATGGATCCCCGCTGGTCAGTTCCAGTTTGTCAGTACGGAAAGCTTCAGTATCGAACTGATCAACGTTCACATCCAATACACCGCCGTT
AGTTACGGTGATAGTATTGGCGTACAGGTTGTAATCTGATCTTACCCAGCAATAGTGGACACGCGGCTAAGTGAGTAAACTCTCAGTCA
GAGGTGACTCACATGACAAAAACAGTATCAACCAGTAAAAAACCCCGTAAACAGCATTCGCCTGAATTTCGCAGTGAAGCCCTGAAGCT
TGCTGACTCGAGAGATCCTCTAGATTTAAGAAGGAGATATACATATG
>helD
GAGGCAATTCCGACGTCTAAGAAACCATTATTATCATGACATTAACCTATAAAAATAGGCGTATCACGAGGCCCTTTCGTCTTCACCTC
GAGACCAGAGTCGCCAACAGCCAGCCCAGAGTGGTGGCAAATCCGCCAAGCACAAAGTTCAGAATGTTCAAAACGGTACGCATAAAACC
TCGCTTTACTGTGGTTTTCAGTAATGGGCGGCAATTGTAACGTTTTTTTGGGCTGGAGCACGTTTTCTCTGACGGTTACACTGATAAGA
AATAATTTCGTGTGGATCTACAGAGTCATGGAACTGAAAGCGACAACGCTTGGAAAACGTCTGGCACAGCACCCTTACGATCGGGCGGT
GATCCTCAATGCCGGGATTAGGATCCTCTAGATTTAAGAAGGAGATATACATATG
>yqiC
GAGGCAATTCCGACGTCTAAGAAACCATTATTATCATGACATTAACCTATAAAAATAGGCGTATCACGAGGCCCTTTCGTCTTCACCTC
GAGAGCACCATTACACCGCGTCCTTCACGCAGCGCAGCCAGTGCATTTTCAACACGTTCGAAAGGCGTACCAAAAGAGGAAAGTAGCGT
CTGATTCATGGTAAAAAAACCTCACTAAAATTATGGTTACCAGAATCAGGGCAGTCTTAGGAGTGGCGGCATATAGCCAAAATAACGTG
AGCGGGTCCATGCCCGACAGAATCGTTACTCTCTCCCATCCGGACTCTAACCGTCGGCCCCGGAATTACACCGGATCTGCTGTCCTTTG
AGTTCGCACCCAAAGCGCTCGCGGGCTTTCAACTGAGTTGATTTACCGCCGGTGGGGAATTTCGCCCCGCCCTGAGAATAAGCGGATTC
ACTATAACGCTAATGATTAGCGGCAGCAACGCATAGCTTCACATAATTCTGGTTTATGACTTACCCTTATCGCACTACAATGGCACTCA
ACACCTATCAGTACAGGGAAACTACAATGATTGACCCGAAAAAAATTGAGCAAATCGCTCGCGGATCCCGGGATCCTCTAGATTTAAGA
AGGAGATATACATATG
>polB
GAGGCAATTCCGACGTCTAAGAAACCATTATTATCATGACATTAACCTATAAAAATAGGCGTATCACGAGGCCCTTTCGTCTTCACCTC
GATGGATCCGTCAGTTAGCGCCGCAGTTACCGGATATGCAGCAAACGCTGCTGGATAAACACTATCTGCGTAAGCATGGCGCGAAGGCA
TATTACGGGCAGTAATGACTGTATAAAACCACAGCCAATCAAACGAAACCAGGCTATACTCAAGCCTGGTTTTTTGATGGATTTTCAGC
GTGGCGCAGGCAGGTTTTATCTTAACCCGACACTGGCGGGACACCCCGCAAGGGACAGAAGTCTCCTTCTGGCTGGCCTCGAGAGATCC
TCTAGATTTAAGAAGGAGATATACATATG
>polA
GAGGCAATTCCGACGTCTAAGAAACCATTATTATCATGACATTAACCTATAAAAATAGGCGTATCACGAGGCCCTTTCGTCTTCACCTC
GAGTTAACCCTTCCAGCCCGTGAACTTCCCATTGAAGGTGTGGGTTCAGATGCAGTAATACCGCCAGACCTTCACACCAGCAATACATC
ATAAAATCACAAAAACGCGAAACCTTTCGCCAAATGACTGGCACAGGTAGCAATAGTTTTACAATCCCGGCAATGATGATCGGGACGGA
ACAAAAAATAGTGACCAAAATGGTCAATACGATACTCAGCAAAAGGGTTATCGCAGCGAGTATTCTCGTCATAATGAATTTATTCAAAA
GGTTAGCCATAGACAGTGCGCTAAAGAAGCGCAAGGCGCAGATTTTAGCAGAAAACGACCCAAATAACGGATGATCCTTAAGGAGAAAA
ATAATTCATATCTATCCACATTAGAAAAAATCCCATTATCTCAATTATTAGGGATGGATTTATTTTTAACTGCATGAAAAACAAAGACA
AACATCATGCTGTAAAAAGCATGATAATAAATTAAAAGCGATGTAAATAATTTATGCACAAAGTTATCCACATGACGATTTGCGAGGAT
CCTCTAGATTTAAGAAGGAGATATACATATG
>yqiJ
GAGGCAATTCCGACGTCTAAGAAACCATTATTATCATGACATTAACCTATAAAAATAGGCGTATCACGAGGCCCTTTCGTCTTCACCTC
GAGCTTCTGCGTGCATTCTGGCAAAACTACGCGCCAGGAAATCGAAATTATTTAAAGAATTAAGACTATGATCCATAAGCACATCCTCC
GGTTATCGCATACCTTTTAAGGCCATATGCTTTTTCTGATTTCAGTATTCCCCAATCAGTGTAGTGCCTTTAAACATAACGTGCGTAAA
TATATTGTCCCCGATCACACTTTTTAGTGAAAAATAAAAAACCATATTAAAAGACTTTCTCTTCCAGAACTTCTCTAACCTGTGTTATT
ATTAACACAGATAAATGTAAGCAAGGAAACTATTACAAAGTAAATATTGCGTCACTAAATGGACATTGGAGTGATATATGATTTTATTC
GCCGACTATAACACCCCTTATCTTTTCGCTATTTCCTTTGTACTGTTGATCGGCCTGTTAGAAATTTTTGCGCTTATCTGCGGTGGATC
CTCTAGATTTAAGAAGGAGATATACATATG
>ecpD
GAGGCAATTCCGACGTCTAAGAAACCATTATTATCATGACATTAACCTATAAAAATAGGCGTATCACGAGGCCCTTTCGTCTTCACCTC
GATGGATCCTCGCAGACCAAACAGCAACTGCTGGTTATGTAAAAACTAACACTGCATACACCATTACTTATCAGTAAGTATTCACCAGT
TAAATTGATTGAATGAATATACAGGGAATAATAATTTCTATTTTATATTATTCCCTGTTTTAATTAACTCTATCAGGGATGGTTTATGT
TTTTTAACACCAAACATACAACAGCTTTATGCTTCGTAACCTGTATGGCTTTTAGTTCATCGTCTATTGCGGACATTGTCATTTCGGGT
ACTCGCGTCTCGAGAGATCCTCTAGATTTAAGAAGGAGATATACATATG
>yjhF
GAGGCAATTCCGACGTCTAAGAAACCATTATTATCATGACATTAACCTATAAAAATAGGCGTATCACGAGGCCCTTTCGTCTTCACCTC
GATGGATCCTGTGGCAGGGATTGCTTTACTCCTGCTTTTAACCATAAAAATTAAGCTTAATACGTTTGTTTCGTTAATTATTGTCTCGA
TTGCTGTCGCCATCGCCAGTGGTATGGATTTGAGTAAAGTCGTCACCTCAGTCGAATCCGGACTTGGCGGTACGCTGGGCCATATTGGT
```

Data Set S4 - MG1655 library reference file.fasta
TTGATATTTGGCTTCGGTGTTATGCTCGGTCGCTTGCTGGCCGATGCGGGTGGTGCGCAGCGAATAGCGCTGACCATGCTGAATTCTCG
AGAGATCCTCTAGATTTAAGAAGGAGATATACATATG
>yjhI
GAGGCAATTCCGACGTCTAAGAAACCATTATTATCATGACATTAACCTATAAAAATAGGCGTATCACGAGGCCCTTTCGTCTTCACCTC
GATGGATCCGCACAGCGTTATCAGCGAGCCGGATGTCAGTTGTTTTTACCTCATTCAATAAAGTGATAAGTAAGTTCATTCGAGAGGGA
TTTCAAGCAAAAATAATCAATGGCACCCAATAGAAAATATTGGCGATGCGCTCGAACGAATAAAGAAGCTCTAGGCGCAATCCACACAC
TACGATGTTGCAACAACACGCCATCTACTTTTTATTCTCATTCACTAAATGTGGCTGTTCTGGATTCATTATTCAAAGTGTGTACAAGA
TCACATTTAATCACATCATTACGGTTCAGCATGCTGAACAAAGCATATTTTCCACTATGTAATGCCGATACCATTTATTCCATGAGCAA
GGAGGAGCCATTATGGTTCGCAAAGGGTGTAATTCATTAGTCCGTGCTGAAAAAATATTAACCCACATTGCATGGGTTGGTATGGCTCG
AGAGATCCTCTAGATTTAAGAAGGAGATATACATATG
>yjhU
GAGGCAATTCCGACGTCTAAGAAACCATTATTATCATGACATTAACCTATAAAAATAGGCGTATCACGAGGCCCTTTCGTCTTCACCTC
GATGGATCCCGCTGTTATCCTGGACGCTGATGTCCACACTCTTATCCATCAGTGGGCTGATTTTTATTCTGCTCGCCAGCTTAGTTTTA
TAATCGGGTTCTTGCCAGTGAGCCAACCTGTTGTTCACTGTCCCCCATAATTTAGTGAGTGACGGCAACCGTCTGCACGGTGACGAATC
GCTGGCGATCAAAAAGGGGCGATATTGAACCTTCGATTATGCTTGATGGGTGTATTCTCAGTAATACTGTAAAGCCGGAGCATGTCTCC
GGCTTTACAGTATGGGGGAACAGCGGGCGATCGCTCAAATGGTATGCTGGTTTATTTCATGTAAAAGAATAAAATATCTGTATAAAATC
ATGCCACTGGCAATCTGCATCAACTCTCCGGGTAAACAGGGTTAAGATGAGCATTTCCCCGGTTTAATACAGGTATACAAAAGTGGATA
GAGATCAAACAAACAGCAGCCTGTTCAATGATGATCCCGTACTTCATGCAACATGGCTGTACTATCAGGAAGGCAAAAGCCAGACTGAA
GTTGCAGCTATCATGGGGGTGTCCCGCGTCACGGTTGTCAAATACCTGCAAACAGCCCGGGAAAATGGACTGGTTCATATCAACCTGGA
TGTGAATGTCTTTGGCTCTATCGACGCTGCTTTGCAAATTCGCGATAAATTCAATCTCCAGAGAGTGATCATCGTGCCGCTCGAGAGAT
CCTCTAGATTTAAGAAGGAGATATACATATG
>yjhS
GAGGCAATTCCGACGTCTAAGAAACCATTATTATCATGACATTAACCTATAAAAATAGGCGTATCACGAGGCCCTTTCGTCTTCACCTC
GATGGATCCAAAGCGGTGACGGATTCAGTTTTGATCACTGTGAAGGATAATAAAGTCACAGTACAAAACTAACGCTTCAGGGCCCCGGT
AAAGGGGCCCTGCTATCAATTTGTCATTTCAATTACGCGAAATTTATATGAACGCAATAATATCGCCCGATTATTACTATGTTCTTACC
GTTGCTGGTCAGTCTAATGCCATGGCGTATGGCGAAGGACTGCCATTACTCGAGAGATCCTCTAGATTTAAGAAGGAGATATACATATG
>hda
GAGGCAATTCCGACGTCTAAGAAACCATTATTATCATGACATTAACCTATAAAAATAGGCGTATCACGAGGCCCTTTCGTCTTCACCTC
GATGGATCCAGAAGTGGTGCTGGACGCAGAAGACGCGGATATAACAGACAAATAATGTACCGACCGGGCAGTGTTGCTGCCCGGTTCAA
ACATCATGGGATTCTGTGGTAAACTTCTCGCGATTTTGTGAAATCCTGGTTGAGGTATCTCTGAACACACCGGCACAGCTCTCTTCTCG
AGAGATCCTCTAGATTTAAGAAGGAGATATACATATG
>pepN
GAGGCAATTCCGACGTCTAAGAAACCATTATTATCATGACATTAACCTATAAAAATAGGCGTATCACGAGGCCCTTTCGTCTTCACCTC
GAGAAACACGGCTTGCTGCATATGCAACTTATAAGCATCTGTATCCAGCAACGAGTGCAGAACAGGAGAAGCGAATTGTGTCATAGGTG
CGCAGTAGCGTCCTCGTACAGGAGCGTTTAGTACAATAAACATCTTCAGGAACTGCTGGAGTATACCTTGTTTCGCAATTTATTGAACC
CCGATCACACCATATGCCACCTTTCTGGTCGATGGCATTACGCGGTGTATGTTATAAAAATGTAGCAATAAAGGCGTTTGTACCTGAAA
AGATGAAGATTCTGCATAGCGCGATTTACGCAACAGGAATAGACTGAACACCAGACTCTATAAAAGATGCTAAAGGTTATTTATGACTC
AACAGCCACAAGCCAAATACCGTCACGATTATCGTGCGCCGGATTACCAGATTACTGATATTGACTTGACCTTTGACCTCGACGCGGAT
CCTCTAGATTTAAGAAGGAGATATACATATG
>pepB
GAGGCAATTCCGACGTCTAAGAAACCATTATTATCATGACATTAACCTATAAAAATAGGCGTATCACGAGGCCCTTTCGTCTTCACCTC
GATAGATCTGCAGGCATCCAACGAGAAAATCCTCGAAGCGATTTTGTTAGTCTGGCTGGACGAGGCCGAATAATTTCCGCGTCATGCTT
CACGCCGCAGATGCGTTGGCTGCGCTCGTTCACCCCAGTCACATAGTTATCTATGCTCCTGGGGCTTCACTCACTTGCCGCCTTCCTGC
AACGCGAATCATTTAGCGGAAAAATCCTGGGGCTGCCAACTGGCGGCCCTTTTACAAAGACTCGAGAGATCCTCTAGATTTAAGAAGGA
GATATACATATG
>pepE
GAGGCAATTCCGACGTCTAAGAAACCATTATTATCATGACATTAACCTATAAAAATAGGCGTATCACGAGGCCCTTTCGTCTTCACCTC
GATGGATCCCGTGAGCAAATTCCGCTTTCGCTGATGTATTTATTTAAACGGACTGATGAGTCGGGCAGCATAAATTCTCCTGTAAAAAA
CGGAATATACCGTACCTTTGGGTTGATAAAAAATAGTCATGAGCGAACTATTTGTTATTTCCATTGGCTTGAATTCGTTACCTGCTCAA
ATCACTTTCCAGAAGCAATGTGCGACATCGTGAGCCATGTCAGTAAGTGGTTAAATTCGCAGTGGTAGCATGCCTCCCGTGATTTCTTA
CTAACCGGAGAAGTAATGGAACTGCTTTTATTGAGTAACTCGACGCTGCCGGGTAAAGCCTGGCTGGAACATGCACTGCCGCTAATTGC
TGAACAGTTGCAGGGTCGCCTCGAGAGATCCTCTAGATTTAAGAAGGAGATATACATATG
>pepD
GAGGCAATTCCGACGTCTAAGAAACCATTATTATCATGACATTAACCTATAAAAATAGGCGTATCACGAGGCCCTTTCGTCTTCACCTC
GATGGATCCAAGGCATCAGTCGGCTTGCGAGTTTACGTGCATGGATCTGCAACATGTCCCAGGTGACGATGTATTTTTCGCTCATGTGA
AGTGTCCCAGCCTGTTTATCTACGGCTTAAAAAGTGTTCGAGGGGAAAATAGGTTGCGCGAGATTATAGAGATCTGGCGCACTAAAAAC
CAGTATTTCACATGAGTCCGCGTCTTTTTACGCACTGCCTCTCCCTGACGCGGGATAAAGTGGTATTCTCAAACATATCTCGCAAGCCT
GTCTTGTGTTGACAACATTTTCTGCTAACCCTGTGACCTGCAATACTGTTTTGCGGGTGATCGACAAGGAGACTTAACGTGTCTGAACT
GTCTCAATTATCTCCACAGCCGCTGTGGGATATTTTTGCCAAAATCTGTTCTATTCCTCACCCGTCCTATCACTCGAGAGATCCTCTAG
ATTTAAGAAGGAGATATACATATG
>mqo
GAGGCAATTCCGACGTCTAAGAAACCATTATTATCATGACATTAACCTATAAAAATAGGCGTATCACGAGGCCCTTTCGTCTTCACCTC
GATGGATCCGGGCAACTTAGCGAGCTGACGGGCGAAGAGCGCGATGCCGCTTCGCGTGATGCCGTTGCCCGGACGGCATAATATTACGA

CGCTGACCTGGCGGACTGCTGCCGTCAGGTCAATATGCTAAAAATCCCTTCATTTCTAATAACCCTATAATTAATCAACGAAATTATAA
TGTTTCTAAAATTAGAATATAATTTATAAACATTATTTAAATGTTGTTACTTAAGTGTTACCGTTGATGCCGCGCAAATCTCACCTGCA
ATAAAGCGACTAAAAGTAAGGCATTAACAAGATGAAAAAAGTGACTGCCATGCTCTTCTCGATGGCCGTGGGGCTTAATGCCGTTTCGA
TGGCGGCAAAAGCGAAAGCGTCCGAGGAGCAGGAAACTGCTCGAGAGATCCTCTAGATTTAAGAAGGAGATATACATATG
>pepQ
GAGGCAATTCCGACGTCTAAGAAACCATTATTATCATGACATTAACCTATAAAAATAGGCGTATCACGAGGCCCTTTCGTCTTCACCTC
GAGACTGAACCTGGGGCATCAAATACCAGTTCGGCAATGCCATCTTCCAGCCAGTCAAGGTACAGGGTGTCGCCTTTGTAAAGCATGTC
AGTCTCCTGAATCCGCAAGGTGATCTGGTCGTACCAGATGAGTCGAAGTGTGTATTTTGTGTTAAAAATATGCAAACAAAAGATTAAAG
AAATGCCGATCTGATCACGCTCGGCAGAAATCACGCTCTGGATGAACGATGTGCTAAGATGCGGAGACTTAAGGTCAAAAAAACAGAAG
GGTAAAAAATGGAATCACTGGCCTCGCTCTATAAAAATCATATAGCTACCTTACAAGAACGGACTCGCGATGCGGATCCTCTAGATTTA
AGAAGGAGATATACATATG
>pepT
GAGGCAATTCCGACGTCTAAGAAACCATTATTATCATGACATTAACCTATAAAAATAGGCGTATCACGAGGCCCTTTCGTCTTCACCTC
GAGCCATCAAAGCATTTGCGAATTCCCGCCAATTGCACCAGCGGTGAAAGCGAACTCGGTTGTTTATTCAATTTTTTACTCTGTCCCAT
GTAAACGCAACGGATGGCTTACCGATGCGGGGTTTGTGGTTAACCACCTTGGTGACTCTTAATGAGGGCGGTAATTCTACGGCAAACCG
CTTGAATCGCCAATCTTTGTTGTGAATTACTGGCTTAGCTTTATATTCATTAAGGTAATGCTGATAAATATTCCCGCTTGCAGGGGTAA
AAGTGACCTGACGCAATATTTGTCTTTTCTTGCTTCTTAATAATGTTGTCACAAAAAGTGAGGGTGACTACATGGATAAACTACTTGAG
CGATTTTTGAACTACGTGTCTCTGGATACCCAATCAAAAGCAGGGGTGAGACAGGTTCCCAGCACGGGATCCTCTAGATTTAAGAAGGA
GATATACATATG
>yidC
GAGGCAATTCCGACGTCTAAGAAACCATTATTATCATGACATTAACCTATAAAAATAGGCGTATCACGAGGCCCTTTCGTCTTCACCTC
GAGCCGACCTCGATAACCGTGCTCTCTCGGAAGCGTTGGAAAAATTATGGCGCCGCCACTGTCGCCTGGCTCGCGGGTCCTGATAGCCC
TCATTCGGGTCTATCAACGCCTGATTAGTCCGCTACTCGGGCCGCATTGTCGTTTCACTCCAACCTGTTCAAGCTACGGAATTGAGGCA
TTGCGCAGGTTTGGAGTGATAAAAGGCAGTTGGTTGACGGTGAAACGCGTATTAAAATGCCACCCTTTACACCCTGGTGGTGACGATCC
CGTCCCGCCCGGACCATTTGATACCAGAGAACACTAACGATGGATTCGCAACGCAATCTTTTAGTCATCGCTTTGCTGTTCGTGTCTTT
CATGATCTGGCAAGCCTGGGGATCCTCTAGATTTAAGAAGGAGATATACATATG
>ybeZ
GAGGCAATTCCGACGTCTAAGAAACCATTATTATCATGACATTAACCTATAAAAATAGGCGTATCACGAGGCCCTTTCGTCTTCACCTC
GATGGATCCCAGTGATTGCCCGTACCCGCAAAGAAAACGACCTTGGCGTGGGTTATTATCAGCCGTAATTCTCAGGCCTGCCGTTCCGG
TGGGCCTTGTTTTTTCTCTCTTTATCCCCATCTTTATTGCAATGCTTGCGCCTTTTTCCTGTGGCGTGAATAATTTCCTTATAGGCCGG
TGAATGTTCAGCTTCCGGCGGCATACAGATAAGAGGAACGGTTTGAACATAGACACTCGCGAAATCACCCTGGAGCCACTCGAGAGATC
CTCTAGATTTAAGAAGGAGATATACATATG
>yidE
GAGGCAATTCCGACGTCTAAGAAACCATTATTATCATGACATTAACCTATAAAAATAGGCGTATCACGAGGCCCTTTCGTCTTCACCTC
GATGGATCCCGCAGCGTATCGCTATCAGCGAACGTCCCGCGTTAAATAGCTAACTAGCTATTCTCTTTGCCCCGCCATTCTGGTGGGGC
TTTTTTTGTCTTACTTACTCAACAATGCGTTAAATTTCGACTGTTTAAGATATTTCGGCACGTTCTTTCCCGTGATAATGTGCGCTATA
AGGCAAATCTGCTTCACCGATTCAGCCATAATCCGTAGCAATTAAAATGTTATCTCAAGAGAAATGGACGATGAGTGATATAGCATTAA
CGGTCAGTATTCTGGCTTTGGTGGCACTCGAGAGATCCTCTAGATTTAAGAAGGAGATATACATATG
>yidK
GAGGCAATTCCGACGTCTAAGAAACCATTATTATCATGACATTAACCTATAAAAATAGGCGTATCACGAGGCCCTTTCGTCTTCACCTC
GATGGATCCCCGGCGAATTATGCTCACAACCTGAACTTAACAGAAAACAATCCGTTCAGGGATACGACGTTTGTATTTTAAGAATCTGA
CTGCCTGACCCGACGCATTTTACCTCTCCCTATATTCATGCGTCCGGGACATAAACATAAATAAGGGCTATGAGATGAATTCGTTACAA
ATCTTGAGTTTTGTCGGTTTTACGCTGCTGGTGGCGGTGATCACCTGGTGGAAGGTCCGCAAAACAGATACCGGATCGCAACAAGGCTA
TTTTCTTGCCGGACGCTCACTAAAAGCGCCGGTTATTGCCGCTTCGTTAATGCTAACCAACCTTTCCACGGAACAACTGGTCGGCCTTT
CCGGGCAGGCCTACAAAAGCGGCATGTCGGTGATGGGCTGGGAAGTGCTCGAGAGATCCTCTAGATTTAAGAAGGAGATATACATATG
>yidH
GAGGCAATTCCGACGTCTAAGAAACCATTATTATCATGACATTAACCTATAAAAATAGGCGTATCACGAGGCCCTTTCGTCTTCACCTC
GATGGATCCGGCCGAATGAAAAGTGGATAATCCCCATCATTAAAGCGATGGCACCGAACAGCATGCCCAATGAACTGATTTTATTTTGA
GCTATGATACCCATCTGACCTCCTTGTAGCGGATGGCAAAAGCATATCAGGAAACGGGCGGACGAAATTCTTAAAAAAACCGCTAAAGG
TAGAGCGTTAGTGAAATTGCAAAGGAGCAACAATGAAAATTTCCCGCCTCGGAGAAGCGCCTGATTACCGCTTCTCGCTGGCTAATGAG
CGTACTTTTCTGGCGTGGATCCGTACTCTCGAGAGATCCTCTAGATTTAAGAAGGAGATATACATATG
>ybeU
GAGGCAATTCCGACGTCTAAGAAACCATTATTATCATGACATTAACCTATAAAAATAGGCGTATCACGAGGCCCTTTCGTCTTCACCTC
GAGTCGTTTTCTTGGGTTGGCTAATACAGGTAATTGCTGCAACTATAAAAATTGCGAATATATACATTAATTTTTTATTCATGTGTGAA
TGCTTCGATTTTCTTTTTATTTTGAAAGCGTCAAGTTTATACTCATCAATAATAAAAAAGTAGTCAGTTAATGCACTTAAGGGATTATT
AACAAATGAATAAGGAAGAACAATATCTTCTTTTTGCGCTTTCTGCGCCCATGGAAATACTTAATCAGGGTTGCAAACCCGCGCATGAC
TCGCCAAAAATGGGATCCTCTAGATTTAAGAAGGAGATATACATATG
>ybeR
GAGGCAATTCCGACGTCTAAGAAACCATTATTATCATGACATTAACCTATAAAAATAGGCGTATCACGAGGCCCTTTCGTCTTCACCTC
GAGCGCCTTTTTCAGCACGTTCGATAATCTCATCTATTGATAAATTATCGCAGCAACTTGACGTGAAAATCATTATCATTTCCTTATGG
TTGCAGATAAAACATATTATATTTTTATGTGTTCCATTTTTGCGTAAGCCTTATTTTATGCGTATTATGTCGCGTCAGAAAAGTATAAA
TAAACGAGCCTCGCAATGCGGCTAATATTCATTTAATGAATATTTAAGGATAAATTATATGGACATGGAGTCGCAAAAAATATTGTTTG

CACTTTCCACTCCGATGGAAATACGCAATGAATGCTGTTTACCTTCGCATTCATCACCCGGATCCTCTAGATTTAAGAAGGAGATATAC
ATATG
>yidL
GAGGCAATTCCGACGTCTAAGAAACCATTATTATCATGACATTAACCTATAAAAATAGGCGTATCACGAGGCCCTTTCGTCTTCACCTC
GAGCCGGTATCTGTTTTGCGGACCTTCCACCAGGTGATCACCGCCACCAGCAGCGTAAAACCGACAAAACTCAAGATTTGTAACGAATT
CATCTCATAGCCCTTATTTATGTTTATGTCCCGGACGCATGAATATAGGGAGAGGTAAAATGCGTCGGGTCAGGCAGTCAGATTCTTAA
AATACAAACGTCGTATCCCTGAACGGATTGTTTTCTGTTAAGTTCAGGTTGTGAGCATAATTCGCCGGAGGGATATTAAAATGAATGGA
AAATTGCAAAGTTCAGATGTAAAAAACGAAACTCCGTACAATATTCCCTTATTGATTAACGAAAATGTGATCTCCAGCGGAATTTCTCT
GATCTCGCTCTGGCATACCTACGCCGACGAGCATTACCGGGTGGATCCTCTAGATTTAAGAAGGAGATATACATATG
>ybeQ
GAGGCAATTCCGACGTCTAAGAAACCATTATTATCATGACATTAACCTATAAAAATAGGCGTATCACGAGGCCCTTTCGTCTTCACCTC
GATGGATCCGGGTGATGAATGCGAAGGTAAACAGCATTCATTGCGTATTTCCATCGGAGTGGAAAGTGCAAACAATATTTTTTGCGACT
CCATGTCCATATAATTTATCCTTAAATATTCATTAAATGAATATTAGCCGCATTGCGAGGCTCGTTTATTTATACTTTTCTGACGCGAC
ATAATACGCATAAAATAAGGCTTACGCAAAAATGGAACACATAAAAATATAATATGTTTTATCTGCAACCATAAGGAAATGATAATGAT
TTTCACGTCAAGTTGCTGCGATAATTTATCAATAGATGAGATTATCGAACGTGCTGAAAAAGGCGCTCGAGAGATCCTCTAGATTTAAG
AAGGAGATATACATATG
>yidR
GAGGCAATTCCGACGTCTAAGAAACCATTATTATCATGACATTAACCTATAAAAATAGGCGTATCACGAGGCCCTTTCGTCTTCACCTC
GATGGATCCGCCAATAATCGCCACGTCGAAATGTTCCATCTTTTCACTCACATTCATCACGATGGCAAAAGACTATTGTCTGAAAGTGA
AGTCCTTGTGAAGTATGATGGCGGGACCCGTTTTGTATCAGGACCGCGCCATGAAACAGATAACCTTTGCTCCCCGTAATCACCTGCTC
ACCAATACCAATACCTGGACGCCCCTCGAGAGATCCTCTAGATTTAAGAAGGAGATATACATATG
>yidS
GAGGCAATTCCGACGTCTAAGAAACCATTATTATCATGACATTAACCTATAAAAATAGGCGTATCACGAGGCCCTTTCGTCTTCACCTC
GAGGGTGATTACGGGGAGCAAAGGTTATCTGTTTCATGGCGCGGTCCTGATACAAAACGGGTCCCGCCATCATACTTCACAAGGACTTC
ACTTTCAGACAATAGTCTTTTGCCATCGTGATGAATGTGAGTGAAAAGATGGAACATTTCGACGTGGCGATTATTGGCGGATCCTCTAG
ATTTAAGAAGGAGATATACATATG
>ybeL
GAGGCAATTCCGACGTCTAAGAAACCATTATTATCATGACATTAACCTATAAAAATAGGCGTATCACGAGGCCCTTTCGTCTTCACCTC
GAGATGTGCGCTTCTCATCCCAATGAAGCTGTACTTTGGATTCTATCTCTTCCGGGCGGTATTGCTCTTGCATGGCAGCCAGTGGTCCT
GTTTTCAATACGGCTACAAATGTAGCGTTGAGGTGGTTTTTCAGATCCGCATAGCATAGCCCAAACGTCCGCGTCAAAACAGCCTTTCG
CGCACTCGACGTTGAAATGATGCCCGGATTATTCATACATTAATTTACAGAGTTTGTGGGCGTATTAGCAAAGCAAGGAACAAAGAACG
TCTATTATTATAGTCAGTTAACGACCCGGGAGATGAAACGATGAACAAGGTTGCTCAATATTACCGTGAACTGGTTGCGTCACTGAGCG
AACGCCTGGATCCTCTAGATTTAAGAAGGAGATATACATATG
>ybeF
GAGGCAATTCCGACGTCTAAGAAACCATTATTATCATGACATTAACCTATAAAAATAGGCGTATCACGAGGCCCTTTCGTCTTCACCTC
GATGGATCCACCCGAAGCGACGACTAATAATATTGCTCCACGTTTACTGGAAAATATTTTAGCGCTACTAAACAATCCGGACTTCGAAT
ATATTACCGCTTAATTCCATACATCAATGGCCCAATTTACACTGGGTCATTACTCATAATATCTGCTAATGCTTTACATCTCCTTCGCG
TTTGTATATTTTCTTTATCACTTGCAGAAATGCGGTGCCACGCCTTGCTCCGTAACTCACCCCATTGATTAACCTATTGAAATAGTTGA
TTATGGTGAGCAAAAGCGTTCCGTTCCAATTAGACATAAGATTATTGTAATTTTTAAATATTATTTTTCCATGAATAAAATTGGAGTA
AGTGCGTGGATAGTAATAATCAAATTGAACCCTGTTTAAGCAGAAAGTCGTCAGAAGGTAAACCACAAATATTTACAACCTTACGAAAT
ATTGATCTTAACCTTCTGACTATTTTTGAAGCTGTATATGTACATAAAGGGATCGTTAATGCAGCGAAAGTGCTTAATCTGACCCCCTC
GGCAATCAGTCAGTCTATTCAGAAACTGCGCGTTATATTCCCTGACCCATTGTTTATTCGCAAAGGCCAGGGTGTCACCTCGAGAGATC
CTCTAGATTTAAGAAGGAGATATACATATG
>yidX
GAGGCAATTCCGACGTCTAAGAAACCATTATTATCATGACATTAACCTATAAAAATAGGCGTATCACGAGGCCCTTTCGTCTTCACCTC
GAGCGAGCCTGGCACGTATTTGCCGTGAACGATCTGTTTACCCAGCGTAATGACAATGCGATCGGTTTTATTGAGAGTCATGGAGAGTC
CTTGTGCTCTGTATGTTCTTCTCTACTTTACCCCGATCGATGCATAACGCGGCAACTTTGTAGTACCAGCGTGATGACGTTCGCGTTTG
CCGTGCGTGTAATGTAGTACAAACTTATATTGTTGTACTACAATTTAGATCACAAAAAGAACAATGCATAAAAAAATGACATGCGTCGG
GCAGAAATCTGAAAAGGGATATCAGGCGCTAAACAGGAGGGAAAGAAGAGTATGCTTTCAACGGCTTAGCTACTCGTTTAAAGGATTAA
TCATGAAGTTGAATTTTAAGGGATTTTTTAAGGCTGCCGGTTTATTCCCACTGGCGCTGATGCTTTCAGGCTGTATCTCGTATGCTCTG
GTTTCCCATACCGCAAAGGGTAGTTCAGGAAAGTATCAATCGCAGTCAGACACCATCACTGGGCTATCGCGGATCCTCTAGATTTAAGA
AGGAGATATACATATG
>ybeB
GAGGCAATTCCGACGTCTAAGAAACCATTATTATCATGACATTAACCTATAAAAATAGGCGTATCACGAGGCCCTTTCGTCTTCACCTC
GATGGATCCAAAATTCGCGACGCTACGCGTCCTCAATAGCCGTGCCATCGGGGTCGAGAATGCATGACTTTTCTGTTTTTTTACGGGTA
AGCCGCAACGACCATTGACAGACCCGGGCAGGCTGATATTCTCCGCAGCCAGACTTTTTCCGCCAGACACGACTTTGTAGAAATTGTTT
TACAAAAATGGCGATGCAATCTGCGGCGCGGGGTGGGATGATAGCCCACTTTCGAAAGCCGATTCGGCGACAATTGTCCCGAAATCGCC
TCTGGTTCAGGTATACTGACAGACCATTTTTATCTATTTGATTCACCCAGGGGGAAAACTTGCAGGGTAAAGCACTCCAGGATTTTGTT
ATCGACAAAATTGATGACCTCAAAGGTCAGGACATCATCGCCTTAGACGTTCAGGGCAAATCCAGCATCACCGACTGCATGATCATCTG
TACGGGTACGTCCAGCCGTCATGTTATGTCCATTGCTGACCACGTTGTGCAGGAGTCTCGCGCAGCGGGCCTGTTACCGCTCGGCGTAG
ACTCGAGAGATCCTCTAGATTTAAGAAGGAGATATACATATG
>chaB

GAGGCAATTCCGACGTCTAAGAAACCATTATTATCATGACATTAACCTATAAAAATAGGCGTATCACGAGGCCCTTTCGTCTTCACCTC
GAGAACAGCACTACCAGCGCCAGAACCGGGAAAATAAGCGAAGTCTCCTTGTGGCGGGTTTTTACCGCCTCTTGAGCATTTGACATTAT
GGTTATCCCTTTGCAGATGAATTTATCGAAAATGTAAAAAATAGGTAGGAAAAATAACAGAAAGTGTCTGGATATCGGTAACATTTTAC
GAATTTTTACCCCGCTGTCGATTTTTTACTCATTGGGGCATAAAAATAAGTATTACGTTTAGACAATGTTTGTTTTAGCGTCTCTTAAG
AAGAGTCTGACCTGAAAATTCTTATGTTTTGGCAAGAGTAGATATTGTTGACCACACTTAATGTTCAACTTTGTAAAAGGAGTCAACGA
TGCCGTATAAAACGAAAAGCGATCTGCCGGAAAGCGTAAAGGGATCCTCTAGATTTAAGAAGGAGATATACATATG
>cvpA
GAGGCAATTCCGACGTCTAAGAAACCATTATTATCATGACATTAACCTATAAAAATAGGCGTATCACGAGGCCCTTTCGTCTTCACCTC
GATGGATCCGAAAGGTTCGCTGGGTGAGTTGAAGCAACTTTCTGGCTTAAGTGGCGTGGTAATGGGCTATACGCCGAATTAATACGGTC
TTGCCTGATGCGACGCTGGCGCGTCTTATCAGGCCTACGCAGGGGTAGAACCGTAGGTCGGATAAGGCGTTTACGCCGCATCCGACACG
CATTGCCCGATGCCGCAAAGGCATAAAAAGTCGATGGCGTTGAATATTTTTTCAGCGCCATTTTTATTGATGCGCGGGAAGGAAATCCC
TACGCAAACGTTTTCTTTTTCTGTTAGAATGCGCCCCGAACAGGATGACAGGGCGTAAAATCGTGGGACACATATGGTCTGGATTGATT
ACGCCATAATCGCGGTGATTGCTTTTTCCTCTCTGGTTAGCCTGATCCGCGGCTTTGTTCGTGAAGGTCGACAGATCCTCTAGATTTAA
GAAGGAGATATACATATG
>ycgX
GAGGCAATTCCGACGTCTAAGAAACCATTATTATCATGACATTAACCTATAAAAATAGGCGTATCACGAGGCCCTTTCGTCTTCACCTC
GATGGATCCCACTGGGGTGAGCAAGGTTATAAAGTCATCTTTCATGCCCCCTAATTTCACCTTTATCCCGCCCCTTAGACAGAATGTTA
TAATCTGAACCCACAACAGCGTTTGTACCGGAGCTGGTCCGTTGCAAAATGCTCACCGGGAGAATTCAGATATTCCCTCTATCACACTC
TGTACATATCACAGCCAAAGGACAGACCAGATTGAAGTGGTAAAACCGATCGTGCACACTACGCAGTCTCACAACTAACAAGAGGTTAT
TATCCTGATGGATCAGGTCGTTGTTTTTCAAAAGATGTTTGAGCAAGTGAGAAAAGAGCAAAACTTCTCCTGGTTTTATTCAGAATTAA
AACATCACCGTATTGCACACTCGAGAGATCCTCTAGATTTAAGAAGGAGATATACATATG
>kdgR
GAGGCAATTCCGACGTCTAAGAAACCATTATTATCATGACATTAACCTATAAAAATAGGCGTATCACGAGGCCCTTTCGTCTTCACCTC
GATGGATCCCCGGCAAAAGATGGATTGCGACCACCCTGGTGCGAAATTATAATAAAAAACAGTTCTGATTTTTATAAAACACTCGCAAT
GAGGTGATAAATGGCTAACGCAGATCTGGATAAACAGCCTGATTCTGTATCTTCCGTGCTAAAAGTTTTTGGCATTTTGCAGGCGCTGG
GTGAAGAGCTCGAGAGATCCTCTAGATTTAAGAAGGAGATATACATATG
>lpxP
GAGGCAATTCCGACGTCTAAGAAACCATTATTATCATGACATTAACCTATAAAAATAGGCGTATCACGAGGCCCTTTCGTCTTCACCTC
GAGAAGAGACCGCCGATACCACGTTCAGTACCTGACCGATGTAGCCGCTTACGCAGACAATACACAGGGCGAGAAACATCCATAAATTA
ATCATAGCTCAACCTCCTGCCGCGCTAAAAAACCTGTAACTTCAGTCACTTTAATAGTAAGACAATCCTTAACCTCTGGCAATTTTTGC
TAAGCATCAATGAAAACAGCATGTTAAATGCAAGACTGTTGTGTACGGAAAAATATTTACTTTGCACGATTAATAATCAGTAGCTGAAA
GCAGTCAGCGAGTGATAAAGTACAGATACCGCATTCATGCTCCTCGTGGTTATGTCCTGACTAGTCTTTACACTCTTTACAGGAACCAT
TGTCGTACATGATGGCCCAACCAATTGAGTGTACCGCTGTCGACATGGAATCTTCTGCGATACAACAATTCGTATCTACAGAAGGTAAC
TATGTTTCCACAATGCAAATTTTCCCGCGGATCCTCTAGATTTAAGAAGGAGATATACATATG
>ssuE
GAGGCAATTCCGACGTCTAAGAAACCATTATTATCATGACATTAACCTATAAAAATAGGCGTATCACGAGGCCCTTTCGTCTTCACCTC
GATGGATCCGCAGCCATAACGCTGGACGTTGCACATACCACAGTTATAAACGCCGTCAATACACTTTTTTTCATCGTTATCATCCTGAT
CTCTTCGTAAATATGATAATGCCTTTAAATGACATTAGACTTAATAATAACAATAATTGTCCGCCATTTAAATCAAGCATACTAAATCC
AATAAGTGACAAATCTTTAATTTATAGATTTCATGAATATTTAAATATTAACTGAACGAGTAATTCATTGATTCAACATCTAATGACTA
TCCAATAAATAAATATGCGTGATATATAAAGCCCTTTCTTTAGTTTATTTCAGAAAAAGATACACAAAATTTATATTTGGAATTTTCT
TGTCTCTCCGACAGACTGACCGAATTGTTATCAATGTTAACAAAAAAAGAACAATTGGTTATAAGGAGAGAGTATGCGTGTCATCACCC
TGGCGGGTAGTCCTCGCTTTCCTTCTCGCTCCAGCTCCTTGCTGGAATATGCGCGGGAAAAACTAAACTCGAGAGATCCTCTAGATTTA
AGAAGGAGATATACATATG
>hyaA
GAGGCAATTCCGACGTCTAAGAAACCATTATTATCATGACATTAACCTATAAAAATAGGCGTATCACGAGGCCCTTTCGTCTTCACCTC
GAGTCTTATTTGGCGGAAGCGCAGAGATTCGAACTCTGGAACCCTTTCGGGTCGCCGGTTTTCAAGACCGGTGCCTTCAACCGCTCGGC
CACACTTCCGGAATGACGCGCACTATAAACATCCCGATGCGGCGTGTAAACCCCTAATTTGTTTGTTTGCCTGAAAAACAGCCAAAAGT
GCATTGATAGCGTGAAATAACAGCAGATTGATCATTTCATCACCATGAATTCCTTCTCTTTTACTCGTTTAGCAACCGGCTAAACATCC
CCACCGCCCGGCCAAAAGAAAAATAGGTCCATTTTTATCGCTAAAAGATAAATCCACACAGTTTGTATTGTTTTGTGCAAAAGTTTCAC
TACGCTTTATTAACAATACTTTCTGGCGACGTGCGCCAGTGCAGAAGGATGAGCTTTCGTTTTCAGCATCTCACGTGAAGCGATGGTTT
GCCTTGCTACAGGGACGTCGCTTGCCGACCATAAGCGCCCGGTGTCCTGCCGGTGTCGCAAGGAGGAGAGACGTGCGATATGAATAACG
AGGAAACATTTTACCAGGCCATGCGGCGTCAGGGCGTTACCCGGCGCAGCTTTCTCAAATATTGTAGTCTGGCTGCCACGGGATCCTCT
AGATTTAAGAAGGAGATATACATATG
>yaaH
GAGGCAATTCCGACGTCTAAGAAACCATTATTATCATGACATTAACCTATAAAAATAGGCGTATCACGAGGCCCTTTCGTCTTCACCTC
GATGGATCCCTGGGAGGCCCTGTAGGTGCGGCACTAAATGGGGTTAAAGCGGTCAGCGGCAGCGCCTATCGCGTGACGATTCCAGCCGT
ACTGCAAATCGCCTGCCTGCGCCGGATGGTTAGCGCCACTCAGGTCTGATACCGGAAACCGGGGAAAATTTTCATTTTCCCCTCAAAAG
ATCGTAGACACTGCCCCACTGGCTGATTATTATGCCGCGCCCTGAAAACACTACAGTTATTCAGGGAAATTATTTCACCATTCATTCGA
TGATGATTTTTGAGGAATTATGGGCAACACTAAGTTGGCTAATCCGGCACCGCTGGGCCTGATGGGCTTCGGCATGACCACCATTCTGC
TTAACCTGCACAACGTGGGTCTCGAGAGATCCTCTAGATTTAAGAAGGAGATATACATATG
>fimD
GAGGCAATTCCGACGTCTAAGAAACCATTATTATCATGACATTAACCTATAAAAATAGGCGTATCACGAGGCCCTTTCGTCTTCACCTC
GAGCCTTCTGATGCAGGAAGCAATATTACTTACCGAACAATAAATGATTATGGCGCACTTACCCCCAAAATGACGGGCGTAATGGAATA

ACGCAGGGGGAATTTTTCGCCTGAATAAAAAGAATTGACTGCCGGGTGATTTTAAGCCGGAGGAATAATGTCATATCTGAATTTAAGAC
TTTACCAGCGAAACACACAATGCTTGCATATTCGTAAGCATCGTTTGGCTGGTTTGGATCCTCTAGATTTAAGAAGGAGATATACATAT
G
>cca
GAGGCAATTCCGACGTCTAAGAAACCATTATTATCATGACATTAACCTATAAAAATAGGCGTATCACGAGGCCCTTTCGTCTTCACCTC
GAGCTTGCTGCTCGGTCTGGTACTGCCGCACCTGATCCCAAGCCGCAAACGCAAAGATCGCTGGATGAACTAAATCGCCTTCTTCGTTG
CACTGTCTTATTATTAGGAATTGTGTATTTGACGAAGGGATGATGGCGTGAAGATTTATCTGGTCGGTGGTGCTGTTCGGGATGCATTG
TTAGGGCTACCGGTCAAAGACAGAGATTGGGTGGTGGTCGGGATCCTCTAGATTTAAGAAGGAGATATACATATG
>ylcE
GAGGCAATTCCGACGTCTAAGAAACCATTATTATCATGACATTAACCTATAAAAATAGGCGTATCACGAGGCCCTTTCGTCTTCACCTC
GAGGCGGGATAAGGTTCAAAAACATCATGGCTAATTTTATGTTTAATTTTTGATTCCCCAGCCCTTGTAGATGCCGCGTTCAGGCTAGC
TTCGTTCAAATCCATTAAAGATATCAGACTACTCTCAGGTACGTGAGTAAGGTAAAACCCAGTTCCAACACCAATATCCAGATGGTTGT
TACCTAAATGTTCCAGAAAGTGTGGAAGAAGGTGTTCCTTTGTAGGACATCCCCATGCAAGCCGATTTGATACTCCCAAAACCCACCAG
TCATAAAGCTTTAGGGTAAGTGGTGTGTAAATTCTAGCCCCATCATCTGTGTTTTTTTATTAATTTCACCATGTTATAGTTTTATTTGT
GAATTAAATCAATTATGGCAATGAATTACAAGGGGTTAAATGCTGCCGCAGCATAGCGATATTGAAATAGCCTGGTATGCTTCAATACA
GCAGGAGCCGAATGGCTGGAAGACCGTCAGGATCCTCTAGATTTAAGAAGGAGATATACATATG
>fumB
GAGGCAATTCCGACGTCTAAGAAACCATTATTATCATGACATTAACCTATAAAAATAGGCGTATCACGAGGCCCTTTCGTCTTCACCTC
GATGGATCCTTATTCTGCCGGGGTTGATTGGTGTGAGCGTATCGTGCGTCTTCGGCTGGATCTTCGCCGCGATGTACGGGTTCTTATAA
ATGCACTTTGCGTGCCGCCCGTGACTACGCGGCACGCCATTTTCGAATAACAAATACAGAGTTACAGGCTGGAAGCTATGTCAAACAAA
CCCTTTATCTACCAGGCACCTTTCCCGATGGGGAAAGACAATACCGAATACTATCTACTCACTTCCGATTACGTTAGCGTTGCCGACTC
GAGAGATCCTCTAGATTTAAGAAGGAGATATACATATG
>ycgL
GAGGCAATTCCGACGTCTAAGAAACCATTATTATCATGACATTAACCTATAAAAATAGGCGTATCACGAGGCCCTTTCGTCTTCACCTC
GAGTTACCCGCAGCAAAGCTCGTGGAGGTTAGCAGTGCAAGCAGCAATACAGCCTTCCTGATGCTCTTGATTTTCATTGGTAGTCTCCT
GTTCAGAATGAGTACGATGAACTGTTATAATATAACAATCCCTAACGGGAGAATTCCAGTGGCTTTCCTGCGTTCGGTAAAATCACGTT
GCGGCATTTTTGCCGGGGGGATTACTTATAAAGGGGGCATGGACTCAACAAACCGTAGAAATCTACAGCTGGCTGGCTTGTGGCGCGGG
TTTCATTGATGTTAGCTTATGCCAAAACCTGGCATCTTAAAGAGTAAAAGTATGTTTTGTGTGATTTATCGAAGCAGCAAGCGTGACCA
GACCGGATCCTCTAGATTTAAGAAGGAGATATACATATG
>rrnB
GAGGCAATTCCGACGTCTAAGAAACCATTATTATCATGACATTAACCTATAAAAATAGGCGTATCACGAGGCCCTTTCGTCTTCACCTC
GAGCGCCAGGAGCTGAACAATTATTGCCCGTTTTACAGCGTTACGGCTTCGAAACGCTCGAAAAACTGGCAGTTTTAGGCTGATTTGGT
TGAATGTTGCGCGGTCAGAAAATTATTTTAAATTTCCTCTTGTCAGGCCGGAATAACTCCCTATAATGCGCCACCACTGACACGGAACA
ACGGCAAACACGCCGCCGGGTCAGCGGGGTTCTCCTGAGAACTCCGGCAGAGAAAGCAAAAATAAATGCTTGACTCTGTAGCGGGAAGG
CGTATTATGCACACCCCGCGCCGCTGAGAAAAAGCGAAGCGGCACTGCTCTTTAACAATTTATCAGACAATCTGTGTGGGCACTCGAAG
ATACGGATTCTTAACGTCGCAAGACGAAAAATGAATACCAAGTCTCAAGAGTGAACACGTAATTCATTACGAAGTTTAATTCTTTGAGC
GTCAAACTTTTAAATTGAAGAGTTTGATCATGGCTCAGATTGAACGCTGGCGGCAGGCCTAACACATGCAAGTCGAACGGTAACAGGAA
ACAGCTTGCTGTTTCGCTGACGAGTGGCGGACGGGTGAGTAATGCTCGAGAGATCCTCTAGATTTAAGAAGGAGATATACATATG
>rrnC
GAGGCAATTCCGACGTCTAAGAAACCATTATTATCATGACATTAACCTATAAAAATAGGCGTATCACGAGGCCCTTTCGTCTTCACCTC
GAGCGAGTACCAATTCGCGGTCGCGGTAAAACCATCCCTTTTGCCGTTAACGTTTTGACCGCTTCGCGTACCGCTGTACGACTCACTCC
AAATTGCTCGCCCAGCTCAATCTCACCGGGCAAAATGGTGCCGGGTTCATATTCACCTTTTAAGATCCGTTGCGCCAGCTTCTCAGCCA
GAACATACGAAAGGTTTTTCTGTGCAGCTAACTGTTGTGCGCTTAAAGGCATTACTTATCTTCCTTTTTCTTTTTATTCCTCCTTAGTA
TGCCACCAGGAAGTGTGATTACGGTTGCAAAAACGGCAAATTGCTTGTTTTATGGCACATTAACGGGGCTTTTGCTGAAAAAATGCGCG
GTCAGAAAATTATTTTAAATTTCCTCTTGTCAGGCCGGAATAACTCCCTATAATGCGCCACCACTGACACGGAACAACGGCAAACACGC
CGCCGGGTCAGCGGGGTTCTCCTGAGAACTCCGGCAGAGAAAGCAAAAATAAATGCTTGACTCTGTAGCGGGAAAGCGTATTATGCACA
CCCCGCGCCGCTGAGAAAAAGCAAAGCGGCACTGCTCTTTAACAATTTATCAGACAATCTGTGTGGGCACTCGAAGATACGGATTCTTA
ACGTCGCAAGACGAAAAATGAATACCAAGTCTCAAGAGTGAACACGTAATTCATTACGAAGTTTAATTCTTTGAGCATCAAACTTTTAA
ATTGAAGAGTTTGATCATGGCTCAGATTGAACGCTGGCGGCAGGCCTAACACATGCAAGTCGAACGGTAACAGGAAACAGCTTGCTGTT
TCGCTGACGAGTGGCGGACGGGTGAGTAATGCTCGAGAGATCCTCTAGATTTAAGAAGGAGATATACATATG
>rrnA
GAGGCAATTCCGACGTCTAAGAAACCATTATTATCATGACATTAACCTATAAAAATAGGCGTATCACGAGGCCCTTTCGTCTTCACCTC
GAGAAACGGATACGCGCAAAGAAGTTGTCTATACCGATTGGGAGCAGGTGGCGAATTTCGCCCGAGAAATCGCCCATTTAACCGACAAA
CCGACGCTGAAATAAGCATAAAGAATAAAAAATGCGCGGTCAGAAAATTATTTTAAATTTCCTCTTGTCAGGCCGGAATAACTCCCTAT
AATGCGCCACCACTGACACGGAACAACGGCAAACACGCCGCCGGGTCAGCGGGGTTCTCCTGAGAACTCCGGCAGAGAAAGCAAAAATA
AATGCTTGACTCTGTAGCGGGAAGGCGTATTATGCACACCCCGCGCCGCTGAGAAAAAGCGAAGCGGCACTGCTCTTTAACAATTTATC
AGACAATCTGTGTGGGCACTCGAAGATACGGATTCTTAACGTCGCAAGACGAAAAATGAATACCAAGTCTCAAGAGTGAACACGTAATT
CATTACGAAGTTTAATTCTTTGAGCGTCAAACTTTTAAATTGAAGAGTTTGATCATGGCTCAGATTGAACGCTGGCGGCAGGCCTAACA
CATGCAAGTCGAACGGTAACAGGAAACAGCTTGCTGTTTCGCTGACGAGTGGCGGACGGGTGAGTAATGCTCGAGAGATCCTCTAGATT
TAAGAAGGAGATATACATATG
>rrnG
GAGGCAATTCCGACGTCTAAGAAACCATTATTATCATGACATTAACCTATAAAAATAGGCGTATCACGAGGCCCTTTCGTCTTCACCTC
GATGGATCCGGTGAATTGGTTCCGGGTAAAGTGATTCGCCTGGAAGTTAATGAAGACCGGATTGTCGCCGTCCAGTAAATGATAAAACG

```
AGCCCTTCGGGGCTCGTTTTTGTCTATAAGTTAGACGGAAAAGACTATATTTAAGATGTTTTGCCTGAAAAGTGAGCGAACGATAAAGT
TTTTATATTTTTCGCTTGTCAGGCCGGAATAACTCCCTATAATGCGCCACCACTGACACGGAACAACGGCAAACACGCCGCCGGGTCGG
CGGGGTTCTCCTGAGAATCTCAACAGAGAAAAGCAAAGAAATGCTTGACTCTGTAGCGGGAAGGCGTATTATGCACACCCCGCGCCGCT
GAGAAAAAGCGAAGCGGCACTGCTCTTTAACAATTTATCAGACAATCTGTGTGGGCACTCGAAGATACGGATTCTTAACGTCGCAAGAC
GAAAAATGAATACCAAGTCTCAAGAGTGAACACGTAATTCATTACGAAGTTTAATTCTTTGAGCATCAAACTTTTAAATTGAAGAGTTT
GATCATGGCTCAGATTGAACGCTGGCGGCAGGCCTAACACATGCAAGTCGAACGGTAACAGGAAGCAGCTTGCTGCTTCGCTGACCTCG
AGAGATCCTCTAGATTTAAGAAGGAGATATACATATG
>rrnD
GAGGCAATTCCGACGTCTAAGAAACCATTATTATCATGACATTAACCTATAAAAATAGGCGTATCACGAGGCCCTTTCGTCTTCACCTC
GATGGATCCATAATGCGCCTCCGTTGAGACGACAACGTGAAACACTTCACAGGATGGTCGGAACAACGAAGAGAAAAAAATCCTGAAAT
TCAGGGTTGACTCTGAAAGAGGAAAGCGTAATATACGCCACCTCGCGACAGTGAGCTGAAAGCCGCGTCGCAACTGCTCTTTAACAATT
TATCAGACAATCTGTGTGGGCACTCGAAGATACGGATTCTTAACGTCGCAAGACGAAAAATGAATACCAAGTCTCAAGAGTGAACACGT
AATTCATTACGAAGTTTAATTCTTTGAGCATCAAACTTTTAAATTGAAGAGTTTGATCATGGCTCAGATTGAACGCTGGCGGCAGGCCT
AACACATGCAAGTCGAACGGTAACAGGAAACAGCTTGCTGTTTCGCTGACGAGTGGCGGACGGGTGAGTAATGCTCGAGAGATCCTCTA
GATTTAAGAAGGAGATATACATATG
>amn
GAGGCAATTCCGACGTCTAAGAAACCATTATTATCATGACATTAACCTATAAAAATAGGCGTATCACGAGGCCCTTTCGTCTTCACCTC
GAGCGCCATTTATTTGCTGGCTGGATGCCTGATTTCCGCAATGACCGCTTTGTTGATGAAAGACAGTCAACGCGCTTGATAGCCTGGCG
AAGATCATCCGATCTTCGCCTTACACTTTTGTTTCACATTTCTGTGACATACTATCGGATGTGCGGTAATTGTATGGAACAGGAGACAC
ACATGAATAATAAGGGCTCCGGTCTGACCCCAGCTCAGGCACTGGATAAACTCGACGCGCTGTATGAGCAAGGATCCTCTAGATTTAAG
AAGGAGATATACATATG
>rrnH
GAGGCAATTCCGACGTCTAAGAAACCATTATTATCATGACATTAACCTATAAAAATAGGCGTATCACGAGGCCCTTTCGTCTTCACCTC
GAGGCCTGAAGCAGAAAACGCGGCGGATTGGGTGTTAAATAGCCTGGCAGACCTGCCGCAAGCGATAAAAAAGCAGCAAAAACCGGCAC
AATGATTAAAAGATGAGCGGTTGAAATAAAAATGCATTTTTCCGCTTGTCTTCCTGAGCCGACTCCCTATAATGCGCCTCCATCGACAC
GGCGGATGTGAATCACTTCACACAAACAGCCGGTTCGGTTGAAGAGAAAAATCCTGAAATTCAGGGTTGACTCTGAAAGAGGAAAGCGT
AATATACGCCACCTCGCGACAGTGAGCTGAAAGCCGCGTCGCAACTGCTCTTTAACAATTTATCAGACAATCTGTGTGGGCACTCGAAG
ATACGGATTCTTAACGTCGCAAGACGAAAAATGAATACCAAGTCTCAAGAGTGAACACGTAATTCATTACAAAGTTTAATTCTTTGAGC
ATCAAACTTTTAAATTGAAGAGTTTGATCATGGCTCAGATTGAACGCTGGCGGCAGGCCTAACACATGCAAGTCGAACGGTAACAGGAA
ACAGCTTGCTGTTTCGCTGACGAGTGGCGGACGGGTGAGTAATGCTCGAGAGATCCTCTAGATTTAAGAAGGAGATATACATATG
>psd
GAGGCAATTCCGACGTCTAAGAAACCATTATTATCATGACATTAACCTATAAAAATAGGCGTATCACGAGGCCCTTTCGTCTTCACCTC
GATGGATCCAGGGAAAATCGCGGAAACCCGTTTCGAAAACTATCACCGTATTCTGGAAAGCATGGCGCAGGTAAAAACGCGTAAAAACT
TTTCTGATACGGATGACTGACAATTAAGCTAACCGTCGTTAGAATCGTCCCCTTTTTTTCAGGATCCGGCATGTAGGCCGGATCAGGAA
CGACAAAACAATGGCCTGGAGGCTACCTTGTTAAATTCATTTAAACTTTCGCTACAGTACATTCTGCCGAAACTATGGCTTACTCGCCT
GGCGGGTTGGGGCGCAAGCAAGCGGGCAGGATCTCGAGAGATCCTCTAGATTTAAGAAGGAGATATACATATG
>b1142
GAGGCAATTCCGACGTCTAAGAAACCATTATTATCATGACATTAACCTATAAAAATAGGCGTATCACGAGGCCCTTTCGTCTTCACCTC
GATGGATCCTTTCCTTGCTTGCTGGCTGTGTGAGAACTCCAGCATACCACCGAGCCTGAAGTGGTGAAAAGACAGGCACATAACAGCTA
AGTATTTTCAACCAGAGAGAATCCTTAGCGTTGTGGTGAATGCGGCTCAGCGCACGCGGGTTAAGGTTGAGGCTGACAATCGACCTTCT
GTGGATACCCACCCGCCTGGTGTGCAACCTTCCTCGAGAGATCCTCTAGATTTAAGAAGGAGATATACATATG
>mog
GAGGCAATTCCGACGTCTAAGAAACCATTATTATCATGACATTAACCTATAAAAATAGGCGTATCACGAGGCCCTTTCGTCTTCACCTC
GAGTGGCGGAAGGTATCCGTAAGTTTGCTATTGACCAGGAAAAACTGGAAAAAATGATCGGCGATCTGCTGTAATCATTCTTAGCGTGA
CCGGGAAGTCGGTCACGCTACCTCTTCTGAAGCCTGTCTGTCACTCCCTTCGCAGTGTATCATTCTGTTTAACGAGACTGTTTAAACGG
AAAAATCTTGATGAATACTTTACGTATTGGCTTAGTTTCCATCTCTGATCGCGCATCCAGCGGCGTTTATCAGGATAAAGGCATCCCTG
CGAGATCCTCTAGATTTAAGAAGGAGATATACATATG
>ycgE
GAGGCAATTCCGACGTCTAAGAAACCATTATTATCATGACATTAACCTATAAAAATAGGCGTATCACGAGGCCCTTTCGTCTTCACCTC
GATGGATCCAGCGTGGCCAGAGAAAAAATAATTTTCAGCACATTCTTTCACATGATTTCAGTAAATCATCGTGAAAATGAATACGTTAC
GTTTAAATTTAATCCACATTAAAAATAATGTTTCATCCAATTTAGCTGAAATCTTATTTATACTTTGTACAATTTTAGCGTATATTGTA
CAAAGTATTATTGGGTCGTGTACAGGCGACGGAGATTTGTGACCGCAAGGAGGAATTGTGGCTTATTACAGCATTGGTGATGTTGCTGA
ACGTTGCGGGATTAATCCTGTCACTCTCCGGGCCTGGCAACGCCGCTACGGTTTGTTACTCGAGAGATCCTCTAGATTTAAGAAGGAGA
TATACATATG
>wrbA
GAGGCAATTCCGACGTCTAAGAAACCATTATTATCATGACATTAACCTATAAAAATAGGCGTATCACGAGGCCCTTTCGTCTTCACCTC
GATGGATCCCAGACCCGCGTTTGGTAATATCCTGCAACAGCTTCTTAGCGAGAATAGATATCATCTCCAATAATTATCCATAAGCCGCT
CTTATGATTAAAAAAACGCGATGAGATAAAAATGTGTCATTTTGCGACAAAATTACGTGCTTGTGAAAGTTGTTATAAATCAAATAAGT
GGTTGTGAAATTTGCACTCTGAAAAGGACGTCTTATCTTTAAATAAGTGGTAGCGAATCGCTACGGAATAGAGATAACACGAGGAGTGG
TTAGAAATGGCTAAAGTTCTGGTGCTTTATTATTCCATGTACGGACATATTGAAACGATGGCACGCGCAGTCGCTGAGGGTGCAAGCAA
AGTGGATGGCGCTGAAGCTCGAGAGATCCTCTAGATTTAAGAAGGAGATATACATATG
>fldA
```

GAGGCAATTCCGACGTCTAAGAAACCATTATTATCATGACATTAACCTATAAAAATAGGCGTATCACGAGGCCCTTTCGTCTTCACCTC
GATGGATCCGCTGCGTAGTCAGGGCATCGTTTAAATTTCCACTTTCATGTAGCACAGTGTGCAGTCCTGCTCGTTTGCTGATTCCGCAA
GACTGCCTGTTCTGCTATGATTGCCTTTATCCGTGGGCAATTTTCCACCCCCATTTCAATAAGTTTCAAGAGGTTATTTCACTCATGGC
TATCACTGGCATCTTTTTCGGCAGCGACACCGGTAATACCGAAAATATCGCAAAAATGATTCAAAAACAGCTTGGTAAAGACGTTGCCG
ATGTCCCTCGAGAGATCCTCTAGATTTAAGAAGGAGATATACATATG
>ynaI
GAGGCAATTCCGACGTCTAAGAAACCATTATTATCATGACATTAACCTATAAAAATAGGCGTATCACGAGGCCCTTTCGTCTTCACCTC
GATGGATCCCGGCGCTTACTGCTGAATTCACTGTCGGCGAAGGTAAGTTGATGACTCATGATGAACCCTGTTCTATGGCTCCAGATGAC
AAACATGATCTCATATCAGGGACTTGTTCGCACCTTCCTTAGCTAATGTTGCTACTTCCTTCCCTGACTATTGATGATCTGAATGAATT
GTCGCGGTAAAATAGCCCCGCCAGGGGTTTCCGGCATGACGACAATTATCTGGAGGAGCCTGATAAAACAATGATCGCTGAACTGTTTA
CAAATAATGCGCTTAATCTGGTCATTATTTTCGGTAGCTGCGCAGCATTGATTCTGATGAGCTTTTGGTTTCGCCCTCGAGAGATCCTC
TAGATTTAAGAAGGAGATATACATATG
>b1367
GAGGCAATTCCGACGTCTAAGAAACCATTATTATCATGACATTAACCTATAAAAATAGGCGTATCACGAGGCCCTTTCGTCTTCACCTC
GAGCCTTGCAGAAGCAGCCAGCATCGCCCAGGGTAAAGAGCAGCAGAAGCAGGATACGATAAAAAAGCCATAGCTGCCCAGCGGTGGTT
ATCGGAGTTCGGGGGAGAAAGGGGGGAAAAGGCAAGATGGAAGCGAGAAAAACTCAGGTTGCCACCGATACCGGAACCAGAAATAGACC
CGGTGCTTAAGGAGTTGTTGTACGCCTATTCGGTAATATCCCGTGCCCGACGTTATGCTGGAAGGATCCTCTAGATTTAAGAAGGAGAT
ATACATATG
>ftsJ
GAGGCAATTCCGACGTCTAAGAAACCATTATTATCATGACATTAACCTATAAAAATAGGCGTATCACGAGGCCCTTTCGTCTTCACCTC
GATGGATCCAACAACTGGCTTGAGCGGATGTGCCAGACCTTTCAGGTGCTGTTTTTGTTTAGTACTCAGATTCATCGTATTTTTTGCTT
ACGTTGGGATTGAAAACGGGTCATTCTACCGCCATCTCCCATATATCACCAAATAGGCGCGTAAAAATTTACGCAATTGGTTACGATGA
GTTATCCCCATGGGAAAGTTAAATGACAGGTAAGAAGCGTTCTGCCAGCTCCAGCCGCTGGCTTCAGGAACACTTTAGCGATAAATATG
TTCAACAGGCACAGAAAAAGGGGTTACGTTCCCGCTCGAGAGATCCTCTAGATTTAAGAAGGAGATATACATATG
>yccR
GAGGCAATTCCGACGTCTAAGAAACCATTATTATCATGACATTAACCTATAAAAATAGGCGTATCACGAGGCCCTTTCGTCTTCACCTC
GAGAGCCCGGCTGTAGTGTTTTCCGTAGAGACACGCGCAATTTTACTTGCTGCGGATGAGAACGACGAAGAACGATGTGCATAGCCTGA
AGTGTACATAATCAATCCAGCCCCTGTGAGTTACTGTATGGATGTACAGTACATCCAGTGACAACAAAGATCAACCCTATTTTCGGAAA
GAGCCTCGCAAATTTTGTCGTTGGTGACGGGAAAACATAAATTAATCTTGCCCCTTAAGAATAAGTTGCCTATTTTCGTAGTTAACGGA
TCCGTTAATGTGAATCATTCTTTTATGTTATGATTTTAAAAGGAATTTTATGAAAAGCCTCTCCTATAAGCGGATCTATAAATCACAAG
AATACCTGGCAACGTTGGGCACAATTGAATACCGATCATTGTTTGGCAGAGATCCTCTAGATTTAAGAAGGAGATATACATATG
>b1331
GAGGCAATTCCGACGTCTAAGAAACCATTATTATCATGACATTAACCTATAAAAATAGGCGTATCACGAGGCCCTTTCGTCTTCACCTC
GAGGGCGAAACCAAAAGCTCATCAGAATCAATGCTGCGCAGCTACCGAAAATAATGACCAGATTAAGCGCATTATTTGTAAACAGTTCA
GCGATCATTGTTTTATCAGGCTCCTCCAGATAATTGTCGTCATGCCGGAAACCCCTGGCGGGGCTATTTTACCGCGACAATTCATTCAG
ATCATCAATAGTCAGGGAAGGAAGTAGCAACATTAGCTAAGGAAGGTGCGAACAAGTCCCTGATATGAGATCATGTTTGTCATCTGGAG
CCATAGAACAGGGTTCATCATGAGTCATCAACTTACCTTCGCCGACAGTGAATTCAGCAGTAAGCGCCGCTCGAGAGATCCTCTAGATT
TAAGAAGGAGATATACATATG
>yccE
GAGGCAATTCCGACGTCTAAGAAACCATTATTATCATGACATTAACCTATAAAAATAGGCGTATCACGAGGCCCTTTCGTCTTCACCTC
GAGGTATTTGCGGGCAAGTCGACGATAGGCGGTCTTGATTGTCTTGAGATCGTCCGTCGGTTTCACGCCCATGATGGCGTAATAATCCT
TTAATTCCATAGCGTTATCTCGCGTAAATCAACACAAATTGAAGGAACCCCTGTAAGGTAACTCCTATAAGTGTAGGGTAATCCTCAAA
ATTTCATATGCCAACACAGAATATGTTATTGAAATCATCGCGGAGAGGAGGTCGCCATCAAGATGGGTTGCTGAACATATTTTAAACAG
GTGAAAAAGGGTGAGCGATTTTTGATAGTTGAACCAGGCACTTTAAGTTTAACTAGGGCGTCATTATTTATTAAATTTTATAGACGCTA
TATATGGGTAGTAATATACATGGAATTAGTTGCACTGCAAATAATTATTTGAAACAGGCCTGGAACGATATAAAAAATGAGTACGAAAA
AAATCAAACATATTCAATCACGCTTTTTGAAAACACACTGGTGTGTTTTATGCGGTTGGATCCTCTAGATTTAAGAAGGAGATATACAT
ATG
>yccF
GAGGCAATTCCGACGTCTAAGAAACCATTATTATCATGACATTAACCTATAAAAATAGGCGTATCACGAGGCCCTTTCGTCTTCACCTC
GATGGATCCTAATCCCGGCATTGAGGATCACCGCCCGATCGTAAGGGTGCTGTGCCAGACGTTTTCCAAGCGTTGTCGCTTTCAGTTCC
ATGACTCTGTAGATCCACACGAAATTATTTCTTATCAGTGTAACCGTCAGAGAAAACGTGCTCCAGCCCAAAAAAACGTTACAATTGCC
GCCCATTACTGAAAACCACAGTAAAGCGAGGTTTTATGCGTACCGTTTTGAACATTCTGAACTTTGTGCTTGGCGGATTTGCCACCACT
CTGGGCTGGCTGTTGGCGACTCTGGTCTCGAGAGATCCTCTAGATTTAAGAAGGAGATATACATATG
>yccA
GAGGCAATTCCGACGTCTAAGAAACCATTATTATCATGACATTAACCTATAAAAATAGGCGTATCACGAGGCCCTTTCGTCTTCACCTC
GATGGATCCTCGAATCTCTGCGCTTCCGCCAAATAAGATAAGGGGTTAGCTAAATGCTAACCCCTTTTTCTTTTGCCTGTCGAAATTCT
CAGGGCGTTATATTTGCTTAATGACCTGATAATCCGCTGTTAAACCTGTTCCAGATCAAATGCGTAAAGATGGGTAAAACTTCTGGGTG
CCCTTACGCATTATCATTATGCTGCTTAATTAATTACATCTGTCATAGAGAGTGACTCAATGGATCGTATTGTTAGTTCTTCACATGAC
CGTACATCACTGCTTAGCACCCATAAGGTGCTGCGTAATACCTATTTTCTGCTGAGCCTGACGCTCGAGAGATCCTCTAGATTTAAGAA
GGAGATATACATATG
>b1966
GAGGCAATTCCGACGTCTAAGAAACCATTATTATCATGACATTAACCTATAAAAATAGGCGTATCACGAGGCCCTTTCGTCTTCACCTC
GAGACGACTTCGACTTCGGTTTAAGTCTGGGTGCTGCATATCAAACTCTGATCGTTCAGATCGTCAGGTTGGAGTTGGTTTAAATGATC

GTAACCATAGCAACAGAAATGCTGGTGGTGAAACCGCTGAAGCATGGACTGTTGGTGCAAAATATGATGCTAATAATGTGTATTTAGCA
GCTATGTACGCTGAAACTCGCAATATGACCCCATATGGTGGTGGCGAGTTTGATAATGGTGAATCAAGATCAGCCATCGCTGGATCCTC
TAGATTTAAGAAGGAGATATACATATG
>b1788
GAGGCAATTCCGACGTCTAAGAAACCATTATTATCATGACATTAACCTATAAAAATAGGCGTATCACGAGGCCCTTTCGTCTTCACCTC
GATAGATCTGGGGCATGCTATGAACGATCAAATGTTTGTCGAGACACTGATTATCACGTCATCGTTTTTTGCCATTGCTGTTGTACTGG
TTTTGTCCGTTCTTCTTATCGAACGGACCGGCTAGTTACGCAGTGCGGCGGAAATTGACCAGTTCTGGTTGCGCAATGCGCAAATAATC
CGCGGTTTTCAAAATAACGGATTTATCCAGCATGCCTGCATTGAAGGCGATTTCATCAAATCGTTCGAAGAGTAGAGGGTCGGCAACCA
GCTCGAGAGATCCTCTAGATTTAAGAAGGAGATATACATATG
>yccM
GAGGCAATTCCGACGTCTAAGAAACCATTATTATCATGACATTAACCTATAAAAATAGGCGTATCACGAGGCCCTTTCGTCTTCACCTC
GATGGATCCCCCCTTCCACACGAAGAAATTACACGCAGTGTTGCCGCTCTGGAAGCATGGTTGCATAAGAAAGACCTGAACGCGATTTG
AATCAATATCAGCCAGTTAAGAACAACTAGAATAGTGGCATTTGGTATACAACTTAACGGACTGGCAAAATCATGGCAGAGAATAAAAG
AACCCGCTGGCAGCGGCGGCCAGGCACGACGGGCGGCAAATTACCGTGGAATGACTGGCGTAATGCCACGACCTGGCCTCGAGAGATCC
TCTAGATTTAAGAAGGAGATATACATATG
>cydD
GAGGCAATTCCGACGTCTAAGAAACCATTATTATCATGACATTAACCTATAAAAATAGGCGTATCACGAGGCCCTTTCGTCTTCACCTC
GATGGATCCTGCATGGCAGCACTTGATGCGGAACGCTACCTCGATGGTTTAGCTGACGCAAAATAATTTTACAAATCAGTAACAAAAGT
AAAGAAGGCGACACCATGCGACTATGGGTCGCCTTTATTTTTTCCCCGTTGTAACATTGCTCTGCAAATAATTCTGATAACTCACCTGC
TAAGCGTGCAATGAATAAATCTCGTCAAAAAGAGTTAACCCGCTGGTTAAAACAGCAAAGCGTCATCTCCCAACGTTGGCTGAATATTT
CTCGTCTGCTGGGCTCTCGAGAGATCCTCTAGATTTAAGAAGGAGATATACATATG
>ushA
GAGGCAATTCCGACGTCTAAGAAACCATTATTATCATGACATTAACCTATAAAAATAGGCGTATCACGAGGCCCTTTCGTCTTCACCTC
GAGAAACGATGTTCGGGCCTTGGTCGTTGACGCAGCCGCGCCCGCCACAGGCTGGGGTTGTTCACTCATTGCCATAGGAAAGTCACTTT
TTCAGGGTTGCGATGTAAAGAATGATCTTATTTGTGATTATTACCAGACTAACATACCTGTATGCGTCGTCTGAAGGAAGTCTCAACGC
CGAATACAGAATTTCTAATCTGGATGCAGATTTATCTTCACCGGACGCAGACTTGTCTATGATGTCGCGTCATACTATTTTTCAACACG
TTGAAATCAGGTCAGGGAGAGAAGTATGAAATTATTGCAGCGGGGCGTGGCGTTAGCGCTGTTAACCACATTTACACTGGCGAGTGAAA
CTGCTCTGGCGTATGAGCAGGAGGATCCTCTAGATTTAAGAAGGAGATATACATATG
>rmuC
GAGGCAATTCCGACGTCTAAGAAACCATTATTATCATGACATTAACCTATAAAAATAGGCGTATCACGAGGCCCTTTCGTCTTCACCTC
GAGCCAGCAAGAGATCCCGAATGCTGAGACGATGAAACAAACCGAAAGCCATGCGGTGAAAATCGTGGTGGAAGCGGCGCGTCGTCTGC
TGTAATTCTCTTCTCCTGTCTGAAGGCCGACGCGTTCGGCCTTTTGTATTTTTGCGTAGCGCCTCGCAGGAAATGCCTTTCCAACTGGA
CGTTTGTACAGCACAATTCTATTTTGTGCGGGTAAGTTGTTGCGTCAGGAGGCGTTGTGGATTTCTCAATCATGGTTTACGCAGTTATT
GCGTTGGTGGGTGTGGCAATTGGCTGGCTGTTTGCCAGTTGGATCCTCTAGATTTAAGAAGGAGATATACATATG
>dpiB
GAGGCAATTCCGACGTCTAAGAAACCATTATTATCATGACATTAACCTATAAAAATAGGCGTATCACGAGGCCCTTTCGTCTTCACCTC
GAGTCCTGAAGGTGCATGTTGTTATCGATTTGCAACGAATGTTGTTCAATGTTGCAAACTGATAACCTTTTATTTTCACTTGGGAGAAA
GGGGGTGATCGAGGTATATCTTTTTCTCCTTTCGCTATACATCCTAAGGAGTATTTCGGCGTGAAATTTTGATTTATTTCACATAGAGT
TAGTGGTTTTTTATTTATTTAATGATTTTAAGTTTTTTAATTAATGTAATTACGAAATGACTCGCAGGTTTAAGTGATTTAATTGATTT
AATGAATAAAATTTGCCACGATCATAATTAATATCTATGTATTTTGATTCAACATTTTAATTACATCCGTCAAAGAGGCTCGGGACAAC
CCGCAAGGAAAACAATGTTGCAGCTTAACGAGAATAAACAGTTTGCATTTTTCCAAAGACTGGCATTTCCGCTGCGTATCTTTTTGGGA
TCCTCTAGATTTAAGAAGGAGATATACATATG
>dapA
GAGGCAATTCCGACGTCTAAGAAACCATTATTATCATGACATTAACCTATAAAAATAGGCGTATCACGAGGCCCTTTCGTCTTCACCTC
GATGGATCCCACCAGATAATGTTGCGATGACAGTGTCAAACTGGTTATTCCTTTAAGGGGTGAGTTGTTCTTAAGGAAAGCATAAAAAA
AACATGCATACAACAATCAGAACGGTTCTGTCTGCTTGCTTTTAATGCCATACCAAACGTACCATTGAGACACTTGTTTGCACAGAGGA
TGGCCCATGTTCACGGGAAGTATTGTCGCGATTGTTACTCCGATGGATGAAAAAGGTAATGTCTGTCGGGCTAGCTTGCTCGAGAGATC
CTCTAGATTTAAGAAGGAGATATACATATG
>mngR
GAGGCAATTCCGACGTCTAAGAAACCATTATTATCATGACATTAACCTATAAAAATAGGCGTATCACGAGGCCCTTTCGTCTTCACCTC
GATGGATCCCGGTGGGTCAGAGTCGTCAGATTCATCATGATCCTCACTTGATCGTTTTTATAGTCGCGCCAGTGTGCCCGATAAAACAA
TACCATTATGTGATCAATTTCACTTGATTGAAATTATTTGTATTGTATTTGTATTATTTACAGGATGAATTTATCGGGCATAATGGTTA
AATTCGTATTAATGAGATACTAAAAATGGGACACAAGCCCTTATACCGGCAGATTGCCGATCGCATTCGTGAACAAATTGCGCGTGGCG
AGCTTAAACCTGGCTCGAGAGATCCTCTAGATTTAAGAAGGAGATATACATATG
>lolB
GAGGCAATTCCGACGTCTAAGAAACCATTATTATCATGACATTAACCTATAAAAATAGGCGTATCACGAGGCCCTTTCGTCTTCACCTC
GATGGATCCCCGGCGAAAACGATACACGTTCTCGCAGCGATACAGGTGCCGTTTTATGGTTGATACCGAGTGCTAAAAGGGTCATGTCT
GCGGGAAATAATACCAACGTTGATAGGGTTAGTCTGCTTGCATCATACAGGATGCGTAAGATCAATAAAAGAGAGCGCCCCCTTTTGGA
GTAATTGCCGAAAGCCGTTAGATTCTGGCAATTAAGACAACTTGAACATAGACGATAGCGGACGGTAACGCTAGCATTAAGGGTTATAA
CTGCAACGTATCTCAAGGACTTGTCATCACTATGCCCCTGCCCGATTTTCGTCTTATCCGCCTGCTACCGCTGGCTGCTCTTGTGCTCA
CTGCCTGTTCCGTCTCGAGAGATCCTCTAGATTTAAGAAGGAGATATACATATG
>b0360

GAGGCAATTCCGACGTCTAAGAAACCATTATTATCATGACATTAACCTATAAAAATAGGCGTATCACGAGGCCCTTTCGTCTTCACCTC
GAGATCTCGGTGGCAAAACCAATTTTTACGCCATTGCTGATTATTGTGCCAGGACGAATAAACGCATAATTACTAGACTGGCCCCCTGA
ATCTCCAGACAACCAATATCACTTAAATAAGTGATAGTCTTAATACTAGTTTTTAGACTAGTCATTGGAGAACAGATGATTGATGTCTT
AGGGCCGGAGAAACGCAGACGGCGTACCACACAGGATCCTCTAGATTTAAGAAGGAGATATACATATG
>rnhA
GAGGCAATTCCGACGTCTAAGAAACCATTATTATCATGACATTAACCTATAAAAATAGGCGTATCACGAGGCCCTTTCGTCTTCACCTC
GATGGATCCATCGAGAACGATCTGGCGTGTAATTGCAGTGCTCATAGCGGTCATTTATGTCAGACTTGTCGTTTTACAGTTCGATTCAA
TTACAGGAAGTCTACCAGAGATGCTTAAACAGGTAGAAATTTTCACCGATGGTTCGTGTCTGGGCAATCCAGGACCTGGGGGTTACGGC
GCTATTTCTCGAGAGATCCTCTAGATTTAAGAAGGAGATATACATATG
>yegE
GAGGCAATTCCGACGTCTAAGAAACCATTATTATCATGACATTAACCTATAAAAATAGGCGTATCACGAGGCCCTTTCGTCTTCACCTC
GAGCGCCAGCGATACCGATAATGACGCACTGATGAGACTGATCAGTCATATATTTAGCGACCTGATTAACCTGGATGTTAGGAAGGGGG
CGACGAAGCGCCAAACGCGGCAATTATAGGGATTTCATCCGTCTGATACCAGTCGAATAGCGTTGCCGCGCGCTCAGAGTTAATTGTTG
ACAAAGAATTCCCGGGGGCAAATTACGTTGATCAGTTTTATGTAAGGTAAAAAATGTTAAACGCAGTTGCGCAAATTATCCGCCTTTAC
GTCACTTTATGAGCAATTCGCATATAAAATGTAAAACTTTTGTACTAGCATAAACACAGAAACGAATACTGGCGACCAGGTCTTGCGGA
TAAAGCGGTAATGAGCAAACAATCACAGCATGTATTAATTGCCCTGCCCCACCCGCTGCTTCACCTGGTCAGTTTAGGTTTAGTCTCGT
TTATCTTTACCCTTTTCTCGCTTGAGCTTTCGCAGTTTGGCACCCAACAGATCCTCTAGATTTAAGAAGGAGATATACATATG
>garD
GAGGCAATTCCGACGTCTAAGAAACCATTATTATCATGACATTAACCTATAAAAATAGGCGTATCACGAGGCCCTTTCGTCTTCACCTC
GAGCGATCGGCGTAGTTAACGGCGGTAACAATAAAAATAATCAGTAATATTAAATAGCGGGTATGCACGCCTTTCTTTTTTTCGTCAAC
GGTGTCCAGAATCATTTTATTTACCTCGGGTACTTATGCTGATTTTTATTATTATGGGGAAGGTGTTATTTATGAGTTTCATTTATGCC
GTAACGACAATGAACTCGGGAATTAGTATAAGCAGCGCGAGAATAATAATCATTGTGCAAATGCTAATTTAATTAATACTATTTAAATA
TTATTTTGAGCATATGCACATAAGGTTGCGCGCTAAAGCACAGATTTGCGCTTTACCTTACCGGGCGGCACTGCAATCCCTGAAATGAT
TGACATTGATCACATTTCTGCGTTTAAACTCCTGACATTCTTATTTCACCCAATGAAGTCATTTATTTTTAAATGAGACCAGGTCCTCA
TTTTAATAACCCCTGGCTGGAGAATATTGCACAATGGCCAACATCGAAATCAGACAAGAAACGCCAACTGCGTAGATCTTCTAGATTTA
AGAAGGAGATATACATATG
>ybiU
GAGGCAATTCCGACGTCTAAGAAACCATTATTATCATGACATTAACCTATAAAAATAGGCGTATCACGAGGCCCTTTCGTCTTCACCTC
GATGGATCCGGCGGTGCTGGATAACACATCCCCTTTTAACAGCTGACCTCTCTACCGGAGCCGTGCGCTCCGGTTTTCCCTTCTTTTTA
CATCCTGTACATCACATCAAATCATTAAACTTGCATTAACTTATTTATAACTTATATTGTCACATGCAAATCAGAATGGTTTCGAATGA
AAGTTAATGAGAGGTGGTTATGGCTTCTACTTTTACCAGCGACACATTGCCTGCCGATCACAAAGCAGCTATCCGTCAGATGAAGCACG
CGCTGCTCGAGAGATCCTCTAGATTTAAGAAGGAGATATACATATG
>lon
GAGGCAATTCCGACGTCTAAGAAACCATTATTATCATGACATTAACCTATAAAAATAGGCGTATCACGAGGCCCTTTCGTCTTCACCTC
GAGCGACGAGTCGGTAATTGATGGTCAAAGCAAACCGTTGCTGATTTATGGCAAGCCGGAAGCGCAACAGGCATCTGGTGAATAATTAA
CCATTCCCATACAATTAGTTAACCAAAAAGGGGGGATTTTATCTCCCCTTTAATTTTTCCTCTATTCTCGGCGTTGAATGTGGGGGAAA
CATCCCCATATACTGACGTACATGTTAATAGATGGCGTGAAGCACAGTCGTGTCATCTGATTACCTGGCGGAAATTAAACTAAGAGAGA
GCTCTATGAATCCTGAGCGTTCTGAACGCATTGAAATCCCCGTATTGCCGCTGCGCGATGTGGTGGTTTATCCGCACATGGTCATCCGG
ATCCTCTAGATTTAAGAAGGAGATATACATATG
>nrfA
GAGGCAATTCCGACGTCTAAGAAACCATTATTATCATGACATTAACCTATAAAAATAGGCGTATCACGAGGCCCTTTCGTCTTCACCTC
GAGGCCTCGTACTGCTGAGGGTTTATCAGGCAACGGTCTGCGATGTTGGCAGGAATGGTGTGTTTGTGAATTTGGCTCATGCTTTTGTT
CTCCTTGTAGGATGTTAATAATATGTGGCATAAGCGTTAAATGTAGGGGTATTGGCAGTTTTGTTTAGTATTTGGGCGACAGATCACGC
AAAAGTAGAATTGTGCAAATAAACGGCAGGGTAATTTTTGAAGGTCAGCAACAAAAGTTGATTAATTCTTTGAGGAACATGCAGTTATG
CATGCTGTTGAAAAAGAGGAAGATACTGACTAACTCTAAAGTGGTATTTTACATGCACTTACAATTGATTAAAGACAACATTCACAGTG
TGGTTATTTGTTACACATAGGGGCGAGCAATGTCATGACAGTGTAGGTGCGGTTACTGTCGTGAAAAACAATAAAAACCGCCATTGCAA
CAATGGCGCAATTCGGATGAAGCCCCTATGACAAGGATAAAAATAAACGCACGCCGTATCTTCAGCTTATTGATTCCTTTTTTCTTTTT
CACTTCTGTTCACGCTGAACAAACGGCTGCTGGATCCTCTAGATTTAAGAAGGAGATATACATATG
>yegP
GAGGCAATTCCGACGTCTAAGAAACCATTATTATCATGACATTAACCTATAAAAATAGGCGTATCACGAGGCCCTTTCGTCTTCACCTC
GAGCTCTCGACGCCGAACAGTCATTTATCCGCGCCGTTTATGGCGTCGGTTACCGCTGGGAAGCCGACGCCTGCCGCATCGTTTAGTTT
TAGCGACATTATTTTGTTAGCCGGAGATGGCGTTCAGACCAGATCCGGCAACATTATCCCACGCATGGTCAGCAAACTGACATTATCTC
TCCGCCCTGGCAGGCATCAGAAAATCGCTCATACTTTAATCGGTAAACAGCACCTTTAGATGCTGTTTTCGATACACAATTTCAATCAA
GGAGTCATTATGGCTGGTTGGTTTGAACTCAGTAAGAGCAGTGATAATCAGTTCCGGTTTGTGCTAAAAGCGGATCCTCTAGATTTAAG
AAGGAGATATACATATG
>yegW
GAGGCAATTCCGACGTCTAAGAAACCATTATTATCATGACATTAACCTATAAAAATAGGCGTATCACGAGGCCCTTTCGTCTTCACCTC
GATGGATCCAGGCATTCGTTGATGGGATTAAAGAAACGCCTTAAATAAGAAGCAGTGATATACTCCCACCAGACTCCCCGGCAGGAAAA
AGACAAATGGAACAAGCGCATACCCAGCTGATTGCCCAACTGAACGAGCGTATTTTAGCGGCGGATAACACGCCGCTTCTCGAGAGATC
CTCTAGATTTAAGAAGGAGATATACATATG
>yegT
GAGGCAATTCCGACGTCTAAGAAACCATTATTATCATGACATTAACCTATAAAAATAGGCGTATCACGAGGCCCTTTCGTCTTCACCTC
GAGGCAACTGCGCAATATCTGTCATTACTATGCTCCCGTAAATTCCGATTGGATATCGGCTATGGATTGTCCTGGCCCGCCTTTTGCGG

GCAATCATAATCCTGGTCGTTACGGGCGAGTTTTTCCATCTTTTGGATGTTTTTTCAGCGTTTTCTGTTGGCTCGATTCATCAGAAAAA
ATGTTAGCGCGGTCAAGTTTTCACCGCAAAGGTATTTAAAAGGTATTATTAAGTGGTATTGTCATCGCGTACCTTACATTACCTGTCAT
GAAGGAATTAAAAGATGAAAACAACAGCAAAGCTGTCGTTCATGATGTTTGTTGAATGGTTTATCTGGGGCGCGTGGTTTGTGCCATTG
TGGGGATCCTCTAGATTTAAGAAGGAGATATACATATG
>ygaD
GAGGCAATTCCGACGTCTAAGAAACCATTATTATCATGACATTAACCTATAAAAATAGGCGTATCACGAGGCCCTTTCGTCTTCACCTC
GATGGATCCGCGGTCTGGCAGTTAGGACAAGCCGTGGCGCTGGCGCGAGTACAGTAGCTATTTACAAGAGGGGGGCGTGAGCTTCCCTC
TGGCTAATCAGCTTTTTTCTGAATCCTCCTCGTAAAATTGCAACGCCAACACCATCTTCCTGACGAAAGTGCTATCTTGTCCGGCATAA
ATTTTTGACTGACAGAGGTTGTGATGACTGACAGTGAACTGATGCAGTTAAGTGAACAGGTTGGGCAGGCGCTGAAAGCCCGTGGCGCA
ACCGTAACAACTCTCGAGAGATCCTCTAGATTTAAGAAGGAGATATACATATG
>ygaZ
GAGGCAATTCCGACGTCTAAGAAACCATTATTATCATGACATTAACCTATAAAAATAGGCGTATCACGAGGCCCTTTCGTCTTCACCTC
GAGTGGCGTTTGTCTGGCTGGCGCGACGATTGCCCTGGTTAACTTACTGGTCTGGTGGCGAGGTTTTCATCGTCAGGAAGCCGCAAATT
AAGCAATCAGCAACCGTTTTCGTGGCTTTACACTTATAAGGGTGTTAAGAAGCCCATCAGTCTGATAAGGTTAAGATATTCATTCAGTC
TATTTATAATATTAACAATCGTTAAGCGTACACTCTATGGAAAGCCCTACTCCACAGCCTGCTCCTGGTTCGGCGACCTTCATGGAAGG
ATGCAAAGACAGTTTACCGATTGTTATTAGTTATATTCCGGTGGCCTTGGATCCTCTAGATTTAAGAAGGAGATATACATATG
>zntA
GAGGCAATTCCGACGTCTAAGAAACCATTATTATCATGACATTAACCTATAAAAATAGGCGTATCACGAGGCCCTTTCGTCTTCACCTC
GAGGGATAACGCGATTGCTGCGGCCTGCTACTTTGCCGGTCACTTCCTGATCGTCCGCTCGCTGTATCTCTGATAAAACTTGACTCTGG
AGTCGACTCCAGAGTGTATCCTTCGGTTAATGAGAAAAAACTTAACCGGAGGATGCCATGTCGACTCCTGACAATCACGGCAAGAAAGC
CCCTCAATTTGCTGCGTTCAAACCGCTAACCACGGTGGATCCTCTAGATTTAAGAAGGAGATATACATATG
>soxR
GAGGCAATTCCGACGTCTAAGAAACCATTATTATCATGACATTAACCTATAAAAATAGGCGTATCACGAGGCCCTTTCGTCTTCACCTC
GAGTCAATGTTAAGCGGCTGGTCAATATGCTCGTCAATCCATGCGATAAGATCCTGAATAATTTTCTGATGGGACATAAATCTGCCTCT
TTTCAGTGTTCAGTTCGTTAATTCATCTGTTGGGGAGTATAATTCCTCAAGTTAACTTGAGGTAAAGCGATTTATGGAAAAGAAATTAC
CCCGCATTAAAGCGCTGCTAACCCCCGGCGAAGTGGCGAAACGCAGCGGTGTGGCGGTATCGGCGCTGCATTTCGGATCCTCTAGATTT
AAGAAGGAGATATACATATG
>soxS
GAGGCAATTCCGACGTCTAAGAAACCATTATTATCATGACATTAACCTATAAAAATAGGCGTATCACGAGGCCCTTTCGTCTTCACCTC
GATAGATCTGAAATGCAGCGCCGATACCGCCACACCGCTGCGTTTCGCCACTTCGCCGGGGGTTAGCAGCGCTTTAATGCGGGGTAATT
TCTTTTCCATAAATCGCTTTACCTCAAGTTAACTTGAGGAATTATACTCCCCAACAGATGAATTAACGAACTGAACACTGAAAAGAGGC
AGATTTATGTCCCATCAGAAAATTATTCAGGATCTTATCGCATGGATTGACGAGCATATTGACCAGCCGCTTAACATTGACTCGAGAGA
TCCTCTAGATTTAAGAAGGAGATATACATATG
>ygaQ
GAGGCAATTCCGACGTCTAAGAAACCATTATTATCATGACATTAACCTATAAAAATAGGCGTATCACGAGGCCCTTTCGTCTTCACCTC
GAGCAGATTTTTATGCGTTCCCATTCACCTTGGGTCGTGATTACACGCGACATAAAACCCGACACCGCCTCCATTCGCAAAGTCGATAC
TCGCAGTCAACAAGCAAATGTTAATAATTAGCACTATCTATAGTTATCATCGATTCAATGATACTTTGTAATGATTTTGTATCTAATAA
TATAACTTTATTACATTAGCTGAAGAGTTTTCGCATCATTATGATATCTGTTACTTTTCACTCCATAAAAATAAACTTCGTATAGCAAT
ATATTCTTTCATAGATCTTATTCTGCTAATCATTAGTTTCGTATGAGCGATTTTTGACAGTTGCCTCTCCAGACCACATCGATAATTAA
TAAAACAGATTTAAGCATTATCCTTTTCCATATAAATATTGGATAAAAGTAGGACATCTGTTTGCAATTACTTTCACAACAATTAAACA
TTTTTATGTTTCCGTATACATCATATTACTCTACCATTAGAGGAACTTTATTATGTTTTCTATAAAACCAGGACCCAGAAATTTACCTA
TCGACAACCCCACATTGTTATCATGGAACATTACTGACGGGGGGATCCTCTAGATTTAAGAAGGAGATATACATATG
>aceK
GAGGCAATTCCGACGTCTAAGAAACCATTATTATCATGACATTAACCTATAAAAATAGGCGTATCACGAGGCCCTTTCGTCTTCACCTC
GAGCGACTATTATTCAGGGCGGCACGTCTTCAGTCACCGCGCTGACCGGCTCCACTGAAGAATCGCAGTTCTAAGCAACAACAACCGTT
GCTGACTGTAGGCCGGATAAGGCGTTCACGCCGCATCCGGCAATCGGTGCACGATGCCTGATGCGACGCTTGCGCGTCTTATCATGCCT
ACAGCCGTTGCCGAACGTAGGCTGGATAAGGCGTTTACGCCGCATCCGGCAATTCTCTGCTCCTGATGAGGGCGCTAAATGCCGCGTGG
CCTGGAATTATTGATTGCTCAAACCATTTTGCAAGGCTTCGATGCTCAGTATGGTCGATTCCTCGAAGTGACCTCCGGGATCCTCTAGA
TTTAAGAAGGAGATATACATATG
>ssrA
GAGGCAATTCCGACGTCTAAGAAACCATTATTATCATGACATTAACCTATAAAAATAGGCGTATCACGAGGCCCTTTCGTCTTCACCTC
GAGGCGTCGCCAAAGGTAAGAAACAGCACGATAAACGTTCAGATATCAAAGAGCGCGAATGGCAGGTGGATAAAGCACGTATCATGAAA
AACGCCCACCGTTAAACCTGCACTCCAATTATTGACCAGTTCCTCACCGCGCCTCCCTCTCCGGCGGCGCGAATGAACATCTTATTGGC
TATCACATCCGACACAAATGTTGCCATCCCATTGCTTAATCGAATAAAAATCAGGCTACATGGGTGCTAAATCTTTAACGATAACGCCA
TTGAGGCTGGTCATGGCGCTCATAAGGATCCTCTAGATTTAAGAAGGAGATATACATATG
>aceB
GAGGCAATTCCGACGTCTAAGAAACCATTATTATCATGACATTAACCTATAAAAATAGGCGTATCACGAGGCCCTTTCGTCTTCACCTC
GAGGACGGAAGAAGGGGATGCATATCTGTTTGCCAGTAAAGATAAGCGCATTGCCTTTGTGACGGGCCATCCCGAATATGATGCGCAAA
CGCTGGCGCAGGAATTTTTCCGCGATGTGGAAGCCGGACTAGACCCGGATGTACCGTATAACTATTTCCCGCACAATGATCCGCAAAAT
ACACCGCGAGCGAGCTGGCGTAGTCACGGTAATTTACTGTTTACCAACTGGCTCAACTATTACGTCTACCAGATCACGCCATACGATCT
ACGGCACATGAATCCAACGCTGGATTAATCTTCTGTGATAGTCGATCGTTAAGCGATTCAGCACCTTACCTCAGGCACCTTCGGGTGCC
TTTTTTATTTCCGAAACGTACCTCAGCAGGATCCTCTAGATTTAAGAAGGAGATATACATATG
>fryB

GAGGCAATTCCGACGTCTAAGAAACCATTATTATCATGACATTAACCTATAAAAATAGGCGTATCACGAGGCCCTTTCGTCTTCACCTC
GATGGATCCGACAATCCGGGCCTTCTCGGTTCCGGTGCACATTTACGCCAGACCTTAGGTCACATTCTGTAAATCCTTCCTTTTATATC
GGGAGGTAACTCTCCCGATAATCTTTTAAATCATACAGTTTATTCAATTTTCTTTGTGTCCCCTCACAAGGTCGACCTGCGTCACACT
TCCGTACAGCGGGATTAATTCTCCAGTAAATGCATTATTTGTCTGGTAACGGCGATTTGTTTTGCACGTTCATAATTTCACTCGTCAAC
TATCACTGAACGAGGAATTAACGATGAGTAAGAAACTGATTGCCTTATGTGCCTGCCCGATGGGCCTGGCTCACACCTTTATGGCCGCT
CAGGCGCTGGAAGAAGCTCGAGAGATCCTCTAGATTTAAGAAGGAGATATACATATG
>ypjD
GAGGCAATTCCGACGTCTAAGAAACCATTATTATCATGACATTAACCTATAAAAATAGGCGTATCACGAGGCCCTTTCGTCTTCACCTC
GAGCTTCGCGCAGCGTATCTTTTACGTTGTCTTCAGTGAGGCGTCCACGGCCACTGATATTGCGCAGCGTGCGCGACAAACGATCGGTT
AAATTATCAAACATTGTCTCTCGCCTGGGGTGGAAACGGTTGGCCGCAATCGCGACACATCATCAGTATTTTGCCGCAGTATAACATGA
AGGCGTCTTTGTTGTTATGCAACGGTTGGAGCAGCGTTCACCTGACGCTATACTGCTTCTCTTTCTTATTGCTCAAACTGTCGACATCA
CTATGCCCGTTTTTGCTCTGGGATCCTCTAGATTTAAGAAGGAGATATACATATG
>ypjA
GAGGCAATTCCGACGTCTAAGAAACCATTATTATCATGACATTAACCTATAAAAATAGGCGTATCACGAGGCCCTTTCGTCTTCACCTC
GATGGATCCGGGGGACGAATGCTGTAAAAGTAAAACAATCTGCGATCTCAAAGTTATTGTGTGATTTTTATGTGAGCAGAAGATATTCA
TCAGAAACGATTATGTAAATCATTTTATTTTGCCAACGGCCTCATTGTCGAAAGAAAAGAATACGCCCGTATTTCCTGAATGGATTGAC
TTTTTTATCCAACCACACTTCAGCGCACTGCGTTTAAAAAATGCCTCATTCTTATGCGGAATATCATCATTTCATCATGATGTCTTTGA
TGAGCGGTGAACACAATACACTTGCGCTGTCTCTTCAGGATGAATCCCCTGGTCTGGTGCCTGTGGGCTGATGTTGCAGCTCGAGAGAT
CCTCTAGATTTAAGAAGGAGATATACATATG
>rihC
GAGGCAATTCCGACGTCTAAGAAACCATTATTATCATGACATTAACCTATAAAAATAGGCGTATCACGAGGCCCTTTCGTCTTCACCTC
GAGCTGGAAGGCCGTGAAGAAAACATTGTTTTCGAAGTGCCGAAAGAGCTGCGTGTCGATATTCGTGAAGTCGATTAAGTCATTAGCAG
CCTAAGTTATGCGAAAATGCCGGTCTTGTTACCGGCATTTTTTATGGAGAAAACATGCGTTTACCTATCTTCCTCGATACTGACCCCGG
CATTGACGATGCCGTCGCCATTGCCGCCGCGATTTTTGCACCCGAACGGATCCCGGGATCCTCTAGATTTAAGAAGGAGATATACATAT
G
>rihA
GAGGCAATTCCGACGTCTAAGAAACCATTATTATCATGACATTAACCTATAAAAATAGGCGTATCACGAGGCCCTTTCGTCTTCACCTC
GATGGATCCATTGTCGAAGACTCGCCGAAAGACGCTTTCTTCGATGATCCGAAATCGGACCGCGCAAAAGACTTCCTCGCGAAAATCCT
GCATTAATCACTCTGGCGCGCATACCCAATTGCGCGCCATCTCACGCTTTATCGTTTCGCCCCTCGTATCTGTTACATCCTGGCGTTAC
CCTTGTCGCAAAGAAGCACACAACAAGGAGCAACAATGGCACTGCCAATTCTGTTAGATTGCGACCCAGGTCATGACGACGCTATCGCA
ATAGTTCTCGCCCTCGCCCTCGAGAGATCCTCTAGATTTAAGAAGGAGATATACATATG
>hsdM
GAGGCAATTCCGACGTCTAAGAAACCATTATTATCATGACATTAACCTATAAAAATAGGCGTATCACGAGGCCCTTTCGTCTTCACCTC
GATGGATCCTATCTGGGACGAGCTGGCCTGACACGTATACACTTCATCCTTCAGGCTGCCTCTGCGTTGGCTGCGCTCGTTCACCCCGG
TCACGTACTTCTGTACGCTCCCGGGGATTCACTCACTTGCCGCCTTGATGCAACCTGAATGATTTTGTGTATATTACCCTCGGCAATTT
CTTCTTCTGCGGCTCGATGAATTTGGGCCGCTGCTTAATTTACGGAACTCACAATGAACAATAACGATCTGGTCGCGAAGCTGTGGAAG
CTGTGCGACAACCTGCGCGATGGCGGCGTTTCCTATCAAAACTACGTCAATGAACTCGCCTCGCTCTCGAGAGATCCTCTAGATTTAAG
AAGGAGATATACATATG
>lysA
GAGGCAATTCCGACGTCTAAGAAACCATTATTATCATGACATTAACCTATAAAAATAGGCGTATCACGAGGCCCTTTCGTCTTCACCTC
GATGGATCCCCGCCTCAGTCAGGCTTCCGGCGGTCATTACCGCATGAAAAATTTCAATATGACGTAAGTTAACGGCGGCCATTAGCGCT
CTCTCGCAATCCGGTAATCCATATCATTTTTGCATAGACTCGACATAAATCGATATTTTTTATTCTTTTTATGATGTGGCGTAATCATA
AAAAAGCACTTATCTGGAGTTTGTTATGCCACATTCACTGTTCAGCACCGATACCGATCTCACCGCCGAAAATCTGCTGCGTTTGCCCG
CTGAATTTGCTCGAGAGATCCTCTAGATTTAAGAAGGAGATATACATATG
>hsdR
GAGGCAATTCCGACGTCTAAGAAACCATTATTATCATGACATTAACCTATAAAAATAGGCGTATCACGAGGCCCTTTCGTCTTCACCTC
GATGGATCCTGCACCTTCCGGTTTTGTTGCCAGATAACGCAGAACAGGTTCAATAAATTTGTCATAGGTAGGAACCGTCATAGTACATC
CTTGCAGAATCAGGTAGATGTTTTTCGGCTACTATAGCACTACAAAAATAGACGAACACGTTAGAAATGAGTCAGTTGCTGTGACCGTG
GTCATTGCCCGGAAAGGTACAGAAAGCTAAGATGAGATGTTATGGGCCTTAAATATTTGGACAGGCCCGCACAGCAATGGATTAATAAC
AATGATGAATAAATCCAATTTTGAATTCCTGAAGGGCGCTCGAGAGATCCTCTAGATTTAAGAAGGAGATATACATATG
>lysR
GAGGCAATTCCGACGTCTAAGAAACCATTATTATCATGACATTAACCTATAAAAATAGGCGTATCACGAGGCCCTTTCGTCTTCACCTC
GAGCAAATTCAGCGGGCAAACGCAGCAGATTTTCGGCGGTGAGATCGGTATCGGTGCTGAACAGTGAATGTGGCATAACAAACTCCAGA
TAAGTGCTTTTTTATGATTACGCCACATCATAAAAAGAATAAAAAATATCGATTTATGTCGAGTCTATGCAAAAATGATATGGATTACC
GGATTGCGAGAGAGCGCTAATGGCCGCCGTTAACTTACGTCATATTGAAATTTTTCATGCGGTAATGACCGCCGGAAGCCTGACTGAGG
CGGGGATCCTCTAGATTTAAGAAGGAGATATACATATG
>lysU
GAGGCAATTCCGACGTCTAAGAAACCATTATTATCATGACATTAACCTATAAAAATAGGCGTATCACGAGGCCCTTTCGTCTTCACCTC
GATAGATCTGGCCTTTGCTACCCGTTTTCTGTTTAGCACGCCGACGAATATGATACAGGAGAGCAACGATTAACTGTCGTAAAAACAAT
TGGTTATGAATAAATTAACTTTCAGTTTATAACACTAACAAATCGAGATTAATAAGTCATTTAAACCATTTTGATGGTTATTTATTAGT
GATATCAACTTGAGGTAAGCGTTAGTTTCGATAAGATAAACTGAGTTACTAATAGTCGAGGCAGATAATACAGTGTACCGATCTGTCTC
TTTATCTACGCTAAATTGAAAGCTGGATTTAGAGGAACCAAAATGTCTGAACAAGAAACACGGGGAGCCAATGAGGCTATTGATTTTAA
CGATGAACTGAGAAATCGCCGCGAAAAACTGGCGGCACTACGCTCGAGAGATCCTCTAGATTTAAGAAGGAGATATACATATG

```
>eaeH
GAGGCAATTCCGACGTCTAAGAAACCATTATTATCATGACATTAACCTATAAAAATAGGCGTATCACGAGGCCCTTTCGTCTTCACCTC
GAGCTGTTTTGATAGTCGAGCCGATTTTAAAGTACTCATCAACACTGGTGTCGTGGAACACCACAGTACGATACTCAGGATGGATATTG
GGCTTCATCATTTTTACCTGTTATGTTATAACATAACCATAAAGATACATGCTCATCCGGATCATTTCAACACTCAATCCCTGTGACTC
TTTGTGGATGTGGAATTTTCGCAACGGGGAACTCTGGCATTGTTAGCAGCCATTGGTGACGCTCCCATAAACGCCTGGATGAGTGATTG
ATGATGGTTTTGCTGAGAGTATGGATTTTTCGAGAATGGAGGTTATCTCCACTTTACTGGCCGATAATTTATCGCAGTATGATTGATTA
CCTGACAGCGTGTATTCCTCTCCCACGGATATAATAATTACTTACAAAAAAGGGAGAGGATGCATATTTTAAATATCACTGAAGTGAAC
AGTTTATTTCCGTTATTAATAGAAATGGAGAAATAAATAGGCGTATTCTACAATTGCGACAAAAACAACGATATTAATCAGTTTATGAC
TGATTTGCTGTACTTTATTCTCTTTCATTGGTACTTCCTCGCTTTAAAAAAGAGTGCACTTCGTAAGTGCCCTTATATAAATAACGAGT
TTGGTCAACCAATTTTTTGACATGTATCACAAATTTGAATAGATGTATTACATCAACTATCTTTTATTGCACCAACGTCATTGATATAT
GTCGCCTGAAGTCAGTTCCGGGAATGAGTCTGATCTCAAGACTGGCCCAGTCCGGGCGTTGATTGGTGCTGAGGAGCATATCGCATCTC
ATCATAATGTCGTATCTCCTGGGGTGTTATACAAGATATCGTTGTTGGTGACCTGGGAGAGGAATTGAGTTCTATTAAACCGTCAACTA
TGCCGGATACATACTGGATTACACTGCAGGCACGCCTTATGAGAGAACGTGCCGCAGTGACGGGTTAATTATCTGAAAGAATTTGTGAG
GCTGTATCGGTTACTCATTGATTTGATAGTTTTACTCTCGGGAGAATAATAGATATTTAATCCATTAACGGAAACCAGCCAGTTCCTTT
CGATGCCTGAATTTGATCCCATAGTTTATCCGGAATGGTTAATTCATGAACAAGTTGTGTTATCGTTCATGAGAAGCATAACGTAAAGG
GAAAAGCTCGATTAGACGGCAGAATTTGTCAGGGGTTATGAACGAAATTCATAAATCTGTTTGAGTGTTGCGATGGGTAGTGCAAGTTC
GATATCTCCGCAATTTACAGTCCGATGAAGGAAAATGAATATCCATAAAAAATATATTGGTTTATCCTGGCATATATACCTATTTCGAC
GTATTTCCAATAGTTTTAATTAAAGGCAGGTCATTGTTATTCACTCTGAATAGTGAATTATTCACTGTCCGCAGAGTAAGAAATATAAC
TTAGGTATCTATTTAATGACTTGCACAAAAAGCTAAATTTTCCCCCATAAATAAAAATATAATCCCGCGCCCAACCACCTGATGAGTGG
CTATAGGCACTGGATATATTAGGTGGCGGTGCACTTTCTTACATAAAGGTATTTCCTTTTCTGCGGAAAAGGAAATCGGGAAATCCCCG
GTTTTTCTGACAAGCAGACGCCATTATTTGTGTCTGCCTATGTTCGTTAATTCGTTCATCAGGAAATTATCTCAATGTCACATTATAAA
ACAGGTCATAAACAACCACGATTTCGTTATTCAGTTCTGGCCCGCTGCGTGGCGTGGGCAAATATCTGGATCCTCTAGATTTAAGAAGG
AGATATACATATG
>ansA
GAGGCAATTCCGACGTCTAAGAAACCATTATTATCATGACATTAACCTATAAAAATAGGCGTATCACGAGGCCCTTTCGTCTTCACCTC
GAGAAAGTGACAAGCTGGCCGCGTTTAATGACCCACAAAACCGTTATGCGTTTTGCCTGACCTGCGCCAACATGCGTTAAGTCTTGTAC
TGAGTGGCCGACAGATCGTCGGCCACATTATTTTTTACGTCGACGAATCCTCTTCCCGCTGTTTCGCCCCATATTTCCTTATCCGCATA
GTATCAGGTGCGCTCCCCCTGCCTCACGTATATACTTTTGCTCTTTCGATATCATTCATATCAATATCATGCAAAAGAAATCAATTTAC
GTTGCCTACACGGGCGGGACCATCGGGATGCAGCGTTCCGAGCAGGGTTATGGATCCTCTAGATTTAAGAAGGAGATATACATATG
>ansB
GAGGCAATTCCGACGTCTAAGAAACCATTATTATCATGACATTAACCTATAAAAATAGGCGTATCACGAGGCCCTTTCGTCTTCACCTC
GATGGATCCAGATAGCCGCAAAGCCCTGGTCGGGAATTTAAAATAATCCTCTATTTTAAGACGGCATAATACTTTTTTATGCCGTTTAA
TTCTTCGTTTTGTTACCTGCCTCTAACTTTGTAGATCTCCAAAATATATTCACGTTGTAAATTGTTTAACGTCAAATTTCCCATACAGA
GCTAAGGGATAATGCGTAGCGTTCACGTAACTGGAGGAATGAAATGGAGTTTTTCAAAAAGACGGCACTTGCCGCACTGGTTATGGGTT
TTAGTGGTGCAGCATTGGCATTACCCCTCGAGAGATCCTCTAGATTTAAGAAGGAGATATACATATG
>mgsA
GAGGCAATTCCGACGTCTAAGAAACCATTATTATCATGACATTAACCTATAAAAATAGGCGTATCACGAGGCCCTTTCGTCTTCACCTC
GATGGATCCTGCTCACAGAACAACGGCTGAAGTATTGGTTTAAACTGGCTGACCCACAAACGCGAAATACTTTCCTGCAGTGGGCGGAA
AAACAACCATCTTCCTGAGATTTTTGTGCCTGTGCGCAGGCTTTTTCGGTCTTTATCTTGCAGCGATAAGTGCTTACAGTAATCTGTAG
GAAAGTTAACTACGGATGTACATTATGGAACTGACGACTCGCACTTTACCTGCGCGGAAACATATTGCGCTGGTGGCACACGATCACCT
CGAGAGATCCTCTAGATTTAAGAAGGAGATATACATATG
>yqfA
GAGGCAATTCCGACGTCTAAGAAACCATTATTATCATGACATTAACCTATAAAAATAGGCGTATCACGAGGCCCTTTCGTCTTCACCTC
GATGGATCCTGCTGACATCTATCCCGGTCAGACACAATTTTATGTGATTGAATTTAAATGTCTTTAAATTGAACTCACGCCTGTTAGTT
GAAATTTCGCATTTAATTTCATGATTTTTAGGAGATTAGTCTCAAATCAGAATTTATTTTAGCTAACAGGTGTTCACTGGAACTATTCT
CAGTTACGCTGGAGAGGTATACATGTGCGTGTACGTTTCTCCGGAGTAAGTTATGGTTCAGAAGCCCCTCATTAAGCAGGGATATTCAC
TGGCAGAGGAAATAGCCAACAGCGTCAGTCACGCTCGAGAGATCCTCTAGATTTAAGAAGGAGATATACATATG
>norR
GAGGCAATTCCGACGTCTAAGAAACCATTATTATCATGACATTAACCTATAAAAATAGGCGTATCACGAGGCCCTTTCGTCTTCACCTC
GATGGATCCTCACGCACTTCCCAGTCACGTTGACCAACCCAATGAATGTTATTTTTCACCACAATAGACATAGCAACCTCAATTTATTC
AGCGTGTTCTAAAAAGATGTCTTGCTAATTGCAGATTGCGTGCCAGTTTTTTATCTGATTGTTTTTACTCTATAAATAAAATTATGCCC
ATTAATGAGTAGTCAAAATGACTATCAATCAAATGGTCGATATGACAATATCTATAGTCAAATTGACAGTGAGGCAAAGATGAGTTTTT
CCGTTGATGTGCTGGCTCGAGAGATCCTCTAGATTTAAGAAGGAGATATACATATG
>norV
GAGGCAATTCCGACGTCTAAGAAACCATTATTATCATGACATTAACCTATAAAAATAGGCGTATCACGAGGCCCTTTCGTCTTCACCTC
GAGCCAGCACATCAACGGAAAAACTCATCTTTGCCTCACTGTCAATTTGACTATAGATATTGTCATATCGACCATTTGATTGATAGTCA
TTTTGACTACTCATTAATGGGCATAATTTTATTTATAGAGTAAAAACAATCAGATAAAAAACTGGCACGCAATCTGCAATTAGCAAGAC
ATCTTTTTAGAACACGCTGAATAAATTGAGGTTGCTATGTCTATTGTGGTGAAAAATAACATTCATTGGGTTGGTCAACGTGACTGGGA
AGTGCGTGATTTTCACGGCACGGAAGGATCCTCTAGATTTAAGAAGGAGATATACATATG
>pth
GAGGCAATTCCGACGTCTAAGAAACCATTATTATCATGACATTAACCTATAAAAATAGGCGTATCACGAGGCCCTTTCGTCTTCACCTC
GATGGATCCCGAAGCGTTTTTGCGTTTCATATGTCACCTCCGGAACTTTCTGGGTTGTAACAGGGAATACCCCTCTTCCTTATGTGTAA
AGTATAGACAACACCCAGTGGGTTATGTGCGGGCGTGATCACAATTACAACCCTTATTTCAACAAAACTTTACAAATAAACGCCTGAAC
```

Data Set S4 - MG1655 library reference file.fasta

ACCTTGACTTTCTAATAGTCCACTCAGGCAATTATTACGTTATCTGCATTTGATACGCAGTTTTTTGTCAAGCGGGGCCGCAACCAGTA
AACTACGCGCCAGTTATGTACACACTCAGGACAAAAAAACGTGACGATTAAATTGATTGTCGGCCTGGCGAACCCCGGTGCTGAATACG
CCGCAACGCGACATAATGCTGGTGCCTGGTTCGTTGACCTCGAGAGATCCTCTAGATTTAAGAAGGAGATATACATATG
>lacA
GAGGCAATTCCGACGTCTAAGAAACCATTATTATCATGACATTAACCTATAAAAATAGGCGTATCACGAGGCCCTTTCGTCTTCACCTC
GATGGATCCCGCTGGGCTTCACCTTAATTTCCGTGTTCACGCTTAGCGGCCCCGGCCCGCTTTCCCTGCTGCGTCGTCAGGTGAATGAA
GTCGCTTAAGCAATCAATGTCGGATGCGGCGCGAGCGCCTTATCCGACCAACATATCATAACGGAGTGATCGCATTGAACATGCCAATG
ACCGAAAGAATAAGAGCAGGCAAGCTATTTACCGATATGTGCGAAGGCTTACCGGTCGACAGATCCTCTAGATTTAAGAAGGAGATATA
CATATG
>lacI
GAGGCAATTCCGACGTCTAAGAAACCATTATTATCATGACATTAACCTATAAAAATAGGCGTATCACGAGGCCCTTTCGTCTTCACCTC
GATGGATCCGGGTTGAATCGCAGGCTATTCTGGTGGCCGGAAGGCGAAGCGGCATGCATTTACGTTGACACCATCGAATGGCGCAAAAC
CTTTCGCGGTATGGCATGATAGCGCCCGGAAGAGAGTCAATTCAGGGTGGTGAATGTGAAACCAGTAACGTTATACGATGTCGCAGAGT
ATGCCGGTGTCTCTTATCAGACCGTTTCCCGCGTGGTGAACCAGGCCAGCCACGTTTCTGCGAAAACTCGAGAGATCCTCTAGATTTAA
GAAGGAGATATACATATG
>lacY
GAGGCAATTCCGACGTCTAAGAAACCATTATTATCATGACATTAACCTATAAAAATAGGCGTATCACGAGGCCCTTTCGTCTTCACCTC
GATGGATCCTCCTGGAGCCCGTCAGTATCGGCGGAATTCCAGCTGAGCGCCGGTCGCTACCATTACCAGTTGGTCTGGTGTCAAAAATA
ATAATAACCGGGCAGGCCATGTCTGCCCGTATTTCGCGTAAGGAAATCCATTATGTACTATTTAAAAAACACAAACTTTTGGATGTTCG
GTTTATTCTTTTTCTTTTACTTTTTTATCATGGGAGCCTACTTCCCGCTCGAGAGATCCTCTAGATTTAAGAAGGAGATATACATATG
>lacZ
GAGGCAATTCCGACGTCTAAGAAACCATTATTATCATGACATTAACCTATAAAAATAGGCGTATCACGAGGCCCTTTCGTCTTCACCTC
GATGGATCCCACGACAGGTTTCCCGACTGGAAAGCGGGCAGTGAGCGCAACGCAATTAATGTGAGTTAGCTCACTCATTAGGCACCCCA
GGCTTTACACTTTATGCTTCCGGCTCGTATGTTGTGTGGAATTGTGAGCGGATAACAATTTCACACAGGAAACAGCTATGACCATGATT
ACGGATTCACTGGCCGTCGTTTTACAACGTCGTGACTGGGAAAACCCTGGCGTTACCCAACTTAATCGCCTTGCAGCACAAGATCCTCT
AGATTTAAGAAGGAGATATACATATG
>atpI
GAGGCAATTCCGACGTCTAAGAAACCATTATTATCATGACATTAACCTATAAAAATAGGCGTATCACGAGGCCCTTTCGTCTTCACCTC
GATGGATCCATGGCGAACGTCATCTGGTGGTGATTAAAGCAAATAAAATTTAATTTTTATCAAAAAAATCATAAAAAAATTGACCGGTTA
GACTGTTAACAACAACCAGGTTTTCTACTGATATAACTGGTTACATTTAACGCCACGTTCACTCTTTTGCATCAACAAGATAACGTGGC
TTTTTTTGGTAAGCAGAAAATAAGTCATTAGTGAAAATATCAGTCTGCTAAAAATCGGCGCTAAGAACCATCATTGGCTGTTAAAACAT
TATTAAAAATGTCAATGGGTGGTTTTTGTTGTGTAAATGTCATTTATTAAAACAGTATCTGTTTTTAGACTGAAATATCATAAACTTGC
AAAGGCATCATTTGCCAAGTAAATAAATATGCTGTGCGCGAACATGCGCAATATGTGATCTGAAGCACGCTTTATCACCAGTGTTTACG
CGTTATTTACAGTTTTTCATGATCGAACAGGGTTAGCAGAAAAGTCGCAATTGTATGCACTGGAAAAATATTTAAACATTTATTCACCT
TTTGGCTACTTATTGTTTGAAATCACGGGGGCGCACCGTATAATTTGACCGCTTTTTGATGCTTGACTCTAAGCCTTAAAGAAAGTTTT
ATACGACACGCGGCATACCTCGAAGGGAGCAGGAGTGAAAAACGTGATGTCTGTGTCGCTCGTGAGTCGAAACGTTGCTCGGAAGCTTC
TGCTCGTTCAGTTACTGGTGGTGATAGCAAGTGGACTCGAGCGGGTGGTGATAGCTAGTGGACTCGAGAGATCCTCTAGATTTAAGAAG
GAGATATACATATG
>sanA
GAGGCAATTCCGACGTCTAAGAAACCATTATTATCATGACATTAACCTATAAAAATAGGCGTATCACGAGGCCCTTTCGTCTTCACCTC
GAGATGCGCCGTTGATTCAGTGGGATGCCACCTCCGCAACGCTGAAAGCTCTCGGCTGTCACAGTATCGACCGAGTGCTTCTCGCTTAA
GCCTGGTGCCGGATGCGGCGTGAACGCCTTGTCCGGCTTGCCAGCCCTCTCCTGGTGTCGAAATTCCCGGCAAACAGTTTGCCGTTTCT
TGCGCAAAACCAGCGGGTAAAGTAGCCTGATGGAAATTTTCCTTAGATCGAGTCTCCTGCATGTTAAAGCGCGTGTTCCTCAGCCTGTT
AGTCCTGATCGGCTTGCTGCTGTTGACTGTGCTCGGCCTGGATCCTCTAGATTTAAGAAGGAGATATACATATG
>pfkB
GAGGCAATTCCGACGTCTAAGAAACCATTATTATCATGACATTAACCTATAAAAATAGGCGTATCACGAGGCCCTTTCGTCTTCACCTC
GAGAAAACGGAATCATCGGCAGCTGCATTCAGTGACGCAAACATGCCCCCCGCCAACATAACGATGGCAGGAACTGTCTTCAAAAGCTT
CATTTATCAAGAGTCCGTACAACAAAAAAAGAGACCATCGCGGTCCCGGAAACTTTCTTAAGGATCAAAGATTAGCGTCCCTGGAAAGG
TAACGAATTATAAAAAGGCGCGAATAACTTAGCAATGTATTCTTATTTCATTTTTTGAATAAGCATGTGGCGAAAACAGATTTTTATTT
ATATATATTTATCTGCAAAATTTTAAATAAAGCTCCAATAAATCATATTGTTAATTTCTTCACTTTCCGCTGATTCGGTGCCAGACTGA
AATCAGCCTATAGGAGGAAATGATGGTACGTATCTATACGTTGACACTTGCGCCCTCTCTCGATAGCGCAACAATTACCCCGCAAATTT
ATCCCGAAGGAAAACTGCGCTGTACCGCGGATCCTCTAGATTTAAGAAGGAGATATACATATG
>pfkA
GAGGCAATTCCGACGTCTAAGAAACCATTATTATCATGACATTAACCTATAAAAATAGGCGTATCACGAGGCCCTTTCGTCTTCACCTC
GAGTTCCGGGATCGGATGTAATTATCCATCAGGACCCCTGTTCCGTCGTACCCAGGGAGGGTAAACGGTCTATGCTTTCATAATCAGTA
TAAAAGAGAGCCAGACCCGCATTTTGTGTATAAAATACCGCCATTTGGCCTGACCTGAATCAATTCAGCAGGAAGTGATTGTTATACTA
TTTGCACATTCGTTGGATCACTTCGATGTGCAAGAAGACTTCCGGCAACAGATTTCATTTTGCATTCCAAAGTTCAGAGGTAGTCATGA
TTAAGAAAATCGGTGTGTTGACAAGCGGCGGTGATGCGCCAGGCATGAACGCCGCAATTCGCGGGGTTGTTCGTTCTGCGCTGGATCCT
CTAGATTTAAGAAGGAGATATACATATG
>hemH
GAGGCAATTCCGACGTCTAAGAAACCATTATTATCATGACATTAACCTATAAAAATAGGCGTATCACGAGGCCCTTTCGTCTTCACCTC
GAGCAAAGAAGCAGAAGCGGGTAATACCAAATACGCGAAAGTTGACGGCACCAAGCCGGTTGCTGAAGTTCGCGCTGATCTGGAAAAAA
TCCTCGGCTAATTCGAAAGCGCGCACGGACAGTCCCCTCGCCCCCTCGGGGAGAGGGTTAGGGTGAGGGGAACAGGCCCGCACAAGCAA

```
ACTTATCAGCAATCTCAGGCCGGATATTCATTCGGCCTTTTACAAAAACCTCAATCTGCGCTATTCCTTTTTCTGATTTGACCTCTCAC
AGCAATTAGTTCTTCTTCCTCACTTTTCCGCTACAATTATCAACAAGTTGAATCGATAAGAGGCGGTAATGCGTCAGACTAAAACCGGT
ATCCTGCTGGCAAACCTGGGTACGCCCGATGCCCCCACACCTGAAGCGGTAGGATCCTCTAGATTTAAGAAGGAGATATACATATG
>hemN
GAGGCAATTCCGACGTCTAAGAAACCATTATTATCATGACATTAACCTATAAAAATAGGCGTATCACGAGGCCCTTTCGTCTTCACCTC
GAGTGCAGAAACTCGGCCTCTCTTATGACGATGACGAAGAAGAGGAAGAAGACGAGAAGCAAGAAGACATGATGCGTCTGCTGCGGGGC
AACTAACGGATTGCCGCCGTGGGCCTTCCCGTTCTGCTGATAACCCTTCCGGTTATATGTTATCTGGTGTGGTTATTCGTTAAACTACA
GCGGTTGTCGCGGCGACAAAAGTGGCTGCGCAACCGGCTGATGACCCGAAACGGGCATCCGCCGGTACGCCGTAGCCGCCAGAGACGCC
ATCGGAAGGAGTGAGCATGTCTGTACAGCAAATCGACTGGGATCTGGCCCTGATCCAGAAATATAACTATTCCGGGCCACGATGGATCC
TCTAGATTTAAGAAGGAGATATACATATG
>yieH
GAGGCAATTCCGACGTCTAAGAAACCATTATTATCATGACATTAACCTATAAAAATAGGCGTATCACGAGGCCCTTTCGTCTTCACCTC
GAGGATCACTTCGGTCCGTGCCGTCGTGCCATGTTCGCGCAGTTTAAACACGCGTTCCAGCATCCCCTGACCAGAAGTCTGGGTGGTAT
GTTGATGACTCATTATCTATTTCCGAACAAGGAGGGAAAATCCGTCGCTATCGTATACCAAAATGCGACAATAGGCGCGTTTGTGAGAG
ACTTTTTTATTGGATTTACTTATACGGCAACGATTGCGTTGCGCAAATCGGCTCTACGAAAACGTTAAACTGATTAAAAAGGAAAGGCA
TGTCCCGGATAGAAGCGGTATTTTTCGACTGCGACGGTACGCTGGTCGACAGTGAAGTCATTTGCTCTCGCGCGGATCCTCTAGATTTA
AGAAGGAGATATACATATG
>hemL
GAGGCAATTCCGACGTCTAAGAAACCATTATTATCATGACATTAACCTATAAAAATAGGCGTATCACGAGGCCCTTTCGTCTTCACCTC
GATGGATCCGGCCTGCGGTGTTTCTAAAGAGGGAGTATCAGTTTTCATCCAATGATCACTTATTGGTCATACAAATAAGATGACGGCAT
TTTACTCGCACGTTCCATGAAAATCCTTTGCAAAGCGTAATGTTTCAAATGCACAAAACTTTTATACTCTAAGAAATCAGGATTGCACC
AGTACAAGCAGCCTGATGTTTGACGAGTATTTAACTTGTTATGAATAACATAGAATAGCAGCCATTCACTTTTTCTACGAATCAGGAAC
CCTCCATGAGTAAGTCTGAAAATCTTTACAGCGCAGCGCGCGAGCTGATCCCTGGCGGTGTGAACTCCCCTGTTCGCCTCGAGAGATCC
TCTAGATTTAAGAAGGAGATATACATATG
>rhaD
GAGGCAATTCCGACGTCTAAGAAACCATTATTATCATGACATTAACCTATAAAAATAGGCGTATCACGAGGCCCTTTCGTCTTCACCTC
GATGGATCCCAACGTCACGATACGCCAGCAGGTAGCGAATGGCTGGAGAGCGTGCGGGCTTATGAGAAAGAAATTTTGAGTCGCCGCGG
GTAAACACTGCCGGATGCGGCGCGAGCGCCTTATCCGGCCTACGGGTCGGCAACAGTTGTAGGCCTGATAAGACGCGACAGCGTCGCAT
CAGGCATTGATTGCCGGATGCGGCACAAGTGCCTTATCCGGCCTACAGGTCGGCAATAGTTGTAGGCCTGATAAGACGCGACAGCGTCG
CATCAGGCATTGATTGCCGGATGCGGCACAAGTGCCTTATCCGGCCTACAGGTCGGCAACAGTTGTAGGCCTGATAAGACGCGACAGCG
TCGCATCAGGCATTGATTGCCGGATGCGGCACAAGTGCCTTATCAGGCCTACAGGTCGGCAATAGTTGTAGGCCTGATAAGACGCGACA
GCGTCGCATCAGGCATTGATTGCCGGATGCGGCGCGAGCGCCTTATCCGGCCTACGGGTCGTGCATCCGACAACACCGAATTTACAGGA
ACACAGAACATGCAAAACATTACTCAGTCCTGGTTTGTCCAGGGAATGATCAAAGCCACCACCGACGCCTGGCTGAAAGGCTGGGATGA
GCCTCGAGAGATCCTCTAGATTTAAGAAGGAGATATACATATG
>hemB
GAGGCAATTCCGACGTCTAAGAAACCATTATTATCATGACATTAACCTATAAAAATAGGCGTATCACGAGGCCCTTTCGTCTTCACCTC
GATGGATCCAGCAGCGCAACCAAACTCCAAAAAATAATAACCAACAGCTATAAATAAAGATTAAAAATATTTTCAGTGAATATAAAATT
ATTGCTGTTTCTATTAAAAAGATGTCCTGGCCTCTCTTCCATTCTCTTCTTGTCAACATCGCGACAACTTTCGTAAAACATCCCTACCC
TGCTTCAGGTATACTATGCCCCTCGATTCCACAAACATCAGGCAGACCATGACAGACTTAATCCAACGCCCTCGTCGCCTGCGCAAATC
TCCTGCGCTGCGCGCTATGTTTGAAGAGACAACACTTAGCCTTAACGACCTGGTGTTGCCGATCCTCGAGAGATCCTCTAGATTTAAGA
AGGAGATATACATATG
>yieG
GAGGCAATTCCGACGTCTAAGAAACCATTATTATCATGACATTAACCTATAAAAATAGGCGTATCACGAGGCCCTTTCGTCTTCACCTC
GATGGATCCACCGCTTCTATCCGGGACATGCCTTTCCTTTTTAATCAGTTTAACGTTTTCGTAGAGCCGATTTGCGCAACGCAATCGTT
GCCGTATAAGTAAATCCAATAAAAAAGTCTCTCACAAACGCGCCTATTGTCGCATTTTGGTATACGATAGCGACGGATTTTCCCTCCTT
GTTCGGAAATAGATAATGAGTCATCAACATACCACCCAGACTTCTGGTCAGGGGATGCTGGAACGCGTGTTTAAACTGCGCGAACATGG
CACGACGGCACGGACCGAAGTGATCCTCGAGAGATCCTCTAGATTTAAGAAGGAGATATACATATG
>ybdR
GAGGCAATTCCGACGTCTAAGAAACCATTATTATCATGACATTAACCTATAAAAATAGGCGTATCACGAGGCCCTTTCGTCTTCACCTC
GAGGAATTCAGCGTGGCGAACAGCTTTGTCGCTCAATTCCATTTCAAATACATCAACTGGCATAATGATTGTCTTATACATAACCCTTT
CTCCCTGTTAATCATGAACAAATCATTCGCCATGATTATAATATTTATCCCTGATATTTGTCTGGTTCTTTTTCCTTACGAACTGTTTC
TGTGATGAATATATTCTCACTGAACACCAGGAATTCTCCCAAAACCTGTGGTACCGCCCGTTTTCCCGCTGTGATAGCTACCCTTAAAG
ACTGACTCTTTTTTGAACTGTCTCTGGAGGTTGCATGAAAGCATTGACTTATCACGGCCCACATCACGTTCAGGTAGAAAATGTTCCCG
ATCCGGGCGTTGAACAGGCAGATGGATCCTCTAGATTTAAGAAGGAGATATACATATG
>hemG
GAGGCAATTCCGACGTCTAAGAAACCATTATTATCATGACATTAACCTATAAAAATAGGCGTATCACGAGGCCCTTTCGTCTTCACCTC
GAGCGCCTATGCATTGGTGTTTATTGTCAGTATGCTGGCGATTATCGCCACGGGCGTGGATGACTTTTCTGCCTTTGCGTCGGTTGTTG
CGACATTGAATAACCTGGGGCCAGGGCTTGGCGTGGTTGCTGATAACTTTACCAGTATGAACCCGGTGGCTAAATGGATCCTGATTGCC
AACATGCTGTTTGGTCGTCTCGAGGTCTTTACATTGCTGGTGCTCTTTACCCCGACTTTCTGGCGTGAATGATGGAGTAATACGTGAAA
ACATTAATTCTTTTCTCAACAAGGGACGGACAAACGCGCGAGATTGCCTCCTACCTGGCTTCGGAACTGAAAGAACTGGGGATCCAGGC
AGATCCTCTAGATTTAAGAAGGAGATATACATATG
>ybdH
```

GAGGCAATTCCGACGTCTAAGAAACCATTATTATCATGACATTAACCTATAAAAATAGGCGTATCACGAGGCCCTTTCGTCTTCACCTC
GATGGATCCTGGTGCCAAGTTGTGGAAGTTTGCTTTGTGGAATCAGAGGGTTATTTGTCATTTTTATTGTACCTGACGAAGTTGCGTGT
TGAAGGCACTATAACACGATGCTAGTATTTGGCAATCAAGACGTTTAGATGTCTAAATAAAACAATAAGGACAACACAACATGCCTCAC
AATCCTATCCGCGTGGTCGTCGGCCCGGCTAACTACTTTTCACATCCAGGAAGTTTCAATCACCTGCACGACTCGAGAGATCCTCTAGA
TTTAAGAAGGAGATATACATATG
>ybdK
GAGGCAATTCCGACGTCTAAGAAACCATTATTATCATGACATTAACCTATAAAAATAGGCGTATCACGAGGCCCTTTCGTCTTCACCTC
GATGGATCCGGCATTACGTGCCGAAGTATCCAGTTCTTCGGGCTTACCAATATGGGCCAGAATTGCTGACCAGTTATCGTGAGAGTAAT
TCATCGGCACGTTAAATCATATCAGGCGTAATACCACAACCCTTAAGTTAGCGCTTATGGGATTACTCCCCGCACAACGGCTACTTCTT
CGGTTCGTAAGCGAGAACAGCCTTAAACTCAATATTACGTTCCTTCACCGTAAACTCACACAGCGAGTCTCCGACCAGAAGCGTAAATC
CCAGCACCGTTAAACACAGCACTATGACTGCCGCAAGGGCATATTTCGTCAGCATGTTTATATTGCCTCCATGCGAAGAAGGGACTACC
ATCCAAATTGTTCGGGTTTGGAGTGGCAGCCCCAGGGTTGATAGAAATATCTCCTGGGGCTTCATCTTTCCAGACCTCAGAGTAAACCT
GAAACCGGAAAGCTTCAGGCACCCGCCGTTATCTTACCCTTCCTCAACAAAATAGCTGTTTATTTATACAGTGAGTCCTGTTTCGGACT
TCTCACTCCTCATTTAATCAACTGAAGCTGAACCCGTTTTTTTCTGCATGACTCAATCGTCAAAATTCTTCGCCACCAGTCAGCACTGC
CATAACTTGTCACATTGCAAAAGTAGCCAAATTGCTTCAGGCACCATCAGTTCACCATCACCAAAAAAAAGAAACATTCCAGAACTTCC
GCATCTCGCCCAGCCGCTCCGACTTCTATACTGAATAGAAAACGCCAACATAAGAGAAACCTATGCCATTACCCGATTTTCATGTTTCT
GAACCTTTTACCCTCGGTATTGAACTGGAAATGCAGGTGGTTAATCCGCCGGGCTATCTCGAGAGATCCTCTAGATTTAAGAAGGAGAT
ATACATATG
>ybdL
GAGGCAATTCCGACGTCTAAGAAACCATTATTATCATGACATTAACCTATAAAAATAGGCGTATCACGAGGCCCTTTCGTCTTCACCTC
GAGTCGTGCAGGTGATTGAAACTTCCTGGATGTGAAAAGTAGTTAGCCGGGCCGACGACCACGCGGATAGGATTGTGAGGCATGTTGTG
TTGTCCTTATTGTTTTATTTAGACATCTAAACGTCTTGATTGCCAAATACTAGCATCGTGTTATAGTGCCTTCAACACGCAACTTCGTC
AGGTACAATAAAAATGACAAATAACCCTCTGATTCCACAAAGCAAACTTCCACAACTTGGCACCACTATTTTCACCCAGATGAGCGCGC
TGGCGCAGCAACACCAGGCGATTGGATCCTCTAGATTTAAGAAGGAGATATACATATG
>ybdN
GAGGCAATTCCGACGTCTAAGAAACCATTATTATCATGACATTAACCTATAAAAATAGGCGTATCACGAGGCCCTTTCGTCTTCACCTC
GATGGATCCTTCTCCCGATTTATTGACCGTCTAAATGAGAGTTTTTGATATATAACTACAGACAGCTACTATAACTTTCATCTATTTAT
TCACAGCGCGAATGAAAAATAAAACACTCTGCTTTACAGTGTTTACTTTATAATTAACAGACTTATCTCACTGATCACCCTGTAACGTT
CAGAGAGCGTCTTCAGATGTCTATTTATAAAATTCCTCTTCCGCTCAATATTCTGGAAGCTGCACGAGAGCGTATTACCTGGACACTCA
ACACCCTACCCCGCGTTTGTGTCTCATTTTCTGGCGGCAAAGACTCTGGTTTAATGCTGCATCTGGCAGCCGAACTTGCCCGACAAATG
GGCAAAAAAATCTGCGTTTTGTTTATCGACTGGGAGGCGCTCGAGAGATCCTCTAGATTTAAGAAGGAGATATACATATG
>ykiA
GAGGCAATTCCGACGTCTAAGAAACCATTATTATCATGACATTAACCTATAAAAATAGGCGTATCACGAGGCCCTTTCGTCTTCACCTC
GAGGGCAGGTGTATGAAGCCGGTTCGGTGTTTAATGTTCCCGGTCACAGTGAGTTTCATCTGCAAGTTGCCGAACCCACCTCTTATCTG
TGCCGCTATCTGTAATTCCTCGCCTTCCCCTTGAACGGGAGGGCATTTTTCTGAAATATCCTTTCTTTAGCCCATAATAATATTTCCTT
TGCTGCGATTTTTTCAATTTCCGATATATTCATAATTTATCAAGGTTGATATAAATATCAGTGAAGATCTCCAGATATTGTTGCGGAAC
TGGCTACGATAAAAGATAAATCAGATGATGAATGGTGGCGTGCATTGCTGCAAAGTCGCAATGATCATTTACGCCAAACGGCATTGCGC
AACGCACATACCCCGGCGTTGTTGTTAACGGATCCTCTAGATTTAAGAAGGAGATATACATATG
>ybdF
GAGGCAATTCCGACGTCTAAGAAACCATTATTATCATGACATTAACCTATAAAAATAGGCGTATCACGAGGCCCTTTCGTCTTCACCTC
GATGGATCCTTATCTGGCGACGCTGGTTCTCGCTGGCGGATTAAACGTCGATTATCGCCGTACTTCGCGGCATACTTTGCTTATTCTCC
TTCGCCTAAAGGAACGTTTATGGATAAGCAATCACTGCACGAAACGGCGAAACGCCTGGCCCTTGAGTTACCCTTTGTCGAGCTTTGCC
TCGAGAGATCCTCTAGATTTAAGAAGGAGATATACATATG
>cueR
GAGGCAATTCCGACGTCTAAGAAACCATTATTATCATGACATTAACCTATAAAAATAGGCGTATCACGAGGCCCTTTCGTCTTCACCTC
GAGAGCCGTGCGTTGATATGGATTATTGGCTCACTCTTACTCAGCCTTATTGTGGCAATGGTCATGAAGCGCAATAAAACCGTATAACA
TCTCTCTGTGCGCAGTACTTCCTGTATTATTGTGGTGGCGGTCGATATTCGCACTGGCAAAAAAACGTGCTTGAATATCTGTTGAAACC
CTTTAACAAAGCACAGGAGGCGTTGCGCGAACGATGAACATCAGCGATGTAGCAAAAATTACCGGCCTGACCAGCAAAGCCATTCGCTT
CTATGAAGAGAAGGGGCTGGTGACGGGATCCTCTAGATTTAAGAAGGAGATATACATATG
>frlA
GAGGCAATTCCGACGTCTAAGAAACCATTATTATCATGACATTAACCTATAAAAATAGGCGTATCACGAGGCCCTTTCGTCTTCACCTC
GAGCGACTACCTGCTGCACCCGGCAATTTCTATTTCTAAAATCATTGGCGGTTGCGGTCCTACTGCACAGTAAGTCTCTGACAGAAATC
GGCTAACACCGGTTTAATTGCCCGACGCCCTCTGGATTTTCCGAGGGCGTATTTGTTTCTATGTCCTTCTTGCCCCCGCCCGTAAAAAT
ATTTTCACTTTAAATTCAATTTGATAACTACATCCATTAGCCACAGCGCGATCCTGTGCGAAATTTTGTGATCTTCCTCCACATTACAT
AACATCATATGTTGTTATATTCATCATGCATTGTCATGTTACCTTTTAAATGACTGCAAACTCTCCCCTACAACGTATTGGACAAGAAA
AAGGTATCGCTATGGGAAGCCAGGAACTCCAACGCAAGCTCGGATTTGGATCCTCTAGATTTAAGAAGGAGATATACATATG
>nanA
GAGGCAATTCCGACGTCTAAGAAACCATTATTATCATGACATTAACCTATAAAAATAGGCGTATCACGAGGCCCTTTCGTCTTCACCTC
GATGGATCCCTGCTACCTGGCACGCTTTCGGTCAGACCACCAACAAAAAGAAATAATGCCACTTTAGTGAAGCAGATCGCATTATAAGC
TTTCTGTATGGGGTGTTGCTTAATTGATCTGGTATAACAGGTATAAAGGTATATCGTTTATCAGACAAGCATCACTTCAGAGGTATTTA
TGGCAACGAATTTACGTGGCGTAATGGCTGCACTCCTGACTCCTTTTGACCAACAACAAGCACTGGATAAAGCGAGTCTGCGTCGCCTG
GTTCAGTTCCTCGAGAGATCCTCTAGATTTAAGAAGGAGATATACATATG
>cueO

GAGGCAATTCCGACGTCTAAGAAACCATTATTATCATGACATTAACCTATAAAAATAGGCGTATCACGAGGCCCTTTCGTCTTCACCTC
GAGTAGCACACTGCGAGCGAGGCGACGACCATTATTGTCGATTAAATTGTGTCTGCGGCTTGACCTTCCCGTAAGGGGAAGGACTATGC
TCAACGTTTGATTTTGTTTCGCCTGCTTAAGAATAAGGAAATAACTATGCAACGTCGTGATTTCTTAAAATATTCCGTCGCGCTGGGTG
TGGCTTCGGCTTTGCCGCTGTGGAGCCGCGCAGTATTTGCGGCAGAACGCCCAACGTTAGGGGATCCTCTAGATTTAAGAAGGAGATAT
ACATATG
>nanR
GAGGCAATTCCGACGTCTAAGAAACCATTATTATCATGACATTAACCTATAAAAATAGGCGTATCACGAGGCCCTTTCGTCTTCACCTC
GATGGATCCCAGCCAGCACCACCTGAGGTTTGTAGTTTTTCAGGATCAAATAGCCCATCGTAATTAATACGATGACAGATATAATTATG
CCGAACATTGTTGAACTCCGTGTCAAAAGAAAACGGTCAATCCCATAAACGGCAGATTGAAAACAACGATGTTATATTTTTTGCAAGGC
TATTTATGGTGCGGATGTCGTGTTTTTAATTGTAGGTGAGGTGATTTTTCATTAAAAAATATGCGCTTATGATTATTTTGTAAGAACAC
ATTCATAATATTCATAATGCTCGTGAATAGTCTTATAAATAATTCAAACGGGATGTTTTTATCTGCGTTACATTAATTTTTCGCAATAG
TTAATTATTCCGTTAATTATGGTAATGATGAGGCACAAAGAGAAAACCCTGCCATTTTCCCCTACTTTCAATCCTGTGATAGGATGTCA
CTGATGATGTTAATCACACTGACCTTACAGAATGGGCCTTATGAACGCATTTGATTCGCAAACCGAAGATTCTTCACCTGCAATTGGTC
GCAACTTGCGTAGCCGCCCCTCGAGAGATCCTCTAGATTTAAGAAGGAGATATACATATG
>nanT
GAGGCAATTCCGACGTCTAAGAAACCATTATTATCATGACATTAACCTATAAAAATAGGCGTATCACGAGGCCCTTTCGTCTTCACCTC
GATGGATCCCAGAACTGAAGGCGCTGGCCCAGCAGTTGATGCAAGAGCGCGGGTGAGTTGTTTCCCCTCGCTCGCCCCTACCGGGTGAG
GGGAAATAAACGCATCTGTACCCTACAATTTTCATACCAAAGCGTGTGGGCATCGCCCACCGCGGGAGACTCACAATGAGTACTACAAC
CCAGAATATCCCGTGGTATCGCCATCTCCTCGAGGAGATGGCGATACCACGGCCGCTCGAGAGATCCTCTAGATTTAAGAAGGAGATAT
ACATATG
>trmU
GAGGCAATTCCGACGTCTAAGAAACCATTATTATCATGACATTAACCTATAAAAATAGGCGTATCACGAGGCCCTTTCGTCTTCACCTC
GATGGATCCCAAAGCGGGCAACGTTATCCGCTGGAGATGATTGGCGATTTTAACTGGCCTTTTACAAAGGGTGTCATCTAAAAGGCTGC
GTGATAGAATACGCCGCCTTGAAGTTCAATGTCGTGAGTGATCCAATGTCTGAAACCGCAAAAAAAGTAATCGTCGGCATGTCCGGCGG
TGTCGATTCCTCCTCGAGAGATCCTCTAGATTTAAGAAGGAGATATACATATG
>trmA
GAGGCAATTCCGACGTCTAAGAAACCATTATTATCATGACATTAACCTATAAAAATAGGCGTATCACGAGGCCCTTTCGTCTTCACCTC
GATGGATCCGTATCCTGTGCCCAAGCGGAAAATGCCGTGACGGAACACGCCGTCAGCAGCGAAGCTTTTTTAATCATTGTAAAGCATCC
ACAATAGAAGAAGGATGCCGCAGGTTTCATCAATATTACGCGATGATGAGAACCAGATGCGACGTTGGCCGGCAGGTCTTCGGGCTTGG
AGGGGTATCTAAGATACTAAGAGATGATGACTTCCCACCGAATGGCAGTGTCCGCATAACGCAATCATCGCACCTTTCCTTACCGCTGC
GCGTCAGCTCCAGATTCGCACTGGATTCCCTATTAACTCACAGGACCGGCAAGTGGATGCTACAGGTTGTAACAAGTTACTGTCCAGAC
GTAGCTCACAAATAGGAATTCATCAAGATCTGGACATCTGATGAGCAATCCCTACAATCGCCGCGTACTTTAATTTTTCAGGATACATC
ATGACCCCCGAACACCTTCCAACAGAACAGTATGAAGCGCAGTTAGCCGAAAAAGTGGTACGTTTGCAAAGTATGATGGCACCGTTTTC
TGACCCTCGAGAGATCCTCTAGATTTAAGAAGGAGATATACATATG
>trmE
GAGGCAATTCCGACGTCTAAGAAACCATTATTATCATGACATTAACCTATAAAAATAGGCGTATCACGAGGCCCTTTCGTCTTCACCTC
GAGTCGTCAGCAACCTGGTAACCATTATTCAGCAGCAGCTGATTTACCGTGGTCTGGAAAAACGTGGCCTGCATAGCCGCGAGAAGAAA
AAATCCTGATTCGGTGAGTTTTCGCTAAAATAAGGGCGGTCAGTTGACCGCCTTTTTTCTTTTCGTAGGGCGGATAAGCACCGCGCATC
CGCCACACAAAGCAACAGGAACATCATGAGCGATAATGACACTATCGTAGCCCAGGCCACGCCTCCGGGACGTGGCGGCGTTGGCATCC
TGCGCATCTCCGGCTTCAAAGCCGGATCCTCTAGATTTAAGAAGGAGATATACATATG
>dgoR
GAGGCAATTCCGACGTCTAAGAAACCATTATTATCATGACATTAACCTATAAAAATAGGCGTATCACGAGGCCCTTTCGTCTTCACCTC
GATGGATCCGCGATAGCCCAGTGATGGTGTCTGACTGCGATTGATACTTTCCTGAACTACCCTTTGCGGTATGGGAAACCAGAGCATAC
GAGATACAGCCTGAAAGCATCAGCGCCAGTGGGAATAAACCGGCAGCCTTAAAAAATCCCTTAAAATTCAACTTCATGATTAATCCTTT
AAACGAGTAGCTAAGCCGTTGAAAGCATACTCTTCTTTCCCTCCTGTTTAGCGCCTGATATCCCTTTTCAGATTTCTGCCCGACGCATG
TCATTTTTTTATGCATTGTTCTTTTTGTGATCTAAATTGTAGTACAACAATATAAGTTTGTACTACATTACACGCACGGCAAACGCGAA
CGTCATCACGCTGGTACTACAAAGTTGCCGCGTTATGCATCGATCGGGGTAAAGTAGAGAAGAACATACAGAGCACAAGGACTCTCCAT
GACTCTCAATAAAACCGATCGCATTGTCATTACGCTGGGTAAACAGATCGTTCACGGCAAATACGTGCCAGGCTCGCTCGAGAGATCCT
CTAGATTTAAGAAGGAGATATACATATG
>ggt
GAGGCAATTCCGACGTCTAAGAAACCATTATTATCATGACATTAACCTATAAAAATAGGCGTATCACGAGGCCCTTTCGTCTTCACCTC
GATGGATCCGGCAGGAGTGCCGTAAGAATCAGAAGTCGTTTCATCGCGTATCCTCCTCTGAAGATATCCTTTAAGTTTACTCGCTTCCC
GACAAAACGATGATTAATTCAGAGTTATATACCAGGCTTAGCTGGGGTTGCCCCTTAATCTCTGGAGAATAACGATGATAAAACCGACG
TTTTTACGCCGGGTGGCCATTGCTGCTCTGCTCTCAGGAAGTTGTTTTAGCGCCGCCTCGAGAGATCCTCTAGATTTAAGAAGGAGATA
TACATATG
>ybjS
GAGGCAATTCCGACGTCTAAGAAACCATTATTATCATGACATTAACCTATAAAAATAGGCGTATCACGAGGCCCTTTCGTCTTCACCTC
GATGGATCCGGCAAAACAAATCGCCAGACTGGCAGAACAAAGCACAGATTAATCGACAAAACCTGCGTTATTCTTTCATGTTTCCCATT
GCATCATGGGGGAAATCACGGAAGAATGCGCACGTTATTTTCTCAACAACAGTGGATTGATATGAAGGTACTGGTTACCGGCGCCACCA
GCGGCTTAGGTCGAAACTCGAGAGATCCTCTAGATTTAAGAAGGAGATATACATATG
>ybjQ
GAGGCAATTCCGACGTCTAAGAAACCATTATTATCATGACATTAACCTATAAAAATAGGCGTATCACGAGGCCCTTTCGTCTTCACCTC
GAGCACTGCGTGTGCCGTTATCTTTATAAGCTGGAGTGACTGTGGTGCAGGCACTGAGGAGCAGTGCGCAGGGGATAAGCATTGTCAAT

TTGCTGTAGCGCATAATGATTTCCTTATAAGCGATCGCTCTGAAAGCGTTCTACGATAATAATGATATCCTTTCAATAATAGCGTATCA
GTCTGATAATGCTTTTGAGATCGAAGGCTTAGCAAACAAGGAGATCGATCATGCAATTTTCTACAACCCCAACTCTGGAAGGCCAGACC
ATCGTTGAATATTGCGGTGTGGTGACCGGCGAAGCGATTTAGGATCCTCTAGATTTAAGAAGGAGATATACATATG
>ybjP
GAGGCAATTCCGACGTCTAAGAAACCATTATTATCATGACATTAACCTATAAAAATAGGCGTATCACGAGGCCCTTTCGTCTTCACCTC
GATGGATCCGCCTTCCAGAGTTGGGGTTGTAGAAAATTGCATGATCGATCTCCTTGTTTGCTAAGCCTTCGATCTCAAAAGCATTATCA
GACTGATACGCTATTATTGAAAGGATATCATTATTATCGTAGAACGCTTTCAGAGCGATCGCTTATAAGGAAATCATTATGCGCTACAG
CAAATTGACAATGCTTATCCCCTGCGCACTGCTCCTCAGTGCCTGCACCACAGTCACTCCAGCTTATAAAGATAACGGCACACGCAGTG
CTCGAGAGATCCTCTAGATTTAAGAAGGAGATATACATATG
>ybjG
GAGGCAATTCCGACGTCTAAGAAACCATTATTATCATGACATTAACCTATAAAAATAGGCGTATCACGAGGCCCTTTCGTCTTCACCTC
GATGGATCCGCCTGACGTCCAAGCCTGGCACCGGAAGCTAATTTATTTTGCATGCAATTTCTTCGCCAATAATAATCGCGCAGAGTTTA
ATAAAAGCGCAGCTAACGAGAAAGCGAATTTTGTAGCTGAAACCACGGTTAAGCACATTCTTACATTATTACGAGTATAGCTACGCTTT
CTTTAAGTTTTATTTAACCTATGCCCGTTACAATCACCCACCGTAAACAGGCCGCTTGAGGGAAATAAGACGATGCCGCTTTACCCAGT
TTAACCTGCACTTTATTCTCAACGACTTGCCTGTATTGGCTCCCTTTTAATCACTTTGCGTCGGGAAGTTATGCTGGAAAATTTGAATC
TCTCTCTATTCTCTCTTATTAACGCGACGCCAGACTCGGCTCCGTGGATGATCTCGCTCGAGAGATCCTCTAGATTTAAGAAGGAGATA
TACATATG
>ybjE
GAGGCAATTCCGACGTCTAAGAAACCATTATTATCATGACATTAACCTATAAAAATAGGCGTATCACGAGGCCCTTTCGTCTTCACCTC
GATGGATCCCGGCATTATCGGTGGTCTGATTTACCGGACCCTGCTGGAAAAGCGTGATTAATTAACGCAGAAGGCCTGGCGTAATGTCA
GGCCTTTTTTTAAGATGTTGTTATATGTTGATTCATCATCTTGTTGCGCTCCTTTCATGAGCTAAGCAACATATTGCCACTGGCGCAAG
GAGCGCGCAGGGGGCGGCCAATCGCCGCCGCCCCCTGCACCCCCGGGCTCTGGCGAACAAAATCGCCGCTGCGCGGTGCCCTCGACTTA
TCCCTTACGGCTACCGGGTCGGGCGCGAGGTAACATCCCTGTAAAACGCGCCCTCAGCCCACATCCATGTGGGCTGCCCCGGCCTTCAG
GGAACGCCTCGGCGATTTTGACGCCACCAACATCGGTGCAGTTTGTAATTAAAGCGCAAGAATATCTTTAAATGAAAACGAATACTCTA
CTGCTCCTTATGCATAACATTTCCTTTTAATTGCAACTTTACTCGTCCTTCCGCTTTCAGTAGTGTGTCATCGCGCATTTCGACTTCAT
CTTTGCAAGGACTGGGTTTTCATGTTTTCTGGGCTGTTAATCATTCTGGTTCCCCTGATTGTGGGTTACCCTCGAGAGATCCTCTAGAT
TTAAGAAGGAGATATACATATG
>ybjC
GAGGCAATTCCGACGTCTAAGAAACCATTATTATCATGACATTAACCTATAAAAATAGGCGTATCACGAGGCCCTTTCGTCTTCACCTC
GAGCATCGCGTTCATTGCTCAATTTCTCAGCCAGATCTTTTGCACGCACACAGTAAGGGCAACCCGAACGACCAAAAATAACGGTTTGC
ATTATTTCTCTCCTCATAGATTTATGCCTGTAATGATCACGCTAAAATGTATTCGCTGAAAGTAGGTTTAACCTGTTGCATTAATTGCT
AAAAGCTATAACTGTTAAACACAATACAGTGAAAAGTTTTAGACTGAAGGCTCACTTTGCAGAGGGAAGCGTATGCGCGCGATCGGTAA
ATTGCCTAAAGGCGTGTTGATACTGGAATTTATCGGAATGATGCTACTGGCGGTGGCGCTGCTGTCGGTAAGCGACTCGGATCCTCTAG
ATTTAAGAAGGAGATATACATATG
>ybjO
GAGGCAATTCCGACGTCTAAGAAACCATTATTATCATGACATTAACCTATAAAAATAGGCGTATCACGAGGCCCTTTCGTCTTCACCTC
GAGCGCCTGGTATCTGATGGCTCGCGCAGAAAAACAGCGGATACGCGACATCCAGCGTGCAAGACGTGGCTGAAGACACTAAAATTTGC
CAACCTGGCTACATAATGCCGCGCATGTCGCGGCATTGTTTTCATGGAAGACGAAACGTTGGGATTTTTTAAGAAAACATCTTCATCTC
ATGCTCGCCTGAATGTGCCTGGGATCCTCTAGATTTAAGAAGGAGATATACATATG
>ybjN
GAGGCAATTCCGACGTCTAAGAAACCATTATTATCATGACATTAACCTATAAAAATAGGCGTATCACGAGGCCCTTTCGTCTTCACCTC
GAGATCGACATCGCGGGTAAAATGATCCGCTGGATCGAACGCCACGCTACGACAGAATATTGCCTGAAAACGGGTGGTTAGTCGCAATC
ACATTACTGATCATGGTTTTGCCTGCGCTTTTTGCGTAAGCTGTGCCGGTCTTTTTATCGAAAGAGGTTGTACAAAATTATGACATCGC
TGGTCGTTCCTGGTCTGGATACGCTGCGTCAATGGCTCGATGACCTGGGGATGAGTTTTTGGATCCTCTAGATTTAAGAAGGAGATATA
CATATG
>ybjM
GAGGCAATTCCGACGTCTAAGAAACCATTATTATCATGACATTAACCTATAAAAATAGGCGTATCACGAGGCCCTTTCGTCTTCACCTC
GAGCGAGACATAGCCCAAGCGCGAGGACCACAAATAATAACAGAATGTAATTCCCATTTAACAATTCGGCGACGTTTATATTCACGGAG
GCTAACTTCTTGTTTACTAGTAAGCTGTTGAAAGAAATGGTAATTTACGATAATGTTTTTTACCAGAATTCAGGGCGCAGATTCATTCA
GCGCACCTAAACGATAGTAAAGTAACAATATATTTTACTAGTGTAATCACATTAGGTATCAACGGCTATATGAATTGCGTTGGCCTATA
TTAGCATGGAATGCGAAGCGGCTTTATCTTACTGAACGCCACACTGGCGAAAAATGTGTTCGATAGACGCAGTGTCAGGAGGAACGAGT
GAAACATAAACAACGTTGGGCGGGGGCAATCTGCTGTTTTGTCCTCTTCATTGTGGTGTGCCTTTTTCTGGCGACGCACATGAAAGGCG
CTTTGGATCCTCTAGATTTAAGAAGGAGATATACATATG
>ybjL
GAGGCAATTCCGACGTCTAAGAAACCATTATTATCATGACATTAACCTATAAAAATAGGCGTATCACGAGGCCCTTTCGTCTTCACCTC
GATGGATCCAAAGCGCCTTTCATGTGCGTCGCCAGAAAAAGGCACACCACAATGAAGAGGACAAAACAGCAGATTGCCCCCGCCCAACG
TTGTTTATGTTTCACTCGTTCCTCCTGACACTGCGTCTATCGAACACATTTTTCGCCAGTGTGGCGTTCAGTAAGATAAAGCCGCTTCG
CATTCCATGCTAATATAGGCCAACGCAATTCATATAGCCGTTGATACCTAATGTGATTACACTAGTAAAATATATTGTTACTTTACTAT
CGTTTAGGTGCGCTGAATGAATCTGCGCCCTGAATTCTGGTAAAAAACATTATCGTAAATTACCATTTCTTTCAACAGCTTACTAGTAA
ACAAGAAGTTAGCCTCCGTGAATATAAACGTCGCCGAATTGTTAAATGGGAATTACATTCTGTTATTATTTGTGGTCCTCGCGCTTGGG
CTATGTCTCGCTCGAGAGATCCTCTAGATTTAAGAAGGAGATATACATATG
>ybjK

Data Set S4 - MG1655 library reference file.fasta
GAGGCAATTCCGACGTCTAAGAAACCATTATTATCATGACATTAACCTATAAAAATAGGCGTATCACGAGGCCCTTTCGTCTTCACCTC
GAGCAGGCCTGGCAAAAAGAAGAACATAAACAGCGCCCAGGTTCGGCGTTTCAATGCATTACGTGAAGAATTTACGGTCATAGATCACG
TCAAAATAAGAAGAGGGAAGACAACATTAGCAAGGTTGTGTACATTTGTACACAATTGCAGAGAGAGGAAATGACATGCGTCGCGCTAA
CGATCCGCAACGGCGAGAAAAAATTATCCAGGCCACGCTGGAGGCGGTGAAACTTTACGGAATACATGCTGTTACGCACCGCAAAAGGA
TCCTCTAGATTTAAGAAGGAGATATACATATG
>ybjJ
GAGGCAATTCCGACGTCTAAGAAACCATTATTATCATGACATTAACCTATAAAAATAGGCGTATCACGAGGCCCTTTCGTCTTCACCTC
GATGGATCCCAGCGTGGCCTGGATAATTTTTTCTCGCCGTTGCGGATCGTTAGCGCGACGCATGTCATTTCCTCTCTCTGCAATTGTGT
ACAAATGTACACAACCTTGCTAATGTTGTCTTCCCTCTTCTTATTTTGACGTGATCTATGACCGTAAATTCTTCACGTAATGCATTGAA
ACGCCGAACCTGGGCGCTGTTTATGTTCTTCTTTTTGCCAGGCCTGCTCGAGAGATCCTCTAGATTTAAGAAGGAGATATACATATG
>ybjH
GAGGCAATTCCGACGTCTAAGAAACCATTATTATCATGACATTAACCTATAAAAATAGGCGTATCACGAGGCCCTTTCGTCTTCACCTC
GATGGATCCCCAATAACCGTGAAGGCGTACTGGATGTGATCGATAAAGTTCTTAAACACGAAGCGCCATTTGACCAATAAAACCGGCAG
CACGCCCGGCAACACAATGCCTGAATGAAGGCTGCATAATTCGCGGAAATTAACCCGCTGCATTATTTCAGGCGAAATGATTATGAAAA
ATTGTCTACTGTTGGGCGCGCTTTTAATGGGCTTTACTGGTGTGGCGATGGCGCAAAGTGTCACCGTCGACTCGAGAGATCCTCTAGAT
TTAAGAAGGAGATATACATATG
>ylbE
GAGGCAATTCCGACGTCTAAGAAACCATTATTATCATGACATTAACCTATAAAAATAGGCGTATCACGAGGCCCTTTCGTCTTCACCTC
GAGGCATTACCAATGGTCGCCAGTCGCCGGTGGCAATAAAAAACTGGCTCGTTTATTAGAACGTTTGCAATAAGGGGTTCCCATGTTTA
CATCAGTGGCGCAAGCCAATGCTGCGGTTATCGAACAAATTCGTCGCGCTCGTCCACACTGGCTGGATGTGCAACCGGCTTCTTCACTT
ATCAGCGAACTAAACGAGGGCAAAACACTGCTTCACGCCGGGCCGCCAATGCGCTGGCAGGAGATGACCGGACCCATGAAAGGGGCGTG
CGTGGGCGCATGTCTGTTCGAAGGTTGGGCGAAAGATGAAGCGCAGGCGCTGGCAATACTGGAGCAGGGGGAAGTGAACTTCATTCCTT
GTCACCATGTGAATGCCGTCGGGCCAATGGGCGGTATTACTTCTGCCAGTATGCCGATGCTGGTGGTTGAGAACGGGATCCTCTAGATT
TAAGAAGGAGATATACATATG
>yjjY
GAGGCAATTCCGACGTCTAAGAAACCATTATTATCATGACATTAACCTATAAAAATAGGCGTATCACGAGGCCCTTTCGTCTTCACCTC
GAGTCATAGCCTTCCGCTTCGAAAATACTTTTCAACGTGTTGCGTGTTACCAACTCGTCTTCAACGATAAGAATGTGCGGGGTCTGCAT
GTTTGCTACCTAAATTGCCAACTAAATCGAAACAGGAAGTACAAAAGTCCCTGACCTGCCTGATGCATGCTGCAAATTAACATGATCGG
CGTAACATGACTAAAGTACGTAATTGCGTTCTTGATGCACTTTCCATCAACGTCAACAACATCATTAGCTTGGTCGTGGGTACTTTCCC
TCAGGACCCGGGATCCTCTAGATTTAAGAAGGAGATATACATATG
>ylbG
GAGGCAATTCCGACGTCTAAGAAACCATTATTATCATGACATTAACCTATAAAAATAGGCGTATCACGAGGCCCTTTCGTCTTCACCTC
GATGGATCCGTCTTTCGCGGTGAGTGGGCGGAAGTGGCGGAATGGGTTAAGGCGCGGTAATGAGGGACGTATTTGTATTAGTCGCGACT
TGATCTGACACATGGCCTTGAAAGGTTGAGAGTTACCGGTTTTGATATGAGTGTCGAATCCTTATCCAAAACAAGAGGTAACTCTCATG
CTTCATACTGCCAATCCCGTCATCAAACACAAAGCCGGTTTGCCTCGAGAGATCCTCTAGATTTAAGAAGGAGATATACATATG
>ylbF
GAGGCAATTCCGACGTCTAAGAAACCATTATTATCATGACATTAACCTATAAAAATAGGCGTATCACGAGGCCCTTTCGTCTTCACCTC
GAGAGATTGGCGCAGGCACCGTGCGGGCACCGCTGGCGTGCTTTGAACAGGCGCTGGAAGCACTGGCTGAAAGCATGGGTATTGGTTGA
GGAACGCGCAATGACGATCATCCATCCTCTGCTTGCCAGTAGTAGCGCACCGAATTATCGCCAGTCCTGGCGGTTAGCGGGAGTGTGGC
GGCGGGCGATTAACCTGATGGGATCCTCTAGATTTAAGAAGGAGATATACATATG
>fklB
GAGGCAATTCCGACGTCTAAGAAACCATTATTATCATGACATTAACCTATAAAAATAGGCGTATCACGAGGCCCTTTCGTCTTCACCTC
GAGGCGTGCCAGACTTTCTCCAGGGTTGGTTTTAGTTCAAAGCGCCCGGGCATGGTTTCCTCCTGCTCCGCGTCTTGTTCCTCAATCAT
AGCCTATGAATAAGCTAACGCTATGATGTCCGTGGTAAACCCGCCTTTATTATATTCATACGGGATATTGCTATTGTTTCTTTTTCCCT
GGGATTTGTCATCATTCCCGCGACAAAGTGGATCCTCTAGATTTAAGAAGGAGATATACATATG
>valV
GAGGCAATTCCGACGTCTAAGAAACCATTATTATCATGACATTAACCTATAAAAATAGGCGTATCACGAGGCCCTTTCGTCTTCACCTC
GAGTCGCCTTCAGTAAGTTGCGTCGCGCCAATGACCTCGCCGGTCTTAACAATTTGCACGTTCCCGTCGCTCATCAAGTTTTTCGCATC
AGATCCCATGATTTATTCCTTTGCTGCATCTGTGTGCCTTTATTGCTACCTAAGTGTAAAGGCTACGGAGGATTTATCCACGACAGATT
TGAGATGGTGGCAAACAACTCTGTTTAAACTCTGATACACGAATTATTGGGTTGTATCAGATGTAAATGCGATCCTGAATAAAAATCAC
CCTTGCAAATCAACAAAATATAGCCAATTTGCTTAAAAGCTCTGCAAACGGTCGAAATAGCGCAGAAAATTACGTTTTGCCTCTTGCCA
CCTTCCCACTCTGCCGCTAATATTCGTCCCCGTTGTCACCTACAACGTTGCGTTCATAGCTCAGTTGGTTAGAGCACCACCTTGACATG
GTGGGGGTCGTTGGTTCGAGTCCAATTGAACGCACCATCCTGCGTCCGTAGCTGGATCCTCTAGATTTAAGAAGGAGATATACATATG
>valT
GAGGCAATTCCGACGTCTAAGAAACCATTATTATCATGACATTAACCTATAAAAATAGGCGTATCACGAGGCCCTTTCGTCTTCACCTC
GAGCAAACCGAAAGCAACGAAAAAGTGGGTCGTTAGCTCAGTTGGTAGAGCAGTTGACTTTTAATCAATTGGTCGCAGGTTCGAATCCT
GCACGACCCACCAATCGCTAAGGTGGAAGCGGTAGTAAAACGTGAAGGATAACGTTGCATGAGCAACGGCCCGAAGGGCGAGACGAAGT
CGAGTCATCCTGCACGACCCACCACTAACATAGTTAGTTGTAGTATCCAGCGTAGTATCGGGTGATTAGCTCAGCTGGGAGAGCACCTC
CCTTACAAGGAGGGGGTCGGCGGTTCGATCCCGTCATCACCGGATCCTCTAGATTTAAGAAGGAGATATACATATG
>zraS
GAGGCAATTCCGACGTCTAAGAAACCATTATTATCATGACATTAACCTATAAAAATAGGCGTATCACGAGGCCCTTTCGTCTTCACCTC
GAGCCAACGATGAAACGCTGATATGACGGGTAATCTGGCATGATAAACGAGTAAAAATGACTCGCCTGCTGCGGGTAGCGAGTCATTTT
TACTCATTGAAACTTAAGCCTTTGTGTTACAGCGCAGGGTAAGCGCTGATAAAAGATGGCATGATTTCTGCTGTCAGAAAGGGATGAGC

AGGCAAAGAAGAAGATGCGTTTTATGCAACGTTCTAAAGACTCCTTAGCTAAATGGTTAAGCGCGATCCTCCCCGTGGTCATTGTTGGG
CTGGTGGGATTGTTTGCGGATCCTCTAGATTTAAGAAGGAGATATACATATG
>zraP
GAGGCAATTCCGACGTCTAAGAAACCATTATTATCATGACATTAACCTATAAAAATAGGCGTATCACGAGGCCCTTTCGTCTTCACCTC
GATGGATCCGCAAACAATCCCACCAGCCCAACAATGACCACGGGGAGGATCGCGCTTAACCATTTAGCTAAGGAGTCTTTAGAACGTTG
CATAAAACGCATCTTCTTCTTTGCCTGCTCATCCCTTTCTGACAGCAGAAATCATGCCATCTTTTATCAGCGCTTACCCTGCGCTGTAA
CACAAAGGCTTAAGTTTCAATGAGTAAAAATGACTCGCTACCCGCAGCAGGCGAGTCATTTTTACTCGTTTATCATGCCAGATTACCCG
TCATATCAGCGTTTCATCGTTGGCTCGAGAGATCCTCTAGATTTAAGAAGGAGATATACATATG
>pntA
GAGGCAATTCCGACGTCTAAGAAACCATTATTATCATGACATTAACCTATAAAAATAGGCGTATCACGAGGCCCTTTCGTCTTCACCTC
GATGGATCCGCTTGCTCCGGTGTCAGTTCTGTTGCTGCGTTAACGGAGAATGCCATAGCAGAAAGCAGTGCCGACGCCAGGAGGGTGTT
CTTAAGCTTCATAAAAATAATCCTTCGCCTTGCGCAAACCAGGTACTGGTATTGTTATTAACGAGAAACGTGGCTGATTATTGCATTTA
AACGGTGTAACTGTCTGCGTCATTTTTCATATCACATTCCTTAAGCCAATTTTAATCCTGCTCAAATGACCGTCTATGCTTAAAAAACA
GCCGTATCAGCATCATTACTACTGAAGCAACTGAATTGTATAAGTTAATTTAATGTTAAGTAGTGATTCGTGCCGGGGCGATGTCTCGT
TTTACCCGACCGTCGAAGACAATTATCAGTCTTTATCCGGCGTTCTAAGGTGTTTATCCCACTATCACGGCTGAATCGTTAATATTTTG
CGAGTTCACGCCGAAATACTGATTTTTGGCGCTAGATCACAGGCATAATTTTCAGTACGTTATAGGGCGTTTGTTACTAATTTATTTA
ACGGAGTAACATTTAGCTCGTACATGAGCAGCTTGTGTGGCTCCTGACACAGGCAAACCATCATCAATAAAACCGATGGAAGGGAATAT
CATGCGAATTGGCATACCAAGAGAACGGTTAACCAATGAAACCCGTGTTGCAGCAACGCCAAAAACAGTGGAACAGCTGCTGAAACTGG
GTTTTACCGTCGCCTCGAGAGATCCTCTAGATTTAAGAAGGAGATATACATATG
>aphA
GAGGCAATTCCGACGTCTAAGAAACCATTATTATCATGACATTAACCTATAAAAATAGGCGTATCACGAGGCCCTTTCGTCTTCACCTC
GAGCTATCTCATCGCCAGCGGTCGCATGTGTGTCGCCGGGTTAAATACGGCAAATGTACAACGTGTGGCAAAGGCGTTTGCTGCGGTGA
TGTAATGCAGGAAAGCAGGCTGGAGCTACCCAGCCTGCAGTGAAATTAAACTGTCGTCGCTTTCACTCTTTCTTTATAGATGATTTTTT
TGATGCCATCGTTCTACGTGAGAGATAATAAACGTTGTTAGTTCTTTTATTGTTAAGCTTATCCCAATTATCTGGAATTCCTTATCCTG
TTTTTTTGGGTGGAGTAATAATTTCTGCACATCTAACCATTATTAAATTTAATAATAAGTATCGGGTTTGTTACGATAGAATATTTCTA
TATCCTCAAGAATATGCTTTGCCTTATTGCCTGTGACTTTCTGAGATTCAATTATAGTATCGTTAAATTCTAATGTTAAAGAGAACTCT
TTTTTCCGCTTTGTGTAAGTTTAACCTTATATTGATTACCAATATCAAATAAAAAGGTCCTGTAATGAGAATAATCACCTTTAACCTCC
TCGGCAAAACAGCAAAGCGTGCAAATATACATAAAGTCTTTCTTACCCCTGATTTTGTTAACTAAATCAATAAATGCGTGAAAAACTTT
ACTTGCAATAGAACTTGATACTTCTTCTCCAGGTCTAAAAAATCTTTTGCAATTTTTAACTGTGTTTCTATGTATAAAAAGAAATCTTT
GGCATAAGAGATATCTTTATTATTTAGTGCATCTAGCTCAACAGGTAAAGATGTCGTTTTGTTGCTATTCACATATGATATATTCATAT
CATCATCTCCATCGCGAGTAATAAAATTAATCACCATTGTAGGGTAGGGGGCTGGTCAATCAGAAATCATCTTTATAAACTTCGATTGT
TTTTGTAATGCTGTATCATTAAGTTCATTAAATCGTACAGCAGATAAATGTTCTATCAAATTTCGCTCATTTGCCGAGGATTCATCATA
ATAAACGTAAAATTAATGTATCCTTACATCGAGTAATAAACATTTTTTATACAAAAAAAGACAGGAACGTATTTACTGGGTTAAATATA
ATCATCCTGCTTTTCATCACAAAAACCGCAGATAATCCTTCCTTTCCCCGGCAGCTGGCGTTATGGTCAGATGGTTTTTGCAACAAATC
TCACAATAAAAAGTTTCAACATACTGACTATTTAGGGAAAAATATGCGCAAGATCACACAGGCAATCAGTGCCGTTTGCTTATTGTTCG
CTCTAAACAGTTCCGCTGTTGCCCTGGCCTCATCTCCTTGGATCCTCTAGATTTAAGAAGGAGATATACATATG
>yjjK
GAGGCAATTCCGACGTCTAAGAAACCATTATTATCATGACATTAACCTATAAAAATAGGCGTATCACGAGGCCCTTTCGTCTTCACCTC
GATGGATCCCAACAGCCGCCAGGTAACTTGTTTGGCTTTTTCCACAAGCGCATCCTCTAAGTGTAAATACATCAATGCAGCGTCATGCC
GCGTAATAGATTTATGCTAACCAGTCATTGCCGTTTACGCCACGTTACGGACACTTTTTTACTTTTACTGCGAGGGTGATCGACCGGCT
GGGATTAGGCGGGGAAAAAGGCTACACTTCGCCTTTGAAAACTCATTATCACGATAAACATAGAGGCGAAGTCCAACGTGGCTCAATTC
GTTTATACCATGCATCGTGTCGGCAAAGTTGTTCCGCCGAAACGTCATATTTTGAAAAACATCTCTCTGAGTTTCTTCCCTGGGGCCTC
GAGAGATCCTCTAGATTTAAGAAGGAGATATACATATG
>yjjJ
GAGGCAATTCCGACGTCTAAGAAACCATTATTATCATGACATTAACCTATAAAAATAGGCGTATCACGAGGCCCTTTCGTCTTCACCTC
GAGGCACTCACGAGCAGACCACTGCCGCTGAGCGTCAGACTACCTTCAACGACATGATCAAAATCGCACTGGAATCCGTTCTGCTGGGC
GATAAAGAGTAATTGTGTTTCGCTGCAAGGCGATTGCCTTGTGAAGCCGGAGCGGGAGACTGCTCCGGCTTTTTAGTATCTATTCATTT
TTCTCTCCAGCTTGAATATTTTCGCTATACTTTTCAGTGAAATGTGTTAATAAATCTATTCAAGTATCTATTCACGAATCTATTCATTA
ATGAGCGAGCTGACTGATCTTTTACTGCAAGGGCCGCGTTCTGCCCCGGAATTGCGCCAGCGTCTGGCAATCAGTCAGGATCCTCTAGA
TTTAAGAAGGAGATATACATATG
>tpiA
GAGGCAATTCCGACGTCTAAGAAACCATTATTATCATGACATTAACCTATAAAAATAGGCGTATCACGAGGCCCTTTCGTCTTCACCTC
GATGGATCCCGAAAAGGGGCTGACCTTCGCTGTTGAACCGATTAAGCTGGCGCTATCTGAATCGCTTGAAGGTTTGAATAAATGACAAA
AAGCAAAGCCTTTGTGCCGATGAATCTCTATACTGTTTCACAGACCTGCTGCCCTGCGGGGCGGCCATCTTCCTTTATTCGCTTATAAG
CGTGGAGAATTAAAATGCGACATCCTTTAGTGATGGGTAACTGGAAACTGAACGGCAGCCGCCACATGGTTCACGAGCTGGTTTCTAAC
CTGCGTAAAGAGCTGGCCTCGAGAGATCCTCTAGATTTAAGAAGGAGATATACATATG
>pldB
GAGGCAATTCCGACGTCTAAGAAACCATTATTATCATGACATTAACCTATAAAAATAGGCGTATCACGAGGCCCTTTCGTCTTCACCTC
GAGAGGTCAATTGCACCAGCGGCGCGCCACTGCTGGATTCCCAGCCAAATCAAGTAAGCCGCGCCTGCCCACTTCAACACTTCAAACGC
AATCACTGAGCGGGAAAATAGCGTCCCCAACCCCACGCCAACCAGCACAATATGAATCGCCAGTCCGGTCTGAAGCCCAGCAATAGACG
CCACCGCGCCGCGATAACCGTGGTTGAGCGAGGTGGTCATAGTGTTGATTGCACCAGAGCCTGGCGACAGGCTTAAAATGATCGATGTC
AGCAGGTAGGCAAACCACCATTCTAAGGTCATGATGAACTCCCGGTGTGTCTATTTTTGTGCCACAATACGCTACTGTCGCAGCGTTGT

GTCAGGCACGCTAAAAAAAACGATTTTACGTGGTTTAAGAGGCAGATTACCCGATGTTTCAGCAGCAAAAAGACTGGGAAACAAGAGAA
AACGCGTTTGCTGCTTTTACCATGGGACCGCTGACTGATTTCTGGCGTCAGCGTGGATCCTCTAGATTTAAGAAGGAGATATACATATG
>malX
GAGGCAATTCCGACGTCTAAGAAACCATTATTATCATGACATTAACCTATAAAAATAGGCGTATCACGAGGCCCTTTCGTCTTCACCTC
GAGGCCCTTTGCCACTAAGCACCAGCGAAACGGTACTTACCGACACGCCCGCAGCCAGCGCAACATCATGAATGGTTATTTTTTTGGCG
GTAGCCATGCAGATGACCTACTCCCTGATTATGTGACAGATAAAACGTTTTACCTTTTATTTATCTTATACCCGCTATTATCGTTGCGT
AATGTGATTTATGCCTCACTAAAATTTGATAAAACGTTTTATCTTCTCGCGCAATTTACTGAATCCAGATTGTTCTCTACGAGGAGTCG
TTTTATGACGGCGAAAACAGCACCGAAAGTCACGCTGTGGGAGTTCTTCCAGCAGTTAGGCAAAACCTTCATGTTACCCGTGGCATTAT
TGTCGTTCTGCGGCAGGATCCTCTAGATTTAAGAAGGAGATATACATATG
>malS
GAGGCAATTCCGACGTCTAAGAAACCATTATTATCATGACATTAACCTATAAAAATAGGCGTATCACGAGGCCCTTTCGTCTTCACCTC
GAGTGTTGTGTGCGTCTTCGCTAACACCTCACTCGAAAACACCAGAGTGAGTAACATAAGAATCATCGCCCCATATCGTCGTATGGGAG
TCAAAATCATCAGGTCTCCTGGTCGGATTTAATCATTCCAACACCTTATATTTTTCACAAATTTGAGAGTTGAATCTCAAATCATATCA
AAAATAGCTGTCAAGAGCACCCCAAGGAATAGTCCAAATCTGAAACTATGTCACGTGTTAACGATTCAGATTGGCGCTAAATCGCAGAA
AATGTGGGGGTTATCGCAAAATTCAGCCGTTTTTTGCGCGAGATCGCTCACCCTTGCTTCTCATCCTGTGGACTTACCGCTCAGGGATG
AGTTTTGTTTGGCTTATCGCTGGCAAACTGTCTGAAATCGCAGCAATAAGGACTCATCCGCCATGAAACTCGCCGCCTGTTTTCTGACA
CTCCTTCCTGGCTTCGCCGTTGCCGCCAGCTGGACTTCTCCGGGGTTGGATCCTCTAGATTTAAGAAGGAGATATACATATG
>malI
GAGGCAATTCCGACGTCTAAGAAACCATTATTATCATGACATTAACCTATAAAAATAGGCGTATCACGAGGCCCTTTCGTCTTCACCTC
GATGGATCCTGCCGCAGAACGACAATAATGCCACGGGTAACATGAAGGTTTTGCCTAACTGCTGGAAGAACTCCCACAGCGTGACTTTC
GGTGCTGTTTTCGCCGTCATAAAACGACTCCTCGTAGAGAACAATCTGGATTCAGTAAATTGCGCGAGAAGATAAAACGTTTTATCAAA
TTTTAGTGAGGCATAAATCACATTACGCAACGATAATAGCGGGTATAAGATAAATAAAAGGTAAAACGTTTTATCTGTCACATAATCAG
GGAGTAGGTCATCTGCATGGCTACCGCCAAAAAAATAACCATTCATGATGTTGCGCTGGCTGCGGGCGTGTCGGTAAGTACCGTTTCGC
TGGTGCTTAGTGGCAAAGGGCCTCGAGAGATCCTCTAGATTTAAGAAGGAGATATACATATG
>malF
GAGGCAATTCCGACGTCTAAGAAACCATTATTATCATGACATTAACCTATAAAAATAGGCGTATCACGAGGCCCTTTCGTCTTCACCTC
GATGGATCCTGAAGCCCTGAAAGACGCGCAGACTCGTATCACCAAGTAATGCTGTGAAATGCCGGATGCGGCGTGAACGCCTTGTCCGG
CCTACAAAACCGAAACGTATGTAGGCCTGATAAGACGCGTCAGCGTCGCATCAGGCAGTTGTTGTCGGATAAGGCGTGAAAGCCTTATC
CGTCCTGGAATGAGGAAGAACCCCATGGATGTCATTAAAAAGAAACATTGGTGGCAAAGCGACGCGCTGAAATGGTCAGTGCTAGGTCT
GCTCGGCCTCGAGAGATCCTCTAGATTTAAGAAGGAGATATACATATG
>malE
GAGGCAATTCCGACGTCTAAGAAACCATTATTATCATGACATTAACCTATAAAAATAGGCGTATCACGAGGCCCTTTCGTCTTCACCTC
GATAGATCTGCCCCAGGCTTTCGTTACATTTTGCAGCTGTACGCTCGCCATGCCCTTCTCCCTTTGTAACAACCTGTCATCGACAGCAA
CATTCATGATGGGCTGACTATGCGTCATCAGGAGATGGCTTAAATCCTCCACCCCCTGGCTTTTTTATGGGGGAGGAGGCGGGAGGATG
AGAACACGGCTTCTGTGAACTAAACCGAGGTCATGTAAGGAATTTCGTGATGTTGCTTGCAAAAATCGTGGCGATTTTATGTGCGCATC
TCCACATTACCGCCAATTCTGTAACAGAGATCACACAAAGCGACGGTGGGGCGTAGGGGCAAGGAGGATGGAAAGAGGTTGCCGTATAA
AGAAACTAGAGTCCGTTTAGGTGTTTTCACGAGCACTTCACCAACAAGGACCATAGATTATGAAAATAAAAACAGGTGCACGCATCCTC
GCATTATCCGCATTAACGACGATGATGTTTTCCGCCTCGGCTCTCGCCAAAATCGAACTCGAGAGATCCTCTAGATTTAAGAAGGAGAT
ATACATATG
>crcB
GAGGCAATTCCGACGTCTAAGAAACCATTATTATCATGACATTAACCTATAAAAATAGGCGTATCACGAGGCCCTTTCGTCTTCACCTC
GATGGATCCAAACTGTCCTGCGGCAACCAGCATTGCCACCTCCTGTCAGAGTCGTAAAATTCGCCAACGACGATTATTCGGCGTTACAC
TTATCACTCATACAAATCAAATAGCAGGATTTTGCAGTGTTACAACTTCTTTTAGCAGTTTTTATTGGCGGTGGTACGGGAAGCGTGGC
GAGATGGCTGTTAAGTATGCGATTTAACCCACTGCATCAGGCGATTCCCTCGAGAGATCCTCTAGATTTAAGAAGGAGATATACATATG
>yahD
GAGGCAATTCCGACGTCTAAGAAACCATTATTATCATGACATTAACCTATAAAAATAGGCGTATCACGAGGCCCTTTCGTCTTCACCTC
GAGCCTTGAGAGAGCCCGACATGTGAAGACACTAACTGCCAAAGCCCTGCACAAATACCGACGGTGACACCCGTTGCCGTGAGTCCGTT
CATCTTGTTCCCCTCTGGTTAGACATGGACGAAAGCTACTCTTACATGTGCAGACTATAGAGAAATAAAACCTGATTGAGTGAAAGGGT
ATCGGGTCAAAGAAACAAATATTGCACTACCGCACACTGCGAAAAGATTGTTGAATATTATTCAACAAAAAGGCGAGATACGGCGAATT
AAATTTAAGCGGGTTCTTTTATCAGATTATCGTTGCTATTCTCAATTCATCGAAAACAGCCACAAAAACCAGGGGAATGTGAATGAGTA
TAAAAAATCTACCTGCCGACTATTTATTGGCTGCACAACAGGGTGATATCGATAAAGTAAAAACCTGCCTTGCGCGGATCCTCTAGATT
TAAGAAGGAGATATACATATG
>yahB
GAGGCAATTCCGACGTCTAAGAAACCATTATTATCATGACATTAACCTATAAAAATAGGCGTATCACGAGGCCCTTTCGTCTTCACCTC
GATGGATCCTACTGGGTTTTCTGGCGGGTAACTGCGTGGGGGTAATTTCAGAATATGGTGGACAAAAATTAAGCGAAGCGACGACTAAA
CGCGACGGGTATTAAAAGTTATTCAAAAATGATTATCTATCTTTAACCGATACCTCTACAGTCATAGCTGGAGAAATTCATAGTAGATA
ATGGCGAGAATGGTTTAATGAACTCAATTTTTACCGAAGAGAATCTGTTGGCATTTACTACCGCTGCGCGTTTTGGCAGCTTCAGTAAA
GCCGCAGAAGAGTTGGCTCGAGAGATCCTCTAGATTTAAGAAGGAGATATACATATG
>yahC
GAGGCAATTCCGACGTCTAAGAAACCATTATTATCATGACATTAACCTATAAAAATAGGCGTATCACGAGGCCCTTTCGTCTTCACCTC
GATGGATCCGCGCAAGGCAGGTTTTTACTTTATCGATATCACCCTGTTGTGCAGCCAATAAATAGTCGGCAGGTAGATTTTTTATACTC
ATTCACATTCCCCTGGTTTTTGTGGCTGTTTTCGATGAATTGAGAATAGCAACGATAATCTGATAAAAGAACCCGCTTAAATTTAATTC
GCCGTATCTCGCCTTTTTGTTGAATAATATTCAACAATCTTTTCGCAGTGTGCGGTAGTGCAATATTTGTTTCTTTGACCCGATACCCT

TTCACTCAATCAGGTTTTATTTCTCTATAGTCTGCACATGTAAGAGTAGCTTTCGTCCATGTCTAACCAGAGGGGAACAAGATGAACGG
ACTCACGGCAACGGGTGTCACCGTCGGTATTTGTGCAGGGCTTTGGCAGTTAGTGTCTTCACATGTCGGGCTCTCTCAAGGCTCGAGAG
ATCCTCTAGATTTAAGAAGGAGATATACATATG
>yahN
GAGGCAATTCCGACGTCTAAGAAACCATTATTATCATGACATTAACCTATAAAAATAGGCGTATCACGAGGCCCTTTCGTCTTCACCTC
GATGGATCCGGTCATCAATTCAGCGGCATAGACATTGGTAACGGCTAACGCTAACGCACCGACTAACATTTTAGAGATTATTTTCATAT
TACGGCTCCAGAGTTGTTATCACACTATCACCCACCCACATTATTGGGTTGATTAATAATTAGTTCAGTCAGGGCATAAACGCCAGTAA
CCGGAAATAAGACACTTTGTGATCTCTCTCGCACCCGATTATTTTTGGTAGAAATATGATGCAGTTAGTTCACTTATTTATGGATGAAA
TCACTATGGATCCTTTGCATGCCGTTTACCTGACCGTCTCGAGAGATCCTCTAGATTTAAGAAGGAGATATACATATG
>yahO
GAGGCAATTCCGACGTCTAAGAAACCATTATTATCATGACATTAACCTATAAAAATAGGCGTATCACGAGGCCCTTTCGTCTTCACCTC
GAGACGGTCAGGTAAACGGCATGCAAAGGATCCATAGTGATTTCATCCATAAATAAGTGAACTAACTGCATCATATTTCTACCAAAAAT
AATCGGGTGCGAGAGAGATCACAAAGTGTCTTATTTCCGGTTACTGGCGTTTATGCCCTGACTGAACTAATTATTAATCAACCCAATAA
TGTGGGTGGGTGATAGTGTGATAACAACTCTGGAGCCGTAATATGAAAATAATCTCTAAAATGTTAGTCGGTGCGTTAGCGTTAGCCGT
TACCAATGTCTATGCCGCTGAATTGATGACCAGATCCTCTAGATTTAAGAAGGAGATATACATATG
>ycdZ
GAGGCAATTCCGACGTCTAAGAAACCATTATTATCATGACATTAACCTATAAAAATAGGCGTATCACGAGGCCCTTTCGTCTTCACCTC
GAGCACCATGGCACCGCTAACCCGCGATGCCATTAGTGCAATGTGGGACGAGCTGGAAGAAGATTCTGAAGAATAAGTGTGATCTACGT
CACTCATAACTGCAACGGATAATTTGTTGTTGCATAAAATGTGTGCTCGATCTCATTCATGGCCGCGTTTTCTGCTATCATGCGCGGCA
TGAACATACTTCTCTCTATTGCAATCACAACAGGCATTCTCTCCGGGGATCCTCTAGATTTAAGAAGGAGATATACATATG
>yahM
GAGGCAATTCCGACGTCTAAGAAACCATTATTATCATGACATTAACCTATAAAAATAGGCGTATCACGAGGCCCTTTCGTCTTCACCTC
GAGGCGCGATTAAATGAATGGATATCTAATAGAGTGCAGGAATTAGGTGACAGATATTATCCTGTAAATAAGCACGTTAAAATCCGCTA
CACTCTTTAGTTCAGAAAGGGTTCCACCTGCAAGTTGTTATTCCACCTGAGAGTGAATCGCAATGCAGGTAATCATTTCATCTGTATAT
CTGTATGCCAACTGGCATTCCATCTCTTTTTTCGCCGATATTTTGTTGTGATCGGCGACACTTCGCTTAAAAAAGCACCAGTAGTGGTT
TCGCAGCCATGCGGTGTATAAAAAATGATCTCATGCAGATGTTTTGTGAATGTGTTGGTTGACATTCATATGAAAAAAATCATAATTCC
ATCATGTTTGTGTGGGGTCTTTTCTGTATCTTACGCATCGCACTCAAGCCTGACAGAAAATATGCTGTAAGGCTCATATCAAAAACCGC
CATTAGCTCATCAGGAAGAGCAGACGACAACCAGGATTGTTGTATGGTACGGGGTTCGAGGCCTCGATGGCGGTCCAACTTTTCAAGGA
TCCTCTAGATTTAAGAAGGAGATATACATATG
>yahK
GAGGCAATTCCGACGTCTAAGAAACCATTATTATCATGACATTAACCTATAAAAATAGGCGTATCACGAGGCCCTTTCGTCTTCACCTC
GAGGGTGGACGCCTCCTGTTCCGCCGAGGCGGTGGCGCGTATCTCGCCGAGAACCGCAACGTTCCATAAAGGGCAACTGGTGTGGGGGA
GTGTGGCAGGTTGATGCGCATTTGAAACGCCTGCAACGGTGAATAGTAAGAGATTTAAGCCCCAGGGAGTGGGGCAATCTGAATATGAG
TGTGTCGGCAGAAATGGACATTATCTTTGAGGAATATGCCCTTATTGACGTTGTGTAGATAACTATTTGTCTACGTGAAAAGATCATCA
GTTTTGCCGATTTTAGTCTTTTACAGATAGCAAATATCACACTTACAGGCGCTCGCCTTATGGGGAACGTTGCTGCTCTGTTTCATTCA
GGCTCGCGGCATGTTGGGGCTGGATCGGGTGGTTTAAATATTGGCTATATTCAATGGACGCGTTTTGCCGCGATGACATATCAGGCGTT
GCCAAATACACATAGCTAATCAGGAGTAAACACAATGAAGATCAAAGCTGTTGGTGCATATTCCGCTAAACAACCACTTGAACCGATGG
ATATCACCCGGCGTGAACCGGGACCGAATGATGGATCCTCTAGATTTAAGAAGGAGATATACATATG
>yahI
GAGGCAATTCCGACGTCTAAGAAACCATTATTATCATGACATTAACCTATAAAAATAGGCGTATCACGAGGCCCTTTCGTCTTCACCTC
GAGCGCATCAGGCATTTGTCACCATTGCCGGATGCGGCGTGAACGCCTTATCCGGCCTACGAATGGCGCAAGAATCTGTAGGCCTGATA
AGCGTAGCGCATCAGGCATTTGCCACCACTGCCGGATGCGGCGTGGACGCCTTATCCGGCCTACGAGTGGCGCGAGAATCTGTAGGCCT
GATAAGCGTAGCGCATCAGGCATTTGTCACCATTGCCGGATGCGGCGTGAACGCCTTATCCGGCCTACGAGTGGCGCGAGAATTGTAGG
CCTGATAAGCGTAGCGCATCAGGCAGTCTGGCGTTGGTCATAACCCCATCACCCTCTGTAGCGGACATAACAACATGAAAGAGCTTGTG
GTCGTTGCCATTGGGATCCTCTAGATTTAAGAAGGAGATATACATATG
>ycdC
GAGGCAATTCCGACGTCTAAGAAACCATTATTATCATGACATTAACCTATAAAAATAGGCGTATCACGAGGCCCTTTCGTCTTCACCTC
GAGCAGCCGTTGTTGCCAATAGGTACGAATACGCCAATTTTCATCATCAACCTCTCTTCGTCTCGTAAAGTGAAAGTCAGACGGGGCGC
TGCATCCTGCATATCCTTTTCAGCCGCGTATTGGCTTGTTTGCAAAGCGGATGCCAGTTTTTAAAAAGTTAATGTTATTAATCTGTTAA
CATTACGTTATCTAAAATATCTGGTAAAAAGTGGACTAAACGGTCAAAACAGTTGCACATAAAACATGCATCTGTGCGCGATGAGAGTG
CAGAAGGTCGAGGCCGGGCGGGGGTTTTGCTATCCTGTTGCCAATCTACAAGAGGGGAGAGCGCATGACGCAAGGCGCAGTGAAAACAA
CGGGTAAACGTTCGCGCGCAGTAAGCGCGAAGAAAAAAGCGATTCTTAGCGCAGCACTGGACACGGATCCTCTAGATTTAAGAAGGAGA
TATACATATG
>ibpB
GAGGCAATTCCGACGTCTAAGAAACCATTATTATCATGACATTAACCTATAAAAATAGGCGTATCACGAGGCCCTTTCGTCTTCACCTC
GATGGATCCGTGATTCCGGAAGCGAAAAAACCGCGCCGTATCGAAATCAACTAATTCCCTAAGGCCGCCTGGCGCGGCCTGACATCTCC
ATGCTCGCCGTCAGGGAGCATATGCGAATCTTCGGATTTGCAGGTACTTACTCGCTTCTTAGAAGGAGAAATGACTATGCGTAACTTCG
ATTTATCCCCACTGATGCGTCAATGGATCGGTTTTGACAAACTGGCCAACGCACCTCGAGAGATCCTCTAGATTTAAGAAGGAGATATA
CATATG
>eutR
GAGGCAATTCCGACGTCTAAGAAACCATTATTATCATGACATTAACCTATAAAAATAGGCGTATCACGAGGCCCTTTCGTCTTCACCTC
GATGGATCCCGCTGGAAAAAGCCAGAAATGCGCTCGAACAGCTCTTTTCTGCCGGGACGTTACGTAAACGCAGTAGTCGTTATCGTCTC

AAGCCCCATTAACCTGTCGGAGGTGCCGGGTGTCCTACAACACCCGGCATTAACATCATGAAAAAGACCCGTACAGCCAATTTGCACCA
TCTTTATCATGAACCCTTACCCGAAAACCTGAAGCTCACGCCGAAGCTCGAGAGATCCTCTAGATTTAAGAAGGAGATATACATATG
>eutS
GAGGCAATTCCGACGTCTAAGAAACCATTATTATCATGACATTAACCTATAAAAATAGGCGTATCACGAGGCCCTTTCGTCTTCACCTC
GATGGATCCTGGTAGAAGCGCAAACCCAACCGCTGTAATTTTTTTTAACTCTCACGCTTACCCTGAATATTCAGGGTAAGCAGTTTAGC
TGCAATATATTAGTAAAGCTTATTACTGAGTTTGCGAATAATAAAAAAAAGCAGTCTATATAATATCTCGATATTATTTTATTTATATT
CATGCGTTGCATATGAAAGTTTATGCACCACAGCGAATATCTCTCCATTCTTAGTGATCTACCTCACCTTTTAAACGCGCTTGCCGAAT
TTTGTTATTTACTCTGACGAAAAATTGTCACGATACACGAAAGTTTTTCACAGGCGGCGACTCATGGATAACTCGAGAGATCCTCTAGA
TTTAAGAAGGAGATATACATATG
>csgC
GAGGCAATTCCGACGTCTAAGAAACCATTATTATCATGACATTAACCTATAAAAATAGGCGTATCACGAGGCCCTTTCGTCTTCACCTC
GAGCAACGGTGCTGCAGTTGACCAGACTGCATCTAACTCCTCCGTCAACGTGACTCAGGTTGGCTTTGGTAACAACGCGACCGCTCATC
AGTACTAATACATCATTTGTATTACAGAAACAGGGCGCAAGCCCTGTTTTTTTTCGGGAGAAGAATATGAATACGTTATTACTCCTTGC
GGCACTTTCCAGTGGATCCTCTAGATTTAAGAAGGAGATATACATATG
>ycdM
GAGGCAATTCCGACGTCTAAGAAACCATTATTATCATGACATTAACCTATAAAAATAGGCGTATCACGAGGCCCTTTCGTCTTCACCTC
GATGGATCCGTGTCCAGTGCTGCGCTAAGAATCGCTTTTTTCTTCGCGCTTACTGCGCGCGAACGTTTACCCGTTGTTTTCACTGCGCC
TTGCGTCATGCGCTCTCCCCTCTTGTAGATTGGCAACAGGATAGCAAAACCCCCGCCCGGCCTCGACCTTCTGCACTCTCATCGCGCAC
AGATGCATGTTTTATGTGCAACTGTTTTGACCGTTTAGTCCACTTTTTACCAGATATTTTAGATAACGTAATGTTAACAGATTAATAAC
ATTAACTTTTTAAAAACTGGCATCCGCTTTGCAAACAAGCCAATACGCGGCTGAAAAGGATATGCAGGATGCAGCGCCCCGTCTGACTT
TCACTTTACGAGACGAAGAGAGGTTGATGATGAAAATTGGCGTATTCGTACCTATTGGCAACAACGGCTGCTCGAGAGATCCTCTAGAT
TTAAGAAGGAGATATACATATG
>ycdN
GAGGCAATTCCGACGTCTAAGAAACCATTATTATCATGACATTAACCTATAAAAATAGGCGTATCACGAGGCCCTTTCGTCTTCACCTC
GAGCAAAAACGCTTTGCCGAGGCCGATGCGCACTATCATTCGGCTCCGCCGTCACGGTTGCAGGAAAGCTAAGGGACTTAGCCTGCGGC
GGTTTTGTTTGGCTTCAGCAGCGGGTTGCGCTCCCTTAATGTGCCTCGCCATATAAATTGAATGGTGCAGGGAGCGCGCAGGGGGCGGC
CAATCGCCGCCGCCCCCTGCTGTCCCGGCCTTCGGGGAACGCTTCAGCGATTTTGACGCCACCAACACCCGAGCTGTTATTATGTTCCG
GGCAAAAAGTTAGATTTGATAATCGCGGATGGACGAAATTGCTTGATACACCCGCTTATCAGTTTTACATGGAAGCTCTGATGCATTGA
GTCTGGACAGTTTTGTCGGCTGGATACGGCGTTTACGCGGCATCCGGCAAGAACACATGGTTCTTTGCAAACAATCCCATCTTTCTACC
CTGGAATAATCGTTTATATCCCTTGGCATTACCTCTCTTTGTTTACATTCCAACATCATTTTATAAACATTCCGCTTGTGTTTTTCTTT
GCCGTAATGATAATCGCTATCACTGCGATTTACTTTTCTTTGCATAGATTGACTCAGAAAAACGTTTAAGGGTGGGTGGCATGTTTGTT
CCGTTTCTCATTATGTTGCGCGAAGGACTTGAAGCCGCGCTGATTGTCAGTTTGATTGCCAGCGGATCCCGGGATCCTCTAGATTTAAG
AAGGAGATATACATATG
>ycdO
GAGGCAATTCCGACGTCTAAGAAACCATTATTATCATGACATTAACCTATAAAAATAGGCGTATCACGAGGCCCTTTCGTCTTCACCTC
GAGGAAGTCGCCGTCTGGTTTATTTATCTCATCCCGGCGCTGGTGGCATTTGCTCTGCCACCACGCGCAGGGGCGACAGCGTCTCGCTC
CGCGTAACAAATACGACGCAAACTCTTGCTTAGTTACAACATACTTTAAAGGGATAGTCTCGTCATGACCATTAACTTCCGCCGTAACG
CATTGCAGTTGAGCGTGGCTGCGCTGTTTTCTTCTGCTTTTATGGCTAACGCCGCGGATCCTCTAGATTTAAGAAGGAGATATACATAT
G
>dsrA
GAGGCAATTCCGACGTCTAAGAAACCATTATTATCATGACATTAACCTATAAAAATAGGCGTATCACGAGGCCCTTTCGTCTTCACCTC
GATGGATCCAGCAGGGTGCCATCAAGATCGCTAAAAACCAGTAGTGGTTGTTGAATTGAAAACATAGTCGCGCAGTACTCCTCTTACCA
GGATGATTCTCAAAACCGCACAAATTATCAAAGATGATTTTTTCGGGAGCGATCCCGCTGACGCCAGAGCCACCAAGGGGTGACGTGCG
TCACATTTCTATTCATAAGTAGCGTTAATCATTCATATGGCGAATATTTTCTTGTCAGCGAAAAAAATTGCGGATAAGGTGATGAACAC
ATCAGATTTCCTGGTGTAACGAATTTTTTAAGTGCTTCTTGCTTAAGCAAGTTTCATCCCGACCCCCTCGAGAGATCCTCTAGATTTAA
GAAGGAGATATACATATG
>sseA
GAGGCAATTCCGACGTCTAAGAAACCATTATTATCATGACATTAACCTATAAAAATAGGCGTATCACGAGGCCCTTTCGTCTTCACCTC
GAGTTTTTCACCGCTGTTGGCGCGTTATCGTTGTTGTCGCACCCTGCCAGCGCCAGCATTAGCATGCAGGCGGCTACGCGTAACTTTTT
CATTTTTCATCCCTGCCACAATGGCCCGTTAGCAACGTCGAATAATTATTATGCGTGAGAATTTACGTTATGTAATTCAGTATCACGCC
CATACAATCCGAATTTTAGTGAGAATTGTGGCTCAGACAGCCAGGATCCTCTAGATTTAAGAAGGAGATATACATATG
>sseB
GAGGCAATTCCGACGTCTAAGAAACCATTATTATCATGACATTAACCTATAAAAATAGGCGTATCACGAGGCCCTTTCGTCTTCACCTC
GATGGATCCCGCTACGGGACTTGGTGTGCGCACGATAGCTAATCTGTTAACGGCGTAAACTAATATGCCGGATGCGGCGTAAACGCCTT
ATCCGGCCTACAAATTCTGCAAATTCAACAAATTGCGAATCCCTTGTAGGCCTGATAAGCGTAGCGCATCAGGCATTTGCGCCCCGAAC
GTTGAATGAATTGTGGTTCTTATGTCCGAAACAAAAAACGAACTTGAAGACCTGCTGGAAAAAGCAGCAACTGAACCGGCGCACCGCCC
GGCCTTTTTCCGTACTCTCTCGAGAGATCCTCTAGATTTAAGAAGGAGATATACATATG
>mntR
GAGGCAATTCCGACGTCTAAGAAACCATTATTATCATGACATTAACCTATAAAAATAGGCGTATCACGAGGCCCTTTCGTCTTCACCTC
GAGCGCCGCTATTTATCGGAAGGTTTATCTTGCTGCGGTTTGTTGTTGACCATATCGCACAACATAGAGAGCAGCATTAACCGTACTTT
AAAGGGAGAATGACTAAACACGCGCATACACCTCTTGAACTCATTCATAAGACCTCCTGACTTGCTAATCCCGTCGATCCTTGAGGGAT
GATTGCATTACATACAGATATAGCACAGGCTATATTATATAGCTATTGCTAAAACGTTAATTTTTTGTACCTTCGCAACTCTGGTTTAC
AATGTGCGCACGAAATGAGAACGCTATGTATGCGTCACCATAATGAGGAAGCACAATGAGTCGTCGCGCAGGTACGCCAACAGCAAAAA

AAGTGACGCAGTTAGTGAACGTGGAAGAGCACGTTGAAGGGTTCCGCCAGGTCAGAGAGGGATCCTCTAGATTTAAGAAGGAGATATAC
ATATG
>yqgA
GAGGCAATTCCGACGTCTAAGAAACCATTATTATCATGACATTAACCTATAAAAATAGGCGTATCACGAGGCCCTTTCGTCTTCACCTC
GAGAGTTCACTACTGGCGGCAATATTCATTGATTTCATAAGCGCAAACCCGTTTCGGGGAAGTAAGAATACCGGACAAGGTGGAAAACC
CTGCCCCATGAGATATGGGTCAAACTGGTCACCAGCTCCGACCGCCAGACATCAGTAAAAGCAGAAACGCTCTGATTTTACTGTTGTCC
TGCAGTGAGCGTGCGTTAACTTCACCGCATGAGCAGTAACATAGAAAGGGAAACGTTTCGCGCGAAACGGCGATAAGCGAGAGAATGTA
AGGAGATGGCGTGCATCGGGCAAACTCCGTACAGAGAGGAGAAAATTCGCGCAATCATGGCACCTTTCGCTAAAGCGTGCAAGCCAGGA
CTTTGCGAACAAACAAGCCATCGACGGCACCAAACGGTCATAATAAGAAAATCAAACAATACAGCTAACAGGAACTCTTGTGGTCATCG
GCCCTTTTATCAACGCTAGTGCAGTCTTACTGGGTGGCGTTCTCGGCGCACTGCTCAGCCAACGCTTACCGGAACGTATCCGCGTCTCC
AGGATCCTCTAGATTTAAGAAGGAGATATACATATG
>secA
GAGGCAATTCCGACGTCTAAGAAACCATTATTATCATGACATTAACCTATAAAAATAGGCGTATCACGAGGCCCTTTCGTCTTCACCTC
GAGAGCAAAATTCAGCACGCCCGTCTGGATAAGCCAGGCGCAAGGCATCCGTGCTGGCCCTCAACGCCTCACCTAACAACAATAAACCT
TTACTTCATTTTATTAACTCCGCAACGCGGGGCGTTTGAGATTTTATTATGCTAATCAAATTGTTAACTAAAGTTTTCGGTAGTCGTAA
CGATCGCACCCTGCGCCGGATGCGCAAAGTGGTCAAGGATCCTCTAGATTTAAGAAGGAGATATACATATG
>secB
GAGGCAATTCCGACGTCTAAGAAACCATTATTATCATGACATTAACCTATAAAAATAGGCGTATCACGAGGCCCTTTCGTCTTCACCTC
GATGGATCCGATGCACGTGGTGGACTGGATCCCCTGCTGAAATAACGTGTGAACGTTGGCATTACATTGCGCAGTATTTAAGGACAACA
CTTAAGGGTTTTCTACACATGTCAGAACAAAACAACACTGAAATGACTTTCCAGATCCAACGTATTTATACCAAGGATATCTCTTTCGA
AGCGCCGCTCGAGAGATCCTCTAGATTTAAGAAGGAGATATACATATG
>secE
GAGGCAATTCCGACGTCTAAGAAACCATTATTATCATGACATTAACCTATAAAAATAGGCGTATCACGAGGCCCTTTCGTCTTCACCTC
GAGGTCTGCGTTTCGCAATCCGTGAAGGCGGCCGTACCGTTGGCGCGGGCGTTGTAGCAAAAGTTCTGAGCTAATTGCCGATAACATTT
GACGCAATGCGCACTAAAAGGGCATCATTTGATGCCCTTTTTGCACGCTTTCGTACCAGAACCTGGCTCATCAGTGATTTTCTTTGTCA
TAATCATTGCTGAGACAGGCTCTGTTGAGGGCGTATAATCCGAAAAGCTAATACGCGTTTCGATTTGGTTTGCCTCGCGATCGCGGGGT
GAAAATGTTTGTAGAAAACTTCTGACAGGTTGGTTTATGAGTGCGAATACCGAAGCTCAAGGAAGCGGGCGCGGCCTGGAAGCGATGAA
GTGGGTCGTTGTGGTGGCATTGCTCCTGGTGGCGATTGTCGGATCCTCTAGATTTAAGAAGGAGATATACATATG
>secM
GAGGCAATTCCGACGTCTAAGAAACCATTATTATCATGACATTAACCTATAAAAATAGGCGTATCACGAGGCCCTTTCGTCTTCACCTC
GAGTCCTGGCGAAACAGGAAGCCTGGGAATATGTGACCTTCCAGGACGACGCAGAACTGCCGTTGGCCTTCAAAGCGCCTTCAGCTGTA
CTGGCATAACGACATTTATACTGTCGTATAAAATTCGACTGGCAAATCTGGCACTCTCTCCGGCCAGGTGAACCAGTCGTTTTTTTTTG
AATTTTATAAGAGCTATAAAAAACGGTGCGAACGCTGTTTTCTTAAGCACTTTTCCGCACAACTTATCTTCATTCGTGCTGTGGACTGC
AGGCTTTAATGATAAGATTTGTGCGCTAAATACGTTTGAATATGATCGGGATGGCAATAACGTGGATCCTCTAGATTTAAGAAGGAGAT
ATACATATG
>ymfI
GAGGCAATTCCGACGTCTAAGAAACCATTATTATCATGACATTAACCTATAAAAATAGGCGTATCACGAGGCCCTTTCGTCTTCACCTC
GAGGAAGGTTGCACACCAGGCGGGTGGGTATCCACAGAAGGTCGATTGTCAGCCTCAACCTTAACCCGCGTGCGCTGAGCCGCATTCAC
CACAACGCTAAGGATTCTCTCTGGTTGAAAATACTTAGCTGTTATGTGCCTGTCTTTTCACCACTTCAGGCTCGGTGGTATGCTGGAGT
TCTCACACAGCCAGCAAGCAAGGAAACTTAATGAACCAGTTTTATGTTCACGTTCGTCTATTTGAAGACACAGCCGAACAGACCAAGGA
TCCTCTAGATTTAAGAAGGAGATATACATATG
>ilvI
GAGGCAATTCCGACGTCTAAGAAACCATTATTATCATGACATTAACCTATAAAAATAGGCGTATCACGAGGCCCTTTCGTCTTCACCTC
GAGCTCTCCTGGCATGAAGCTGCCGGGCGCGATAAAGGCCATCAGTGGATGGAAGAGCAATTAGTCTCAATTTGCAAACGCTAACTGAT
TGCAGAATAGGTCAGACATGAATGTCTGGTTTATTCTGCATTTTTTATTGAATGTAGAATTTTATTCTGAATGTGTGGGCTCTCTATTT
TAGGATTAATTAAAAAAATAGAGAAATTGCTGTAAGTTGTGGGATTCAGCCGATTTATTATCAATTTAATCCTCTGTAATGGAGGATTT
TATCGTTTCTTTTCACCTTTCCTCCTGTTTATTCTTATTACCCCGTGTTTATGTCTCTGGCTGCCAATTGCTTAAGCAAGATCGGACGG
TTAATGTGTTTTACACATTTTTTCCGTCAAACAGTGAGGCAGGCCAGGATCCTCTAGATTTAAGAAGGAGATATACATATG
>coaE
GAGGCAATTCCGACGTCTAAGAAACCATTATTATCATGACATTAACCTATAAAAATAGGCGTATCACGAGGCCCTTTCGTCTTCACCTC
GATGGATCCGTTCAACATCGGAACGGCTTTTAAGAGTGGAGCGTTTAGGGCGGATGAGAACGTCTTTAAAACCTAACTTCAGATCTTCT
TCAATACGCATGTGCGGATTCCTGGGGTTAATGGCTATAAGGGTAAGAGCGCAACTCCAGTGACGCTATCATACGCACTAATCAATGCG
GCGCAAGACTGCGAAATTGTCTTTTTTTACGCTACAATCCCATAAATTTACCTGGTAAATAGTGAGCGAACGTTTCATCAATTTCGCTT
TCGTGCACTCATTTTTAACGTTTTGATAATCATGATGAATTCCCGGGAATATAAATTATGAGGTATATAGTTGCCTTAACGGGAGGCAT
TGGCAGTGGCAAGAGTACCGTTGCCAATGCGTTTGCTCTCGAGAGATCCTCTAGATTTAAGAAGGAGATATACATATG
>ilvD
GAGGCAATTCCGACGTCTAAGAAACCATTATTATCATGACATTAACCTATAAAAATAGGCGTATCACGAGGCCCTTTCGTCTTCACCTC
GAGAAACGCATTCAGCAAGCCTTCTTCGGCCTCTTCACTGGCGAAACCGAAGATAAATGGGGCTGGTTAGATCAAGTTAATCAATAAAT
ACAAAAAATGGGACGGCACGCACCGTCCCATTTACGAGACAGACACTGGGAGTAAATAAAGTATGCCTAAGTACCGTTCCGCCACCACC
ACTCATGGTCGTAATATGGCGGGTGCTCGTGCGCTGTGGCGCGCCACCGGAATGACCGACGCCGATTTCGGTAAGCCGATTATCGCGGG
GATCCTCTAGATTTAAGAAGGAGATATACATATG
>ymfE

GAGGCAATTCCGACGTCTAAGAAACCATTATTATCATGACATTAACCTATAAAAATAGGCGTATCACGAGGCCCTTTCGTCTTCACCTC
GATGGATCCGGCACACTGGGAACGATTAAGGTAAGCATGTCGTAAGACACCATCCGTGCCCACTGGACACTCTGTCATCATTTCATCAT
TTCATCATTTCATCATTTCATCATTTCATCATTTCATTATTTCATTATTTCATTTCTATCAATTTCAGATTCTCACCCAGCGTCAAATT
TGTAGCGCTACTCCTCCTCGATACGTGCTACTGATCACAATATTGTCTGTCAGCGAAGCTAGGTCAGCCTGAACAGATACAAAAAATTG
TAACTAACCTACTGAAAAGTATATAATGACACTTAATTAAATATTACCATTACACTATCCTAAAGCGTGACTCAGGTGAATAATATGTT
CGAACCCCCCAAAAATTATAATGAAATGTTGCCTAAACTTCATAAAGCAACTTTCTTAAATACGCTAATATATTGCATACTTCTAGTTA
TTTACGAATACATCCCTTTAATAACATTACCAACCAAGTATGTCCCACCTATTAAAGATCATGAGAGCTTTATTAATTGGGCACTATCT
TTTGGTATATTACCTTGTGCTTTTGCGATTTTTGCATATTTAATTAGCGGTGCGTTAGACCTACATAACAATGCAGCCAAACTACTTCG
GGTGCGATATCTTTGGCTCGAGAGATCCTCTAGATTTAAGAAGGAGATATACATATG
>ilvC
GAGGCAATTCCGACGTCTAAGAAACCATTATTATCATGACATTAACCTATAAAAATAGGCGTATCACGAGGCCCTTTCGTCTTCACCTC
GAGGTGGATGGGCTAACGTGCATCGCCCGCGCGCTGCGGCCAAAATGGCGGCTTTCCGCCAGATGCAGGAAGGTTTTCAGATCGCGTAA
ATCCACAGGGACAGCCCTCGATGTTGACGTTGCAAAAATTGCAATGTGACGTTGTGAATATATCAATTTCCGCAATAAATTTCCTGTCA
TATAGTGAATTCAATCTCGCAAACGCGAACCGAACAATAAGAAGCACAACATCACGAGGAATCACCATGGCTAACTACTTCAATACACT
GAATCTGCGCCAGCAGCTGGCACAGCTGGGCAAATGTCGCTTTATGGGCCGGATCCTCTAGATTTAAGAAGGAGATATACATATG
>hfq
GAGGCAATTCCGACGTCTAAGAAACCATTATTATCATGACATTAACCTATAAAAATAGGCGTATCACGAGGCCCTTTCGTCTTCACCTC
GAGGGAAGGGGTTCACTGGCTTGACAGTGAAAAACCAGAACAGGCGCGTGACGAAGTATTACAGGTTGTTGGTGCTATCGCAGGCTGAA
TGTGTACAATTGAGACGTATCGTGCGCAATTTTTTCAGAATCGAAAGGTTCAAAGTACAAATAAGCATATAAGGAAAAGAGAGAATGGC
TAAGGGGCAATCTTTACAAGATCCGTTCCTGAACGCGGATCCTCTAGATTTAAGAAGGAGATATACATATG
>ego
GAGGCAATTCCGACGTCTAAGAAACCATTATTATCATGACATTAACCTATAAAAATAGGCGTATCACGAGGCCCTTTCGTCTTCACCTC
GAGGGTCAGCCCGTCGTGATAGTAAAACCACGCGATCCGCGCGACCTGTTCTTCTTCACACATTCCCTGTTCTGAAATTGCCGAATCGT
TGATTGTCATAATTCATTCTTCACTTTGAACATATTTAAATCTTTAATGCAATTGTTCAGTTCTTGCTCATTTATATCTGTGATGGCAA
CCACAGTTTGACTCTACGAGCATGAACAAACGCAACCGTGAAAATCAAAATAGCATAAATTGTGATCTATTCGTCGGAAATATGTGCAA
TGTCCACCTAAGGTTATGAACAAATTAAAAGCAGAAATACATTTGTTCAAAACTCACCTGCAAAACTGAACGGGGGAAATATGCAAACG
AGTGATACCCGCGCGTTACCGCTACTTTGCGCCCGCTCGGTTTATAAACAGTATTCAGGGGTCAATGTCCTGAGGATCCTCTAGATTTA
AGAAGGAGATATACATATG
>yedX
GAGGCAATTCCGACGTCTAAGAAACCATTATTATCATGACATTAACCTATAAAAATAGGCGTATCACGAGGCCCTTTCGTCTTCACCTC
GAGCCCATTCCTGGGTCCTTTGATTATCTTCAATAAGTAGAATCTTCATGAAAATAATATGCCATAAATCATATGTTATAGAAACAGCC
TGGTTCATTACAAAATTGTAATGCTGCTGTAAGGTTACCCTGGACGCTTTTTCGCTATCCTCAAAACTCATTCACATGACAAGGATATA
AACATGTTAAAGCGTTATTTAGTACTCTCGGTAGCAACGGCAGCATTTTCATTACCTTCTTTGGTTAATGCCGCACAACAAAACAGGAT
CCTCTAGATTTAAGAAGGAGATATACATATG
>yedY
GAGGCAATTCCGACGTCTAAGAAACCATTATTATCATGACATTAACCTATAAAAATAGGCGTATCACGAGGCCCTTTCGTCTTCACCTC
GAGCTGAGATCCCCGTTGAATTTCATATTAATAAAGTGAACGAGCATTATCATGTGCCTTTATTACTTAGCCAATATGGGTATTCAACC
TATCGTGGCAGTTAATGCATAAATATGTAAAAGCAGATCTCTGCTACTGTTAGGGAGGACACTCCCTGACAGATTAACAGTAAACGGCT
CTTGCTGGCTAACGACAAAAAAGTGTGATGGCTTATGAAAAAGAATCAATTTTTAAAAGAATCAGATGTTACGGCCGAGTCGGTATTCT
TTATGAAGCGTCGGCAGGTGGATCCTCTAGATTTAAGAAGGAGATATACATATG
>yedW
GAGGCAATTCCGACGTCTAAGAAACCATTATTATCATGACATTAACCTATAAAAATAGGCGTATCACGAGGCCCTTTCGTCTTCACCTC
GATGGATCCGAAAATGCTGCCGTTGCTACCGAGAGTACTAAATAACGCTTTAACATGTTTATATCCTTGTCATGTGAATGAGTTTTGAG
GATAGCGAAAAAGCGTCCAGGGTAACCTTACAGCAGCATTACAATTTTGTAATGAACCAGGCTGTTTCTATAACATATGATTTATGGCA
TATTATTTTCATGAAGATTCTACTTATTGAAGATAATCAAAGGACCCAGGAATGGGCTCGAGAGATCCTCTAGATTTAAGAAGGAGATA
TACATATG
>ybeT
GAGGCAATTCCGACGTCTAAGAAACCATTATTATCATGACATTAACCTATAAAAATAGGCGTATCACGAGGCCCTTTCGTCTTCACCTC
GATGGATCCGTATTTCCATGGGCGCAGAAAGCGCAAAAAGAAGATATTGTTCTTCCTTATTCATTTGTTAATAATCCCTTAAGTGCATT
AACTGACTACTTTTTTATTATTGATGAGTATAAACTTGACGCTTTCAAAATAAAAAGAAAATCGAAGCATTCACACATGAATAAAAAAT
TAATGTATATATTCGCAATTTTTATAGTTGCAGCAATTACCTGTATTAGCCAACCCAAGAAAACGACTCGAGAGATCCTCTAGATTTAA
GAAGGAGATATACATATG
>yedP
GAGGCAATTCCGACGTCTAAGAAACCATTATTATCATGACATTAACCTATAAAAATAGGCGTATCACGAGGCCCTTTCGTCTTCACCTC
GAGGGGGTCGGGATGAAACTTGCTTAAGCAAGAAGCACTTAAAAAATTCGTTACACCAGGAAATCTGATGTGTTCATCACCTTATCCGC
AATTTTTTTCGCTGACAAGAAAATATTCGCCATATGAATGATTAACGCTACTTATGAATAGAAATGTGACGCACGTCACCCCTTGGTGG
CTCTGGCGTCAGCGGGATCGCTCCCGAAAAAATCATCTTTGATAATTTGTGCGGTTTTGAGAATCATCCTGGTAAGAGGAGTACTGCGC
GACTATGTTTTCAATTCAACAACCACTACTGGTTTTTAGCGATCTTGATGGCACCCTGCTGGACAGTCATAGTTATGACTGGCAACCGG
CGGATCCTCTAGATTTAAGAAGGAGATATACATATG
>yedQ
GAGGCAATTCCGACGTCTAAGAAACCATTATTATCATGACATTAACCTATAAAAATAGGCGTATCACGAGGCCCTTTCGTCTTCACCTC
GATGGATCCCAAGCGGGAAGCTTTTCATGAAACCGAGCTTGTTCCGGGGGAGGGGAGTCGTTCTGCGGAAGAAGTGTGAAAGGGTATCA
TTAACATCAGGCCGGATCACATCCGGCCTGGTGTGCTAATGCCTGCAATGAATGATAGTCGCGTCTTATCATGCCACCATTATAAAATT

```
ATCGCTGTTTTGCGGTACGCTATCGCCAGGTTATGCCAGAATCATAAAAAAGCAGGTTGGGAGTCGTCAGGGTGCAGCACGAGACAAAA
ATGGAAAACCAGAGCTGGTTGAAAAAACTCGCACGCCGCCTGGGGCCTGGTCATGTCTCGAGAGATCCTCTAGATTTAAGAAGGAGATA
TACATATG
>allD
GAGGCAATTCCGACGTCTAAGAAACCATTATTATCATGACATTAACCTATAAAAATAGGCGTATCACGAGGCCCTTTCGTCTTCACCTC
GATAGATCTCCGAATCCTGAAAACACCCTTTTTTAATAAAGGCGTGGATCATAACAACTCCTTAAAAATTGCACCCATGGCAATTGCAC
GAGCAGTGTTAATGTGGCGATTACGGTATCGATGACAACGAGTAAATAAGGTGATGAATATCACACAGGAATTATTAACAAAAGAATTC
AAATAACCACCAGGCTATTACTCATTATTTTTTAATAAGCGTTGTTGTTTTAAAAATCAGGTTAAAAGTGAGCGGATTATCAAATAAAT
TCTAACTATATTTTTTTGCCTGTCTGGATCACATAATCCAGATATTTTCCCTGTATGTTAATTGCAGTCATGCTTCACACCGTCGTTAA
AAAGGAAGACAGATGAAAATCAGTCGGGAAACACTCCACCAGCTAATTGAGAATAAACTCTGCCAGGCTGGGTTAAAACGTGAGCACGC
TGCCTCGAGAGATCCTCTAGATTTAAGAAGGAGATATACATATG
>allA
GAGGCAATTCCGACGTCTAAGAAACCATTATTATCATGACATTAACCTATAAAAATAGGCGTATCACGAGGCCCTTTCGTCTTCACCTC
GAGATCGTTCTGCCGCTTTTGAAAAACTTCCTGTTTCAGCAACCGCAATGAAAGTCCGCAAGGTTTCTGGATCGAACATCTTCAGGTAT
CCCCTTTTAAATCCGCAAGTTGCGTGATTTTCTTATCCTCTGATTTATCAGTATTTTTACATGATAACCCTGTTCAATTTGTGGACTAA
ATCTAGTTTTGGAAAAATATTCCAACTTTTGTATTGATGTTGTTCTCTTAAGGTTTTAGATTGCCTGTTATTGAAACCAAGCTGACCGG
TCGGCGGTGGTTGAACGGAATTATGTTACAAGGACAAAAAGATGAAACTTCAGGTATTACCGTTAAGTCAGGAAGCCTTTAGTGCTTAT
GGCGACGTAATCGAAACGCAGGATCCTCTAGATTTAAGAAGGAGATATACATATG
>poxB
GAGGCAATTCCGACGTCTAAGAAACCATTATTATCATGACATTAACCTATAAAAATAGGCGTATCACGAGGCCCTTTCGTCTTCACCTC
GATGGATCCCACAATGACGCTGACCGACGCCGAAATCGCTGAAGGTTACGTACTGGCCTGCTCCTGCCATCCGCAGGGGGATTTGGTTC
TCGCATAATCGCCTTATGCCCGATGATATTCCTTTCATCGGGCTATTTAACCGTTAGTGCCTCCTTTCTCTCCCATCCCTTCCCCCTCC
GTCAGATGAACTAAACTTGTTACCGTTATCACATTCAGGAGATGGAGAACCATGAAACAAACGGTTGCAGCTTATATCGCCAAAACACT
CGAATCGGCAGGGGTGAAACGCATCTGGGGAGTCACTCGAGAGATCCTCTAGATTTAAGAAGGAGATATACATATG
>emrR
GAGGCAATTCCGACGTCTAAGAAACCATTATTATCATGACATTAACCTATAAAAATAGGCGTATCACGAGGCCCTTTCGTCTTCACCTC
GAGCGCAGCATTATCATCCCAACACTGCTTAGTGCGCTGGCCTATGGGCTCGCCTGGAAAGTGATGGCGATTATATAACCCACAAGAAT
CATTTTTCTAAAACAATACATTTACTTTATTTGTCACTGTCGTTACTATATCGGCTGAAATTAATGAGGTCATACCCAAATGGATAGTT
CGTTTACGCCCATTGAACAAATGCTAAAATTTCGCGCCAGCCGCCACGAAGATTTTCCTTGGATCCTCTAGATTTAAGAAGGAGATATA
CATATG
>ugpA
GAGGCAATTCCGACGTCTAAGAAACCATTATTATCATGACATTAACCTATAAAAATAGGCGTATCACGAGGCCCTTTCGTCTTCACCTC
GATGGATCCACTGGATACCGCCGTTGAGCGTGGAAATCAGTTGCTGCGCCGCTTTGAGAAATCGACGAAGTCTTAATCAGTGTAATGTC
GGATGCGTTTCGCTTATCTGACCTGGCATCGCGTGTAGGCCGGATAAGCGAAGCGCATCCGGCACAGTTCAGGAATTAACCGTAATGTC
ATCATCCCGTCCGGTGTTCCGCTCGCGCTGGCTGCCTTATCTGCTGGTCGCGCCGCAGCTCATCATCACCGTTATCTTTTTTATCTGGC
CTGCGGCTCGAGAGATCCTCTAGATTTAAGAAGGAGATATACATATG
>ugpB
GAGGCAATTCCGACGTCTAAGAAACCATTATTATCATGACATTAACCTATAAAAATAGGCGTATCACGAGGCCCTTTCGTCTTCACCTC
GATGGATCCAAGCGGTGAGAAGTGCGTATTTAGGCGGGTAATAACACGTTGATTGATAGGGAGTCAAAAGACTCCTTTGAGACAGGTGA
CAAATGTAAAATTGCCTGATGCGCTGCGCTTATCAGGCCTACTGGGTGAGTGGCAATATGTTGAATTTGCACGATCTTGTAGGCCTGAT
AAGGCGTTTACGCCGCATCCGGCATGAAACGATGAGCAATCTGTAGAGTTTGATTCAGACCTTCTATATTTTCCCGCTTATCCGTGCCC
CATCTCCCATTTTCCCTCACCCACGCCGTCACCGCCTTGTCATCTTTCTGACACCTTACTATCTTACAAATGTAACAAAAAAGTTATTT
TTCTGTAATTCGAGCATGTCATGTTACCCCGCGAGCATAAAACGCGTGAATTCGCGCATTCGGTACAACAAGAGAGATAAACGATGAAA
CCGTTACATTATACAGCTTCAGCACTGGCGCTCGGACTGGCGTTAATGGGGAATGCACAGGCAGTGACGACTCGAGAGATCCTCTAGAT
TTAAGAAGGAGATATACATATG
>allS
GAGGCAATTCCGACGTCTAAGAAACCATTATTATCATGACATTAACCTATAAAAATAGGCGTATCACGAGGCCCTTTCGTCTTCACCTC
GATGGATCCCGTTTCGATTACGTCGCCATAAGCACTAAAGGCTTCCTGACTTAACGGTAATACCTGAAGTTTCATCTTTTTGTCCTTGT
AACATAATTCCGTTCAACCACCGCCGACCGGTCAGCTTGGTTTCAATAACAGGCAATCTAAAACCTTAAGAGAACAACATCAATACAAA
AGTTGGAATATTTTTCCAAAACTAGATTTAGTCCACAAATTGAACAGGGTTATCATGTAAAAATACTGATAAATCAGAGGATAAGAAAA
TCACGCAACTTGCGGATTTAAAAGGGGATACCTGAAGATGTTCGATCCAGAAACCTTGCGGACTTTCATTGCGGTTGCTGAAACAGGAA
GTTTTTCAAAAGCGGCAGAACGATCTCGAGAGATCCTCTAGATTTAAGAAGGAGATATACATATG
>allR
GAGGCAATTCCGACGTCTAAGAAACCATTATTATCATGACATTAACCTATAAAAATAGGCGTATCACGAGGCCCTTTCGTCTTCACCTC
GAGAGCGCGTCACCGATTTTCTGACCATCGATCGCGGCGGCAGTGACAACTGTGATGTTGAAAGTATTCCTGAACAGGAACTCTGTTTT
GCGTGACGCCTGCAACCGACTTGCATAAGATAAACTAATTGTTCATTGTTTATGCTCACTTGTAGGTCGGAGTTAACGTAGGTATGACG
GAAGTTAGACGGCGCGGCAGGCCAGGACAGGCGGAGCCTGTGGCACAGAAGGGCGCACAGGCGTTAGAGCGGGGAATTGGATCCTCTAG
ATTTAAGAAGGAGATATACATATG
>emrA
GAGGCAATTCCGACGTCTAAGAAACCATTATTATCATGACATTAACCTATAAAAATAGGCGTATCACGAGGCCCTTTCGTCTTCACCTC
GAGCACCCGCAAATTGCTCTCCCGTCTCGACCAGATGGAACAAGACGGTGTGGTTCTCGAAGCGATGAGCTAACGCGTCATCTCGCTCA
AAAATCCAGATTTATAAAAGAAAAAATGACTGGCCAGCATCGCAACATGCTGGCCTTTTTGGCAAGCAGGTCGGCTCAGCCGATGAGTT
```

```
AAGAAGATCGTGGAGAACAATATGAGCGCAAATGCGGAGACTCAAACCCCGCAGCAACCGGTAAAGAAGAGCGGCAAACGTAAGCGTCT
GGATCCTCTAGATTTAAGAAGGAGATATACATATG
>emrE
GAGGCAATTCCGACGTCTAAGAAACCATTATTATCATGACATTAACCTATAAAAATAGGCGTATCACGAGGCCCTTTCGTCTTCACCTC
GAGGAGTGTCGCCAGAGTGCCGCGATGAAACGGGTATTGAGGGTATATAAAAGAACATCAATGGGAACACAATGATGAAACAGGTGAGT
TGAGTTCAAACTGTAGTACAATTCTCTCCAGTTTGAACAGGAAAGAATATGCTATGAACCCTTATATTTATCTTGGTGGTGCAATACTT
GCAGAGGTCATTGGTACAACCTTAATGAAGTTTTCAGAAGGTTTTACACGGTTATGGCCATCTGGGATCCTCTAGATTTAAGAAGGAGA
TATACATATG
>pcnB
GAGGCAATTCCGACGTCTAAGAAACCATTATTATCATGACATTAACCTATAAAAATAGGCGTATCACGAGGCCCTTTCGTCTTCACCTC
GATGGATCCCAGCCGTCAAAAACTGGCGGCTTACCGCCGTGCCCGAGTCGGCAATTGTAAATTCAACATTCTCAAATGCGTCATGCTGA
GCTATGATTAGCCGCTATTTTTTTGTCCTGAATGATGTTTGACACTACCGAGGTGTACTATTTTTACCCGAGTCGCTAATTTTTGCCGC
AAGGTGCTAAGCCGCGAGGAAAGCGAGGCTGAACAGGCAGTCGCCCGTCCACAGGTGACGGTGATCCCGCGTGAGCAGCATGCTATTCT
CGAGAGATCCTCTAGATTTAAGAAGGAGATATACATATG
>yibL
GAGGCAATTCCGACGTCTAAGAAACCATTATTATCATGACATTAACCTATAAAAATAGGCGTATCACGAGGCCCTTTCGTCTTCACCTC
GAGCAACAGGCTGTGCGATCAACAATGGTCCTTTCACTGACTGAGCTGATTTCCAAAATCAGCTTAAAAAAAGCCTTTCAAAAGTAAGC
AACGTCTGCTTACTGCCCCTCTACCTGCTTCGGCCGATAAAGCCGACGATAATACTCCAGACGTTGTAGATATAACGGCACGCTTTCAT
CGGGTATTCCGGACGGAATCGCGTTACGGGGAGGAAGTTTTTTCAGATACTCCCGGAACGCCTGGCTTGATGCCATGAAATCTACGGCT
TTATCGATAAGAAGCGGAACGTTTTCACCTAATTTGCCCATGATGATATCTCCGTATTACCCCGCGCCGGGAATGCGCGGCCGCCAATT
TTAGTTTAGGTTCCCGGTAAAGATGACATTAAGGAAAACGTGCTGAATCCTCAATAATCATCCGCTGCTGTTTATTATCTTTTTTCTAT
AGATTTCTTAATTAATCAACGAATTGCGCAGTTTTTGAGCATGTTAATGATTACGTTACAGAACTATTACAATGTCATTAGTTGTGAAC
AAAGCACCTGGTCGCGCATACTAGGGGCTATAAATTTATCTTTATCAGAAGCCATCACATGAAAGAAGTCGAAAAAAACGAAATCAAAC
GCCTCAGCGGGATCCTCTAGATTTAAGAAGGAGATATACATATG
>yibF
GAGGCAATTCCGACGTCTAAGAAACCATTATTATCATGACATTAACCTATAAAAATAGGCGTATCACGAGGCCCTTTCGTCTTCACCTC
GATGGATCCGCCCTTTTTCCCATCACCAGGCCATACTGATATTTTTCCTCGCACAAAGGACTCGGCATGAAACTCGTCGGTAGCTACAC
CAGCCCGTTTGTACGCAAACTTTCTATTCTGTTGTTAGAAAAGGGCATAACTTTCGAATTTATTAATGAACTGCCCTATAACGCGGACA
ACGGCGTGGCGCAATTTAACCCGTTAGGAAAAGTGCCGGTGCTGGTGACCGAAGAGGGCGAATGCTGGTTTGATTCGCCGATCATCGCT
GAATATATTGAATTAATGAATGTCGCTCCGGCGATGTTGCCGCGCGATCCGCTGGAGTCGTTGCGGGTGCGCAAAATTGAGGCACTGGC
GGTCGACAGATCCTCTAGATTTAAGAAGGAGATATACATATG
>panB
GAGGCAATTCCGACGTCTAAGAAACCATTATTATCATGACATTAACCTATAAAAATAGGCGTATCACGAGGCCCTTTCGTCTTCACCTC
GATGGATCCACCCGGCGAATTTAAAGCCACCAGTACTTTCCAGGTAACCTATCCCTGATAACGTAGCAGCACTGTCACCTTCTGGCAGT
GCTGCTACGGATAAATCATTGCCCCCACCAATGTGACTGATAAAATGCCAGACAGCACAACATCAATTTATCAGGATACGTTATGAAAC
CGACCACCATCTCCTTACTGCAGAAGTACAAACAGGAAAAAAAACGTTTCGCGACCATCACCGCTTATCTCGAGAGATCCTCTAGATTT
AAGAAGGAGATATACATATG
>panE
GAGGCAATTCCGACGTCTAAGAAACCATTATTATCATGACATTAACCTATAAAAATAGGCGTATCACGAGGCCCTTTCGTCTTCACCTC
GATGGATCCGACCGCGTTACGTGCTTCCTGAAGATCAACTTCAGAGACAATATCGAAAGATGGCATCTTTTCCTCTCCCTTCATTTTTG
ATGCGAAGCATAATACCCGCAAAGTTAAATAACCGACAGTCCTGCACCAGGACGTATCCCTTAATTGTGCATAGTCGCAACAGCATGCA
CAATTAAGGGATAGCCTTATTGAACACCCGGAGTGGTTGCGGGTGAGGAGGAACAATGAAAATTACCGTATTGGGATGCGGTGCCTTAG
GGCAATTATGGCTTACAGCACTTTGCAAACAGGGTCATGAAGTTCAGGGCTGGCTGCCTCGAGAGATCCTCTAGATTTAAGAAGGAGAT
ATACATATG
>panD
GAGGCAATTCCGACGTCTAAGAAACCATTATTATCATGACATTAACCTATAAAAATAGGCGTATCACGAGGCCCTTTCGTCTTCACCTC
GATGGATCCCATGCGGTGTGGTACTCGGTGCATCCATCTTCCCTGTCTTCCGTGTGGCTTTGCGTGAAGGCAGTATCCAGTCAAGGCGC
GCTATCGTCAGCCACCTCTTTATTGTTTACCGAGCAGCGTTCAGAGAAATATCAATTCGTTACAGGCGATACATGGCACGCTTCGGCGC
GGTACGCAAACTGCTGACGCAAATGCCGTCAGATGAGTAGACACTAAACAAAAATCGGGCAATACTGCGTGAGAATTTCCTCAAAGCAG
GCATCGCCTGCTTCGTTAACGACAGGGTAGAAAGGTAGAAGTTATGATTCGCACGATGCTGCAGGGCAAACTCCACCGCGTGAAAGTGA
CTCATGCGGACCTGCACTATGAAGGTTCTTGCGCCATTGCTCGAGAGATCCTCTAGATTTAAGAAGGAGATATACATATG
>yejG
GAGGCAATTCCGACGTCTAAGAAACCATTATTATCATGACATTAACCTATAAAAATAGGCGTATCACGAGGCCCTTTCGTCTTCACCTC
GATGGATCCCGCAACCAGCTCCATTCTCTTCTGTCTGTACGCCAGTCGGCCGAAAAAACGGTGATCTATTGCACAACGAGGAAGCTAAA
AGGCTTCCTTTGTTGATGCATGTCAACCACAAATCTATCATTCCCCCGATATATGTTTATTTTATGTAAAATCAATTTATGTAAAAAGT
CACATCATTGTAGTTAAAAAGGTTGAGTTAGATCGCAGAAACGGGTACATATAGCCCCGCAAACGTGACCACGCCCGCAGATATTACTT
AAATCAGAGCCATAGAGGCCACGCAGGCGAGGCATCAATCTTTACGATCTGTATAAAGACGGATTGTTGATGATGTGTTAAAATTGATG
TAAACAAATTGTGAAGTGAATGTGCTTCCGGGGAAAATAAGTGACTTCATTACAACTCTCAATCGTCCATCGACTGCCGCAGAACTATC
GCTGGTCTGCTGGTTTCGCAGGTTCTCGAGAGATCCTCTAGATTTAAGAAGGAGATATACATATG
>yejA
GAGGCAATTCCGACGTCTAAGAAACCATTATTATCATGACATTAACCTATAAAAATAGGCGTATCACGAGGCCCTTTCGTCTTCACCTC
GAGGATTAGCCGCCCGTTACCGCTGGACGATTTTGTTCGCTGGCTAAAGAAACCGTATACGCCGCAGTGGTAAGGTGTGCTTACGTCCC
TTATTATTCATAGTGAAAGCATGCCGGATTGCGGCTAATGATGAGTAAAAGGAAATCCGTTGCAGATGATTGTGCGCATACTGCTGCTG
```

TTTATCGCTCTGTTCACCTTTGGTGTGCAGGCGCAGGCTATCAAGGAAAGCTATGCCTTTGCCGTGCTGGATCCTCTAGATTTAAGAAG
GAGATATACATATG
>yejL
GAGGCAATTCCGACGTCTAAGAAACCATTATTATCATGACATTAACCTATAAAAATAGGCGTATCACGAGGCCCTTTCGTCTTCACCTC
GAGCAATGAATCGCGCAACACCAGCTCAAGGTTCTGCTCATCACGCTTGATAAGCTGGTGCAGAGCAATCTGGTTGATATCCAGACTCA
TGATAAACTCTCCTTTAAGACCGGGCGGTATTCAACCACCGCCGGTTAAGCGACGCAATAAAAGATGGCGGTCATTTTACAGAAGGCAG
CGTAACGAGAATGATTTAACGCATAAAAAAGCAGAAAAAAAAACCGTTGCTACGGTAATATGTTGCCCTTTCATCAACTAACTGATTTCG
ATTTATGCCACAAATTTCCCGCTACAGTGATGAACAGGTTGAACAATTGCTCGCAGAGCTGCTCAACGTACTGGAAAAACATAAGGCTC
CGACCGACCGGATCCTCTAGATTTAAGAAGGAGATATACATATG
>yejO
GAGGCAATTCCGACGTCTAAGAAACCATTATTATCATGACATTAACCTATAAAAATAGGCGTATCACGAGGCCCTTTCGTCTTCACCTC
GATGGATCCATTTCCCCGCTTATTCGCACCTTCCTTATATTCACCCATAGCATTGGCATCAACTGTTGAGTATGGTGAGACAGTTGATG
GTGTTGTCCTGGAAAAAGATATCCAGCTGGTTTATGGGACCGCCAATAATACGAAAATCAATCCTGGCGCTCGAGAGATCCTCTAGATT
TAAGAAGGAGATATACATATG
>yejH
GAGGCAATTCCGACGTCTAAGAAACCATTATTATCATGACATTAACCTATAAAAATAGGCGTATCACGAGGCCCTTTCGTCTTCACCTC
GAGTCGACGGTGACACGATTGCCGCGGATTTCACGCCCGGCAATAGCACGGCTAACGCCGAGTTGCTGTGCGATAAATTTATCAAGTCG
CATGTGTGTGATTTTGCCTGTAAAAACGGAGGTCGGGCATTGACCCGAAAATCTGAACTGTTGTCTGCCCAGTATAGCGGTCTAATAAC
ATCCCTCAAGGGAAAAAGATCCATGGCATACTATTAGCAGAATAATCTACCTACGCGAGACCATGATTTTTACACTTCGCCCATATCAG
CAAGAAGCCGTGGATGCCACGCTCAACCATTTTCGTCGTCATAAAACCCCTGCCGTTATCGTGCTGCCCACCGGATCCTCTAGATTTAA
GAAGGAGATATACATATG
>rbsD
GAGGCAATTCCGACGTCTAAGAAACCATTATTATCATGACATTAACCTATAAAAATAGGCGTATCACGAGGCCCTTTCGTCTTCACCTC
GAGTGCGCTGATGCATAACCTTAAACATAACAAGGTATTGCATGAGCGGGTGATTCTGTTAACTCTGCGCACCGAAGACGCTCCATATG
TCCATAACGTCCGTCGGGTACAGATTGAACAACTGTCGCCCACTTTCTGGCGCGTGGTGGCAAGTTATGGTTGGCGAGAAACGCCAAAC
GTAGAAGAAGTTTTCCACCGCTGCGGTCTGGAAGGATTAAGTTGCCGGATGATGGAAACCTCCTTCTTTATGTCGCATGAGTCGTTGAT
CCTCGGCAAACGCCCGTGGTATTTGCGTCTGCGCGGCAAGCTGTACTTGCTGCTGCAACGTAATGCGCTGCGTGCACCAGATCAATTTG
AAATCCCGCCAAACAGGGTTATCGAACTGGGTACTCAGGTCGAAATCTAACGCCAGACGCCTCCTTTCTTCATAAGGGGGCGTTTTTGT
TTTCATGGTTAATCACCATGTAAAACGTTTCGAGGTTGATCACATTTCCGTAACGTCACGATGGTTTTCCCAACTCAGTCAGGATTAAA
CTGTGGGTCAGCGAAACGTTTCGCTGATGGAGAAAAAAATGAAAAAAGGCACCGTTCTTAATTCTGATATTTCATCGGTGATCTCCCGT
CTGGGACGGATCCTCTAGATTTAAGAAGGAGATATACATATG
>yejK
GAGGCAATTCCGACGTCTAAGAAACCATTATTATCATGACATTAACCTATAAAAATAGGCGTATCACGAGGCCCTTTCGTCTTCACCTC
GATGGATCCAGCAGCTCTGCGAGCAATTGTTCAACCTGTTCATCACTGTAGCGGGAAATTTGTGGCATAAATCGAAATCAGTTAGTTGA
TGAAAGGGCAACATATTACCGTAGCAACGGTTTTTTTTCTGCTTTTTTATGCGTTAAATCATTCTCGTTACGCTGCCTTCTGTAAAATG
ACCGCCATCTTTTATTGCGTCGCTTAACCGGCGGTGGTTGAATACCGCCCGGTCTTAAAGGAGAGTTTATCATGAGTCTGGATATCAAC
CAGATTGCTCTGCACCAGCTTATCAAGCGTGATGAGCAGAACCTTGAGCTGGTGTTGCGCGATTCATTGCTCGAGAGATCCTCTAGATT
TAAGAAGGAGATATACATATG
>lsrR
GAGGCAATTCCGACGTCTAAGAAACCATTATTATCATGACATTAACCTATAAAAATAGGCGTATCACGAGGCCCTTTCGTCTTCACCTC
GATGGATCCGGCGCAAAGTAGCGGTAACGCGCGGGTATCACTCGTTTGCATATTTCCCCCGTTCAGTTTTGCAGGTGAGTTTTGAACAA
ATGTATTTCTGCTTTTAATTTGTTCATAACCTTAGGTGGACATTGCACATATTTCCGACGAATAGATCACAATTTATGCTATTTTGATT
TTCACGGTTGCGTTTGTTCATGCTCGTAGAGTCAAACTGTGGTTGCCATCACAGATATAAATGAGCAAGAACTGAACAATTGCATTAAA
GATTTAAATATGTTCAAAGTGAAGAATGAATTATGACAATCAACGATTCGGCAATTTCAGAACAGGGAATGTGTGAAGAAGAACAGGTC
GCGCGGATCGCGTGGTTTTACTATCACGACGGGCTGACCCTCGAGAGATCCTCTAGATTTAAGAAGGAGATATACATATG
>ydiF
GAGGCAATTCCGACGTCTAAGAAACCATTATTATCATGACATTAACCTATAAAAATAGGCGTATCACGAGGCCCTTTCGTCTTCACCTC
GAGGCAACTTTTGGTGCGGTAAAAAAAGCGTCTGCGCCAGGGCAAATCTCGGTAAATGATTTGCGCACGGTATTAACTATTTTACACCA
GGCATAAGCAATAATATTTCGGCGGGAACACCCTCCCCGCCGAACTAAAAAATATATTCAATCGTATTTAATAAAAATATTTCGTGAGT
CTCTGTGCGCTAATTCTCCATTTGGCGTAGGGAAAATCACATCTGAATCAGGAATTAACAATGAAACCTGTAAAACCACCTCGTATTAA
TGGACGAGTGCCGGTCCTGTCGGCACAGGAAGCGGTGAATTATATTCCCGACGAAGCAACACTTGGATCCTCTAGATTTAAGAAGGAGA
TATACATATG
>ygbA
GAGGCAATTCCGACGTCTAAGAAACCATTATTATCATGACATTAACCTATAAAAATAGGCGTATCACGAGGCCCTTTCGTCTTCACCTC
GATGGATCCGCTGCATCATGGGCGTATGGGCGTCGAAATTTTCTATTGCACTCATGGGGTTATGTCCGGTTCCCTGATATTAAATGGGG
TGTGATGGTTTTTATTTTCGTTGTAATCACATAAGGCGCACATGATACCGGAGTTAATCATTTCAGGGGAAGAAGCGAACGGCGAGTTC
TTGATGCTGGTTCCGGGAAAGATGCTGTTTTGCAGCTTTTGCGTGATGCTTATGAAGCAAGTACGCAAAATTGTATCGGTTGGCTAACC
CTTCTTCTTTGGAGTCATAGTGCCCTTTAAGATTAAGATGTAATATAAATACATCTTAATGAGGTGGAGGCAGCATGTCCGGCAAGCGT
ATCTCTCGTGAAAAACTGACGATTAAAAAAATGATCGATCTTTATCAAGCGAAATGCCCGCTCGAGAGATCCTCTAGATTTAAGAAGGA
GATATACATATG
>ygbN
GAGGCAATTCCGACGTCTAAGAAACCATTATTATCATGACATTAACCTATAAAAATAGGCGTATCACGAGGCCCTTTCGTCTTCACCTC
GAGCCTGTTCGATGAGGTGGGATATCAGGGCTGGATCGGTTGTGAATATAAACCTCGTGGCCTCACCGAAGAAGGGCTTGGCTGGTTTG

Data Set S4 - MG1655 library reference file.fasta
ACGCCTGGCGCTAATTCGTTAAATCCCTAATTACAACGTACCCATACATCCCCCCATCTACGGATGGGGTAGGTTAACTATAACTTTCA
GACAGGGTTTCCCATGTCCACAATTACATTGTTATGCATTGCGTTAGCTGGCGTAATCATGCTGTTGCTGCTGGTCATCAAGGCGGATC
CCGGGATCCTCTAGATTTAAGAAGGAGATATACATATG
>ygbJ
GAGGCAATTCCGACGTCTAAGAAACCATTATTATCATGACATTAACCTATAAAAATAGGCGTATCACGAGGCCCTTTCGTCTTCACCTC
GAGTGACAATGCCTTTTTCAGCTACCATCTCAAGGATGATTTGGCGACGCTCTACGGGTATCAACTTTTGCTCCTTCCTTTGTCCTGCT
GACATTCTACGCTATTTGCCTGCGAAACGTGCGCGGCGCAACTAACGCTTAGTTCACATAAAATAACACACAATGTTAATTTATGTGAA
TCAGATCACCATACCGTTATCTTCCAGCGCTTATATTCACAATATCAAACAAAATATCACTTAAATTAACAAGGAGAGCAGATGAAAAC
GGGATCTGAGTTTCATGTCGGTATCGTTGGCTTAGGGTCAATGGGAATGGGAGCAGCACTGGGATCCTCTAGATTTAAGAAGGAGATAT
ACATATG
>ygbI
GAGGCAATTCCGACGTCTAAGAAACCATTATTATCATGACATTAACCTATAAAAATAGGCGTATCACGAGGCCCTTTCGTCTTCACCTC
GATGGATCCCTGCTCCCATTCCCATTGACCCTAAGCCAACGATACCGACATGAAACTCAGATCCCGTTTTCATCTGCTCTCCTTGTTAA
TTTAAGTGATATTTTGTTTGATATTGTGAATATAAGCGCTGGAAGATAACGGTATGGTGATCTGATTCACATAAATTAACATTGTGTGT
TATTTTATGTGAACTAAGCGTTAGTTGCGCCGCGCACGTTTCGCAGGCAAATAGCGTAGAATGTCAGCAGGACAAAGGAAGGAGCAAAA
GTTGATACCCGTAGAGCGTCGCCAAATCATCCTTGAGATGGTAGCTGAAAAAGGCATTGTCACTCGAGAGATCCTCTAGATTTAAGAAG
GAGATATACATATG
>adrA
GAGGCAATTCCGACGTCTAAGAAACCATTATTATCATGACATTAACCTATAAAAATAGGCGTATCACGAGGCCCTTTCGTCTTCACCTC
GAGTGCGCGAATGCAATAATCAAGCAACACAACAATCTCTGAAAGGTGATGATCGTAATAAGTTTATGAGTGCCTGCCTCAAGAAAGCC
GCCTGATACCTGATAGTGCTAACGGGTGAGCTACGAAAATGGCTCACCCGAAATATCATACTTCTGCCTTTAGCTCCGTCTCTATAATT
TGGGAAAATTGTTTCTGAATGTTCCCAAAAATAATGAATGATGAAAACTTTTTCAAAAAAGCGGCGGCGCACGGGGAGGAACCTCCTTT
AACTCCTCAAAACGAACATCAGCGGTCCGGATCCTCTAGATTTAAGAAGGAGATATACATATG
>dedA
GAGGCAATTCCGACGTCTAAGAAACCATTATTATCATGACATTAACCTATAAAAATAGGCGTATCACGAGGCCCTTTCGTCTTCACCTC
GATGGATCCCTTCCAAAACCGCCAATGGGCCCGCTATTTCTGGCGGACTAACGAGAATAATGCTCGGAACATTTCAGGCTGTAGCCAAC
GTAGTTACAGCCTGAAAGATGACGAGTACAAGGCATAGGCAAATATGGACCTGATTTATTTCCTCATCGATTTTATCCTGCACATTGAC
GTGCATCTGGCAGAGCTGGTCGCGGAGTACGGCGTCTGGGTTTATGCCACTCGAGAGATCCTCTAGATTTAAGAAGGAGATATACATAT
G
>lolC
GAGGCAATTCCGACGTCTAAGAAACCATTATTATCATGACATTAACCTATAAAAATAGGCGTATCACGAGGCCCTTTCGTCTTCACCTC
GAGGGTCAGATGCGGATAAAAGGTAATGACCGAGTGATAAATCACCACCAGACAAATACATAACCCTTTGATCTGGTTAATCCATAGCT
CTTTTTGTTTCATCAGGACAACACGCTCTTTTTTAATTTACATGCGGTTGATCCTGCGGGTTATGACCCGAGGTGTAATGGGTATCTGT
CTGTATTGGATGATTTTTCAGATTAAGATCAGGCGGCAAGATTGATGATAAAACATGGCAATTTAGCCGATTGATTTACGGGGGCTTTT
CAGATTAGCCCTGACGATCACTTACAGTTCAGACGTTTACCCATCTTGCTTTCGCTTATATACTCGTGTCTTTGCTACAGCAACCAGAC
GGATTTCATGTACCAACCTGTCGCTCTATTTATTGGCCTGCGTTACATGCGTGGGCGTGCAGCGGATCGCTTCGGTCGTTTCGTCTCCG
GATCCTCTAGATTTAAGAAGGAGATATACATATG
>ybiS
GAGGCAATTCCGACGTCTAAGAAACCATTATTATCATGACATTAACCTATAAAAATAGGCGTATCACGAGGCCCTTTCGTCTTCACCTC
GATGGATCCACTGCCGAACTGCATGGTGACGTTACTGGAAACTAACACAGGCGTATCCTGAAAAGAGATATGACAAACCGCGCATTATG
CCATAAGTCTGAAAGAAGATCCTGCTCCTCGCTTAAGGTCTACACTCAAGCCAAAGGCAAAAAATGTGATTTCGTACACATCTGATTTC
ACTGTGAGCTGGAATGAACTTATAATGCGCTTCCAATACTCTAATATTCTCAACCCAATGGCCTGCCAGGCACAAAATCTCGCTTAACA
TGAATATGAAATTGAAAACATTATTCGCAGCGGCCTTCGCTGTTGTCGGCTTTTGCAGTACCGCCTCTGCGGTACTCGAGAGATCCTCT
AGATTTAAGAAGGAGATATACATATG
>lolA
GAGGCAATTCCGACGTCTAAGAAACCATTATTATCATGACATTAACCTATAAAAATAGGCGTATCACGAGGCCCTTTCGTCTTCACCTC
GAGAACAGATGGAAGCGCAGGGGATTGTCAGCGAACAGGGGCACAACGGTAATCGTGAAGTGCTGGCCCCACCGCCGTTTGACTAACTA
ATGCATCGTATGCCGGATAAGGCGCGGTAGCGTCGCATCCGGCACTCTATCAACTGAAAATTCAGTATTTTCTTCTTTCCTCAAGCTGA
TTATTAGCCTGGAATAGAGAGTAGAGGGAACTCCCGATCGGGAGTGACGTAATTTGAGGAATAATGATGAAAAAAATTGCCATCACCTG
TGCATTACTCTCAAGCTTAGTAGCAAGCAGCGTTTGGGCTGATGCCGCAAGCGATCTGAAAAGCCGCCTGGATAGGATCCTCTAGATTT
AAGAAGGAGATATACATATG
>ybiV
GAGGCAATTCCGACGTCTAAGAAACCATTATTATCATGACATTAACCTATAAAAATAGGCGTATCACGAGGCCCTTTCGTCTTCACCTC
GATAGATCTCGCTCTCTCCAGACGCTCAGGACGATATCATCGCCCGTACTGAACATATGCTGTAATCTCCTCGGCCCGGCGGCGTCGTC
GCCGGGCTAAATCATTCACATCATCAATTTCATCCTTACTTTCATTCGAAATATAATTTGTGCTCTGCGTCACATTGTTATTACACTGT
TACGGTTTCATTTTCGAGCCAGGAGCACACCTATGAGCGTAAAAGTTATCGTCACAGACATGGACGGTACTTTTCTTAACGACGCCAAA
ACGTACAACCAACCACGTTTTATGGCGCCTCGAGAGATCCTCTAGATTTAAGAAGGAGATATACATATG
>dapF
GAGGCAATTCCGACGTCTAAGAAACCATTATTATCATGACATTAACCTATAAAAATAGGCGTATCACGAGGCCCTTTCGTCTTCACCTC
GAGTTTACCGAGCGAGCATCCTCACGCTGACGGAACTAAAAAAGACAACAAACAAACCACATTGCGATAGTGCATAAAGCCATCCTGGC
GCGAGGTGCCGATCACGAAACTACCAGCAAAACATAAATCCCCACGAGTAAGCGTTATACTCGCAGCATTTCCTCACTTTTCAGACTTC
ATAAAGAGTCGCTAAACGCTTGCTTTTACGTCTTCTCCTGCGATGATAGAAAGCAGAAAGCGATGAACTTTACAGGCAATCCATAATGA
AAAACGTGTTTAAGGCACTCACTGTATTACTTACTCTCTTCAGCCTGACGGGCTGCGGTCTGAAAGGTCCGCTCTATTTCCCGCCTGCA

Data Set S4 - MG1655 library reference file.fasta
GATAAAAACGCACCGCCGCCGACCAAACCGGTAGAGACGCAAACGCAATCCACGGTGCCGGATAAAAACGATCGCGCCACTGGCGATGG
TCCATCCCAGGTGAATTACTAAAAGTCAGTTTCTGTACCCGCGTGATTGGAGTAAATGATGCAGTTCTCGAAAATGCATGGCCTTGGCA
ACGATTTTATGGTCGTCGACGCGGTAACGCAGAATGGATCCTCTAGATTTAAGAAGGAGATATACATATG
>ybiA
GAGGCAATTCCGACGTCTAAGAAACCATTATTATCATGACATTAACCTATAAAAATAGGCGTATCACGAGGCCCTTTCGTCTTCACCTC
GATGGATCCTCGGGGATCTGTTCCTGAAGCGCCTTATACCAGGCGGCAATTTGCGCTTTAAGCGCGGCGGTTAATGCCATGAGAAAACC
TGAAATACTGTATAAACAGCCAATATTGTGGCATTTTTCGGTTTTCAGGGGTAATGGTTTTTCTGTTTCTGGAAAGCGGCTCGCAATAA
CCACCAACATAATGCGGTTTAATGCCCGTTCGAGCACAAAGGATCCAACACGTCATGCAAGACACCATCATAAATTTCTACAGCACCAG
CGATGACTACGCTCGAGAGATCCTCTAGATTTAAGAAGGAGATATACATATG
>ybiF
GAGGCAATTCCGACGTCTAAGAAACCATTATTATCATGACATTAACCTATAAAAATAGGCGTATCACGAGGCCCTTTCGTCTTCACCTC
GATGGATCCACGGAAGTACCTGCGGTGAAAGCCAGAACTGCGGCCAGTGCTGAAAGACATGCAATTTTTTTCATAACCACCTCAAATGT
GATTCAAATAAGTCCTAAGTTTTAAATATATCAAAAATTAATGGGAAACTCTTCGCGATTTGTGATGTCTAACGGGCCATTTCATGTAA
CAGAACGTTTCCATACACCGCTATCCATCTAAATTTAAATCACTTTTTCAGAGAACTGCGTAAGTATTACGCATGTTTTCCCTGTCATT
CATCCAGATTATTCCTAATCACCAGACTAATGATTCCATCAATCCTGGCGCATTTTAGTCAAAACGGGGGAAAATTTTTTCAACAAATG
CTCAACCAGCATTGGGTATATCCAGTACACTCCACGCTTTACTTAAGTCTAGATATTTGTGGGAGAAAGGATGCCTGGTTCATTACGTA
AAATGCCGGTCTGGTTACCAATAGTCATATTGCTCGTTGCCATGGCGTCTATTCAGGGTGGAGCCTCGTTACTCGAGAGATCCTCTAGA
TTTAAGAAGGAGATATACATATG
>ybiJ
GAGGCAATTCCGACGTCTAAGAAACCATTATTATCATGACATTAACCTATAAAAATAGGCGTATCACGAGGCCCTTTCGTCTTCACCTC
GATGGATCCCGCAGAGGTTCGAAAGACAGCCAGTTACGTTAACTACAACTGGCGAGATGCATAGCGAGTATAATTTGTATTTTGCGTGT
CATTCCGTGCCTTTAACGCTGCTGTGACTGCCACGCGGTATAAAACAAGTTCATAAGTACAACAAATAAATGGTTTATCAGTAGGTTAG
ATATCAATCAATTTATTTGAACAAGGCGGTCAATTCTCTTCGATTTTATCTCTCGTAAAAAAACGTGATACTCATCACATCGACGAAAC
AACGTCACTTATACAAAAATCACCTGCGAGAGATTAATTATGAAAACTATCAATACTGTTGTTGCTGCTATGGCTCTTTCAACTCTGTC
ATTTGGCGTGTTCGCGGCGGAACCGGTAACGGCATCCCAGGCACAGAACATGAATAAAATCGGCGTGGTTTCTGCCGATGGCGCATCCA
CCCTCGATGCCCTGGAAGCGAAACTGGTGACTCGAGAGATCCTCTAGATTTAAGAAGGAGATATACATATG
>ybiH
GAGGCAATTCCGACGTCTAAGAAACCATTATTATCATGACATTAACCTATAAAAATAGGCGTATCACGAGGCCCTTTCGTCTTCACCTC
GATGGATCCGCCTGCTGCTGAATAGGGGTGGGTTCACGGTAACCCTGCTCGGCAACGGCGCGCAGGATATCAGGGCTTAAACCCAAAGA
ATCGAAAGACATAACTACTCCGAACCGCCCCGACCGTTACCGGTGTAGTTTTCAGGGAGATACTGAAAAGGAAAATGACAAAAACCACA
ATGTCATGAGGGAGCGGAGTGTAGCAGTTTTTGTGACGCAGCGCATAAATTATCCCTGTGACTGGCGTTGCTAAAATTCTTAGCCATAC
TGGACAACTCCCCTATGGAGTCATAATCTTAATCAATCATTTGATTAAGAATGGACTGTGCGATGAATAATCCTGCCATGACAATCAAG
GGTGAACAGGCGAAAAAACAGCTGATTGCTGCCGCACTGGCGCAGTTTGGTGAATCTCGAGAGATCCTCTAGATTTAAGAAGGAGATAT
ACATATG
>ybiN
GAGGCAATTCCGACGTCTAAGAAACCATTATTATCATGACATTAACCTATAAAAATAGGCGTATCACGAGGCCCTTTCGTCTTCACCTC
GAGCGATGCAAATGACGCAAAAAATTGACAGAGATGTGCTTATTTTTTGAACGATATATTTTTACAAAATAAGACAAATCAATGACATG
CATAATGCTGATTTTTATGATGAATTGCCATAAACCATAGCCATGTTAAGGTTTTGGGGAATGATAAACGCAGCCAGATATTTACCCTG
TCGCGTTATGCGGGTATCTTACGCCGCTGCTTAAAGGAGAATGCTATGTCCGCCCAGAGGATCCTCTAGATTTAAGAAGGAGATATACA
TATG
>ybiO
GAGGCAATTCCGACGTCTAAGAAACCATTATTATCATGACATTAACCTATAAAAATAGGCGTATCACGAGGCCCTTTCGTCTTCACCTC
GATGGATCCCAGCGCTTGCAGGAATTTTTGCAGCACGTCTCTTAATAAGACACATTGCCTGATGCGCTACGCTTATCAGGCCTACAGGA
TATCTGGCAACTTATTAAAATTGCATGAACTTGTAGGACGGATAAGGCGTTCACGCCGCATCCGGCAAAAAAGCCCGCACGTTGTCAGC
AACCTGCTTAATATCCCTTCCTCCCTTTCACCCGAAAGGGAGGCACACCAGATTCCTCTCATTTAAAATCGCCCCTCCTCCCTCGAGAG
ATCCTCTAGATTTAAGAAGGAGATATACATATG
>ybiM
GAGGCAATTCCGACGTCTAAGAAACCATTATTATCATGACATTAACCTATAAAAATAGGCGTATCACGAGGCCCTTTCGTCTTCACCTC
GATGGATCCTCTGGGCGGACATAGCATTCTCCTTTAAGCAGCGGCGTAAGATACCCGCATAACGCGACAGGGTAAATATCTGGCTGCGT
TTATCATTCCCCAAAACCTTAACATGGCTATGGTTTATGGCAATTCATCATAAAAATCAGCATTATGCATGTCATTGATTTGTCTTATT
TTGTAAAAATATATCGTTCAAAAAATAAGCACATCTCTGTCAATTTTTTGCGTCATTTGCATCGCTCGAGAGATCCTCTAGATTTAAGA
AGGAGATATACATATG
>nac
GAGGCAATTCCGACGTCTAAGAAACCATTATTATCATGACATTAACCTATAAAAATAGGCGTATCACGAGGCCCTTTCGTCTTCACCTC
GATGGATCCCGCCGTTCTACCGACTGAACTACAGAGGAATCGTGTGAACGGGGCGCATATTAATGGCAGTGCATGGTGATGTCAAAGGG
TGAAATGCATTTTTTTGTTTGTTTGCTGTCAATTTCAACAAAGTGGTGAGCTTTCAATCTTATTGGGTAATGAACCATCGTGGTGCATA
CCCTCCTTTTATAGGGCAGGGGAACGCGACAGCTGATTAAAGGAGCAAATGACGAATCTATCATAATCTCTTATTTGACAGTTGGTTAG
CTTGTACATCAACACCAAAATAAAACTGGCAAGCATCTTGCAATCTGGTTGTAAGTAATGGCGGCACTTGGGCCGATTCTTAAAAACCG
GAGGCAACATGAACTTCAGACGCCTGAAATACTTCGTAAAAATTGTAGATATTGGTAGCCTGACCCAGGCTGCTGAAGTATTGCATATC
GCACAACCAGCGCTCGAGAGATCCTCTAGATTTAAGAAGGAGATATACATATG
>b2680
GAGGCAATTCCGACGTCTAAGAAACCATTATTATCATGACATTAACCTATAAAAATAGGCGTATCACGAGGCCCTTTCGTCTTCACCTC
GAGGGTTGGATCAAAGCCCACCAGCAGCAGTTCGATGGCTGGGTGAATGAGGCGCTGGCAGCGCAGAAGTAATTTTTATTCGGGCGGAT

AAGGCGTTTTTGCCGCATCCGCCGTTCTGTGCACAATGCCTGATGCGACGCTGGCGCGTCTTATCATGCCTACAAACCACATCGCACAT
TTCAACAATCTATCTTTCATCCCATATTCATCAACATCCGCTATTATTGATTTCCAGCTTAATCATCACCTGATGAACAAAAATAATGA
CTAAACCTAATCATGAGCTTAGCCCGGCGCTGATCGTGCTGATGTCTATCGCCACCGGTCTGGCGGTAGCCAGTGGATCCCGGGATCCT
CTAGATTTAAGAAGGAGATATACATATG
>asr
GAGGCAATTCCGACGTCTAAGAAACCATTATTATCATGACATTAACCTATAAAAATAGGCGTATCACGAGGCCCTTTCGTCTTCACCTC
GAGCGGCGCATTTATTGCTCTGATGCTGGTCATTGCTCTGCTGGTCGGGACGCGTTTGCATCGTCGTCTGCACGCCTGAAAAATAAGTC
CGGACTGCGGTAAATACCCGTCCGGACTTATTGCCAGCTCAAACCAACGTTAATAGCCATCCTAAAATAGACGAAGCGCCAGCCAATTC
CCGCAGCGCGTCTAGCGTCATCAGGATTATAAGTACCCAAATAAACGGATTCATTTTGCTGTGTGTCATTTATTACTGATGCGCAGTTA
TTCTACTGCTTTGTAAGTAGTAAAATAGTTAACCCGATCAAGACTACTATTATTGGTAGCTAAATTTCCCTTAAGTCACAATACGTTAT
TATCAACGCTGTAATTTATTCAGCGTTTGTACATATCGTTACACGCTGAAACCAACCACTCACGGAAGTCTGCCATTCCCAGGGATATA
GTTATTTCAACGGCCCCGCAGTGGGGTTAAATGAAAAAACAAATTGAGGGTATGACAATGAAAAAAGTATTAGCTCTGGTTGTTGCCGC
TGCTATGGGTCTGTGGATCCTCTAGATTTAAGAAGGAGATATACATATG
>oppA
GAGGCAATTCCGACGTCTAAGAAACCATTATTATCATGACATTAACCTATAAAAATAGGCGTATCACGAGGCCCTTTCGTCTTCACCTC
GAGATTGCTGATGGCATTGGGGATTGAATTTATCGTTACTGGTATTAAGGGGATTTTCCCCGGCCTGCTTAATTAATTCCTTTCAAATG
AAACGGAGCTGCCATGCTCCGTTTACTTCGTCATTATTTTTACTTTGTTCCCGCGCAGTTATCAAAAGCAAAAGGAATAGGTAAAAATA
TTCTTCTCAAATTACAGTTAGTTATAAGGATTTCCTTAACTGCTTCTCCTCACCATCATGTTATTTTCGCCACATCATAATCCTGGGCT
TGCTGAAGAATAATTGAAATGATATTATTAATTCCACTGCCTTTGGTAGAGGAAAGTGCTAAGGAAGGTGCGAACAAGTCCCTGATATG
AGATCATGTTTGTCATCTGGAGCCATAGAACAGGGTTCATCATGAGTCATCAACTTACCTTCGCCGACAGTGAATTCAGCAGTAAGCGC
CGTCAGACCAGAAAAGAGATTTTCTTGTCCCGCATGGAGCAGATTCTGCCATGGCAAAACATGGTGGAAGTCATCGAGCCGTTTTACCC
CAAGGCTGGTAATGGCCGGCGACCTTATCCGCTGGAAACCATGCTACGCATTCACTGCATGCAGCATTGGTACAACCTGAGCGATGGCG
CGATGGAAGATGCTCTGTACGAAATCGCCTCCATGCGTCTGTTTGCCCGGTTATCCCTGGATAGCGCCTTGCCGGACCGCACCACCATC
ATGAATTTCCGCCACCTGCTGGAGCAGCATCAACTGGCCCGCCAATTGTTCAAGACCATCAATCGCTGGCTGGCCGAAGCAGGCGTCAT
GATGACTCAAGGCACCTTGGTCGATGCCACCATCATTGAGGCACCCAGCTCGACCAAGAACAAAGAGCAGCAACGCGATCCGGAGATGC
ATCAGACCAAGAAAGGCAATCAGTGGCACTTTGGCATGAAGGCCCACATTGGTGTCGATGCCAAGAGTGGCCTGACCCACAGCCTGGTC
ACCACCGCGGCCAACGAGCATGACCTCAATCAGCTGGGTAATCTGCTGCATGGAGAGGAGCAATTTGTCTCAGCCGATGCCGGCTACCA
AGGGGCGCCACAGCGCGAGGAGCTGGCCGAGGTGGATGTGGACTGGCTGATCGCCGAGCGCCCCGGCAAGGTAAGAACCTTGAAACAGC
ATCCACGCAAGAACAAAACGGCCATCAACATCGAATACATGAAAGCCAGCATCCGGGCCAGGGTGGAGCACCCATTTCGCATCATCAAG
CGACAGTTCGGCTTCGTGAAAGCCAGATACAAGGGGTTGCTGAAAAACGATAACCAACTGGCGATGTTATTCACGCTGGCCAACCTGTT
TCGGGCGGACCAAATGATACGTCAGTGGGAGAGATCTCACTAAAAACTGGGGATAACGCCTTAAATGGCGAAGAAACGGTCTAAATAGG
CTGATTCAAGGCATTTACGGGAGAAAAAATCGGCTCAAACATGAAGAAATGAAATGACTGAGTCAGCCGAGAAGAATTTCCCCGCTTAT
TCGCACCTTCCCTAAATAATAATCAATTGTTAAATTATTGTGCATTTCACTACTGGAACTGTAATCAGAAAAGATAGACATGCTTAGCC
AATCTCTATTTGATTGAATTGAAAGATGTTTGTTAAGGCATGGATGCAAGCTATAGATTCTGATACGGTCAATAAAAGAGAATTGCTTA
ACAATTTTGCAAAATGTATTGGCGAGTAAGAACCGCATTTGGTACTTTCCGGGCAACCGCCAGACGATTCTTTATTGGTAATGAGAATA
ATTAACAATTAAAGAGCGTCGCGAAAGAATAATGTGTCTCGACAGGGGAGACACAGTACGAATCGACATAAGGTGATCGTCTGAATCAC
CAGAATAAATAAAGTCGGTGATAGTAATACGTAACGATAAAGTAACCTGACAGCAGAAAGTCTCCGAGCCTGTGCAGGGTCCCAATCCG
GGATTACACATGCTGGTTAATACCAGTAATTATAATGAGGGAGTCCAAAAAACAATGACCAACATCACCAAGAGAAGTTTAGTAGCAGC
TGGCGTTCTGGCTGCGCTAATGGCAGGGAATGTCGCGCTGGCAGCTGATGTACCCGCAGGCGTGGATCCTCTAGATTTAAGAAGGAGAT
ATACATATG
>oppB
GAGGCAATTCCGACGTCTAAGAAACCATTATTATCATGACATTAACCTATAAAAATAGGCGTATCACGAGGCCCTTTCGTCTTCACCTC
GAGGTCTGGTGAAACCGTGGGTTGGTGGCTATACCGGCAAAGATCCGCTGGATAATACCTATACCCGGAATATGTACATTGTGAAGCAC
TAATGGCAATACGTGGGGCAGGAGTGTCCTGCTCCACGGTGTCTGATTTTTATCGCATTACAGAAGGCACAGGCCAGAAGGTAGGGCAA
TGTTAAAATTTATTCTACGTCGCTGTCTGGAAGCGATTCCGACGCTATTTATTCTTATTACTATTTCGTTCTTTATGATGCGCCTCGCG
GATCCTCTAGATTTAAGAAGGAGATATACATATG
>asd
GAGGCAATTCCGACGTCTAAGAAACCATTATTATCATGACATTAACCTATAAAAATAGGCGTATCACGAGGCCCTTTCGTCTTCACCTC
GATGGATCCTTCCGAGCGGATCCATAATCAGGATCAATAAAACTGCTGCAGAAATGATTTCATTCATAACTCAAATTCCCTGATAATTG
CCGCGGACTTTCTGCGTGCTAACAAAGCAGGATAAGTCGCATTACTGATGGCTTCGCTATCATTGATTAATTTCACTTGCGACTTTGGC
TGCTTTTTGTATGGTGAAAGATGTGCCAAGAGGAGACCGGCACATTTATACAGCACACATCTTTGCAGGAAAAAAACGCTTATGAAAAA
TGTTGGTTTTATCGGCTGGCGCGGTATGGTCGGCTCCGTTCTCATGCAACGCATGGTTGAAGAGCGCGACCTCGAGAGATCCTCTAGAT
TTAAGAAGGAGATATACATATG
>yaiE
GAGGCAATTCCGACGTCTAAGAAACCATTATTATCATGACATTAACCTATAAAAATAGGCGTATCACGAGGCCCTTTCGTCTTCACCTC
GAGCGCGATTTACTGCAAAAACAGCTCGATGTTCCTGTCTTGCTGTCTAACGTATTGATTGCACGGCTGGCTGCGGAATTACTGGTGTA
ATTTTGCGTGACAGCCAGCGCCTCTGGCCCCTATAGTGAAGTAGATGTTCAACTACCAAACAGGGCCAGTTTATGCTTCAAAGTAATGA
GTACTTTTCCGGCAAAGTGAAATCAATCGGCTTTTCCAGCGGATCCCGGGATCCTCTAGATTTAAGAAGGAGATATACATATG
>dinB
GAGGCAATTCCGACGTCTAAGAAACCATTATTATCATGACATTAACCTATAAAAATAGGCGTATCACGAGGCCCTTTCGTCTTCACCTC
GAGTTTGGTCAGCATGGGGATAAAGTGGTGCAGCCGCTGGTGCAAAAGCTGGATAAGCAGCAGGTGCTTTCGCAGCGAACGCGTTAAAT
GCTGAATCTTTACGCATTTCTCAAACCCTGAAATCACTGTATACTTTACCAGTGTTGAGAGGTGAGCAATGCGTAAAATCATTCATGTG

```
GATATGGACTGCTTTTTCGCCGCAGTGGAGATGCGCGACAATCCCGCCCTGCGCGATATCCCTATTGGATCCTCTAGATTTAAGAAGGA
GATATACATATG
>recA
GAGGCAATTCCGACGTCTAAGAAACCATTATTATCATGACATTAACCTATAAAAATAGGCGTATCACGAGGCCCTTTCGTCTTCACCTC
GATGGATCCTTGCAGACCTTGTGGCAACAATTTCTACAAAACACTTGATACTGTATGAGCATACAGTATAATTGCTTCAACAGAACATA
TTGACTATCCGGTATTACCCGGCATGACAGGAGTAAAAATGGCTATCGACGAAAACAAACAGAAAGCGTTGGCGGCAGCACTGGGCCAG
ATTGAGAAACAATTTGGTAAAGGCTCCATCATGCGCTCGAGAGATCCTCTAGATTTAAGAAGGAGATATACATATG
>recE
GAGGCAATTCCGACGTCTAAGAAACCATTATTATCATGACATTAACCTATAAAAATAGGCGTATCACGAGGCCCTTTCGTCTTCACCTC
GATGGATCCGAATCTGCCCGCAGAAAAAAAAGCCGAATTTCTGGCAACTCTAATTGCAATGAAATTAAAAGGTGAAATCTGAAAGAAAT
AGCCTGCGTATGGCGCAGGCTATGAACAGTGTGTATCCGGCAAGATCATTCACTGAACAAAACGAATTTTAATCTGAGTTGAGGTTAAA
AAACAATGAGCACAAAACCACTCTTCCTGTTACGGAAAGCGAAAAAATCATCCGGTGAACCTGACGTCGTCCTGTGGGCAAGCAACGAT
TTTGCTCGAGAGATCCTCTAGATTTAAGAAGGAGATATACATATG
>recX
GAGGCAATTCCGACGTCTAAGAAACCATTATTATCATGACATTAACCTATAAAAATAGGCGTATCACGAGGCCCTTTCGTCTTCACCTC
GATGGATCCAGCAACCCGAACTCAACGCCGGATTTCTCTGTAGATGATAGCGAAGGCGTAGCAGAAACTAACGAAGATTTTTAATCGTC
TTGTTTGATACACAAGGGTCGCATCTGCGGCCCTTTTGCTTTTTTAAGTTGTAAGGATATGCCATGACAGAATCAACATCCCGTCGCCC
GGCATATGCTCGCCTGTTGGATCGTGCGGTACGCATTCTGGCGGTGCGCGATCACAGTGAGCAAGAACTGCGACGCTCGAGAGATCCTC
TAGATTTAAGAAGGAGATATACATATG
>ispB
GAGGCAATTCCGACGTCTAAGAAACCATTATTATCATGACATTAACCTATAAAAATAGGCGTATCACGAGGCCCTTTCGTCTTCACCTC
GAGTGTCCAGCTTTTCCAGGCGAACGGTCTGACCTTCGCTTACTCGGTGTTGTTTACCACCACTTTGGAAAACCGCGTACATATAAAAC
TCCGCTTCCGCGCACACCTTTTCAATGATTCAGAGTGCGCTATAAATATTCACAATAGGGCGCGAATATTACGCAAAACGCACGCCTTT
GACAAGTGCTACAGTCAATACACGAAGAAAAAAAACACAACTTGTACGGTAACGTTTATCTGTGCCATTTTTTCAGTACAATCACCCTA
TATTCCTAACCATAAACCCTAAGTTGCCTTTGTTCACAGTAAGGTAATCGGGGCGAAAAGCCCGGCTTTTGCGATGAATTTAGAAAAAA
TCAATGAGTTAACCGCGCAAGATATGGCGGGTGTTAATGCGGCAATCCTTGAGCAGCTTAATTCCGACGTCCAACTGATCAGGATCCTC
TAGATTTAAGAAGGAGATATACATATG
>hscB
GAGGCAATTCCGACGTCTAAGAAACCATTATTATCATGACATTAACCTATAAAAATAGGCGTATCACGAGGCCCTTTCGTCTTCACCTC
GATGGATCCGGCCTGAACGAAGGGTTTAAATTCACCAACCCGAACGTCAAAGATGAGTGTGGTTGCGGCGAAAGCTTCCACGTTTGATG
CGCATACCGATAACCCCACCGTGGTCGCCTGCGCGTGGGGTTTGTTTTACCTGATTCGCCGTTATCGCGGCGGATCGCAGCCCTGAGAA
TGTTATGGATTACTTCACCCTCTTTGGCTTGCCTGCCCGCTATCAACTCGATACCCAGGCGCTGAGCCTGCGTTTTCAGGACTCGAGAG
ATCCTCTAGATTTAAGAAGGAGATATACATATG
>ydiL
GAGGCAATTCCGACGTCTAAGAAACCATTATTATCATGACATTAACCTATAAAAATAGGCGTATCACGAGGCCCTTTCGTCTTCACCTC
GAGGCGTGGGTGGAAGAAGTCCCGCCGCCGACGGACCAACCGGAAGAAATTCTCGAAGAACTTGGCGAAATCGAGAAACCGAATAAGTA
ATTTCTCATCAGGCGGCTCTGCCGCCTGATTGTTAACCACCGCTAATTGATATTTGCTTGCTCTTCCCCATATTTTCCTGCTTACCGCC
ATTCAGCTGGTAGTACCTGTCGCAAATTCTTTACAGTTTTTAAACTAATGAGACGAATCTGATCGACGCAAAAAGTCCGTATGCCTACT
ATTAGCTCACGGTTATAAATCAACATATTGATTTATAAGCATGGAAATCCCCTGAGTGAAACAACGAATTGCTGTGTGTAGTCTTTGCC
CATCTCCCACGATGGGCTTTTTTTTAACATTTTTCCGCTTCGCTACCTCGCCCCTCACTCTTTCCAATAATGTTGCACAAGATACTGTG
ACTCTCTTCAAGCCGTCAGCAGGCTGATAATGTTATTTATATTTTATTGTTTATTTCATGCAGCCATTACACGATGAACGCTTATGAAC
TCCAGGCACTCCGCCATATTTTTGCCATGGATCCTCTAGATTTAAGAAGGAGATATACATATG
>ydiM
GAGGCAATTCCGACGTCTAAGAAACCATTATTATCATGACATTAACCTATAAAAATAGGCGTATCACGAGGCCCTTTCGTCTTCACGTC
GACCGAGTCTACCCTGATGGCAATTTTATTGACTGGAAGATCTATCAATCGGTAGCTGCTGAATTATATGCACACGATCTGGAGCGGCT
TTGTTAATTTTTCCACAGAAAGGAATTGTCGTTGTTACAACAATAATGAACGGATGCTGACACAACATCGCTTCACTTTTTAAAGCACC
TTTGCTAAGTAGAACCTATGAAAAATCCCTATTTCCCTACCGCACTTGGGTTGTATTTTAATTACCTGGTGCATGGTATGGGCGTCCTT
TTGATGAGCCTGAATATGGCCTCGCTGGAGGATCCTCTAGATTTAAGAAGGAGATATACATATG
>tatE
GAGGCAATTCCGACGTCTAAGAAACCATTATTATCATGACATTAACCTATAAAAATAGGCGTATCACGAGGCCCTTTCGTCTTCACCTC
GAGGAACGTGTGCGTCAGGTGCGCGCGCAACTGCCCGTCTTAAACAACCGTCGCTTTGCGCCGCCGCAATTATTATGATGTTTTTTTAC
TCGGCGCTTGATTCACCTTGTTACAGATTGCTATTGTGTGCGCGCGTCGAATGACCGTTAATATTCTCTGGTTTTTAAGGCGCGTTCTG
TTGCCGGTTATATGTCAAGAAGGTATCTATGGGTGAGATTAGTATTACCAAACTGCTGGTAGTTGCGGCGCTGGTCGTTCTGCTGTTTG
GGACTAAGAAGTTACGTACGCTGGGCGGAGACCTTGGATCCTCTAGATTTAAGAAGGAGATATACATATG
>pspF
GAGGCAATTCCGACGTCTAAGAAACCATTATTATCATGACATTAACCTATAAAAATAGGCGTATCACGAGGCCCTTTCGTCTTCACCTC
GATGGATCCGTCGGCAAAGCGAGAAAAAAATACCCATAATGTTGTCCTCTTGATTTCTGCGTTCAGGATTGTCCTGCTGAACTGTTAATA
CAATTTGCGTGCCAATTTTTTATCTTTTTGATTTATAAAGATCTGATTGAAGAATCAACAGCAACATGCCAGGATGAGTTAGCGAATTA
CACTAACAAGTGGCGAATTTCATCATGGCAGAATACAAAGATAATTTACTTGGTGAGGCGAACAGCTTTCTCGAAGTGCTGGAACAGGT
TTCGCATCTCGCACCTCGAGAGATCCTCTAGATTTAAGAAGGAGATATACATATG
>pspE
GAGGCAATTCCGACGTCTAAGAAACCATTATTATCATGACATTAACCTATAAAAATAGGCGTATCACGAGGCCCTTTCGTCTTCACCTC
GAGAATCAGTTGCTCGCCGACCGCTGAAAATGTTGCTGGCTGTGGCGCTGGAACCGCTGTTAAGTCGGGCTGCTAATAAACTGGCACAG
```

CGTTATAAAAGGTGAGGGGAGTTTCGCAAAAATTGTTAAATCTCAGGCGTATAATGGATGGCAATTTTCATCCATAGAAGGACGCTTAC
ATGTTTAAAAAAGGCTTACTTGCTCTGGCACTGGTGTTTTCACTGCCCGTTTTCGCCGCTGAACACTGGATCGGATCCTCTAGATTTAA
GAAGGAGATATACATATG
>pspB
GAGGCAATTCCGACGTCTAAGAAACCATTATTATCATGACATTAACCTATAAAAATAGGCGTATCACGAGGCCCTTTCGTCTTCACCTC
GAGTTTGCCGAACTGAAAGCCGATGATGCAATCAGCGAACAACTGGCACAATTAAAAGCCAAAATGAAGCAAGACAATCAATAATAACA
TCCAGGCGGCGTCCGAACGCGCCGCCGCTCATCGTCTAAGGAGTACTTATGAGCGCGCTATTTCTGGCTATTCCGTTAACCATTTTTGT
GCTGTTTGTTTTACCGATCTGGTTATGGCTGCATTACAGCAATCGTTCTGGTCGCGGATCCTCTAGATTTAAGAAGGAGATATACATAT
G
>moaA
GAGGCAATTCCGACGTCTAAGAAACCATTATTATCATGACATTAACCTATAAAAATAGGCGTATCACGAGGCCCTTTCGTCTTCACCTC
GAGATGAGAGAACGCGTCCCAGTCCATGCCCTCCGCCGAGAGCAACGACACGATCAAGATCAGCCAGCGTACGATTGCGCATATAGTTT
CCTGGAGTCAGATTATCCGCGCTACAGTAGCGCAAATTGCGGGGAAACAGCAATTAACCTGCCAATATTGAGGATACAAACTATTTTCT
TCTACCTCTAAAGGACGATGCACGCTATGCCTCCCTGATGATGTATATCAAAGTAAAACCGCCATTTTCCCTTATTCTGTAGCGAAATA
GCACGATCATGACGCTATATACATGATTACATAGCGAAAGTGTGGATGGTAAAAATCTCATTTACACGCTAGTATCGGCATAACCACTA
AACACTCTAGCCTCTGCACCTGGGTCAACTGATACGGTGCTTTGGCCGTGACAATGCTCGTAAAGATTGCCACCAGGGCGAAGGAAGAA
ATGACTTCGCCTCCCGTATCTGGAAAGGTGTACATGGCTTCACAACTGACTGATGCATTTGCGCGTAAGTTTTACTACTTGCGCCTGTC
GATTACCGATGTGTGTAACTTTCGTTGCACCTACTGCCTGCCGGATCCTCTAGATTTAAGAAGGAGATATACATATG
>dinJ
GAGGCAATTCCGACGTCTAAGAAACCATTATTATCATGACATTAACCTATAAAAATAGGCGTATCACGAGGCCCTTTCGTCTTCACCTC
GATGGATCCGATGGAATCGACGGCAAGGACATACTCAATAATGGCTGGTCATGCTTTAATGTTGCAGTGTATCCCGGATCGTTTCTGAA
CAAACTTAATAAACCATTATTATAAAATTAACATATATATTTCAGTTAATTATGTAAAAACATCCGTTTATTAAATTACTCATGGATTG
TCGCTGTTGCTCATTTGAGCTACAATTCAAGCTGAATAAATATACAGCACAGGAGATACCCCAATGGCTGCTAACGCGTTTGTTCGCGC
CCGAATCGATGAAGATCTGAAGAATCAGGCAGCGGACGTACTGGCCGGGATGGGGCTGACCATCTCTGACTCGAGAGATCCTCTAGATT
TAAGAAGGAGATATACATATG
>ydiY
GAGGCAATTCCGACGTCTAAGAAACCATTATTATCATGACATTAACCTATAAAAATAGGCGTATCACGAGGCCCTTTCGTCTTCACCTC
GATGGATCCGCGGTACAGCGCAGTTTTCCTTCGGGATAAATTTGCGGGGTAATTGTTGCGCTATCGAGAGAGGGCGCAAGTGTCAACGT
ATAGATACGTACCATCATTTCCTCCTATAGGCTGATTTCAGTCTGGCACCGAATCAGCGGAAAGTGAAGAAATTAACAATATGATTTAT
TGGAGCTTTATTTAAAATTTTGCAGATAAATATATATAAATAAAAATCTGTTTTCGCCACATGCTTATTCAAAAAATGAAATAAGAATA
CATTGCTAAGTTATTCGCGCCTTTTTATAATTCGTTACCTTTCCAGGGACGCTAATCTTTGATCCTTAAGAAAGTTTCCGGGACCGCGA
TGGTCTCTTTTTTTGTTGTACGGACTCTTGATAAATGAAGCTTTTGAAGACAGTTCCTGCCATCGTTATGTTGGCGGGGGGCATGTTTG
CGTCACTGAATGCAGCTGCCGATGATTCCGTTTTCTCGAGAAAACGGAATCATCGGCCGCTCGAGAGATCCTCTAGATTTAAGAAGGAG
ATATACATATG
>ydiP
GAGGCAATTCCGACGTCTAAGAAACCATTATTATCATGACATTAACCTATAAAAATAGGCGTATCACGAGGCCCTTTCGTCTTCACCTC
GATGGATCCATTTTGGCGTCGGCATTGTCGAAATTCAGGGTGTATTCTGGAGTGACAACAATGTCCTGTTCTTCAGGCACCAGCTTAAA
GCAGGTTATTATTTTCATGATTTCTCCTGTGAGTCGCATACGGAACCATCTCCGATGGAAGGTAAAAATGGCCTGAAAATAGTGTGCGA
TAGCGGTGCGGGGTTTTCGACGGAGCAAGCTGTTTTCCTTGCCAACAATCTTGCAAAAAAAATGCGATCGCTAAAACAAAAGTGCAATT
TGCGGCAATATAGTTATTCCATGACCGTACAATCTTGCATTCAGTGTAAATAATCTTGTTTAGCTCTCGCTAAACGCGGTTGTATAACC
GCAATACACGCCGTATTTTATCCGGCATATATTGTTTTTGCTTGCTCATGGAATAGGGAGGTTGTCATGTATCAACGCTGTTTTGATAA
CGCCAGTGAAACGCTGTTTGTCGCCGGTAAAACGCCACGGCTTTCACGTTTTGCATTTAGCGATGATCCAAAATGGGCTCGAGAGATCC
TCTAGATTTAAGAAGGAGATATACATATG
>ydiQ
GAGGCAATTCCGACGTCTAAGAAACCATTATTATCATGACATTAACCTATAAAAATAGGCGTATCACGAGGCCCTTTCGTCTTCACCTC
GAGCCCATTTTGGATCATCGCTAAATGCAAAACGTGAAAGCCGTGGCGTTTTACCGGCGACAAACAGCGTTTCACTGGCGTTATCAAAA
CAGCGTTGATACATGACAACCTCCCTATTCCATGAGCAAGCAAAAACAATATATGCCGGATAAAATACGGCGTGTATTGCGGTTATACA
ACCGCGTTTAGCGAGAGCTAAACAAGATTATTTACACTGAATGCAAGATTGTACGGTCATGGAATAACTATATTGCCGCAAATTGCACT
TTTGTTTTAGCGATCGCATTTTTTTTGCAAGATTGTTGGCAAGGAAAACAGCTTGCTCCGTCGAAAACCCCGCACCGCTATCGCACACT
ATTTTCAGGCCATTTTTACCTTCCATCGGAGATGGTTCCGTATGCGACTCACAGGAGAAATCATGAAAATAATAACCTGCTTTAAGCTG
GTGCCTGAAGAACAGGACATTGTTGTCACTCCAGAATACACCCTGAATTTCGACAATGCCGACGCCAAAATGGATCCTCTAGATTTAAG
AAGGAGATATACATATG
>ydiS
GAGGCAATTCCGACGTCTAAGAAACCATTATTATCATGACATTAACCTATAAAAATAGGCGTATCACGAGGCCCTTTCGTCTTCACCTC
GAGTGCCGACTACGGTCTGGTGGGCGATATCTACAAAGTCGTCCCTGCCCTGATTAGCCAGTTGAGCCGCTAATTCCCCTTTCAACCAC
ATCTCCCGCTGTAAACGCAGCGGGAAGGGAGCATAACGCATGTCGGATGACAAATTTGATGCCATTGTGGTCGGTGCGGGCGTTGCTGG
TAGCGTTGCCGCACTGGTCATGGCACGAGGGATCCTCTAGATTTAAGAAGGAGATATACATATG
>ydiU
GAGGCAATTCCGACGTCTAAGAAACCATTATTATCATGACATTAACCTATAAAAATAGGCGTATCACGAGGCCCTTTCGTCTTCACCTC
GATGGATCCTTTCCATGCTCTCCAGGGATGCCTGTGGCCTGCCGTTCCCATAAATCAGATAACGACGTTGGTTCAGCGATAACCCTTCT
GTTTGCTGGTGTTTAAGACGAGAGTAACCGTCTACACTATCAAACAGGAGGATCTATGACCCTGTCTTTTGTTACCCGCTGGCGCGATG
AATTGCCAGAAACCTATACAGCACTTTCCCCTACGCCCTCGAGAGATCCTCTAGATTTAAGAAGGAGATATACATATG
>ydiV

```
GAGGCAATTCCGACGTCTAAGAAACCATTATTATCATGACATTAACCTATAAAAATAGGCGTATCACGAGGCCCTTTCGTCTTCACCTC
GATGGATCCGCAAGGGAGTGATGCGTTCCTCACTTGATAATGTCTATTGGCAGAAAAATTTCTGGCAGGCGAGACGAATCTAGTATGTC
GAAAAATGGACGGCGATGATACATCGCCGTCCATTTACTCCTGAATACAGTAGTATTTTGCAGACAAAAATTCCGTAAATGGATTATTC
GTTTATAATCAGGATAATCTAAAAAAAGGACCCCTGAAAAAGTGGTTAACATGGAACAAATGAAAGTTACTCCTTCTGTAATTGTTATG
TCTTATGGCCTCATCCCTTTTAGCCGGATACTGAAAAACATCCTTCGAGAGGGACGGTTACCATGAAGATTTTTTTGGAGAATCTTTAT
CATTCTGATTGCTATTTCCTGCCGATAAGAGACAACCAGCAAGTCCTTGTCGGCGTTGAGTTCTCGAGAGATCCTCTAGATTTAAGAAG
GAGATATACATATG
>dinG
GAGGCAATTCCGACGTCTAAGAAACCATTATTATCATGACATTAACCTATAAAAATAGGCGTATCACGAGGCCCTTTCGTCTTCACCTC
GAGCGTAGTCATCGCTGGTGCTGTAGAAATTTATGATGGTGTCTTGCATGACGTGTTGGATCCTTTGTGCTCGAACGGGCATTAAACCG
CATTATGTTGGTGGTTATTGCGAGCCGCTTTCCAGAAACAGAAAAACCATTACCCCTGAAAACCGAAAAATGCCACAATATTGGCTGTT
TATACAGTATTTCAGGTTTTCTCATGGCATTAACCGCCGCGCTTAAAGCGCAAATTGCCGCCTGGTATAAGGCGCTTCAGGAACAGATC
CCCGAAGATCCTCTAGATTTAAGAAGGAGATATACATATG
>yceP
GAGGCAATTCCGACGTCTAAGAAACCATTATTATCATGACATTAACCTATAAAAATAGGCGTATCACGAGGCCCTTTCGTCTTCACCTC
GATGGATCCTCCAGGAAACGTGGGAAAGCGCCGATGACTGGTTTGTCAGCGAATAATATGCAGTGATTTTTTTTGCCGGGTCGCCCCGG
CTTTTTTATTTTACTTACGAATTTTGAGTACAAAATTTCTACACTTCCTCAAAATAATTGCACAATATGCTAAAAAAATATGAATGACA
TGCTGTTTTATTGTTCATATAAATCTTAAAAGCAGAAATTTATCTGTTTCCCCCTTTAAAAATTCGGTGAATACCCTTACTTATTGGTA
TACTGAAGTTGCTTCAGATAAAAACGCATTGAACCTCGAATAACGTTGTCTAGTAACACGAATTAGGGGGCCATGATGGAAAAAAATAA
TGAAGTCATTCAGACTCATCCGCTCGTAGGGTGGGACATCAGCACCGTTGATAGCTATGATGCGCTGATGTTGCGTTTGCACTCGAGAG
ATCCTCTAGATTTAAGAAGGAGATATACATATG
>guaC
GAGGCAATTCCGACGTCTAAGAAACCATTATTATCATGACATTAACCTATAAAAATAGGCGTATCACGAGGCCCTTTCGTCTTCACCTC
GAGAGCAAACGCATTGGCAACGGTACTCTTGCCACTGCCAATGCCTCCCGTTAAGGCAACTATATACCTCATAATTTATATTCCCGGGA
ATTCATCATGATTATCAAAACGTTAAAAATGAGTGCACGAAAGCGAAATTGATGAAACGTTCGCTCACTATTTACCAGGTAAATTTATG
GGATTGTAGCGTAAAAAAAGACAATTTCGCAGTCTTGCGCCGCATTGATTAGTGCGTATGATAGCGTCACTGGAGTTGCGCTCTTACCC
TTATAGCCATTAACCCCAGGAATCCGCACATGCGTATTGAAGAAGATCTGAAGTTAGGTTTTAAAGACGTTCTCATCCGCCCTAAACGC
TCCACTCTTAAAAGCCGTTCCGATGTTGAACGGATCCTCTAGATTTAAGAAGGAGATATACATATG
>guaB
GAGGCAATTCCGACGTCTAAGAAACCATTATTATCATGACATTAACCTATAAAAATAGGCGTATCACGAGGCCCTTTCGTCTTCACCTC
GATGGATCCCCGTTTGATTCAGGCGACTAACGGTAAAAATTGCAGGGGATTGAGAAGGTAACATGTGAGCGAGATCAAATTCTAAATCA
GCAGGTTATTCAGTCGATAGTAACCCGCCCTTCGGGGATAGCAAGCATTTTTTGCAAAAAGGGGTAGATGCAATCGGTTACGCTCTGTA
TAATGCCGCGGCAATATTTATTAACCACTCTGGTCGAGATATTGCCCATGCTACGTATCGCTAAAGAAGCTCTGACGTTTGACGACGTT
CTCCTCGTTCCTGCTCACTCTACCGTTCTGCCGACTCGAGAGATCCTCTAGATTTAAGAAGGAGATATACATATG
>guaA
GAGGCAATTCCGACGTCTAAGAAACCATTATTATCATGACATTAACCTATAAAAATAGGCGTATCACGAGGCCCTTTCGTCTTCACCTC
GATGGATCCGCATTCAGGAAAGCCACGTTCACGACGTGACCATTACTAAAGAGTCCCCGAACTACCGTCTGGGCTCCTGATTCTCTTCG
CCCGACTTCATGTCGGGCGATTTATATTATCTGTTTCACTTGCCTCGGAATAAGCGTCAATGACGGAAAACATTCATAAGCATCGCATC
CTCATTCTGGACTTCGGTTCTCAGTACACTCAACTGGTTGCGCGCCGCGTGCGTGAGCTGGGTGTTCTCGAGAGATCCTCTAGATTTAA
GAAGGAGATATACATATG
>cadA
GAGGCAATTCCGACGTCTAAGAAACCATTATTATCATGACATTAACCTATAAAAATAGGCGTATCACGAGGCCCTTTCGTCTTCACCTC
GATAGATCTGGATAACCACACCGCGTCTAACGCACATTAATTAAAAGTATTTTCCGAGGCTCCTCCTTTCATTTTGTCCCATGTGTTGG
GAGGGGCCTTTTTTACCTGGAGATATGACTATGAACGTTATTGCAATATTGAATCACATGGGGGTTTATTTTAAAGAAGAACCCATCCG
TGAACTTCATCGCTCGAGAGATCCTCTAGATTTAAGAAGGAGATATACATATG
>yaiI
GAGGCAATTCCGACGTCTAAGAAACCATTATTATCATGACATTAACCTATAAAAATAGGCGTATCACGAGGCCCTTTCGTCTTCACCTC
GAGGACCGCTGGCAATCAGACCGCCGAGAATGGCTTTTCCCATATTGCCGCAGCCAATAAAACCGATTTTCTTTTCCATTGCCTCACTC
CTGCCGTGAAATTCATTGTTTTGATAATCGCTGGCAGAAGCATAAACAGAACTATGCCGGAAGGCAAAAGGATCCTCTAGATTTAAGAA
GGAGATATACATATG
>yaiM
GAGGCAATTCCGACGTCTAAGAAACCATTATTATCATGACATTAACCTATAAAAATAGGCGTATCACGAGGCCCTTTCGTCTTCACCTC
GATGGATCCAATGACGCCTTCGACCTGATGCATGAAGGCAAATCCATTCGAACCGTAATTCGTTACTGATTTCCCGCAGGTTTACCCCG
TCCACTTCAGGCGGGGGTTCTTAATACTCTCCCCGGGCAGCCGTCCGGGGGATTAACCCTGAGATAATGACTGATGGAACTCATTGAAA
AACATGTCAGCTTTGGCGGCTGGCAAAATATGTATCGGCATTATTCCCAATCACTGAAATGTGAAATGAATGTCGGCGCTCGAGAGATC
CTCTAGATTTAAGAAGGAGATATACATATG
>dfp
GAGGCAATTCCGACGTCTAAGAAACCATTATTATCATGACATTAACCTATAAAAATAGGCGTATCACGAGGCCCTTTCGTCTTCACCTC
GAGAGCAGCTCGACATCCGTTAAGGCGCTAATACCAAACTTCAGCATTTTTTCGCGCGGCATCAACAGCTGTGAATTGTTTTTCACCTT
CACCTCCTTTGTGGTGCTCGCATCCTGTCATAGCCGTTTCGTTAAATCGACGGCCAGTTTTCAATCCTGGAAAGCGCCTCGCAAAGTGA
ATCTCAGGTGAAGGCACGACAACAAAAGTATTGTGATAAAATCGCCAACTTCTGGTGTCACCCTACAGGAAAAATCATCATGAGCCTGG
CCGGTAAAAAGGATCCTCTAGATTTAAGAAGGAGATATACATATG
>yaiN
```

Data Set S4 - MG1655 library reference file.fasta
GAGGCAATTCCGACGTCTAAGAAACCATTATTATCATGACATTAACCTATAAAAATAGGCGTATCACGAGGCCCTTTCGTCTTCACCTC
GATGGATCCTGGTACGACACCCCAACCGATGATTTTAACGGTCTGCAACTTGCAGCCCGTCTGACCTGGAAATTCTGATTCCTTCTGCC
GCCCGCTATCCGGGGCGGCCTTCCCTGCCGATTAGCCCCCCCCCCTTTCCTCTTTGTTTTCCGACCACATTCACCGGATAAATTTTATT
CTCCAGTGTTATATACTATAGGGGGGTATGCATTGACATATAGAATACCCCCCTATAGTATATTGCATGCAGATGATGAGGTGCGAAAT
GCCCAGTACTCCGGAAGAGAAGAAAAAGGTCCTTACTCGAGTTCGTCGTATTCGGGGGCAGATTGACTCGAGAGATCCTCTAGATTTAA
GAAGGAGATATACATATG
>yaiP
GAGGCAATTCCGACGTCTAAGAAACCATTATTATCATGACATTAACCTATAAAAATAGGCGTATCACGAGGCCCTTTCGTCTTCACCTC
GATGGATCCCCCCAAGTACCTGGCTTTCATTTATGCCTCAGGTTTTTGAATCCGTTAAAGAAGAATATTTCACAGTCAAACTTGCTGCA
TTAAAAAAACATAAAAGCTAGGAACGACGCGATTATATGCGTCATGATCGCCTGCGTGCAGTTGCACAATTTCGCGGACAACAGGTCAA
TAGCGATCTGGGTGAAGGCTTTGTAATTCATAAAATGATTCTTTGAGCAAATTGCATGAAAACCTGGATATTTATCTGTATGTCCATAG
CAATGTTGCTATGGTTTTTAAGTACGCTAAGACGTAAACCCAGTCAAAAGAAAGGCTGCTCGAGAGATCCTCTAGATTTAAGAAGGAGA
TATACATATG
>cadB
GAGGCAATTCCGACGTCTAAGAAACCATTATTATCATGACATTAACCTATAAAAATAGGCGTATCACGAGGCCCTTTCGTCTTCACCTC
GATGGATCCCAGGGGCAAACACCCTTTACTGGATTGAAAATGGTATATTCCAGACTTCTGTTCCTTATGTTGTACCTTATCTCGACAAA
TTTCTTGCTTCAGAATAAGTAACTCCGGGTTGATTTATGCTCGGAAATATTTGTTGTTGAGTTTTTGTATGTTCCTGTTGGTATAATAT
GTTGCGGCAATTTATTTGCCGCATAATTTTTATTACATAAATTTAACCAGAGAATGTCACGCAATCCATTGTAAACATTAAATGTTTAT
CTTTTCATGATATCAACTTGCGATCCTGATGTGTTAATAAAAAACCTCAAGTTCTCACTTACAGAAACTTTTGTGTTATTTCACCTAAT
CTTTAGGATTAATCCTTTTTTCGTGAGTAATCTTATCGCCAGTTTGGTCTGGTCAGGAAATAGTTATACATCATGACCCGGACTCCAAA
TTCAAAAATGAAATTAGGAGAAGAGCATGAGTTCTGCCAAGAAGATCGGGCTATTTGCCTGTACCGGTGTTGTTGCCGGTAATATGATG
GGGAGCGGTATTGCTCGAGAGATCCTCTAGATTTAAGAAGGAGATATACATATG
>yceF
GAGGCAATTCCGACGTCTAAGAAACCATTATTATCATGACATTAACCTATAAAAATAGGCGTATCACGAGGCCCTTTCGTCTTCACCTC
GATGGATCCTCAAGGCGTTTTTGAGCCGTACGAACCGGATCGAGAGTCAGGGGTAATTTTACCTTTTGCATAGGGCGCGCATATTAACT
TTGTAACGTCATAGAGTCAAAGAAAAAGGCAGCCTGTGGTTGCCTTTTGCCAATAATTCGCACACATTGCGGGTTACTGCTTTATTTTC
GTTCAGTGCTGATAACAAACATCAACCTGCAAAACGAAGCGCATAGTTTAAAATGGCTATCATCAATACGCTATATCTGATGGAAAAAA
ATATGCCTAAACTTATTTAGCCTCCACCTCGCCATGGCGTCGCGCTCTTCTGGAAACTCGAGAGATCCTCTAGATTTAAGAAGGAGAT
ATACATATG
>yaiW
GAGGCAATTCCGACGTCTAAGAAACCATTATTATCATGACATTAACCTATAAAAATAGGCGTATCACGAGGCCCTTTCGTCTTCACCTC
GAGACGTCTGCGCAGCTTTGAACATGAGCTGGATGGTGACAAAATTCAGGAAGTAACCCATACCTTGAGCTAAAAAGGAGAAGTAATGT
CGCGCGTCAATCCTCTATCTTCGCTGTCATTGTTGGCGGTTCTGGTTTTAGCGGGGTGTAGTAGTCAGGCTCCGCAACCGGATCCTCTA
GATTTAAGAAGGAGATATACATATG
>gpt
GAGGCAATTCCGACGTCTAAGAAACCATTATTATCATGACATTAACCTATAAAAATAGGCGTATCACGAGGCCCTTTCGTCTTCACCTC
GAGTGATAGGACGGGTGAGGAATAGAACAGATTTTGGCAAAAATATCCCACAGCGGCTGTGGAGATAATTGAGACAGTTCAGACACGTT
AAGTCTCCTTGTCGATCACCCGCAAAACAGTATTGCAGGTCACAGGGTTAGCAGAAAATGTTGTCAACACAAGACAGGCTTGCGAGATA
TGTTTGAGAATACCACTTTATCCCGCGTCAGGGAGAGGCAGTGCGTAAAAAGACGCGGACTCATGTGAAATACTGGTTTTTAGTGCGCC
AGATCTCTATAATCTCGCGCAACCTATTTTCCCCTCGAACACTTTTTAAGCCGTAGATAAACAGGCTGGGACACTTCACATGAGCGAAA
AATACATCGTCACCTGGGACATGTTGCAGATCCATGCACGTAAACTCGCAAGCCGACTGATGCCTTGGATCCTCTAGATTTAAGAAGGA
GATATACATATG
>yaiX
GAGGCAATTCCGACGTCTAAGAAACCATTATTATCATGACATTAACCTATAAAAATAGGCGTATCACGAGGCCCTTTCGTCTTCACCTC
GATGGATCCTCCGGCCCTAAGACATCAATCATCTGTTCTCCAATGACTAGTCTAAAAACTAGTATTAAGACTATCACTTATTTAAGTGA
TATTGGTTGTCTGGAGATTCAGGGGGCCAGTCTAGTAATTATGCGTTTATTCGTCCTGGCACAATAATCAGCAATGGCGTAAAAATTGG
TTTTGCCACCGAGATCTCGAGAGATCCTCTAGATTTAAGAAGGAGATATACATATG
>yaiZ
GAGGCAATTCCGACGTCTAAGAAACCATTATTATCATGACATTAACCTATAAAAATAGGCGTATCACGAGGCCCTTTCGTCTTCACCTC
GAGGTTGCCAGGCGTTGAAGAGGCATTGCGATATTTTCCGCTAAACAGCGATTTTGACAGGGTGAAATCAGCCATAAAAACCTCCGGAA
GTCATCTGCTTCTCTTCTGGTGTGATACCAGCGGCTTATGGCAAGTCTGACAGGTCATTTAGATTGTCAATATCAGAATCATGGTAAAT
TGATGTTGGGAATATTCCCGAAGCGTGAATCTTCGTTGTGAGTCACAAAATGGCAACCTTAAGTTAAATTTATTAACAATTACGAAAAT
GTTGTCTACACCACAGACGGTTGTTGCAGAGGATCCTCTAGATTTAAGAAGGAGATATACATATG
>gpp
GAGGCAATTCCGACGTCTAAGAAACCATTATTATCATGACATTAACCTATAAAAATAGGCGTATCACGAGGCCCTTTCGTCTTCACCTC
GATGGATCCGATCTGCCAAAACCGCTGCGCCTCACGCGCCCGCGCACAGGCAATGGTCCGCGTCGTACTGGCGCTCCGCGTAATCGTCG
TCGTTCAGGTTAAATAAAAAATGTCATACTCAAAATCATTCAAACTGTATCAAGATGGCAAGAGAATGAATCCCGATGAGCTTACTGTA
GTAAGTGATTCAGGTGAAAGAATGCAGCCAACACAGAGACAGATTGAAGGATGAAGAGTATGGGTTCCACCTCGTCGCTGTATGCAGCC
ATTGATCTCGGTTCGAATAGTTTTCATATGCTGGTTGTGCGCGAGGTGGCTGGAAGCATCCTCGAGAGATCCTCTAGATTTAAGAAGGA
GATATACATATG
>gntT
GAGGCAATTCCGACGTCTAAGAAACCATTATTATCATGACATTAACCTATAAAAATAGGCGTATCACGAGGCCCTTTCGTCTTCACCTC
GAGCTGCTGAACGAATTCCCGGAGCTGAAAGGTGTGCGCGATCTCACCGAACACCAGCGCGGCGAACACTCCTACTACTAAGTTATCCC

```
CTCTTTTGTAGATTGCCCGATGCGACGCTAAAGCGTCATTTCGGGCCTACGTTGAACATGTGCCAGACGTTGGCGCAGCATTGCTTTGT
CAAATCCTCGGTTCGTTTCCCCGCGATAATATGACCAACCTCTCATAATTTAAATTTACCCCGCTCTGGTGATTCTCAAACGCCAGATG
TTACCCGTATCATTCACATGGGTACCAAACATACTCCTGACATCTGACTACAATAATTAGTTTTAGTGGGTATCAGTCGTGGTGCCGCA
ATATCTCTGTTCCCGATTGGGATAATTAGAGTTTGTCGTCAGAAAATTGACGTTACCCATAACAAATGAAAGGCCAGGTAAATCATGCC
ATTAGTCATTGTTGCTATCGGTGTAATCTTGTTGTTGCTCCTGATGATCCGCTTCAAAATGAACGGCTTCATCGCTCTCGTCCTCGTGG
ATCCTCTAGATTTAAGAAGGAGATATACATATG
>ubiX
GAGGCAATTCCGACGTCTAAGAAACCATTATTATCATGACATTAACCTATAAAAATAGGCGTATCACGAGGCCCTTTCGTCTTCACCTC
GATGGATCCACGCCAAAGCAGTGCAACGTCAGAACGAAGTGGAAAATCTCGAAATGCATAACGAAGGATGATGCCTTCGCTGAGGGTGC
CGGTCTGGCACCCTGACTTGCAACTCCCGCCGAAATCCTGCAAAGTCTGCCTGCAAGTCTGACAGGGCAACTATTTATGAAACGACTCA
TTGTAGGCATCAGCGGTGCCAGCGGCGCGATTTATGGCGTGCGCTTATTACAGGTTCTGCGCGATGTCTCGAGAGATCCTCTAGATTTA
AGAAGGAGATATACATATG
>b0501
GAGGCAATTCCGACGTCTAAGAAACCATTATTATCATGACATTAACCTATAAAAATAGGCGTATCACGAGGCCCTTTCGTCTTCACCTC
GAGCGCACGTAGCGGTGGATATATCAATATACACAAATGTAGAAGATGATTATTTTTTTCTTATTTTTCCCTAACAAATGATGATGGGG
GTAGGTTTAAGAAAACATTGACAGTGAGGGCCAGGGAACAAGTGGCGAAAATCGTATCAAAGAATGATCCAGATACAAAAAAAGTGTGG
TGTAAATATGGTAAGATACCAGGGCAAGGGGGGATCCTCTAGATTTAAGAAGGAGATATACATATG
>yceK
GAGGCAATTCCGACGTCTAAGAAACCATTATTATCATGACATTAACCTATAAAAATAGGCGTATCACGAGGCCCTTTCGTCTTCACCTC
GAGTTAACGCTCTGGCAACCGCAATGGCGACCGCGCGTCACCGCGCCAGTAAGGTGCTGGAAATCGCCCGTGACCGCCACGTTGAACAG
GCGCTGAACGAGACGCCAGAGAAGCTGAATCGCGATCGTCGCCTGGTGCTGCTAAGCGATCCGGTGACGATGGCCCGTCTGCATTTCCG
TGTCTGGAATTCCCCGGAGAGATATTCTTCATGGGTGAGTTATTACGAAGGGATAAAGCTCAATCCACTGGCATTGCGTAAACCGGATG
CGGCTTCGCAATAAAAAGGATCCTCTAGATTTAAGAAGGAGATATACATATG
>greA
GAGGCAATTCCGACGTCTAAGAAACCATTATTATCATGACATTAACCTATAAAAATAGGCGTATCACGAGGCCCTTTCGTCTTCACCTC
GATGGATCCCCAGCGGGGAGTTGAGTAATGTACTCATCAACATTTGCGGCCTGAACACTGAACGCTATACAGCTGGTCAATCCGATGAT
AAATCTGGAAAATCGCATAATCTCGCGCTAACAACCTGGAATCGAGCCGTCATACTACGGCGCAACGCCCTATAAAGTAAACGATGACC
CTTCGGGAACTTCAGGGTAAAATGACTATCAAAATGTGAATTGTAGCTGACCTGGGACTTGTACCCGGGTCGGTATTTTTTTGCTTCTG
GTCCCGGTAAGGAGTTATGCCGGGCAGGCCGAACAGCCGGGGTGGGTGAAGACTTGCCCTATCAGGAATATTCAAGAGGTATAACAAAT
GCAAGCTATTCCGATGACCTTACGCGGCGCTGAAAAATTACGCGAAGAGCTGGATTTTCTGAAATCTGTGCGCCGTCCTGAAATCATTC
TCGAGAGATCCTCTAGATTTAAGAAGGAGATATACATATG
>greB
GAGGCAATTCCGACGTCTAAGAAACCATTATTATCATGACATTAACCTATAAAAATAGGCGTATCACGAGGCCCTTTCGTCTTCACCTC
GAGTCGAACCTGGAAGCCTTGTTCGGTGAGATAACGTTCCAGCAGCGCACGCAGGCGCATGTCGTCATCGACCACCAGAATCTTGTAGT
TCTCTTGCATTGTTTGTACTCCCAAAGGTTCGCAACAATTTGTAAGCGTGTATTCTTAAAAAAGCTCACGTTCGTCACCAGCTAAATCT
GGTATGAATTTCAGCCTAAATTGTTACAAAGCATATTAAACAGCAGCTTAAGTATACAATTTATTCGGCGAAACATTATTGATTCTGTT
GATATGATCACGTTATACCCAATGTGCGCATTATCAAACAGACAAAGGGAATCAACGAGATGAAAACGCCCCTGGTTAGGATCCTCTAG
ATTTAAGAAGGAGATATACATATG
>gntR
GAGGCAATTCCGACGTCTAAGAAACCATTATTATCATGACATTAACCTATAAAAATAGGCGTATCACGAGGCCCTTTCGTCTTCACCTC
GATGGATCCGGCAATCTCCATCCATGCGGATAGCGACAGCGAAGTGTTACTGTTCGATCTGCCGCCGGTTTAAAACTCAACGGCATCTT
TGAAGCCTGCCTTTTTGCAGGCTTCCTCGCTCGCTCCGGTCGCGTGCTGATGTACGCTTCCGGGGTTTTGCCGCGTCGTTAGCACAATG
CGATGCAATTGATTTAGTGTATATCTAATAATTGTAAACAAGCACTGTTTTCCCCGGGAAGGTTCTGATGTGTCCGTGTTAAACTAAGA
GAATCTATCTCTTTTGTACCTTCAGGACGATGAAAAAGAAAAGACCCGTACTTCAGGATGTGGCTGACCGTGTAGGCGTGACCAAAATG
ACGGTCAGCCCTCGAGAGATCCTCTAGATTTAAGAAGGAGATATACATATG
>ubiF
GAGGCAATTCCGACGTCTAAGAAACCATTATTATCATGACATTAACCTATAAAAATAGGCGTATCACGAGGCCCTTTCGTCTTCACCTC
GAGGTTGATAGCCGTGGGTGGCATCCAGCAGATCGGCCATCTTCGATGAATCGTACTCGTTCATCTGACAGCCCCAGGTTTTAATATGG
AGTTTTTTGGTCATCGACTTGCTCTTGCGAAATAGTAGCCAGGAATGCAGGGCGCATAGTGTAATGCTTTGCTGCCGTTGTGACCAGTA
TGAGCGTTATCAGCCCTTAGGGGTAAAAATCCTGTAAACTTAAAGCATATTGCTAACAGGATGATTGACCATGACAAATCAACCAACGG
AAATTGCCATTGTCGGCGGAGGAATGGTCGGCGGCGCACTGGCGCTGGGGCTGGCACAGCACGGATTTGCGGTAACGGTGGGATCCTCT
AGATTTAAGAAGGAGATATACATATG
>ubiG
GAGGCAATTCCGACGTCTAAGAAACCATTATTATCATGACATTAACCTATAAAAATAGGCGTATCACGAGGCCCTTTCGTCTTCACCTC
GAGACAATGACCGACATCGCATAATCCAGATAGGAGCTCTTCAGCTCTTCCTCAATGTTGACCGGTGTAATTTCTCTCGCAAGGTCGCT
CATCTAACCGCTATCCCTCTACTGTATCCCGGATTCAAAGGTCGCAAATTATAACACAGCCGCGCAGTTTGAGGTAAACCTATACGCTT
TATTCACATCCAATGCCTGATATACTCGTTTGTCTTGCCAATTACGGAGTAGAAGTGCCAATGAATGCCGAAAAATCGCCGGTAAACCA
TAACGGGATCCTCTAGATTTAAGAAGGAGATATACATATG
>ubiE
GAGGCAATTCCGACGTCTAAGAAACCATTATTATCATGACATTAACCTATAAAAATAGGCGTATCACGAGGCCCTTTCGTCTTCACCTC
GAGCGGTGAGCCAGGATGAAGAGTATCGACTTCGGTCGGTTCCGGAGCAGCCGAATGATGAAGCTTATCAACGCGATGATGAATATAAT
CAGCAGTCGCGCTAGCCCATTGGGAGTAGTTAAGCCGGGTAGAAATCTAGGGCATCGACGCCCAATCTGTTACACTTCTGGAACAATTT
```

TTTGATGAGCAGGCATTGAGATGGTGGATAAGTCACAAGAAACGACGCACTTTGGTTTTCAGACCGTCGCGAAGGAACAAAAAGCGGAT
ATGGTCGCCCACGTTTTGGATCCTCTAGATTTAAGAAGGAGATATACATATG
>ubiC
GAGGCAATTCCGACGTCTAAGAAACCATTATTATCATGACATTAACCTATAAAAATAGGCGTATCACGAGGCCCTTTCGTCTTCACCTC
GAGTTCAGCTCCAAATCTCATGGTAAATATGCGCTTTTCCCCATTTTTACTGCGAGTTGGCGAGCTCATCGTATAATGAATAAGGGTGT
TTAAGTAAAGGAAAACATCACCGTTCCTGGCATCCTGGACGGTGATGCCCCTACGGTTGCCCTCGCCAGCACGGGCATCGGTAAAGCGT
AAGGTTCAACATCGTTTTACCACTTCATGCGATTGTTGCGTTTTTGTTGCGTATTAGATCACTTAATTTGCTTTACATCTCCCGTAAAC
ACTTTTCTGCGATACAATGCCTTTACGGGATCCTCTAGATTTAAGAAGGAGATATACATATG
>gntP
GAGGCAATTCCGACGTCTAAGAAACCATTATTATCATGACATTAACCTATAAAAATAGGCGTATCACGAGGCCCTTTCGTCTTCACCTC
GATGGATCCGGTAACCACGCCAGTTGCGCCCGCCTGACGGACATCAGCTAAAGAAACCGGATCGTTTGGGCCGTACCAGCGCCAGGTCT
GTTCCATGATTTTTCCTCTTTCACATAGTGGGTTCCACAGCAAAGAGACGCGGAACCCTGGAGTTGTTTAGGTATCCGGATGAACAACC
AGACCGCAGCCGTGGCGTTGCATCGGATGAGATGGCGTATAAGTTCTACCTTTTGGTAAGACATGTCAACCCGAATTCTGAAATTGGTT
AACCACATCACAAGAATTTCACTCTCATGGTTGTTGCTGCAAATTGCAGCAAAGGGTAATCGATGGTTATTGATCAGAGGATGTGACAT
TCATCGCAACAATGGTTGACCAATTTACATAACATATCGCCCAAAATAACAACGGTTCAACCAGGTAAGGTAAAACGTTATGCATGTGC
TTAACATTCTCTGGGTGGTATTCGGCATTGGTCTGATGCTGGTACTGAATTTGAAGTTCAAAATCAATTCAATGGTGGCTTTGTTGGCT
CGAGAGATCCTCTAGATTTAAGAAGGAGATATACATATG
>parC
GAGGCAATTCCGACGTCTAAGAAACCATTATTATCATGACATTAACCTATAAAAATAGGCGTATCACGAGGCCCTTTCGTCTTCACCTC
GATGGATCCACCCTACGTTGGCGGTTTTCCGCTGCATAATCCCCAGGATTATGTCTACAGCAAAGAGTTGTATATCAAGGCACATTGAT
GCCTTTATCGCCGTCGGAGCACGTCCGGCGGCGCTATTCACGTTGATCTCCTGTGACTCGACGCGGCAGATAATGTAGTATCTCCGGCA
ATATTGCCCCTTTGAAGGCTGGCGAATAAGTTGAGGAATCAGAATTAATGAGCGATATGGCAGAGCGCCTTGCGCTACATGAATTTACG
GAAAACGCCTACTTAAACTACTCCATGTACGTGATCATGGACCGTGCGTTGCTCGAGAGATCCTCTAGATTTAAGAAGGAGATATACAT
ATG
>bglA
GAGGCAATTCCGACGTCTAAGAAACCATTATTATCATGACATTAACCTATAAAAATAGGCGTATCACGAGGCCCTTTCGTCTTCACCTC
GAGTGCGATTCAGACTCGTCGCGGATGGTGATGGTTTTACGCCCAGCCAGAATGTCATCCTGGAAACGTTGAAAAAAAGTGATGTCGTT
TGGCTGCATGGTACTATTTCCTGTAAGAATTGACTCATCTGGAGCCTATGATAGTGAAAAAACTCACCTTACCGAAAGATTTCTTATGG
GGCGGCGCAGTTGCCGCTCATCAGGTCGAAGGCGGCTGGAACAAAGGCGGAAAAGGGGGATCCTCTAGATTTAAGAAGGAGATATACAT
ATG
>ascG
GAGGCAATTCCGACGTCTAAGAAACCATTATTATCATGACATTAACCTATAAAAATAGGCGTATCACGAGGCCCTTTCGTCTTCACCTC
GATGGATCCAGTGCCGCTATCACCGAGCGTGCCAGCGCCGCATAATTTTTGGCCATCGGATTTTATCCTGTTATCAGTAGGGTAGATGT
AGTTGTTATCAGTCGACGTGACGGAAAAATGCCGATGCTGAAACGGAATTAAGAAACCGGTTTCACAAAATCATGAATAGTTGTCACTA
AATAAGCAAGAGAGTGAATTGTTCATTTTTTATATTTGTGAAACCGGTCACCTGAATATTTTTCCCGCCGATTGTGCAAGGACAATCGG
CAAACGGTGCAGTACACTGCGTTGATTGCGTGTAAAGGGATGAAAATGATGACGACGATGCTGGAAGTGGCGAAGCGCGCCGGGGTTTC
AAAAGCGACCGTTTCCCGCGTGCTTTCAGGTAACTCGAGAGATCCTCTAGATTTAAGAAGGAGATATACATATG
>b0165
GAGGCAATTCCGACGTCTAAGAAACCATTATTATCATGACATTAACCTATAAAAATAGGCGTATCACGAGGCCCTTTCGTCTTCACCTC
GAGTAGCCTGCAAAAATCGGCGGCGTGAAATCATCGCTCATCCCTGCAAAAAAATCGGGCAGCGTCGTGCTGCCCGTGTGCATACTTTT
AGTCGATGGTACGCAGCAGTTCGTTAATGCCGACTTTGCCGCGAGTTTTGGATCCATCGAGGTGAAGACGAAAGGGCCTACGGGGATCT
ACACCTTATTCTCGCTTCTACAGCAACAGGATCCTCTAGATTTAAGAAGGAGATATACATATG
>selD
GAGGCAATTCCGACGTCTAAGAAACCATTATTATCATGACATTAACCTATAAAAATAGGCGTATCACGAGGCCCTTTCGTCTTCACCTC
GATGGATCCGATTAACGTCCCGGACCCGACGCCGTTTGTAACTTATTTCTGATAATTGTCGCGAAACTGTCTGGATTGTGAGCAAAGGC
GCAGGAATTCAGACACTCTCACTTATCACTTCACGGAATGAGGGTTACCATACCCGGATTGCATTGACAGGAGATGTCCATGAGCGAGA
ACTCGATTCGTTTGACCCAATACAGCCACGGAGCTGGTTGCGGCTGTAAAATTTCCCCAAAAGTGTTGGAAACCATCCTGCCTCGAGAG
ATCCTCTAGATTTAAGAAGGAGATATACATATG
>alsR
GAGGCAATTCCGACGTCTAAGAAACCATTATTATCATGACATTAACCTATAAAAATAGGCGTATCACGAGGCCCTTTCGTCTTCACCTC
GATGGATCCCCGACATGATCACAGCCAAATGCAATCTTTTTCATTCATCTTCCCCGTCCGTGCAAAACTTCACAATCAAAAAACATTTG
TGATGTTAATGAATTAAAAACACCCAAAATCGATGAATTACGCCTTCCTGATCTGTTTTACAGAGTTACATCTGACAGCGCAATAGCGT
TAAAGACACTCACCATCCAGCAACATCTATCATCTAAAAAACCAGAAAAACAAATAACATCATGTTTTTAAACTAATTAAATGAAATAA
AATTTTAAGCCACTCGCCATTGTTCACAATAAAATAAACTTTATAAATTTTATTTTTTTGTGAAGTCGCCAGCATCTTTTCTGTTCTTG
CTGTGGTGATATAGTGGCGTCTTCAATTCAAGGACAAGAGAACGTGATGAGCCAGTCAGAGTTTGATTCAGCGCTTCCGAACCTCGAGA
GATCCTCTAGATTTAAGAAGGAGATATACATATG
>selA
GAGGCAATTCCGACGTCTAAGAAACCATTATTATCATGACATTAACCTATAAAAATAGGCGTATCACGAGGCCCTTTCGTCTTCACCTC
GATGGATCCAAAGTTTTGCTCGCACCGAACCGCCAAAGGCTTGATGCGCGATATGTCCTCCTGACCCATCTCACGTTACAATCCGTGGT
TATGTTAAACGCCCTTCTCCGTGTGAGAGGGCCTTGATCAGCCAGGTTTCCTATGACAACCGAAACGCGTTCCCTCTATAGTCAACTTC
CGGCTATTGATCGCTTATTGCGCGATAGCTCCTTCCTTTCTTTGCGCTCGAGAGATCCTCTAGATTTAAGAAGGAGATATACATATG
>selC

```
GAGGCAATTCCGACGTCTAAGAAACCATTATTATCATGACATTAACCTATAAAAATAGGCGTATCACGAGGCCCTTTCGTCTTCACCTC
GAGCGAAATTTACCCCAGCGAGAGCGCGTTCGATCGGCCAACAACCCCATGCAAGGATCGGAAATCGCATCCAGTGCACGAGCGACCAA
AAACATGGTTCCGACAAAACCGGCAGGAATGCCAAAAATATCGGTATAAAAGAACATCATATATAACATTACGTTATCGAAAATAATGT
GGCTGGCGGCGTCTCCCATGCCATAACCAATTTTCTCTTTAACGGACAACACTTCACTCTTCATCTGCTTTTCCTTGATGCTATAGGGG
TGCTGAGACCGATAACGCATTTTGTAAACCATCCCTGTGAGAGCGAATATTCCGATATCTGGTTATTAATTTATGATTCTTGTTTTATG
TGATCGTGGTAGCGTTAATTCCGGGATCCTCTAGATTTAAGAAGGAGATATACATATG
>alsI
GAGGCAATTCCGACGTCTAAGAAACCATTATTATCATGACATTAACCTATAAAAATAGGCGTATCACGAGGCCCTTTCGTCTTCACCTC
GAGGTTCGGAAGCGCTGAATCAAACTCTGACTGGCTCATCACGTTCTCTTGTCCTTGAATTGAAGACGCCACTATATCACCACAGCAAG
AACAGAAAAGATGCTGGCGACTTCACAAAAAAATAAAATTTATAAAGTTTATTTTATTGTGAACAATGGCGAGTGGCTTAAAATTTTAT
TTCATTTAATTAGTTTAAAAACATGATGTTATTTGTTTTTCTGGTTTTTAGATGATAGATGTTGCTGGATGGTGAGTGTCTTTAACGC
TATTGCGCTGTCAGATGTAACTCTGTAAAACAGATCAGGAAGGCGTAATTCATCGATTTTGGGTGTTTTTAATTCATTAACATCACAAA
TGTTTTTTGATTGTGAAGTTTTGCACGGACGGGGAAGATGAATGAAAAAGATTGCATTTGGCTGTGATCATGTCGGGGATCCTCTAGAT
TTAAGAAGGAGATATACATATG
>alsB
GAGGCAATTCCGACGTCTAAGAAACCATTATTATCATGACATTAACCTATAAAAATAGGCGTATCACGAGGCCCTTTCGTCTTCACCTC
GAGACAGCATATTCGGCGGCAGCAAAAGCGCTGGTTGACAACATTAAGCCCACGAGTGTGCCGCTGAAATATTTCAGATATTTATTCAT
AATAATGTTCTCACGGTAAGGGGACAGGTAACTCCGCGATGGCGGAATTACCTTTTTCTTATTTCAGCGCGCCTGGTGAGAAGAAATCA
ACAATTGCGCCGGTTTTTTGCATATTAATATTAGCTTGTTCAATGTTGAGCTGGGCGACGGATCCTCTAGATTTAAGAAGGAGATATAC
ATATG
>ymgE
GAGGCAATTCCGACGTCTAAGAAACCATTATTATCATGACATTAACCTATAAAAATAGGCGTATCACGAGGCCCTTTCGTCTTCACCTC
GAGAGGAATTGCGGCTTTGTGCAAATCGCGTAACACGCCCAGGACGGCTAACGGATTTTGTTTCAGGAACTGCTCATGGTAATGACTCA
CGCGGAAAAACTCCTGACTGACAAACTGTCTTTGGTTTATCGGTCACAGTTAACAAAACTTAATACAAATGCGTGAATATTTTTTACAT
GTTGTTCTTAAATCAGCCGACATACGCCCAGCATCGATCCCGCCCTTGCCTATACTTAGAGCGTTGATGTAAGCATTTCTTGCGTCCGA
TTCATCGAAACGAGGGCCTGAACATGGGAATTATTGCCTGGATTATTTTTGACCTGATAGCCGGCATTATCGCCAAGCTAATCATGCCG
GGGGATCCTCTAGATTTAAGAAGGAGATATACATATG
>alsA
GAGGCAATTCCGACGTCTAAGAAACCATTATTATCATGACATTAACCTATAAAAATAGGCGTATCACGAGGCCCTTTCGTCTTCACCTC
GATGGATCCGCTGGATAAAGCACCGGAATTTAAACTGGTCGATTCAATCCTGGTCACTCAATAAAACTGAATGATGCCGGATGATGATG
CTAATGAAGTGTCTTATCCGGCCAACAAAATAACCTGAACGTAGACCTGATAAATGTAGCCATCAGGCAATAATACGATATTTCGTGCA
GAGAGGTTATTTATGGCCACGCCATATATATCGATGGCGGGGATCGGCAAGTCCTTTGGTCCGGTTCACGCATTAAAGTCGGTCTCGAG
AGATCCTCTAGATTTAAGAAGGAGATATACATATG
>yeeI
GAGGCAATTCCGACGTCTAAGAAACCATTATTATCATGACATTAACCTATAAAAATAGGCGTATCACGAGGCCCTTTCGTCTTCACCTC
GAGAAAGTGGCGGAGAGAGGGGGATTTGAACCCCCGGTAGAGTTGCCCCTACTCCGGTTTTCGAGACCGGTCCGTTCAGCCGCTCCGGC
ATCTCTCCGTTCAGATGGTTGCCATGATGCCAGGAAATTTGGCATTTTAACAGTCCCTGTCCGTGCAATTTTGTTCAAGTGACGAGTTT
GCGAGCAAAACGATGATTAAGTGGCCCTGGAAAGTACAAGAATCAGCACATCAAACTGCCCTTCCCTGGGATCCTCTAGATTTAAGAAG
GAGATATACATATG
>galU
GAGGCAATTCCGACGTCTAAGAAACCATTATTATCATGACATTAACCTATAAAAATAGGCGTATCACGAGGCCCTTTCGTCTTCACCTC
GAGGAATCAATTGAGCCAGCGATGCGATGCCTGGCAATGCCAAATATGGGGAACCGGTGGTCGACTGCGCTTGATGTTGTCTGCAGAAT
GAGCAAACGATAACGCGGGCTAAATTTGCATTACCTGCTAATGTCGGCTGGTGGTACTATCGTCGCCATTCGTATAAGTAATTGTCTTA
ATTATGCTAACTCGCCTCCTTTTCAGAACTTAGCCCCTTCGGGGTGCTGATATACTGGGATGCGATACAGAAATATGAACACGTTCAAA
ACACGAACAGTCCAGGAGAATTTAAATGGCTGCCATTAATACGAAAGTCAAAAAAGCCGTTATCCCCGTTGCGGGATTAGGAACCAGGA
TGTTGCCGGCGACGAAAGCCATCCCGAAAGAGATGCGGATCCTCTAGATTTAAGAAGGAGATATACATATG
>yeeJ
GAGGCAATTCCGACGTCTAAGAAACCATTATTATCATGACATTAACCTATAAAAATAGGCGTATCACGAGGCCCTTTCGTCTTCACCTC
GAGATATTCGCCCCGTTCACACGATTCCTCTGTAGTTCAGTCGGTAGAACGGCGGACTGTTAATCCGTATGTCACTGGTTCGAGTCCAG
TCAGAGGAGCCAAATTCTAAAAATTCGCTTTTTAGCGCAATGTCACTGACCTTAGTTGAACATTGTTTTTAACGGATAGCGGGTTTT
TAACATCTTAAGCGCCCTCGACCTTTATGGTTGAGGGCGTTTTGCTATGAACGCCATCACCATTTTCCCCTCGATTATAAAACTTGAGT
TATTCAGTAGTCTCCCCTCTTGCAACTCACACCCAAAACTGCCTAACGAAAAGTTATTAATTTTCAATCATATTGCTATCAGTATTTAC
ATTTTTTCGCTGTGCTAGAAAGGGCGCATTTATGTTAGCTCGTTCAGGGAAGGTAAGCATGGCGGATCCTCTAGATTTAAGAAGGAGAT
ATACATATG
>galP
GAGGCAATTCCGACGTCTAAGAAACCATTATTATCATGACATTAACCTATAAAAATAGGCGTATCACGAGGCCCTTTCGTCTTCACCTC
GAGACCGCAGCATACGGTCACTTTGGTCGTGAACATTTCCCGTGGGAAAAAACCGACAAAGCGCAGCTGCTGCGCGATGCTGCCGGTCT
GAAGTAATCTTTCTTCACCTGCGTTCAAAGGCCAGCCTCGCGCTGGCCTTTTTCTTTTGGATAGGCGTTCACGCCGCATCCGGCAAAAA
AACCGCCCGCACAATAACATCATTCTTCCTGATCACGTTTCACCGCAGATTATCATCACAACTGAAACCGATTACACCAACCACAACAG
ACAAAGATTTGTAATATTTTCATATTATTATTCGGTTTTCACAGTTGTTACATTTCTTTTCAGTAAAGTCTTAATTGCAGATAACAGCG
TTTAATCTATGATGATATAACTCAATTATTTTCATGCACTTAAATCATAACTAAGATAAATGTTAGTGTAAGCGATTACACTGATGTGA
TTTGCTTCACATCTTTTTACGTCGTACTCACCTATCTTAATTCACAATAAAAAATAACCATATTGGAGGGCATCATGCCTGACGCTAAA
AAACAGGGGCGGTCAAACAAGGCAATGACGTTTTTCGTCTGCTTCCTTGCCGGATCCTCTAGATTTAAGAAGGAGATATACATATG
```

```
>galR
GAGGCAATTCCGACGTCTAAGAAACCATTATTATCATGACATTAACCTATAAAAATAGGCGTATCACGAGGCCCTTTCGTCTTCACCTC
GAGTTCAGTGCCTGGGTGTCACCCGTAACGCGAACGCGATACAAAACACGGCACAAATTTCGAAAAAAGCTAAAAAGCATTCCAACTCC
CTTTGCTCTGATTCAGTAAAAGCGAATGGAGGGAGATTACACGAGATAAAGAACGCGAGCGACAGTAAATTAGGTGCGAAAAAAAACCT
GCGCATCCGCGCAGGTTGGTGCAAGAGACAGGGTACGAAGAGCGTACCGAATAATCTCACCAATCAATACCTCTGGGATCTTGATTGTG
GTCTGCACGACGACTCTTCGCCAGCGAGAAAACGCAAAGGAATGAAGGGAAATGCAACGAGGTGTGTAAATTGTCGGTTACTGTTACAG
ATTGATGACCGGCAAAAAAAACCTGCGCATCTGCGCAGGCTGGTGTAATTCATGTGCTCAACCCGAAGTTGACTTCACCTATCAATACC
TCTGGGATCACCACTTTAGCAACCTGAAGCCAAACGCCACCAGCGGTCAATCGCAACAGCGTTTCGCAAAGTGTAACCAAAGGTTTGCA
ATCTCTTTTTTTGTCTTACTTATTTGCGTTTGGCTCACATTCCCACGATGAAAACACGCCACCCCTTGAACCAACGGGCGTTTTCCGTA
ACACTGAAAGAATGTAAGCGTTTACCCACTAAGGTATTTTCATGGCGACCATAAAGGATGTAGCCCGACTGGCAGGCGTTTCAGTCGCC
ACCGTTTCCCGCGTCATTAATAATTCACCCAAAGCCAGCGAAGCTTCGGATCCTCTAGATTTAAGAAGGAGATATACATATG
>galS
GAGGCAATTCCGACGTCTAAGAAACCATTATTATCATGACATTAACCTATAAAAATAGGCGTATCACGAGGCCCTTTCGTCTTCACCTC
GATGGATCCCCGGACCGGCAATATCTAAAACATCAAGATAACGATCTGGATCACATTCGTTAACAAAACGGCTGTAACCGTTTCCATTG
CTGTGACTCGATTCACGAAGTCCTGTATTCAGTGCTGACAAAATAGCCGCCAGCAAGCAGTCATTTACTGCAATCTCATAACAGGTAGT
GAATATGATCACCATTCGTGATGTAGCGCGTCAGGCTGGCGTCTCTGTGGCAACGGTTTCCCGGGTGCTCAATACTCGAGAGATCCTCT
AGATTTAAGAAGGAGATATACATATG
>yeeA
GAGGCAATTCCGACGTCTAAGAAACCATTATTATCATGACATTAACCTATAAAAATAGGCGTATCACGAGGCCCTTTCGTCTTCACCTC
GATGGATCCAGCCCTGCTTCGAGGTTTATTTGAACAATGGCGCGGAAGATGGGTACTGGGATATCGAAATGTATGTTGCGGTGCAGCCA
AAACATCACTAATTCATCTCAGGGCGGTGTGTTAACGCGATGACCACTCTTTTTTTTGAAAGCGAAAAGAGTAAGATGCGCCTTTCAAT
TTTTTCGCTCCTGCCGGGAAATTACACTGTTCCCGGTTTGTCCGTCGGATAATTCAGAGGCGCGCCTTCTGGCCGACAGATGAGTTATG
AGCGCTTTTAATCTCATTACGGAGTTTCTGCGTGCGTGCCGATAAGTCATTAAGCCCGTTTGAAATCCGGGTATACCGCCATTACCGCA
TTGTGCATGGTACTCGAGAGATCCTCTAGATTTAAGAAGGAGATATACATATG
>yeeE
GAGGCAATTCCGACGTCTAAGAAACCATTATTATCATGACATTAACCTATAAAAATAGGCGTATCACGAGGCCCTTTCGTCTTCACCTC
GATGGATCCTCTGGGCTGCTATCGGTCTGATTTACCTGGCTTGCGTCACCAAGAGCTTCCGTAATCCGGTTCCGCAGTACGAAGACGTA
GCATAATCATCTTTTCTAATAATAAACCGGAGTCGAATGAACATTTGGCTCCGGTTTTTTTATTCTTTTGTATCGAACTCCTCATTAAT
AACCATTTGTTCTAATTTAGAGAATTTGGTTATTGGTCTGCATTCCACCAGTCATGAATAATTTCCTCATTAACATTCATCTGGTTAAT
TAATTTATGTTTTCAATGATATTAAGCGGGCTAATTTGTGGTGCTCTGCTGGGATTTGTCATGCAGCGTGGGCGTTTTTGCCTGACAGC
TCGAGAGATCCTCTAGATTTAAGAAGGAGATATACATATG
>lit
GAGGCAATTCCGACGTCTAAGAAACCATTATTATCATGACATTAACCTATAAAAATAGGCGTATCACGAGGCCCTTTCGTCTTCACCTC
GAGCCAAAGATATCGCACCCGAAGTAGTTTGGCTGCATTGTTATGTAGGTCTAACGCACCGCTAATTAAATATGCAAAAATCGCAAAAG
CACAAGGTAATATACCAAAAGATAGTGCCCAATTAATAAAGCTCTCATGATCTTTAATAGGTGGGACATACTTGGTTGGTAATGTTATT
AAAGGGATGTATTCGTAAATAACTAGAAGTATGCAATATATTAGCGTATTTAAGAAAGTTGCTTTATGAAGTTTAGGCAACATTTCATT
ATAATTTTTGGGGGGTTCGAACATATTATTCACCTGAGTCACGCTTTAGGATAGTGTAATGGTAATATTTAATTAAGTGTCATTATATA
CTTTTCAGTAGGTTAGTTACAATTTTTTGTATCTGTTCAGGCTGACCTAGCTTCGCTGACAGACAATATTGTGATCAGTAGCACGTATC
GAGGAGGAGTAGCGCTACAAATTTGACGCTGGGTGAGAATCTGAAATTGATAGAAATGAAATAATGAAATAATGAAATGATGAAATGAT
GAAATGATGAAATGATGAAATGATGAAATGATGACAGAGTGTCCAGTGGGCACGGATGGTGTCTTACGACATGCTTACCTTAATCGTTC
CCAGTGTGCCGGATCCTCTAGATTTAAGAAGGAGATATACATATG
>galE
GAGGCAATTCCGACGTCTAAGAAACCATTATTATCATGACATTAACCTATAAAAATAGGCGTATCACGAGGCCCTTTCGTCTTCACCTC
GATGGATCCGATGCGCCTGCCTGTATTACCCATCGTCTTGAGTTCGTGCCGGACGGTGGACTCTATCGCTATGTGCTGACAAAAATATA
TTGAGTCGGTAGTGCTGACCTTGCCGGAGGCGGCCTTAGCACCCTCTCCGGCCAACGGTTCGACGCATGCAGGCATGAAACCGCGTCTT
TTTTCAGATAAAAAGCGCAATCATTCATAAACCCTCTGTTTTATAATCACTTAATCGCGCATAAAAAACGGCTAAATTCTTGTGTAAAC
GATTCCACTAATTTATTCCATGTCACACTTTTCGCATCTTTGTTATGCTATGGTTATTTCATACCATAAGCCTAATGGAGCGAATTATG
AGAGTTCTGGTTACCGGTGGTAGCGGTTACATTGGAAGTCATACCTGTGTGCAATTACTGCAAAACGGTCATGATGCTCGAGAGATCCT
CTAGATTTAAGAAGGAGATATACATATG
>yeeR
GAGGCAATTCCGACGTCTAAGAAACCATTATTATCATGACATTAACCTATAAAAATAGGCGTATCACGAGGCCCTTTCGTCTTCACCTC
GAGGGTTATGCCCACAGCGTCAGCGGCAGCAGCGCTGAAGGGTATAACGGTCAGGCCACACTGAATGTGACCTTCTGACAGAACCATCG
CCTCTCTGTGGTCCCGGTCATCATGACCGGGACCCGGACCGGCGCAACGGATCTTCAACGCCACATTCGCTGGCATTAACAATAACATG
ATATTCATCACGGAGTGACTATGTTACAGATAGTCGGCGCGCTGATCCTGCTGATCGCAGGATTTGCCATTCTTCGCCTTTTGTTCAGA
GCATTAATCAGCACGGCTTCTGCGGATCCCGCCTCTAGATTTAAGAAGGAGATATACATATG
>yeeU
GAGGCAATTCCGACGTCTAAGAAACCATTATTATCATGACATTAACCTATAAAAATAGGCGTATCACGAGGCCCTTTCGTCTTCACCTC
GAGTGCAGGATATTCCCGGTGTGCTGGCCGTGTTTGCTGAACGCCGTAAGGACAGTTTTGGTCCGTATGTCCGGCTGATGAGCGTCACC
CTGAACTGAGTGGGAATTCTGATGAGCAGAATTATCGCCACTACCGGACCATTCTTAGCCGATTTTCTGTAAGGATTTTATCGTGTCAG
ACACACTCCCCGGGACAACACTTCCCGACGACAATCACGACCGCCCCTGGTGGGGGCTGCCCTGCACCGTGACGCCCTGTTTGGATCCT
CTAGATTTAAGAAGGAGATATACATATG
>yeeV
```

```
GAGGCAATTCCGACGTCTAAGAAACCATTATTATCATGACATTAACCTATAAAAATAGGCGTATCACGAGGCCCTTTCGTCTTCACCTC
GAGCTGACCTGCAAAGCCGATACCCTCAGCAGTTGTGATTACGTTTATCTGGCTGTTTATCCGACGCCCGAAATGAAAAATTAACTCTC
CAGAATAGCCTTCTGCTACGGCCTGGTGTTTTCACCACGCCACTTTTCCATTTTTATATCTGCATATCAGGAAAATCTTCAGTATGAAA
ACATTACCTGTATTACCCGGGCAGGCGGCCAGTTCTCGCCCGTCTCCTGTTGAAATCTGGCAGATACTGCTGTCCCGGATCCTCTAGAT
TTAAGAAGGAGATATACATATG
>hipB
GAGGCAATTCCGACGTCTAAGAAACCATTATTATCATGACATTAACCTATAAAAATAGGCGTATCACGAGGCCCTTTCGTCTTCACCTC
GATGGATCCAACCTTAGCCTGTGGGGGAATGTCGGTGTGCAACTAGGTGATAAAGGCTATAGCGATACTCAGGGCATGCTGGGAGTGAA
ATATAGCTGGTAAACCGTATAAGCCGCATGTCGAGATGGCATGCGGCTTAATATTGCCGACTTCAAACGGCGCATCAACGCCTTATTTA
AATCCTCCTTTTTATCCGCGATCGCGGATATCGCAGCGTTTATCCCGTAGAGCGGATAAGATGTGTTTCCAGATTGACTTATCCTCACT
AAAGGATAAAACTTATAATATCCCCTTAAGCGGATAAACTTGCTGTGGACGTATGACATGATGAGCTTTCAGAAGATCTATAGCCCAAC
GCAATTGGCGAATGCAATGAAACTGGTTCGCCAGCAAAATGGCTGGACGCAGAGCTCGAGAGATCCTCTAGATTTAAGAAGGAGATATA
CATATG
>rho
GAGGCAATTCCGACGTCTAAGAAACCATTATTATCATGACATTAACCTATAAAAATAGGCGTATCACGAGGCCCTTTCGTCTTCACCTC
GAGGGCTCGTCACTCAATCCGTCTTGTCGTTTCAGTTCTGCGTACTCTCCTGTGACCAGGCAGCGAAAAGACATGAGTCGATGACCGTA
AACAGGCATGGATGATCCTGCCATACCATTCACAACATTAAGTTCGAGATTTACCCCAAGTTTAAGAACTCACACCACTATGAATCTTA
CCGAATTAAAGAATACGCCGGTTTCTGAGCTGATCACTCTCGGCGAAAATATGGGGCTGGAAAACCTGGCTCGTATGCGGGATCCTCTA
GATTTAAGAAGGAGATATACATATG
>msrA
GAGGCAATTCCGACGTCTAAGAAACCATTATTATCATGACATTAACCTATAAAAATAGGCGTATCACGAGGCCCTTTCGTCTTCACCTC
GATGGATCCCCGATAACCCCTCGACCTGTAGACGGACGTTCGCGGCGACGGCAGATCCGCTTAAGCAGAGTAAGCTTACACAGCATAAC
TGTCGGATATAGCGCACATTTTCTCCTGAATATCCTTTTTTCCTGCCCCTGGAACGCCGTTAAACGGCTTAACAAAAATCCAGTAATAT
GGATTAAAAAGCAGACTAAACCCCAAATATTTCTTATGTTTTACTTTAGACCTATTCACGGTGGTTATTGTGTGCAAATACGCCTCTTG
TTACAACCTTAACCCCAATGACCGATTTTCGGGAGAGCGACACCATGAGTTTATTTGATAAAAAGCATCTGGTTTCCCCCGCCGATGCC
CTGCCTGGACGTAACACCCCGATGCCTCGAGAGATCCTCTAGATTTAAGAAGGAGATATACATATG
>hofM
GAGGCAATTCCGACGTCTAAGAAACCATTATTATCATGACATTAACCTATAAAAATAGGCGTATCACGAGGCCCTTTCGTCTTCACCTC
GATGGATCCTCCCAGCAGAATGCAACAGACTGCAAGGATCAAAAAATACTTTACGAACTTCACTGGAAATTTCCCATTTAGTTTCATTT
GGGCAGTTTATAAACAAACGCGCGGTAGTATAAAGGCAAGCCAGACGCATTGATATACCCGTCAGAGTGACGGGTGATAAGGAGATCAT
CACAATGGCATTTAAGATCTGGCAAATTGGTTTGCATTTACAACAGCAAGAAGCGGTAGCGGTTGCTCGAGAGATCCTCTAGATTTAAG
AAGGAGATATACATATG
>yhaH
GAGGCAATTCCGACGTCTAAGAAACCATTATTATCATGACATTAACCTATAAAAATAGGCGTATCACGAGGCCCTTTCGTCTTCACCTC
GAGTTCCGCAGCCATAAGACCATCAACCCTACGGGGATTATTTCAATTGGTCCGTGGCAGGATCTCGATGAACCGCATGGACGAGATGT
TCGCTTCGGTTAAAAAAAGGCACCTTACGAGGTGCCTCTGATAATTAGCTAAGTGCATGTTGTTCATGTCGGATGTGGCATGAACGCCT
TATCTGACATACAAAACATTGTACATTCAATAGATTGCATAACCCGCGCAGACCAGATAAGCGTAGCGCATCTGGCGATTTTGCGCTTG
TCATCAGTCTTAGCGCGCCTTCACGGCACTTTATTAATTCCTTGAACGAATATTTACTGCCATTTCATTCACGTCTATTCTTAATTTGC
TGCTTTAAAAACAAGTGATTGAGCAAAATTGAGGCAAAAATGGACTGGTATCTGAAAGTACTAAAAAATTATGTCGGTTTCCGGGGTCG
TGCGCGGCGCAAAGAGTACTGGGGATCCTCTAGATTTAAGAAGGAGATATACATATG
>yhaJ
GAGGCAATTCCGACGTCTAAGAAACCATTATTATCATGACATTAACCTATAAAAATAGGCGTATCACGAGGCCCTTTCGTCTTCACCTC
GATGGATCCGCCTGCAACCATCCGTAGTCTGCTTGTCCACACTGCCTGGCAGTTCGGGTAGTAATCATAATTCTTGACCTCTCTTTACT
TCCTTTTATGGTAAGGGGCTGGACGCTACATTGTTAGCCAGATATTCTGCCCGGTATGTTCAAATTTCCTGAATGAGAACGAAATGGCC
AAAGAAAGGGCATTAACGCTGGAAGCACTACGGGTTATGGATGCGATCGATCGCCGGGGCAGTTTTGCGGCGGCGGCGGATGAGCTGGG
ACGCGTCTCGAGAGATCCTCTAGATTTAAGAAGGAGATATACATATG
>yhaK
GAGGCAATTCCGACGTCTAAGAAACCATTATTATCATGACATTAACCTATAAAAATAGGCGTATCACGAGGCCCTTTCGTCTTCACCTC
GAGACGCGTCCCAGCTCATCCGCCGCCGCCGCAAAACTGCCCCGGCGATCGATCGCATCCATAACCCGTAGTGCTTCCAGCGTTAATGC
CCTTTCTTTGGCCATTTCGTTCTCATTCAGGAAATTTGAACATACCGGGCAGAATATCTGGCTAACAATGTAGCGTCCAGCCCCTTACC
ATAAAAGGAAGTAAAGAGAGGTCAAGAATTATGATTACTACCCGAACTGCCAGGCAGTGTGGACAAGCAGACTACGGATGGTTGCAGGC
CCGGTATACTGGATCCTCTAGATTTAAGAAGGAGATATACATATG
>cdd
GAGGCAATTCCGACGTCTAAGAAACCATTATTATCATGACATTAACCTATAAAAATAGGCGTATCACGAGGCCCTTTCGTCTTCACCTC
GAGTAGCGCTGGTGTTATGCGGGATAATTACTTCGCTGATCGCACCGTTCCTTTTCCCGATTATTCTGGCAGTAATGGGCTAAAATTTG
CGATGCGTCGCGCATTTTTGATGTATGTTTCACGCGTTGCATAATTAATGAGATTCAGATCACATATAAAGCCACAACGGGTTCGTAAA
CTGTTATCCCATTACATGATTATGAGGCAACGCCATGCATCCACGTTTTCAAACCGCTTTTGCCCAACTTGCGGATAACTTGCAATCTG
CACTGGAACCTATTCTGGCAGACAAGTACTTCCCCGCTTTGTTGACGGATCCTCTAGATTTAAGAAGGAGATATACATATG
>yoaB
GAGGCAATTCCGACGTCTAAGAAACCATTATTATCATGACATTAACCTATAAAAATAGGCGTATCACGAGGCCCTTTCGTCTTCACCTC
GAGGCCATCTGTCGCTGTGGTTCTCGCGGCTTAAAGCCTGGTATCGCTTTCGCCAGCTGACCGTCTGGTGCAAAATCGTCCGTCACACT
ACCCCCTGTTGATTTGAACAGGGATTATGTCAGGATGAGGGCGCTTTCGCCAGTTGAAGTGGTGACGGCGACCTCACATTGTGGCAGTC
```

```
TTTGCAGCACGAAAATGGAAGTATAATGAGGACAAAATGATGACTATCGTTCGTATCGATGCTGAAGCCCGCTGGTCTGATGTAGGATC
CTCTAGATTTAAGAAGGAGATATACATATG
>yoaA
GAGGCAATTCCGACGTCTAAGAAACCATTATTATCATGACATTAACCTATAAAAATAGGCGTATCACGAGGCCCTTTCGTCTTCACCTC
GATGGATCCTCCATTTTCGTGCTGCAAAGACTGCCACAATGTGAGGTCGCCGTCACCACTTCAACTGGCGAAAGCGCCCTCATCCTGAC
ATAATCCCTGTTCAAATCAACAGGGGGTAGTGTGACGGACGATTTTGCACCAGACGGTCAGCTGGCGAAAGCGATACCAGGCTTTAAGC
CGCGAGAACCACAGCGACAGATGGCCTCGAGAGATCCTCTAGATTTAAGAAGGAGATATACATATG
>prpB
GAGGCAATTCCGACGTCTAAGAAACCATTATTATCATGACATTAACCTATAAAAATAGGCGTATCACGAGGCCCTTTCGTCTTCACCTC
GAGAAACAGCTCGAACAGGCGCGTTACAGATACCGTCCAGATAACCGGTTTGTCGTCATTAAGCCGTGGTGGATGTGCCATAGCGCACC
GCAAAGTTAAGAAACCGAATATTGGGTTTAGTCTTGTTTCATAATTGTTGCAATGAAACGCGGTGAAACATTGCCTGAAACGTTAACTG
AAACGCATATTTGCGGATTAGTTCATGACTTTATCTCTAACAAATTGAAATTAAACATTTAATTTTATTAAGGCAATTGTGGCACACCC
CTTGCTTTGTCTTTATCAACGCAAATAACAAGTTGATAACAAAGGATGGGCTATGTCTCTACACTCTCCAGGTAAAGCGTTTCGCGCTG
CACTGACTAAAGAAAATCCATTGCAGATTGTTGGCACCATCAACGCTAGGATCCTCTAGATTTAAGAAGGAGATATACATATG
>prpC
GAGGCAATTCCGACGTCTAAGAAACCATTATTATCATGACATTAACCTATAAAAATAGGCGTATCACGAGGCCCTTTCGTCTTCACCTC
GAGACAACCGATGCCTGATGCGCCGCTGACGTGACTTATCAGGCCTACGGGGTGCACTGAACTGTAGGTCGGATAAGACGCATAGCGTC
GCATCCGACAACCGATGCCTGATGCGCCGCTGACGCGACTTATCAGGCCTACGGGGTGCACTGAACTGTAGGTCGGATAAGACGCATAG
CGTCGCATCCGACAACCGATGCCTGATGCGCCGCTGACGCGACTTATCAGGCCTACGGGGTGAACTGAACTGTAGGTCGGATAAGACGC
ATAGCGTCGCATCCGACAACAATCTCGACCCTACAAATGATAACAATGACGAGGACAACATGAGCGACACAACGATCCTGCAAAACAGT
ACCCATGTCATTAAACCGAAAAAATCTGTGGCACTTTCTGGCGTTCGGATCCTCTAGATTTAAGAAGGAGATATACATATG
>yoaE
GAGGCAATTCCGACGTCTAAGAAACCATTATTATCATGACATTAACCTATAAAAATAGGCGTATCACGAGGCCCTTTCGTCTTCACCTC
GATGGATCCCGTTTTCCTGCTCGCCTAACAGCATTTCTGCCGTTTTAAGCAACTGCTCTGCAGCCCAACCATGTGTGCCTATAACAATA
GCAATGGTCACTTGCTACCTCCTTTATTATCGTTAACACCTCAACGTGCCAGATGTATTTTTGAATCGTCGTCCACAATCGAATCGATT
CAGATAAGGGCTACAACCAAAAACCCCTGCGTTTCGCGATTTATTTTAGATATCGAAAAAATTATTTTATGTGATGAAGATCCGTAATT
TAACTTTCGATTAGCAGAAATTTCGAAAGGTAAAATATCCTTGTCACATTCGTTTGCAAAGGAAGGTAAATCTTTGCCAAATCAGAGGC
GTCTCTGATATGTTTAACTCCCGTTTAACAACCATGGAGTATAGGGCAGTAGCCCGCAGTATGGATCGTCACCGACGTCATTTCAGCAT
CAGGCCTTTTAACGCCTGCCTTTCTGGCACTCTATGCCGCACCTTTCGTTTGCATTTTGTCGTTACGCCTGCATTATTTCTGGCGTCGA
ATAGCTATTCCTTAAGCAGGAGCTTGTCATGGAATTCTTAATGGACCCCTCAATTTGGGCGGGGCTACTCACGCTTGTTGTTCTCGAAA
TTGTGCTGGGTATCGATAACCTGGTCTTCATCGCCATCTCGAGAGATCCTCTAGATTTAAGAAGGAGATATACATATG
>yoaD
GAGGCAATTCCGACGTCTAAGAAACCATTATTATCATGACATTAACCTATAAAAATAGGCGTATCACGAGGCCCTTTCGTCTTCACCTC
GAGGGTAAGGACATGAACGCCAAATACCGCGAAACCTCACGCGGTGGTCTGGCAATCAAAGTCCAGTGTGACTAATACTTCTTACTCGC
CCATCTGCAACGGATGGGCGAATTTATACCCGCTTTCTCGTCTGCTGTAATATTCCCCACTACACTTCCACTGTTGCGTCAGGCGTTTG
TCGCCATACGCTTACAGGGTGGCCCGCATGCAAAAAGCACGGATCCTCTAGATTTAAGAAGGAGATATACATATG
>yoaH
GAGGCAATTCCGACGTCTAAGAAACCATTATTATCATGACATTAACCTATAAAAATAGGCGTATCACGAGGCCCTTTCGTCTTCACCTC
GATGGATCCTAAAATTCAGCGGCGTCCTGACGCCAGAGTAAAGTAATCACAGCGGGAGATAACGTCTTCATCAGTCCTGACTCTACTGG
CTATGTGCCGGGTAGCATGAAATAATTAGCGCCAACAATTTAGCAGGAGTTAACAATGTTTGCAGGTTTACCTTCACTCACCCATGAAC
AGCAGCAAAAAGCTGTCGAGCGGATCCAGGAACTGATGGCACAAGGGATGAGCAGCTCGAGAGATCCTCTAGATTTAAGAAGGAGATAT
ACATATG
>ugd
GAGGCAATTCCGACGTCTAAGAAACCATTATTATCATGACATTAACCTATAAAAATAGGCGTATCACGAGGCCCTTTCGTCTTCACCTC
GATGGATCCACCTGATCCAGGCACAGCGTGACTATTTTGGTGCGCATACTTATAAGCGTATTGATAAAGAAGGTGTGTTCCATACCGAA
TGGCTGGATTAATCTGATTTAACCAACAATAAAATTGAGGCCCGGCGTATATTGCACCGGGCTTTTTTTTGCCAAATATCTTATAGACT
AAATTTCACTGCTTAATATTAACTTAATAAATATCAGCTATTCTTATAAAGAAAATCTGAATTGTTTTTCGTTGCGTTGACCATCGAAC
AACGTAGCGTTAAAACTTTTAGCTCTTATCAGGATGCTAAAAACATCATGATTCACAGTTAAGTTAATTCTGAGAGCATGAAATGAAAA
TCACCATTTCCGGTACTGGCTATGTAGGCTTGTCAAACGGGCTTCTAATCGCACAAAATCATGAGGTTGCTCGAGAGATCCTCTAGATT
TAAGAAGGAGATATACATATG
>ygcB
GAGGCAATTCCGACGTCTAAGAAACCATTATTATCATGACATTAACCTATAAAAATAGGCGTATCACGAGGCCCTTTCGTCTTCACCTC
GATGGATCCGCCCCAGGCGATATTTCTATCAACCTGAGGCCAGCGTTCGAACCCAAACAATTCGAATGTTAGTCTCTTCTTTGCAGGGA
GGCAAGACATGTGTATATCACTGTAATTCGATATTTATGAGCAGCATCGAAAAATAGCCCGCTGATATCATCGATAATACTAAAAAAAC
AGGGAGGCTATTAATGGAACCTTTTAAATATATATGCCATTACTGGGGAAAATCCTCAAAAAGCTTGACGAAAGGAAATGATATTCATC
TGTTAATTTATCATTGCCTTGATGTTGCTGCTGTTGCAGATTGCTGGTGGGATCAATCAGTCGTACTGCAAAATACTTTTTGCCGAAAT
GAAATGCTATCAAAACAGAGGGTGAAGGCCTGGCTGTTATTTTTCATTGCTCTTCATGATATTGGAAAGTTTGATATACGATTCCAATA
TAAATCAGCAGAAAGTTGGCTGAAATTAAATCCTGCAACGCCATCCTCGAGAGATCCTCTAGATTTAAGAAGGAGATATACATATG
>prpR
GAGGCAATTCCGACGTCTAAGAAACCATTATTATCATGACATTAACCTATAAAAATAGGCGTATCACGAGGCCCTTTCGTCTTCACCTC
GATGGATCCTAGCGTTGATGGTGCCAACAATCTGCAATGGATTTTCTTTAGTCAGTGCAGCGCGAAACGCTTTACCTGGAGAGTGTAGA
GACATAGCCCATCCTTTGTTATCAACTTGTTATTTGCGTTGATAAAGACAAAGCAAGGGGTGTGCCACAATTGCCTTAATAAAATTAAA
TGTTTAATTTCAATTTGTTAGAGATAAAGTCATGAACTAATCCGCAAATATGCGTTTCAGTTAACGTTTCAGGCAATGTTTCACCGCGT
```

```
TTCATTGCAACAATTATGAAACAAGACTAAACCCAATATTCGGTTTCTTAACTTTGCGGTGCGCTATGGCACATCCACCACGGCTTAAT
GACGACAAACCGGTTATCTGGACGGTATCTGTAACGCGCCTGTTCGAGCTGTTTCGCGATATCAGCCTCGAGAGATCCTCTAGATTTAA
GAAGGAGATATACATATG
>ygcF
GAGGCAATTCCGACGTCTAAGAAACCATTATTATCATGACATTAACCTATAAAAATAGGCGTATCACGAGGCCCTTTCGTCTTCACCTC
GATGGATCCGGCGGCATCGGTAATTTTATATTCGTTATCATTAAAAACTAATACAGAAAAAATAAATATAATTTTGGTGAATGTTTTGA
CATGATTAAATCCATTTTCTTTATTTTCTTCTGACATAATACGTTACTCCCTTTAATTATTAACGATATGTTTCGCATTTTTCTTTAAA
CCTGCCTGTAACAGAATGTTAATAAGACATGATTGTTGTGGCAGGATCTGCAATAATTAACAATTATCCACTTCACAGAGAATGCTATG
CAGTACCCGATTAACGAGATGTTCCAGACCCTGCAAGGTGAGGGTTACTTTACCGGCGTTCCCGCCATTTTTATTCGTTTACAGGGATG
CCCCTCGAGAGATCCTCTAGATTTAAGAAGGAGATATACATATG
>clpX
GAGGCAATTCCGACGTCTAAGAAACCATTATTATCATGACATTAACCTATAAAAATAGGCGTATCACGAGGCCCTTTCGTCTTCACCTC
GAGTGAACGTGATACCGAGCGCGATCGCTTCCTTTCCGCCCCTGAAGCGGTGGAATACGGTCTGGTCGATTCGATTCTGACCCATCGTA
ATTGATGCCAGAGGCGCAACTGTGCCGCTATACTTATCCAGGGCGGCACAACGCTGTAAGCGGCTTGCGCCTGAGAATGGCATTTGCGT
CGTCGTGTGCGGCACAAAGAACAAAGAAGAGGTTTTGACCCATGACAGATAAACGCAAAGATGGCTCAGGGGATCCTCTAGATTTAAGA
AGGAGATATACATATG
>stfE
GAGGCAATTCCGACGTCTAAGAAACCATTATTATCATGACATTAACCTATAAAAATAGGCGTATCACGAGGCCCTTTCGTCTTCACCTC
GATGGATCCAATCAGCTCGCATCCTGCACAGTTCAGCGCATTACGTTGTAGATCTGTGTTCTGGTCATTTGTTGATACGCGTACATAGC
CAATAAGCATGTTAAATCCCCCTGGTAAAAGCAGGAATGATGCCATTTGCTTGTTATTTCTTCATTTTCATAAACGTTGGTTTGGGAGA
AGGCTCTGCATTACCTGTTGGTGTCCCGGTTCCTCGAGAGATCCTCTAGATTTAAGAAGGAGATATACATATG
>bfd
GAGGCAATTCCGACGTCTAAGAAACCATTATTATCATGACATTAACCTATAAAAATAGGCGTATCACGAGGCCCTTTCGTCTTCACCTC
GATGGATCCACAGGCACCGAAGTACGCATTTAGCGAAATCTTCGCGCCGTTTACTAAGCAATAAATGTGTTGGCGATACCTTCCAGTAT
CGCCGCATTTTTTGTCAATGGGCTACTTTTGGTAGCCCATTATTTTTATTCATATTTCTATTCTTTGCTTAAAAAAGAGTCTTTCCGTA
ATTGAAATAAGAACTATTTTCATTTATTTTTGAAGAAAGTATACGGGAGTCTCTATGTACGTTTGTCTTTGTAATGGTATCAGCGATAA
AAAAATTCGTCAGGCTGTGCGCCAGTTTTCCCCTCACTCGTTCCACTCGAGAGATCCTCTAGATTTAAGAAGGAGATATACATATG
>clpP
GAGGCAATTCCGACGTCTAAGAAACCATTATTATCATGACATTAACCTATAAAAATAGGCGTATCACGAGGCCCTTTCGTCTTCACCTC
GAGGCTGTACTGGCGAAAGCGAAAGTGACTGAAAAAGAAACCACTTTCAACGAGCTGATGAACCAGCAGGCGTAATTTACGCAGCATAA
CGCGCTAAATTCGCACAAAGGCCCGTCACCGCCAGGTGGTGGGCTTTTTTTTGTCATGAATTTTGCATGGAACCGTGCGAAAAGCCTCT
TTCGGTGTTAGCGTAACAACAAAAGATTGTTATGCTTGAAATATGGTGATGCCGTACCCATAACACAGGGACTAGCTGATAATCCGTCC
ATAAGGTTACAATCGGTACAGCAGGTTTTTTCAATTTTATCCAGGAGACGGAAATGTCATACAGCGGCGAACGAGATAACTTTGCACCC
CATATGGCGGATCCTCTAGATTTAAGAAGGAGATATACATATG
>clpS
GAGGCAATTCCGACGTCTAAGAAACCATTATTATCATGACATTAACCTATAAAAATAGGCGTATCACGAGGCCCTTTCGTCTTCACCTC
GAGGAAAATATCTTCGCCGCCGCCTTCAGGGCAGATGAAACCAAACCCTTTGGCATTGTTGAACCACTTAACAGTACCCTTTTCCATGC
TTCGACATCCTTCGCAAATCTTATACAAGTAAGATGGAATAAACCGGGGTCAGAGAGGGGGCTGTTCAAAACCTCGCCAACTCTAGAAA
TACAATTTAGAGAATTAGGGCGAGCCGTCAAGCATTTGACAGGGGACAAGGGGCAGGTATGAATCAAAAATTTGAAGCAGTTAACGCTA
TTGACAGGAATGTGACAGATGTCGCTGATGCCAACGATAGATGATAGTTATCTATCATGTGGAGTAGATTGGTCAGGCAAATAAGCTCT
TGTCAGCGGCAGGGCGTTCTGCCGATAACCGTAACCGAAGATGATAACTGACAATGGGTAAAACGAACGACTGGCTGGACTTTGATCAA
CTGGCGGAAGAAAAAGTTCGCGACGCGCTAAAACCGGGATCCTCTAGATTTAAGAAGGAGATATACATATG
>gshA
GAGGCAATTCCGACGTCTAAGAAACCATTATTATCATGACATTAACCTATAAAAATAGGCGTATCACGAGGCCCTTTCGTCTTCACCTC
GATGGATCCTCAGGGCATGATGTGGTGGCACTAATTGTAGGCCTGCACATATGGTCACCATTACAGTTATGCTAATTAAAACGATTTTG
ACAGGCGGGAGGTCAATTTGATCCCGGACGTATCACAGGCGCTGGCCTGGCTGGAAAAACATCCTCAGGCGTTAAAGGGGATACAGCGT
GGGCTGGAGCTCGAGAGATCCTCTAGATTTAAGAAGGAGATATACATATG
>sspA
GAGGCAATTCCGACGTCTAAGAAACCATTATTATCATGACATTAACCTATAAAAATAGGCGTATCACGAGGCCCTTTCGTCTTCACCTC
GATGGATCCGTCGTCGTCCGCAGTTCTCCAAACGTTAATTGGCTTCTGCTCCGGCAGAAAACAATTTTCGAAAAAACCCGCTTCGGCGG
GTTTTTTTATAGCTAAAATCTGAATCAGCGTAAAAACTGGAAAGTTGCTTTTTGCTGCCACCTGACAGACAGGTAAAACAAAAACCACA
TCGCCAATAAGGGACTAAGTCAACTATTTCAGACTAAAGCGCATCTCTTTTTCCCCATTTCCGGCATCGACTCACCACAATGGTCGCAA
AATCTGGTAAACTATCATCCAATTTTCTGCCCAAATGTCGGGTATTGCTCATTTTTTGTTTGATTTTCGAACAAAGAGAGTCGTTCCTT
ATTGGGTAACACAACTTCTGACTGGCCACCTGGTGGCTGGTAGCAGTAAAAATTCTGACTATACCTGGAGGTTTTCATGGCTGTCGCTG
CCAACAAACGTTCGGTAATGACGCTGTTTTCCGGTCCTACTGACATCTATAGCCATCAGGTCCGCATCTCGAGAGATCCTCTAGATTTA
AGAAGGAGATATACATATG
>stfR
GAGGCAATTCCGACGTCTAAGAAACCATTATTATCATGACATTAACCTATAAAAATAGGCGTATCACGAGGCCCTTTCGTCTTCACCTC
GAGTCCGTGGCCATTGATATTGCTTATGAAGGCCCCGGCAGTGGCGACTGGCGCACTGACGGTTTCATCGTGGGTGTCGGTTATAAGTT
CTGATTAGCCAGGTAACACAGTGTTATGACAGCCCGCCGGTTCAGGCGGGCTTTTTTGTGGGGTGAATATGGCAGTAAAGATTTCAGGT
GTACTGAAAGACGGCACAGGAAAACCGGTACAGAACTGCACAATCCAGCTGAAAGCGGATCCTCTAGATTTAAGAAGGAGATATACATA
TG
>hyfR
```

```
GAGGCAATTCCGACGTCTAAGAAACCATTATTATCATGACATTAACCTATAAAAATAGGCGTATCACGAGGCCCTTTCGTCTTCACCTC
GAGGACGTCTGTGTGGCAGGCGCGTAGCATTCAATTCATTCATCTGTTGGATGAAATTGTGCAGGAACCGGCCATCTATCTGATGGCCA
GAAAAATTGCGTGAGAAGGATTTCTCATTAATAAGGACTGTTGATGGCTATGTCAGACGAGGCGATGTTTGCCCCGCCACAAGGAATAA
CAATTGAAGCGGTAAACGGAATGCTCGCGGAGCGGTTAGCACAGAAACACGGCAAGGCGGATCCTCTAGATTTAAGAAGGAGATATACA
TATG
>clpB
GAGGCAATTCCGACGTCTAAGAAACCATTATTATCATGACATTAACCTATAAAAATAGGCGTATCACGAGGCCCTTTCGTCTTCACCTC
GATGGATCCCCGGTCGTATGGCAAGTTTCATTTGGCTGATATAACCTAAAGAATCAAGACGATCCGGTACGCGTGATTTTCTTTTCACA
TTAATCTGGTCAATAACCTTGAATAATTGAGGGATGACCTCATTTAATCTCCAGTAGCAACTTTGATCCGTTATGGGAGGAGTTATGCG
TCTGGATCGTCTTACTAATAAATTCCAGCTTGCTCTTGCCGATGCCCAATCACTTGCACTCGGGCCTCGAGAGATCCTCTAGATTTAAG
AAGGAGATATACATATG
>sthA
GAGGCAATTCCGACGTCTAAGAAACCATTATTATCATGACATTAACCTATAAAAATAGGCGTATCACGAGGCCCTTTCGTCTTCACCTC
GATGGATCCACGCCCATCACGATGTCTGAATCCTTGCCAAAAGTACTGCTGGCACTATACCAGAGAATGAACATACTGGATTGCTCCAG
TATTGTGAATGAACGGTAACGCAATAAAACATATTTACTGCAATTCTTTGGCCGGTTCTTTTACGTACAGCGGAAACCTGCCGCTTAAA
CGGAGAGTATCGTCGATAAAAATCCAATAAAACGTCAGGGCAAAAGTAAGAAACAGACAAAGCAAAGGCCGCTCAGGATATAGCCAGAT
AAATGACGGGGATCAATTGGCTTACCCGCGATAAAATGTTACCATTCTGTTGCTTTTATGTATAAGAACAGGTAAGCCCTACCATGCCA
CATTCCTACGATTACGATGCCATAGTAATAGGTTCCGGCCCCGGCGGCGAAGGCGCTGCAATGGGCCTGGTTAAGCAAGGTGCGCGCGT
CGCAGTTATCGAGCGTTATCAAAATGTTGGCGGCGGTTGCACCCACTGGGGCACCATCCCGTCGAAAGCTCTCGAGAGATCCTCTAGAT
TTAAGAAGGAGATATACATATG
>plsX
GAGGCAATTCCGACGTCTAAGAAACCATTATTATCATGACATTAACCTATAAAAATAGGCGTATCACGAGGCCCTTTCGTCTTCACCTC
GAGTGGTGAAAAACACCTGCGTCACCACATCACTGCCGACGGTTACTACCGCGGCCGCAAGGTCATCGCTAAGTAATCACGCATCTGCG
TGATGAAGCTTAGTGAGGATTTTCCCCAGGCAACTGGGGAAAGACCAAACCGGGCGGCGACGATACCTTGACACGTCTAACCCTGGCGT
TAGATGTCATGGGAGGGGATTTTGGCCCTTCCGTGACAGTGCCTGCAGCATTGCAGGCACTGAATTCTAATTCGCAACTCACTCTTCTT
TTAGTCGGCAATTCCGAGGATCCTCTAGATTTAAGAAGGAGATATACATATG
>hrpB
GAGGCAATTCCGACGTCTAAGAAACCATTATTATCATGACATTAACCTATAAAAATAGGCGTATCACGAGGCCCTTTCGTCTTCACCTC
GAGCGCGCCAATGGATAATCTGTTCGCGGATTTCTGCAGGTAAGTCGATAGCAAAGAACAGACGTTGCGGTTCAGACATGTGAGGCACT
CGGTTATGAATTACCGCGATGCTACAATGTGGCGCGAAGAATGTTAACCCTCTGGAGCGTTTTGTGTCGTCGTTGCCCGTTGCTGCCGT
CTTACCTGAATTACTTACCGCTCTCGATTGTGCGGATCCTCTAGATTTAAGAAGGAGATATACATATG
>hrpA
GAGGCAATTCCGACGTCTAAGAAACCATTATTATCATGACATTAACCTATAAAAATAGGCGTATCACGAGGCCCTTTCGTCTTCACCTC
GAGCTTTTCGCGCCATTGTTCAACAAAATAATCGGACAACTGATTAGACTGAGAGTACCCTGCCAGGATGCTGGATTTAAGAACTAATA
CCTTGCTCATGGTGTTTCCTTATAGATGTTTGAATGGGCGATGCCCCGTTGCTTGTTGACACTTTATTCACAATCCTGCCACAGAGATA
GCGCAATAAATCGAAGCCTATGTTCGAATTTATTGAACAACGCATAGAAAGCCGCGATGTGGTACTCTATATCTATCATTTAAAAGAAA
ATTAATCAGGCAGACTACTGCCCACTAACGTTATGACAGAACAACAAAAATTGACCTTTACGGCCTTGCAGCAGCGGCTGGATTCGCTG
ATGCTGCGTGACAGACTGCGTTTTTCTCGCCGTCTGCACGGCGTGAAGAAGGTTAAAAATCCTGATGCACAACAGGCCATTTTCCAGGA
GATGGCGAAAGAGGATCCTCTAGATTTAAGAAGGAGATATACATATG
>plsC
GAGGCAATTCCGACGTCTAAGAAACCATTATTATCATGACATTAACCTATAAAAATAGGCGTATCACGAGGCCCTTTCGTCTTCACCTC
GATGGATCCGCAGCGGTGATAGCGAAGAGTAATATGCCGGGGAGGGCGCTGTTTCCTCCCGCTTGCAAGGAACGCCGGATGAAACGTTA
AGTCGTCATCCGGCAGCGTATTCAACCCCCGGAAACGATAAAAATTCCCTTGAAACCGTTGTTTATTCATGCGTTGCGATTAACAATAC
GCTTTTCCAGAGAGCGGCTTTTAACAATGCCCTAACCTGATTTCAGGTGACGTACAATGCCAGTCTCTCAGACCTTCAGAGGGTGTTAT
GCTATATATCTTTCGTCTTATTATTACCGTGATTTACAGCATCTTAGTCTGTGTATTCGGCTCCATTTACTGCCTTTTCAGCCCGCTCG
AGAGATCCTCTAGATTTAAGAAGGAGATATACATATG
>plsB
GAGGCAATTCCGACGTCTAAGAAACCATTATTATCATGACATTAACCTATAAAAATAGGCGTATCACGAGGCCCTTTCGTCTTCACCTC
GATGGATCCTCGGGTGAATCCAGTGGTATTATTGGCCATTGAAAGAACCTTTTTACATTATGAGCGTCAATATCAGTGTACCGTTAAAT
CCGATGATTCGGGTTTGCGTAGCGCCGCGCGAAACATGAGCGGATACCACAGAATTTCCCATGACTTTCTGCTATCCTTGCCGCGCATT
TGCATTATTAACCAGAGGCTTTACATCGTTTATGTCCGGCTGGCTCGAGAGATCCTCTAGATTTAAGAAGGAGATATACATATG
>rpiA
GAGGCAATTCCGACGTCTAAGAAACCATTATTATCATGACATTAACCTATAAAAATAGGCGTATCACGAGGCCCTTTCGTCTTCACCTC
GATGGATCCATGAGTTAACCACGCGGCTTGCCAACGGGGTCTGAATCGCTTTTTTTGTATATAATGCGTGTGAAATTTCATACCACAGG
CGAAACGATCATGACGCAGGATGAATTGAAAAAAGCAGTAGGATGGGCGGCACTTCAGTATGTTCAGCCCGGCACCATTGTTGGTGTAG
GTACAGGTTCCACCGCCTCGAGAGATCCTCTAGATTTAAGAAGGAGATATACATATG
>pck
GAGGCAATTCCGACGTCTAAGAAACCATTATTATCATGACATTAACCTATAAAAATAGGCGTATCACGAGGCCCTTTCGTCTTCACCTC
GAGCCAGGCGATCAGCAGGTAGCAAACCAGACCCTGAAGTAATCCGGTAGCAATCATCCCCCAGCGTGTTGCGGGTGAAAGTTCAACGT
TATCCATGTTGGTTATCCAGAATCAAAAGGTGGGTTAATTATCGCATCCGGGCAGTAGTATTTTGCTTTTTTCAGAAAATAATCAAAAA
AAGTTAGCGTGGTGAATCGATACTTTACCGGTTGAATTTGCATCAATTTCATTCAGGAATGCGATTCCACTCACAATATTCCCGCCATA
TAAACCAAGATTTAACCTTTTGAGAACATTTTCCACACCTAAAATGCTATTTCTGCGATAATAGCAACCGTTTCGTGACAGGAATCACG
GAGTTTTTTGTCAAATATGAATTTCTCCAGATACGTAAATCTATGAGCCTTGTCGCGGTTAACACCCCCAAAAAGACTTTACTATTCAG
```

GCAATACATATTGGCTAAGGAGCAGTGAAATGCGCGTTAACAATGGTTTGACCCCGCAAGAACTCGGATCCCGGGATCCTCTAGATTTA
AGAAGGAGATATACATATG
>ldhA
GAGGCAATTCCGACGTCTAAGAAACCATTATTATCATGACATTAACCTATAAAAATAGGCGTATCACGAGGCCCTTTCGTCTTCACCTC
GATGGATCCATAACAGCGCGAGAACGGCTTTATATTTACCCAGCATGGGTAGTTAATATCCTGATTTAGCGAAAAATTAAGCATTCAAT
ACGGGTATTGTGGCATGTTTAACCGTTCAGTTGAAGGTTGCGCCTACACTAAGCATAGTTGTTGATGAATTTTTCAATATCGCCATAGC
TTTCAATTAAATTTGAAATTTTGTAAAATATTTTTAGTAGCTTAAATGTGATTCAACATCACTGGAGAAAGTCTTATGAAACTCGCCGT
TTATAGCACAAAACAGTACGACAAGAAGTACCTGCAACAGGTGAACGAGTCCTTTGGCTTTGAGCTGGCTCGAGAGATCCTCTAGATTT
AAGAAGGAGATATACATATG
>yjcD
GAGGCAATTCCGACGTCTAAGAAACCATTATTATCATGACATTAACCTATAAAAATAGGCGTATCACGAGGCCCTTTCGTCTTCACCTC
GAGCAGTGATTGCCCGTTGCGTAACCCGGGCGACCGCTTAGGAGAAGAAGGTACCGGCGCACGCTTGCTGGAAGATGAACAAAACTAAA
GCGCCACAAGGGCGCTTTAGTTTGTTTTCCGGTCTTTGTCTTTCTCTCTATCCCGCTGGTACACAGGAGGGTTTCCCCCGACGTCAACA
CACCTCATTCGAGCACGTGGTGGAGGTTCCGGTTGGTGTTGATGCTTTAATTGTATGTCACCGACGTTTCTTCGCCAGTGTAAAAGTAT
ACTTTTTAACCGCAATATTTTTGTCATCTCAGACGATTTTTTATCGCAATCCTGAACGGTATACGGCTCGATAACGCTGCAATCTTGCG
CACCGACGATAACGTTTGCGCATCAATTGCCTGGTTTTTCATCGTCAAGACAATAAAAGAGAAAAAAGCAGCAAACTTCGGTTGAAAAA
GCCGCTATGATCGCCGGATAATCGTTTGCTTTTTTTACCACCCGTTTTGTATGCGCGGAGCTAAACGTTTGCTTTTTTTGCGACGCAGCA
AATTGTCGCAAACCTGGAGCAGGAAGATAACGTTTCGCTGGCAGGGGATTGTCCGCCACGCATCTTGACGAAAATTAAACTCTCAGGGG
ATGTTTTCTTATGTCTACGCCATCAGCGCGTACCGGCGGTTCACTCGACGCCTGGTTTAAAATTTCACAACGTGGAAGCACTGTCCGTC
AGGATCCTCTAGATTTAAGAAGGAGATATACATATG
>yjcE
GAGGCAATTCCGACGTCTAAGAAACCATTATTATCATGACATTAACCTATAAAAATAGGCGTATCACGAGGCCCTTTCGTCTTCACCTC
GAGTTGGTCGTCTGGTTTCCGGCGAATGGCGCAAGTTGAACATCGGTACGGTCGTTATCGCCGTGGCGCTGGTGACCTTCTATGCGGGT
GGCTGGGCTATCTAATCTTTCATCGGATTCTGAAAACGGGTGGCAATGGCTGCCCGTTTTTATTTTCTCCGCACATTGTGTGAGCTTTT
GCGATATTCTGAAAAAAATGAGAATTCAGGCATAACGTCTTTCTCAGATCATAAAAAACATCGCAAACAGGGAACGCATGGAAATCTTC
TTCACCATACTGATAATGACCCTCGTGGTCTCGCTGTCCGGGGTGGTCACTCGTGTCGGATCCTCTAGATTTAAGAAGGAGATATACAT
ATG
>crp
GAGGCAATTCCGACGTCTAAGAAACCATTATTATCATGACATTAACCTATAAAAATAGGCGTATCACGAGGCCCTTTCGTCTTCACCTC
GAGCCTGAGTTGCCGTCCATTAAAATCTGATGACCAGAGGCGGATTCGCCCAGAAAAGTTAACCCTTCGACCCACTTCACTCGCGCTTG
CATTTTTGCTACTCCACTGCGTCAATTTTCCTGACAGAGTACGCGTACTAACCAAATCGCGCAACGGAAGGCGACCTGGGTCATGCTGA
AGCGAGACACCAGGAGACACAAAGCGAAAGCTATGCTAAAACAGTCAGGATGCTACAGTAATACATTGATGTACTGCATGTATGCAAAG
GACGTCACATTACCGTGCAGTACAGTTGATAGCCCCTTCCCAGGTAGCGGGAAGCATATTTCGGCAATCCAGAGACAGCGGCGTTATCT
GGCTCTGGAGAAAGCTTATAACAGAGGATAACCGCGCATGGTGCTTGGCAAACCGCAAACAGACCCGACTCTCGAATGGTTCTTGTCTC
ATTGCCACATTCATAAGTACCCATCCAAGAGCACGCTTGGATCCTCTAGATTTAAGAAGGAGATATACATATG
>fsaA
GAGGCAATTCCGACGTCTAAGAAACCATTATTATCATGACATTAACCTATAAAAATAGGCGTATCACGAGGCCCTTTCGTCTTCACCTC
GAGTACGACCGTGCGGATACCGGGGCCATCATGGGTCGAGTAGCGCTGAATATTGAAAATCATAGTTGCCTCTCTATTTCGTTCAAGCA
TTAAAATACTTTCGAATGAAAGTTAGATTGATGTGCGTCAACTGTTCAGAGAGTTTTCCCGTGATAGTCTACATTCAGACAAAAAGTAC
ATTTTGAGGATGGTTATGGAACTGTATCTGGATACTTCAGACGTTGTTGCGGTGGGATCCTCTAGATTTAAGAAGGAGATATACATATG
>fkpB
GAGGCAATTCCGACGTCTAAGAAACCATTATTATCATGACATTAACCTATAAAAATAGGCGTATCACGAGGCCCTTTCGTCTTCACCTC
GAGCTGCCATCTGTGTCGGTGCGGCACTGATTGTGCTGGAAGGTTTTTTGCCTTCTAGAGCGAAAAAACAATAATAAACCCTGCCGGAT
GCGATGCTGACGCATCTTATCCGGCCTACAGATTGCTGCGAAATCGTAGGCCGGATAAGGCGTTTACGCCGCATCCGGCAAAAATCCTT
AAATATAAGAGCAAACCTGCATGTCTGAATCTGTACAGAGCAATAGCGCCGTCCTGGTGCACTTCACGCTAAAACTCGACGATGGCACC
ACCGCCGAGTCTACCCGCAACAACGGTAAAGGATCCTCTAGATTTAAGAAGGAGATATACATATG
>yjcO
GAGGCAATTCCGACGTCTAAGAAACCATTATTATCATGACATTAACCTATAAAAATAGGCGTATCACGAGGCCCTTTCGTCTTCACCTC
GATGGATCCCGCCGAGCAGTACGTGATTGACGAGTACAACAAGTTGAAAACTCGCCTGCGCGAAGCGGCACTGGCGTAATACCGTCCTT
TCTACAGCCTCCTTTCGGAGGCTGTTTTTTTATCCATTCGAACTCTTTATACTGGTTACTCCCTACCCAATCGTATTATCAAAATGAAA
AAAATTATCGCATTGATGTTGTTTTTGACATTCTTTGCCCACGCCAACGACTCCGAGCCTGGCTCGAGAGATCCTCTAGATTTAAGAAG
GAGATATACATATG
>yjcH
GAGGCAATTCCGACGTCTAAGAAACCATTATTATCATGACATTAACCTATAAAAATAGGCGTATCACGAGGCCCTTTCGTCTTCACCTC
GATGGATCCCGATACCAGCAACCTGGGCGATACCTCGACGCTTGCCGATCCTGGCGTAGTCGAGAAGCTGCTTGAAGAGAAGCAGGCTA
TCGCGATGCCATCGTAACCCACAATTGCCCGATGCGAGTCGGTAACGGTTTGTAGGCCTGATAAGACGCGACAGCGTCGCATCAGGCAT
TGATTGCCGGATGCGGCGTAAACGCCTTATCCGGCCTACATTCGGCAAGGGTTACCCGAGCGTTAACCTTCTCCCATAAGGGAGCGGGA
ATTAAAACAATCCCTACATTACCTCTGGAGAATCTGTGATGAATGGCACTATTTATCAGCGGATAGAAGACAATGCGCATTTCAGGGAG
TTAGTCGAAAAACGGCAACGCTCGAGAGATCCTCTAGATTTAAGAAGGAGATATACATATG
>rfe
GAGGCAATTCCGACGTCTAAGAAACCATTATTATCATGACATTAACCTATAAAAATAGGCGTATCACGAGGCCCTTTCGTCTTCACCTC
GAGGCGAAATCGATGCAATGGAATTCCTCATTAATAAACTGGCAATGACCAAGACCAATGACGATTTCTTCGAAATGATGAAACGCTCA
TAAATTTGTCTTATGCCAAAAACGCCACGTGTTTACGTGGCGTTTTGCTTTTATATCTGTAATCTTAATGCCGCGCTGGCGATGTTAGG

```
AAAATTCCTGGAATTTGCTGGCATGTTATGCAATTTGCATATCAAATGGTTAATTTTTGCACAGGACTGGTGGGTTTGGAACGGACTTT
CCCTTCTGAATAAAGGTCTTCGTGGTTATACTTCTGCTAATAATTTTCTCTGAGAGCATGCATTGTGAATTTACTGACAGTGAGTACTG
ATCTCATCAGTATTTTTTTATTCACGACACTGTTTCTGTTTTTTGCCCGTAAGGTGGGGATCCCGGGATCCTCTAGATTTAAGAAGGAG
ATATACATATG
>aaeR
GAGGCAATTCCGACGTCTAAGAAACCATTATTATCATGACATTAACCTATAAAAATAGGCGTATCACGAGGCCCTTTCGTCTTCACCTC
GAGCCAAACACCACGATAACGGGAAACAGACTCATGTTGACCTTGGTTGTAAAGAGAGAGCAGGCGTTATTATTTTCAGCATCTGTCGC
CGCAGAGAAGGGCATGGAAAGCCGGGCGAGAGCAACATTGCTGTAGATTGATATTTAATATATTAGCGTAACTGTTATGCTGTTATCTA
TATTATGTGATCTAAATCACTTTTAAGTCAGAGTGAATAATGGAACGACTAAAACGCATGTCGGTGTTTGCCAAAGTAGTTGAATTTGG
CTCTTTTACCGCCGCCGCCAGACAGCTACAGATGAGCGTTTCGTCCATCAGGATCCTCTAGATTTAAGAAGGAGATATACATATG
>crl
GAGGCAATTCCGACGTCTAAGAAACCATTATTATCATGACATTAACCTATAAAAATAGGCGTATCACGAGGCCCTTTCGTCTTCACCTC
GAGGATCCCATTTAACCCGGTGTATCGGAATTTTGACAAAGGTCTTCAGGAAATCACCGACTGGATCGAAAAACGCTTGTGTTAAAAAT
TTGCTAAATTTTGCCAATTTGGTAAAACAGTTGCATCACAACAGGAGATAGCAATGACGTTACCGAGTGGACACCCGAAGAGCAGATTG
ATCAAAAAATTTACCGCACTAGGCCCGTATATTCGGATCCTCTAGATTTAAGAAGGAGATATACATATG
>agn43
GAGGCAATTCCGACGTCTAAGAAACCATTATTATCATGACATTAACCTATAAAAATAGGCGTATCACGAGGCCCTTTCGTCTTCACCTC
GAGATTGCCTCTGTTGCACGCACGGTTCTGCGTGCCGTCCGTGACACGGTGGTCTCTGTTGCCCGTGCGGTATGGAACTGGATTTTCTT
CTGAACCTGTCGTGACTGATGCCCTCCCTGACTCTGAGTCTGCTCACAAAAGCACTGTTTTCGTTACTGTCTCTCTTGTCCGTGCAATA
GCTCAATAATAGAATAAAACGATCAATATCTATTTTATCGATCGTTTATATCGATCGATAAGCTAATAATAACCTTTGTCAGTAACATG
CACAGATACGTACAGAAAGACATTCAGGGAACAACAGAACCACAATTCAGAAACTCCCACAGCCGGGATCCTCTAGATTTAAGAAGGAG
ATATACATATG
>uxuA
GAGGCAATTCCGACGTCTAAGAAACCATTATTATCATGACATTAACCTATAAAAATAGGCGTATCACGAGGCCCTTTCGTCTTCACCTC
GAGCCAACAAAGCCACCATTGAATTGATTTTGAACTTCAAATTCAGTACCAGCATCAGACCAATGCCGAATACCACCCAGAGAATGTTA
AGCACATGCATAACGTTTTACCTTACCTGGTTGAACCGTTGTTATTTTGGGCGATATGTTATGTAAATTGGTCAACCATTGTTGCGATG
AATGTCACATCCTCTGATCAATAACCATCGATTACCCTTTGCTGCAATTTGCAGCAACAACCATGAGAGTGAAATTCTTGTGATGTGGT
TAACCAATTTCAGAATTCGGGTTGACATGTCTTACCAAAAGGTAGAACTTATACGCCATCTCATCCGATGCAACGCCACGGCTGCGGTC
TGGTTGTTCATCCGGATACCTAAACAACTCCAGGGTTCCGCGTCTCTTTGCTGTGGAACCCACTATGTGAAAGAGGAAAAATCATGGAA
CAGACCTGGCGCTGGTACGGCCCAAACGATCCGGTTTCTTTAGCTGATGTCCGTCAGGCGGGCGCAACTGGCGTGGTTACCGGATCCTC
TAGATTTAAGAAGGAGATATACATATG
>cysK
GAGGCAATTCCGACGTCTAAGAAACCATTATTATCATGACATTAACCTATAAAAATAGGCGTATCACGAGGCCCTTTCGTCTTCACCTC
GAGTGTTCATCATGCCCGTTGCCGTTTGTGGCGCGACGGCGATGTGGGTCGATTGCTATCGCGATAAACACGCGATGTGGCGGTAACAA
TCTACCGGTTATTTTGTAAACCGTTTGTGTGAAACAGGGGTGGCTTATGCCGCCCCTTATTCCATCTTGCATGTCATTATTTCCCTTCT
GTATATAGATATGCTAAATCCTTACTTCCGCATATTCTCTGAGCGGGTATGCTACCTGTTGTATCCCAATTTCATACAGTTAAGGACAG
GCCATGAGTAAGATTTTTGAAGATAACTCGCTGACTATCGGTCACACGCCGCTGGTTCGCCTGAATCGCATCGGTAACGGAGGATCCTC
TAGATTTAAGAAGGAGATATACATATG
>barA
GAGGCAATTCCGACGTCTAAGAAACCATTATTATCATGACATTAACCTATAAAAATAGGCGTATCACGAGGCCCTTTCGTCTTCACCTC
GAGCACGCCCTGACCAAAAGAGTCGAGGTCGTTGACTGAAACGGTTATGATCTGACGCGTCGTCGTGCGTCGTTTTGCAGAGTAGAATT
GCGCCATTGGCGAGACTTTCTCAATTTAACAGTGTGACCTTAATTGTCCCATAACGGAACTCCATGACCAACTACAGCCTGCGCGCACG
CATGATGATTCTGATCCTGGCACCGACCGTCCTTATTGGTTTATTGCTGAGTATCTTTTTCGTCGTGCATCGCTAGGATCCTCTAGATT
TAAGAAGGAGATATACATATG
>cbpA
GAGGCAATTCCGACGTCTAAGAAACCATTATTATCATGACATTAACCTATAAAAATAGGCGTATCACGAGGCCCTTTCGTCTTCACCTC
GATGGATCCCGTTCCAGGCCTGTTTCAAATAATTATTTGCAGTGCAACTAATTCCATGTATATTACTACCCATATATAGCGTCTATAAA
ATTTAATAAATAATGACGCCCTAGTTAAACTTAAAGTGCCTGGTTCAACTATCAAAAATCGCTCACCCTTTTTCACCTGTTTAAAATAT
GTTCAGCAACCCATCTTGATGGCGACCTCCTCTCCGCGATGATTTCAATAACATATTCTGTGTTGGCATATGAAATTTTGAGGATTACC
CTACACTTATAGGAGTTACCTTACAGGGGTTCCTTCAATTTGTGTTGATTTACGCGAGATAACGCTATGGAATTAAAGGATTATTACGC
CATCATGGGCGTGAAACCGACGGACGATCTCAAGACAATCAAGACCGCCTATCGTCGACTTGCCCGCAAATACCATCCTGATGTCAGCA
AAGAACCGGATGCCGAAGCCCGCTTCAAAGAGGTCGCTGAAGCCTGGGAAGTGTTAAGTGATGAACAACGTCGCGCTGAGTATGATCAG
ATGTGGCAACATCGCAACGATCCGCAATTTAACCGCTCGAGAGATCCTCTAGATTTAAGAAGGAGATATACATATG
>imp
GAGGCAATTCCGACGTCTAAGAAACCATTATTATCATGACATTAACCTATAAAAATAGGCGTATCACGAGGCCCTTTCGTCTTCACCTC
GATGGATCCCCATCAGTAAGGCCACGGCCACGCCAATGATTTTTCCCCAATACTGCATATATTCCCCAAATCGACACACGGATATCAGG
GCTATCTCCCACAATATAAAGGTGCTTTTACCGTTTTCCGGCTGCGGTCAATTGTGGGACATCGCCTATAATTTGCATTATCATACCTG
TCATTCAATGCCGTGCCTAACACCACTGACGCTATTCGGACAGGATTAACACTAGCGTAGAGATGACGAGTACGTTAGTCTCTGACCGT
TTGTCACGCGCAACGTTACCGATGATGGAACAATAAAATCAACGTATGAAAAAACGTATCCCCACTCTCCTGGCCACCATGATTGCCAC
CGCCCTTTATAGTCAACAGGGACTGGCAGCCGACCTCGCCTCACAGTGCATGTCTCGAGAGATCCTCTAGATTTAAGAAGGAGATATAC
ATATG
>mhpR
```

GAGGCAATTCCGACGTCTAAGAAACCATTATTATCATGACATTAACCTATAAAAATAGGCGTATCACGAGGCCCTTTCGTCTTCACCTC
GATGGATCCAGCAGGCTGGATGTCAGGGTGTTGTATTGCCATTTCAGTACCTCACGACTCGGACAAAATGTCGTTGCGCGCACAGTACA
GCGCAACTTATTTGTTAAAAACATGTAAATGATTTTTTATTGTGCGCTCAGTATAGGAAGGGTGTTTTCGGCTACAATCAAAACATGC
CCGAATGTGCCTCGAGAGATCCTCTAGATTTAAGAAGGAGATATACATATG
>yicH
GAGGCAATTCCGACGTCTAAGAAACCATTATTATCATGACATTAACCTATAAAAATAGGCGTATCACGAGGCCCTTTCGTCTTCACCTC
GAGCCCCTGAATGGCTGAAAAACCTGCTCTCCTCCGGGATCGCCGCGGGCGGTATTACTGCCATCGTGCTGAATCTGATTTTCCCACCA
GAAAAACAGTAATTCACTGGCGGCGGTAAGATAATTACCGCCGCAGTCTCACCTCTTTAACCATTCCCGCCTTGAGGATTGCGCGTAAA
TCGTGCATAACTCCCTTATGTGCATTTCACGGGATGGAAGACCATGAAATTTATTGGGAAGCTGCTTCTCTACATTCTCATCGCTCTGT
TAGTGGCGATCGCTGGCCTCTATTTTCTTCTGCAAACCCGCGGATCCTCTAGATTTAAGAAGGAGATATACATATG
>yicC
GAGGCAATTCCGACGTCTAAGAAACCATTATTATCATGACATTAACCTATAAAAATAGGCGTATCACGAGGCCCTTTCGTCTTCACCTC
GAGCCAAATTCGACCAGCACCGAGCCTTCTGCATGTTTTGTATAGTTACGAGTCAGGGTAACGGGACGCACCTGATTATTGCTACGGCC
TGCTGGACGCATATTGAAATCTCCGGCTTGAAACAAATGTGGCTGCGCATTATACGGACTTCCGGCGGTTATTCCTATCCTGACAAGGC
ATCGATGGCTATAATCCTTCCACCTCTCCTTTTATAAACAGGAACGTCTATGATCCGCAGTATGACCGCCTACGCCCGGCGTGAAATCA
AGGGTGAATGGGGGAGCGCAACCTGGGAAATGCGCTCGGTAAACCAGGGATCCCTGGTTTACCGAGCGCATCGGGATCCTCTAGATTTA
AGAAGGAGATATACATATG
>solA
GAGGCAATTCCGACGTCTAAGAAACCATTATTATCATGACATTAACCTATAAAAATAGGCGTATCACGAGGCCCTTTCGTCTTCACCTC
GATGGATCCGAAGCAGGAATCGCCAAAATTGAATCCGGTGATTTCCAGGTAAACGAGTATCGGCGTCATTAACATGACATGACCATCCA
CCCAAAAGGCACCTCACCGGTGCCTTTACCATTTCTGGGTTGTATATCTGCATTCAGTTAATTACTCTGCTTAAACCATGAGAGCAGAG
AGAAAAAAATGAAATACGATCTCATCATTATTGGCAGCGGTTCCGTAGGCGCTGCCGCCGGGTATTATGCAACCCGCGCCTCGAGAGAT
CCTCTAGATTTAAGAAGGAGATATACATATG
>fdhF
GAGGCAATTCCGACGTCTAAGAAACCATTATTATCATGACATTAACCTATAAAAATAGGCGTATCACGAGGCCCTTTCGTCTTCACCTC
GATGGATCCGGGTATCAGGCAGGTCCCGTCGTCGAGAAAAAATAAAATGTCTGCCGCGTGATGGCTGTCACGCGGTATTTCGTTTCGTC
ACGTCAAAACTGACGACAGCCTGTTTTTCGTCAGAGTTTTGAATAAATAGTGCCCGTAATATCAGGGAATGACCCCACATAAAATGTGG
CATAAAAGATGCATACTGTAGTCGAGAGCGCGTATGCGTGATTTGATTAACTGGAGCGAGACCGATGAAAAAAGTCGTCACGGTTTGCC
CCTATTGCGCATCAGGTTGCAAAATCAACCTGGTCGTCGATAACGGCTCGAGAGATCCTCTAGATTTAAGAAGGAGATATACATATG
>fdhD
GAGGCAATTCCGACGTCTAAGAAACCATTATTATCATGACATTAACCTATAAAAATAGGCGTATCACGAGGCCCTTTCGTCTTCACCTC
GAGCTGGGTGCAAAACCCAGTGCCGCTGCCGTGGTGCCTGCCATACCGCCAGCGCAGATCTTAAAGAACTGCCTTCTGCTGACCTGCAT
GGATTGCTCCTTGTTTCGACATTGTCACGTCCCATTTACATTCGCTTGCTGCGTGTGCAGGGAGTGGGAGTTATTTTTCTTTGCGGAAG
GGGCCGCAAAGGTCCAGAATTGGCTCAATTTCCCTCCATCCAGGAAGGGTTTGTAACAGAATACCATAATGTTGGTGTGTGTGTTCTTA
TCTGGTTAAGAGAAAGTGAAAAAAACACAGCGAAAAGAAATCGAAAATGTGACAAATATCACAGGTGTTCGTCAAATTGAGTTATGGCG
GCGTGATGATTTACGGATCCTCTAGATTTAAGAAGGAGATATACATATG
>tgt
GAGGCAATTCCGACGTCTAAGAAACCATTATTATCATGACATTAACCTATAAAAATAGGCGTATCACGAGGCCCTTTCGTCTTCACCTC
GAGAAGCAGCGGTAGAAGAGAAATATCGCTTTTTTAGTTACGGTGATGCGATGTTTATCACGTACAATCCGCAGGCAATTAATGAGCGC
GTCGGGGAGTAATTCCGCGGCGCTGGTTTAAAACGTTGGACTGTTTTTCTGACGTAGTGGAGAAAAAATGAAATTTGAACTGGACACCA
CCGACGGTCGCGCACGCCGTGGCCGCCTGGTCTTTGATCGTGGCGTAGTGGAGGATCCTCTAGATTTAAGAAGGAGATATACATATG
>napF
GAGGCAATTCCGACGTCTAAGAAACCATTATTATCATGACATTAACCTATAAAAATAGGCGTATCACGAGGCCCTTTCGTCTTCACCTC
GATGGATCCGCGGAAGTGGTAGCGAAAGCGGCAAACAATACTGCAGGTAGAATGGTCTTCATTTTTTTGCTTTCCTTCTTCATTTATAG
GTTTTCACCGATGATGTCGCTGAAGGTTAACATATTCATCAGATTTATTAATTTGCTTTATGAATAACTTAAAGAACATTCAGACAATT
TTACCTGCTCTTTTAGTGTTAAATTCTAATGAGAGAGCAAACCAGCAGAAATATCTCTGGTGAATCAATCGCTATATAAATATATTTAT
AACCATTTGAAATGTGAGCAAAAGCCCGTTTTTGCCACGCTTCCCGCCACTCTTTTGATCCTGCTACAGGTTTTACCCCGATCGGGGTA
TGCATCTTTGACACATCCTTTAATATCTTAGCGGCTATAAAAATGGCTTATTAATTATGCGGTTTATTTGGTCGCTCTCAATTTTCAGA
GCGCGTTAATGATGGAAGGTCAATGTGAAGATTGATGCATCCCGTCGGGGCATACTCACTGGTCGCTGGCGCAAAGCCAGTAACGGTAT
CCGTCCGCCCTGGTCCGGTGATGAATCTCGAGAGATCCTCTAGATTTAAGAAGGAGATATACATATG
>sppA
GAGGCAATTCCGACGTCTAAGAAACCATTATTATCATGACATTAACCTATAAAAATAGGCGTATCACGAGGCCCTTTCGTCTTCACCTC
GAGAGTTGTTCACCCGTTGGCGCGGGTTCAGCCAAGCGGGAGGCGCTACGGCGATTGATCAATAGTTCGAGTGCATCCATTTGATTAAC
TCCTGTCGTGATATTTATTCACAAAATTAACACGAGAGTGGATTTTGTTACAGCACAGTCCGCAATTCCTGCTGACAAGTACCGGTTGG
GTCATTACGATAACCACATCTATTGCGCCTGTGACAGGTGTGACCTTAAGTTGGGAGAATACATGCGAACCCTTTGGCGATTTATTGCC
GGATTTTTTAAATGGACGTGGCGTCTGCTGAATTTCGTCCGTGGGATCCTCTAGATTTAAGAAGGAGATATACATATG
>ybhQ
GAGGCAATTCCGACGTCTAAGAAACCATTATTATCATGACATTAACCTATAAAAATAGGCGTATCACGAGGCCCTTTCGTCTTCACCTC
GAGGCAAAATGAAGCGTCGGTTAAACGCGGTAAAGCCTTTGTGAATATTGATGGTGAGCACCTTGAACGAAAATTGTTGTGTTTGATCG
GGCATAATTTTCCTGTCTTTGCCTCTTATCTCATTGAAATAGTGTAGTCGGCGTCACAAAAAGGTGCGGTCTTACGGAATTTTCCGTAA
AGTTCGGTACTCTGAGTAAGTAGAGATAAATTCTTCAGGAGAGAAGCCATGAAGTGGCAACAACGTGTTCGTGTCGCAACGGGTCTAAG
TTGCTGGCAGATTATGTTGCATTTACTGGTAGTGGCGCTGCTGGATCCTCTAGATTTAAGAAGGAGATATACATATG
>ybhM

GAGGCAATTCCGACGTCTAAGAAACCATTATTATCATGACATTAACCTATAAAAATAGGCGTATCACGAGGCCCTTTCGTCTTCACCTC
GAGTTGGCGCGTTAACCTTGTATCTGGACTTCATCAACCTGTTCCTGATGTTGTTGCGGATCTTCGGCAACCGCCGTTAATTCTTTTTC
ACCATCACTATCCCGGCCATGTGTCGGGATTTTTTTCTTACTAAAAACGACACTTAACAAGTACCAGGTATAAAATATGCCTTATGGAC
GAGCGGAAGCGTTTTCTGTATCCATAAATGCAAAAGTATTTTGTATGCGTGGTTTTAAATTTATTGATGGTGAATATTAATATTTTTTC
TAAGGATAACTAAAAGAATGGAATCATACTCGCAAAACAGCAATAAATTAGATTTTCAGCACGAGGCCAGGATATTAAACGGTATATGG
CTCATTACCGCTTTAGGTTTGGTGGGGATCCTCTAGATTTAAGAAGGAGATATACATATG
>ybhL
GAGGCAATTCCGACGTCTAAGAAACCATTATTATCATGACATTAACCTATAAAAATAGGCGTATCACGAGGCCCTTTCGTCTTCACCTC
GAGTCTGGAAGCGCGAAGCCACGCCGGAAGGCGACCGCTGGGTTGAAGCTCGGGAGAGCGATCAGCAGGCGGCAAAACGCTGGTAGTTT
TTTGTTAGCCGGATAAGGCACCGCTGCGTCCGGCAATTTCACCGCTCAGTACATACGTTTACGCTGCGTCGATGCACAGCCTCATCATT
TTGCAGTATCCTTAAGATATTCCTTATATCTTCAGGAGATCGTCATGGACAGATTCCCACGTTCTGATTCAATCGTACAACCCCGGGCT
GGCTTGCAAACTTATATGGCTCAAGTCTATGGCTGGATGACCGTTGGCTTGTTGCTGGATCCTCTAGATTTAAGAAGGAGATATACATA
TG
>ybhI
GAGGCAATTCCGACGTCTAAGAAACCATTATTATCATGACATTAACCTATAAAAATAGGCGTATCACGAGGCCCTTTCGTCTTCACCTC
GAGAAGGTCAGGATGCCACGACGTTACGCGCATCTGTTATTCGGACGACCAGAAAAATATTTTCCGGTGAAGTTTATCTTCCCTGAAAA
AATTCGTTGTCAGGATAAGGACAATCAATAAAGGACTTCTGTATGAGTCATACAGAAAGAACAGGATTTTAAATGAATAAGAAATCGTT
ATGGAAGCTAATTCTGATATTAGCGATCCCATGTATTATTGGTTTTATGCCAGCTCCGGCAGGATTAAGGATCCTCTAGATTTAAGAAG
GAGATATACATATG
>ybhK
GAGGCAATTCCGACGTCTAAGAAACCATTATTATCATGACATTAACCTATAAAAATAGGCGTATCACGAGGCCCTTTCGTCTTCACCTC
GATGGATCCATCGGTAATCGACAGGCGCAAGTAGTAAAACTTACGCGCAAATGCATCAGTCAGTTGTGAAGCCATGTACACCTTTCCAG
ATACGGGAGGCGAAGTCATTTCTTCCTTCGCCCTGGTGGCAATCTTTACGAGCATTGTCACGGCCAAAGCACCGTATCAGTTGACCCAG
GTGCAGAGGCTAGAGTGTTTAGTGGTTATGCCGATACTAGCGTGTAAATGAGATTTTTACCATCCACACTTTCGCTATGTAATCATGTA
TATAGCGTCATGATCGTGCTATTTCGCTACAGAATAAGGGAAAATGGCGGTTTTACTTTGATATACATCATCAGGGAGGCATAGCGTGC
ATCGTCCTTTAGAGGTAGAAGAAAATAGTTTGTATCCTCAATATTGGCAGGTTAATTGCTGTTTCCCCGCAATTTGCGCTACTGTAGCG
CGGATAATCTGACTCCAGGAAACTATATGCGCAATCGTACGCTGGCTGATCTTGATCGTGTCGTTGCTCTCGGCGGAGGGCATGGACTG
GGACGCGTTCTCTCATCTCGAGAGATCCTCTAGATTTAAGAAGGAGATATACATATG
>ybhJ
GAGGCAATTCCGACGTCTAAGAAACCATTATTATCATGACATTAACCTATAAAAATAGGCGTATCACGAGGCCCTTTCGTCTTCACCTC
GAGGTTGGTCGGTGCGGTACTGACGATATTAACCTTCCTGGTGCATATCACCCTCGGCGTGTGGTGGTGGAATATGCTGATCGGCTGGA
ACATGCTGTAAATATTCTCGTCATACTTCAAGTTGCATGTGCTGCGTCTGCGTTCGCTCACCCCAGTCACTTACTTATGTAAGCTCCTG
GGGATTCACTCGCTTGTCGCCTTCCTGCAACTCGAATTATTTAGAGTATATCCATTTATTATCTTTCTGCGCACTTCACGGTGCGCAGA
TATCTGGAGCATTTGATGATCAAGTTATCTGAAAAAGGCGTGTTTCTCGCCAGTAATAACGAAATAATTGCCGAAGAACATTTCACCGG
CGAGGATCCTCTAGATTTAAGAAGGAGATATACATATG
>ybhE
GAGGCAATTCCGACGTCTAAGAAACCATTATTATCATGACATTAACCTATAAAAATAGGCGTATCACGAGGCCCTTTCGTCTTCACCTC
GAGTCGATGAAGGAAGCAGGGTCTTTTTCGGGGTCAATAAGGTGCCGTCTAAGTCGAGAGCAATCACGCGTGTGGTCATGGCGTATTCC
AGATTAAGGTTAAGAATTTTCGTCTGCGCGAATGGTACACCGATACCACTATCGGACAAAATTCTGCGTTTTAATTCAGCATTCACCGC
CAAAAGCGACTAATTTTAGCTGTTACAGTCAGTTGCTAAATGCAAAGGAGCATTCATGAAGCAAACAGTTTATATCGCCAGCCCTGAGA
GCCAGCAAATTCACGTCTGGAATCTGAATCATGAAGGCGCACTGACGCTGACACAGGTTGTCGGGATCCTCTAGATTTAAGAAGGAGAT
ATACATATG
>ybhD
GAGGCAATTCCGACGTCTAAGAAACCATTATTATCATGACATTAACCTATAAAAATAGGCGTATCACGAGGCCCTTTCGTCTTCACCTC
GATGGATCCCAGCCACCCAAAATCGTCGCTTCCATTTTCCGGAGTAATAATGAAGCATGAATTATCAAGTATGAAGGCATTTGTCATAC
TGGCAGAGTCCAGTTCGTTTAACAATGCCGCTAAATTACTCAATATTACGCAACCTGCATTAACGCGAAGAATAAAAAAGATGGAGGAA
GATTTACATGTTCAACTATTTGAGCGTACAACTCGCAAGGTTACGTTAACAAAAGCAGGAAAAAGGTTGCTCCCGGAGGTCGAGAGATC
CTCTAGATTTAAGAAGGAGATATACATATG
>ybhA
GAGGCAATTCCGACGTCTAAGAAACCATTATTATCATGACATTAACCTATAAAAATAGGCGTATCACGAGGCCCTTTCGTCTTCACCTC
GATGGATCCCGACAACCTGTGTCAGCGTCAGTGCGCCTTCATGATTCAGATTCCAGACGTGAATTTGCTGGCTCTCAGGGCTGGCGATA
TAAACTGTTTGCTTCATGAATGCTCCTTTGCATTTAGCAACTGACTGTAACAGCTAAAATTAGTCGCTTTTGGCGGTGAATGCTGAATT
AAAACGCAGAATTTTGTCCGATAGTGGTATCGGTGTACCATTCGCGCAGACGAAAATTCTTAACCTTAATCTGGAATACGCCATGACCA
CACGCGTGATTGCTCTCGACTTAGACGGCACCTTATTGACCCCGAAAAAGACCCTGCTTCCTTCATCGACTCGAGAGATCCTCTAGATT
TAAGAAGGAGATATACATATG
>ybhC
GAGGCAATTCCGACGTCTAAGAAACCATTATTATCATGACATTAACCTATAAAAATAGGCGTATCACGAGGCCCTTTCGTCTTCACCTC
GATGGATCCCGATGGTCGGGTTTAACGTTCATTTCCACTCTCTGGCAAGCGCCTCGATTACTGCGATGTTTAGTTAATCACTCTGCCAG
ATGGCGCAATGCCATCTGGTATCACTTAAAGGTATTAAAAACAACTTTTTGTCTTTTTACCTTCCCGTTTCGCTCAAGTTAGTATAAAA
AAGCAGGCTTCAACGGATTCATTTTTCTATTTCATAGCCCGGAGCAACCTGTGAACACATTTTCAGTTTCCCGTCTGGCGCTGGCATTG
GCTTTTGGCGTGACGCTGACCGCCTGTAGCTCAACTCGAGAGATCCTCTAGATTTAAGAAGGAGATATACATATG
>mlc

```
GAGGCAATTCCGACGTCTAAGAAACCATTATTATCATGACATTAACCTATAAAAATAGGCGTATCACGAGGCCCTTTCGTCTTCACCTC
GATGGATCCTTGGTCTGGCCGAAACATCATGAACAAAGTCCGGCTGCGCGTAACTTTCGTATTCATCTGCTGAATGCTCTCAGGTGAGG
GAAATTTCAGCGAAAAAGCCCGAAAAATGTGCTGTTAATCACATGCCTAAGTAAAAATTTGACGACACGTATTGAAGTGCTTCACCATA
GCCTACAGATTATTTCGGAGCGCGAAAATATAGGGAGTATGCGGTGGTTGCTGAAAACCAGCCTGGGCACATTGATCAAATAAAGCAGA
CCAACGCGGGCGCGGTTTATCGCCTGATTGATCAGCTTGGTCCAGTCTCGCGTCTCGAGAGATCCTCTAGATTTAAGAAGGAGATATAC
ATATG
>fur
GAGGCAATTCCGACGTCTAAGAAACCATTATTATCATGACATTAACCTATAAAAATAGGCGTATCACGAGGCCCTTTCGTCTTCACCTC
GATGGATCCCGGAACTGACCGCTGAACGTGTAGAAAAATGGGTTAAACAGATTTCTGAAGAGTTGCATCTCGACGAAATTCTCAATGCC
TGATGTGATGCGGCGTAGACTCATGTCTACGCCGTATTAATAGATAATGCCAATCAAAATAATTGCTACAAATTTGTAACTTTTGCTGT
TGTACCTGTACAATGTCCCGGTGTTCAAGTGGCCTTGCCGTTGTAAATGTAAGCTGTGCCACGTTTTTATTAACAATATTTGCCAGGGA
CTTGTGGTTTTCATTTAGGCGTGGCAATTCTATAATGATACGCATTATCTCAAGAGCAAATTCTGTCACTTCTTCTAATGAAGTGAACC
GCTTAGTAACAGGACAGATTCCGCATGACTGATAACAATACCGCCCTAAAGAAAGCTGGCCTGAAAGTAACGCTTCCTCGTTTAAAAAT
CCTGGAAGTTCTTCAGGAGCCGGACAACCATCACCTCGAGAGATCCTCTAGATTTAAGAAGGAGATATACATATG
>tdcB
GAGGCAATTCCGACGTCTAAGAAACCATTATTATCATGACATTAACCTATAAAAATAGGCGTATCACGAGGCCCTTTCGTCTTCACCTC
GATGGATCCAGCATCGGTTTTGGTGGAATTAGCCAAAGAGTATTCATCTTATAATGGGTGTAGACGAAGGCAATTAATAGAAGTTGGTT
AGTTATTTGTTTTTATTTAAACATAAATAATCCACCTGTCTGTCTTGCAGGTGTCGGTTACGGTTACCTACATATTTAATTCAGGCGAA
GAGGTTTTATAATGCATATTACATACGATCTGCCGGTTGCTATTGATGACATTATTGAAGCGAAACAACGACTGGCTGGGCTCGAGAGA
TCCTCTAGATTTAAGAAGGAGATATACATATG
>b1172
GAGGCAATTCCGACGTCTAAGAAACCATTATTATCATGACATTAACCTATAAAAATAGGCGTATCACGAGGCCCTTTCGTCTTCACCTC
GATGGATCCACGTCTGCCCGACAAATCATTGACGAAATTACCGTCATTATTCACCTTCACCCGGTTATCATCGACAAATTCTTTGCGCA
CAGCCGCCTTCAGCCAGGGTTCATTTCCATACCGTTCCCCAGGCGCATGTTGTAACTCAGCGTTGCACCCAGTTCACGATATATACTGC
GGGTATCGACTGATTTCGATTCCATGCCATTGGATAAATGATAGTAGCGTCTAAGTGACCTAAGCTCTGCTGCAACATCAGCCCGCAGG
CACCAGACCAGGGGATTCATCCTGAAGAGACAGCGCAGGTGTATTGTGTTCACCGCTCACCAAAGACATCATGATGAAATGATGATATC
CCGCATAAGAAAGAGGCATTTTTTACACGCAGTGCGCTGAAGTGTAGTTGGATAAAAAATCACTCTTTTCTGATAATACTGTCGTACGC
TTATCTTTCGACAATGAGGCCGTCGGCAAAATAAAATGACTAATGTAATCGTTGCTGATGAATATCTTCTGCTCACATAAAAATCACAC
AATAATATTGATAAATCAGATTATCTTACCTTTACAACATTGCGTTAACAGTCAATGACACTTCTTGAATAAAAATATTGTCAATCAAT
GAGTTATTCCTCATTTAAGATTATTCTCATTCACGTAAAAAACATAATTCCTATCATCACAGCCACGCTCATTTTGAATTATCTGAATA
ATAGCGAGCGTTCTTTAGTTAAGCAGATCTTAATTGAAGACGAAATTATTGTGTGTGCAACTTACTTAATACCGGACATCTAAGTTGAA
GAAAAAAATACTTGCCTTCGGGCTTATCTCAGCATTGTTCTGCTCCACGCCAGCGATGGCGGATATGAACCGCACCACGAAAGGTGCAT
TGCTAGGCGCAGGCGTTGGTTTACTTACTGGCAATGGTGTTAATGGCCTCGAGAGATCCTCTAGATTTAAGAAGGAGATATACATATG
>glgB
GAGGCAATTCCGACGTCTAAGAAACCATTATTATCATGACATTAACCTATAAAAATAGGCGTATCACGAGGCCCTTTCGTCTTCACCTC
GATGGATCCGTCAGCCTTTACCGTGGGCGACCAGCTGCTGTGGGGGGCCGCGGAGCCGCTGCGTCGGATGCTTCGTCAACTGGCGTAAT
CTTTATTCATTAAATCTGGGGCGCGATGCCGCCCCTGTTAGTGCGTAATACAGGAGTAAGCGCAGATGTTTCATGATTTACCGGGAGTT
AAATAGAGCATTGGCTATTCTTTAAGGGTGGCTGAATACATGAGTATTCACAGCCTTACCTGAAGTGAGGACGACGCAGAGAGGATGCA
CAGAGTGCTGCGCCGTTCAGGTCAAAAAAATGTCACAACCAGAAGTCAAAAATCCAATTGGATGGGGTGACACAATAAAACAGGAAGAC
AAGCATGTCCGATCGTATCGATAGAGACGTGATTAACGCGCTAATTGCAGGCCATTTTGCGGATCCTTTTTCCGTACTGGGAATGCATA
AAACCACCGCGGGACCTCGAGAGATCCTCTAGATTTAAGAAGGAGATATACATATG
>glgS
GAGGCAATTCCGACGTCTAAGAAACCATTATTATCATGACATTAACCTATAAAAATAGGCGTATCACGAGGCCCTTTCGTCTTCACCTC
GATGGATCCACCGCAGATAAGCGCAAAAATTTCTAACAGGCCGATCAACAGTACAAAGGAAATAGCGAAAAGATAAGGGGTGTTATAGT
CGGCGAATAAAATCATATATCACTCCAATGTCCATTTAGTGACGCAATATTTACTTTGTAATAGTTTCCTTGCTTACATTTATCTGTGT
TAATAATAACACAGGTTAGAGAAGTTCTGGAAGAGAAAGTCTTTTAATATGGTTTTTTATTTTTCACTAAAAAGTGTGATCGGGGACAA
TATATTTACGCACGTTATGTTTAAAGGCACTACACTGATTGGGGAATACTGAAATCAGAAAAAGCATATGGCCTTAAAAGGTATGCGAT
AACCGGAGGATGTGCTTATGGATCATAGTCTTAATTCTTTAAATAATTTCGATTTCCTGGCGCGTAGTTTTGCCAGAATGCACGCAGAA
GCTCGAGAGATCCTCTAGATTTAAGAAGGAGATATACATATG
>motA
GAGGCAATTCCGACGTCTAAGAAACCATTATTATCATGACATTAACCTATAAAAATAGGCGTATCACGAGGCCCTTTCGTCTTCACCTC
GATGGATCCCCCAGAATCCTGCCGATATTATCCCACAACTGCTGGATGAACAGAGAGTACAGGCTGTTTAACTGATACGGTGAGGCGCA
ACATTCCAGCAGCGGTAACGACGTACCGCTGCTTTTTTTTGCCCCAATCGCGCGTTAACGCCTGACGACTGAACATCCTGTCATGGTCA
ACAGTGGAAGGATGATGTCGTGCTTATCTTATTAGGTTACCTGGTTGTTCTCGGTACAGTTTTCGGCGGTTATTTGATGACCGGTGGAA
GCCTTCTCGAGAGATCCTCTAGATTTAAGAAGGAGATATACATATG
>ptrB
GAGGCAATTCCGACGTCTAAGAAACCATTATTATCATGACATTAACCTATAAAAATAGGCGTATCACGAGGCCCTTTCGTCTTCACCTC
GATGGATCCAACGCGATCTCGAACAGCAAAAACGCACTTTGGCGGAATAATTCCGCTCTCTGGATAGCCTGAAGTTGACTATTTCGGGC
TATCCAGAATCGCACAGCCATTTGAACAGGCTGATTTTCTCTGTCTGCATAACATTCGATATCCGCGCCATTACCTGTCATCATCTAAG
CAATGACTCCCCTGTTTCGCTTGCATCCCCGGTGAGTTTTGCCACCCTTATAAGATGTTTCAACCAGAAAGAACAATAACATGCTACCA
AAAGCCGCCCGCATTCCCCACGCCATGACGCTTCATGGCGATACGCGCATCGATAATTACTACTGGCTGCGGGACGATACGCGTTCTCC
TCGAGAGATCCTCTAGATTTAAGAAGGAGATATACATATG
>dsbG
```

GAGGCAATTCCGACGTCTAAGAAACCATTATTATCATGACATTAACCTATAAAAATAGGCGTATCACGAGGCCCTTTCGTCTTCACCTC
GATGGATCCCCGTTTTTGAATGCCTGGTTTTTAAAAGGTTTAATTTTGGTGTTAATCAAGGACATCTATACTTCCTCCGTGTTTTCGAT
GAGATGTAAGGTAACCAATTTTTGCCGATTCGGCTAATGCGTTTCCATTATCAAATCGATAAGTTTTACCTTACAACCTTCGTAAGACA
ACTTTGTGAACTTTGCACCGCCAGCGGCGATAAAAAGTAAAATGGCCCCCTTTCCAGGTGACCGTGTAATCTGAACTACCCTTTGGTAT
CTTCAGTGCCTGCCTGCAGGCTGAGGTGTTGCCCTTACAAATGCAACAACGACATGGATTACAACACCCTCATAAACAAAGGGCAATCA
CCTGATCTAAGCTCTTACCTATGACAGTGATAGGTTATGCCTTTTACTCGACTTTTGCACTGACTGAAAAGGACAAATTAATGTTAAAA
AAGATACTTTTACTGGCTCTGCTTCCTGCCTCGAGAGATCCTCTAGATTTAAGAAGGAGATATACATATG
>ampH
GAGGCAATTCCGACGTCTAAGAAACCATTATTATCATGACATTAACCTATAAAAATAGGCGTATCACGAGGCCCTTTCGTCTTCACCTC
GATGGATCCCAAACGAAGGCCGAGAGAAAAAACGTTCCCGGCTTTGGGAAAAAAGACTTAAACATATGCACTCCTGCTAACTTCTTATC
GTTTTAACAACTGCTGTGTATTAACCGCATGACCGCTACAAAGCGTCAGCCAAATGAAACCGCCCATTGTGACCCATAGCAAGGCGCTT
AGTTCGCCAGCCTGACCATAAGAATTGTTTCGACGGATTTCACAAACCAAAGATTGTTTACATAAGCGGATTAACACGCGCAGGATCAC
AAATGCAGGCATTGGTGAGCAGGAAAAAGATCGTGTTTACGCGCCGTTACGTTTTTAACACTATTATTGCACTAATTCGCCCTTTGCAA
TCTATCAATGAGTAGTATAAATACGCTCAGTTACCTTCATTCAATCTATGGACACCACCGTTGAAACGTAGTCTGCTTTTTTCTGCCGT
GCTGTGTGCGGCGTCATTGACCTCCGTCCATGCGGCGCAGCCGATCACTGAACCGGAGTTTGCCTCCTCGAGAGATCCTCTAGATTTAA
GAAGGAGATATACATATG
>ydhH
GAGGCAATTCCGACGTCTAAGAAACCATTATTATCATGACATTAACCTATAAAAATAGGCGTATCACGAGGCCCTTTCGTCTTCACCTC
GATGGATCCGCGGTATAAACATCCCCTGACAGGGTGTCGTTATTAACACAACCGACAAGAGACAGACCTACCATTGAAACAACCAATAC
GCGTTTAATCATTGAAAAATCTCCTGTTCACCATGAAACGCCACGCAAGCATCCCTCATGGTCAAAGTATATGGCATATGTGAATCCAT
GGTTACATGTTCTACCAAATATGAGCAAATCATAATGAAAGTAGGATCAACCAATTATAAACAAAACATTCATGATGATTATTGATTTG
AAGATAATGTTTTACTACCAACGGTAAAAATCGGTCATTCCGCATGGTATTCACGAAAAGTTTATGTTTGAGTGGTGCGCAAACGTCAG
ATTGAAAAAGGATGAACTATGAAATCGGGCCGCTTTATTGGCGTTATGTCAGGCACCAGCCTTGATGGTGTTGATGTTGTGTTGGCGAC
AATTGATGAACACCGGGTCGCTCGAGAGATCCTCTAGATTTAAGAAGGAGATATACATATG
>ydhO
GAGGCAATTCCGACGTCTAAGAAACCATTATTATCATGACATTAACCTATAAAAATAGGCGTATCACGAGGCCCTTTCGTCTTCACCTC
GAGGCAGAGAAACCGCAGCTCGGCAGTTTCGGTGAACCTTTCATGTACAGCAGGATCGGGTTTTCAGCAATCTGGCGTTGGATTTTTTC
GATAGTGGTGCTCATTGTCTTGCTTCCTCAAACTTCTTTACGGCAGTAATCTGACATTGTAGCGGGTCAGTGCGGCATCGGAAAATAAC
ATTTTCATCACGCTTTTGCTATTTTATCCCTTTGCTCTATTTGTTGCATTTCAAATATTCGTTTCTTCTCTGATGCCGGCAAACTGGCG
TATTTATAACCATTTTTAGCTTTCACTGCTGCATTTTTTTGCGCTCGCCAACGAAACGTATTTTTTAACAATAAAAGCTATTAACTTTC
TCTTCTTCTATGCATTAGAATCATCAAGTTTTGTAAATCAGACGCAGGCATGATAGACCTGCCTTTACAGAGGGACGCTCAGTGGCGCG
GATAAACCGTATTTCGATCACGCTCTGTGCTTTACTTTTTACCACCCTGCCTTTAACGCCTATGGCCCAGGATCCTCTAGATTTAAGAA
GGAGATATACATATG
>syd
GAGGCAATTCCGACGTCTAAGAAACCATTATTATCATGACATTAACCTATAAAAATAGGCGTATCACGAGGCCCTTTCGTCTTCACCTC
GATGGATCCGCAGTAGGCTGGCGTCATAGGTATCCCGGTAATCGGTTGATTTTCCAAGAGTCAGGCCCGCAAGTGCCTGATGGTTTGCA
TAAGAAGACATGTTTCACCGTCATAAACCAGATAAACTAAACGACAAGTGTATCTGGTTTAAGTGAAGAGAGAAATTAGTGGACGATTT
GACCGCACAAGCCCTGAAAGATTTTACTGCACGCTACTGCGATGCGTGGCATGAAGAGCATAAAAGCTGGCCGCTCGAGAGATCCTCTA
GATTTAAGAAGGAGATATACATATG
>ydhB
GAGGCAATTCCGACGTCTAAGAAACCATTATTATCATGACATTAACCTATAAAAATAGGCGTATCACGAGGCCCTTTCGTCTTCACCTC
GATGGATCCGGCGAAAGCAGGCAGATACATATCGGTTGCCAGAAAACCGAGTACGCTCAAACCCGCCAGCCAGACTAAAAATCTTTTCC
CAGGTTGCATGTATATTCTCTAAATCTCTCAATTAAACACCGCCGCAAAGTGTAGGGAGTGAAATTAGTGTTGTGAAACGGTAATATTT
GATGACTGGTTTCAAAAATTTTGCAGGCAGACTATGTGGTCAGAATATTCACTCGAAGTTGTTGATGCGGTAGCGCGTAATGGTAGTTT
TAGCGCTGCGGCACAGGAGCTGCATCGCGTTCCTTCTCTCGAGAGATCCTCTAGATTTAAGAAGGAGATATACATATG
>mobA
GAGGCAATTCCGACGTCTAAGAAACCATTATTATCATGACATTAACCTATAAAAATAGGCGTATCACGAGGCCCTTTCGTCTTCACCTC
GATGGATCCAGGCTGGCTGGAGGAGTTCAATAACTTCATTCAGACGTTTACATTTCATAGTGATGCTCCTTACTCTTGAGACAGACACG
TTAGCAGGGTCAATCCCACAATAAAAGAGGCGATATCGGTGAATCTGATGACGACGATAACAGGCGTTGTGCTGGCAGGCGGTAAAGCC
AGACGAATGGGCGGCGTAGATAAAGCTCGAGAGATCCTCTAGATTTAAGAAGGAGATATACATATG
>ampD
GAGGCAATTCCGACGTCTAAGAAACCATTATTATCATGACATTAACCTATAAAAATAGGCGTATCACGAGGCCCTTTCGTCTTCACCTC
GAGGCCGGGGATATCGAGATTAATGCGTTCCAGCAGCTCGTCACGTCGGGTGTCAGGGTTATAGCGGCGAGGCGGCATGTTAAAACTCC
AGATAGCTAACGAATCATAAGGTAGAAACATGCTACTCTGAACCGGGTATTAGCACCACATATAAGGAGATCCTGCATGTTGTTAGAAC
AGGGGTGGCTGGTTGGCGCGCGCCGCGTTCCCTCACCACATTACGATTGCCGCCCGGATGACGAAACACCCACCCTGGATCCTCTAGAT
TTAAGAAGGAGATATACATATG
>ydhD
GAGGCAATTCCGACGTCTAAGAAACCATTATTATCATGACATTAACCTATAAAAATAGGCGTATCACGAGGCCCTTTCGTCTTCACCTC
GATGGATCCTGGGCCATAGGCGTTAAAGGCAGGGTGGTAAAAAGTAAAGCACAGAGCGTGATCGAAATACGGTTTATCCGCGCCACTGA
GCGTCCCTCTGTAAAGGCAGGTCTATCATGCCTGCGTCTGATTTACAAAACTTGATGATTCTAATGCATAGAAGAAGAGAAAGTTAATA
GCTTTTATTGTTAAAAAATACGTTTCGTTGGCGAGCGCAAAAAAATGCAGCAGTGAAAGCTAAAAATGGTTATAAATACGCCAGTTTGC
CGGCATCAGAGAAGAAACGAATATTTGAAATGCAACAAATAGAGCAAAGGGATAAAATAGCAAAAGCGTGATGAAAATGTTATTTTCCG
ATGCCGCACTGACCCGCTACAATGTCAGATTACTGCCGTAAAGAAGTTTGAGGAAGCAAGACAATGAGCACCACTATCGAAAAAATCCA

```
ACGCCAGATTGCTGAAAACCCGATCCTGCTGTACATGAAAGGTTCACCGAAACTGCCGAGCTGCGGTTTCTCTGCCTCGAGAGATCCTC
TAGATTTAAGAAGGAGATATACATATG
>ydhZ
GAGGCAATTCCGACGTCTAAGAAACCATTATTATCATGACATTAACCTATAAAAATAGGCGTATCACGAGGCCCTTTCGTCTTCACCTC
GATGGATCCGTTTTCGGTCCGATGGTGCAAACAATTTTGGTCTTTTTCATGACAGTCTTAGTCTTTAAGTTGAGAAGGATGGGAGAAAC
TTGCTTTCTGGGCGCACTGCCGGAAAGCGAATCCGGTATTACGAAAGTGGTGGTGCCTCGCTCTAAGGATAGGTGACATCGAAAAAGCG
TGCAGAGGAATGTGTGCTTGTGTCTCAGCCCAACGTGAGTGCCTGTTTTTGAGTCGTTGAGTTCTACAGTCCAAAGTGCTGAAACCATT
CAAGAGTCAATTGGCGCGCATTATACGCTGAAATTAATCCAAAAGGAAAACAAAAACAGCGTTTCAAAAAATATACAACTTGTTAGACA
ATAAATTGTTATCAAAGATTTAACGTTCCAGGGAAAAGGTTACGCAAAAACTTCACTGTTTGCGCAACGTCTGGTGCCGAAGTTGTCAG
TGTGAGGATAAGTGAAAGTTCATCCGGTGTATAAATATATCATCGCTCAATTGCTACAGTTAAAATAATTTATCGCGTTATTTGTCTGA
TTTTGTTTGATTTTTGTTTAATTGGTTAGATTGATATATGCTGATAGTTGGCATGATAAATAGGTATAAAGAATGGGAAATCGGACAAA
AGAGGATGAACTGTACCGAGAAATGTGCAGAGTTGTCGGTAAAGTTGTGCTGGAAATGCGCGATCTGGGGCAGGAACCACTCGAGAGAT
CCTCTAGATTTAAGAAGGAGATATACATATG
>dgt
GAGGCAATTCCGACGTCTAAGAAACCATTATTATCATGACATTAACCTATAAAAATAGGCGTATCACGAGGCCCTTTCGTCTTCACCTC
GAGCAACCGCCGAGACTGATAGTTTGACGGTTTTCGATTTTGTCACGCAGCAGCGTAACTTCTTCTTCCATTGCACCAATGATGCCGAT
TTTCATAGATTTACTCGCGATAAGCCCGATTTGAAGGCATAGTTTACCATGCGCTTACGGGGAAGCGTATTTCTCACGCGGGAGAGGAC
ATGGCACAGATTGATTTCCGAAAAAAAATAAACTGGCATCGTCGTTACCGTTCACCGCAGGGCGTTAAAACCGAAGGATCCCGGATCCT
CTAGATTTAAGAAGGAGATATACATATG
>ydhS
GAGGCAATTCCGACGTCTAAGAAACCATTATTATCATGACATTAACCTATAAAAATAGGCGTATCACGAGGCCCTTTCGTCTTCACCTC
GAGCGGCGTTGAGGAAGTTGTCGCCAAAGTTTTCGATGTCAATGAGCCACTTAGTCAAATCAATCAGGCAAAACTCGCCTGACAGAATT
TAATCAAGGGCGGTTAGCGCCCTTTTCATCCCTGTCTGAAATTTCTCAAATTCTAAAAATCTCAACCAAACTTATCTGATAACACTAAA
TTCGAAAGAATGCGTACAGGTAAGTAACAATGAAAAAAATTGCTATTGTGGGTGCCGGGCCTACGGGGATCTACACCTTATTCTCGCTT
CTACAGCAACAGGATCCTCTAGATTTAAGAAGGAGATATACATATG
>ydhR
GAGGCAATTCCGACGTCTAAGAAACCATTATTATCATGACATTAACCTATAAAAATAGGCGTATCACGAGGCCCTTTCGTCTTCACCTC
GAGGTCCAATTGAACGCACCATCCTGCGTCCGTAGCTCAGTTGGTTAGAGCACCACCTTGACATGGTGGGGGTCGGTGGTTCGAGTCCA
CTCGGACGCACCAGATTTTCTTAATCTGGTCTTCTCCTTTTTCCCTCTGTTTCTTCTCTGTATCCAATACGTTAAAAGATTTACACTGT
CTTCGTATGCGTTATCAGAAGGAGAATCGCTATGGCAACTTTGTTACAACTTCATTTTGCTTTTAATGGCCCATTTGGCGACGCAATGG
CTGAGCAGCTTAAACCGGATCCTCTAGATTTAAGAAGGAGATATACATATG
>ydhQ
GAGGCAATTCCGACGTCTAAGAAACCATTATTATCATGACATTAACCTATAAAAATAGGCGTATCACGAGGCCCTTTCGTCTTCACCTC
GATGGATCCAGCTACGGACGCAGGATGGTGCGTTCAATTGGACTCGAACCAACGACCCCCACCATGTCAAGGTGGTGCTCTAACCAACT
GAGCTATGAACGCAACGTTGTAGGTGACAACGGGGACGAATATTAGCGGCAGAGTGGGAAGGTGGCAAGAGGCAAAACGTAATTTTCTG
CGCTATTTCGACCGTTTGCAGAGCTTTTAAGCAAATTGGCTATATTTTGTTGATTTGCAAGGGTGATTTTTATTCAGGATCGCATTTAC
ATCTGATACAACCCAATAATTCGTGTATCAGAGTTTAAACAGAGTTGTTTGCCACCATCTCAAATCTGTCGTGGATAAATCCTCCGTAG
CCTTTACACTTAGGTAGCAATAAAGGCACACAGATGCAGCAAAGGAATAAATCATGGGATCTGATGCGAAAAACTTGATGAGCGACGGG
AACGTGCAAATTGTTAAGACCGGCGAGGTCATTGGCGCGACGCAACTTACTGAAGGCGACTCGAGAGATCCTCTAGATTTAAGAAGGAG
ATATACATATG
>ydhP
GAGGCAATTCCGACGTCTAAGAAACCATTATTATCATGACATTAACCTATAAAAATAGGCGTATCACGAGGCCCTTTCGTCTTCACCTC
GATGGATCCAACGTTTGCTCGTTTCGCTACATCTTTTATTGTTGCCATTCCATTTCACTCCAGACCCTAACTTCAGCTCCTGCAAAATC
GCAAGGTAAACGTTTGCCTTTACACACCTTACGCACAACAAAAGTGTTGCGGTACGCCGGAAAAACGACACGGAGCGTCAGGAAGGGGT
CAATGGCCAGTACGCAAATAGTGTAAGCGTGGAATTTTGGCCGATCTTGACGAAAAGGGGAAGTGGTAAAAACACTTATCATTATAAAT
GACGGGAGATTTTTTCATCACAGTGTGTAAAAATGCGTAATATCACTTTTTGTGTGGTAATAAAAAAGGAGAAAACTTGATGAGCACCG
ACCTTAAATTTTCACTGGTAACAACGATTATCGTCCTCGGTTTGATCGTAGCCGTGGGTTTGACTGCCGCGCTGCACTGATTTCCTCTC
GAGGGAGCGCGTTCTCCCTCGCATCTCCTTCCTGTTTGTTAGCAAAATGGCAAAAGTGTTTTCTCTTTTTGTTATTTTCATTTTTTTGT
GATTGCTGTCATGCTTTCGCCACTTTAATGAATATCGCGTAATGACGCGAAGAGGTGGAGATAAGGCATGAAAATTAACTATCCGTTGC
TGGCGCTGGCGATTGGCGCGTTTGGTATCGGGACAACGGAGTTCTCGCCAATGCTCGAGAGATCCTCTAGATTTAAGAAGGAGATATAC
ATATG
>yafL
GAGGCAATTCCGACGTCTAAGAAACCATTATTATCATGACATTAACCTATAAAAATAGGCGTATCACGAGGCCCTTTCGTCTTCACCTC
GAGTCAGAGATGGTCAGCCCCATCCCGGCCAGTACGTCCGCTGCCTGATTCTTCAGATCTTCATCGATTCGGGCGCGAACAAACGCGTT
AGCAGCCATTGGGGTATCTCCTGTGCTGTATATTTATTCAGCTTGAATTGTAGCTCAAATGAGCAACAGCGACAATCCATGAGTAATTT
AATAAACGGATGTTTTTACATAATTAACTGAAATATATATGTTAATTTTATAATAATGGTTTATTAAGTTTGTTCAGAAACGATCCGGG
ATACACTGCAACATTAAAGCATGACCAGCCATTATTGAGTATGTCCTTGCCGTCGATTCCATCGGATCCTCTAGATTTAAGAAGGAGAT
ATACATATG
>yafM
GAGGCAATTCCGACGTCTAAGAAACCATTATTATCATGACATTAACCTATAAAAATAGGCGTATCACGAGGCCCTTTCGTCTTCACCTC
GAGAAGCCGATTAGCCGAACCTTTCTGGCAGGACCATTTTTTGGGCGCGCGCAGGATTTTGACGGAAGAGACGATTTTGTAGGACGGAT
AAGGCGTTTACGCCGCATCCGGCAGTTGTACGCAGGTGCCTGATGCGACGCTGGCGCGTCTTATCATGCCTACGAGCCCGCGAATATTT
GCGAGCCGCTTTCCCGATATAAAACAACCTCATTGCCAACCTTTCCTTTTCTTCTTACCGTTGAGAAAAAGGAGTCGCCATGTCTGAAT
```

Data Set S4 - MG1655 library reference file.fasta

```
ATCGTCGTTATTACATCAAGGGGGGAACATGGTTTTTCACGGTGAATTTACGAAATCGTCGAAGCCAACTTTTGACCACCCGGATCCTC
TAGATTTAAGAAGGAGATATACATATG
>yafN
GAGGCAATTCCGACGTCTAAGAAACCATTATTATCATGACATTAACCTATAAAAATAGGCGTATCACGAGGCCCTTTCGTCTTCACCTC
GAGGCGGTGTGCGTCTGGTGGGGCTGCATGTGACGTTGCTTGACCCGCAAATGGAAAGACAACTGGTGCTGGGATTATGATGTATACTA
TTATGTATATTCTGGTGTGCATTATTATGAGGGTATCACTGTATGCATCGAATTCTCGCTGAAAAATCGGTCAGGATCCTCTAGATTTA
AGAAGGAGATATACATATG
>zitB
GAGGCAATTCCGACGTCTAAGAAACCATTATTATCATGACATTAACCTATAAAAATAGGCGTATCACGAGGCCCTTTCGTCTTCACCTC
GATGGATCCAAAGATGGTCGCTGTCCGGATATCAATAAAAAGGTACAAACCGGTGATGGCATCAATAATGATGTCGATACCAAAACCGA
CGGTACCACGCAGTAACCCTGCAAAATAGCCAATAATGGGAGAGCGGAGTCTCTCCCTTTTAATTTTTCACTTCTAAAAGCAGTATGAT
GTCATAAAGTTAAGTGAGAATAATTAAGGGATGACTCGTCATGGCGCACTCACACTCACACACGTCCTCACACCTGCCAGAAGATAATA
ATGCTCGTCGCTTGTTGTATGCTTTCGGCGTGACTCGAGAGATCCTCTAGATTTAAGAAGGAGATATACATATG
>yafJ
GAGGCAATTCCGACGTCTAAGAAACCATTATTATCATGACATTAACCTATAAAAATAGGCGTATCACGAGGCCCTTTCGTCTTCACCTC
GAGGTTATGCCGACCGCATTCAGGAGATTCACATTAAAGTGATCCATATCCTGATCCAGTTGATTGAAAAAGAGATGGTTAAGTAAGTC
TGGCGTAGGCCGGATAAGACGTTTACGCCGCATCCGGCATTTGTGCGCTGATGCCTGATGCGACGCTGACGCGTCTTATCATGCCTACA
AATCTGTACGCGAACCGTAGGCCGAATAATGCGTTCACGCCGCATCCGACCTGAAAATTCTTAAATCAATCTTCGCCGGGGGCCATGCG
CTCCCGCTGTTGTGGAGGTTACCCATGTGCGAATTGCTCGGGATGAGCGCCAACGTCCCTACCGATATCTGCTTTAGTTTCACCGGGCT
TGTACAGCGTGGTGGATCCTCTAGATTTAAGAAGGAGATATACATATG
>yafK
GAGGCAATTCCGACGTCTAAGAAACCATTATTATCATGACATTAACCTATAAAAATAGGCGTATCACGAGGCCCTTTCGTCTTCACCTC
GATGGATCCTGTCGAACCGGACTGGATCCTGATTTACAAACTTACCGATAAACTTTTACGATTTGAGAGAACTGGAACTCACGCGGCGC
TCTTTGGGTAAAGCCCGCTTCCGATGAAATTTGTAAACCCTGCGCCGACAGGCTTGCGTAAAACTCTTTTGGTGATAATGTGAGCCTTC
CAAATCCAGACTATTCCGATTTGGGATCCATTTTGCCGTCCTGGCGTGTGTAACCGTTTTATCAAGGAATAAGCAGTATGCGTAAAATC
GCATTAATTCTTGCGATGTTGTTGATCCCGTGCGTTTCATTTGCCGGTTTGCTGGGCAGCAGTAGCTCTACCACGCCTGTGAGCAACTC
GAGAGATCCTCTAGATTTAAGAAGGAGATATACATATG
>yafD
GAGGCAATTCCGACGTCTAAGAAACCATTATTATCATGACATTAACCTATAAAAATAGGCGTATCACGAGGCCCTTTCGTCTTCACCTC
GAGACCGCTGAGTTTGCTTGCCCTAATTGTTCCGCTGCCCGGCTAAAGCTGCCGCTTTCGACGACCGAAACAAAAATGGCGAGTTCTTC
CGACGTGGCTTTCATTTTTGCTCCTGTTGCAAAATAGAAGAGATATTTTGAATTTATTTGTCATTAAACCATCAGGATGCGTGATATGT
CATGCGATTTAATGTTCTCCATAATGAGCAAAATTCTGACCGGTGTAAGCACTTGCTTACATAACAATATACAATTGCTCGTTGAAAGA
GTGAGCTAAAATCCCTATAACAGTAGAACCCTCCCGAGTGCGGAAGGGTTGACGTAATAGAGGTTTCAAAGTCAAAAGTGCGAAAAAAC
ACCTATGCCATGCGCTATGTTGCCGGACAACCTGCGGAAAGGATCTTACCGCCGGGGTCTTTTGCGAGCATCGGGATCCCGGGATCCTC
TAGATTTAAGAAGGAGATATACATATG
>ycfT
GAGGCAATTCCGACGTCTAAGAAACCATTATTATCATGACATTAACCTATAAAAATAGGCGTATCACGAGGCCCTTTCGTCTTCACCTC
GATGGATCCGGAGACGAAACGACCGAAGCGATCCGCTGCACGCCCACGCATGTAACGCAGGCCAATAAATAGAGCGACAGGTTGGTACA
TGAAATCCGTCTGGTTGCTGTAGCAAAGACACGAGTATATAAGCGAAAGCAAGATGGGTAAACGTCTGAACTGTAAGTGATCGTCAGGG
CTAATCTGAAAAGCCCCCGTAAATCAATCGGCTAAATTGCCATGTTTTATCATCAATCTTGCCGCCTGATCTTAATCTGAAAAATCATC
CAATACAGACAGATACCCATTACACCTCGGGTCATAACCCGCAGGATCAACCGCATGTAAATTAAAAAAGAGCGTGTTGTCCTGATGAA
ACAAAAAGAGCTATGGATTAACCAGATCAAAGGGTTATGTATTTGTCTGGTGGTGATTTATCACTCGGTCATTACCTTTTATCCGCATC
TGACCCTCGAGAGATCCTCTAGATTTAAGAAGGAGATATACATATG
>mipA
GAGGCAATTCCGACGTCTAAGAAACCATTATTATCATGACATTAACCTATAAAAATAGGCGTATCACGAGGCCCTTTCGTCTTCACCTC
GATGGATCCCGTCCTTGGCAGCTTCATATCGCTGGCGATAGTGATCGAATATATTCATGGTATGCCGTCCTTTCGTTTTTTAGCACAGG
TTAAGAGCCGTTCATATGAATAGTAGAGGCTCCCGGAAGAGGTAAACTGAACGACGTCTCTTGCTAAAACACAGCAACTCTCATGCCAT
TTTATCGACGATGAGAACAACGTGCCCGGTGATACACCTTCAAAATTAAGCGTAGATGGCAAATGGAAAACTTGCCTGCACGGCTTTCT
TAATTTCAATGGCATATCAACAACTCATCCATAAATTTTGCATAATTAATGTAAAGACCAGGCTCGCCAGTAACGCTAAATTCATTTGG
CTGTAAGCGCGGTGTCATCCGCGTCAGGAAAATTAAACAGTTACTTTAAAAAATGAAAACGTAAAAAGGTTGGGTTTCGATGTATTGAC
GGGTAAACTTTGTCGCCCGCTAAACATTTGTTTAAGGAATGATTAATTGTGACCAAACTCAAACTTCTGGCACTTGGAGTGCTTATCGC
AACGTCTGCAGGCGTAGCGCACGCTGAAGGTAAATTCTCGAGAGATCCTCTAGATTTAAGAAGGAGATATACATATG
>ycfR
GAGGCAATTCCGACGTCTAAGAAACCATTATTATCATGACATTAACCTATAAAAATAGGCGTATCACGAGGCCCTTTCGTCTTCACCTC
GAGAGTTGCCATGATGCCGTTGTACCTGGTGACTGTGAATGAAAGGTTATTATAAAAATAATCACCTCCGTTCACCAGTCCAGATCCCA
TAAAAATAATTGCTTTCTATTTAACTGAAATTTAAAGATTTTTAAATTAATTAATGATTGTTATAAAAAATATCTTGTATGTGATCCAG
ATCACATCTATCATTTAGTTATCGATCGTTAAGTAATTGCTTGCGACGTCATTCATCTGCATAAGGCCACTATTATGAAAAACGTAAAA
ACCCTCATCGCTGCGGCGATTTTAAGCTCCATGTCATTTGCCAGCTTTGCGGCTGTCGAAGTTGGATCCCGGATCCTCTAGATTTAAGA
AGGAGATATACATATG
>ycfS
GAGGCAATTCCGACGTCTAAGAAACCATTATTATCATGACATTAACCTATAAAAATAGGCGTATCACGAGGCCCTTTCGTCTTCACCTC
GATGGATCCAAACGCGTATCGAATGGGTACGCCAGTTTATGCGTGAACTGGAAGAGAACGCGATCGCTTAATGGTAAATCGGGGGCGTT
TCTGCGCCCCCATCTGCAACATTTACAAATTCTTTGCACTTCCCTGCACTATCCGACACTGTCACCATCCATAATTCAGGCTTATCTGT
```

TTATTACAATAACCTTATATTTATTATGGATTTTTGGTGATGATCAAAACGCGTTTTTCTCGCTGGCTAACGTTTTTTACGTTCGCCGC
TGCCGTGGCGCTGGCGCTACCGGCAAAAGCCAACACCTGGCCTCGAGAGATCCTCTAGATTTAAGAAGGAGATATACATATG
>slyX
GAGGCAATTCCGACGTCTAAGAAACCATTATTATCATGACATTAACCTATAAAAATAGGCGTATCACGAGGCCCTTTCGTCTTCACCTC
GAGTGCAGCTTCAGCAGCAAAAGTGATTGGTGCATGCAGGGCAACGGCCATTGTGGTCGCCAGCAGCGTTACTTTAAACAGTGATTTCA
TCCATATCTCCAGGATCGGGGCATCTCACCCCAGGGTTAACTATTATCAGAAGGGTACTATAAAGCGTTGTCGAACAAATCTACATACA
GACACGCCCTATTATCATCTATTTTCAGACTCTTTTTGTTTAAATTAGTTTCGATGACCGCGAAATGAGTGCTGTCTCGGGCAGCAAAG
TTAAGTAGAATCCGCGGCGACCATTCGACAAAAGAGGTGAATCATGCAGGATTTATCATTGGAAGCACGCCTGGCAGAGCTGGAGAGCC
GACTGGCTTTTCAGGAAATCACCATTGAAGAACTGAACGTCACGGGGATCCTCTAGATTTAAGAAGGAGATATACATATG
>yafC
GAGGCAATTCCGACGTCTAAGAAACCATTATTATCATGACATTAACCTATAAAAATAGGCGTATCACGAGGCCCTTTCGTCTTCACCTC
GATGGATCCCCGGCGGTAAGATCCTTTCCGCAGGTTGTCCGGCAACATAGCGCATGGCATAGGTGTTTTTTCGCACTTTTGACTTTGAA
ACCTCTATTACGTCAACCCTTCCGCACTCGGGAGGGTTCTACTGTTATAGGGATTTTAGCTCACTCTTTCAACGAGCAATTGTATATTG
TTATGTAAGCAAGTGCTTACACCGGTCAGAATTTTGCTCATTATGGAGAACATTAAATCGCATGACATATCACGCATCCTGATGGTTTA
ATGACAAATAAATTCAAAATATCTCTTCTATTTTGCAACAGGAGCAAAAATGAAAGCCACGTCGGAAGAACTCGCCATTTTTGTTTCGG
TCGTCGAAAGCGGCAGCTTTAGCCGGGCAGCGGAACAATTAGGGCAAGCAAACTCAGCGGTCTCGAGAGATCCTCTAGATTTAAGAAGG
AGATATACATATG
>efp
GAGGCAATTCCGACGTCTAAGAAACCATTATTATCATGACATTAACCTATAAAAATAGGCGTATCACGAGGCCCTTTCGTCTTCACGTC
GACATCGGTCACAACATCGGCAAGTTGCGTTAACCAATCTTCTCTGGATGGGGTATTTAGGGTTACAATATGCGCCATTTTGTGGCTTA
GCTACCAATTAACAAATTTCAGAGGGCCTTATGGCAACGTACTATAGCAACGATTTTCGTGCTGGTCTTAAAATCATGTTAGACGGCGA
ACCTTACGCGGTTGAAGCGAGTGGATCCTCTAGATTTAAGAAGGAGATATACATATG
>slyD
GAGGCAATTCCGACGTCTAAGAAACCATTATTATCATGACATTAACCTATAAAAATAGGCGTATCACGAGGCCCTTTCGTCTTCACCTC
GATGGATCCAACAAGAACGACGCCAGCTGGACGGCTGGGATGAATTTGAGTAGAGGGTAAAGATGGCAATCCGAAAACGTTTTATTGCG
GGCGCAAAATGCCCGGCCTGTCAGGCGCAGGATTCAATGGCGATGTGGCGCGAAAATAATATTGATATTGTTGAATGTGTTAAGTGCGG
ACATCAGATGCGAGAAGCAGACAAAGAAGCCCGCGATCACGTTCGCAAAGATGAGCAAGTGATCGGGATTTTTCATCCGGACTAGCGAT
ATGCGCCGAGTTTTTTTAAGCTAGTGAGTACACGGCTGCAGAATTCCGCTACAATCTGCGCCACTATTCTTCCCATGCTCAGGAGATAT
CATGAAAGTAGCAAAAGACCTGGTGGTCAGCCTGGCCTATCAGGTACGTACAGAAGACGGTGTGTTGGTTGATGAGTCTCCGGCTCGAG
AGATCCTCTAGATTTAAGAAGGAGATATACATATG
>ycfJ
GAGGCAATTCCGACGTCTAAGAAACCATTATTATCATGACATTAACCTATAAAAATAGGCGTATCACGAGGCCCTTTCGTCTTCACCTC
GAGAGATTGCGCTGCATGGTTACTTTAAAACCGGATTAATGATGCTGGTGGGGAGTATTAACCGCGTTATCCGTCCGCGTTTGAAGTTG
CATTAATCGACGTACACTGGCGGATGTGGCATAAACGCCTCATCCGCCCTTGAGGAACAGCGCGATCGGCAGCCGCGTTGTATCAGGCA
TCCTTTCAGACTCCTCCGAATCCTTAAGTATTTCCAGCCATTCCCGCGCTTTTCATCTTCTGTCTGATAGCTGCTTTTCTCCTTCGCTT
GCATGATTGGCATAACTGCAAAGAAGGAGGTGTTCCCGTGAATAAATCAATGTTGGCGGGTATCGGGATTGGTGTCGCAGCTGCGCTGG
GCGTAGCGGCAGTGGCCAGTCTGAACGTGTTTGAACGGGGGGATCCTCTAGATTTAAGAAGGAGATATACATATG
>yafZ
GAGGCAATTCCGACGTCTAAGAAACCATTATTATCATGACATTAACCTATAAAAATAGGCGTATCACGAGGCCCTTTCGTCTTCACCTC
GATGGATCCCCTTTCCCCTGATACCCGCCCCGGTGCGGGCCGTCATTCAGGCCGTGCGCACCACGGTGGTGTCAGTGGCCCGCGCCGTC
TGGGATTTCTTCTTCTGAGTATTTAACCCATCCCTGTTACTTCCCTGCCCCTGTCGCCGATGCGACGGGGATCCCTTTTATTTGTTTTC
CCGTTATGAGGAGTCTGCTTATGACCCGTCTGGCTTCGCGCTTTGGCGCAGCAAACCTTATCCGTCGCGACCGTCCGTTAACCCGTGAA
CTCGAGAGATCCTCTAGATTTAAGAAGGAGATATACATATG
>slyA
GAGGCAATTCCGACGTCTAAGAAACCATTATTATCATGACATTAACCTATAAAAATAGGCGTATCACGAGGCCCTTTCGTCTTCACCTC
GATGGATCCACGCACCGAACACCAGATCTTGTAAGGGCAATCCTGTTGCATTGAGCATAAACTTCACAAAGTAGATTCCTTTACGACCG
AATATATTGCGTTAGATTAATAAATATTCTTTAAGTGCGAAAAATTTACGCGCAATTTCTGAAAAATATGTCTGACGGTAACCAAATGC
AGCAATACATTTGTTTTAGCAATACAATTGCTGCACACTATTCTAAAAGCCGCATAATATCTTAGCAAGCTAATTATAAGGAGATGAAA
TTGGAATCGCCACTAGGTTCTGATCTGGCACGGTTGGTGCGCATATGGCGTGCTCTGATAGACCCTCGAGAGATCCTCTAGATTTAAGA
AGGAGATATACATATG
>yafT
GAGGCAATTCCGACGTCTAAGAAACCATTATTATCATGACATTAACCTATAAAAATAGGCGTATCACGAGGCCCTTTCGTCTTCACCTC
GAGTCGTTCCCAACGGAACACAACGCGGAGCGGTAGTTCAGTCGGTTAGAATACCTGCCTGTCACGCAGGGGGTCGCGGGTTCGAGTCC
CGTCCGTTCCGCCACTATTCACTCATGAAAATGAGTTCAGAGAGCCGCAAGATTTTTAATTTTGCGGTTTTTTTGTATTTGAATTCCAC
CATTTCTCTGTTCAATGATTTTACTCTGGCGTAGGTGCGTGATTCTCGCTTGTTGTCTCATTCATTAAAATTCAATAATGATATCGAAC
CATTCAGCTTAAATATATTTCTAGAGAATAAATTTATATTGATTAAATGAATGTATATTTCAAATTGATTTTGTTTGTTATTATTTAAG
TGAGGTATATAATTAGAGTCCGTTATCAATGCTAAATATTCTAATCATTATGACAGGCGAGGGAGTGTCCAATTATGAATTCAAAAAAG
CTTTGTTGCATATGTGTGTTATTCTCGCTGCTTGCAGGATGTGCCTCTGAATCTTCGGATCCTCTAGATTTAAGAAGGAGATATACATA
TG
>yafU
GAGGCAATTCCGACGTCTAAGAAACCATTATTATCATGACATTAACCTATAAAAATAGGCGTATCACGAGGCCCTTTCGTCTTCACCTC
GATGGATCCCTGTCGATGGCAGCATTGCGGGAATATAGAGAAAAGTTTCCGGCATGGCAGGATGCGGATGAGTTTAGGTTGTGGTGAGT
GGGGGTTGTGTTTAAGGACGGGGAGAGTCGGGGTATTACTACGAAAGCCCGCTCCCCGCAAGGACTGACGCCAGGTAGTTTCTGTCCAT

Data Set S4 - MG1655 library reference file.fasta

GGCTGCTTTTCGCATCTTACGTCTTAACCCTGCCTTGAATACCTTCTCATTCGTCAAAATATTAATAGCAATGTGCCGTATCCCTGAAA
ATAATTCTGCTGCGTTTCCTCTTCTTATTTGTAGTCGTCTTCATTCATTACCACGTCCAGACGCCTGTGCAGCTTATTCTCCACGTGC
CAGTGGTTTCGGATTGCTGTGGCGAACTTCTCTGCGGTTAAATCAGCAGAACCATAGAATTTATTATGAGTTCAGAGCGAGATCTGGTT
AATTTTCTTGGCGATTTTTCAATGGATGTGGCCAAAGCAGTTATAGCCGGTGGTGTTGCAACCGCTACTCGAGAGATCCTCTAGATTTA
AGAAGGAGATATACATATG
>yafV
GAGGCAATTCCGACGTCTAAGAAACCATTATTATCATGACATTAACCTATAAAAATAGGCGTATCACGAGGCCCTTTCGTCTTCACCTC
GATGGATCCAACATCATTCCTCCCGAGCTTATCCTGCCCATGTTATTAACCTCCTAAAGCCTGGCGATAACACGATAATTTGCAAATCA
GATATTTCCAATATCAGCATTTGTTATCAATTTATTCTGCGCTAAGGCAACTGCCTGATAAGGTGTTAAAGTTATAAAACCCTTCGCAT
AAAGGAGTTGTACACCGTGCCTGGTTTGAAGATTACGCTTTTGCAGCAACCACTGGTGTGGATGGATGGTCCTGCCAACCTGCGTCATT
TTCTCGAGAGATCCTCTAGATTTAAGAAGGAGATATACATATG
>uxaC
GAGGCAATTCCGACGTCTAAGAAACCATTATTATCATGACATTAACCTATAAAAATAGGCGTATCACGAGGCCCTTTCGTCTTCACCTC
GATGGATCCTACGACCGGGTAACCGAAAAAAAACCGGCACGCCCCGAACGTTGCCATGAAAGTTTCGTGCTAATTCGGCTTCCGTACCG
GTTAACCCGATTCATCTAACAAATGAATGAATCGGCAATTTGCATTCCGTCGACTTATGATTTGCGACGGCAGAAAGATAACTTGTCAT
ACAACTTTAAAAGGTGAGAGCCATCACAAATGTGGGAATATTTGTAGGGACATTACCTGACGACAGCAAGGCCAGTACTGGCGCGGCCT
GCAGCGAGATTTACCACTTTGAGAGTAATTTTTTTAACTACGTTTATTGATCTAACTCACGAAAATATCTTCGGACTCTGGAAATTGGT
GTGATAACTTTGTCAGCATCGCACCATAAGCAAGCTAGCTCACTCGTTGAGAGGAAGACGAAAATGACTCCGTTTATGACTGAAGATTT
CCTGTTAGATACCGAATTTGCCCGCCGTCTGTATCACGACTACGCAAAAGACCAGCCGATTTTCGACTCGAGAGATCCTCTAGATTTAA
GAAGGAGATATACATATG
>flgM
GAGGCAATTCCGACGTCTAAGAAACCATTATTATCATGACATTAACCTATAAAAATAGGCGTATCACGAGGCCCTTTCGTCTTCACCTC
GATGGATCCCAGGTAGTCAGCGGCGTTGTTGATGCAGATGGGAATATTCTTATAAACCTGTAAGCTGTTAAAGATTACCCGTCCCTTGC
CGATAAATAAGCAACACATGATAAAAGCGCCCTCAATGAGGAATAAACCATGAGTATTGATCGCACTTCGCCTCTGAAGCCTGTAAGCA
CCGTTCAACCGCGCGAAACCACTGACGCGCCGGTAACGAACAGCCGGGCTCGAGAGATCCTCTAGATTTAAGAAGGAGATATACATATG
>yafS
GAGGCAATTCCGACGTCTAAGAAACCATTATTATCATGACATTAACCTATAAAAATAGGCGTATCACGAGGCCCTTTCGTCTTCACGTC
GACCAGCGACCTGCTTCATCATTCAAAACCCAGATGTAATTGTCATCAAAGGCGGGAATACTGTTAAGATTCATAGATTACCTCTCAGT
GTGAAACGGAAGGTTGTGATGAAACCGGCAAGAGTCCCTCAAACTGTCGTGGCTCCTGATTGCTGGGGCGATTTGCCCTGGGGAAAGCT
TTAGGATCCTCTAGATTTAAGAAGGAGATATACATATG
>sscR
GAGGCAATTCCGACGTCTAAGAAACCATTATTATCATGACATTAACCTATAAAAATAGGCGTATCACGAGGCCCTTTCGTCTTCACCTC
GAGGCAAGCTGGGTGGGAGTTAAATCGGTCGTGGCCGCCTGAAGGCGTGCCAGTTGCTCCGGGTTCAACGGAAGCAACGCGGAAGGTGG
GACCTGTGTCGTCATGCGTCGTTATGTTCCAGTAAGCAAAGCTGTTTCTGCGCCCTGTCAGCGCCCATAAAACAGAAGAGAAGGTAAGG
TTAACGGGGCAAACGGTGTGGATTAAAGACGGGATAGCGATAACTAATAACCAAATCGACTAACCTGTTTTAGCAATAGTCTTTAACAA
CAAAATAGATTAACCAACCTAATGAAAAACAAATGAATTTAGCCAATCATTAAGATAAATCAGCGATTTTGCGCAACAAAGTCGTTTTA
GATAATGCGAAAAAACAGCCTTTCCGGTACTCTACGGCGGTTTTATCGTCCTGTAGAGAAATTATGATGTCCACCACGTTATTTAAAGA
TTTCACCTTCGAAGCCGCTCACCGCTTACCACACGTCCCGGAAGGGCATAAATGTGGTCGCGGATCCTCTAGATTTAAGAAGGAGATAT
ACATATG
>dhaM
GAGGCAATTCCGACGTCTAAGAAACCATTATTATCATGACATTAACCTATAAAAATAGGCGTATCACGAGGCCCTTTCGTCTTCACCTC
GATGGATCCATGATGCAAATGTTGGCGTTAGCCGCAAAAGAGTAAGGAATTGGTGATGGTAAACCTGGTCATAGTTTCACATAGCAGCC
GACTGGGAGAAGGTGTCGGTGAATTAGCCCGTCCTCGAGAGATCCTCTAGATTTAAGAAGGAGATATACATATG
>dhaR
GAGGCAATTCCGACGTCTAAGAAACCATTATTATCATGACATTAACCTATAAAAATAGGCGTATCACGAGGCCCTTTCGTCTTCACCTC
GAGGCCAGTCCTGCCAGTTGTTCGTCCAGTACGTCTTGCACATCATTGATCAATTTTTTCATTATTTTGCTCCAGCAATTACGGTAGGG
CATGGATGATGTTCAACGACACGGCGACCACTGACTGCCGATGAATCCATTGTGCATCAGGCAAGGGAAAAGAAAATTAGCCCAAATTA
TGTTTCATAGTGAAACATATGCTTTAATGAATGTTCCATATTGAAACTTTTACGTGTATTAATACTTAAAATTGCGAGCCGGAACACCT
TTTGTCATAAGGGATGCGGGATATGAGTGGCGCTTTTAACAACGATGGTCGGGGCATATCTCCCTTAATTGCAACCTCCCGGGAGCGAT
GCAATAAGCGGATCCTCTAGATTTAAGAAGGAGATATACATATG
>yjbF
GAGGCAATTCCGACGTCTAAGAAACCATTATTATCATGACATTAACCTATAAAAATAGGCGTATCACGAGGCCCTTTCGTCTTCACCTC
GAGCGTAAGCTCGGCGGTGGGTGTCGCGCTTGCGGCAACCGGTGGCGGTGATGGTTCTAATACCGGGACCACCACAACCACGACCACCA
GTACCCAGTAATAAAGTATGTATCCCCAAAATAATTCGAGTCATTGCATCTGTGGCTAGAAGTATGAAGGGATTAACCATAACCACACT
CCGGTGTGGTTATTCTGCCCCTCTGGAGAAGAGTCGTGAAGCGACCTGCACTCATTCTTATCTGCCTGTTATTACAGGCCTGTTCAGCC
ACGACTAAAGAGCTGGGGATCCTCTAGATTTAAGAAGGAGATATACATATG
>yjbE
GAGGCAATTCCGACGTCTAAGAAACCATTATTATCATGACATTAACCTATAAAAATAGGCGTATCACGAGGCCCTTTCGTCTTCACCTC
GAGGAACCGTATTCTGCCAGAGCTGAAAGATGATAAAGAAATCAGCAGCCACGATAGCTCGACCAATGGTCTGATTAACCGCTATAAAG
CGTGGCGCGGTTAATCATCGTCGATATGTAGGCCGGATAAGGCGTTCACGCCGCATCCGGCAACCGATGCCTGATGCGACGCGGTCGCG
TCTTATCAGGCCTACAGGTCGATGCCGATATGTACATCGTATTCGGCAATTAATACATAGCACGATTGATTAAATAACCTTAATAACAA
TGCCGACGTTATGTCGGCATTTTTTTATCAGATAAATCCCCTTGTCTGTAATTTAACGGAAATCATACCGTGAGGTTAATCCTAAAATA
GATTTTTAATCGTTGTTTATTTCGGAAAATACGCAGATTAATTGCTTTTGTTTTTATTTAAGTTTATGATTTTTATTGTTATTTAAAT

ATAAGTTGAAACTTATATTTGATATTCATTCCAATTATCCTAAAACGCCATCGCTAATTCCCCGCGCCGTAATTCGCATGCTTTAGTTG
TGTATACTCGATCCCGCCCGAAATGTTTTTGGGTAAATCTCCATTCATTCAATGAAGGGAAATTGTTATGAAAAAAGTTCTGTATGGCA
TTTTTGCCATATCTGCGCTTGCGGCGACTTCTGCGTGGGCTGCACCTGTACAGGGATCCTCTAGATTTAAGAAGGAGATATACATATG
>yjbB
GAGGCAATTCCGACGTCTAAGAAACCATTATTATCATGACATTAACCTATAAAAATAGGCGTATCACGAGGCCCTTTCGTCTTCACGTC
GACAGCGCGATCAGGTTGAAGATTATGCCCGCCGTAAAGGTATGAGCGTTACCGAAGTTGAGCGCTGGCTGGCACCGAATCTGGGGTAT
GACGCGGACTGATTCACAAATCTGTCACTTTTCCTTACAACAAACAGGGCGCTCAATGAGTGCCCTGTCTCTTTATTAATATGAAACAC
TTATACTGGAAACAGGCTGGAATAATCTTAGCCGGAAAAACGAATGTCAAAAGGCACAGTAAAGATGCCAGTCAGCGTGTTTCCGGTCC
TTTGGTCATTCAATTGCTTCAGGAACATATAACGATAAGGAGAACCTGAACTCGTGTTAACGCTGCTTCACCTGCTTTCTGCCGTCGCC
CTGCTGGTCTGGGGGACTCATATTGTTCGAACCGGCGTGGATCCTCTAGATTTAAGAAGGAGATATACATATG
>yeeX
GAGGCAATTCCGACGTCTAAGAAACCATTATTATCATGACATTAACCTATAAAAATAGGCGTATCACGAGGCCCTTTCGTCTTCACCTC
GATGGATCCGGCTGAACATGGAAACGGCGCATCAGCTTGAGTTGCTATCGAATCTGATTTGCCGGGCCTTGCGCAAATAATTCCTGAAC
TTCAGAATCATCTTGCTGCTGCTTCGATTCAGCAAGGATAAAGGGTATGATAGTGAAAAGGGATAAAAGCATTGTCATCTGCGGCAGCT
ATGAGTAATGTTGGCCCTAACGAATAGCGGTTGCTTAAACGAATCCGACTCTCACATTATCAGGGGTATAAAAATGGAAACTACCAAGC
CTTCATTCCAGGCTCGAGAGATCCTCTAGATTTAAGAAGGAGATATACATATG
>mreC
GAGGCAATTCCGACGTCTAAGAAACCATTATTATCATGACATTAACCTATAAAAATAGGCGTATCACGAGGCCCTTTCGTCTTCACCTC
GATGGATCCAAGCGCTGGAAATGATCGACATGCACGGCGGCGACCTGTTCAGCGAAGAGTAATCGGATGCAGGCAGGGGAAGTGTCTGT
TTACCCTGCCTGGTCTGATACGAGAATACGCATAACTTATGAAGCCAATTTTTAGCCGTGGCCCGTCGCTACAGATTCGCCTTATTCTG
GCGGTGCTGGTGGCGCTCGGCATTATTATTCTCGAGAGATCCTCTAGATTTAAGAAGGAGATATACATATG
>yjbJ
GAGGCAATTCCGACGTCTAAGAAACCATTATTATCATGACATTAACCTATAAAAATAGGCGTATCACGAGGCCCTTTCGTCTTCACCTC
GAGACCGTCTTTCTGGCGTTGCGCGGGCTTTCTCTGGCGGCTATCTGGCGGCGTCACTGGCGCAATGGTACCTGGTTTGCCGCAACGTG
ACGGTTAAAAATTCTGAATAAATAATCCTAAGCCAAATTGCTGACTACACTTAATCTCACGTTCAGAAGAAAAGTGAACGTACTCTCAT
TCACAACCTAACGATGAGGTCTTGATTATGAATAAAGATGAAGCCGGCGGTAACTGGAAACAGTTTAAAGGTAAAGTGAAAGAGCAATG
GGGCAAACTGACGGATCCTCTAGATTTAAGAAGGAGATATACATATG
>yjbI
GAGGCAATTCCGACGTCTAAGAAACCATTATTATCATGACATTAACCTATAAAAATAGGCGTATCACGAGGCCCTTTCGTCTTCACCTC
GAGATGAAGCTGAACGCCTGGGATCGACATCTGCCCGTTCCACCTTTATCAGCAGTGTAAAAGGCAAGGGGTAATTACGCCCCACAGTG
CTGATTTTGCAACAACTGGTGCGTCTCCTGGCGCACCTTTTTTTATGCTTCCTTCCTGGGATATGAGCGATTTTTTATAGTAACTCACT
TCTTCTTCACTAAGAATATCCATTATCTCAATGCCTTATCAGAGATTCTTTTCCTTTCGCCGGTAGTGTCTGGACATTCAGGCTACTTT
TCCAGGTTATTTTATTTCTGTTATGCAGAGGTTTTATGATAAGTCATATCCTAAATTCTGGCGGCAATAACTCTTTGATGAAACATGAT
GTGGTGCAAGGAAATAATATAGTAGATCTTGATTTACTACGTAATTTAAATGGGGTGCCAGGTTTAAACAGAGATAACTTTATTTATAT
CAGCAATATTTTTTCAAATATAAAACAACGGAACGAAAAAATCATGCAATAAATATGTTTCGTGAAGTCTCAATCAGTAATGATACTAT
AAGTGTAAAATTCTACAGAAATGAAAAAAATTGAATGCGCTTGCAATTTTCTGATGGATAAAGATGCGCAGGGGTATATCGACCTGTCT
GATTTGGATTGGATCCTCTAGATTTAAGAAGGAGATATACATATG
>yjdB
GAGGCAATTCCGACGTCTAAGAAACCATTATTATCATGACATTAACCTATAAAAATAGGCGTATCACGAGGCCCTTTCGTCTTCACCTC
GATTGATCACGCTGAATTACAACCGGCTACATAAAAACCCGTATCCCTTAGATGCACCAATAAGCAAAGATTAATTCCCCTTAATCCAG
CAAACATAAAAGCCAACCTTAAGAACTTAAGGTTGGCTTAATTTTGCTTTGCGAGCATATGCGCACTTTGTTCGATGGAAACACCGTGA
TGTTGAAGCGCCTACTAAAAAGACCCTCTTTGAATTTACTCGCCTGGCTATTGTTGGCCGCTTTCTCGAGAGATCCTCTAGATTTAAGA
AGGAGATATACATATG
>yjbQ
GAGGCAATTCCGACGTCTAAGAAACCATTATTATCATGACATTAACCTATAAAAATAGGCGTATCACGAGGCCCTTTCGTCTTCACCTC
GAGCGTGGTATCCGCATTCTGCGCGCCTCCAACTCTACCTACAAACCCTTGCCACAAGCGGGTGCGTTTGGTGAAGAGGTGATCGTCAA
TTCAGAATACTGACAGAGCGGGAGAGCGTGATGCTCTCCCGAATGCTGTTTTTTAATCACACCTTTATCCTTTCGCTGTCTTGCTGCA
AACTGATTAAGAGAGTTTTATCAAGGAGCAGCACATGTGGTATCAAAAGACGCTCACGCTTAGCGCCAAATCTCGTGGGTTTCATCTGG
TAACGGATGAAATTCTGAATCAGCTGGCTGATATGCCGCGCGTTAAGGATCCTCTAGATTTAAGAAGGAGATATACATATG
>deoR
GAGGCAATTCCGACGTCTAAGAAACCATTATTATCATGACATTAACCTATAAAAATAGGCGTATCACGAGGCCCTTTCGTCTTCACCTC
GATGGATCCCAAGCGATGGGGCATAAACTCTATCAACGTCTGCAATCGTGGTATCGCGTCTGTTTTGCATTACCGATCCGCAAAGGCTG
GGTGCGTGACTGATCTCTGCTAAAAAGTGTAGTATTGAGCGGCTCGCTTCAATAACTATTCAGAGGGATTATGGAAACACGTCGCGAAG
AGCGTATCGGGCAGCTGCTGCAAGAATTAAAACGCAGCGATAAGTTACATCTTAAAGACGCCGCCCTCGAGAGATCCTCTAGATTTAAG
AAGGAGATATACATATG
>yhhQ
GAGGCAATTCCGACGTCTAAGAAACCATTATTATCATGACATTAACCTATAAAAATAGGCGTATCACGAGGCCCTTTCGTCTTCACCTC
GAGGGTTTTGCGCACCATCATCACCGGTTCCGGGCAGCGCAGGCCAAGCGCGTCGAGTGTGTGGTCAGGGCTGGAAAAGAGATCGGTCA
TTTTCTTCTCATCACTTAAAAAAACGGCGCTAGTTTACGCCCTGTGAGTCCGTAAGCAAACTAGGTTAACGATTGCGTGAAAATTAACC
ATTGCATTGTCAACGTAAAGCAGTATCATGCGGCGGCTCGAAAAAAGGGTAAGCACGTTATTATGTTAAGGTAACAGACGTGTCGTACG
TATTGGGTTCCCTCACCCCAATGGTTAATCAAAAAGGTACAATATGAACGTTTTCTCGCAAACTCAACGCTATAAGGCGTTGTTCTGGT
TATCGTTATTTCATCTGCTGGTGATCACGGATCCTCTAGATTTAAGAAGGAGATATACATATG
>ychF

GAGGCAATTCCGACGTCTAAGAAACCATTATTATCATGACATTAACCTATAAAAATAGGCGTATCACGAGGCCCTTTCGTCTTCACCTC
GATGGATCCACCGATTGCACGCCTTTAAAGCGCAATAAGTCGTTGTCTGCGGCATTTTTGCCGAGTGCCGTGTATAATAGGCAAAGTTA
TTTCCATTTCTGCAATCTGTTAGCAATAACAGGTTGATTATTAAGATATTAAGGTGATTTAAATCATGGGATTCAAATGCGGTATCGTC
GGTTTGCCCAACGTCGGGAAATCTACCCTGTTCAACGCGCTGACCAAAGCCGGTATTGAAGCGGCCAACTTTCCTCGAGAGATCCTCTA
GATTTAAGAAGGAGATATACATATG
>ptsA
GAGGCAATTCCGACGTCTAAGAAACCATTATTATCATGACATTAACCTATAAAAATAGGCGTATCACGAGGCCCTTTCGTCTTCACCTC
GATGGATCCCTGCCGGTTTCACTGACCAATCTGAATCCGCAAATTATCCGCGCCCGCACCGTGTGCAGCGGTAGTGCAGGCGGCATTCT
GACGCCGATCTCTTCTTTAGATCTCAATGCGCTGGGTAATCTTCCCGCAGCCAAAGGCGTTGACGCCGAGCAATCCGCACTGGAAACTC
GAGAGATCCTCTAGATTTAAGAAGGAGATATACATATG
>ychM
GAGGCAATTCCGACGTCTAAGAAACCATTATTATCATGACATTAACCTATAAAAATAGGCGTATCACGAGGCCCTTTCGTCTTCACCTC
GATGGATCCGCCGAACGTGCGTACTCTGACCCTGTCAGGTATGCTGGCCGAAGCGATTCGTCGTATCAGCAACGAAGAATCGATCTCTG
CCATGTTCGAACACTAATCGAACCCGGCTCAAAGACCCGCTGCGGCGGGTTTTTTTGTCTGTAATATCCATTTGTATGACCTATGCCTC
CTTCACCTGCCATTTAGTTGACAGATGATGCGCTCACGGATGAAACATTATTGTGAACAAAATATTTTCCTCACATGTGATGCCTTTCC
GCGCTCTGATCGACGCTTGCTGGAAAGAAAAATATACTGCCGCACGGTTTACCCGTGACCTGATTGCCGGGATAACCGTCGGGATTCTC
GAGAGATCCTCTAGATTTAAGAAGGAGATATACATATG
>ychH
GAGGCAATTCCGACGTCTAAGAAACCATTATTATCATGACATTAACCTATAAAAATAGGCGTATCACGAGGCCCTTTCGTCTTCACCTC
GAGGTCAACGAACCAGGCACCAGCATTATGTCGCGTTGCGGCGTATTCAGCACCGGGGTTCGCCAGGCCGACAATCAATTTAATCGTCA
CGTTTTTTTGTCCTGAGTGTGTACATAACTGGCGCGTAGTTTACTGGTTGCGGCCCCGCTTGACAAAAAACTGCGTATCAAATGCAGAT
AACGTAATAATTGCCTGAGTGGACTATTAGAAAGTCAAGGTGTTCAGGCGTTTATTTGTAAAGTTTTGTTGAAATAAGGGTTGTAATTG
TGATCACGCCCGCACATAACCCACTGGGTGTTGTCTATACTTTACACATAAGGAAGAGGGGTATTCCCTGTTACAACCCAGAAAGTTCC
GGAGGTGACATATGAAACGCAAAAACGCTTCGGGATCCCGGGGATCCTCTAGATTTAAGAAGGAGATATACATATG
>ptsG
GAGGCAATTCCGACGTCTAAGAAACCATTATTATCATGACATTAACCTATAAAAATAGGCGTATCACGAGGCCCTTTCGTCTTCACCTC
GAGACTGGCGCAGGTAACCACCGATAACTTCGCCCGTCTGTTTCACATCGACGCTTCCCGCCTTCAATCCATCCGTTGAATGAGTTTTT
TTAAAGCTCGTAATTAATGGCTAAAACGAGTAAAGTTCACCGCCGAAAATTGGGCGGTGAATAACCACGTTTGAAATATTGTGACATAT
GTTTTGTCAAAATGTGCAACTTCTCCAATGATCTGAAGTTGAAACGTGATAGCCGTCAAACAAATTGGCACTGAATTATTTTACTCTGT
GTAATAAATAAAGGGCGCTTAGATGCCCTGTACACGGCGAGGCTCTCCCCCCTTGCCACGCGTGAGAACGTAAAAAAAGCACCCATACT
CAGGAGCACTCTCAATTATGTTTAAGAATGCATTTGCTAACCTGCAAAAGGTCGGTAAATCGCTGATGCTGCCGGTATCCGTAGGATCC
TCTAGATTTAAGAAGGAGATATACATATG
>ymdB
GAGGCAATTCCGACGTCTAAGAAACCATTATTATCATGACATTAACCTATAAAAATAGGCGTATCACGAGGCCCTTTCGTCTTCACCTC
GAGTGAGCGGCAATGTCAAAAACGCGCAAATAGCCTCAGTAAAAGTGCAGTATCTTGATAAACAGAAAAAGCTGGCTGTAATGAACATC
GAATATAACTAAGTTTCTGAACAACTCACGTTGCTGAGCAGAAAAATGCGATTTAACCAAAAAGCCTGCTGTACACTTAAGAAACAAGA
AGGTGTAAGGAGGCGTTATGAAAACGCGTATTCATGTTGTGCAGGGTGATATTACCAAACTGGCCGTTGATGTGATTGTGAATGCGGCT
AATCCGGGATCCTCTAGATTTAAGAAGGAGATATACATATG
>dppA
GAGGCAATTCCGACGTCTAAGAAACCATTATTATCATGACATTAACCTATAAAAATAGGCGTATCACGAGGCCCTTTCGTCTTCACCTC
GATGGATCCCCTGGTAGCGCACTTCGTTCGGGACGAAGGGGTCGGAGGTTCGAATCCTCTATCACCGACCAAATTCGAAAAGCCTGCTC
AACGAGCAGGCTTTTTTGCATCTGCAGCTCATGAGGATGAGAACCTCCGGGGGCAGGGAGGTTCGACTCGAGCGAAGCGAGAGAACGTT
GCGCCAGCAACGGCCCGCAGGGCGAGCCACGAAGTGGCGAGTAATCCTCTATCACCGACCAAATTCGAAAAGCCTGCTCAACGAGCAGG
CTTTTTTGCATCTGCAGCTCATGAGGATGAGAACCTCCGGGGGCAGGGAGGTTCGACTCGACGTAGGCCTGATAAGACGCGTCAGCGTC
GCATCAGGCAAATGTGCCGGCCATTGTCGGATGCGGCGCGACCGCCTTATTCGACCTACACGACACACATTTCCCTGCCAAATTTGCGC
CCCATATCTCACTATCCATTAACGGATTTGTGACAGAAACCATTGTCTTTTTTTATATTTGCTTAAATCTTTTTTTGCGTTACTTATGG
CTGGCCCTGGCATTTTCCGCTGTGCGTTTTAGCGTTCATGGTATTAATCGCTCAGATATCCACCTTTTCCTCAGATTTTTTTCATTAAA
TGCATAAAACTTAATCACCAAATGTTGCTAAATATTAAGCTGATTGTTCTGGTCATGTTGGAGTAGCACAAATTTCTCTGCTGCAAATA
GCTGTTTGAAGTTTTTTTAATCTTTGTGTGTGGATTCTCACCGTTGGTGTAAATCCCGGTTGGCTGTATTGACGTTTTCACATTCTGTT
GACAGATTGTAGGTCACGAGGGGCATTTTATGGAGGATCCGCACTGTTACACTGATGTTAATTAGTACGGCATCCCCACCTCATAACGT
TGACCCGACCGGGCAAAAAACAAAAAAGGTCAGGCAGCGACAACCCACTGCAAAGGGTTAAAACAACAAACATCACAATTGGAGCAGAA
TAATGCGTATTTCCTTGAAAAAGTCAGGGATGCTGAAGCTTGGTCTCAGCCTGGTGGCTATGACCGTCGCAGCAAGTGTTCAGGGTCGA
GAGATCCTCTAGATTTAAGAAGGAGATATACATATG
>dppB
GAGGCAATTCCGACGTCTAAGAAACCATTATTATCATGACATTAACCTATAAAAATAGGCGTATCACGAGGCCCTTTCGTCTTCACCTC
GATGGATCCCGCTCACTCCACCGTGTTTGAACCGGTACGTAAAGAAGTTAAAGGCTATGTGGTTGATCCATTAGGCAAACATCACTTCG
AAAACGTCTCTATCGAATAATTAAAAGCCATACAAGACTGATGGCAAAGGCAAAAATGCCTGATGCGCTCCGCTTATCAGGCCTACGAA
AATTCTGCAATGTATTGAATTTGCACGATTTTGTAGGCCGGATAAGGCGTTAACGCCGCATCCGGCATAAACAAAGCGCACTTTGTCAA
CAATCTGTATACCCGGTGGCGCTGTGCCTCTGCGTGCAGCGCCATCCGGCAGCAATACTTCATTCCCTGTCCGATAAGGGCGGGAATTG
ATTTGTGAGCAATACAGACACGCAGTTCCAGGCTGCGGGTCACTATAGAGAATCCGGGTTATGTTGCAGTTTATTCTCCGACGTTTGGG
ACTCGTCATCCCCACGTTTATCGGTATTACCCTTCTCACATTTGCCTTTGTCCACATGATCCCGCTCGAGAGATCCTCTAGATTTAAGA
AGGAGATATACATATG
>uhpT

GAGGCAATTCCGACGTCTAAGAAACCATTATTATCATGACATTAACCTATAAAAATAGGCGTATCACGAGGCCCTTTCGTCTTCACCTC
GATGGATCCCCTTTTTGAACGCCCAGACACCGCGCGAAGCGTGATGCATCTCACCTTTTCACTTCATATCCGGCAAAACTAAGAAATTT
TCCAGGTTTTGCCTGGACGCTATCTCAGGCCTGATTTGCTGCTGATTTTTACAATGCATGCCTCACGCAGGTATTCATTTCAGGAGTAA
CCCATGCTGGCTTTCTTAAACCAGGTTCGCAAGCCGACCCTGGACCTTCCGCTCGAAGTGCGGCGCAAAATGTGGTTCAAACCGTTCAT
GCAATCCCTCGAGAGATCCTCTAGATTTAAGAAGGAGATATACATATG
>fhuF
GAGGCAATTCCGACGTCTAAGAAACCATTATTATCATGACATTAACCTATAAAAATAGGCGTATCACGAGGCCCTTTCGTCTTCACCTC
GATGGATCCCGCGCTAGCTTCAGGTGTTAGTGTCCTTACGGACGTGGGGGTTCAAGTCCCCCCCCTCGCACCAATTATCTTTACTTCCT
TTCTTGTTTCTTCCTTGATTTTTCATCCCTGATTCAAATTCATTATCATCAGCATGATGCCAGCAATAAGTGCCAGGCACGATGGTAGC
AAAATGTAATGCCGCAACTGTTTGTGATACAGCATCATCGGCGTCAATACTAATCCCATCAGAATGACAATCCGCCACTGATCTGACGA
TAAATCCGCATACATCAGCCCTGCAATCAGCAATCCCGGCAGCAACACTCCCCAGCCACTGCCCAGCGTACGTTGCAACATGATTTCAT
CTCTTTCATTGATAATGATAACCAATATCATATGATAATTTTTATCATTTGCAAGCCAGATAAATCCCTTGCTATCGGGTAAACCTATC
GCTATGATTAGCAATCATTATCATTTAGATTACTATCCCGATTATGGCCTATCGTTCCGCACCGCTCTATGAAGATGTCATCTGGCGAA
CGCATCTCCAGCCGCAGGATCCCACGCTTGCCTCGAGAGATCCTCTAGATTTAAGAAGGAGATATACATATG
>fhuA
GAGGCAATTCCGACGTCTAAGAAACCATTATTATCATGACATTAACCTATAAAAATAGGCGTATCACGAGGCCCTTTCGTCTTCACCTC
GAGGCAACAGCAACCTGCTCAGCAAGAGCAGAAAGACAGCGACGGTGTAGCCGGTTGGATCAAGGATATGTTTGGTAGTAATTAACATC
TAAGCGTGAAATACCGGATGGCGAGTTGCCATCCGGTAAAATAACATCCCATCTAAGATATTAACCCTTTCTTTTCATCTGGTTGTTTA
TTAACCCTTCAGGAACGCTCAGATTGCGTACCGCTTGCGAACCCGCCAGCGTTTCGAATATTATCTTATCTTTATAATAATCATTCTCG
TTTACGTTATCATTCACTTTACATCAGAGATATACCAATGGCGCGTTCCAAAACTGCTCAGCCAAAACACTCACTGCGTAAAATCGCAG
TTGTAGTAGCCACAGCGGTTAGCGGCATGTCTGTTTATGCGGATCCTCTAGATTTAAGAAGGAGATATACATATG
>fhuC
GAGGCAATTCCGACGTCTAAGAAACCATTATTATCATGACATTAACCTATAAAAATAGGCGTATCACGAGGCCCTTTCGTCTTCACCTC
GAGATACGTCGCCAGCTGCTTTAACACTTATGGCTGCTTCTGGGGCGCAGAACGTCAGGTCGTTGCAACCGCAACCTTCCGTTTCTAAT
TTCTCTTTTGGGGCACGGATTTCCGTGCCCATTTCACAAGTTGGCTGTTATGCAGGAATACACGAATCATTCCGATACCACTTTTGCAC
TGCGTAATATCTCCTTTCGTGTGCCCGGGCGCACGCTTTTGCATCCGCTGTCGTTAGGATCCTCTAGATTTAAGAAGGAGATATACATA
TG
>uhpA
GAGGCAATTCCGACGTCTAAGAAACCATTATTATCATGACATTAACCTATAAAAATAGGCGTATCACGAGGCCCTTTCGTCTTCACCTC
GATGGATCCTCCGATCCGACGATGTTTAACAAGATCGCGGTGTTTTTTCAGTAACCGCTCAAGGCTTGAACAACATCGCGCTTATCGTT
AAGGTAAGCGCGTATTTTTTTTACCCGCCAGGACAAGACCATGATCACCGTTGCCCTTATAGACGATCACCTCATCGTCCGCTCCGGCT
TTGCGCAGCTGCTGGGGCTGGAACCTGATTTCTCGAGAGATCCTCTAGATTTAAGAAGGAGATATACATATG
>nuoM
GAGGCAATTCCGACGTCTAAGAAACCATTATTATCATGACATTAACCTATAAAAATAGGCGTATCACGAGGCCCTTTCGTCTTCACCTC
GATGGATCCGTGGTGCTGGCACTGTTGATGGTACTGCGTTGAGTTAAGGATTGTGGGATTGCCCCTGGGCATATAAGAACGTAGGTCGG
ATAAGCCGCCTTCGCGGTGCATCCGACACTACCCCCAGTCCGAAGAGAATTTTCTGTCGTGAGAATTCGTTGAAAATCCGGTCCTGACG
GGACTTTTACAAGGAATAAAGATCGCCATGTTACTACCCTGGCTAATATTAATTCCCTTTATTGGCGGCTTCCTGTGCTGGCAGACCGA
ACGCTTTGGCGTCAAGCTCGAGAGATCCTCTAGATTTAAGAAGGAGATATACATATG
>proX
GAGGCAATTCCGACGTCTAAGAAACCATTATTATCATGACATTAACCTATAAAAATAGGCGTATCACGAGGCCCTTTCGTCTTCACCTC
GAGGTCTGACGCAGGCCGTTGGGCGCGACTCACGCAGTCGCGGCAACCGTCGCTGGTACACCACTGGCCCTGTTGGTCTGCTGACCCGC
CCATTCATTAAGTAACTCTGCACTTGCCCGGTGACGCCGGGCATTATCACCCTGCCAAAAAAAGGAATAACAATGCGACATAGCGTACT
TTTTGCGACAGCGTTTGCCACGCTTATCTCTACACAAACTTTTGCTGCCGATCTGCCGGGCAAAGGCATTACTGGGATCCTCTAGATTT
AAGAAGGAGATATACATATG
>proS
GAGGCAATTCCGACGTCTAAGAAACCATTATTATCATGACATTAACCTATAAAAATAGGCGTATCACGAGGCCCTTTCGTCTTCACCTC
GATGGATCCTGCCGTCTGGCTACATGATTTTAACGTTCGCTGGCGCGTCACCGACGCAGGTTTTGAAGTCTTTGCGCTGGAACCGCGTT
AAATTCACGCCCTTCTCTTTTGACATTTCTTTTGCACTGGTAAACTAAATCACTTTTTTTTGTCCCAGGCTCGCCTTGAGCCTGTTCTA
CCTTCCAACTGGAACCGTAACAACATGCGTACTAGCCAATACCTGCTCTCCACTCTCAAGGAGACACCTGCCGACGCCGAGGTGATCAG
CCATCCTCGAGAGATCCTCTAGATTTAAGAAGGAGATATACATATG
>proQ
GAGGCAATTCCGACGTCTAAGAAACCATTATTATCATGACATTAACCTATAAAAATAGGCGTATCACGAGGCCCTTTCGTCTTCACCTC
GATGGATCCGCTTGTGGCACAGCTTGAAAAAGTGCTTGCAACGACGGATTACAAAAAATTCTTTGCGAGCGTCGCAGGATAATCAACGG
ATAACGTAGCATTTACTGATAGCGTCATTATAATGACGCCTGTTCATGCCTGCGCTTGTTGGCTACGTCCGTTGTAATCAGGAAATTTC
ATGGAAAATCAACCTAAGTTGAATAGCAGTAAAGAAGTAATCGCGTTTCTGGCCGAACGTTTTCCCCACTGTTTCAGTGCGGAAGGTGA
AGCGCGTCCGCTGAAAATCGGTATTTTTCAGGATTTGGTCGATCGTGTTGCTGGGGAAATGAACCTGAGCAAAACGCAATTGCGATCCG
CTTTACGTCTCTACACTTCGAGCTGGCGTTATCTTTACGGTGTTAAACCCGGCGCAACGCGTGTCGATCTTGACGGCAACCCATGCGGT
GAGCTGGACGAGCAACATGTAGAGCATGCTCGCAAGCAGCTTGAAGAAGCGAAAGCGCGTGTTCAGGCACAGCGTGCTGAACAGCAAGC
GAAAAAACGCGAAGCTGCCGCAACTGCTGGTGAGAAAGAAGACGCACCGCGCCGCGAACGCAAGCCACGTCCGACTACGCCACGCCGCA
AAGAAGGCGCTGAACGTAAACCTCGTGCGCAAAAGCCGGTAGAGAAAGCGCCAAAAACAGTAAAAGCACCTCGCGAAGAACAGCACACC
CCGGTTTCTGACATTTCAGCTCTGACTGTCGGACAAGCCCTGAAGGTGAAAGCGGGTCAAAACGCGATGGATGCCACCGTATTAGAAAT
CACCAAAGACGGCGTCCGCGTCCAGCTGAATTCGGGTATGTCTTTGATTGTGCGCGCTCGAGAGATCCTCTAGATTTAAGAAGGAGATA
TACATATG

Data Set S4 - MG1655 library reference file.fasta

```
>proL
GAGGCAATTCCGACGTCTAAGAAACCATTATTATCATGACATTAACCTATAAAAATAGGCGTATCACGAGGCCCTTTCGTCTTCACCTC
GAGATGACGACGCTGATGCAACGCCTGCTACATGTCAGCACACCTGCCAGCGAATATTCGCAAGGTCAGGATTTGTTCAACCCTCAACG
CCGTCATTACTGGGTTACCGCAGCGGATAACGATACGCTGGCAATTACCACCCCGAAAAAGACGCTGGTACTGAACAATAACGGTAAAT
ACCGCACTTACAACTTACGTGGTGAAAGAGTGAAAGATGAAAAACCACAGTTAAGTTTGTTATTGCAAGTGCTGACAGACGAGAAGCGG
GATCCTCTAGATTTAAGAAGGAGATATACATATG
>phnC
GAGGCAATTCCGACGTCTAAGAAACCATTATTATCATGACATTAACCTATAAAAATAGGCGTATCACGAGGCCCTTTCGTCTTCACCTC
GATGGATCCAATTTGGCGTACCGTGGATGATTAATGTCGTCAAACAACAACCAACGCAATAACCCGCCGGGAGGCCCGCCCTCCCGCAC
TGTCATCGAATTCCCGTTAACTCTTCATCTGTTAGTCACTTTTAATTAACCAAATCGTCACAATAATCCGCCACGATGGAGCCACTTTT
TTAGGGAGGCTGCATCATGCAAACGATTATCCGTGTCGAGAAGCTCGCCAAAACCTTCAATCAGCATCAGGCGCTGCATGCGGTTGATC
TGAACATTCATCACGGTGAAATGGTGGCTCGAGAGATCCTCTAGATTTAAGAAGGAGATATACATATG
>luxS
GAGGCAATTCCGACGTCTAAGAAACCATTATTATCATGACATTAACCTATAAAAATAGGCGTATCACGAGGCCCTTTCGTCTTCACCTC
GATGGATCCCGGTGTGGCTGGAAAAACACGCCTGACAGAAAAGAAAAAGGCCACTCGTGAGTGGCCAAAATTTCATCTCTGAATTCAGG
GATGATGATAACAAATGCGCGTCTTTCATATACTCAGACTCGCCTGGGAAGAAAGAGTTCAGAAAATTTTTAAAAAAATTACCGGAGGT
GGCTAAATGCCGTTGTTAGATAGCTTCACAGTCGATCATACCCGGATGGAAGCGCCTGCAGTTCGGGTGGCGAAAACAATGAACACCCC
GCATCTCGAGAGATCCTCTAGATTTAAGAAGGAGATATACATATG
>phnL
GAGGCAATTCCGACGTCTAAGAAACCATTATTATCATGACATTAACCTATAAAAATAGGCGTATCACGAGGCCCTTTCGTCTTCACCTC
GATGGATCCCAGCTGCTGGTGTCATCGGTTTTGCAGAATTGAGCCGGTGCCGGATGCGGCGTAAACGCCTTATCCGGCCTACAAATGCG
CTCCCCGTAGGTCGGATAAGACGCGTCAGCGTCGCATCCGACACCCGAACCACGAGGCGAAAAATGATTAACGTACAAAACGTCAGTAA
AACCTTCATCCTGCACCAGCAAAACGGCGTGCGCCTGCCCGTCCTCAATCGCGCCTCGCTCACCGTCAACGCTCGAGAGATCCTCTAGA
TTTAAGAAGGAGATATACATATG
>proB
GAGGCAATTCCGACGTCTAAGAAACCATTATTATCATGACATTAACCTATAAAAATAGGCGTATCACGAGGCCCTTTCGTCTTCACCTC
GAGCGGCAAAAGCGAGACGCGTTTTTTGCTGTGCAGTATCACTCTCTGCTTTATTACCGGCAAACTCTGCTTCCCAACGACCATAACCA
GTCAGTTGATCGTTAATTTGTGTTTCGCCTTTGAAACCAAAACGGATATAACTCTGGTCGCCATCTTTACTGGCGTTATCACTCATATA
ATGCATGGCTTTAACTTTGCCATAGACATCCAGTTTATTACCGTCTTTATTATATATTTCTGCAGCCTGTACAGATGCAGATGCCACAA
TGCCCATCACCACTAATGCCAGAGTGCTCTTTTTCATTTTCATTCCTGATTTTAATTAACGCGCGAATATTCAGCGGGAGAGTCCCGTT
GAAAACAGGAAAGTTTTTAACCTGAGATTGTTAAAGATATATTACAGATTAATAATATTCTTAAAATGTGGTAATTTATTAAATCTGTA
ATAAAAGCGTAAACAACTGCCGCGGATCCTCTAGATTTAAGAAGGAGATATACATATG
>lipA
GAGGCAATTCCGACGTCTAAGAAACCATTATTATCATGACATTAACCTATAAAAATAGGCGTATCACGAGGCCCTTTCGTCTTCACCTC
GATGGATCCATTCAGCCTGCGTGATCCCATCCTGCACGGAGTAATAGATGTTATCCGTAATGCATTTTGAAAAAGTATTGTTACAGCCA
GATCACAGCGCAGAACGATAAAATGGCACAATCCAACAACAATTTTACATTTTCGCGACCGCTTTGGCTGCTTTCAGGTCCGTTTCAAT
GATATACTGCCAGTCGTTAATTCAAAAATAGTTGATAATTACAACAATCTATTGAATTGAAACGCTTTCCTTCGTAATTCGCAACTGGA
ACACGCACGCTATGAGTAAACCCATTGTGATGGAACGCGGTGTTAAATACCGCGATGCCGATAAGATGGCCCTTATCCCGCTCGAGAGA
TCCTCTAGATTTAAGAAGGAGATATACATATG
>lipB
GAGGCAATTCCGACGTCTAAGAAACCATTATTATCATGACATTAACCTATAAAAATAGGCGTATCACGAGGCCCTTTCGTCTTCACCTC
GATGGATCCCGCCACTCATATCGAGCAGGTTGAAACACTGTATGAAGAACTGGGCAAAATCGATATTGTCCGCATGGTTCTGTAACTCG
CTTCTCCGTTACCCGGCTCCTGGTCGGGTAACTCCCCTCTCCTCGCTGTGATATACTTTTCCCCCACTTTTACTCATTCTCCACGGAGA
TGCCGTTTTGTATCAGGATAAAATTCTTGTCCGCCAGCTCGGTCTTCAGCCTTACGAGCCAATCTCCCAGGCTATGCATGAATTCACCG
ATACCCGCGATGATAGTACCCTTGATGAAATCTGGCTGGTCGAGCACTATCCGGTATTCACCCAAGGTCAGGCAGGAAACTCGAGAGAT
CCTCTAGATTTAAGAAGGAGATATACATATG
>bacA
GAGGCAATTCCGACGTCTAAGAAACCATTATTATCATGACATTAACCTATAAAAATAGGCGTATCACGAGGCCCTTTCGTCTTCACCTC
GATGGATCCGGCGAATGTTGGCGTAATCATTGAGCGTGGCAATAATCTGAAAGAAAATAATTAATTTTACAGCTGTTAAACCAAACGGT
TATAACCTGGTCATACGCAGTAGTTCGGACAAGCGGTACATTTTAATAATTTAGGGGTTTATTGATGAGCGATATGCACTCGCTGCTGA
TAGCGGCAATATTGGGTGTGGTCGAAGGATTGACAGAATTTCTGCCGGTATCCAGCCTCGAGAGATCCTCTAGATTTAAGAAGGAGATA
TACATATG
>somA_macA
GAGGCAATTCCGACGTCTAAGAAACCATTATTATCATGACATTAACCTATAAAAATAGGCGTATCACGAGGCCCTTTCGTCTTCACCTC
GAGTGGCGTAAAGGTCCGTTCAGTTAGCTGCGACATATTCATGAAATCAATGGTTATACATGGCGTCGATTTCACCATTGCGTATCTTA
ACCAAACATCAATAGTGTGATTACTAACGTAAATTTTAGGGTTTTGTTGATATTTCGTTGAAGTTAATGACCCGGATTGGCATATGGAG
TATTCAGAAAATTTATGAAAAAGCGGAAAACCGTGAAGAAGCGTTACGTTATTGCGCTGGTGATAGTCATCGCCGGACTGATTACGTGG
ATCCTCTAGATTTAAGAAGGAGATATACATATG
>bcsE
GAGGCAATTCCGACGTCTAAGAAACCATTATTATCATGACATTAACCTATAAAAATAGGCGTATCACGAGGCCCTTTCGTCTTCACCTC
GAGGCCTGCTTTAACGCCACAATATCATTCTGGAAGATATAGCCTATCGCGGGATCAGGCAGAGTATCTGGTTCATTGTTATTCATTTT
GCTTGATCCCTATGAAATAACCGTACCGTTCATGAGACAATTCTAACCGTGAGCATTTCGCCGGTTTATTGTTTACGGCGAGACGGGAA
TTTGCTCACTTTTTGACTTTCCGTTACGCAACAGAAAAAGTTAAAAACGAAAAGACTAAAATCACACGTCTTGTGACCTTCTAAATTGA
```

TTCACATGCTAAATCTGATAAGTTTTAATTTCAATGGTAGGTTTATTTCTTAGCTTTCGCTAGTAAACTGATAATCAGACAAAATGGTG
ACATGAGGGACATTGTGGACCCTGGGATCCTCTAGATTTAAGAAGGAGATATACATATG
>b3044
GAGGCAATTCCGACGTCTAAGAAACCATTATTATCATGACATTAACCTATAAAAATAGGCGTATCACGAGGCCCTTTCGTCTTCACCTC
GAGTGGAATCCCCAAGCAAAGATGCCACCCCGGGCAACGTTTACGCTCAGGCTGACTACAAGATTGCTTACGAGTAATCGCATTTGGCC
GGGTTCGCCCGGCCACATCATTAACGGATTAATGATAAGTGGATCAGATGTATAAAAAATTAAAATTAACCACAATAAGCGAATTGATT
AAAAATATTTATTGTTCATTATCCGTTATTAGACTGGCCCCCTGAATCTCCAGACAACCAATATCACTTAAATAAGTGATAGTCTTAAT
ACTAGTTTTTAGACTAGTCATTGGAGAACAGATGATTGATGTCTTAGGGCCGGAGAAACGCAGACGGCGTACCACACAGGAAAAGATCG
CAATTGTTCAGCAGAGCTTTGAACCGGGGATGGATCCTCTAGATTTAAGAAGGAGATATACATATG
>yqeG
GAGGCAATTCCGACGTCTAAGAAACCATTATTATCATGACATTAACCTATAAAAATAGGCGTATCACGAGGCCCTTTCGTCTTCACCTC
GAGCTACCAAGTTCCACGGCGGAATGACCCGCTAACGCACCACGAAAGCAGCCGATAGGTGTCCGTAACGCCCCGACAATCACAACGTC
TTTCATCACAACCTCGCACCAATTAACAGTGCAATAGTAAATGATTGTTAATTACAGTTATCTTAATTGTTTAAAAAAAGTGATTTTTA
TCACAAAGGAAATATGCCTGAGCAGCAGTCAGAGACATAACTGGCACGTAAGGTTTGCAACCACTAACCCACCAATAGAGGGGTAGATA
GGGCGTTAATCTCCCATACTTAACCTGGTTTATGGTAAATTGCCCTCCATTTTGTTTAATTTGTAGATGATACGTTCAGATAATGTCTA
ATATTTGGTCTAAAGAAGAAACTCTGTGGAGTTTCGCGCTCTACGGCACAGCCGTTGGTGCAGGCACGCTCTTCCTTCGGATCCGAAGG
AAGAGCGTGCCTGCGGGATCCTCTAGATTTAAGAAGGAGATATACATATG
>yehD
GAGGCAATTCCGACGTCTAAGAAACCATTATTATCATGACATTAACCTATAAAAATAGGCGTATCACGAGGCCCTTTCGTCTTCACCTC
GATGGATCCGTTTTCTTCGCCCGTCAAAGGTGTAACGACAGAAGTGGTTGTTGCAGGTAGCGATAGCAAACGTCGAATCGTGTTAAATC
GCTACGATTAATTTGGCAACCTGATTTTAATTGCATTTAGAAAGCCGAAATCATTTATATAAATGATTTCGGCTTTTTTATTGATATCA
ACAATACCATTTACATAAATCTACATTAATTGATAGATGCGATAAATAGTTAATACTCTTAAATATTTTTTGTATTTATATTGCAATGG
TTAATTTGTTTTAAATCAATAAAACAATGTGTTTACATTGCGGCAACTTTTTTGATGTGAGAAATTTGTTTCCAAGAGATACAGACTCT
TAACAACCAAAAATAAATTTTATTAAGGAACATATTATGAAACGTTCAATTATTGCTGCCGCTGTCTTTTCTTCTTTTTTTATGAGCGC
TGGAGTATTTGCTGCAGACGTTGATACCGGAACTCGAGAGATCCTCTAGATTTAAGAAGGAGATATACATATG
>yifE
GAGGCAATTCCGACGTCTAAGAAACCATTATTATCATGACATTAACCTATAAAAATAGGCGTATCACGAGGCCCTTTCGTCTTCACCTC
GAGATAGAGCGATTCAGCCGCTCGACCAAAGTGACGCGTTCGGCTAACTTCCAGGAAAGTTTTTAACAATTCCGTATCCACAGTGCTCT
CCGCAAAATTATTTGTCGTTATGATTTAAATGTTTTGTTTTACACTCTGTCAAGCGTAACTAATACTCCGCGCCATAACTAGCTCGGTC
AAAGAATTAGGAGCGTGCAGGATGGCGGAAAGCTTTACGACGACTAATCGATATTTCGACAATAAACATTATCCACGTGGATTCTCTCG
TCATGGTGATTTCACCATCAAAGAGGCACAAGGATCCCGGGATCCTCTAGATTTAAGAAGGAGATATACATATG
>yobF
GAGGCAATTCCGACGTCTAAGAAACCATTATTATCATGACATTAACCTATAAAAATAGGCGTATCACGAGGCCCTTTCGTCTTCACCTC
GATGGATCCGTGCGAAGCAAACGAACCAGAGAAAGCAGATAAAAAGACCGTTGAGAGTCAAAAATCGGTTGAAGACGAAGATATTATTC
GTCTGGTCGCCGAACGATAATCCTTCCTCAATCTTGAGTAACCCCGTCGTTGAAGGGTTACTCAATTCATCCTGATTATACGCACTGTC
ATGGTGCAACTGCTTCAAATTAAAACTCTGGTCTGACTTCAAATAATTATCATATTATTTATTTGCGTAACGGGATATCAGAAAAAGAT
ATTTACCCCATTAATTTATTAGGATGTTTACATCGGATTTGTGATTAAGCGTGGTATTATTTATTAGCGAAACGTTTCTCTCTTGATTT
TTTTGCTCATTCATCAATTTTTCTTATTTTAAATTTACAATCCCTTTGGGGATTGACTTCTCTTTAGGGTAATTAATAGCCGTTAACTG
ACTGTTTTATGAGAAAAAGTGATATAACTTTTTATTCATTGCATAGCAAAAAATGTGATATTGCACGCACTATGTAATAACTTCTCCCA
CTGGCCTGGAACAACTGAACTTATTGAACTATGTTAGAAAATACGCCAGTTTAAGTATCTGCCTGAACTGGCAAGGTTAAGCACAATGA
TATATCGGCGCGTATTCCGTTGCATAAGTGTGCAAAAAAAGTGGAAGACGTATCGAGATTTGTGCGTCTGATCGAGACATGTTTAAAAA
TGGCTTGCCATAATTAACGTTGTATGTGATAACAGATTTCGGGTTAAACGAGGTACAGTTCTGTTTATGTGTGGCATTTTCAGTAAAGA
AGTCCTGAGTAAACACGTTGACGTTGAATACCGCTTCTCTGCCGAGCTCGAGAGATCCTCTAGATTTAAGAAGGAGATATACATATG
>b2641
GAGGCAATTCCGACGTCTAAGAAACCATTATTATCATGACATTAACCTATAAAAATAGGCGTATCACGAGGCCCTTTCGTCTTCACCTC
GATGGATCCAGAGCGCTGTTCACCTGGAAAGTCTTGCTCGGTGCCGCTGTTGATGCGTTGTTTGCCAATGACGGCGCCGCGCTGATCCT
GACGCCGATTGTGATTGCGATGCTGATCGCACTGGGTTCAGCCAGGGCACGACACTGGCCTTTGTCATGGCTGCAGGATTTATTGCAGA
TACGGCCAGCCTGCCGCTCATTGTTTCTAACCTGGTGAATATCGTCTCGGCGGACTTCTTCGGTCTGAGCTTTGCGCAGTACGCCTCCG
TTATGATCTCCGTGGATGCGGCAGCGATTGCGGCCACGCTGATTATGCTGTATCTCTTCTTTCGCAGAGTTATTCCGGCAACGTATTAA
GTTTCGCTGCTGAAGACGCCTGCCAGTGCGATCAAGGATCTCGAGAGATCCTCTAGATTTAAGAAGGAGATATACATATG
>yqeI
GAGGCAATTCCGACGTCTAAGAAACCATTATTATCATGACATTAACCTATAAAAATAGGCGTATCACGAGGCCCTTTCGTCTTCACCTC
GAGACGCGTACAAGCAACGAATCGAAAAGAGAATGGGTGGTAAAATAAACGAATTATTTATTCGTTCACATTCGGTCCAACATTGATAC
TCAACGACCAGCCAGAATCATACTCTGTTTATACGGGCATTTATTGAAATGTCCGTATTTTATAAATAGGCCGTCCCCTCAAGAGTAAA
CACCATAGTATTTAAAACCATCACACGAAAAATTCAGAAGCATTACGAAATATGAATTTACAAACAATAGTGGCATAAATGTTAACCAT
GTTAATTTACGTAAAGTTTTACGTTGCAACATTAAAGCCTCATTTCAATCATCATGATAAATATAAAATTAATATATATTTATGCCGTA
AATACCAATATACTTAGCAAACTATGTGATCTCCATTTCGATTGATTTAGTGTTTATTGACGTATGTACTGGATTATTAACGATAATAT
CGAGTTCTGGCCTGAGCACCGAAAATTAATATCGGTACATAACGCCGATCTTAACGTCGGGATCCTCTAGATTTAAGAAGGAGATATAC
ATATG
>yqeH
GAGGCAATTCCGACGTCTAAGAAACCATTATTATCATGACATTAACCTATAAAAATAGGCGTATCACGAGGCCCTTTCGTCTTCACCTC
GAGTTAAACCCTGGCGTTCCATCGGAAATCTGATTACGCTGATCGTGGGTATCCTGTGCGTATCGGTAATGTTCTTTAGCTAATATCAG
TTGTGGATGGTTCAAAATTGCTACAATTTTGAACCATTCCATTTTTCATAATATTCATTTGTTATTATTCCACCCTATCACTTCTCTCG

TCATATCCTAGTATTATCAGTTAGTTAAACACACAAAGAGATCTGAACATCTTAAATAAGACTATTTAAGATGCATAACTTAGATTCGC
AAGATATCTTCTGGTCATTCAAAAACAATTTGCGATATAGATATTCTTATACCATGAGATAGTTATCTGGACCGCTCTCAACCATGCAA
GATTAAATCAGTGAAACATAATCATATTTGATAACGAGAAATGCATTTTTAAATGCCTTACTCAGAATTATAGCAAATACAGATTAATC
GCAATTTTTCCCGGCAAAATTACAAAATCATGGTGTATTGACATCATCAATAACCAATGAAATTATGCAATTATATACGGATAGGGAGG
TTCTTAACATGGGGCTTTGTAGTCGGGATCCCGGGATCCTCTAGATTTAAGAAGGAGATATACATATG
>yqeK
GAGGCAATTCCGACGTCTAAGAAACCATTATTATCATGACATTAACCTATAAAAATAGGCGTATCACGAGGCCCTTTCGTCTTCACCTC
GATGGATCCTGGTTGCAGTGGTAGGCTATTATTAATATTTCGTGGCTTCATTTTACATTTCTCTCATTTGATTAGAGCAAAATCCGAAT
TCGCAGCTTCTGATAATGTCAAATATCACCTTATGGAGACGTTACGTAGAATGACTATATTTGTTATCGGCACATTAGGGGTATTAATT
TTAGTATTGATGGAAGGTGATTATTGAACAGTAACATCGAGTCACGGTAAAAATGTGAGGTATACAATGGACATTGAATTTTCGCAGAT
TCATGAAATGGTCTATATGCATGATATTGTAAATTCGGACTCGAAAAAGAAACCACTCGAGAGATCCTCTAGATTTAAGAAGGAGATAT
ACATATG
>yehS
GAGGCAATTCCGACGTCTAAGAAACCATTATTATCATGACATTAACCTATAAAAATAGGCGTATCACGAGGCCCTTTCGTCTTCACCTC
GATGGATCCGCCGAGTTGATTTTGCGTAATGGCTTAACCGTGCCGGTCAGCCGCCGTTATCTGAAAAGCTTAAAAGAGGCGATTGGCCT
GTAAAAGACTGCTAAAATGGCTTTTTGCCTCATCAACACCTGAAGGCCTCATGCTAAGTAACGATATTCTGCGCAGCGTGCGCTACATT
TTGAAAGCCAATAATAATGACCTGGTGCGTATTCTGGCGCTGGGTAATGTCTCGAGAGATCCTCTAGATTTAAGAAGGAGATATACATA
TG
>yhfG
GAGGCAATTCCGACGTCTAAGAAACCATTATTATCATGACATTAACCTATAAAAATAGGCGTATCACGAGGCCCTTTCGTCTTCACCTC
GATGGATCCAATGTGCCGTCAAAACCGGTAGTTATCCTTTCCGCTAAAGTCCTGCCGTAATGATTTCTCGCGCGGGCAATCTTGCCCGC
GCTTCTGCTCTCCCGGCGTAACCCGGATTTGCCGCTTATACTTGTGGCAAATGGACACGTTCAGGGAGGCATCAAGTGAAGAAACTCAC
CGATAAGCAAAAGTCCCGTCTCTGGGAGCTTCAGCGTAATCGTAATTTTCAGGCCAGTCGCCGTCTTGAAGGCGTCGAGATGCCTCGAG
AGATCCTCTAGATTTAAGAAGGAGATATACATATG
>yehZ
GAGGCAATTCCGACGTCTAAGAAACCATTATTATCATGACATTAACCTATAAAAATAGGCGTATCACGAGGCCCTTTCGTCTTCACCTC
GATGGATCCAATATGACGCCGAGCCTGGCAAGTTCAATGTCTTTATCGGCACTGATTCCGCACGCGTTAAGAAAGGCGAGTTTGAGTTG
CTGTAATGCTTTAGTAAGGGCGCGACGTTTGCGCCCTTTGTAGGCCGGATAAGGCGCTCGCGCCGCATCCGGCAGGAAACGCCGGACCG
AATATGCCTCTTAAGTATTTATCTGATTCTGCAACTGAATCCTTCCGCTCAAGCTAACCCCGCCATTATCAACTATGCTTTTCTCTTAA
TTCGCTGAAAACAACAGTAAAAGAATGAGGAAAGCATCATGCCACTCTTAAAGCTCTGGGCAGGTTCACTGGTTATGTTGGCAGCCGTG
AGCCTGCCGCTACAGGCGGCTTCCCCCGTTAAAGTCGGTCTCGAGAGATCCTCTAGATTTAAGAAGGAGATATACATATG
>bisC
GAGGCAATTCCGACGTCTAAGAAACCATTATTATCATGACATTAACCTATAAAAATAGGCGTATCACGAGGCCCTTTCGTCTTCACCTC
GATGGATCCTCGCCGGTGTAAGGGTTAGTTGTGCAGCCAGATACCGCCAGAGCGCCACTCACTACGGCGGCAATAAGATAAACACGTTT
CTTCATTGTTAATCCTTAATAACCTTTTTATTCTTTGCCACGGGTTCCGTGGCGGGAGATTATGCCGCGTGAACATGAAGATTATTCCT
GGGAATACTCGGAAATTTGTAAGTAATATTTAACTGCTCAATACATCTAACCTTTCAGGAGTCTTCGGTTTGGCCAACTCATCCTCACG
ATATTCCGTTCTGACTGCCGCCCATTGGGGGCCCATGCTGGTTGAAACCGACGGCGAAACCGTGTTTAGCTCGCGTGGCGCGTTAGCCA
CAGGAATGGAAAACTCCTTGCAGAGCGCGGTTCGCGACCAGGTTCACAGCAATACGCGGGTACGATTTCCAATGGTGCGAAAAGGCTTT
CCTCGAGAGATCCTCTAGATTTAAGAAGGAGATATACATATG
>pbpG
GAGGCAATTCCGACGTCTAAGAAACCATTATTATCATGACATTAACCTATAAAAATAGGCGTATCACGAGGCCCTTTCGTCTTCACCTC
GATGGATCCGAAGTGCTGCTCGCCCTAACCGTTATACTATGGGGCTACGGTTATCGTTTGTTCTAGAAAGTTCTTTGACGTTGCATTGC
TGGCGTAAATGTCAGCAATGCAGCATTTCCTCACGATTCTCCTTTGACGATCGCACTTTTTGCTCGTTATGATGGCGTCGCTAGCCTCA
GTAAATCCTTACTCCGGCGGTGCGCAACCCGTGCGCGTGAACCACTATCTGAATGCTCATCATGCCGAAATTTCGAGTTTCTTTATTTA
GCCTGGCCCTGATGCTGGCTGTGCCTTTTGCACCGCAGGCCGTTGCGAAAACGGCAGCCGCTACCTCGAGAGATCCTCTAGATTTAAGA
AGGAGATATACATATG
>idi
GAGGCAATTCCGACGTCTAAGAAACCATTATTATCATGACATTAACCTATAAAAATAGGCGTATCACGAGGCCCTTTCGTCTTCACCTC
GAGACCTGGTGAAGGAATCCGTCTCTGATAAAGATTTAAAAGTCAGGACAGTACGTATGTGGCTTCTGATGCGCAAGCTGAAGAAAAAT
GAGCATGGAGAATAATATGAATTTTTTAATGCGCGCTATATTCAGTCTGTTGTTGCTTTTTACTCTCTCTATTCCTGTCATTTCTGACT
GTGTTGCAATGGCCATTGAAAGTCGCTTCAAATATATGATGCTACTTTTTTAAATGGTTTTTACCTGTCGGCATCCGCTCAAAACGGGC
GGTTGTCGATAAACGCTCACTTGGTTAATCATTTCACTCTTCAATTATCTATAATGATGAGTGATCAGAATTACATGTGAGAAATTATG
CAAACGGAACACGTCATTTTATTGAATGCACAGGGAGTTCCCACGGGTACGCTGGAAAAGTATGCCGCACACACGGCAGACACCCGCTT
ACATCTCGCGGATCCTCTAGATTTAAGAAGGAGATATACATATG
>bioB
GAGGCAATTCCGACGTCTAAGAAACCATTATTATCATGACATTAACCTATAAAAATAGGCGTATCACGAGGCCCTTTCGTCTTCACCTC
GATGGATCCACCTGCTGCGCTTCAAACAGCAGATCCAGCAACGGTTTTTCAAATAATTCTGTGACTTGCGACAATGTCCAGCGTGGGCG
GTGAGCCATGGGGCTTCTCCAAAACGTGTTTTTTGTTGTTAATTCGGTGTAGACTTGTAAACCTAAATCTTTTCAATTTGGTTTACAAG
TCGATTATGACAACGGACGATCTTGCCTTTGACCAACGCCATATCTGGCACCCATACACATCCATGACCTCCCCTCTGCCGGTTTATCT
CGAGAGATCCTCTAGATTTAAGAAGGAGATATACATATG
>b0332
GAGGCAATTCCGACGTCTAAGAAACCATTATTATCATGACATTAACCTATAAAAATAGGCGTATCACGAGGCCCTTTCGTCTTCACCTC
GATGGATCCTGCAGGATCGTTGTGTCGCTCATGTTGTCCTCGTCATTGTTATCATTTGTAGGGTCGAGATTGTTGTCGGATGCGACGCT

Data Set S4 - MG1655 library reference file.fasta

ATGCGTCTTATCCGACCTACAGTTCAGTTCACCCCGTAGGCCTGATAAGTCGCGTCAGCGGCGCATCAGGCATCGGTTGTCGGATGCGA
CGCTATGCGTCTTATCCGACCTACAGTTCAGTGCACCCCGTAGGCCTGATAAGTCGCGTCAGCGGCTCGAGAGATCCTCTAGATTTAAG
AAGGAGATATACATATG
>pabC
GAGGCAATTCCGACGTCTAAGAAACCATTATTATCATGACATTAACCTATAAAAATAGGCGTATCACGAGGCCCTTTCGTCTTCACCTC
GAGGAAGCGCGTCAGGTTAGCGGAATGGAATACACTCTGTGTAACTCCTTCGGCTTCGGTGGCACTAATGGTTCTTTGATCTTTAAAAA
GATCTAAGTTGTCATTTTCCACCCTTATAAAAGGTCCGCTTGCGGGCCTTTTTTCTTAGCTTTTATTCCGACTTGTTCCGTAGTGAACA
TGCTGCCACACTAACAATTCTCTGATAAGGAGCCGGTATGTTCTTAATTAACGGTCATAAGCAGGAATCGCTGGCAGTAAGCGATCGGG
CAACGCAGTTTGGTGATGGTTGTTTTACCACCGCCAGAGTTATCGGGATCCTCTAGATTTAAGAAGGAGATATACATATG
>yeeO
GAGGCAATTCCGACGTCTAAGAAACCATTATTATCATGACATTAACCTATAAAAATAGGCGTATCACGAGGCCCTTTCGTCTTCACCTC
GATGGATCCAGCGGGCTTTTCAGGAATTTGGCTCCTCTGACTGGACTCGAACCAGTGACATACGGATTAACAGTCCGCCGTTCTACCGA
CTGAACTACAGAGGAATCGTTGTTGAGGCACATCTTAACGGCGAAAAATCTTTTGTCAAACCCGATTTTTAAATTCCCCAACTGTTTGC
CGTTTCTATCAACAGTTTGTTGCATTTGCAGACAATTTGTTGGCCTCGAGAGATCCTCTAGATTTAAGAAGGAGATATACATATG
>intF
GAGGCAATTCCGACGTCTAAGAAACCATTATTATCATGACATTAACCTATAAAAATAGGCGTATCACGAGGCCCTTTCGTCTTCACCTC
GATGGATCCCGCGCCCTGATTGACTACGTCAAAGAATGGCGGCAGCAATTGGCTTAAATGTCTGCACCTGCACTGCCTGATGTCAGAAC
AGTATTTTGATGAATTGCCAAGGTTACAATGGCACAAATACGGCACAGGAGGAAACGTGGTGTATTTAAATATGGGGTAACTCTTTGAT
TTTAATGGTGCCGATAATAGGAGTCGAACCTACGACCTTCGCATTACGAATGCGATGTTAATGCATTTTTAACTACTTGAGCAATCAAT
CAAAACAACATACTTTATATTATAAAACACCAAGTTATAGCAATAAACCCGTTGATTGAAATTAACAAAAGTTGATGTTTATTCCCTCC
ATTTACTTACACCAGCAGTTACATTATTGTAAGACAGCAATTCTCAACTGGAGCCGAAAAATGGCGCTTTCACTCGAGAGATCCTCTAG
ATTTAAGAAGGAGATATACATATG
>intA
GAGGCAATTCCGACGTCTAAGAAACCATTATTATCATGACATTAACCTATAAAAATAGGCGTATCACGAGGCCCTTTCGTCTTCACCTC
GAGGCTGGCAAGCGAATGTAAAGACTGACTAAGCATGTAGTACCGAGGATGTAGGAATTTCGGACGCGGGTTCAACTCCCGCCAGCTCC
ACCAAAATTCTCCATCGGTGATTACCAGAGTCATCCGATGAAGTCCTAAGAGCCCGCACGGCGCAAGCCCTGCGGGCTTTTTTGTGCCC
TCAATTTGTCCCGCGAAGTCCGAAGAGAACTAATTAAATCCGAACCTTTTAGGCCCATTGATAGGCCCAACGAAAAGCTCTATTGTTTA
CGTTGGGCCTAAACGCAGGGAGACTCCCCATGGCAAGAAAAACCAAGCCGTTAACTGATACGGAAATCAAAGCCGCCAAACCTAAAGAT
GCCGATTACCAGCTTTATGACGGTGACGGGCTGGATCCTCTAGATTTAAGAAGGAGATATACATATG
>ybgA
GAGGCAATTCCGACGTCTAAGAAACCATTATTATCATGACATTAACCTATAAAAATAGGCGTATCACGAGGCCCTTTCGTCTTCACCTC
GAGGGCAGAGAAGAAACCCGTCTTCATATTGTTTGCGATGTCCCTGATGAGCTTATTGATTTCACGTTTGAATAGAAAGGGCAGCTATG
GTTAGAAACTACCTGACGTCAGTCCTTGCGGGGAGCAGGCTTTCGTAAATTTGTCCTGCTACAACAGGATTAACTTCACAAATATCATT
TCTCAACGTCTACACTTACTCCTGTAAACCGCTCAGGAGCAGTAATGAATCTACAACGATTTGATGACAGCACCCTAATCCGTATCTTT
GCCCTTCATGAGTTACATCGACTGAAAGAACATGGCTTAACGCGCGGATCCTCTAGATTTAAGAAGGAGATATACATATG
>yfdQ
GAGGCAATTCCGACGTCTAAGAAACCATTATTATCATGACATTAACCTATAAAAATAGGCGTATCACGAGGCCCTTTCGTCTTCACCTC
GAGCTGCAGGAAGACCAGGGACAATTACTGACTCTTGCCGGTGAGCCTGACGGAAAACTCCGCGCAGCAGGTCATTAATATCATTCTTA
ATTAACTAATTATTTATCTCATCACTGAATATCTTAATATAGTGAGGACTTATTATGTCTCAGAACTTAGACGCAACCGCAATTAATCA
AATCCATGCCCTTATTTCTGCTCAGGGTGTTAATGAAATTATCAGTAAGATTGGTGCCGATGCTGTGGATCCCGGGATCCTCTAGATTT
AAGAAGGAGATATACATATG
>intS
GAGGCAATTCCGACGTCTAAGAAACCATTATTATCATGACATTAACCTATAAAAATAGGCGTATCACGAGGCCCTTTCGTCTTCACCTC
GAGGCGGGTATACTCATGCCGCATTGTCCTCTTAGTTAAATGGATATAACGAGCCCCTCCTAAGGGCTAATTGCAGGTTCGATTCCTGC
AGGGGACACCATTTATCAGTTCGCTCCCATCCGTACCAGTCCGCAAAATCCCCTGAATATCAAGCATTCCGTAGATTTACAGTTCGTCA
TGGTTCGCTTCAGATCGTTGACAGCCGCACTCCATGACGGGTAAAAAGTGGATAAAATAATTTTACCCACCGGATTTTTACCCATGCTC
ACCGTTAAGCAGATTGAAGCAGCAAAGCCGAAAGAAAAACCATACCGCCTTCTCGATGGTAAGGATCCTCTAGATTTAAGAAGGAGATA
TACATATG
>intZ
GAGGCAATTCCGACGTCTAAGAAACCATTATTATCATGACATTAACCTATAAAAATAGGCGTATCACGAGGCCCTTTCGTCTTCACCTC
GAGCATTGCCGAACAATGTGGTCTTTAGTTTCATAAGTCGTTCCCTCAGGAAGAATCGATGATTGGCATTTTTCACAGCATTCTAACAA
TCACGTTTCATCGTCAGCCCTTGTCGTGTAAGGTGGTTGCCTAAACACGCCCGTTATTCATCACGCCGAACGCGCCGGATACATGATCG
GGGTTATCCAGTCGTTAAATCAAGGTATCCGGTTTTGAGCAAGACAGGATCCTCTAGATTTAAGAAGGAGATATACATATG
>infA
GAGGCAATTCCGACGTCTAAGAAACCATTATTATCATGACATTAACCTATAAAAATAGGCGTATCACGAGGCCCTTTCGTCTTCACCTC
GATGGATCCCGGACGATTGCCGAATAATTTCTGGGTACCACGATGCTTGTTTTCACCACAAGAATGAATGTTTTCGGCACATTTCTCCC
CAGAGTGTTATAATTGCGGTCGCAGAGTTGGTTACGCTCATTACCCCGCTGCCGATAAGGAATTTTTCGCGTCAGGTAACGCCCATCGT
TTATCTCACCGCTCCCTTATACGTTGCGCTTTTGGTGCGGCTTAGCCGTGTGTTTTCGGAGTAATGTGCCGAACCTGTTTGTTGCGATT
TAGCGCGCAAATCTTTACTTATTTACAGAACTTCGGCATTATCTTGCCGGTTCAAATTACGGTAGTGATACCCCAGAGGATTAGATGGC
CAAAGAAGACAATATTGAAATGCAAGGTACCGTTCTTGAAACGTTGCCTAATACCATGTTCCGCGTAGAGTTAGAAAACGGTCACGTGG
TTACTGCCTCGAGAGATCCTCTAGATTTAAGAAGGAGATATACATATG
>rlmB

GAGGCAATTCCGACGTCTAAGAAACCATTATTATCATGACATTAACCTATAAAAATAGGCGTATCACGAGGCCCTTTCGTCTTCACCTC
GAGAGACGCGAGAAAAGCGAAAAAGCCATCGGCGAAAACGCAGAAAATAGCTGCAGCGACCAAAGCGAAGCGTGCGGCGAAGAAAAAAG
TGGCAGAGTGATCAATACCCTCTTTAAAAGAAGAGGGTTAGATTGCTGACAAAATGCGCTTTGTTCATGCCGGATGCGGCGTGAACGCC
TTATCCGGCCTACATAATCACGCAAATTCAATATATTGCAGAGATCATGTAGGCCGGATAAACGTAGTGCATCAGGCAAAACGTAAACA
ACGAGTACATTAATGAGCGAAATGATTTACGGCATCCACGCAGTGCAGGCCCTGCTGGAGCGCGCCCCTGAACGTTTTCAGGGATCCTC
TAGATTTAAGAAGGAGATATACATATG
>ihfB
GAGGCAATTCCGACGTCTAAGAAACCATTATTATCATGACATTAACCTATAAAAATAGGCGTATCACGAGGCCCTTTCGTCTTCACCTC
GAGTGACGAGAAAGATGCAATCGCAACTGTTAACAAACAGGAAGATGCAAACTTCTCCAACAACGCAATGGCTGAAGCTTTCAAAGCAG
CTAAAGGCGAGTAATTCTCTGACTCTTCGGGATTTTTATTCCGAAGTTTGTTGAGTTTACTTGACAGATTGCAGGTTTCGTCCTGTAAT
CAAGCACTAAGGGCGGCTACGGCCGCCCTTAATCAATGCAGCAACAGCAGCCGCTTAATTTGCCTTTAAGGAACCGGAGGAATCATGAC
CAAGTCAGAATTGATAGAAAGACTTGCCACCCAGCAATGGATCCTCTAGATTTAAGAAGGAGATATACATATG
>atoS
GAGGCAATTCCGACGTCTAAGAAACCATTATTATCATGACATTAACCTATAAAAATAGGCGTATCACGAGGCCCTTTCGTCTTCACCTC
GAGCGAATTTCCGATTCTCGCTGATGTAACGCATTAACGATGTAAAAAACGGATGAAAAAGCAATCAACAGCCAGAGCACTAACGCCAA
TGCTCTGAACATGTAGCGCGAGGCTTTCAGGGTTGTACGAAAAGAAGCAAGGTATTTCAAAGGGGCGAAGCTCCGCCTCAGGTGACCGA
TGGAGTGTGGTTAAGGTAGCGGTAAAAGCGTGTTACCGCAATGTTCTCTCTTCTCTGGAATATGATACACCGCCGAGAAATCATCACCT
TAACCTCTGATAATCGTCATATACCGGACAAGACTAGTGGATTTCAGCATGCATTATATGAAGTGGATTTATCCACGCCGCTTACGCAA
TCAAATGATCCTGATGGCAATCCTGATGGGGATCCTCTAGATTTAAGAAGGAGATATACATATG
>yhcG
GAGGCAATTCCGACGTCTAAGAAACCATTATTATCATGACATTAACCTATAAAAATAGGCGTATCACGAGGCCCTTTCGTCTTCACCTC
GAGTCGCGAATGATTTAGGCGGTTCAGAAAAAAATCGAACTTGATGGGAAAGCTGTGATAAGCGTAGTCTACTTGTGATAACAATGCCGG
ACAGGAGTACTATCAGGCATTCGTTTTATACGATCGGTTAATGTTTCAGGAAAGCTCTTCGCATATCTTGGGGAAAGTAAATGTAATAC
GCTGTTCTTAATTGTTCAGTGCTCACCCCCCAGAGTATAGCTCATTCCATTATTCAACTTAGCTCCATATAAAGGGACAGATAATGGAA
TCGCTCTCAGAAGGAACCACAGCAGGCTACCAGCAAATCCACGACGGTATTATTCATCTGGTCGATAGCGCCCGGACGGAAACGGTACG
TAGCGGGATCCTCTAGATTTAAGAAGGAGATATACATATG
>exuR
GAGGCAATTCCGACGTCTAAGAAACCATTATTATCATGACATTAACCTATAAAAATAGGCGTATCACGAGGCCCTTTCGTCTTCACCTC
GAGCGACCTGTTAGGTGCGCTGGTTATCTGGACCGTGTTGCAGAACAAACCGGCGATTGAGGTGGCGCAGGAAACACATAACGATCCCG
CACCGCAACATTAACGTTTCGTGTAAGTTTTAGTTTCCCTTCAAAGCCGCCTTCTCAGGCGGCTTTTTCATCACTGCGAGTAGAGCTAA
ACTCGCAAAAGTGGTATAACAAATATAGTCTGCCGTATCATGCCTGGAGCGCATATGGAAATCACTGAACCACGCCGTTTGTATCAACA
ACTTGCCGCTGACCTGAAAGAGCGCATCGAACAGGGGATCCTCTAGATTTAAGAAGGAGATATACATATG
>tufB
GAGGCAATTCCGACGTCTAAGAAACCATTATTATCATGACATTAACCTATAAAAATAGGCGTATCACGAGGCCCTTTCGTCTTCACCTC
GAGCGCTCCAAGATGTGCTGATATAGCTCAGTTGGTAGAGCGCACCCTTGGTAAGGGTGAGGTCGGCAGTTCGAATCTGCCTATCAGCA
CCACTTCTTTTCTCCTCCCTGTTTTTTCCTTCTGTTTATTGCATTCAACAAGTCGGGCATGTTGCCTGGTTGATGTGGTGATATCACCG
ATTTATCCGTGTCTTAGAGGGACAATCGATGTCTAAAGAAAAGTTTGAACGTACAAAACCGCACGTTAACGTCGGTACTATCGGCCACG
TTGACCACGGTAAAACTACTCTGACCGCTGCAATCACTCGAGAGATCCTCTAGATTTAAGAAGGAGATATACATATG
>tufA
GAGGCAATTCCGACGTCTAAGAAACCATTATTATCATGACATTAACCTATAAAAATAGGCGTATCACGAGGCCCTTTCGTCTTCACCTC
GATGGATCCATGAAGCGCCGAGTAACGTTGCTCAGGCCGTAATTGAAGCCCGTGGTAAATAAGCCTAAGGGTTAATACCAAAGTCCCGT
GCTCTCTCCTGAAGGGGAGAGCACTATAGTAAGGAATATAGCCGTGTCTAAAGAAAAATTTGAACGTACAAAACCGCACGTTAACGTTG
GTACTATCGGCCACGTTGACCACGGTAAAACTACTCTGACCGCTGCAATCACTCGAGAGATCCTCTAGATTTAAGAAGGAGATATACAT
ATG
>ycgY
GAGGCAATTCCGACGTCTAAGAAACCATTATTATCATGACATTAACCTATAAAAATAGGCGTATCACGAGGCCCTTTCGTCTTCACGTC
GACCGGCTCCATCAGTGGTTTTAATCTGCACAGCTTCCTGGTGGCGGTGGTGGGAGCTATTCTCGTTCTGGGCATATTCCGCCTCCTGC
GAAGAGAATAAGATTTTCATAAGGCGGATAGCGATACAGATGCCGCTATCCGCTTTCACATCAGAACGTATACTCGACACCTGCTTTAC
GGGTGAAAAAAATCAATATAGCACTTCGATATCCGATAAAGTCGGTTCCCGGTAACAACAACAGAATCCTTTTTTAATTATTGTTTCGT
TGTTTTTGGTGTGATTCAGAAAATAAATATTTTAGACAGTAGCCAGTCAGGCGTTAAAAATGGGGCGGATAGTGAGGCGGTTTTGGCGG
ATCCTCTAGATTTAAGAAGGAGATATACATATG
>chaC
GAGGCAATTCCGACGTCTAAGAAACCATTATTATCATGACATTAACCTATAAAAATAGGCGTATCACGAGGCCCTTTCGTCTTCACCTC
GAGCGAAGAAACCGCGCATAAAGTGGCCTGGGCTGCTGTGAAGCATGAATATGCCAAAGGGGATGATGATAAATGGCATAAAAAATCGT
AAAACCGGTCGCTTAGTTAAAGCTATTCGTGCGGTGTTGCCTTGCAAGTGGTCCGTGGATTGCATATTGTCCCGTTAGTGGTTTCAAAA
TGAGCAGTAAAAATGTCCGGAAGACACCAAAAAGTTGTCGCAGGGAAGTATGCAGTGGCGGAAGTGTAAGGTGATAACGCGTGATTTCT
TGATGAATGCCGATTGTAAAACGGCATTTGGTGCCATTGAAGGATCCTCTAGATTTAAGAAGGAGATATACATATG
>yagM
GAGGCAATTCCGACGTCTAAGAAACCATTATTATCATGACATTAACCTATAAAAATAGGCGTATCACGAGGCCCTTTCGTCTTCACCTC
GATGGATCCAACGTTTCCTTCTGCCTTCATTACTACCCCTACTAATTCAATAATGCTTAAAGCTTTCTTTGCCAATATCTCAATCACTG
AGGTGGCAAAATGAGCGCATTCAAACTCCCGGATACATCTCAATCACAGCTCATTTCAACAGCTGAGTTAGCTAAAATCATTAGCTACA
AATCTCAAACCATTCGTAAATGGCTTTGTCAGGACAAATTGCCTGAGGGGCTACCTCGCCCAAAACAAATCAATGGCCGCCATTACTGG
TTACGTAAAGATGTCCTCGATTTTATAGATACATTTTCTGTACGAGAAAGTCTGTAATAAATTACAGATTTAATTTTATTGATTTATAG

Data Set S4 - MG1655 library reference file.fasta
CGATGTTGCCCCGAGAAAAATGGGGCAACACTGAGAAATTTCAGATAGTAGTTTTATATTGAGATAACAAAGAGGTTTCCTTAAAAATG
TCTAATAGTGTTACTAATTTTGAGATGAGCAGCGTTCTACCAGGAAAAAAACCTTGTCAAGGCAAAAACAATGACTCGAGAGATCCTCT
AGATTTAAGAAGGAGATATACATATG
>yagI
GAGGCAATTCCGACGTCTAAGAAACCATTATTATCATGACATTAACCTATAAAAATAGGCGTATCACGAGGCCCTTTCGTCTTCACCTC
GATGGATCCGAGGTGTTTGAGTCGGCGGCGAGCATCGTGTTCGACCAGGCGGAAAACCGGATGCATACGATTAAGGCGGTGATGATGGC
AACGCTTGGGGAGTGATTGGGTCCGCTGCGCGTTGGTGCCCTCACCCCGGCCCTCTCCCACAGGGAGAGGGAGAACACCGGCTCCATTT
CATTGATTTTTCATCCCGAAAAAGGTACGTTTTCGCCTTAATTCCAGCGTGGACATGCCAGCATTATGCCGATTATTCAGTCTGTTGAA
CGTGCGTTGCAGATCCTCGACCTGTTCAACGAGCAGGCCACCGAGCTTAAGCTCGAGAGATCCTCTAGATTTAAGAAGGAGATATACAT
ATG
>yagG
GAGGCAATTCCGACGTCTAAGAAACCATTATTATCATGACATTAACCTATAAAAATAGGCGTATCACGAGGCCCTTTCGTCTTCACCTC
GAGGGCACCTGGAAAGGCTGTATTTATGACACCGATAAAATTATCGAGGTAATTAACGCCGGTAAAAAAGCGCTCGGAATTTAATTATT
TTAAGAGATAAAACCGTCTGCGGAATATTTCCCGCAGACGGCTTTGTTGTTTTTGAAATTTATTAATTTAAACAATTAGTTGAGATATA
TCGTTGGCGTCACAAAAGCAAAATAACGTAATTCGGAAATAAGATATGACCATTGCTGGTTAATTGAATAGCTCATTACACTCCATTAA
CACGATGTTGTAATTCGGCACACTACATAAGGGTGTAATTCTGATGACGCAATTAACCATGAAAGACAAAATTGGCTACGGGCTGGGAG
ACACCGCCTGCGGCTTCGTCTGGCAGGCCACGATGTTCCTGCTGGGGATCCTCTAGATTTAAGAAGGAGATATACATATG
>yagE
GAGGCAATTCCGACGTCTAAGAAACCATTATTATCATGACATTAACCTATAAAAATAGGCGTATCACGAGGCCCTTTCGTCTTCACCTC
GAGCTGCGAGGCGAACAAAACAAACTCAGTACGTAATGACATGGTATCTCTCGCATCCCAGGGCATAAGCGACTCCATAAACGGGTTCT
TATGCCTTAGTTGTAAGTGTCTACCATGTCCCCGAACAAGTGTTCACTATGTCCCCGGACCGTACACCCCAAAGGGGAGAGGGGACTGC
ACCGAGCCATCTTTTCCCCCTCGCCCCTTTGGGGAGAGGGCCGGGGTGAGGGGCAATATGTGATCCAGCTTAAATTTCCCGCACTCCCT
CTTCCCTTCCGATTTACCTCTCCTTGTTCTGCGTCATAGTATGATCGTTAAATAAACGAACGCTGTTCTATAATGTAGAACAAAATGAT
TCAGCAAGGAGATCTCATGCCGCAGTCCGCGTTGTTCACGGGAATCATTCCCCCTGTCTCCACCGGATCCTCTAGATTTAAGAAGGAGA
TATACATATG
>ycgR
GAGGCAATTCCGACGTCTAAGAAACCATTATTATCATGACATTAACCTATAAAAATAGGCGTATCACGAGGCCCTTTCGTCTTCACCTC
GATGGATCCCCGGCATGATTAGCTTGGCGATAATGCCGGCTATCAGGTCAAAAATAATCCAGGCAATAATTCCCATGTTCAGGCCCTCG
TTTCGATGAATCGGACGCAAGAAATGCTTACATCAACGCTCTAAGTATAGGCAAGGGCGGGATCGATGCTGGGCGTATGTCGGCTGATT
TAAGAACAACATGTAAAAAATATTCACGCATTTGTATTAAGTTTTGTTAACTGTGACCGATAAACCAAAGACAGTTTGTCAGTCAGGAG
TTTTTCCGCGTGAGTCATTACCATGAGCAGTTCCTGAAACAAAATCCGTTAGCCGTCCTGGGCGTGTTACGCGATTTGCACAAAGCCGC
AATTCCTCTCGAGAGATCCTCTAGATTTAAGAAGGAGATATACATATG
>yagB
GAGGCAATTCCGACGTCTAAGAAACCATTATTATCATGACATTAACCTATAAAAATAGGCGTATCACGAGGCCCTTTCGTCTTCACCTC
GATGGATCCGGAAGACGGCAGCTACGAGGTGTGGTGGTACAGCACGAAAGTGGGGGTGATCGACCTGAAGAAAAAGTCGATCACCATGG
GTAAAGGATGTTAAAAAGTGTTCACCATGTCCCCGACCACCTGATTGTCGGCGGCAGGCAAATATTCTCATTCGCCGAACATGGTCTTC
TTTAACAGGAATAAAATTATGAATAAAGACTCTACTCAAACTTGGGGGCTTAAACGAGATATCACACCGTGTTTCGGAGCACGACTGGT
CTCGAGAGATCCTCTAGATTTAAGAAGGAGATATACATATG
>ycgI
GAGGCAATTCCGACGTCTAAGAAACCATTATTATCATGACATTAACCTATAAAAATAGGCGTATCACGAGGCCCTTTCGTCTTCACCTC
GAGGCCATTAACACCATTGCCAGTAAGTAAACCAACGCCTGCGCCTAGCAATGCACCTTTCGTGGTGCGGTTCATATCCGCCATCGCTG
GCGTGGAGCAGAACAATGCTGAGATAAGCCCGAAGGCAAGTATTTTTTTCTTCAACTTAGATGTCCGGTATTAAGTAAGTTGCACACAC
AATAATTTCGTCTTCAATTAAGATCTGCTTAACTAAAGAACGCTCGCTATTATTCAGATAATTCAAAATGAGCGTGGCTGTGATGATAG
GAATTATGTTTTTTACGTGAATGAGAATAATCTTAAATGAGGAATAACTCATTGATTGACAATATTTTTATTCAAGAAGTGTCATTGAC
TGTTAACGCAATGTTGTAAAGGTAAGATAATCTGATTTATCAATATTATTGTGTGATTTTTATGTGAGCAGAAGATATTCATCAGCAAC
GATTACATTAGTCATTTTATTTTGCCGACGGCCTCATTGTCGAAAGATAAGCGTACGACAGTATTATCAGAAAAGAGTGATTTTTTATC
CAACTACACTTCAGCGCACTGCGTGTAAAAAATGCCTCTTTCTTATGCGGGATATCATCATTTCATCATGATGTCTTTGGTGAGCGGTG
AACACAATACACCTGCGCTGTCTCTTCAGGATGAATCCCCTGGTCTGGTGCCTGCGGGCTGATGTTGCAGCAGAGCTTAGGTCACTTAG
ACGCTACTATCATTTATCCAATGGCATGGAATCGAAATCAGTCGATACCCGCAGTATATATCGTGAACTGGGTGCAACGCTGAGTTACA
ACATGCGCCTGGGGAACGGTATGGAAATGAACCCTGGCTGAAGGCGGCTGTGCGCAAAGAATTTGTCGATGATAACCGGGTGAAGGTGA
ATAATGACGGTAATTTCGTCAATGATTTGTCGGGCAGACGTGGATCCTCTAGATTTAAGAAGGAGATATACATATG
>ycgH
GAGGCAATTCCGACGTCTAAGAAACCATTATTATCATGACATTAACCTATAAAAATAGGCGTATCACGAGGCCCTTTCGTCTTCACCTC
GAGAAAATGCTGGAGGAAGCGGCGCGGCCACGCTAAACGGGATCGAACTCATAACTGTGAATGGCGATGAATCTCCAGCAGATGCCTTC
AGATAAGGTGATGCAAGGATTGCAGCTGGAGCTTTCGAATATCAACTAAAACAACAAGGCAAAAACTGGTATTTGACAAGTTATCAGTC
AGTGAATGAGGAGGATAACAGCTCAGAGGGAAATTCAGAATCAACAGAAACGCCTACCCCCGTCCTGCGCCCCGAAGCTGGAAGTTAGG
ATCCTCTAGATTTAAGAAGGAGATATACATATG
>ycgK
GAGGCAATTCCGACGTCTAAGAAACCATTATTATCATGACATTAACCTATAAAAATAGGCGTATCACGAGGCCCTTTCGTCTTCACCTC
GATGGATCCGGTCTGGTCACGCTTGCTGCTTCGATAAATCACACAAAACATACTTTTACTCTTTAAGATGCCAGGTTTTGGCATAAGCT
AACATCAATGAAACCCGCGCCACAAGCCAGCCAGCTGTAGATTTCTACGGTTTGTTGAGTCCATGCCCCCTTTATAAGTAATCCCCCCG
GCAAAAATGCCGCAACGTGATTTTACCGAACGCAGGAAAGCCACTGGAATTCTCCCGTTAGGGATTGTTATATTATAACAGTTCATCGT

ACTCATTCTGAACAGGAGACTACCAATGAAAATCAAGAGCATCAGGAAGGCTGTATTGCTGCTTGCACTGCTAACCTCCACGAGCTTTG
CTGCGGGTAACTCGAGAGATCCTCTAGATTTAAGAAGGAGATATACATATG
>ycgJ
GAGGCAATTCCGACGTCTAAGAAACCATTATTATCATGACATTAACCTATAAAAATAGGCGTATCACGAGGCCCTTTCGTCTTCACCTC
GAGCCAGCGCCTGATGGATAACCTTAGGTTCTGCCTCATGCAGATGAACCACAGATAAAGTGAAGCTACTGCCTTTAAGCTCGATTGGC
GTGTTTGACATCCTGGCCTTACTCAATTAGCTATTAATCATCGCCAGCGCGCGATGATGTTCCGAAGACTATAAGGCATGTTATAGTCT
GGATTATATTGAGGCAAGTCACCCTCCCATTTATTCAGAGTAAAAGTCTATTCTGTGATAAATGGCGCTGATTCATAGCTTAAAAAATA
CCCTTGTCAATCAACCCATTGCCGTCGTACTTTTGATTGTTCTTATTTACGCTTCCTTTTTCCGCACCCTAATCGTGACAGCATCTCCA
GACCTGTTTAAATATCCAGTATTGAGGCAAGTGTATTTTTTCACCACGCCAGGCTTTCTTAACTTTCCATTTTTTGCGGCATCATTAAA
GAACAGATACCCGTTATCTTAATATCCGCCACAGAATTGCTCTATCATAGAAACCGCAGAAGACGTCTGATACTTCTGTCAGGTCATCG
TAACTCGTCCAGGTCAGCGGTTTTGATTACGGATTGATGATGAAATGAGCATAAGTGTTGTACGGTTGGATTATCAGGATGTGCTATGA
ATATGATGCGTATTTTTTATATCGGATTGTCAGGTGTGGGGATGATGTTCTCATCAATGGCGTCTGGAAATGACGCTGGCGGACTTCAA
TCTCCGGATCCTCTAGATTTAAGAAGGAGATATACATATG
>ycgM
GAGGCAATTCCGACGTCTAAGAAACCATTATTATCATGACATTAACCTATAAAAATAGGCGTATCACGAGGCCCTTTCGTCTTCACCTC
GAGCACCACCCGAAGATTTGCTGAAGCAACATCTTTCCGTCATGGGGCAGAAAACAGACGACACTAACAAATAACCGATATCCGGCGGT
GGCATTATCTTTGTCGGCGCGGGTTTTCATATCCACGATAAGGTGAGGGGAACGTTATGTATCAACATCACAACTGGCAAGGTGCGCTG
CTGGATTATCCGGTGAGTAAAGTAGTCTGTGTTGGCAGTAACTATGCCAGGATCCTCTAGATTTAAGAAGGAGATATACATATG
>yagZ
GAGGCAATTCCGACGTCTAAGAAACCATTATTATCATGACATTAACCTATAAAAATAGGCGTATCACGAGGCCCTTTCGTCTTCACCTC
GATGGATCCAATGCTGAGGCCAAGCTGTACTCAAAAATATATAAGTTGGTTCAGTAAACTCCAGGCAAGTTAGTTTTAAAAAATGACTC
ACTGGGACATCACGTCCTCAATTCAACTCGGGAAGAAATGCAATGAAAAAAAAGGTTCTGGCAATAGCTCTGGTAACGGTGTTTACCGG
CATGGGTGTGGCGCAGGCTGCTGACGTAACAGCTCAGGCTGTAGCGACCTGGTCCTCGAGAGATCCTCTAGATTTAAGAAGGAGATATA
CATATG
>yagY
GAGGCAATTCCGACGTCTAAGAAACCATTATTATCATGACATTAACCTATAAAAATAGGCGTATCACGAGGCCCTTTCGTCTTCACCTC
GATGGATCCGCACTCTACCGGAAGGCATCTGGAGCGGCGACGTTAGCGTACAGTTCGACGCGACCTGGACCAGTTAATCTCTCTGATGT
ACCAGCAGGGGTAGCCCCCCTGCTTTATTCCTGGACGGAATTGTTATGAAAAAGCACCTTCTGCCTCTCGCTCTGCTGTTTTCCGGAAT
ATCTCCGGCCCAGGCGCTGGATGTCGGCGATATCTCGAGAGATCCTCTAGATTTAAGAAGGAGATATACATATG
>ycgN
GAGGCAATTCCGACGTCTAAGAAACCATTATTATCATGACATTAACCTATAAAAATAGGCGTATCACGAGGCCCTTTCGTCTTCACCTC
GAGTTGTGTTGACAGGCACGCCTGATGGCGTCGGCCCGTTGCAAAGCGGTGATGAGCTGACAGTCACTTTCGATGGGCATTCTTTGACA
ACTCGCGTTTTGTAATACTTTTTGCCGCCTGAAAGCGGCGGCAAAACTTGCATCGCTGTGCCAGACTGGTTATAAGGTGCGTTTTAACG
TAATGGCGGAACACCTGATGAGCGATGTACCTTTCTGGCAAAGTAAAACCCTGGACGAAATGAGCGATGCGGATCCTCTAGATTTAAGA
AGGAGATATACATATG
>yagU
GAGGCAATTCCGACGTCTAAGAAACCATTATTATCATGACATTAACCTATAAAAATAGGCGTATCACGAGGCCCTTTCGTCTTCACCTC
GAGGCGGCGCTCACTTTAATCAGATCGCGACGGGTCAAACTTAAATCGTGCGGCTCGTGCTTCCCAACCCGATTATCTTCGGGGTATTC
GCCTTGGTTGCTCATACCAGGCCTCCGGTATTCTAAAGGGGAAAATAAGAGTGCGTTGCGAGAGTGCTGATGCTTTTTTAAACGTTAAG
CATAGTCGGCGACAGCAGGAGGATTATTCTTAAAACGCGAATGCAGCTATGAGCCCGGCTAATAAATTCATGTTCGCCGGGATGTTGAT
GATGATGGGAGCTGGTTTATTGCTTTAGTTGTACGATGCAAAAACCAATAAGGAAACCTGTGATTTTCAGCTCTACATCACCCTGCAAA
TCTCTGTCACTTCTAATATAAAAATAGGGAGAAATGATGGAGCTTATATTCATTGGCGATTAGGAAACTATCTTGTTATACAAAACAAT
ACAGTTCTTTACATTTGCCTTGTTTTATGAATACTCCTGAAGAGGTGTATAACATAATGGTACAAGCAGGGTAGATATGAATATATTTG
AACAAACTCCACCGAACCGCAGACGTTATGGTCTTGCTGCATTCATTGGGCTGATTGCTGGCGTTGTTTCCGCATTCGTGAGGATCCCG
CACGAATGCGGAAACGGGATCCTCTAGATTTAAGAAGGAGATATACATATG
>yagT
GAGGCAATTCCGACGTCTAAGAAACCATTATTATCATGACATTAACCTATAAAAATAGGCGTATCACGAGGCCCTTTCGTCTTCACCTC
GATGGATCCTCACGAATGCGGAAACAACGCCAGCAATCAGCCCAATGAATGCAGCAAGACCATAACGTCTGCGGTTCGGTGGAGTTTGT
TCAAATATATTCATATCTACCCTGCTTGTACCATTATGTTATACACCTCTTCAGGAGTATTCATAAAACAAGGCAAATGTAAAGAACTG
TATTGTTTTGTATAACAAGATAGTTTCCTAATCGCCAATGAATATAAGCTCCATCATTTCTCCCTATTTTTATATTAGAAGTGACAGAG
ATTTGCAGGGTGATGTAGAGCTGAAAATCACAGGTTTCCTTATTGGTTTTTGCATCGTACAACTAAAGCAATAAACCAGCTCCCATCAT
CATCAACATCCCGGCGAACATGAATTTATTAGCCGGGCTCATAGCTGCATTCGCGTTTTAAGAATAATCCTCCTGCTGTCGCCGACTAT
GCTTAACGTTTAAAAAAGCATCAGCACTCTCGCAACGCACTCTTATTTTCCCCTTTAGAATACCGGAGGCCTGGTATGAGCAACCAAGG
CGAATACCCCGAAGATAATCGGGTTGGGAAGCACGAGCCGCACGATTTAAGTTTGACCCGTCGCGATCTGATTAAAGTGAGCGCCGCCT
CGAGAGATCCTCTAGATTTAAGAAGGAGATATACATATG
>fldB
GAGGCAATTCCGACGTCTAAGAAACCATTATTATCATGACATTAACCTATAAAAATAGGCGTATCACGAGGCCCTTTCGTCTTCACCTC
GAGCAGATCGCGACGGTAAGCGTTCAACGTATTTTCAGCCAGATTTTTTTCCAGCCACAGAGCATCAAGAAACTGCTCGATGCGTGCCA
GATCCTGTTTCACTTGCGCCCCTTATGGTCACTCATTTGATCCATTATGCCTTATTGTGCCGTGACTAAAGCGATTCTGATACACTAGC
CGCAAAAGCCACAGCAGAATCGAGAAGCTTACGTTATGAATATGGGTCTTTTTTACGGTTCCAGCACCTGTTACACCGAAATGGCGGCA
GAAAAAATCCGCGATATTATCGGCCCAGAACTGGTGACCTTGGATCCGGGGATCCTCTAGATTTAAGAAGGAGATATACATATG
>ycgF

GAGGCAATTCCGACGTCTAAGAAACCATTATTATCATGACATTAACCTATAAAAATAGGCGTATCACGAGGCCCTTTCGTCTTCACCTC
GATAGATCTAACGAGTGATTGCTCCCGCAGAATCTAAAGTCACTGAATTTTGATGCATGCTACGCCTCTGTTAAAAATGTTAATCAATG
TTAAGTGAAGTAACAATAGCAAACTTGTACGAAATATGTGTACAGAATGATAAATTAATTTTTTGCAATTAGTGCTAATGCATATAAAA
CAATGTGTTATGATAAAACATATTGCATCATTTAACTGACAATCTATACAAGATTTTTCGCCAACTAGTGTTCAACTTTTTCCATTTTT
TATCAATGTAAAGAAACGGTTATGTTTTGGTATTAAAAGAAAGACACAAGAAGATATGTCTGTTACCGTCTTACTCTCGCCTCACCCAT
TACCCTGGATTGTTAATGCTTACCACCCTTATTTATCGTAGCCATATACGTGACGACGAACCTGTCAAAAAAATCGAAGAAATGGTTTC
GATAGCAAATCGCAGGAACATGCAGTCCTCGAGAGATCCTCTAGATTTAAGAAGGAGATATACATATG
>yhbP
GAGGCAATTCCGACGTCTAAGAAACCATTATTATCATGACATTAACCTATAAAAATAGGCGTATCACGAGGCCCTTTCGTCTTCACCTC
GATGGATCCATAGCGGCGTTCGACATCCGTGGTGATCCCGGTATAAAGCTTATTGTCGGCGGTACGGATCAAGTAAAGAAACCAGGGTG
TCATCGTCTGCGTCGCATGTTAAGGTCAGGCCCACAGTATAAAAGAGAACAGCAATGGAAACACTCATCGCCATCAGCCGTTGGCTGGC
AAAACAACATGTTGTCACTTGGTGTGTGCAGCAGGCTCGAGAGATCCTCTAGATTTAAGAAGGAGATATACATATG
>pyrH
GAGGCAATTCCGACGTCTAAGAAACCATTATTATCATGACATTAACCTATAAAAATAGGCGTATCACGAGGCCCTTTCGTCTTCACCTC
GAGGTGGGTGAAGGCATCGAGAAAGTTGAGACTGACTTTGCAGCAGAAGTTGCTGCGATGTCCAAGCAGTCTTAATTATCAAAAAGGAG
CCGCCTGAGGGCGGCTTCTTTTTGTGCCCATCTTGTAAATTCAGCTAACCCTTGTGGGGCTGCGCTGAAAAGCGACGTACAATGTCGCT
AGTATTAATTCATTTCAATCGTTGACAGTCTCAGGAAAGAAACATGGCTACCAATGCAAAACCCGTCTATAAACGCATTCTGCTTAAGT
TGAGTGGCGAAGCTCTGCAGGGCACTGAAGGCTTCGGTATTGATGCAAGGATCCTCTAGATTTAAGAAGGAGATATACATATG
>pyrG
GAGGCAATTCCGACGTCTAAGAAACCATTATTATCATGACATTAACCTATAAAAATAGGCGTATCACGAGGCCCTTTCGTCTTCACCTC
GATGGATCCGGCAACAGGTAAAACGGCAGGAAATTGATCTCTAAGGGAATTACGCGGTCAAGCGCCATTTGTGTCATTTTTTAAATGAC
AAGCGCTTGATTTGCGTCAAAAACATTTACCCCAAAGGGGCTATTTTCTCACTCCTGATTTCAATAGTGCGCTGGCGAAGAGGAGGGAT
AATGAAAGTTTGTGGCACAGGTCATGTTCGGGTATACTGCTTTCCCGTCTTGGTTATTCCATCGTCTTTTCAACCTAACTTCTCAGGTT
CAGCATGACAACGAACTATATTTTTGTGACCGGCGGGGTCGTATCCTCTCTGGGTAAAGGCATTGCCGCAGCCTCCCTCGCAGCCATTC
TTGAAGCCCGTGCTCGAGAGATCCTCTAGATTTAAGAAGGAGATATACATATG
>pyrC
GAGGCAATTCCGACGTCTAAGAAACCATTATTATCATGACATTAACCTATAAAAATAGGCGTATCACGAGGCCCTTTCGTCTTCACCTC
GATGGATCCGAAGTGAAGCCAGGCGAAATTGTCATTCCATTTACTGATTAATCACGAGGGCGCATTCGCGCCCTTTATTTTTCGTGCAA
AGGAAAACGTTTCCGCTTATCCTTTGTGTCCGGCAAAAACATCCCTTCAGCCGGAGCATAGAGATTAATGACTGCACCATCCCAGGTAT
TAAAGATCCGCCGCCCAGACGACTGGCACCTTCACCTCCGCGATGGCTCGAGAGATCCTCTAGATTTAAGAAGGAGATATACATATG
>pyrB
GAGGCAATTCCGACGTCTAAGAAACCATTATTATCATGACATTAACCTATAAAAATAGGCGTATCACGAGGCCCTTTCGTCTTCACCTC
GATGGATCCGCCGTTTTTCTTCCCGTTGATCACCCATTCCCAGCCCCTCAATCGAGGGGCTTTTTTTGCCCAGGCGTCAGGAGATAAA
AGATGGCTAATCCGCTATATCAGAAACATATCATTTCCATAAACGACCTTAGTCGCGATGACCTTAATCTGGTGCTGGCGACAGCGGCG
AAACTGAAACTCGAGAGATCCTCTAGATTTAAGAAGGAGATATACATATG
>glpX
GAGGCAATTCCGACGTCTAAGAAACCATTATTATCATGACATTAACCTATAAAAATAGGCGTATCACGAGGCCCTTTCGTCTTCACCTC
GATGGATCCGCGTGGGAAGAACACGACGAATAATGTAAATGCCGAATGAAGCGTTTATGCCGCATCCGGTAGTCCCGAAACGTGCGGGG
GCAACCCCGCACACATCAATAATCCCTCCCTTCCCCTGTGCTACACTTCGCGCCATTCCTTACTGCTTAGAGTTTGCTATGAGACGAGA
ACTTGCCATCGAATTTTCCCGCGTCACCGAATCAGCGGCGCTGGCTGGCTACAAATGGTTAGGACGCGGCTCGAGAGATCCTCTAGATT
TAAGAAGGAGATATACATATG
>glpE
GAGGCAATTCCGACGTCTAAGAAACCATTATTATCATGACATTAACCTATAAAAATAGGCGTATCACGAGGCCCTTTCGTCTTCACCTC
GATGGATCCCGCCCCCTATCACAATCAGATCTTTGGTTTCCATGCTGCCCTCATTCACTTTCGTTAAAGCTCATAAATGTTCGTTATCG
AACATATTAGCAAAGAATCGCGCTTTAGGTAACATTGAAAAAACATTTTAGAGTGATATGTATAACATTATGGCGTTTATCTGCCGCTT
CGACGTAAACTGTGCGGTAAATTTGCCCACTTGTTTGTAAAGAAAGAGAGACGCATGGATCAGTTCGAATGTATTAACGTTGCCGACGC
GCACCAGAAGTTGCAGGAAAAAGAGGCGGTGCTGGTCGATATTCTCGAGAGATCCTCTAGATTTAAGAAGGAGATATACATATG
>glpD
GAGGCAATTCCGACGTCTAAGAAACCATTATTATCATGACATTAACCTATAAAAATAGGCGTATCACGAGGCCCTTTCGTCTTCACCTC
GAGAATATCGACCAGCACCGCCTCTTTTTCCTGCAACTTCTGGTGCGCGTCGGCAACGTTAATACATTCGAACTGATCCATGCGTCTCT
CTTTCTTTACAAACAAGTGGGCAAATTTACCGCACAGTTTACGTCGAAGCGGCAGATAAACGCCATAATGTTATACATATCACTCTAAA
ATGTTTTTTCAATGTTACCTAAAGCGCGATTCTTTGCTAATATGTTCGATAACGAACATTTATGAGCTTTAACGAAAGTGAATGAGGGC
AGCATGGAAACCAAAGATCTGATTGTGATAGGGGGCGGCATCAATGGTGCTGGTATCGCGGCAGACGCCGCTGGACGCGGTTTATCCGT
GCTGATGCTGGGGATCCTCTAGATTTAAGAAGGAGATATACATATG
>glpA
GAGGCAATTCCGACGTCTAAGAAACCATTATTATCATGACATTAACCTATAAAAATAGGCGTATCACGAGGCCCTTTCGTCTTCACCTC
GAGCAGCGCAATCGACGATAAGTCGGATCGATCTCCGCGGCAGGTAAGCGCGCTTTGTGTGGCGCTGGTTTAAAAATACTCAACATTGA
TAGCCTCCGTGGCCCGTGGTCTTATTTATGATTAACAGCCTGATTCAGTGAGAGAACCTGCCGTTTCTTGAGTTGCCGCGATGTTAAGA
AAACATTCATAAATTAAATGTGAATTGCCGCACACATTATTAAATAAGATTTACAAAATGTTCAAAATGACGCATGAAATCACGTTTCA
CTTTCGAATTATGAGCGAATATGCGCGAAATCAAACAATTCATGTTTTTACTATGGCTAAATGGTAAAAAACGAACTTCAGAGGGATAA
CAATGAAAACTCGCGACTCGCAATGGATCCTCTAGATTTAAGAAGGAGATATACATATG
>yciW

GAGGCAATTCCGACGTCTAAGAAACCATTATTATCATGACATTAACCTATAAAAATAGGCGTATCACGAGGCCCTTTCGTCTTCACCTC
GATGGATCCTCTCTGCCGGTATCTCCGGTGAAGTGGTCCACGTTGACGGCGGTTTCAGCATTGCTGCAATGAACGAACTCGAACTGAAA
TAATCGTTCTGTTGGTAAAGATGGGCGGCGTTCTGCCGCCCGTTATCTCTGTTATACCTTTCTGATATTTGTTATCGCCGATCCGTCTT
TCTCCCCTTCCCGCCTTGCGTCAGGATAACGATTTCCTTTACGACCAAGGAGCGCCTCGAGAGATCCTCTAGATTTAAGAAGGAGATAT
ACATATG
>fxsA
GAGGCAATTCCGACGTCTAAGAAACCATTATTATCATGACATTAACCTATAAAAATAGGCGTATCACGAGGCCCTTTCGTCTTCACCTC
GAGCTTCCCTGGTACCCAACAGATCTTCTTCGATACGAATGTTGTTTGACATGTGAACCTTCTTTTTCAAGCTGCCAATGATTTGCTTT
AAACACACAGAATATATGTGGTTTCGAATGTTTTTCGACCGACGATTATCCCCTGCATCGACCGAATACCCGAGATCATATGCTGCTTG
AGGATTTCTACCGTAATCTGGATCACTTTAAGTGTCGGTTTTTACCCCTTAATTATTAATTTGTGAAATAGATCACCGCTTTGGGATTA
CTACCAAAAATAGTTGCGCAAACATCTTGAAATTTTGCTAATGACCACAATATAAGCTAAACGCGATTCGCAACCCATTCAGGTAGCCG
GGGTTAACCGGCTGCTATTACAGGAGAAACCTTTGCGCTGGTTACCTTTTATTGCCATTTTCCTTTATGTCTATATTGAGATTTCAATC
TTTATTCAGGTTGCCCATGTATTGGGGGTATTGCTGACCCTCGTGCTGGTTATATTCACGTCAGTTATCGGTATGTCACTGGTACGTAA
CCAGGGCTTTAAGAATTTCGTGCTGATGCAGCAAGGATCCTCTAGATTTAAGAAGGAGATATACATATG
>yciT
GAGGCAATTCCGACGTCTAAGAAACCATTATTATCATGACATTAACCTATAAAAATAGGCGTATCACGAGGCCCTTTCGTCTTCACCTC
GATGGATCCGAACGGGATCAATGAGCGGCAAGGATTTTTGTTTGCCAAACCGATGCCCGCCGTCGCCTTCGAACGCTGGTATAAACGCT
ATCTGAAGCGCGCATAAGTAGATGAATATATATACTTAGCGCTATATCCCGTCATTATATATCGTATTCCTGGATGATAATTAGCCGTT
GCTGTTAAACACTTCTCTTTTAGCGGCAATTTTTGCATTATTCATTTTCCCCCTTTTGCTATAAGGATGTCCTTGATGTCTGAATTTGA
TGCCCAGCGCGTTGCCGAACGTATTGATATTGTGTTAGATATCCTTGTGGCAGGCGATTACCACTCTGCGATCCATAATCTCGAGATAC
TTAAAGCAGAACTCTTGCGCCAGGTTGCTGAATCTACGCCAGACATCCCGAAGGCACCGTGGGAGATTTAACAATCGTGCCTTCAGGTT
TTCCTATTTGAGGATGCCCGGCGCGTCAGTATTTTTTTATTAAGTATTATAACGTTATAAGAATTACAGCGATGAATTCCCGACAACAA
ACTATTCTACAGATGGTCATTGACCAGGGTCAGGTTAGCGTAACCGATCTGGCAAAAGCCACGTCGAGAGATCCTCTAGATTTAAGAAG
GAGATATACATATG
>yciS
GAGGCAATTCCGACGTCTAAGAAACCATTATTATCATGACATTAACCTATAAAAATAGGCGTATCACGAGGCCCTTTCGTCTTCACCTC
GAGAACGAATTTGTGGGCCATTAACACCACCTGCGGAAGAAAATCGCGAAATAGCGCAACGAGAACAAGAAAGTTAACGCTGGTTGATT
TTCCGAATTTAGCCCTTAAATCATCAACAATGCGTGTGGATGCCATTTCGCAGACGGCGCGAAAATGGTACTTTAAAGGGCTATTGCGG
TAAGTTGACCATAATTTATTCGCTCTAACCACATAACGGGAAGTAATGTGAAATATTTACTCATTTTCTTACTGGTGTTAGCGATCTTC
GTGATTTCGGTCACGTTGGGTGCGCAGAACGATCAACAGGGGATCCTCTAGATTTAAGAAGGAGATATACATATG
>edd
GAGGCAATTCCGACGTCTAAGAAACCATTATTATCATGACATTAACCTATAAAAATAGGCGTATCACGAGGCCCTTTCGTCTTCACCTC
GATGGATCCGGCGATGATTACCCGTGATGGTCGTTCCTGGAATGAGTTTGAGTAATATCTGCGCTTATCCTTTATGGTTATTTTACCGG
TAACATGATCTTGCGCAGATTGTAGAACAATTTTTACACTTTCAGGCCTCGTGCGGATTCACCCACGAGGCTTTTTTTATTACACTGAC
TGAAACGTTTTTGCCCTATGAGCTCCGGTTACAGGCGTTTCAGTCATAAATCCTCTGAATGAAACGCGTTGTGAATCATCCTGCTCTGA
CAACTCAATTTCAGGAGCCTTTATGAATCCACAATTGTTACGCGTAACAAATCGAATCATTGAACGTTCGCGCGAGACTCGCTCTGCTT
ATCTCGCCGTCGAGGTGAAGACGAAAGGGCCTCGAGAGATCCTCTAGATTTAAGAAGGAGATATACATATG
>yciG
GAGGCAATTCCGACGTCTAAGAAACCATTATTATCATGACATTAACCTATAAAAATAGGCGTATCACGAGGCCCTTTCGTCTTCACCTC
GATGGATCCCTGGCGGCACTGAAAGTTTTTGTACAACCGATGAAAGCGGCGACGCGCAGTTAATCCCACAGCCGCCAGTTCCGCTGGCG
GCATTTTAACTTTCTTTAATGAAGCCGGAAAAATCCTAAATTCATTTAATATTTATCTTTTTACCGTTTCGCTTACCCCGGTCGAACGT
CAACTTACGTCATTTTTCCGCCCAACAGTAATATAATCAAACAAATTAATCCCGCAACATAACACCAGTAAAATCAATAATTTTCTCTA
AGTCACTTATTCCTCAGGTAATTGTTAATATATCCAGAATGTTCCTCAAAATATATTTTCCCTCTATCTTCTCGTTGCGCTTAATTTGA
CTAATTCTCATTAGCGACTAATTTTAATGAGTGTCGACACACAACACTCATATTAATGAAACAATGCAACGCAACGGGAGAAATAACAT
GGCCGAACATCGTCTCGAGAGATCCTCTAGATTTAAGAAGGAGATATACATATG
>b1437
GAGGCAATTCCGACGTCTAAGAAACCATTATTATCATGACATTAACCTATAAAAATAGGCGTATCACGAGGCCCTTTCGTCTTCACCTC
GATGGATCCATATAACCGCCTTCCGGCGCGGGTGTAAGAGTGACGGGATAACGCATAATTTCGACTGTCTCTCGCATGTCATTAACCTT
TTTAATTGGCGATTAACTCAAACCGAGTTGTTTCAGGATTGCTTTACGCAATGGTTCTTTAATCTCATCGCAGGGGTGACGCGGCATGA
CACTGCGCCTCCCATGAAACCTGAGTTTCAAATGGTTGCTGCCACTCGAGAGATCCTCTAGATTTAAGAAGGAGATATACATATG
>yciC
GAGGCAATTCCGACGTCTAAGAAACCATTATTATCATGACATTAACCTATAAAAATAGGCGTATCACGAGGCCCTTTCGTCTTCACCTC
GATGGATCCAATTCGCCTGCTTCATGCGCAAAGGCACTGCCAGAGAGAAGAGTTGTTACTGCCAAAGCCGCCACTGTTAACTTTTTCAT
ATCCGCTCCGTCGTTATGGTTATAAAAACGTGACGAATATACCTACATTGGAGTAACAAATGATCCTACATAGATCACATTACATATGA
TAATTCACCCTTTATTGATCTGGATTAATTTAGAAGTTGCATTGAGAATCCTGGTTAATGTGATTTAAATCAATTTTTAAATATTTTCG
CTACGCAAATATTAACTCGGTCGCCACTTGCCGCGATTTGCCCATGCCTGAAGGCCAGCAATTACCTCCGCTTTACAAAGATATGCTTT
TCCATACAAGACCTGATGCTGCCACCCGAGGCTTAACCAGGGTACAATTGCCCGCTAATTAACACCTGCATAAAACTCAAGGAGAGTGC
ATGTCTATCACGGCGCAGTCCGTATACCGTGACACCGGAAATTTCTTCCGTAATCAATTTATGACCATTCTGTTGGTATCGTTGCTATG
TGCGCTCGAGAGATCCTCTAGATTTAAGAAGGAGATATACATATG
>yciA
GAGGCAATTCCGACGTCTAAGAAACCATTATTATCATGACATTAACCTATAAAAATAGGCGTATCACGAGGCCCTTTCGTCTTCACCTC
GATGGATCCCCTGACCGCCCTTACCTTAATCTTCACATTGTTAAGCGGTATCTATATCTACCGCCACATGCCGCAGGAAGATAAATCCT
AACCAGACTGCCAGTCCTTCGGGGCTGGCGCTGTATCTCCTCTGTTCTCCTCAGGAAAATCATAGTAGCATCGCGCCTGTGATTTTCCT

```
TTTAAGTCGGTTTTACCATGTCTACAACACATAACGTCCCTCAGGGCGATCTTGTTTTACGTACTTTAGCCATGCCCGCCGATACCAAT
GCCAATCTCGAGAGATCCTCTAGATTTAAGAAGGAGATATACATATG
>phoH
GAGGCAATTCCGACGTCTAAGAAACCATTATTATCATGACATTAACCTATAAAAATAGGCGTATCACGAGGCCCTTTCGTCTTCACCTC
GAGCTATCGGCGGCGGTTATTTTTTTGCGCTGCCGGGGGTGAAGGACGCGAACGATTATTTCGGAAGCGCGTTATTGCGGGTTTAATGT
TTTTAGGCGGATAAGGCATTTGTGCGCAGATGCCTGATGCGACGCTTGCGCGTCTTATCATGCCTACAATCAGTGCGGGTTTGGTAGGC
TGGATAAGGCGTTCACGCCGCATCCGGCGATCGTGCACTGATGCCTGATGCAAATCCTGCTGAAAGCACACAGCTTTTTTCATCACTGT
CATCACTCTGTCATCTTTCCAGTAGAAACTAATGTCACTGAAATGGTGTTTTATAGTTAAATATAAGTAAATATATTGTTGCAATAAAT
GCGAGATCTGTTGTACTTATTAAGTAGCAGCGGAAGTTCCCGGCAGTGATAGTCAGTCACTATGGAGATCGCGGATGGTAACGTCCTGT
ACTGGACATGTTTTAGATAATCAACGCGCCACCACTCGCGGAGTTTTCTCCTCCGGGGATCCTCTAGATTTAAGAAGGAGATATACATA
TG
>ygcU
GAGGCAATTCCGACGTCTAAGAAACCATTATTATCATGACATTAACCTATAAAAATAGGCGTATCACGAGGCCCTTTCGTCTTCACCTC
GATGGATCCTTGGTGCAGATCGCGTAATTACCGATGAAACAGTATTAAAGAAAAACAGTATTGACCGTTTTCGTAAATTTCCGGATATT
CATGGCATTTATACTTTGCCGATTCCGGCAGCGGTCGTAAAACTCGGTTCCACAGAGCAAGTATCCCGTGTGCTGAATTTTATGAATGC
GCACAAAATTAACGGTGTGCCGCGTACCGGTGCTTCCGCCACCGAAGGTGGGCTGGAAACTGTTGTAGAAAACTCGGTGGTGCTCGCTC
GAGAGATCCTCTAGATTTAAGAAGGAGATATACATATG
>yciK
GAGGCAATTCCGACGTCTAAGAAACCATTATTATCATGACATTAACCTATAAAAATAGGCGTATCACGAGGCCCTTTCGTCTTCACCTC
GATGGATCCTGACAATAATGGCGGCAATCGCCGCAATCGCTAGCACAACGGTAACGATTTTCGCCAAAAACAAACCATATTCAGACAAC
AATTCCACGAGTCCCCACCTTGGTTAAACAACGCTTTGGCTCACAGTTTACAACAGGCATGTTAAGCCGTCTCGCAGAGATGTTATTGC
AGCGAAAGTATGACAAAGTATCCATTTTGCGGTGTTATCGCCCCAGGCTGATTGAAATGCCCTTCTGTTTCAGGCATAAAGCCCCAAAG
TCATAAAGTACACTGGCAGCGCGACAGACAACGTCGTTGAGGAGTAGCCATGCATTACCAGCCAAAACAAGATTTACTCAATGATCGCA
TTATCCTGGTGACGGGAGCCAGCGATGGTATTGGTCGTGAAGCCGCGATGACGTATCTCGAGCGGGATCCTGACAATAATGGCGGCAAT
GACGTATCTCGAGAGATCCTCTAGATTTAAGAAGGAGATATACATATG
>yciI
GAGGCAATTCCGACGTCTAAGAAACCATTATTATCATGACATTAACCTATAAAAATAGGCGTATCACGAGGCCCTTTCGTCTTCACCTC
GATGGATCCGGAAGCGGCGAGGTAAATCAAGGGTCATTGAAGTCATAATCATTTCAGTAGAAAAACCAGGTCTCGATTTTAAATGCAAA
TAGCAATCATATTCAATAAGGCAAGACGATCAGGATCAGTGAAAATGTCATTTTTGGCGTAATTACCCTTACTTTATGCATGGCCTTAA
CAAAGACATCATCTTTCCGTTGCAGATCTTCGCCCTTTCAAATAACGTACTTTACAACTTTCCCGAACAAGGAGTTGTGCCCGCTCGAG
AGATCCTCTAGATTTAAGAAGGAGATATACATATG
>yoaF
GAGGCAATTCCGACGTCTAAGAAACCATTATTATCATGACATTAACCTATAAAAATAGGCGTATCACGAGGCCCTTTCGTCTTCACCTC
GATGGATCCTTTGCAGGCATGATAAAACATCATCAACAAAGATGAAACAAGACGTAATATGTTTTGAGTGAATGTAATAAAAATAGTGC
TGCAAAACAAAACGATACCGCGATACTCTAATCTTGGATAAATATGTGAGATCAATAAATGAAAATTATCAGTTTTGTTCTGCCTTGCT
TGCTGGTCCTGGCTGGTTGTTCAACCCCTTCTCAGCCACTCGAGAGATCCTCTAGATTTAAGAAGGAGATATACATATG
>mdtH
GAGGCAATTCCGACGTCTAAGAAACCATTATTATCATGACATTAACCTATAAAAATAGGCGTATCACGAGGCCCTTTCGTCTTCACCTC
GATGGATCCACCAGACGCACGACCAGTCGGTCTGTGGTTAAGCGCACTTTTGGCACGTTACTGCGATAGCCAAACATTCTATACCTACT
CCTTCCCGTAACGTCTTTAATACGCTGTAAGTATTACTATACCTGCGCAACCAGACCCTGTGAAAAGTCCAAAATGCCATTTTCTGCAT
TATTCACATATTTGATAAAAACTCTCTATCAAAGCGACGTTTTGATAAAAAAATATTGTCCCCGAAGTGATGGGAAAAGTCTTAGCGGC
AATGCAGAATAGAAAGTGCTCATCTTTTATTTCCCGGGAGGGGAAATGTCCCGCGTGTCGCAGGCGAGGAACCTGGGTAAATATTTCCT
GCTCATCGATAATATGCTGGTCGTGCCTCGAGAGATCCTCTAGATTTAAGAAGGAGATATACATATG
>codB
GAGGCAATTCCGACGTCTAAGAAACCATTATTATCATGACATTAACCTATAAAAATAGGCGTATCACGAGGCCCTTTCGTCTTCACCTC
GAGAGGACGCGATCCTGGAGATCTGACGACCATTGATGATCCTGCGTCGTTGGATCAGATCCGCCAGGCGATGGAAGAGTAGGTTATTG
TCGGATGCGTCGCGCGGTGCATCCGGCACTGTGTGCCGATGCCTGATGCGACGCTGACGCGTTTTATCATGCCTACGGACCTGAACCGT
AGGTCGGATAAGGCGCTCGCGTCGCATCCGACACCATGCTCAGATGCCTGATGCGACGCTGACGCGTCTTATCAGGCCTACCCACTGTT
TTTACACCGATAATTTTTCCCCCACCTTTTTGCACTCATTCATATAAAAAATATATTTCCCCACGAAAACGATTGCTTTTTATCTTCAG
ATGAATAGAATGCGGCGGATTTTTTGGGTTTCAAACAGCAAAAAGGGGGAATTTCGTGTCGCAAGATAACAACTTTAGCCAGGGGCCAG
TCCCGCAGTCGGCGCGGAAAGGGGTATTGGCATTGACGGGATCCCGCTCGAGAGGACGCGATCCTGGAGCGGGGGGTATTGGCATTGAC
GGGATCCTCTAGATTTAAGAAGGAGATATACATATG
>ygcW
GAGGCAATTCCGACGTCTAAGAAACCATTATTATCATGACATTAACCTATAAAAATAGGCGTATCACGAGGCCCTTTCGTCTTCACCTC
GATGGATCCTCGCGAGGGTTATCCAACGGCGATATGAGTTGTGTTGCATATCGTATTCCCATTTGTAATAGACAAAAAAAAGAGAAAAG
TAAGCCGCCCTTTCAGGTGGATTACTTTTCTCAGGTCTCTAGTAATTGAAATAATAAATAACATGGAAAGCAGAGGTGTTGTGTGAGAG
TAATCACGCTTCTGCTTTTTAAAATTATTTGACAATACGAATATCACACTGTGATTACGATCACATTAGCAGTTATGCCCTTATGCTAA
AAATTATCTCCAGTTACCAGAGATAATGAGAAAGGTAACTTCCCTCTCGCGGGGGAAGTTACCTTTCTTTTTTTTTGCCTTACGGAAAT
TGGAGCCTCGAGAGATCCTCTAGATTTAAGAAGGAGATATACATATG
>yihR
GAGGCAATTCCGACGTCTAAGAAACCATTATTATCATGACATTAACCTATAAAAATAGGCGTATCACGAGGCCCTTTCGTCTTCACCTC
GATGGATCCGCTGCATTGCCTGGTGATCCCGCGTATCCCGTTAGCGCCCGGCATGGCTCCAGCGGTTGCAGCGGGTCTGCTGGATATTA
ATGCGAAATAACTGATCTGAAAATAGGGTGCGGGTTTTCCCCTCTCCCTAACCCTCTCCCCAAAGGGGCGAGGGGACCGTCCGGTGCAG
```

Data Set S4 - MG1655 library reference file.fasta
TTTTTGGGTTTGACATCAGCCTCACGGGAGGCTGTATGTCGTTAATTAAGGTGCCTTGTATGCAAATCACCAATATGCATTGCAGTGGA
CAGACCGTTTCACTTGCCGCTCGAGAGATCCTCTAGATTTAAGAAGGAGATATACATATG
>zupT
GAGGCAATTCCGACGTCTAAGAAACCATTATTATCATGACATTAACCTATAAAAATAGGCGTATCACGAGGCCCTTTCGTCTTCACCTC
GAGCGTTCATCGGACTACCGTGACCTAAAAACAATGCTGGCATACGTGTTGAAGACATGATGATATCCTTAACTAAAGGTGTCATTTTG
ATATCCTCACAATACGCTTGTTCGGCGGAGTAAGAACCCGGATAACAATGATGATGATCATCAGTTATTTTGACGATCTGCCTGAAGGT
GAAGATTTATAAGGAGTTGTCGATGTCAGTACCTCTCATTCTGACCATACTGGCGGGGGCAGCCACGTTTATTGGCGCGTTTCTCGGCG
TTCTCGGGCAAAAACCCTCGAACCGCTGGATCCTCTAGATTTAAGAAGGAGATATACATATG
>katE
GAGGCAATTCCGACGTCTAAGAAACCATTATTATCATGACATTAACCTATAAAAATAGGCGTATCACGAGGCCCTTTCGTCTTCACCTC
GAGCGTGCGTGGGACATAGCTAATAATCTGGCGGTTTTGCTGGCGGAGCGGTTTCTTCATTACTGGCTTCACTAAACGCATATTAAAAA
TCAGAAAAACTGTAGTTTAGCCGATTTAGCCCCTGTACGTCCCGCTTTGCGTGTATTTCATAACACCGTTTCCAGAATAGTCTCCGAAG
CGGGATCTGGCTGGTGGTCTATAGTTAGAGAGTTTTTTGACCAAAACAGCGGCCCTTTCAGTAATAAATTAAGGAGACGAGTTCAATGT
CGCAACATAACGAAAAGAACCCACATCAGCACCAGTCACCACTACACGATTCCAGCGAAGCGAAAGGATCCTCTAGATTTAAGAAGGAG
ATATACATATG
>katG
GAGGCAATTCCGACGTCTAAGAAACCATTATTATCATGACATTAACCTATAAAAATAGGCGTATCACGAGGCCCTTTCGTCTTCACCTC
GAGAGCCGTGAAGGAGTGAAAGATTTCCACTTCTATACGCTTAACCGTGCTGAAATGAGTTACGCGATTTGCCATACGCTGGGGGTTCG
ACCTGGTTTATAAATAGTGTGGCTTTTGTGAAAATCACACAGTGATCACAAATTTTAAACAGAGCACAAAATGCTGCCTCGAAATGAGG
GCGGGAAAATAAGGTTATCAGCCTTGTTTTCTCCCTCATTACTTGAAGGATATGAAGCTAAAACCCTTTTTTATAAAGCATTTGTCCGA
ATTCGGACATAATCAAAAAAGCTTAATTAAGATCAATTTGATCTACATCTCTTTAACCAACAATATGTAAGATCTCAACTATCGCATCC
GTGGATTAATTCAATTATAACTTCTCTCTAACGCTGTGTATCGTAACGGTAACACTGTAGAGGGGAGCACATTGATGAGCACGTCAGAC
GATATCCATAACACCACAGCCACTGGCAAATGGGATCCTCTAGATTTAAGAAGGAGATATACATATG
>racR
GAGGCAATTCCGACGTCTAAGAAACCATTATTATCATGACATTAACCTATAAAAATAGGCGTATCACGAGGCCCTTTCGTCTTCACCTC
GATGGATCCCCACCGACAACAGCACAAGCTTGCTTGAATGAATAGTTCTCTTTTTTCATCGAATGAACTCCAAAAACACACAGAAATAT
TAGGCGACGCCTAACGCAATTGTCAATAGGCTGTGCCTAATGCAGTAAGGGTAGGGATTGCCTAATGTAATGCGCATAGGAGAATATTA
AGCAATGCTTAGTGGTAAAGACTTAGGCCGAGCGATAGAGCAGGCCATTAACAAAAAAATCGCATCGGGATCCGTCAAATCAAAGGCGG
AGGCTCGAGAGATCCTCTAGATTTAAGAAGGAGATATACATATG
>acnB
GAGGCAATTCCGACGTCTAAGAAACCATTATTATCATGACATTAACCTATAAAAATAGGCGTATCACGAGGCCCTTTCGTCTTCACCTC
GAGGTAGAGGCGGCACAAAGCCCGGCACTGCAAATTAGCGCCAGCACATGGGGTTTAAACGGCAAAGTCATTTTCATAATTCGGATCTC
AAGGAAATCGCAATGGTCGGCGAACTGCCACCCGCAGGTGCTGTGAATCCGAGTATAAAGAGGCGGTAGTTTAAATTTTGACTAATCTT
GGGATTCGTTGAGAAAGGTGATTATCACCATGCGAATTAACGAAGTTTTTACGGAGGGAAACAATCTCTAGACCATCCTTAACGATTCA
GCCACTTTTTTATGTTGCTTTTTTGTAAACAGATTAACACCTCGTCAAAATCCTGCTATTCTGCCCGTTGCGGTACTGGGCATTTACCC
TACAAACTGCTGTCTCACAGGAGCGTGAAGAGAATCGCCTGCCGCACTATGACAATGAGAGCGAGGAGAACCGTCGTGCTAGAAGAATA
CCGTAAGCACGTAGCTGAGCGTGCCGCTGAGGGGATTGCGCCCAAACCCCTGGATGCAAACCAAATGGCCGGATCCTCTAGATTTAAGA
AGGAGATATACATATG
>kch
GAGGCAATTCCGACGTCTAAGAAACCATTATTATCATGACATTAACCTATAAAAATAGGCGTATCACGAGGCCCTTTCGTCTTCACCTC
GATGGATCCCGTACGTAGCGGCAGGAGTCTATGAACACGTTTCAGTGAAACCATTTAAGAAAGTGTTCTGAATAGAGATTCAATATTTA
AGGGCACGGTTTTTGCAACCGTGTTCTTGTTTGGCCTTTCCTTATAGCCGGTCAGTGCATTCACCACCACAAGCAGATAATAACTTTAT
CTGTATTTCTTTATATGATGTGAGCATATTCAGGAAAAATGACCTTTCCTGAGGTACAGTTGCTCGCTCAAATCCTTTGATTACTGCGG
ATAAAGTCATCTACTATTGTTTAACACATTCACGCGCTGCATGTATGTTCAGAAAATGTGCACGCTAAGATAATTAAACAAAATTGTGG
TTACAGAGGTATCAGTGAGTCACTGGGCTACATTCAAACAAACAGCAACAAATTTATGGGTGACCCTTCGGCACGACATTCTCGCGCTG
GCCGTCTTTTTAACTCGAGAGATCCTCTAGATTTAAGAAGGAGATATACATATG
>b0259
GAGGCAATTCCGACGTCTAAGAAACCATTATTATCATGACATTAACCTATAAAAATAGGCGTATCACGAGGCCCTTTCGTCTTCACCTC
GATGGATCCCATTGTGCGCTTCAGTGGCGCATTTTGTTGTGTTGTTTGCATATAACCTTCTAAACGTATGGACATCTAAACATCCAAAG
AAGGTTTGTTATACCGCGTTTGTTAACAGAATGCAAAGTCAACCCTGCACCAGACGCAGATGCCCGCGCCGGGCGCTTAATAGCGGGTG
GGGATTAATAATCAGATACAAAAAGGGCCGCCAGCTCATAACGCCTCCGGAAGGTGCGAACAAGTCCCTGATATGAGATCATGTTTGTC
ATCTGGAGCCATAGAACAGGGTTCATCATGAGTCATCAACTTACCTTCGCCGACAGTGAATTCAGCAGTAAGCGCCGCTCGAGAGATCC
TCTAGATTTAAGAAGGAGATATACATATG
>racC
GAGGCAATTCCGACGTCTAAGAAACCATTATTATCATGACATTAACCTATAAAAATAGGCGTATCACGAGGCCCTTTCGTCTTCACCTC
GATGGATCCTCCTCGGTCGTGATGGTTTACCTGTAAAGGCGGAGTCATGGTGATGACTAAGAAAATAAAATGTGCTTACCACCTTTGCA
AAAAAGACGTTGAAGAAAGCAAAGCTATTGAAAGAATGCTTCACTTCATGCACGGGATTTTATCAAAAGACGAACCGAGAAAATATTGC
AGTGAAGCTTGTGCCGAAAAAGACCAGATGGCACATGAACTTTAATTAATTGACTATTCGAAACTGAATTTATGCCAGAAATGGCAGGT
ATTCGCTCAACCTTAATTAAGGAGAAAAACATGATTACCAATTATGAAGCCACTGTTGTAACTACCGATGACATTGTTCACGAGGTGAA
TCTGGAAGGAAAGCGCACTCGAGAGATCCTCTAGATTTAAGAAGGAGATATACATATG
>b0257
GAGGCAATTCCGACGTCTAAGAAACCATTATTATCATGACATTAACCTATAAAAATAGGCGTATCACGAGGCCCTTTCGTCTTCACCTC
GAGTCAGCTAAACAACAGACCGAGAAAGACACTGAAGTTCAAAACACCGAAAGAGATAATTGAAAGGGGTGTTGCATTGACAGATTGAA

TCTACAACCGCGCTCTTGATGTCAGACTCCCTGAACAGTTCTCGATAATCGGGAAACTCAGGGCGCGTTATCCTGTGGCCACTCTCTGC
CATGTGTTCGGGGTTCATCGCAGCAGCTACAAATACTGGAAAAACCGTCCTGAAAAGCCAGACGGCAGAGGATCCTCTAGATTTAAGAA
GGAGATATACATATG
>b0255
GAGGCAATTCCGACGTCTAAGAAACCATTATTATCATGACATTAACCTATAAAAATAGGCGTATCACGAGGCCCTTTCGTCTTCACCTC
GAGGCAAAAGCTCCGGTACGACCAGCCGCTTCAAAGGCGCGTAACAGATTTAGAGGTGCTTTTGCTAAGAGCTTCATTCATGAATCCTT
TTCACTAGGGATGAAGTGGTTTACTGAATTTGGCCACCTGAACAGAGGTGATATGCTCACCTCAGAACAACACAGGTGTCATAATGAAA
AAAAGAAATTTCAGCGCAGAGTTTAAACGCGAATCCGCTCAACTGGTCGGGATCCTCTAGATTTAAGAAGGAGATATACATATG
>yhcO
GAGGCAATTCCGACGTCTAAGAAACCATTATTATCATGACATTAACCTATAAAAATAGGCGTATCACGAGGCCCTTTCGTCTTCACCTC
GATGGATCCGGTGACGGAACCGGTAAATCGGCTGGCGGGGATGCTCCATAAGTATCAACATGCGTTGACCGATAGTTAAGTCAAAACCG
ACGCCAAAAGCGTCGGTTTTTTCATGGCTATACTTAGCGATGCACGGCAGAACTCGCCGCGAAACGTGACGGTGGCAACAGATGAATAT
TTATACCTTTGATTTTGATGAGATTGAGAGTCAGGAGGATTTTTATCGTGACTTTAGCCAAACCTTTGGTCTGGCGAACTCGAGAGATC
CTCTAGATTTAAGAAGGAGATATACATATG
>yhcM
GAGGCAATTCCGACGTCTAAGAAACCATTATTATCATGACATTAACCTATAAAAATAGGCGTATCACGAGGCCCTTTCGTCTTCACCTC
GATGGATCCATGCCGACGACTAACCCAATTAGCGCATATTCCCAGGTCATGAACATCTCCCGTTGTCTTTTGTTTCCGTAGGGTGTTGG
CTTCAGGCTCCATGCCTGCGGCTGATTATGCCACTATAACCGTTAATTCCACAGAAGTGGAATCCCGACTGCATATCGCGTAGTGTAGA
ACGGCCTTTTTTTCGTCAACCGTGAACAACGGCGCACCGATTATTCAAGGAATAACAATAAGATCATGCAAAGCGTTACCCCAACATCG
CAATACCTGAAGGCGCTTAATGAAGGCAGCCATCAACCCGACGACGTTCAAAACTCGAGAGATCCTCTAGATTTAAGAAGGAGATATAC
ATATG
>yhcB
GAGGCAATTCCGACGTCTAAGAAACCATTATTATCATGACATTAACCTATAAAAATAGGCGTATCACGAGGCCCTTTCGTCTTCACCTC
GAGTTTTGAACGTCGTCGGGTTGATGGCTGCCTTCATTAAGCGCCTTCAGGTATTGCGATGTTGGGGTAACGCTTTGCATGATCTTATT
GTTATTCCTTGAATAATCGGTGCGCCGTTGTTCACGGTTGACGAAAAAAAGGCCGTTCTACACTACGCGATATGCAGTCGGGATTCCAC
TTCTGTGGAATTAACGGTTATAGTGGCATAATCAGCCGCAGGCATGGAGCCTGAAGCCAACACCCTACGGAAACAAAAGACAACGGGAG
ATGTTCATGACCTGGGAATATGCGCTAATTGGGTTAGTCGTCGGCATCATTATTGGTGCTGTGGCCATGCGTTTTGGTAATCGGGATCC
TCTAGATTTAAGAAGGAGATATACATATG
>yhcA
GAGGCAATTCCGACGTCTAAGAAACCATTATTATCATGACATTAACCTATAAAAATAGGCGTATCACGAGGCCCTTTCGTCTTCACCTC
GAGCTGATTGATGCTGCGGTAAATGACAATACTATTCTTGGTTCTGATGAAACGATCAAACTGGATGGTAACGTCACTATCAGCGTTCA
GTATTTATAAGAACTTGTAATACAGTTTTTATTTTTACGCATTAACCAGTTAGCTATAACTCGGACAGATTATGCCCTATGACATGGTC
TGTCTTAATCAATCTCATCGCAAACAGTGCATAACATATATTAACAATATGTTTCTGCGTTATTTATCATTATTACAGGAGTTTTAATT
TAAATGCTCAGACACATTACATTCACTGTATTTATAACAACATCAATGAATACCTTAGCCACAGGGATGGTTCCTGGGATCCTCTAGAT
TTAAGAAGGAGATATACATATG
>yhjX
GAGGCAATTCCGACGTCTAAGAAACCATTATTATCATGACATTAACCTATAAAAATAGGCGTATCACGAGGCCCTTTCGTCTTCACCTC
GATGGATCCGGGGTTAGCGCCAGATTTTAACGTAACAGTCACAATTGAAACCATTAAATAACAATAGTTGTGGCGATAGTGGGTGCTAA
CTTACCAAATAATAAATTTGGTGAATAATTGTCGCGTCATTCATTCCTGAACTAAGGCATTTCATTCCGTTCTGATGGCATTTCATGCC
GTTTTTCCCCAGGCATAAAGTGCACTTCGTTATGGTTGTCGGCAGAGATTTTTCCTTTTTATTACTGCAGGAATACTGCCATGACACCT
TCAAATTATCAGCGTACCCGCTGGCTGACACTCATCGGTACTATCATTACCCAGTTTGCGCTGGGGTCGGTTTATACTCGAGAGATCCT
CTAGATTTAAGAAGGAGATATACATATG
>yeiM
GAGGCAATTCCGACGTCTAAGAAACCATTATTATCATGACATTAACCTATAAAAATAGGCGTATCACGAGGCCCTTTCGTCTTCACCTC
GATGGATCCGCGGTGACAGCCTGAAATCCAACATCCAGCTGGTGTTCAACAACGCCATTCTGGCGAGCGAAATTGCCAAAGAATATCAG
CGTCTCGCGGGTTAATTTCTGACGGGCAGAAAGCCGTCGCCCGTACTCATCAAGCATACCAAAACCTGGCGCTGACGCCGGGTTTACCG
GCATGAAGGGAAAATCATTATGGATATAATGAGAAGTGTTGTGGGGATGGTGGTGTTACTGGCAATAGCATTTCTGTTGTCAGTGAATA
AAAAGAGCATCAGTTTGCGCACGGTTCTCGAGAGATCCTCTAGATTTAAGAAGGAGATATACATATG
>yeiE
GAGGCAATTCCGACGTCTAAGAAACCATTATTATCATGACATTAACCTATAAAAATAGGCGTATCACGAGGCCCTTTCGTCTTCACCTC
GATGGATCCCCCCGGTGATAACTGCACTCAGGGCTAACCCCGGAATAAAATGCCACAGTGTACGATGTTGTTTCTGTAAGGTGATATTC
GTCATAACCCTCTCCTTTAACCGGGCATAAGGTTACGACTAACTGGTTTAAAAATAAAATTGATTATATATTTATAATTAATCTTTATA
AGTGGTAAGCGACTATGCACATCACCCTCCGGCAGTTGGAAGTTTTTGCAGAAGTATTGAAAAGTGGATCAACCACCCAGGCGTCGGTG
ATGCTGGCGTTGTCGCAATCAGCAGCTCGAGAGATCCTCTAGATTTAAGAAGGAGATATACATATG
>yeiG
GAGGCAATTCCGACGTCTAAGAAACCATTATTATCATGACATTAACCTATAAAAATAGGCGTATCACGAGGCCCTTTCGTCTTCACCTC
GAGGCAGCGGTGTTTCCAGTCCTCGCGCAACTAACGCTTCATGAACCAGGGCCGCTTCTTTACTGAGTGATGGCATTTATGATTTCTCC
TGCAGGTGTGACGCCTCCGCCCTGCGTGGGGGCAAAGTTATTAAGCTGATTTACAGCCTGATTATTGTGCGTGAGGCGGCGCACATAAT
CCAGTATTCACAGCGATAATTATTGTAATTGCCGCTGCCTTTCATCAGCAGATGTTAAAACATCGTTATGCAAATACGGAAGTGAAAGT
TACTCACAGCACATTGAATAAACGGTATGATGAAGAAATTGCAAACAACACAACAAGGAGCCACGCATGGAAATGCTCGAAGAGCACCG
CTGTTTTGAAGGCTGGCAGCAACGCTGGCGACACGACTCCAGTACCTTAAACTGCCCGATGGCGGGTGGCCATTGCTGCTCTGCTCTCA
GGAAGTTGTTTTAGCGCCGCCTCGAGAGATCCTCTAGATTTAAGAAGGAGATATACATATG
>sapA

GAGGCAATTCCGACGTCTAAGAAACCATTATTATCATGACATTAACCTATAAAAATAGGCGTATCACGAGGCCCTTTCGTCTTCACCTC
GATGGATCCTCAGCGAAGAAGCCCAACATGAAATGGCTGCTGAAGCGGGGATTAATCCCGTACGCATTGATGAGATCGCAGAGTTTTTG
AATCAATGGGGTAATGAGTAATGATTTAATTGCAGACGATCGGTCAGTATCGTTGTCCGGATAAAGCGTTCACGCCGCATCTGGCTAAC
AATGACACGCACTGCAGCGTGCTTTAATGGGTTTCTCACGATCGTACCGCTCTGCTCCTCGCTAAGTTGTTCTGTGAGTTATAGTTATA
GAGAAATCACGGGTACGAATGTTGAACAGATGGCGACGTCACCTGTTCAACGACCAACCACACGGCCCTCATTACGACCCCTGAACACA
AACGGTAATGGCAAATGTCATTAAATATTAGTAAGATAGCCTCTTTACATAAAATCCCTTAATATTATGCGCCAGGTATTATCGTCTCT
TTTGGTGATTGCTGGACTTGTGAGTGGTCAGGCAATCGCCGCGCCTGAATCTCCCCCGCATGCTGATATCTCGAGAGATCCTCTAGATT
TAAGAAGGAGATATACATATG
>yeiC
GAGGCAATTCCGACGTCTAAGAAACCATTATTATCATGACATTAACCTATAAAAATAGGCGTATCACGAGGCCCTTTCGTCTTCACCTC
GATGGATCCTATTGCCGGTACGCTGGTGGCGGGTTTGGCCTATGCCTTCCTGAAACGTCCGGAAGTGGACGCAGTAGCGAAAGCAGCGT
AATAAAAGGTGTGTTACAGGGCAGAAATTTCCTGTCCTGTAACATTTGATGAGCGCAAAGAGAATGACCGGTTTGTGAATGTAATTTTA
TGAATTTAAATGGTTTTATCTTTCAGTAGAAACTTCAAACGTTCATCAAACACTCAGGTTATTTCCTCAAGGGCAGATTATTTTCTGCC
CTTTTTTATTGTTCTGTTCTGTGCATCCCGTCACAAATTCCTCGCTTTTGTAATTCAGGCATTTAATCGCATTTTGTGAGCCTGCTCAA
ATCTGCCCATTTTGCCTCTCTGGCGGTAATTTACTTTACGTAAAACATGCATACTATGAGCACATGTTTAAATATGAACACATGTTTAA
ACCGTCTTTGCAGTATCTATATTCTTGTGGATTGACAGGGTTGGTAATTTTTTACCAGAGGACGTGCTATGCGCGAAAAGGATTATGTC
GTAATTATAGGTTCGGCGAATATTGATGTCGCCGGATATTCACATGAATCATTAAATTATGCGGATTCAAATCCAGGCTCGAGAGATCC
TCTAGATTTAAGAAGGAGATATACATATG
>nhaA
GAGGCAATTCCGACGTCTAAGAAACCATTATTATCATGACATTAACCTATAAAAATAGGCGTATCACGAGGCCCTTTCGTCTTCACCTC
GAGGATGACGATCAGGGCGACAATCATCGCCTTATGCTGCTTCATTGCTCTCTTCTCCTTGACCTTACGGTCAGTAAGAGGCACTCTAC
ATGTGTTCAGCATATAGGAGGCCTCGGGTTGATGGTAAAATATCACTCGGGGCTTTTCTCTATCTGCCGTTCAGCTAATGCCTGAGACA
GACAGCCTCAAGCACCCGCCGCTATTATATCGCTCTCTTTAACCCATTTTGTTTTATCGATTCTAATCCTGAAGACGCCTCGCATTTTT
GTGGCGTAATTTTTTAATGATTTAATTATTTAACTTTAATTTATCTCTTCATCGCAATTATTGACGACAAGCTGGATTATTTTTGAAAT
ATTGGCCTAACAAGCATCGCCGACTGACAACAAATTAATTATTACTTTTCCTAATTAATCCCTCAGGAATCCTCACCTTAAGCTATGAT
TATCTAGGCTTAGGGTCACTCGTGAGCGCTTACAGCCGTCAAAAACGCATCTCACCGCTGATGGCGCAAATTCTTCAATAGCTCGTAAA
AAACGAATTATTCCTACACTATAATCTGATTTTAACGATGATTCGTGCGGGGTAAAATAGTAAAAACGATCTATTCACCTGAAAGAGAA
ATAAAAAGTGAAACATCTGCATCGATTCTTTAGCAGTGATGCCTCGGGAGGCATTATTCTTATCATTGCCGCTATCCTGGCGATGATTA
TGGCCGGATCCTCTAGATTTAAGAAGGAGATATACATATG
>yeiT
GAGGCAATTCCGACGTCTAAGAAACCATTATTATCATGACATTAACCTATAAAAATAGGCGTATCACGAGGCCCTTTCGTCTTCACCTC
GAGCCCGGACTCTTTTTCGCTTATATGGTGGCATATATCGGCTTCGGAATTTTATCTATCGGCATGATTGTTTATCTTATTTTCTATCG
TTAATATTCCCCTCTCCAGTTAATTATTGAGAATAATTATTACTTCACCTGATAAGCTGCGGATATCACATTCCTAACCGCAGCTATTT
GTGAATCTTTTCACAGTTTAAATTCCCCCGCACGCTTAGCCTTAATATCAGTACATTATTATTTACTAAACGCTCGCCTTAATTACCTA
TAGCATTAAGGAAGATCACATGCCGCAACAAAATTATCTGGATGAACTCACTCCGGCTTTTACGTCTTTACTGGCGATTAAAGAAGCCT
CTCGCTGTTTATTATGTCACGACGCTCCCTGGGATCCTCTAGATTTAAGAAGGAGATATACATATG
>yeiP
GAGGCAATTCCGACGTCTAAGAAACCATTATTATCATGACATTAACCTATAAAAATAGGCGTATCACGAGGCCCTTTCGTCTTCACCTC
GAGGCTGTGTTCTGGTTTGCGATGGTGATGATTATCGCCACTCTGTTTTGCTTACTGCGGATTAAAGATGTTTAAGGCGCGGTCAGCAT
TTCCAGATCGATAAGCCAGGTCATGGCCTGCTGACGCGAGTTACCGCACATTTCAGCGCTGGCTTGTAAGCCTGCACAGACTTTCGGTC
GCAGTGGCGAGGTGAAAATTTTACAGCGTTGCTGTTCATCAAGCTGAATGCAGGGCGTATTGGCGGGCTTGCCATCTGGCATACCGGGA
ATAGGGCTGGAAATTGAAGGGGCAGTGCAAGGATCCTCTAGATTTAAGAAGGAGATATACATATG
>nfrB
GAGGCAATTCCGACGTCTAAGAAACCATTATTATCATGACATTAACCTATAAAAATAGGCGTATCACGAGGCCCTTTCGTCTTCACCTC
GATGGATCCGCATAAAGGCACGGTTGCGGTAACGTCGGATGCGCGGGGGACAAGGTTTGTTATCACATTACCCGCTTAATCCTTCAGCA
AACGGCAACTTTTATAACCAGTGTAAAAATAACGTGCCGCAATAGTCATACCTATTAATGGTAAAAAGCTGTCACAATTCATAAAAAAC
CTTAATATACGCCACCCTAAACATAACCAGCGTTAATGTAAGGTTTTTGTGTGGACTGGCTTCTTGATGTTTTTGCTACCTGGCTCTAC
GGCTTAAAAGTAATCGCGATAACGTTAGCGGTCATCATGTTCATCAGCGGGCTGCTCGAGAGATCCTCTAGATTTAAGAAGGAGATATA
CATATG
>folX
GAGGCAATTCCGACGTCTAAGAAACCATTATTATCATGACATTAACCTATAAAAATAGGCGTATCACGAGGCCCTTTCGTCTTCACCTC
GAGAAGAACTGGCATGAGCGGATCCGTTCGCGCCCTGCCACCGGGCAGGCACTGCTAAAAGCACAACTCGGTGATGAGCGTTCGGATAG
TTAACAGAAACAGGTTCTCGTGTATTATTTCATCCTAAGTAAAACAACGGAGAACCTGCAATGGCACAACCTGCCGCTATTGGATCCTC
TAGATTTAAGAAGGAGATATACATATG
>folE
GAGGCAATTCCGACGTCTAAGAAACCATTATTATCATGACATTAACCTATAAAAATAGGCGTATCACGAGGCCCTTTCGTCTTCACCTC
GATGGATCCAAACAGCGGTGCTCTTCGAGCATTTCCATGCGTGGCTCCTTGTTGTGTTGTTTGCAATTTCTTCATCATACCGTTTATTC
AATGTGCTGTGAGTAACTTTCACTTCCGTATTTGCATAACGATGTTTTAACATCTGCTGATGAAAGGCAGCGGCAATTACAATAATTAT
CGCTGTGAATACTGGATTATGTGCGCCGCCTCACGCACAATAATCAGGCTGTAAATCAGCTTAATAACTTTGCCCCCACGCAGGGCGGA
GGCGTCACACCTGCAGGAGAAATCATAAATGCCATCACTCAGTAAAGAAGCGGCCCTGGTTCATGAAGCGTTAGTTGCGCGAGGACTGG
AAACACCGCTGCCTCGAGAGATCCTCTAGATTTAAGAAGGAGATATACATATG
>folD

GAGGCAATTCCGACGTCTAAGAAACCATTATTATCATGACATTAACCTATAAAAATAGGCGTATCACGAGGCCCTTTCGTCTTCACCTC
GATGGATCCTAACTACCCGTAGCGGCGAATAGTGCGGCAGCCAGCGCAGACGAAATAAATCTTAATTTCATATATATTCCTTCAATCTC
ATTTATCGACTCCACATCCGTATATAACCGATTACTTTATTTAAGACACTGATAGTAGTAAATTCCTTTTTATCCTCTAAGAATGTCTT
AATTGAAAATATGCACTCTATTCTAAAAAATAGAGAGCCCCGTTAGATGAATACTTCCGCGCAAAATATATTCAACACAAATATAGACC
TGAAGCGGTAAATTACCAGGCTGAAAATTCTTTTTATATTGTCAGGTATTTCTTAAATTATCTTAATCCTTAGACAAGGAAATAAATCA
GTTCCAGATTTACAACGCCATCATGGACGAAAAATGAAGCTTTCAGTCTCAGCGACGGTGCGCCTCACCTTCGCAAGAGGTCGCTTCAC
GCGATAAATCTGAAACGAAACCTGACAGCGCGCCCCGCTTCTGACAAAATAGGCGCATCCCCTTCGATCTACGTAACAGATGGAATCCT
CTCTCTGATGGCAGCAAAGATTATTGACGGTAAAACGATTGCGCAGCAGGTGCGCTCTGAAGTTGCTCAAAAAGTTCAGGCGCGTCTCG
AGAGATCCTCTAGATTTAAGAAGGAGATATACATATG
>folA
GAGGCAATTCCGACGTCTAAGAAACCATTATTATCATGACATTAACCTATAAAAATAGGCGTATCACGAGGCCCTTTCGTCTTCACCTC
GAGGCAGGGAACCGAAGAAGGTAAACATACCGGCAACATGGCGGATGAACCGGAAACGAAACCCTCATCCTAATAAAGAGTGACGTAAA
TCACACTTTACAGCTAACTGTTTGTTTTTGTTTCATTGTAATGCGGCGAGTCCAGGGAGAGAGCGTGGACTCGCCAGCAGAATATAAAA
TTTTCCTCAACATCATCCTCGCACCAGTCGACGACGGTTTACGCTTTACGTATAGTGGCGACAATTTTTTTTATCGGGAAATCTCAATG
ATCAGTCTGATTGCGGCGTTAGCGGTAGATCGCGTTATCGGCATGGAAAACGCCATGCCGTGGAACCTGCCTGCCGATCTCGCCTGGTT
TAGGATCCTCTAGATTTAAGAAGGAGATATACATATG
>folC
GAGGCAATTCCGACGTCTAAGAAACCATTATTATCATGACATTAACCTATAAAAATAGGCGTATCACGAGGCCCTTTCGTCTTCACCTC
GATGGATCCGCGTGAAGGCGTAGTGGTACCCCCGGTACCGGATCAGGAACCTGAGGCCTGATAACTGATAAGGGCAGGGCCACTGGCTC
TGCCCTTTTGCTATTCTCACCGTAACGAATCAGCGGATACCATGATTATCAAACGCACTCCTCAAGCCGCGTCGCCTCTGGCTTCGTGG
CTTTCTTATCTGGAAAACCTGCACAGTAAAACTATCGATCTCGGCCTTGAGCTCGAGAGATCCTCTAGATTTAAGAAGGAGATATACAT
ATG
>folB
GAGGCAATTCCGACGTCTAAGAAACCATTATTATCATGACATTAACCTATAAAAATAGGCGTATCACGAGGCCCTTTCGTCTTCACCTC
GATGGATCCGGCAAACCAGAATGGCACTGGAAATGGAGCCGCAGAGGTACGCGATGAGGATCATTCCAGGCGCGATTGCACTCATAAGC
TGTTCCGTTTTGAAAATTCGTGTTCAACGATGAATCTGTGGATAATACGCACATTTCGCCGGAAGTGGTATCCGGTTAGCCAAAAAGCA
GGCAGGACGTGATGGATATTGTATTTATAGAGCAACTTTCGGTAATCACCACTATTGGTGTTTACGACTGGGAACAGACCATCGAACTC
GAGAGATCCTCTAGATTTAAGAAGGAGATATACATATG
>hydN
GAGGCAATTCCGACGTCTAAGAAACCATTATTATCATGACATTAACCTATAAAAATAGGCGTATCACGAGGCCCTTTCGTCTTCACCTC
GATGGATCCTTTCTCACCGCCGAAAACCTTCAGCGGAAAACTGATCCGCCGCGACTCACTCATTGCTCCTTCGCGATAAACATTGTCAT
CACATGTCATCGTCACTAGTGTCGATGACATCGTTTTTCCTGCATTACAGAAATTAATTTCCTTAAATAACAGTAAATTAAAAACTGGC
ATGATTTGTGAATGTATCGGCGCATTAACTGTCATTGCTGGAGAATTTGATGAACCGTTTCATCATTGCTGACGCGAGTAAATGTATTG
GTTGCCGTACCTGTGAAGTAGCGTGCGTGGTTTCTCATCTCGAGAGATCCTCTAGATTTAAGAAGGAGATATACATATG
>rpoS
GAGGCAATTCCGACGTCTAAGAAACCATTATTATCATGACATTAACCTATAAAAATAGGCGTATCACGAGGCCCTTTCGTCTTCACCTC
GATGGATCCCATGGGTAGCACCGGAACCAGTTCAACACGCTTGCATTTTGAAATTCGTTACAAGGGGAAATCCGTAAACCCGCTGCGTT
ATTTGCCGCAGCGATAAATCGGCGGAACCAGGCTTTTGCTTGAATGTTCCGTCAAGGGATCACGGGTAGGAGCCACCTTATGAGTCAGA
ATACGCTGAAAGTTCATGATTTAAATGAAGATGCGGAATTTGATGAGAACGGAGTTGAGGTTTTTGACGAAAAGGCCCTCGAGAGATCC
TCTAGATTTAAGAAGGAGATATACATATG
>kdtA
GAGGCAATTCCGACGTCTAAGAAACCATTATTATCATGACATTAACCTATAAAAATAGGCGTATCACGAGGCCCTTTCGTCTTCACCTC
GAGGCATATCGATTTTTGCATCAGGATAATTCTGCTTGAGCGTACTGATGACAGGAGTAGTTAATAACATATCCCCATGATATCGCATC
TTTATGACCAGGATTTTTCGAAATGGCTTTTCCACTAGCGACTCTTTTGTGTGATTGTCTGGTTAAGTTAAGCAGAAAAAAGCACGCTA
CCGCCCCAGGCTCAACAGCTACCTGAATACTGATAACGCACTTACTTTGACGTGCTAGTGTATCATTTCTTTCGCAGCAATCCGACCCG
AATGATTTTTATACACAAAATATACTTTAATCATAAAAATCAAATAGATAAAAAATAGCATTTCGATAATGTCAGTACGGCCTATGTAG
ATTTGAAAATGGGGCCGAGTTTCAATGAATCGCATAAAAATACAAAAAAAATGAAGATGTAGCGGCAGATCAGACCAAACTGAAGCTAT
TTAAGTCAAAAACGGGAAAAGTAATGGTAAAGCCACAGCTAAATACATAGAATCCCCAGCACATCCATAAGTCAGCTATTTACTATGCT
CGAATTGCTTTACACCGCCCTTCTCTACCTTATTCAGCCGCTGATCTGGATACGGCTCTGGGTGCGCGGACGTAAGGCTCCGGCCTATG
GATCCTCTAGATTTAAGAAGGAGATATACATATG
>rpoE
GAGGCAATTCCGACGTCTAAGAAACCATTATTATCATGACATTAACCTATAAAAATAGGCGTATCACGAGGCCCTTTCGTCTTCACCTC
GATGGATCCCCAGTGAAAGTCCGGCTGCGCCGCTACCGATAATCAACACGTCACATGAATGTTCAGGGAGAGTATTCATTTTCTTTGTT
TAATTTACTAAACATGGTTTGGTCAGCATAGCATCATGTTGTGCGGATAAACACCTGCTATTTTAATATTTGTTACAGTTGCTAAACAC
GCTGACGCAGGGCGGCGAGAAAAAGAGAAGTTACTGGCTGGTGGAGGATTAGGTGGTGAAATAAAAAGGCCGTTGGGTTACTCTTCAGG
CAGTTAAATGGGCATTTCTACACAGATAATGCGATGTTCAGATTCTGTAGACTTATAATGATAGATAATGATCCGTCTACAGCATGACA
AACAAAAACAGATGCGTTACGGAACTTTACAAAAACGAGACACTCTAACCCTTTGCTTGCTCAAATTGCAGCTAATGGAGTGGCGTTTC
GATAGCGCGTGGAAATTTGGTTTGGGGAGACTTTACCTCGGATGAGCGAGCAGTTAACGGACCAGGTCCTGGTTGAACGGGTCCAGAAG
GGAGATCAGAAAGCCTTTAACTTACTGGTAGTGCGCTATCAGCATAAAGTGGCCTCGAGAGATCCTCTAGATTTAAGAAGGAGATATAC
ATATG
>rpoD
GAGGCAATTCCGACGTCTAAGAAACCATTATTATCATGACATTAACCTATAAAAATAGGCGTATCACGAGGCCCTTTCGTCTTCACCTC
GAGAGCGCACGCATGGTTTAAGCAACGAAGAACGCCTGGAGCTCTGGACATTAAACCAGGAGCTGGCGAAAAAGTGATTTAACGGCTTA

Data Set S4 - MG1655 library reference file.fasta
AGTGCCGAAGAGCGATCGGGAAGCCCCCGACAGCCGCACTGAGAGGCAGCGGCAAATATATAAGTACGCCCTCGTAATTATCGTTGGCG
GTAAACAACCGTTGGATTTCAGCGTTAACGGCTGAAGGACATCGGGTCAATCGCCCAACACCAACCTCATGAAATAAGTGTGGATACCG
TCTTATGGAGCAAAACCCGCAGTCACAGCTGAAACTTCTTGTCACCCGTGGTAAGGAGCAAGGCTATCTGACCTATGCCGAGGTCAATG
ACCATCTGCCGGAAGGATCCTCTAGATTTAAGAAGGAGATATACATATG
>rpoH
GAGGCAATTCCGACGTCTAAGAAACCATTATTATCATGACATTAACCTATAAAAATAGGCGTATCACGAGGCCCTTTCGTCTTCACCTC
GATGGATCCGGCTTGCCACGGTACAACATTTACGCCACTTTACGCCTGAATAATAAAAGCGTGTTATACTCTTTCCCTGCAATGGGTTC
CGTAGCAGGGAAAGAGACCCCGTTGTCTCTTCCCGGTATTTCATCTCTATGTCACATTTTGTGCGTAATTTATTCACAAGCTTGCATTG
AACTTGTGGATAAAATCACGGTCTGATAAAACAGTGAATGATAACCTCGTTGCTCTTAAGCTCTGGCACAGTTGTTGCTACCACTGAAG
CGCCAGAAGATATCGATTGAGAGGATTTGAATGACTGACAAAATGCAAAGTTTAGCTTTAGCCCCAGTTGGCAACCTGGATTCCTACAT
CCGGGCAGCTACTCGAGAGATCCTCTAGATTTAAGAAGGAGATATACATATG
>yjaB
GAGGCAATTCCGACGTCTAAGAAACCATTATTATCATGACATTAACCTATAAAAATAGGCGTATCACGAGGCCCTTTCGTCTTCACCTC
GATGGATCCTGACCAGACGCACGAGAAGTTGTCATCACAAAGACGTTTTCTTCACGCAAGAAATTGACGGCGGGTAGCTCGTCCGGCAC
ACGAATCGGCATAACCTGATTACCTCACTACATACGCTTATACGTTTAGACATCCAGATAGCTGAAGATAACCAGAAAATTTGCCAATG
TCGAGCCTGCATGTTGAAGGTGAGAAAGTTTCAAGGAAAAGACTACGTTAGAATATAAGAATGACTAAAAGGAGAGAAAATGGTTATTA
GTATTCGCCGCTCACGGCATGAGGAAGGGGAGGAACTCGTTGCGATTTGGTGTCGTTCTGTCGATGCCACTCACGCTCGAGAGATCCTC
TAGATTTAAGAAGGAGATATACATATG
>yjaH
GAGGCAATTCCGACGTCTAAGAAACCATTATTATCATGACATTAACCTATAAAAATAGGCGTATCACGAGGCCCTTTCGTCTTCACCTC
GAGCCGCAGACCGGTAAAGAAATCAAAATTGCCGCAGCTAACGTACCGGCATTTGTTTCTGGCAAGGCACTGAAAGACGCAGTTAAGTA
AGATTGCGTGGCAGTGAACAGTTTTAACGAAGGGGTGGTTTCACCCCTTTTGTCTTTCTGGCGTCGATCATTGAGGATCCTCTAGATTT
AAGAAGGAGATATACATATG
>nohB
GAGGCAATTCCGACGTCTAAGAAACCATTATTATCATGACATTAACCTATAAAAATAGGCGTATCACGAGGCCCTTTCGTCTTCACCTC
GAGTCTGGAAGCATTCAGGGCAATTGAGGCAGCGTTGGTGAAGCACGATAATAATATGAAGGATTATTCCCTGGTGGTTGACTGATCAC
CATAACTGCTAATCATTCAAACTACTTAACCTGTGACAGAGCCAACACGCAGTCTGTCACTGTCAGGAAAGTGGTAAAACTGCAACTCA
ATTACTGCAATGCCCTCGTAATTAAGTGAATTTACAATATCGTCCTGTTCGGAGGGAAGAACGCGGGATGTTCATTCTTCATCACTTTT
AATTGATGTATATGCTCTCTTTTCTGACGTTAGCCTCCGACGGCAGGCTTCAATGACCCAGGCTGAGAAATTCCCGGACCCTTTTTGCT
CAAGAGCGATGTTAATTTGTTCAATCATTTGGTTAGGAAAGCGGATGTTGCGGGTTGTTGTTCTGCGGGTTCTGTTCTTAGTTGACATG
AGGTTGCCCCGTATTCAGTGTCGCTGATTTGTATTGTCTGAAGTTGTTTTTACGTTAAGTTGATGCAGATCAATTAATACGATACCTGC
GTCATAATTGATTATTTGACGTGGTTTGATGGCGTAGATGCACGTTGTGACATGTAGATGATAATTATTATCATTTTGTGGGTCCTTTC
CGGCGATCCGACAGGTTACGGGGCGGCGACCTCGCGGGTTTTCGCTATTTATGAAAATTTTCCGGTTTAAGGTGTTTCCGTTCTTCTTC
GTCGTAACTTAATGTATTTATTTAAAATACCCCCTGAAAAGAAAGGAAACGACAGGTGCTGAAAGCGAGCTTTTTGGCCTCTGTCGTTT
CCTTTCTCTGTTTTTGTCCGTGGAATGTGCAATGGAAGTCAACAAAAAGCAGCTGGCTGACATTTTCGGTGCGGATCCTCTAGATTTAA
GAAGGAGATATACATATG
>rpsG
GAGGCAATTCCGACGTCTAAGAAACCATTATTATCATGACATTAACCTATAAAAATAGGCGTATCACGAGGCCCTTTCGTCTTCACCTC
GATGGATCCGACTGCTCCGGCGTTAAAGACCGTAAGCAGGCTCGTTCCAAGTATGGCGTGAAGCGTCCTAAGGCTTAATGGTTCTCCGT
TAAGTAAGGCCAAACGTTTTAACTTAAATGTCAAACTAAACTCGTAGAGTTTTGGACAATCCTGAATTAACAACGGAGTATTTCCATGC
CACGTCGTCGCGTCATTGGTCAGCGTAAAATTCTGCCGGATCCGAAGTTCGGATCAGAACTGCTGGCTAACTCGAGAGATCCTCTAGAT
TTAAGAAGGAGATATACATATG
>rpsB
GAGGCAATTCCGACGTCTAAGAAACCATTATTATCATGACATTAACCTATAAAAATAGGCGTATCACGAGGCCCTTTCGTCTTCACCTC
GAGATCTCCAGCACTTCGGCAGCCAGTCGGCCAGCGACGCGCATTTTTTCGATATCTTCTGGGGTCTTGATTGAGATAGCCATTAATTC
TGTCCATCAGCGTCGGTGATACCGACAATATATATGTAAGTGCCGTCAATGGTATCACACCCGGGCAAATTGAGAATCATTCTGAATTT
CGCCAAACGTGCCACTGAAGGTTTTCTATAATAGAAAATTCGACGTCTGATGCTGTACACAGCGCCAACAATTATTGGTGTCCACGACG
TATTTGTGGTATAAAGCGCGCCGGACTTCCGATCCATTTCGTATACACAGACTGGACGGAAGCGACAATCTCACTTTGTGTAACAACAC
ACACGTATCGGCACATATTCCGGGGTGCCCTTTGGGGTCGGTAATATGGGATACGTGGAGGCATAACCCCAACTTTTATATAGAGGTTT
TAATCATGGCAACTGTTTCCATGCGCGACATGCTCAAGGCTGGTGTTCACTTCGGTCACCAGACCCGTTACTGGAACCCGAAAATGAAG
CCGTTCATCTTCGGTGGGATCCTCTAGATTTAAGAAGGAGATATACATATG
>rpsA
GAGGCAATTCCGACGTCTAAGAAACCATTATTATCATGACATTAACCTATAAAAATAGGCGTATCACGAGGCCCTTTCGTCTTCACCTC
GAGGGCAGCCGATGCTTTAGTGTTGGATTCCACCACCTTAAGCATTGAGCAAGTGATTGAAAAAGCGCTACAATACGCGCGCCAGAAAT
TGGCTCTCGCATAAGCGACCGAATTTGCAGTACCCCCGTTGCAATGGAATGACAGCGGGTATGTTAAACAACCCCATCCGGCATGGAGC
CAGGTGGACGTTAAATATAAACCTGAAGATTAAACATGACTGAATCTTTTGCTCAACTCTTTGAAGAGTCCTTAAAAGAAATCGAAACC
CGCCCGGGTTCTATCGTTCGTGGCGTTGTTGGATCCTCTAGATTTAAGAAGGAGATATACATATG
>fdnG
GAGGCAATTCCGACGTCTAAGAAACCATTATTATCATGACATTAACCTATAAAAATAGGCGTATCACGAGGCCCTTTCGTCTTCACCTC
GAGGACACCGCGAATCAATCCTACCATCGTGCTCCACAGGACGATCGCTATCAGCCCTATGAGCGTTGCTTTTTGTCGTGTCATGCTCG
CTGTTTTGTCTCTCTTGCCGTTAAAAATTAAGCTGAATTTTATAGCATTTTTTTAACTGGCCTGTCAGGCAGTGGTGCGTTTTTCTACC
GCTATTGAGGTAGGTCAATTTGCGAAGGCGGATTATTTTGTGGCAAACAGATGTTCTTTTTGATTTCGCGCAAAAAGATTCAGAATTTT

ACTGTTAGTTTCCTCGCGCAGTAATACCCCTGAAAAAAGAGGAAAGCAATGGACGTCAGTCGCAGACAATTTTTTAAAATCTGCGCGGG
CGGTATGGCTGGAACAACAGTAGCGGCATTGGGGGATCCTCTAGATTTAAGAAGGAGATATACATATG
>rpsO
GAGGCAATTCCGACGTCTAAGAAACCATTATTATCATGACATTAACCTATAAAAATAGGCGTATCACGAGGCCCTTTCGTCTTCACCTC
GATGGATCCCACGGAAGGTGAGAACGGCAAATTTATCGGTATGGGCGAAATTGACGATGAAGGCCGCGTTGCGCCTCGTCGCCTGGTGG
TTGAATACCCGGCGTAATGTTAACCGTCTTGCGATAACAGGTCGCTACGAGTAGAATACTGCCGCTTAACGTCGCGTAAATTGTTTAAC
ACTTTGCGTAACGTACACTGGGATCGCTGAATTAGAGATCGGCGTCCTTTCATTCTATATACTTTGGAGTTTTAAAATGTCTCTAAGTA
CTGAAGCAACAGCTAAAATCGTTTCTGAGTTTGGTCGTGACGCAAACGACACCGGTTCTCTCGAGAGATCCTCTAGATTTAAGAAGGAG
ATATACATATG
>rpsM
GAGGCAATTCCGACGTCTAAGAAACCATTATTATCATGACATTAACCTATAAAAATAGGCGTATCACGAGGCCCTTTCGTCTTCACCTC
GATGGATCCGTGTGATTTGCAGTGCCGAGCCGAAGCATAAACAGCGCCAAGGCTGATTTTTTCGCATATTTTTCTTGCAAAGTTGGGTT
GAGCTGGCTAGATTAGCCAGCCAATCTTTTGTATGTCTGTGCGTTTCCATTTGAGTATCCTGAAAACGGGCTTTTCAGCATGGAACGTA
CATATTAAATAGTAGGAGTGCATAGTGGCCCGTATAGCAGGCATTAACATTCCTGATCATAAGCATGCCGTAATCGCATTAACTTCGAT
TTATGGCGTCGGCAAGACCCGTTCTAACTCGAGAGATCCTCTAGATTTAAGAAGGAGATATACATATG
>rpsL
GAGGCAATTCCGACGTCTAAGAAACCATTATTATCATGACATTAACCTATAAAAATAGGCGTATCACGAGGCCCTTTCGTCTTCACCTC
GATGGATCCTTAAGCACCCCAGCCAGATGGCCTGGTGATGGCGGGATCGTTGTATATTTCTTGACACCTTTTCGGCATCGCCCTAAAAT
TCGGCGTCCTCATATTGTGTGAGGACGTTTTATTACGTGTTTACGAAGCAAAAGCTAAAACCAGGAGCTATTTAATGGCAACAGTTAAC
CAGCTGGTACGCAAACCACGTGCTCGCAAAGTTGCGAAAAGCAACGTGCCTGCGCTGGAAGCATGCCCGCAAAAACGTGGCGTATCTCG
AGAGATCCTCTAGATTTAAGAAGGAGATATACATATG
>rpsJ
GAGGCAATTCCGACGTCTAAGAAACCATTATTATCATGACATTAACCTATAAAAATAGGCGTATCACGAGGCCCTTTCGTCTTCACCTC
GATGGATCCAGCGAAGAGAAAGCCGGTTTAAGAGAGCGTGTCAAAAATGCACTGAACGAACTTGAGAGATAACCCGAAGGCTGTTTACT
TTACTAGCAATACGCTTGCGTTCGGTGGTTAAGTATGTATAATGCGCGGGCTTGTCGTAGTTGACAGCAGGTTCAATCTGAACCTCAGT
TAGCCGAATTTGGCTACCTAACAATGCTCCCAATCGGGGAGCTACGTAAGAACGGTTACACTCTCCCATCAATCGTAATGGGTCTGAGG
AGTAATCATTTTCGTTTATAAAATAATTGGAGCTCTGGTCTCATGCAGAACCAAAGAATCCGTATCCGCCTGAAAGCGTTTGATCATCG
TCTGATCGATCAAGCAACCGCGGAAATCGTCGAGACTGCCAAGCGCACTGTCGAGAGATCCTCTAGATTTAAGAAGGAGATATACATAT
G
>apt
GAGGCAATTCCGACGTCTAAGAAACCATTATTATCATGACATTAACCTATAAAAATAGGCGTATCACGAGGCCCTTTCGTCTTCACCTC
GAGTGCTGGTAATTCTCGCCTGTTTGCTTTTCTATATGTGGCGAATTCCGGTGATTGATGAAAAGCAAGAAAAGCACTGAAGCACAACA
ATCGCAGTTGCAATTATTGCGTACAGCCAGTACATTCTGGCGTTTTCGAGCACAGGCGCAGGCGGTCAAAGGTTAAACAACTGTTACTT
TTGATACGTTTAAAACGCGCCGTGAGTACCACCGTAACAAGCAGGCATACACTTATGACCGCGACTGCACAGCAGCTTGAGTATCTCAA
AAATAGCATCAAAAGCATTCAGGACTACCCAAAACCCGGCATTCTTTTCCGGATCCTCTAGATTTAAGAAGGAGATATACATATG
>rpsU
GAGGCAATTCCGACGTCTAAGAAACCATTATTATCATGACATTAACCTATAAAAATAGGCGTATCACGAGGCCCTTTCGTCTTCACCTC
GAGCAAACCTTTTTCATCGTCGTAAATGGCGATGCCGGTTTCATCGCAGGAAGTTTCAATACCCAGTACACGCATGACTTGTTTTACCT
CGCTTTATTACCGCGCAGTGTAGGACCAATGCGGGTTGATGTAAAACTTTGTTCGCCCCTGGAGAAAGCCTCGTGTATACTCCTCACCC
TTATAAAAGTCCCTTTCAAAAAAGGCCGCGGTGCTTTACAAAGCAGCAGCAATTGCAGTAAAATTCCGCACCATTTTGAAATAAGCTGG
CGTTGATGCCAGCGGCAAACCGAATTAATCAAAGGTGAGAGGCACATGCCGGTAATTAAAGTACGTGAAAACGAGCCGTTCGACGTAGC
TCTGCGTCGCTTCAAGCGTTCCTGCGAAAAAGCAGGTGTTCTGGCGGGATCCTCTAGATTTAAGAAGGAGATATACATATG
>rpsT
GAGGCAATTCCGACGTCTAAGAAACCATTATTATCATGACATTAACCTATAAAAATAGGCGTATCACGAGGCCCTTTCGTCTTCACCTC
GATGGATCCGCTGTATTCACACCCCGCAAGCTGGTAGAATCCTGCGCCATCACTACGTAACGAGTGCCGGCACATTAACGGCGCTTATT
TGCACAAATCCATTGACAAAAGAAGGCTAAAAGGGCATATTCCTCGGCCTTTGAATTGTCCATATAGAACACATTTGGGAGTTGGACCT
TGGCTAATATCAAATCAGCTAAGAAGCGCGCCATTCAGTCTGAAAAGGCTCGTAAGCACAACGCAAGCCGTCGCTCTATGATGCCTCGA
GAGATCCTCTAGATTTAAGAAGGAGATATACATATG
>rpsP
GAGGCAATTCCGACGTCTAAGAAACCATTATTATCATGACATTAACCTATAAAAATAGGCGTATCACGAGGCCCTTTCGTCTTCACCTC
GATGGATCCGAAGGGTATGATGCCCCCAGGCTTCCCTGGTCGCTAATCGGCCTGGCTTGTTTGCCATTTTGGCACCGGGATGTGCTCGT
CATCCTCACGTACTGCGTGTACGCTCCGGTGACTGCGCGCACTCCGGCACCAAACTGGCTGCACCGCCGACGGCCTTTTACAGCAGAGT
TAGCAACTGTTGATTGCAATTCCCGCAAAAATGAGTAAAATTTTCGGGCTTTTAATATGACACCGGACTCCGTTCCTCGATGGGGTCCG
GTTGTTTTATTCACACAAGAGGATGTTATGGTAACTATTCGTTTAGCACGTCACGGCGCTAAAAAGCGTCCGTTCTACCAGGTTGTTGT
CGCTGACAGCCCTCGAGAGATCCTCTAGATTTAAGAAGGAGATATACATATG
>deoA
GAGGCAATTCCGACGTCTAAGAAACCATTATTATCATGACATTAACCTATAAAAATAGGCGTATCACGAGGCCCTTTCGTCTTCACCTC
GAGGCGTCACTACCGCTTTGGCGCTTCCAGCCTGCTGGCAAGCCTGCTGAAAGCGCTGGGTCACGGCGACGGTAAGAGCGCCAGCAGCT
ACTAAGTAAGATGCTTTACGCCTGATGCGCTGCGCTTATCAGGCCTACGAGACGTATCTACCCGTAGGCCGGATAAGGCGTAGACGCAT
CCGGCAAAAGCCGCCTCATACTCTTTTCCTCGGGAGGTTACCTTGTTTCTCGCACAAGAAATTATTCGTAAAAAACGTGATGGTCATGC
GCTGAGCGATGAAGAAATTCGTTTCTTTATCAACGGTATTCGCGACAGGATCCTCTAGATTTAAGAAGGAGATATACATATG
>deoB

GAGGCAATTCCGACGTCTAAGAAACCATTATTATCATGACATTAACCTATAAAAATAGGCGTATCACGAGGCCCTTTCGTCTTCACCTC
GAGCGGTGAAAGCGGCAATTAAACTTGCCGATAAAGCACCGGAAAGCACACCAACTGTCTATCGCCGTATCAGCGAATAACGGTATACT
GATCTGATCATTTAAATTTGAAGCACTGAGTACGGAGAACATATGAAACGTGCATTTATTATGGTGCTGGACTCATTCGGCATCGGCGC
TACAGAAGATGCAGAACGCTTTGGTGACGTCGGGGCTGACACCCTGGGTCATGGATCCTCTAGATTTAAGAAGGAGATATACATATG
>deoC
GAGGCAATTCCGACGTCTAAGAAACCATTATTATCATGACATTAACCTATAAAAATAGGCGTATCACGAGGCCCTTTCGTCTTCACCTC
GAGCCAGTGCCAGAAAATGGCGCTTCTGCTCCGGGCTAAGCACTGGGCTGGTGACAATTTGCTGGCAACGTTGTTGCAGTGCATTTTCA
TGAGAAGTGGGCATCTTCTTTTCCTTTTATGCCGAAGGTGATGCGCCATTGTAAGAAGTTTCGTGATGTTCACTTTGATCCTGATGCGT
TTGCCACCACTGACGCATTCATTTGAAAGTGAATTATTTGAACCAGATCGCATTACAGTGATGCAAACTTGTAAGTAGATTTCCTTAAT
TGTGATGTGTATCGAAGTGTGTTGCGGAGTAGATGTTAGAATACTAACAAACTCGCAAGGTGAATTTTATTGGCGACAAGCCAGGAGAA
TGAAATGACTGATCTGAAAGCAAGCAGCCTGCGTGCACTGAAATTGATGGACCTGACCACCCTGAATGACGACGACACCGGGATCCTCT
AGATTTAAGAAGGAGATATACATATG
>deoD
GAGGCAATTCCGACGTCTAAGAAACCATTATTATCATGACATTAACCTATAAAAATAGGCGTATCACGAGGCCCTTTCGTCTTCACCTC
GAGGGCTCACTGGGTCATCGTGAAACCTTCGCGGATATCGGCCAGACTCTGGCAAAATATTTGGTACTTCTGATATGGAATATGGCAA
AGCCATGTTCTGATGGATTTGGGCGGAGCGTTGACTCCGCCTTTGTTATGTCACAAAAAGGATAAAACAATGGCTACCCCACACATTAA
TGCAGAAATGGGCGATTTCGCTGACGTAGTTTTGATGCCAGGCGACCGGATCCTCTAGATTTAAGAAGGAGATATACATATG
>usg
GAGGCAATTCCGACGTCTAAGAAACCATTATTATCATGACATTAACCTATAAAAATAGGCGTATCACGAGGCCCTTTCGTCTTCACCTC
GATGGATCCTTGTGATGACGCCAGTGCGGCATCATTGCTGTGTAAACTGGGTTTTAACGCCGTTCATCATCCGGCACGTTAATCTCTTC
TTCATGCTCTCTGCTGTAACATTGGCAGGGAGCTTTGCTATTTCTGGAGTAAACCACCATGTCTGAAGGCTGGAACATTGCCGTCCTGG
GCGCAACTGGCGCTGTGGGCGAAGCCCTGCTTGAAACGCTGGCTGAACGTCCTCGAGAGATCCTCTAGATTTAAGAAGGAGATATACAT
ATG
>sirA
GAGGCAATTCCGACGTCTAAGAAACCATTATTATCATGACATTAACCTATAAAAATAGGCGTATCACGAGGCCCTTTCGTCTTCACCTC
GATGGATCCAGCGTTGAGTTTGCGAGAAAACGTTCATATTGTACCTTTTTGATTAACCATTGGGGTGAGGGAACCCAATACGTACGACA
CGTCTGTTACCTTAACATAATAACGTGCTTACCCTTTTTTCGAGCCGCCGCATGATACTGCTTTACGTTGACAATGCAATGGTTAATTT
TCACGCAATCGTTAACCTAGTTTGCTTACGGACTCACAGGGCGTAAACTAGCGCCGTTTTTTTAAGTGATGAGAAGAAAATGACCGATC
TCTTTTCCAGCCCTGACCACACACTCGACGCGCTTGGCCTGCGCTGCCCGGAACCGGTGATGATGGTGCGCAAAACCCTCGAGAGATCC
TCTAGATTTAAGAAGGAGATATACATATG
>fes
GAGGCAATTCCGACGTCTAAGAAACCATTATTATCATGACATTAACCTATAAAAATAGGCGTATCACGAGGCCCTTTCGTCTTCACCTC
GAGGGTCGGCTCTTGTGCCTGCGCTACCCCATAAATCCCCAGATTGACCAACAAGGCCAGGGAATGAATCTTCTTGTTCATTGTTTTAT
TCCTGCATTTTTGCCACGAATTGCAACTGTCGGGCATGGTCGTCATCAACACGACGCATCCCGCTACCGCGAAAACCTTTGATCCTGAA
AGACACGCAGTGCAGTTGGTTAATTAATGTCCGCGCTTCCCACGGCGCGCCATTACGCTATTGCAAATGCAAATAGTTATCAATAATAT
TATCAATATATTTCTGCAATCAATGAAAAATTGCACAGTAAACATGGGGTTATGGTGTGACGGCGTTAAAAGTAGGAAGTGAGAGCTGG
TGGCAGTCGAAACATGGCCCGGAATGGCAGCGTCTGAATGACGAAATGTTTGAGGTCACTTTCTGGTGGCGTGATCCCCAAGGTTCTGA
AGAATACTCGACGATAAAGCGCGTGGATCCTCTAGATTTAAGAAGGAGATATACATATG
>xasA
GAGGCAATTCCGACGTCTAAGAAACCATTATTATCATGACATTAACCTATAAAAATAGGCGTATCACGAGGCCCTTTCGTCTTCACCTC
GATGGATCCACTGCAGGGTATTGCCCAACAGAACAGCTTTAAACATACCTGATAACGTTTAACGGTAACGGTGTCCCGAAACGAACCCG
TTTCGGGACAATTTCCAAAGTCTGTTCACTGGCATTAGCAACGGAAAATATTGTTCTGAATACGCTTCAGAACAAAACAGGTGCGGTTC
CGACAGGAATACCGTTTTAGGGGGATAATATGGCTACATCAGTACAGACAGGTAAAGCTAAGCAGCTCACATTACTTGGATTCTTTGCC
ATAACGGCATCGATGGTAATGGCTCGAGAGATCCTCTAGATTTAAGAAGGAGATATACATATG
>pheL
GAGGCAATTCCGACGTCTAAGAAACCATTATTATCATGACATTAACCTATAAAAATAGGCGTATCACGAGGCCCTTTCGTCTTCACCTC
GAGACAAAGGCGAAGCACGTCGTGCCGCAACATCGGTGAAAGACGCCAACTTCGTCGAAGAAGTTGAAGAAGAGTAGTCCTTTATATTG
AGTGTATCGCCAACGCGCCTTCGGGCGCGTTTTTTGTTGACAGCGTGAAAACAGTACGGGTACTGTACTAAAGTCACTTAAGGAAACAA
ACATGAAACACATACCGTTTTTCTTCGCATTCTTTTTTACCTTCCCCTGAATGGGAGGATCCTCTAGATTTAAGAAGGAGATATACATA
TG
>phoP
GAGGCAATTCCGACGTCTAAGAAACCATTATTATCATGACATTAACCTATAAAAATAGGCGTATCACGAGGCCCTTTCGTCTTCACCTC
GATGGATCCAGAAAGCCCGCCTGAAAGCGATGACGCCGGCTAACTATATTGGTCGAGCTATCACGATGGTTGATGAGCTGAAATAAACC
TCGTATCAGTGCCGGATGGCGATGCTGTCCGGCCTGCTTATTAAGATTATCCGCTTTTTATTTTTTCACTTTACCTCCCCTCCCCGCTG
GTTTATTTAATGTTTACCCCCATAACCACATAATCGCGTTACACTATTTTAATAATTAAGACAGGGAGAAATAAAAATGCGCGTACTGG
TTGTTGAAGACAATGCGTTGTTACGTCACCACCTTAAAGTTCAGATTCAGGATGCTGGTCATCAGGTCGATGACGCAGAAGATGCCAAC
TCGAGTTGGCATCTTCTGCGTCCCGCTCGAGAGATCCTCTAGATTTAAGAAGGAGATATACATATG
>phoR
GAGGCAATTCCGACGTCTAAGAAACCATTATTATCATGACATTAACCTATAAAAATAGGCGTATCACGAGGCCCTTTCGTCTTCACCTC
GAGATTCGTCGCCTGCGTAAAGCACTGGAGCCCGGCGGGCATGACCGCATGGTGCAGACCGTGCGCGGTACAGGATATCGTTTTTCAAC
CCGCTTTTAACGCCTTGCTCATCGGACGCAGAGCAGGGCTTATGATTTCTTAACTGGAGTATCTTACGTGCTGGAACGGCTGTCGTGGA
AAAGGCTGGTGCTGGAGGATCCTCTAGATTTAAGAAGGAGATATACATATG
>ndh

GAGGCAATTCCGACGTCTAAGAAACCATTATTATCATGACATTAACCTATAAAAATAGGCGTATCACGAGGCCCTTTCGTCTTCACCTC
GAGCGAAGAGCAGACGCACAAATTCAAGAATATCTCCCCGCATTTACAGCGCATTAAAGCGTTCAAAACCCTCGGGTAAATGCCCTCGT
CGCATCAGGTAACCTTGCCGGTACCTGATGCGCTCCGAATTCTGTGGGTCGGATAAGGCGTCCACGCCGCATCCGACAGTCGAGCATCA
ATGCCTGATGCGCTTCTTATCAGGCCTACCGAACGCCCTGCATACACCCCTCACTCTATATCACTCTCACAAATTCGCTCAAATAATAA
ACAATAAACTCTGTTTTTGATCTCACCCGGTAAAGTCGCCTATCTTTTCAGCAACAAAACTTGATTAACATCAATTTTGGTATGACCA
ATGCACCATTCATGTTATTCTCAATAGCGAAGAACATTTTCATTGCTGTAACCTGTTGTTAATTAAGAGCTATGTTAATAACCATTAAT
TAACAATTGGTTAATAAATTTAAGGGGGTCACGTTGACTACGCCATTGAAAAAGATTGTGATTGTCGGCGGCGGTGCTGGTGGGCTGGA
AATGGCAACACAGCTGGGGCATAAGCTGGGACGCAAGAAAAAGGATCCTCTAGATTTAAGAAGGAGATATACATATG
>yfhK
GAGGCAATTCCGACGTCTAAGAAACCATTATTATCATGACATTAACCTATAAAAATAGGCGTATCACGAGGCCCTTTCGTCTTCACCTC
GATGGATCCGAAAACTGGGGCGAGGATGGCCCATGGATGCGCATTTTCCGCAATGCGCGTAAGCAGTTGGGGTAAGTCGTCAGCCCATT
GGTTTTGCAGCCCGGATGCGGCATTAAGCACCAACCACTATCATAAGCTCCCAACTCCGGGAGCTTTTTTGTGTCTGTAAATCACGACA
ATGGGTGGTTTGCCGTGTCGCTTTCTGGCGACACTTAACTCATTGATTTTAATATTATCTAATAAGTTTATCTTAAGTTGTCTCTTTTT
AGCGACACAGTGGCTGATTTACATTCAAACAGCGGAGGAGACATAAACCCCTCAAAGCAACGTATAATCAGTACGTTACCAAACTATTT
TCTTTATTGGCACAGTTACTGCATAATAGTAACCAGTGGCTCATTCACCGACTTATGTCAGCCCCTTCGGGACGTGCTACATAAAATAC
GAATGACGCACAACAAGGTGCCTGCCGTCCAACTTCTGATATCAGCGTAGCTATATCAACCATCGGGCGAAACGTCGAGTTAGGCACCG
CCTTATTCCATAACAAAGCCGGGTAATTCCCGGCTTTGTTGTATCTGAACTTCCCCTCGGTTAGCATCAGGCTATTCGCGTCTGACGAG
AGTAACACCTTGAAACGCTGGCCCGTTTTTCCCCGCTCATTACGACTCGAGAGATCCTCTAGATTTAAGAAGGAGATATACATATG
>phoA
GAGGCAATTCCGACGTCTAAGAAACCATTATTATCATGACATTAACCTATAAAAATAGGCGTATCACGAGGCCCTTTCGTCTTCACCTC
GAGGGGCGCTGTACGAGGTAAAGCCCGATGCCAGCATTCCTGACGACGATACGGAGCTGCTGCGCGATTACGTAAAGAAGTTATTGAAG
CATCCTCGTCAGTAAAAAGTTAATCTTTTCAACAGCTGTCATAAAGTTGTCACGGCCGAGACTTATAGTCGCTTTGTTTTTATTTTTTA
ATGTATTTGTACATGGAGAAAATAAAGTGAAACAAAGCACTATTGCACTGGCAGGATCCTCTAGATTTAAGAAGGAGATATACATATG
>thiI
GAGGCAATTCCGACGTCTAAGAAACCATTATTATCATGACATTAACCTATAAAAATAGGCGTATCACGAGGCCCTTTCGTCTTCACCTC
GAGCAGGTCGCCACTTTCCAGACGGGTTACAATCTGTTCCAGCTCGCTCAGCGCCTTTTCAAAGCTGGCGGGCGCCTCATTTTTCTTCG
GCATAATGAATGTCTGACTCTCAATATTTTTCGCCCCGTCATGGTAACGGACTCAGGGCAAATAGCAAATAACGCGCAATGGTAAGGTG
ATGTGCACAGCAAAGCGATGTTAGTGGTATACTTCCGCGCCTGGATGCAGCCGCAGGTGTGGGCTGCTGTATTTTTCCCTATACAAGTC
GCTTAAGGCTTGCCAACGAACCATTGCCGCCATGAAGTTTATCATTAAATTGTTCCCGGAAATCACCATCAAAAGCCAATCTGTGCGCT
TGCGCTTTATAAAAATCCTTACCGGGAACATTCGGGATCCCGCCTTACCGGGAACATTCGGGATCCTCTAGATTTAAGAAGGAGATATA
CATATG
>phoB
GAGGCAATTCCGACGTCTAAGAAACCATTATTATCATGACATTAACCTATAAAAATAGGCGTATCACGAGGCCCTTTCGTCTTCACCTC
GAGTCTCCAGCAGCCAGTCAAGAAAAGCCTGATGTTCAGCTTCGCGGCTTTTACTGTAGAAGTTCTGGCCGAGATGCCAGTCTGAGGTG
TGAAGGATGCGCATAACGGTTCCCTGGCGAAAAAGCATGGGCGCGATTATACCCAAACAGATGTGCCATTTGCTTTTTTCTGCGCCACG
GAAATCAATAACCTGAAGATATGTGCGACGAGCTTTTCATAAATCTGTCATAAATCTGACGCATAATGACGTCGCATTAATGATCGCAA
CCTATTTATTACAACAGGGCAAATCATGGCGAGACGTATTCTGGTCGTAGAAGATGAAGCTCCAATTCGCGAAATGGTCTGCTTCGTGC
TCGAACAAAATGGCTTTCAGCCGGGATCCTCTAGATTTAAGAAGGAGATATACATATG
>pinE
GAGGCAATTCCGACGTCTAAGAAACCATTATTATCATGACATTAACCTATAAAAATAGGCGTATCACGAGGCCCTTTCGTCTTCACCTC
GAGGAACCGGGACACCAACAGGTAATGCAGAGCCTTCTCCCAAACCAACGTTTATGAAAATGAAGAAATAACAAGCAAATGGCATCATT
CCTGCTTTTACCAGGGGGATTTAACATGCTTATTGGCTATGTACGCGTATCAACAAATGACCAGAACACAGATCTACAACGTAATGCGC
TGAACTGTGCAGGATGCGAGCTGATTGGATCCTCTAGATTTAAGAAGGAGATATACATATG
>thiM
GAGGCAATTCCGACGTCTAAGAAACCATTATTATCATGACATTAACCTATAAAAATAGGCGTATCACGAGGCCCTTTCGTCTTCACCTC
GATGGATCCATCGTTCACCAGGGGGATGAGCTAAAACGTGAAGAAGATCTGGATGTCGTTCTGAAGGTGCTGGATTCATATATCAAATA
ATTTATTAACGCGATTGTAAAACTGCCGTTTTTCCTCGTTTACAACGCGTGCGCTGGACATTACCATCCTCCTCTGCGATTTATCATCG
CAACCAAACGACTCGGGGTGCCCTTCTGCGTGAAGGCTGAGAAATACCCGTATCACCTGATCTGGATAATGCCAGCGTAGGGAAGTCAC
GGACCACCAGGTCATTGCTTCTTCACGTTATGGCAGGAGCAAACTATGCAAGTCGACCTGCTGGGTTCAGCGCAATCTGCGCACGCGTT
ACACCTTTTTCACCAACATTCCCCTCTTGTGCACTGCCTCGAGAGATCCTCTAGATTTAAGAAGGAGATATACATATG
>thiL
GAGGCAATTCCGACGTCTAAGAAACCATTATTATCATGACATTAACCTATAAAAATAGGCGTATCACGAGGCCCTTTCGTCTTCACCTC
GAGCGAACTGGCGAAATCGTTCGGCGCAGAAGACAGCCATAAGTTCGTCAACGGCGTACTCGATAAAGCAGCACCTGTGATTCGCCCTA
ACAAAAAGTGATATCCAGGCCGGTAGATTCACGGAAGACCGTTCCATGATCGCCGGCCTTTTCTTTTTTACCTGCTGAGGCATAACGTA
TGGCATGTGGCGAGTTCTCCCTGATTGCCCGTTATTTTGACCGTGTAAGAAGTTCTCGTCTTGATGTCGAACTGGGCAGGATCCTCTAG
ATTTAAGAAGGAGATATACATATG
>thiC
GAGGCAATTCCGACGTCTAAGAAACCATTATTATCATGACATTAACCTATAAAAATAGGCGTATCACGAGGCCCTTTCGTCTTCACCTC
GATGGATCCTTCTGCTGGTGCTTGACGCCGCCCGCGTCAAACATCCTGCTTGAGTTCTGCGCTGTTAACGCGTAATTTACATTCAATGC
CCCATTTGCGGGGCTAATTTCTTGTCGGAGTGCCTTAACTGGCTGAGACCGTTTATTCGGGATCCGCGGAACCTGATCAGGCTAATACC
TGCGAAGGGAACAAGAGTTAATCTGCTATCGCATCGCCCCTGCGGCGATCGTCTCTTGCTTCATCCGTCGTCTGACAAGCCACGTCCTT
AACTTTTTGGAATGAGCTATGTCTGCAACAAAACTGACCCGCCGCGAACAACGCGCCCGGGCCCAACATTTTATCGACACCCTGGAAGG
CACCGCCTTTCCCAACTCAAACTCGAGAGATCCTCTAGATTTAAGAAGGAGATATACATATG

Data Set S4 - MG1655 library reference file.fasta

```
>prlC
GAGGCAATTCCGACGTCTAAGAAACCATTATTATCATGACATTAACCTATAAAAATAGGCGTATCACGAGGCCCTTTCGTCTTCACCTC
GATGGATCCCGTGAAAGCTGTGGCGATAACTGAGCATGGGTAAAGGTGTTCCGGTAAGTAAAAATCGGCCCGTATTCGGGGCGCACAAA
AGCGTATCAGGACAGTATACCGAAAAGAGGCCGCCGCCGCGAAAGCGTAACGTTTCTCATTGAAATTCACTACACTTAACCCCATGCTA
CACACATTATGTAAAGCGCCTGTTGAGCGCTTCCTTAACCTCTTTAACCAGGACTGCGCGAATGACGAATCCGTTACTGACTCCCTTTG
AATTGCCTCCGTTTTCTAAAATTCTCCCGGAACATGTCGTTCCAGCCGTGACTAAGGCCTCGAGAGATCCTCTAGATTTAAGAAGGAGA
TATACATATG
>thiG
GAGGCAATTCCGACGTCTAAGAAACCATTATTATCATGACATTAACCTATAAAAATAGGCGTATCACGAGGCCCTTTCGTCTTCACCTC
GATGGATCCTCGACGGTAAATCGAGCCAGTGGCGCAGCCTGGCGTTGCGCCGCGCCAGTGGTTGCCCGGTATGCGGAGGAAGCAATGCA
GATCCTGTTTAACGATCAAGCGATGCAGTGCGCCGCCGGGCAAACTGTTCACGAACTACTGGAGCAACTCGACCAACGACAAGCGGGCG
CGGCTCTGGCGATTAATCAGCAAATCGTCCCGCGTGAGCAGTGGGCGCAACATATCGTGCAGGATGGCGACCAGATCCTGCTTTTTCAG
GTTATTGCAGGGGGTTGAAATGCTCGAGAGATCCTCTAGATTTAAGAAGGAGATATACATATG
>yiaU
GAGGCAATTCCGACGTCTAAGAAACCATTATTATCATGACATTAACCTATAAAAATAGGCGTATCACGAGGCCCTTTCGTCTTCACCTC
GAGGCACCAGCACCAATGGATAATTCAGAAGCAGTTGCGCTTGCCATAAAAGGCAACGCAAATAACGCCACAATATTGCGATTAATTAA
CATGTCCGGTATTCCATTTTAAAATAAGATAAAAAAGAGTCGGCATAATAATGCTTACAATTTCTCCGGTGCAAGATTCATATTTATAT
AGTAATATATAAAATTATATATAATTGGGCTGTTGCGATGACAAAATTACAACTTAAGTACCGGGAGTTAAAAATTATCTCGGTAATCG
CTGCCAGTGAAAATATCAGCCATGCCGCGGATCCTCTAGATTTAAGAAGGAGATATACATATG
>yiaT
GAGGCAATTCCGACGTCTAAGAAACCATTATTATCATGACATTAACCTATAAAAATAGGCGTATCACGAGGCCCTTTCGTCTTCACCTC
GATGGATCCACTGGCAGCGATTACCGAGATAATTTTTAACTCCCGGTACTTAAGTTGTAATTTTGTCATCGCAACAGCCCAATTATATA
TAATTTTATATATTACTATATAAATATGAATCTTGCACCGGAGAAATTGTAAGCATTATTATGCCGACTCTTTTTTATCTTATTTTAAA
ATGGAATACCGGACATGTTAATTAATCGCAATATTGTGGCGTTATTTGCGTTGCCTTTTATGGCAAGCGCAACTGCTTCTGAATTATCC
ATTGGTGCTGGTGCCTCGAGAGATCCTCTAGATTTAAGAAGGAGATATACATATG
>yiaW
GAGGCAATTCCGACGTCTAAGAAACCATTATTATCATGACATTAACCTATAAAAATAGGCGTATCACGAGGCCCTTTCGTCTTCACCTC
GATGGATCCGCAAACCAAGTGCCTGCTGGTGAGCTACTCGGATAAACCGTTGGGGCTGTTCTGATATAAGAAGCTGGTCGCATTGGGTA
TTCATTGCCTGATGCGACGCTTACGCGTCTTATCATGCCTACGGGAACCTGACCGTAGGCCGGATAAGGCGTACACGCCGCATCCGGCA
TAAACAAAGCGCACGTTGTTACAATTTACGTTTTAAACCTACGTGCGCAATAACCCAGACTACGCCAACAGAAGTACGTTTCTGGGGCG
TGTTTTGTCAGGAGTTTAATCTCGCGTTTCGATGGTGAGCGGGCGAACCAGGCCGTAAATCACCATTGGTGATCCGAGGCCAATCAGCA
ATAAACGGTAGATAATTGAAGGTTTAATACGCTTAATCTGTACTGTGGATGTTTTACTGGAGTTATTCATTAGTTTGGCCAATAGTTCC
ACTATTTCAATATGATAAACCTCGTCCTGTTGAAAAACCAGACAGCACATTTAAGAAAGTATATAAAAAACAATATGTCGTTATATAAA
TAAGGAAATTGCACTGCATCATGGCATTTCATAATATGCGTTAATTGTTCTTTTTTATATTCTTTGAGGGAAATATGTTCCTGGACTAT
TTTGCACTGGGAGTGCTTATTTTTGTATTTCTGGTGATCTTCTATGGGATCATTATTTTACATGATATTCCCTACCTGATTGCCAACTC
GAGAGATCCTCTAGATTTAAGAAGGAGATATACATATG
>fabF
GAGGCAATTCCGACGTCTAAGAAACCATTATTATCATGACATTAACCTATAAAAATAGGCGTATCACGAGGCCCTTTCGTCTTCACCTC
GAGTTCCGGACGAAGAAGCTGAGAAAATCACCACCGTTCAGGCTGCCATTGATTACATCAACGGCCACCAGGCGTAAGTGAACATCTCC
AGGCGGTCGTTCGACCGCCTGAGTTTTATCTTTTTGTCCCACTAGAATCATTTTTTCCCTCCCTGGAGGACAAACGTGTCTAAGCGTCG
TGTAGTTGTGACCGGACTGGGCATGTTGTCTCCTGTCGGCAATACCGTAGAGTCTACCTGGAAAGCTCTGCTTGCCGGATCCTCTAGAT
TTAAGAAGGAGATATACATATG
>yiaD
GAGGCAATTCCGACGTCTAAGAAACCATTATTATCATGACATTAACCTATAAAAATAGGCGTATCACGAGGCCCTTTCGTCTTCACCTC
GAGGAAAGCCTTTTCGCACCATTGGAAATCGTACCCGCGTATTGCTGTGAACCTGGTCGCGAACCGCGCTCTGCAAGGAGTTTTCCATT
CCTGTGGCTAACGCGCCACGCGAGCTAAACACGGTTTCGCCGTCGGTTTCAACCAGCATGGGCCCCCAATGGGCGGCAGTCAGAACGGA
ATATCGTGAGGATGAGTTGGCCAAACCGAAGACTCCTGAAAGGTTAGATGTATTGAGCAGTTAAATATTACTTACAAATTTCCGAGTAT
TCCCAGGAATAATCTTCATGTTCACGCGGCATAATCTCCCGCCACGGAACCCGTGGCAAAGAATAAAAAGGTTATTAAGGATTAACAAT
GAAGAAACGTGTTTATCTTATTGCCGCCGTAGTGAGTGGCGCTCTGGCGGTATCTGGCTGCACAACTAACCCTTACACCGGCGAGGATC
CTCTAGATTTAAGAAGGAGATATACATATG
>yiaG
GAGGCAATTCCGACGTCTAAGAAACCATTATTATCATGACATTAACCTATAAAAATAGGCGTATCACGAGGCCCTTTCGTCTTCACCTC
GAGAAAGCGCACAACGACACACGCGTGTCTGACGTGACAAGGATAGTCCAGGACAGGCTTGCAGGATACCCTTTCTACGTAATCTTCCT
GGTAAAATCATGGAGAAAATGAAACAAGGTTAATTTGTTGTTTCATTGTTAAAAAATGATAGTGCCGTTGCCTTTTTTGTAGCCAGAGA
TTTACGCCTTCGCAGAAGGTTAGATTTTCGAAATCTTTACAAAACAGGAGTGGTAAGAGATATATCGGCATTTATCAGATCGTTAGGTG
GCTATAAGTCACGCTAAATGACAGGCTGAATCGAATCATAGCCAGAGCATGCCCTGACTTCACCCCGCTGTGTCTGCTTTTCCCGACTA
TTCTTAATGAGCTTCGATGCAATTCACGATCCCGCAGTGTGATTTGAGGAGTTTTCAATGGAATATAAAGATCCAATGCATGAGCTGTT
GAGCAGCCTGGAACAGATTGTTTTTAAAGATGAAACGCAGAAAATTACCCTGACGCACAGAACAAGGATCCTCTAGATTTAAGAAGGAG
ATATACATATG
>yiaF
GAGGCAATTCCGACGTCTAAGAAACCATTATTATCATGACATTAACCTATAAAAATAGGCGTATCACGAGGCCCTTTCGTCTTCACCTC
GATGGATCCTTGTTCTGTGCGTCAGGGTAATTTTCTGCGTTTCATCTTTAAAAACAATCTGTTCCAGGCTGCTCAACAGCTCATGCATT
GGATCTTTATATTCCATTGAAAACTCCTCAAATCACACTGCGGGATCGTGAATTGCATCGAAGCTCATTAAGAATAGTCGGGAAAAGCA
```

```
GACACAGCGGGGTGAAGTCAGGGCATGCTCTGGCTATGATTCGATTCAGCCTGTCATTTAGCGTGACTTATAGCCACCTAACGATCTGA
TAAATGCCGATATATCTCTTACCACTCCTGTTTTGTAAAGATTTCGAAAATCTAACCTTCTGCGAAGGCGTAAATCTCTGGCTACAAAA
AAGGCAACGGCACTATCATTTTTTAACAATGAAACAACAAATTAACCTTGTTTCATTTTCTCCATGATTTTACCAGGAAGATTACGTAG
AAAGGGTATCCTGCAAGCCTGTCCTGGACTATCCTTGTCACGTCAGACACGCGTGTGTCGTTGTGCGCTTTCTCGAGAGATCCTCTAGA
TTTAAGAAGGAGATATACATATG
>yiaJ
GAGGCAATTCCGACGTCTAAGAAACCATTATTATCATGACATTAACCTATAAAAATAGGCGTATCACGAGGCCCTTTCGTCTTCACCTC
GATGGATCCAGCCGTTTCGCTGTCAACGCCGCGTGAAATTAAGACCCGATTAAAGGCTGCTTTTAACTGCTCAAATGTCACTTTCATCC
CAGGCTTCCTTGTTTTTTTGATCGCTTTTTTGGCTTAAAATTTCAATATGCGAAACTTGATTTCAAATATATCGATCACTTTTTAACAA
GGCAATCTGATTGATGAATTTCAAAAGACATAAAAATCAATTGGTTATAAATTATCTGTCGAGATCGTGAACTACGGCACACTTTGCGC
TACCATCAGGACGCGACAAAATGGGGAAAGAAGTGATGGGAAAAAAAGAGAACGAGATGGCGCAGGAAAAAGAGCGTCCAGCCGGAAGC
CAGAGTCTCTCGAGAGATCCTCTAGATTTAAGAAGGAGATATACATATG
>yqjH
GAGGCAATTCCGACGTCTAAGAAACCATTATTATCATGACATTAACCTATAAAAATAGGCGTATCACGAGGCCCTTTCGTCTTCACCTC
GATGGATCCCTTTGCAGCAGCCCTCATGGCGTGGCTGGCCTTCATGTTTACAACACCCTTCGTGATGATGGCTCATATCAGCCTCCTTT
TGCTTAATTAGATATATCTAATTTATATCTGATATGAGCGTTCTTGCAAGTGTAAAAATGATTTTTAAACGTGGACTCATCTCACACGT
AGGCCGGATAAGGCGTTCACGCCGCATCCGGCGTTGAGTAATGATGCCTGAGAAGAGGCGGTTCGTCTTATCAGACTATAGACATATCA
GTTCTACAAATCGCTTGCATTTATCATGATTAACAATCATTATCATTTGCGAGTTTTATTTAGATATATCTGATTACCATCACGAAGGC
GATAACAATGAATAACACCCCCCGCTACCCGCAGCGCGTTCGCAATGATCTGCGCTTCCGTGAACTGACTGTGTTACGCGTTGAGCGCT
CGAGAGATCCTCTAGATTTAAGAAGGAGATATACATATG
>narX
GAGGCAATTCCGACGTCTAAGAAACCATTATTATCATGACATTAACCTATAAAAATAGGCGTATCACGAGGCCCTTTCGTCTTCACCTC
GATGGATCCGTTGCCAGAACGCAGGATCTTCCGGTCGCCAATCTGTAATGACAGCTCCAGTAGCCCTTTCGGGGGCGGATGAGTGACTC
ATAGACACCTCTGATACTCGTTTCGATGTCTGCCACCTTAGTGTCTGTAGCTAAAGGCAATTTGATGTAAATCAAACGATAAGATAACT
TTATCTATCATTGATATTTATCATTACCCATAGTGAGTACAGTGACTTCATAAAAATTATGAGATTTTTCACGGTGCTGTAAAATCCCT
ACCCTTACCGATGTAAAGCGACTAACCACACGGCAAATAAGGAGTAACTCTTTCCGGGTATGGGTATACTTCAGCCAATAGCCGAGAAT
ACTGCCATTCCAGAATGTATCGTCACATTCATTAAGGTTATTGCTCATTTAAAGCCTGAAGGAAGAGGTTTACATGCTTAAACGTTGTC
TCTCTCCGCTCACCCTGGTTAATCAGGTTGCGCTTATTGTGTTGCTTTCTACTGCTATTGGACTGGCAGGGATGCTCGAGAGATCCTCT
AGATTTAAGAAGGAGATATACATATG
>tar
GAGGCAATTCCGACGTCTAAGAAACCATTATTATCATGACATTAACCTATAAAAATAGGCGTATCACGAGGCCCTTTCGTCTTCACCTC
GATGGATCCGCGACCGGATGTTGATTCTGGTGAACATCGAAAAACTGCTGAACAGCGAAGAGATGGCGCTGTTAGATAGCGCGGCGTCA
GAAGTGGCGTAATTCTCCCGATTTCCTCAATTGAAATGAACCCGATGATCTGCGCATCGGGTTTTTTATTTCAATTTCGCGGCGGGTGG
CATCAGCAATAAAGTTTCCCCCCTCCTTGCCGATAACGAGATCAACTTGTTTTCAGGAAGGTGCCTTATGATTAACCGTATCCGCGTAG
TCACGCTGTTGGTAATGGTGCTGGGGGTATTCGCACTGTTACAGCTTATTTCCGGCAGTCTCTCGAGAGATCCTCTAGATTTAAGAAGG
AGATATACATATG
>narZ
GAGGCAATTCCGACGTCTAAGAAACCATTATTATCATGACATTAACCTATAAAAATAGGCGTATCACGAGGCCCTTTCGTCTTCACCTC
GATGGATCCCTGCTGACCTGGCTGGTTTATGGTCGGCGGAAGTTCAGCCAAAAATAAGAGATCGATGCCTGATGTACAACCGTGTCGCA
TCCGACAAGGCATCAGATCCCAATTCATTCCTGGAGCAGGAGTCATGTCATGAGTAAACTTTTGGATCGCTTTCGCTACTTCAAACAAA
AGGGCGAAACCTTTGCCGATGGTCACGGACAGGTCTCGAGAGATCCTCTAGATTTAAGAAGGAGATATACATATG
>narU
GAGGCAATTCCGACGTCTAAGAAACCATTATTATCATGACATTAACCTATAAAAATAGGCGTATCACGAGGCCCTTTCGTCTTCACCTC
GATGGATCCGGACAGATCGTTTTCCTCCAGTTCATGGCCTTTTTAGTGAATAAATCATCACTAATTCTATAAGTGTCTGGTCTTGTTGA
TAATATAGTTAATATGTAACCCTTTAGATTCACGATCTGAAGGGTTGAAACATGACATACGTTCAAAGATTAACCCCGTTTGAAGGTTA
GCTTGTTAGTTACATTTAGTAACACATATCTGGCTACTTAATCCCCATTCTGCCCCCGCGGTTACGCTTTCTTATCTATTCTTTCAATC
AGCACTTCACCCAATTCATCCTTAATTCTAATAACTGGCTATTGCGCCTTACCTCGTGTTTTGTCTTTTTTTCATTACCAATGTGTGCA
TGTGAGGAACAATATGGCACTGCAAAATGAGAAAAATAGTCGTTATCTTTTGCGCGACTGGAAACCAGAAAATCCGGCCTTCTGGCTCG
AGAGATCCTCTAGATTTAAGAAGGAGATATACATATG
>cbdA
GAGGCAATTCCGACGTCTAAGAAACCATTATTATCATGACATTAACCTATAAAAATAGGCGTATCACGAGGCCCTTTCGTCTTCACCTC
GAGGCAGCGCCAGAAGATTTGGTCGACTCTGCGCAGCGGCTTAGCGAGGTATGTCAGTGGCTGGCGGAAGCTGCACCGACGTAAAAAGA
CGGTAAGTATCGCTTTCAGTCTTATGAATATCGCAATCGGCGAATACCTCTGGTCGTAGAGTTTCAGGATAAAGAGGGAGATCTACCAT
TATCGGGTTATTTTTCTCTCTTCGCCTACAGGAGTGCGCATGTGGGATGTCATTGATTTATCGCGCTGGCAGTTTGCTCTGACCGCGCT
GTATCACTTTTTATTTGTACCCCTTACCCTGGGGCTGATGGATCCTCTAGATTTAAGAAGGAGATATACATATG
>narQ
GAGGCAATTCCGACGTCTAAGAAACCATTATTATCATGACATTAACCTATAAAAATAGGCGTATCACGAGGCCCTTTCGTCTTCACCTC
GAGAGCCATGACACAGGCGATTTCACAAGCATGACAACCCAGACATTGCTGACTGTTGGCCATAATAAAACGATTCATAACGACCTTCT
TTTTTGGTTGTAAAAACCTTATTCTTTATATGAGTGTTGTTATTACCCGACTTACAGGGGAATCGGCAATGTTCATGTGCCCAGAATAA
GTAACTATTTCGCTATAAACTGTGGCAGATCAAATAATCCCATCACTGACTAAATTGCGTTTCAGCGAACTGGAACATTAATGATTTTT
TGTGGAGAAGACGCGTGTGATTGTTAAACGACCCGTCTCGGCCAGTCTGGCCCGGGCCTTTTTTTACATTGTGCTGCTGTCGATTCTTT
CCACGGGTATCGCTCTGCTGGATCCTCTAGATTTAAGAAGGAGATATACATATG
>tam
```

GAGGCAATTCCGACGTCTAAGAAACCATTATTATCATGACATTAACCTATAAAAATAGGCGTATCACGAGGCCCTTTCGTCTTCACCTC
GAGAAACCACGCCCCACTACAAAACCTGTGTCGCGAAACTGGAATCTTTAATGACCGGGCCGCGTAAAAAACGTCTGTTCAATGGTTTG
ATGCCGTGAGGCGAATTTATCAATTTTATCTACAATTGGGGTAACGCGCTGACGGGAGTAAAAAAATGTCTGACTGGAACCCCTCTTTA
TATCTACACTTTTCCGCTGAACGATCGCGTCCGGCGGTGGAGCTGCTTGCCAGAGTGCCGCTGGAAAATGTCGGATCCTCTAGATTTAA
GAAGGAGATATACATATG
>gcl
GAGGCAATTCCGACGTCTAAGAAACCATTATTATCATGACATTAACCTATAAAAATAGGCGTATCACGAGGCCCTTTCGTCTTCACCTC
GAGTCAGGGTGAGCTGGTCAGAGACACCGCCCGCGATATCAGCACGGCGTTGGGACTGAAAGCACATCCATAATGTCTGTCGCATCCCG
CTCTGCGGAGCGGGTTTTTTTGACAAAATTTGAAAGTTGGAAAAATTTTCCAATAAATAGAGGTAGGAATAAAATGGCAAAAATGAGAG
CCGTTGACGCGGCAATGTATGTGCTGGAGAAAGAAGGTATCACTACCGCCTTCGGTGTGGATCCTCTAGATTTAAGAAGGAGATATACA
TATG
>yneF
GAGGCAATTCCGACGTCTAAGAAACCATTATTATCATGACATTAACCTATAAAAATAGGCGTATCACGAGGCCCTTTCGTCTTCACCTC
GATGGATCCGCATAACATTCCGCAGGGTGTTTCGTTAAGCGAAGAACAACAACGTTATATCGTGGCGGTAATCTATCACTGGCTGGTTG
TACAGATGAATCAGCCATGAGAATTGCGGAAAACGACAAAATAGCATTCCCAGGGTACCTTTATTTGCCGCCGCTGTCAGCGAACATGC
TATCTTGGTTCGGTCAGTATTGATAGTCTGGATTTCTGGCCTTTGCTTGCGGACTGGTTCAGTGAGCAATTCTCAACGGGCGTGTTGAT
CGTGCCTTGTATGCTGACGTTGGCTATTCCCTCGAGAGATCCTCTAGATTTAAGAAGGAGATATACATATG
>paaX
GAGGCAATTCCGACGTCTAAGAAACCATTATTATCATGACATTAACCTATAAAAATAGGCGTATCACGAGGCCCTTTCGTCTTCACCTC
GAGGGCAGTATCCCGCGTTCAGAAGGCAAGGCGTGTCGGGTGTTTGATCTGCGCAATATTGTTGGTGCCTGACGTATCGTCTGGCCCTG
GTGGGGTAAAGCGCCAGGGCCAGAAGTCGATACGACCTGTGCTATGATTCATAAATCACAACAATAACAACAGACTGAAGCGAATGAGT
AAACTTGATACTTTTATCCAACATGCTGTAAACGCTGTTCCGGTCAGGATCCTCTAGATTTAAGAAGGAGATATACATATG
>yrbL
GAGGCAATTCCGACGTCTAAGAAACCATTATTATCATGACATTAACCTATAAAAATAGGCGTATCACGAGGCCCTTTCGTCTTCACCTC
GAGCCAAAGGACGGCAGATTGAAGTTGAAGCGACCGGTCCACAGGAAGAGGAAGCACTGGCCGCCGTTATCGCCCTCTTTAATTCTGGT
TTTGATGAAGATTAATCTTCATCACTTTGACATACAACTCCCTTCAAACTCCCCCCGACAATAAGAAAATCACGTACTGAAATCGTTCT
CAATCAACGTCATTTGTACATTTTGTGCGCTTTTCACTTTCAGAAGAACCTTAAGAAAACCTTAAGAGGCATTGTTTAGGTTTTGTTTA
AGTTAATCGACCATACTGGAGATCGTCAGAAAATATTTCCAGGAGATGGCATGATTCGCTTATCTGAACAAAGTCCCCTGGGCACCGGA
AGACATCGGAAATGTTATGCGCATCCGGAAGATGCCCAACGCTGTGGATCCTCTAGATTTAAGAAGGAGATATACATATG
>yrbG
GAGGCAATTCCGACGTCTAAGAAACCATTATTATCATGACATTAACCTATAAAAATAGGCGTATCACGAGGCCCTTTCGTCTTCACCTC
GAGTCGAAGATGCAGCGATTGCCACGCGTAAAACTGACATCGCGCATATCGACTAAATTCGCCACAGACTGCTCCATAATTCACCCTTC
GTCTTGCGTTGATTTTCTAAGCATGGCGCTCAATTTAACCTTGAACCCAACATATTTACAGAATATTACCCGCCGTGGTTAGCGAAAGC
TGGCATTTGTTTTACTTTTTAGCCGCATAAAGTCAAAATTAAGCATCCGTTACGGCTTTCTGAAAATCTTCAGCGGACCGGCGAGTATA
CCTGAAGAAAGGACGTTAGATGCTTTTAGCTACGGCACTGTTAATTGTTGGTTTACTTTTGGTCGTTTACAGTGCCGACCGCCTGGTTT
TTGCCGCGTCTATTCTTTGCCGAACCTTTGGGGATCCTCTAGATTTAAGAAGGAGATATACATATG
>yrbF
GAGGCAATTCCGACGTCTAAGAAACCATTATTATCATGACATTAACCTATAAAAATAGGCGTATCACGAGGCCCTTTCGTCTTCACCTC
GATGGATCCCCAAAGGTTCGGCAAAGAATAGACGCGGCAAAAACCAGGCGGTCGGCACTGTAAACGACCAAAAGTAAACCAACAATTAA
CAGTGCCGTAGCTAAAAGCATCTAACGTCCTTTCTTCAGGTATACTCGCCGGTCCGCTGAAGATTTTCAGAAAGCCGTAACGGATGCTT
AATTTTGACTTTATGCGGCTAAAAAGTAAAACAAATGCCAGCTTTCGCTAACCACGGCGGGTAATATTCTGTAAATATGTTGGGTTCAA
GGTTAAATTGAGCGCCATGCTTAGAAAATCAACGCAAGACGAAGGGTGAATTATGGAGCAGTCTGTGGCGAATTTAGTCGATATGCGCG
ATGTCAGTTTTACGCGTGGCAATCGCTGCATCTTCGACTCGAGAGATCCTCTAGATTTAAGAAGGAGATATACATATG
>yrbA
GAGGCAATTCCGACGTCTAAGAAACCATTATTATCATGACATTAACCTATAAAAATAGGCGTATCACGAGGCCCTTTCGTCTTCACCTC
GATGGATCCTGACGCTTCAGGGGGTAAACGACAAAGTGTATACCCTGGCGAAATTGTATAATTTGCCTGCTGATGTTCTGCCTCGTTAA
ATTTTTTCAGGCCGGATATGGCGTTAACCGCCTGTCCGGCAAACCTGCCTGCAAGTCGAGCCTTGTCTGATTTATCAGCGAGGCTTTTT
GCTTATTTATGCCAGCGCAACTTTGCTCTAAGATGTTTCGCTGGTTTATTAACTGATGATTGAAGATCCCATGGAAAATAATGAAATTC
AGAGCGTGTTGATGAACGCTCTCTCCCTCCAGGAAGTCCACGTTTCCGCTCGAGAGATCCTCTAGATTTAAGAAGGAGATATACATATG
>kbl
GAGGCAATTCCGACGTCTAAGAAACCATTATTATCATGACATTAACCTATAAAAATAGGCGTATCACGAGGCCCTTTCGTCTTCACCTC
GATGGATCCAAAAATGCAGGCGCTGAGAATTTTCCTTTCAAGAAAATAGGTTTCCCTCCATTAGTGGGTAGCGTTTTGATACCAACTGA
GATAGGAAAAATGATGCTTTGGCATCATTTTTTCTTTTTATATCAATGCAGTAAAATCAATCATCATTATGCCCAGCCAAAACTGTACA
GTTTTTCTACTATCTGTAGGTGTTGAGAGACATCAAGTTACACGTTATTTATCCTGAATTTTGCAGAAGTGTTAACGCGTTATCTCGTC
GCGACCTATAAGTTTGGGTAATATGTGCTGGAATTTGCCCTGTCTGGAGAATCGCAATGCGTGGAGAATTTTATCAGCAGTTAACCAAC
GATCTGGAAACCGCACGGGCGGAAGGGTTGTTTAAAGAAGAGCGCATTATTACGTCTGCGCAGCAAGCTCGAGAGATCCTCTAGATTTA
AGAAGGAGATATACATATG
>dmsA
GAGGCAATTCCGACGTCTAAGAAACCATTATTATCATGACATTAACCTATAAAAATAGGCGTATCACGAGGCCCTTTCGTCTTCACCTC
GAGCAGCAGGCTGATGGTCGTATTGAAGTACCAGAAGTTCTGCGTCCGTATATGAACGGACTGGAATATATTGGCTAATACCCAATTTT
TCTGAATCTAAAAAGCGCCTGCGGGCGCTTTTTTTTGTCTCCCTTTGATACCGAACAATAATTACTCCTCACTTACACGTAATACTACTT
TCGAGTGAAAATCTACCTATCTCTTTGATTTTCAAATTATTCGATGTATACAAGCCTATATAGCGAACTGCTATAGAAATAATTACACA
ATACGGTTTGTTACTGGAATCAATCGTGAGCAAGCTTGAGTGAGCCATTATGAAAACGAAAATCCCTGATGCGGTATTGGCTGCTGAGG

```
TGAGTCGCCGTGGTTTGGTAAAAACGACAGCGATCGGCGGCCTGGCAATGGCCAGCAGCGCATTAACATTACCTTTTAGTCGGATTGCG
CACGCTGTCGATAGCGCCATTCCAACAAAAGGATCCTCTAGATTTAAGAAGGAGATATACATATG
>ruvA
GAGGCAATTCCGACGTCTAAGAAACCATTATTATCATGACATTAACCTATAAAAATAGGCGTATCACGAGGCCCTTTCGTCTTCACCTC
GATAGATCTTTTCAGGAGGCGCAATGTATATTTTTATTACGCATTTCTTCACTGAATATGTAATATTAAAATATTTGCTTCCAATATAA
CCTGTAGAATAAATTATACTGTGCCATTTTTCAGTTCATCGAGACACCTCGCAAGTTTTCTTCATCCTTCGCTGGATATCTATCCAGCA
TTTTTTTATCATACAGCATTATCTTTGATTCATTACGCAGGAGCGTCATGTGATAGGCAGACTCAGAGGCATCATCATTGAAAAACAAC
CCCCGCTGGTGTTAATTGAAGTGGGCGGCGTAGGCTATGAAGCTCGAGAGATCCTCTAGATTTAAGAAGGAGATATACATATG
>tsf
GAGGCAATTCCGACGTCTAAGAAACCATTATTATCATGACATTAACCTATAAAAATAGGCGTATCACGAGGCCCTTTCGTCTTCACGTC
GACCTGTACCTGGGCGCTGTTGCTGCAACCGTACGTGAAGGCCGTTCTCAGGATCTGGCTTCCCAGGCGGAAGAAAGCTTCGTAGAAGC
TGAGTAATAAGGCTTGATAACTCCCCCAAAATAGTTCGAGTTGCAGAAAGGCGGCAAGCTCGAGAATTCCCGGGAGCTTACATCAGTAA
GTGACCGGGATGAGCGAGCGAAGATAACGCATCTGCGGCGCGAAATATGAAGGGGGAGAGCCCTTATAGACCAGGTAGTACACGTTTGG
TTAGGGGGCCTGCATATGGCCCCCTTTTTCACTTTTATATCTGTGCGGTTTAATGCCGGGCAGATCACATCTCCGAGGATTTTAGAATG
GCTGAAATTACCGCATCCCTGGTAAAAGAGCTGCGTGAGCGTACTGGCGCAGGCATGATGGATTGCAAAGGATCCTCTAGATTTAAGAA
GGAGATATACATATG
>mutH
GAGGCAATTCCGACGTCTAAGAAACCATTATTATCATGACATTAACCTATAAAAATAGGCGTATCACGAGGCCCTTTCGTCTTCACCTC
GAGCCACATTACCTGCCCCTGGCGATTACAAATCACGATACCTACGTTTGGGCGGTAGCCATCGTCATCAATCACCGGACTACCTCAAA
ATAAAGCTTTATATACGAATGATTGTTTCATACTCCAGGAAGACGGTAAACCACTCTCTGCAGGGCATTACACACTAATAACAATTGAA
TAACTCACAGTTATGTGCAGAGTTATAAACAGAGGAAGGGGTGGATAGCCGTTTTTATTCACCTTTTCTGTGGATAGAGTTGTGAAGAA
CTACGGAATTACTACGGGAAAACCCGGAGCATTCTGAATAAGCCCGATATAGACATTTTAAAATATTCATTATCAGTCAATGCATTACA
TGTAATTTTAGTAATGAAAAAGAGTAATTCGTGACCCAGGTCACACCTCTCATTTACGGGTTGGCGAAAGATCAACCAATGCCGTATTT
ATCCACAGAATGTGCCACTAAGTTAAGCACTGAACCACTAAAAACTGGAGTTTCGTCGCACGTCAAGGCTGTAAATGGAAACAGTAGTG
GAGGTTTTTCACAGTTATCCCAGCTTTCTGTGGATAACATGGTGTAAGATCCTGTTTATTTTCAGTGACCAGATTTGGAAAACCCGTTG
CAGTGTTGCGCAACTCGATTACCGGCAACCTAAAAAGCAATATAAATCAGAGAATTGAACAACGCATGTGGAAAAAGTTACACTGCGAA
TATTCGGCACATAATTGCTGTTTGTTTTTAATCAAGGTATCATGACATGTCCCAACCTCGCCCACTGCTCTCTCCTCCCGAAACTGAA
GAACAGTTGTTAGCGCAAGCACAGCAACTTTCTGGTTATACATTGGGAGAACTGGCGGGATCCTCTAGATTTAAGAAGGAGATATACAT
ATG
>ynjF
GAGGCAATTCCGACGTCTAAGAAACCATTATTATCATGACATTAACCTATAAAAATAGGCGTATCACGAGGCCCTTTCGTCTTCACCTC
GATGGATCCCATAATCCCGCCTGATCGCTCTGGGCGGGGCGTTGCGCGAGTAAAATTTTGCCATCACGTTCAATGATGGCGGCAACAAC
TTCAATCATTTTCATAAATTACACACGGGCAGATAAGCGAAAAATGCCATAATATCCGCACTGTTGCTCAATCAGAAAGGCTTTATATT
TAAGGAGTGCCTGTGCTAGACCGCCATCTTCATCCCCGGATTAAACCGTTGTTGCATCAGTGCGTGCGGGTTCTTGATACTCGAGAGAT
CCTCTAGATTTAAGAAGGAGATATACATATG
>mutM
GAGGCAATTCCGACGTCTAAGAAACCATTATTATCATGACATTAACCTATAAAAATAGGCGTATCACGAGGCCCTTTCGTCTTCACCTC
GATGGATCCTCGCCAGCACGTGATCTACAAAGAAGCGAAAATCAAATAATTCTCGCTTTGATGTAACAAAAAACCTCGCTCCGGCGGGG
TTTTTTGTTATCTGCTTGCCCCCATATTGACTGCATCTGTTCATTCCTGGAGATGCTATGCCTGAATTACCCGAAGTTGAAACCAGCCG
CCGCGGCATAGAACCGCATCTCGTTGGTGCAACCATTCCTCGAGAGATCCTCTAGATTTAAGAAGGAGATATACATATG
>mutS
GAGGCAATTCCGACGTCTAAGAAACCATTATTATCATGACATTAACCTATAAAAATAGGCGTATCACGAGGCCCTTTCGTCTTCACCTC
GAGCGGGCATTTCGCTTGATAAAGATCGATCATTTTTTTAATCGTCAGTTTTTCACGAGAGATACGCTTGCCGGACATGCTGCCTCCAC
CTCATTAAGATGTATTTATATTACATCTTAATCTTAAAGGGCACTATGACTCCAAAGAAGAAGGGTTAGCCAACCGATACAATTTTGCG
TACTTGCTTCATAAGCATCACGCAAAAGCTGCAAAACAGCATCTTTCCCGGAACCAGCATCAAGAACTCGCCGTTCGCTTCTTCCCCTG
AAATGATTAACTCCGGTATCATGTGCGCCTTATGTGATTACAACGAAAATAAAAACCATCACACCCCATTTAATATCAGGGAACCGGAC
ATAACCCCATGAGTGCAATAGAAAATTTCGACGCCCATACGCCCATGATGCAGCAGTATCTCAGGCTGAAAGCCCAGCATGGATCCTCT
AGATTTAAGAAGGAGATATACATATG
>mutT
GAGGCAATTCCGACGTCTAAGAAACCATTATTATCATGACATTAACCTATAAAAATAGGCGTATCACGAGGCCCTTTCGTCTTCACCTC
GAGCGGAGAGCGCAAAGTAGGACGTAACGATCCTTGCCCGTGCGGTTCTGGTAAAAAATACAAGCAGTGCCATGGCCGCCTGCAATAAA
AGCTAACTGTTGAAGTAAAAGGCGCAGGATTCTGCGCCTTTTTTATAGGTTTAAGACAATGAAAAAGCTGCAAATTGCGGTAGGTATTA
TTCGCAACGAGAACAATGAAATCTTTACAACGCGTCGCGCAGCAGATGCGCACATGGGGATCCTCTAGATTTAAGAAGGAGATATACAT
ATG
>mutY
GAGGCAATTCCGACGTCTAAGAAACCATTATTATCATGACATTAACCTATAAAAATAGGCGTATCACGAGGCCCTTTCGTCTTCACCTC
GAGGGCGCACAAAACTACGGATACGGCGCAGTGGGCGGCCGTTTTCATCAAATTCCGGTGAAATGACGTCGTTTTTCATAAAGGTTTAG
TCGCTTGTGAAAGTGTTCTGAAAACGGGCATTATCCAAAGTTAGTTGCCGGATGCAAGCATGATAAGGCCGTGGCTGCGGAAAGTTCCG
GTTTACACCCTGCCGTCGCTGTGCTGCAATCTTGCCCCCAACAACAGTGAATTCGGTGACCATGCAAGCGTCGCAATTTTCAGCCCAGG
TTCTGGACTGGTACGATAAATACGGGCGAAAAACTCTGCCCTGGCAGGATCCTCTAGATTTAAGAAGGAGATATACATATG
>ydjX
GAGGCAATTCCGACGTCTAAGAAACCATTATTATCATGACATTAACCTATAAAAATAGGCGTATCACGAGGCCCTTTCGTCTTCACCTC
GAGTTGCGTAGAAACCGGCATCGACTATGAAATCCGCAGCATGGAAAAACCGTCCGATCACGCCCCCGTCTGGGCGACCTTCCGCCGCT
```

AATTTAGCAGCTCTCCTGGCTCAAACTGGGTCAGGAGAATTAACCTTGAGAAAAATCAACAAACTGTCAGTAATGATTTGTTGCCTGCC
GTCCTTTGTTATACCGTCTCTGCGTTTTTAGTTGTCTGACCACTTCTCTATTATCAAGTTTGATATAGGAAACTCCACGATGAACGCTG
AGCGTAAATTTCTTTTTGCCTGTCTTATTTTTGCGCGGATCCTCTAGATTTAAGAAGGAGATATACATATG
>ydjA
GAGGCAATTCCGACGTCTAAGAAACCATTATTATCATGACATTAACCTATAAAAATAGGCGTATCACGAGGCCCTTTCGTCTTCACCTC
GATGGATCCCGAAATTCAGCAGACGCCACGTCCATTTAAAAAATCCGGCAATAAATCGCCAAAGGGTTCGCATGTATTCTCCCAACTTA
AGGTCACACCTGTCACAGGCGCAATAGATGTGGTTATCGTAATGACCCAACCGGTACTTGTCAGCAGGAATTGCGGACTGTGCTGTAAC
AAAATCCACTCTCGTGTTAATTTTGTGAATAAATATCACGACAGGAGTTAATCAAATGGATGCACTCGAACTATTGATCAATCGCCGTA
GCGCCTCCCGCTTGGCTGAACCCGCGCCAACGGGTGAACAACTCTCGAGAGATCCTCTAGATTTAAGAAGGAGATATACATATG
>ydjE
GAGGCAATTCCGACGTCTAAGAAACCATTATTATCATGACATTAACCTATAAAAATAGGCGTATCACGAGGCCCTTTCGTCTTCACCTC
GATGGATCCTTTGGCCCGGTTTCATTATCGTATATTTGGTATTATAAGCTTTAGTCTGTTATTAACAGGGTTTTTGAGTTACTCCGGTA
ATGTCGTCTTAGCAAAGCTGGTAAGCAATGGATGGTCAAATAATTTCCTCAATGCCGCCTTTACCTCGGCATTAATGTTTGGTTATTTC
ATCGGCTCACTTACTGGTGGGTTTATTGGTGACTACTTTGGGCGGCGCAGGGCGTTTCGCATAAACTCGAGAGATCCTCTAGATTTAAG
AAGGAGATATACATATG
>b1420
GAGGCAATTCCGACGTCTAAGAAACCATTATTATCATGACATTAACCTATAAAAATAGGCGTATCACGAGGCCCTTTCGTCTTCACCTC
GATGGATCCCGCCTAGAATGCAGGCAAATAACGGAACCAACCTGATGTGATGCAAAAAACCTAATCGCTGTGAGGGAGTTGTATTCATT
ACCGTTGTCTCTCGTCCAGGTTTACTCCTGATAGAAAGTCATCGGCACTTGACGGTAATTACTTATGACCAAAAACCTCTTTGCGTCAC
ATTTTTACAACAATTATTGCATAGAAAATTAATGCTATTGGAATGAAGAGAGTAACGAATGGCTAAACTGGCTAACCCGAATGACGAAA
AATTCTGGAAAACAGCACGCAATTTCGTGAAGTGATTATTTTCGTCCCTAAGGAAACCTCATAGGATGCCTGCGGGTGCCTGAGACTTT
CTGTTTTCAGGCTAACCGGGAATCAGAAAGACGAAGGCCCCGACCGATATTTCTATCAACCGAGGCCAACGAATGAACCTAGCAATTCA
GATCTTAGCCTCTTACCCGCCATCCGGCAAGGAGAAAGGCTATGAAGCACAACCCTCTGGTGGCTCGAGAGATCCTCTAGATTTAAGAA
GGAGATATACATATG
>ydjF
GAGGCAATTCCGACGTCTAAGAAACCATTATTATCATGACATTAACCTATAAAAATAGGCGTATCACGAGGCCCTTTCGTCTTCACCTC
GATGGATCCTATCGGATGCAGACGCAACATTGATGAGGGAAATGGCAGAGGCCCTGGAGCGTTAAATACTCAACACGATTCACCTCTGT
TTCTGTGAAAACTGTGATGAAAATCTGTGATTCAAACAGGTTATTTTGAAAGTAAACATCGGTTATGCGATAATCGCGCTAATGTGATG
TAAAAAGATACAGGGGTGTAATCGTGGCGGCAAAAGACAGGATTCAGGCAATTAAGCAAATGGTTGCCAACGATAAAAAGGTGACAGTC
TCAAATTTGAGTGGGATTTTTCAGGTAACCGCTCGAGAGATCCTCTAGATTTAAGAAGGAGATATACATATG
>ydjL
GAGGCAATTCCGACGTCTAAGAAACCATTATTATCATGACATTAACCTATAAAAATAGGCGTATCACGAGGCCCTTTCGTCTTCACCTC
GATGGATCCCAGCAAAGCCTATTGCCATGTTTAAGTAAGTGGTTCATCCCCTTTTTGCCTCGGCTATGAAGGGGATAGCCAACAATGAC
CCCACGACTCATCTCAATTATCTCGCCATGACCTGGCTAAGTCCCAGTCCGATCTCTTGCCGTCCCGTTCACACCTGAATGACTTCTCC
TGAACTTAAGTCTCAGACCTATTTGGCCGGTAATCCCTCTCGAATTTGCTCTTTTTGCGCCCCGCCTTCCATCAACCTGTAACAGACCC
AGCCATTTTCTGCGTAATTGTGATCTGAGTTGTAAAACGCAGATCATTATCTGTGTTTTACAGATTTAATGTTGACCTCAAAAATCATG
TGGGTAAGTATGATGGCCATCTAAGCAAAACTAAACACAGAGGATGACGCAATGAAAGCACTGGCTCGGTTTGGCAAGGCCTTTGGCGG
CTACAAGATGATTGATGTCCCACAACCCATGTGTGCTCGAGAGATCCTCTAGATTTAAGAAGGAGATATACATATG
>ydjN
GAGGCAATTCCGACGTCTAAGAAACCATTATTATCATGACATTAACCTATAAAAATAGGCGTATCACGAGGCCCTTTCGTCTTCACCTC
GAGTTCGTTGGTCATCGCAAATGATCAATACCTTGCAGATCCAGTTTCATCGGCTTATTAAGCATCAGGTTGAATACTAAAAAGGCAAA
AATCACCTTTCTGGAATAAGCAATTCCATTTGAATATAAGAGCCAGCTCACAGTTCTGTTAATCTTGCGCCAACACTATGACTGCTACG
CAGTGATAGAAATAATAAGATCAGGAGAACGGGGATGAACTTTCCATTAATTGCGAACATCGTGGTGTTCGTTGTACTGCTGTTTGCGC
TGGCTCAGACCCGCCATAAAGGATCCTCTAGATTTAAGAAGGAGATATACATATG
>nadK
GAGGCAATTCCGACGTCTAAGAAACCATTATTATCATGACATTAACCTATAAAAATAGGCGTATCACGAGGCCCTTTCGTCTTCACCTC
GAGGCTTCTGGCTCAACTGCCTCAATCTCTTCGTGCTGATCCATGATAATTTCTTCCGGGGCTTGCCCCTCAGGCGTTTTCTGTTCTTT
ACTACTCATGAATTTCTCCGCGTTTTTTTCGCATTCATCTCGCTAACTTCGCTTATTATGGGGATCAGTTTCAGGGTTTCAAGGGAAGC
ACTCACATTGTCATCAATCTTCGCAACAAGGACCTCGGAAAAATGAATAATCATTTCAAGTGTATTGGCATTGTGGGACACCCACGGCA
CCCCACTGCACTGACAACACATGAAATGCTCTACCGCTGGCTGGGATCCTCTAGATTTAAGAAGGAGATATACATATG
>yadK
GAGGCAATTCCGACGTCTAAGAAACCATTATTATCATGACATTAACCTATAAAAATAGGCGTATCACGAGGCCCTTTCGTCTTCACCTC
GATGGATCCTTTTCTGCACCAGCCACATTTACAGTAACCTATAACTAACAGACTTACAGATTCAGGAACTATATATGCATCCCACTCAA
CGTAAGCTGATGAAGAGAATAATTCTGTTTCTGTCATTACTGTTTTGCATCGCCTGTCCCTCGAGAGATCCTCTAGATTTAAGAAGGAG
ATATACATATG
>cusR
GAGGCAATTCCGACGTCTAAGAAACCATTATTATCATGACATTAACCTATAAAAATAGGCGTATCACGAGGCCCTTTCGTCTTCACCTC
GATGGATCCACAACCGGTTAGCGCAAGGGCCACACAAAATGGCAGAAGTTTACAAGGAGACATAGGCTCATAATTTCTGGTGATTTTAT
GCCGCCAACTTTACTCGCCAGGCTCTGATTTTCCGGTGACAGAAAAATGACAAAATTGTCATTTTGCCAATAAGCGATTGCCATCTGAT
CCCGCTACTCTAGAATTGCCCGGGCAACATGCGGAGGAAATATGAAACTGTTGATTGTCGAAGATGAAAAGAAAACCGGAGAATACTTG
ACCAAAGGGTTAACCGAAGCCGGTTTTGTGGTCGATTTGGCCGACAACCTCGAGAGATCCTCTAGATTTAAGAAGGAGATATACATATG
>yadN

```
GAGGCAATTCCGACGTCTAAGAAACCATTATTATCATGACATTAACCTATAAAAATAGGCGTATCACGAGGCCCTTTCGTCTTCACCTC
GATGGATCCTGGTGTTTCCTGATGGGGAGATGTTGCGTCAAATCTTACATACAAGAGCATTTGACAAATTAAACAAATGGTAATTTATT
TATATTGCGAATATAAATATTTTCTGAAAATCATATCTCATCTCTTTCCTTAATTACGATTTAGTTTAAACATCTCTTTGCGGAAATA
ATATAACTTCCTATAAATAAATATCAATCCCATCTCCATACCCTAACCAATTGCACTTGTGAACTTATTTTTACTTAATTTTGAATTAA
GTAAATTTAATTTGTTCAAAAGCTTCTAAAGTGCACAATAGGCGACCTAATGAACAATGGTTGTAAATTTGGGTAACAAATATCATTGC
TCATTAATATCTAACATTTGTTTCTCCTATTTGTGTCATGAATAGTCATTCTTCAAACATGTAGTCTGACTAATGGAATTTTATTAAAG
GATTTTTATGTCTAAAAAATTAGGTTTTGCCCTGTCTGGTCTTATGCTGGCGATGGTTGCAGGTACTGCTTCTGCTGATATGGACGGCG
GTCAGCTCGAGAGATCCTCTAGATTTAAGAAGGAGATATACATATG
>nfo
GAGGCAATTCCGACGTCTAAGAAACCATTATTATCATGACATTAACCTATAAAAATAGGCGTATCACGAGGCCCTTTCGTCTTCACCTC
GAGCTCTGCTGATGGCACTGGTACTGTTTGCCTGGCTGATTGTTGGTGGTGGTGCGATTAACTATGTGATTCAAAGCGTCATCGCATAA
ACCACTACATCTTGCTCCTGTTAACCCGCTATCATTACCGTTTTCCTCCAGCGGGTTTAACAGGAGTCCTCGCATGAAATACATTGGAG
CGCACGTTAGTGCTGCTGGCGGTCTGGCAAATGCCGCAATTCGCGCCGCCGAAATCGACGCAACCGCGTTTGCCTTGTTCACCGGATCC
TCTAGATTTAAGAAGGAGATATACATATG
>yadC
GAGGCAATTCCGACGTCTAAGAAACCATTATTATCATGACATTAACCTATAAAAATAGGCGTATCACGAGGCCCTTTCGTCTTCACCTC
GATGGATCCAGCACCCGAATGCAAAAAATCGATCCTGCATTGAGTGTTACATCTGGTCAACTTATGAGCCATGTGTTAGTGGATATTTA
CTACGAATAACAAACAAAAACAGCGCAAACCATAACAATAAAGAAACATTGGAGTAAAAATGAAGACTATCTTCAGGTACATTCTTTTT
TTAGCACTGTATTCTTGTTGTAATACAGTCAGTGCATATACAAGTTTTATTGTGGGAAATAATGCAGGACTCGAGAGATCCTCTAGATT
TAAGAAGGAGATATACATATG
>yfjI
GAGGCAATTCCGACGTCTAAGAAACCATTATTATCATGACATTAACCTATAAAAATAGGCGTATCACGAGGCCCTTTCGTCTTCACCTC
GAGTGCCACCTTTCCCAAAAAGCGAATGCTCGGCGTGAAATCTGTCGGATGGATTGAGGCCGAGATTGATGAGTGGTTATCACAACGCT
GTAAACTTATTTGAGGGTGTTACATGAATTCATACTCAATTGCTGTCATCGGAGTGTAAACATCCTGCTATTTCCGCCATTAATAGCGG
AATAGCCATATTTGCTAAACAATTTCTTAATTTAATTAAATAAGAGGCATTGCATGTTCAATGGTCGTCCTTTCCCTGTAGATGCATTT
CCTAAAATTATCAGGAATGCAATTTATGAAGTGGAACAGCATACGCAGGCGGATCCTCTAGATTTAAGAAGGAGATATACATATG
>yadF
GAGGCAATTCCGACGTCTAAGAAACCATTATTATCATGACATTAACCTATAAAAATAGGCGTATCACGAGGCCCTTTCGTCTTCACCTC
GATAGATCTGCCGCCTGGATAGGTTTTTTTAAGCTGTTGAAGTTCCAGTGCAATGGTCATAAATTTTTACTTACCTTACGTTCTTACAC
TTTATCTGTGGTTTAAATCGTCCCGGAGTTGCCCTATATTAGCCAAACGTAATTATTTGGTTACAGGTCGTTAACCTCCATGAAAGACA
TAGATACACTCATCAGCAACAATGCACTATGGTCAAAAATGCTGGTGGAAGAGGATCCCGGGTTTTTTGAGAAACTGGCACAAGCGCCT
CGAGAGATCCTCTAGATTTAAGAAGGAGATATACATATG
>yfjN
GAGGCAATTCCGACGTCTAAGAAACCATTATTATCATGACATTAACCTATAAAAATAGGCGTATCACGAGGCCCTTTCGTCTTCACCTC
GAGTTTCTTGTCTGCTGGCGAATAATTACTAAGTATCGAGTTGCTGCTTGGTGAGATCAGGAAGAGGTTACCGAAACGATGGACATCAT
CCCCCATTTTGTGGACTGCCCCCCCGGTAGACGATCCTGCCCTATAGTTGGAGTGCCCCCTATTCAGGTGGCTAAATTTAGCAAATCAA
TACACTTCAGGGGGGTATTATTGTAGAGTTTCCCCATATGTTTCTATGGGATCCAGGAAATGACAATCAGGAGTTACAAAAACTTAAAT
CTGGTCAGGGCAAATATCGAGACTGAATCCAGACAATTCATTGAAAATAAAAACTATTCAATCCAATCAATTGGTCCTATGCCAGGGTC
AAGGGCTGGGCTTCGGGTCGTATTTAGATCTAAATACGACCCGAAGCCCGATGGGCAAGGCATCAACCTCGAGGGGATCCTCTAGATTT
AAGAAGGAGATATACATATG
>yfjM
GAGGCAATTCCGACGTCTAAGAAACCATTATTATCATGACATTAACCTATAAAAATAGGCGTATCACGAGGCCCTTTCGTCTTCACCTC
GATGGATCCTCGATATTTGCCCTGACCAGATTTAAGTTTTTGTAACTCCTGATTGTCATTTCCTGGATCCCATAGAAACATATGGGGAA
ACTCTACAATAATACCCCCCTGAAGTGTATTGATTTGCTAAATTTAGCCACCTGAATAGGGGGCACTCCAACTATAGGGCAGGATCGTC
TACCGGGGGGGCAGTCCACAAAATGGGGGATGATGTCCATCGTTTCGGTAACCTCTTCCTGATCTCACCAAGCAGCAACTCGATACTTA
GTAATTATTCGCCAGCAGACAAGAAAAAATTCTACGTCGAAACAGAACGAGCAGAGAGCCCAAAACAAGCCATCATGATGAGCTATAAG
GAGTGGGGTCCGGATGGGCAAGGCATCAACCTCGAGAGATCCTCTAGATTTAAGAAGGAGATATACATATG
>yfjL
GAGGCAATTCCGACGTCTAAGAAACCATTATTATCATGACATTAACCTATAAAAATAGGCGTATCACGAGGCCCTTTCGTCTTCACCTC
GATGGATCCGATGGGCAAGGCATCAACAATATAGAAAGCCATGAACACGCGATGTTGACATTGCTCAAAGAGCATCGAGATATGACCCT
ACCAACCAGGAAATAATGCGTGCAACTGTGTTTTATAATATCAAGTGTGATTACAATCGCTGGCGGTGCCATAGTGCCATAGTGCCATA
GTGCCATAGTGCCATAGTGCCATAGTGCGGTTGGCGGTCTCAGCTCTGAACTATTTGAAAACCAAAACCTTGATTAAGGCTGTGCCTAC
CTTACGTTAGTTGGGGCAGTATGTTATATAACCAAATTAGACGTCAGGGTTACAATATTTGTGGAATCCTGAAAAGCTTGATACCTGCG
ACCGAGAATGGCTGTGAAGTATTTAGATCACTGGTGATGATGCAGTGATCCGATTGGACTGTGTCATAACATCTCCTTAATTTTCGCAA
AATTGTCACGGATGTCTAAATGGAGTCGAACGATTCAGGCGGAGTAGCCGCCAAGCACGGTTTCTTATTTCAGGACTGCGTAGCCGCTT
ACCATGTCACCCTCGAGAGATCCTCTAGATTTAAGAAGGAGATATACATATG
>yfjR
GAGGCAATTCCGACGTCTAAGAAACCATTATTATCATGACATTAACCTATAAAAATAGGCGTATCACGAGGCCCTTTCGTCTTCACCTC
GAGTTGACGGGGATGTGAAGCTTAACCGTGCGCTGTGGGTGATGGCAGAAACACTGCTCACGCAACTGCAGTAGACGTTTCATGTTGCC
ACGTTGTTAATATCGGACACCACCTGTCCGCATCGCTATGTGCTCGTGTGCCTCAATCCCCCGGTTATAGCTTTTAACCCCCGTTACAT
CTGGCTTTTGCAGAAATAAAAAATAGTTTCTGCGTTGTCCATACCCTGTCCGCCCCCCTCTTTAAAGTAATCACATCATTTTCAGTCAG
TTAACTTTCCTGGAGAATCTCTCATGACACAGGCAGAACGCCGCCATGACCGGCTGGCTGTCAGGCTGTCACTGATAATCAGCCGTCTG
GTTGCGGGTGAAACGTTGAGCGTGGATCCTCTAGATTTAAGAAGGAGATATACATATG
```

Data Set S4 - MG1655 library reference file.fasta

```
>cusC
GAGGCAATTCCGACGTCTAAGAAACCATTATTATCATGACATTAACCTATAAAAATAGGCGTATCACGAGGCCCTTTCGTCTTCACCTC
GAGGTTGTCGGCCAAATCGACCACAAAACCGGCTTCGGTTAACCCTTTGGTCAAGTATTCTCCGGTTTTCTTTTCATCTTCGACAATCA
ACAGTTTCATATTTCCTCCGCATGTTGCCCGGGCAATTCTAGAGTAGCGGGATCAGATGGCAATCGCTTATTGGCAAAATGACAATTTT
GTCATTTTTCTGTCACCGGAAAATCAGAGCCTGGCGAGTAAAGTTGGCGGCATAAAATCACCAGAAATTATGAGCCTATGTCTCCTTGT
AAACTTCTGCCATTTTGTGTGGCCCTTGCGCTAACCGGTTGTTCACTGGCACCGGATTATCAGCGTCCGGCAATGGATCCTCTAGATTT
AAGAAGGAGATATACATATG
>yfjP
GAGGCAATTCCGACGTCTAAGAAACCATTATTATCATGACATTAACCTATAAAAATAGGCGTATCACGAGGCCCTTTCGTCTTCACCTC
GAGCGAAACTTTCTGCCCTACTGGGTAGTGAGACCACTGGTGAAATCCTTTTTGACCTCTTGTGCGCTAACGGTCCAGAGTGGAACCGC
TTCGTTACGCTGGAAATGAAATATGGCCGGATCATGCTGGATACGGCCAAAATTATTGATGAGCAGGACGTTCCGACACATATCCTGAG
CAAACTGACTTTTACTTTACGTAATCATCCTGAGTACCTAGAAGCCTCTGTTTTGTCTCCCGATGATGTACGCCAAGTCCTGTCAATGG
ATTTTTGAAACTCTTCTTTCTAAACGAACCAGTATAGAATTCGCCCTTTTGGCATGATATTAACTTGTCCAATATGATCAAATAGCATT
AACCCCCCCTCACAACGTCCTGCATAGGGAACACGTTTTCCCCTGTGCACCCACGACTAAATTTCCCCCCCTTTAAACTTCTCAATAAT
GTCACCAATTTTCTGGAAATCCTATGATGCCTTTCCCGGTAAATGGTGCTGTGAGTTCAGAAAGATTCTGCAATCTATGTGCATTGAAA
AAATCTTCCTTGCGGACTGGTTCAGTGAGCAATTCTCAACGGGCGTGTTGATCGTGCCTTGTATGCTGACGTTGGCTATTCCCTCGAGA
GATCCTCTAGATTTAAGAAGGAGATATACATATG
>yfjW
GAGGCAATTCCGACGTCTAAGAAACCATTATTATCATGACATTAACCTATAAAAATAGGCGTATCACGAGGCCCTTTCGTCTTCACCTC
GAGATCCTTGATCGCACTGGCAGGCGTCTTCAGCAGCGAAACTTAATACGTTGCCGGAATAACTCTGCGAAAGAAGAGATACAGCATAA
TCAGCGTGGCCGCAATCGCTGCCGCATCCACGGAGATCATAACGGAGGCGTACTGCGCAAAGCTCAGACCGAAGAAGTCCGCCGAGACG
ATATTCACCAGGTTAGAAACAATGAGCGGCAGGCTGGCCGTATCTGCAATAAATCCTGCAGCCATGACAAAGGCCAGTGTCGTGCCCTG
GCTGAACCCAGTGCGATCAGCATCGCAATCACAATCGGCGTCAGGATCAGCGCGGCGCCGTCATTGGCAAACAACGCATCAACAGCGGC
ACCGAGCAAGACTTTCCAGGTGAACAGCGCTCTGGATCCTCTAGATTTAAGAAGGAGATATACATATG
>yadR
GAGGCAATTCCGACGTCTAAGAAACCATTATTATCATGACATTAACCTATAAAAATAGGCGTATCACGAGGCCCTTTCGTCTTCACCTC
GAGACGCTGGCAAAACAGGAAGCTGAGCAACTGGCGCGAAGCAAGGCCGCATCAGCCAGCGAGAATACTTGAACGAAATACCAGGGTAT
TAGATAATGGCGATTATTATTGGGTTAGAATTTGCCCAATTGCCGATGTCGTTTGGAGCAAAATATGAGTGATGACGTAGCACTGCCGC
TGGAGTTTACCGACGCAGCAGCCAACAAAGTTAAAAGCCTGATCGCTGACGGGATCCTCTAGATTTAAGAAGGAGATATACATATG
>yadS
GAGGCAATTCCGACGTCTAAGAAACCATTATTATCATGACATTAACCTATAAAAATAGGCGTATCACGAGGCCCTTTCGTCTTCACCTC
GATGGATCCACGTGCAAGCCCACGTATTATCCTCGCTGCACAACAGCTCTGTAATGCGCTTTCACAGGTAGATTAGCGCCTGACCTTTT
CAGGTGCTCAGTGGGGATTGAACAATGCTCGTCTATTGGCTGGATATAGTCGGCACAGCGGTATTTGCCATCTCCGGCGTTTTGTTAGC
CGGAAAATTGCGTATGCTCGAGAGATCCTCTAGATTTAAGAAGGAGATATACATATG
>psiE
GAGGCAATTCCGACGTCTAAGAAACCATTATTATCATGACATTAACCTATAAAAATAGGCGTATCACGAGGCCCTTTCGTCTTCACGTC
GACGGTGGTCAGCAACTTGGGCGTAAGTTCCAGTTGTATGACATGACCAGCGACCGTAGCGTCAATTTCCGCTAAGTCATGGGAAAGGT
GCCAGTTTTCGCACTCACTGGCACCTTCATTCTTAATGGCCTGTCGAGCCATCAGGAACGCAGTTCACTCCCGGTTTGCGATCTGGTGT
GTAAAGGCGAAAACATTCTGTCTTCCTGTCCGTCGATATCGGTTGCAGGTCTGGTTGCCTGAGCGACTGGGCCATCATTGGGCTGGAGA
CATTGCTTTGCGGTAGGTCGGCTTTATCAGCAGTACCGAGCGGACCAGCATAAGCAGGAAGAACAGAGACTGATAACATCAAAGCAGCA
AAATAAGGCTTCATTTTTACCACCTTTATCAGGTTACGTTTCATTTGTTCCAGAGGAACATTGTCGATTTTTCGCGCATTGCTGGTGGC
TGGGAATCACCTGAATGGGTGATTTTTGAATTACCGGCTTTGGTGCGGTTTGTCTTGCCGGATGCGCCGCCAGGCGCGGCTTATCCGGC
CTACGGGTAGGTATATCCGGCTTTGGTGGAGGCGCGCTCCAAATCCAGGTTGAACAAAACATACACAAAAAATATAGATCTCCGTCACA
TTTTTGCGTTATACAGGAAGCTCGCCACTGTGAAGGAGGTACTGCTATGACGTCACTCTCTCGTCCGCGCGTGGAGTTTATCTCCACCA
TTTTGCAGACCGTACTCAATCTTGGCCTGCTGTGCCTCGGCCTGATTTTGGTTGTGGATCCTCTAGATTTAAGAAGGAGATATACATAT
G
>psiF
GAGGCAATTCCGACGTCTAAGAAACCATTATTATCATGACATTAACCTATAAAAATAGGCGTATCACGAGGCCCTTTCGTCTTCACCTC
GAGCCGCCAATGTTGTTGGACTGACCGACCAGACCGATCTCTTCTACACCATGAAAGCCGCTCTGGGGCTGAAATAAAACCGCGCCCGG
CAGTGAATTTTCGCTGCCGGGTGGTTTTTTTGCTGTTAGCAACCAGACTTAATGGCAGATCACGGGCGCATACGCTCATGGTTAAAACA
TGAAGAGGGATGGTGCTATGAAAATAACATTACTGGTTACCTTGCTTTTCGGTCTGGTTTTTTTAACCACCGTCGGCGGATCCTCTAGA
TTTAAGAAGGAGATATACATATG
>rplY
GAGGCAATTCCGACGTCTAAGAAACCATTATTATCATGACATTAACCTATAAAAATAGGCGTATCACGAGGCCCTTTCGTCTTCACCTC
GAGTACTGGCAGGTGCGTGAAAAAGTGTTCGATTACGAAGGTCGTTTTCGTCTGGCGCACGAATTACGCGGTTAATGGTAGTTTTGATT
GATGTATAAGGCGATTGAGTATAGAATCTCGCCCGCTTTTGCATACGCAAAGCAGATCACTTACCTGTTGCTGGGTCGCCTGTAGCAGG
ATTAATTTAAGAGAGAAAGAAATGTTTACTATCAACGCAGAAGTACGTAAAGAGCAGGGTAAGGGTGCGAGCCGCCGCCTGCGTGCCGC
TAACAAGTTCCCGGCAATCAGGATCCTCTAGATTTAAGAAGGAGATATACATATG
>rplT
GAGGCAATTCCGACGTCTAAGAAACCATTATTATCATGACATTAACCTATAAAAATAGGCGTATCACGAGGCCCTTTCGTCTTCACCTC
GATGGATCCTTTCCAAAGGCGATCTGGGCCTGGTAATCGCGTGCCTGCCGTACGCATAAGCCGTTAACGTTTTTAACTTTTTAATTAGA
ATATAGATACAGGAGAGCACATATGGCTCGCGTAAAACGTGGTGTTATTGCACGTGCACGTCACAAGAAAATTTTGAAACAAGCTAAAG
GCTACTACGGTGCGCGTTCTCCTCGAGAGATCCTCTAGATTTAAGAAGGAGATATACATATG
```

Data Set S4 - MG1655 library reference file.fasta

```
>hlyE
GAGGCAATTCCGACGTCTAAGAAACCATTATTATCATGACATTAACCTATAAAAATAGGCGTATCACGAGGCCCTTTCGTCTTCACCTC
GATGGATCCAATCTGCTGCCGGTGAAGGAAAGCCACACTGAACAAGATCGCTAAATAGCGGAAAAGTCACAATTTCGCGGAGATCCGCA
GGCTTGATAAACAACATAATAATCTGCCTGAAGTTATACTGTTTTTATATACAGTAGTCTGTTCTTGCCAGCAGATCAATACTGATTCA
GGCTATCAATATTTGTCGCTGCATAGGCTGCTGATTTTTCGTTCTCTTATCTTGTGCTCACGTGGCCTTCTGGCGACGACGCTCATCCA
GCAGAAATGAAAAATATCACCCGGCTAAAAAATAGAATAGAAGCATTCGCCATAATGACATTAAACATTGTTTGATATTTATCATATTA
ATAGAAATAAAGACATTGACGCATCCCGCCCGGCTAACTATGAATTAGATGAAGTAAAATTTATTAATAGTTGTAAAACAGGAGTTTCA
TTACAATTTATATATTTAAAGAGGCGAATGATTATGACTGAAATCGTTGCAGATAAAACGGTAGAAGTAGTTAAAAACGCAATCGAAAC
CGCAGATGGACTCGAGAGATCCTCTAGATTTAAGAAGGAGATATACATATG
>rplN
GAGGCAATTCCGACGTCTAAGAAACCATTATTATCATGACATTAACCTATAAAAATAGGCGTATCACGAGGCCCTTTCGTCTTCACCTC
GATGGATCCCGTCCGCTGTCCAAGACTAAATCCTGGACGCTGGTTCGCGTTGTAGAGAAAGCGGTTCTGTAATACAGTACACTCTCTCA
ATACGAATAAACGGCTCAGAAATGAGCCGTTTATTTTTTCTACCCATATCCTTGAAGCGGTGTTATAATGCCGCGCCCTCGATATGGGG
ATTTTTAACGACCTGATTTTCGGGTCTCAGTAGTAGTTGACATTAGCGGAGCACTAAAATGATCCAAGAACAGACTATGCTGAACGTCG
CCGACAACTCCGGTGCACGTCGCGTAATGTGTATCAAGGTTCTGGGTGGCTCTCGAGAGATCCTCTAGATTTAAGAAGGAGATATACAT
ATG
>kdsB
GAGGCAATTCCGACGTCTAAGAAACCATTATTATCATGACATTAACCTATAAAAATAGGCGTATCACGAGGCCCTTTCGTCTTCACCTC
GAGGGACAACCTCGCTTTCCCGCTGCGTGATGGCATTCCAGTGTTGCTGGAAACGGAAGCCCGCGTGCTGACTGCTGATGAGAGTAAAT
CATGAGTTTTGTGGTCATTATTCCCGCGCGCTACGCGTCCACGCGTCTGCCCGGTAAACCATTGGGATCCTCTAGATTTAAGAAGGAGA
TATACATATG
>metZ
GAGGCAATTCCGACGTCTAAGAAACCATTATTATCATGACATTAACCTATAAAAATAGGCGTATCACGAGGCCCTTTCGTCTTCACCTC
GAGTGGTTTGGAAGAGCAGGCGGCAAGCATTGCCACAACCGTGCCCATAAGAAGGTACTTTACCCAACGTCCTTTCATTGTTCTCTTCT
TCAGGTTAAAAAATAAGGCGCAACGAAGATAACAAACCGCCGCGTTTAATGAAATGAGCAGCCTCTCCCTGACGCAAATTTTGCACAAA
AAATAGGCTTTAGTGATTTGTTTTTGTTCAAAATCATGCCAAATCCGTGATCGGGGTAAAAAAAAGGTTGCATGAAAACGCGAGCGGAG
TATAGTGCGCATCCACGGACGCGGGGTGGAGCAGCCTGGTAGCTCGTCGGGCTCATAACCCGAAGGTCGTCGGTTCAAATCCGAGATCT
TCTAGATTTAAGAAGGAGATATACATATG
>metB
GAGGCAATTCCGACGTCTAAGAAACCATTATTATCATGACATTAACCTATAAAAATAGGCGTATCACGAGGCCCTTTCGTCTTCACCTC
GAGCCGGGTTAATCCCCATCTCACGCATGATCTCTTTTGCCGCTTCCGGGATTTCGTCGCTGCGCTCTTTACGCAGATCGGCATCATCC
GGCAAAGGTTGCCCGGTAAAGGCATGCAGAAACGCTTCGCACAGCAGCTCGCTGTTGGTAGCGTGACGCAGGTTGTTCACCTGACGACG
CGTGCGTTCATCGGTGAGGATTTTTAACACCTTAAGAGGAATGGAAACCGTAATCTTTTTGACTTGTTCACTCTTCTTGCCGTGCTCAG
CGTATGGGCTGATATATTCGCCGCTCCATTCAGCCATGAGATACTTAATCCTCTTCGTCAATAAATTGAGACCAGACCACACAGTTGAT
GTGGGTACTGACCGTAAACCCGCATAGTTTACCGTACAGGCGTTACCGGGATCCTCTAGATTTAAGAAGGAGATATACATATG
>metC
GAGGCAATTCCGACGTCTAAGAAACCATTATTATCATGACATTAACCTATAAAAATAGGCGTATCACGAGGCCCTTTCGTCTTCACCTC
GAGGAAGAAGATTGCCCAGGTGACTACGGAGGCCAAAATAAGCCCAATCATCACGCACTTAACGACAATATCGGCGTGCTGATACATAC
CCCAGACGGAAAGGTCCGTCTGCATTAAATTATTACCCACTGTGTATCTCCAGGACGCAAGTCACAAAATCTGCGCATAATAATATCAA
AACGACGTCGAATTGATAGTCGTTCTCATTACTATTTGCATACTGCCGTACCTTTGCTTTCTTTTCCTTGCGTTTACGCAGTAAAAAAG
TCACCAGCACGCCAGATCTTCTAGATTTAAGAAGGAGATATACATATG
>mrcB
GAGGCAATTCCGACGTCTAAGAAACCATTATTATCATGACATTAACCTATAAAAATAGGCGTATCACGAGGCCCTTTCGTCTTCACCTC
GAGTGAAAGGGCGTTATCCCAAACATGTCTGGCCGGACGACCCGGCAAATACTGCACCGACGCGACGGACGAAAAAGTATTCGTAAGGT
TGGTTTTCTCCCTCTCCCTGTGGGAGAGGGCCGGGGTGAGGGCATCAGCGCGCACGTTCACCCTCACCCTAACCCTCTCCCTCAAGGGA
GAGGGGACCGATCGAGCACAAATTTTGAGAGATATCTTCTTCTGTCTTGTAACAGAAGAACAGAAAATCGGGCTTTTGCGCCTGAATAT
TGCGGAGAAAAAGCATGGCCGGGAATGACCGCGAGCCAATTGGACGCAAAGGGAAACCGACGCGTCCGGTCAAACAAAAGGTAAGCCGT
CGTCGTTACGAAGGATCCTCTAGATTTAAGAAGGAGATATACATATG
>metA
GAGGCAATTCCGACGTCTAAGAAACCATTATTATCATGACATTAACCTATAAAAATAGGCGTATCACGAGGCCCTTTCGTCTTCACCTC
GAGCGTGAGTGGCATCGACAGAACGACACCAAATCGCAACGAGTTCCTCCCCTTCCTCATGCCGTGAGCGGCGAATACTAATAACCATT
TTCTCTCCTTTTAGTCATTCTTATATTCTAACGTAGTCTTTTCCTTGAAACTTTCTCACCTTCAACATGCAGGCTCGACATTGGCAAAT
TTTCTGGTTATCTTCAGCTATCTGGATGTCTAAACGTATAAGCGTATGTAGTGAGGTAATCAGGTTATGCCGATTCGTGTGCCGGACGA
GCTACCCGCCGTCAATTTCTTGCGTGAAGAAAACGTCTTTGTGATGACAACTTCTCGTGCGTCTGGTCAGGATCCTCTAGATTTAAGAA
GGAGATATACATATG
>metF
GAGGCAATTCCGACGTCTAAGAAACCATTATTATCATGACATTAACCTATAAAAATAGGCGTATCACGAGGCCCTTTCGTCTTCACCTC
GAGCGCGATAACCCTCTGGTGATCCGCGGACCTGGCGCTGGGCGCGACGTCACCGCCGGGGCGATTCAGTCGGATATCAACCGGCTGGC
ACAGTTGTTGTAATTTCAGAAATTTAATAATGCCCGGTACTCATGTTTTCGGGTTTATGGTTTCTAATGAAATATATTGAATTATCATA
GGATTAGGCCGGATTAAGCGTTTACGACGAATCCGGCAAGAAGCAATAAGTACATGGTTAGTTTATATTTGCAGTCCGGTTTGCTTTGC
ATACCGGATTTTCTTTTTCTTACCATCCTGAAGTTTTTTCATCTTCCCTGATTTTTCCTCACCATCATTGGTCATTTTTCGGTTGACGC
CCTTCGGCTTTTCCTTCATCTTTACATCTGGACGTCTAAACGGATAGATGTGCACAACACAACATATAACTACAAGCGATTGATGAGGT
```

AAGGTATGAGCTTTTTCACGCCAGCCAGCGGGATGCCCTGAATCAGAGCCTGGCAGAAGTCCAGGGGCAGATTAACGGGATCCTCTAG
ATTTAAGAAGGAGATATACATATG
>metG
GAGGCAATTCCGACGTCTAAGAAACCATTATTATCATGACATTAACCTATAAAAATAGGCGTATCACGAGGCCCTTTCGTCTTCACCTC
GAGCGCAAGGCTTCTGGTGATTTGGCCTGGGATTGTTCGTTCATCCCGACTCCTTTTTTGTATAGATAAACCATCAGCTGATAGTTTAC
CTGAAGAATATAGAGAAGTACTTACTTAACATTTTCCCATTTGGTACTATCTAACCCCTTTTCACTATTAAGAAGTAATGCCTACTATG
ACTCAAGTCGCGAAGAAAATTCTGGTGACGTGCGCACTGCCGTACGCTAACGGCTCAATCCACCTCGGCCATAGGATCCTCTAGATTTA
AGAAGGAGATATACATATG
>metJ
GAGGCAATTCCGACGTCTAAGAAACCATTATTATCATGACATTAACCTATAAAAATAGGCGTATCACGAGGCCCTTTCGTCTTCACCTC
GATGGATCCCGGTAACGCCTGTACGGTAAACTATGCGGGTTTACGGTCAGTACCCACATCAACTGTGTGGTCTGGTCTCAATTTATTGA
CGAAGAGGATTAAGTATCTCATGGCTGAATGGAGCGGCGAATATATCAGCCCATACGCTGAGCACGGCAAGAAGAGTGAACAAGTCAAA
AAGATTACGGTTTCCATTCCTCTTAAGGTGTTAAAAATCCTCACCGATGAACGCACGCGTCGTCAGGTGAACAACCTGCGTCACGCTAC
CAACAGCGAGCTGCTGTGCGAAGCGTTTCTGCATGCCTTTACCGGGCAACCTTTGCCGGATGATGCCGATCTGCGTAAAGAGCGCAGCG
ACGAAATCCCGGAAGCGGCAAAAGAGATCATGCGTGAGATGGGGATTAACCCGGCTCGAGAGATCCTCTAGATTTAAGAAGGAGATATA
CATATG
>metK
GAGGCAATTCCGACGTCTAAGAAACCATTATTATCATGACATTAACCTATAAAAATAGGCGTATCACGAGGCCCTTTCGTCTTCACCTC
GAGGCCATCCAGATGTTAATCCATCCATACCGATTAACACTCAGACTGCCAGTGTTTTTAACCTGCAGAGTCGTGGTAGGATCCGCTAC
CACAGAAAATCCACACAACAGTTTGAGCTAACCAAATTCTCTTTAGGTGATATTAAATATGGCAAAACACCTTTTTACGTCCGAGTCCG
TCTCTGAAGGGCATCCTGACAAAATTGCTGACCAAATTTCTGATGCGTTTTAGACGCGAGATCCTCTAGATTTAAGAAGGAGATATACA
TATG
>yhbE
GAGGCAATTCCGACGTCTAAGAAACCATTATTATCATGACATTAACCTATAAAAATAGGCGTATCACGAGGCCCTTTCGTCTTCACCTC
GATGGATCCAGGCCCGAAAAACCGTAAATTTATCAGCATCGAAGCTGAATAAGTTTTTCGCGTCCCGGTAACGGATGAAAGCCCCGCAA
CACGTTGCGGGGCTTTTTACATTCGATAACCGGTAAAAATGCGCATGGTGCGACGTTAATCCGCAAATGCGCTTTGCAGGGAATACGGT
ATGAAGCAGCAGGCAGGCATTGGCATTCTTTTGGCGCTCACCACAGCAATTTGCTGGGGGGCGTTGCCAATCGCAATGAAGCAGGTGCT
GGAGGTGATGGCTCGAGAGATCCTCTAGATTTAAGAAGGAGATATACATATG
>ycjW
GAGGCAATTCCGACGTCTAAGAAACCATTATTATCATGACATTAACCTATAAAAATAGGCGTATCACGAGGCCCTTTCGTCTTCACCTC
GATGGATCCTCTGTCGACACCCCGATTCACCAGCGCATTGAGTTCATTTTTAAGTCGCTTCATTGCTGTCCTGTGCTGCTCTGAAGACC
AGAAAAAAAAATCATTCGTAAGGATATGAATGTGGGGGCCAGGCCATTGATTATCAACTGCCCGTGTTGTCATCGACCGGCGAACTCCAT
AAGATGTGCAGCATTCACGTTCGGACGGAGTTCCTGACCTATGTCCCCTACTATTTATGATATTGCCAGGGTTGCAGGCGTATCAAAAT
CCACCGTATCACGCGTGCTGAATAAGCAAACTAATATCTCCCCGGAAGCGCTCGAGAGATCCTCTAGATTTAAGAAGGAGATATACATA
TG
>yhbC
GAGGCAATTCCGACGTCTAAGAAACCATTATTATCATGACATTAACCTATAAAAATAGGCGTATCACGAGGCCCTTTCGTCTTCACCTC
GATGGATCCGAGCAGCCTGGTAGCTCGTCGGGCTCATAACCCGAAGATCGTCGGTTCAAATCCGGCCCCCGCAACCACTTTCCCTTAGA
GTCCTTTTTCAAATATACTGTGAAGACTTCGGCCTTCGTAGTGGGATTTGAAAAAATCCTTCTGGAAAGTGCTCCAGACCGCAGTTGCG
GTTATAGGGTTCAGTTATATAAAGCCCCGATTTATCGGGGTTTTTTGTTATCTGACTACAGAATAACTGGGCTTTAGGCCCTTTTTTTA
TGTCTTGGGGGTGGGCTTGTCCACATTAGAGCAAAAATTAACAGAGATGATTACTGCGCCAGTTGAGGCCCTGGGTTTTCTCGAGAGAT
CCTCTAGATTTAAGAAGGAGATATACATATG
>ycjZ
GAGGCAATTCCGACGTCTAAGAAACCATTATTATCATGACATTAACCTATAAAAATAGGCGTATCACGAGGCCCTTTCGTCTTCACCTC
GAGGCAGACAAAGAGATGGTTGGATTATTGCTGTTAGTGAAGCTGACTTTATTATTCATCATTCCGTTCCTCATGGAGTTGTCGGTTCG
TTTTAACGGTTGGTGATATCACTATAGATATTGATCATTAAGTTGATTAGACCCAAAATCATGATTAGACCTATCATTTAAATGATTAA
TAGATTGCATGGAGATACTGAATGAAGCGTGAAGAAATCGCTGATCTGATGGCGTTTGTCGTCGTTGCAGAGGAGCGTAGCTTCACTCG
TGCAGCAGCCCGCCTGAGCATGGCGCAGTCAGGGATCCTCTAGATTTAAGAAGGAGATATACATATG
>ycjX
GAGGCAATTCCGACGTCTAAGAAACCATTATTATCATGACATTAACCTATAAAAATAGGCGTATCACGAGGCCCTTTCGTCTTCACCTC
GAGCGCTTCCGGGGAGATATTAGTTTGCTTATTCAGCACGCGTGATACGGTGGATTTTGATACGCCTGCAACCCTGGCAATATCATAAA
TAGTAGGGGACATAGGTCAGGAACTCCGTCCGAACGTGAATGCTGCACATCTTATGGAGTTCGCCGGTCGATGACAACACGGGCAGTTG
ATAATCAATGGCCTGGCCCCCACATTCATATCCTTACGAATGATTTTTTTTCTGGTCTTCAGAGCAGCACAGGACAGCAATGAAGCGAC
TTAAAAATGAACTCAATGCGCTGGTGAATCGGGGTGTCGACAGACATCTGCGCCTCGCTGTAACCGGACTTAGCCGCAGGATCCTCTAG
ATTTAAGAAGGAGATATACATATG
>ycjY
GAGGCAATTCCGACGTCTAAGAAACCATTATTATCATGACATTAACCTATAAAAATAGGCGTATCACGAGGCCCTTTCGTCTTCACCTC
GATGGATCCCTGCAACGACGACAAACGCCATCAGATCAGCGATTTCTTCACGCTTCATTCAGTATCTCCATGCAATCTATTAATCATTT
AAATGATAGGTCTAATCATGATTTTGGGTCTAATCAACTTAATGATCAATATCTATAGTGATATCACCAACCGTTAAAACGAACCGACA
ACTCCATGAGGAACGGAATGATGAATAATAAAGTCAGCTTCACTAACAGCAATAATCCAACCATCTCTTTGTCTGCCTCGAGAGATCCT
CTAGATTTAAGAAGGAGATATACATATG
>yhbU

GAGGCAATTCCGACGTCTAAGAAACCATTATTATCATGACATTAACCTATAAAAATAGGCGTATCACGAGGCCCTTTCGTCTTCACCTC
GAGTTTTAGCGCAAATGGCGTCAGTTTTACCGGTACACTCAACAGAGATGGCCCCAAATGCACAATACGGGAACGCAGTTTATCCAACA
CGAGTTTTACTCCCTGTTTCAACAATCATCCTATTTTGCCATATCAGAAAAATAACATAGCGGTATAAATCAACAATTCCATATGAAAT
TGCTGCTACCACCAATACAACTTTAACTGCCTTAAATCAAAAATTGTCGCAGCAAGGTTAACTAAAATCCCAGTTCGTTAACATTTTTG
CGTTTTGATAGCGCAACCTTCAGGAAAAATTATGGAGCTGCTCTGCCCTGCCGGAAATCTCCCGGCGCTTAAGGCGGCCATCGAAAACG
GCGCAGATGCTGGATCCCGGGATCCTCTAGATTTAAGAAGGAGATATACATATG
>yhbT
GAGGCAATTCCGACGTCTAAGAAACCATTATTATCATGACATTAACCTATAAAAATAGGCGTATCACGAGGCCCTTTCGTCTTCACCTC
GATGGATCCTTAAGCGCCGGGAGATTTCCGGCAGGGCAGAGCAGCTCCATAATTTTTCCTGAAGGTTGCGCTATCAAAACGCAAAAATG
TTAACGAACTGGGATTTTAGTTAACCTTGCTGCGACAATTTTTGATTTAAGGCAGTTAAAGTTGTATTGGTGGTAGCAGCAATTTCATA
TGGAATTGTTGATTTATACCGCTATGTTATTTTTCTGATATGGCAAAATAGGATGATTGTTGAAACAGGGAGTAAAACTCGTGTTGGAT
AAACTGCGTTCCCGTATTGTGCATTTGGGGCCATCTCTGTTGAGTGTACCGGTAAAACTGACGCCATTTGCGCTAAAACGCCAGGTTCC
TCGAGAGATCCTCTAGATTTAAGAAGGAGATATACATATG
>yhbW
GAGGCAATTCCGACGTCTAAGAAACCATTATTATCATGACATTAACCTATAAAAATAGGCGTATCACGAGGCCCTTTCGTCTTCACCTC
GAGCCGCGCTAATGAAAATGGCGCTGCACCACTGCCGTTGACGGCGAACAGCGACTGTAACGGCTACTGGCGGCGGCTGGCAGGACTGG
AGCTGCAAGCCTAAGTAAATAGCTCACTTTGTTAACAACTTTAACTACTCTTTAATGCAGTATTAAAGATTAATCGGTAACAAAGTGAG
CTGTTATGACTGATAAAACCATTGCGTTTTCGCTACTCGATCTGGCCCCCATTCCCGAAGGTTCTTCAGCGCGAGAGGATCCTCTAGAT
TTAAGAAGGAGATATACATATG
>yhbQ
GAGGCAATTCCGACGTCTAAGAAACCATTATTATCATGACATTAACCTATAAAAATAGGCGTATCACGAGGCCCTTTCGTCTTCACCTC
GAGCCTGCTGCACACACCAAGTGACAACATGTTGTTTTGCCAGCCAACGGCTGATGGCGATGAGTGTTTCCATTGCTGTTCTCTTTTAT
ACTGTGGGCCTGACCTTAACATGCGACGCAGACGATGACACCCTGGTTTCTTTACTTGATCCGTACCGCCGACAATAAGCTTTATACCG
GGATCACCACGGATGTCGAACGCCGCTATGGATCCTCTAGATTTAAGAAGGAGATATACATATG
>ycjG
GAGGCAATTCCGACGTCTAAGAAACCATTATTATCATGACATTAACCTATAAAAATAGGCGTATCACGAGGCCCTTTCGTCTTCACCTC
GAGGTCTGCGCTTTGCTACCCGCCTGCGGGATGGAATTGGCGACTGTAACCGGGTTGCCCTGGAAATGAACGGTTTGTGACATGATTAT
CTTTCCTGTTTACATATAGTTAACGTCACACCTAGTTTATGCTAACTGTCAATAACACAGCAAACGCTATTTGCGCTTAATCCGCAGAC
CACCGCGACAACAAGGAGTAAAGATGAGAACCGTTAAGGTATTCGAGGAAGCCTGGCCCTTAGGATCCTCTAGATTTAAGAAGGAGATA
TACATATG
>ycjK
GAGGCAATTCCGACGTCTAAGAAACCATTATTATCATGACATTAACCTATAAAAATAGGCGTATCACGAGGCCCTTTCGTCTTCACCTC
GATGGATCCGCAGAGTCTGGGTCGCATGACCCTTAAGCCTGTTCCTGCACATTACGACACCGATAACCGGATTGTTCATTATATTTTCC
ATGCTCAATCTCACAAAGTGGACTAAATTATCGCCATTACTGCGCTTTAAATGACCTTTATGTTCAATATTTTTTCAATCTAGCAGTGG
GTTTTCATTTTTGCAAACTCAATTTAACATTTGACAAACATTTAGTTTGCATACAGATTCGAATGGTGGTCATTATATTTTACGCTTTG
ATAACGAGCGGAAAACAAACCAAAGGCGCTCGAGAGATCCTCTAGATTTAAGAAGGAGATATACATATG
>yhbX
GAGGCAATTCCGACGTCTAAGAAACCATTATTATCATGACATTAACCTATAAAAATAGGCGTATCACGAGGCCCTTTCGTCTTCACCTC
GATGGATCCCCTTGAAACGATGGTGCCGGTACGCCTTAGTTATAAATTTTTCCCTTCTACTGACATAATTGGCACTTTATTGTGTCAAC
TCCAGTTCCGGTACGAATACGGATATGTTTACTTCTCGATAAACCATCCTGCTCCTGTCAGTATAACGACTGATGGCTACGTTCTGCGT
AATGTTTAGAGACGCATTGATAATGTTATTTAATTACTGAGAACGCAATGGCCGCATCATTTAAAAAGAAAAAACTAACAGTGTTTCGT
TCTGATATGGATCAGAGTTGAAAAACATGTCTGATATAAAATTAAACATTGTTGATAATTGTTTATATTGGTTGTGAACTTGGATTTCT
GGATGACAGTATTCAACAAATTTGCTAGAACTTTTAAATCTCATTGGTTGTTGTATCTTTGTGTTATTGTTTTTGGTATTACGAACTTA
GTCGCTTCTTCCGGAGCGCATATGGTTCAGCGCTTGCTGTTCTTCGTTCTGACCATCCTGGTTGTAAAACGTATATCATCCCTTCCGCT
TCGCCTGCTTGTTGCCGCACCATTTGTGTTACTGACTGCGGCAGACATGAGTATTAGCCTCTATTCATGGTGTACCTTTGGTACAACTT
TCAATGATGGATTTGCGACTCGAGAGATCCTCTAGATTTAAGAAGGAGATATACATATG
>ycjL
GAGGCAATTCCGACGTCTAAGAAACCATTATTATCATGACATTAACCTATAAAAATAGGCGTATCACGAGGCCCTTTCGTCTTCACCTC
GAGCGCCTTTGGTTTGTTTTCCGCTCGTTATCAAAGCGTAAAATATAATGACCACCATTCGAATCTGTATGCAAACTAAATGTTTGTCA
AATGTTAAATTGAGTTTGCAAAAATGAAAACCCACTGCTAGATTGAAAAAATATTGAACATAAAGGTCATTTAAAGCGCAGTAATGGCG
ATAATTTAGTCCACTTTGTGAGATTGAGCATGGAAAATATAATGAACAATCCGGTTATCGGTGTCGTAATGTGCAGGAACAGGCTTAAG
GGTCATGCGACCCAGACTCTGCGGATCCTCTAGATTTAAGAAGGAGATATACATATG
>ycjM
GAGGCAATTCCGACGTCTAAGAAACCATTATTATCATGACATTAACCTATAAAAATAGGCGTATCACGAGGCCCTTTCGTCTTCACCTC
GAGGGGCAAGCAAAAGAGATCCTTAGCGAGATGGGATATACCCACGTTGAGAACGCCGGTGGCCTGAAAGACATCGCAATGCCGAAGGT
CAAAGGTTAAAAGAATTCACCATGAGCGGCGCTTATGCGCCGTTTTTTTTTCTGTCACACCTTATTTACATCCCCATAGATTATTTGCG
TCAGCTCACAAATACGCTTTTTCCCTGGTAAAAAATGATTTCCTGCGTGACTAAAACCCTTGTGCTCAATTGACAGTTTATTTTCTGCG
GAGTAGTCTCTCGTTTCATGGGACCGCTACCACGGAAAGGCAACATGAAACAGAAAATTACGGATTACCTGGACGAAATCTACGGTGGG
GATCCTCTAGATTTAAGAAGGAGATATACATATG
>ymjA
GAGGCAATTCCGACGTCTAAGAAACCATTATTATCATGACATTAACCTATAAAAATAGGCGTATCACGAGGCCCTTTCGTCTTCACCTC
GATGGATCCTCGCAAAGTGCCGTTGTATGATGGTGACAGAACGCCGGTGAGTGAAACGTAATAGTCGGATGCGCGGCGCATTATCCTGC
CTGAGGGTAACGCGCCGCGCCATTAACTATAAAAAGATGTATGACCCGGATCACGAGTGCTATCATCCGCCCTTATGCTTAGCGCATTA

ACCCCAGGGGCAAATAATGAACCACGATATCCCGCTAAAATATTTTGATATTGCCGATGAGTACGCGACCGAATGCGCAGAGCCTGTCG
CAGAGGCTGAACGCACACCCTCGAGAGATCCTCTAGATTTAAGAAGGAGATATACATATG
>rhoL
GAGGCAATTCCGACGTCTAAGAAACCATTATTATCATGACATTAACCTATAAAAATAGGCGTATCACGAGGCCCTTTCGTCTTCACCTC
GAGAAAAACGGTGAAGTGGCGGCAACCAAAGTGGGTGCACTGTCTAAAGGTCAGTTGAAAGAGTTCCTCGACGCTAACCTGGCGTAAGG
GAATTTCATGTTCGGGTGCCCCGTCGCTAAAAACTGGACGCCCGGCGTGAGTCATGCTAACTTAGTGTTGACTTCGTATTAAACATACC
TTATTAAGTTTGAATCTTGTAATTTCCAACGCTTCCCGTTTTATCTTAAATGCGAAGTGAACAGATTTCTGGCTCGTCACTCAATCCGT
CTTGTCGTTTCAGTTCTGCGTACTCTCCTGTGACCAGGCAGCGAAAAGAGGATCCCGGGATCCTCTAGATTTAAGAAGGAGATATACAT
ATG
>nrfE
GAGGCAATTCCGACGTCTAAGAAACCATTATTATCATGACATTAACCTATAAAAATAGGCGTATCACGAGGCCCTTTCGTCTTCACCTC
GAGGCATTCCTGCCGTGCTGGCGGCGTGTGGGGCCAGTCTGGTCGGCGTGTTGATGCTGCGCTTTTTCATTCTCTACGCCGGACAGTTG
ACGGTGGCGTAAGCCAGAAAAGAGGTGGTTTCTGGACGTATTCCTTCCTGAAGTCGGTTTTCTGGCGTTGTTGTTAAGTCTCGGGGTCA
ACGTGTTGACCCCGTTGACGGCCTTCGCGGGAGTGCGGTTGCGCTGGCCTGCCATGATGCGACTCACTTGCATCGGCAGGATCCTCTAG
ATTTAAGAAGGAGATATACATATG
>pgk
GAGGCAATTCCGACGTCTAAGAAACCATTATTATCATGACATTAACCTATAAAAATAGGCGTATCACGAGGCCCTTTCGTCTTCACCTC
GATGGATCCGTGCGATAACGAATGGGGCTTTGCTAACCGAATGCTCGACACGACGTTAGCTATGGCTACTGTTGCTTTCAGGTAAGACG
CAAGCAGCGTCTGCAAAACTTTTAGAATCAACGAGAGGATTCACCATGTCTGTAATTAAGATGACCGATCTGGATCTTGCTGGGAAACG
TGTATTTATCCGTGCGGATCTGAACGTACCAGTAAAAGACGGGAAAGTAACCAGCGACTCGAGAGATCCTCTAGATTTAAGAAGGAGAT
ATACATATG
>pgi
GAGGCAATTCCGACGTCTAAGAAACCATTATTATCATGACATTAACCTATAAAAATAGGCGTATCACGAGGCCCTTTCGTCTTCACCTC
GAGATATCAGCGCTGCGGTTCATGGCGTCAAAATCAGCTACGCTGGTACCGCCAAATTTGGAGACAACAATTTCAGACATAACTACCTC
GTGTCAGGGGATCCATTTTCAGCCTTGGCACAAGGGAAGAGCGGAAGACGGGTGGGCGCAGAGCGATACTTCGCTACTATTTTCACCCA
GAAGTGCTCCACCACTTGCGAAACGCCCGACTGCGAACGCTTCTGGTGACAACCCAGGGGATTCAGCCCCTGTAGCCGATGATGAACGT
GGCCAGCCGTTCAATCACCTCGGCGATGCACCCCCTCAGGTGTTATCACAGGACTGGCTCCTCCAACACCGTTACTTGGGCAACGCGCC
TCTTCTGGCCTGCGCTAGCGCAGGTAGTACATTTATAAATAAAGGGTGAGCGGGGCGGTTGTCAACGATGGGGTCATGCGGATTTTTCA
TCCACTCCTGGCGGTCAGTAGTTCAGCTAATAAATGCTTCACTGCGCTAAGGGTTTACACTCAACATTACGCTAACGGCACTAAAACCA
TCACATTTTTCTGTGACTGGCGCTACAATCTTCCAAAGTCACAATTCTCAAAATCAGAAGAGTATTGCTAATGAAAAACATCAATCCAA
CGCAGACCGCTGCCTGGCAGGCACTACAGAAACACTTCGATGAAATGAAAGACGTTACGATCGCCGAGATCCGAATTCCTCGAGAGATC
CTCTAGATTTAAGAAGGAGATATACATATG
>potF
GAGGCAATTCCGACGTCTAAGAAACCATTATTATCATGACATTAACCTATAAAAATAGGCGTATCACGAGGCCCTTTCGTCTTCACCTC
GAGGCTAATGCCCATCAACTGCTGTTACCGACTGATGATGAAGGGCAAAACAACGTTACCGAAAACTATTTCCTCCACTGATAACTCCT
TTCGAGCACGCAGTCGCTGGTGCAGTGGCTGCGCGCTGCAAAATTATCTGCTGTTTTTAACCTTTTCTTAAAGATTATTTCACTTCTCT
TGTGTCGATTTGGCTTTATCACATAGAGCAAATATGCATAAAAAATTTGTTAAATACCGTTTTTTAATCCGAGCTATAGTCTCAAACCCT
GGCTAAAGTTATTCTTGCGATGCTTTTATATAGCGAGCAGTGCTGGCCGGGAGAAAGTTCTCTTTTCTTACACCGCGCCGATAAAAAAT
ATGCACGTTTATTGCATATCTTTCAGTGTGACAACTTTTGTTCGTTTGTTAACGAACTTTCAGAAGGAAAGAGATATGACCGCCTTAAA
TAAAAAATGGCTATCGGGTCTGGTTGCGGGTGCTCTGATGGCCGTCTCTGTCGGCACGCTCGGATCCTCTAGATTTAAGAAGGAGATAT
ACATATG
>potA
GAGGCAATTCCGACGTCTAAGAAACCATTATTATCATGACATTAACCTATAAAAATAGGCGTATCACGAGGCCCTTTCGTCTTCACCTC
GATGGATCCCGTGCTGGGAACCTGTCTCACCCCTGCTTTTGATTGGGTATCCAGAGACACGTAGTTCAAAAATCGCTCAAGTAGTTTAT
CCATGTAGTCACCCTCACTTTTTGTGACAACATTATTAAGAAGCAAGAAAAGACAAATATTGCGTCAGGTCACTTTTACCCCTGCAAGC
GGGAATATTTATCAGCATTACCTTAATGAATATAAAGCTAAGCCAGTAATTCACAACAAAGATTGGCGATTCAAGCGGTTTGCCGTAGA
ATTACCGCCCTCATTAAGAGTCACCAAGGTGGTTAACCACAAACCCCGCATCGGTAAGCCATCCGTTGCGTTTACATGGGACAGAGTAA
AAAATTGAATAAACAACCGAGTTCGCTTTCACCGCTGGTGCAATTGGCGGGAATTCGCAAATGCTTTGATGGCTCGAGAGATCCTCTAG
ATTTAAGAAGGAGATATACATATG
>mviN
GAGGCAATTCCGACGTCTAAGAAACCATTATTATCATGACATTAACCTATAAAAATAGGCGTATCACGAGGCCCTTTCGTCTTCACCTC
GAGAGACAGTTCCGCAAACCGCCGGCGAACAGGCCGTGCTGGCGCAACGTATCGTTGACAAGATCTGGCGCGATGCGATGAGTGAATAA
ACCCTGTAACATCTGGCGGTAGCAATTCACCATAATCCAGGTAGACTATTCGCCTCTTTCAGCGCCTGCCTTGCAGGCGTTTTGCCCGT
GGGTCTGGATTAGAACACCGATGAATTTATTAAAATCGCTGGCCGCCGTCAGCTCGATGACCATGTTTTCGCGTGTGCTTGGCTTCGCA
CGAGACGCAATTGTCGGGATCCTCTAGATTTAAGAAGGAGATATACATATG
>gatR1
GAGGCAATTCCGACGTCTAAGAAACCATTATTATCATGACATTAACCTATAAAAATAGGCGTATCACGAGGCCCTTTCGTCTTCACCTC
GAGGCTGCCGGGAGACTGTTGGTAATCACCGTGATATTTTTGGCGGTCATTAATCCTTCGGCGATAAGCATTGTCGTACTTCCGCTGTC
GAGAATAACGGTCTGATCTTACCCAGCAATAGTGGACACGCGGCTAAGTGAGTAAACTCTCAGTCAGAGGTGACTCACATGACAAAAAC
AGTATCAACCAGTAAAAAACCCCGTAAACAGCATTCGCCTGAATTTCGCAGTGAAGCCCTGAAGCTTGCCTCGAGCGGAGATCCTCTAG
ATTTAAGAAGGAGATATACATATG
>pflA

GAGGCAATTCCGACGTCTAAGAAACCATTATTATCATGACATTAACCTATAAAAATAGGCGTATCACGAGGCCCTTTCGTCTTCACCTC
GATGGATCCAAAGAACAGCAGCAGGACGTTATTACTCGTACCTTCACTCAATCTATGTAATTAGATTTGACTGAAATCGTACAGTAAAA
AGCGTACAATAAAGGCTCCACGAAAGTGGGGCCTTTTTAGCGCGAGAGCCTTTTTTGTCAGCTATCTATACTTTAAGGTGACTGCCAA
AACAGACTCGACGTAGCCTTCGAGCTGCGCACCAACACGGCCTCAGATGGGCCACATCTGGAGAAACACCGCAATGTCAGTTATTGGTC
GCATTCACTCCTTTGAATCCTGTGGAACCGTAGACGGCCCAGGTATTCGCTTTATCACCTTTTTCCAGGGCTGCCTCGAGAGATCCTCT
AGATTTAAGAAGGAGATATACATATG
>ahpF
GAGGCAATTCCGACGTCTAAGAAACCATTATTATCATGACATTAACCTATAAAAATAGGCGTATCACGAGGCCCTTTCGTCTTCACCTC
GAGGTGAAGTTTGCCCGGCTAAATGGAAAGAAGGTGAAGCAACTCTGGCTCCGTCTCTGGACCTGGTTGGTAAAATCTAAATTTCCTTC
GTCTTTCACGCCATAGCGGCGTTGGCGTCGCCCGCTCACCCCGGTCACTTACTTGTGTAAGCTCCCGGGGATTCACAGGCTAGCCGCCT
TGCTCTGACGCGAAATACTTCGGAAATTCACCTAATTCTTCGGGTGCTGCGGCACCCGATTTCTTCCCCGCACCATGATGCAAGCTGC
ATCCAGGTAGCCGCAGAGGCCGCTTGCATGATGATGTTTAAAGCCCAGGAGATAAACATGCGGATCCTCTAGATTTAAGAAGGAGATAT
ACATATG
>ahpC
GAGGCAATTCCGACGTCTAAGAAACCATTATTATCATGACATTAACCTATAAAAATAGGCGTATCACGAGGCCCTTTCGTCTTCACCTC
GAGGCAGGAAGCAGAGCCAGTAAAAGTATCTTTTTTAACATTAATTTGTCCTTTTCAGTCAGTGCAAAAGTCGAGTAAAAGGCATAACC
TATCACTGTCATAGGTAAGAGCTTAGATCAGGTGATTGCCCTTTGTTTATGAGGGTGTTGTAATCCATGTCGTTGTTGCATTTGTAAGG
GCAACACCTCAGCCTGCAGGCAGGCACTGAAGATACCAAAGGGTAGTTCAGATTACACGGTCACCTGGAAAGGGGGCCATTTTACTTTT
TATCGCCGCTGGCGGTGCAAAGTTCACAAAGTTGTCTTACGAAGGTTGTAAGGTAAAACTTATCGATTTGATAATGGAAACGCATTAGC
CGAATCGGCAAAAAATTGGTTACCTTACATCTCATCGAAAACACGGAGGAAGTATAGATGTCCTTGATTAACACCAAAATTAAACCTTTT
AAAAACCAGGCATTCAAAAACGGCGAATTCATCGAGGATCCTCTAGATTTAAGAAGGAGATATACATATG
>udp
GAGGCAATTCCGACGTCTAAGAAACCATTATTATCATGACATTAACCTATAAAAATAGGCGTATCACGAGGCCCTTTCGTCTTCACCTC
GAGGCAGGCATGTTATCCCCTTGTGAAGGGATAGAGGTGAAGCCCGCCACAATCGCGTCGTCCGGGGTCTGAACGATGGTCGAAGCGAG
AGGCGATGCAGCAGGTGCAAATCCAGATTGTTGTGTTGTTGCCATGGTATTCTCCGTACCCTTATAAAAATGTTGCGCAATGTTAACTA
TAGTCAGCATGCAACAAATCACATTGCCTGAATCGGCTCATCTTTTATGCAGTCCTGCAGAATGAAGGGTGATTTATGTGATTTGCATC
ACTTTTGGTGGGTAAATTTATGCAACGCATTTGCGTCATGGTGATGAGTATCACGAAAAAATGTTAAACCCTTCGGTAAAGTGTCTTTT
TGCTTCTTCTGACTAAACCGATTCACAGAGGAGTTGTATATGTCCAAGTCTGATGTTTTTCATCTCGGCCTCACTAAAAACGATTTACA
AGGGGCTACGCTTGCCATCGTCCGGATCCTCTAGATTTAAGAAGGAGATATACATATG
>yqcE
GAGGCAATTCCGACGTCTAAGAAACCATTATTATCATGACATTAACCTATAAAAATAGGCGTATCACGAGGCCCTTTCGTCTTCACCTC
GAGGCTCCAATTTCCGTAAGGCAAAAAAAAAGAAAGGTAACTTCCCCCGCGAGAGGGAAGTTACCTTTCTCATTATCTCTGGTAACTGG
AGATAATTTTTAGCATAAGGGCATAACTGCTAATGTGATCGTAATCACAGTGTGATATTCGTATTGTCAAATAATTTTAAAAAGCAGAA
GCGTGATTACTCTCACACAACACCTCTGCTTTCCATGTTATTTATTATTTCAATTACTAGAGACCTGAGAAAAGTAATCCACCTGAAAG
GGCGGCTTACTTTTCTCTTTTTTTTTGTCTATTACAAATGGGAATACGATATGCAACACAACTCATATCGCCGTTGGATAACCCTCGCGA
TAATTAGTTTTAGCGGCGGCGTTAGTTTCGACCTGGATCCTCTAGATTTAAGAAGGAGATATACATATG
>yqcD
GAGGCAATTCCGACGTCTAAGAAACCATTATTATCATGACATTAACCTATAAAAATAGGCGTATCACGAGGCCCTTTCGTCTTCACCTC
GAGCGGCCAGCTTTTATGCTCTTCATGCCACGCATCGCAGTAGCGTGCAGTAAAATCTTTCAGGGCTTGTGCGGTCAAATCGTCCACTA
ATTTCTCTCTTCACTTAAACCAGATACACTTGTCGTTTAGTTTATCTGGTTTATGACGGTGAAACATGTCTTCTTATGCAAACCATCAG
GCACTTGCGGGCCTGACTCTTGGAAAATCAACCGATTACCGGGATACCTATGACGCCAGCCTACTGCGGATCCTCTAGATTTAAGAAGG
AGATATACATATG
>rsd
GAGGCAATTCCGACGTCTAAGAAACCATTATTATCATGACATTAACCTATAAAAATAGGCGTATCACGAGGCCCTTTCGTCTTCACCTC
GATGGATCCTCCCTTCGGCAACCATAATTTTTGTTCATGGCTGACGACCCACCAGCCGTGATCTAATTTTTCAATTATACGATCCATAG
CTCTTGCACTACCTTTGCATCACTGGCATGTTTAACATGGTTTTTACATTTCTCACTGAGCAGTTTTTGAATACAAACTTGCGGAGTCA
ATCATGCTTAACCAGCTCGATAACCTGACGGAGCGCGTCAGAGGAAGTAACAAACTGGTTGATCGCTGGCTACATGTACGTAAGCATCT
GCTCGTGGCTCTCGAGAGATCCTCTAGATTTAAGAAGGAGATATACATATG
>udk
GAGGCAATTCCGACGTCTAAGAAACCATTATTATCATGACATTAACCTATAAAAATAGGCGTATCACGAGGCCCTTTCGTCTTCACCTC
GATGGATCCGTGGGGCAGGGCAATTAATACATGCTGTGATTGTTTGCTCATTACCGCTTTATCCGCAAGACCTGGTCGCCAGTATTCGT
TTCTGTGTTTATGCTAGTACAAAAGTTTTACATTTTATATGCGAATTGCTCATAAAGTGACGTAAAGGCGGATAATTTGCGCAACTGCG
TTTAACATTTTTTACCTTACATAAAACTGATCAACGTAATTTGCCCCCGGGAATTCTTTGTCAACAATTAACTCTGAGCGCGCGGCAAC
GCTATTCGACTGGTATCAGACGGATGAAATCCCTATAATTGCCGCGTTTGGCGCTTCGTCGCCCCCTTCCTAACATCCAGGTTAATCAG
GTCGCTAAATATATGACTGATCAGTCTCATCAGTGCGTCATTATCGGTATCGCTGGCGCATCGGCTTCCGGCAAGAGTCTTATTGCCAG
TACCCTTTATCGTGAATTGCGTGAGCAAGTCGGTGATGAACACATCGGCGTAATTCCCCTCGAGAGATCCTCTAGATTTAAGAAGGAGA
TATACATATG
>xylB
GAGGCAATTCCGACGTCTAAGAAACCATTATTATCATGACATTAACCTATAAAAATAGGCGTATCACGAGGCCCTTTCGTCTTCACCTC
GATGGATCCGTGGTCGCCAGGAACAACTGGAAAATCTGGTAAACCATTATCTGTTCGACAAATAACGGCTAACTGTGCAGTCCGTTGGC
CCGGTTATCGGTAGCGATACCGGGCATTTTTTTAAGGAACGATCGATATGTATATCGGGATAGATCTTGGCACCTCGGGCGTAAAAGTT
ATTTTGCTCAACGAGCAGGGTGAGGTGCTCGAGAGATCCTCTAGATTTAAGAAGGAGATATACATATG
>xylE

```
GAGGCAATTCCGACGTCTAAGAAACCATTATTATCATGACATTAACCTATAAAAATAGGCGTATCACGAGGCCCTTTCGTCTTCACCTC
GATGGATCCGGCAGGTGGTGTGAAAGGTTAAAGATGTTGTTCTGCCAATGTTATGCCGCTGCACCCTCAACTTACGTTATCCCAACTTG
TGACTGTTATTCGGCGCTCCACGGAGCGCCTTTTTTTCTTTCGTCTGCAATCTGAATCGTTCGCCGGTTAATATTTCCATCATAGAGCT
TATTATTTTTACGTTATTTGTTTTCCCACTTACGATAATTCTCTTTCGTGCTCTGAGTCACGGCAATAGTATTGTTTTTATCAATTTTG
GATAATTATCACAATTAAGATCACAGAAAAGACATTACGTAAACGCATTGTAAAAAATGATAATTGCCTTAACTGCCTGACAATTCCAA
CATCAATGCACTGATAAAAGATCAGAATGGTCTAAGGCAGGTCTGAATGAATACCCAGTATAATTCCAGTTATATATTTTCGATTACCT
TAGTCGCTACATTAGGTGGTTTATTATTTGGCTACGACACCGCCGTTATTTCCTCGAGAGATCCTCTAGATTTAAGAAGGAGATATACA
TATG
>xylG
GAGGCAATTCCGACGTCTAAGAAACCATTATTATCATGACATTAACCTATAAAAATAGGCGTATCACGAGGCCCTTTCGTCTTCACCTC
GAGCTCCCGCCTCCTGACACCGATCGATGTGAATAAAAACAACATCAAAGATACGGTAATTAAAGACGGATTCCACAAAGAGAGCGAGC
TGTAAGCGTTACGCCCCAGCGCGGAGCGGGGGCGTGATTTCTCTCCATGCCGCGTGAATGAATTGGCTTAGGTGGAGTCGTTATGCCTT
ATCTACTTGAAATGAAGAACATTACCAAAACCTTCGGCAGTGTGAAGGCGATTGATAACGTCTGCTTGCGGTTGAATGCTGGCGAAATC
GTCGGATCCTCTAGATTTAAGAAGGAGATATACATATG
>xylF
GAGGCAATTCCGACGTCTAAGAAACCATTATTATCATGACATTAACCTATAAAAATAGGCGTATCACGAGGCCCTTTCGTCTTCACCTC
GAGAACACCAGTTCGTCGGGATTGTAGTGACGGAATGCTAACGGGTTTGAGGATTTTGAGCCTTCATAACGAACGCGATCGAGCTGGTC
AAAATAGGCTTGCATATTGAACTCCATAATCAGGTAATGCCGCGGGTGATGGATGATGTCGTAATATTGGGCACTCCCTTTCAGTTGCT
CAATTATGTTATTTCACACTGCTATTGAGATAATTCACAAGTGTGCGCTCGCTCGCAAAATAAAATGGAATGATGAAACTGGGTAATTC
CTCGAAGAGAAAAATGCAATAAGTACAATTGCGCAACAAAAGTAAGATCTCGGTCATAAATCAAGAAATAAACCAAAAATCGTAATCGA
AAGATAAAAATCTGTAATTGTTTTCCCCTGTTTAGTTGCTAAAAATTGGTTACGTTTATCGCGGTGATTGTTACTTATTAAAACTGTCC
TCTAACTACAGAAGGCCCTACACCATGAAAATAAAGAACATTCTACTCACCCTTTGCACCTCACTCCTGCTTACCAACGTTGCTGCACA
CGCCAAAGAAGGGATCCTCTAGATTTAAGAAGGAGATATACATATG
>dnaQ
GAGGCAATTCCGACGTCTAAGAAACCATTATTATCATGACATTAACCTATAAAAATAGGCGTATCACGAGGCCCTTTCGTCTTCACCTC
GAGAAATAGCGCCGTAACCCCCAGGTCCTGGATTGCCCAGACACGAACCATCGGTGAAAATTTCTACCTGTTTAAGCATCTCTGGTAGA
CTTCCTGTAATTGAATCGAACTGTAAAACGACAAGTCTGACATAAATGACCGCTATGAGCACTGCAATTACACGCCAGATCGTTCTCGA
TACCGAAACCACCGGTATGAACCAGATTGGTGCGCACTATGAAGGCCACAGGATCCTCTAGATTTAAGAAGGAGATATACATATG
>dnaX
GAGGCAATTCCGACGTCTAAGAAACCATTATTATCATGACATTAACCTATAAAAATAGGCGTATCACGAGGCCCTTTCGTCTTCACCTC
GAGTCAACCTGTTCGATCTCGGCGGCGAACAGCGTCTCGAAAAACAGGGCATTACCAGCTACAGCCTTGTCCCGTTCCCGGGCCATTAA
TTATCGCCAGTCTTGTGCTGCCCACGCTACGGACAGCACAAGATGTGCATTCAGCCTCGCCGTTCTGACGGGGCTGTGTTAGCATTACC
CCTTCGTGAATCCACCTTCCAGCGTTTCAGAGCCTGCCAATGAGTTATCAGGTCTTAGCCCGAAAATGGCGCCCACAAACCTTTGCTGA
CGTCGTCGGCCAGGAACATGTGCTGAGGATCCTCTAGATTTAAGAAGGAGATATACATATG
>rdoA
GAGGCAATTCCGACGTCTAAGAAACCATTATTATCATGACATTAACCTATAAAAATAGGCGTATCACGAGGCCCTTTCGTCTTCACCTC
GAGCGAAAGATGCGGTGATCCCGGGTTTGCAGAAAGATTATGAAGAAGACTTCAAAACGGCGCTGTTACGCGCTCGCGGCGTAATTAAA
GAGTAAAAGCTTGTAAGCGGCGCCACCAAAATCATCGTGAAATGATATCCTTCGTCATTCGTAATGTTTTCCGGATGATGGGATGAATA
ACAGCGCTTTTACTTTCCAGACACTACACCCGGATACCATCATGGACGCTCTGTTTGAGCATGGGATCCGGGATCCTCTAGATTTAAGA
AGGAGATATACATATG
>dnaK
GAGGCAATTCCGACGTCTAAGAAACCATTATTATCATGACATTAACCTATAAAAATAGGCGTATCACGAGGCCCTTTCGTCTTCACCTC
GAGTTTATGGTCGTTTGCCTGCGCCGTGCAGCACAGCATCAGGCTAATCGCCAGGCTGGCGGAAATCGTAAAAACGGATTTCATAAGGA
TTCTCTTAGTGGGAAGAGGTAGGGGGATGAATACCCACTAGTTTACTGCTGATAAAGAGAAGATTCAGGCACGTAATCTTTTCTTTTTA
TTACAATTTTTTGATGAATGCCTTGGCTGCGATTCATTCTTTATATGAATAAAATTGCTGTCAATTTTACGTCTTGTCCTGCCATATCG
CGAAATTTCTGCGCAAAAGCACAAAAAAATTTTTGCATCTCCCCCTTGATGACGTGGTTTACGACCCCATTTAGTAGTCAACCGCAGTGA
GTGAGTCTGCAAAAAAATGAAATTGGGCAGTTGAAACCAGACGTTTCGCCCCTATTACAGACTCACAACCACATGATGACCGAATATAT
AGTGGAGACGTTTAGATGGGTAAAATAATTGGTATCGACCTGGGTACTACCAACTCTTGTGTAGCGATTATGGATGGCACCGGATCCTC
TAGATTTAAGAAGGAGATATACATATG
>dnaJ
GAGGCAATTCCGACGTCTAAGAAACCATTATTATCATGACATTAACCTATAAAAATAGGCGTATCACGAGGCCCTTTCGTCTTCACCTC
GAGTTCTGCAAACAACGCGAAAGATGACGATGTTGTCGACGCTGAATTTGAAGAAGTCAAAGACAAAAAAATAATCGCCCTATAAACGGG
TAATTATACTGACACGGGCGAAGGGGAATTTCCTCTCCGCCCGTGCATTCATCTAGGGGCAATTTAAAAAAGATGGCTAAGCAAGATTA
TTACGAGATTTTAGGCGTTTCCAAAACAGCGGAAGAGCGTGAAATCAGAAAGGCCTACAAACGCCTGGCCATGAAATACGGATCCTCTA
GATTTAAGAAGGAGATATACATATG
>sdhC
GAGGCAATTCCGACGTCTAAGAAACCATTATTATCATGACATTAACCTATAAAAATAGGCGTATCACGAGGCCCTTTCGTCTTCACCTC
GAGTGCCTTTCAGCACATCCAGTTCAACAGCTGTATCCCCGTTGAGGGTGAGTTTTGCTTTTGTATCAGCCATTTAAGGTCTCCTTAGC
GCCTTATTGCGTAAGACTGCCGGAACTTAAATTTGCCTTCGCACATCAACCTGGCTTTACCCGTTTTTTATTTGGCTCGCCGCTCTGTG
AAAGAGGGGAAAACCTGGGTACAGAGCTCTGGGCGCTTGCAGGTAAAGGATCCATTGATGACGAATAAATGGCGAATCAAGTACTTAGC
AATCCGAATTATTAAACTTGTCTACCACTAATAACTGTCCCGAATGAATTGGTCAATACTTCCACACTGTTACATAAGTTAATCTTAGG
TGAAATACCGACTTCATAACTTTTACGCATTATATGCTTTTCCTGGTAATGTTTGTAACAACTTTGTTGAATGATTGTCAAATTAGATG
ATTAAAAATTAAATAAATGTTGTTATCGTGACCTGGATCACTGTTCAGGATAAAACCCGACAAACTATATGTAGGTTAATTGTAATGAT
```

TTTGTGAACAGCCTATACTGCCGCCAGGTCTCCGGAACACCCTGCAATCCCGAGCCACCCAGCGTTGTAACGTGTCGTTTTCGCATCTG
GAAGCAGTGTTTTGCATGACGCGCAGTTATAGAAAGGACGCTGTCTGACCCGCAAGCAGACCGGAGGAAGGAAATCCCGACGTCTCCAG
GTAACAGAAAGTTAACCTCTGTGCCCGTAGTCCCCAGGGAATAATAAGAACAGCATGTGGGCGTTATTCATGATAAGAAATGTGAAAAA
ACAAAGACCTGTTAATCTGGACCTACAGACCATCCGGTTCCCCATCACGGCGATAGCGTCCATTCTCCATCGAGATCCTCTAGATTTAA
GAAGGAGATATACATATG
>trpL
GAGGCAATTCCGACGTCTAAGAAACCATTATTATCATGACATTAACCTATAAAAATAGGCGTATCACGAGGCCCTTTCGTCTTCACCTC
GATGGATCCTGGCTGTGCAGGTCGTAAATCACTGCATAATTCGTGTCGCTCAAGGCGCACTCCCGTTCTGGATAATGTTTTTTGCGCCG
ACATCATAACGGTTCTGGCAAATATTCTGAAATGAGCTGTTGACAATTAATCATCGAACTAGTTAACTAGTACGCAAGTTCACGTAAAA
AGGGTATCGACAATGAAAGCAATTTTCGTACTGAAAGGTTGGTGGCGCACTTCCTGAAACGGGCAGTGTATTCACCATGCGTAAAGCAA
TCAGATACCCAGCCCGCCTACTCGAGAGATCCTCTAGATTTAAGAAGGAGATATACATATG
>yraR
GAGGCAATTCCGACGTCTAAGAAACCATTATTATCATGACATTAACCTATAAAAATAGGCGTATCACGAGGCCCTTTCGTCTTCACCTC
GATGGATCCGCGTTGTTGCATTGGATGAGGCGAATTTCCCACAATCGGGGATAATTACCTCATGGTCTATTAAGCCTGGTGCATGAGCC
GTGGAGCGCAAGGCAGACAATTCTGTAAAGGAGCGAAGATGAGTCAGGTACTGATTACAGGCGCAACGGGGCTGGTGGGCGGTCACCTG
CTGCGGATGTCTCGAGAGATCCTCTAGATTTAAGAAGGAGATATACATATG
>yraQ
GAGGCAATTCCGACGTCTAAGAAACCATTATTATCATGACATTAACCTATAAAAATAGGCGTATCACGAGGCCCTTTCGTCTTCACCTC
GATGGATCCGGGTCATGCTGGCAGAGTCGATGCGGCCTGAGCATGAAGGCGTGACGATTTTAAGCTCTTCAGAATTACGCAAAAGAGCT
GAATAATTATTCCCAAGAATTGATAGTGAGCGTAATATTCACGCGCTCAATTATCATAACTTATTCCTGGGGAATGCTATGACTGGTCA
GTCTTCATCTCAGGCGGCAACGCCCATTCAGTGGTGGAAACCCGCGCTTTTCTTTCTCGTTGTCATTGCCGGTCTCTGGTCTCGAGAGA
TCCTCTAGATTTAAGAAGGAGATATACATATG
>vacJ
GAGGCAATTCCGACGTCTAAGAAACCATTATTATCATGACATTAACCTATAAAAATAGGCGTATCACGAGGCCCTTTCGTCTTCACCTC
GATGGATCCGCCGCGCTCTTTTTCTTCGCTCTCAACTTCAATTTCGTCGCTGTGTTGATCAATTTTGTCGTTGTCCATGGTCTCTTCTC
TTGAATTAGCACGTATAGCTAAAGCGTAGCGGCTTTTTTGCTCGCAACTGACGGGAGTTACTCTGAAAATGTAGAAAAGGCTGCGTTTG
CCTTTTTCTGTTTCTATAGAATCAAGTAGCCTACAGGGCGGCGATTACCAGGCTATGATCAAATCAGCAAATCAGGGCGTCTGGACATC
AGTTGACGTGCTGTTACAATCGCCCACACCTAAACAGGCGGATACGGTATCGTTCCGTCATGGATGGCAAACTGCATAAGCCATAAAAA
AACAGGGAGACATTTATGAAGCTTCGCCTGTCGGCGCTTGCTCTGGGAACTACGCTTCTGGTGGGGTGTGCGAGTTCCGGTACAGATCA
GCAAGGGCGTTCTGCTCGAGAGATCCTCTAGATTTAAGAAGGAGATATACATATG
>yliE
GAGGCAATTCCGACGTCTAAGAAACCATTATTATCATGACATTAACCTATAAAAATAGGCGTATCACGAGGCCCTTTCGTCTTCACCTC
GAGTGACCGTACTGGCGTTCAATTTGTTGGGCGATGGTTTACGCGATGCGCTGGATCCGAAAATTAAAGGATAGTTACGTTTGAATATT
GCTTGAAAGGGTAATCACCTCACAGGAAATTATTGCCCTAAGCAAGTGTTGTAACTTTCTGCTGATTTTGTAGAATCGGGTAATTTGGT
TAAAAAGCCGCAGCAAGGGACAATTTTTGCAGCGGCACAGCGTTCAGATAGTTATTTTGTTAAATGTATTAACATGCTGAGTTTATACG
AAAAGATAAAGATAAGGCTGATAATTTTATTTTTATTGGCAGCACTGTCAGGATCCTCTAGATTTAAGAAGGAGATATACATATG
>yraL
GAGGCAATTCCGACGTCTAAGAAACCATTATTATCATGACATTAACCTATAAAAATAGGCGTATCACGAGGCCCTTTCGTCTTCACCTC
GATGGATCCGCGGCTTTCAAACGAGAAAATGTTGAGGGTACCATAATGTATCCAGTGATATTTTTTTTACGCAATGCTCAATATTAAAT
CGGCAATACGGACGACACAATGAAACAACACCAATCGGCGGATAATTCTCAGGGCCAGCTTTACATTGTACCGACGCCAATCGGCAATC
TGGCGGATATCACCCAGCGTGCGTTAGAGGTCTCGAGAGATCCTCTAGATTTAAGAAGGAGATATACATATG
>yraM
GAGGCAATTCCGACGTCTAAGAAACCATTATTATCATGACATTAACCTATAAAAATAGGCGTATCACGAGGCCCTTTCGTCTTCACCTC
GAGACCTCTAACGCACGCTGGGTGATATCCGCCAGATTGCCGATTGGCGTCGGTACAATGTAAAGCTGGCCCTGAGAATTATCCGCCGA
TTGGTGTTGTTTCATTGTGTCGTCCGTATTGCCGATTTAATATTGAGCATTGCGTAAAAAAAATATCACTGGATACATTATGGTACCCT
CAACATTTTCTCGTTTGAAAGCCGCGCGTTGTCTGCCTGTTGTTCTGGCAGCCCTGATTTTCGCCGGTTGTGGCACCCATACTCCCGGA
TCCTCTAGATTTAAGAAGGAGATATACATATG
>trpR
GAGGCAATTCCGACGTCTAAGAAACCATTATTATCATGACATTAACCTATAAAAATAGGCGTATCACGAGGCCCTTTCGTCTTCACCTC
GAGTGTGAAGAACGTGCTGGCTTATGACGCTTACTACCGCTATTTCATGGGGGATAAACCGACGTTGATGAGCGCCACGGAATGGGGAC
GTCGTTACTGATCCGCACGTTTATGATATGCTATCGTACTCTTTAGCGAGTACAACCGGGGGAGGCATTTTGCTTCCCCCGCTAACAAT
GGCGACATATTATGGCCCAACAATCACCCTATTCAGCAGCGATGGCAGAACAGCGTCACCAGGAGTGGTTACGTTTTGTCGACCTGCTT
GGATCCTCTAGATTTAAGAAGGAGATATACATATG
>yliI
GAGGCAATTCCGACGTCTAAGAAACCATTATTATCATGACATTAACCTATAAAAATAGGCGTATCACGAGGCCCTTTCGTCTTCACCTC
GAGACCGTCAACGCCGATAAGCTTGAAAATTGTCTGATGAGCTTAACGCTGAGTATCCAGAGCCTGCAGAAGCACGCCATGCTTGAGAA
GGCCTGAAAACTAAGGGGGAGAAAGCGTCTCCCCCTTCATGTTTAAGTTTTTGTAAAAATGAATTTGTTATCTCCTCCACTGACTACGC
TTTAAGCCAGAGTCAATCCGGAGGCGTTATGCATCGACAATCCTTTTTCCTTGTGCCCCTTATTTGTCTTTCTTCCGCTCTCTGGGCGG
CTCCTGCAACGGTAAAGGATCCTCTAGATTTAAGAAGGAGATATACATATG
>fdoG
GAGGCAATTCCGACGTCTAAGAAACCATTATTATCATGACATTAACCTATAAAAATAGGCGTATCACGAGGCCCTTTCGTCTTCACCTC
GATGGATCCGTAAATCATCACGCCGCCATAACTCAATTTGACGAACACCTGTGATATTTGTCACATTTTCGATTTCTTTTCGCTGTGTT
TTTTTCACTTTCTCTTAACCAGATAAGAACACACACACCAACATTATGGTATTCTGTTACAAACCCTTCCTGGATGGAGGGAAATTGAG

CCAATTCTGGACCTTTGCGGCCCCTTCCGCAAAGAAAAATAACTCCCACTCCCTGCACACGCAGCAAGCGAATGTAAATGGGACGTGAC
AATGTCGAAACAAGGAGCAATCCATGCAGGTCAGCAGAAGGCAGTTCTTTAAGATCTGCGCTGGCGGTATGGCAGGCACCACGGCAGCG
GCACTGGGTTTTGCACCCAGCTCGAGAGATCCTCTAGATTTAAGAAGGAGATATACATATG
>rfaQ
GAGGCAATTCCGACGTCTAAGAAACCATTATTATCATGACATTAACCTATAAAAATAGGCGTATCACGAGGCCCTTTCGTCTTCACCTC
GATGGATCCATAGGCCGGAGCCTTACGTCCGCGCACCCAGAGCCGTATCCAGATCAGCGGCTGAATAAGGTAGAGAAGGGCGGTGTAAA
GCAATTCGAGCATAGTAAATAGCTGACTTATGGATGTGCTGGGGATTCTATGTATTTAGCTGTGGCTTTACCATTACTTTTCCCGTTTT
TGACTTAAATAGCTTCAGTTTGGTCTGATCTGCCGCTACATCTTCATTTTTTTTGTATTTTTATGCGATTCATTGAAACTCGGCCCCAT
TTTCAAATCTACATAGGCCGTACTGACATTATCGAAATGCTATTTTTTATCTATTTGATTTTTATGATTAAAGTATATTTTGTGTATAA
AAATCATTCGGGTCGGATTGCTGCGAAAGAAATGATACACTAGCACGTCAAAGTAAGTGCGTTATCAGTATTCAGGTAGCTGTTGAGCC
TGGGGCGGTAGCGTGCTTTTTTCTGCTTAACTTAACCAGACAATCACACAAAAGAGTCGCTAGTGGAAAAGCCATTTCGAAAAATCCTG
GTCATAAAGATGCGATATCATGGGGATATGTTATTAACTACTCCTGTCATCAGTACGCTCAAGCAGAATTATCCTGATGCAAAAATCGA
TATGCCTCGAGAGATCCTCTAGATTTAAGAAGGAGATATACATATG
>rfaZ
GAGGCAATTCCGACGTCTAAGAAACCATTATTATCATGACATTAACCTATAAAAATAGGCGTATCACGAGGCCCTTTCGTCTTCACCTC
GATGGATCCTTGAAAGGGAAACCAGCGCGCTAAAAAAAATCCCACAATTACATGTCTTCACCAGGACAGTTTTAAGCGGAGTTATTTAT
TTCACAGGTTTATCATGAAGAATATTAGATACATTGACAAAAAAGATGTTGAAAACCTGATTGAAAATAAAATATCTGATGATGTTATT
ATTTTTCTTTCGGGCCCTACATCGCAAAAAACACCTTTGTCAGTATTACGAACAAAGGATATTATTGCTGTCAATGGTTCTGCGCAACT
CGAGAGATCCTCTAGATTTAAGAAGGAGATATACATATG
>rfaD
GAGGCAATTCCGACGTCTAAGAAACCATTATTATCATGACATTAACCTATAAAAATAGGCGTATCACGAGGCCCTTTCGTCTTCACCTC
GAGCAGTCCAATCCCCTCTACCAATGTCAAAATATGCAGTGATAATTGTGGTGGATGATTTCATTACTAATAAGACCATTAATATAAAA
AAAGAATTATATCATAAATATTAATAGAATGTATACTGTATATTATCCATTCAAATGCTTCCGTTATTTATCATACATCGGAATATTGA
TACTAAAGCACTATTCACATGCAAAACCAACATCCGCCATGAAGGACTAGCTAAAACCCAAACTAGTTTGTTGCAATTAGCATCCTTGC
ACCTCTATGTAAAGGGCTGAAGGGATTCGGATGTGATGGTATGATTACAGACATTCGTGTCTGAGATTGTCTCTGACTCCATAATTCGA
AGGTTACAGTTATGATCATCGTTACCGGCGGCGCGGGCTTTATCGGCAGCAACATCGTTAAAGCCCTGAATGATAAAGGCATCACCGAT
ATTCTGGTGGTGGACAACCTGGGATCCCGGGATCCTCTAGATTTAAGAAGGAGATATACATATG
>rfaH
GAGGCAATTCCGACGTCTAAGAAACCATTATTATCATGACATTAACCTATAAAAATAGGCGTATCACGAGGCCCTTTCGTCTTCACCTC
GATGGATCCTGCTGTTCAAGCAGCGTCAAGAAGTCGCGTAAATCGTTATATTTCATGGCGTCCATTGTAGCCTCTTAATCTGCGCCCAT
TATACGGCGTTCATCTTTGCGATGCTGTAAATTTGTTAAATTAGCGTGAACTCTGACGGTATAACGCAAACCGGGGAATATAATTAACT
TAGCGTAAAGCTTTTGCTATCCTTGCGCCCCGATTAAACGGATAAGAGTCATTATGCAATCCTGGTATTTACTGTACTGCAAGCGCGGG
CAACTTCAACGTGCCCAGGAACACCTCGAAAGACAGGCTGTGAATTGCCTGGCACCGATGATCACCTCGAGAGATCCTCTAGATTTAAG
AAGGAGATATACATATG
>pstS
GAGGCAATTCCGACGTCTAAGAAACCATTATTATCATGACATTAACCTATAAAAATAGGCGTATCACGAGGCCCTTTCGTCTTCACCTC
GATGGATCCCGCGTAACCTGGCAAAATCGGTTACGGTTGAGTAATAAATGGATGCCCTGCGTAAGCGGGGCATTTTTCTTCCTGTTATG
TTTTTAATCAAACATCCTGCCAACTCCATGTGACAAACCGTCATCTTCGGCTACTTTTTCTCTGTCACAGAATGAAAATTTTTCTGTCA
TCTCTTCGTTATTAATGTTTGTAATTGACTGAATATCAACGCTTATTTAAATCAGACTGAAGACTTTATCTCTCTGTCATAAAACTGTC
ATATTCCTTACATATAACTGTCACCTGTTTGTCCTATTTTGCTTCTCGTAGCCAACAAACAATGCTTTATGAATCCTCCCAGGAGACAT
TATGAAAGTTATGCGTACCACCGTCGCAACTGTTGTCGCCGCGACCTTATCGATGAGTGCTTTCTCTGTGTTTGCAGAAGCAAGCCTGA
CAGGTGCAGGCTCGAGAGATCCTCTAGATTTAAGAAGGAGATATACATATG
>evgA
GAGGCAATTCCGACGTCTAAGAAACCATTATTATCATGACATTAACCTATAAAAATAGGCGTATCACGAGGCCCTTTCGTCTTCACCTC
GAGGGCACCTGAAAACGCAATAAATAAAACTACCGCCAATAAAGAAAAGTATTTTCTTCTGTTAGAATGTTTTTTATTTGAATTAATCT
GTTCCACTATTATCTCTCATTTCTCATAGATGAAATTTATGAATTGAATGAAAAGAGCATTCATTCTTCATATCAAACACAGTGCATGT
GTTGCACATTATCCCAGGAACGTTGGAGGCCTAATTTAAATGAGGTGCAAAAAAAACAACAGAGGACTAAACCGTGGCTTTTGCAATAC
AATTCTTACGCCTGTAGGATTAGTAAGAAGACTTATAGTGCCAACTTGAAACTATAAATCATCGGTACAATCCCTGATTTTATTGTTGA
CATTTCATTTATGCCGACTATTTATATGGTATACTTGTCGAATTATCTTAAAGGAAGCTCAGATTTTCTTATTTTTATTGAGAAAATGA
GATGACGCCTTATGTCTGTATTACTACAGGGAGAAGGGAGATGCTTCATTGCAAAGGGAATAATCTATGAACGCAATAATTATTGATGA
CCATCCTCTTGCTATCGCAGGATCCTCTAGATTTAAGAAGGAGATATACATATG
>envY
GAGGCAATTCCGACGTCTAAGAAACCATTATTATCATGACATTAACCTATAAAAATAGGCGTATCACGAGGCCCTTTCGTCTTCACCTC
GATGGATCCGCCTCGCTGTTAGCGAAACGTTCCTCTTGCGGTAAAGACACTCTCTATTTCTCTCTGCGCTATTTGCCCGTCGCGCACGG
CATTCTCGAGTATTAAATCTCCTCCAGGTAAGTCGGGTACGACCTGGCTTACCTCTTTCGCTCTCATCAGAATTCCATCAAGATTAATT
GCTAAAAAGCGTGCTAAAGAAAGAAGATCGTTATCGGCGAACGGGCCGATCACTCATGCAGGAAGCATAGATCCTTTGAGCGGAGTTTC
TGATATCTGAGGAGTGCGAAATGCAATTGAGCAGCAGTGAACCTTGCGTGGTGATCCTGACCGAAAAAGAGGTAGAGGTAAGCGTCAAT
AACCATGCTACGTTTACCCTTCCGGTCGAGAGATCCTCTAGATTTAAGAAGGAGATATACATATG
>ydeV
GAGGCAATTCCGACGTCTAAGAAACCATTATTATCATGACATTAACCTATAAAAATAGGCGTATCACGAGGCCCTTTCGTCTTCACCTC
GATGGATCCCTGCAATGAAAGGCGGTTATATCAACGCACTGGTTACCGATCAGGACACAGCAGCGGCGATTTTACGTAGTTAAATTTGC
ATGACCTTATTACCGGGAATGATGGAGAACGTATAGAGGAACAGACAGAACGCATAAGCCGAGGATAATCTAATGGCTCGACTCTTTAC

CCTTTCAGAATCAAAGTACTACCTGATGGCGCTGGATGCAGGCACCGGAAGTACTCGAGAGATCCTCTAGATTTAAGAAGGAGATATAC
ATATG
>ydeP
GAGGCAATTCCGACGTCTAAGAAACCATTATTATCATGACATTAACCTATAAAAATAGGCGTATCACGAGGCCCTTTCGTCTTCACCTC
GATGGATCCCGGAACCGTCCAGTCAGTCATTGGTGTCACTTTTATCTATGAATAATTGTGATTTTGACCTTAGTCTATGCGCTCCATTC
ACGAAAATTGCGCGGATAATTATTTTGTGAAGGCTATTAGCCTACACCTGTAAGAAAATCCGCGCACAATTGAAGCCAGAGTCTTATTT
TTTATATTGAAAAATATGTTAATTTAAACACTATGAGAGATGATGCTTAACTTATCGTATTTAATCTATTGTTTAACGATAATAGATTA
AAAATAGAAAGAGAGTAGTCCCCTACTCTTATATATCCATGTTGGCGATAATCCCTTTGTATGTACTGTGCATCATCGCTATTACAAAT
CCTAATAATTCATTTCCACACAGGATAAGTAGATGAAGAAAAAAATTGAATCCTACCAGGGTGCTGCAGGTGGTTGGGGTGCTGTTAAA
TCCGTAGCGAATGCAGTACGTAAGCACTCGAGAGATCCTCTAGATTTAAGAAGGAGATATACATATG
>ydeQ
GAGGCAATTCCGACGTCTAAGAAACCATTATTATCATGACATTAACCTATAAAAATAGGCGTATCACGAGGCCCTTTCGTCTTCACCTC
GATGGATCCGAACGATCGAGGCGCTAATCAATGTGATCTACACCTGGCAATAACGTCGATTCGATTTTCTTAGCCTTCTTTTTAAATCA
ATTTTTAACCTTAGGATATATCATGGGTAAAACAATCAGTATAAAGGTCCTGTTCGGTATTTATCTTCTATTAATGGCAGGCAAAGTTT
TTGCCTTCAGTTGTAATGTCGATGGCGGGTCCTCGAGAGATCCTCTAGATTTAAGAAGGAGATATACATATG
>btuC
GAGGCAATTCCGACGTCTAAGAAACCATTATTATCATGACATTAACCTATAAAAATAGGCGTATCACGAGGCCCTTTCGTCTTCACCTC
GATGGATCCAAAAGCCGGGTCGAAAACGCTTCGCCCAAAGACGAGTAATCTGATCTAACTAAAAAGGCCGCTCTGCGGCCTTTTTTCTT
TTCACTGTCGAAGAGTCACCGTAAAATCAACGCCATGACACTTCAGCAGAACGGATACCATGCTGACACTTGCCCGCCAACAACAGCGA
CAAAATATTCGCTGGTTATTATGCCTGTCAGTTTTGATGCTGCTGGCGCTTCTCTTAAGCCCTCGAGAGATCCTCTAGATTTAAGAAGG
AGATATACATATG
>btuB
GAGGCAATTCCGACGTCTAAGAAACCATTATTATCATGACATTAACCTATAAAAATAGGCGTATCACGAGGCCCTTTCGTCTTCACCTC
GAGGGTCAGAAAACGGTGCCATCATACTTTGCAAACGTACCACTTTTTCGGCTAACTGCGCTTCATACTGTTCTGTTGGAAGGTGTTCG
GGGGTCATGATGTATCCTGAAAAATTAAAGTACGCGGCGATTGTAGGGATTGCTCATCAGATGTCCAGATCTTGATGAATTCCTATTTG
TGAGCTACGTCTGGACAGTAACTTGTTACAACCTGTAGCATCCACTTGCCGGTCCTGTGAGTTAATAGGGAATCCAGTGCGAATCTGGA
GCTGACGCGCAGCGGTAAGGAAAGGTGCGATGATTGCGTTATGCGGACACTGCCATTCGGTGGGAAGTCATCATCTCTTAGTATCTTAG
ATACCCCTCCAAGCCCGAAGACCTGCCGGCCAACGTCGCATCTGGTTCTCATCATCGCGTAATATTGATGAAACCTGCGGCATCCTTCT
TCTATTGTGGATGCTTTACAATGATTAAAAAAGCTTCGCTGCTGACGGCGTGTTCCGTCACGGCATTTTCCGCTTGGGCACAGGATACG
GATCCTCTAGATTTAAGAAGGAGATATACATATG
>pstC
GAGGCAATTCCGACGTCTAAGAAACCATTATTATCATGACATTAACCTATAAAAATAGGCGTATCACGAGGCCCTTTCGTCTTCACCTC
GATGGATCCGTTGAACAGGTTCGCGCTGCGTGGAAGACCAATATTAAAGACAGTAGCGGTAAGCCGCTGTACTAATAAAACTCCAGGCC
GGGTACGGTGTTTTACGCCGCATCCGGCATTACAAAATGACTTTGTAAACGCGTTTAACTGAAGAGTAACTTATGGCTGCAACCAAGCC
TGCTTTTAACCCACCGGGTAAAAAGGGCGACATAATTTTCAGCGTGCTGGTAAAACTGGCGGCTCGAGAGATCCTCTAGATTTAAGAAG
GAGATATACATATG
>moeA
GAGGCAATTCCGACGTCTAAGAAACCATTATTATCATGACATTAACCTATAAAAATAGGCGTATCACGAGGCCCTTTCGTCTTCACCTC
GATGGATCCGCAATGACTGCTTTGCCCATAGTATTCGTCCATTAAGGGTGGTTCGCCACTTTTGCTATAAATATCAGAATTTTTATGAA
TTACTCATTCAATTTTTGAATATAGAAAGATGGATATGGCATGTAAAGGCAGGACCCAGCGCTACGCCCGAGGTGGCATACAGTCTGTG
GCGTTTTTCTGACATAATAGGCAAATTCGATTTTGCCTCCGCAGGAGTGTTTTCATGGAATTTACCACCGGATTGATGTCGCTCGACAC
CGCGCTTAATGAGATGCTTTCTCGCGTCACCCCACTGACCGCCCAGGAAACCTCGAGAGATCCTCTAGATTTAAGAAGGAGATATACAT
ATG
>ydeO
GAGGCAATTCCGACGTCTAAGAAACCATTATTATCATGACATTAACCTATAAAAATAGGCGTATCACGAGGCCCTTTCGTCTTCACCTC
GATGGATCCATGGGTCGCATTGCAAGAGATCTTACACATTTCATTATCAATAGTTTGCAAGGCTGAAAACAAACAGCAAATTATAAATA
TGAAATGTTAAAAAAGTATCGATAAAAACTTTATTGTTTTAAGGAGATAAAATGTCGCTCGTTTGTTCTGTTATATTTATTCATCATGC
CTTCAACGCTAACATTTTAGATAAAGATTACGCCTTCTCTGACGGCCTCGAGAGATCCTCTAGATTTAAGAAGGAGATATACATATG
>ydeI
GAGGCAATTCCGACGTCTAAGAAACCATTATTATCATGACATTAACCTATAAAAATAGGCGTATCACGAGGCCCTTTCGTCTTCACCTC
GATGGATCCCGCGGATGTCAGGTTTTCAATAGTATCGGTATCTACAAGTTGTACAATTTTATCTCTATTAATGTTCACTGTTTTTTGTC
CACGCTTTTATATGTAAAGGTGATTTGCTTTTTCTGTATGCTCGTCCAATATAATAATTTTCAAACATATAAATCCGATTATTCTGAAA
ATATGTTTAAGACTTTTCAATCTTCTTAACTTTAGTGAGTTAAATTCTCTGTATATTTGACAGCTAGTTGGCTGATTTTAAATGCCTTA
TAACAATTGGCAATATTCTTATTTGGCAATTATAAAAATAGCAACCAGACTTAATAAGTCAAACCAAACGCAATTAAGGAGTTAGTATG
AAATTTCAGGCCATTGTATTAGCAAGTTTTCTTGTCATGCCTTATGCGTTGGCAGATGACCAGGGCGGTTTAAACTCGAGAGATCCTCT
AGATTTAAGAAGGAGATATACATATG
>ydeJ
GAGGCAATTCCGACGTCTAAGAAACCATTATTATCATGACATTAACCTATAAAAATAGGCGTATCACGAGGCCCTTTCGTCTTCACCTC
GAGTTTAAACCGCCCTGGTCATCTGCCAACGCATAAGGCATGACAAGAAAACTTGCTAATACAATGGCCTGAAATTTCATACTAACTCC
TTAATTGCGTTTGGTTTGACTTATTAAGTCTGGTTGCTATTTTTATAATTGCCAAATAAGAATATTGCCAATTGTTATAAGGCATTTAA
AATCAGCCAACTAGCTGTCAAATATACAGAGAATTTAACTCACTAAAGTTAAGAAGATTGAAAAGTCTTAAACATATTTTCAGAATAAT
CGGATTTATATGTTTGAAAATTATTATATTGGACGAGCATACAGAAAAAGCAAATCACCTTTACATATAAAAGCGTGGACAAAAAACAG

TGAACATTAATAGAGATAAAATTGTACAACTTGTAGATACCGATACTATTGAAAACCTGACATCCGCGGGATCCTCTAGATTTAAGAAG
GAGATATACATATG
>ydeA
GAGGCAATTCCGACGTCTAAGAAACCATTATTATCATGACATTAACCTATAAAAATAGGCGTATCACGAGGCCCTTTCGTCTTCACCTC
GAGAAAAACAGGCCTCGCTCCCCTGTCAGACCACCTATACCCGCAACCGCCAAAACGCCAGACCTCCCTGAACGTCATTAAACCGTGAT
GTTACCGACTCTCTGACGCGTGAAAGAATCAGCGTCAGAGAAACGGAAAACGCGATCCAGATCACAAATGCATTGTATTCACATCATTA
ACCGTTTTAAGATCATTTCATCACTTTTTCGCAACTCACCCGATAATCTGTTATGACAACAAACACTGTTTCCCGCAAAGTGGCGTGGC
TACGGGTCGTTACGCTGGCAGTCGCCGCCTTCATCTTCAACACCACCGAATTTGTCCCTGGGATCCTCTAGATTTAAGAAGGAGATATA
CATATG
>yhdX
GAGGCAATTCCGACGTCTAAGAAACCATTATTATCATGACATTAACCTATAAAAATAGGCGTATCACGAGGCCCTTTCGTCTTCACCTC
GAGCAGAAAGCCCGCTGAAAATTAAACGTGGGCAAAATAATCTCTGGAATAACGGCGGTATTCAGTACGCACCGCCCGTGCGTTAAGTC
GGTGTGAGGTAACGAGCGTCGCTACGGCGGCGCTCTCTTCAGCGTTATGCTTTCCGAGGTTTCTATGTCTCATCGCCGCTCAACCGTTA
AAGGCTCACTCTCTTTTGCCAACCCTACGGTTCGCGCCTGGTTATTCGGATCCTCTAGATTTAAGAAGGAGATATACATATG
>yaeQ
GAGGCAATTCCGACGTCTAAGAAACCATTATTATCATGACATTAACCTATAAAAATAGGCGTATCACGAGGCCCTTTCGTCTTCACGTC
GACTTAAATGATGCTGGCAGGCGAGCTCAAGATTATCGTAATCATCACAATTGATTGGTTGATACGTATCATTCATTGACATAATCGCT
CACCAGTAAGTTTGCCGCAGCGTATGCTGCTTTTTCCCTGACAGCCTCAGAAAGGGCGTCGTCGGCAGCCATTTCATTCAGCACTTTCA
AAACGCAGCCCAGCGCGTCCGGAACGTATCCTAAGTCTCCGCTGGCGATTTCCGCGTACCGCTTGCGTATTAACTCACAATATTTTTCC
ACATGCCCTCCTGTCAGCACTCTGACTTAACCGTGGATGCAAGTCTAAGCCTACGAAGATAAACTCTGTTTCGCAAGGTGACTATACCA
CACTCATTTCTGCAATATCAGCGCCGCAACTGCACGTATTCCGTTACAATGGCCTCCTGATTCGAAAGGAGTTTTCTTATGGCGCTTAA
AGCGACAATTTATAAAGCGACGGTTAATGTGGCCGATCTCGACCGCAACCAGTTTCTCGATGCCTCTCTGACGCTGGGGATCCTCTAGA
TTTAAGAAGGAGATATACATATG
>yfiF
GAGGCAATTCCGACGTCTAAGAAACCATTATTATCATGACATTAACCTATAAAAATAGGCGTATCACGAGGCCCTTTCGTCTTCACCTC
GATGGATCCTTTTGCCGCATCTTCGATCCGATCGTCGGGAATGCGATTGATGGCCTGACAATGGGTACAAACGGTATTCATAACTAACC
TCGGGATGAGTAAGGAGACTTATCGGCAACGACGCCGGTATGTTTCTAATATGTTACATATTATCGGGGACATTGTTCCAGACAACAAT
CTGTTTTATTCAATAGGTATGATAGTCGCAGGCTTTGGTACGAAATCATGGCGAACCGCGCGGACTTCGGGTAATCTGCGCGCTTCGCG
CAGCGCTGGTGGAGAAAAGCATGAACGATGAAATGAAAGGTAAAAGCGGCAAGGTCAAAGTGATGTATGTCCGCAGTGATGATGATTCT
GATAAACGTACCCACAACCCGCGTACCTCGAGAGATCCTCTAGATTTAAGAAGGAGATATACATATG
>feaB
GAGGCAATTCCGACGTCTAAGAAACCATTATTATCATGACATTAACCTATAAAAATAGGCGTATCACGAGGCCCTTTCGTCTTCACCTC
GAGAACGCTCAGTCAGCAGGCGTCCGGTAAAATTTCCGCATACCTGATTGATTTGGGAAAGCCATTGCTGAAACTCATTATCCACTGCG
GGGTTCATGGCACGTTTTCGCTCTGTGAAATGTATTTTTATTGTTGCATTTGTGTTGCAATAAACGAAGCTAATGAGCCTGACTATAGG
AAATAAGTCTTGTCAGGCATAGAGACATAAGCGGTTATTGTCACGATTTGCGGAGCTTGTCACAGCTGACAAAGCGAATGTCACAGCGA
AAAAAGTGACTTTTCTTGTCGCTGCGTACACTGAAATCACACTGGGTAAATAATAAGGAAAAGTGATGACAGAGCCGCATGTAGCAGTA
TTAAGCCAGGTCCAACAGTTTCTCGATCGTCAACACGGTCTTTATATTGATGGTCGTCCTGGCGGATCCTCTAGATTTAAGAAGGAGAT
ATACATATG
>yfiE
GAGGCAATTCCGACGTCTAAGAAACCATTATTATCATGACATTAACCTATAAAAATAGGCGTATCACGAGGCCCTTTCGTCTTCACCTC
GATGGATCCTATTGTTCGGTCCTGGCGTCATAGCGGTAATCAGGGTGTAAGTCCAAAAAGCACTTAAAAGGGTCGGTGTCACTTGTTAT
CTCCTCTGTTTTGGTAGACAGTACCCGACACGGAACATTTAAAAAAATGAATGTTTTTATATTCTAGATTCGATATTTTCGATAGGTTT
GGGGTTATGGATCTGCGCCGTTTTATTACGCCTCGAGAGATCCTCTAGATTTAAGAAGGAGATATACATATG
>yfiC
GAGGCAATTCCGACGTCTAAGAAACCATTATTATCATGACATTAACCTATAAAAATAGGCGTATCACGAGGCCCTTTCGTCTTCACCTC
GATGGATCCTTCCAGCAGGCTTTCGTCGAGTTCAAGTTCGGAAAAAGTCGTTACAGTCATGTTCTACCTCTGTGTGGGGCGCTGATTAT
AGACGTTACGGCTGCAATCTTCATCTGTTTGTATGGATATCGCTTCTCGGGTATTGTTCTCACCCGGTGCTATCGCCGTTTTTCCTGCC
TCGAGAGATCCTCTAGATTTAAGAAGGAGATATACATATG
>nagE
GAGGCAATTCCGACGTCTAAGAAACCATTATTATCATGACATTAACCTATAAAAATAGGCGTATCACGAGGCCCTTTCGTCTTCACCTC
GAGATCGGCAGTCGGTTTGAACGCATTGATACGATTGACGATATGGCGAGCAGCCCATTTGCCGACCTGTTCAGCGGTAGTCAGGGGGA
TCAGTCTCATTATTCACCTCAATAAGTAAAATGTAAGCCGTTGGCGGATTAGGCATCTTTAAGCGTAACCTGGATTTGCGCAGACAGGC
GTCAATCCGACCTGATTTTTTGAATGATAAAATAAGTTTTCTGGTTTAGCCAGTAAAAGGGAGTGATAATAACGATATTTGGTGACAAA
ACTCACAAAAGACACGCGTTTAATTTGCGATACGAATTAAATTTTCACACACTCTGTAGCAGATGATCTAACAATCTGATTACAGAACA
TCGGCAGTACAATTTGCAGCAAAATAAAAATACGGCTTGAAACGAGCCAAATAGGGTTCTCGTAGGGGGAATAAGATGAATATTTTAGG
TTTTTTCCAGCGACTCGGTAGGGCGTTACAGCTCCCTATCGCGGTGCTGCCGGTGGCGGCACTGTTGCTGCGATTCGGGATCCTCTAGA
TTTAAGAAGGAGATATACATATG
>nagB
GAGGCAATTCCGACGTCTAAGAAACCATTATTATCATGACATTAACCTATAAAAATAGGCGTATCACGAGGCCCTTTCGTCTTCACCTC
GATGGATCCCCTACCGAGTCGCTGGAAAAAACCTAAAATATTCATCTTATTCCCCCTACGAGAACCCTATTTGGCTCGTTTCAAGCCGT
ATTTTTATTTTGCTGCAAATTGTACTGCCGATGTTCTGTAATCAGATTGTTAGATCATCTGCTACAGAGTGTGTGAAAATTTAATTCGT
ATCGCAAATTAAACGCGTGTCTTTTGTGAGTTTTGTCACCAAATATCGTTATTATCACTCCCTTTTACTGGCTAAACCAGAAAACTTAT
TTTATCATTCAAAAAATCAGGTCGGATTGACGCCTGTCTGCGCAAATCCAGGTTACGCTTAAAGATGCCTAATCCGCCAACGGCTTACA

TTTTACTTATTGAGGTGAATAATGAGACTGATCCCCCTGACTACCGCTGAACAGGTCGGCAAATGGGCTGCTCGCCATATCGTCAATCG
TATCAATGCGTTCAAACCGACTGCCGATCTCGAGAGATCCTCTAGATTTAAGAAGGAGATATACATATG
>nagA
GAGGCAATTCCGACGTCTAAGAAACCATTATTATCATGACATTAACCTATAAAAATAGGCGTATCACGAGGCCCTTTCGTCTTCACCTC
GATGGATCCCCTTCCACCATGGAGCTGAAAGTTAAGACTTTAAGATATTTCAATGAATTAGAAGCAGAAAATATCAAAGGTCTGTAATT
GTTATCCCTGCCCTGGCTCCTTGCTCAGGGCAATATTTTTTAAAATCGGGGGTCAGAATGTATGCATTAACCCAGGGCCGGATCTTTAC
CGGCCACGAATTTCTTGATGACCACGCGGTTGTTATCGCTGCTCGAGAGATCCTCTAGATTTAAGAAGGAGATATACATATG
>feaR
GAGGCAATTCCGACGTCTAAGAAACCATTATTATCATGACATTAACCTATAAAAATAGGCGTATCACGAGGCCCTTTCGTCTTCACCTC
GATGGATCCCATGCGGCTCTGTCATCACTTTTCCTTATTATTTACCCAGTGTGATTTCAGTGTACGCAGCGACAAGAAAAGTCACTTTT
TTCGCTGTGACATTCGCTTTGTCAGCTGTGACAAGCTCCGCAAATCGTGACAATAACCGCTTATGTCTCTATGCCTGACAAGACTTATT
TCCTATAGTCAGGCTCATTAGCTTCGTTTATTGCAACACAAATGCAACAATAAAAATACATTTCACAGAGCGAAAACGTGCCATGAACC
CCGCAGTGGATAATGAGTTTCAGCAATGGCTTTCCCAAATCAATCAGGTATGCGGAAATTTTACCGGACGCCTGCTGACTGAGCGTTCT
CGAGAGATCCTCTAGATTTAAGAAGGAGATATACATATG
>yaeH
GAGGCAATTCCGACGTCTAAGAAACCATTATTATCATGACATTAACCTATAAAAATAGGCGTATCACGAGGCCCTTTCGTCTTCACCTC
GATGGATCCGGCGTGGGCAGTTTGTATGGTTTGCGTCTGAACTGCCGCCCGGAAGTAACGATGCTGGAACTGGTGTAGATCGTGAATTT
TATCACCAGAATAAAGCAGGTAATTTTTTGATTTTTACGCCATGATAGAAGTCTGGAAGAAAAAGGCTTTCTGCGATCACAGTCGAGCC
GGGCGGGCCAGGCCAAATGCTAAAATAAGCACAATTTACTGACTCATTTTGAGGTAACGCTATGTACGACAATCTGAAAAGTCTGGGTA
TTACCAATCCTGAAGAAATTGATCGTTACAGCCTCCGGCAGGAAGCCAACAACGACTCGAGAGATCCTCTAGATTTAAGAAGGAGATAT
ACATATG
>uidC
GAGGCAATTCCGACGTCTAAGAAACCATTATTATCATGACATTAACCTATAAAAATAGGCGTATCACGAGGCCCTTTCGTCTTCACCTC
GATGGATCCGGTTTTATCCGCTCACGGATAAAAAATTCAAAGAAATCGTGGTTGAAATTGATAATCGTAAAAAAGTGCAGCAGCAATTA
ATCAGCGATATCACTAATTAATATTCAATAAAAATAATCAGAACATCAAAGGTGCAACTATGAGAAAAATAGTGGCCATGGCCGTTATT
TGCCTGACGGCTGCCTCTGGCCTTACCTCTGCTTATGCGGCGCAACTGGCTGACGATGAAGCGGGACTACGCATCAGCTCGAGAGATCC
TCTAGATTTAAGAAGGAGATATACATATG
>uidA
GAGGCAATTCCGACGTCTAAGAAACCATTATTATCATGACATTAACCTATAAAAATAGGCGTATCACGAGGCCCTTTCGTCTTCACCTC
GATGGATCCGCGATGATTGGCGGTATCTTAACCGCATCCTGATTCTCTCTCTTTTTCGGCGGGCTGGTGATAACTGTGCCCGCGTTTCA
TATCGTAATTTCTCTGTGCAAAAATTATCCTTCCCGGCTTCGGAGAATTCCCCCCAAAATATTCACTGTAGCCATATGTCATGAGAGTT
TATCGTTCCCAATACGCTCGAACGAACGTTCGGTTGCTTATTTTATGGCTTCTGTCAACGCTGTTTTAAAGATTAATGCGATCTATATC
ACGCTGTGGGTATTGCAGTTTTTGGTTTTTTGATCGCGGTGTCAGTTCTTTTTATTTCCATTTCTCTTCCATGGGTTTCTCACAGATAA
CTGTGTGCAACACAGAATTGGTTAACTAATCAGATTAAAGGTTGACCAGTATTATTATCTTAATGAGGAGTCCCTTATGTTACGTCCTG
TAGAAACCCCAACCCGTGAAATCAAAAAACTCGACGGCCTGTGGGCATTCAGTCTGGATCGCGAAAACTGTGGCTCGAGAGATCCTCTA
GATTTAAGAAGGAGATATACATATG
>aes
GAGGCAATTCCGACGTCTAAGAAACCATTATTATCATGACATTAACCTATAAAAATAGGCGTATCACGAGGCCCTTTCGTCTTCACCTC
GATGGATCCCAGCGCTGGTTTCGTTTTCTGGCTGGAATTGCTGAAGCAGCGGATCGCGTGCGTTTACGGGGAAGTAATGTTTGGATTTA
CGTTTACCGGGAAATTTCATGGTTGTTTTTACGGGTGGTTAACTGAGCGGCGAATGTTAACACAATCTGACGGGAGCGCGCGAGACGCA
GCCAGTTTCAATATCAGCAGGCGGGATCGCATTTTCATGGCAGTGGTTCCTTACAATGACGTAATTTGAAAGGAGTTTTTGTATGAAGC
CGGAAAACAAACTACCTGTTCTGGACCTTATTTCTGCTGAAATGAAGACCGTTGTGAATACTCTTCAGCCGGATTTACCGCTCGAGAGA
TCCTCTAGATTTAAGAAGGAGATATACATATG
>aer
GAGGCAATTCCGACGTCTAAGAAACCATTATTATCATGACATTAACCTATAAAAATAGGCGTATCACGAGGCCCTTTCGTCTTCACCTC
GATGGATCCTCGCGAAAATCGGGATAAACACAAATTCGGTTATCATTTGTGCAATATTTAATGCTATCTGTTAACATTTGTTGTTATTT
TGCACGGTTTTTGTCGGGTATGGATTTGATGATTGCTGCGCAAATGCACGTTTTCAGGGCAATAATTGCCTGATTTTTGGGCAAGATAA
TTAACGTTTTCGTATAGTAACAGAGTGAAAGTATTTTAAATCAAGGAATTAGATTTGTGTTTAAATCGCAAATTGCGATCTAAATCAAA
TTAATCGGTTAAAGATAACCGCAGCGGGGCCGACATAAACTCTGACAAGAAGTTAACAACCATATAACCTGCACAGGACGCGAACATGT
CTTCTCATCCGTATGTCACCCAGCAAAATACCCCGCTGGCGGACGATACCACTCTGATGTCCACTACCGATCTGCACTCGAGAGATCCT
CTAGATTTAAGAAGGAGATATACATATG
>dps
GAGGCAATTCCGACGTCTAAGAAACCATTATTATCATGACATTAACCTATAAAAATAGGCGTATCACGAGGCCCTTTCGTCTTCACCTC
GATGGATCCCCGCTTCAATGGGGTCTACGCTGACAGTACGCAAAGAGAGCAAAATAAAAGAATTAGACATTAATTAAATTTACATTTCT
GCATGGTTATGCATAACCATGCAGAATTTCTCGCTACTTTTCCTCTACACCGTCTTTATATATCGAATTATGCAAAAGCATATTTATTC
CGAAAATTCCTGGCGAGCAGATAAATAAGAATTGTTCTTATCAATATATCTAACTCATTGAATCTTTATTAGTTTTGTTTTTCACGCTT
GTTACCACTATTAGTGTGATAGGAACAGCCAGAATAGCGGAACACATAGCCGGTGCTATACTTAATCTCGTTAATTACTGGGACATAAC
ATCAAGAGGATATGAAATTATGAGTACCGCTAAATTAGTTAAATCAAAAGCGACCAATCTGCTTTATACCCGCAACGATGTCTCCGACA
CTCGAGAGATCCTCTAGATTTAAGAAGGAGATATACATATG
>b0725
GAGGCAATTCCGACGTCTAAGAAACCATTATTATCATGACATTAACCTATAAAAATAGGCGTATCACGAGGCCCTTTCGTCTTCACCTC
GAGTATGTCCGAAGGGGCTGAACCCGACGCGCGCCATCGGCCATATCAAGTCGATGTTGTTGCAACGTAATGCGTAAACCGTAGGCCTG

```
ATAAGACGCGCAAGCGTCGCATCAGGCAACCAGTGCCGGATGCGGCGTGAACGCCTTATCCGGCCTACAAGTCATTACCCGTAGGCCTG
ATAAGCGCAGCGCATCAGGCGTAACGGATCCTCTAGATTTAAGAAGGAGATATACATATG
>aroP
GAGGCAATTCCGACGTCTAAGAAACCATTATTATCATGACATTAACCTATAAAAATAGGCGTATCACGAGGCCCTTTCGTCTTCACCTC
GATGGATCCGTTGGCGGATTTTGCTGTAGGCCATGAGTTCCTGTCTTAAGCCACTTGCCGAAGTCAATTGGTCTTACCAATTTCATGTC
TGTGACGCTAAAGTAACAAAGTATTCACCTTATGTCCATACAGGTTTTGATTGAAATCATGAAACTGTGCACATTTTAACAACTTGACA
TATATAACGTTTCAAAGTTGTAACTATGCACAAATGTAGACTTTACGTAGGAAAGGAGTTTTTTAACGATTATTAAACTATAAAAATCC
GAATTGAACCGATTCACTTACCAATTTTGTGATTTTTAATTCAATTAAAACGAATTTAAATTCATTCTACATATTGAGAGGGGTTGAGG
CTGAGCTTTACAAACGGTTTCTTTTTAAGCAACTCATCTTCAACCATGCATAAAGCGGGTGCATTCGCTGCCGCATACCATTATTCTTG
ATCTGACGGAAGTCTTTTTGTAACAATTCAAACTTCTTTGATGTAAACAAATTAATACAACAAACGGAATTGCAAACTTACACACGCAT
CACTGCGTAGATCAAAAAAACAACCACCGCACGAGGTTTCATGATGGAAGGTCAACAGCACGGCGAGCAGCTAAAGCGCGGCCTTAAAA
ACCGCCATATTCAGCTTATCGCGCTGGGTGGCGCGATAGGGACCGGGTTACTCGAGAGATCCTCTAGATTTAAGAAGGAGATATACATA
TG
>tyrP
GAGGCAATTCCGACGTCTAAGAAACCATTATTATCATGACATTAACCTATAAAAATAGGCGTATCACGAGGCCCTTTCGTCTTCACCTC
GAGTCCAGACTGGCATGCGTATATTGCTCGCCTGATTCAATCATCATATTTAACACTTCATGACCGTGAATAGAGTCCATCGTCCCTCC
TCAAAAAAAGCCTAGCGTAGCGATTGCCGCTTATGAAGACTTTGCGCCAGCGCAGGACTGAATGCTTTTTATTGTACATTTATATTTAC
ACCATATGTAACGTCGGTTTGACGAAGCAGCCGTTATGCCTTAACCTGCGCCGCAGATATCACTCATAAAGATCGTCAGGACAGAAGAA
AGCGTGAAAAACAGAACCCTGGGAAGTGTTTTTATCGTGGCGGGAACCACAATTGGCGCAGGCATGCTGGCAATGCCGCTGGCTGGATC
CTCTAGATTTAAGAAGGAGATATACATATG
>yfgA
GAGGCAATTCCGACGTCTAAGAAACCATTATTATCATGACATTAACCTATAAAAATAGGCGTATCACGAGGCCCTTTCGTCTTCACCTC
GATGGATCCAAACGGATGCAGGGTGAAGCCATCGACATTAAAGCGGTCTGATAGCCATTACGCCACGGTTACTTTATGTACCGTGGCGT
AATGTTTTATGAATGAAGTATTAAGTTGCAATTGTCTGGTCAATATTAACGGTGCGTTTTTGCTGACTTTAAGGCAGTATGTAACGATG
CAACAGTAAGTTAGCCTTAGTGAATGTGGGCTTTGTCACGAGCACACAGACGGTCTTATACTGTATGATAACGGTTAACTTAACGGATG
TTTCGCGGTGTGGGTGAGCATGATGGTTCACCGGCATCTCAATTCTCATTTAAACGTACCTGCAGCGAATGAATACTGAAGCCACGCAC
GACCAAAATGAAGCACTTACTACCGGCGCTCGCCTGCGTAATGCTCGCGAACAACTCGAGAGATCCTCTAGATTTAAGAAGGAGATATA
CATATG
>yfgB
GAGGCAATTCCGACGTCTAAGAAACCATTATTATCATGACATTAACCTATAAAAATAGGCGTATCACGAGGCCCTTTCGTCTTCACCTC
GATGGATCCACGGTTCTGATTCCGTCGAATCTGCCGCTCGCGAAATCGCTTATTTCTTTGGCGAAGGCGAAGTGTGCCCGCGCACCCGT
TAATAATTTCGTAAATGCCGCGTGCAAACGTGGCATCCGTGCGCCAGAATTTGTACAATGCAGCGCCCCCGGACGAGCAGCCGCTCACC
GGGGCGTTTCTTTTTTCAACCCTCCAGGGGCCATAACGTGTAATAACGAGGCCGGAATAGCATATGTCTGAACAATTAGTCACACCTGA
AAACGTCACCACGAAAGATGGAAAAATCAACCTGCTGGATCTCAACCGTCAGCAGATGCGGGAGTCTCGAGAGATCCTCTAGATTTAAG
AAGGAGATATACATATG
>yfgH
GAGGCAATTCCGACGTCTAAGAAACCATTATTATCATGACATTAACCTATAAAAATAGGCGTATCACGAGGCCCTTTCGTCTTCACCTC
GAGGCAGCACCATCAGAGCAAAAAAGAAGCTCAGCAATCCACACTCTCCGTCGAATCACCGGTACAACGTTAGCGGTTACCTTCTCCAC
TTTCACAGAACATAACGGCACTTCGCTGTCGGATGCTTTTGCTGTTTGGGATTATCAAAGCGGCAGATATTCTTTCATCTTAAATTTTA
CGTCTTTATCCTGACTGATGTTTATCCTGTTTGGCTGCGAAATAAATATAAAATTAATATATATGTTGTAATGATATATTTTTATAAAT
TATTCCCTGCGTGAATTTTAATAAATTTAATCTATCCCTTTATACGCAATACATTTACTTTCCTCTTTTGATGATCTTAAATGTCTTAT
TTTTCGTAATGTGTATAACAAGGAATAGTGATGAAATTTAAAAAATGTCTTCTGCCTGTGGCAGGATCCTCTAGATTTAAGAAGGAGAT
ATACATATG
>yheO
GAGGCAATTCCGACGTCTAAGAAACCATTATTATCATGACATTAACCTATAAAAATAGGCGTATCACGAGGCCCTTTCGTCTTCACCTC
GATGGATCCGAAGCTGATGCGAAAGCCGCAGATTCTGCTAAAAAATAAGCATTAAGAACCGCCGCCTGACCAGGCGGCGGTTTTTTTAT
TACAGGCCGGATATAATTAGTGCTGGAAAGCGGAACCTCCGCTGTATTAATTTAGTTACCCGCATCATTAATGAGCCTGCCCTGAAAAG
TTAACGACAGGCTCCTGAAAAGGAGTGTTTTTTTTCATGTCCAGGTCGCTTTTAACCAACGAAACCAGTGAGTTGGATTTACTGGATCA
ACGTCCTTTCGACCAGACCGACTCGAGAGATCCTCTAGATTTAAGAAGGAGATATACATATG
>tyrB
GAGGCAATTCCGACGTCTAAGAAACCATTATTATCATGACATTAACCTATAAAAATAGGCGTATCACGAGGCCCTTTCGTCTTCACCTC
GAGGCCCGTAGAACGTATCGCTGAAATGACGAAAGTAAGCGCTTACGAACTTATTACGCGCCTGACTTCAAGGGTCGCGATGAAATACG
TGGATTAATCGTTCTGTAATATTTGATTGTCTGTGCCGGATGCGGCGTGAATGCCTTATCCGGCCAATAAAATCCTAAAAATTCAATAA
GTTGATGTTCTTTCATGCTCTTATAAAGGTCGTGCCTCTGGCGGATGTACGTTTGTCATGAGTCTCACTCTGTTGCTAATTGCCGTTCG
CTCCTGAACATCCACTCGATCTTCGCCTTCTTCCGGTTTATTGTGTTTTAACCACCTGCCCGTAAACCTGGAGAACCATCGCGTGTTTC
AAAAAGTTGACGCCTACGCTGGCGACCCGATTCTTACGCTTATGGAGCGTTTTAAAGAAGACCCTCGCAGCGACGGATCCTCTAGATTT
AAGAAGGAGATATACATATG
>yheS
GAGGCAATTCCGACGTCTAAGAAACCATTATTATCATGACATTAACCTATAAAAATAGGCGTATCACGAGGCCCTTTCGTCTTCACCTC
GAGCAGTACCCGGTTTGCCACCGAGTCCTGAGATTCCGGATGGGCATACAGCAGCAAAACTTTCGCTGGCTGAGACATCATCCCCCTCC
CGGTATGGTTTTTGTGTATAGTCGCTGTTTTGGGCTACCATTGCGCCCGGTGCGGCAGCTCGCCCATACATTACATTATCATAATGATA
AGTTAACATAGTCTGAACATACGGCACCTTATGATTGTTTTCTCCTCGTTACAAATTCGTCGCGGCGTGCGCGTCCTGCTGGATAATGC
CACCGCCACCATCAACCCTGGGCAGAAAGTCGGCCTGGTGGGATCCTCTAGATTTAAGAAGGAGATATACATATG
```

```
>aroF
GAGGCAATTCCGACGTCTAAGAAACCATTATTATCATGACATTAACCTATAAAAATAGGCGTATCACGAGGCCCTTTCGTCTTCACCTC
GATGGATCCCAACAAGGGGGCGATAAACTTTTTCATCATTTCTTTCTCCTTTTTCAAAGCATAGCGGATTGTTTTCAAAGGGAGTGTAA
ATTTATCTATACAGAGGTAAGGGTTGAAAGCGCGACTAAATTGCCTGTGTAAATAAAAATGTACGAAATATGGATTGAAAACTTTACTT
TATGTGTTATCGTTACGTCATCCTCGCTGAGGATCAACTATCGCAAACGAGCATAAACAGGATCGCCATCATGCAAAAAGACGCGCTGA
ATAACGTACATATTACCGACGAACAGGTTTTAATGACTCCGGAACAACTGAAGGCCGCTTTTCCATTCTCGAGAGATCCTCTAGATTTA
AGAAGGAGATATACATATG
>aroH
GAGGCAATTCCGACGTCTAAGAAACCATTATTATCATGACATTAACCTATAAAAATAGGCGTATCACGAGGCCCTTTCGTCTTCACCTC
GAGTCAGATCCCGTGGATTAACAGTACCAATTATTCGGTAGAAGAGATTGCCACCAAGATCCTCGATATCATGGGCCTTAGTCGCCGAA
TGTACTAGAGAACTAGTGCATTAGCTTATTTTTTTGTTATCATGCTAACCACCCGGCGAGGTGTGACACACCTCGCACTTGAAATCAGC
AGCGATTGGTTTATCGTGATGCGCATCACTTCCCGGCAGTCCTGCCGTAGAAGCAACAAATTTCTGAGACTTGTAATGAACAGAACTGA
CGAACTCCGTACTGCGCGTATTGAGAGCCTGGTAACGCCCGCCGAACTCGCGCTACGGTATCCCGTAACGCCTGGGGATCCTCTAGATT
TAAGAAGGAGATATACATATG
>aroK
GAGGCAATTCCGACGTCTAAGAAACCATTATTATCATGACATTAACCTATAAAAATAGGCGTATCACGAGGCCCTTTCGTCTTCACCTC
GATGGATCCATCACGCCACGACTGGTTTCCAGTGAGTAAACAGCCGTAAAAGCGGTAATGTTTTTACGCTGAACGTGTTTCATCTATTT
GACGCGCGCAGGTATTTAGCATACAAGGAGTACCGATTTGAGAGTTGGTGCTCTTCGCTGCCTGCGTTCCATGATGATGATTTATCATT
CAGGCGGCATTTTGCTGTCTTTTTTACGCTAATCTTACCCGGTGATTTATCGCCAGAGCGGTGGTAGCAAGGCAGCGCGCTTGCAGCGA
CCAGATATGCAGAGGGATGGGTGATTTATTCAGTTGCCAAACCCGCTCGAGAGATCCTCTAGATTTAAGAAGGAGATATACATATG
>kefF
GAGGCAATTCCGACGTCTAAGAAACCATTATTATCATGACATTAACCTATAAAAATAGGCGTATCACGAGGCCCTTTCGTCTTCACGTC
GACTGATTTCCGTAGCGTTTGCCCCGGAGACTCGAGGGATGTCACTGGCGCAGACCAGCAATATGACGATCCGCGGGCAGAGAATGGGG
TAAATTGTTCAGATTTCTCTCTTTTCTGAATCAATATTATTGACTATAAGCCGCGTGAATATATGACTACACTTTGTGGGAAAACAAAG
GCGTAATCACGCGGGCTACCTATGATTCTTATAATTTATGCGCATCCGTATCCGCATCATTCCCATGCGAATAAACGGATGCTTGAACA
GGCGGATCCTCTAGATTTAAGAAGGAGATATACATATG
>kefG
GAGGCAATTCCGACGTCTAAGAAACCATTATTATCATGACATTAACCTATAAAAATAGGCGTATCACGAGGCCCTTTCGTCTTCACCTC
GATGGATCCCACCAGGCCGACTTTCTGCCCAGGGTTGATGGTGGCGGTGGCATTATCCAGCAGGACGCGCACGCCGCGACGAATTTGTA
ACGAGGAGAAAACAATCATAAGGTGCCGTATGTTCAGACTATGTTAACTTATCATTATGATAATGTAATGTATGGGCGAGCTGCCGCAC
CGGGCGCAATGGTAGCCCAAAACAGCGACTATACACAAAAACCATACCGGGAGGGGGATGATGTCTCAGCCAGCGAAAGTTTTGCTGCT
GTATGCCCATCCGGAATCTCAGGACTCGGTGGCAAACCGGGTACTGCTCGAGAGATCCTCTAGATTTAAGAAGGAGATATACATATG
>uvrD
GAGGCAATTCCGACGTCTAAGAAACCATTATTATCATGACATTAACCTATAAAAATAGGCGTATCACGAGGCCCTTTCGTCTTCACCTC
GAGGGCGATCTGATGCAAACCTGGGACAGCCGTTTACTGCCGCATCTGGAAATTTCCCGGTTGGCATCTCTGACCTCGCTGATATAATC
AGCAAATCTGTATATATACCCAGCTTTTTGGCGGAGGGCGTTGCGCTTCTCCGCCCAACCTATTTTTACGCGGCGGTGCCAATGGACGT
TTCTTACCTGCTCGACAGCCTTAATGACAAACAGCGCGAAGCGGTGGCCGCGCCACGCAGCAACCTTCTGGTGCTGGCGGATCCTCTAG
ATTTAAGAAGGAGATATACATATG
>uvrC
GAGGCAATTCCGACGTCTAAGAAACCATTATTATCATGACATTAACCTATAAAAATAGGCGTATCACGAGGCCCTTTCGTCTTCACCTC
GATGGATCCCATGGCGATGTTGAGCTGACTCACCTGGCAATTCGCCATGGTCTGTGTAATGCGGAGACATTATCAAGTCAGTGAGTGAT
CAGTTTGACGCAAAAGCGTTTTTAAAAACCGTAACCAGCCAGCCAGGCGTTTATCGCATGTACGATGCTGGTGGTACGGTTATCTATGT
CGGCAAAGCGAAAGACCTGAAAAAACGGCTTTCCAGCTATTTCCGTAGCAACCTCGCCTCGAGAGATCCTCTAGATTTAAGAAGGAGAT
ATACATATG
>b4283
GAGGCAATTCCGACGTCTAAGAAACCATTATTATCATGACATTAACCTATAAAAATAGGCGTATCACGAGGCCCTTTCGTCTTCACCTC
GAGTCTACGGTTCCGTCGCATTGCTTGTTATTGGTATATTCCTGACGTTCTTCATGCGTCCAGATAAGTCATTGTAAAAGTCGAATCAG
GGCTGAAGTGGCACACTGAATTTGGCCACCTGAACAGAGGTGATATGCTCACCTCAGAACAACACAGGTGCTCCAATGAAAAAAAGAAA
TTTCAGCGCAGAGTTTAAACGCGAATCCGCTCAACTGGTCGGGATCCTCTAGATTTAAGAAGGAGATATACATATG
>yidI
GAGGCAATTCCGACGTCTAAGAAACCATTATTATCATGACATTAACCTATAAAAATAGGCGTATCACGAGGCCCTTTCGTCTTCACCTC
GAGAGTACGGATCCACGCCAGAAAAGTACGCTCATTAGCCAGCGAGAAGCGGTAATCAGGCGCTTCTCCGAGGCGGGAAATTTTCATTG
TTGCTCCTTTGCAATTTCACTAACGCTCTACCTTTAGCGGTTTTTTTAAGAATTTCGTCCGCCCGTTTCCTGATATGCTTTTGCCATCC
GCTACAAGGAGGTCAGATGGGTATCATAGCTCAAAATAAAATCAGTTCATTGGGCATGCTGTTCGGTGCCATCGCTTTAATGATGGGGA
TTATCCACTTTTCATTCGGCCAGATCCTCTAGATTTAAGAAGGAGATATACATATG
>ycfQ
GAGGCAATTCCGACGTCTAAGAAACCATTATTATCATGACATTAACCTATAAAAATAGGCGTATCACGAGGCCCTTTCGTCTTCACCTC
GATGGATCCGAGCTTAAAATCGCCGCAGCGATGAGGGTTTTTACGTTTTTCATAATAGTGGCCTTATGCAGATGAATGACGTCGCAAGC
AATTACTTAACGATCGATAACTAAATGATAGATGTGATCTGGATCACATACAAGATATTTTTTATAACAATCATTAATTAATTTAAAAA
TCTTTAAATTTCAGTTAAATAGAAAGCAATTATTTTTATGGGATCTGGACTGGTGAACGGAGGTGATTATTTTTATAATAACCTTTCAT
TCACAGTCACCAGGTACAACGGCATCATGGCAACTCTCGAGAGATCCTCTAGATTTAAGAAGGAGATATACATATG
>rsuA
```

Data Set S4 - MG1655 library reference file.fasta

GAGGCAATTCCGACGTCTAAGAAACCATTATTATCATGACATTAACCTATAAAAATAGGCGTATCACGAGGCCCTTTCGTCTTCACCTC
GATGGATCCCATCCACGGCTTCTTGCTGATATGGGCGAAGTGTAAAAATCATGGTCTCGCGTAGGTAGATTATTCTGCTAATAGTATGC
CATGGATCTTTTTCCCTTGAGGGATGTTATTAGACCGCTATACTGGGCAGACAACAGTTCAGATTTTCGGGTCAATGCCCGACCTCCGT
TTTTACAGGCAAAATCACACACATGCGACTTGATAAATTTATCGCACAGCAACTCGGCGTTAGCCGTGCTATTGCCGGGCGTGAAATCC
GCGGCAATCGTGTCACCGTCGACTCGAGAGATCCTCTAGATTTAAGAAGGAGATATACATATG
>aegA
GAGGCAATTCCGACGTCTAAGAAACCATTATTATCATGACATTAACCTATAAAAATAGGCGTATCACGAGGCCCTTTCGTCTTCACCTC
GATGGATCCAGCAGAGCGATACCCGTGGAAAGAATCGACAGCAGCACAATGTAAAAAAAGGCCCGGGCCAGACTGGCCGAGACGGGTCG
TTTAACAATCACACGCGTCTTCTCCACAAAAAATCATTAATGTTCCAGTTCGCTGAAACGCAATTTAGTCAGTGATGGGATTATTTGAT
CTGCCACAGTTTATAGCGAAATAGTTACTTATTCTGGGCACATGAACATTGCCGATTCCCCTGTAAGTCGGGTAATAACAACACTCATA
TAAAGAATAAGGTTTTTACAACCAAAAAAGAAGGTCGTTATGAATCGTTTTATTATGGCCAACAGTCAGCAATGTCTGGGTTGTCATGC
TTGTGAAATCGCCTGTGTCATGGCTCTCGAGAGATCCTCTAGATTTAAGAAGGAGATATACATATG
>yniC
GAGGCAATTCCGACGTCTAAGAAACCATTATTATCATGACATTAACCTATAAAAATAGGCGTATCACGAGGCCCTTTCGTCTTCACGTC
GACACGCCAGTCCGACAAAAATCAGGATGAAAATCACCCAGAACATAACGTTCACGCCACCATTAAAATTGGGCGTCGGGGAGTTATAC
CAGAATAGATTCAAAAAAGGCGTATTCGCCTGCATCATATCGATCATGACATGAGTAAAGTCGAGCATGACCGCATTAATGCCTTCCTT
GTTTTCCTGCCGGGTATTCATAAATTTCAGCAAAGAAATCAATGTAGAGATCAATGCAGGTATAAAAATCACCCAACCCAAAATCCTTT
TCAAAACAGCAATGCGTCCAGCGGATCCTCTAGATTTAAGAAGGAGATATACATATG
>yniA
GAGGCAATTCCGACGTCTAAGAAACCATTATTATCATGACATTAACCTATAAAAATAGGCGTATCACGAGGCCCTTTCGTCTTCACCTC
GAGGAACTGGTTTCGGGGCTTACCCAAAGTGCAACCGGCAAAGATCCTGAAATCACCATCACGACCTGGGAGGAATGGAATAGCAACAG
AAAATAAATGGTTTTTGGGCAATAATCAGTCTGTGGTGTGCGTTAGCTCGTGTTTTTACACCGCATTCTTGCGCTAACCTTATGATCTG
GCAGACAACATGGGAGAGACATCATGTGGCAGGCAATCAGTCGTCTTTTGAGCGAGCAGTTAGGTGAAGGCGAAATCGAACTGCGTAAT
GAACTGCCTGGCGGAGAAGTCCATGCCGCATGGATCCTCTAGATTTAAGAAGGAGATATACATATG
>caiC
GAGGCAATTCCGACGTCTAAGAAACCATTATTATCATGACATTAACCTATAAAAATAGGCGTATCACGAGGCCCTTTCGTCTTCACCTC
GATGGATCCGGTCAGCAAAGGTCTGGCCAAAGTTGAGGACTAAACGCGTGGCGTCAGTCGGCTTGCCTGACTAGCGCCCGCCCAATAAC
AAACGAACAAAAATGGATAGAGGTGCAATGGATATCATTGGCGGACAACATCTACGTCAAATGTGGGACGATCTTGCGGACGTTCTCGA
GAGATCCTCTAGATTTAAGAAGGAGATATACATATG
>rumA
GAGGCAATTCCGACGTCTAAGAAACCATTATTATCATGACATTAACCTATAAAAATAGGCGTATCACGAGGCCCTTTCGTCTTCACCTC
GATGGATCCGCGCAGGCTGTAGTTGGTCATGGAGTTCCGTTATGGGACAATTAAGGTCACACTGTTAAATTGAGAAAGTCTCGCCAATG
GCGCAATTCTACTCTGCAAAACGACGCACGACGACGCGTCAGATCATAACCGTTTCAGTCAACGACCTCGACTCTTTTGGTCAGGGCGT
GCTCGAGAGATCCTCTAGATTTAAGAAGGAGATATACATATG
>ttdA
GAGGCAATTCCGACGTCTAAGAAACCATTATTATCATGACATTAACCTATAAAAATAGGCGTATCACGAGGCCCTTTCGTCTTCACCTC
GAGCGCTAAAACTCCCGCTGTGAACGATTTCTACCAGTACCTGCAAGTCTTTGGCTAAAGGCCAGCTATTCAGCATCGTTAATTATCCG
CAGTTGTGATAAGCGCAGTGTATTTCGCAAAACATTGCCATTCCCTGCTCTGGCGCAATATTCATTCTTATTTTCGCGTTGATTCACGA
ACTGTTAATTACATATTCACGGTGGCAAAAAATATAAAACCACATTTTTAGTGGTAGTTTGTGGCGGTGAATTTTTCCAGACAAATACA
AAAACTGGAGTTGCCATGATGAGCGAAAGTAATAAGCAACAGGCAGTGAATAAGTTGACAGAGATTGTCGCTAACTTTACCGCCATGAT
TTCTACCCGGGATCCTCTAGATTTAAGAAGGAGATATACATATG
>rpe
GAGGCAATTCCGACGTCTAAGAAACCATTATTATCATGACATTAACCTATAAAAATAGGCGTATCACGAGGCCCTTTCGTCTTCACCTC
GATGGATCCGTGGACGAACTGCTGGCTTTGTACAAACCAGGAGTCGTTTCACCCGCGAAAAAATAATTCTCAAGGAGAAGCGGATGAAA
CAGTATTTGATTGCCCCCTCAATTCTGTCGGCTGATTTTGCCCGCCTGGGTGAAGATACCGCAAAAGCCCTGCTCGAGAGATCCTCTAG
ATTTAAGAAGGAGATATACATATG
>rph
GAGGCAATTCCGACGTCTAAGAAACCATTATTATCATGACATTAACCTATAAAAATAGGCGTATCACGAGGCCCTTTCGTCTTCACCTC
GATGGATCCCACCCTTGATTTCACGCCGGGCGTAGGCGGTCATACTGCGGATCATAGACGTTCCTGTTTATAAAAGGAGAGGTGGAAGG
ATTATAGCCATCGATGCCTTGTCAGGATAGGAATAACCGCCGGAAGTCCGTATAATGCGCAGCCACATTTGTTTCAAGCCGGAGATTTC
AATATGCGTCCAGCAGGCCGTAGCAATAATCAGGTGCGTCCCGTTACCCTGACTCGTAACTATACAAAACATGCAGAAGGCTCGGTGCT
GGTCGAATTTGGCTCGAGAGATCCTCTAGATTTAAGAAGGAGATATACATATG
>nhoA
GAGGCAATTCCGACGTCTAAGAAACCATTATTATCATGACATTAACCTATAAAAATAGGCGTATCACGAGGCCCTTTCGTCTTCACCTC
GAGCGGCTGGCATGGTGTAATTCTATGGGGATGAATCGGCTCATGGTATCGCCTCGCGAAGAAGGTTTTTTTAAGCGTAGTCCGTAACG
CAATAAGTAACGAAATTAACGGGATTGGCGATTTGCGAACGTGATGCATGTCCGCGATCGCACAAAATAGCCGGTGCGGCGTCTATTCC
AGGTTATAAGTTGAGAAAACCACTAAGGGAAACGCCTGATGACGCCCATTCTGAATCACTATTTTGCCCGTATTAACTGGTCGGGAGCT
GCTGCGGTCAATATTGATACGCTTCGTGCATTGCAGGATCCTCTAGATTTAAGAAGGAGATATACATATG
>rhlE
GAGGCAATTCCGACGTCTAAGAAACCATTATTATCATGACATTAACCTATAAAAATAGGCGTATCACGAGGCCCTTTCGTCTTCACCTC
GAGATTCACCAAACTGCGCCAGTGCGGCAGCAATCAGCTGTTTTTTCGCCTGTTCACCCTTGATTGTCATGGCAGGATTATTCATCGCA
CAGTCCATTCTTAATCAAATGATTGATTAAGATTATGACTCCATAGGGGAGTTGTCCAGTATGGCTAAGAATTTTAGCAACGCCAGTCA
CAGGGATAATTTATGCGCTGCGTCACAAAAACTGCTACACTCCGCTCCCTCATGACATTGTGGTTTTTGTCATTTTCCTTTTCAGTATC

TCCCTGAAAACTACACCGGTAACGGTCGGGGCGGTTCGGAGTAGTTATGTCTTTCGATTCTTTGGGTTTAAGCCCTGATATCCTGCGCG
CCGTTGCCGAGCAGGGTTACCGTGAACCCACCCCTATTCAGCAGCAGGCGGATCCCGGGGATCCTCTAGATTTAAGAAGGAGATATACA
TATG
>rhlB
GAGGCAATTCCGACGTCTAAGAAACCATTATTATCATGACATTAACCTATAAAAATAGGCGTATCACGAGGCCCTTTCGTCTTCACCTC
GATGGATCCTAAGCCTGGCGTGTTGGTTTCGTTGTTGGTGTAACATTAACCAACTAAAGGTTGACTTTATTTCACCGGATACGCTTTCG
TAAAGCAATAGTAAGCTGATATTCTACCACACTATGAGCAAAACACATTTAACAGAACAGAAGTTTTCCGACTTCGCCCTGCATCCGAA
GGTTGTAGAAGCCCCTCGAGAGATCCTCTAGATTTAAGAAGGAGATATACATATG
>rpmI
GAGGCAATTCCGACGTCTAAGAAACCATTATTATCATGACATTAACCTATAAAAATAGGCGTATCACGAGGCCCTTTCGTCTTCACCTC
GATGGATCCTGGTCGAATCCTTCCCAACGAAGATCGAAGGCCGCCAGATGATCATGGTGCTCGCTCCTAAGAAGAAACAGTAAGGCCTT
CAAGTAGCACTGTCTGTGGAGCCTTCGGGCTTCACAGGCTTTGTTCGCCTTTGTTTCGTTTATTAACAATGCGAAGTGGAAGTTATTAA
AATGCCAAAAATTAAGACCGTACGCGGTGCTGCTAAGCGCTTCAAAAAAACCGGTAAAGGTGGTTTTAAGCACAAGCACGCTAACCTGC
GTCACACTCGAGAGATCCTCTAGATTTAAGAAGGAGATATACATATG
>agaR
GAGGCAATTCCGACGTCTAAGAAACCATTATTATCATGACATTAACCTATAAAAATAGGCGTATCACGAGGCCCTTTCGTCTTCACCTC
GATGGATCCATTGCAGCTTCCAGCACCAGCGGATGTGCGGAACAAACGGCATAAATTCCATTTGTTTTGCCCGCTTTGTGCTGTCTCAC
CATTTCTGTCAGATGTTTCACTTTCCTCTCCAGTTCAGATTACTGCGATATTAATCTTTCGTTTTGTTTCAGTTGATACTGCGTCATGG
TGAGATAAAAATAAAACGAAAGATAATGGTTTTCTGATCTGATTCACAAAAGAAACATAATGAAAGTCACTGAAACGAGAGTAGCTTGC
GTCAATGGGCAAGGTGGGCTTGCATTTGCTTAATAGAAAGGCGTTAATAGGCAAAACGAAATGAAACGAAAGTTTTACGAAAGGACTTA
CTATGAGTAATACCGACGCTTCAGGTGAGAAGCGAGTGACTGGCACCAGCGAGCGACGAGAACAGATCATTCAGCGTCTGCGACAGCAA
GGGAGTGTCTCGAGAGATCCTCTAGATTTAAGAAGGAGATATACATATG
>rpmB
GAGGCAATTCCGACGTCTAAGAAACCATTATTATCATGACATTAACCTATAAAAATAGGCGTATCACGAGGCCCTTTCGTCTTCACCTC
GATGGATCCTACGCGTGCTCGACCATATCGTGATTGGGCGTGGAGAGTATGTTTCTTTTGCCGAACGCGGCTGGATTTAACCCGCTATG
CGCGATCCTTCGGGATCTTTGTCTGTTCGGGACTTGAGCACATCGCTGAGTCAGCGTATACTACGCCACCTTTGAGAATCTCGGGTTTG
GCATTTGGGCCTGGCAATCGAGAGTTCACATAGAACTGCGATGACCGGGCTGTAAAGCCTGACGAGGCGCCAATACCCCATACGAAGCT
CGAGCTAATTTGATTTTTGGAGAATAGACATGTCCCGAGTCTGCCAAGTTACTGGCAAGCGTCCGGTGACCGGTAACAACCGTTCCCAC
GCACTGAACGCGACTAAGTCGAGAGATCCTCTAGATTTAAGAAGGAGATATACATATG
>rpmE
GAGGCAATTCCGACGTCTAAGAAACCATTATTATCATGACATTAACCTATAAAAATAGGCGTATCACGAGGCCCTTTCGTCTTCACCTC
GAGAACCGTCATGCCTTCTGGCAGCAGATAGTCAAAGGTACGAGGAAGCGGAACGGGCAAGGCAACGTGGGCAACGGGCATAGCATCAT
CCTGACTTGAAATTCGGTGGGTTAGTATACACATTGCCGTAGAAGAGTGCGGATCAGTTTGCATACGCTGGTTAATTTCTGTATGATTT
CGCGCCTTCGTACGAAATGATCGTATTGAAGCTATACTTTTAACATCGCGTGGTGTCTGGCGTTAGGGCTGGAAGAGCGACGCGGCCTT
AAACCGAGGTTTTCCCATGAAAAAAGATATTCACCCGAAATACGAAGAAATTACTGCTAGCTGCTCTTGCGGTAACGTAATGAAAATCC
GCTCCACCGTTGGATCCTCTAGATTTAAGAAGGAGATATACATATG
>lgt
GAGGCAATTCCGACGTCTAAGAAACCATTATTATCATGACATTAACCTATAAAAATAGGCGTATCACGAGGCCCTTTCGTCTTCACCTC
GATGGATCCAGTTCGCCATCAGGTTGCAGCCTTTATGGAGCGTCGCGGCATGGGCGGGCTGATTCGCGGAGGGTTATAGCGCGGATCAT
ATACATATCTTTTAACGGTATCCGGCAACCAGCCAGGTCCCCTTGTGCTATTATTCGCACCTTTGGAGCGCCTGAAACCTGCGGCGCGC
ATTTCAATCGCTGTTCTCTTTCAGCGAAATAACAAGAACTTGTGGTGACAGATGACCAGTAGCTATCTGCATTTTCCGGAGTTTGATCC
GGTCATTTTCTCAATAGGACCCGTGGCGCTTCACTGGTACGGCCTGATGTATCTGGTGCTCGAGAGATCCTCTAGATTTAAGAAGGAGA
TATACATATG
>yjgH
GAGGCAATTCCGACGTCTAAGAAACCATTATTATCATGACATTAACCTATAAAAATAGGCGTATCACGAGGCCCTTTCGTCTTCACCTC
GATGGATCCTGCATACCATTGATGGCGCGTTTGGCGCATAACCGACTACGCTCAATTAAGCCCAGCCATTATCCATGATGTCTGGGTTT
TGTTTACTCACGTCGTCCGCTAAAAGCGGCTCCCGGTAAAGAGAAATCTTCTGTTCATGTCGTAAATCTTGAGGATAACCAAATGGTAG
AAAGAACCGCTGTTTTCCCTGCTGGCCGACATTCACTATATGCTGAGCATCGTTATTCTGCGGCTATTCGTTCCCTCGAGAGATCCTCT
AGATTTAAGAAGGAGATATACATATG
>yjgJ
GAGGCAATTCCGACGTCTAAGAAACCATTATTATCATGACATTAACCTATAAAAATAGGCGTATCACGAGGCCCTTTCGTCTTCACCTC
GAGTTTTGGCAGTAAAGCTGGGTTATTTAGCCGTGTACTCAATGAATACGTCGGTACGGAAGCTATTCCGCTTGCCGATATTCTTCGTG
ATGATCGTCCGGTAGGCGAGTGCCTGGTTGAGGTATTAAAAGAAGCGGCGCGCAGATATAGCCAAAACGGCGGCTGCGCTGGCTGTATG
GTTCTTGAAGGTATTCATAGTCATGATCCACAAGCGCGTGATATTGCCGTTCAATATTATCACGCCGCAGAAACGACCATTTGGATCCT
CTAGATTTAAGAAGGAGATATACATATG
>yddA
GAGGCAATTCCGACGTCTAAGAAACCATTATTATCATGACATTAACCTATAAAAATAGGCGTATCACGAGGCCCTTTCGTCTTCACCTC
GATGGATCCGGATGTTGCAAAACAAATGGAGAATAATTAATAACGCTCTCTTTTAACTCCATTCTTAGCCTCCTTCTTATGACCTTAAC
AAGGCACCCGACATAAATTCCTGTCGGGTGTTTCGTCATCATCATCTTCCCTTGCGAAAATCCTGGCTTACGTGGTATGCATCAGGGCA
TTAGCAATATCCTTGATTATCAATAGCATAACCATAACCACTGAAAATGAAATATTCCTTTATATTTCATATTGTTAAATGATACTTAA
ATAAATTTCATCAGTAAATCATTGCCATAGATAATCATTGGATGGCACAATCAATTTTCGCTAATGATAACCATTCCCATTACGCTTCG
TATGTTGATAGCAAAGTATCTCTGTCTGTTGAAGCCCTTTTGGCTGCTCGAGAGATCCTCTAGATTTAAGAAGGAGATATACATATG
>yjgA

GAGGCAATTCCGACGTCTAAGAAACCATTATTATCATGACATTAACCTATAAAAATAGGCGTATCACGAGGCCCTTTCGTCTTCACCTC
GATAGATCTACTGGCTCGACGACGTTCCCGGTGATGACATCGAAGACGAAGACGATGAAATTATCTGGGTCAGTAAAAGTGAAATTAAA
CGTGATGCCGAGGAGCTAAAACGCCTTGGCGCGGAAATTGTTGATCTGGGGAAAAACGCGCTGGATAAGATCCCGTTAGATGCGGATCT
GCGCGCGGCTATTGAGCTCGCCCAGCGTATTAAGATGGAAGGTCGCCGCCGCCAGTTGCAGCTCATCGGTAAGATGCTGCGCGTCGAGA
GATCCTCTAGATTTAAGAAGGAGATATACATATG
>yjgR
GAGGCAATTCCGACGTCTAAGAAACCATTATTATCATGACATTAACCTATAAAAATAGGCGTATCACGAGGCCCTTTCGTCTTCACCTC
GATGGATCCTCACGGCAACACGCTTTAGGTTATTTTCTCTTCATTACACGTATTCTCTTTGCCGGACGTTGCCTATACTCCAGGCAGGA
AACTGGAGGATATCTCATGAGTGAACCCCTGTTAATTGCCCGCACGCCGGACACAGAACTGTTTTTACTGCCGGGAATGGCTAACCGTC
ATGGGCTGATTACCGGCTCGAGAGATCCTCTAGATTTAAGAAGGAGATATACATATG
>fabR
GAGGCAATTCCGACGTCTAAGAAACCATTATTATCATGACATTAACCTATAAAAATAGGCGTATCACGAGGCCCTTTCGTCTTCACCTC
GAGAGCTTTCGACGGGATGGTGCCCCAGTGGGTGCAACCGCCGCCAACATTTTGATAACGCTCGATAACTGCGACGCGCGCACCTTGCT
TAACCAGGCCCATTGCAGCGCCTTCGCCGCCGGGGCCGGAACCTATTACTATGGCATCGTAATCGTAGGAATGTGGCATGGTAGGGCTT
ACCTGTTCTTATACATAAAAGCAACAGAATGGTAACATTTTATCGCGGGTAAGCCAATTGATCCCCGTCATTTATCTGGCTATATCCTG
AGCGGCCTTTGCTTTGTCTGTTTCTTACTTTTGCCCTGACGTTTTATTGGATTTTTATCGACGATACTCTCCGTTTAAGCGGCAGGTTT
CCGCTGTACGTAAAAGAACCGGCCAAAGAATTGCAGTAAATATGTTTTATTGCGTTACCGTTCATTCACAATACTGGAGCAATCCAGTA
TGTTCATTCTCTGGTATAGTGCCAGCAGTACTTTTGGCAAGGATTCAGACATCGTGATGGGCGTAAGAGCGCAACAAGGATCCTCTAGA
TTTAAGAAGGAGATATACATATG
>fabZ
GAGGCAATTCCGACGTCTAAGAAACCATTATTATCATGACATTAACCTATAAAAATAGGCGTATCACGAGGCCCTTTCGTCTTCACCTC
GAGAACCGCTGCACTGGTGATGAACATTGATGACATGAGCAAGCGTCTGAAATCGCTTGAGCGCAAGGTTAATCAACAAGACTAACGTT
CCATCTTTTGTTCGCCAAACTTTACGGCCTGTCTCATTCTTACGATTGCGGCAGGCCGTGTTATTATTGTCGTTTCTTATATTTTGACA
GGAAGAGTATCTTGACTACTAACACTCATACTCTGCAGATTGAAGAGATTTTAGAACTTCTGCCGCACCGTTTCCCGTTCTTACTGGTG
GATCGCGTGCTGGATTTTGGGATCCTCTAGATTTAAGAAGGAGATATACATATG
>rfbB
GAGGCAATTCCGACGTCTAAGAAACCATTATTATCATGACATTAACCTATAAAAATAGGCGTATCACGAGGCCCTTTCGTCTTCACCTC
GATGGATCCTCCTTGTTTCCAGAGCGGATTGGTAAGACAATTAGCGTTTGAATTTTTCGGGTTTAGCGCGAGTGGGTAACGCTCGTCAC
ATCATAGGCATGCATGCAGTGCTCTGGTAGCTGTAAAGCCAGGGGCGGTAGCGTGCATTAATACCTCTATTAATCAAACTGAGAGCCGC
TTATTTCACAGCATGCTCTGAAGTAATATGGAATAAATTAAGTGAAAATACTTGTTACTGGTGGCGCAGGATTTATTGGTTCAGCTGTA
GTTCGTCACATTATAAATAATACGCAGGATAGTGTTGTTAATGTCGATAAATTAACGTACGCCGGAAACCGGGAATCACTTGCTGATGT
TTCTGATTCTGAACGCTATGTTTTTGAACATGCGGATATTTGCGATGCACCTGCAATGGCACGGATTTTTGCTCAGCATCAGCCGGATG
CAGTGATGCACCTGGCTGCTGAAAGCCATGTTGACCGTTCAATTACAGGCCCTGCGGCATTTATTGACTCGAGAGATCCTCTAGATTTA
AGAAGGAGATATACATATG
>rfbA
GAGGCAATTCCGACGTCTAAGAAACCATTATTATCATGACATTAACCTATAAAAATAGGCGTATCACGAGGCCCTTTCGTCTTCACCTC
GATGGATCCAGGTTGGCGTGAAACGAATGCTCAATGAATTATTTACGACTACAGCAATTTAATAGTTTTTGCATCTTGTTCGTGATGGT
GGAGCAAGATGAATTAAAAGGAATGATGAAATGAAAATGCGTAAAGGTATTATTTTAGCGGGTGGTTCTGGTACACGTCTTTATCCTGT
GACTATGGCTGTCACTCGAGAGATCCTCTAGATTTAAGAAGGAGATATACATATG
>cpsG
GAGGCAATTCCGACGTCTAAGAAACCATTATTATCATGACATTAACCTATAAAAATAGGCGTATCACGAGGCCCTTTCGTCTTCACCTC
GATGGATCCGGGAAAATTCCGCTCGATTTAATTGAAGTGCGCTCCGGCTCTTATCTCGAAGAGGATGATGTGGTGCGTTTCGCGGATCG
CTACGGACGGGTGTAAACGTCGCATCAGGCAATGAATGCGAAACCGCGGTGTAAATAACGACAAAAATAAAATTGGCCGCTTCGGTCAG
GGCCAACTATTGCCTGAAAAAGGGTAACGATATGAAAAAATTAACCTGCTTTAAAGCCTATGATATTCGCGGGAAATTAGGCGAAGAAC
TGAATGAAGATATCGCCTGGCGCTCGAGAGATCCTCTAGATTTAAGAAGGAGATATACATATG
>ygaV
GAGGCAATTCCGACGTCTAAGAAACCATTATTATCATGACATTAACCTATAAAAATAGGCGTATCACGAGGCCCTTTCGTCTTCACCTC
GAGCGAACCCTTTACCGAGCAGCACGCCGAGCGGCGGCAGAATGATGGTGATGACGATTCTCCAGAAACCCATATGTACTCCCTATAAG
AAAATTACTCATTGTTTAAAAAGAGATTTTATCTCTTAAGCGTAGTAATTTTAGCGGAGCCTGCCAGTTTCAGCTTACGCTTTTACTCT
CATTTACGCTGCTGAGGCTGGTATATGTTGCATTTTTAGTTTAGAGTTTACTTAATTTAGAAAATACTTAAATAAATATGACTGAACTC
GCGCAATTACAGGCCAGTGCCGAACAGGCAGCGGCCTTATTGAAAGCAATGAGCCCCCGGATCCTCTAGATTTAAGAAGGAGATATACA
TATG
>fkpA
GAGGCAATTCCGACGTCTAAGAAACCATTATTATCATGACATTAACCTATAAAAATAGGCGTATCACGAGGCCCTTTCGTCTTCACCTC
GATGGATCCTGAAAAGCCAGTCGGCTCTCCAGCTCTGCCAGGCGTGCTTCCAATGATAAATCCTGCATGATTCACCTCTTTTGTCGAAT
GGTCGCCGCGGATTCTACTTAACTTTGCTGCCCGAGACAGCACTCATTTCGCGGTCATCGAAACTAATTTAAACAAAAAGAGTCTGAAA
ATAGATGATAATAGGGCGTGTCTGTATGTAGATTTGTTCGACAACGCTTTATAGTACCCTTCTGATAATAGTTAACCCTGGGGTGAGAT
GCCCCGATCCTGGAGATATGGATGAAATCACTGTTTAAAGTAACGCTGCTGGCGACCACAATGGCCGTTGCCCTGCATGCACCAATCAC
TTTTGCTGCTGAAGCTGCACTCGAGAGATCCTCTAGATTTAAGAAGGAGATATACATATG
>dam
GAGGCAATTCCGACGTCTAAGAAACCATTATTATCATGACATTAACCTATAAAAATAGGCGTATCACGAGGCCCTTTCGTCTTCACCTC
GATGGATCCCGTCAGGTACAGGCCGATCTGAAGTAATCAAGGTTATCTCCCGCAATGGTTTATCGTTGCGGGAGTTGCCTGAAGCGCTG
GATGCTGTCGGAGCTTTCTCCACAGCCGGAGAAGGTGTAATTAGTTAGTCAGCATGAAGAAAAATCGCGCTTTTTTGAAGTGGGCAGGG

GGCAAGTATCCCCTGCTTGATGATATTAAACGGCATTTGCCCAAGGGCGAATGTCTGGTCTCGAGAGATCCTCTAGATTTAAGAAGGAG
ATATACATATG
>ackA
GAGGCAATTCCGACGTCTAAGAAACCATTATTATCATGACATTAACCTATAAAAATAGGCGTATCACGAGGCCCTTTCGTCTTCACCTC
GAGCGGCCACGTCTTTGAGTAATGCTGTCCCCGGCGAAACAAGCTAAAAAAATTAACAGAACGATTATCCGGCGTTGACATGCTTCACC
TCAACTTCACATATAAAGATTCAAAAATTTGTGCAAATTCACAACTCAGCGGGACAACGTTCAAAACATTTTGTCTTCCATACCCACTA
TCAGGTATCCTTTAGCAGCCTGAAGGCCTAAGTAGTACATATTCATTGAGTCGTCAAATTCATATACATTATGCCATTGGCTGAAAATT
ACGCAAAATGGCATAGACTCAAGATATTTCTTCCATCATGCAAAAAAAATTTGCAGTGCATGATGTTAATCATAAATGTCGGTGTCATC
ATGCGCTACGCTCTATGGCTCCCTGACGTTTTTTTAGCCACGTATCAATTATAGGTACTTCCATGTCGAGTAAGTTAGTACTGGTTCTG
AACTGCGGTAGTTCTTCACTGAAATTTGCCATCATCGATGCAGGATCCTCTAGATTTAAGAAGGAGATATACATATG
>setA
GAGGCAATTCCGACGTCTAAGAAACCATTATTATCATGACATTAACCTATAAAAATAGGCGTATCACGAGGCCCTTTCGTCTTCACCTC
GAGGCCAGTTCGTTGAGCGTTGTGTCCTGCGATTTACCCTCGCAGCATTGCCACAGGCGGATGAACTGTTGTTGCAGACGAGCAGATGG
CATAAAAGGGGAACTCCTGTGCAAAAGACAGCAATTTTATTTTCCCTATATTAAGTCAATAATTCCTAACGATGAAGCAAGGGGGTGCC
CCATGCGTCAGTTTTATCAGCACTATTTTACCGCGACAGCGAAGTTGTGCTGGTTGCGTTGGTTAAGCGTCCCACAACGATTAACCATG
CTTGAAGGACTGATGCAGTGGGATGACCGCAATTCTGAAAGTTGACTTGCCTGCATCATGTGTGACTGAGTATTGGTGTAAAATCACCC
GCCAGCAGATTATACCTGCTGGTTTTTTTTATTCTCGCCGCGCTAAAAAGGGAACGTATGATCTGGATAATGACGATGGCTCGCCGTAT
GAACGGTGTTTACGCGGCATTTATGCTGGTCGCTTTTATGATGGGGGGGATCCTCTAGATTTAAGAAGGAGATATACATATG
>setB
GAGGCAATTCCGACGTCTAAGAAACCATTATTATCATGACATTAACCTATAAAAATAGGCGTATCACGAGGCCCTTTCGTCTTCACCTC
GAGGCGAATCGCCTCTTCTTTGTCTCCGGCCTTTTCGCCCGGATGGATGTCCTGTACGGATAACTGGAACATAGTTCTCCTCTCTTGCT
GAATTGAAACGATTCAGCCTCTATGAGAAAAAAAGCGCCAACCTGGCTTAGGGTTAAAGACAAGATCGCGCTGAAACGTTTCAAGAAAG
CATAATACTTCTGTTTCAGCACGCAAGGAAAGCCGTAAAGTTATGAGCAAAAGTTTGATGTGCTGCACAATTTTTTGCCATTTTTCCTA
ATTGCTGATGGGAAAATCCGTTATCAGCAATTTCATTTCAGCAGCGTCAGCAGCCAATCAGCAACAGTGAAATGGCATTTTGACTATTC
GTGGCGCTTTTGATTTCGCTGTCTGTTTATTTTCTGACAAGCAGCGTAAACTCCGCGTCTTCCTCTTCCAGTGATCGACCAGCATGCAT
AACTCCCCCGCAGTCTCCAGCGCGAAATCGTTTGACGGATCCTCTAGATTTAAGAAGGAGATATACATATG
>mglB
GAGGCAATTCCGACGTCTAAGAAACCATTATTATCATGACATTAACCTATAAAAATAGGCGTATCACGAGGCCCTTTCGTCTTCACCTC
GATGGATCCATGCCGACGTTAGTGCGTCGCCATTCTGTCGCAACACGCCAGAATGCGGCGGCGATCACTAACTCAACAAATCAGGCGAT
GTAACCGCTTTCAATCTGTGAGTGATTTCACAGTATCTTAACAATGTGATAGCTATGATTGCACCGTTTTAACGTTGTAACCCGTATGT
AACAGTGAATAATCACTTTTGCCGAGGTAACAGCGTCATAACAACAATTAAAGCCGTTTTCTGGAGCGTTACCGGGCATGGAAGAACGA
ATTTTAAAAAGTGAGCTTCGGCGTTCAGTAACACTTCATTAACTCTACTGCCCCGCCGAGCATTTATCTCAAGCACTACCCTGCATAAG
AAAAACCGGAGATACCATGAATAAGAAGGTGTTAACCCTGTCTGCTGTGATGGCCAGCATGTTATTCGGTGCCGCTGCACACGCTGCTG
ATACTCGCACTCGAGAGATCCTCTAGATTTAAGAAGGAGATATACATATG
>fhlA
GAGGCAATTCCGACGTCTAAGAAACCATTATTATCATGACATTAACCTATAAAAATAGGCGTATCACGAGGCCCTTTCGTCTTCACCTC
GAGAAGGTGTTCGTCTTGCCGGTCTGTATGGCGTGAAACGAACCCTCGATTTACCACACGCCGAACCGCTTCCGCGTATATGCTAATAA
AATTCTAAATCTCCTATAGTTAGTCAATGACCTTTTGCACCGCTTTGCGGTGCTTTCCTGGAAGAACAAAATGTCATATACACCGATGA
GTGATCTCGGACAACAAGGGTTGTTCGACATCACTCGGACACTATTGCAGCAGCCCGAGGATCCTCTAGATTTAAGAAGGAGATATACA
TATG
>iscS
GAGGCAATTCCGACGTCTAAGAAACCATTATTATCATGACATTAACCTATAAAAATAGGCGTATCACGAGGCCCTTTCGTCTTCACCTC
GATAGATCTATACTCACGACGCGCCACGCACCCGCACACAAGACGCGATCGACGTTAAGTTACGCGCTTAATAAAAAGAATTCAGAATC
AGGCCGGAGTGCTAAATACTCCGTAAACACGGTCGTACATCCAGCCGGTAGCCTGATTCCTTGCATTGAGTGATGTACGGAGTTTATAG
AGCAATGAAATTACCGATTTATCTCGACTACTCCGCAACCACGCCGGTGGACCCGCGTGTTGCCGAGAAAATGATGCACTCGAGAGATC
CTCTAGATTTAAGAAGGAGATATACATATG
>iscR
GAGGCAATTCCGACGTCTAAGAAACCATTATTATCATGACATTAACCTATAAAAATAGGCGTATCACGAGGCCCTTTCGTCTTCACCTC
GATGGATCCCGGAAAGCCAGGAGTTGAATATCCTGCGCGGGATTCTGGCTTCTATTGAGCAGCAGAATAAAGGTAACAAGGCCGAATAA
CAGCCGTTGCCTGATGCGACGCGTAATGCGTCTTATCAGGCCTACAGTGAACAGAACCGTAGGTCGGATAAGGCGTTCACGCCGCATCC
GACAGCCGTTGCCTGATGCGACGCGTAATGCGTCTTATCAGGCCTACAGTGAACAGAACCGTAGGTCGGATAAGGCGTTCACGCCGCAT
CCGACAGCCGTTGCCTGATGCGACGCGTAATGCGTCTTATCAGGCCTACAGTGTACAGAACCCCAGGGCGGATATGGCGTTCACGCCGC
ATCCGACAACAGGTACAAACGCCACGATAAAAAAATGGCACTGAAGGTTAAATACCCGACTAAATCAGTCAAGTAAATAGTTGACCAAT
TTACTCGGGAATGTCAGACTTGACCCTGCTATGCAATACCCCCACTTTTACAATAAAAAACCCCGGGCAGGGGCGAGTTTGAGGTGAAG
TAAGACATGAGACTGACATCTAAAGGGCGCTATGCCGTGACCGCAATGCTTGACGTTGCGCTCAACTCTGAAGCGGGCCTCGAGAGATC
CTCTAGATTTAAGAAGGAGATATACATATG
>dusA
GAGGCAATTCCGACGTCTAAGAAACCATTATTATCATGACATTAACCTATAAAAATAGGCGTATCACGAGGCCCTTTCGTCTTCACCTC
GAGGCGAAGTCGTTTGACCCAACCTCAAAGGAAGAGATTAAAATACTTACAGACACTTCTAAATTAATTTATGATTTGTTCGGGGTTAA
ATCGGAGAAATAATATGTGAAGTTCTTCGATAGTATGGAAGGCATTATATAAAAGGACCCAATATTTATTGGGTTCTTTTTTCTCTATC
AATACTATTAGCAGGGAGATATATCACCAGAGTTTAATGTGTGATTTTTTATTTATCGTCGAACCTGGATTGTTTATCATTGGCCTTAA
CAAAGTTAACGGCTAATAAGATTATTTCCATCACTTCGTGAGAGCTTCATGCCTTGAGAGGATCTCAATTTTCTTTTGCAATGAGACAG
GCGCTTCCTGTTGTTATGGTATAGTACCCCGCTATTGAGCCTCCTGAATAGTGATGCTGAATAACATAACCCCATGATATATCGATAAA

ATAATCTCTACATTTGAAAATGCACGGTAATTCTGAAATGCAAAAAATCAACCAAACCAGCGCAATGCCTGAAAAAACTGACGTTCACT
GGAGTGGTCGGTTTAGCGTTGCACCAATGCGGATCCTCTAGATTTAAGAAGGAGATATACATATG
>dusB
GAGGCAATTCCGACGTCTAAGAAACCATTATTATCATGACATTAACCTATAAAAATAGGCGTATCACGAGGCCCTTTCGTCTTCACCTC
GAGCCGATAGCTTCGCACTGGACCCGGTCGTGGAAAAAGAAGAGTGGTGCCGTATTACCGGTCGTAAGAATTAACCTTCGCATCGCCGT
AGGGTGACGCGGGGCGAAGTGCGAGCAAGCTCACAAAAGGCACGTAAATTTGCCGATTATTTACGCAAATTTGCGTGCCAAAATTTTCA
TTCATAAAGAAAAATTGAGAACTTACTCAAATTTCTTTGAGTGTAAATTTTAGTCACTATTTTCTAATATGATGATTTTTATGAGTAAT
TATCGCACCACGCTCATTTTAAATGCAATTCTTTGATCCATCTCAGAGGATTGGTCAAAGTTTGGCCTTTCATCTCGTGCAAAAAATGC
GTAATATACGCCGCCTTGCAGTCACAGTATGGTCATTTCTTAACTCATGCGCATCGGACAATATCAGCTCAGAAATCGCCTGATCGCAG
CGCCCATGGCTGGCATTACAGACAGACCTTTTCGGACGTTGTGCTACGAGGATCCTCTAGATTTAAGAAGGAGATATACATATG
>dusC
GAGGCAATTCCGACGTCTAAGAAACCATTATTATCATGACATTAACCTATAAAAATAGGCGTATCACGAGGCCCTTTCGTCTTCACCTC
GATGGATCCAGGACGAAAGCGCGTAAATATTGCCAGATAATGTTCAGTGTCTTACTCATAATCAACACCCCAGATGAGATGGATTCATC
ATACAATTAATTCGTAAAACGTGCTACCGATCACATTAAGAAATCAGCGCTTATTTTTTCGCTCCCCGGCATTACGCAATGCGCAGCGA
ACTCAGGCATACATCAAGCATAGAGAGGGATTGCAGTTTGCAGCAATCCCTCTTTGCAGACCTTTAATCCGCTGACCATAGGCGCTAAC
ATAGCGCCTCATTTTTTTGCGGGTGATGATATGCGTGTGTTACTGGCACCGATGGAGGGAGTGCTTGACTCTCTGGTGCGTGAATTGCT
GACCGAAGTTAACGACTACGATCTGTGCATCACCGACTCGAGAGATCCTCTAGATTTAAGAAGGAGATATACATATG
>nfnB
GAGGCAATTCCGACGTCTAAGAAACCATTATTATCATGACATTAACCTATAAAAATAGGCGTATCACGAGGCCCTTTCGTCTTCACCTC
GATGGATCCTCCGAAGCGTTACTTCGCGATCTGATCAACGATTCGTGGAATCTGGTGGTTGATGGTCTGGCTAAACGCGATCAAAAAAG
AGTGCGTCCAGGCTAAAGCGGAAATCTATAGCGCATTTTTCTCGCTTACCATTTCTCGTTGAACCTTGTAATCTGCTGGCACGCAAAAT
TACTTTCACATGGAGTCTTTATGGATATCATTTCTGTCGCCTTAAAGCGTCATTCCACTAAGGCATTTGATGCCAGCAAAAAACTTACC
CCGGAACAGGCCTCGAGAGATCCTCTAGATTTAAGAAGGAGATATACATATG
>b2639
GAGGCAATTCCGACGTCTAAGAAACCATTATTATCATGACATTAACCTATAAAAATAGGCGTATCACGAGGCCCTTTCGTCTTCACCTC
GATGGATCCGGCCACGCTGATTATGCTGTATCTCTTCTTTCGCAGAGTTATTCCGGCAACGTATTAAGTTTCGCTGCTGAAGACGCCTG
CCAGTGCGATCAAGGATCTGGCAACCTTCAGGGCGGGCTGGATTGTCCTGTTATTGCTGCTTGTCGGTTTCTTCTTTCTGGAGCCGCAG
GGAATCCTGGTCAGTGCGATAGCGGCCGCTGGCGCAGCAGTGCTGTTTGTGGTGGCGAAAAGAGGTCATTCGATCAACACCGGAAAAGT
CCTGCGCGGTGCGCCATGGCAGACTCGAGAGATCCTCTAGATTTAAGAAGGAGATATACATATG
>ycfD
GAGGCAATTCCGACGTCTAAGAAACCATTATTATCATGACATTAACCTATAAAAATAGGCGTATCACGAGGCCCTTTCGTCTTCACCTC
GATGGATCCCTGCGCCGAAAGATGAATAAATATGCCCATACTTCACGCATTACGTTAAGCATCCGTTATAATCGGTTGCAGATACCAGC
CTGTGGATGCTTAACATGGAATACCAACTCACTCTTAACTGGCCCGATTTTCTTGAACGTCACTGGCAGAAACGCCCGGTGGTGTTAAA
ACGCGGCTCGAGAGATCCTCTAGATTTAAGAAGGAGATATACATATG
>mfd
GAGGCAATTCCGACGTCTAAGAAACCATTATTATCATGACATTAACCTATAAAAATAGGCGTATCACGAGGCCCTTTCGTCTTCACCTC
GATGGATCCCGGGCTTGCTGGTAAGAAAACTTTCACAGCGTGCATTCAGCGATCTGTTGTTCTCCCCGCCTTCACTGCCCGCTGCCGTC
AGTTACTCCCGCTAAGCCATCGCCCCAGCCGGGGCGATTCAATTTGCTAAACCATGTCGATTACGGATAATGAACAGCATTGCTTATCA
GAGATATGCCCCCATATGTTGAGGCATATCCTAACGAGAATCTGACAACCGTTATGCCTGAACAATATCGTTATACGCTGCCCGTCAAA
GCGGGTGAGCAGCGTCTGCTGGGCGAGTTAACCCTCGAGAGATCCTCTAGATTTAAGAAGGAGATATACATATG
>modF
GAGGCAATTCCGACGTCTAAGAAACCATTATTATCATGACATTAACCTATAAAAATAGGCGTATCACGAGGCCCTTTCGTCTTCACCTC
GATGGATCCTTAATGCCGACAGCGTGATTATCGCCACGCTGTGCTAAGCGTGTTGACAATTTGTTATGAAACACGTATCCCTGTCAGTA
ATCGCTGCACAAAGTGGGATATAAAATGTCATCGTTGCAAATTTTGCAAGGCACGTTTCGTCTTAGCGACACAAAAACGCTGCAATTGC
CTCAGCTAACGTTAAACGCGGGTGATAGTTGGCTCGAGAGATCCTCTAGATTTAAGAAGGAGATATACATATG
>gutM
GAGGCAATTCCGACGTCTAAGAAACCATTATTATCATGACATTAACCTATAAAAATAGGCGTATCACGAGGCCCTTTCGTCTTCACCTC
GAGTGCTGCTGTTCTACGCCAGTCCTAAGGCGTCGTACTGCACCGGACAGTCGATCAATGTCACCGGCGGTCAGGTGATGTTCTGATCA
ACAGCGGAGATCCATTAAGGATCTCCGTGAGACTATAGAATGCCTGATGCGCTACGCTCATCAGGCATACAGGACTTCCGCCACTACAT
TAAGGAAAAGTTATGGTATCCGCACTCATCACCGTCGCCGTTATCGCCTGGTGTGCGCAACTGGCCTTAGGCGGCTGGCAAATTTCTCG
TTTTAACCGTGCCTTCGACACACGGATCCTCTAGATTTAAGAAGGAGATATACATATG
>gmk
GAGGCAATTCCGACGTCTAAGAAACCATTATTATCATGACATTAACCTATAAAAATAGGCGTATCACGAGGCCCTTTCGTCTTCACCTC
GAGAGACCGCCCACACAGATGATTGCCAGCACAAGATACCTATTAATATCGCCATCCATACTTTCATCATCACCTCCGCTGTAGATAGT
CAGGCAGATATACAACGTGATGAAAGCAAAGCCGAGTGGCAAAAACGGAGTCTGCGAGGACGCTTCCTGAAAATCGTCTTTATTGCAGT
GAATGACAGGCAAATGCGGAAGCAGCTACGCAAAACGCAACAACTTTGCGCAAAAAGTGTGAGCAAGGGCTACGTCACATGGCCGCGCC
GTGTATAATAAGCTCGTATGTAGGCTTTATTTCGCTAATCACATACGAAAGATACTCATGGCTCAAGGCACGCTTTATATTGTTTCTGC
CCCCAGTGGCGCGGGTAAATCCAGCCTGAGGATCCTCTAGATTTAAGAAGGAGATATACATATG
>skp
GAGGCAATTCCGACGTCTAAGAAACCATTATTATCATGACATTAACCTATAAAAATAGGCGTATCACGAGGCCCTTTCGTCTTCACCTC
GAGTGTTCTCCTACGCCCAGCCGTTCAAAAAGTACGATGGAGACAAGGCAGAACAGTTCCAGTTTAACATCGGTAAAACCTGGTAAGTG
TTCTCCACAAAGGAATGTAGTGGTAGTGTAGCGATGACTTTAGGCGATCAATATAAGATCGCCGGGCCACGCAAAGAACTGCACCCTCC

GGTGCAAATGGGATGGTAAGGAGTTTATTGTGAAAAAGTGGTTATTAGCTGCAGGTCTCGGTTTAGCACTGGCAACTTCTGCTCAGGCG
GCTGACAAAATTGCAATCGTCAACATGGGCAGGATCCTCTAGATTTAAGAAGGAGATATACATATG
>sufI
GAGGCAATTCCGACGTCTAAGAAACCATTATTATCATGACATTAACCTATAAAAATAGGCGTATCACGAGGCCCTTTCGTCTTCACCTC
GATGGATCCAGGTTCGTGAGCTGGCTGCCCATTGTCGTTCGATAATGGAACAAAAAATCGCCGAGCTCGATAAAGAAGTCGCAGAACGC
GAAGCCGCCGGAAAAGTTTAAGTCGGCAATCTGTATTTTTGCGGGGAACACTTTCCTGCACGGTATTACTTTAGCCAGTTTTACATGGA
GCAAATATGTCACTCAGTCGGCGTCAGTTCATTCAGGCATCGGGGATTGCACTTTGTGCAGGCGCTGTTCCCCTGAAGGGTCGAGAGAT
CCTCTAGATTTAAGAAGGAGATATACATATG
>add
GAGGCAATTCCGACGTCTAAGAAACCATTATTATCATGACATTAACCTATAAAAATAGGCGTATCACGAGGCCCTTTCGTCTTCACCTC
GAGAGGTCGTGGTTTTGTCCGTCTCAATGCCGGCTGCCCACGTTCGAAACTGGAAAAAGGTGTGGCTGGATTAATTAACGCCATCCGCG
CTGTTCGTTAACCCCAATTGCGCAACGTAAAAAATCGTTGCGCAATCGTGGATTTTTACCCTGCTTTGTTTTTATAATGGTGCGCACTT
TTATATCCAGAAAAAGAGTGCGACCATGATTGATACCACCCTGCCATTAACTGATATCCATCGCCACCTTGATGGCAACATTCGTCCCC
AGACCAGGATCCTCTAGATTTAAGAAGGAGATATACATATG
>ada
GAGGCAATTCCGACGTCTAAGAAACCATTATTATCATGACATTAACCTATAAAAATAGGCGTATCACGAGGCCCTTTCGTCTTCACCTC
GATGGATCCGAAGGGCTGGCGGTTTATATGATCACCAAAGAAGGCGATAGCTTTAAAACCTGGATGTCACCACAGTTTAAAAGCTTCCT
TGTCAGCGAAAAAAATTAAAGCGCAAGATTGTTGGTTTTTGCGTGATGGTGACCGGGCAGCCTAAAGGCTATCCTTAACCAGGGAGCTG
ATTATGAAAAAAGCCACATGCTTAACTGACGATCAACGCTGGCAATCTGTCTTAGCCCGCGACCCGAATGCCGACGGCGAATTCGTTTT
CGCCGTGCCTCGAGAGATCCTCTAGATTTAAGAAGGAGATATACATATG
>gmr
GAGGCAATTCCGACGTCTAAGAAACCATTATTATCATGACATTAACCTATAAAAATAGGCGTATCACGAGGCCCTTTCGTCTTCACCTC
GATGGATCCCGAAGTCCGCATGGAAACCCGCAGCATTATTGCGCGCCCGGTCGCGTAATCTCCTTTCACGGCCCATTCCTCATGGATGG
GCCGTTTATTTCCCCGCTCTACCTTCGTTATATCCAGGCAGATTTTTTAACTGACTTTCGTTTGAAAACTGGCGTTTTTCCAAAACTGG
ATTACAAATAAATATATAAAAAAAACAATGCGTTTAGCCGCCGTTAATTTCATGTAAGTCATGTTGTTCCTTTTATTTAACCGCGGACT
CCGCTGTTAACCGGAGGATATGCATCATGAAAACCGTTAGGGAGTCCACAACGTTGTACAACTTTCTCGGATCGCACAATCCATACTGG
CGGTTGACGGAAAGCAGCGATGTTTTGCGCTTTTCTACCACCGAAACCACAGAACCTGATCGTACATTGCAGTTATCTGCCGAACAGGC
TGCTCGCATCAGGGAAATGACGGCTCGAGAGATCCTCTAGATTTAAGAAGGAGATATACATATG
>adk
GAGGCAATTCCGACGTCTAAGAAACCATTATTATCATGACATTAACCTATAAAAATAGGCGTATCACGAGGCCCTTTCGTCTTCACCTC
GAGGAACTGCTGCTGGATCAGGCGCTGCTGGCAGAACGCGGCACGCTGGAAGATCCGAACCTGTTTATTCGTCGTATGAACCAGCTGCT
GGTTTCCTGATGTAATGCCGGATGACCTTCGTGTCATCCGGCATTTTTCTTTTCATCATCTGCACTTTCCGCAAATTATCTCGCCATTA
ACCGTTTCAGCCCCAGGTGCCTTTCTTGAGGCAATCGCCTGTTGGTGGTATCGTTTATCGCTTTTTCAAAAAATTCGACACATTTTAAG
GGGATTTTCGCAATGCGTATCATTCTGCTTGGCGCTGGATCCTCTAGATTTAAGAAGGAGATATACATATG
>eutN
GAGGCAATTCCGACGTCTAAGAAACCATTATTATCATGACATTAACCTATAAAAATAGGCGTATCACGAGGCCCTTTCGTCTTCACCTC
GATGGATCCCGGCGAGCTGGTATCTGTACACGTTATCCCGCGTCCGCACGGCGACCTGGAAGAAGTCTTCCCGATCGGCCTGAAAGGCG
ATAGCAGCAACCTGTAATCACCCTGTGTGACGTCGATTCGGGCGTCACGCTCTCTCCTTTTTCATTTACCTTCTGCGGAGGGTAGGGG
GAGCTTTTACCTGAAATACGCCACGGAGGCGGGTATGAAACTGGCAGTCGTCACTGGACAAATTGTTTGTACCGTACGCCATCACGGAC
TGGCGCATGACAAATTGCTCGAGAGATCCTCTAGATTTAAGAAGGAGATATACATATG
>ppdD
GAGGCAATTCCGACGTCTAAGAAACCATTATTATCATGACATTAACCTATAAAAATAGGCGTATCACGAGGCCCTTTCGTCTTCACCTC
GATGGATCCGTGAATTTGCCGAAACGGGCGTGGACTTTATCTCCGTCGGTGCGCTAACTAAACACGTACAAGCACTCGACCTTTCAATG
CGTTTTCGCTAATAGTTGACAGATTTATCCGCTCCATCGCGGGCGGATAATCTCCCCTTCCCCAACTTTCTTCGTAACGCCTCGCAAAT
TTCTTGTCTTTCGCTGCAACGAAATAAAAAACCTCGGAATCTGCTTTCCCGTTTCGTTTTTTGCCCCTCGCCTGTGCTGATGGCGTTTG
CCAAAGTAGCGCCAACCAAATCAAGGAGCGAAACAGATGGACAAGCAACGCGGTTTTACACTTATCGAACTGATGGTGGTTATTGGCAT
CATTGCCATTTTAAGCGCCATCTCGAGAGATCCTCTAGATTTAAGAAGGAGATATACATATG
>otsB
GAGGCAATTCCGACGTCTAAGAAACCATTATTATCATGACATTAACCTATAAAAATAGGCGTATCACGAGGCCCTTTCGTCTTCACCTC
GATGGATCCGCGAAACGCACTGTCTGATGCTTTTTTCTGCAACAATTTAGCGTTTTTTCCCACCATAGCCAACCGCCATAACGGTTGGC
TGTTCTTCGTTGCAAATGGCGACCCCCGTCACACTGTCTATACTTACATGTCTGTAAAGCGCGTTCTGCGCAACACAATAAGAAAAGAG
AAGGAGGAGAACCGGGTGACAGAACCGTTAACCGAAACCCCTGAACTATCCGCGAAATATGCCTGGTTTTTTGATCTTGATGGAACGCT
GGCGGAAATCAAACCGCATCCTCGAGAGATCCTCTAGATTTAAGAAGGAGATATACATATG
>yfjG
GAGGCAATTCCGACGTCTAAGAAACCATTATTATCATGACATTAACCTATAAAAATAGGCGTATCACGAGGCCCTTTCGTCTTCACCTC
GATGGATCCGATGGTCGCTGAACCAGGTTTATGTGCTTTTTTCTTCGTCATAAGCGTCGTGAATCATCGGTAATCTGAAATCGAAAACA
CCCCATATCAATCCTGCGGGGGACAAGGCTCTATCTTAGCATGAACCCGATGTTACCCAGCGCCGGGATAGCGTTTTTTTTACAGCAGG
ATAAATGATATTATTTGTTGGATTTTTGTTGATGGAAATTGTTATGCCTCAGATTAGCCGGACCGCACTGGTACCCTACAGCGCGCTCG
AGAGATCCTCTAGATTTAAGAAGGAGATATACATATG
>pagB
GAGGCAATTCCGACGTCTAAGAAACCATTATTATCATGACATTAACCTATAAAAATAGGCGTATCACGAGGCCCTTTCGTCTTCACCTC
GATGGATCCACCTGTGCTTCCTGGTGCCCGGCGAAGATGCCGAGCAGGTGGTGCAAAAACTGCATAGTAATTTGTTTGAGTAAATACTG
TATGGCCTGGAAGCTATATTTCGGGCCGTATTGATTTTCTTGTCACTATGCTCATCAATAAACGAGCCTGTACTCTGTTAACCAGCGTC

TTTATCGGAGAATAATTGCCTTTAATTTTTTTATCTGCATCTCTAATTAATTATCGAAAGAGATAAATAGTTAAGAGAAGGCAAAATGA
ATATTATCAGTTCTGCTCGCAAACGAATTCCCGTACACTGAACTCTCTGGCTCACTGAGGGCACCATCATGGCACTCCCCCGTATTACC
CAAAAAGAGATGACTGAACGCGAGCAGCGTGAACTTAAAACACTGCTGGATCGGGCTCGAGAGATCCTCTAGATTTAAGAAGGAGATAT
ACATATG
>nadC
GAGGCAATTCCGACGTCTAAGAAACCATTATTATCATGACATTAACCTATAAAAATAGGCGTATCACGAGGCCCTTTCGTCTTCACCTC
GATGGATCCAGGGTGGGTGTTTCGTCATCCGGGCGGCAATCGTAATGTGGTGAGGGAACGCGGCGCGCGCCAACCAGCCACCCCTGTTC
TAACAACATGCAGGATCTCCTTATATGTGGTGCTAATACCCGGTTCAGAGTAGCATGTTTCTACCTTATGATTCGTTAGCTATCTGGAG
TTTTAACATGCCGCCTCGCCGCTATAACCCTGACACCCGACGTGACGAGCTGCTGGAACGCATTAATCTCGATATCCCCGGCCTCGAGA
GATCCTCTAGATTTAAGAAGGAGATATACATATG
>b0663
GAGGCAATTCCGACGTCTAAGAAACCATTATTATCATGACATTAACCTATAAAAATAGGCGTATCACGAGGCCCTTTCGTCTTCACCTC
GAGTGCCACCACTGCGTTTTATGCGTAATCTCGGGTTAATGGCGGCGGAGCGTGCTGGCGTGTTGAAACGTCAGGCGCTGAAATATGCG
TTAGGGTTGTAGCCTTACAACATTGCCGGATGCGTGCCAACCGTAGGTCGGATAAGACGCGGCAGCGTCGCATCCGACATTGAAGGATA
AGACGTGTCAACATCGCATTCGACATTGAATGAACGCAGAAAAGCAAAAAGCTCGCCGAAGCGAGCTTTTTAATGTGGCTGGGGTACG
AGGATTCGAGGATCCTCTAGATTTAAGAAGGAGATATACATATG
>yadG
GAGGCAATTCCGACGTCTAAGAAACCATTATTATCATGACATTAACCTATAAAAATAGGCGTATCACGAGGCCCTTTCGTCTTCACCTC
GAGGCGCTTGTGCCAGTTTCTCAAAAAACCCGGGATCCTCTTCCACCAGCATTTTTGACCATAGTGCATTGTTGCTGATGAGTGTATCT
ATGTCTTTCATGGAGGTTAACGACCTGTAACCAAATAATTACGTTTGGCTAATATAGGGCAACTCCGGGACGATTTAAACCACAGATAA
AGTGTAAGAACGTAAGGTAAGTAAAAATTTATGACCATTGCACTGGAACTTCAACAGCTTAAAAAAACCTATCCAGGCGGCGTTCAGGC
GCTTCGTGGGATAGATTTGCAGGTCGAAGCGGGTGATTAGATCCTCTAGATTTAAGAAGGAGATATACATATG
>rstA
GAGGCAATTCCGACGTCTAAGAAACCATTATTATCATGACATTAACCTATAAAAATAGGCGTATCACGAGGCCCTTTCGTCTTCACCTC
GAGATCAGCCCGGCGATATAATAATTTTTCGTTTTTGCTAAAACACCAATCAACAGCACTACCAGCGCACCGAGCGCGGCTTTGATTAC
CAGCCCCATCTTTTTACCTTAACACTTCCATAACAAGTCATCAGTAGAATACCTGATGAAAACTTGTTTAGAAACGATTGATAGTAAGT
AAAAACAGCGCGGTGTATTGTGACGTTTTTATATCTACCGTGAATGTTATGAACACTATCGTATTTGTGGAAGATGATGCGGAAGTCGG
TTCACTGATTGCCGCGTAGGATCCTCTAGATTTAAGAAGGAGATATACATATG
>grpE
GAGGCAATTCCGACGTCTAAGAAACCATTATTATCATGACATTAACCTATAAAAATAGGCGTATCACGAGGCCCTTTCGTCTTCACCTC
GATGGATCCCAGCCAGCGGTAGAGCATTTCATGTGTTGTCAGTGCAGTGGGGTGCCGTGGGTGTCCCACAATGCCAATACACTTGAAAT
GATTATTCATTTTTCCGAGGTCCTTGTTGCGAAGATTGATGACAATGTGAGTGCTTCCCTTGAAACCCTGAAACTGATCCCCATAATAA
GCGAAGTTAGCGAGATGAATGCGAAAAAAACGCGGAGAAATTCATGAGTAGTAAAGAACAGAAAACGCCTGAGGGGCAAGCCCCGGAAG
AAATTATCATGGATCAGCACGAAGAGATTGAGGCAGTTGAGCCAGAAGCCTCGAGAGATCCTCTAGATTTAAGAAGGAGATATACATAT
G
>murA
GAGGCAATTCCGACGTCTAAGAAACCATTATTATCATGACATTAACCTATAAAAATAGGCGTATCACGAGGCCCTTTCGTCTTCACCTC
GATGGATCCCGCAAACTGAACGGCTTTTGAGCTATGGGCGATTCGCCGGTAGCGGATATGAATTGTTAACTGAGAACAAACTAAATGGA
TAAATTTCGTGTTCAGGGGCCAACGAAGCTCCAGGGCGAAGTCACAATTTCCGGCGCTAAAAATGCTGCTCTGCCTATCCTTTTTGCCG
CTCGAGAGATCCTCTAGATTTAAGAAGGAGATATACATATG
>murC
GAGGCAATTCCGACGTCTAAGAAACCATTATTATCATGACATTAACCTATAAAAATAGGCGTATCACGAGGCCCTTTCGTCTTCACCTC
GAGTTAACCATGGCAGAACGCGCCCGCGCTGCATCCATTCCGGATGCCACCGAGCGAGTGGCAAATGAAGTGAGCCGGGTTGCCCGGGC
GTAATTGTAGCGATGCCTTTTGCATCGTATGAATTTAAGAAGTTAATGGCGTAAAGAATGAATACACAACAATTGGCAAAACTGCGTTC
CATCGTGCCCGAAATGCGTCGCGTTCGGCACATACATTTTGTCGGCATTGGTGGTGCCGGTATGGGATCCTCTAGATTTAAGAAGGAGA
TATACATATG
>molR2
GAGGCAATTCCGACGTCTAAGAAACCATTATTATCATGACATTAACCTATAAAAATAGGCGTATCACGAGGCCCTTTCGTCTTCACCTC
GAGTTGAATGGGATTATGCGAATAATGTCATTATTATTCTGCCGTCTGGCATTGCACCTAGCTACTTGGAGCAATATTCCAGATTTGAA
TGACGCCTACGTAAACATTTATCACTGACGGAAGAGTCTCTCTGGCAAAAATGTGCACAAAAACTTATTGCCGCAATTCCACATATTCC
AGAATGGCGGCAACCATTAATTGCTTTGTTATTACCCGAAAAACCAGAAATTGCACATGAAATTGCCCAGCGTCTACTGGGGCAAAAAA
AATTACCCTCGCTTGAGTGGTTAAGGATCCTCTAGATTTAAGAAGGAGATATACATATG
>dcm
GAGGCAATTCCGACGTCTAAGAAACCATTATTATCATGACATTAACCTATAAAAATAGGCGTATCACGAGGCCCTTTCGTCTTCACCTC
GATGGATCCAGATGCGTTTTCATCCGCTGGCTGAGCGCGTGGCCCTAAATGGCTGTAATTATGTTAACCTGTCGGCCATCTCAGATGGC
CGGTGAAATCTATGCAGGAAAATATATCAGTAACCGATTCATACAGCACCGGGAATGCCGCACAGGCAATGCTGGAGAAACTGCTGCAC
TCGAGAGATCCTCTAGATTTAAGAAGGAGATATACATATG
>yddW
GAGGCAATTCCGACGTCTAAGAAACCATTATTATCATGACATTAACCTATAAAAATAGGCGTATCACGAGGCCCTTTCGTCTTCACCTC
GATGGATCCCACTTCTTCCTGCACCCGCGTGCACGTTCACCACACTATATTGTGATGAATGACAAGAAACACTAACCATGTCTGAATGG
TTATAAGACAAGGGAGCGATAATTCATCGCTCCCTTTTTCGTGCTTGCTGTCGTATTGACTTAACGGGGATAAGTACGAGAATGAGCGC
ACATCTGTTTACCGGAAACCAGCACATATGGATATCTGCTCCCGAAACAAGAAATTAACGATTAGAAGACCAGCGATACTAGTTGCACT
GGCACTTTTACTGTGTAGTTGTAAAAGCACGCCTCCAGCTCGAGAGATCCTCTAGATTTAAGAAGGAGATATACATATG

```
>dcd
GAGGCAATTCCGACGTCTAAGAAACCATTATTATCATGACATTAACCTATAAAAATAGGCGTATCACGAGGCCCTTTCGTCTTCACCTC
GATGGATCCTGCGGACATTATCGTGCCGCGCGGCGGGAAAAACCGCATCGCGATCGATATATTGAAAGCGAAAATAAGTCAGTTCTTTG
AATAAGCTTGATAAATTGTGTACCGTTCAGTGATAACCTAGTATGCCCTTGACGTAATTGGCGTTAAGGGAGTGATGCGCGAAAGGAGA
AAATGCCATGCGTCTGTGTGACCGAGATATTGAAGCCTGGCTTGATGAAGGCCGTTTGTCGATCAACCCACGTCCGCCAGTGGAGCGTA
TTAACGGCGCTCGAGAGATCCTCTAGATTTAAGAAGGAGATATACATATG
>yqaD
GAGGCAATTCCGACGTCTAAGAAACCATTATTATCATGACATTAACCTATAAAAATAGGCGTATCACGAGGCCCTTTCGTCTTCACCTC
GAGCGTTCAGCAGGCCTTTAGTTTACTGGGATGGCCAAGTATTAAAATCAACGACTGAAATTAAAAATTATGATGGGTCCACGCGTGTC
GGCGGTGAGGCGTAGCTTAATAAAGGTTGCTCTACCTATCAGCAGCTCTACAATGAATTCCACGCATTATATATAGATAAAGCAGGAAT
ATGGAGCGCATTTGAGGGTGTATCTGCAACTAAAAACGTTCTGGCCTTTAGTCGTGGGGATTCTGTGAACATTAATCACTCTCCTCATG
ATGGACTAGTTATAATAAATAAAGGAAACGAAGAAGTTGAAGGTACCTGGCCTAACAAATTGCAGGATCCTCTAGATTTAAGAAGGAGA
TATACATATG
>nohA
GAGGCAATTCCGACGTCTAAGAAACCATTATTATCATGACATTAACCTATAAAAATAGGCGTATCACGAGGCCCTTTCGTCTTCACCTC
GATGGATCCATTCCCGGACCCTTTTTGCTCAAGAGCGATGTTAATTTGTTCAATCATTTGGTTAGGAAAGCGGATGTTGCGGGTTGTTG
TTCTGCGGGTTCTGTTCTTCGTTGACATGAGGTTGCCCCGTATTCAGTGTCGCTGATTTGTATTGTCTGAAGTTGTTTTTACGTTAAGT
TGATGCAGATCAATTAATACGATACCTGCGTCATAATTGATTATTTGACGTGGTTTGATGGCCTCCACGCACGTTGTGATATGTAGATG
ATAATCATTATCACTTTACGGGTCCTTTCCGGTGATCCGACAGGTTACGGGGCGGCGACCTCGCGGGTTTTCGCTATTTATGAAAATTT
TCCGGTTTAAGGCATTTCCGTTCTTCTTCGTCGTAACTTAATGTTTTTATTTAAAATACCCCCTGAAAAGAAAGGAAACGACAGGTGCT
GAAAACGAGCTTTTGGGCCTCTGTCGTTTCCTTTCTCTGTTTTTGGCCGTGGAATGAACAATGGAAGTCAACAAAAAGCAGCTGGCTGA
CATTTTCGGTGCGAGTATCCGTACCATTCAGAACTGGCAGGAACAGGGAATGCCCGTTCTGCGAGGCGGTGGCAAGGGTAATGAGGTGC
TTTATGACTCTGCCGCCGTTATAAGATGGTATGCCGAAAGGGATGCTGAAATTGAGAACGAAAAGCTGCGCCGGGAAGTTGAAGAACTG
CGGCAGGCCAGCGAGACAGATCTCCAGCCAGGGACTATTGAGTACGAACGCCATCGACTTACGCCTCGAGAGATCCTCTAGATTTAAGA
AGGAGATATACATATG
>gudP
GAGGCAATTCCGACGTCTAAGAAACCATTATTATCATGACATTAACCTATAAAAATAGGCGTATCACGAGGCCCTTTCGTCTTCACCTC
GATGGATCCTGCCTGGAGTCAAGCGATCCATTTCATACTCTTCTTTATTTCTTCGTTTTAACCCTTCCTTTCTTGTTCTTGTTTTCATT
TCCGTGAAGTGGATTCCACCGTCCAGGGCTAATGCCAAAATCGGGCCTCATTGAACGCATTAATGTTGTGTTGTTGCACGGTGAGCCGC
TATGGCGCGCTTTTTATACTGCTATTGCCAGATATAAACACGCGCCGTATTCGGCGAACGACCTATAAAAACGGCAAAAAACACCCTAC
GTCACCTCTGATTTCCTGGCGATGTCGCAGTCCAGAGTGAGCGTGGCTAACGCGAATTTTCAGGAGTGCAACAATGAGTTCTTTAAGTC
AGGCTGCGAGCAGTGTGGAAAAACGCACAAATGCTCGTTACTGGATAGTGGTGATGTTGTTTATCGTCACATCCTTCAACTACGGCGAC
CGCGCTACGCTCTCTATCGCCGGTTCGGAAATGGCCAAAGATATCGGCCTTGATCCCGTGGGAATGGGCTATGTGTTCTCTGCTTTCTC
ATGGGCTTATGTTATCGGGCAGATCCCTGGTGGCTGGTTGCTGGACCGTTTTGGTTCAAAACGCGTCTACTTCTGGTCGATCTTTATCT
GGTCGATGTTTACCTTGCCTCGAGAGATCCTCTAGATTTAAGAAGGAGATATACATATG
>yddM
GAGGCAATTCCGACGTCTAAGAAACCATTATTATCATGACATTAACCTATAAAAATAGGCGTATCACGAGGCCCTTTCGTCTTCACCTC
GATGGATCCGCCGCATGGTGATTGATTTCCGTCACTAAGAGGCCTTTGCTGCGACTGCCATGTTCGGGTCGCAGCATCGCACACTCTCC
AACATGAAATGGCTAAATGGATTATTGTTGCGTAGGAAGGAACAATAATGATAGTGTTCCCAGTGAAGGAACGCTACTGAGAGTAAAGA
CACTAATGAAAATGGCCAATCATCCCTCGAGAGATCCTCTAGATTTAAGAAGGAGATATACATATG
>yddL
GAGGCAATTCCGACGTCTAAGAAACCATTATTATCATGACATTAACCTATAAAAATAGGCGTATCACGAGGCCCTTTCGTCTTCACCTC
GATGGATCCTTCTCGTTCTGGCAAGGCGCGCTGATGGTCTGCGGCGGTTCCCTGCTCTGCTGGCTGGCGACACGTCGTGGTTAAAATAA
CTGCCGGGTCTACGGCCCGGTAGTTAATCAGATAAATTCCGACAATATTAAAAAAATAAATTTCAAATACCAGTAATTCACCTTATATC
TCATTTCGACTTCGATCACATATCGATAAATAATATTATTTCATCTATTTTTCTCATTGTTTTTAATCACCATTCTATTTTAAAAACAG
TATGTCATTTATAAATGACGCCGTTTAAAATTCGTTTAGAAAATACGTTTAACGATAATTAAAAGGATTCTGTTAAATGAAATTAAAAA
TAGTTGCGGTGGTTGTAACTGGTTTGTTAGCTGCGAACGTAGCACACGCTGCCGAAGTCCTCGAGAGATCCTCTAGATTTAAGAAGGAG
ATATACATATG
>yddH
GAGGCAATTCCGACGTCTAAGAAACCATTATTATCATGACATTAACCTATAAAAATAGGCGTATCACGAGGCCCTTTCGTCTTCACCTC
GATGGATCCGCAGCTCCCGACCAGTTAATACGGGCAAAATAGTGATTCAGAATGGGCGTCATCAGGCGTTTCCCTTAGTGGTTTTCTCA
ACTTATAACCTGGAATAGACGCCGCACCGGCTATTTTGTGCGATCGCGGACATGCATCACGTTCGCAAATCGCCAATCCCGTTAATTTC
GTTACTTATTGCGTTACGGACTACGCTTAAAAAAACCTTCTTCGCGAGGCGATACCATGAGCCGATTCATCCCCATAGAATTACACCAT
GCCAGCCGCTCGAGAGATCCTCTAGATTTAAGAAGGAGATATACATATG
>yddE
GAGGCAATTCCGACGTCTAAGAAACCATTATTATCATGACATTAACCTATAAAAATAGGCGTATCACGAGGCCCTTTCGTCTTCACCTC
GATGGATCCCTGGTGCATGTGTGGAGCGCGCCGTTTGAGTACTTTACTCGTCGATATCAAATTGTGCGTTCGCGGCGATAATTTCACCC
TGGCCGGATAGCCCTTCGCTCTCCGGCCACTTGCAGGCTATGATAGTTCTCCAGTCACAGGAGAACGTTAATGAAACCGCAGGTGTACC
ACGTCGATGCTTTTACCTCACAACCGTTTCGCGGCAATTCTGCCGGTGTTGTTTTCCCTCGAGAGATCCTCTAGATTTAAGAAGGAGAT
ATACATATG
>dcp
GAGGCAATTCCGACGTCTAAGAAACCATTATTATCATGACATTAACCTATAAAAATAGGCGTATCACGAGGCCCTTTCGTCTTCACCTC
GATGGATCCTTCACCAAAACCTGCCGTTGCTCCAGTTACTAAAACGATCATCTCACTGTTCCTCAACGCTTTTGTGTGTCAATAACATA
```

GCACGGGATGCTTAAGAGCGGGTAATACTGGATTGCAAGATTGCACTCCGTTATCAGTAAGCCTACGCTATGGAACACCTCGCGTTCAG
GAGAAGAAAATGACAACAATGAATCCTTTCCTTGTGCAAAGCACACTGCCGTATCTGGCTCCCCATTTTGATCAAATTGCCAATCATCA
CTATCGCCCGGCTCGAGAGATCCTCTAGATTTAAGAAGGAGATATACATATG
>b1403
GAGGCAATTCCGACGTCTAAGAAACCATTATTATCATGACATTAACCTATAAAAATAGGCGTATCACGAGGCCCTTTCGTCTTCACCTC
GATGGATCCGCCTGTTCCGTTCTTCCATAGTTCAAAAACAGAGGCTTTTTCCTCTGCTGTAAATGTTCGTCTCATTCAAAAAACCTCCG
CAACCCCATGTTTTCACATAACTGTTGCGTTGACCAATTGAATCTACAAGTACCTTTATTAACAATTTTCGCACCGGAATCAGCATAAG
TAAGTTCTGGGGTTTAGACTGGCCCCCTGAATCTCCAGACAACCAATATCACTTAAATAAGTGATAGTCTTAATACTAGTTTTTAGACT
AGTCATTGGAGAACAGATGATTGATGTCTTAGGGCCGGAGAAACGCAGACGGCGTACCACACAAGATCCTCTAGATTTAAGAAGGAGAT
ATACATATG
>yabP
GAGGCAATTCCGACGTCTAAGAAACCATTATTATCATGACATTAACCTATAAAAATAGGCGTATCACGAGGCCCTTTCGTCTTCACCTC
GAGGGAGATGGCGAAGCAGAAAGCGCAGGAAATTCAGCAGGCATATGAGCTGATAAAGCAGCAGAAAGGGTTTAAATGACCCTGTAAAT
GATGCTGAGTAACTGCCCACGATTAAAGGTGGCCGCCCTGGCGGTCACTTCTTTGAGAAAAGGCGTTTACTCAGAATGGTGGACAGGCT
CAATGCACGGTTTACGGGAGGGGTTCTGTAGGTTTTATCGCGTTGACCCTGCTTAAGGTTGAGAGCTTTACGACGAGCGGAATTATATT
TTTACGTCTTAAAAATAAAAAACACATACCTGAATGAGCGATTTTTGAAAGTATATTTATTCAGAACGCGCATCATGAGTTTTTAACTC
AATGCGAGGCTATTACCATGAAAGTATCAGTTCCAGGCATGCCGGTTACACTTTTAAATATGAGCAAGAACGATATTTATAAGATGGTG
AGCGGGGACAAGATGGACGTGGGATCCTCTAGATTTAAGAAGGAGATATACATATG
>yfhH
GAGGCAATTCCGACGTCTAAGAAACCATTATTATCATGACATTAACCTATAAAAATAGGCGTATCACGAGGCCCTTTCGTCTTCACCTC
GAGCGGTAACAACGGCAGGACAAGTAACGCATTCAGCGGTTGGCGACGTAGTAAATAGCGCAGAAAACTGCCGAACATATCCTGCTGAT
GCAATGTTCCATCTAAGTCAAAAAACACCACACGACGCTCGTGAGTTGCCAAACCATACTCCTCTGGACTGAAACATCCTGATTAATTT
TCGCTACTACATAGCCTAACAGATAGATCATCACTTTTCCGGCAACAATCGACCGCATCATGGCTAACTGGAATTTTTAATTCATGGCA
ATTAGCGGCAATGGAATATAAAATTCACTCGCGTGTGTCTCATATTGCCTGTTGTCGCCACTATTTCATCGCGAGGAAAGATATGAACG
GCTTACTTCGTATCCGTCGGATCCTCTAGATTTAAGAAGGAGATATACATATG
>ndk
GAGGCAATTCCGACGTCTAAGAAACCATTATTATCATGACATTAACCTATAAAAATAGGCGTATCACGAGGCCCTTTCGTCTTCACCTC
GATGGATCCTGGTGATGGATGACGTGGGGCAAATCGCGACAGTGAAATTTGTCATGCAATAGTCTGTTTTGTCCCTGATTGTTGCTAAA
ACTCATTTTATTTTAAAAAAATGTTACCTTCATCAATAGTCAACGGCCCTGTTGCTCATTATAATCCGCGCCATCTCGTACGCTGGTAC
AGACAACAACAGAACAATTTACAGAGGTAAAAATGGCTATTGAACGTACTTTTTCCATCATCAAACCGAACGCGGTAGCAAAAAACGTC
ATTGGTAATATCTTTGCGCGCTTTGAACTCGAGAGATCCTCTAGATTTAAGAAGGAGATATACATATG
>yfhM
GAGGCAATTCCGACGTCTAAGAAACCATTATTATCATGACATTAACCTATAAAAATAGGCGTATCACGAGGCCCTTTCGTCTTCACCTC
GATGGATCCTGGCTGTCTGAGCCACAATTCTCACTAAAATTCGGATTGTATGGGCGTGATACTGAATTACATAACGTAAATTCTCACGC
ATAATAATTATTCGACGTTGCTAACGGGCCATTGTGGCAGGGATGAAAAATGAAAAGTTACGCGTAGCCGCCTGCATGCTAATGCTGG
CGCTGGCAGGGTGCGACAACAACGATAACGCGCCAACAGCGGTGAAAAACTCGAGAGATCCTCTAGATTTAAGAAGGAGATATACATAT
G
>yfhL
GAGGCAATTCCGACGTCTAAGAAACCATTATTATCATGACATTAACCTATAAAAATAGGCGTATCACGAGGCCCTTTCGTCTTCACCTC
GAGTTGCTGTTCATTGCGCTGATTCAGCAGGATCTGGAACTGGCACCAGAACGTATTCGTCATAGTGAAGCGCTGGTGAAAAAACTGGT
CTGAGTAAAGAATGCGCGTATAATGCCCGCCCGGTTTGTGTTGTTTTGAGAGTTTCCTTATGGCGTTGTTAATCACTAAAAAATGCATC
AATTGTGATATGTGTGAACCCGAATGCCCGAATGAGGCGATTTCAATGGGAGGATCCCGGATCCTCTAGATTTAAGAAGGAGATATACA
TATG
>yfhB
GAGGCAATTCCGACGTCTAAGAAACCATTATTATCATGACATTAACCTATAAAAATAGGCGTATCACGAGGCCCTTTCGTCTTCACCTC
GATGGATCCTAGTGGCGACAACAGGCAATATGAGACACACGCGAGTGAATTTTATATTCCATTGCCGCTAATTGCCATGAATTAAAAAT
TCCAGTTAGCCATGATGCGGTCGATTGTTGCCGGAAAAGTGATGATCTATCTGTTAGGCTATGTAGTAGCGAAAATTAATCAGGATGTT
TCAGTCCAGAGGAGTATGGTTTGGCAACTCACGAGCGTCGTGTGGTGTTTTTTGACTTAGATGGAACATTGCATCAGCAGGATATGTTC
GGCAGTTTTCTGCGCTATTTACTACGTCGCCAACCGCTGAATGCGTTACTTGTCCTGCCGTTGTTACCGCTCGAGAGATCCTCTAGATT
TAAGAAGGAGATATACATATG
>yfhG
GAGGCAATTCCGACGTCTAAGAAACCATTATTATCATGACATTAACCTATAAAAATAGGCGTATCACGAGGCCCTTTCGTCTTCACCTC
GATGGATCCCGGGCAAGACGTTTGTTTCCGCATTGAATTACCGTCGTCGAAAAACACGAAATAATGCTAACAGCCTATCCTATCAGGCT
ATTTTATTTGTCAGTTCGGCCCCGGGCAGTGCTCGAAATCTTCACGTACGATATGTACGCTCCGGTTTCTCCGCGCTGTCCATGTCCAG
ACTGTCTGCAACAATTACGCCCACTGGGATAGGCTTTAAGTCTGGTGAATATGCGACACATTTTTCAACGATTATTGCCCCGACGGCTG
TGGCTGGCAGGGCTGCCATGTCTGGCATTGCTGGGTTCTCGAGAGATCCTCTAGATTTAAGAAGGAGATATACATATG
>yabI
GAGGCAATTCCGACGTCTAAGAAACCATTATTATCATGACATTAACCTATAAAAATAGGCGTATCACGAGGCCCTTTCGTCTTCACCTC
GAGCCAGCCCGAGCGAGTTTCGTGCCGGTTGTGAAGAAAAAGTGAATGATGTAGCCGTCAAGTTGTCATAATTGGTAACGAATCAGACA
ATTGACGGCTTGACGGAGTAGCATAGGGTTTGCAGAATCCCTGCTTCGTCCATTTGACAGGCACATTATGCAAGCATTGCTGGAACACT
TTATTACCCAATCCACCGTGTATTCATTGATGGCGGTGGTGTTGGTGGCCTTTCTGGAGGGATCCTCTAGATTTAAGAAGGAGATATAC
ATATG
>yfhR

GAGGCAATTCCGACGTCTAAGAAACCATTATTATCATGACATTAACCTATAAAAATAGGCGTATCACGAGGCCCTTTCGTCTTCACCTC
GAGTGCTGACCGGTAACATCGTTGCTGGTAACCCGCGCGTTGTTAAAGCCATGCTGGCGAACATGCGTGACGAGTTAAGCGACGCTCTG
AAGCGTTAATGACTCAGGCGGGTGATATCACTCACCCGCCCTCGCCTTTCAGGCGCTATTCCGAAATACTTCCTCACTGCTTTACTTTC
TTTCGTCACTCTCCCACCATCTTTCCCCGATTAATGGATGATAAAGAAGTCGCAAGGAAAGGTTATGGCACTGCCAGTGAACAAACGCG
TTCCCAAAATTCTGTTTATTCTCTTTGTTGTTGCCTTCTGCGGGATCCCGCAGAAGGCAACAACAACCGCTCGATGTTACCGGTCAGCA
CTCGAGCGGGGATCCTCTAGATTTAAGAAGGAGATATACATATG
>yabN
GAGGCAATTCCGACGTCTAAGAAACCATTATTATCATGACATTAACCTATAAAAATAGGCGTATCACGAGGCCCTTTCGTCTTCACCTC
GATGGATCCAAATGCCGCGTAAACACCGTTCATACGGCGAGCCATCGTCATTATCCAGATCATACGTTCCCTTTTTAGCGCGGCGAGAA
TAAAAAAAACCAGCAGGTATAATCTGCTGGCGGGTGATTTTACACCAATACTCAGTCACACATGATGCAGGCAAGTCAACTTTCAGAAT
TGCGGTCATCCCACTGCATCAGTCCTTCAAGCATGGTTAATCGTTGTGGGACGCTTAACCAACGCAACCAGCACAACTTCGCTGTCGCG
GTAAAATAGTGCTGATAAAACTGACGCATGGGGCACCCCCTTGCTTCATCGTTAGGAATTATTGACTTAATATAGGGAAAATAAAATTG
CTGTCTTTTGCACAGGAGTTCCCCTTTTATGCCATCTGCTCGTCTGCAACAACAGTTCATCCGCCTGTGGCAATGCTGCGAGGGTAAAT
CGCAGGACACAACGCTCAACGAACTGGCCTCGAGAGATCCTCTAGATTTAAGAAGGAGATATACATATG
>cvrA
GAGGCAATTCCGACGTCTAAGAAACCATTATTATCATGACATTAACCTATAAAAATAGGCGTATCACGAGGCCCTTTCGTCTTCACCTC
GATGGATCCAATAACACCCGGGAAGGCACTCAGTTGACGATTAGCGGTCATCCTGTTCTTAAAATGTAAAAAAATCACGCTTCAGGGCT
AAGTAAAACGCTGTCTCTGCCGCTAATATGGTAGGGTTTATATAAGAGACAGCGTAATCAGGAGTAACCGACCGTTGGATGCCACAACA
ATAATTAGCCTTTTCATCTTAGGATCCATCCTTGTCACCAGCAGTATATTACTTAGTTCATTTTCTTCCCGTCTTGGCATTCCTATTCT
GGTTATTTTTCTGGCGATCGGCATGCTGGCAGGAGTCGATGGCGTCGGCGGTATCCCGTTTGATAATTACCCCTTCGCCTACATGGTAA
GTAACCTGGCACTGGCGCTCGAGAGATCCTCTAGATTTAAGAAGGAGATATACATATG
>yaaU
GAGGCAATTCCGACGTCTAAGAAACCATTATTATCATGACATTAACCTATAAAAATAGGCGTATCACGAGGCCCTTTCGTCTTCACCTC
GAGCACCTGTCGCATTCTGGGGCTGGGGTCGGCGCTGGAACAGTGGGAATACCCGCGCGGCACCTTTGGTGTGGAGTTCCGTTACGGCT
GATGTTGGTTTGATACGTAACGCCGCACTGACTCTCATTGCAAAAAACAGGAATAACCATGCAACCGTCCAGAAACTTTGACGATCTCA
AATTCTCCTCTATTCACCGCCGCATTTTGCTGTGGGGAAGCGGTGGTCCGTTTCTGGATGGGGATCCTCTAGATTTAAGAAGGAGATAT
ACATATG
>mazE
GAGGCAATTCCGACGTCTAAGAAACCATTATTATCATGACATTAACCTATAAAAATAGGCGTATCACGAGGCCCTTTCGTCTTCACCTC
GATGGATCCACCAGGTGCCGGATGTTATCGACGCGCGTCGGTTGCACGGGAGTTAGGCCGAAATTTGCTCGTATCTACAATGTAGATTG
ATATATACTGTATCTACATATGATAGCGGTTTGAGGAAAGGGTTATGATCCACAGTAGCGTAAAGCGTTGGGAAATTCACCGGCGGTG
CGGATCCCGGCTACGTTAATGCAGGCGCTCAATCTCTCGAGAGATCCTCTAGATTTAAGAAGGAGATATACATATG
>mazG
GAGGCAATTCCGACGTCTAAGAAACCATTATTATCATGACATTAACCTATAAAAATAGGCGTATCACGAGGCCCTTTCGTCTTCACCTC
GATGGATCCGGCAAGAGGAGCAACGAAGAAAGGAACAGTTGCCCCAGAGGAATTACAACTCATTAAAGCCAAAATTAACGTACTGATTG
GGTAGTGTTACTAACTCACTTCTATTCTGGTCACAGGTTTCCACCTGACAGACCCGCAATTTTCAGGACAATTCAATGAATCAAATCGA
CCGTTTGCTCACTATTATGCAGCGCCTGCGCGATCCGGAAAACGGCTGCCCGTGGGATAAAGACTCGAGAGATCCTCTAGATTTAAGAA
GGAGATATACATATG
>appA
GAGGCAATTCCGACGTCTAAGAAACCATTATTATCATGACATTAACCTATAAAAATAGGCGTATCACGAGGCCCTTTCGTCTTCACCTC
GAGTTGCCCATTGTGTTGCTCTACACTCTCTGGAGCTACTACAAAATGTGGGGGCGCATGACAACAGAAACTCTCCGCCGTAACGAAAA
CGAGTTGTACTAAGGAGCAGAAACAATGTGGTATTTACTTTGGTTCGTCGGCATTTTGTTGATGTGTTCGCTCTCCACCCTTGTGTTGG
TATGGCTGGACCCGCGTCTGAAAAGTTAACGAACGTAGGCCTGATGCGGCGCATTAGCATCGCATCAGGCAATCAATAATGTCAGATAT
GAAAAGCGGAAACATATCGATGAAAGCGATCTTAATCCCATTTTTATCTCTTCTGATTCCGTTAACCCCGCAATCTGCATTCGCTCAGA
GTGAGCCGGAGCTGAGGATCCTCTAGATTTAAGAAGGAGATATACATATG
>yjfK
GAGGCAATTCCGACGTCTAAGAAACCATTATTATCATGACATTAACCTATAAAAATAGGCGTATCACGAGGCCCTTTCGTCTTCACCTC
GAGGCGACCTTGACGAAAAGCTGGCGGAAGCCGGAATTGGCGGTAGCAATAAAAGTAGCGCCCAGGATGTATTAGCAAGACTGCAACGC
CAACAGGGCGAGTAATTTTTTTGCCAGCCTCGCTAAAAGGCTGGCAACTATTTTAAGGATAAAATATGTCTGGTTTTTTCCAGCGTCTG
TTTGGCAAGGATAATAAGCCAGCTATCGCTCGTGGTCCGCTGGGACTTCATCTCGGATCCTCTAGATTTAAGAAGGAGATATACATATG
>yjfI
GAGGCAATTCCGACGTCTAAGAAACCATTATTATCATGACATTAACCTATAAAAATAGGCGTATCACGAGGCCCTTTCGTCTTCACCTC
GAGTGGCTGGAAGCGTTTCTTCCCTGAACGTTTCGGTTGCGACCGGAATTTGCTTATTTGAAGCGGTGCGCCAGCGCAGCTAATTTCTC
AGAATTATGAAAAGCCATCCAGATTTGGATGGTTTTTTTTTGTCTATAGCTGGTAAGATAATTACGTATTGCAAATATTCATTTTATTA
ATATTTAAACTTAATCTATATGGAATAATATATGACATGGAATCCGTTGGCGCTAGCGACGGCGCTGCAAACTGTACCTGAACAAAATA
TTGATGTAACAAATAGCGAGAACGCAGGATCCTCTAGATTTAAGAAGGAGATATACATATG
>mraZ
GAGGCAATTCCGACGTCTAAGAAACCATTATTATCATGACATTAACCTATAAAAATAGGCGTATCACGAGGCCCTTTCGTCTTCACCTC
GAGGAACCGCGTAAGCCAAAACCTGGTTTAACGCGCATTAAACGTAATCTCTATCGCCGCGGCGTGCTCAGCCGTAGCTAAGCCGCGAA
CAAAAATACGCGCCAGGTGAATTTCCCTCTGGCGCGTAGAGTACGGGACTGGACATCAATATGCTTAAAGTAAATAAGACTATTCCTGA
CTATTATTGATAAATGCTTTTAAACCCGCCCGTTAATTAACTCACCAGCTGAAATTCACAATAATTAAGTGATATCGACAGCGCGTTTT
TGCATTATTTTGTTACATGCGGCGATGAATTGCCGATTTAACAAACACTTTTCTTTGCTTTTGCGCAAACCCGCTGGCATCAAGCGCCA
CACAGACGTAACAAGGACTGTTAACCGGGGAAGATATGTCCTAAAATGCCGCTCGCGTCGCAAACTGACACTTTATATTTGCTGTGGAA

AATAGTGAGTCATTTTAAAACGGTGATGACGATGAGGGATTTTTTCTTACAGCTATTCATAACGTTAATTTGCTTCGCACGTTGGACGT
AAAATAAACAACGCTGATATTAGCCGTAAACATCGGGTTTTTTACCTCGGTATGCCTTGTGACTGGCTTGACAAGCTTTTCCTCAGCTC
CGTAAACTCCTTTCAGTGGGAAATTGTGGGGCAAAGTGGGAATAAGGGGTGAGGCTGGCATGTTCCGGGGAGCAACGTTAGTCAATCTC
GACAGCAAAGGGCGCTTATCAGTGCCTACCCGTTATCGGGAACAGCTGCTTGGGATCCTCTAGATTTAAGAAGGAGATATACATATG
>ydaQ
GAGGCAATTCCGACGTCTAAGAAACCATTATTATCATGACATTAACCTATAAAAATAGGCGTATCACGAGGCCCTTTCGTCTTCACCTC
GATGGATCCGAAGCAGGCGTTTCCAGAGAGCAGAAAGTAAAACTGATAAATTTTAATTGCGAGAAACTTCAGTCCTCGAGAATTGCATT
GTATTCAAATTAAAACTTCATAGCTGATTATTAATAATCAACATCGGGCGTCAATTTCAGTCTAACATTGGCGCCTGCCAGAGGTGATG
CGATGGCACAAGTAATCTTTAATGAAGAGTGGATGGTTGAATACGGCCTGATGCTTCGCACTGGTCGTCGAGAGATCCTCTAGATTTAA
GAAGGAGATATACATATG
>hdhA
GAGGCAATTCCGACGTCTAAGAAACCATTATTATCATGACATTAACCTATAAAAATAGGCGTATCACGAGGCCCTTTCGTCTTCACCTC
GATGGATCCCACCCATGAAGAGACGCATTCACGCAATCTTATTATTCCCGCCCGGCTCATTGCAGCGAAATAATCCTCTCTTTATCTGC
TATACCTGGTAGTGTCCTTTCCTCAAGGTTAATGATAAAAAATGTGGCACACAGGCGGTGCCTTTTATTTTTACTGAAGTTCACAGGAG
GTTTATGTGTTTAATTCTGACAACCTGAGACTCGACGGAAAATGCGCCATCATCACAGGTGCGGGTGCAGGTATTGGTAAAGAAATCGC
CATTACATTCGCGACACTCGAGAGATCCTCTAGATTTAAGAAGGAGATATACATATG
>yjfY
GAGGCAATTCCGACGTCTAAGAAACCATTATTATCATGACATTAACCTATAAAAATAGGCGTATCACGAGGCCCTTTCGTCTTCACCTC
GATGGATCCCACCAGTGATGGCAGCAGTGTAGCGCTCGATCATGCCCGGAACCTGTTCGCTCTGGTCAGGATGGACCATAAAAACGATT
TCGTAATGACGCATCGAATTGCTCCTTACGGATTATTCAGCCTCCTGTCTGGGTCAGCCGAATCCCGGGGAGGCAAGGAACGTGTTAAA
GGTCGGCTGAAAAATGACGCGTTATAGTACTTGTATGCCTCGCTAAACTCAAGGGCAGAGCCGAAATAAATCTCCCCTTTCGCAATTAA
CCTCTAACACGCTGATTTGAAGTTCATCACACTTCATCATGGTGTTTTTTTGAACAACTGCATCAGAAATGGTCACCATCAACTTCTTT
ATCATTGATTAAAATTAAATCAGAAGATCATCACAGAGGAGAGCACACAAAGTCAGCCGTCGAGGAGGTATCTATGTTCAGTCGTGTTT
TAGCCCTTCTGGCTGTGCTTTTGCTAAGTGCAAATACATGGGCAGCCATTGAAATTCTCGAGAGATCCTCTAGATTTAAGAAGGAGATA
TACATATG
>yjfQ
GAGGCAATTCCGACGTCTAAGAAACCATTATTATCATGACATTAACCTATAAAAATAGGCGTATCACGAGGCCCTTTCGTCTTCACCTC
GATGGATCCCTGGCCGCTGGATAAAGACAACTTCGAGTACCACTATCCGCGCGGTTTCGATGATTGCTTCACTATTGAACCGGATCTGC
CGTTCAAGTCATTCCTGTAATCTGTCGTTTATACCGGGTAGCGTGGAGAAGTTCGCGCTATCCGGCGTAGTATTTCCTGGCTATTTCAA
ATATCATCTAAAAAAATCAAATTTTATCGGAATAGCTCATGACTGAAGCACAAAGACATCAAATCCTCCTGGAAATGCTCGCACAATTG
GGCTTTGTGACCGTTGAGAAAGTCGTTGAGCGTCTGGGAATCTCGAGAGATCCTCTAGATTTAAGAAGGAGATATACATATG
>yhdA
GAGGCAATTCCGACGTCTAAGAAACCATTATTATCATGACATTAACCTATAAAAATAGGCGTATCACGAGGCCCTTTCGTCTTCACCTC
GATGGATCCAGTTTTGCCGTCCTGCTGTTCTAAAAGTAACGCCTGCATAAGTGGTCCTCATGTGCATGATGGATTGGAAAATAATTTCT
GAAGACTATACTCGCTAATGGAAAAGCAAAGCCGATGAAGCGCAATAAATTGCGAGATAAATCTGATTTGCTAGTATGCCCGCTTCCTC
ACTATCGGAGTTAACACAAGGATGAGATTAACGACGAAATTTTCGGCCTTTGTTACGCTGCTCACCGGGTTAACAATTTTTGTGACTTT
GCTGGGCTGTTCGCTCGAGAGATCCTCTAGATTTAAGAAGGAGATATACATATG
>fliY
GAGGCAATTCCGACGTCTAAGAAACCATTATTATCATGACATTAACCTATAAAAATAGGCGTATCACGAGGCCCTTTCGTCTTCACCTC
GATGGATCCTGCAGAAATCCTCCAGCCTGCCTTCTTCTGATATATATTAAATAAGAATAAGATGTAGCGGAGTTGTTTTTGTGTTTACA
AACAATGGCTCTACACTGCAAACAGACATAACAACATTCGGGGTGAATATGAAATTAGCACATCTGGGACGTCAGGCATTGATGGGTGT
GATGGCCGTGGCGCTGGTTGCGGGCATGAGCGTTACTCGAGAGATCCTCTAGATTTAAGAAGGAGATATACATATG
>yhdJ
GAGGCAATTCCGACGTCTAAGAAACCATTATTATCATGACATTAACCTATAAAAATAGGCGTATCACGAGGCCCTTTCGTCTTCACCTC
GAGTGGTAACCAGACCCGTGCTGCGCTGATGATGGGCATCAACCGTGGTACGCTGCGTAAAAAATTGAAAAAATACGGCATGAACTAAT
TCAGGTTAGCTAAATGCTTGATTAAAAAGGCGCTACTCGGCATGGGGAAGCGCCTTTTTTATAGGTGTCACAAAGGGAGTGACCATGAG
AACAGGATGTGAACCGACCCGGTTTGGTAATGAAGCTAAGACCATTATTCACGGTGATGCGCGGATCCTCTAGATTTAAGAAGGAGATA
TACATATG
>yhdH
GAGGCAATTCCGACGTCTAAGAAACCATTATTATCATGACATTAACCTATAAAAATAGGCGTATCACGAGGCCCTTTCGTCTTCACCTC
GAGCGAACAGCCCAGCAAAGTCACAAAAATTGTTAACCCGGTGAGCAGCGTAACAAAGGCCGAAAATTTCGTCGTTAATCTCATCCTTG
TGTTAACTCCGATAGTGAGGAAGCGGGCATACTAGCAAATCAGATTTATCTCGCAATTTATTGCGCTTCATCGGCTTTGCTTTTCCATT
AGCGAGTATAGTCTTCAGAAATTATTTTCCAATCCATCATGCACATGAGGACCACTTATGCAGGCGTTACTTTTAGAACAGCAGGACGG
CAAAACTCTCGCATCAGTACAGACTCTGGACGAAAGTCGCCTGGATCCCGGGATCCTCTAGATTTAAGAAGGAGATATACATATG
>yhdN
GAGGCAATTCCGACGTCTAAGAAACCATTATTATCATGACATTAACCTATAAAAATAGGCGTATCACGAGGCCCTTTCGTCTTCACCTC
GATGGATCCGAAGTGTGGCTTCCGTGCAGGCGACAACGCGCCGATGGCTTACATCGAGCTGGTTGATCGTTCAGAGAAAGCAGAAGCTG
CTGCAGAGTAATCTGAAGCAACGTAAAAAAACCCGCCCCGGCGGGTTTTTTATACCCGTAGTATCCCCACTTATCTACAATAGCTGTA
CTCTTTTTGTTCATCCCCTGGAGTATTTATGTGGTTACTTGACCAGTGGGCAGAGCGCCATATAGCAGAAGCGCAAGCGAAAGGTGAGT
TTGATAACCTGGCAGGTAGCGGCGAACCATTGATACCTCGAGAGATCCTCTAGATTTAAGAAGGAGATATACATATG
>yffH
GAGGCAATTCCGACGTCTAAGAAACCATTATTATCATGACATTAACCTATAAAAATAGGCGTATCACGAGGCCCTTTCGTCTTCACCTC
GATGGATCCTGACTGGCTGGGGGTAAAATCAGTCAAATCTCACTGATAGCCTGCGCAGACAAACCCGACTTCACAGCGTAAGATAATTG

Data Set S4 - MG1655 library reference file.fasta
TTCATTTCGCGCTGTGGAGTCGGTATGACGCAACAAATCACCCTCATTAAAGACAAAATTCTCTCCGATAACTATTTCACCCTGCACAA
CATTACTTACGATCTCACGCGCAAACTCGAGAGATCCTCTAGATTTAAGAAGGAGATATACATATG
>fliL
GAGGCAATTCCGACGTCTAAGAAACCATTATTATCATGACATTAACCTATAAAAATAGGCGTATCACGAGGCCCTTTCGTCTTCACCTC
GAGGACGACGATACGCTTCCGGTTCCCGTCTCTTTACAAGGGCGTGTAACAGGCAACAGCGGCGTTGATATTTTCGCCTAACGTCAGAG
GTAGCACCGTAATCCGCGTCTTTTCCCCGCTTTGTTGCGCTCAAGACGCAGGATAATTAGCCGATAAGCAGTAGCGACACAGGAAGACC
GCAACACATGACTGATTACGCGATAAGCAAGAAAAGCAAGCGATCGCAGATCCTCTAGATTTAAGAAGGAGATATACATATG
>yffS
GAGGCAATTCCGACGTCTAAGAAACCATTATTATCATGACATTAACCTATAAAAATAGGCGTATCACGAGGCCCTTTCGTCTTCACCTC
GAGAGAATGCGCCAACGTAGCAGAAAACAACGGCTTTGTTTTGGTAGGTAGAGTATTTGAAAAACGTAATTGATTGCGAATAGCAACAA
CTAAACAACCAATACACCCGGCCCCTCGCCGGGTTTTTTGTGATCTGCGTCAATATTCCCTTCCGTTACGCTTACCCTCCACCAATACG
CGTGTTAACCTCCCAATGGATTCTATGAGATGGGAGATAAAAATCATGAGTTACGAAATCAAAATTTGCGACATCCTCAAAGGCGCTGG
GATCCTCTAGATTTAAGAAGGAGATATACATATG
>fliE
GAGGCAATTCCGACGTCTAAGAAACCATTATTATCATGACATTAACCTATAAAAATAGGCGTATCACGAGGCCCTTTCGTCTTCACCTC
GATGGATCCAATTTTCGGATTCGCACGCAGGCGATTAAGCCACTCAAGAGATTTTGTCTGGGCTGCAGTCGCATTCATCGCGCACCTCG
TGGCTGATTATGGACGGCGTTATTGGCAAAGTGGAACAAGACTCACTCCCGGGTCAGCAAACTCGAAAAATTGACGCTCCCATTATTTG
TGCTTTCGTTTTTTTCTATCCGCCAATAAACCCGTTTTTTTGTTGCTATTTAGCGCCTTTGTCTTATTGACTTACTGGTAGGCTTTGCT
ACCAGAAATTATCCGGGAGACGAGAATGTCAGCGATACAGGGGATTGAAGGGGTTATCAGCCAGTTACAGGCTACGGCGATGAGTGCGC
GTGCGCAGGAATCACTGCCGCAACCTCGAGAGATCCTCTAGATTTAAGAAGGAGATATACATATG
>yhdY
GAGGCAATTCCGACGTCTAAGAAACCATTATTATCATGACATTAACCTATAAAAATAGGCGTATCACGAGGCCCTTTCGTCTTCACCTC
GAGCGCTATTGGCTATCCCGATATGGTTTCGCTGTTTGCCGGCACCGTGCTGAATCAGACGGGGCAAGCCATCGAGACGATAGCCATGA
CCATGTCGGTCTATCTGATTATCAGCCTGACTATCTCGCTGCTGATGAATATCTATAACCGCCGCATCGCGATCGTTGAACGCTAAGGA
GTCATGATGACAAAAGTATTGCTGTCTCATCCCCCGCGCCCGGCGAGCCATAACTCAAGGATCCTCTAGATTTAAGAAGGAGATATACA
TATG
>fliF
GAGGCAATTCCGACGTCTAAGAAACCATTATTATCATGACATTAACCTATAAAAATAGGCGTATCACGAGGCCCTTTCGTCTTCACCTC
GAGGTTGCGGCAGTGATTCCTGCGCACGCGCACTCATCGCCGTAGCCTGTAACTGGCTGATAACCCCTTCAATCCCCTGTATCGCTGAC
ATTCTCGTCTCCCGGATAATTTCTGGTAGCAAAGCCTACCAGTAAGTCAATAAGACAAAGGCGCTAAATAGCAACAAAAAAACGGGTTT
ATTGGCGGATAGAAAAAAACGAAAGCACAAATAATGGGAGCGTCAATTTTTCGAGTTTGCTGACCCGGGAGTGAGTCTTGTTCCACTTT
GCCAATAACGCCGTCCATAATCAGCCACGAGGTGCGCGATGAATGCGACTGCAGCCCAGACAAAATCTCTTGAGTGGCTTAATCGCCTG
CGTGCGAATCCGAAAATTGGATCCTCTAGATTTAAGAAGGAGATATACATATG
>fliA
GAGGCAATTCCGACGTCTAAGAAACCATTATTATCATGACATTAACCTATAAAAATAGGCGTATCACGAGGCCCTTTCGTCTTCACCTC
GATGGATCCCCGGTAACTCCGTGTTGGCAAAAGCTAACCAGGTACCGCAGCAGGTTCTGTCTCTGCTGCAGGGTTAATCGTTGTAACCT
GATTAACTGAGACTGACGGCAACGCCAAATTGCCTGATGCGCTGCGCTTATCAGGCCTACAAGTTGAATTGCAATTTATTGAATTTGCA
CATTTTTGTAGGCCGGATAAGGCGTTTACGCCGCATCCGGCAACATAAAGCGCAATTTGTCAGCAACGTGCTTCCCCGCCACCGGCGGG
GTTTTTTTCTGCCTGGAATTTACCTGTAACCCCCAAATAACCCCTCATTTCACCCACTAATCGTCCGATTAAAAACCCTGCAGAAACGG
ATAATCATGCCGATAACTCATATAACGCAGGGCTGTTTATCGTGAATTCACTCTATACCGCTGAAGGTGTAATGGATAAACACTCGCTG
TGGCAGCGTTATGTCCCGCTGGTGCCTCGAGAGATCCTCTAGATTTAAGAAGGAGATATACATATG
>fliC
GAGGCAATTCCGACGTCTAAGAAACCATTATTATCATGACATTAACCTATAAAAATAGGCGTATCACGAGGCCCTTTCGTCTTCACCTC
GATGGATCCAGACCTGACCCGACTCCCAGCGATGAAATACTTGCCATGCGATTTCCTTTTATCTTTCGACACGTAAAACGAATACCGGG
GTTATCGGTCTGAATTGCGCAAAGTTTACGTTTAATTGTTTTTTTTAATAGCGGGAATAAGGGGCAGAGAAAAGAGTATTTCGGCGACT
AACAAAAAATGGCTGTTTTTGAAAAAAATTCTAAAGGTTGTTTTACGACAGACGATAACAGGGTTGACGGCGATTGAGCCGACGGGTGG
AAACCCAATACGTAATCAACGACTTGCAATATAGGATAACGAATCATGGCACAAGTCATTAATACCAACAGCCTCTCGCTGATCACTCA
AAATAATATCAACAAGAACCAGTCTGCGCTGTCGCTCGAGAGATCCTCTAGATTTAAGAAGGAGATATACATATG
>yaiA
GAGGCAATTCCGACGTCTAAGAAACCATTATTATCATGACATTAACCTATAAAAATAGGCGTATCACGAGGCCCTTTCGTCTTCACCTC
GAGGACGCAACAAACGAACCCAGCCAGGTGATTTCTGAAATTCGCAGCGCCCTGGCACAGACGATCAATTGTTGATTTTCGAGCGCCTA
TACTTAACGTTCATCCCGTGAAATAAGGAAGAACGATGCCAACGAAACCGCCTTATCCTCGTGAAGCATATATAGTGACGATTGAAAAA
GGAAAGCCAGGACAGACGGGATCCTCTAGATTTAAGAAGGAGATATACATATG
>rplL
GAGGCAATTCCGACGTCTAAGAAACCATTATTATCATGACATTAACCTATAAAAATAGGCGTATCACGAGGCCCTTTCGTCTTCACCTC
GAGAAGAAGCAATTGCACGCCTGATGGCAACCATGAAAGAAGCTTCGGCTGGCAAACTGGTTCGTACTCTGGCTGCTGTACGCGATGCG
AAAGAAGCTGCTTAATCGCAGTTATCTTTTTAACGCATTCGCTTACGTATAAACTTATTCTGATATTCAGGAACAATTTAAATGTCTAT
CACTAAAGATCAAATCATTGAAGCAGTTGCAGCTATGTCTGTAATGGACGTTGTAGAACTGATCTCTGCAATGGAAGAAAAATTCGGGG
ATCCTCTAGATTTAAGAAGGAGATATACATATG
>srmB
GAGGCAATTCCGACGTCTAAGAAACCATTATTATCATGACATTAACCTATAAAAATAGGCGTATCACGAGGCCCTTTCGTCTTCACCTC
GAGAGCACCGGGTGAGAACAATACCCGAGAAGCGATATCCATACAAACAGATGAAGATTGCAGCCGTAACGTCTATAATCAGCGCCCCA

CACAGAGGTAGAACATGACTGTAACGACTTTTTCCGAACTTGAACTCGACGAAAGCCTGCTGGAAGCCCTCCAGGATAAAGGTTTCACT
CGCCCGACCGCCATTCAGGCTGGGATCCTCTAGATTTAAGAAGGAGATATACATATG
>mglA
GAGGCAATTCCGACGTCTAAGAAACCATTATTATCATGACATTAACCTATAAAAATAGGCGTATCACGAGGCCCTTTCGTCTTCACCTC
GATGGATCCTGATGGCACCAACTGGAAAATCGACAACAAAGTGGTCCGCGTACCTTATGTTGGCGTAGATAAAGACAACCTGGCTGAAT
TCAGCAAGAAATAAGACTGATTGGGCGCAGTCTATTACTGCGCCCTTATTGTATGGCCAACAAGGTAAAATTATATGGTCAGCTCAACG
ACTCCGTCCTCCGGGGAATACTTGTTGGAAATGAGCGGTATCAACAAGTCCTTTCCTGGTGTTAAGGCACTCGAGAGATCCTCTAGATT
TAAGAAGGAGATATACATATG
>yohJ
GAGGCAATTCCGACGTCTAAGAAACCATTATTATCATGACATTAACCTATAAAAATAGGCGTATCACGAGGCCCTTTCGTCTTCACCTC
GAGTCGGTGATGCACAGATCGTAGTCGTTAACTTCGGTCAGCAATTCACGCACCAGAGAGTCAAGCACTCCCTCCATCGGTGCCAGTAA
CACACGCATATCATCACCCGCAAAAAAATGAGGCGCTATGTTAGCGCCTATGGTCAGCGGATTAAAGGTCTGCAAAGAGGGATTGCTGC
AAACTGCAATCCCTCTCTATGCTTGATGTATGCCTGAGTTCGCTGCGCATTGCGTAATGCCGGGGAGCGAAAAAATAAGCGCTGATTTC
TTAATGTGATCGGTAGCACGTTTTACGAATTAATTGTATGATGAATCCATCTCATCTGGGGTGTTGATTATGAGTAAGACACTGAACAT
TATCTGGCAATATTTACGCGCTTTCGTCCTGATTTATGCCTGCCTGTATGCAGGCATTTTCATTGCTTCCCTGCTACCGGGATCCTCTA
GATTTAAGAAGGAGATATACATATG
>yohL
GAGGCAATTCCGACGTCTAAGAAACCATTATTATCATGACATTAACCTATAAAAATAGGCGTATCACGAGGCCCTTTCGTCTTCACCTC
GATGGATCCAGGCGTTTCCTTGCTGAAGAAGAGTTGTAAATTCGGTCATGATAATAATTCTTAGTATTAATTCGGCAATCTGATTCTAC
TCCCCCCCAGTACCTGATACTACCCCCCAGTAGATTAATAGTGCTATGATTTTTCATATTCTTGTTAACCAGGTGTTGCCATGTCTCAT
ACAATCCGTGATAAACAGAAACTGAAAGCGCGTGCCAGTAAGATTCAGGGCCAGGTCGTGGCGCTCAAGAAAATGCTCTCGAGAGATCC
TCTAGATTTAAGAAGGAGATATACATATG
>yohM
GAGGCAATTCCGACGTCTAAGAAACCATTATTATCATGACATTAACCTATAAAAATAGGCGTATCACGAGGCCCTTTCGTCTTCACCTC
GAGAGCATTTTCTTGAGCGCCACGACCTGGCCCTGAATCTTACTGGCACGCGCTTTCAGTTTCTGTTTATCACGGATTGTATGAGACAT
GGCAACACCTGGTTAACAAGAATATGAAAAATCATAGCACTATTAATCTACTGGGGGGTAGTATCAGGTACTGGGGGGGAGTAGAATCA
GATTGCCGAATTAATACTAAGAATTATTATCATGACCGAATTTACAACTCTTCTTCAGCAAGGAAACGCCTGGTTCTTCATCCCCAGCG
CCATCTTACTTGGTGCGCTTCATGGGGATCCTCTAGATTTAAGAAGGAGATATACATATG
>pdxY
GAGGCAATTCCGACGTCTAAGAAACCATTATTATCATGACATTAACCTATAAAAATAGGCGTATCACGAGGCCCTTTCGTCTTCACCTC
GATGGATCCGCGTCGCGGTAAAAAGAATTACTGTCTGATTTGCTGGAAATAATGCATTAAGTGGAAAGGGGGAGTGAGAAATCACTCCC
CCTGGTTTTTATACAGGGAACATGATGAAAAATATTCTCGCTATCCAGTCTCACGTTGTTTATGGTCATGCGGGTAACAGTGCGGCAGA
GTTTCCCTCGAGAGATCCTCTAGATTTAAGAAGGAGATATACATATG
>yohC
GAGGCAATTCCGACGTCTAAGAAACCATTATTATCATGACATTAACCTATAAAAATAGGCGTATCACGAGGCCCTTTCGTCTTCACCTC
GATGGATCCTCTCCCTTTGGTTGCCAGTAAGCATACACCCTGTGAATCATTCACGCTGAAAAAGGACGTTCATCAGACAGTGAAAATTA
CATGGGCGAACAAAAAATAATCACAAGCTGACTTATACTTGAACTGTCAGGTTCATGAACTCACAACAAGGGGGCGGTATGAGCCATGT
CTGGGGACTGTTTTCCCATCCCGATCGTGAAATGCAGGTGATTAATCGCGAAAACGAAACGCTCGAGAGATCCTCTAGATTTAAGAAGG
AGATATACATATG
>yohD
GAGGCAATTCCGACGTCTAAGAAACCATTATTATCATGACATTAACCTATAAAAATAGGCGTATCACGAGGCCCTTTCGTCTTCACCTC
GAGCGTTTCGTTTTCGCGATTAATCACCTGCATTTCACGATCGGGATGGGAAAACAGTCCCCAGACATGGCTCATACCGCCCCCTTGTT
GTGAGTTCATGAACCTGACAGTTCAAGTATAAGTCAGCTTGTGATTATTTTTTGTTCGCCCATGTAATTTTCACTGTCTGATGAACGTC
CTTTTTCAGCGTGAATGATTCACAGGGTGTATGCTTACTGGCAACCAAAGGGAGACAGACTGGCCTATGGATCTCAATACACTTATCTC
ACAATATGGTTATGCCGCGCGGATCCTCTAGATTTAAGAAGGAGATATACATATG
>yaiB
GAGGCAATTCCGACGTCTAAGAAACCATTATTATCATGACATTAACCTATAAAAATAGGCGTATCACGAGGCCCTTTCGTCTTCACCTC
GAGTTGCCGATTGCAGAGACACTTCATGTTCCGCTGATTTACCACCAAAAACGATTCCTACCCGCAGTTTTTCCATCTTAAAAACCTAT
CCCGTCTAACACAAAGTGCATACATTACCACGACAAAACGGGGGATTCGCGGCCTTCTGAAAGATTGTTGCAATCTTCTGCTGACAAAG
CGTGCAACGTACTGGTGAAGAAAGTGCGTTATCTCAAAGATGTGCGCAAGATCACAAAAAATGATGAACGGGAAGCTAATTTATTCCTGG
CTTAAATGGCCATGCGGTGAGTTTTTTTCTCTTAATTATAAGTTAACGAAGAGAATATATTTCATAACTTTTATTTATAATAAAGGTTG
ATAATTAAAAGCCTATATTTTGTGTGGGTAATTATTTAAATAAGAGAAACGTTTCGCTGGTAATCAAACAAAAAATATTTGCGCAAAGT
ATTTCCTTTGTCATAAAAATAATACTTCCAGACACTATGAAGTTGTGAAACATAATGTTAACTTCTCCATACTTTGGATAAGGAAATAC
AGACATGAAAAATCTCATTGCTGAGTTGTTATTTAAGCTTGCCCAAAAAGAAGAAGAGTCGAAAGAACTGTGTGCGCGGATCCTCTAGA
TTTAAGAAGGAGATATACATATG
>yeaE
GAGGCAATTCCGACGTCTAAGAAACCATTATTATCATGACATTAACCTATAAAAATAGGCGTATCACGAGGCCCTTTCGTCTTCACCTC
GATGGATCCACAGCCCGATGGTGGATAAATCCTGGACTGGCCTGATTTCTACCGGGATCACCTACAAATTCTGATAATGCATCTTTCCA
GGGCATTTTGTGCATAAAAACAGGGCGTCACGCCCTGTTTTGCATTACGGTGGTGCAATACCTGGGAGGAAACAATGCAACAAAAAATG
ATTCAATTTAGTGGCGATGTCTCACTGCCAGCCGTAGGGCAGGGAACATGGTATATGGGCGAAGATGCCAGTCAGCTCGAGAGATCCTC
TAGATTTAAGAAGGAGATATACATATG
>yqaC

```
GAGGCAATTCCGACGTCTAAGAAACCATTATTATCATGACATTAACCTATAAAAATAGGCGTATCACGAGGCCCTTTCGTCTTCACCTC
GAGGGGAATTGGGGGTAGTGGTAAAAATGACCTGGATTATCCCTCTGCTGATATACGAGCGAAGATCAGTAAAGATATAGAAGGTGGTA
TCCCTGGCTATTAGCAAGGTCAGGTTTTGATTCCATTCATTAAAGATCCAGTAACAAAAGAACGTAAACAAATCCATCCAGAAGATATA
CATCTCACTGCAAAAGACTTCGAAGCAAGTAAAGATAACATCTCTAAGGATGAATGGGAAAACCTCCATGCATTAAAAGAAAAGCGTTT
AAATGGAATGCCTAAAACAACACCCAAAAGTGACCAGGTTATTATGTTGCAAAATCAATACGTTCGTGAAATGCGAAAGGATCCTCTAG
ATTTAAGAAGGAGATATACATATG
>yqaB
GAGGCAATTCCGACGTCTAAGAAACCATTATTATCATGACATTAACCTATAAAAATAGGCGTATCACGAGGCCCTTTCGTCTTCACCTC
GATGGATCCCGATGCATCCGTAGCTCAGCTGGATAGAGTACTCGGCTACGAACCGAGCGGTCGGAGGTTCGAATCCTCCCGGATGCACC
ATATTCTCCGTAACCTTCAGCAATGAAGGTATCGAAGAGATAGCGGTATTAACCGCAGGCACCAGGAAGGATAACGTTGCGCCAGCAAC
GGCCCGAAGGGCGAACGTCAGTGAGTCATCCTCCCGGATGCACCATCTTCTCCAAAAAATTTCCTTCAAAAATATCTCATTTCTAAATA
ACTCCTTTATCCATTTCCTGAATTTTCTTCTCCTTCGCCATCAGCGACAAAATCAAGCAATTACGCTAAAGTAGCGTCATATTTTTCGA
CTTGCGAGAAGACCATGTACGAGCGTTATGCAGGTTTAATTTTGATATGGATGGCACAATCCTGGATACGGAGCCTACGCACCGTAAA
GCGCTCGAGAGATCCTCTAGATTTAAGAAGGAGATATACATATG
>yqaE
GAGGCAATTCCGACGTCTAAGAAACCATTATTATCATGACATTAACCTATAAAAATAGGCGTATCACGAGGCCCTTTCGTCTTCACCTC
GATGGATCCTGCTTTCAATAAGGCCGCTGCCTGTTCGGCACTGGCCTGTAATTGCGCGAGTTCAGTCATATTTATTTAAGTATTTTCTA
AATTAAGTAAACTCTAAACTAAAAATGCAACATATACCAGCCTCAGCAGCGTAAATGAGAGTAAAAGCGTAAGCTGAAACTGGCAGGCT
CCGCTAAAATTACTACGCTTAAGAGATAAAATCTCTTTTTAAACAATGAGTAATTTTCTTATAGGGAGTACATATGGGTTTCTGGAGAA
TCGTCATCACCATCATTCTGCCGCCGCTCGGCGTGCTGCTCGGTAAAGGGTTCGCTCGAGAGATCCTCTAGATTTAAGAAGGAGATATA
CATATG
>nrdB
GAGGCAATTCCGACGTCTAAGAAACCATTATTATCATGACATTAACCTATAAAAATAGGCGTATCACGAGGCCCTTTCGTCTTCACCTC
GAGCGCTGAAGACGCACAAGACGATCTGGTGCCGTCAATCCAGGACGATGGCTGCGAAAGCGGCGCATGTAAGATCTGATATTGAGATG
CCGGATGCGGCGTAAACGCCTTATCCGGCCTACGGCTCGGTTTGTAGGCCTGATAAGACGCGCCAGCGTCGCATCAGGCTCCGGGTGCC
GGATGCAGCGTGAACGCCTTATCCGGCCTACGGCTCGGATTTGTAGGCCTGATAAGACGCGCCAGCGTCGCATCAGGCACAGGATGCGG
CGTAAAATGCCTTATCCGGCATTAAACTCCCAACAGGACACACTCATGGCATATACCACCTTTTCACAGACGAAAAATGATCAGCTCAA
AGAACCGATGTTCTTTGGTCAGCCGGTCGGGGATCCTCTAGATTTAAGAAGGAGATATACATATG
>nrdD
GAGGCAATTCCGACGTCTAAGAAACCATTATTATCATGACATTAACCTATAAAAATAGGCGTATCACGAGGCCCTTTCGTCTTCACCTC
GATGGATCCCACAACCCTGTGCCATGAATTTACGGCCTTTTGAGGCTGTCTGGTGGTTACAGAAGTAAATCTTCCCTAAGCCCCGGTAA
TGCCGGTCATTCCGGGGTTTTGTCTGTCAGTATGTTGTTTTTGTTGATTTTTCAACCAGCAAATTCATTAAAAAATTTACATATCGCTG
TAGCGCCCGTCATCCGTACGCTCTGCTTTTTACTTTGAGCTACATCAAAAAAAGCTCAAACATCCTTGATGCAAAGCACTATATATAGA
CTTTAAAATGCGTCCCAACCCAATATGTTGTATTAATCGACTATAATTGCTACTACAGCTCCCCACGAAAAAGGTGCGGCGTTGTGGAT
AAGCGGATGGCGATTGCGGAAAGCACCGGAAAACGAAACGAAAAAACCGGAAAACGCCTTTCCCAATTTCTGTGGATAACCTGTTCTTA
AAAATATGGAGCGATCATGACACCGCATGTGATGAAACGAGACGGCTGCAAAGTGCCGTTTAAATCAGAGCGCATCAAAGAAGCGATTC
TGCGTGCAGCTACTCGAGAGATCCTCTAGATTTAAGAAGGAGATATACATATG
>rhsE
GAGGCAATTCCGACGTCTAAGAAACCATTATTATCATGACATTAACCTATAAAAATAGGCGTATCACGAGGCCCTTTCGTCTTCACCTC
GAGTGATGGATATGCTCGGAGTTGGAGATGCAATTAATATTAATACAAATAAAAATATCCGGCAGGTATGCTAAAAGGATAAATGGCGT
TCTCTGTATTTATTCGCAATTATCTGTAGGACAAATATGGAGAGCTTGATGAGTCAGAGCTAAAACAGCAATTGATTAATAAACATGTT
GATTTATTCAAGAAAATGGATGTATTTTAAATGTTGATTCTTAATATAACCTAATCAATAGATTATCAAGTGAGAGCTACACATTATTG
GAGGGGCAGGGGAAAACACCATTTTGGGAACAATGCATCAGGTACTCTTACCACTGAAGCAAGATCATCCGGCACCAGGTGGTGGTGCA
GTGCTTTGGAAATGGTCCCCTCGGGGGAGTTGTCATTATTTTCTATGCTATTTGCTTATTATATTCATCAGGTGAATTTAATAAATTCC
TAATTTGAAATATCCACTATTAAGCTAGTGTTTAACGACGTTAAGATGGCAATGTGCAGATCATGTTATCAGGGTGGGGCGGTAGCGTA
TATTCCTTATACAAACCATCATTCATTTCAATGATTGTTATTGAATTATTTATTGTTGTGAATAAAAAGACATTATGAATATTGGTTTG
ACTTTGCATATCAGGTTTTTTATTACTCGCTGTAATGTACGAGCCTGTCGTGAATAACGTCGAGAACCCATCCGTTGCGTTTGTGATTA
TTTTGTTGACTAAACAGACACCCGTTTCTCTGAAGTAAAATCCCAGACTAAATCATCACATAACCATGACATTTTTCTGATATTCCCCG
GTAACGCCAGATGTCGACTCGCTTAACCACCCTCAGCCATACCAGTGAAGGCCACCGCGTGTCGGTCCACTACGGCGGATCCTCTAGAT
TTAAGAAGGAGATATACATATG
>rhsD
GAGGCAATTCCGACGTCTAAGAAACCATTATTATCATGACATTAACCTATAAAAATAGGCGTATCACGAGGCCCTTTCGTCTTCACCTC
GAGGCTGCTGCTGTCGCTTTTCGGCGGCTGGCTGGGTGCGCGACTGGTTAAGGGTAAGGCGCTGTTCAGGCAGTTTGCGGGGTGATGAG
AGTAAATAAGGGGCGTTCTGGTTGAATGGAACGCCTTGTTTAAATTAATTCTACAATATGTTTCATTGATGATGTTATTGAATTGGTGA
TTTCCTATCTTTCTATTGCTGATATTAATCTGAATCGGTGGAGTTTATTGATTTTAAATATAGCCCTACAAAAATAACTCAGACTAATA
AAATCATAAATCATATGCGTTGAATGGATATTATCCATATAGTGAATTTGTTGATGATGAATTCATCTGTGCTAAAAATGTTAGTTTAA
TAAAATATTGAAAGTGACCTGTAATAACAGTTGTTGTTGATTGAGAACAAATAAGTTTATGTGAAAAATATATAAATACATTAGCTGGT
CTTGTGTGTCATTTTATTTTTTTTTGTTGCTAACACAGGGATATGAACAATAACTAAAAGGGCACTTTATATGAGCGGAAAACCAGCGG
CGCGTCAGGGAGATATGACTCAGTATGGCGGTCCCATGGATCCTCTAGATTTAAGAAGGAGATATACATATG
>rhsB
GAGGCAATTCCGACGTCTAAGAAACCATTATTATCATGACATTAACCTATAAAAATAGGCGTATCACGAGGCCCTTTCGTCTTCACCTC
GAGGACATGGGTGACGTGCAGCATTTTGCCGATGACGTTATCGCCCAGCGCGGCGTGCGGCACGGGCATTTGCAGTGCTTGCCGAAGGA
AGATTGAGTCTATTATTGGGTCCTTTGGTGAACGATCAACAAAGGGCCACTAGCACACCTGATTCCTCTCTAAATACCCTATCCGAACC
```

TTCTCTTTTGTAACGTTCTAAATATATTCCTAAAAATCTTCAATTCATTGTGACCACAAGTTTTTCTTCGCTTTTTCGTATGAAGATAC
TGTCATTAAAATAATAGAAAAGGATTTTACGATGAGCGGAAAACCGGCGGCGCGTCAGGGTGACATGACGCAGTATGGCGGTAGCATTG
TTCAGGGTTCAGCCGGGGTACGTATTGGGATCCTCTAGATTTAAGAAGGAGATATACATATG
>rhsA
GAGGCAATTCCGACGTCTAAGAAACCATTATTATCATGACATTAACCTATAAAAATAGGCGTATCACGAGGCCCTTTCGTCTTCACGTC
GACCGCCAGTGCCTCAATTTTGCGCACCCGCAACGACTCCAGCGGATCGCGCGGCAACATCGCCGGAGCGACATTCATTAATTCAATAT
ATTCAGCGATGATCGGCGAATCAAACCAGCATTCGCCCTCTTCGGTCACCAGCACCGGCACTTTTCCTAACGGGTTAAATTGCGCCACG
CCGTTGTCCGCGTTATAGGGCAGTTCATTAATAAATTCGAAAGTTATGCCCTTTTCTAACAACAGAATAGAAAGTTTGCGTACAAACGG
GCTGGTGTAGCTACCGACGAGTTTCATGCCGAGTCCTTTGTGCGAGGAAAAATATCAGTATGGCCTGGTGATGGGAAAAAGGGCGGATC
CTCTAGATTTAAGAAGGAGATATACATATG
>pdxH
GAGGCAATTCCGACGTCTAAGAAACCATTATTATCATGACATTAACCTATAAAAATAGGCGTATCACGAGGCCCTTTCGTCTTCACCTC
GATGGATCCTTTTCTGGTCGAAAGGCGATGAAGCGACTGTCTATAAACGCGACCGCATCGTCTTGAATAACTGTCAGTTACAAAATCCA
CAGCGTTGAGATTTTTCCAGGGGCGGCGCACAATAGCGTCACCCACTGACAATCCGTAAAGAAAACCATGTCTGATAACGACGAATTGC
AGCAAATCGCGCATCTGCGCCGTGAATACACCAAAGGCGGGTTACGCCGCCGCGATCTTCCCGCCGATCCATTAACCCTCTCGAGAGAT
CCTCTAGATTTAAGAAGGAGATATACATATG
>pdxK
GAGGCAATTCCGACGTCTAAGAAACCATTATTATCATGACATTAACCTATAAAAATAGGCGTATCACGAGGCCCTTTCGTCTTCACCTC
GATGGATCCTGATGCGGGCTTCTTCATGGGCAGTTAACTTCGCGTAAACGGGGAGTGTCATTAGTAGTGTAATGACCAGACAAATGATT
TTCTTCATTGAATTACCTTATCCTTGTTTTGTGATAGCCGATTTGACTGCCAGAATCAGGTAAAGTTTCGGTTGGATTACGATGAGGAT
CATCGGCAAGGAGAAAAAATTAAATGAGTAGTTTGTTGTTGTTTAACGATAAGAGTAGGGCACTGCAGGCGGATATCGTCGCCGTGCAG
TCGCAGGTGGTTTACGGCTCGAGAGATCCTCTAGATTTAAGAAGGAGATATACATATG
>pdxB
GAGGCAATTCCGACGTCTAAGAAACCATTATTATCATGACATTAACCTATAAAAATAGGCGTATCACGAGGCCCTTTCGTCTTCACCTC
GATGGATCCTCCCTGACCTGGTGGTTGCCCAGGAGGAGGGCCGGAAATAGGTTGTATCATTACGTATCCTTATACCTGAAATCTTCGCA
AGTATGCCTGGCCGCGAGATTATGGCACACTTGTCCGGTTAACTCTCGTCTCATACAGGTAACACAAACGTGAAAATCCTTGTTGATGA
AAATATGCCTTATGCCCGCGACTTATTTAGCCGTTTGGGTGAGGTGACCGCTCGAGAGATCCTCTAGATTTAAGAAGGAGATATACATA
TG
>nrdG
GAGGCAATTCCGACGTCTAAGAAACCATTATTATCATGACATTAACCTATAAAAATAGGCGTATCACGAGGCCCTTTCGTCTTCACCTC
GATGGATCCGCCGCGTTAAACATTTGGGGAATGGGCAGATAGGTTAATCTTCACTCTGAGCTAAGCAAATGCCGGATGCGGCGTAAACG
CCTTATCCGGCCTACATTTGGCAATTGCTCCAGGTCTGATAAGACGCGACAGCGTCGCATCAGGCATTGATTACCGGATGCAGCCTACA
CGCGGACTGATAAGCCAGGCGAAATAATGAATTATCATCAGTACTATCCTGTCGACATCGTCAACGGCCCCGGCACTCGTTGCACCCTG
TTTGTCTCCGGCTCGAGAGATCCTCTAGATTTAAGAAGGAGATATACATATG
>nrdH
GAGGCAATTCCGACGTCTAAGAAACCATTATTATCATGACATTAACCTATAAAAATAGGCGTATCACGAGGCCCTTTCGTCTTCACCTC
GAGGACCCTGGTGTAGCGTGGGTACAGCAGCTGCGGTAGGTATTTTTATCGGCGCACTGTTAAGCATGCGCAAATCGTAGTGCAAAAAT
GATAATAAATACGCGTCTTTGACCCCGAAGCCTGTCTTCGGGGTTTCTTTTTGCCTGGTGAATCACAAAAATCCCCCTACCCCGTCACG
CTCATATCCAGGGTAATTTCGACCACTATTTGCTATATATTGTGTGGTTGAATCTTTTTTCAACTACATCTAGTATCTCTGTATCAACA
GAGAGACAACCCGACGCGTATCATCGCGCCGTATCTTCATTTTAAACGGAAATACGAATCATGCGCATTACTATTTACACTCGTAACGA
TTGCGTTCAGTGCCACGCCACCAAACGGGCGATGGAAAACCGGGGCTTTGATTTTGAAGGATCCCGGGATCCTCTAGATTTAAGAAGGA
GATATACATATG
>yghU
GAGGCAATTCCGACGTCTAAGAAACCATTATTATCATGACATTAACCTATAAAAATAGGCGTATCACGAGGCCCTTTCGTCTTCACCTC
GAGCCCACCTGGGGCGTAGCCCAATAATGTCCCGAACGGGGCATCCTGGCTGGTCGTTCCTTTGCTCATCACTTTACCTTTAACAATAC
AACCTAAGCAGTTGGCGTAATTGTTGTGCTCTGATTACCTGCTCAGCGACATTAACCGGACAGAGGTCAGACTAATCATACACCTCATC
GCAGCGGCTGGCGCAAAGGGTTAAAAAATTCACATTCTGTACAGCAAGTGACCTGCTACACTGCTTCAACACTACCACTCAGAAGGCAA
CTCACTATGACAGACAATACTTATCAGCCCGCGAAAGTCGGATCCTCTAGATTTAAGAAGGAGATATACATATG
>mdoB
GAGGCAATTCCGACGTCTAAGAAACCATTATTATCATGACATTAACCTATAAAAATAGGCGTATCACGAGGCCCTTTCGTCTTCACCTC
GATGGATCCACCAAAGCCTGCGATCTGGTGTTAAAACAGGGGCTGAACTTCATTTCCTGAACCGCGAAACGCCGCGTTTTAGCTGTTAC
CCGGCTGACGCGGCTCTCTCCTGCCCTTTTCCCTGCTCCTGGACGGTTTTACCCCTGTTACACGGGAAGTGACCGTAATCAAAAAAGTC
TGTTTATAAACTAATTCCTAACAACATCGCATTTTTTTGACACTAGAATTGCAGCAATATGGCGACGCCATTACATTGTGCTTCTCAAA
AAGCTTAAACGTTGGCTAAACGGCCCGGTTTAGTGAGGATCGCCCGTTGTCAGAACTACTCTCTTTCGCCCTTTTTCTCGCCTCTGTGC
TGATTTACGCATGGAAAGCGGGACGTAACACCTGGTGGTTTGCAGCCACGTTAACGGTGCTGGGGCTATTCTCGAGAGATCCTCTAGAT
TTAAGAAGGAGATATACATATG
>negCtrl
TTAAACTAGCGTTGAGTTAGTTAACTTTGCAATTTGTTCTAGTGTTTTTATGGTTGGATCTGATTTTCCTGATTCTATTCGTGAATAAT
TTGATCTACTCATTTCTAATTCTTGGGAGAATTTTTCTTTTGATTTGGTACCTCTCATCTTTTTTAAGCTTTCTCTAAATGCTAGTTCC
AAGTTTCTCCCCCCTCTTTTTGTGTAATAAAATTCACGCTTTATTTAAATTATAGCTTTTAATATCTTAATAAACAAGTGATTAATCAC
TTGTGATTTGTGATTAATCACTTGTGATTTGTGATTATTCCAAAAGTAGGTAATTTCACCAACCTTTTTCAAAAGTTTTTATCGACGAT
TATTGGACTGGATCGTGTTTCCTAAAGCGATTCTAAGCGCTAGGGACCTCTTTAGCTCGAAC

Data Set S5 - Nissle 1917 library reference file.fasta

```
>cydAWTcI
GAGGAGTTACCAGTGGCTCATGATTCACACATAATTAACATATGTGGTTGATTTTGATATAAATCAATTTTTAATTCTTGAAAAAAATC
TGCAAAACACGATAAATTGTTGTAGATCAAATTGGTGAGATCGTGACAATTTCGCTATAAATTGATCACTGTCGAAAAATGCAAATTTG
CTCAATAAAAACCTGTTTATTGTAAGGATTTTGCGGCGTAATATAATTACGGGATCAATTTGGTTTATTCATTAACACATTTGTAACTT
TTTTATCATTGTCTTTTCCTCTTTTGATGAAATGGCAACGATAGAGGCGAGCAAATAATTTTGTATTGGGGCATGCGTGTGACCCTTTC
TAACGGGGTTCACTCTCGGAGTCTTCATGCGATGAGCAAGGAGTCATTATG
>cydA5cI
GAGGGTTCATACGACCTCCTAATCCATCCAGCCTCCGAATCACGCCTCGTCTCAACCAAAGCAATCAACCCATCAACCACCTGGCATTA
CTCGCATCCATTCTCAGGCTGTCTCGTCTCGTCTCGCACTGAAGGTCCTCAATCGCACTGGAAACATCAAGGTCGGTTCCAGCAGTCAT
AGAGTTACCAGTGGCTCATGATTCACACATAATTAACATATGTGGTTGATTTTGATATAAATCAATTTTTAATTCTTGAAAAAAATCTG
CAAAACACGATAAATTGTTGTAGATCAAATTGGTGAGATCGTGACAATTTCGCTATAAATTGATCACTGTCGAAAAATGCAAATTTGCT
CAATAAAAACCTGTTTATTGTAAGGATTTTGCGGCGTAATATAATTACTCTTAATCATGCAGGGGAGGGTTTCTTATG
>cydA10cI
GAGGGTTCATACGACCTCCTAATCCATCCAGCCTCCGAATCACGCCTCGTCTCAACCAAAGCAATCAACCCATCAACCACCTGGCATTA
CTCGCATCCATTCTCAGGCTGTCTCGTCTCGTCTCGCACTGAAGGTCCTCAATCGCACTGGAAACATCAAGGTCGGTTCCAGCAGTCAT
AGAGTTACCAGTGGCTCATGATTCACACATAATTAACATATGTGGTTGATTTTGATATAAATCAATTTTTAATTCTTGAAAAAAATCTG
CAAAACACGATAAATTGTTGTAGATCAAATTGGTGAGATCGTGACAATTTCGCTATAAATTGATCACTGTCGAAAAATGCAAATTTGCT
CAATAAAAACCTGTTTATTGTAAGGATTTTGCGGCGTAATATAATTACTCTTAATCATGCGGAGGATCGTTTCTTATG
>cydA15cI
GAGGGTTCATACGACCTCCTAATCCATCCAGCCTCCGAATCACGCCTCGTCTCAACCAAAGCAATCAACCCATCAACCACCTGGCATTA
CTCGCATCCATTCTCAGGCTGTCTCGTCTCGTCTCGCACTGAAGGTCCTCAATCGCACTGGAAACATCAAGGTCGGTTCCAGCAGTCAT
AGAGTTACCAGTGGCTCATGATTCACACATAATTAACATATGTGGTTGATTTTGATATAAATCAATTTTTAATTCTTGAAAAAAATCTG
CAAAACACGATAAATTGTTGTAGATCAAATTGGTGAGATCGTGACAATTTCGCTATAAATTGATCACTGTCGAAAAATGCAAATTTGCT
CAATAAAAACCTGTTTATTGTAAGGATTTTGCGGCGTAATATAATTACTCTTAATCATGCGGGGGAGTCTTTCTTATG
>cydA17cI
GAGGGTTCATACGACCTCCTAATCCATCCAGCCTCCGAATCACGCCTCGTCTCAACCAAAGCAATCAACCCATCAACCACCTGGCATTA
CTCGCATCCATTCTCAGGCTGTCTCGTCTCGTCTCGCACTGAAGGTCCTCAATCGCACTGGAAACATCAAGGTCGGTTCCAGCAGTCAT
AGAGTTACCAGTGGCTCATGATTCACACATAATTAACATATGTGGTTGATTTTGATATAAATCAATTTTTAATTCTTGAAAAAAATCTG
CAAAACACGATAAATTGTTGTAGATCAAATTGGTGAGATCGTGACAATTTCGCTATAAATTGATCACTGTCGAAAAATGCAAATTTGCT
CAATAAAAACCTGTTTATTGTAAGGATTTTGCGGCGTAATATAATTACTCTTAATCATGCGGAGGAGGGTTTCTTATG
>cydA23cI
GAGGGTTCATACGACCTCCTAATCCATCCAGCCTCCGAATCACGCCTCGTCTCAACCAAAGCAATCAACCCATCAACCACCTGGCATTA
CTCGCATCCATTCTCAGGCTGTCTCGTCTCGTCTCGCACTGAAGGTCCTCAATCGCACTGGAAACATCAAGGTCGGTTCCAGCAGTCAT
AGAGTTACCAGTGGCTCATGATTCACACATAATTAACATATGTGGTTGATTTTGATATAAATCAATTTTTAATTCTTGAAAAAAATCTG
CAAAACACGATAAATTGTTGTAGATCAAATTGGTGAGATCGTGACAATTTCGCTATAAATTGATCACTGTCGAAAAATGCAAATTTGCT
CAATAAAAACCTGTTTATTGTAAGGATTTTGCGGCGTAATATAATTACTCTTAATCATGCGAAGGACAGTTTCTTATG
>fdnGWTcI
GAGGCATTACTCGCATCCATTCTCAGGCTGTCTCGTCTCGTCTCGCTGGGAGTTCGTAGACGGAAACAAACGCAGAATCCAAGCGCACT
GAAGGTCCTCAATCGCACTGGAAACATCAAGGTCGGACACACTGTATCTGACAACGACGCACTCTGGTAACCCACCTCGTTCGGCTCGC
TGTTTTGTCTCTCTTGCCGTTAAAAATTAAGCTGAATTTTATAGCATTTTTTTAACTGGCCTGTCAGGCAGTGGTGCGTTTTTCTACC
GCTATTGAGGTAGGTCAATTTGCGCAGGCGGATTATTTTGTGGCAAACACGTGTTCTTTTTGATTTCGCGCAAAAAGATTCAGAATTTT
ACTGTTAGTTTCCTCGCGCAGTAATACCCCTGAAAAAAGAGGAAAGCTATG
>fdnG5cI
GAGGCATTACTCGCATCCATTCTCAGGCTGTCTCGTCTCGTCTCCATCATCTGTAACCACTACTAACGCTCATCTCTGTCGCAGCCAGG
ATACATAGATTACCACAACTCCGAGCCCTTCCACCGTCCGAACTACCGTATCACATCCACCAAGCGTTACACAGCGACACACTGTATCT
GACAACGACGCACTCTGGTAACCGCTCGCTGTTTTGTCTCTCTTGCCGTTAAAAATTAAGCTGAATTTTATAGCATTTTTTTAACTGG
CCTGTCAGGCAGTGGTGCGTTTTTCTACCGCTATTGAGGTAGGTCAATTTGCGCAGGCGGATTATTTTGTGGCAAACACGTGTTCTTTT
TGATTTCGCGCAAAAAGATTCAGAATTTTACTGTTAGTTTCCTCGCGCTCTTAATCATGCAGGGGAGGGTTTCTTATG
>fdnG10cI
GAGGCATTACTCGCATCCATTCTCAGGCTGTCTCGTCTCGTCTCCATCATCTGTAACCACTACTAACGCTCATCTCTGTCGCAGCCAGG
ATACATAGATTACCACAACTCCGAGCCCTTCCACCGTCCGAACTACCGTATCACATCCACCAAGCGTTACACAGCGACACACTGTATCT
GACAACGACGCACTCTGGTAACCGCTCGCTGTTTTGTCTCTCTTGCCGTTAAAAATTAAGCTGAATTTTATAGCATTTTTTTAACTGG
CCTGTCAGGCAGTGGTGCGTTTTTCTACCGCTATTGAGGTAGGTCAATTTGCGCAGGCGGATTATTTTGTGGCAAACACGTGTTCTTTT
TGATTTCGCGCAAAAAGATTCAGAATTTTACTGTTAGTTTCCTCGCGCTCTTAATCATGCGGAGGATCGTTTCTTATG
>fdnG15cI
GAGGCATTACTCGCATCCATTCTCAGGCTGTCTCGTCTCGTCTCCATCATCTGTAACCACTACTAACGCTCATCTCTGTCGCAGCCAGG
ATACATAGATTACCACAACTCCGAGCCCTTCCACCGTCCGAACTACCGTATCACATCCACCAAGCGTTACACAGCGACACACTGTATCT
GACAACGACGCACTCTGGTAACCGCTCGCTGTTTTGTCTCTCTTGCCGTTAAAAATTAAGCTGAATTTTATAGCATTTTTTTAACTGG
CCTGTCAGGCAGTGGTGCGTTTTTCTACCGCTATTGAGGTAGGTCAATTTGCGCAGGCGGATTATTTTGTGGCAAACACGTGTTCTTTT
TGATTTCGCGCAAAAAGATTCAGAATTTTACTGTTAGTTTCCTCGCGCTCTTAATCATGCGGGGGAGTCTTTCTTATG
>fdnG17cI
GAGGCATTACTCGCATCCATTCTCAGGCTGTCTCGTCTCGTCTCCATCATCTGTAACCACTACTAACGCTCATCTCTGTCGCAGCCAGG
ATACATAGATTACCACAACTCCGAGCCCTTCCACCGTCCGAACTACCGTATCACATCCACCAAGCGTTACACAGCGACACACTGTATCT
```

GACAACGACGCACTCTGGTAACCGCTCGCTGTTTTGTCTCTCTTGCCGTTAAAAATTAAGCTGAATTTTATAGCATTTTTTTAACTGG
CCTGTCAGGCAGTGGTGCGTTTTTCTACCGCTATTGAGGTAGGTCAATTTGCGCAGGCGGATTATTTTGTGGCAAACACGTGTTCTTTT
TGATTTCGCGCAAAAAGATTCAGAATTTTACTGTTAGTTTCCTCGCGCTCTTAATCATGCGGAGGAGGGTTTCTTATG
>fdnG23cI
GAGGCATTACTCGCATCCATTCTCAGGCTGTCTCGTCTCGTCTCCATCATCTGTAACCACTACTAACGCTCATCTCTGTCGCAGCCAGG
ATACATAGATTACCACAACTCCGAGCCCTTCCACCGTCCGAACTACCGTATCACATCCACCAAGCGTTACACAGCGACACACTGTATCT
GACAACGACGCACTCTGGTAACCGCTCGCTGTTTTGTCTCTCTTGCCGTTAAAAATTAAGCTGAATTTTATAGCATTTTTTTAACTGG
CCTGTCAGGCAGTGGTGCGTTTTTCTACCGCTATTGAGGTAGGTCAATTTGCGCAGGCGGATTATTTTGTGGCAAACACGTGTTCTTTT
TGATTTCGCGCAAAAAGATTCAGAATTTTACTGTTAGTTTCCTCGCGCTCTTAATCATGCGAAGGACAGTTTCTTATG
>hycAWTcI
GAGGCCTCGTCTCAACCAAAGCAATCAACCCATCAACCACCTGGCATTCGCACTCTTTCGCCAAGCCAACCTCTTAGACCCTACGTTCC
ACTTAGACACCGTATCTTTCCCTCCAATCCAGAGCCTTACTGGTCTGTCTGGTCAAATCTACGGGTTCCCGCAGTGCCCGTTGGCAGAG
GGTTATTTCGTGCATATCGCCTCCCATTAACTATTGCCAGCTACAAGCAATAATTGTGCCAGTGTTGATTATCCCTGCGGTGAATAATG
TCGATGATGTCGAAATGACACGTCGACACGGCGACGAAATTCATCTTTAGCTTAGAAATCGCTTTAATAACAATGAATTAAAAGTTGGC
ACAGAAAATGCTTAAAGCTGGCATCTCTGTTAAACGGGTAACCTGACTATG
>hycA5cI
GAGGCCTCGTCTCAACCAAAGCAATCAACCCATCAACCACCTGGCATTCGCACTCTTTCGCCAAGCCAACCTCTTAGACCCTACGCTGG
GAGTTCGTAGACGGAAACAAACGCAGAATCCAAGCGTTCCAGCAGTCATAGTCAAGTTTCACACGCAAAGCCAGCCTGACCTCCTGCCA
GCAATAGTCAGTGCCCGTTGGCAGAGGGTTATTTCGTGCATATCGCCTCCCATTAACTATTGCCAGCTACAAGCAATAATTGTGCCAGT
GTTGATTATCCCTGCGGTGAATAATGTCGATGATGTCGAAATGACACGTCGACACGGCGACGAAATTCATCTTTAGCTTAGAAATCGCT
TTAATAACAATGAATTAAAAGTTGGCACAGAAAATGCTTAAAGCTGGCTCTTAATCATGCAGGGGAGGGTTTCTTATG
>hycA10cI
GAGGCCTCGTCTCAACCAAAGCAATCAACCCATCAACCACCTGGCATTCGCACTCTTTCGCCAAGCCAACCTCTTAGACCCTACGCTGG
GAGTTCGTAGACGGAAACAAACGCAGAATCCAAGCGTTCCAGCAGTCATAGTCAAGTTTCACACGCAAAGCCAGCCTGACCTCCTGCCA
GCAATAGTCAGTGCCCGTTGGCAGAGGGTTATTTCGTGCATATCGCCTCCCATTAACTATTGCCAGCTACAAGCAATAATTGTGCCAGT
GTTGATTATCCCTGCGGTGAATAATGTCGATGATGTCGAAATGACACGTCGACACGGCGACGAAATTCATCTTTAGCTTAGAAATCGCT
TTAATAACAATGAATTAAAAGTTGGCACAGAAAATGCTTAAAGCTGGCTCTTAATCATGCGGAGGATCGTTTCTTATG
>hycA15cI
GAGGCCTCGTCTCAACCAAAGCAATCAACCCATCAACCACCTGGCATTCGCACTCTTTCGCCAAGCCAACCTCTTAGACCCTACGCTGG
GAGTTCGTAGACGGAAACAAACGCAGAATCCAAGCGTTCCAGCAGTCATAGTCAAGTTTCACACGCAAAGCCAGCCTGACCTCCTGCCA
GCAATAGTCAGTGCCCGTTGGCAGAGGGTTATTTCGTGCATATCGCCTCCCATTAACTATTGCCAGCTACAAGCAATAATTGTGCCAGT
GTTGATTATCCCTGCGGTGAATAATGTCGATGATGTCGAAATGACACGTCGACACGGCGACGAAATTCATCTTTAGCTTAGAAATCGCT
TTAATAACAATGAATTAAAAGTTGGCACAGAAAATGCTTAAAGCTGGCTCTTAATCATGCGGGGGAGTCTTTCTTATG
>hycA17cI
GAGGCCTCGTCTCAACCAAAGCAATCAACCCATCAACCACCTGGCATTCGCACTCTTTCGCCAAGCCAACCTCTTAGACCCTACGCTGG
GAGTTCGTAGACGGAAACAAACGCAGAATCCAAGCGTTCCAGCAGTCATAGTCAAGTTTCACACGCAAAGCCAGCCTGACCTCCTGCCA
GCAATAGTCAGTGCCCGTTGGCAGAGGGTTATTTCGTGCATATCGCCTCCCATTAACTATTGCCAGCTACAAGCAATAATTGTGCCAGT
GTTGATTATCCCTGCGGTGAATAATGTCGATGATGTCGAAATGACACGTCGACACGGCGACGAAATTCATCTTTAGCTTAGAAATCGCT
TTAATAACAATGAATTAAAAGTTGGCACAGAAAATGCTTAAAGCTGGCTCTTAATCATGCGGAGGAGGGTTTCTTATG
>hycA23cI
GAGGCCTCGTCTCAACCAAAGCAATCAACCCATCAACCACCTGGCATTCGCACTCTTTCGCCAAGCCAACCTCTTAGACCCTACGCTGG
GAGTTCGTAGACGGAAACAAACGCAGAATCCAAGCGTTCCAGCAGTCATAGTCAAGTTTCACACGCAAAGCCAGCCTGACCTCCTGCCA
GCAATAGTCAGTGCCCGTTGGCAGAGGGTTATTTCGTGCATATCGCCTCCCATTAACTATTGCCAGCTACAAGCAATAATTGTGCCAGT
GTTGATTATCCCTGCGGTGAATAATGTCGATGATGTCGAAATGACACGTCGACACGGCGACGAAATTCATCTTTAGCTTAGAAATCGCT
TTAATAACAATGAATTAAAAGTTGGCACAGAAAATGCTTAAAGCTGGCTCTTAATCATGCGAAGGACAGTTTCTTATG
>moaAWTcI
GAGGGAGCATAGTTTCCTGGAGTCAGATTATCCGCGCTACAGTAGCGCAAATTGCGAGGAAACAGCAATTAACCTGCCAATATTGAGGA
TACAAACTATTTTCTTCTACCTCTAAAGGACGATGCACGCTATGCCTCCCTGATGATGTATATCAAAGTAAAACCGCCATTTTCCCTTA
TTCTGTAGCGAAATAGCACGATCATGACGCTATATACATGATTACATAGCGAAAGTGTGGATGGTAAAAATCTCATTTACACGCTAGTA
TCGGCTTAACCACTAAACACTCTAGCCTCTGCACCTGGGTCAACTGATACGGTGCTTTGGCCGTGACAATGCTCGTAAAGATTGCCACC
AGGGCGAAGGAAGAAATGACTTCGCCTCCCGTATCTGGAAAGGTGTATATG
>moaA5cI
GAGGGAGCCAACTCCCTTTACAACCTCACTCAAGTCCGTTAGAGCAGAGCCCTTTAGTTACCGTCATACTTCCGAAACCGATAGGCACT
GAAGGTCCTCAATCGCACTGGAAACATCAAGGTCGCTCCCATCACTCATAGTTTCCTGGAGTCAGATTATCCGCGCTACAGTAGCGCAA
ATTGCGAGGAAACAGCAATTAACCTGCCAATATTGAGGATACAAACTATTTTCTTCTACCTCTAAAGGACGATGCACGCTATGCCTCCC
TGATGATGTATATCAAAGTAAAACCGCCATTTTCCCTTATTCTGTAGCGAAATAGCACGATCATGACGCTATATACATGATTACATAGC
GAAAGTGTGGATGGTAAAAATCTCATTTACACGCTAGTATCGGCTTTATCTTAATCATGCAGGGGAGGGTTTCTTATG
>moaA10cI
GAGGGAGCCAACTCCCTTTACAACCTCACTCAAGTCCGTTAGAGCAGAGCCCTTTAGTTACCGTCATACTTCCGAAACCGATAGGACTG
AAGGTCCTCAATCGCACTGGAAACATCAAGGTCGCTCCCATCACTCATAGTTTCCTGGAGTCAGATTATCCGCGCTACAGTAGCGCAAA
TTGCGAGGAAACAGCAATTAACCTGCCAATATTGAGGATACAAACTATTTTCTTCTACCTCTAAAGGACGATGCACGCTATGCCTCCCT
GATGATGTATATCAAAGTAAAACCGCCATTTTCCCTTATTCTGTAGCGAAATAGCACGATCATGACGCTATATACATGATTACATAGCG
AAAGTGTGGATGGTAAAAATCTCATTTACACGCTAGTATCGGCTTTATCTTAATCATGCGGAGGATCGTTTCTTATG

Data Set S5 - Nissle 1917 library reference file.fasta

```
>moaA15cI
GAGGGAGCCAACTCCCTTTACAACCTCACTCAAGTCCGTTAGAGCAGAGCCCTTTAGTTACCGTCATACTTCCGAAACCGATAGGCACT
GAAGGTCCTCAATCGCACTGGAAACATCAAGGTCGCTCCCATCACTCATAGTTTCCTGGAGTCAGATTATCCGCGCTACAGTAGCGCAA
ATTGCGAGGAAACAGCAATTAACCTGCCAATATTGAGGATACAAACTATTTTCTTCTACCTCTAAAGGACGATGCACGCTATGCCTCCC
TGATGATGTATATCAAAGTAAAACCGCCATTTTCCCTTATTCTGTAGCGAAATAGCACGATCATGACGCTATATACATGATTACATAGC
GAAAGTGTGGATGGTAAAAATCTCATTTACACGCTAGTATCGGCTTTATCTTAATCATGCGGGGGAGTCTTTCTTATG
>moaA17cI
GAGGGAGCCAACTCCCTTTACAACCTCACTCAAGTCCGTTAGAGCAGAGCCCTTTAGTTACCGTCATACTTCCGAAACCGATAGGCACT
GAAGGTCCTCAATCGCACTGGAAACATCAAGGTCGCTCCCATCACTCATAGTTTCCTGGAGTCAGATTATCCGCGCTACAGTAGCGCAA
ATTGCGAGGAAACAGCAATTAACCTGCCAATATTGAGGATACAAACTATTTTCTTCTACCTCTAAAGGACGATGCACGCTATGCCTCCC
TGATGATGTATATCAAAGTAAAACCGCCATTTTCCCTTATTCTGTAGCGAAATAGCACGATCATGACGCTATATACATGATTACATAGC
GAAAGTGTGGATGGTAAAAATCTCATTTACACGCTAGTATCGGCTTTATCTTAATCATGCGGAGGAGGGTTTCTTATG
>moaA23cI
GAGGGAGCCAACTCCCTTTACAACCTCACTCAAGTCCGTTAGAGCAGAGCCCTTTAGTTACCGTCATACTTCCGAAACCGATAGGCACT
GAAGGTCCTCAATCGCACTGGAAACATCAAGGTCGCTCCCATCACTCATAGTTTCCTGGAGTCAGATTATCCGCGCTACAGTAGCGCAA
ATTGCGAGGAAACAGCAATTAACCTGCCAATATTGAGGATACAAACTATTTTCTTCTACCTCTAAAGGACGATGCACGCTATGCCTCCC
TGATGATGTATATCAAAGTAAAACCGCCATTTTCCCTTATTCTGTAGCGAAATAGCACGATCATGACGCTATATACATGATTACATAGC
GAAAGTGTGGATGGTAAAAATCTCATTTACACGCTAGTATCGGCTTTATCTTAATCATGCGAAGGACAGTTTCTTATG
>napFWTcI
GAGGTGTTAAATTCTGATGAGAAAACAAACCAGCAGAAATATCTCTGGCGAATTAATCGCTATATAAATATGTTTATAACCATTTGAAA
TGTGAGCAAAAGCCCGTTTTTGCCACGCTTCCCGCCACTCTTTTGATCCTGCTACAGGTTTTACCCCGATCGGGGTATGCATCTTTGAC
ACATCCTTTAATATCTTAGCGGCTTATG
>napF5cI
GAGGTGTTAAATTCTGATGAGAAAACAAACCAGCAGAAATATCTCTGGCGAATTAATCGCTATATAAATATGTTTATAACCATTTGAAA
TGTGAGCAAAAGCCCGTTTTTGCCACGCTTCCCGCCACTCTTTTGATCCTGCTACAGGTTTTACCCCGATCGGGGTATGCATCTTTGAC
ACATCCTTTAATATCTTAGCGGCTTCTTAATCATGCTAAGGAGGTTTTCTTTATG
>napF10cI
GAGGTGTTAAATTCTGATGAGAAAACAAACCAGCAGAAATATCTCTGGCGAATTAATCGCTATATAAATATGTTTATAACCATTTGAAA
TGTGAGCAAAAGCCCGTTTTTGCCACGCTTCCCGCCACTCTTTTGATCCTGCTACAGGTTTTACCCCGATCGGGGTATGCATCTTTGAC
ACATCCTTTAATATCTTAGCGGCTTCTTAATCATGCGGAGGATCGTTTCTTTATG
>napF15cI
GAGGTGTTAAATTCTGATGAGAAAACAAACCAGCAGAAATATCTCTGGCGAATTAATCGCTATATAAATATGTTTATAACCATTTGAAA
TGTGAGCAAAAGCCCGTTTTTGCCACGCTTCCCGCCACTCTTTTGATCCTGCTACAGGTTTTACCCCGATCGGGGTATGCATCTTTGAC
ACATCCTTTAATATCTTAGCGGCTTCTTAATCATGCGGGGGAGTCTTTCTTTATG
>napF17cI
GAGGTGTTAAATTCTGATGAGAAAACAAACCAGCAGAAATATCTCTGGCGAATTAATCGCTATATAAATATGTTTATAACCATTTGAAA
TGTGAGCAAAAGCCCGTTTTTGCCACGCTTCCCGCCACTCTTTTGATCCTGCTACAGGTTTTACCCCGATCGGGGTATGCATCTTTGAC
ACATCCTTTAATATCTTAGCGGCTTCTTAATCATGCGGAGGAGGGTTTCTTTATG
>napF23cI
GAGGTGTTAAATTCTGATGAGAAAACAAACCAGCAGAAATATCTCTGGCGAATTAATCGCTATATAAATATGTTTATAACCATTTGAAA
TGTGAGCAAAAGCCCGTTTTTGCCACGCTTCCCGCCACTCTTTTGATCCTGCTACAGGTTTTACCCCGATCGGGGTATGCATCTTTGAC
ACATCCTTTAATATCTTAGCGGCTTCTTAATCATGCGAAGGACAGTTTCTTTATG
>narKWTcI
GAGGCTCGTTCGCTGCCACCTAAGAATACTCTACGGTCACATACCCAGGATACATAGATTACCACAACTCCGAGCCCTTCCACCGTTCC
AGCAGTCATAGTCAAGTTTCACACGCAAAGCCAGCGGACACTGGCTGGATAAGAATACTCTGACTACAATCGCAGCATTACTCGCATCC
ATTCTCAGGCTGTCTCGTCTCGTCTCCATCTGCGTCACATCGGTAAGGGTAGGGATTTTACAGTACCGTGAAAAATCTCATAATTTTTA
TGAAATCACTGTACTCACTATGGATAATGATAAATATCAATGATAGATAAAGTTATCTTATCGTTTGATTTACATCAAATTGCCTTTAG
CTACAGACACTAAGGTGGCAGACATCGAAACGAGTATCAGAGGTGTCTATG
>narK5cI
GAGGCTCGTTCGCTGCCACCTAAGAATACTCTACGGTCACATACCTCCCATCACTCGTTATCCCACATCTCACGGAAACACTCGCAGAG
CCCTTTAGTTACCGTCATACTTCCGAAACCGATAGCATCTGCGTCCCGAACTCATACAGCATACAACAAGCAACCGCTGGGAGTTCGTA
GACGGAAACAAACGCAGAATCCAAGCGTTCATACGACCTCCTAATCCATCCAGCCTCCGAAACATCGGTAAGGGTAGGGATTTTACAGT
ACCGTGAAAAATCTCATAATTTTTATGAAATCACTGTACTCACTATGGATAATGATAAATATCAATGATAGATAAAGTTATCTTATCGT
TTGATTTACATCAAATTGCCTTTAGCTACAGACACTAAGGTGGCAGACTCTTAATCATGCAGGGGAGGGTTTCTTATG
>narK10cI
GAGGCTCGTTCGCTGCCACCTAAGAATACTCTACGGTCACATACCTCCCATCACTCGTTATCCCACATCTCACGGAAACACTCGCAGAG
CCCTTTAGTTACCGTCATACTTCCGAAACCGATAGCATCTGCGTCCCGAACTCATACAGCATACAACAAGCAACCGCTGGGAGTTCGTA
GACGGAAACAAACGCAGAATCCAAGCGTTCATACGACCTCCTAATCCATCCAGCCTCCGAAACATCGGTAAGGGTAGGGATTTTACAGT
ACCGTGAAAAATCTCATAATTTTTATGAAATCACTGTACTCACTATGGATAATGATAAATATCAATGATAGATAAAGTTATCTTATCGT
TTGATTTACATCAAATTGCCTTTAGCTACAGACACTAAGGTGGCAGACTCTTAATCATGCGGAGGATCGTTTCTTATG
>narK15cI
GAGGCTCGTTCGCTGCCACCTAAGAATACTCTACGGTCACATACCTCCCATCACTCGTTATCCCACATCTCACGGAAACACTCGCAGAG
CCCTTTAGTTACCGTCATACTTCCGAAACCGATAGCATCTGCGTCCCGAACTCATACAGCATACAACAAGCAACCGCTGGGAGTTCGTA
```

GACGGAAACAAACGCAGAATCCAAGCGTTCATACGACCTCCTAATCCATCCAGCCTCCGAAACATCGGTAAGGGTAGGGATTTTACAGT
ACCGTGAAAAATCTCATAATTTTTATGAAATCACTGTACTCACTATGGATAATGATAAATATCAATGATAGATAAAGTTATCTTATCGT
TTGATTTACATCAAATTGCCTTTAGCTACAGACACTAAGGTGGCAGACTCTTAATCATGCGGGGGAGTCTTTCTTATG
>narK17cI
GAGGCTCGTTCGCTGCCACCTAAGAATACTCTACGGTCACATACCTCCCATCACTCGTTATCCCACATCTCACGGAAACACTCGCAGAG
CCCTTTAGTTACCGTCATACTTCCGAAACCGATAGCATCTGCGTCCCGAACTCATACAGCATACAACAAGCAACCGCTGGGAGTTCGTA
GACGGAAACAAACGCAGAATCCAAGCGTTCATACGACCTCCTAATCCATCCAGCCTCCGAAACATCGGTAAGGGTAGGGATTTTACAGT
ACCGTGAAAAATCTCATAATTTTTATGAAATCACTGTACTCACTATGGATAATGATAAATATCAATGATAGATAAAGTTATCTTATCGT
TTGATTTACATCAAATTGCCTTTAGCTACAGACACTAAGGTGGCAGACTCTTAATCATGCGGAGGAGGGTTTCTTATG
>narK23cI
GAGGCTCGTTCGCTGCCACCTAAGAATACTCTACGGTCACATACCTCCCATCACTCGTTATCCCACATCTCACGGAAACACTCGCAGAG
CCCTTTAGTTACCGTCATACTTCCGAAACCGATAGCATCTGCGTCCCGAACTCATACAGCATACAACAAGCAACCGCTGGGAGTTCGTA
GACGGAAACAAACGCAGAATCCAAGCGTTCATACGACCTCCTAATCCATCCAGCCTCCGAAACATCGGTAAGGGTAGGGATTTTACAGT
ACCGTGAAAAATCTCATAATTTTTATGAAATCACTGTACTCACTATGGATAATGATAAATATCAATGATAGATAAAGTTATCTTATCGT
TTGATTTACATCAAATTGCCTTTAGCTACAGACACTAAGGTGGCAGACTCTTAATCATGCGAAGGACAGTTTCTTATG
>narGWTcI
GAGGGAGCCAACATCGTAATATTTGATTATCCTTTGCGCGGCATGATGTCGCGCTTTTTTTATGCGTCATTTAGTTACAACATACTAAT
GTTGTATGGCTTATTTCGCTGGATTTCATTAAGAGTCATTAATATGTTACCCATGGGGAATACTCCTTAATACCCATCTGCATAAAAAT
CTTAATGGTTTAAATAACTACAGGTGTAAAACGTCTTAATTTACAGTCTGTTATGCTATAAATGTTAATTATCCTAAAGGGGTATTCCA
GGAATTTACTTTATTTTTCATCCCCATCACTCTTGATCGTTATCAATTCCCACGCTGTTTCAGAGCGTTACCTTGCCCTTAAACATTAG
CAATGTCGATTTATCAGAGGGCCGACAGGCTCCCACAGGAGAAAACCTATG
>narG5cI
GAGGGAGCCAACTCCCTTTACAACCTCACTCAAGTCCGTTAGAGGTTCCAGCAGTCATAGTCAAGTTTCATCGTAATATTTGATTATCC
TTTGCGCGGCATGATGTCGCGCTTTTTTTATGCGTCATTTAGTTACAACATACTAATGTTGTATGGCTTATTTCGCTGGATTTCATTAA
GAGTCATTAATATGTTACCCATGGGGAATACTCCTTAATACCCATCTGCATAAAAATCTTAATGGTTTAAATAACTACAGGTGTAAAAC
GTCTTAATTTACAGTCTGTTATGCTATAAATGTTAATTATCCTAAAGGGGTATTCCAGGAATTTACTTTATTTTTCATCCCCATCACTC
TTGATCGTTATCAATTCCCACGCTGTTTCAGAGCGTTACCTTGCCCTTTCTTAATCATGCAGGGGAGGGTTTCTTATG
>narG10cI
GAGGGAGCCAACTCCCTTTACAACCTCACTCAAGTCCGTTAGAGGTTCCAGCAGTCATAGTCAAGTTTCATCGTAATATTTGATTATCC
TTTGCGCGGCATGATGTCGCGCTTTTTTTATGCGTCATTTAGTTACAACATACTAATGTTGTATGGCTTATTTCGCTGGATTTCATTAA
GAGTCATTAATATGTTACCCATGGGGAATACTCCTTAATACCCATCTGCATAAAAATCTTAATGGTTTAAATAACTACAGGTGTAAAAC
GTCTTAATTTACAGTCTGTTATGCTATAAATGTTAATTATCCTAAAGGGGTATTCCAGGAATTTACTTTATTTTTCATCCCCATCACTC
TTGATCGTTATCAATTCCCACGCTGTTTCAGAGCGTTACCTTGCCCTTTCTTAATCATGCGGAGGATCGTTTCTTATG
>narG15cI
GAGGGAGCCAACTCCCTTTACAACCTCACTCAAGTCCGTTAGAGGTTCCAGCAGTCATAGTCAAGTTTCATCGTAATATTTGATTATCC
TTTGCGCGGCATGATGTCGCGCTTTTTTTATGCGTCATTTAGTTACAACATACTAATGTTGTATGGCTTATTTCGCTGGATTTCATTAA
GAGTCATTAATATGTTACCCATGGGGAATACTCCTTAATACCCATCTGCATAAAAATCTTAATGGTTTAAATAACTACAGGTGTAAAAC
GTCTTAATTTACAGTCTGTTATGCTATAAATGTTAATTATCCTAAAGGGGTATTCCAGGAATTTACTTTATTTTTCATCCCCATCACTC
TTGATCGTTATCAATTCCCACGCTGTTTCAGAGCGTTACCTTGCCCTTTCTTAATCATGCGGGGGAGTCTTTCTTATG
>narG17cI
GAGGGAGCCAACTCCCTTTACAACCTCACTCAAGTCCGTTAGAGGTTCCAGCAGTCATAGTCAAGTTTCATCGTAATATTTGATTATCC
TTTGCGCGGCATGATGTCGCGCTTTTTTTATGCGTCATTTAGTTACAACATACTAATGTTGTATGGCTTATTTCGCTGGATTTCATTAA
GAGTCATTAATATGTTACCCATGGGGAATACTCCTTAATACCCATCTGCATAAAAATCTTAATGGTTTAAATAACTACAGGTGTAAAAC
GTCTTAATTTACAGTCTGTTATGCTATAAATGTTAATTATCCTAAAGGGGTATTCCAGGAATTTACTTTATTTTTCATCCCCATCACTC
TTGATCGTTATCAATTCCCACGCTGTTTCAGAGCGTTACCTTGCCCTTTCTTAATCATGCGGAGGAGGGTTTCTTATG
>narG23cI
GAGGGAGCCAACTCCCTTTACAACCTCACTCAAGTCCGTTAGAGGTTCCAGCAGTCATAGTCAAGTTTCATCGTAATATTTGATTATCC
TTTGCGCGGCATGATGTCGCGCTTTTTTTATGCGTCATTTAGTTACAACATACTAATGTTGTATGGCTTATTTCGCTGGATTTCATTAA
GAGTCATTAATATGTTACCCATGGGGAATACTCCTTAATACCCATCTGCATAAAAATCTTAATGGTTTAAATAACTACAGGTGTAAAAC
GTCTTAATTTACAGTCTGTTATGCTATAAATGTTAATTATCCTAAAGGGGTATTCCAGGAATTTACTTTATTTTTCATCCCCATCACTC
TTGATCGTTATCAATTCCCACGCTGTTTCAGAGCGTTACCTTGCCCTTTCTTAATCATGCGAAGGACAGTTTCTTATG
>torCWTcI
GAGGCAGAGGGTTTTACTCATTCTGTTCATATCTGTTCATATTCTGCCGTAAGCCGTTCATCCTGACCAGTGCCGCTGTTCATATTTGC
TCATTAAGATCGCTTCTATG
>torC5cI
GAGGCAGAGGGTTTTACTCATTCTGTTCATATCTGTTCATATTCTGCCGTAAGCCGTTCATCCTGACCAGTGCCGCTGTTCATATTTGC
TCATTAAGATCGCTTCTCTTAATCATGCTAAGGAGGTTTTCTTTATG
>torC10cI
GAGGCAGAGGGTTTTACTCATTCTGTTCATATCTGTTCATATTCTGCCGTAAGCCGTTCATCCTGACCAGTGCCGCTGTTCATATTTGC
TCATTAAGATCGCTTCTCTTAATCATGCGGAGGATCGTTTCTTTATG
>torC15cI
GAGGCAGAGGGTTTTACTCATTCTGTTCATATCTGTTCATATTCTGCCGTAAGCCGTTCATCCTGACCAGTGCCGCTGTTCATATTTGC
TCATTAAGATCGCTTCTCTTAATCATGCGGGGGAGTCTTTCTTTATG

>torC17cI
GAGGCAGAGGGTTTTACTCATTCTGTTCATATCTGTTCATATTCTGCCGTAAGCCGTTCATCCTGACCAGTGCCGCTGTTCATATTTGC
TCATTAAGATCGCTTCTCTTAATCATGCGGAGGAGGGTTTCTTTATG
>torC23cI
GAGGCAGAGGGTTTTACTCATTCTGTTCATATCTGTTCATATTCTGCCGTAAGCCGTTCATCCTGACCAGTGCCGCTGTTCATATTTGC
TCATTAAGATCGCTTCTCTTAATCATGCGAAGGACAGTTTCTTTATG
>yeaRWTcI
GAGGGTTCCAGCAGTCATAGTCAAGTTTCACACGCAAAGCCAGCCTCGTTCGCTGCCACCTAAGAATACTCTACGGTCACATACGTTCC
ACTTAGACACCGTATCTTTCCCTCCAATCCAGAGCGCTGGGAGTTCGTAGACGGAAACAAACGCAGAATCCAAGCGGACACTGGCTGGA
TAAAATACTCTGACTACAATCGCAGGACACACTGTATCTGACAACGACGCACCTGCTTTCAGCCCGCGTTGCCGCGGGCTTCCCACCTG
TAATCCTCCCTGATTCTTCTCTGATCCGGTGCTAAAAAGTAACCAATAAATGGTATTTAAAATGCAAATTATCAGGCGTACCCTGAAAC
GGCTGGAATAAACCGTTTTCAGCGCATTCACCGAAGGAGGGAAAAGTATG
>yeaR5cI
GAGGGTTCCAGCAGTCATAGTCAAGTTTCACACGCAAAGCCAGCCATCTGCGTCCCGAACTCATACAGCATACAACAAGCAACCCATCA
TCTGTAACCACTACTAACGCTCATCTCTGTCGCAGCTCTCACTTCTTCACCATTACAACCATACCCATTCCGCAGCATTCGCACTCTTT
CGCCAAGCCAACCTCTTAGACCCTACGCTGGGAGTTCGTAGACGGAAACAAACGCAGAATCCAAGCCTCGTTCGCTGCCACCTAAGAAT
ACTCTACGGGTTGCTTTCAGCCCGCGTTGCCGCGGGCTTCCCACCTGTAATCCTCCCTGATTCTTCTCTGATCCGGTGCTAAAAAAACC
AATAAATGGTATTTAAAATGCAAATTATCAGGCGTACCCTGAAACGTCTTAATCATGCAGGGGAGGGTTTCTTATG
>yeaR10cI
GAGGGTTCCAGCAGTCATAGTCAAGTTTCACACGCAAAGCCAGCCATCTGCGTCCCGAACTCATACAGCATACAACAAGCAACCCATCA
TCTGTAACCACTACTAACGCTCATCTCTGTCGCAGCTCTCACTTCTTCACCATTACAACCATACCCATTCCGCAGCATTCGCACTCTTT
CGCCAAGCCAACCTCTTAGACCCTACGCTGGGAGTTCGTAGACGGAAACAAACGCAGAATCCAAGCCTCGTTCGCTGCCACCTAAGAAT
ACTCTACGGGTTGCTTTCAGCCCGCGTTGCCGCGGGCTTCCCACCTGTAATCCTCCCTGATTCTTCTCTGATCCGGTGCTAAAAAGTAA
CCAATAAATGGTATTTAAAATGCAAATTATCAGGCGTACCCTGAAACGTCTTAATCATGCGGAGGATCGTTTCTTATG
>yeaR15cI
GAGGGTTCCAGCAGTCATAGTCAAGTTTCACACGCAAAGCCAGCCATCTGCGTCCCGAACTATACAGCATACAACAAGCAACCCATCAT
CTGTAACCACTACTAACGCTCATCTCTGTCGCAGCTCTCACTTCTTCACCATTACAACCATACCCATTCCGCAGCATTCGCACTCTTTC
GCCAAGCCAACCTCTTAGACCCTACGCTGGGAGTTCGTAGACGGAAACAAACGCAGAATCCAAGCCTCGTTCGCTGCCACCTAAGAATA
CTCTACGGGTTGCTTTCAGCCCGCGTTGCCGCGGGCTTCCCACCTGTAATCCTCCCTGATTCTTCTCTGATCCGGTGCTAAAAAGTAAC
CAATAAATGGTATTTAAAATGCAAATTATCAGGCGTACCCTGAAACGTCTTAATCATGCGGGGGAGTCTTTCTTATG
>yeaR17cI
GAGGGTTCCAGCAGTCATAGTCAAGTTTCACACGCAAAGCCAGCCATCTGCGTCCCGAACTCATACAGCATACAACAAGCAACCCATCA
TCTGTAACCACTACTAACGCTCATCTCTGTCGCAGCTCTCACTTCTTCACCATTACAACCATACCCATTCCGCAGCATTCGCACTCTTT
CGCCAAGCCAACCTCTTAGACCCTACGCTGGGAGTTCGTAGACGGAAACAAACGCAGAATCCAAGCCTCGTCAGAATCCAAGCCTCGTT
CGCTGCCACCTAAGAATACTCTACGGGTTGCTTTCAGCCCGCGTTGCCGCGGGCTTCCCACCTGTAATCCTCCCTGATTCTTCTCTGAT
CCGGTGCTAAAAAGTAACCAATAAATGGTATTTAAAATGCAAATTATCAGGCGTACCCTGAAACGTCTTAATCATGCGGAGGAGGGTTT
CTTATG
>yeaR23cI
GAGGGTTCCAGCAGTCATAGTCAAGTTTCACACGCAAAGCCAGCCATCTGCGTCCCGAACTCATACAGCATACAACAAGCAACCCATCA
TCTGTAACCACTACTAACGCTCATCTCTGTCGCAGCTCTCACTTCTTCACCATTACAACCATACCCATTCCGCAGCATTCGCACTCTTT
CGCCAAGCCAACCTCTTAGACCCTACGCTGGGAGTTCGTAGACGGAAACAAACGCAGAATCCAAGCCTCGTTCGCTGCCACCTAAGAAT
ACTCTACGGGTTGCTTTCAGCCCGCGTTGCCGCGGGCTTCCCACCTGTAATCCTCCCTGATTCTTCTCTGATCCGGTGCTAAAAAGTAA
CCAATAAATGGTATTTAAAATGCAAATTATCAGGCGTACCCTGAAACGTCTTAATCATGCGAAGGACAGTTTCTTATG
>yedYWTcI
GAGGCATCTGCGTCCCGAACTCATACAGCATACAACAAGCAACCCTGACCTCCTGCCAGCAATAGTAAGACAACACGCAAAGTCCATTC
GCACTCTTTCGCCAAGCCAACCTCTTAGACCCTACCAGAGCCCTTTAGTTACCGTCATACTTCCGAAACCGATAGCAAGACGCTGGCTC
TGACATTTCCGCTACTGAACTACTCGCATTACTCGCATCCATTCTCAGGCTGTCTCGTCTCGTCTCGTCCGAACTACCGATCACATCCA
CCAAGCGTTACACAGCGCTGGGAGTTCGTGCGTAAATATGTAAAAGCTGATCTCTGCTACTGTTAGGGAGGTGACTCCCTGACAGATTA
ACAGTAAACGGCTCTTGTTGGCTAACGACAAAAAAGTGTGATGGCTTATG
>yedY5cI
GAGGCATCTGCGTCCCGAACTCATACAGCATACAACAAGCAACCCTTACTGGTCTGTCTGGTCAAATCTACGGGTTCCCTAACGCATTC
GCACTCTTTCGCCAAGCCAACCTCTTAGACCCTACGGACACTGGCTGGATAAGAATACTCTGACTACAATCGCAGCAGAGCCCTTTAGT
TACCGTCATACTTCCGAAACCGATAGGCACTGAAGGTCCTCAATCGCACTGGAAACATCAAGGTCGCAAGACGCTGGCTCTGACATTTC
CGCTACTGAACTACTCGCTCTCACTTCTTCACCATTACAACCATACCCATTCCGCAGGCTGGGAGTTCGTAGACTGCGTAAATATGTAA
AAGCTGATCTCTGCTACTGTTAGGGAGGTGACTCCCTGACAGATTAACTCTTAATCATGCAGGGGAGGGTTTCTTATG
>yedY10cI
GAGGCATCTGCGTCCCGAACTCATACAGCATACAACAAGCAACCCTTACTGGTCTGTCTGGTCAAATCTACGGGTTCCCTAACGCATTC
GCACTCTTTCGCCAAGCCAACCTCTTAGACCCTACGGACACTGGCTGGATAAGAATACTCTGACTACAATCGCAGCAGAGCCCTTTAGT
TACCGTCATACTTCCGAAACCGATAGGCACTGAAGGTCCTCAATCGCACTGGAAACATCAAGGTCGCAAGACGCTGGCTCTGACATTTC
CGCTACTGAACTACTCGCTCTCACTTCTTCACCATTACAACCATACCCATTCCGCAGGCTGGGAGTTCGTAGACTGCGTAAATATGTAA
AAGCTGATCTCTGCTACTGTTAGGGAGGTGACTCCCTGACAGATTAACTCTTAATCATGCGGAGGATCGTTTCTTATG
>yedY15cI

```
GAGGCATCTGCGTCCCGAACTCATACAGCATACAACAAGCAACCCTTACTGGTCTGTCTGGTCAAATCTACGGGTTCCCTAACGCATTC
GCACTCTTTCGCCAAGCCAACCTCTTAGACCCTACGGACACTGGCTGGATAAGAATACTCTGACTACAATCGCAGCAGAGCCCTTTAGT
TACCGTCATACTTCCGAAACCGATAGGCACTGAAGGTCCTCAATCGCACTGGAAACATCAAGGTCGCAAGACGCTGGCTCTGACATTTC
CGCTACTGAACTACTCGCTCTCACTTCTTCACCATTACAACCATACCCATTCCGCAGGCTGGGAGTTCGTAGACTGCGTAAATATGTAA
AAGCTGATCTCTGCTACTGTTAGGGAGGTGACTCCCTGACAGATTAACTCTTAATCATGCGGGGGAGTCTTTCTTATG
>yedY17cI
GAGGCATCTGCGTCCCGAACTCATACAGCATACAACAAGCAACCCTTACTGGTCTGTCTGGTCAAATCTACGGGTTCCCTAACGCATTC
GCACTCTTTCGCCAAGCCAACCTCTTAGACCCTACGGACACTGGCTGGATAAGAATACTCTGACTACAATCGCAGCAGAGCCCTTTAGT
TACCGTCATACTTCCGAAACCGATAGGCACTGAAGGTCCTCAATCGCACTGGAAACATCAAGGTCGCAAGACGCTGGCTCTGACATTTC
CGCTACTGAACTACTCGCTCTCACTTCTTCACCATTACAACCATACCCATTCCGCAGGCTGGGAGTTCGTAGACTGCGTAAATATGTAA
AAGCTGATCTCTGCTACTGTTAGGGAGGTGACTCCCTGACAGATTAACTCTTAATCATGCGGAGGAGGGTTTCTTATG
>yedY23cI
GAGGCATCTGCGTCCCGAACTCATACAGCATACAACAAGCAACCCTTACTGGTCTGTCTGGTCAAATCTACGGGTTCCCTAACGCATTC
GCACTCTTTCGCCAAGCCAACCTCTTAGACCCTACGGACACTGGCTGGATAAGAATACTCTGACTACAATCGCAGCAGAGCCCTTTAGT
TACCGTCATACTTCCGAAACCGATAGGCACTGAAGGTCCTCAATCGCACTGGAAACATCAAGGTCGCAAGACGCTGGCTCTGACATTTC
CGCTACTGAACTACTCGCTCTCACTTCTTCACCATTACAACCATACCCATTCCGCAGGCTGGGAGTTCGTAGACTGCGTAAATATGTAA
AAGCTGATCTCTGCTACTGTTAGGGAGGTGACTCCCTGACAGATTAACTCTTAATCATGCGAAGGACAGTTTCTTATG
>ynfEWTcI
GAGGGGACACTGGCTGGATAAGAATACTCTGACTACAATCGCAGCCAGGATACATAGATTACCACAACTCCGAGCCCTTCCACCCAAGA
CGCTGGCTCTGACATTTCCGCTACTGAACTACTCGGCACTGAAGGTCCTCAATCGCACTGGAAACATCAAGGTCGCCTCGTCTCAACCA
AAGCAATCAACCCATCAACCACCTGGCCCACATTATCGACTGAACGCCGGATACGACACATCCGGCGATTTGAACATACAACATAATCC
CACCTTATTACTCATACCCTTCTATTGATATGGATTAATAATTATTAACCCCAAATGGGTAGACTCTCTCTATTGTTAGCGCGCTAAAT
ATTCAATATATAAACATTTATATAACGATAAAGAACAGGGAGTGAGTTATG
>ynfE5cI
GAGGGGACACTGGCTGGATAAGAATACTCTGACTACAATCGCAGCTCTCACTTCTTCACCATTACAACCATACCCATTCCGCAGCAGAG
CCCTTTAGTTACCGTCATACTTCCGAAACCGATAGCATTACTCGCATCCATTCTCAGGCTGTCTCGTCTCGTCTCCATCTGCGTCCCGA
ACTCATACAGCATACAACAAGCAACCGACACACTGTATCTGACAACGACGCACTCTGGTAACCCACCTCCCATCACTCGTTATCCCACA
TCTCACACATTATCGACTGAACGCCGGATACGACACATCCGGCGATTTGAACATACAACATAATCCCACCTTATTACTCATACCCTTCT
ATTGATATGGATTAATAATTATTAACCCCAAATGGGTAGACTCTCTCTTCTTAATCATGCAGGGGAGGGTTTCTTATG
>ynfE10cI
GAGGGGACACTGGCTGGATAAGAATACTCTGACTACAATCGCAGCTCTCACTTCTTCACCATTACAACCATACCCATTCCGCAGCAGAG
CCCTTTAGTTACCGTCATACTTCCGAAACCGATAGCATTACTCGCATCCATTCTCAGGCTGTCTCGTCTCGTCTCCATCTGCGTCCCGA
ACTCATACAGCATACAACAAGCAACCGACACACTGTATCTGACAACGACGCACTCTGGTAACCCACCTCCCATCACTCGTTATCCCACA
TCTCACACATTATCGACTGAACGCCGGATACGACACATCCGGCGATTTGAACATACAACATAATCCCACCTTATTACTCATACCCTTCT
ATTGATATGGATTAATAATTATTAACCCCAAATGGGTAGACTCTCTCTTCTTAATCATGCGGAGGATCGTTTCTTATG
>ynfE15cI
GAGGGGACACTGGCTGGATAAGAATACTCTGACTACAATCGCAGCTCTCACTTCTTCACCATTACAACCATACCCATTCCGCAGCAGAG
CCCTTTAGTTACCGTCATACTTCCGAAACCGATAGCATTACTCGCATCCATTCTCAGGCTGTCTCGTCTCGTCTCCATCTGCGTCCCGA
ACTCATACAGCATACAACAAGCAACCGACACACTGTATCTGACAACGACGCACTCTGGTAACCCACCTCCCATCACTCGTTATCCCACA
TCTCACACATTATCGACTGAACGCCGGATACGACACATCCGGCGATTTGAACATACAACATAATCCCACCTTATTACTCATACCCTTCT
ATTGATATGGATTAATAATTATTAACCCCAAATGGGTAGACTCTCTCTTCTTAATCATGCGGGGGAGTCTTTCTTATG
>ynfE17cI
GAGGGGACACTGGCTGGATAAGAATACTCTGACTACAATCGCAGCTCTCACTTCTTCACCATTACAACCATACCCATTCCGCAGCAGAG
CCCTTTAGTTACCGTCATACTTCCGAAACCGATAGCATTACTCGCATCCATTCTCAGGCTGTCTCGTCTCGTCTCCATCTGCGTCCCGA
ACTCATACAGCATACAACAAGCAACCGACACACTGTATCTGACAACGACGCACTCTGGTAACCCACCTCCCATCACTCGTTATCCCACA
TCTCACACATTATCGACTGAACGCCGGATACGACACATCCGGCGATTTGAACATACAACATAATCCCACCTTATTACTCATACCCTTCT
ATTGATATGGATTAATAATTATTAACCCCAAATGGGTAGACTCTCTCTTCTTAATCATGCGGAGGAGGGTTTCTTATG
>ynfE23cI
GAGGGGACACTGGCTGGATAAGAATACTCTGACTACAATCGCAGCGACACACTGTATCTGACAACGACGCACTCTGGTAACCCACCTCC
CATCACTCGTTATCCCACATCTCACACATTATCGACTGAACGCCGGATACGACACATCCGGCGATTTGAACATACAACATAATCCCACC
TTATTACTCATACCCTTCTATTGATATGGATTAATAATTATTAACCCCAAATGGGTAGACTCTCTCTTCTTAATCATGCGAAGGACAGT
TTCTTATG
>fabR
GAGGCAATTCCGACGTCTAAGAAACCATTATTATCATGACATTAACCTATAAAAATAGGCGTATCACGAGGCCCTTTCGTCTTCACCTC
GAGAGCTTTCGACGGGATGGTGCCCCAGTGGGTGCAACCGCCGCCAACATTTTGATAACGCTCGATAACTGCGACGCGCGCACCTTGCT
TAACCAGGCCCATTGCAGCGCCTTCGCCGCCGGGGCCGGAACCTATTACTATGGCATCGTAATCGTAGGAATGTGGCATGGTAGGGCTT
ACCTGTTCTTATACATAAAAGCAACAGAATGGTAACATTTTATCGCGGGTAAGCCAATTGATCCCCGTCATTTATCTGGCTATATCCTG
AGCGGCCTTTGCTTTGTCTGTTTCTTACTTTTGCCCTGACGTTTTATTGGATTTTTATCGACGATACTCTCCGTTTAAGCGGCAGGTTT
CCGCTGTACGTAAAAGAACCGGCCAAAGAATTGCAGTAAATATGTTTTATTGCGTTACCGTTCATTCACAATACTGGAGCAATCCAGTA
TGTTCATTCTCTGGTATAGTGCCAGCAGTACTTTTGGCAAGGATTCAGACATCGTGATGGGCGTAAGAGCGCAACAAGGATCCTCTAGA
TTTAAGAAGGAGATATACATATG
```
